# The motif composition of variable-number tandem repeats impacts gene expression

**DOI:** 10.1101/2022.03.17.484784

**Authors:** Tsung-Yu Lu, Paulina N. Smaruj, Geoffrey Fudenberg, Nicholas Mancuso, Mark J.P. Chaisson

## Abstract

Understanding the impact of DNA variation on human traits is a fundamental question in human genetics. Variable number tandem repeats (VNTRs) make up roughly 3% of the human genome but are often excluded from association analysis due to poor read mappability or divergent repeat content. While methods exist to estimate VNTR length from short-read data, it is known that VNTRs vary in both length and repeat (motif) composition. Here, we use a repeat-pangenome graph (RPGG) constructed on 35 haplotype-resolved assemblies to detect variation in both VNTR length and repeat composition. We align population scale data from the Genotype-Tissue Expression (GTEx) Consortium to examine how variations in sequence composition may be linked to expression, including cases independent of overall VNTR length. We find that 9,422 out of 39,125 VNTRs are associated with nearby gene expression through motif variations, of which only 23.4% associations are accessible from length. Fine-mapping identifies 174 genes to be likely driven by variation in certain VNTR motifs and not overall length. We highlight two genes, *CACNA1C* and *RNF213* that have expression associated with motif variation, demonstrating the utility of RPGG analysis as a new approach for trait association in multiallelic and highly variable loci.

## Introduction

Variable number tandem repeats (VNTRs) are repetitive DNA sequences with the size of a repeat unit greater than six nucleotides. The copy number of a repeat unit is hypervariable due to its susceptibility to replication slippage caused by strand mispairing between the same (Viguera et al. 2001) or across haplotypes (Jeffreys et al. 1994). At the sequence level, single-nucleotide variants (SNVs) or short indels are also prevalent along a repeat sequence and can greatly expand the number of identified alleles relative to classification by length (Novroski et al. 2016). Altogether, copy number variations, SNVs and short indels contribute to the full spectrum of VNTR polymorphism. Missing heritability (Eichler et al. 2010) that cannot be explained by SNVs can be partially attributed to VNTR polymorphisms (Hannan 2018; Mitra et al. 2021; Lu and Chaisson 2021). Accumulating evidence indicates that VNTRs are associated with a diverse array of human traits and are casual to several diseases at the copy number level or sequence level (Beyter et al. 2021; Bakhtiari et al. 2021; Mukamel et al. 2021). Furthermore, significant enrichment of VNTRs in subtelomeric genes that are mostly expressed in the brain suggests further exploration of their roles in shaping behavioral/cognitive polymorphisms and modulating neurological disease risks (Linthorst et al. 2020).

VNTR length polymorphism can modulate human traits through several mechanisms, including changing the number of protein domains (Desseyn et al. 2000), the distance between gene and gene regulators (Sun et al. 2018), the number of regulator binding sites (Tsuge et al. 2005), and the number of CpG sites (DeJesus-Hernandez et al. 2011; Renton et al. 2011). Abundant associations between repeat copy number and human traits have been widely reported (Mukamel et al. 2021) and provide insights to functional annotations. However, it is impossible to fully understand the biological functions of VNTRs without examining variation at the sequence level. For example, a single cytosine insertion in *MUC1* VNTR was identified to be causal to medullary cystic kidney disease type 1 by adding a premature stop in translation (Kirby et al. 2013). In addition, certain repeat motifs in *CACNA1C* but not the total repeat copy number were reported to be associated with schizophrenia risk by tuning gene expression activity (Song et al. 2018). In both cases, long-read sequencing such as single-molecule real-time sequencing or capillary sequencing has been useful to resolve the full sequence of VNTRs and yield meaningful clinical interpretations.

Currently large-scale sequencing efforts use high-throughput short-read sequencing (SRS) (Taliun et al. 2021), however, VNTR analysis with SRS suffers from ambiguity in read alignment, allelic bias of reference and the hypermutability of repeat sequences. Single-nucleotide and small indel variant calls from VNTR regions using short-read alignments are error-prone and blacklisted by ENCODE (Amemiya et al. 2019). Recently, several methods have been developed specifically to estimate VNTR length from short-read data using Hidden Markov Models (Bakhtiari et al. 2021), read-depth (Garg et al. 2021), and repeat-pangenome graphs (Lu and Chaisson 2021). These approaches have found an association between estimated VNTR length and gene expression. In this study, we use a reference pangenome graph (PGG) to reduce allelic bias when mapping short reads to a reference, and to improve variant inference for motif composition. The PGG is a graph-based data structure that summarizes sequence variations from a collection of samples by representing variants as alternative paths or “bubbles” from the reference (Eizenga et al. 2020). One of the most common implementations of PGG is to use a sequence graph. In this model, each node represents an allele; each edge points to a downstream allele; a traversal through the graph matches an observed haplotype. This allows sequencing reads to be placed more accurately across the genome and significantly improves variant calling accuracy in regions containing SVs (Chen et al. 2019; Eggertsson et al. 2019; Li et al. 2020; Sirén et al. 2020), with the majority of which coming from indel events within VNTRs (Li et al. 2020). However, variant calling remains challenging for multiallelic VNTR regions as the position of calls varies (Li et al. 2020); an extra processing step is needed to reveal the multiallelic property of a locus.

Another commonly used graph model is the de Bruijn graph (dBG). The main distinction is that each node is a unique *k*-mer derived from one or more *k*-base substrings present in the input sequences. By augmenting with additional haplotype or distance information, dBG-based models have been useful in genome assembly (Iqbal et al. 2012; Muggli et al. 2017) and variant calling (Cameron et al. 2017; Narzisi et al. 2018). Furthermore, this formulation groups all occurrences of repetitive *k*-mers across input sequences by construction, which can be a particularly desirable property when studying the biological implication of VNTR motifs.

By leveraging the advantages of PGG and dBG, genotyping VNTR from SRS samples at a population scale has been made possible with danbing-tk (Lu and Chaisson 2021). The method constructs a repeat-pangenome graph (RPGG) that consists of disjoint locus-RPGGs, each representing a single VNTR locus and encoding observed VNTR alleles with a dBG. Read mapping to RPGG reveals the coverage of each *k*-mer and can be accumulated as a copy number estimate, allowing associations between repeat copy number and human trait to be identified.

In this work, we extend the application danbing-tk to examine the association between each path in the graph, or VNTR “motif”, and gene expression using the complete read-mapping output i.e. the coverages of all *k*-mers. The extension includes new depth normalization approaches that accurately show motif usage/repeat count in a RPGG, and that this may be used to compare motif composition between individuals. To demonstrate the utility of this method, we profiled motif composition from genomes sequenced by the Genotype-Tissue Expression (GTEx) project and Geuvaids to identify motif *cis*-eQTLs. We envision this as a framework for genetics studies to associate tandem repeat variation to traits when only short-read data are available.

## Results

### Repeat pangenome graphs enable accurate profiling of motif composition

We developed an extended computational analysis pipeline based on the previously published danbing-tk method to map read depth and identify eQTLs from individual paths in an RPGG (Fig. 1). The RPGG is constructed using 35 haplotype-resolved assemblies including three trios released by the Human Genome Structural Variation Consortium (HGSVC) (Ebert et al. 2021). Orthologous boundaries of 80,478 VNTR loci were annotated using danbing-tk (Lu and Chaisson 2021) from a set of 84,411 VNTRs (Methods). We further augment the VNTR annotations with additional 40 clinically relevant loci (Supplemental Table S1) and removed 40,204 loci that contain tandem mobile elements, giving a total of 40,314 loci (Supplemental Data S1) for subsequent analyses. The VNTRs annotated have a mean length of 403 bp across assemblies versus 371 bp in GRCh38 (Fig. 2A, Supplemental Fig. S1). The assemblies give a limited estimate of VNTR diversity. Among the 70 haplotypes, a VNTR has an average of 8.7 alleles per locus when defining an allele based on exact length (Supplemental Fig. S2). Each locus has an average of 3.9 alleles that are observed only once, denoted as private allele count, and 513 loci that have at least half the alleles (N≥35) as private (Supplemental Fig. S2). The number of alleles per locus is positively correlated with VNTR length (Pearson’s *r* = 0.43, P<10^-323^). As VNTR length increases, the private allele count also increases (Supplemental Fig. S2), e.g. private allele count = 8.9 when VNTR length > 500 bp.

**Figure 1.**
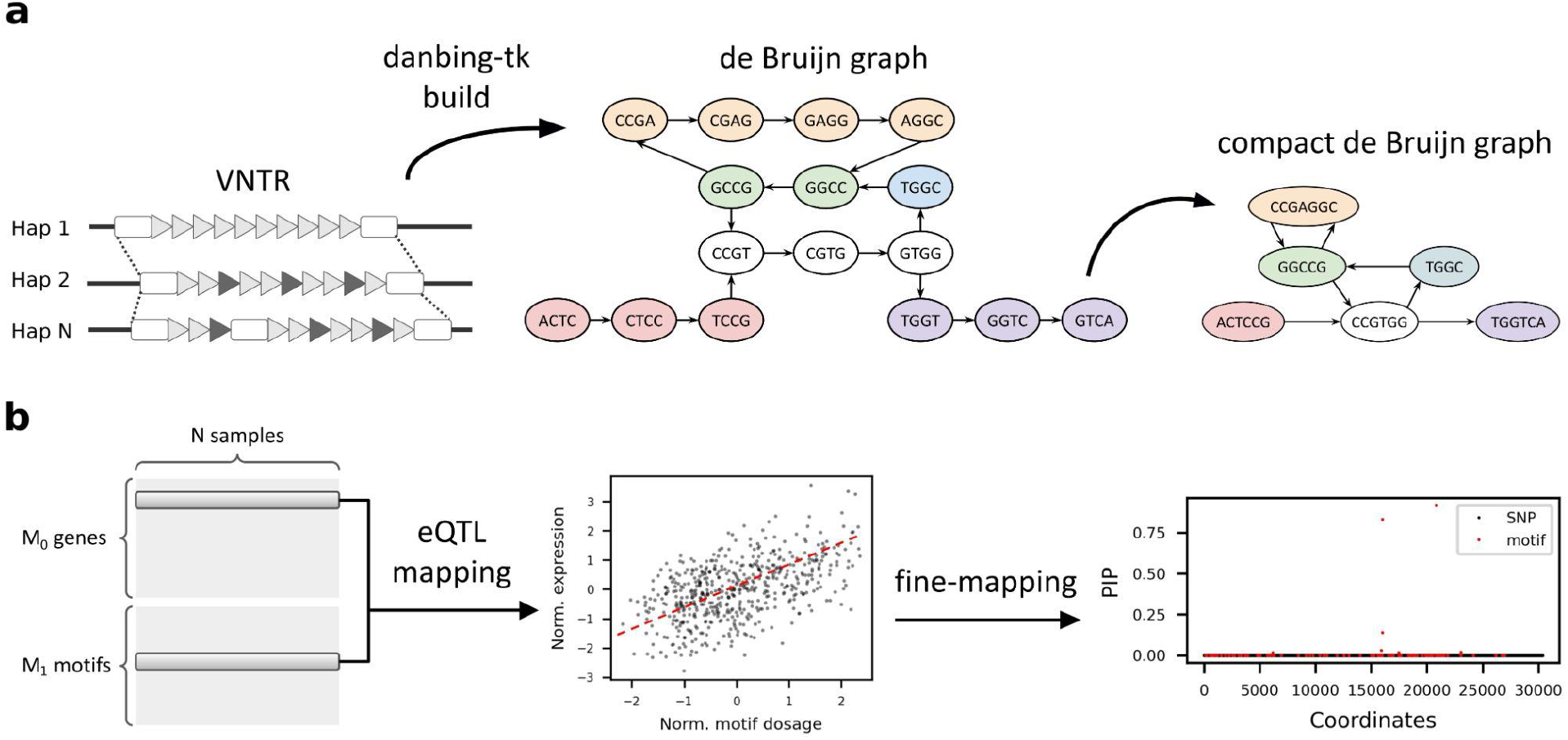
Methods overview. **a,** Estimating the dosages of VNTR motifs using a locus-RPGG. A locus-RPGG is built from haplotype-resolved assemblies by first annotating the orthology mapping of VNTR boundaries and then encoding the VNTR alleles with a de Bruijn graph (dBG), or locus-RPGG. A compact dBG is constructed by merging nodes on a non-branching path into a unitig, denoted as a motif in this context. Motif dosages of a VNTR can be computed by aligning short reads to an RPGG and averaging the coverage of nodes corresponding to the same motif. **b,** Identifying likely causal eMotifs. The dosage of each motif is tested against the expression level of a nearby gene. Genes in significant association with at least one motif (denoted as eMotif) are fine-mapped using susieR (Wang et al. 2020) in order to identify eMotifs that are likely causal to nearby gene expression. All GTEx variants (Methods) and lead motifs from each gene-VNTR pair are included in the fine-mapping model.

**Figure 2.**
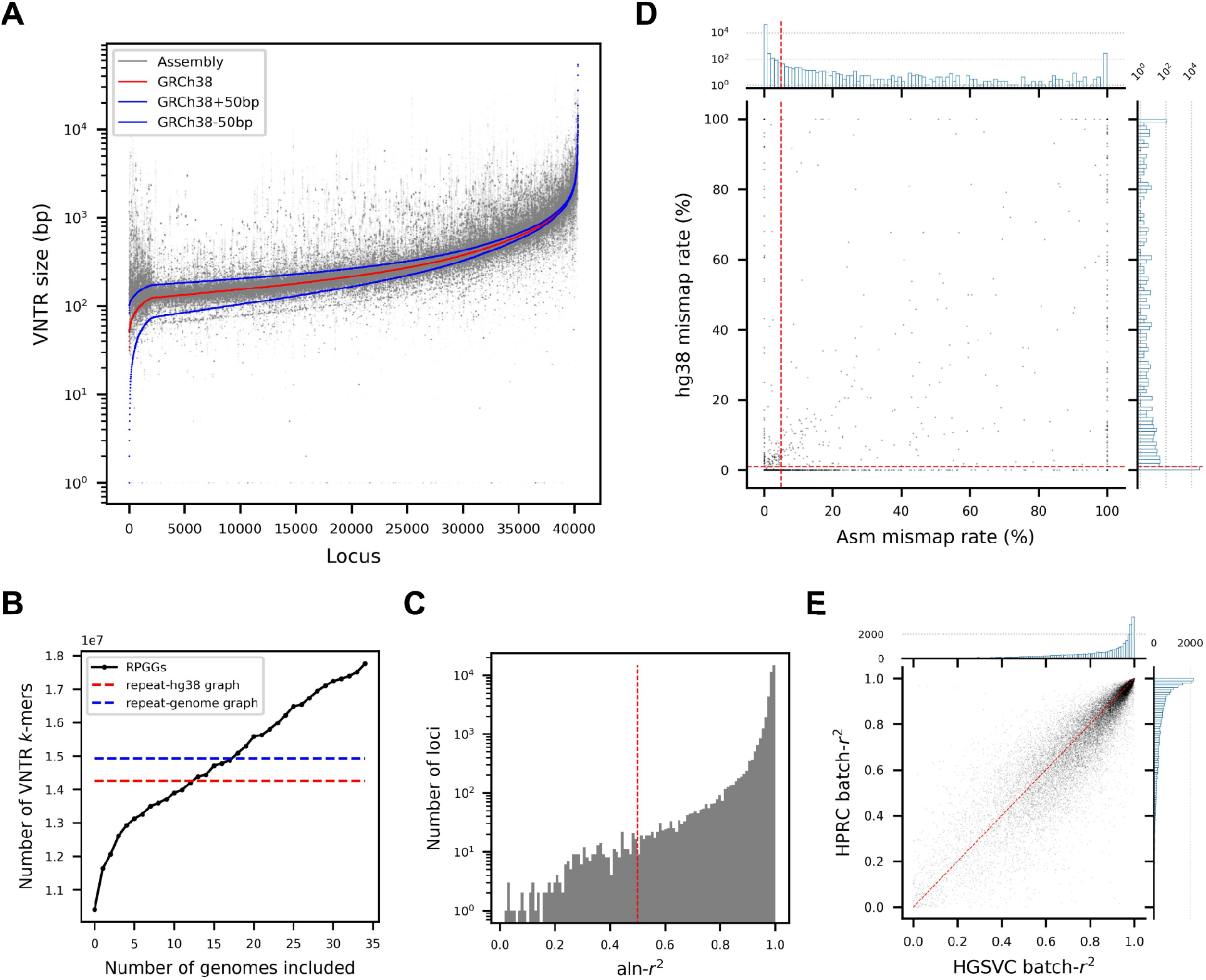
Characteristics of VNTRs and the RPGG. (A) Size distribution of VNTR alleles across 35 HGSVC assemblies. Each dot represents the size of a VNTR locus in an assembly. The order of 40,314 VNTR loci were sorted according to size in GRCh38. (B) Cumulative graph sizes. A total of 35 repeat-genome graphs were incrementally added to the RPGG in the order of their respective graph size. The red dash line denotes the size of the repeat graph built from GRCh38. The blue dash line denotes the average size of the graphs built from assemblies. (C) Distribution of aln-*r*^2^ for all locus-RPGGs. The aln-*r*^2^ of each locus was computed by regressing the assembly *k*-mer counts against the read *k*-mer counts from graph alignments. The cutoff for aln-*r*^2^ (0.5) is shown in red dotted lines. (D) Distribution of assembly and GRCh38 mismap rates for all locus-RPGGs. The cutoffs for assembly (5%) and GRCh38 (1%) mismap rates are shown in red dotted lines. The histograms represent the marginal distribution for each variable. (E) Distribution of VNTR dosage between-sample correlation batch-*r*^2^ on the HGSVC and HPRC datasets, with the diagonal with slope of 1 in red.

We used danbing-tk (Lu and Chaisson 2021) to encode the allele information across haplotypes in an RPGG, consisting of 40,314 locus subgraphs, or locus-RPGGs. The graph has a total of 173,944,578 *k*-mers (*k*=21) or nodes, and 176,697,311 edges, with an average of 4,315 nodes in each locus-RPGG when including 700 bp flanking sequences on both sides. Each locus-RPGG has an average of 102 nodes or 13.4% nodes that are observed only in one assembly. The repeat region of RPGG (excluding flanking sequences) has 17,762,872 *k*-mers (Fig. 2B), which is 25% greater than the graph built from GRCh38 alone (n=14,257,939) and 71% greater than the smallest graph built from an assembly (HG00864, n=10,413,896).

We evaluated the quality of the alignments to each locus-RPGG by measuring the consistency of *k*-mer counts from a diploid assembly versus from mapping of short-read data from the same sample, denoted as aln-*r*^2^ (Methods). Overall, there is a slightly higher aln-*r*^2^ of 0.96 (Fig. 2C) compared to the aln-*r*^2^ (0.93) on the previously published 19 haplotype-resolved assemblies, with enrichment of loci with higher aln-*r*^2^ (Supplemental Fig. S3). We also measured read mismapping rates from reads simulated from and aligned back to each assembly and comparing the reads incorrectly mapped to a VNTR to those mapping correctly. The majority of loci (n=39,560) had less than 5% mismap rate (Fig. 2D). To account for sequences missing from the HGSVC assemblies, we also considered the mismap rate from GRCh38-only simulations to filter problematic loci, selecting mismapping rate thresholds based on variance of mismap rate among loci (Fig. 2D). VNTR loci with aln-*r*^2^ > 0.5, assembly mismap rate < 5%, and GRCh38 mismap rate < 1% were retained (n=39,125) for further analysis. The above approaches removed 775 loci from segmental duplications with high similarity to another copy and retained 544 loci within regions that are still distinguishable from other copies. To ensure data quality, we genotyped all 40,314 loci so that low quality loci can act as “baits” for reads that tend to mismap.

Even though *k*-mer compositions in the majority of VNTRs show high consistency with ground truths within each sample (37,840 loci with aln-*r*^2^ > 0.8), we observed that read depth variation due to unknown bias or sampling stochasticity could cause *k*-mer dosages to vary across samples even for loci where ground truth VNTR lengths are identical (Supplemental Fig. S4). We corrected these biases by searching for VNTR *k*-mers that occur the same number of times throughout the 70 HGSVC haplotypes, denoted as invariant *k*-mers. For each locus in an individual, all *k*-mer dosages were scaled by a constant factor (Supplemental Data S2) so that the average dosage of invariant *k*-mers were the same across samples. We assessed the effect of bias correction by measuring the *k*-mer dosage consistency across samples, denoted as batch-*r*^2^. When summing up all *k*-mer dosages as an estimate for VNTR length, we saw an increase in batch-*r*^2^ from 0.531 to 0.820 after bias correction (Supplemental Fig. S5). Applying the same procedure to 36 Human Pangenome Reference Consortium (HPRC) assemblies (Liao et al. 2022) with matching short-read samples in the 1000 Genomes Project (1KGP) (Byrska-Bishop et al. 2022), we also observed higher batch-*r*^2^ after bias correction (0.521 versus 0.795, Supplemental Fig. S5). The aln-*r*^2^ metric indicates that we can accurately genotype VNTR *k*-mer compositions within a sample while the batch-*r*^2^ further confirms that VNTR dosages are scaled properly across samples (Fig. 2E), warranting analyses at a population level.

### VNTR motif composition has pervasive *cis*-effects on gene expression

Using the short-read alignment module in danbing-tk, we estimated the VNTR content of 39,125 loci as graph genotypes and processed 838 GTEx genomes (GTEx Consortium 2020) using ~12 cpu hours and ~29 Gb memory per sample. The read alignments to each subgraph are summarized as a vector of the number of reads mapped to each node/*k*-mer. When normalized by global read depth these represent mapping dosage used as input for eQTL mapping of the 100 kb *cis*-windows of genes.

In Lu 2021 (Lu and Chaisson 2021), *cis*-eQTL mapping using an RPGG was reported using the sum of the dosage vector for each locus-RPGG as an estimate of VNTR length. Applying this approach on the 35-genome RPGG and adding an additional bias correction step, we discovered 2,870 VNTRs in association with nearby gene expression (denoted as eVNTR and eGene, respectively; Supplemental Fig. S6, Supplemental Data S3), which is an 8-fold increase over the number (n=346) reported from the previous RPGG built on 32,138 VNTRs (Lu and Chaisson 2021). Of the original eVNTRs, 39.6% (137/346) were reproduced in this version. Among the 209 eVNTRs not reproduced, 70% were removed in the more stringent QC filtering in this version.

In addition to VNTR length, our previous work (Lu and Chaisson 2021) identified motifs enriched in certain populations sequenced by the 1KGP (The 1000 Genomes Project Consortium 2015). We hypothesized that differential motif usage across individuals, possibly independent of overall VNTR length variation, can modulate nearby gene expression. To test this, we converted each locus-RPGG to a compact de Bruijn graph (Chikhi et al. 2016), and considered each path as a locus to test in eQTL mapping. The original RPGG contains on average 400 nodes in each locus-RPGG, which is reduced to 46 paths (referred to as motifs hereafter) after compaction (Supplemental Fig. S7), with a total of 1,810,042 motifs.

To ensure the quality of read-mapping to each motif in each locus-RPGG, we evaluated the “mappability” of each motif by measuring the consistency between the dosage from short reads and the dosage from the ground-truth assemblies using mean absolute percentage error (MAPE, Methods). We removed 48.7% (880,895/1,810,042) of the motifs with MAPE ≥ 0.25 (Supplemental Fig. S8). The number of motifs with zero variance in absolute percentage error (n=313,034) is equivalent to the number paths private to one genome among the 35 HGSVC assemblies (Supplemental Fig. S8) and were retained for subsequent analyses. Similar to eQTL mapping on SNVs where homozygous variants are removed, we avoid testing “invariant” motifs that appear the same number of times in a repeat across all assemblies. This further removes 17.1% (159,212/929,147) of motifs. By construction, our RPGG could contain loci with multiple VNTRs in close proximity but spaced apart by short flanking sequences. We removed additional 0.04% (319/769,935) motifs that are derived from those sequences and could be simply explained by SNVs, leaving a total of 769,616 motifs for eQTL mapping. We evaluated how well we can genotype these motifs following the same procedure for VNTR dosage using batch-*r*^2^. Bias correction moderately improved batch-*r*^2^ on both the HGSVC (0.656 versus 0.701) and the HPRC (0.653 versus 0.697) datasets (Supplemental Fig. 9). For each motif, we consider only GTEx samples of which the dosages are within two standard deviations from the mean, avoiding the discoveries of associations whose effects are mainly driven by outliers. This on average removes 32 from the 813 samples per motif (Supplemental Fig. S10).

We ran *cis*-eQTL mapping for each VNTR motif and discovered 9,422 eVNTRs, including 25,031 motifs associated with nearby gene expression, denoted as eMotifs (Fig. 3A, Supplemental Data S4). While 79.8% (2,291/2,870) of the eVNTRs discovered using length estimates were also reported using motif dosage, 75.7% (7,131/9,422) of the eVNTRs discovered from motif were undetectable with length-based eQTL mapping. We noted a consistent trend of a higher batch-*r*^2^ in eMotifs compared to the genome-wide values, with the mean batch-*r*^2^ being 0.842 and 0.835 in HGSVC and HPRC, respectively (Supplemental Fig. S11). The genomic correction values λ_GC_ of eMotifs (0.953-1.226, depending on the tissue) are consistently lower than the ones from GTEx eQTLs (Supplemental Fig. S12).

**Figure 3.**
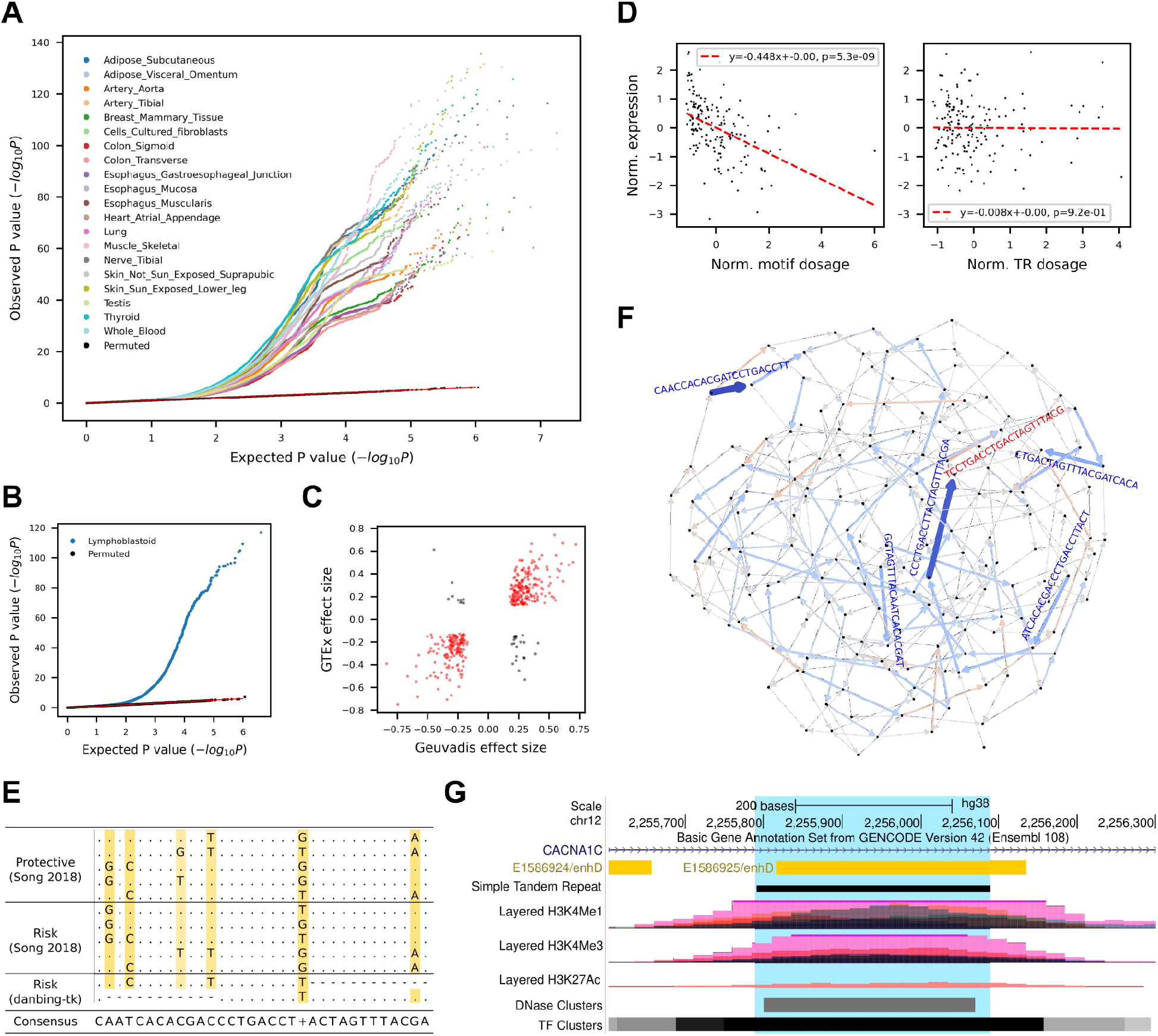
*cis*-eQTL mapping of VNTR motifs. (A) Quantile-quantile plot of gene-level eMotif discoveries across 20 human tissues from GTEx datasets. The expected P-values (x-axis) were drawn from Unif(0,1) and plotted against observed nominal association P-values. (B-C) Replication on Geuvadis dataset. (B) The quantile-quantile plot shows the observed P-value of each association test (two-sided t-test) versus the P-value drawn from the expected uniform distribution. Black dots indicate the permutation results from the top 5% associated (gene, motif) pairs. (C) Correlation of eMotif effect sizes between Geuvadis and GTEx whole blood tissue. Only eGenes significant in both datasets were shown. Each pair of gene and motif that has the same/opposite sign across datasets were colored in red/black. (D-F) Identification of risk motifs for *CACNA1C* expression. The motifs in *CACNA1C* VNTR at chr12:2,255,789-2,256,088 were analyzed. (D) Association of *CACNA1C* VNTR motif CAACCACACGATCCTGACCTT (left) or VNTR length (right) with gene expression in brain cerebellar hemisphere. (E) Multiple sequence alignment of known *CACNA1C* risk motifs (Song et al. 2018) and the likely causal eMotifs reported in this study. (F) Graph visualization of motif effect sizes from the *CACNA1C* VNTR. Each edge denotes a motif and is colored blue/red if having a negative/positive effect on gene expression. Color saturation and edge width both scale with the absolute value of effect size. The sequence of a motif is shown parallel to the edge and colored in dark blue if having a significant effect or colored in light red/blue if borderline significant. (G) UCSC Genome Browser view (Kent et al. 2002) of *CACNA1C* VNTR.

To further assess the reproducibility of our methods, we performed eQTL mapping on the Geuvadis dataset (Lappalainen et al. 2013) (Fig. 3B) and compared the discoveries with the GTEx results (Supplemental Fig. S13). We found that 68.5% (862/1,259) of the eMotifs and 90.0% (1,039/1,154) of the eVNTRs from Geuvadis were also observed in at least one GTEx tissue. Unreplicated eGene-eMotif pairs (ePairs) tended to have lower significance in Geuvadis, especially when P > 10^-6.6^ (odds ratio = 3.35, Fisher’s exact P = 4.0×10^-21^, Supplemental Fig. S14). When comparing the whole blood tissue from GTEx and the lymphoblastoid from Geuvadis, the effect sizes from the two datasets had a correlation coefficient of 0.81 (P=1.5× 10^-94^) (Fig. 3C). Among the replicated ePairs, 92% (363/394) had the same sign of effect (Fig. 3C). When comparing with the EBV-transformed lymphocytes from GTEx, we observed a stronger correlation (Pearson’s *r* = 0.96, P=2.1 × 10^-120^) and 100% of the ePairs (n=208) with the same sign of effect (Supplemental Fig. S15), suggesting motif variations as a common explanatory variable in gene expression.

### Disease relevance of eMotifs

Among the 40 additional disease-relevant tandem repeats (matched with 36 genes) included in the RPGG, 17 of them (Supplemental Table S2) were identified as eQTLs and were associated with their original disease-linked genes, including *ATN1, ATXN7, AVPR1A, C9orf72, CACNA1C, CEL, DRD4, HTT, IL1RN, JPH3, MAOA, MUC1, MUC21, PER3*, and *SLC6A4*. Additionally, at least one eVNTR was detected for 17 of the 21 remaining genes and was different from the originally annotated disease-relevant tandem repeat.

We investigated two examples of motifs associated with disease to see if they had associations with expression in healthy individuals. Landefeld et al. (Landefeld et al. 2018) report that the “RS1” short tandem repeat (STR) at the 5’UTR of *AVPR1A* and the “AVR” STR in the intron are associated with externalizing behaviors while Vollebregt et al. (Vollebregt et al. 2021) report that the “RS3” STR but not RS1 is associated with childhood onset aggression. No association was found between *AVPR1A* expression and the lengths of the three STRs, however we found eMotifs for *AVPR1A* in healthy individuals that correspond to RS1 ((GATA)_5_AATA(GATA)_4_G, b=-0.17, P=1.0×10^-4^, artery tibial), AVR ((GA)_9_A_4_, b=-0.17, P=3.1×10^-4^, fibroblast), and RS3 (C(AG)_10_, b=0.28, P=7.5×10^-5^, artery coronary). The RS3 STR (chr12:63,156,354-63,156,429) has a nested repeat structure with an average size of 701 bp in assemblies. It consists of two slightly divergent copies that each carries a highly repetitive (CT)nTT(CT)n(GT)n core motif at chr12:63,156,354-63,156,394 and chr12:63,156,701-63,156,751 (Supplemental Fig. S16). Other tandem repeat annotation approaches might consider this region as two separate STRs of which the length of the core motif is associated with *AVPR1A* expression.

The decreased expression of *CACNA1C* was known to be a risk factor for schizophrenia and has been reported to be associated with several 30 bp risk motifs at chr12:2,255,789-2,256,090 based on a case-control study (Song et al. 2018). Here, we found that the expression of *CACNA1C* in brain cerebellar hemisphere was associated with a risk eMotif CAACCACACGATCCTGACCTT (denoted as motif 1, b=-0.45, P=5.3×10^-9^, Fig. 3D left) but not the VNTR length (P = 0.92, Fig. 3D right). The eMotif covers five of the six mutation sites (Fig. 3E) and is novel to all of the risk motifs reported previously (Song et al. 2018). In addition, we were able to replicate findings from the case-control study (Song et al. 2018) in healthy populations. The risk eMotif CCCTGACCTTACTAGTTTACGA (b=-0.41, P=6.9×10^-8^, denoted as motif 2) that covers three of the mutation sites (Fig. 3E) matches two of the risk motifs and none of the protective motifs, indicating the prevalence of risk-modulating motifs even among healthy populations. When examining the frequency of the two risk eMotifs in the 35 HGSVC assemblies, 28 haplotypes carry motif 1 while 38 haplotypes carry motif 2 (Supplemental Fig. S17). Most of the individuals carry only few copies of the risk motifs but some could carry up to 191 copies of motif 1 in one haplotype (Supplemental Fig. S18). Annotating the locus-RPGG with the eQTL effect sizes also reveals that motifs with minor risk and protective effects are pervasive within the locus, e.g. CTGACTAGTTTACGATCACACGA (b=-0.23, P=2.8×10^-3^) and CGTAAACTAGTCAGGTCAGGA (b=0.16, P=4.2×10^-2^) (Fig. 3F). The size of this VNTR locus in GRCh38 is underrepresented with only 301 bp compared to an average size of 6,247 bp across assemblies. In addition, immense histone modification, DNase clusters and TF clusters signals can also be found in this locus (Fig. 3G), necessitating future investigations to fully understand the regulation mechanism of *CACNA1C*.

To further narrow down eMotifs that are likely causal to gene expression, we used susieR (Wang et al. 2020) to fine-map the 1Mb *cis*-window of each eGene. We discovered 179 out of 12,894 eGenes of which the highest eVNTR posterior inclusion probability (PIP) is greater than 0.8, suggesting the likely causal roles of these eMotifs and VNTR lengths (Supplemental Fig. S19, Supplemental Data S5). The expression of these 179 eGenes are likely modulated by 163 eVNTRs, through the total length of 3 VNTRs or the overall dosage of 206 eMotifs (Fig. 4A). Among these, 90 VNTRs are associated by motif composition independent of length (Fig. 4A). On average, 56.2% of the eGenes are shared across tissues while 58.7% of the eVNTRs and 38.1% of the eMotifs are observed across multiple tissues (Supplemental Figs. S20-S22). The genotyping accuracy of these likely causal variants are high on the HGSVC dataset (mean batch-*r*^2^ = 0.857) and reproducible in the HPRC dataset (mean batch-*r*^2^ = 0.851, Fig. 4B). While no single regulatory element was found to be enriched in the fine-mapped motif sequences, we found that 54/140 VNTR loci with fine-mapped motifs overlap ENCODE *cis*-regulatory elements, a 2.3-fold increase over what is expected by chance (P < .001, permutation test).

**Figure 4.**
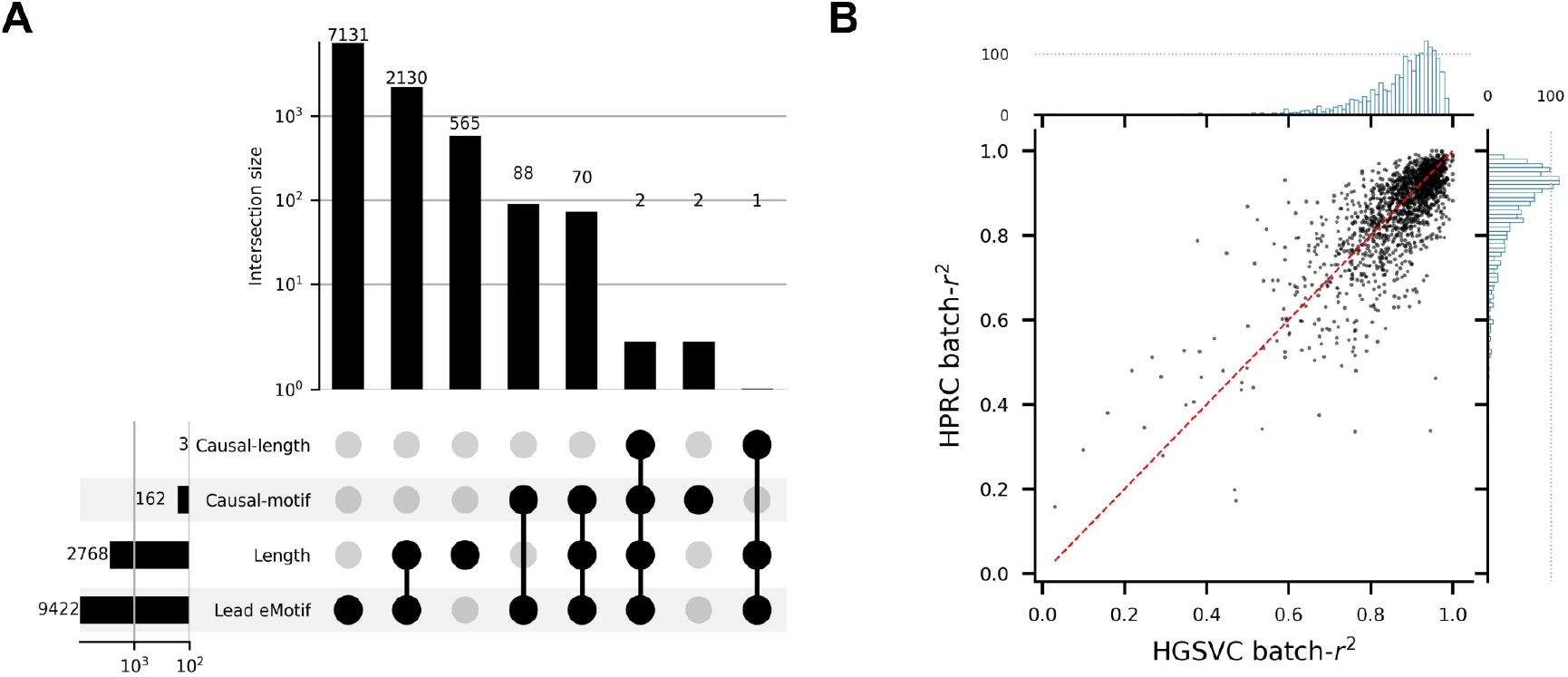
Fine-mapping of VNTR variants. (A) VNTR-centric view of gene-level eQTL discoveries and fine-mapping. Lead eMotif denotes any VNTRs that are associated with gene expression through at least one eMotif under gene-level discoveries. Length denotes any VNTRs that are associated with gene expression through length under gene-level discoveries. Causal-motif denotes any VNTRs of which a motif passes the fine-mapping procedure with a posterior inclusion probability ≥ 0.8 while being a significant eQTL under genome-wide P-value cutoff. Motifs with the lowest P-value for each VNTR-gene pair are included in the fine-mapping model in addition to VNTR length. (B) The batch-*r*^2^ of likely causal motifs on the HGSVC and HPRC datasets. Each bin on the marginal histogram spans 0.01.

We identified a fine-mapped VNTR at chr1:152,213,243-152,221,044, within an exon of *HRNR*, a large polymorphic VNTR (7.8 kbp in GRCh38 versus an average of 13.5 kbp in assemblies; Fig. 5A-B), previously identified as an eQTL using read depth analysis (Garg et al. 2021). The VNTR length as well as 16 other motifs are reported to be likely causal across 24 tissues for the expression of three different genes – *HRNR, FLG-AS1*, and *FLG* (Fig. 5E, Supplemental Fig. S23, Supplemental Data S5), which are responsible for ichthyosis vulgaris (Smith et al. 2006) and atopic dermatitis (Fallon et al. 2009). The repeat overlaps almost the entire exon 3 of *HRNR*, a shared feature for the S100 fused type protein family including *FLG*. While it is expected that repeat expansion of the *HRNR* exon lead to a higher read count, *HRNR* repeat expansions were associated with reduced expression of *FLG-AS1* and *FLG*. Given that one CTCF binding site (ENCODE ID: EH38E1384952) was annotated on GRCh38 and the critical role of CTCF in genome folding (Merkenschlager and Nora 2016), we conjecture that repeat expansion creates more CTCF binding sites, alters nearby chromatin contacts, and perturbs the expression of *FLG-AS1* and *FLG* (Fig. 5C). We found several potential CTCF binding sites matching a 34 bp two-core motif (Soochit et al. 2021) (JASPAR ID: MA1929.1) when scanning each VNTR allele with FIMO (Grant et al. 2011) (Fig. 5D; Supplemental Fig. S24). The number of detected binding sites tightly correlates with the size of the repeat (Supplemental Fig. S25, Pearson’s *r* = 0.996, P = 2.4×10^-160^) and estimates the magnitude of perturbation in local genome folding predicted by the *in-silico* DNA folding software Akita (Fudenberg et al. 2020) (Fig. 5F, Pearson’s *r* = -0.838, P = 8.2×10^-42^; Supplemental Fig. S26). Overall, *HNHR* repeat expansion could perturb nearby gene expression – *FLG-AS1* and *FLG* in this case – with a potential interplay with CTCF binding.

**Figure 5.**
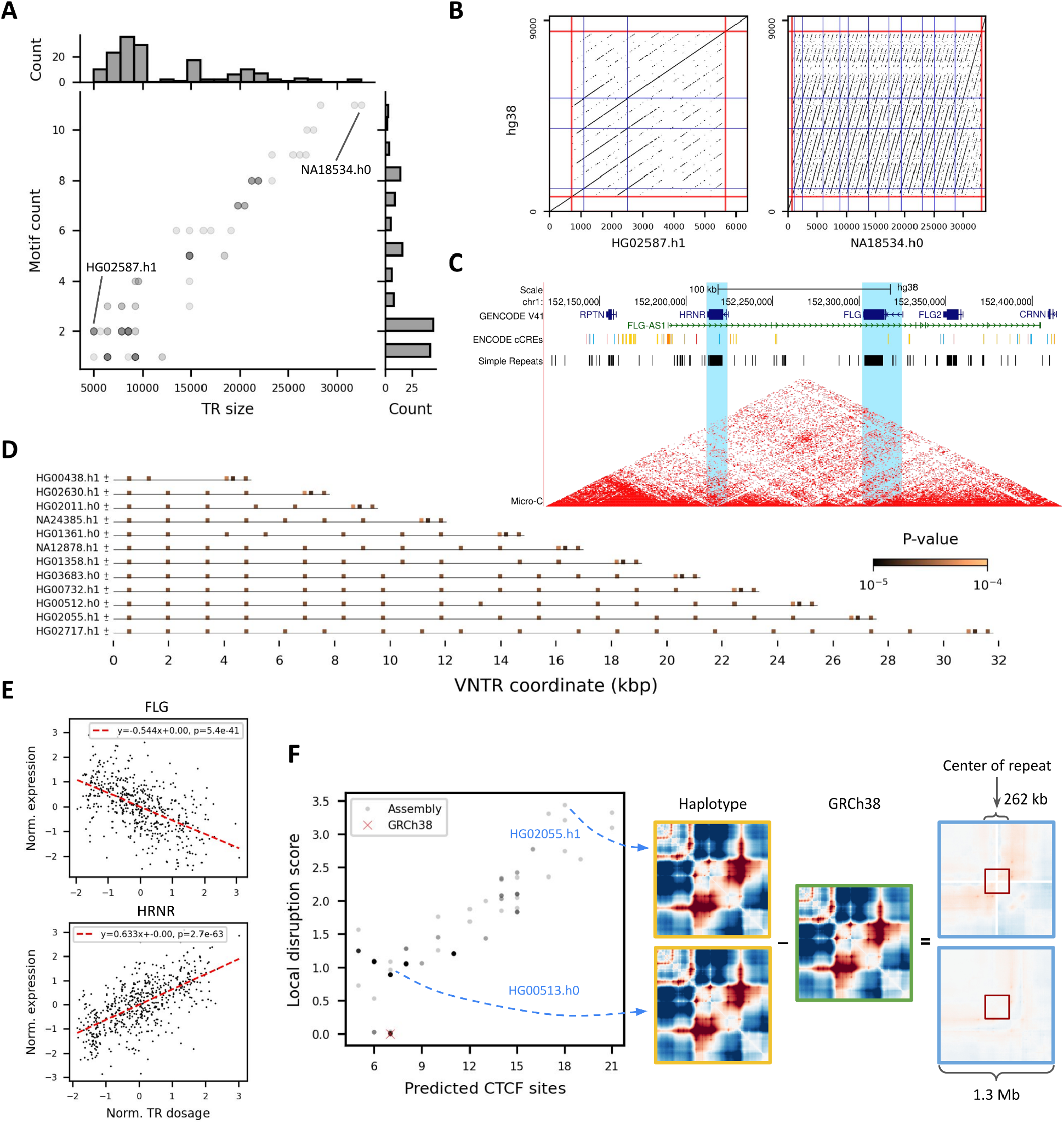
*HRNR* repeat expansion impacts nearby gene expression. (A-B) VNTR allele diversity of the *HRNR* repeat (chr1:152,213,243-152,221,044). (A) The VNTR lengths and counts for motif AGGAGTGCCCCAAACCGGACCCATGTCGGCCG in HGSVC and HPRC assemblies are shown (matplotlib alpha = 0.2). (B) Two divergent haplotypes in (A) are highlighted with dot plots. Red lines indicate the boundary of the repeat, flanked by 700 bp sequences on both sides. The locations of the motif in GRCh38 and assemblies are shown as blue lines. Each dot denotes an exact match of 21-mers. (C) UCSC Genome Browser (Kent et al. 2002) view of *HRNR, FLG*, and *FLG-AS1*. Blue, red, and yellow lines in the ENCODE cCRE track denote CTCF sites, promoters, and enhancers, respectively. Micro-C chromatin structure from HFFc6 cell line was shown. *HRNR* and *FLG* are highlighted in light blue. (D) Predicted CTCF binding sites across 13 length-divergent haplotypes. Each haplotype was scanned for matches with a 34 bp, two-core CTCF motif (MA1929.1) using FIMO with a cutoff of P < 10^-4^. (E) Association of the estimated *HRNR* repeat size in GTEx genomes with *FLG* (in fibroblast) and *HRNR* (in thyroid) expression. Red dashed lines indicate the best fit under simple linear regression. (F) (*left*) The number of predicted CTCF sites versus disruption of local genome folding predicted by Akita for alternate VNTRs among 83 assemblies. Each variant VNTR is shown as a gray dot with a shade reflecting the multiplicity of alleles that have the similar numbers of CTCF sites and disruption scores. A higher local disruption score reflects greater changes in contact frequencies relative to GRCh38 in a 262 kb window. (*right*) Illustration how local disruption score is calculated comparing predicted folding in a haplotype to GRCh38.

While likely causal motifs in some cases are strongly correlated with VNTR length such as the above example, they can also be very distinct from length such as the ones in chr17:80260506-80260846, a VNTR immediate upstream of *RNF213* (Fig. 6A, Supplemental Figs. 27-28). The VNTR adjoins the promoter and a proximal enhancer (ENCODE ID: EH38E2146500) of *RNF213*, and overlaps abundant histone marks, DNaseI hypersensitivity, and TF clusters (Supplemental Fig. S29). A search for TF binding motif matching the likely causal motif GCGGGGCCGGCGGCGGCGGCGG using TOMTOM (Gupta et al. 2007) indicates a strong match with ZNF93 (JASPAR ID: MA1721.1, P = 9.0×10^-7^, Supplemental Fig. S30). Scanning each haplotype sequence using FIMO suggests that there are up to 16 ZNF93 binding sites in the repeat (Supplemental Fig. S31). Intriguingly, repeat expansion does not create new binding sites in all cases since the motif count better correlates with the number of TF binding sites than the repeat length (Supplemental Fig. S31). In some haplotypes, repeats expand through alternative motifs (Fig. 6B-C) that have a poorer match with the ZNF93 motif (Supplemental Fig. S32) and minor effects on gene expression (Supplemental Fig. S33). In this case, the likely causal VNTR variant for gene expression is motif and not length.

**Figure 6.**
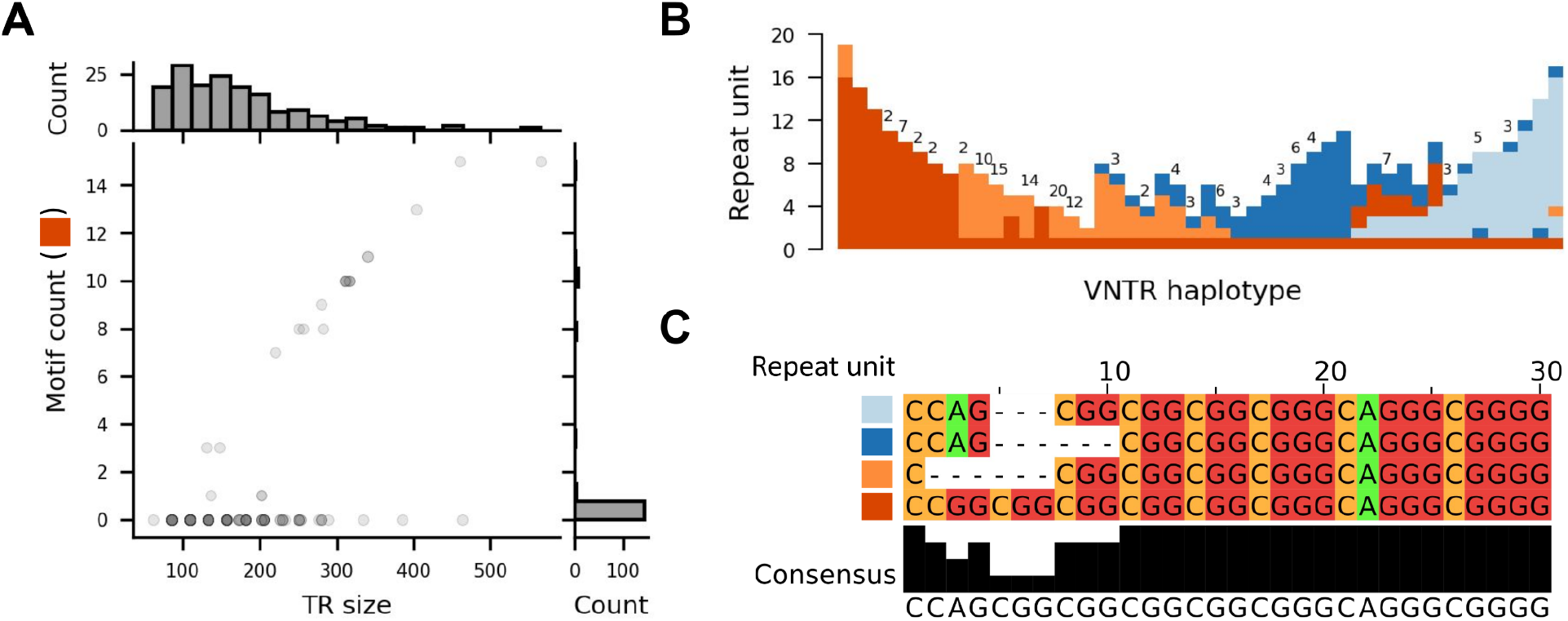
Motif composition but not length explains *RNF213* expression changes. (A) VNTR allele diversity of the *RNF213* promoter repeat (chr17:80260506-80260846). The VNTR lengths and counts for the likely causal motif GCGGGGCCGGCGGCGGCGGCGG, corresponding to the red repeat unit in (B), in HGSVC and HPRC assemblies are shown (matplotlib alpha = 0.2). (B) Repeat expansion of the *RNF213* VNTR from diverse repeat units. VNTR haplotypes from all assemblies were annotated using vamos (Ren et al. 2022). Different repeat units are illustrated by different colors; the repeat unit containing a ZNF93 binding site is colored dark orange. Only unique annotations were shown, with the frequency of each entry shown on top if greater than one. (C) Multiple sequence alignment of the four repeat units shown in (B). Alignments were generated using MUSCLE (Edgar 2004) and visualized with Jalview (Waterhouse et al. 2009).

## Discussion

Genomic variant discovery serves to link genetic and phenotypic variation. Using gene expression as a phenotypic measure, diverse classes of variation have been found to have an effect on gene expression including single-nucleotide variants (GTEx Consortium 2020), structural variation (Chiang et al. 2017; Ebert et al. 2021; Sudmant et al. 2015), STRs (Gymrek et al. 2016), and VNTRs (Bakhtiari et al. 2021; Eslami Rasekh et al. 2021; Garg et al. 2021). Here we show that in addition to association of VNTR length with expression, a more nuanced measurement of VNTR variation that takes into account sequence composition reveals eMotifs that influence gene expression.

Overall, we find 20,834 VNTR loci containing at least one eMotif. In contrast, previous studies that used associations based on length estimate alone ranged between 163-2,980 eVNTRs (Bakhtiari et al. 2021; Lu and Chaisson 2021; Garg et al. 2021), with the number roughly correlating with the number of loci each study profiled. Although more tests per VNTR locus are performed, the fine-mapping analysis finds that the majority of variants are linked with nearby eQTLs. After applying fine-mapping, 229 (1.1%) eVNTRs contain motifs determined as likely causal. In contrast, 0.18% of the 4.3M eQTL variants discovered in the GTEx (v8) are fine-mapped (GTEx Consortium 2020).

We observe that most eVNTRs have different motifs positively and negatively associated with expression of the same nearby gene. Most eQTL mapping pipelines are based on biallelic variants. When encoding a variant, the reference allele is usually treated as zero while the alternative allele is treated as one. Alternatively, this can be viewed as encoding the alternative allele as its copy number, which is simply one for a biallelic variant, while keeping the reference allele as zero. When the same encoding method is applied to a VNTR locus consisting of a reference motif and an alternative motif, the only difference is that the alternative allele becomes a continuous value representing the adjusted motif depth, and may take on a positive or negative association depending on the relation to the reference motif.

This study profiles 39,125 VNTR loci, a 1.2-fold increase over our previous analysis of VNTR variation using repeat-pangenome graphs (Lu and Chaisson 2021) that is largely attributable to the high-quality haplotype-resolved assemblies used to construct the pangenome. The size of the graph increases sequentially with the number of assemblies included in the graph, and is consistent with the increasing number of structural variants discovered in VNTRs by whole-genome alignment (Ebert et al. 2021). The inclusion of additional genomes from large-scale sequencing projects such as the Human Pangenome Reference Consortium will yield an improved estimate of saturation of VNTR variation.

The use of repeat-pangenome graphs in this study differs from other implementations of pangenom-graphs including those constructed by progressive whole-genome alignment (Li et al. 2020) and variant inclusion (Sirén et al. 2021), both of which preserve haplotype information from the genomes or variants used to construct the pangenome graph. While systematic analysis of variation in VNTRs and association with expression has not yet been conducted using these approaches, we anticipate the repeat-pangenome graph will provide complementary analysis. In particular, variant genotyping in graphs that preserve haplotype as implemented by Giraffe (Sirén et al. 2021) and PanGenie (Ebler et al. 2022) corresponds to associating read data with haplotypes (paths in a pangenome graph) covering variants. These approaches provide highly accurate genotyping of variants shared with the graph, however hypervariable VNTR sequences are more likely to have differences from genomes represented in the graph, and additional analysis is required to quantify motif usage in addition to genotype.

The implementation of the pangenome as a de Bruijn graph is an elegant approach to identifying the composition of identical motif repeats, however small differences in motif composition can make the graph complex, and additional development is necessary to identify graph topologies that naturally reflect VNTR repeat composition. One result of this complexity is our number of eMotifs that are deemed likely causal using fine-mapping is possibly an underestimate. Many motifs have highly correlated read dosage, however we use a conservative approach of considering each motif as an independent variable for fine-mapping. Future development that merges similar motifs to the same edge both aggregate depth otherwise split on several edges, and reduce the correlated motifs tested during fine-mapping.

In summary, this study demonstrates how VNTR composition has a pervasive influence on gene expression, and highlights the need to profile variation in complex, repetitive regions of the genome. We anticipate this approach will be useful for future expression and association studies.

## Methods

### Data retrieval

HGSVC haplotype-resolved assemblies (n=35; HG00096, HG00171, HG00512, HG00513, HG00514, HG00731, HG00732, HG00733, HG00864, HG01114, HG01505, HG01596, HG02011, HG02492, HG02587, HG02818, HG03009, HG03065, HG03125, HG03371, HG03486, HG03683, HG03732, NA12329, NA12878, NA18534, NA18939, NA19238, NA19239, NA19240, NA19650, NA19983, NA20509, NA20847, NA24385) were downloaded from (https://www.internationalgenome.org/data-portal/data-collection/hgsvc2) (Ebert et al. 2021). HGSVC whole-genome sequencing (WGS) samples are downloaded from 1000 Genomes Project phase 3 (https://www.internationalgenome.org/data-portal/data-collection/hgsvc2) (The 1000 Genomes Project Consortium 2015).

HPRC freeze 1 (v2) haplotype-resolved assemblies (n=47) (Liao et al. 2022) were downloaded from https://zenodo.org/record/5826274#.ZAAaa9LMKEI. CHM13 telomere-to-telomere assembly (v1.0) (Nurk et al. 2022) was retrieved from GenBank accession GCA_009914755.2. WGS samples (n=36) matching HPRC genomes were downloaded from https://www.internationalgenome.org/data-portal/data-collection/30x-grch38 (study accession PRJEB36890) (Byrska-Bishop et al. 2022).

Maternal and paternal haplotypes were referred to as h0 and h1 in the text for brevity.

WGS samples (n=879), gene expression matrices of 49 tissues, and covariates for eQTL mapping of GTEx genomes were retrieved from GTEx Analysis Release V8 (dbGaP accession phs000424.v8.p2) (GTEx Consortium 2020). WGS samples for Geuvadis genomes (n=445) were downloaded from 1000 Genomes Project phase 3 (https://www.internationalgenome.org/data-portal/data-collection/hgsvc2), and the matching residualized gene expression matrix for lymphoblastoid cell lines (GD462.GeneQuantRPKM.50FN.samplename.resk10.txt.gz) were retrieved directly from the Geuvadis portal https://www.ebi.ac.uk/biostudies/arrayexpress/studies/E-GEUV-1 (Lappalainen et al. 2013).

### Repeat pangenome graph construction

A set of 88,441 VNTR coordinates were retrieved from danbing-tk v1.3 (Lu and Chaisson 2021). The VNTR set was obtained by (1) detecting VNTRs over the five haplotype-resolved assemblies (AK1, HG00514, HG00733, NA19240, NA24385) released by Lu and Chaisson (Lu and Chaisson 2021) using Tandem Repeat Finder v4.09.1 (Benson 1999), (2) selecting for VNTRs with size between 100 bp and 10 kbp and motif size > 6 bp, and (3) applying danbing-tk to the VNTRs in the five genomes to identify 88,441 loci with proper orthology mapping. VNTR annotations on the 35 HGSVC assemblies and the corresponding RPGG were generated using the build module of danbing-tk, giving a total of 80,518 loci.

### VNTR genotyping

WGS samples from GTEx (GTEx Consortium 2020) and Geuvadis (Lappalainen et al. 2013) with a total of 879 and 445 samples respectively. Both cohorts are primarily composed of individuals from European/European-American populations: 85.3% (GTEx), and 83% (Geuvadis). The remaining individuals in GTEx are African American (12.5%), Asian American (1.4%), and Hispanic or Latino (1.9%), and West African (Yoruban) in the Geuvadis cohort. VNTRs were genotyped using danbing-tk v1.3 with options “-ae -kf 4 1 -gc 85 -k 21 -cth 45”. While each locus-RPGG encodes the population diversity of a VNTR along with the 700 bp flanking sequences, which is crucial for read alignment, only the *k*-mer counts in repeat regions are output by danbing-tk. The output *k*-mer counts were adjusted by the coverage of each sample before subsequent analyses.

### Graph compaction and motif dosage computation

Locus-RPGG built from *k=21* contains abundant contiguous paths without branches. It is desirable to reduce the number of nodes to be tested in eQTL mapping by merging nodes on this type of paths. This is essentially a problem of converting dBGs to compact dBGs (Chikhi et al. 2016) where nodes on a non-branching path are merged into a unitig, or referred to as a motif in this context. For each motif, we recorded the mapping relation from its constituent nodes and computed the motif dosage by averaging the *k*-mer counts from constituent nodes. In practice, when given a matrix of VNTR genotype where each column represents a *k*-mer in a locus-RPGG and each row represents a sample, the matrix of motif dosages can be simply computed by column operations using the mapping relations.

### Quality control of VNTRs

To ensure the VNTR set in the database are mainly composed of simple repeats instead of tandemly duplicated mobile elements, and can be genotyped with high accuracy, we applied the following four filters: (1) mobile element mask, (2) aln-*r*^2^ > 0.5, (3) assembly mismap rate < 5%, (4) GRCh38 mismap rate < 1%, and (5) length variable on the HGSVC assemblies.

All of the genomes (AK1, HG00514, HG00733, NA19240, NA24385) used for calling VNTRs were processed with RepeatMasker v4.1.2-p1 (http://repeatmasker.org). VNTR annotations that overlapped with any of the mobile elements detected by repeatmasker were masked. This removed 40,199 out of the 80,518 loci.

The aln-*r*^2^ statistic was used to evaluate how well a VNTR can be genotyped within a sample. It is the *r*^2^ computed by regressing the *k*-mer counts from assemblies against the counts from reads aligned to the locus-RPGG. Since VNTRs were genotyped using the RPGG, any read *k*-mers not present in the original assembly were ignored.

A VNTR within segmental duplications (segdups) has the chance to be genotyped with high aln-*r*^2^ but contain extensive mismapped reads from other genomic regions. A perfectly duplicated VNTR creates the following scenario. All reads from the two loci will be deterministically mapped to one locus. The locus without any reads will be dropped since aln-*r*^2^ is not a value. However, the other locus will have the same aln-*r*^2^, and the only difference is that the slope doubled in the linear fit. To avoid genotyping these loci, we ran simulations to detect loci with mismapping from or to other genomic regions. Error-free 150 bp paired-end reads (fragment size = 500 bp) were simulated from 35 HGSVC diploid assemblies. Contigs < 50kb were ignored since a short contig can contain a partially assembled VNTR sequence that is fully assembled in another longer contig. Reads were mapped to the RPGG using the same options (-ae -kf 4 1 -gc 85 -k 21 -cth 45). The loci of aligned reads were extracted from danbing-tk’s alignment output and were compared to the origin of the reads, restricting to reads with VNTR *k*-mers only. The number of reads from other loci or untracked regions aligned to this locus is denoted as *N_G_*. The number of reads from this locus aligned to other loci or untracked regions is denoted as *N_L_*. The number of VNTR reads from a locus is *N_0_*. Mismap rate is computed as (*N_G_*+*N_L_*)/*N_0_*. The average mismap rate from all assemblies was used as the final metric for filtering.

To account for loci missing from HGSVC assemblies but present on GRCh38, we ran another simulation on GRCh38 alone. The overall procedure is similar except that reads were simulated from GRCh38 without alt contigs. Read mapping was performed on the graph built from GRCh38 alone. VNTR boundaries on GRCh38 were adjusted such that the *k*-mer set was consistent with the annotations from other assemblies. This expanded 34,637 out of the 80,518 loci and dropped four loci with unresolved inconsistency.

Length-invariant VNTRs based on the observation from the 35 HGSVC assemblies were removed from eQTL mapping by estimated VNTR length to avoid associating gene expression with length variation predominantly driven by noise in sequencing read depth and length estimation.

All loci passing the mobile element mask were retained (n=40,314) for genotyping so that loci with higher mismap rates can act as “baits” for reads from problematic regions. Only the results for loci passing all filters (n=39,125) were used for eMotif mapping and fine-mapping.

### Quality control of motifs

To ensure the quality of motifs tested in eQTL mapping, we applied three filters to remove motifs (1) with mean absolute percentage error (MAPE) > 0.25, (2) with dosage invariant across HGSVC haplotypes, or (3) that were derived from an inter-VNTR region (denoted as “flank-like”) but were included in the repeat region of a locus-RPGG due to the distance between the upstream and downstream VNTRs being < 700 bp. For the first filter, we computed mean absolute percentage error (APE) for each motif by measuring the error size of each motif to the linear fit for aln-*r*^2^. Formally, let **x** = (*x*_1_, *x*_2_, …, *x_P_*) be the motif dosages from assemblies and **y** = (*y*_1_, *y*_2_, …, *y_P_*) be the motif dosages from short reads, where *P* is the number of motifs in the locus-RPGG. For each genome *g*, 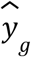 is the fitted value for the dosage of a motif from the linear fit between **x** and **y**. The MAPE of a motif can be computed as follows:

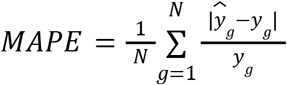

where *N* is the number of genomes with the motif. For the second filter, a motif was removed if the dosage of a motif was the same across all 70 haplotypes. The dosage was set to zero if the motif was not present in a haplotype. For the third filter, any *k*-mers derived from the inter-VNTR regions before the VNTR merging step were extracted. Any motifs overlapping with these *k*-mers were removed.

### Bias correction and batch-r^2^

While aln-*r*^2^ indicates that most loci can be accurately genotyped in terms of the relative *k*-mer compositions in a locus when examining each individual separately. We noticed that loci with high aln-*r*^2^ and not overlapping segdup regions can have huge variations in the slopes of the linear fits when comparing across individuals due to the stochastic nature of read coverage or unknown technical biases. This imposes a challenge for population-scale analysis such as eQTL mapping. To correct for this, we searched for invariant *k*-mers in each locus that appear the same number of times across all HGSVC VNTR haplotypes. For each locus, raw VNTR genotypes (motif counts, or sum of *k*-mer counts for length) were adjusted by coverage and then by a bias term such that the final *k*-mer coverage of these invariant *k*-mers are the same across individuals. More formally, let ***k*** be a vector of invariant *k*-mers, and let ***c*** ∈ be the counts associated with these *k*-mers in the diploid assembly. If short reads sampled from the whole genome were perfectly uniform such that the only factor changing the observed *k*-mer counts ***c’*** ∈ ℝ^N^ were read coverage, or *d* ∈ ℝ^1^. In this case, *k*-mer coverage, or *c_i_*‘/*c_i_*, is *d* for all *k*-mers. Now we extend this procedure to a stochastic sampling process. For an individual *j* and read coverage *d_j_*, let 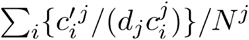 be the bias term *b_j_* for this VNTR locus, reflecting the magnitude of a sample-specific deviation of *k*-mer coverage from *d_j_*. Let ***b*** ∈ ℝ^M^ be the bias vector for *M* individuals, and **T** ∈ ℝ^*N*’×*M*^ be the *k*-mer count matrix collated from danbing-tk’s outputs, where each row represents a *k*-mer from a locus with *N*’ *k*-mers, and each column represents an individual from *M* samples. The final *k*-mer dosage table, or **T**’, can be computed by dividing each column in **T** with *d_j_b_j_*, or 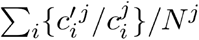 removing the read depth and bias factors at the same time as an estimate for diploid *k*-mer copy number. Corrected motif dosage can be computed by averaging the values from *k*-mers corresponding to the motif, as described in the “Graph compaction and motif dosage computation” section.

To measure the performance of this correction, we define batch-*r*^2^ for length estimation as the *r*^2^ between assembly VNTR size and estimated VNTR dosage (the sum of each column in **T’**) across a batch of samples; the batch-*r*^2^ for motif dosage estimation is defined as as the *r*^2^ between assembly motif count and corrected motif dosage.

### Validation of estimated VNTR and motif dosage

Using the batch-*r*^2^ defined in the previous section, we evaluated the accuracy of our genotyping approach using the assemblies from HGSVC and HPRC. VNTR annotations on the 48 HPRC assemblies including CHM13 were generated using the build module of danbing-tk, giving a total of 81,261 loci. Only loci with one-to-one mapping between HGSVC and HPRC annotations were retained when comparing between the two sets, leaving a total of 79,626 loci. For VNTRs with invariable length and motifs with invariable count in the assemblies of each dataset, batch-*r*^2^ is not computed. If a VNTR locus does not have any invariant *k*-mers in HGSVC for bias correction, the batch-*r*^2^ is also not computed. Motifs with at least one overlapping *k*-mer between the two datasets were retained for analysis.

### eQTL mapping

Gene expression data was processed as previously described (Lu and Chaisson 2021) unless stated otherwise. Briefly, fully processed, filtered and normalized gene expression matrices and covariates were downloaded from GTEx portal as described in the “Data retrieval” section. Confounding factors were removed using covariates including sex, sequencing platform, amplification method, PEER factors, and top 10 principal components (PCs) from the joint single-nucleotide polymorphism matrix with 1KGP samples. Residualization of the gene expression matrices was done with the following formula: where Y is the residualized expression matrix; Y’ is the normalized expression matrix; I is the identity matrix; C is the covariate matrix where each column corresponds to a covariate mentioned above. Fully processed, filtered, and residualized expression matrices of Geuvadis samples were obtained from the Geuvadis portal as described in the “Data retrieval” section.

For eQTL mapping using motif dosage, samples with motif dosage being two standard deviations away from the mean were removed for each motif. For eQTL mapping using VNTR length, samples with VNTR length being three standard deviations away from the mean were removed for each VNTR. The motif dosages, VNTR lengths, and the residualized expression counts for the remaining samples were z-score normalized before testing for association. VNTRs that are less than 100 kb from a gene body were tested for association.

The nominal P-values were obtained from *t*-test on the slope of each linear model consisting of expression (response variable) versus motif dosage or estimated VNTR length (explanatory variables) using Python statsmodels v0.13.2 (Seabold and Perktold 2010). For gene-level *cis*-eQTL discoveries, the P-value of each test was adjusted according to the number of variants tested for each gene using Bonferroni correction. The minimal P-values from all the tests against each gene were extracted and controlled at 5% false discovery rate using the Benjamini–Hochberg procedure. Only one eMotif (if using motif dosage) or one eVNTR (if using VNTR length) was reported for each eGene. For *cis*-eQTL discoveries using a genome-wide P-value cutoff, all P-values from all tests were recorded and controlled at 5% false discovery rate using the Benjamini–Hochberg procedure. The P-value cutoffs range from 2.9 × 10^-5^ (Kidney) to 1.7 × 10^-3^ (Thyroid), depending on the power in each tissue (Supplemental Table S3). There can be more than one eMotif to be reported as significantly associated with a gene when reporting discoveries with this approach, which is required for analyses that investigate all motifs in a VNTR locus. A VNTR that contains an eMotif was also regarded as an eVNTR. Consequently, an eVNTR can be associated with gene expression through length or motif dosage depending on the type of tests performed.

### Comparing λ_GC_ across datasets and variant types

λ_GC_ for GTEx variants were computed from the P-values of all tests. The full tables ($tissue.allpairs.txt.gz) containing all P-values were downloaded from the GTEx portal. λ_GC_ was computed using using the scipy.stats v1.9.1 (Virtanen et al. 2020) and Numpy (Harris et al. 2020) module in Python as follows:

1. med = stats.chi2.ppf(0.5, 1) # expected median of chi-squared distribution
2. chi2 = np.median(stats.chi2.ppf(1-ps, 1)) # ps are the P-values from GTEx variants
3. lambda_gc = chi2/med

The λ_GC_ for VNTR motifs in each GTEx tissue or in Geuvadis is computed in the same way as above, taking the P-values from our eMotif mapping. However, to make fair comparisons, we reran eQTL mapping on GTEx whole blood and Geuvadis with a 100 kb *cis*-window using QTLtools v1.3.1 (Delaneau et al. 2017) to be consistent with the setting in our model.

### Fine-mapping

To evaluate whether the eMotifs are causal to gene expression, we used susieR v0.11.92 (Wang et al. 2020) to fine-map the *cis*-window around the transcription start site of each gene. All variants in GTEx’s catalog (GTEx_Analysis_2017-06-05_v8_WholeGenomeSeq_838Indiv_Analysis_Freeze.vcf.gz), including SNVs, indels or structural variants, were extracted if within the 1 Mb *cis*-windows of transcription start sites. For each tissue, all motifs that have the lowest P-value for each gene-VNTR pair were extracted. The extracted GTEx variants and motifs were taken as input for fine-mapping. Susie was run using L=5 to allow up to five sets of causal variants within the whole region. Motifs with posterior inclusion probability (PIP) ≥ 0.8 while passing the genome-wide P-value cutoff as described in the previous section were reported as likely causal eMotifs.

Enrichment analysis was performed using 1000 permutations of ENCODE *cis*-regulatory element regions defined by the encodeCcreCombined (CRE) UCSC Genome Browser track on GRCh38 (Rosenbloom et al. 2013). The ENCODE CRE elements were randomly shuffled excluding centromere sequences and counted for overlap with fine-mapped VNTR sequences.

### Genome-folding disruption prediction

Genome-folding predictions were made using Akita (Fudenberg et al. 2020), a deep convolutional neural network model that takes one-hot encoded DNA sequence as input and makes predictions for pairwise observed/expected contact frequency maps with 2048bp bins. The trained model is available on github (https://github.com/calico/basenji/tree/master/manuscripts/akita/v2). Predictions were made using tensorflow version 2.6, and model weights were loaded into Basenji (Kelley 2020) (commit d61389dc553aa610544503a3e937c1b53906fe35). The predictions here used the human head (suffix 0) and were averaged across all human targets and models from each train/test/validation split. To obtain the reference prediction, the input sequence was centered on HRNR chr1:152,213,243-152,221,044 (in GRCh38 coordinates). The DNA input sequences for alternate haplotype (35 HGSVC + 47 HPRC assemblies + CHM13) predictions were created by replacing the reference VNTR at this locus with sequence for each haplotype and trimming the resulting sequence from the right to match Akita’s fixed input length. To quantify the difference in local contact frequencies for each haplotype, we subtracted the predicted reference map from the predicted map for each haplotype. Since VNTR haplotype lengths range from 4,984bp to 32,507bp, predicted maps can be shifted by multiple bins as compared with the reference. Because of this, we took a conservative approach to quantification that focused on differences outside of any inserted (or deleted) bins relative to the reference. We achieved this by aligning predicted maps as follows. For alternate haplotypes with insertions relative to the reference, we first replaced the corresponding bins in the predicted map with NaNs. We then inserted an equal number of bins with NaNs into the reference map. For deletions relative to the reference, the corresponding bins in the reference are replaced with NaNs and an equal number of bins with NaNs are inserted in the predicted alternate maps.

We then calculated a local disruption score (LDS) in a 128-bin (262,144 bp) region centered around the center of the aligned maps, i.e.

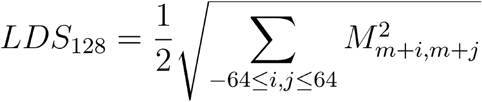

where *M* is the predicted map, *m* is its midpoint, and *i, j* are integers. If the window overlaps bins containing NaN values as a result of map alignment, the window is expanded by a corresponding number of bins on each side of the center of the map.

### Annotating RNF213 repeat with vamos

The repeat sequences for all HGSVC (n=35) and HPRC (n=48, including CHM13) assemblies without flanking sequences were extracted based on danbing-tk’s annotation and saved in fasta format. The efficient motif set was retrieved from https://zenodo.org/record/7155329/files/vntrs_motifs_delta_0.2.bed?download=1. The bed entry corresponding to the *RNF213* repeat was retained. For each fasta entry, vamos v1.1.1 (Ren et al. 2022) was run using the command “vamos --single_seq -b $fasta -r $bed -o $out.vcf -s RNF213”. The annotation following the ALTANNO_H1 tag was extracted.

### Software availability

The source code of danbing-tk v1.3.1-manuscript associated with this study is available on GitHub (https://github.com/ChaissonLab/danbing-tk), on Zenodo (https://zenodo.org/record/7697439#.ZATe99LMKEI), and in Supplemental Code. The bias correction, eQTL mapping, and fine-mapping scripts are available on GitHub (https://github.com/ChaissonLab/eMotif_manuscript_analysis_scripts), on Zenodo (https://sandbox.zenodo.org/record/1169833#.ZATf99LMLmF), and in Supplemental Code.

## Data access

The VNTR annotations from assemblies, RPGG, bias matrices for each datasets, eQTL tables, fine-mapping table, VNTR dosage matrices for GTEx and Geuvadis, danbing-tk source code, and analysis scripts associated with this study are available at https://sandbox.zenodo.org/record/1169833#.ZATf99LMLmF.

## Competing interest statement

The authors declare no competing interests.

## Acknowledgement

This work has been funded by NIH grants U24HG007497, R01HG012133, R01GM140287, R01HG011649. GF and PS are supported by NIGMS R35 GR1057075.

1000 Genomes Acknowledgement for deep coverage of the extended 3202 genomes (or subset thereof): The following cell lines/DNA samples were obtained from the NIGMS Human Genetic Cell Repository at the Coriell Institute for Medical Research: [NA06984, NA06985, NA06986, NA06989, NA06991, NA06993, NA06994, NA06995, NA06997, NA07000, NA07014, NA07019, NA07022, NA07029, NA07031, NA07034, NA07037, NA07045, NA07048, NA07051, NA07055, NA07056, NA07340, NA07345, NA07346, NA07347, NA07348, NA07349, NA07357, NA07435, NA10830, NA10831, NA10835, NA10836, NA10837, NA10838, NA10839, NA10840, NA10842, NA10843, NA10845, NA10846, NA10847, NA10850, NA10851, NA10852, NA10853, NA10854, NA10855, NA10856, NA10857, NA10859, NA10860, NA10861, NA10863, NA10864, NA10865, NA11829, NA11830, NA11831, NA11832, NA11839, NA11840, NA11843, NA11881, NA11882, NA11891, NA11892, NA11893, NA11894, NA11917, NA11918, NA11919, NA11920, NA11930, NA11931, NA11932, NA11933, NA11992, NA11993, NA11994, NA11995, NA12003, NA12004, NA12005, NA12006, NA12043, NA12044, NA12045, NA12046, NA12056, NA12057, NA12058, NA12144, NA12145, NA12146, NA12154, NA12155, NA12156, NA12234, NA12239, NA12248, NA12249, NA12264, NA12272, NA12273, NA12274, NA12275, NA12282, NA12283, NA12286, NA12287, NA12329, NA12335, NA12336, NA12340, NA12341, NA12342, NA12343, NA12344, NA12347, NA12348, NA12375, NA12376, NA12383, NA12386, NA12399, NA12400, NA12413, NA12414, NA12485, NA12489, NA12546, NA12707, NA12708, NA12716, NA12717, NA12718, NA12739, NA12740, NA12748, NA12749, NA12750, NA12751, NA12752, NA12753, NA12760, NA12761, NA12762, NA12763, NA12766, NA12767, NA12775, NA12776, NA12777, NA12778, NA12801, NA12802, NA12812, NA12813, NA12814, NA12815, NA12817, NA12818, NA12827, NA12828, NA12829, NA12830, NA12832, NA12842, NA12843, NA12864, NA12865, NA12872, NA12873, NA12874, NA12875, NA12877, NA12878, NA12889, NA12890, NA12891, NA12892]. These data were generated at the New York Genome Center with funds provided by NHGRI Grants 3UM1HG008901-03S1 and 3UM1HG008901-04S2.

## Author contributions

T.Y.L. and M.J.C. designed the study. T.Y.L. improved danbing-tk and performed association study. T.Y.L. and M.J.C. analyzed association results. P.N.S. and G.F. performed genome folding analysis. N.M. assisted in association study. T.Y.L., P.N.S., G.F., and M.J.C. wrote the manuscript.

## Notes

https://sandbox.zenodo.org/record/1036159#.YjLl_3XMKEB

### Competing Interest Statement

The authors have declared no competing interest.

### Summary of Updates

This is updated according to reviewer comments: - VNTR Loci overlapping Alu elements are removed. - A read normalization step is used to account for sequencing bias. - Additional examples of eQTLs were added.

## References

Amemiya HM, Kundaje A, Boyle AP. 2019. The ENCODE Blacklist: Identification of Problematic Regions of the Genome. Sci Rep 9: 9354.

Bakhtiari M, Park J, Ding Y-C, Shleizer-Burko S, Neuhausen SL, Halldórsson BV, Stefánsson K, Gymrek M, Bafna V. 2021. Variable number tandem repeats mediate the expression of proximal genes. Nat Commun 12: 2075.

Benson G. 1999. Tandem repeats finder: a program to analyze DNA sequences. Nucleic Acids Res 27: 573–580.

Beyter D, Ingimundardottir H, Oddsson A, Eggertsson HP, Bjornsson E, Jonsson H, Atlason BA, Kristmundsdottir S, Mehringer S, Hardarson MT, et al. 2021. Long-read sequencing of 3,622 Icelanders provides insight into the role of structural variants in human diseases and other traits. Nat Genet 53: 779–786.

Byrska-Bishop M, Evani US, Zhao X, Basile AO, Abel HJ, Regier AA, Corvelo A, Clarke WE, Musunuri R, Nagulapalli K, et al. 2022. High-coverage whole-genome sequencing of the expanded 1000 Genomes Project cohort including 602 trios. Cell 185: 3426–3440.e19.

Cameron DL, Schröder J, Penington JS, Do H, Molania R, Dobrovic A, Speed TP, Papenfuss AT.2017. GRIDSS: sensitive and specific genomic rearrangement detection using positional de Bruijn graph assembly. Genome Res 27: 2050–2060.

Chen S, Krusche P, Dolzhenko E, Sherman RM, Petrovski R, Schlesinger F, Kirsche M, Bentley DR, Schatz MC, Sedlazeck FJ, et al. 2019. Paragraph: a graph-based structural variant genotyper for short-read sequence data. Genome Biol 20: 291.

Chiang C, Scott AJ, Davis JR, Tsang EK, Li X, Kim Y, Hadzic T, Damani FN, Ganel L, GTEx Consortium, et al. 2017. The impact of structural variation on human gene expression. Nat Genet 49: 692–699.

Chikhi R, Limasset A, Medvedev P.2016. Compacting de Bruijn graphs from sequencing data quickly and in low memory. Bioinformatics 32: i201–i208.

DeJesus-Hernandez M, Mackenzie IR, Boeve BF, Boxer AL, Baker M, Rutherford NJ, Nicholson AM, Finch NA, Flynn H, Adamson J, et al. 2011. Expanded GGGGCC hexanucleotide repeat in noncoding region of C9ORF72 causes chromosome 9p-linked FTD and ALS. Neuron 72: 245–256.

Delaneau O, Ongen H, Brown AA, Fort A, Panousis NI, Dermitzakis ET. 2017. A complete tool set for molecular QTL discovery and analysis. Nat Commun 8: 15452.

Desseyn J-L, Aubert J-P, Porchet N, Laine A. 2000. Evolution of the Large Secreted Gel-Forming Mucins. Mol Biol Evol 17: 1175–1184.

Ebert P, Audano PA, Zhu Q, Rodriguez-Martin B, Porubsky D, Bonder MJ, Sulovari A, Ebler J, Zhou W, Serra Mari R, et al. 2021. Haplotype-resolved diverse human genomes and integrated analysis of structural variation. Science 372. https://www.ncbi.nlm.nih.gov/pubmed/33632895.

Ebler J, Ebert P, Clarke WE, Rausch T, Audano PA, Houwaart T, Mao Y, Korbel JO, Eichler EE, Zody MC, et al. 2022. Pangenome-based genome inference allows efficient and accurate genotyping across a wide spectrum of variant classes. Nat Genet 54: 518–525.

Edgar RC. 2004. MUSCLE: multiple sequence alignment with high accuracy and high throughput. Nucleic Acids Res 32: 1792–1797.

Eggertsson HP, Kristmundsdottir S, Beyter D, Jonsson H, Skuladottir A, Hardarson MT, Gudbjartsson DF, Stefansson K, Halldorsson BV, Melsted P.2019. GraphTyper2 enables *population-scale genotyping of structural variation using pangenome graphs*. Nat Commun 10: 5402.

Eichler EE, Flint J, Gibson G, Kong A, Leal SM, Moore JH, Nadeau JH. 2010. Missing *heritability and strategies for finding the underlying causes of complex disease*. Nat Rev Genet 11: 446–450.

Eizenga JM, Novak AM, Sibbesen JA, Heumos S, Ghaffaari A, Hickey G, Chang X, Seaman JD, Rounthwaite R, Ebler J, et al. 2020. Pangenome Graphs. Annu Rev Genomics Hum Genet 21: 139–162.

Eslami Rasekh M, Hernández Y, Drinan SD, Fuxman Bass JI, Benson G. 2021. Genome-wide characterization of human minisatellite VNTRs: population-specific alleles and gene expression differences. Nucleic Acids Res 49: 4308–4324.

Fallon PG, Sasaki T, Sandilands A, Campbell LE, Saunders SP, Mangan NE, Callanan JJ, Kawasaki H, Shiohama A, Kubo A, et al. 2009. A homozygous frameshift mutation in the mouse Flg gene facilitates enhanced percutaneous allergen priming. Nat Genet 41: 602–608.

Fudenberg G, Kelley DR, Pollard KS. 2020. Predicting 3D genome folding from DNA sequence with Akita. Nat Methods 17: 1111–1117.

Garg P, Martin-Trujillo A, Rodriguez OL, Gies SJ, Hadelia E, Jadhav B, Jain M, Paten B, Sharp AJ. 2021. Pervasive cis effects of variation in copy number of large tandem repeats on local DNA methylation and gene expression. Am J Hum Genet 108: 809–824.

Grant CE, Bailey TL, Noble WS. 2011. FIMO: scanning for occurrences of a given motif. Bioinformatics 27: 1017–1018.

GTEx Consortium. 2020. The GTEx Consortium atlas of genetic regulatory effects across human tissues. Science 369: 1318–1330.

Gupta S, Stamatoyannopoulos JA, Bailey TL, Noble WS. 2007. Quantifying similarity between motifs. Genome Biol 8: R24.

Gymrek M, Willems T, Guilmatre A, Zeng H, Markus B, Georgiev S, Daly MJ, Price AL, Pritchard JK, Sharp AJ, et al. 2016. Abundant contribution of short tandem repeats to gene expression variation in humans. Nat Genet 48: 22–29.

Hannan AJ. 2018. Tandem repeats mediating genetic plasticity in health and disease. Nat Rev Genet 19: 286–298.

Harris CR, Millman KJ, van der Walt SJ, Gommers R, Virtanen P, Cournapeau D, Wieser E, Taylor J, Berg S, Smith NJ, et al. 2020. Array programming with NumPy. Nature 585: 357–362.

Iqbal Z, Caccamo M, Turner I, Flicek P, McVean G. 2012. De novo assembly and genotyping of variants using colored de Bruijn graphs. Nat Genet 44: 226–232.

Jeffreys AJ, Tamaki K, MacLeod A, Monckton DG, Neil DL, Armour JA. 1994. Complex gene conversion events in germline mutation at human minisatellites. Nat Genet 6: 136–145.

Kelley DR. 2020. Cross-species regulatory sequence activity prediction. PLoS Comput Biol 16: e1008050.

Kent WJ, Sugnet CW, Furey TS, Roskin KM, Pringle TH, Zahler AM, Haussler D. 2002. The human genome browser at UCSC. Genome Res 12: 996–1006.

Kirby A, Gnirke A, Jaffe DB, Barešová V, Pochet N, Blumenstiel B, Ye C, Aird D, Stevens C, Robinson JT, et al. 2013. Mutations causing medullary cystic kidney disease type 1 lie in a large VNTR in MUC1 missed by massively parallel sequencing. Nat Genet 45: 299–303.

Landefeld CC, Hodgkinson CA, Spagnolo PA, Marietta CA, Shen P-H, Sun H, Zhou Z, Lipska BK, Goldman D. 2018. Effects on gene expression and behavior of untagged short tandem repeats: the case of arginine vasopressin receptor 1a (AVPR1a) and externalizing behaviors. Transl Psychiatry 8: 1–10.

Lappalainen T, Sammeth M, Friedländer MR, ‘t Hoen PAC, Monlong J, Rivas MA, Gonzàlez-Porta M, Kurbatova N, Griebel T, Ferreira PG, et al. 2013. Transcriptome and genome sequencing uncovers functional variation in humans. Nature 501: 506–511.

Liao W-W, Asri M, Ebler J, Doerr D, Haukness M, Hickey G, Lu S, Lucas JK, Monlong J, Abel HJ, et al. 2022. A Draft Human Pangenome Reference. bioRxiv 2022.07.09.499321.http://dx.doi.org/10.1101/2022.07.09.499321 (Accessed January 23, 2023).

Li H, Feng X, Chu C. 2020. The design and construction of reference pangenome graphs with minigraph. Genome Biol 21: 265.

Linthorst J, Meert W, Hestand MS, Korlach J, Vermeesch JR, Reinders MJT, Holstege H. 2020. Extreme enrichment of VNTR-associated polymorphicity in human subtelomeres: genes with most VNTRs are predominantly expressed in the brain. Transl Psychiatry 10: 369.

Lu T-Y, Chaisson MJP. 2021. Profiling variable-number tandem repeat variation across populations using repeat-pangenome graphs. Nat Commun 12: 1–12.

Merkenschlager M, Nora EP. 2016. CTCF and Cohesin in Genome Folding and Transcriptional Gene Regulation. Annu Rev Genomics Hum Genet 17: 17–43.

Mitra I, Huang B, Mousavi N, Ma N, Lamkin M, Yanicky R, Shleizer-Burko S, Lohmueller KE,Gymrek M. 2021. Patterns of de novo tandem repeat mutations and their role in autism. Nature 589: 246–250.

Muggli MD, Bowe A, Noyes NR, Morley PS, Belk KE, Raymond R, Gagie T, Puglisi SJ,Boucher C. 2017. Succinct colored de Bruijn graphs. Bioinformatics 33: 3181–3187.

Mukamel RE, Handsaker RE, Sherman MA, Barton AR, Zheng Y, McCarroll SA, Loh P.R.2021. Protein-coding repeat polymorphisms strongly shape diverse human phenotypes. Science 373: 1499–1505.

Narzisi G, Corvelo A, Arora K, Bergmann EA, Shah M, Musunuri R, Emde A-K, Robine N, Vacic V, Zody MC. 2018. Genome-wide somatic variant calling using localized colored de Bruijn graphs. Commun Biol 1: 20.

Novroski NMM, King JL, Churchill JD, Seah LH, Budowle B. 2016. Characterization of genetic *sequence variation of 58 STR loci in four major population groups*. Forensic Sci Int Genet 25: 214–226.

Nurk S, Koren S, Rhie A, Rautiainen M, Bzikadze AV, Mikheenko A, Vollger MR, Altemose N, Uralsky L, Gershman A, et al. 2022. The complete sequence of a human genome. Science 376: 44–53.

Ren J, Gu B, Chaisson MJP. 2022. vamos: VNTR annotation using efficient motif sets. bioRxiv 2022.10.07.511371. http://dx.doi.org/10.1101/2022.10.07.511371 (Accessed February 25,2023).

Renton AE, Majounie E, Waite A, Simón-Sánchez J, Rollinson S, Gibbs JR, Schymick JC, Laaksovirta H, van Swieten JC, Myllykangas L, et al. 2011. A hexanucleotide repeat expansion in C9ORF72 is the cause of chromosome 9p21-linked ALS-FTD. Neuron 72: 257–268.

Rosenbloom KR, Sloan CA, Malladi VS, Dreszer TR, Learned K, Kirkup VM, Wong MC, Maddren M, Fang R, Heitner SG, et al. 2013. ENCODE data in the UCSC Genome Browser: year 5 update. Nucleic Acids Res 41: D56–63.

Seabold S, Perktold J. 2010. Statsmodels: Econometric and Statistical Modeling with Python. Proceedings of the Python in Science Conference.http://dx.doi.org/10.25080/majora-92bf1922-011.

Sirén J, Monlong J, Chang X, Novak AM, Eizenga JM, Markello C, Sibbesen JA, Hickey G, Chang P-C, Carroll A, et al. 2021. Pangenomics enables genotyping of known structural variants in 5202 diverse genomes. Science 374: abg8871.

Sirén J, Monlong J, Chang X, Novak AM, Eizenga JM, Markello C, Sibbesen J, Hickey G, Chang P-C, Carroll A, et al. 2020. Genotyping common, large structural variations in 5,202 *genomes using pangenomes, the Giraffe mapper, and the vg toolkit*. bioRxiv 2020.12.04.412486. http://dx.doi.org/10.1101/2020.12.04.412486 (Accessed February 24,2021).

Smith FJD, Irvine AD, Terron-Kwiatkowski A, Sandilands A, Campbell LE, Zhao Y, Liao H, Evans AT, Goudie DR, Lewis-Jones S, et al. 2006. Loss-of-function mutations in the gene encoding filaggrin cause ichthyosis vulgaris. Nat Genet 38: 337–342.

Song JHT, Lowe CB, Kingsley DM. 2018. Characterization of a Human-Specific Tandem Repeat Associated with Bipolar Disorder and Schizophrenia. Am J Hum Genet 103: 421–430.

Soochit W, Sleutels F, Stik G, Bartkuhn M, Basu S, Hernandez SC, Merzouk S, Vidal E, Boers R, Boers J, et al. 2021. CTCF chromatin residence time controls three-dimensional genome organization, gene expression and DNA methylation in pluripotent cells. Nat Cell Biol 23: 881–893.

Sudmant PH, Rausch T, Gardner EJ, Handsaker RE, Abyzov A, Huddleston J, Zhang Y, Ye K, Jun G, Fritz MH-Y, et al. 2015. An integrated map of structural variation in 2,504 human genomes. Nature 526: 75–81.

Sun JH, Zhou L, Emerson DJ, Phyo SA, Titus KR, Gong W, Gilgenast TG, Beagan JA, Davidson BL, Tassone F, et al. 2018. Disease-Associated Short Tandem Repeats Co-localize with Chromatin Domain Boundaries. Cell 175: 224–238.e15.

Taliun D, Harris DN, Kessler MD, Carlson J, Szpiech ZA, Torres R, Taliun SAG, Corvelo A, Gogarten SM, Kang HM, et al. 2021. Sequencing of 53,831 diverse genomes from the NHLBI TOPMed Program. Nature 590: 290–299.

The 1000 Genomes Project Consortium. 2015. A global reference for human genetic variation. Nature 526: 68–74.

Tsuge M, Hamamoto R, Silva FP, Ohnishi Y, Chayama K, Kamatani N, Furukawa Y, Nakamura Y. 2005. A variable number of tandem repeats polymorphism in an E2F-1 binding element in the 5’ flanking region of SMYD3 is a risk factor for human cancers. Nat Genet 37: 1104–1107.

Viguera E, Canceill D, Ehrlich SD. 2001. Replication slippage involves DNA polymerase pausing and dissociation. EMBO J 20: 2587–2595.

Virtanen P, Gommers R, Oliphant TE, Haberland M, Reddy T, Cournapeau D, Burovski E, Peterson P, Weckesser W, Bright J, et al. 2020. SciPy 1.0: fundamental algorithms for scientific computing in Python. Nat Methods 17: 261–272.

Vollebregt O, Koyama E, Zai CC, Shaikh SA, Lisoway AJ, Kennedy JL, Beitchman JH. 2021. Evidence for association of vasopressin receptor 1A promoter region repeat with childhood onset aggression. J Psychiatr Res 140: 522–528.

Wang G, Sarkar A, Carbonetto P, Stephens M. 2020. A simple new approach to variable *selection in regression, with application to genetic fine mapping*. Journal of the Royal Statistical Society: Series B (Statistical Methodology) 82: 1273–1300.

Waterhouse AM, Procter JB, Martin DMA, Clamp M, Barton GJ. 2009. Jalview Version 2--a multiple sequence alignment editor and analysis workbench. Bioinformatics 25: 1189–1191.

