## Supplemental material for "The motif composition of variable-number tandem repeats impacts gene expression"

### Supplementary information

#### Table of contents


**Supplemental Figures** **4**

Supplemental Fig. S1. Distribution of VNTR size in GRCh38 and in the 70 HGSVC assemblies. 4

Supplemental Fig. S2. Distribution of VNTR size versus allele properties. Private alleles are those observed only once across the 70 haplotypes. 4

Supplemental Fig. S3. Distribution of aln-r2 in this work and Lu 2021. 5

Supplemental Fig. S4. Example of a locus with high mapping quality but low consistency in VNTR dosage across samples. 6

Supplemental Fig. S5. Bias correction for VNTR dosage. 7

Supplemental Fig. S6. eQTL mapping with VNTR length. 8

Supplemental Fig. S7. Size of locus-RPGG before and after compaction. 9

Supplemental Fig. S8. Absolute percentage error of motifs. 9

Supplemental Fig. S9. Bias correction on the HGSVC and HPRC dataset. 10

Supplemental Fig. S10. Distribution of the number of outlier samples across all motifs. 10

Supplemental Fig. S11. Distribution of eMotif batch-r2. 11

Supplemental Fig. S12. Distribution of λGC in GTEx. 11

Supplemental Fig. S13. λGC from biallelic variants versus VNTR motifs. 12

Supplemental Fig. S14. Distribution of eMotif P-values from Geuvadis missing in GTEx. 12

Supplemental Fig. S15. Correlation of eMotif effect sizes between Geuvadis and GTEx EBV-transformed lymphocytes. 13

Supplemental Fig. S16. Dot plot analysis of AVPR1A RS3 VNTR. 13

Supplemental Fig. S17. Frequency of CACNA1C risk motifs across the 35 HGSVC assemblies. 14

Supplemental Fig. S18. CACNA1C VNTR allele diversity. 14

Supplemental Fig. S19. Distribution of posterior inclusion probability (PIP). 15

Supplemental Fig. S20. Number of eGenes with likely causal eMotifs. 16

Supplemental Fig. S21. Number of eVNTRs with likely causal eMotifs. 17

Supplemental Fig. S22. Number of likely causal eMotifs. 18

Supplemental Fig. S23. Correlation between HRNR/FLG/FLG-AS1 expression and HRNR repeat.q 19

Supplemental Fig. S24. Alignment of CTCF motif to the HRNR repeat. 19

Supplemental Fig. S25. Correlation between HRNR repeat size and Number of predicted CTCF sites. 20

Supplemental Fig. S26. Akita's prediction versus the micro-C map from the UCSC Genome Browser. 20

Supplemental Fig. S27. Allelic diversity of the RNF213 promoter VNTR. 21
Supplemental Fig. S28. Effect of VNTR variants on RNF213 expression. 21
Supplemental Fig. S29. UCSC Genome Browser view of the RNF213 VNTR. 22
Supplemental Fig. S30. Best matching transcription factor motif for ZNF93 likely causal motif. 23
Supplemental Fig. S31. Correlation between the number of predicted ZNF93 binding sites and RNF213 VNTR variants. 24
Supplemental Fig. S32. Predicted ZNF93 binding sites for each VNTR motif. 24
Supplemental Fig. S33. Correlation between RNF213 expression level and RNF213 VNTR repeat units. 25

**Supplemental Tables 26**

Supplemental Table S1. Additional disease-relevant tandem repeats included in our reference set. 26
Supplemental Table S2. eMotif discoveries for disease-relevant genes. 28
Supplemental Table S3. eQTL mapping using a genome-wide P-value cutoff. 31

**References 33**

#### Supplemental Figures

**Supplemental Fig. S1.** Distribution of VNTR size in GRCh38 and in the 70 HGSVC assemblies.

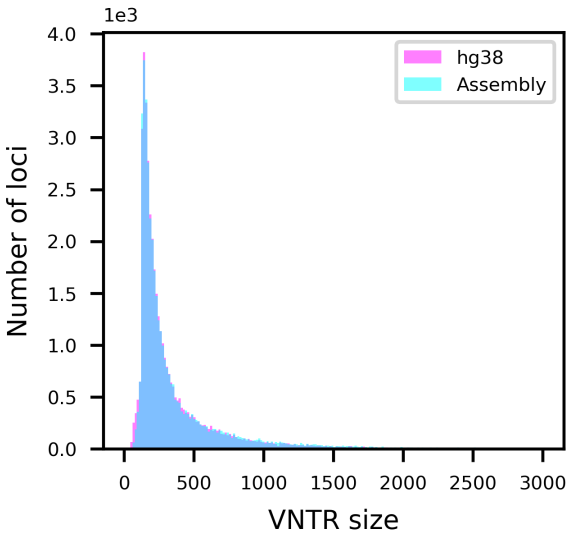


VNTR sizes greater than 3000 bp are not shown, contributing to 0.7% (281/40,314) of loci.

**Supplemental Fig. S2.** Distribution of VNTR size versus allele properties. Private alleles are those observed only once across the 70 haplotypes.

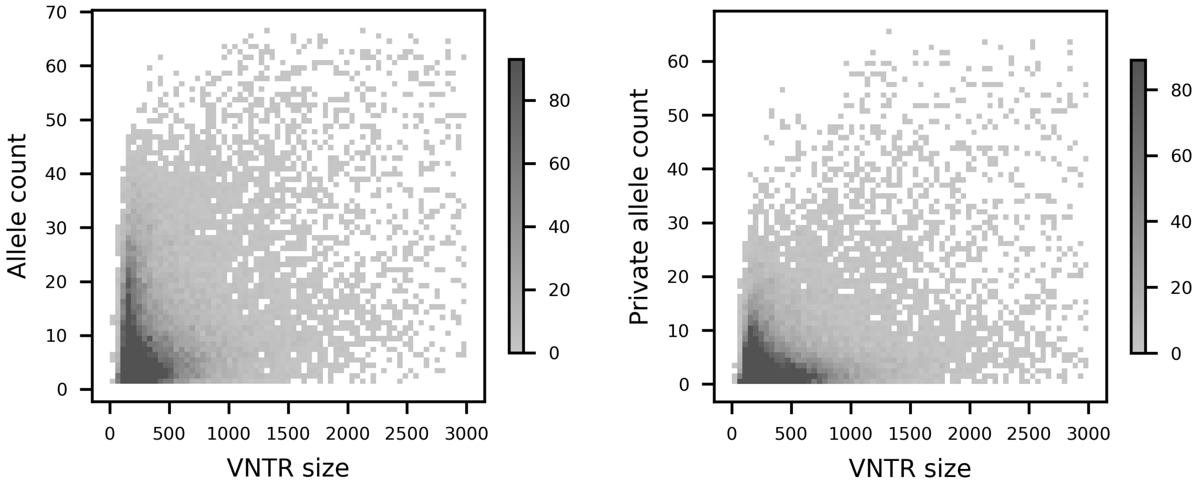


VNTR sizes greater than 3000 bp are not shown in both subpanels, contributing to 0.7% (281/40,314) of loci. Density greater than the limit shown in the color bar is clipped.

**Supplemental Fig. S3.** Distribution of aln-$r^2$ in this work and Lu 2021.

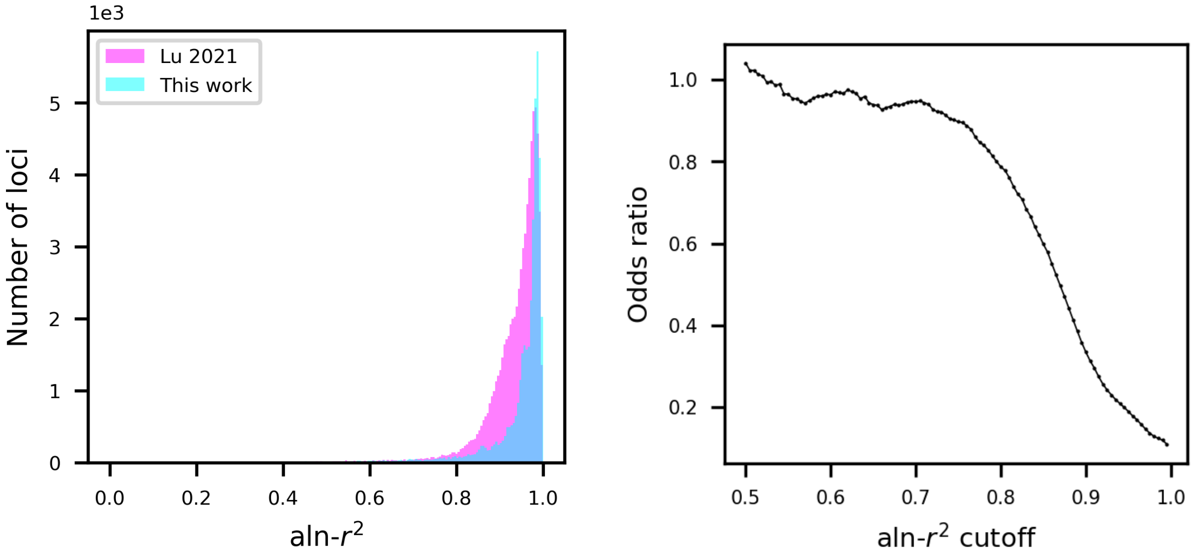


**a,** Comparing the distribution of aln-$r^2$ in both studies. **b,** Enrichment of high aln-$r^2$ loci in this work. Contingency tables with two variables, "≥ aln-$r^2$ cutoff" and "is from Lu 2021", were constructed to compute odds ratio.

**Supplemental Fig. S4.** Example of a locus with high mapping quality but low consistency in VNTR dosage across samples.

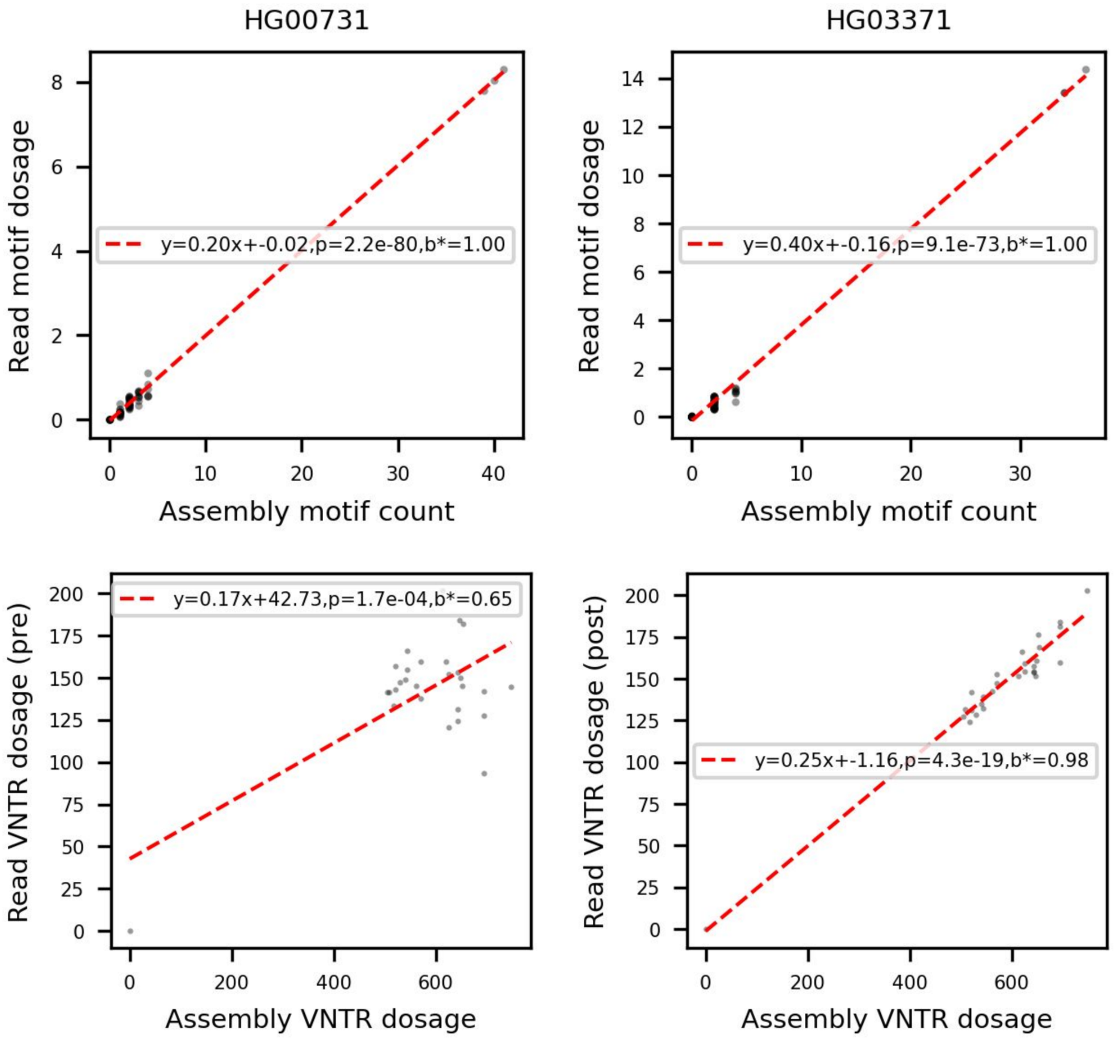


The VNTR at chr1:192,048,505-192,048,821 was selected to highlight the strong influence of bias, even when the average aln-$r^2$ was 0.99. The top two panels showed the two samples with two-fold difference in the slope of the regression lines. The batch-$r^2$ before correction (bottom left) is only 0.35 while the statistic after correction (bottom right) is 0.97.

**Supplemental Fig. S5.** Bias correction for VNTR dosage.

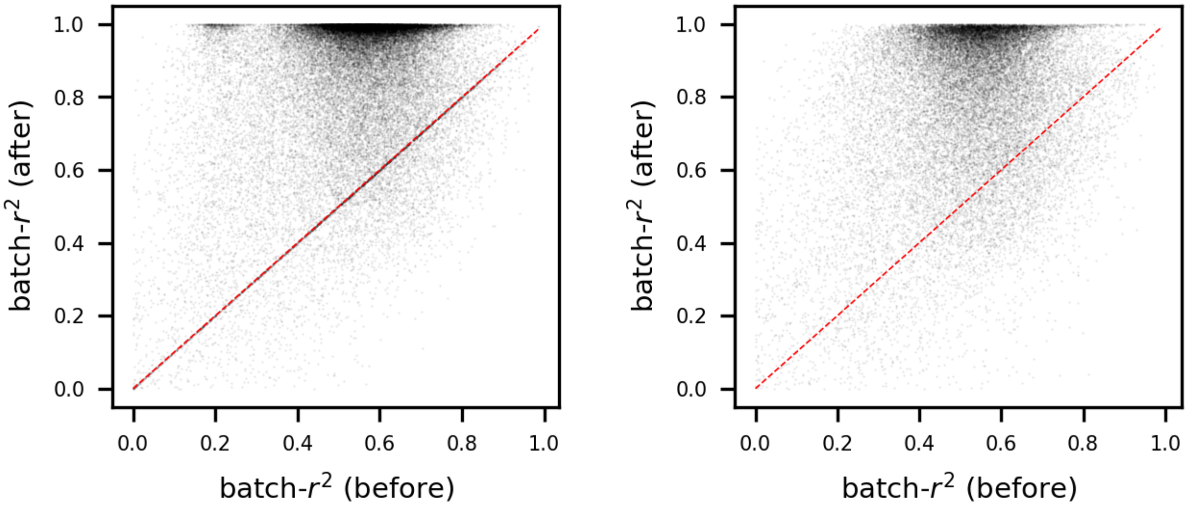


Bias correction on the HGSVC dataset (left panel). The batch-$r^2$ for 29,602 loci (75.7%) are shown. Bias correction on the HPRC dataset (right panel). The batch-$r^2$ for 27,499 loci (70.3%) are shown. The red dashed line indicates no improvement after bias correction.

**Supplemental Fig. S6**. eQTL mapping with VNTR length.

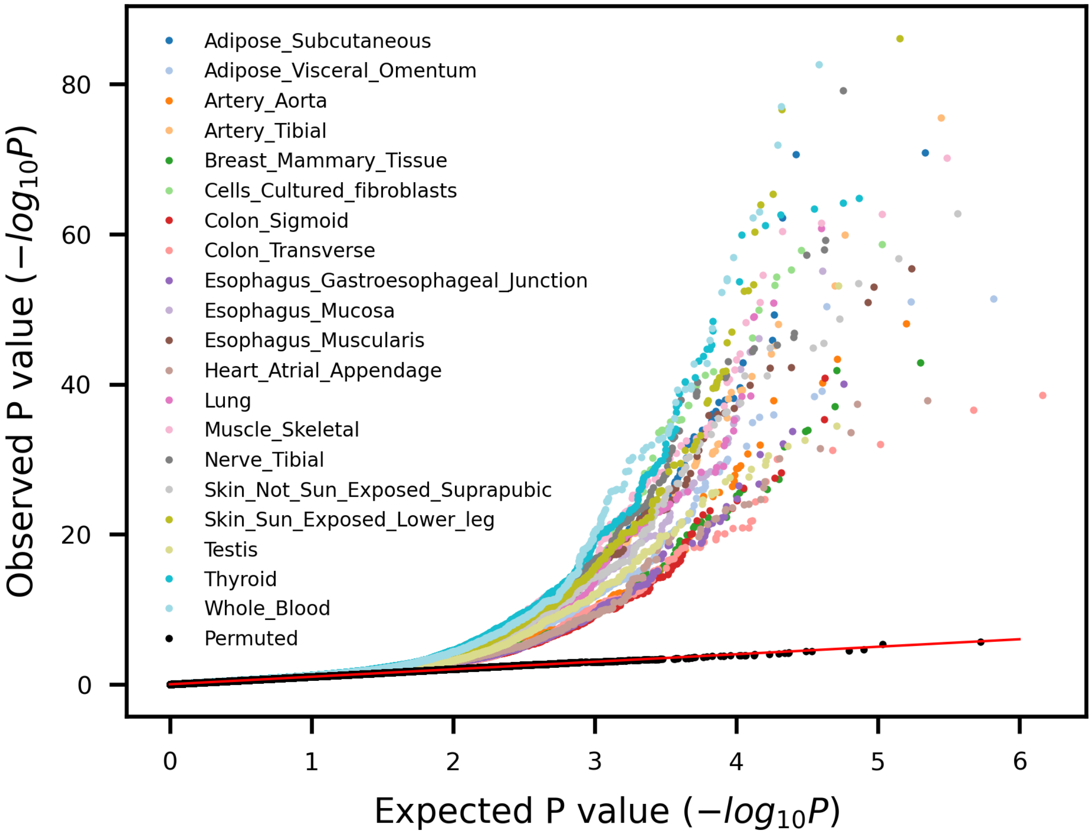


(A) Quantile-quantile plot of gene-level eVNTR discoveries using VNTR length across 20 human tissues from GTEx datasets. The expected P-values (x-axis) were drawn from Unif(0,1) and plotted against observed nominal association P-values.

**Supplemental Fig. S7.** Size of locus-RPGG before and after compaction.

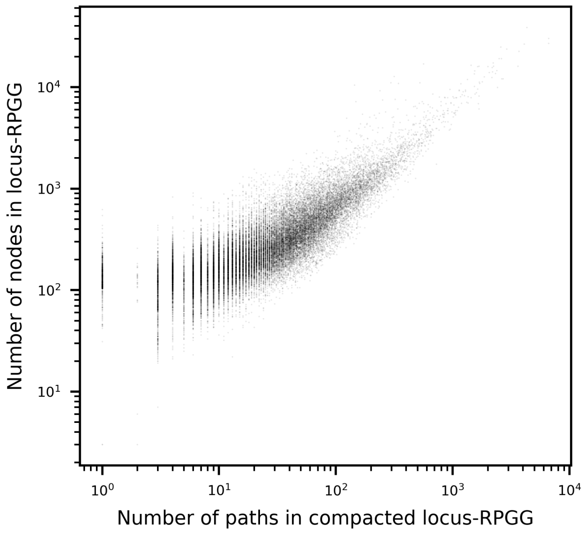


The number of nodes before compaction (y-axis) and the number of paths after compaction (x-axis) were compared for all 80,518 loci in the RPGG, both illustrated on a log scale.

**Supplemental Fig. S8.** Absolute percentage error of motifs.

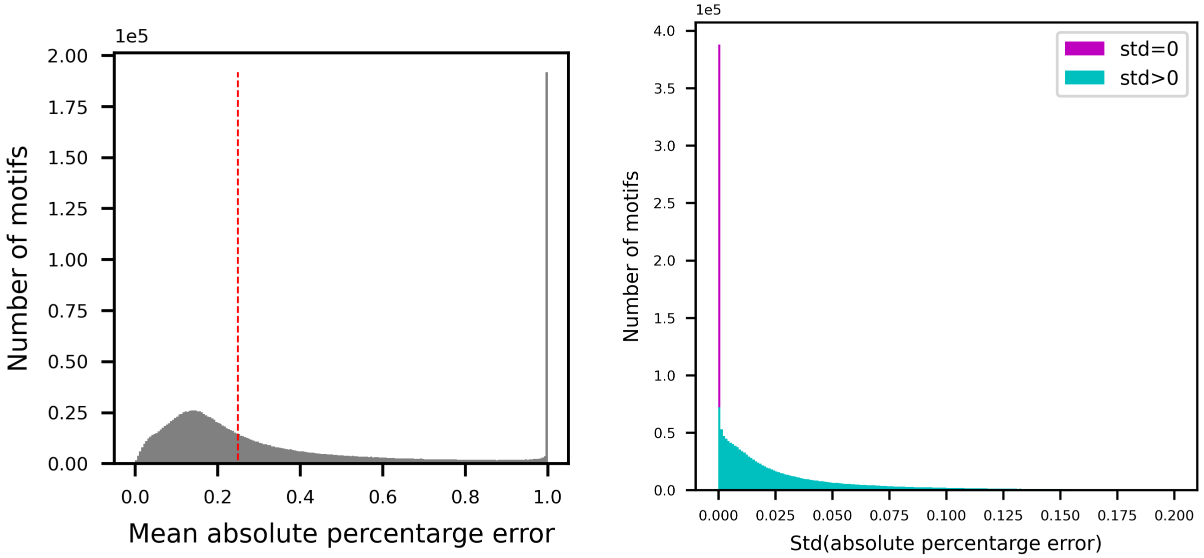


(Left) Distribution of motif-MAPE. For each motif, the mean absolute percentage error (MAPE) was computed by averaging the absolute percentage errors (APEs, Methods) across 35 genomes. A total of 1,724,959 motifs are shown in the plot, excluding the remaining 4.7% with MAPE greater than 1. (Right) Distribution of the standard deviation of motif-MAPE. Ground truth is measured from motif count in assemblies, and absolute percentage error is measured by regression from read depth on edges (motif) to assembly count. The spike at std=0 corresponds to motifs with only one observation across all haplotypes. Motifs with std > 0.2 are not shown, contributing to 7.8% (140,989/1,810,042) of motifs.

**Supplemental Fig. S9.** Bias correction on the HGSVC and HPRC dataset.

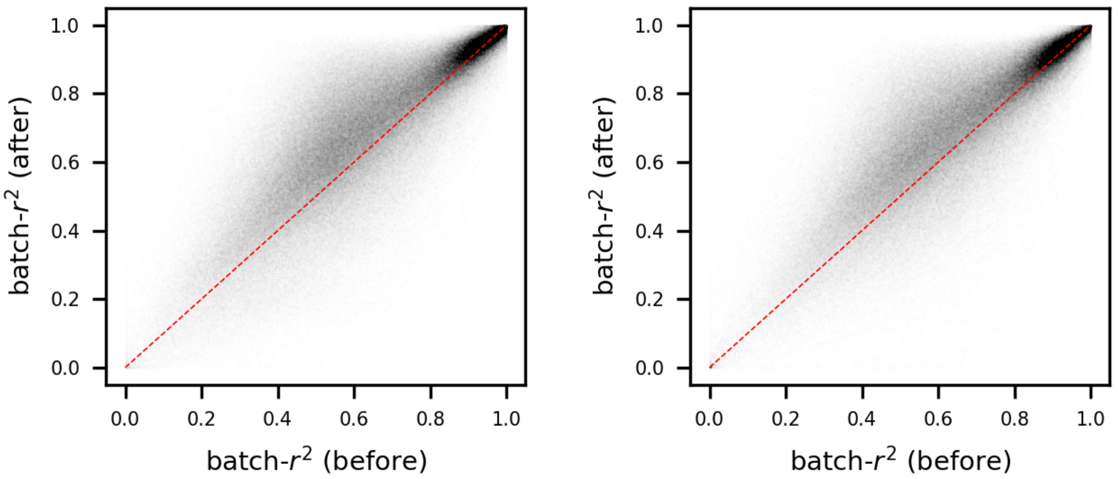


Motif batch-$r^2$ before and after bias correction were compared on the HGSVC (left panel) and the HPRC assemblies (right panel). VNTRs without invariant *k*-mers were removed from analysis; motifs in HPRC that do not overlap any k-mers in the same locus in HGSVC were removed as well, giving a 86.9% (n=668,961) and 76.8% (n=590,873) ascertainment rate in HGSVC and HPRC. The red dashed lines indicate x=y.

**Supplemental Fig. S10.** Distribution of the number of outlier samples across all motifs.

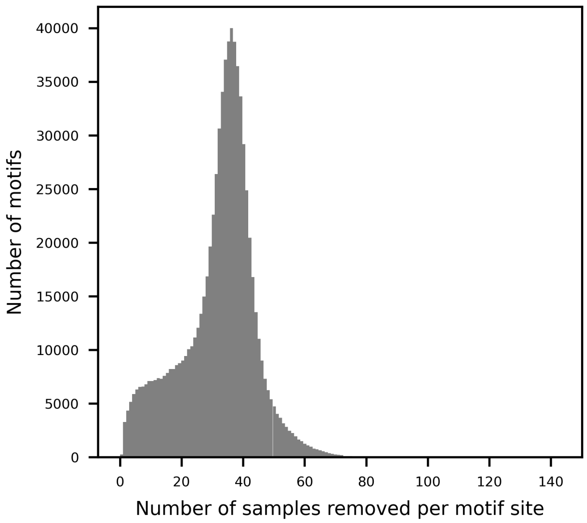


For each motif, samples with motif dosage greater than two standard deviations away from the mean were removed. Data points were collected from the final set of 769,616 motifs and 813 GTEx samples.

**Supplemental Fig. S11.** Distribution of eMotif batch-$r^2$.

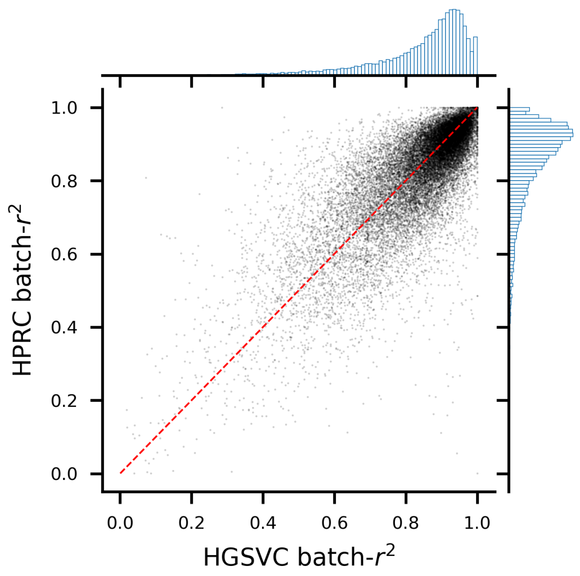


VNTRs without invariant k-mers were removed from analysis; motifs in HPRC that do not overlap any k-mers in the same locus in HGSVC were removed as well, giving a 95.5% ascertainment rate for 25,031 eMotif. The red dashed lines indicate x=y. Each bin in the marginal histogram spans 0.01.

**Supplemental Fig. S12.** Distribution of $\lambda_{GC}$ in GTEx.

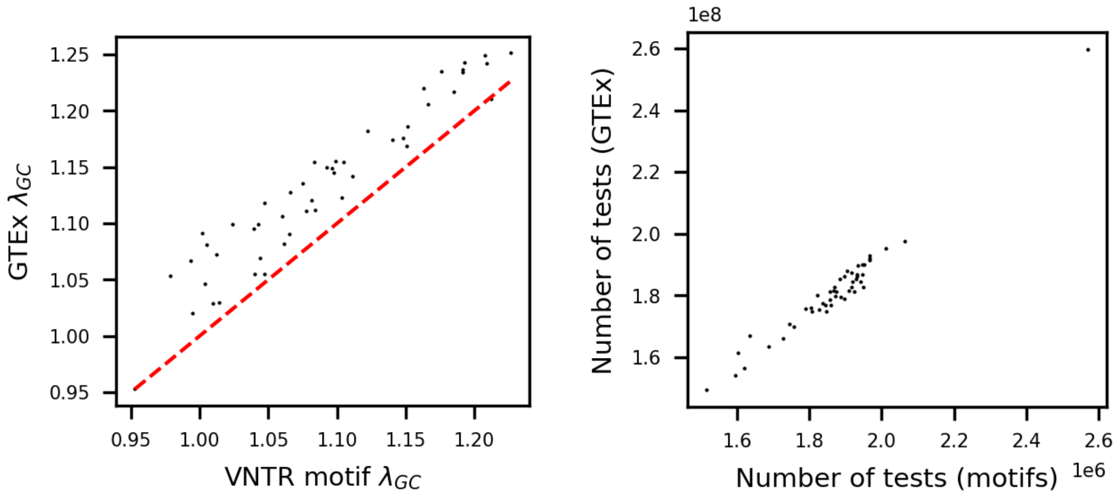


(Left panel) The $\lambda_{GC}$ from GTEx variants in each tissue is shown on the y-axis, computed using the P-values from all tests originally released by the GTEx Consortium. The $\lambda_{GC}$ from VNTR motifs in each GTEx tissue is shown on the x-axis. (Right panel) The number of tests in each tissue is compared.

**Supplemental Fig. S13.** $\lambda_{GC}$ from biallelic variants versus VNTR motifs.

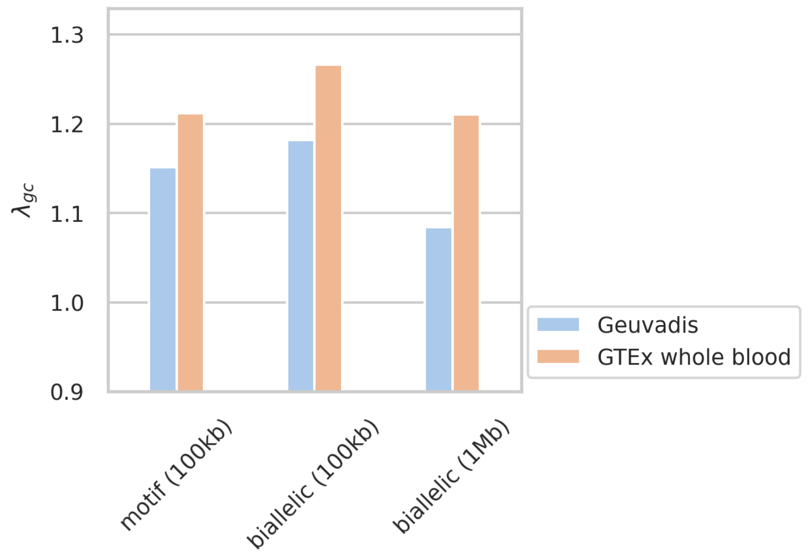


The $\lambda_{GC}$ computed from different variants (biallelic versus VNTR motifs), different datasets (GTEx versus Geuvadis), and different cis-windows (100 kb versus 1Mb) are shown.

**Supplemental Fig. S14.** Distribution of eMotif P-values from Geuvadis missing in GTEx.

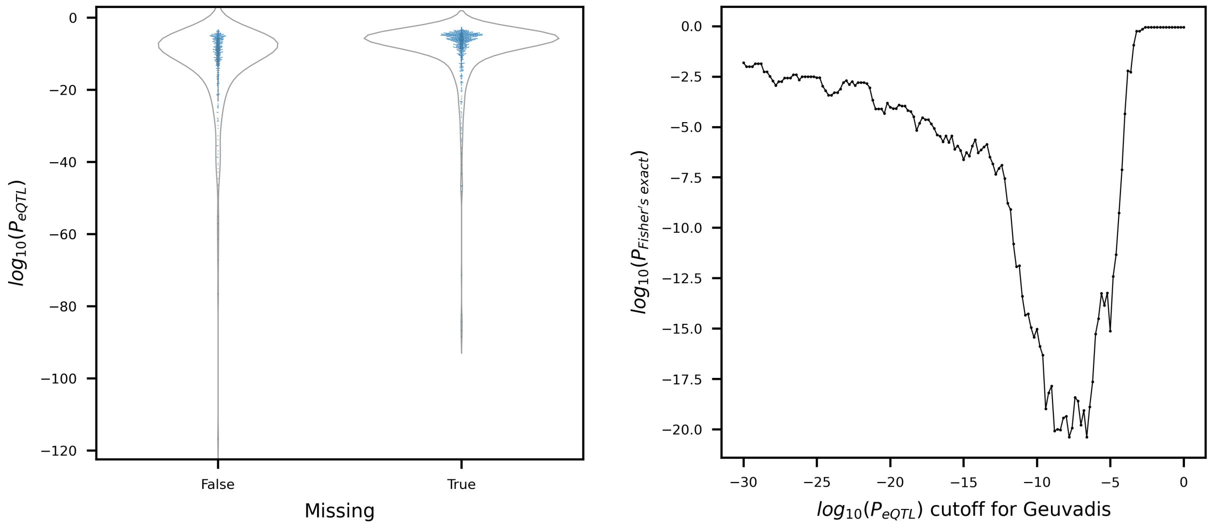


**a,** Distribution of the eMotif P-values from Geuvadis. The x-axis denotes whether the eMotifs are missing from the GTEx whole blood. **b,** Enrichment of missing eMotifs with lower significance. Contingency tables with two variables, "< $P_{eQTL}$ cutoff" and "is missing", were constructed. Significance was computed with one-sided Fisher's exact test.

**Supplemental Fig. S15.** Correlation of eMotif effect sizes between Geuvadis and GTEx EBV-transformed lymphocytes.

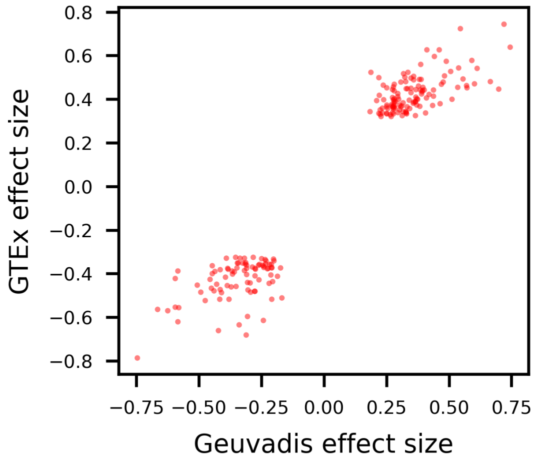


Only eGenes significant in both datasets were shown. Each pair of gene and motif that has the same/opposite sign across datasets were colored in red/black.

**Supplemental Fig. S16.** Dot plot analysis of AVPR1A RS3 VNTR.

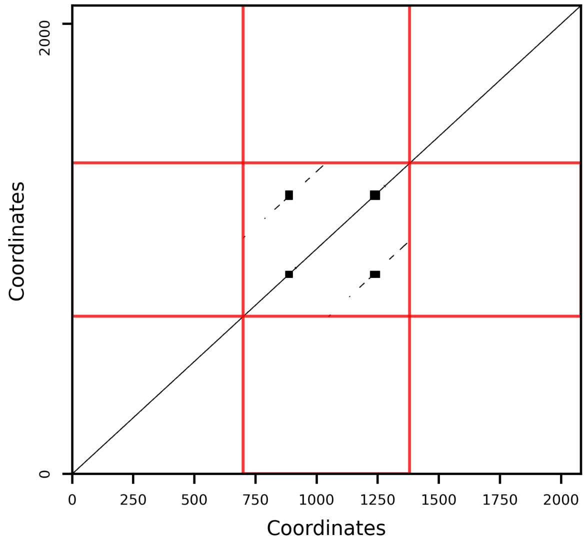


GRCh38 sequence at chr12:63,156,354-63,156,429 was plotted against itself. Each dot represents an exact matching of a 13-mer. The boundaries of 700 bp flanking sequences were indicated with red lines.

**Supplemental Fig. S17.** Frequency of CACNA1C risk motifs across the 35 HGSVC assemblies.

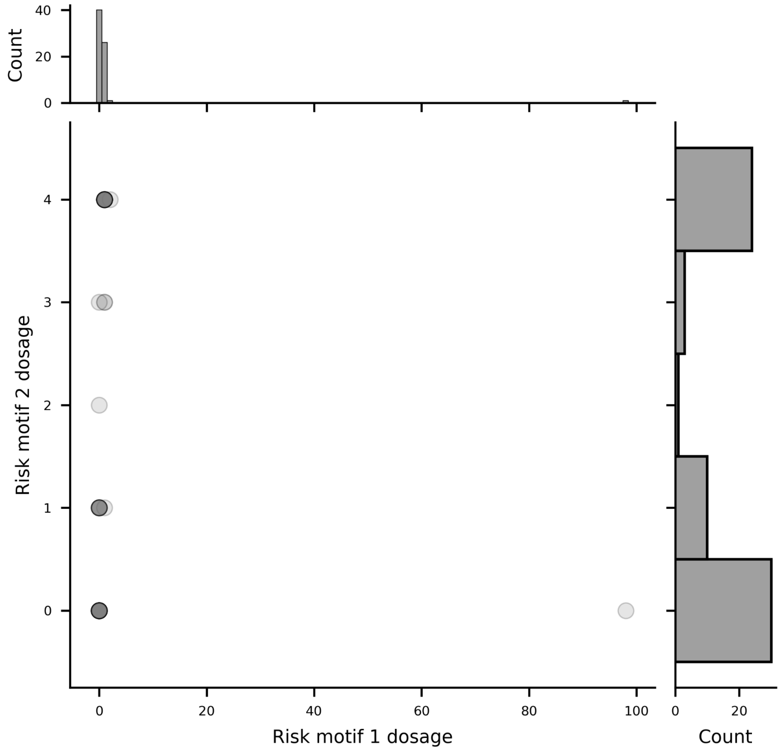


The occurrence of risk motif 1 CAACCACACGATCCTGACCTT and risk motif 2 CCCTGACCTTACTAGTTTACGA were counted considering both the motif and its reverse complement. Bin sizes for both marginal distributions are one.

**Supplemental Fig. S18.** CACNA1C VNTR allele diversity.

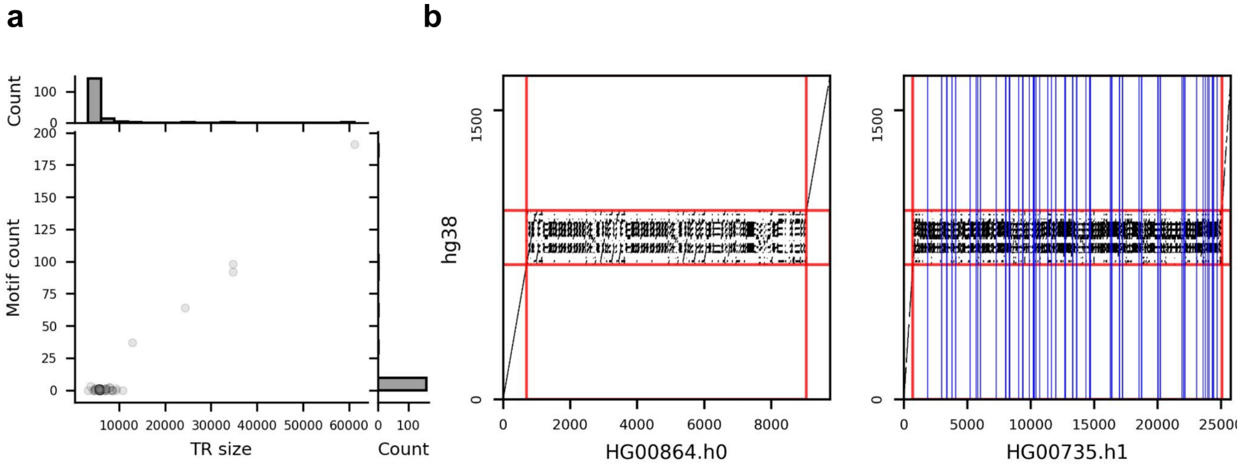


**a**, The count of risk motif 1 CAACCACACGATCCTGACCTT versus VNTR size. **b**, Two divergent haplotypes were compared with the location of risk motif 1 shown as blue lines. Red lines indicate the boundaries of the VNTR, with a 700 bp flank on each side.

**Supplemental Fig. S19.** Distribution of posterior inclusion probability (PIP).

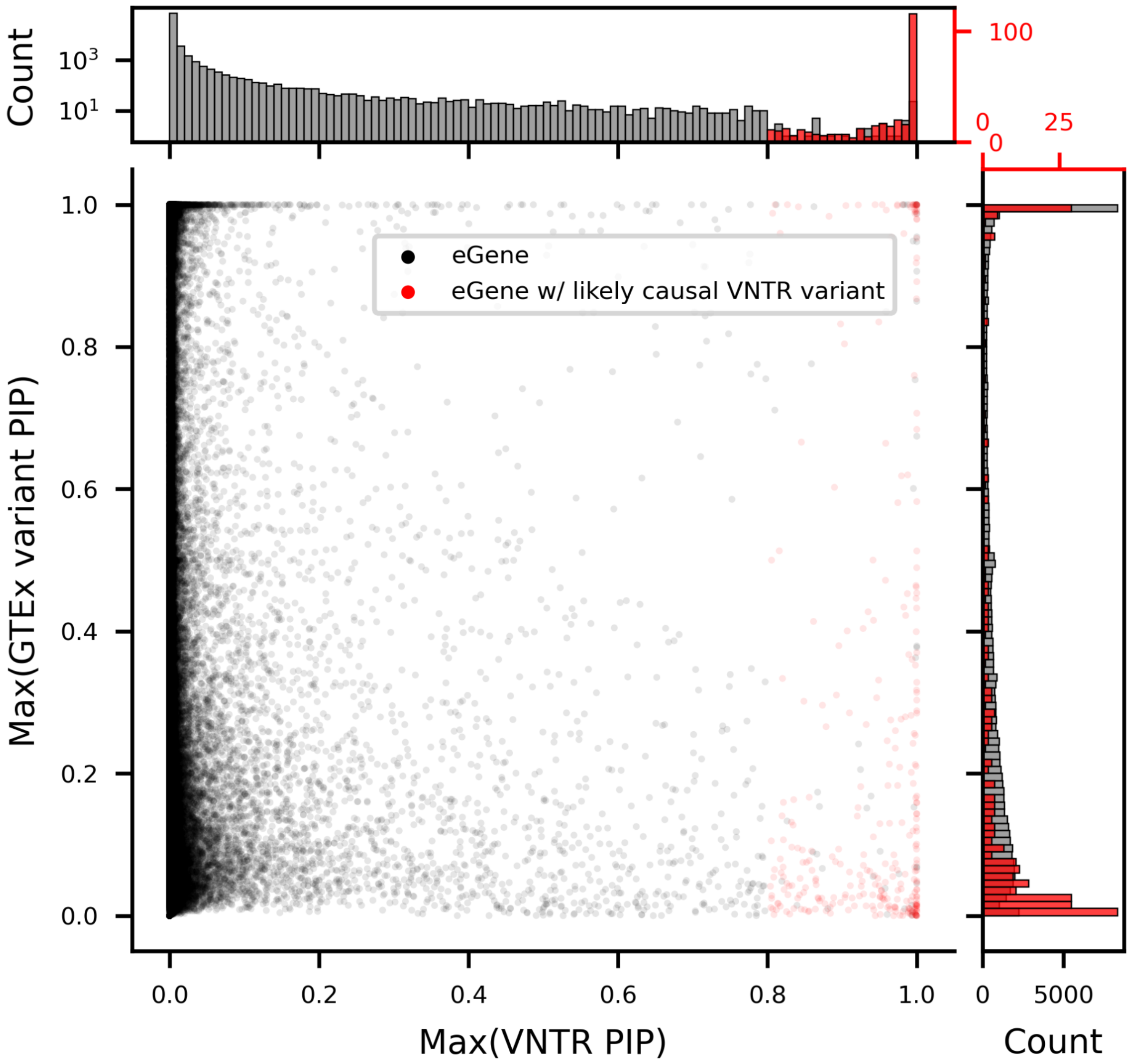


VNTR variants (motif or full VNTR length) with PIP greater than 0.8 were called likely causal. Each dot is from a fine-mapped gene in a tissue. The maximal PIPs of all VNTR/GTEx variants nearby a gene are shown on the x-axis/y-axis. Histograms show the marginal distribution of each variable; the counts of each subset of the data points are shown in histogram in different scales and colors. All counts are shown in linear scale except that the black histogram on top is shown in log scale.

**Supplemental Fig. S20.** Number of eGenes with likely causal eMotifs.

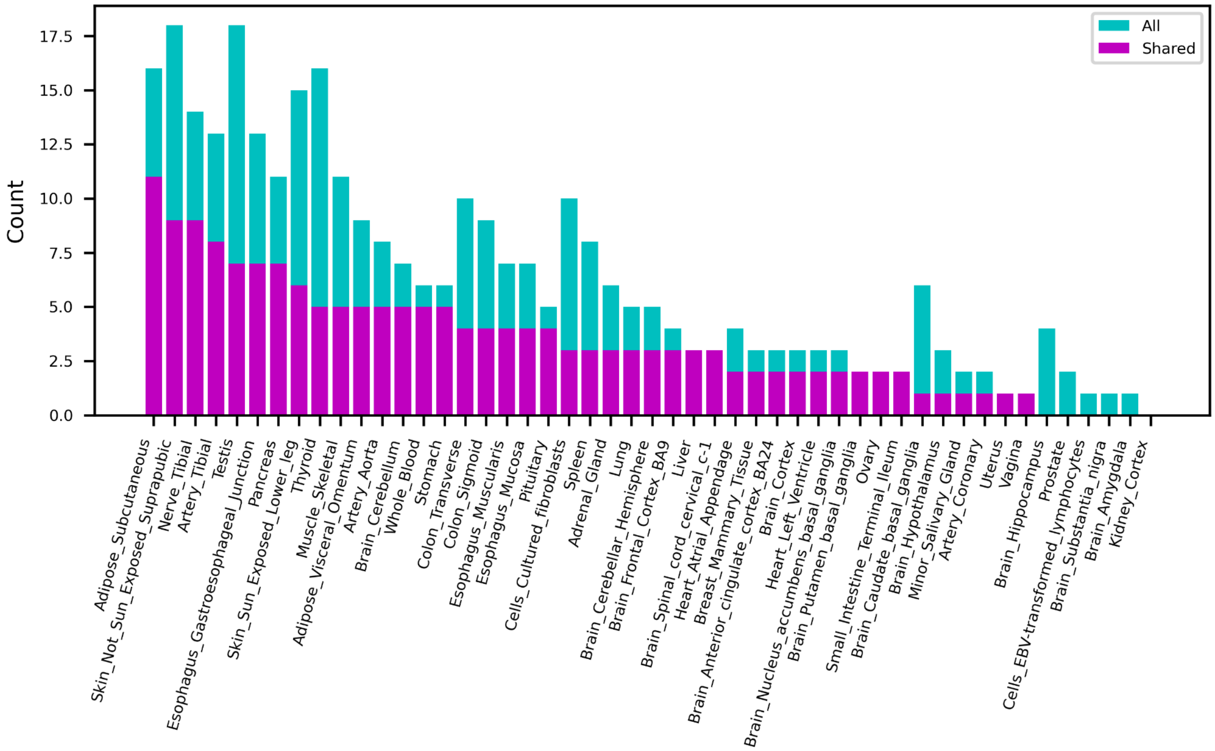


Each eGene reported by eQTL mapping was fine-mapped using all SNPs and eMotifs among the 100 kb window. eGenes of which the highest eMotif posterior inclusion probability (PIP) was greater than 0.8 were reported as containing likely causal eMotifs. An eGene was considered shared across tissues (magenta) if it was reported in at least one other tissue.

**Supplemental Fig. S21.** Number of eVNTRs with likely causal eMotifs.

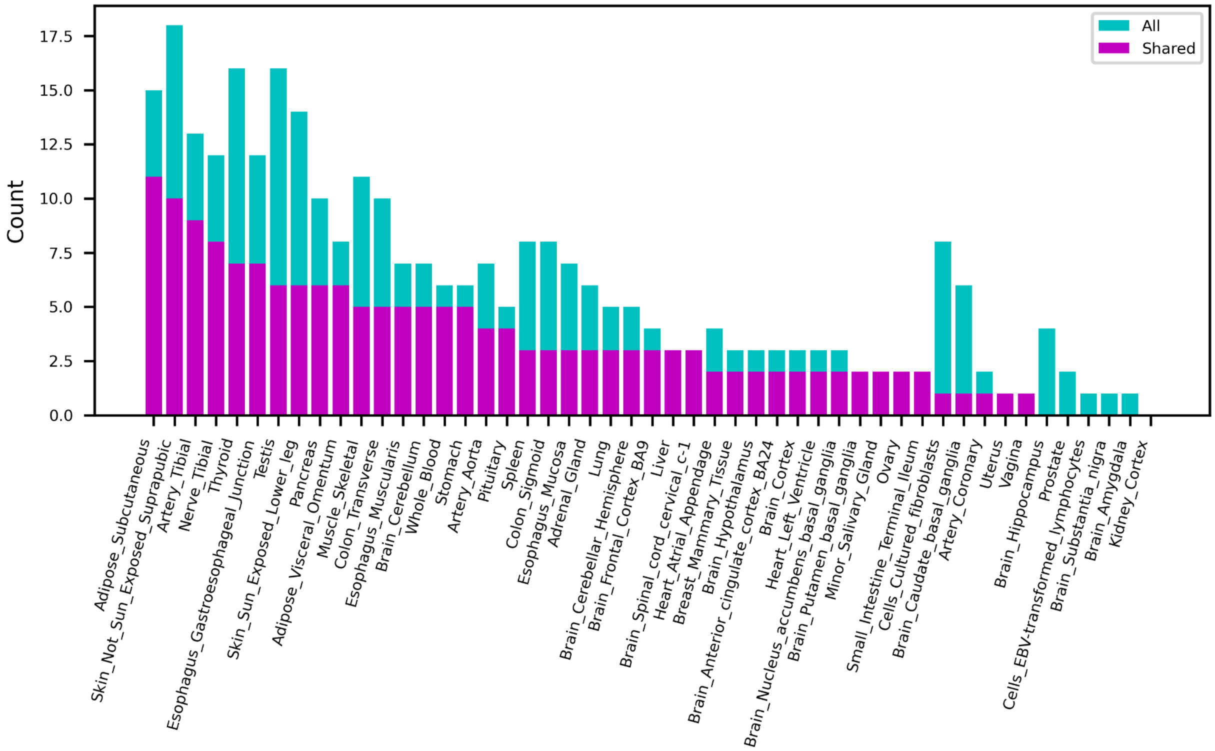


Each eGene reported by eQTL mapping was fine-mapped using all SNPs and eMotifs among the 100 kb window. eGenes of which the highest eMotif posterior inclusion probability (PIP) was greater than 0.8 were reported as containing likely causal eMotifs. An eVNTR was considered shared across tissues (magenta) if any eMotif in the VNTR was reported in at least one other tissue.

**Supplemental Fig. S22.** Number of likely causal eMotifs.

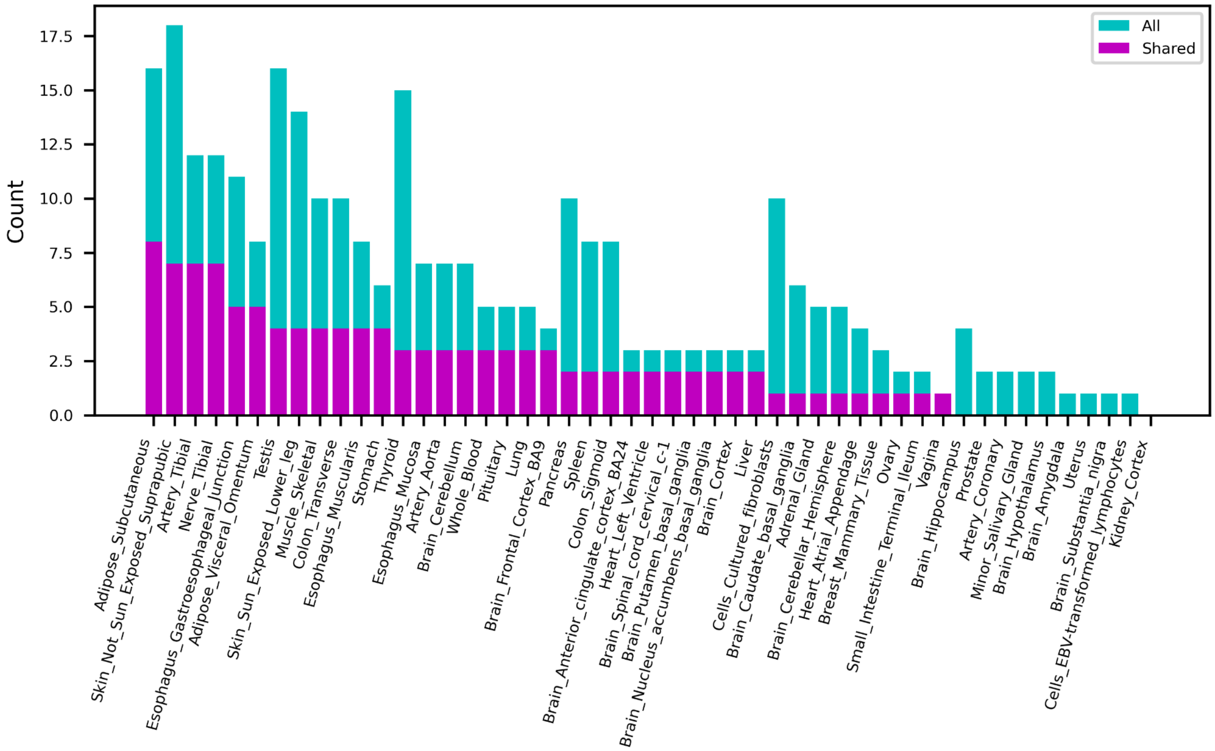


Each eGene reported by eQTL mapping was fine-mapped using all SNPs and eMotifs among the 100 kb window. eGenes of which the highest eMotif posterior inclusion probability (PIP) is greater than 0.8 were reported as containing likely causal eMotifs. An eMotif was considered shared across tissues (magenta) if it was reported in at least one other tissue.

**Supplemental Fig. S23.** Correlation between HRNR/FLG/FLG-AS1 expression and HRNR repeat.q

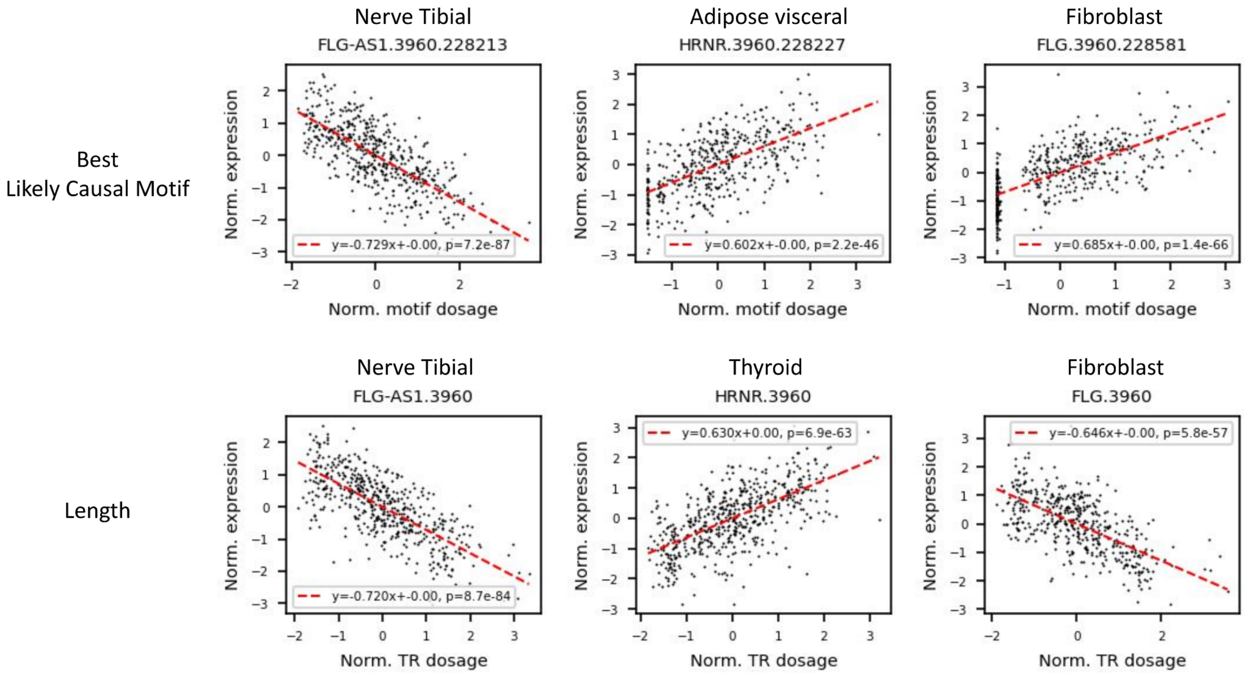


The best likely causal motif from HRNR repeat across all tissues is shown for each gene in the top three panels. The tissue with the most significant length association with each gene is shown on the bottom three panels.

**Supplemental Fig. S24.** Alignment of CTCF motif to the HRNR repeat.

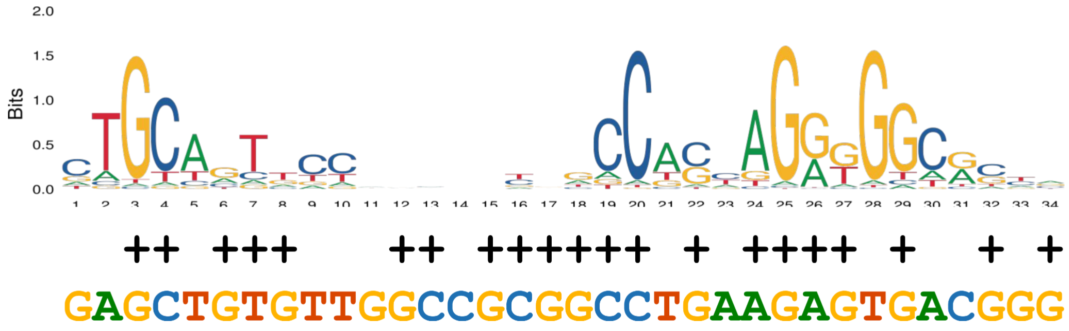


The HRNR repeat on GRCh38 (chr1:152,213,243-152,221,044) was aligned to the 34 bp, two-core CTCF motif (MA1929.1) using the MAST module (Bailey and Gribskov 1998) in the MEME suite. The best alignment at chr1:152,220,381 with a P-value of 4.6e-5 is shown.

**Supplemental Fig. S25.** Correlation between HRNR repeat size and Number of predicted CTCF sites.

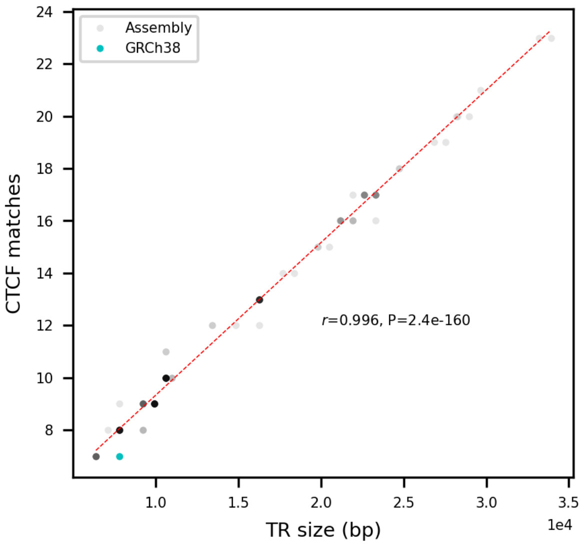


The HRNR repeat size for all HGSVC and HPRC haplotypes are shown on the x-axis. Number of CTCF sites in each haplotype was predicted using FIMO with a cut off of P < 1e-4. Red dashed line indicates the regression line, with the Pearson's *r* and the associated P-value shown.

**Supplemental Fig. S26.** Akita's prediction versus the micro-C map from the UCSC Genome Browser.

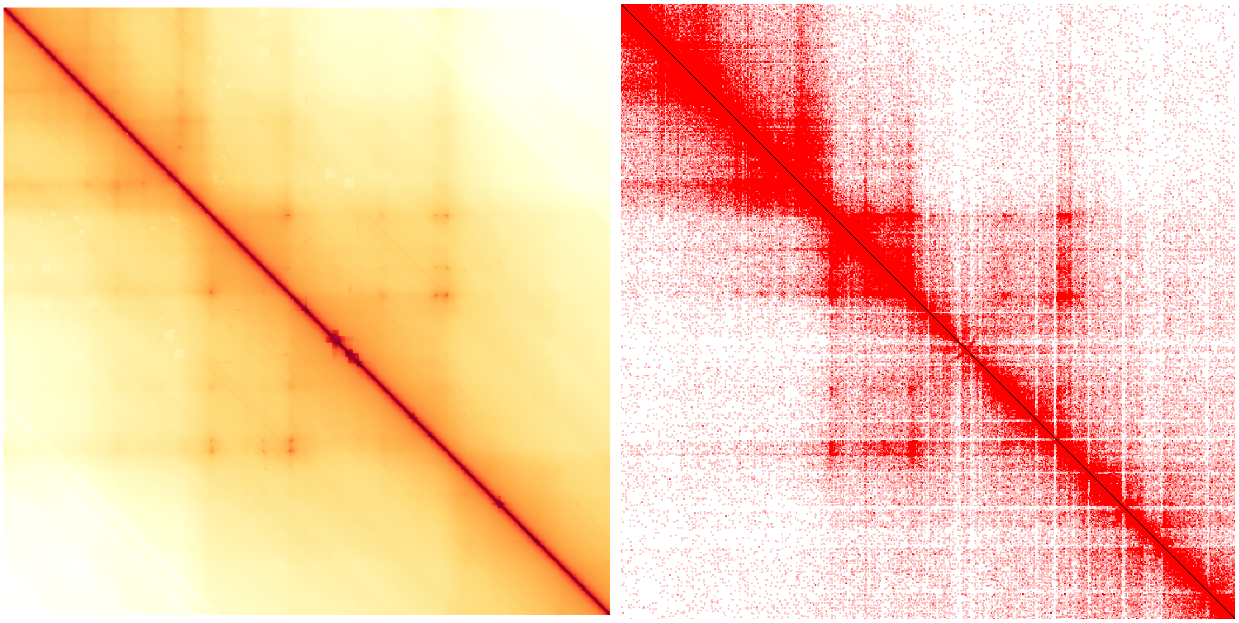

Akita's prediction for chr1:151,693,880-152,740,407 is compared with the HFFc6 micro-C map from UCSC Genome Browser. Akita's initial output "log(observed/expected)" was transformed to an "observed" map to be visually compared with the raw micro-C map. The "expected" map was generated by averaging corrected counts in the region over increasing genomic distances, using cooltools observed_over_expected (Open2C et al. 2022).

**Supplemental Fig. S27.** Allelic diversity of the *RNF213* promoter VNTR.

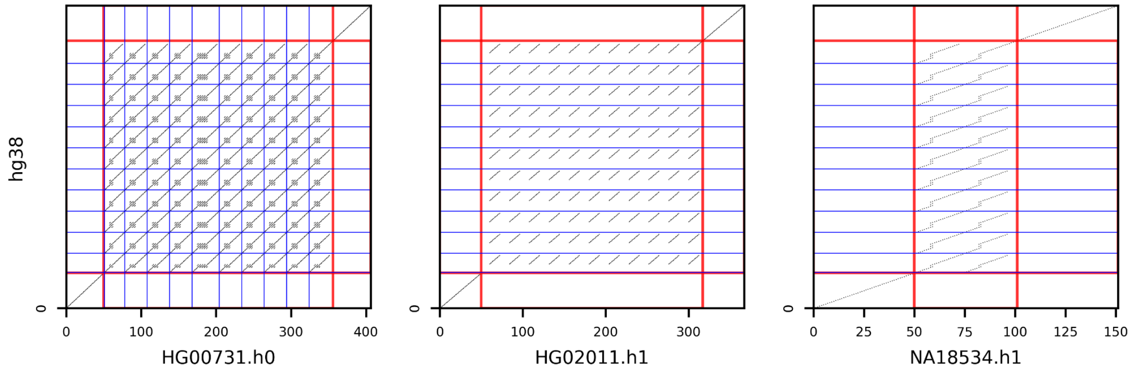


The dot plot of three divergent haplotypes are shown to demonstrate repeat expansion with different motifs. Blue lines represent the red repeat unit in Figure 5. Each dot represents a match of 21-mers. Red lines represent the boundaries of VNTR, with a flank of 50 bp on each side.

**Supplemental Fig. S28.** Effect of VNTR variants on *RNF213* expression.

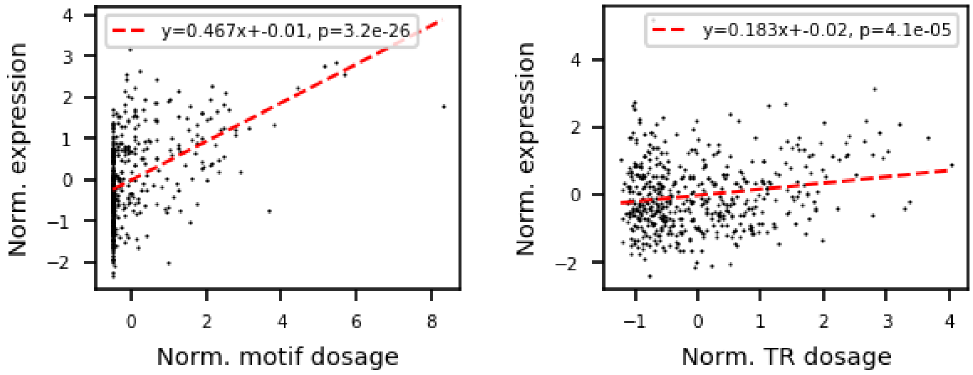


(Left panel) Significant association of the likely causal motif GCGGGGCCGGCGGCGGCGGCGG in the promoter VNTR (chr17:80,260,506-80,260,846) with *RNF213* expression. Outliers are not removed to demonstrate the strong effect from extreme motif expansion. The P-value and effect size for the motif after outlier removal were $6.3 \times 10^{-16}$ and 0.37, respectively. (Right panel) Mild effect from the repeat size of the *RNF213* VNTR. Expression in GTEx thyroid is shown for both panels.

**Supplemental Fig. S29.** UCSC Genome Browser view of the *RNF213* VNTR.

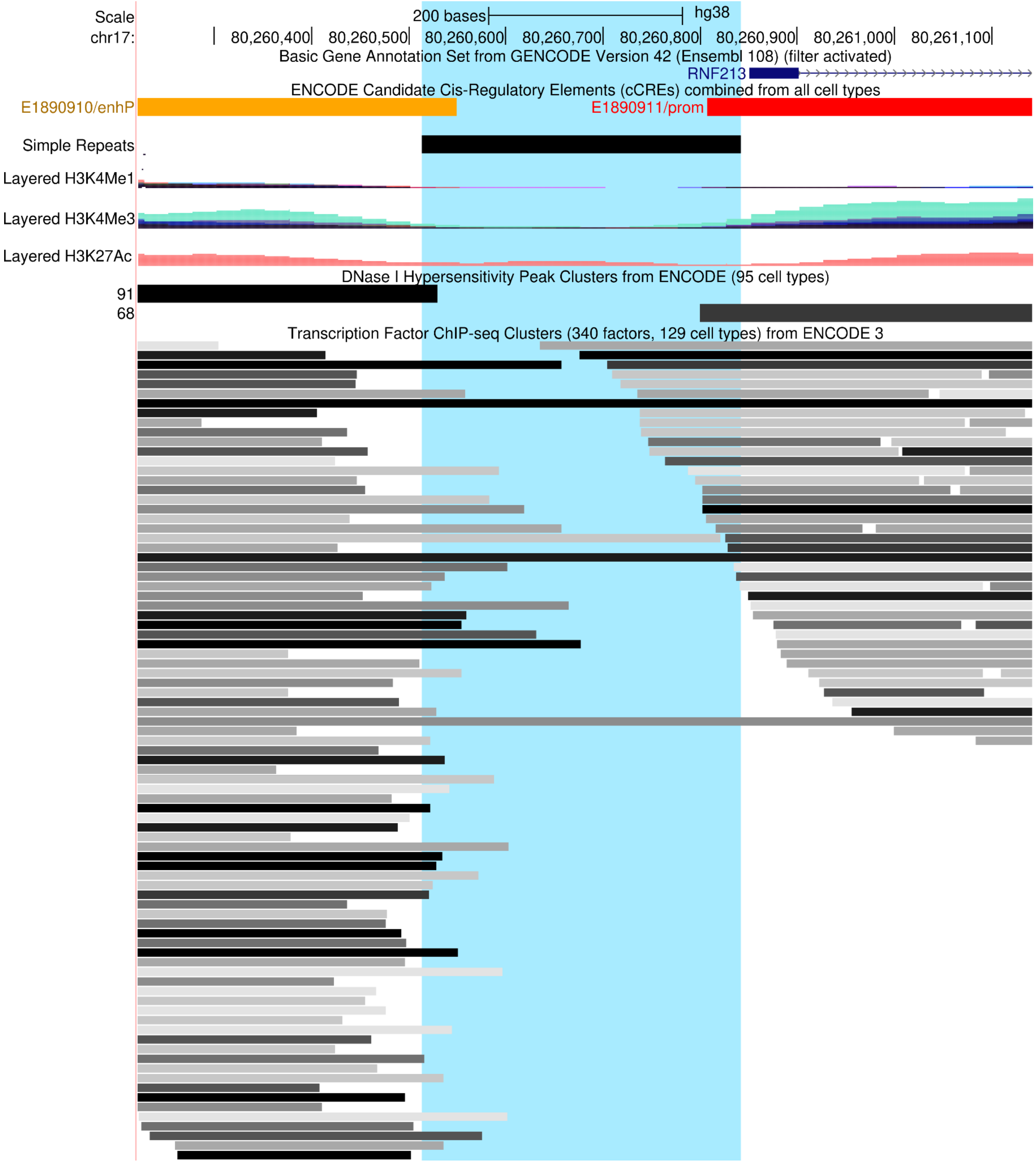


The *RNF* promoter repeat at chr17:80,260,506-80,260,846 was highlighted in light blue.

**Supplemental Fig. S30.** Best matching transcription factor motif for ZNF93 likely causal motif.

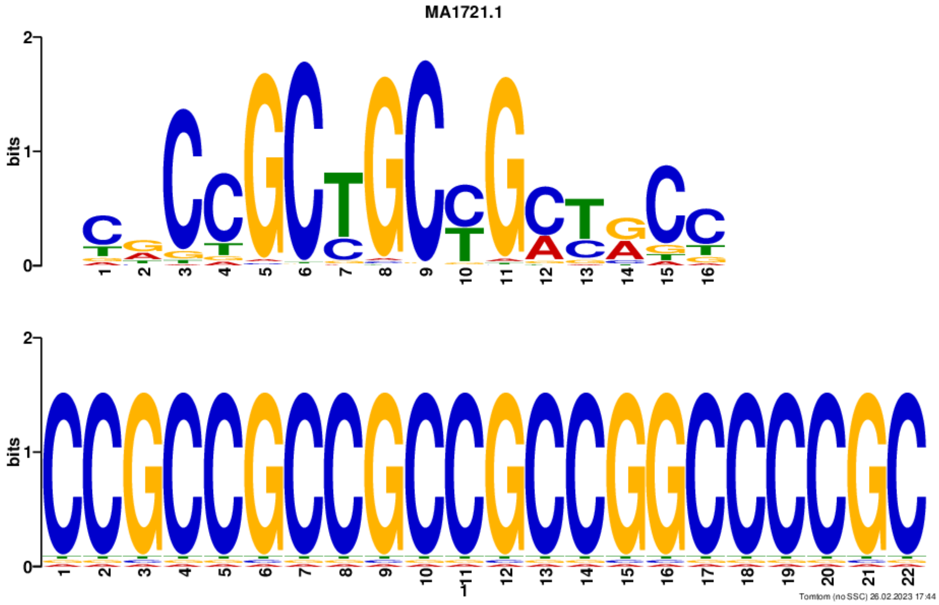


The likely causal motif CCGCCGCCGCCGCCGGCCCCGC (corresponding to the red repeat unit in Fig. 6) was aligned against the vertebrate (in *silico* and in *vivo*) database using TOMTOM with default parameters. The top hit is shown, a ZNF93 binding motif (JASPAR ID: MA1721.1) with p-value = 8.98e-07, E-value = 1.86e-03, and q-value = 3.68e-03.

**Supplemental Fig. S31.** Correlation between the number of predicted ZNF93 binding sites and *RNF213* VNTR variants.

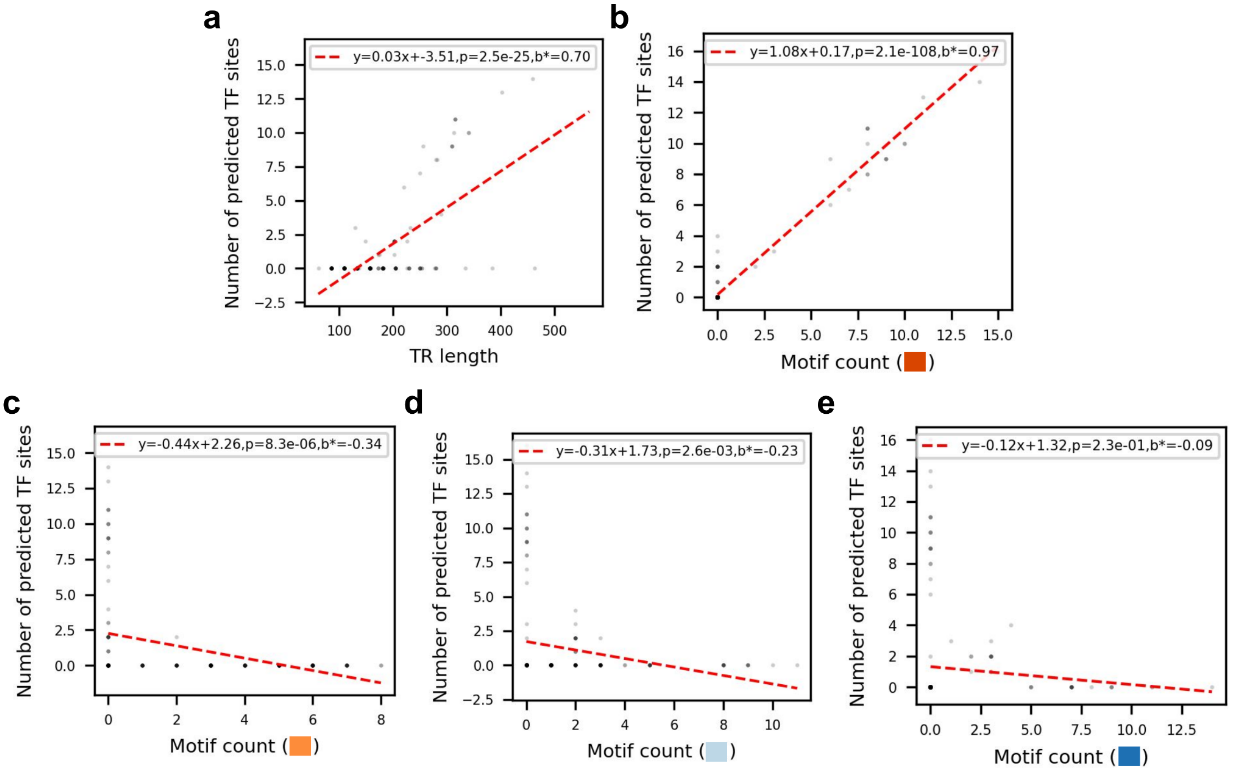


ZNF93 binding sites were predicted using FIMO with the MA1721.1 matrix and a cutoff of P<1e-8. Repeat length (a) and repeat unit counts for each of the efficient motifs annotated by vamos (b-e) are shown . Data points are collected from 35 HGSVC and 48 HPRC assemblies.

**Supplemental Fig. S32.** Predicted ZNF93 binding sites for each VNTR motif.

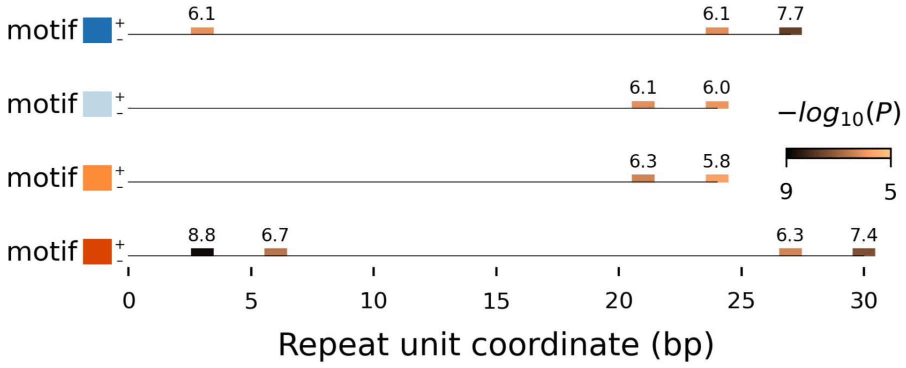


ZNF93 binding sites (MA1721.1) across the four efficient motifs annotated by vamos were predicted using FIMO with a cutoff of P < $10^{-5}$. To detect binding sites that span the boundary of repeat units, two-copy units were taken as inputs. Only one repeat unit is shown such that periodic binding sites are removed. The significance ($-log_{10}P$) of the match is shown above each binding site.

**Supplemental Fig. S33.** Correlation between *RNF213* expression level and *RNF213* VNTR repeat units.

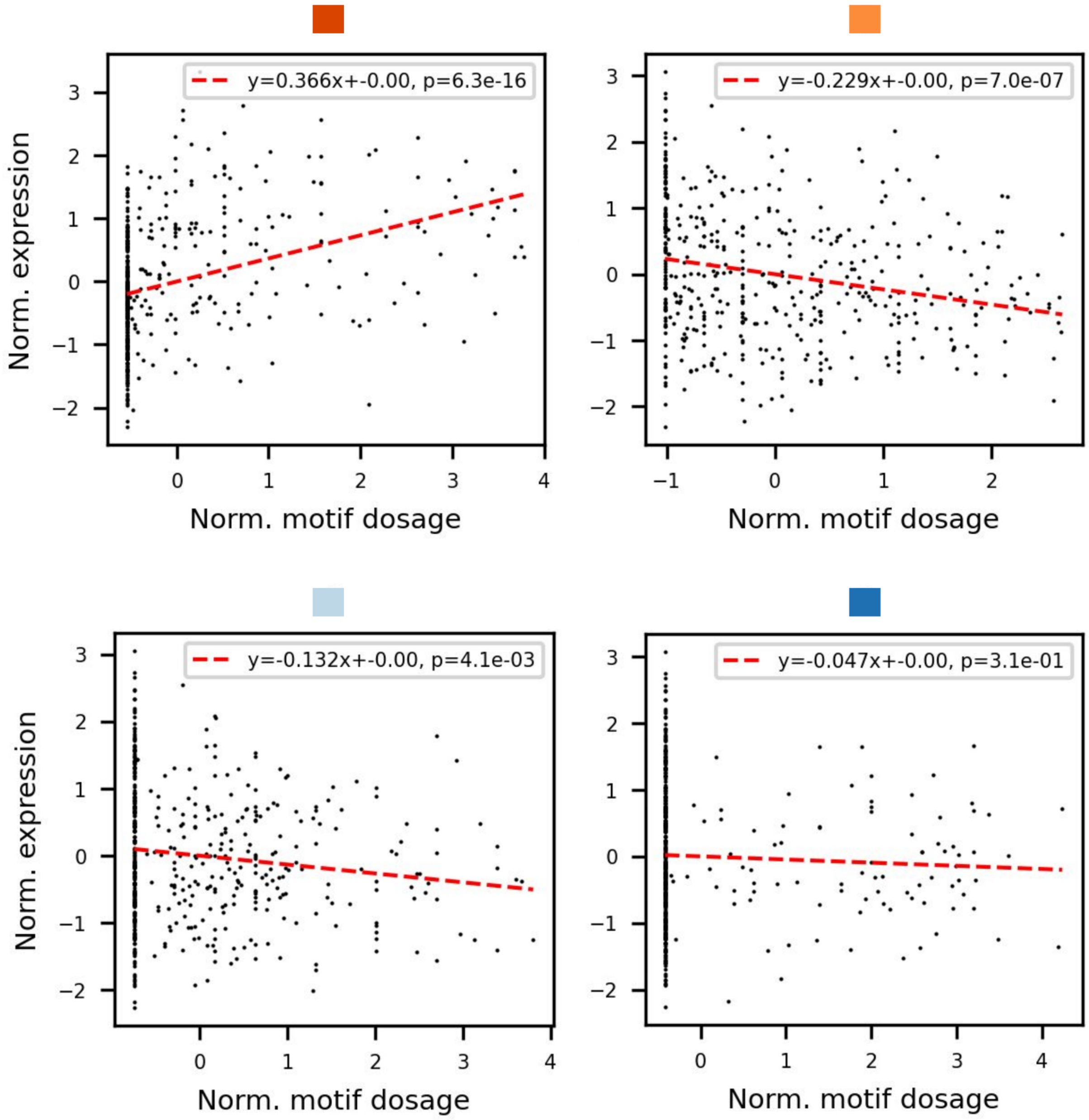


Each *RNF213* VNTR motif in the locus-RPGG that has a unique match with a two-copy unit (among the four repeat units annotated by vamos) was assigned as representative of the repeat unit. The VNTR motif with the lowest P-value in explaining gene expression in thyroid is shown for each repeat unit.

#### Supplemental Tables

**Supplemental Table S1.** Additional disease-relevant tandem repeats included in our reference set.

| Gene | Associated/Causal TR Region | Disease | Note |
|---|---|---|---|
| ACAN | chr15:88855423-88857301 | OSTD | coding |
| AR | chrX:67545317-67545419 | SBMA | coding |
| ATN1 | chr12:6936717-6936775 | DRPLA | coding |
| ATXN1 | chr6:16327634-16327724 | SCA1 | coding |
| ATXN2 | chr12:111598950-111599019 | SCA2 | coding |
| ATXN3 | chr14:92071011-92071052 | SCA3 | coding |
| ATXN7 | chr3:63912685-63912716 | SCA7 | coding |
| ATXN8 | chr13:70,139,352-70,139,429 | SCA8 | noncoding |
| AVPR1A | chr12:63149772-63149849 | EXTB | intron_AVR |
| AVPR1A | chr12:63153304-63153366 | EXTB | 5UTR_RS1 |
| AVPR1A | chr12:63156354-63156429 | EXTB | 5UTR_RS3 |
| C9orf72 | chr9:27573485-27573546 | ALS | intron 1 |
| CACNA1A | chr19:13207859-13207898 | SCA6 | coding |
| CACNA1C | chr12:2255791-2256090 | bipolar, SCZ | intron 3 |
| CEL | chr9:133071170-133071694 | Monogenic diabetes | coding |
| CNBP | chr3:129172568-129172736 | DM2 | intron 1 |
| CSTB | chr21:43776293-43776322 | PMP1A | 5' UTR |
| DMPK | chr19:45770205-45770266 | DM1 | 3' UTR |
| DRD4 | chr11:639989-640194 | OCD, ADHD | coding |
| FMR1 | chrX:147912037-147912111 | FXS, FXTAS | 5' UTR |
| FXN | chr9:69087917-69087957 | FA | noncoding |
| GP1BA | chr17:4933823-4933963 | ATF in stroke | coding |
| HIC1 | chr17:2052378-2053079 | MCC | promoter |
| HTT | chr4:3074877-3074940 | HD | coding |
| IL1RN | chr2:113130529-113130872 | stroke, CAD | intron |

| | | | |
|---|---|---|---|
| INS | chr11:2161570-2161976 | T1D, T2D, Obesity | promoter |
| JPH3 | chr16:87604283-87604329 | HDL2 | intron |
| MAOA | chrX:43655101-43655205 | bipolar | promoter |
| MMP9 | chr20:46013363-46013426 | Kawasaki | coding |
| MUC1 | chr1:155188676-155192051 | MCKD1 | coding |
| MUC21 | chr6:30986289-30987497 | DPB | coding |
| NACA | chr12:56717496-56718739 | atrial fibrillation | coding |
| PER3 | chr1:7829888-7830126 | bipolar | coding |
| PPP2R2B | chr5:146878728-146878759 | SCA12 | noncoding |
| SLC6A3 | chr5:1393582-1393985 | ADHD, PK | 3' UTR |
| SLC6A3 | chr5:1414387-1414518 | ADHD, PK | intron 8 |
| SLC6A4 | chr17:30221385-30221590 | BPSD, AZ | intron |
| SLC6A4 | chr17:30237128-30237450 | OCD, anxiety, SCZ | promoter |
| TBP | chr6:170561907-170562017 | SCA17 | coding |
| TCHH | chr1:152111363-152112285 | Male pattern baldness score | coding |

ALS, amyotrophic lateral sclerosis. DM, myotonic dystrophy. DRPLA, dentatorubral-pallidoluysian atrophy. FTD, frontotemporal dementia. FXS, fragile X syndrome. FXTAS, fragile X tremor-ataxia syndrome. HD, Huntington disease. HDL2, Huntington disease-like 2. SBMA, spinobulbar muscular atrophy. SCA, spinocerebellar ataxia. SCZ, schizophrenia. FA, Friedreich ataxia. DPB, respiratory bronchioles, causing a progressive suppurative and severe obstructive respiratory disorder . AZ, Alzheimer’s disease. MCC, metastatic colorectal cancer. PMP1A, progressive myoclonic epilepsy 1A. PK, Parkinson’s disease. OSTD, osteochondritis dissecans. EXTB, externalizing behaviors. CAD, coronary artery disease.

**Supplemental Table S2.** eMotif discoveries for disease-relevant genes.

| Gene | eGene | | | eGene-eTR pair | | | |
|---|---|---|---|---|---|---|---|
| | Rep.[1] | # of tissues | # of eTRs | Rep[2] | Tissue | Disease match[3] | eMotif (positive/negative effect)[4] |
| PER3 | v | 3 | 1 | v | Cells_Cultured_fibroblasts<br>Whole_Blood | | |
| TCHH | v | 11 | 1 | | | | |
| MUC1 | v | 6 | 1 | v | Artery_Tibial<br>Cells_Cultured_fibroblasts<br>Esophagus_Mucosa<br>Whole_Blood | ? | -0.17:CCACGGTGTCACCTCGGCCCC<br>-0.17:ACACCAGGCCGGCCCCGGGCT<br>-0.17:GCCCACGGTGTCACCTCGGCC<br>-0.16:CCCACGGTGTCACCTCGGCCC<br>-0.16:CAGCCCACGGTGTCACCTCGG<br>-0.16:CACGGTGTCACCTCGGCCCCG<br>-0.16:CACCAGGCCGGCCCCGGGCTCCA<br>-0.16:CCGGCCTGGTGTCCGGGGCCG<br>-0.16:AGCCCACGGTGTCACCTCGGC<br>-0.16:ACCTCGGCCCCGGACACCAGG<br>-0.16:GACACCAGGCCGGCCCCGGGC<br>-0.16:GTGTCCGGGGCCGAGGTGACA<br>-0.16:CCGGGGCCGGCCTGGTGTCCG<br>-0.16:CTGGTGTCCGGGGCCGAGGTGAC<br>-0.16:CCCGGGGCCGGCCTGGTGTCC<br>-0.16:GCCCCGGACACCAGGCCGGCC<br>-0.16:ACGGTGTCACCTCGGCCCCGG<br>-0.16:GTGTCACCTCGGCCCCGGACA<br>-0.16:CGGTGTCACCTCGGCCCCGGA<br>-0.15:CGGCCTGGTGTCCGGGGCCGA<br>-0.15:GGTGTCACCTCGGCCCCGGAC<br>-0.15:CCTCGGCCCCGGACACCAGGC<br>-0.15:CCGGACACCAGGCCGGCCCCG<br>-0.15:CCAGCCCACGGTGTCACCTCG<br>-0.15:CGGCCCCGGGCTCCACCGCCC<br>-0.15:CCCGGACACCAGGCCGGCCCC<br>-0.15:CTCGGCCCCGGACACCAGGCC<br>-0.15:AGGCCGGCCCCGGGCTCCACC<br>-0.15:CCGGCCCCGGGCTCCACCGCC<br>-0.15:GCCGGCCTGGTGTCCGGGGCC<br>-0.15:GCCGGCCCCGGGCTCCACCGC<br>-0.15:CAGGCCGGCCCCGGGCTCCAC<br>-0.15:CCCAGCCCACGGTGTCACCTC<br>-0.14:AGGTGACACCGTGGGCTGGGG<br>-0.14:CCCCCAGCCCACGGTGTCACC<br>-0.14:CCCCGGACACCAGGCCGGCCC<br>-0.14:CGGTGGAGCCCGGGGCCGGCC<br>-0.14:CTCCACCGCCCCCCCAGCCCA |
| DRD4 | v | 28 | 1 | v | Colon_Transverse<br>Esophagus_Mucosa<br>Esophagus_Muscularis<br>Nerve_Tibial<br>Skin_Not_Sun_Exposed_Suprapubic<br>Testis | | |
| INS | v | 7 | 1 | | | | |
| CACNA1C | v | 16 | 1 | v | Brain_Cerebellar_Hemisphere<br>Brain_Cerebellum<br>Cells_Cultured_fibroblasts<br>Colon_Transverse<br>Esophagus_Muscularis | v | -0.45:AAGGTCAGGATCGTGTGGTTG<br>-0.41:CCCTGACCTTACTAGTTTACGA<br>-0.18:ATCACACGATCCTGACCTGAC<br>-0.18:AATCACACGATCCTGACCTGA<br>-0.17:CACACGATCCTGACCTGACTA<br>-0.17:ACACGATCCTGACCTGACTAG<br>-0.17:AAACTAGTCAGGTCAGGATCGTG<br>-0.17:TAGTCAGGTCAGGGTCGTGTAATTGTAAACTAGTCAGGTCA<br>-0.17:ATCCTGACCTGACTAGTTTAC<br>-0.17:CAACCACACGATCCTGACCTGA<br>-0.17:CTGACCTTACTAGTTTACAACCACACGA<br>-0.17:GATCCTGACCTGACTAGTTTA<br>-0.17:CAATCACACGATCCTGACCTG<br>-0.17:ACCACACGATCCTGACCTGAC<br>-0.16:TCCTGACCTGACTAGTTTACA<br>-0.16:CTGACTAGTTTACAATCACAC<br>-0.16:ACCTGACTAGTTTACAATCAC<br>-0.16:ATTGTAAACTAGTCAGGTCAG<br>-0.16:GATTGTAAACTAGTCAGGTCA<br>-0.16:CCTGACTAGTTTACAATCACA |

| | | | | | | | |
|---|---|---|---|---|---|---|---|
| | | | | | | | -0.16:GACCTGACTAGTTTACAATCA<br>-0.16:AGTCAGGTCAGGATCGTGTGG<br>-0.16:AGTCAGGTCAGGATCGTGTGA<br>-0.16:CGTGTGGTTGTAAACTAGTCA<br>-0.16:GACTAGTTTACAACCACACGA<br>-0.16:CAGGTCAGGATCGTGTGATTCTAAAC<br>-0.15:AAGGTCAGGATCGTGTGATTTTAAACTAGCAAGGTCAGGAT<br>-0.15:GACCTGACTAGTTTACAACCA<br>-0.15:GGTTGTAAACTAGTCAGGTCA<br>-0.15:ACCTGACTAGTTTACAACCAC<br>-0.15:CTGACCTGACTAGTTTACAAC<br>-0.15:ACCACACGACCCTGACCTGACT<br>-0.15:ACTAGTTTACAATCACAGGAT<br>+0.16:AGGATCGTGTGGTTGTAAACCAG<br>+0.18:AAACTAGCAAGGTCAGGGTCTTGTGATTG<br>+0.19:CAAGGTCAGGATCGTGTGGTCG<br>+0.21:CTTGCTAGTTTACAATCACAAGAC<br>+0.24:ACCCTGACTTGACTAGTTTACAATCACAC |
| ATN1 | v | 33 | 1 | v | Adipose_Subcutaneous<br>Adipose_Visceral_Omentum<br>Artery_Aorta<br>Artery_Coronary<br>Artery_Tibial<br>Brain_Cerebellar_Hemisphere<br>Brain_Cerebellum<br>Brain_Cortex<br>Breast_Mammary_Tissue<br>Cells_Cultured_fibroblasts<br>Colon_Sigmoid<br>Colon_Transverse<br>Esophagus_Gastroesophageal_Junction<br>Esophagus_Mucosa<br>Esophagus_Muscularis<br>Heart_Atrial_Appendage<br>Heart_Left_Ventricle<br>Liver<br>Lung<br>Muscle_Skeletal<br>Nerve_Tibial<br>Ovary<br>Pancreas<br>Pituitary<br>Prostate<br>Skin_Not_Sun_Exposed_Suprapubic<br>Skin_Sun_Exposed_Lower_leg<br>Small_Intestine_Terminal_Ileum<br>Spleen<br>Stomach<br>Testis<br>Thyroid<br>Vagina | v | +0.59:CAGCAGCAGCAGCAGCAGCA |
| NACA | v | 1 | 1 | | | | |
| AVPR1A | v | 5 | 3 | v | Adipose_Subcutaneous<br>Cells_Cultured_fibroblasts | | |
| | | | | v | Artery_Tibial | | |
| | | | | v | Artery_Coronary<br>Artery_Tibial | | |
| ATXN2 | v | 1 | 1 | | | | |
| ATXN8 | | 0 | 0 | | | | |
| ATXN3 | | 0 | 0 | | | | |
| ACAN | v | 3 | 1 | | | | |
| JPH3 | v | 9 | 1 | v | Testis | | |
| HIC1 | v | 2 | 1 | | | | |
| GP1BA | v | 17 | 1 | | | | |
| SLC6A4 | v | 2 | 2 | v | Esophagus_Muscularis<br>Nerve_Tibial | | |
| SLC6A4 | v | 2 | 2 | | | | |

| | | | | | | | |
|---|---|---|---|---|---|---|---|
| CACNA1A | v | 11 | 1 | | | | |
| DMPK | | 0 | 0 | | | | |
| IL1RN | v | 9 | 1 | v | Breast_Mammary_Tissue<br>Cells_Cultured_fibroblasts<br>Skin_Not_Sun_Exposed_Suprapubic<br>Skin_Sun_Exposed_Lower_leg | | |
| MMP9 | v | 2 | 1 | | | | |
| CSTB | v | 39 | 1 | | | | |
| ATXN7 | v | 9 | 1 | v | Brain_Cerebellar_Hemisphere<br>Esophagus_Muscularis<br>Testis | v | -0.33:GCAGCAGCAGCAGCAGCAGCAGC |
| CNBP | | 0 | 0 | | | | |
| HTT | v | 13 | 1 | v | Muscle_Skeletal | | |
| SLC6A3 | v | 17 | 2 | | | | |
| PPP2R2B | v | 12 | 1 | | | | |
| ATXN1 | v | 2 | 1 | | | | |
| MUC21 | v | 5 | 1 | v | Esophagus_Mucosa<br>Lung<br>Nerve_Tibial | v | -0.21:CACAGCCACCAACTCTGACTCCAGCACA<br>-0.20:ACCTCCAGTGGGGCTAGCACAGCCACCAACTCTGA<br>-0.20:CAGCACAACCTCCAGTGGGGTCAGCAC<br>-0.20:ACTGCCACCAACTCTGAGTCTAG<br>-0.20:CCAACTCTGAGTCTAGCACACTCTCCAGTGGGGCCAGCACA<br>-0.19:GACTCAGAGTTGGTGGCAGTGCTGGC<br>-0.19:GCCACCAACTCTGAGTCTAGCACA<br>-0.19:AGCACAACCTCCAGTGGGGCTAGCAC<br>-0.19:ACAGCCACCAACTCTGAGTCTAG<br>-0.19:GGGGCCAGCACTGCCACCAAC<br>-0.19:ATCCCACTGGACACTGTGCTAGACTCAGAGTTGG<br>-0.18:ACTGCCACCAACTCTGAGTCCAG<br>-0.18:AGCACAGCCACCAACTCTGACT<br>-0.18:GAGTTGGTGGCAGTGCTGGCCC<br>-0.17:CTGTGCTGGCCCCACTGGAGA<br>-0.15:ACCTCCAGTGGGGCCGGCACA<br>+0.15:ACCCAGAGTTGGTGGCTGTGCTG<br>+0.16:AGGTCGTGCTGGACTCAGAGGTGGTGGCTGTGCTGGCCCCA<br>+0.16:CAGAGTTGGTGACTGTGCTGGCCCCACTGGAG<br>+0.17:CACAACCTCCAGTGGGGCCAGCAC |
| TBP | v | 8 | 1 | | | | |
| C9orf72 | v | 35 | 1 | v | Adipose_Subcutaneous<br>Adipose_Visceral_Omentum<br>Artery_Aorta<br>Artery_Tibial<br>Brain_Anterior_cingulate_cortex_BA24<br>Brain_Caudate_basal_ganglia<br>Brain_Cerebellar_Hemisphere<br>Brain_Cerebellum<br>Brain_Cortex<br>Brain_Frontal_Cortex_BA9<br>Brain_Nucleus_accumbens_basal_ganglia<br>Brain_Putamen_basal_ganglia<br>Breast_Mammary_Tissue<br>Colon_Sigmoid<br>Colon_Transverse<br>Esophagus_Gastroesophageal_Junction<br>Esophagus_Mucosa<br>Esophagus_Muscularis<br>Heart_Atrial_Appendage<br>Heart_Left_Ventricle<br>Lung<br>Muscle_Skeletal<br>Nerve_Tibial<br>Pancreas<br>Pituitary<br>Prostate<br>Skin_Not_Sun_Exposed_Suprapubic<br>Skin_Sun_Exposed_Lower_leg<br>Small_Intestine_Terminal_Ileum | v | -0.22:CCGCCCCGGGCCCGCCCCCGGGCCCGCCCCGACCACGCCCCGGCCCCGGCCCC<br>+0.45:CCGGCCCCGGCCCCGGCCCCGGC<br>+0.46:CCCCGGCCCCGGCCCCGGCCCC<br>+0.46:CACGCCCCGGCCCCGGCCCCGGC<br>+0.46:GCCCCGGCCCCGGCCCCGGCC |

| | | | | | | | |
|---|---|---|---|---|---|---|---|
| | | | | | Spleen<br>Stomach<br>Testis<br>Thyroid<br>Whole_Blood | | |
| FXN | v | 6 | 1 | | | | |
| CEL | v | 5 | 1 | v | Brain_Hypothalamus<br>Pancreas | v | +0.20:CCCGGAGTCACCCGTGGGCGG<br>+0.29:ACTCCGGGGCCCCCCCTGTGACCCCCACGGGTGACTC |
| MAOA | v | 4 | 1 | v | Muscle_Skeletal<br>Thyroid<br>Whole_Blood | ? | -0.23:CCAGTACCGGCACCGGCACCAGTAC<br>-0.18:CCGGCACCGGCACCAGTACCC<br>-0.18:ACCGGCACCGGCACCAGTACC<br>-0.17:ACTGGTGCGGGTACTGGTGCCGGTGCCG<br>-0.17:CCCGCACCAGTACCGGCACCGGCACC<br>-0.16:GCACCAGTACCCGCACCAGTACC<br>-0.15:CCAGTACCCGCACCAGTACCGGCACC |
| AR | v | 2 | 1 | | | | |
| FMR1 | v | 2 | 1 | | | | |

[1] eGene replicated in eMotif discoveries using genome-wide P-value cutoffs.
[2] eGene-eTR pair replicated in eMotif discoveries using genome-wide P-value cutoffs. For eGene with multiple disease-relevant repeats, the order shown here is the same as the one listed in Table S1.
[3] The eGene-eTR tissue matches disease ontology. The ”?” denotes cases where the exact disease mechanism is unclear.
[4] Only eMotifs from eGene-eTR pairs matching disease ontology are shown. The largest effect size for each unique motif is shown.

**Supplemental Table S3.** eQTL mapping using a genome-wide P-value cutoff.

| Tissue | P-value cutoff* |
|---|---|
| Kidney_Cortex | 2.86E-05 |
| Brain_Substantia_nigra | 6.64E-05 |
| Vagina | 9.62E-05 |
| Brain_Amygdala | 9.64E-05 |
| Uterus | 1.06E-04 |
| Cells_EBV-transformed_lymphocytes | 1.25E-04 |
| Brain_Spinal_cord_cervical_c-1 | 1.36E-04 |
| Minor_Salivary_Gland | 1.57E-04 |
| Brain_Hippocampus | 1.83E-04 |
| Brain_Hypothalamus | 1.99E-04 |
| Brain_Anterior_cingulate_cortex_BA24 | 2.03E-04 |
| Small_Intestine_Terminal_Ileum | 2.05E-04 |
| Ovary | 2.19E-04 |

| | |
|---|---|
| Brain_Putamen_basal_ganglia | 2.60E-04 |
| Artery_Coronary | 2.79E-04 |
| Liver | 2.80E-04 |
| Brain_Frontal_Cortex_BA9 | 2.84E-04 |
| Prostate | 3.37E-04 |
| Brain_Nucleus_accumbens_basal_ganglia | 3.44E-04 |
| Brain_Caudate_basal_ganglia | 3.46E-04 |
| Adrenal_Gland | 4.10E-04 |
| Brain_Cerebellar_Hemisphere | 4.46E-04 |
| Brain_Cortex | 4.51E-04 |
| Pituitary | 4.87E-04 |
| Spleen | 5.20E-04 |
| Stomach | 5.29E-04 |
| Pancreas | 6.43E-04 |
| Brain_Cerebellum | 6.49E-04 |
| Colon_Transverse | 6.78E-04 |
| Colon_Sigmoid | 6.79E-04 |
| Breast_Mammary_Tissue | 7.13E-04 |
| Esophagus_Gastroesophageal_Junction | 7.34E-04 |
| Heart_Left_Ventricle | 7.74E-04 |
| Heart_Atrial_Appendage | 8.11E-04 |
| Artery_Aorta | 9.59E-04 |
| Testis | 9.69E-04 |
| Adipose_Visceral_Omentum | 9.74E-04 |
| Lung | 1.05E-03 |
| Skin_Not_Sun_Exposed_Suprapubic | 1.23E-03 |
| Esophagus_Mucosa | 1.25E-03 |
| Esophagus_Muscularis | 1.26E-03 |
| Adipose_Subcutaneous | 1.34E-03 |
| Artery_Tibial | 1.44E-03 |
| Cells_Cultured_fibroblasts | 1.46E-03 |

| | |
|---|---|
| Muscle_Skeletal | 1.47E-03 |
| Skin_Sun_Exposed_Lower_leg | 1.48E-03 |
| Nerve_Tibial | 1.58E-03 |
| Whole_Blood | 1.59E-03 |
| Thyroid | 1.71E-03 |

* False discovery rate controlled using the Benjamini–Hochberg procedure taking all P-values from all tests as input. The genome-wide P-value cutoff  was used to allow for more than one eMotif to be reported as significantly associated with a gene.

#### References


Bailey TL, Gribskov M. 1998. Combining evidence using p-values: application to sequence homology searches. *Bioinformatics* **14**: 48–54.

Open2C, Abdennur N, Abraham S, Fudenberg G, Flyamer IM, Galitsyna AA, Goloborodko A, Imakaev M, Oksuz BA, Venev SV. 2022. Cooltools: enabling high-resolution Hi-C analysis in Python. *bioRxiv* 2022.10.31.514564. https://www.biorxiv.org/content/10.1101/2022.10.31.514564v1.abstract (Accessed February 28, 2023).
