## Supplemental code for "The motif composition of variable-number tandem repeats impacts gene expression": Lu2023_akita_preds_environment.yml

```
name: basenji-numba2
channels:
  - defaults
  - conda-forge
  - bioconda
  - nvidia
dependencies:
  - astropy
  - cudatoolkit=11
  - cudnn=8
  - cython
  - h5py
  - intervaltree
  - joblib
  - matplotlib
  - natsort
  - networkx
  - numpy=1.19
  - numba=0.53
  - packaging
  - pandas
  - pillow
  - pip
  - pybedtools
  - pybigwig
  - pysam
  - python==3.8
  - scikit-learn
  - scipy
  - seaborn
  - statsmodels
  - tensorflow-gpu=2.6
  - tabulate
  - pip:
    - cooler
    - cooltools
```
