## Supplemental code for "The motif composition of variable-number tandem repeats impacts gene expression": Lu2023_akita_preds_functions.py

```
import matplotlib.pyplot as plt
import seaborn as sns
import numpy as np
import h5py
import pandas as pd


def read_fasta(fasta_file):

    """
    Reads the fasta file with haplotypes sequences and extracts information necessary on further steps.

    Parameters
    ------------
    fasta_file : str
        A path to a fasta file.

    Returns
    ---------
    summary_df : DataFrame
        Pandas DataFrame collecting information about each haplotype.
    """

    sequences_dict = {"haplotyple_name": [], "length": []}

    seq_identifier = ""

    with open(fasta_file, "r") as fasta_input:
        for line in fasta_input:
            line = line.strip()
            if line[0] == ">":
                seq_identifier = line[1:]
                sequences_dict["haplotyple_name"].append(seq_identifier)
            else:
                seq_length = len(line[700:-700])
                # since sequences are provided with 700bp flanks

                sequences_dict["length"].append(seq_length)

    haplotyples_names = sequences_dict["haplotyple_name"]
    summary_df = pd.DataFrame.from_dict(sequences_dict)

    return summary_df


def average_experiments_multiple_models(
    paths_to_hf,
    haplotyples_names,
    head_index=0,
    pred_length=130305,
    target_indices_list=[0, 1, 2, 3, 4],
):

    """
    Returns a summary matrix where each row represents a single experiment.

    Parameters
    ------------
    paths_to_hf : list
        A list of paths to all h5 files with all the predictions saved.
    haplotyples_names : list
        List of haplotypes' IDs.
    head_index : int
        Index of a head used to get a prediction.
    pred_length : int
        Length of a predicted vector.
    target_indices_list : list
        List of targets' indices that predictions will be averaged over.

    Returns
    ---------
    summary_matrix : matrix
        number_of_haplotypes by pred_length matrix, where each row is a prediction vector averaged over all desired models and targets.
    """

    number_names = len(haplotyples_names)
    num_targets = len(target_indices_list)
    num_models = len(paths_to_hf)

    divider = num_models * num_targets

    summary_matrix = np.zeros((number_names, pred_length))

    for hf_path in paths_to_hf:
        print(hf_path + " - done")
        model_index = int(hf_path.split(".")[-2][-1])

        hf = h5py.File(hf_path, "r")

        for name_index in range(number_names):
            name = haplotyples_names[name_index]
            for target_index in target_indices_list:
                summary_matrix[name_index] += hf[
                    f"{name}_h{head_index}_m{model_index}_t{target_index}"
                ]

    summary_matrix = summary_matrix / divider
    return summary_matrix


def ut_dense(preds_ut, diagonal_offset):

    """
    Constructs symmetric dense prediction matrices from upper triangular vectors.

    Parameters
    -----------
    preds_ut : ( M x O) numpy array
        Upper triangular matrix to convert. M is the number of upper triangular entries,
        and O corresponds to the number of different targets.
    diagonal_offset : int
        Number of diagonals that are added as zeros in the conversion.
        Typically 2 diagonals are ignored in Hi-C data processing.

    Returns
    --------
    preds_dense : (D x D x O) numpy array
        Each output upper-triangular vector is converted to a symmetric D x D matrix.
        Output matrices have zeros at the diagonal for `diagonal_offset` number of diagonals.
    """

    ut_len, num_targets = preds_ut.shape

    # infer original sequence length
    seq_len = int(np.sqrt(2 * ut_len + 0.25) - 0.5)
    seq_len += diagonal_offset

    # get triu indexes
    ut_indexes = np.triu_indices(seq_len, diagonal_offset)
    assert len(ut_indexes[0]) == ut_len

    # assign to dense matrix
    preds_dense = np.zeros(
        shape=(seq_len, seq_len, num_targets), dtype=preds_ut.dtype
    )
    preds_dense[ut_indexes] = preds_ut

    # symmetrize
    preds_dense += np.transpose(preds_dense, axes=[1, 0, 2])

    return preds_dense


def get_map(predicted_vector, diagonal_offset=2):

    """
    Creates a 512x512 map representating changes in the DNA contacts.

    Parameters
    ------------
    predicted_vector : numpy vector
        Akita's output.
    diagonal_offset : int
        Number of diagonals that are added as zeros in the conversion.
        Typically 2 diagonals are ignored in Hi-C data processing.

    Returns
    ---------
    matrix : numpy matrix
        512x512 map
    """

    matrix = ut_dense(np.expand_dims(predicted_vector, 1), diagonal_offset)
    matrix = np.squeeze(matrix, axis=2)

    return matrix


def triple_plot(
    summary_matrix,
    summary_df,
    haplotyple_name,
    reference_length,
    diff_window_start,
    bin_size=2048,
    ref_index=0,
    vmin=-0.6,
    vmax=0.6,
    to_save=False,
    resolution_dpi=300,
    headers_size=11,
    name_rotation=90,
    name_fontsize=11,
    name_labelpad=10,
    tick_size=8,
):

    """
    Plots a triple of maps for a given haplotyple_name:
    (1) predicted changes in the DNA contacts, (2) changes in the DNA contacts in the reference (both maps are adjusted to a possible translation),
    (3) difference map: predicted changes - reference changes.

    Parameters
    ------------
    summary_matrix : numpy array
        Changes in DNA contacts for all haplotyples.
    summary_df : DataFrame
        Pandas DataFrame collecting information about each haplotype.
    haplotyple_name : str
        Name of a desired haplotype.
    reference_length : int
        Length of a reference haplotype.
    diff_window_start : int
        A relative start of the subsequence that may differ between haplotypes and a reference.
    bin_size : int
        The length of each bin.
    ref_index : int
        Index of a reference sequence in the summary_matrix.
    vmin : float
    vmax : float
        Minimum and maximum in the colormap scale.
    to_save : boolean
        True if an image is supposed to be saved.
    resolution : int
        Image's desired resolution.
    headers_size : int
        Fontsize of the text above each plot.
    name_rotation : int
        Rotation applied to the y-axis text being the haplotyple_name.
    name_fontsize : int
        Fontsize of the y-axis text being the haplotyple_name.
    name_labelpad : int
        Labelpad applied to the y-axis text being the haplotyple_name.
    tick_size : int
        Fontsize of the colorbar values.
    """

    haplotype_index = np.where(
        summary_df["haplotyple_name"] == haplotyple_name
    )[0][0]
    difference_length = (
        int(
            summary_df[summary_df["haplotyple_name"] == haplotyple_name][
                "length"
            ]
        )
        - reference_length
    )

    map_matrix = get_map(summary_matrix[haplotype_index])
    ref_matrix = get_map(summary_matrix[ref_index])

    if abs(difference_length) > bin_size:
        # we need to adjust to translation

        if difference_length > 0:
            # this corresponds to an insertion relative to the reference

            bins_overlapped = get_bins(
                diff_window_start, diff_window_start + difference_length
            )
            ref_matrix = insert_nan_bin(ref_matrix, bins_overlapped)
            map_matrix = fill_bin_nans(map_matrix, bins_overlapped)

            adjusted_ref_matrix = mask_nans(ref_matrix)
            adjusted_map_matrix = mask_nans(map_matrix)

            difference_matrix = map_matrix - ref_matrix

        else:
            # this corresponds to a deletion relative to the reference

            bins_overlapped = get_bins(
                diff_window_start + difference_length, diff_window_start
            )
            ref_matrix = fill_bin_nans(ref_matrix, bins_overlapped)
            map_matrix = insert_nan_bin(map_matrix, bins_overlapped)

            # in both matrices nans have to be masked
            adjusted_ref_matrix = mask_nans(ref_matrix)
            adjusted_map_matrix = mask_nans(map_matrix)

            difference_matrix = map_matrix - ref_matrix

    else:
        difference_matrix = map_matrix - ref_matrix

    fig, axs = plt.subplots(1, 3, figsize=(12, 4))

    # plotting the predicted matrix
    sns.heatmap(
        map_matrix,
        ax=axs[0],
        center=0,
        vmin=vmin,
        vmax=vmax,
        cbar=False,
        cmap="RdBu_r",
        square=True,
        xticklabels=False,
        yticklabels=False,
    )

    # plotting the reference matrix
    sns.heatmap(
        ref_matrix,
        ax=axs[1],
        center=0,
        vmin=vmin,
        vmax=vmax,
        cbar=False,
        cmap="RdBu_r",
        square=True,
        xticklabels=False,
        yticklabels=False,
    )

    # plotting the difference matrix
    sns.heatmap(
        difference_matrix,
        ax=axs[2],
        center=0,
        vmin=vmin,
        vmax=vmax,
        cbar=False,
        cmap="RdBu_r",
        square=True,
        xticklabels=False,
        yticklabels=False,
    )

    cols = ["prediction", "reference", "difference"]

    for ax, col in zip(axs, cols):
        ax.set_title(col, size=headers_size)

    ax = axs[0]
    ax.set_ylabel(
        f"{haplotyple_name}",
        rotation=name_rotation,
        fontsize=name_fontsize,
        labelpad=name_labelpad,
    )

    a = np.array([[vmin, vmax]])
    img = plt.imshow(a, cmap="RdBu_r")

    cb_ax = fig.add_axes([0.915, 0.130, 0.015, 0.74])
    cbar = fig.colorbar(img, orientation="vertical", cax=cb_ax)
    for t in cbar.ax.get_yticklabels():
        t.set_fontsize(tick_size)

    if to_save:
        plt.savefig(f"{haplotyple_name}_triple_map.png", dpi=resolution_dpi)

    plt.show()


def get_bins(
    window_start,
    window_end,
    number_bins=512,
    bin_size=2048,
    input_size=1310720,
    target_crop=131072,
    min_bin_coverage=0.50,
):

    """
    Returns a list of bins overlapping in at least min_bin_coverage fraction the given window.

    Parameters
    ------------
    window_start : int
    window_end : int
        Start and end of the window that we aim to find a set of overlapped bins.
        Note, those values are relative.
    number_bins : int
        Number of bins used to create a map.
    bin_size : int
        The length of each bin.
    input_size : int
        Length of model's input sequence.
    target_crop : int
        Length of the sequence cropped from each side.
    min_bin_coverage : float
        If the given window is bigger that bin_size, only those bins whose overlap is greater than (min_bin_coverage * bin_size)
        will be outputted.

    Returns
    ---------
    bin_numbers : list
        A list of bins overlapping the given window.
    """

    window_size = window_end - window_start
    min_bin_length = int(min_bin_coverage * bin_size)

    assert (
        window_size > min_bin_length
    ), "there is no bin covered by the given sequence in more that min_bin_coverage"

    corrected_window_start = window_start - target_crop
    corrected_window_end = window_end - target_crop

    first_bin_covered = corrected_window_start // bin_size
    last_bin_covered = corrected_window_end // bin_size

    relative_start = corrected_window_start % bin_size
    relative_end = corrected_window_end % bin_size

    if window_size < bin_size:
        # return this bin that overlaps in higher proportion
        first_bin_cov = bin_size - relative_start
        second_bin_cov = relative_end
        if first_bin_cov > second_bin_cov:
            return [first_bin_covered]
        else:
            return [last_bin_covered]

    if relative_start > min_bin_length:
        first_bin_covered += 1

    if relative_end < min_bin_length:
        last_bin_covered -= 1

    assert (
        last_bin_covered >= first_bin_covered
    ), f"make sure that window_end > window_start"

    bin_numbers = [
        bin_number
        for bin_number in range(first_bin_covered, (last_bin_covered + 1))
    ]

    return bin_numbers


def mask_nans(matrix):
    """
    Masks bins filled in with nans.
    """

    masked_matrix = np.ma.array(matrix, mask=np.isnan(matrix))
    return masked_matrix


def fill_bin_nans(matrix, bin_list):

    """
    Returns a map with some columns and rows filled in with nans.

    Parameters
    ------------
    matrix : numpy array
        A 512x512 map representing differences in the DNA contacts.
    bin_list : list
        A list of bins corresponding to rows' and columns' indices that should be replaced with nans.

    Returns
    ---------
    naned_matrix : numpy array
        A map-metrix with some rows and columns replaced with nans.
    """

    naned_matrix = matrix.copy()

    bin_indices_list = [(bin_number - 1) for bin_number in bin_list]
    # since python counts from 0

    for bin_index in bin_indices_list:
        naned_matrix[:, bin_index] = np.nan
        naned_matrix[bin_index, :] = np.nan

    return naned_matrix


def insert_nan_bin(matrix, bin_list):

    """
    Returns a map with some rows and columns nans-inserted.

    Parameters
    ------------
    matrix : numpy array
        A 512x512 map representing differences in the DNA contacts.
    bin_list : list
        A list of bins corresponding to rows' and columns' indices that should be inserted with nans.

    Returns
    ---------
    translated_matrix : numpy array
        A map with some rows and columns nans-inserted. To keep the matrix the same size,
        the same number of columns and rows are deleted.
    """

    translated_matrix = matrix.copy()

    bin_indices_list = [(bin_number - 1) for bin_number in bin_list]

    for bin_index in bin_indices_list:
        translated_matrix = np.insert(
            translated_matrix, bin_index, np.nan, axis=1
        )
        translated_matrix = np.delete(translated_matrix, -1, 0)
        translated_matrix = np.insert(
            translated_matrix, bin_index, np.nan, axis=0
        )
        translated_matrix = np.delete(translated_matrix, -1, 1)

    return translated_matrix


def map_scd(matrix):

    """
    Returns an SCD-analogous metric calculated for a given map.
    SCD = squared contact differences

    Parameters
    ------------
    matrix : numpy array
        A 512x512 map representing differences in the DNA contacts.

    Returns
    ---------
    map_scd : float
        SCD metric
    """

    return np.sqrt(np.sum(np.square(matrix))) * (1 / 2)


def lds(matrix, square_size, nan_bins=[], matrix_size=512):

    """
    Returns a local disruption score (LDS) of a square window centered around the map's midpoint.

    Parameters
    ------------
    matrix : numpy array
        A 512x512 map representing differences in the DNA contacts.
    square_size : int
        A size of square centered around the map's midpoint.
    nan_bins : list
        A list of bins corresponding to rows' and columns' indices that were filled in with nans.
    matrix_size :int
        Size of an input matrix.

    Returns
    ---------
    lds : float
        Local disruption score of a central square.
    """

    central_bin = 512 // 2
    bin_start = (512 // 2) - (square_size // 2)

    left_expand = 0
    right_expand = 0

    # checking how the square_size should be exapanded
    if len(nan_bins) != 0:
        for nan_bin_index in nan_bins:
            if (nan_bin_index >= bin_start) and (nan_bin_index <= central_bin):
                left_expand += 1
            elif (nan_bin_index <= (bin_start + square_size)) and (
                nan_bin_index >= central_bin
            ):
                right_expand += 1

    central_matrix = matrix[
        (bin_start - left_expand) : (bin_start + square_size + right_expand),
        (bin_start - left_expand) : (bin_start + square_size + right_expand),
    ]
    lds = map_scd(central_matrix)
    return lds


def add_LDS(
    summary_df,
    summary_matrix,
    haplotyples_names,
    reference_length,
    diff_window_start,
    square_size,
    target_crop=131072,
    input_size=1310720,
    bin_size=2048,
    number_bins=512,
    diagonal_offset=2,
    ref_index=0,
):

    """
    Adds to the summary_df a new column "LDS" containing local disruption score for each haplotype.

    Parameters
    ------------
    summary_df : DataFrame
        Pandas DataFrame collecting information about each haplotype.
    summary_matrix : numpy array
        Changes in DNA contacts for all haplotyples.
    haplotyples_names : list
        List of haplotypes' IDs.
    reference_length : int
        Length of the reference haplotype.
    diff_window_start : int
        A relative start of the subsequence that may differ between haplotypes and a reference.
    square_size : int
        A desired size of a central square.
    target_crop : int
        Length of the sequence cropped from each side.
    input_size : int
        Length of model's input sequence.
    bin_size : int
        The length of each bin.
    number_bins : int
        Number of bins used to create a map.
    diagonal_offset : int
        Number of diagonals that are added as zeros in the conversion.
        Typically 2 diagonals are ignored in Hi-C data processing.
    ref_index : int
        Index of a reference sequence in the summary_matrix.

    Returns
    ---------
    summary_df : DataFrame
        An updated Pandas DataFrame collecting information about each haplotype.
    """

    ref_matrix = get_map(summary_matrix[ref_index])
    LDS = []

    for i in range(len(haplotyples_names)):

        name = haplotyples_names[i]
        difference_length = (
            int(summary_df.iloc[i]["length"]) - reference_length
        )

        map_matrix = get_map(summary_matrix[i], diagonal_offset)

        if abs(difference_length) > bin_size:
            # we need to adjust to translation

            if difference_length > 0:
                # this corresponds to an insertion relative to the reference

                bins_overlapped = get_bins(
                    diff_window_start,
                    diff_window_start + difference_length,
                    number_bins,
                    bin_size,
                    input_size,
                    target_crop,
                )

                adjusted_ref_matrix = ref_matrix.copy()
                adjusted_ref_matrix = insert_nan_bin(
                    adjusted_ref_matrix, bins_overlapped
                )
                adjusted_map_matrix = fill_bin_nans(
                    map_matrix, bins_overlapped
                )

                # in both matrices nans have to be masked
                adjusted_ref_matrix = mask_nans(adjusted_ref_matrix)
                adjusted_map_matrix = mask_nans(adjusted_map_matrix)

                difference_matrix = adjusted_map_matrix - adjusted_ref_matrix
                local_disruption_score = lds(
                    difference_matrix, square_size, nan_bins=bins_overlapped
                )
                LDS.append(local_disruption_score)

            else:
                # this corresponds to a deletion relative to the reference

                bins_overlapped = get_bins(
                    diff_window_start + difference_length,
                    diff_window_start,
                    number_bins,
                    bin_size,
                    input_size,
                    target_crop,
                )

                adjusted_ref_matrix = ref_matrix.copy()
                adjusted_ref_matrix = fill_bin_nans(
                    adjusted_ref_matrix, bins_overlapped
                )
                adjusted_map_matrix = insert_nan_bin(
                    map_matrix, bins_overlapped
                )

                # in both matrices nans have to be masked
                adjusted_ref_matrix = mask_nans(adjusted_ref_matrix)
                adjusted_map_matrix = mask_nans(adjusted_map_matrix)

                difference_matrix = adjusted_map_matrix - adjusted_ref_matrix
                local_disruption_score = lds(
                    difference_matrix, square_size, nan_bins=bins_overlapped
                )
                LDS.append(local_disruption_score)

        else:
            difference_matrix = map_matrix - ref_matrix
            local_disruption_score = lds(difference_matrix, square_size)
            LDS.append(local_disruption_score)

    summary_df["LDS"] = LDS

    return summary_df
```
