## Supplemental code for "The motif composition of variable-number tandem repeats impacts gene expression": Lu2023_akita_preds_notebook.ipynb

```
# %% [code]
import pandas as pd
import numpy as np
import matplotlib.pyplot as plt
import seaborn as sns
from Lu2023_akita_preds_functions import *

# %% [code]
# Akita-dependent global variables 

global BIN_SIZE, HEAD_INDEX, INPUT_LENGTH, TARGET_CROP, PRED_LENGTH, MAP_SIZE, TARGET_INDICES, DIAGONAL_OFFSET

BIN_SIZE = 2048                    #bp
HEAD_INDEX = 0                     #human head
INPUT_LENGTH = 1310720             #constant length of Akita input
TARGET_CROP = 131072               #length of the sequence cropped from each side (Akita input sequence preprocessing)
PRED_LENGTH = 130305               #length of Akita prediction vector
MAP_SIZE = 512                     #map representating changes in the DNA contacts (transformed Akita prediction vector)
TARGET_INDICES = [0, 1, 2, 3, 4]   #5 human target cell lines
DIAGONAL_OFFSET = 2                #number of diagonals that are added as zeros in the conversion (Akita input sequence preprocessing)

# %% [code]
# files to read

fasta_file = "repeat.same_orient.with_hg38.fa"

preds_directory = "/project/fudenber_735/data/Lu2023_akita_preds/"
preds_files = [f"genomic_context_predictions_m{i}.h5" for i in range(8)]
preds_paths = [f"{preds_directory}{file}" for file in preds_files]

fimo_file = "fimo.34bp.MA1929.1e_4.tsv"

# %% [markdown]
# reading a fasta file of alternate VNTR haplotypes

# %% [code]
summary_df = read_fasta(fasta_file)

haplotyples_names = list(summary_df.haplotyple_name)
reference_length = int(summary_df[summary_df["haplotyple_name"] == "GRCh38"].length)

# a relative end of the reference haplotype
# since the reference prediction is centered around it
rel_ref_start = (INPUT_LENGTH // 2) + (reference_length // 2) + 1

# %% [markdown]
# loading Akita predictions

# %% [code]
global summary_matrix
summary_matrix = average_experiments_multiple_models(
    preds_paths,
    haplotyples_names,
    head_index=HEAD_INDEX,
    pred_length=PRED_LENGTH,
    target_indices_list=TARGET_INDICES
)

# %% [markdown]
# plotting example maps

# %% [markdown]
### HGSVC.HG00513.0

# %% [code]
triple_plot(summary_matrix, summary_df, "HGSVC.HG00513.0", reference_length, rel_ref_start)

# %% [markdown]
### HPRC.HG02055.1

# %% [code]
triple_plot(summary_matrix, summary_df, "HPRC.HG02055.1", reference_length, rel_ref_start)

# %% [markdown]
# calculating local disruption score (LDR-128)

# %% [code]
summary_df = add_LDS(summary_df,
                     summary_matrix, 
                     haplotyples_names,
                     reference_length,
                     rel_ref_start,
                     square_size=128,
                     target_crop=TARGET_CROP, 
                     input_size=INPUT_LENGTH, 
                     bin_size=BIN_SIZE,
                     number_bins=MAP_SIZE,
                     diagonal_offset=DIAGONAL_OFFSET,
                     ref_index=0)

# %% [markdown]
# reading a table with CTCF motifs found

# %% [code]
fimo_motifs = pd.read_csv(fimo_file, sep="\t")

summary_df["num_motifs"] = summary_df["haplotyple_name"].apply(lambda haplotype_name : fimo_motifs["sequence_name"].value_counts()[haplotype_name])

# %% [markdown]
# plotting LDR-128 against haplotypes' lengths and predicted number in CTCF motifs

# %% [code]
plt.figure(figsize=(8,6))
plt.scatter(x=summary_df["length"], y=summary_df["LDS"])
plt.xlabel("haplotype_length")
plt.ylabel("local_discruption_score-128")

# %% [code]
plt.figure(figsize=(8,6))
plt.scatter(x=summary_df["num_motifs"], y=summary_df["LDS"])
plt.xlabel("predicted number of CTCF motifs")
plt.ylabel("local_discruption_score-128")

# %% [code]
```
