## Supplemental code for "The motif composition of variable-number tandem repeats impacts gene expression": 20230303.release.bias_correction.GTEx.ipynb

```
# %% [code]
#!/usr/bin/env python3
import sys
srcdir = "/project/mchaisso_100/cmb-16/tsungyul/work/vntr/danbing-tk/script/"
sys.path.insert(0, srcdir)

# %% [code]
import numpy as np
import pandas as pd
import vntrutils as vu
import utils
# import scipy.stats as stats
import matplotlib
import matplotlib.pyplot as plt
%matplotlib inline

import seaborn as sns
# %load_ext autoreload
# %autoreload 2

# %% [code]
from collections import defaultdict
import pickle
# import networkx as nx
# import pygraphviz
# import collections
import gc
# import glob
# import warnings
import statsmodels.api as sm
# from statsmodels.stats.multitest import fdrcorrection as fdr
# from scipy import stats
import gzip
# from sklearn.decomposition import PCA
# from scipy.stats import chi2
# import re
# import json

# %% [code]
matplotlib.rc('font', size=7)
matplotlib.rc('axes', titlesize=7)
matplotlib.rc('xtick', labelsize=5)
matplotlib.rc('ytick', labelsize=5)
%load_ext autoreload
%autoreload 2

# %% [code]


# %% [code]
def test():
    with open("/project/mchaisso_100/cmb-17/vntr_genotyping/rpgg2_k21_84k/hgsvc/clinic/cdbg/v1/ki_tr.ccki_tr.pickle", 'rb') as f:
        ki_tr, ccki_tr = pickle.load(f)
    return ki_tr, ccki_tr
ki_tr, ccki_tr = test()

# %% [code]
def get_ki(tri):
    si = ki_tr[tri-1] if tri else 0
    ei = ki_tr[tri]
    return si, ei
def get_ccki(tri):
    si = ccki_tr[tri-1] if tri else 0
    ei = ccki_tr[tri]
    return si, ei

# %% [markdown]
# Bias correction for GTEx genotypes

# %% [code]
def test():
    with open("/project/mchaisso_100/cmb-17/vntr_genotyping/rpgg2_k21_84k/hgsvc/clinic/bias/invariant_kmer/ikmer.ikstats.pickle", 'rb') as f:
        ikmer, ikstats = pickle.load(f)
    return ikmer, ikstats
ikmer, ikstats = test()

# %% [code]
cov = np.loadtxt("/project/mchaisso_100/cmb-17/vntr_genotyping/analysis/read_depth/ctrl/gtex.cov.tsv", usecols=1)

# %% [code]
def test():
    with open(f"/scratch1/tsungyul/tmp/gt.1.pickle", 'rb') as f:
        gt = pickle.load(f)
    with open(f"/project/mchaisso_100/cmb-17/vntr_genotyping/rpgg2_k21_84k/hgsvc/gtex/gt_pickle/repickle.python3.9/gt.1.pickle", 'wb') as f:
        pickle.dump(gt, f, protocol=pickle.HIGHEST_PROTOCOL)
test()

# %% [code]
def test():
    ng = np.loadtxt("/project/mchaisso_100/cmb-16/tsungyul/work/vntr/hapdb/config/genomes.gtex.txt", dtype=object).size
    with open("/project/mchaisso_100/cmb-17/vntr_genotyping/rpgg2_k21_84k/hgsvc/clinic/cdbg/v1/tr_cck_ns.pickle", 'rb') as f:
        tr_cck_ns = pickle.load(f)
    nk = ki_tr[-1]
    ntr = len(ki_tr)
    
    NB = 10
    BS = ng // NB
    bias = np.full([ng, ntr], np.nan)
    for i in range(NB):
        BS_ = BS if i != NB-1 else ng - BS*i
        i0s = i * BS
        i0e = i0s + BS_
        print(f"batch {i}: loading gt", end="")
        with open(f"/project/mchaisso_100/cmb-17/vntr_genotyping/rpgg2_k21_84k/hgsvc/gtex/gt_pickle/gt.{i}.pickle", 'rb') as f:
            agt = pickle.load(f).astype(float) / cov[i0s:i0e,None]
        for tri in range(ntr):
            if tri % 1000 == 0:
                print(".", end="")
            si, ei = get_ki(tri)
            inv = np.nonzero(ikmer[si:ei])[0] + si
            if inv.size:
                bias[i0s:i0e, tri] = np.mean(agt[:,inv] / ikstats[0,inv], axis=1)
        print()
        # del agt
        # gc.collect()
    bias[bias==0] = np.nan
    bias /= np.nanmean(bias, axis=0)
    n0 = np.sum(~np.isfinite(bias))
    print(f"{n0} cells of nan's")
    return bias

bias = test()

# %% [code]
pd.DataFrame(bias)

# %% [code]
def test():
    with open("/project/mchaisso_100/cmb-17/vntr_genotyping/rpgg2_k21_84k/hgsvc/gtex/bias/bias.v1.pickle", 'wb') as f:
        pickle.dump(bias, f)
# test()

# %% [code]
bias = pickle.load(open("/project/mchaisso_100/cmb-17/vntr_genotyping/rpgg2_k21_84k/hgsvc/gtex/bias/bias.v1.pickle", 'rb'))

# %% [code]
def test():
    gloci = np.all(pd.read_csv("/project/mchaisso_100/cmb-17/vntr_genotyping/rpgg2_k21_84k/hgsvc/clinic/analysis/80518_loci.alu.mismap_1.r2.v2.filter.tsv", sep="\t", index_col=0).iloc[:,3:].values, axis=1)
    droppedloci = ~np.isfinite(bias[0])
    n0 = np.sum(droppedloci)
    n1 = np.sum(gloci & droppedloci)
    print(f"{n0} loci have nan bias")
    print(f"{n1}/{n0} overlapped with loci passing QC")
test()

# %% [code]
def test():
    with open("/project/mchaisso_100/cmb-17/vntr_genotyping/rpgg2_k21_84k/hgsvc/gtex/gt_pickle/acgt.pickle", 'rb') as f:
        return pickle.load(f).astype(float) # ng x ncck
acgt = test()

# %% [code]
def test():
    ntr = len(ccki_tr)
    acgt_bc = np.copy(acgt)
    for tri in range(ntr):
        si, ei = get_ccki(tri)
        acgt_bc[:,si:ei] /= (bias[:,tri][:,None])
    return acgt_bc
acgt_bc = test()

# %% [code]
pd.DataFrame(acgt_bc)

# %% [code]
def test():
    with open("/project/mchaisso_100/cmb-17/vntr_genotyping/rpgg2_k21_84k/hgsvc/gtex/gt_pickle/acgt_bc.v1.pickle", 'wb') as f:
        pickle.dump(acgt_bc, f, protocol=pickle.HIGHEST_PROTOCOL)
# test()

# %% [code]
def test():
    with open("/project/mchaisso_100/cmb-17/vntr_genotyping/rpgg2_k21_84k/hgsvc/gtex/gt_pickle/acgt_bc.v1.pickle", 'rb') as f:
        return pickle.load(f)
acgt_bc = test()

# %% [code]
def test():
    ntr = len(ki_tr)
    for tri in range(ntr):
        si, ei = get_ccki(tri)
        xs = np.sum(acgt_bc[:,si:ei], axis=1)
        utils.hist(xs, "Read TR dosage", "Count", f"{tri}", bins=50)
        input()
test()

# %% [code]
def test():
    with open("/project/mchaisso_100/cmb-17/vntr_genotyping/rpgg2_k21_84k/hgsvc/gtex/genotype/gtex.80518x879.akms.pickle", 'rb') as f:
        return pickle.load(f)
akms = test()

# %% [code]
pd.DataFrame(akms)

# %% [code]
def test():
    akms_bc = np.copy(akms)
    akms_bc /= bias.T
    return akms_bc
akms_bc = test()

# %% [code]
pd.DataFrame(akms_bc)

# %% [code]
def test():
    with open("/project/mchaisso_100/cmb-17/vntr_genotyping/rpgg2_k21_84k/hgsvc/gtex/genotype/gtex.80518x879.akms_bc.v1.pickle", 'wb') as f:
        pickle.dump(akms_bc, f, protocol=pickle.HIGHEST_PROTOCOL)
test()

# %% [code]
```
