## Supplemental code for "The motif composition of variable-number tandem repeats impacts gene expression": 20230303.release.bias_correction.HGSVC.ipynb

```
# %% [code]
#!/usr/bin/env python3
import sys
srcdir = "/project/mchaisso_100/cmb-16/tsungyul/work/vntr/danbing-tk/script/"
sys.path.insert(0, srcdir)

# %% [code]
import numpy as np
import pandas as pd
import vntrutils as vu
import utils
# import scipy.stats as stats
import matplotlib
import matplotlib.pyplot as plt
%matplotlib inline

import seaborn as sns
# %load_ext autoreload
# %autoreload 2

# %% [code]
from collections import defaultdict
import pickle
# import networkx as nx
# import pygraphviz
# import collections
import gc
# import glob
# import warnings
import statsmodels.api as sm
# from statsmodels.stats.multitest import fdrcorrection as fdr
# from scipy import stats
# import gzip
# from sklearn.decomposition import PCA
# from scipy.stats import chi2
# import re
# import json

# %% [code]
matplotlib.rc('font', size=7)
matplotlib.rc('axes', titlesize=7)
matplotlib.rc('xtick', labelsize=5)
matplotlib.rc('ytick', labelsize=5)

# %% [raw]
- procedure
    - HGSVC asm --+--> pbgt --(convert kmers to motifs)--> pbcgt: input for motif batch-r2
                 |
                 +--> trlens: input for length batch-r2
                 
    - HGSVC short read samples --(aln to HGSVC RPGG)--> ilgt --(compute bias with HGSVC invariant kmers)--> bias -->>---+
                                                         |                                                              |
        +--------------------------------------------<<--+                                                              |
        |                                                                                                               |
        +--(summing kmer counts in each locus)--> trdsg: input for length batch-r2 (pre-correction)                     |
        |                                                                                                               |
        +--(convert kmers to motifs + read depth normalization)--> ilcgt: input for motif batch-r2 (pre-correction)     |
        |                                                             |                                                 |
        |  +--------------------------------------------------------------------------------------------------------<<--+
        |  |                                                          |                                                 |
        +--+-->>--(bias correction)--> ilgt_bc --> trdsg_bc: input for length batch-r2 (post-correction)                |
                                                                      |                                                 |
                                                                      |  +------------------------------------------<<--+
                                                                      |  |
                                                                      +--+-->>--(bias correction)--> ilcgt_bc: input for motif batch-r2 (post-correction)

# %% [code]
def test():
    with open("/project/mchaisso_100/cmb-17/vntr_genotyping/rpgg2_k21_84k/hgsvc/clinic/cdbg/v1/ki_tr.ccki_tr.pickle", 'rb') as f:
        ki_tr, ccki_tr = pickle.load(f)
    return ki_tr, ccki_tr
ki_tr, ccki_tr = test()

# %% [code]
def test():
    with open("/project/mchaisso_100/cmb-17/vntr_genotyping/rpgg2_k21_84k/hgsvc/clinic/cdbg/v1/ks.ccks.cck_ns.ki_map.pickle", 'rb') as f:
        ks, ccks, cck_ns, ki_map = pickle.load(f)
    return ks, ccks, cck_ns, ki_map
ks, ccks, cck_ns, ki_map = test()

# %% [code]
def get_ki(tri):
    si = ki_tr[tri-1] if tri else 0
    ei = ki_tr[tri]
    return si, ei
def get_ccki(tri):
    si = ccki_tr[tri-1] if tri else 0
    ei = ccki_tr[tri]
    return si, ei

# %% [code]
def test():
    with open("/project/mchaisso_100/cmb-17/vntr_genotyping/rpgg2_k21_84k/hgsvc/clinic/bias/invariant_kmer/ikmer.ikstats.pickle", 'rb') as f:
        ikmer, ikstats = pickle.load(f)
    return ikmer, ikstats
ikmer, ikstats = test()

# %% [markdown]
# Evaluate using HGSVC samples

# %% [markdown]
## Bias correction for HGSVC genotypes

# %% [code]
cov = np.loadtxt("/project/mchaisso_100/cmb-17/vntr_genotyping/analysis/read_depth/ctrl/HGSVC.cov.tsv", usecols=1)

# %% [code]
# if bias == 0, set it as nan
def test():
    ng = cov.size
    ntr = len(ki_tr)
    
    NB = 1
    BS = ng // NB
    bias = np.zeros([ng, ntr])
    for i in range(NB):
        BS_ = BS if i != NB-1 else ng - BS*i
        i0s = i * BS
        i0e = i0s + BS_
        print(f"batch {i}: loading gt", end="")
        with open(f"/project/mchaisso_100/cmb-17/vntr_genotyping/rpgg2_k21_84k/hgsvc/clinic/gt_pickle/gt.{i}.pickle", 'rb') as f:
            gt = pickle.load(f)
        agt = gt / cov[i0s:i0e,None]
        for tri in range(ntr):
            if tri % 1000 == 0:
                print(".", end="")
            si, ei = get_ki(tri)
            inv = np.nonzero(ikmer[si:ei])[0] + si
            if inv.size:
                bias[i0s:i0e, tri] = np.mean(agt[:,inv] / ikstats[0,inv], axis=1)
        print()
    bias[bias==0] = np.nan
    bias /= np.nanmean(bias, axis=0)
    n0 = np.sum(~np.isfinite(bias))
    print(f"{n0} cells of nan's")
    return bias

bias = test()

# %% [code]
def test():
    with open("/project/mchaisso_100/cmb-17/vntr_genotyping/rpgg2_k21_84k/hgsvc/clinic/bias/invariant_kmer/bias.v2.pickle", 'wb') as f:
        pickle.dump(bias, f)
# test()

# %% [code]
def test():
    with open("/project/mchaisso_100/cmb-17/vntr_genotyping/rpgg2_k21_84k/hgsvc/clinic/bias/invariant_kmer/bias.v2.pickle", 'rb') as f:
        return pickle.load(f)
bias = test()

# %% [code]
np.savetxt("/project/mchaisso_100/cmb-17/vntr_genotyping/rpgg2_k21_84k/hgsvc/clinic/bias/invariant_kmer/bias.v2.32x80518", sep="\t")

# %% [code]
def test():
    with open("/project/mchaisso_100/cmb-17/vntr_genotyping/rpgg2_k21_84k/hgsvc/clinic/cdbg/v1/ks.ccks.cck_ns.ki_map.pickle", 'rb') as f:
        ks, ccks, cck_ns, ki_map = pickle.load(f)
    return ks, ccks, cck_ns, ki_map
ks, ccks, cck_ns, ki_map = test()

# %% [code]
ks = np.array(ks)

# %% [markdown]
## motif batch-$r^2$

# %% [code]
def test():
    omap1 = pd.read_csv("/project/mchaisso_100/cmb-17/vntr_genotyping/rpgg2_k21_84k/hgsvc/clinic/output/OrthoMap.v2.tsv", sep="\t", header=None, na_values=".").values
    
    gs = np.loadtxt("/project/mchaisso_100/cmb-17/vntr_genotyping/rpgg2_k21_84k/hgsvc/clinic/LOO/input/genome.LOO.bam.tsv", usecols=0, dtype=object)
    gs1 = np.loadtxt("/project/mchaisso_100/cmb-16/tsungyul/work/vntr/hapdb/config/genomes.HGSVC2.txt", dtype=object)
    gm1 = np.isin(gs1, gs)
    hm1 = np.repeat(gm1, 2)
    ng = gs.size
    nh = 2*ng
    ntr = omap1.shape[0]
    
    omap = omap1[:,hm1]
    trgmask = np.ones([ntr, ng], dtype=bool)
    i1s = np.arange(0, nh, 2)
    trgmask &= np.isfinite(omap[:,i1s]) # hap0 in not missing in TR annotation
    trgmask &= np.isfinite(omap[:,i1s+1]) # hap1 in not missing in TR annotation
    return trgmask
    
trgmask = test()

# %% [code]
np.savetxt("/project/mchaisso_100/cmb-17/vntr_genotyping/rpgg2_k21_84k/hgsvc/clinic/bias/invariant_kmer/input2/tr.genome.mask.txt", trgmask, fmt="%s", delimiter="\t")

# %% [code]
pd.read_csv("/project/mchaisso_100/cmb-17/vntr_genotyping/rpgg2_k21_84k/hgsvc/clinic/bias/invariant_kmer/input2/tr.genome.mask.txt", sep="\t", header=None)

# %% [code]
def test():
    with open("/project/mchaisso_100/cmb-17/vntr_genotyping/rpgg2_k21_84k/hgsvc/clinic/gt_pickle/pbcgt.pickle", 'rb') as f:
        pbcgt = pickle.load(f)
    return pbcgt
pbcgt = test()

# %% [code]
def test():
    with open("/project/mchaisso_100/cmb-17/vntr_genotyping/rpgg2_k21_84k/hgsvc/clinic/gt_pickle/acgt.pickle", 'rb') as f:
        acgt = pickle.load(f)
    return acgt
acgt = test()

# %% [code]
# compute `r2.{bi}.pickle` using /project/mchaisso_100/cmb-17/vntr_genotyping/rpgg2_k21_84k/hgsvc/clinic/bias/invariant_kmer/log2/batch_r2.sh
def test():
    ng, ntr = bias.shape
    nk = ccki_tr[-1]
    NB = 8
    BS = ntr // NB
    R2 = np.full(nk, np.nan)
    R2a = np.full(nk, np.nan)
    for bi in range(NB):
        with open(f"/project/mchaisso_100/cmb-17/vntr_genotyping/rpgg2_k21_84k/hgsvc/clinic/bias/invariant_kmer/output2/r2.{bi}.pickle", 'rb') as f:
            r2, r2a = pickle.load(f)
        tri0 = bi*BS
        tri1 = (bi+1)*BS-1 if bi != NB-1 else ntr-1
        si = ccki_tr[tri0-1] if tri0 else 0
        ei = ccki_tr[tri1]
        R2[si:ei] = r2[si:ei]
        R2a[si:ei] = r2a[si:ei]
    return R2, R2a
R2_all_mtf, R2a_all_mtf = test()

# %% [code]
def test():
    with open(f"/project/mchaisso_100/cmb-17/vntr_genotyping/rpgg2_k21_84k/hgsvc/clinic/bias/invariant_kmer/output2/R2_all_mtf.R2a_all_mtf.pickle", 'wb') as f:
        pickle.dump((R2_all_mtf, R2a_all_mtf), f)
# test()

# %% [code]
def test():
    with open(f"/project/mchaisso_100/cmb-17/vntr_genotyping/rpgg2_k21_84k/hgsvc/clinic/bias/invariant_kmer/output2/R2_all_mtf.R2a_all_mtf.pickle", 'rb') as f:
        return pickle.load(f)
R2_all_mtf, R2a_all_mtf = test()

# %% [code]
trfilter = np.all(pd.read_csv("/project/mchaisso_100/cmb-17/vntr_genotyping/rpgg2_k21_84k/hgsvc/clinic/analysis/80518_loci.alu.mismap_1.r2.v2.filter.tsv", sep="\t", index_col=0).values[:,3:], axis=1)

# %% [code]
motiffilter = np.all(np.loadtxt("/project/mchaisso_100/cmb-17/vntr_genotyping/rpgg2_k21_84k/hgsvc/clinic/analysis/mape.invariant.flank.alu.mismap_1.r2.v2.filter.tsv.gz", dtype=int, usecols=[3,4,5,6,7,8,9], skiprows=1).astype(bool), axis=1)

# %% [code]
# Peak at 1 due to private motifs observed in only 1 genome.
# Large number of motifs with r2 close to 1 but not exactly due to noise for 0-dosage samples.
def test():
    xs = R2a_all_mtf[motiffilter]
    utils.hist(xs, "batch-$r^2$", "Number of motifs", bins=100, dpi=300)
    xs = xs[xs < 1]
    utils.hist(xs, "batch-$r^2$", "Number of motifs", bins=100, dpi=300)
test()

# %% [code]
def test():
    ng, ntr = bias.shape
    R2 = np.full(ntr, np.nan)
    R2a = np.full(ntr, np.nan)
    for tri, good in enumerate(trfilter):
        if not good: continue
        si = ccki_tr[tri-1] if tri else 0
        ei = ccki_tr[tri]
        m0 = motiffilter[si:ei]
        R2[tri] = np.nanmean(R2_all_mtf[si:ei][m0])
        R2a[tri] = np.nanmean(R2a_all_mtf[si:ei][m0])
    
    n0 = np.sum(np.isfinite(R2a))
    n1 = np.sum(trfilter)
    print(f"{n0:,}/{n1:,} ({n0/n1:.1%}) loci evaluated")
    print(f"{np.nanmean(R2):.3f} -> {np.nanmean(R2a):.3f}")
    xl = [np.nanmin(R2), np.nanmax(R2)]
    plt.figure(dpi=300, figsize=(2,1.8))
    plt.plot(R2, R2a, '.k', mew=0, markersize=1, alpha=0.1)
    plt.plot(xl, xl, '--r', lw=0.4)
    plt.xlabel("batch-$r^2$ (before, locus ave.)")
    plt.ylabel("batch-$r^2$ (after, locus ave.)")
    plt.show(); plt.close()
    return R2, R2a
R2_all_loci, R2a_all_loci = test()

# %% [code]
def test():
    R2, R2a = R2_all_mtf[motiffilter], R2a_all_mtf[motiffilter]
    n0 = np.sum(np.isfinite(R2a))
    n1 = R2.size
    print(f"{n0:,}/{n1:,} ({n0/n1:.1%}) motifs evaluated")
    print(f"{np.nanmean(R2):.3f} -> {np.nanmean(R2a):.3f}")
    xl = [np.nanmin(R2), np.nanmax(R2)]
    plt.figure(dpi=300, figsize=(2,1.8))
    plt.plot(R2, R2a, '.k', mew=0, markersize=1, alpha=0.002)
    plt.plot(xl, xl, '--r', lw=0.4)
    plt.xlabel("batch-$r^2$ (before)")
    plt.ylabel("batch-$r^2$ (after)")
    plt.show(); plt.close()
test()

# %% [markdown]
## length batch-$r^2$

# %% [code]
def test():
    trhlens0 = pd.read_csv("/project/mchaisso_100/cmb-17/vntr_genotyping/rpgg2_k21_84k/hgsvc/clinic/analysis/trlen.tsv", sep="\t", na_values="nan", index_col=0).iloc[:,6:].values.astype(float)
    ntr = trhlens0.shape[0]
    gs = np.loadtxt("/project/mchaisso_100/cmb-17/vntr_genotyping/rpgg2_k21_84k/hgsvc/clinic/LOO/input/genome.LOO.bam.tsv", usecols=0, dtype=object)
    gs0 = np.loadtxt("/project/mchaisso_100/cmb-16/tsungyul/work/vntr/hapdb/config/genomes.HGSVC2.txt", dtype=object)
    gm0 = np.isin(gs0,gs)
    hm0 = np.repeat(gm0, 2)
    ng = gs.size
    trhlens = trhlens0[:,hm0]
    trlens = np.full([ntr, ng], np.nan)
    for gi in range(ng):
        trlens[:,gi] = np.mean(trhlens[:,2*gi:2*gi+2], axis=1)
    return trlens
trlens = test()

# %% [code]
pd.DataFrame(trlens)

# %% [code]
def test():
    with open("/project/mchaisso_100/cmb-17/vntr_genotyping/rpgg2_k21_84k/hgsvc/clinic/gt_pickle/gt.0.pickle", 'rb') as f:
        ilgt = pickle.load(f).astype(float)
    return ilgt / np.loadtxt("/project/mchaisso_100/cmb-17/vntr_genotyping/analysis/read_depth/ctrl/HGSVC.cov.tsv", usecols=1)[:,None]
ilagt = test()

# %% [code]
def test():
    trdsg = np.zeros_like(trlens)
    for tri, good in enumerate(trfilter):
        if not good: continue
        si, ei = get_ki(tri)
        trdsg[tri] = np.sum(ilagt[:,si:ei], axis=1)
    return trdsg
trdsg = test()

# %% [code]
pd.DataFrame(trdsg)

# %% [code]
def test():
    ntr = len(ki_tr)
    
    nscan, nconst, ndrop = 0, 0, 0
    R2 = np.full(ntr, np.nan)
    for tri, good in enumerate(trfilter):
        if not good: continue
        nscan += 1
        xs = trlens[tri]
        ys = trdsg[tri]
        if not np.any(np.isfinite(ys)):
            ndrop += 1
            continue
        m = np.isfinite(xs) & np.isfinite(ys)
        xs = xs[m]
        xs = np.clip(xs-20, 0, None)
        if not np.any(xs): 
            assert False
            continue
        if np.unique(xs).size == 1:
            nconst += 1
            continue
        ys = ys[m]
        R2[tri] = sm.OLS(np.insert(ys,0,0), sm.add_constant(np.insert(xs,0,0))).fit().rsquared
        
    n0 = np.sum(R2>0.8)
    n1 = nscan - nconst
    n2 = nscan
    print(f"{nscan} TRs scanned")
    print(f"{nconst} TRs not tested: no variation in length for this TR in HGSVC")
    print(f"{ndrop} TRs dropped")
    print(f"{n0:,}/{n1:,} ({n0/n1:.1%}) has r2 > 0.8")
    
    nscan, nconst, ndrop = 0, 0, 0
    R2a = np.full(ntr, np.nan)
    for tri, good in enumerate(trfilter):
        if not good: continue
        nscan += 1
        xs = trlens[tri]
        ys = trdsg[tri] / bias[:,tri]
        if not np.any(np.isfinite(ys)):
            ndrop += 1
            continue
        m = np.isfinite(xs) & np.isfinite(ys)
        xs = xs[m]
        xs = np.clip(xs-20, 0, None)
        if not np.any(xs): 
            assert False
            continue
        if np.unique(xs).size == 1:
            nconst += 1
            continue
        ys = ys[m]
        R2a[tri] = sm.OLS(np.insert(ys,0,0), sm.add_constant(np.insert(xs,0,0))).fit().rsquared
        
    n0 = np.sum(R2a>0.8)
    n1 = nscan - nconst
    n2 = nscan
    print()
    print(f"{nscan} TRs scanned")
    print(f"{nconst} TRs not tested: no variation in length for this TR in HGSVC")
    print(f"{ndrop} TRs dropped during bias correction")
    print(f"{n0:,}/{n1:,} ({n0/n1:.1%}) has r2 > 0.8")
    
    n0 = np.sum(np.isfinite(R2a))
    n1 = np.sum(trfilter)
    print()
    print(f"{n0:,}/{n1:,} ({n0/n1:.1%}) loci evaluated")
    print(f"{np.nanmean(R2):.3f} -> {np.nanmean(R2a):.3f}")
    
    xl = [np.nanmin(R2), np.nanmax(R2)]
    plt.figure(dpi=300, figsize=(2,1.8))
    plt.plot(R2, R2a, '.k', mew=0, markersize=1, alpha=0.1)
    plt.plot(xl, xl, '--r', lw=0.4)
    plt.xlabel("batch-$r^2$ (before)")
    plt.ylabel("batch-$r^2$ (after)")
    plt.show(); plt.close()
    
    return R2, R2a
R2_all_loci, R2a_all_loci = test()

# %% [code]
np.savetxt("/project/mchaisso_100/cmb-17/vntr_genotyping/rpgg2_k21_84k/hgsvc/clinic/bias/invariant_kmer/output2/R2_all_loci.R2a_all_loci.txt", np.hstack((R2_all_loci[:,None], R2a_all_loci[:,None])), delimiter="\t", fmt="%s")

# %% [code]
```
