## Supplemental code for "The motif composition of variable-number tandem repeats impacts gene expression": 20230303.release.bias_correction.HPRC.ipynb

```
# %% [code]
#!/usr/bin/env python3
import sys
srcdir = "/project/mchaisso_100/cmb-16/tsungyul/work/vntr/danbing-tk/script/"
sys.path.insert(0, srcdir)

# %% [code]
import numpy as np
import pandas as pd
import vntrutils as vu
import utils
# import scipy.stats as stats
import matplotlib
import matplotlib.pyplot as plt
%matplotlib inline

import seaborn as sns
# %load_ext autoreload
# %autoreload 2

# %% [code]
from collections import defaultdict
import pickle
# import networkx as nx
# import pygraphviz
# import collections
import gc
import glob
# import warnings
import statsmodels.api as sm
# from statsmodels.stats.multitest import fdrcorrection as fdr
# from scipy import stats
import gzip
# from sklearn.decomposition import PCA
# from scipy.stats import chi2
# import re
# import json

# %% [code]
matplotlib.rc('font', size=7)
matplotlib.rc('axes', titlesize=7)
matplotlib.rc('xtick', labelsize=5)
matplotlib.rc('ytick', labelsize=5)
%load_ext autoreload
%autoreload 2

# %% [raw]
- procedure
    - HPRC asm --+--> pbgt --(convert kmers to motifs)--> pbcgt: input for motif batch-r2
                 |
                 +--> trlens: input for length batch-r2
                 
    - HPRC short read samples --(aln to HGSVC RPGG)--> ilgt --(compute bias with HGSVC invariant kmers)--> bias -->>---+
                                                        |                                                              |
        +-------------------------------------------<<--+                                                              |
        |                                                                                                              |
        +--(summing kmer counts in each locus)--> trdsg: input for length batch-r2 (pre-correction)                    |
        |                                                                                                              |
        +--(convert kmers to motifs + read depth normalization)--> ilcgt: input for motif batch-r2 (pre-correction)    |
        |                                                             |                                                |
        |  +-------------------------------------------------------------------------------------------------------<<--+
        |  |                                                          |                                                |
        +--+-->>--(bias correction)--> ilgt_bc --> trdsg_bc: input for length batch-r2 (post-correction)               |
                                                                      |                                                |
                                                                      |  +-----------------------------------------<<--+
                                                                      |  |
                                                                      +--+-->>--(bias correction)--> ilcgt_bc: input for motif batch-r2 (post-correction)

# %% [markdown]
# IL dosage

# %% [code]
def test():
    fn = "/project/mchaisso_100/cmb-17/vntr_genotyping/rpgg2_k21_84k/hgsvc/clinic/cdbg/v1/ks.ccks.cck_ns.ki_map.pickle"
    with open(fn, 'rb') as f:
        ks, ccks, cck_ns, ki_map = pickle.load(f)
    print(len(ks), len(ccks))
    return ki_map
ki_map = test()

# %% [code]
def load_single_gt(fn, gt, i):
    with open(fn) as f:
        j = 0
        for line in f:
            gt[i,j] = int(line)
            j += 1
    
def compute_gt_cgt_batch():
    kmerfs = sorted(glob.glob("/project/mchaisso_100/cmb-17/vntr_genotyping/rpgg2_k21_84k/hgsvc/1kgr/hprc/output/*.tr.kmers"))
    ng = len(kmerfs)
    fn = "/project/mchaisso_100/cmb-17/vntr_genotyping/rpgg2_k21_84k/hgsvc/clinic/cdbg/v1/ks.ccks.cck_ns.ki_map.pickle"
    with open(fn, 'rb') as f:
        ks, ccks, cck_ns, ki_map = pickle.load(f)
    nk = len(ks)
    ncck = len(ccks)
    print(ng, nk, ncck)
    cck_ns = np.array(cck_ns)
    tr_cck_ns = np.zeros(ncck, dtype=int)
    for i0 in range(nk):
        tr_cck_ns[ki_map[i0]] += 1
    
    NB = 1
    BS = ng // NB
    for i in range(NB):
        BS_ = BS if i != NB-1 else ng - BS*i
        print(f"batch {i}: loading gt... ", end="")
        gt = np.zeros([BS_, nk], dtype=np.int32)
        for j in range(BS_):
            load_single_gt(kmerfs[i*BS+j], gt, j)
        print(f"computing cgt... ", end="")
        cgt = np.zeros([BS_,ncck], dtype=np.float32)
        for i0, i1 in ki_map.items():
            cgt[:,i1] += gt[:,i0]
        print(pd.DataFrame(cgt))
        cgt /= tr_cck_ns
        print("dumping gt... ", end="")
        with open(f"/project/mchaisso_100/cmb-17/vntr_genotyping/rpgg2_k21_84k/hgsvc/1kgr/hprc/gt_pickle/gt.{i}.pickle", 'wb') as f:
            pickle.dump(gt, f, protocol=pickle.HIGHEST_PROTOCOL)
        print("dumping cgt...")
        with open(f"/project/mchaisso_100/cmb-17/vntr_genotyping/rpgg2_k21_84k/hgsvc/1kgr/hprc/gt_pickle/cgt.{i}.pickle", 'wb') as f:
            pickle.dump(cgt, f, protocol=pickle.HIGHEST_PROTOCOL)

!mkdir -p /project/mchaisso_100/cmb-17/vntr_genotyping/rpgg2_k21_84k/hgsvc/1kgr/hprc/gt_pickle
compute_gt_cgt_batch()

# %% [code]
def test():
    # correct coverage and X chrom dosage
    genomes = np.loadtxt("/project/mchaisso_100/cmb-16/tsungyul/work/vntr/hapdb/config/genomes.HPRC.notinHGSVC.hasIL.src.txt", usecols=1, dtype=object)
    ng = genomes.size
    g2i = dict([[g,i] for i, g in enumerate(genomes)])
    cov = np.array([float(c) for g, c in np.loadtxt("/project/mchaisso_100/cmb-17/vntr_genotyping/analysis/read_depth/ctrl/HPRC.notinHGSVC.chr1.cov.tsv", dtype=object) if g in genomes])
    sex = np.zeros(cov.size, dtype=int)
    sex2i = {"female":2, "male":1}
    for g, v1 in np.loadtxt("/project/mchaisso_100/cmb-17/vntr_genotyping/rpgg2_k21_84k/hgsvc/1kgr/hprc/input/sample.meta.txt", skiprows=1, usecols=[0,1], dtype=object):
        if g in g2i:
            sex[g2i[g]] = sex2i[v1]
    print("Loading cgt", end="")
    NCCK = 4456881
    NB = 1
    BS = ng//NB
    cgt = np.zeros([ng,NCCK], dtype=np.float32)
    for i in range(NB):
        print(f".",end="")
        BS_ = BS if i != NB-1 else ng - BS*i
        with open(f"/project/mchaisso_100/cmb-17/vntr_genotyping/rpgg2_k21_84k/hgsvc/1kgr/hprc/gt_pickle/cgt.{i}.pickle", 'rb') as f:
            si = i*BS
            ei = i*BS + BS_
            cgt[si:ei] = pickle.load(f)
    print()
    print("Computing acgt")
    with open("/project/mchaisso_100/cmb-17/vntr_genotyping/rpgg2_k21_84k/hgsvc/clinic/cdbg/v1/ki_tr.ccki_tr.pickle", 'rb') as f:
        _, ccki_tr = pickle.load(f)
    acgt = np.zeros_like(cgt, dtype=np.float32)
    ai = ccki_tr[NAU-1]
    acgt[:,:ai] = cgt[:,:ai] / cov[:,None]
    acgt[:,ai:] = cgt[:,ai:] / (cov * sex)[:,None]
    print("Dumping acgt")
    with open("/project/mchaisso_100/cmb-17/vntr_genotyping/rpgg2_k21_84k/hgsvc/1kgr/hprc/gt_pickle/acgt.pickle", 'wb') as f:
        pickle.dump(acgt, f, protocol=pickle.HIGHEST_PROTOCOL)
    return acgt

ilcgt = test()

# %% [code]
ilcgt =  pickle.load(open("/project/mchaisso_100/cmb-17/vntr_genotyping/rpgg2_k21_84k/hgsvc/1kgr/hprc/gt_pickle/acgt.pickle", 'rb'))

# %% [code]
ilcgt.shape

# %% [code]
def test():
    with open(f"/project/mchaisso_100/cmb-17/vntr_genotyping/rpgg2_k21_84k/hgsvc/1kgr/hprc/gt_pickle/gt.0.pickle", 'rb') as f:
        gt = pickle.load(f)
    genomes = np.loadtxt("/project/mchaisso_100/cmb-16/tsungyul/work/vntr/hapdb/config/genomes.HPRC.notinHGSVC.hasIL.src.txt", usecols=1, dtype=object)
    cov = np.array([float(c) for g, c in np.loadtxt("/project/mchaisso_100/cmb-17/vntr_genotyping/analysis/read_depth/ctrl/HPRC.notinHGSVC.chr1.cov.tsv", dtype=object) if g in genomes])
    agt = gt / cov[:,None]
    with open("/project/mchaisso_100/cmb-17/vntr_genotyping/rpgg2_k21_84k/hgsvc/1kgr/hprc/gt_pickle/agt.pickle", 'wb') as f:
        pickle.dump(agt, f, protocol=pickle.HIGHEST_PROTOCOL)
    return agt
# ilgt = test()

# %% [code]
ilgt =  pickle.load(open("/project/mchaisso_100/cmb-17/vntr_genotyping/rpgg2_k21_84k/hgsvc/1kgr/hprc/gt_pickle/agt.pickle", 'rb'))

# %% [code]
ilgt.shape

# %% [markdown]
# PB dosage

# %% [code]
def test():
    tri2km2i = {}
    with open("/project/mchaisso_100/cmb-17/vntr_genotyping/rpgg2_k21_84k/hgsvc/clinic/cdbg/v1/pan.reindex_as_output.tr.kmers") as f:
        tri = -1
        ki = -1
        for line in f:
            if line[0] == ">":
                tri += 1
                tri2km2i[tri] = {}
            else:
                ki += 1
                tri2km2i[tri][int(line.rstrip())] = ki
    return tri2km2i
tri2km2i = test()

# %% [code]
def test():
    with open("/project/mchaisso_100/cmb-17/vntr_genotyping/rpgg2_k21_84k/hgsvc/clinic/cdbg/v1/ki_tr.ccki_tr.pickle", 'rb') as f:
        ki_tr, _ = pickle.load(f)
    return ki_tr
ki_tr = test()

# %% [code]
def test():
    fn = "/project/mchaisso_100/cmb-17/vntr_genotyping/rpgg2_k21_84k/hgsvc/clinic/cdbg/v1/ks.ccks.cck_ns.ki_map.pickle"
    with open(fn, 'rb') as f:
        ks, ccks, cck_ns, ki_map = pickle.load(f)
    return ks, ccks, cck_ns, ki_map
ks, ccks, cck_ns, ki_map = test()

# %% [code]
def test():
    with open("/project/mchaisso_100/cmb-17/vntr_genotyping/rpgg2_k21_84k/hgsvc/clinic/cdbg/v1/ki_tr.ccki_tr.pickle", 'rb') as f:
        _, ccki_tr = pickle.load(f)
    return ccki_tr
ccki_tr = test()

# %% [code]
def test():
    with open("/project/mchaisso_100/cmb-17/vntr_genotyping/rpgg2_k21_84k/hgsvc/clinic/cdbg/v1/tr_cck_ns.pickle", 'rb') as f:
        tr_cck_ns = pickle.load(f)
    return tr_cck_ns
tr_cck_ns = test()

# %% [code]
# generate hprc_in_hgsvc.uniq_map.final.bed
# !/project/mchaisso_100/cmb-17/vntr_genotyping/rpgg2_k21_84k/hprc/analysis/gen.uniq_overlap_bed.sh
def test():
    i1i0 = np.loadtxt("/project/mchaisso_100/cmb-17/vntr_genotyping/rpgg2_k21_84k/hprc/analysis/hprc_in_hgsvc.uniq_map.final.v1.bed", usecols=[3,4], dtype=int)
    tri021, tri120 = {}, {}
    for i1, i0 in i1i0:
        tri021[i0] = i1
        tri120[i1] = i0
    return tri021, tri120
tri021, tri120 = test()

# %% [code]
def test():
    NK = len(ks)
    ng, NCCK = ilgt.shape
    pbgt = np.zeros([ng,NK], dtype=np.int32)
    for gi, gn in enumerate(np.loadtxt("/project/mchaisso_100/cmb-16/tsungyul/work/vntr/hapdb/config/genomes.HPRC.notinHGSVC.hasIL.src.txt", usecols=1, dtype=object)):
        print(".", end="")
        with open(f"/project/mchaisso_100/cmb-17/vntr_genotyping/rpgg2_k21_84k/hprc/output/{gn}.rawPB.tr.kmers") as f:
            tri = -1
            skip = True
            for line in f:
                if line[0] == ">":
                    tri += 1
                    if tri in tri120:
                        tri0 = tri120[tri]
                        skip = False
                    else:
                        skip = True
                else:
                    if not skip:
                        km, ct = [int(v) for v in line.split()]
                        if km in tri2km2i[tri0]:
                            ki = tri2km2i[tri0][km]
                            pbgt[gi,ki] = ct
    with open("/project/mchaisso_100/cmb-17/vntr_genotyping/rpgg2_k21_84k/hgsvc/1kgr/hprc/gt_pickle/pbgt.pickle", 'wb') as f:
        pickle.dump(pbgt, f, protocol=pickle.HIGHEST_PROTOCOL)
# test()

# %% [code]
def test():
    ng, NCCK = ilgt.shape
    with open("/project/mchaisso_100/cmb-17/vntr_genotyping/rpgg2_k21_84k/hgsvc/1kgr/hprc/gt_pickle/pbgt.pickle", 'rb') as f:
        pbgt = pickle.load(f)
    pbcgt = np.zeros([ng,NCCK], dtype=np.float32)
    for i0, i1 in ki_map.items():
        pbcgt[:,i1] += pbgt[:,i0]
    pbcgt /= tr_cck_ns
    with open("/project/mchaisso_100/cmb-17/vntr_genotyping/rpgg2_k21_84k/hgsvc/1kgr/hprc/gt_pickle/pbcgt.pickle", 'wb') as f:
        pickle.dump(pbcgt, f, protocol=pickle.HIGHEST_PROTOCOL)
# test()

# %% [code]
def test():
    with open("/project/mchaisso_100/cmb-17/vntr_genotyping/rpgg2_k21_84k/hgsvc/1kgr/hprc/gt_pickle/pbgt.pickle", 'rb') as f:
        pbgt = pickle.load(f)
    return pd.DataFrame(pbgt)
test()

# %% [code]
def test():
    with open("/project/mchaisso_100/cmb-17/vntr_genotyping/rpgg2_k21_84k/hgsvc/1kgr/hprc/gt_pickle/pbcgt.pickle", 'rb') as f:
        pbcgt = pickle.load(f)
    return pbcgt
pbcgt = test()
pd.DataFrame(pbcgt)

# %% [markdown]
# batch-$r^2$

# %% [code]
def get_ki(tri):
    si = ki_tr[tri-1] if tri else 0
    ei = ki_tr[tri]
    return si, ei
def get_ccki(tri):
    si = ccki_tr[tri-1] if tri else 0
    ei = ccki_tr[tri]
    return si, ei

# %% [code]
i1i0 = np.loadtxt("/project/mchaisso_100/cmb-17/vntr_genotyping/rpgg2_k21_84k/hprc/analysis/hprc_in_hgsvc.uniq_map.final.v1.bed", usecols=[3,4], dtype=int)

# %% [markdown]
## motif

# %% [code]
mtfilter = np.all(np.loadtxt("/project/mchaisso_100/cmb-17/vntr_genotyping/rpgg2_k21_84k/hgsvc/clinic/analysis/mape.invariant.flank.alu.mismap_1.r2.v2.filter.tsv.gz", usecols=[3,4,5,6,7,8,9], dtype=bool, skiprows=1), axis=1)

# %% [code]
np.sum(mtfilter), mtfilter.size

# %% [code]
def test():
    omap1 = pd.read_csv("/project/mchaisso_100/cmb-17/vntr_genotyping/rpgg2_k21_84k/hprc/output/OrthoMap.v2.tsv", sep="\t", header=None, na_values=".").values
    
    gs = np.loadtxt("/project/mchaisso_100/cmb-16/tsungyul/work/vntr/hapdb/config/genomes.HPRC.notinHGSVC.hasIL.src.txt", usecols=1, dtype=object)
    gs1 = np.loadtxt("/project/mchaisso_100/cmb-17/vntr_genotyping/rpgg2_k21_84k/hprc/input/genomes.HPRC.txt", dtype=object)
    gm1 = np.isin(gs1, gs)
    hm1 = np.repeat(gm1, 2)
    ng = gs.size
    nh = 2*ng
    ntr0 = len(pickle.load(open("/project/mchaisso_100/cmb-17/vntr_genotyping/rpgg2_k21_84k/hgsvc/clinic/cdbg/v1/ki_tr.ccki_tr.pickle", 'rb'))[0])
    ntr1 = omap1.shape[0]
    
    omap = np.full([ntr0, nh], np.nan)
    omap[i1i0[:,1]] = omap1[i1i0[:,0]][:,hm1]
    trgmask = np.ones([ntr0, ng], dtype=bool) 
    i1s = np.arange(0, nh, 2)
    trgmask &= np.isfinite(omap[:,i1s]) # hap0 not missing in TR annotation
    trgmask &= np.isfinite(omap[:,i1s+1]) # hap1 not missing in TR annotation
    return trgmask
    
trgmask = test()

# %% [code]
pd.DataFrame(trgmask)

# %% [code]
def test():
    NCCK = mtfilter.size
    nnull = 0
    nconst = 0
    r2s = np.full(NCCK, np.nan)
    ntr = len(pickle.load(open("/project/mchaisso_100/cmb-17/vntr_genotyping/rpgg2_k21_84k/hgsvc/clinic/cdbg/v1/ki_tr.ccki_tr.pickle", 'rb'))[0])
    for tri in range(ntr):
        gm = trgmask[tri]
        si, ei = get_ccki(tri)
        km = mtfilter[si:ei]
        Xs = pbcgt[:,si:ei][gm][:,km]
        Ys = ilcgt[:,si:ei][gm][:,km]
        for i1, ki in enumerate(np.nonzero(km)[0] + si):
            xs = Xs[:,i1]
            if not np.any(xs): 
                nnull += 1
                continue
            if np.unique(xs).size == 1:
                nconst += 1
                continue
            ys = Ys[:,i1]
            r2s[ki] = sm.OLS(np.insert(ys,0,0), sm.add_constant(np.insert(xs,0,0))).fit().rsquared
    print(f"{nnull} motifs not tested: not found in HPRC")
    print(f"{nconst} motifs not tested: no variation for this motif in HPRC")
    return r2s
r2s_mt_pre = test()

# %% [code]
np.savetxt("/project/mchaisso_100/cmb-17/vntr_genotyping/rpgg2_k21_84k/hgsvc/1kgr/hprc/eval/pre_correction.motif.r2.txt", r2s_mt_pre, fmt="%s")

# %% [code]
def test():
    utils.hist(r2s_mt_pre, "batch-$r^2$", "Count", bins=200)
    n0 = np.sum(r2s_mt_pre>0.8)
    n1 = np.sum(np.isfinite(r2s_mt_pre))
    n2 = np.sum(mtfilter)
    print(f"{n1:,}/{n2:,} ({n1/n2:.1%}) motifs evaluated")
    print(f"mean r2 = {np.nanmean(r2s_mt_pre):.3f}")
    print(f"{n0:,}/{n1:,} ({n0/n1:.1%}) has r2 > 0.8")
test()

# %% [code]


# %% [markdown]
## length

# %% [code]
trfilter = np.all(pd.read_csv("/project/mchaisso_100/cmb-17/vntr_genotyping/rpgg2_k21_84k/hgsvc/clinic/analysis/80518_loci.alu.mismap_1.r2.v2.filter.tsv", sep="\t", index_col=0).values[:,3:], axis=1).astype(bool)

# %% [code]
def test():
    n0 = np.sum(trfilter[i1i0[:,1]])
    n1 = np.sum(trfilter)
    print(f"{n0:,}/{n1:,} ({n0/n1:.1%}) loci can be evaluated")
test()

# %% [code]
def test():
    pbed = np.loadtxt("/project/mchaisso_100/cmb-17/vntr_genotyping/rpgg2_k21_84k/hprc/output/pan.tr.mbe.v2.bed", dtype=object, comments=None)
    ntr, ncol = pbed.shape
    nh = (ncol-3) // 4
    ng= nh // 2
    trlens = np.full([ntr, nh], np.nan, dtype=object)
    for tri in range(ntr):
        for hi in range(nh):
            if pbed[tri,3+4*hi] == "None": continue
            trlens[tri,hi] = int(pbed[tri,5+4*hi]) - int(pbed[tri,4+4*hi])
    
    gs = np.loadtxt("/project/mchaisso_100/cmb-17/vntr_genotyping/rpgg2_k21_84k/hprc/input/genomes.HPRC.txt", dtype=object)
    hs = np.array([f"{g}.{h}" for g in gs for h in [0,1]])
    trs = np.array([f"{ch}:{s}-{e}" for ch, s, e in pbed[:,:3]])
    df = pd.DataFrame(trlens, columns=hs, index=trs)
    df.to_csv(f"/project/mchaisso_100/cmb-17/vntr_genotyping/rpgg2_k21_84k/hprc/analysis/trlens.{ntr}x{nh}.txt", sep="\t", na_rep=".")
    
    atrlens = np.full([ntr,ng], np.nan, dtype=object)
    for gi in range(ng):
        atrlens[:,gi] = np.mean(trlens[:,2*gi:2*gi+2], axis=1)
    df = pd.DataFrame(atrlens, columns=gs, index=trs)
    df.to_csv(f"/project/mchaisso_100/cmb-17/vntr_genotyping/rpgg2_k21_84k/hprc/analysis/trlens.{ntr}x{ng}.txt", sep="\t", na_rep=".")
# test()

# %% [code]
def test():
    gs = np.loadtxt("/project/mchaisso_100/cmb-16/tsungyul/work/vntr/hapdb/config/genomes.HPRC.notinHGSVC.hasIL.src.txt", usecols=1, dtype=object)
    gs1 = np.loadtxt("/project/mchaisso_100/cmb-17/vntr_genotyping/rpgg2_k21_84k/hprc/input/genomes.HPRC.txt", dtype=object)
    gm1 = np.isin(gs1, gs)
    
    ng = gs.size
    ntr = len(ki_tr)

    trlens_ = pd.read_csv(f"/project/mchaisso_100/cmb-17/vntr_genotyping/rpgg2_k21_84k/hprc/analysis/trlens.81261x48.txt", sep="\t", na_values=".", index_col=0).values    
    trlens = np.full([ntr,ng], np.nan)
    trlens[i1i0[:,1]] = trlens_[i1i0[:,0]][:,gm1]
    return trlens
trlens = test()

# %% [code]
def test():
    trdsg = np.zeros_like(trlens)
    for tri in i1i0[:,1]:
        si, ei = get_ki(tri)
        trdsg[tri] = np.sum(ilgt[:,si:ei], axis=1)
    return trdsg
trdsg = test()

# %% [code]
def test():
    nscan, nconst = 0, 0
    ntr = len(ki_tr)
    r2s = np.full(ntr, np.nan)
    for tri in i1i0[:,1]:
        if not trfilter[tri]: continue
        nscan += 1
        xs = trlens[tri]
        m = np.isfinite(xs)
        xs = xs[m]
        xs = np.clip(xs-20, 0, None)
        if not np.any(xs): 
            assert False
            continue
        if np.unique(xs).size == 1:
            nconst += 1
            continue
        ys = trdsg[tri,m]
        r2s[tri] = sm.OLS(np.insert(ys,0,0), sm.add_constant(np.insert(xs,0,0))).fit().rsquared
        
    n0 = np.sum(r2s>0.8)
    n1 = nscan - nconst
    n2 = nscan
    print(f"{nscan} TRs scanned")
    print(f"{nconst} TRs not tested: no variation in length for this TR in HPRC")
    print(f"{n1:,}/{n2:,} ({n1/n2:.1%}) TRs evaluated")
    print(f"mean r2 = {np.nanmean(r2s):.3f}")
    print(f"{n0:,}/{n1:,} ({n0/n1:.1%}) has r2 > 0.8")
    utils.hist(r2s, "batch-$r^2$", "Count", bins=200)
    
    return r2s
r2s_len_pre = test()

# %% [code]
np.savetxt("/project/mchaisso_100/cmb-17/vntr_genotyping/rpgg2_k21_84k/hgsvc/1kgr/hprc/eval/pre_correction.length.r2.txt", r2s_len_pre, fmt="%s")

# %% [code]


# %% [markdown]
# batch-$r^2$ after bias correction

# %% [code]
def test():
    with open("/project/mchaisso_100/cmb-17/vntr_genotyping/rpgg2_k21_84k/hgsvc/clinic/bias/invariant_kmer/ikmer.ikstats.pickle", 'rb') as f:
        ikmer, ikstats = pickle.load(f)
    return ikmer, ikstats
ikmer, ikstats = test()

# %% [code]
def test():
    ng, NCCK = ilgt.shape
    ntr = len(ki_tr)
    
    bias = np.zeros([ng, ntr])
    with open(f"/project/mchaisso_100/cmb-17/vntr_genotyping/rpgg2_k21_84k/hgsvc/1kgr/hprc/gt_pickle/agt.pickle", 'rb') as f:
        agt = pickle.load(f)
    for tri in range(ntr):
        if tri % 1000 == 0:
            print(".", end="")
        si, ei = get_ki(tri)
        inv = np.nonzero(ikmer[si:ei])[0] + si
        if inv.size:
            bias[:,tri] = np.mean(agt[:,inv] / ikstats[0,inv], axis=1)
    print()
    bias[bias==0] = np.nan
    bias /= np.nanmean(bias, axis=0)
    n0 = np.sum(~np.isfinite(bias))
    print(f"{n0} cells of nan's")
    return bias

bias = test()

# %% [code]
pd.DataFrame(bias)

# %% [code]
def test():
    with open("/project/mchaisso_100/cmb-17/vntr_genotyping/rpgg2_k21_84k/hgsvc/1kgr/hprc/bias/bias.v1.pickle", 'wb') as f:
        pickle.dump(bias, f)
# test()

# %% [code]
def test():
    ntr = len(ccki_tr)
    ilcgt_bc = np.copy(ilcgt)
    for tri in range(ntr):
        si, ei = get_ccki(tri)
        ilcgt_bc[:,si:ei] /= (bias[:,tri][:,None])
    return ilcgt_bc
ilcgt_bc = test()

# %% [code]
pd.DataFrame(ilcgt_bc)

# %% [code]
def test():
    with open("/project/mchaisso_100/cmb-17/vntr_genotyping/rpgg2_k21_84k/hgsvc/1kgr/hprc/gt_pickle/acgt_bc.pickle", 'wb') as f:
        pickle.dump(ilcgt_bc, f, protocol=pickle.HIGHEST_PROTOCOL)
# test()

# %% [code]
def test():
    with open("/project/mchaisso_100/cmb-17/vntr_genotyping/rpgg2_k21_84k/hgsvc/1kgr/hprc/gt_pickle/acgt_bc.pickle", 'rb') as f:
        return pickle.load(f)
ilcgt_bc = test()

# %% [code]
def test():
    ntr = len(ki_tr)
    ilgt_bc = np.copy(ilgt)
    for tri in range(ntr):
        si, ei = get_ki(tri)
        ilgt_bc[:,si:ei] /= (bias[:,tri][:,None])
    return ilgt_bc
ilgt_bc = test()

# %% [code]
pd.DataFrame(ilgt_bc)

# %% [code]
def test():
    with open("/project/mchaisso_100/cmb-17/vntr_genotyping/rpgg2_k21_84k/hgsvc/1kgr/hprc/gt_pickle/agt_bc.pickle", 'wb') as f:
        pickle.dump(ilgt_bc, f, protocol=pickle.HIGHEST_PROTOCOL)
# test()

# %% [code]
def test():
    with open("/project/mchaisso_100/cmb-17/vntr_genotyping/rpgg2_k21_84k/hgsvc/1kgr/hprc/gt_pickle/agt_bc.pickle", 'rb') as f:
        return pickle.load(f)
ilgt_bc = test()

# %% [code]


# %% [markdown]
## motif

# %% [code]
def test():
    NCCK = mtfilter.size
    nnull = 0
    nconst = 0
    ndrop = 0
    r2s = np.full(NCCK, np.nan)
    ntr = len(ccki_tr)
    for tri in range(ntr):
        gm = trgmask[tri]
        si, ei = get_ccki(tri)
        km = mtfilter[si:ei]
        Xs = pbcgt[:,si:ei][gm][:,km]
        Ys = ilcgt_bc[:,si:ei][gm][:,km]
        for i1, ki in enumerate(np.nonzero(km)[0] + si):
            xs = Xs[:,i1]
            if not np.any(xs): 
                nnull += 1
                continue
            if np.unique(xs).size == 1:
                nconst += 1
                continue
            ys = Ys[:,i1]
            if not np.any(np.isfinite(ys)):
                ndrop += 1
            r2s[ki] = sm.OLS(np.insert(ys,0,0), sm.add_constant(np.insert(xs,0,0))).fit().rsquared
    print(f"{nnull} motifs not tested: not found in HPRC")
    print(f"{nconst} motifs not tested: no variation for this motif in HPRC")
    print(f"{ndrop} motifs dropped during bias correction")
    return r2s
r2s_mt_post = test()

# %% [code]
np.savetxt("/project/mchaisso_100/cmb-17/vntr_genotyping/rpgg2_k21_84k/hgsvc/1kgr/hprc/eval/post_correction.motif.r2.txt", r2s_mt_post, fmt="%s")

# %% [code]
def test():
    utils.hist(r2s_mt_post, "batch-$r^2$", "Count", bins=200)
    n0 = np.sum(r2s_mt_post>0.8)
    n1 = np.sum(np.isfinite(r2s_mt_post))
    n2 = np.sum(mtfilter)
    print(f"{n1:,}/{n2:,} ({n1/n2:.1%}) motifs evaluated")
    print(f"mean r2 = {np.nanmean(r2s_mt_post):.3f}")
    print(f"{n0:,}/{n1:,} ({n0/n1:.1%}) has r2 > 0.8")
test()

# %% [code]


# %% [markdown]
## length

# %% [code]
def test():
    trdsg_bc = np.zeros_like(trlens)
    for tri in i1i0[:,1]:
        si, ei = get_ki(tri)
        trdsg_bc[tri] = np.sum(ilgt_bc[:,si:ei], axis=1)
    return trdsg_bc
trdsg_bc = test()

# %% [code]
def test():
    nscan, nconst, ndrop = 0, 0, 0
    ntr = len(ki_tr)
    r2s = np.full(ntr, np.nan)
    for tri in i1i0[:,1]:
        if not trfilter[tri]: continue
        nscan += 1
        xs = trlens[tri]
        ys = trdsg_bc[tri]
        if not np.any(np.isfinite(ys)):
            ndrop += 1
            continue
        m = np.isfinite(xs) & np.isfinite(ys)
        xs = xs[m]
        xs = np.clip(xs-20, 0, None)
        if not np.any(xs): 
            assert False
            continue
        if np.unique(xs).size == 1:
            nconst += 1
            continue
        ys = ys[m]
        r2s[tri] = sm.OLS(np.insert(ys,0,0), sm.add_constant(np.insert(xs,0,0))).fit().rsquared
        assert np.isfinite(r2s[tri]), print(tri, r2s[tri])
        
    n0 = np.sum(r2s>0.8)
    n1 = nscan - nconst
    n2 = nscan
    print(f"{nscan} TRs scanned")
    print(f"{nconst} TRs not tested: no variation in length for this TR in HPRC")
    print(f"{ndrop} TRs dropped during bias correction")
    print(f"{n1:,}/{n2:,} ({n1/n2:.1%}) TRs evaluated")
    print(f"mean r2 = {np.nanmean(r2s):.3f}")
    print(f"{n0:,}/{n1:,} ({n0/n1:.1%}) has r2 > 0.8")
    utils.hist(r2s, "batch-$r^2$", "Count", bins=200)
    
    return r2s
r2s_len_post = test()

# %% [code]
np.savetxt("/project/mchaisso_100/cmb-17/vntr_genotyping/rpgg2_k21_84k/hgsvc/1kgr/hprc/eval/post_correction.length.r2.txt", r2s_len_post, fmt="%s")

# %% [code]


# %% [markdown]
# Summary

# %% [code]
def test():
    r20 = np.loadtxt("/project/mchaisso_100/cmb-17/vntr_genotyping/rpgg2_k21_84k/hgsvc/1kgr/hprc/eval/pre_correction.motif.r2.txt")
    r21 = np.loadtxt("/project/mchaisso_100/cmb-17/vntr_genotyping/rpgg2_k21_84k/hgsvc/1kgr/hprc/eval/post_correction.motif.r2.txt")
    n0 = np.sum(np.isfinite(r21))
    n1 = np.sum(mtfilter)
    print(f"{n0:,}/{n1:,} ({n0/n1:.1%}) motifs evaluated")
    print(f"{np.nanmean(r20):.3f} -> {np.nanmean(r21):.3f}")
    xl = [np.nanmin(r20), np.nanmax(r20)]
    plt.figure(dpi=300, figsize=(2,1.8))
    plt.plot(r20, r21, '.k', mew=0, markersize=1, alpha=0.002)
    plt.plot(xl, xl, '--r', lw=0.4)
    plt.xlabel("batch-$r^2$ (before)")
    plt.ylabel("batch-$r^2$ (after)")
    plt.show(); plt.close()
    
    ntr = len(ccki_tr)
    r20tr = np.full(ntr, np.nan)
    r21tr = np.full(ntr, np.nan)
    for tri in range(ntr):
        si, ei = get_ccki(tri)
        r20tr[tri] = np.nanmean(r20[si:ei])
        r21tr[tri] = np.nanmean(r21[si:ei])
    n0 = np.sum(np.isfinite(r21tr))
    n1 = np.sum(trfilter)
    print(f"{n0:,}/{n1:,} ({n0/n1:.1%}) loci evaluated")
    print(f"{np.nanmean(r20tr):.3f} -> {np.nanmean(r21tr):.3f}")
    xl = [np.nanmin(r20tr), np.nanmax(r20tr)]
    plt.figure(dpi=300, figsize=(2,1.8))
    plt.plot(r20tr, r21tr, '.k', mew=0, markersize=1, alpha=0.1)
    plt.plot(xl, xl, '--r', lw=0.4)
    plt.xlabel("batch-$r^2$ (before, locus ave.)")
    plt.ylabel("batch-$r^2$ (after, locus ave.)")
    plt.show(); plt.close()

test()

# %% [code]
# allow y == 0
def test():
    r20 = np.loadtxt("/project/mchaisso_100/cmb-17/vntr_genotyping/rpgg2_k21_84k/hgsvc/1kgr/hprc/eval/pre_correction.length.r2.txt")
    r21 = np.loadtxt("/project/mchaisso_100/cmb-17/vntr_genotyping/rpgg2_k21_84k/hgsvc/1kgr/hprc/eval/post_correction.length.r2.txt")
    
    n0 = np.sum(np.isfinite(r21))
    n1 = np.sum(trfilter)
    print(f"{n0:,}/{n1:,} ({n0/n1:.1%}) loci evaluated")
    print(f"{np.nanmean(r20):.3f} -> {np.nanmean(r21):.3f}")
    xl = [np.nanmin(r20), np.nanmax(r20)]
    plt.figure(dpi=300, figsize=(2,1.8))
    plt.plot(r20, r21, '.k', mew=0, markersize=1, alpha=0.1)
    plt.plot(xl, xl, '--r', lw=0.4)
    plt.xlabel("batch-$r^2$ (before)")
    plt.ylabel("batch-$r^2$ (after)")
    plt.show(); plt.close()
test()


# %% [code]
def test():
    r2hgsvc = np.loadtxt("/project/mchaisso_100/cmb-17/vntr_genotyping/rpgg2_k21_84k/hgsvc/clinic/bias/invariant_kmer/output2/R2_all_loci.R2a_all_loci.txt", usecols=1)
    r2hprc = np.loadtxt("/project/mchaisso_100/cmb-17/vntr_genotyping/rpgg2_k21_84k/hgsvc/1kgr/hprc/eval/post_correction.length.r2.txt")
    
    df = pd.DataFrame(r2hgsvc, columns=["HGSVC batch-$r^2$"])
    df["HPRC batch-$r^2$"] = r2hprc
    
    n0 = np.sum((np.isfinite(r2hgsvc) & np.isfinite(r2hprc))[trfilter])
    n1 = r2hgsvc[trfilter].size
    print(f"{n0:,}/{n1:,} ({n0/n1:.1%}) VNTRs evaluated")
    print(f"mean r2 = {np.nanmean(r2hgsvc):.3f} / {np.nanmean(r2hprc):.3f} in HGSVC / HPRC")

    gr = sns.JointGrid(data=df, y="HPRC batch-$r^2$", x="HGSVC batch-$r^2$", height=2.5)
    gr.fig.set(dpi=600)
    gr.plot_joint(sns.scatterplot, s=.1, alpha=.2, color='k', linewidth=0)
    xmin, xmax = gr.ax_joint.get_xlim()
    ymin, ymax = gr.ax_joint.get_ylim()
    js = [min(xmin, ymin), max(xmax, ymax)]
    gr.ax_joint.set_xlim(js)
    gr.ax_joint.set_ylim(js)
    ds = [min(np.nanmin(r2hgsvc),np.nanmin(r2hprc)), max(np.nanmax(r2hgsvc),np.nanmax(r2hprc))]
    gr.ax_joint.plot(ds, ds, '--r', lw=0.3)
    gr.plot_marginals(sns.histplot, bins=100, fill=False, linewidth=0.2)
    
    gr.ax_marg_y.grid(True, axis='x', ls=':', lw=0.4)
    gr.ax_marg_y.tick_params(labeltop=True, labelsize=3)
    gr.ax_marg_x.grid(True, axis='y', ls=':', lw=0.4)
    gr.ax_marg_x.tick_params(labelleft=True, labelsize=3)
    
    
test()

# %% [code]
def test():
    r2hprc = np.loadtxt("/project/mchaisso_100/cmb-17/vntr_genotyping/rpgg2_k21_84k/hgsvc/1kgr/hprc/eval/post_correction.motif.r2.txt")
    with open(f"/project/mchaisso_100/cmb-17/vntr_genotyping/rpgg2_k21_84k/hgsvc/clinic/bias/invariant_kmer/output2/R2_all_mtf.R2a_all_mtf.pickle", 'rb') as f:
        _, r2hgsvc = pickle.load(f)
    df = pd.DataFrame(r2hgsvc, columns=["HGSVC batch-$r^2$"])
    df["HPRC batch-$r^2$"] = r2hprc
    
    n0 = np.sum((np.isfinite(r2hgsvc) & np.isfinite(r2hprc))[mtfilter])
    n1 = r2hgsvc[mtfilter].size
    print(f"{n0:,}/{n1:,} ({n0/n1:.1%}) motifs evaluated")
    print(f"mean r2 = {np.nanmean(r2hgsvc):.3f} / {np.nanmean(r2hprc):.3f} in HGSVC / HPRC")

    gr = sns.JointGrid(data=df, y="HPRC batch-$r^2$", x="HGSVC batch-$r^2$", height=2.5)
    gr.fig.set(dpi=600)
    gr.plot_joint(sns.scatterplot, s=.1, alpha=.05, color='k', linewidth=0)
    xmin, xmax = gr.ax_joint.get_xlim()
    ymin, ymax = gr.ax_joint.get_ylim()
    js = [min(xmin, ymin), max(xmax, ymax)]
    gr.ax_joint.set_xlim(js)
    gr.ax_joint.set_ylim(js)
    ds = [min(np.nanmin(r2hgsvc),np.nanmin(r2hprc)), max(np.nanmax(r2hgsvc),np.nanmax(r2hprc))]
    gr.ax_joint.plot(ds, ds, '--r', lw=0.5)
    gr.plot_marginals(sns.histplot, bins=100, fill=False, linewidth=0.2)
test()

# %% [markdown]
## eMotifs

# %% [code]
def test():
    df = pd.read_csv("/project/mchaisso_100/cmb-17/vntr_genotyping/rpgg2_k21_84k/hgsvc/gtex/emotif/analysis10/Alltissue.egenes.tsv", sep="\t", index_col=0)
    kis = np.sort(list(set(df["lead_ki"])))
    r2hprc = np.loadtxt("/project/mchaisso_100/cmb-17/vntr_genotyping/rpgg2_k21_84k/hgsvc/1kgr/hprc/eval/post_correction.motif.r2.txt")[kis]
    
    with open(f"/project/mchaisso_100/cmb-17/vntr_genotyping/rpgg2_k21_84k/hgsvc/clinic/bias/invariant_kmer/output2/R2_all_mtf.R2a_all_mtf.pickle", 'rb') as f:
        _, r2hgsvc = pickle.load(f)
    r2hgsvc = r2hgsvc[kis]
    df = pd.DataFrame(r2hgsvc, columns=["HGSVC batch-$r^2$"])
    df["HPRC batch-$r^2$"] = r2hprc
    
    n0 = np.sum(np.isfinite(r2hgsvc) & np.isfinite(r2hprc))
    n1 = r2hgsvc.size
    print(f"{n0:,}/{n1:,} ({n0/n1:.1%}) eMotifs evaluated")
    print(f"mean r2 = {np.nanmean(r2hgsvc):.3f} / {np.nanmean(r2hprc):.3f} in HGSVC / HPRC")

    gr = sns.JointGrid(data=df, y="HPRC batch-$r^2$", x="HGSVC batch-$r^2$", height=2.5)
    gr.fig.set(dpi=600)
    gr.plot_joint(sns.scatterplot, s=0.3, alpha=.2, color='k', linewidth=0)
    xmin, xmax = gr.ax_joint.get_xlim()
    ymin, ymax = gr.ax_joint.get_ylim()
    js = [min(xmin, ymin), max(xmax, ymax)]
    gr.ax_joint.set_xlim(js)
    gr.ax_joint.set_ylim(js)
    ds = [min(min(r2hgsvc),min(r2hprc)), max(max(r2hgsvc),max(r2hprc))]
    gr.ax_joint.plot(ds, ds, '--r', lw=0.5)
    gr.plot_marginals(sns.histplot, bins=100, fill=False, linewidth=0.2)
    
    
test()

# %% [markdown]
## likely causal motifs

# %% [code]
def test():
    df = pd.read_csv("/project/mchaisso_100/cmb-17/vntr_genotyping/rpgg2_k21_84k/hgsvc/gtex/susieR/analysis10/Alltissue.egenes.finemap.validated.tsv", sep="\t", index_col=0)
    kis = set()
    for kistr in df["ki"]:
        for ki in kistr.split(","):
            if ki == "-1": continue
            kis.add(int(ki))
    kis = np.sort(list(kis))
    r2hprc = np.loadtxt("/project/mchaisso_100/cmb-17/vntr_genotyping/rpgg2_k21_84k/hgsvc/1kgr/hprc/eval/post_correction.motif.r2.txt")[kis]
    
    
    with open(f"/project/mchaisso_100/cmb-17/vntr_genotyping/rpgg2_k21_84k/hgsvc/clinic/bias/invariant_kmer/output2/R2_all_mtf.R2a_all_mtf.pickle", 'rb') as f:
        _, r2hgsvc = pickle.load(f)
    r2hgsvc = r2hgsvc[kis]
    df = pd.DataFrame(r2hgsvc, columns=["HGSVC batch-$r^2$"])
    df["HPRC batch-$r^2$"] = r2hprc
    
    n0 = np.sum(np.isfinite(r2hgsvc) & np.isfinite(r2hprc))
    n1 = r2hgsvc.size
    print(f"{n0:,}/{n1:,} ({n0/n1:.1%}) likely causal motifs evaluated")
    print(f"mean r2 = {np.nanmean(r2hgsvc):.3f} / {np.nanmean(r2hprc):.3f} in HGSVC / HPRC")

    # dmin = min(min(r2hgsvc), min(r2hprc)) - 0.03
    # dmax = max(max(r2hgsvc), max(r2hprc)) + 0.03
    
    gr = sns.JointGrid(data=df, y="HPRC batch-$r^2$", x="HGSVC batch-$r^2$", height=2.5)
    gr.fig.set(dpi=600)
    gr.plot_joint(sns.scatterplot, s=1, alpha=0.6, color='k', linewidth=0)
    xmin, xmax = gr.ax_joint.get_xlim()
    ymin, ymax = gr.ax_joint.get_ylim()
    js = [min(xmin, ymin), max(xmax, ymax)]
    gr.ax_joint.set_xlim(js)
    gr.ax_joint.set_ylim(js)
    ds = [min(min(r2hgsvc),min(r2hprc)), max(max(r2hgsvc),max(r2hprc))]
    gr.ax_joint.plot(ds, ds, '--r', lw=0.5)
    gr.plot_marginals(sns.histplot, bins=100, fill=False, linewidth=0.2)    
    
test()

# %% [code]
```
