## Supplemental code for "The motif composition of variable-number tandem repeats impacts gene expression": access.hpp

```
/*! \file access.hpp
    \brief Access control and default construction */
/*
  Copyright (c) 2014, Randolph Voorhies, Shane Grant
  All rights reserved.

  Redistribution and use in source and binary forms, with or without
  modification, are permitted provided that the following conditions are met:
      * Redistributions of source code must retain the above copyright
        notice, this list of conditions and the following disclaimer.
      * Redistributions in binary form must reproduce the above copyright
        notice, this list of conditions and the following disclaimer in the
        documentation and/or other materials provided with the distribution.
      * Neither the name of cereal nor the
        names of its contributors may be used to endorse or promote products
        derived from this software without specific prior written permission.

  THIS SOFTWARE IS PROVIDED BY THE COPYRIGHT HOLDERS AND CONTRIBUTORS "AS IS" AND
  ANY EXPRESS OR IMPLIED WARRANTIES, INCLUDING, BUT NOT LIMITED TO, THE IMPLIED
  WARRANTIES OF MERCHANTABILITY AND FITNESS FOR A PARTICULAR PURPOSE ARE
  DISCLAIMED. IN NO EVENT SHALL RANDOLPH VOORHIES OR SHANE GRANT BE LIABLE FOR ANY
  DIRECT, INDIRECT, INCIDENTAL, SPECIAL, EXEMPLARY, OR CONSEQUENTIAL DAMAGES
  (INCLUDING, BUT NOT LIMITED TO, PROCUREMENT OF SUBSTITUTE GOODS OR SERVICES;
  LOSS OF USE, DATA, OR PROFITS; OR BUSINESS INTERRUPTION) HOWEVER CAUSED AND
  ON ANY THEORY OF LIABILITY, WHETHER IN CONTRACT, STRICT LIABILITY, OR TORT
  (INCLUDING NEGLIGENCE OR OTHERWISE) ARISING IN ANY WAY OUT OF THE USE OF THIS
  SOFTWARE, EVEN IF ADVISED OF THE POSSIBILITY OF SUCH DAMAGE.
*/
#ifndef CEREAL_ACCESS_HPP_
#define CEREAL_ACCESS_HPP_

#include <type_traits>
#include <iostream>
#include <cstdint>
#include <functional>

#include "cereal/macros.hpp"
#include "cereal/specialize.hpp"
#include "cereal/details/helpers.hpp"

namespace cereal
{
  // ######################################################################
  //! A class that allows cereal to load smart pointers to types that have no default constructor
  /*! If your class does not have a default constructor, cereal will not be able
      to load any smart pointers to it unless you overload LoadAndConstruct
      for your class, and provide an appropriate load_and_construct method.  You can also
      choose to define a member static function instead of specializing this class.

      The specialization of LoadAndConstruct must be placed within the cereal namespace:

      @code{.cpp}
      struct MyType
      {
        MyType( int x ); // note: no default ctor
        int myX;

        // Define a serialize or load/save pair as you normally would
        template <class Archive>
        void serialize( Archive & ar )
        {
          ar( myX );
        }
      };

      // Provide a specialization for LoadAndConstruct for your type
      namespace cereal
      {
        template <> struct LoadAndConstruct<MyType>
        {
          // load_and_construct will be passed the archive that you will be loading
          // from as well as a construct object which you can use as if it were the
          // constructor for your type.  cereal will handle all memory management for you.
          template <class Archive>
          static void load_and_construct( Archive & ar, cereal::construct<MyType> & construct )
          {
            int x;
            ar( x );
            construct( x );
          }

          // if you require versioning, simply add a const std::uint32_t as the final parameter, e.g.:
          // load_and_construct( Archive & ar, cereal::construct<MyType> & construct, std::uint32_t const version )
        };
      } // end namespace cereal
      @endcode

      Please note that just as in using external serialization functions, you cannot get
      access to non-public members of your class by befriending cereal::access.  If you
      have the ability to modify the class you wish to serialize, it is recommended that you
      use member serialize functions and a static member load_and_construct function.

      load_and_construct functions, regardless of whether they are static members of your class or
      whether you create one in the LoadAndConstruct specialization, have the following signature:

      @code{.cpp}
      // generally Archive will be templated, but it can be specific if desired
      template <class Archive>
      static void load_and_construct( Archive & ar, cereal::construct<MyType> & construct );
      // with an optional last parameter specifying the version: const std::uint32_t version
      @endcode

      Versioning behaves the same way as it does for standard serialization functions.

      @tparam T The type to specialize for
      @ingroup Access */
  template <class T>
  struct LoadAndConstruct
  { };

  // forward decl for construct
  //! @cond PRIVATE_NEVERDEFINED
  namespace memory_detail{ template <class Ar, class T> struct LoadAndConstructLoadWrapper; }
  namespace boost_variant_detail{ template <class Ar, class T> struct LoadAndConstructLoadWrapper; }
  //! @endcond

  //! Used to construct types with no default constructor
  /*! When serializing a type that has no default constructor, cereal
      will attempt to call either the class static function load_and_construct
      or the appropriate template specialization of LoadAndConstruct.  cereal
      will pass that function a reference to the archive as well as a reference
      to a construct object which should be used to perform the allocation once
      data has been appropriately loaded.

      @code{.cpp}
      struct MyType
      {
        // note the lack of default constructor
        MyType( int xx, int yy );

        int x, y;
        double notInConstructor;

        template <class Archive>
        void serialize( Archive & ar )
        {
          ar( x, y );
          ar( notInConstructor );
        }

        template <class Archive>
        static void load_and_construct( Archive & ar, cereal::construct<MyType> & construct )
        {
          int x, y;
          ar( x, y );

          // use construct object to initialize with loaded data
          construct( x, y );

          // access to member variables and functions via -> operator
          ar( construct->notInConstructor );

          // could also do the above section by:
          double z;
          ar( z );
          construct->notInConstructor = z;
        }
      };
      @endcode

      @tparam T The class type being serialized
      */
  template <class T>
  class construct
  {
    public:
      //! Construct and initialize the type T with the given arguments
      /*! This will forward all arguments to the underlying type T,
          calling an appropriate constructor.

          Calling this function more than once will result in an exception
          being thrown.

          @param args The arguments to the constructor for T
          @throw Exception If called more than once */
      template <class ... Args>
      void operator()( Args && ... args );
      // implementation deferred due to reliance on cereal::access

      //! Get a reference to the initialized underlying object
      /*! This must be called after the object has been initialized.

          @return A reference to the initialized object
          @throw Exception If called before initialization */
      T * operator->()
      {
        if( !itsValid )
          throw Exception("Object must be initialized prior to accessing members");

        return itsPtr;
      }

      //! Returns a raw pointer to the initialized underlying object
      /*! This is mainly intended for use with passing an instance of
          a constructed object to cereal::base_class.

          It is strongly recommended to avoid using this function in
          any other circumstance.

          @return A raw pointer to the initialized type */
      T * ptr()
      {
        return operator->();
      }

    private:
      template <class Ar, class TT> friend struct ::cereal::memory_detail::LoadAndConstructLoadWrapper;
      template <class Ar, class TT> friend struct ::cereal::boost_variant_detail::LoadAndConstructLoadWrapper;

      construct( T * p ) : itsPtr( p ), itsEnableSharedRestoreFunction( [](){} ), itsValid( false ) {}
      construct( T * p, std::function<void()> enableSharedFunc ) : // g++4.7 ice with default lambda to std func
        itsPtr( p ), itsEnableSharedRestoreFunction( enableSharedFunc ), itsValid( false ) {}
      construct( construct const & ) = delete;
      construct & operator=( construct const & ) = delete;

      T * itsPtr;
      std::function<void()> itsEnableSharedRestoreFunction;
      bool itsValid;
  };

  // ######################################################################
  //! A class that can be made a friend to give cereal access to non public functions
  /*! If you desire non-public serialization functions within a class, cereal can only
      access these if you declare cereal::access a friend.

      @code{.cpp}
      class MyClass
      {
        private:
          friend class cereal::access; // gives access to the private serialize

          template <class Archive>
          void serialize( Archive & ar )
          {
            // some code
          }
      };
      @endcode
      @ingroup Access */
  class access
  {
    public:
      // ####### Standard Serialization ########################################
      template<class Archive, class T> inline
      static auto member_serialize(Archive & ar, T & t) -> decltype(t.CEREAL_SERIALIZE_FUNCTION_NAME(ar))
      { return t.CEREAL_SERIALIZE_FUNCTION_NAME(ar); }

      template<class Archive, class T> inline
      static auto member_save(Archive & ar, T const & t) -> decltype(t.CEREAL_SAVE_FUNCTION_NAME(ar))
      { return t.CEREAL_SAVE_FUNCTION_NAME(ar); }

      template<class Archive, class T> inline
      static auto member_save_non_const(Archive & ar, T & t) -> decltype(t.CEREAL_SAVE_FUNCTION_NAME(ar))
      { return t.CEREAL_SAVE_FUNCTION_NAME(ar); }

      template<class Archive, class T> inline
      static auto member_load(Archive & ar, T & t) -> decltype(t.CEREAL_LOAD_FUNCTION_NAME(ar))
      { return t.CEREAL_LOAD_FUNCTION_NAME(ar); }

      template<class Archive, class T> inline
      static auto member_save_minimal(Archive const & ar, T const & t) -> decltype(t.CEREAL_SAVE_MINIMAL_FUNCTION_NAME(ar))
      { return t.CEREAL_SAVE_MINIMAL_FUNCTION_NAME(ar); }

      template<class Archive, class T> inline
      static auto member_save_minimal_non_const(Archive const & ar, T & t) -> decltype(t.CEREAL_SAVE_MINIMAL_FUNCTION_NAME(ar))
      { return t.CEREAL_SAVE_MINIMAL_FUNCTION_NAME(ar); }

      template<class Archive, class T, class U> inline
      static auto member_load_minimal(Archive const & ar, T & t, U && u) -> decltype(t.CEREAL_LOAD_MINIMAL_FUNCTION_NAME(ar, std::forward<U>(u)))
      { return t.CEREAL_LOAD_MINIMAL_FUNCTION_NAME(ar, std::forward<U>(u)); }

      // ####### Versioned Serialization #######################################
      template<class Archive, class T> inline
      static auto member_serialize(Archive & ar, T & t, const std::uint32_t version ) -> decltype(t.CEREAL_SERIALIZE_FUNCTION_NAME(ar, version))
      { return t.CEREAL_SERIALIZE_FUNCTION_NAME(ar, version); }

      template<class Archive, class T> inline
      static auto member_save(Archive & ar, T const & t, const std::uint32_t version ) -> decltype(t.CEREAL_SAVE_FUNCTION_NAME(ar, version))
      { return t.CEREAL_SAVE_FUNCTION_NAME(ar, version); }

      template<class Archive, class T> inline
      static auto member_save_non_const(Archive & ar, T & t, const std::uint32_t version ) -> decltype(t.CEREAL_SAVE_FUNCTION_NAME(ar, version))
      { return t.CEREAL_SAVE_FUNCTION_NAME(ar, version); }

      template<class Archive, class T> inline
      static auto member_load(Archive & ar, T & t, const std::uint32_t version ) -> decltype(t.CEREAL_LOAD_FUNCTION_NAME(ar, version))
      { return t.CEREAL_LOAD_FUNCTION_NAME(ar, version); }

      template<class Archive, class T> inline
      static auto member_save_minimal(Archive const & ar, T const & t, const std::uint32_t version) -> decltype(t.CEREAL_SAVE_MINIMAL_FUNCTION_NAME(ar, version))
      { return t.CEREAL_SAVE_MINIMAL_FUNCTION_NAME(ar, version); }

      template<class Archive, class T> inline
      static auto member_save_minimal_non_const(Archive const & ar, T & t, const std::uint32_t version) -> decltype(t.CEREAL_SAVE_MINIMAL_FUNCTION_NAME(ar, version))
      { return t.CEREAL_SAVE_MINIMAL_FUNCTION_NAME(ar, version); }

      template<class Archive, class T, class U> inline
      static auto member_load_minimal(Archive const & ar, T & t, U && u, const std::uint32_t version) -> decltype(t.CEREAL_LOAD_MINIMAL_FUNCTION_NAME(ar, std::forward<U>(u), version))
      { return t.CEREAL_LOAD_MINIMAL_FUNCTION_NAME(ar, std::forward<U>(u), version); }

      // ####### Other Functionality ##########################################
      // for detecting inheritance from enable_shared_from_this
      template <class T> inline
      static auto shared_from_this(T & t) -> decltype(t.shared_from_this());

      // for placement new
      template <class T, class ... Args> inline
      static void construct( T *& ptr, Args && ... args )
      {
        new (ptr) T( std::forward<Args>( args )... );
      }

      // for non-placement new with a default constructor
      template <class T> inline
      static T * construct()
      {
        return new T();
      }

      template <class T> inline
      static std::false_type load_and_construct(...)
      { return std::false_type(); }

      template<class T, class Archive> inline
      static auto load_and_construct(Archive & ar, ::cereal::construct<T> & construct) -> decltype(T::load_and_construct(ar, construct))
      {
        T::load_and_construct( ar, construct );
      }

      template<class T, class Archive> inline
      static auto load_and_construct(Archive & ar, ::cereal::construct<T> & construct, const std::uint32_t version) -> decltype(T::load_and_construct(ar, construct, version))
      {
        T::load_and_construct( ar, construct, version );
      }
  }; // end class access

  // ######################################################################
  // Deferred Implementation, see construct for more information
  template <class T> template <class ... Args> inline
  void construct<T>::operator()( Args && ... args )
  {
    if( itsValid )
      throw Exception("Attempting to construct an already initialized object");

    ::cereal::access::construct( itsPtr, std::forward<Args>( args )... );
    itsEnableSharedRestoreFunction();
    itsValid = true;
  }
} // namespace cereal

#endif // CEREAL_ACCESS_HPP_
```
