## Supplemental code for "The motif composition of variable-number tandem repeats impacts gene expression": adapters.hpp

```
/*! \file adapters.hpp
    \brief Archive adapters that provide additional functionality
           on top of an existing archive */
/*
  Copyright (c) 2014, Randolph Voorhies, Shane Grant
  All rights reserved.

  Redistribution and use in source and binary forms, with or without
  modification, are permitted provided that the following conditions are met:
      * Redistributions of source code must retain the above copyright
        notice, this list of conditions and the following disclaimer.
      * Redistributions in binary form must reproduce the above copyright
        notice, this list of conditions and the following disclaimer in the
        documentation and/or other materials provided with the distribution.
      * Neither the name of cereal nor the
        names of its contributors may be used to endorse or promote products
        derived from this software without specific prior written permission.

  THIS SOFTWARE IS PROVIDED BY THE COPYRIGHT HOLDERS AND CONTRIBUTORS "AS IS" AND
  ANY EXPRESS OR IMPLIED WARRANTIES, INCLUDING, BUT NOT LIMITED TO, THE IMPLIED
  WARRANTIES OF MERCHANTABILITY AND FITNESS FOR A PARTICULAR PURPOSE ARE
  DISCLAIMED. IN NO EVENT SHALL RANDOLPH VOORHIES OR SHANE GRANT BE LIABLE FOR ANY
  DIRECT, INDIRECT, INCIDENTAL, SPECIAL, EXEMPLARY, OR CONSEQUENTIAL DAMAGES
  (INCLUDING, BUT NOT LIMITED TO, PROCUREMENT OF SUBSTITUTE GOODS OR SERVICES;
  LOSS OF USE, DATA, OR PROFITS; OR BUSINESS INTERRUPTION) HOWEVER CAUSED AND
  ON ANY THEORY OF LIABILITY, WHETHER IN CONTRACT, STRICT LIABILITY, OR TORT
  (INCLUDING NEGLIGENCE OR OTHERWISE) ARISING IN ANY WAY OUT OF THE USE OF THIS
  SOFTWARE, EVEN IF ADVISED OF THE POSSIBILITY OF SUCH DAMAGE.
*/
#ifndef CEREAL_ARCHIVES_ADAPTERS_HPP_
#define CEREAL_ARCHIVES_ADAPTERS_HPP_

#include "cereal/details/helpers.hpp"
#include <utility>

namespace cereal
{
  #ifdef CEREAL_FUTURE_EXPERIMENTAL

  // Forward declaration for friend access
  template <class U, class A> U & get_user_data( A & );

  //! Wraps an archive and gives access to user data
  /*! This adapter is useful if you require access to
      either raw pointers or references within your
      serialization functions.

      While cereal does not directly support serialization
      raw pointers or references, it is sometimes the case
      that you may want to supply something such as a raw
      pointer or global reference to some constructor.
      In this situation this adapter would likely be used
      with the construct class to allow for non-default
      constructors.

      @note This feature is experimental and may be altered or removed in a future release. See issue #46.

      @code{.cpp}
      struct MyUserData
      {
        int * myRawPointer;
        std::reference_wrapper<MyOtherType> myReference;
      };

      struct MyClass
      {
        // Note the raw pointer parameter
        MyClass( int xx, int * rawP );

        int x;

        template <class Archive>
        void serialize( Archive & ar )
        { ar( x ); }

        template <class Archive>
        static void load_and_construct( Archive & ar, cereal::construct<MyClass> & construct )
        {
          int xx;
          ar( xx );
          // note the need to use get_user_data to retrieve user data from the archive
          construct( xx, cereal::get_user_data<MyUserData>( ar ).myRawPointer );
        }
      };

      int main()
      {
        {
          MyUserData md;
          md.myRawPointer = &something;
          md.myReference = someInstanceOfType;

          std::ifstream is( "data.xml" );
          cereal::UserDataAdapter<MyUserData, cereal::XMLInputArchive> ar( md, is );

          std::unique_ptr<MyClass> sc;
          ar( sc ); // use as normal
        }

        return 0;
      }
      @endcode

      @relates get_user_data

      @tparam UserData The type to give the archive access to
      @tparam Archive The archive to wrap */
  template <class UserData, class Archive>
  class UserDataAdapter : public Archive
  {
    public:
      //! Construct the archive with some user data struct
      /*! This will forward all arguments (other than the user
          data) to the wrapped archive type.  The UserDataAdapter
          can then be used identically to the wrapped archive type

          @tparam Args The arguments to pass to the constructor of
                       the archive. */
      template <class ... Args>
      UserDataAdapter( UserData & ud, Args && ... args ) :
        Archive( std::forward<Args>( args )... ),
        userdata( ud )
      { }

    private:
      //! Overload the rtti function to enable dynamic_cast
      void rtti() {}
      friend UserData & get_user_data<UserData>( Archive & ar );
      UserData & userdata; //!< The actual user data
  };

  //! Retrieves user data from an archive wrapped by UserDataAdapter
  /*! This will attempt to retrieve the user data associated with
      some archive wrapped by UserDataAdapter.  If this is used on
      an archive that is not wrapped, a run-time exception will occur.

      @note This feature is experimental and may be altered or removed in a future release. See issue #46.

      @note The correct use of this function cannot be enforced at compile
            time.

      @relates UserDataAdapter
      @tparam UserData The data struct contained in the archive
      @tparam Archive The archive, which should be wrapped by UserDataAdapter
      @param ar The archive
      @throws Exception if the archive this is used upon is not wrapped with
                        UserDataAdapter. */
  template <class UserData, class Archive>
  UserData & get_user_data( Archive & ar )
  {
    try
    {
      return dynamic_cast<UserDataAdapter<UserData, Archive> &>( ar ).userdata;
    }
    catch( std::bad_cast const & )
    {
      throw ::cereal::Exception("Attempting to get user data from archive not wrapped in UserDataAdapter");
    }
  }
  #endif // CEREAL_FUTURE_EXPERIMENTAL
} // namespace cereal

#endif // CEREAL_ARCHIVES_ADAPTERS_HPP_
```
