## Supplemental code for "The motif composition of variable-number tandem repeats impacts gene expression": binary.hpp

```
/*! \file binary.hpp
    \brief Binary input and output archives */
/*
  Copyright (c) 2014, Randolph Voorhies, Shane Grant
  All rights reserved.

  Redistribution and use in source and binary forms, with or without
  modification, are permitted provided that the following conditions are met:
      * Redistributions of source code must retain the above copyright
        notice, this list of conditions and the following disclaimer.
      * Redistributions in binary form must reproduce the above copyright
        notice, this list of conditions and the following disclaimer in the
        documentation and/or other materials provided with the distribution.
      * Neither the name of cereal nor the
        names of its contributors may be used to endorse or promote products
        derived from this software without specific prior written permission.

  THIS SOFTWARE IS PROVIDED BY THE COPYRIGHT HOLDERS AND CONTRIBUTORS "AS IS" AND
  ANY EXPRESS OR IMPLIED WARRANTIES, INCLUDING, BUT NOT LIMITED TO, THE IMPLIED
  WARRANTIES OF MERCHANTABILITY AND FITNESS FOR A PARTICULAR PURPOSE ARE
  DISCLAIMED. IN NO EVENT SHALL RANDOLPH VOORHIES OR SHANE GRANT BE LIABLE FOR ANY
  DIRECT, INDIRECT, INCIDENTAL, SPECIAL, EXEMPLARY, OR CONSEQUENTIAL DAMAGES
  (INCLUDING, BUT NOT LIMITED TO, PROCUREMENT OF SUBSTITUTE GOODS OR SERVICES;
  LOSS OF USE, DATA, OR PROFITS; OR BUSINESS INTERRUPTION) HOWEVER CAUSED AND
  ON ANY THEORY OF LIABILITY, WHETHER IN CONTRACT, STRICT LIABILITY, OR TORT
  (INCLUDING NEGLIGENCE OR OTHERWISE) ARISING IN ANY WAY OUT OF THE USE OF THIS
  SOFTWARE, EVEN IF ADVISED OF THE POSSIBILITY OF SUCH DAMAGE.
*/
#ifndef CEREAL_ARCHIVES_BINARY_HPP_
#define CEREAL_ARCHIVES_BINARY_HPP_

#include "cereal/cereal.hpp"
#include <sstream>

namespace cereal
{
  // ######################################################################
  //! An output archive designed to save data in a compact binary representation
  /*! This archive outputs data to a stream in an extremely compact binary
      representation with as little extra metadata as possible.

      This archive does nothing to ensure that the endianness of the saved
      and loaded data is the same.  If you need to have portability over
      architectures with different endianness, use PortableBinaryOutputArchive.

      When using a binary archive and a file stream, you must use the
      std::ios::binary format flag to avoid having your data altered
      inadvertently.

      \ingroup Archives */
  class BinaryOutputArchive : public OutputArchive<BinaryOutputArchive, AllowEmptyClassElision>
  {
    public:
      //! Construct, outputting to the provided stream
      /*! @param stream The stream to output to.  Can be a stringstream, a file stream, or
                        even cout! */
      BinaryOutputArchive(std::ostream & stream) :
        OutputArchive<BinaryOutputArchive, AllowEmptyClassElision>(this),
        itsStream(stream)
      { }

      ~BinaryOutputArchive() CEREAL_NOEXCEPT = default;

      //! Writes size bytes of data to the output stream
      void saveBinary( const void * data, std::streamsize size )
      {
        auto const writtenSize = itsStream.rdbuf()->sputn( reinterpret_cast<const char*>( data ), size );

        if(writtenSize != size)
          throw Exception("Failed to write " + std::to_string(size) + " bytes to output stream! Wrote " + std::to_string(writtenSize));
      }

    private:
      std::ostream & itsStream;
  };

  // ######################################################################
  //! An input archive designed to load data saved using BinaryOutputArchive
  /*  This archive does nothing to ensure that the endianness of the saved
      and loaded data is the same.  If you need to have portability over
      architectures with different endianness, use PortableBinaryOutputArchive.

      When using a binary archive and a file stream, you must use the
      std::ios::binary format flag to avoid having your data altered
      inadvertently.

      \ingroup Archives */
  class BinaryInputArchive : public InputArchive<BinaryInputArchive, AllowEmptyClassElision>
  {
    public:
      //! Construct, loading from the provided stream
      BinaryInputArchive(std::istream & stream) :
        InputArchive<BinaryInputArchive, AllowEmptyClassElision>(this),
        itsStream(stream)
      { }

      ~BinaryInputArchive() CEREAL_NOEXCEPT = default;

      //! Reads size bytes of data from the input stream
      void loadBinary( void * const data, std::streamsize size )
      {
        auto const readSize = itsStream.rdbuf()->sgetn( reinterpret_cast<char*>( data ), size );

        if(readSize != size)
          throw Exception("Failed to read " + std::to_string(size) + " bytes from input stream! Read " + std::to_string(readSize));
      }

    private:
      std::istream & itsStream;
  };

  // ######################################################################
  // Common BinaryArchive serialization functions

  //! Saving for POD types to binary
  template<class T> inline
  typename std::enable_if<std::is_arithmetic<T>::value, void>::type
  CEREAL_SAVE_FUNCTION_NAME(BinaryOutputArchive & ar, T const & t)
  {
    ar.saveBinary(std::addressof(t), sizeof(t));
  }

  //! Loading for POD types from binary
  template<class T> inline
  typename std::enable_if<std::is_arithmetic<T>::value, void>::type
  CEREAL_LOAD_FUNCTION_NAME(BinaryInputArchive & ar, T & t)
  {
    ar.loadBinary(std::addressof(t), sizeof(t));
  }

  //! Serializing NVP types to binary
  template <class Archive, class T> inline
  CEREAL_ARCHIVE_RESTRICT(BinaryInputArchive, BinaryOutputArchive)
  CEREAL_SERIALIZE_FUNCTION_NAME( Archive & ar, NameValuePair<T> & t )
  {
    ar( t.value );
  }

  //! Serializing SizeTags to binary
  template <class Archive, class T> inline
  CEREAL_ARCHIVE_RESTRICT(BinaryInputArchive, BinaryOutputArchive)
  CEREAL_SERIALIZE_FUNCTION_NAME( Archive & ar, SizeTag<T> & t )
  {
    ar( t.size );
  }

  //! Saving binary data
  template <class T> inline
  void CEREAL_SAVE_FUNCTION_NAME(BinaryOutputArchive & ar, BinaryData<T> const & bd)
  {
    ar.saveBinary( bd.data, static_cast<std::streamsize>( bd.size ) );
  }

  //! Loading binary data
  template <class T> inline
  void CEREAL_LOAD_FUNCTION_NAME(BinaryInputArchive & ar, BinaryData<T> & bd)
  {
    ar.loadBinary(bd.data, static_cast<std::streamsize>( bd.size ) );
  }
} // namespace cereal

// register archives for polymorphic support
CEREAL_REGISTER_ARCHIVE(cereal::BinaryOutputArchive)
CEREAL_REGISTER_ARCHIVE(cereal::BinaryInputArchive)

// tie input and output archives together
CEREAL_SETUP_ARCHIVE_TRAITS(cereal::BinaryInputArchive, cereal::BinaryOutputArchive)

#endif // CEREAL_ARCHIVES_BINARY_HPP_
```
