## Supplemental code for "The motif composition of variable-number tandem repeats impacts gene expression": json.hpp

```
/*! \file json.hpp
    \brief JSON input and output archives */
/*
  Copyright (c) 2014, Randolph Voorhies, Shane Grant
  All rights reserved.

  Redistribution and use in source and binary forms, with or without
  modification, are permitted provided that the following conditions are met:
      * Redistributions of source code must retain the above copyright
        notice, this list of conditions and the following disclaimer.
      * Redistributions in binary form must reproduce the above copyright
        notice, this list of conditions and the following disclaimer in the
        documentation and/or other materials provided with the distribution.
      * Neither the name of cereal nor the
        names of its contributors may be used to endorse or promote products
        derived from this software without specific prior written permission.

  THIS SOFTWARE IS PROVIDED BY THE COPYRIGHT HOLDERS AND CONTRIBUTORS "AS IS" AND
  ANY EXPRESS OR IMPLIED WARRANTIES, INCLUDING, BUT NOT LIMITED TO, THE IMPLIED
  WARRANTIES OF MERCHANTABILITY AND FITNESS FOR A PARTICULAR PURPOSE ARE
  DISCLAIMED. IN NO EVENT SHALL RANDOLPH VOORHIES OR SHANE GRANT BE LIABLE FOR ANY
  DIRECT, INDIRECT, INCIDENTAL, SPECIAL, EXEMPLARY, OR CONSEQUENTIAL DAMAGES
  (INCLUDING, BUT NOT LIMITED TO, PROCUREMENT OF SUBSTITUTE GOODS OR SERVICES;
  LOSS OF USE, DATA, OR PROFITS; OR BUSINESS INTERRUPTION) HOWEVER CAUSED AND
  ON ANY THEORY OF LIABILITY, WHETHER IN CONTRACT, STRICT LIABILITY, OR TORT
  (INCLUDING NEGLIGENCE OR OTHERWISE) ARISING IN ANY WAY OUT OF THE USE OF THIS
  SOFTWARE, EVEN IF ADVISED OF THE POSSIBILITY OF SUCH DAMAGE.
*/
#ifndef CEREAL_ARCHIVES_JSON_HPP_
#define CEREAL_ARCHIVES_JSON_HPP_

#include "cereal/cereal.hpp"
#include "cereal/details/util.hpp"

namespace cereal
{
  //! An exception thrown when rapidjson fails an internal assertion
  /*! @ingroup Utility */
  struct RapidJSONException : Exception
  { RapidJSONException( const char * what_ ) : Exception( what_ ) {} };
}

// Inform rapidjson that assert will throw
#ifndef CEREAL_RAPIDJSON_ASSERT_THROWS
#define CEREAL_RAPIDJSON_ASSERT_THROWS
#endif // CEREAL_RAPIDJSON_ASSERT_THROWS

// Override rapidjson assertions to throw exceptions by default
#ifndef CEREAL_RAPIDJSON_ASSERT
#define CEREAL_RAPIDJSON_ASSERT(x) if(!(x)){ \
  throw ::cereal::RapidJSONException("rapidjson internal assertion failure: " #x); }
#endif // RAPIDJSON_ASSERT

// Enable support for parsing of nan, inf, -inf
#ifndef CEREAL_RAPIDJSON_WRITE_DEFAULT_FLAGS
#define CEREAL_RAPIDJSON_WRITE_DEFAULT_FLAGS kWriteNanAndInfFlag
#endif

// Enable support for parsing of nan, inf, -inf
#ifndef CEREAL_RAPIDJSON_PARSE_DEFAULT_FLAGS
#define CEREAL_RAPIDJSON_PARSE_DEFAULT_FLAGS kParseFullPrecisionFlag | kParseNanAndInfFlag
#endif

#include "cereal/external/rapidjson/prettywriter.h"
#include "cereal/external/rapidjson/ostreamwrapper.h"
#include "cereal/external/rapidjson/istreamwrapper.h"
#include "cereal/external/rapidjson/document.h"
#include "cereal/external/base64.hpp"

#include <limits>
#include <sstream>
#include <stack>
#include <vector>
#include <string>

namespace cereal
{
  // ######################################################################
  //! An output archive designed to save data to JSON
  /*! This archive uses RapidJSON to build serialize data to JSON.

      JSON archives provides a human readable output but at decreased
      performance (both in time and space) compared to binary archives.

      JSON archives are only guaranteed to finish flushing their contents
      upon destruction and should thus be used in an RAII fashion.

      JSON benefits greatly from name-value pairs, which if present, will
      name the nodes in the output.  If these are not present, each level
      of the output will be given an automatically generated delimited name.

      The precision of the output archive controls the number of decimals output
      for floating point numbers and should be sufficiently large (i.e. at least 20)
      if there is a desire to have binary equality between the numbers output and
      those read in.  In general you should expect a loss of precision when going
      from floating point to text and back.

      JSON archives do not output the size information for any dynamically sized structure
      and instead infer it from the number of children for a node.  This means that data
      can be hand edited for dynamic sized structures and will still be readable.  This
      is accomplished through the cereal::SizeTag object, which will cause the archive
      to output the data as a JSON array (e.g. marked by [] instead of {}), which indicates
      that the container is variable sized and may be edited.

      \ingroup Archives */
  class JSONOutputArchive : public OutputArchive<JSONOutputArchive>, public traits::TextArchive
  {
    enum class NodeType { StartObject, InObject, StartArray, InArray };

    using WriteStream = CEREAL_RAPIDJSON_NAMESPACE::OStreamWrapper;
    using JSONWriter = CEREAL_RAPIDJSON_NAMESPACE::PrettyWriter<WriteStream>;

    public:
      /*! @name Common Functionality
          Common use cases for directly interacting with an JSONOutputArchive */
      //! @{

      //! A class containing various advanced options for the JSON archive
      class Options
      {
        public:
          //! Default options
          static Options Default(){ return Options(); }

          //! Default options with no indentation
          static Options NoIndent(){ return Options( JSONWriter::kDefaultMaxDecimalPlaces, IndentChar::space, 0 ); }

          //! The character to use for indenting
          enum class IndentChar : char
          {
            space = ' ',
            tab = '\t',
            newline = '\n',
            carriage_return = '\r'
          };

          //! Specify specific options for the JSONOutputArchive
          /*! @param precision The precision used for floating point numbers
              @param indentChar The type of character to indent with
              @param indentLength The number of indentChar to use for indentation
                             (0 corresponds to no indentation) */
          explicit Options( int precision = JSONWriter::kDefaultMaxDecimalPlaces,
                            IndentChar indentChar = IndentChar::space,
                            unsigned int indentLength = 4 ) :
            itsPrecision( precision ),
            itsIndentChar( static_cast<char>(indentChar) ),
            itsIndentLength( indentLength ) { }

        private:
          friend class JSONOutputArchive;
          int itsPrecision;
          char itsIndentChar;
          unsigned int itsIndentLength;
      };

      //! Construct, outputting to the provided stream
      /*! @param stream The stream to output to.
          @param options The JSON specific options to use.  See the Options struct
                         for the values of default parameters */
      JSONOutputArchive(std::ostream & stream, Options const & options = Options::Default() ) :
        OutputArchive<JSONOutputArchive>(this),
        itsWriteStream(stream),
        itsWriter(itsWriteStream),
        itsNextName(nullptr)
      {
        itsWriter.SetMaxDecimalPlaces( options.itsPrecision );
        itsWriter.SetIndent( options.itsIndentChar, options.itsIndentLength );
        itsNameCounter.push(0);
        itsNodeStack.push(NodeType::StartObject);
      }

      //! Destructor, flushes the JSON
      ~JSONOutputArchive() CEREAL_NOEXCEPT
      {
        if (itsNodeStack.top() == NodeType::InObject)
          itsWriter.EndObject();
        else if (itsNodeStack.top() == NodeType::InArray)
          itsWriter.EndArray();
      }

      //! Saves some binary data, encoded as a base64 string, with an optional name
      /*! This will create a new node, optionally named, and insert a value that consists of
          the data encoded as a base64 string */
      void saveBinaryValue( const void * data, size_t size, const char * name = nullptr )
      {
        setNextName( name );
        writeName();

        auto base64string = base64::encode( reinterpret_cast<const unsigned char *>( data ), size );
        saveValue( base64string );
      };

      //! @}
      /*! @name Internal Functionality
          Functionality designed for use by those requiring control over the inner mechanisms of
          the JSONOutputArchive */
      //! @{

      //! Starts a new node in the JSON output
      /*! The node can optionally be given a name by calling setNextName prior
          to creating the node

          Nodes only need to be started for types that are themselves objects or arrays */
      void startNode()
      {
        writeName();
        itsNodeStack.push(NodeType::StartObject);
        itsNameCounter.push(0);
      }

      //! Designates the most recently added node as finished
      void finishNode()
      {
        // if we ended up serializing an empty object or array, writeName
        // will never have been called - so start and then immediately end
        // the object/array.
        //
        // We'll also end any object/arrays we happen to be in
        switch(itsNodeStack.top())
        {
          case NodeType::StartArray:
            itsWriter.StartArray();
            // fall through
          case NodeType::InArray:
            itsWriter.EndArray();
            break;
          case NodeType::StartObject:
            itsWriter.StartObject();
            // fall through
          case NodeType::InObject:
            itsWriter.EndObject();
            break;
        }

        itsNodeStack.pop();
        itsNameCounter.pop();
      }

      //! Sets the name for the next node created with startNode
      void setNextName( const char * name )
      {
        itsNextName = name;
      }

      //! Saves a bool to the current node
      void saveValue(bool b)                { itsWriter.Bool(b);                                                         }
      //! Saves an int to the current node
      void saveValue(int i)                 { itsWriter.Int(i);                                                          }
      //! Saves a uint to the current node
      void saveValue(unsigned u)            { itsWriter.Uint(u);                                                         }
      //! Saves an int64 to the current node
      void saveValue(int64_t i64)           { itsWriter.Int64(i64);                                                      }
      //! Saves a uint64 to the current node
      void saveValue(uint64_t u64)          { itsWriter.Uint64(u64);                                                     }
      //! Saves a double to the current node
      void saveValue(double d)              { itsWriter.Double(d);                                                       }
      //! Saves a string to the current node
      void saveValue(std::string const & s) { itsWriter.String(s.c_str(), static_cast<CEREAL_RAPIDJSON_NAMESPACE::SizeType>( s.size() )); }
      //! Saves a const char * to the current node
      void saveValue(char const * s)        { itsWriter.String(s);                                                       }
      //! Saves a nullptr to the current node
      void saveValue(std::nullptr_t)        { itsWriter.Null();                                                          }

    private:
      // Some compilers/OS have difficulty disambiguating the above for various flavors of longs, so we provide
      // special overloads to handle these cases.

      //! 32 bit signed long saving to current node
      template <class T, traits::EnableIf<sizeof(T) == sizeof(std::int32_t),
                                          std::is_signed<T>::value> = traits::sfinae> inline
      void saveLong(T l){ saveValue( static_cast<std::int32_t>( l ) ); }

      //! non 32 bit signed long saving to current node
      template <class T, traits::EnableIf<sizeof(T) != sizeof(std::int32_t),
                                          std::is_signed<T>::value> = traits::sfinae> inline
      void saveLong(T l){ saveValue( static_cast<std::int64_t>( l ) ); }

      //! 32 bit unsigned long saving to current node
      template <class T, traits::EnableIf<sizeof(T) == sizeof(std::int32_t),
                                          std::is_unsigned<T>::value> = traits::sfinae> inline
      void saveLong(T lu){ saveValue( static_cast<std::uint32_t>( lu ) ); }

      //! non 32 bit unsigned long saving to current node
      template <class T, traits::EnableIf<sizeof(T) != sizeof(std::int32_t),
                                          std::is_unsigned<T>::value> = traits::sfinae> inline
      void saveLong(T lu){ saveValue( static_cast<std::uint64_t>( lu ) ); }

    public:
#if defined(_MSC_VER) && _MSC_VER < 1916
      //! MSVC only long overload to current node
      void saveValue( unsigned long lu ){ saveLong( lu ); };
#else // _MSC_VER
      //! Serialize a long if it would not be caught otherwise
      template <class T, traits::EnableIf<std::is_same<T, long>::value,
                                          !std::is_same<T, int>::value,
                                          !std::is_same<T, std::int64_t>::value> = traits::sfinae> inline
      void saveValue( T t ){ saveLong( t ); }

      //! Serialize an unsigned long if it would not be caught otherwise
      template <class T, traits::EnableIf<std::is_same<T, unsigned long>::value,
                                          !std::is_same<T, unsigned>::value,
                                          !std::is_same<T, std::uint64_t>::value> = traits::sfinae> inline
      void saveValue( T t ){ saveLong( t ); }
#endif // _MSC_VER

      //! Save exotic arithmetic as strings to current node
      /*! Handles long long (if distinct from other types), unsigned long (if distinct), and long double */
      template <class T, traits::EnableIf<std::is_arithmetic<T>::value,
                                          !std::is_same<T, long>::value,
                                          !std::is_same<T, unsigned long>::value,
                                          !std::is_same<T, std::int64_t>::value,
                                          !std::is_same<T, std::uint64_t>::value,
                                          (sizeof(T) >= sizeof(long double) || sizeof(T) >= sizeof(long long))> = traits::sfinae> inline
      void saveValue(T const & t)
      {
        std::stringstream ss; ss.precision( std::numeric_limits<long double>::max_digits10 );
        ss << t;
        saveValue( ss.str() );
      }

      //! Write the name of the upcoming node and prepare object/array state
      /*! Since writeName is called for every value that is output, regardless of
          whether it has a name or not, it is the place where we will do a deferred
          check of our node state and decide whether we are in an array or an object.

          The general workflow of saving to the JSON archive is:

            1. (optional) Set the name for the next node to be created, usually done by an NVP
            2. Start the node
            3. (if there is data to save) Write the name of the node (this function)
            4. (if there is data to save) Save the data (with saveValue)
            5. Finish the node
          */
      void writeName()
      {
        NodeType const & nodeType = itsNodeStack.top();

        // Start up either an object or an array, depending on state
        if(nodeType == NodeType::StartArray)
        {
          itsWriter.StartArray();
          itsNodeStack.top() = NodeType::InArray;
        }
        else if(nodeType == NodeType::StartObject)
        {
          itsNodeStack.top() = NodeType::InObject;
          itsWriter.StartObject();
        }

        // Array types do not output names
        if(nodeType == NodeType::InArray) return;

        if(itsNextName == nullptr)
        {
          std::string name = "value" + std::to_string( itsNameCounter.top()++ ) + "\0";
          saveValue(name);
        }
        else
        {
          saveValue(itsNextName);
          itsNextName = nullptr;
        }
      }

      //! Designates that the current node should be output as an array, not an object
      void makeArray()
      {
        itsNodeStack.top() = NodeType::StartArray;
      }

      //! @}

    private:
      WriteStream itsWriteStream;          //!< Rapidjson write stream
      JSONWriter itsWriter;                //!< Rapidjson writer
      char const * itsNextName;            //!< The next name
      std::stack<uint32_t> itsNameCounter; //!< Counter for creating unique names for unnamed nodes
      std::stack<NodeType> itsNodeStack;
  }; // JSONOutputArchive

  // ######################################################################
  //! An input archive designed to load data from JSON
  /*! This archive uses RapidJSON to read in a JSON archive.

      As with the output JSON archive, the preferred way to use this archive is in
      an RAII fashion, ensuring its destruction after all data has been read.

      Input JSON should have been produced by the JSONOutputArchive.  Data can
      only be added to dynamically sized containers (marked by JSON arrays) -
      the input archive will determine their size by looking at the number of child nodes.
      Only JSON originating from a JSONOutputArchive is officially supported, but data
      from other sources may work if properly formatted.

      The JSONInputArchive does not require that nodes are loaded in the same
      order they were saved by JSONOutputArchive.  Using name value pairs (NVPs),
      it is possible to load in an out of order fashion or otherwise skip/select
      specific nodes to load.

      The default behavior of the input archive is to read sequentially starting
      with the first node and exploring its children.  When a given NVP does
      not match the read in name for a node, the archive will search for that
      node at the current level and load it if it exists.  After loading an out of
      order node, the archive will then proceed back to loading sequentially from
      its new position.

      Consider this simple example where loading of some data is skipped:

      @code{cpp}
      // imagine the input file has someData(1-9) saved in order at the top level node
      ar( someData1, someData2, someData3 );        // XML loads in the order it sees in the file
      ar( cereal::make_nvp( "hello", someData6 ) ); // NVP given does not
                                                    // match expected NVP name, so we search
                                                    // for the given NVP and load that value
      ar( someData7, someData8, someData9 );        // with no NVP given, loading resumes at its
                                                    // current location, proceeding sequentially
      @endcode

      \ingroup Archives */
  class JSONInputArchive : public InputArchive<JSONInputArchive>, public traits::TextArchive
  {
    private:
      using ReadStream = CEREAL_RAPIDJSON_NAMESPACE::IStreamWrapper;
      typedef CEREAL_RAPIDJSON_NAMESPACE::GenericValue<CEREAL_RAPIDJSON_NAMESPACE::UTF8<>> JSONValue;
      typedef JSONValue::ConstMemberIterator MemberIterator;
      typedef JSONValue::ConstValueIterator ValueIterator;
      typedef CEREAL_RAPIDJSON_NAMESPACE::Document::GenericValue GenericValue;

    public:
      /*! @name Common Functionality
          Common use cases for directly interacting with an JSONInputArchive */
      //! @{

      //! Construct, reading from the provided stream
      /*! @param stream The stream to read from */
      JSONInputArchive(std::istream & stream) :
        InputArchive<JSONInputArchive>(this),
        itsNextName( nullptr ),
        itsReadStream(stream)
      {
        itsDocument.ParseStream<>(itsReadStream);
        if (itsDocument.IsArray())
          itsIteratorStack.emplace_back(itsDocument.Begin(), itsDocument.End());
        else
          itsIteratorStack.emplace_back(itsDocument.MemberBegin(), itsDocument.MemberEnd());
      }

      ~JSONInputArchive() CEREAL_NOEXCEPT = default;

      //! Loads some binary data, encoded as a base64 string
      /*! This will automatically start and finish a node to load the data, and can be called directly by
          users.

          Note that this follows the same ordering rules specified in the class description in regards
          to loading in/out of order */
      void loadBinaryValue( void * data, size_t size, const char * name = nullptr )
      {
        itsNextName = name;

        std::string encoded;
        loadValue( encoded );
        auto decoded = base64::decode( encoded );

        if( size != decoded.size() )
          throw Exception("Decoded binary data size does not match specified size");

        std::memcpy( data, decoded.data(), decoded.size() );
        itsNextName = nullptr;
      };

    private:
      //! @}
      /*! @name Internal Functionality
          Functionality designed for use by those requiring control over the inner mechanisms of
          the JSONInputArchive */
      //! @{

      //! An internal iterator that handles both array and object types
      /*! This class is a variant and holds both types of iterators that
          rapidJSON supports - one for arrays and one for objects. */
      class Iterator
      {
        public:
          Iterator() : itsIndex( 0 ), itsType(Null_) {}

          Iterator(MemberIterator begin, MemberIterator end) :
            itsMemberItBegin(begin), itsMemberItEnd(end), itsIndex(0), itsSize(std::distance(begin, end)), itsType(Member)
          {
            if( itsSize == 0 )
              itsType = Null_;
          }

          Iterator(ValueIterator begin, ValueIterator end) :
            itsValueItBegin(begin), itsIndex(0), itsSize(std::distance(begin, end)), itsType(Value)
          {
            if( itsSize == 0 )
              itsType = Null_;
          }

          //! Advance to the next node
          Iterator & operator++()
          {
            ++itsIndex;
            return *this;
          }

          //! Get the value of the current node
          GenericValue const & value()
          {
            if( itsIndex >= itsSize )
              throw cereal::Exception("No more objects in input");

            switch(itsType)
            {
              case Value : return itsValueItBegin[itsIndex];
              case Member: return itsMemberItBegin[itsIndex].value;
              default: throw cereal::Exception("JSONInputArchive internal error: null or empty iterator to object or array!");
            }
          }

          //! Get the name of the current node, or nullptr if it has no name
          const char * name() const
          {
            if( itsType == Member && (itsMemberItBegin + itsIndex) != itsMemberItEnd )
              return itsMemberItBegin[itsIndex].name.GetString();
            else
              return nullptr;
          }

          //! Adjust our position such that we are at the node with the given name
          /*! @throws Exception if no such named node exists */
          inline void search( const char * searchName )
          {
            const auto len = std::strlen( searchName );
            size_t index = 0;
            for( auto it = itsMemberItBegin; it != itsMemberItEnd; ++it, ++index )
            {
              const auto currentName = it->name.GetString();
              if( ( std::strncmp( searchName, currentName, len ) == 0 ) &&
                  ( std::strlen( currentName ) == len ) )
              {
                itsIndex = index;
                return;
              }
            }

            throw Exception("JSON Parsing failed - provided NVP (" + std::string(searchName) + ") not found");
          }

        private:
          MemberIterator itsMemberItBegin, itsMemberItEnd; //!< The member iterator (object)
          ValueIterator itsValueItBegin;                   //!< The value iterator (array)
          size_t itsIndex, itsSize;                        //!< The current index of this iterator
          enum Type {Value, Member, Null_} itsType;        //!< Whether this holds values (array) or members (objects) or nothing
      };

      //! Searches for the expectedName node if it doesn't match the actualName
      /*! This needs to be called before every load or node start occurs.  This function will
          check to see if an NVP has been provided (with setNextName) and if so, see if that name matches the actual
          next name given.  If the names do not match, it will search in the current level of the JSON for that name.
          If the name is not found, an exception will be thrown.

          Resets the NVP name after called.

          @throws Exception if an expectedName is given and not found */
      inline void search()
      {
        // The name an NVP provided with setNextName()
        if( itsNextName )
        {
          // The actual name of the current node
          auto const actualName = itsIteratorStack.back().name();

          // Do a search if we don't see a name coming up, or if the names don't match
          if( !actualName || std::strcmp( itsNextName, actualName ) != 0 )
            itsIteratorStack.back().search( itsNextName );
        }

        itsNextName = nullptr;
      }

    public:
      //! Starts a new node, going into its proper iterator
      /*! This places an iterator for the next node to be parsed onto the iterator stack.  If the next
          node is an array, this will be a value iterator, otherwise it will be a member iterator.

          By default our strategy is to start with the document root node and then recursively iterate through
          all children in the order they show up in the document.
          We don't need to know NVPs to do this; we'll just blindly load in the order things appear in.

          If we were given an NVP, we will search for it if it does not match our the name of the next node
          that would normally be loaded.  This functionality is provided by search(). */
      void startNode()
      {
        search();

        if(itsIteratorStack.back().value().IsArray())
          itsIteratorStack.emplace_back(itsIteratorStack.back().value().Begin(), itsIteratorStack.back().value().End());
        else
          itsIteratorStack.emplace_back(itsIteratorStack.back().value().MemberBegin(), itsIteratorStack.back().value().MemberEnd());
      }

      //! Finishes the most recently started node
      void finishNode()
      {
        itsIteratorStack.pop_back();
        ++itsIteratorStack.back();
      }

      //! Retrieves the current node name
      /*! @return nullptr if no name exists */
      const char * getNodeName() const
      {
        return itsIteratorStack.back().name();
      }

      //! Sets the name for the next node created with startNode
      void setNextName( const char * name )
      {
        itsNextName = name;
      }

      //! Loads a value from the current node - small signed overload
      template <class T, traits::EnableIf<std::is_signed<T>::value,
                                          sizeof(T) < sizeof(int64_t)> = traits::sfinae> inline
      void loadValue(T & val)
      {
        search();

        val = static_cast<T>( itsIteratorStack.back().value().GetInt() );
        ++itsIteratorStack.back();
      }

      //! Loads a value from the current node - small unsigned overload
      template <class T, traits::EnableIf<std::is_unsigned<T>::value,
                                          sizeof(T) < sizeof(uint64_t),
                                          !std::is_same<bool, T>::value> = traits::sfinae> inline
      void loadValue(T & val)
      {
        search();

        val = static_cast<T>( itsIteratorStack.back().value().GetUint() );
        ++itsIteratorStack.back();
      }

      //! Loads a value from the current node - bool overload
      void loadValue(bool & val)        { search(); val = itsIteratorStack.back().value().GetBool(); ++itsIteratorStack.back(); }
      //! Loads a value from the current node - int64 overload
      void loadValue(int64_t & val)     { search(); val = itsIteratorStack.back().value().GetInt64(); ++itsIteratorStack.back(); }
      //! Loads a value from the current node - uint64 overload
      void loadValue(uint64_t & val)    { search(); val = itsIteratorStack.back().value().GetUint64(); ++itsIteratorStack.back(); }
      //! Loads a value from the current node - float overload
      void loadValue(float & val)       { search(); val = static_cast<float>(itsIteratorStack.back().value().GetDouble()); ++itsIteratorStack.back(); }
      //! Loads a value from the current node - double overload
      void loadValue(double & val)      { search(); val = itsIteratorStack.back().value().GetDouble(); ++itsIteratorStack.back(); }
      //! Loads a value from the current node - string overload
      void loadValue(std::string & val) { search(); val = itsIteratorStack.back().value().GetString(); ++itsIteratorStack.back(); }
      //! Loads a nullptr from the current node
      void loadValue(std::nullptr_t&)   { search(); CEREAL_RAPIDJSON_ASSERT(itsIteratorStack.back().value().IsNull()); ++itsIteratorStack.back(); }

      // Special cases to handle various flavors of long, which tend to conflict with
      // the int32_t or int64_t on various compiler/OS combinations.  MSVC doesn't need any of this.
      #ifndef _MSC_VER
    private:
      //! 32 bit signed long loading from current node
      template <class T> inline
      typename std::enable_if<sizeof(T) == sizeof(std::int32_t) && std::is_signed<T>::value, void>::type
      loadLong(T & l){ loadValue( reinterpret_cast<std::int32_t&>( l ) ); }

      //! non 32 bit signed long loading from current node
      template <class T> inline
      typename std::enable_if<sizeof(T) == sizeof(std::int64_t) && std::is_signed<T>::value, void>::type
      loadLong(T & l){ loadValue( reinterpret_cast<std::int64_t&>( l ) ); }

      //! 32 bit unsigned long loading from current node
      template <class T> inline
      typename std::enable_if<sizeof(T) == sizeof(std::uint32_t) && !std::is_signed<T>::value, void>::type
      loadLong(T & lu){ loadValue( reinterpret_cast<std::uint32_t&>( lu ) ); }

      //! non 32 bit unsigned long loading from current node
      template <class T> inline
      typename std::enable_if<sizeof(T) == sizeof(std::uint64_t) && !std::is_signed<T>::value, void>::type
      loadLong(T & lu){ loadValue( reinterpret_cast<std::uint64_t&>( lu ) ); }

    public:
      //! Serialize a long if it would not be caught otherwise
      template <class T> inline
      typename std::enable_if<std::is_same<T, long>::value &&
                              sizeof(T) >= sizeof(std::int64_t) &&
                              !std::is_same<T, std::int64_t>::value, void>::type
      loadValue( T & t ){ loadLong(t); }

      //! Serialize an unsigned long if it would not be caught otherwise
      template <class T> inline
      typename std::enable_if<std::is_same<T, unsigned long>::value &&
                              sizeof(T) >= sizeof(std::uint64_t) &&
                              !std::is_same<T, std::uint64_t>::value, void>::type
      loadValue( T & t ){ loadLong(t); }
      #endif // _MSC_VER

    private:
      //! Convert a string to a long long
      void stringToNumber( std::string const & str, long long & val ) { val = std::stoll( str ); }
      //! Convert a string to an unsigned long long
      void stringToNumber( std::string const & str, unsigned long long & val ) { val = std::stoull( str ); }
      //! Convert a string to a long double
      void stringToNumber( std::string const & str, long double & val ) { val = std::stold( str ); }

    public:
      //! Loads a value from the current node - long double and long long overloads
      template <class T, traits::EnableIf<std::is_arithmetic<T>::value,
                                          !std::is_same<T, long>::value,
                                          !std::is_same<T, unsigned long>::value,
                                          !std::is_same<T, std::int64_t>::value,
                                          !std::is_same<T, std::uint64_t>::value,
                                          (sizeof(T) >= sizeof(long double) || sizeof(T) >= sizeof(long long))> = traits::sfinae>
      inline void loadValue(T & val)
      {
        std::string encoded;
        loadValue( encoded );
        stringToNumber( encoded, val );
      }

      //! Loads the size for a SizeTag
      void loadSize(size_type & size)
      {
        if (itsIteratorStack.size() == 1)
          size = itsDocument.Size();
        else
          size = (itsIteratorStack.rbegin() + 1)->value().Size();
      }

      //! @}

    private:
      const char * itsNextName;               //!< Next name set by NVP
      ReadStream itsReadStream;               //!< Rapidjson write stream
      std::vector<Iterator> itsIteratorStack; //!< 'Stack' of rapidJSON iterators
      CEREAL_RAPIDJSON_NAMESPACE::Document itsDocument; //!< Rapidjson document
  };

  // ######################################################################
  // JSONArchive prologue and epilogue functions
  // ######################################################################

  // ######################################################################
  //! Prologue for NVPs for JSON archives
  /*! NVPs do not start or finish nodes - they just set up the names */
  template <class T> inline
  void prologue( JSONOutputArchive &, NameValuePair<T> const & )
  { }

  //! Prologue for NVPs for JSON archives
  template <class T> inline
  void prologue( JSONInputArchive &, NameValuePair<T> const & )
  { }

  // ######################################################################
  //! Epilogue for NVPs for JSON archives
  /*! NVPs do not start or finish nodes - they just set up the names */
  template <class T> inline
  void epilogue( JSONOutputArchive &, NameValuePair<T> const & )
  { }

  //! Epilogue for NVPs for JSON archives
  /*! NVPs do not start or finish nodes - they just set up the names */
  template <class T> inline
  void epilogue( JSONInputArchive &, NameValuePair<T> const & )
  { }

  // ######################################################################
  //! Prologue for deferred data for JSON archives
  /*! Do nothing for the defer wrapper */
  template <class T> inline
  void prologue( JSONOutputArchive &, DeferredData<T> const & )
  { }

  //! Prologue for deferred data for JSON archives
  template <class T> inline
  void prologue( JSONInputArchive &, DeferredData<T> const & )
  { }

  // ######################################################################
  //! Epilogue for deferred for JSON archives
  /*! NVPs do not start or finish nodes - they just set up the names */
  template <class T> inline
  void epilogue( JSONOutputArchive &, DeferredData<T> const & )
  { }

  //! Epilogue for deferred for JSON archives
  /*! Do nothing for the defer wrapper */
  template <class T> inline
  void epilogue( JSONInputArchive &, DeferredData<T> const & )
  { }

  // ######################################################################
  //! Prologue for SizeTags for JSON archives
  /*! SizeTags are strictly ignored for JSON, they just indicate
      that the current node should be made into an array */
  template <class T> inline
  void prologue( JSONOutputArchive & ar, SizeTag<T> const & )
  {
    ar.makeArray();
  }

  //! Prologue for SizeTags for JSON archives
  template <class T> inline
  void prologue( JSONInputArchive &, SizeTag<T> const & )
  { }

  // ######################################################################
  //! Epilogue for SizeTags for JSON archives
  /*! SizeTags are strictly ignored for JSON */
  template <class T> inline
  void epilogue( JSONOutputArchive &, SizeTag<T> const & )
  { }

  //! Epilogue for SizeTags for JSON archives
  template <class T> inline
  void epilogue( JSONInputArchive &, SizeTag<T> const & )
  { }

  // ######################################################################
  //! Prologue for all other types for JSON archives (except minimal types)
  /*! Starts a new node, named either automatically or by some NVP,
      that may be given data by the type about to be archived

      Minimal types do not start or finish nodes */
  template <class T, traits::EnableIf<!std::is_arithmetic<T>::value,
                                      !traits::has_minimal_base_class_serialization<T, traits::has_minimal_output_serialization, JSONOutputArchive>::value,
                                      !traits::has_minimal_output_serialization<T, JSONOutputArchive>::value> = traits::sfinae>
  inline void prologue( JSONOutputArchive & ar, T const & )
  {
    ar.startNode();
  }

  //! Prologue for all other types for JSON archives
  template <class T, traits::EnableIf<!std::is_arithmetic<T>::value,
                                      !traits::has_minimal_base_class_serialization<T, traits::has_minimal_input_serialization, JSONInputArchive>::value,
                                      !traits::has_minimal_input_serialization<T, JSONInputArchive>::value> = traits::sfinae>
  inline void prologue( JSONInputArchive & ar, T const & )
  {
    ar.startNode();
  }

  // ######################################################################
  //! Epilogue for all other types other for JSON archives (except minimal types)
  /*! Finishes the node created in the prologue

      Minimal types do not start or finish nodes */
  template <class T, traits::EnableIf<!std::is_arithmetic<T>::value,
                                      !traits::has_minimal_base_class_serialization<T, traits::has_minimal_output_serialization, JSONOutputArchive>::value,
                                      !traits::has_minimal_output_serialization<T, JSONOutputArchive>::value> = traits::sfinae>
  inline void epilogue( JSONOutputArchive & ar, T const & )
  {
    ar.finishNode();
  }

  //! Epilogue for all other types other for JSON archives
  template <class T, traits::EnableIf<!std::is_arithmetic<T>::value,
                                      !traits::has_minimal_base_class_serialization<T, traits::has_minimal_input_serialization, JSONInputArchive>::value,
                                      !traits::has_minimal_input_serialization<T, JSONInputArchive>::value> = traits::sfinae>
  inline void epilogue( JSONInputArchive & ar, T const & )
  {
    ar.finishNode();
  }

  // ######################################################################
  //! Prologue for arithmetic types for JSON archives
  inline
  void prologue( JSONOutputArchive & ar, std::nullptr_t const & )
  {
    ar.writeName();
  }

  //! Prologue for arithmetic types for JSON archives
  inline
  void prologue( JSONInputArchive &, std::nullptr_t const & )
  { }

  // ######################################################################
  //! Epilogue for arithmetic types for JSON archives
  inline
  void epilogue( JSONOutputArchive &, std::nullptr_t const & )
  { }

  //! Epilogue for arithmetic types for JSON archives
  inline
  void epilogue( JSONInputArchive &, std::nullptr_t const & )
  { }

  // ######################################################################
  //! Prologue for arithmetic types for JSON archives
  template <class T, traits::EnableIf<std::is_arithmetic<T>::value> = traits::sfinae> inline
  void prologue( JSONOutputArchive & ar, T const & )
  {
    ar.writeName();
  }

  //! Prologue for arithmetic types for JSON archives
  template <class T, traits::EnableIf<std::is_arithmetic<T>::value> = traits::sfinae> inline
  void prologue( JSONInputArchive &, T const & )
  { }

  // ######################################################################
  //! Epilogue for arithmetic types for JSON archives
  template <class T, traits::EnableIf<std::is_arithmetic<T>::value> = traits::sfinae> inline
  void epilogue( JSONOutputArchive &, T const & )
  { }

  //! Epilogue for arithmetic types for JSON archives
  template <class T, traits::EnableIf<std::is_arithmetic<T>::value> = traits::sfinae> inline
  void epilogue( JSONInputArchive &, T const & )
  { }

  // ######################################################################
  //! Prologue for strings for JSON archives
  template<class CharT, class Traits, class Alloc> inline
  void prologue(JSONOutputArchive & ar, std::basic_string<CharT, Traits, Alloc> const &)
  {
    ar.writeName();
  }

  //! Prologue for strings for JSON archives
  template<class CharT, class Traits, class Alloc> inline
  void prologue(JSONInputArchive &, std::basic_string<CharT, Traits, Alloc> const &)
  { }

  // ######################################################################
  //! Epilogue for strings for JSON archives
  template<class CharT, class Traits, class Alloc> inline
  void epilogue(JSONOutputArchive &, std::basic_string<CharT, Traits, Alloc> const &)
  { }

  //! Epilogue for strings for JSON archives
  template<class CharT, class Traits, class Alloc> inline
  void epilogue(JSONInputArchive &, std::basic_string<CharT, Traits, Alloc> const &)
  { }

  // ######################################################################
  // Common JSONArchive serialization functions
  // ######################################################################
  //! Serializing NVP types to JSON
  template <class T> inline
  void CEREAL_SAVE_FUNCTION_NAME( JSONOutputArchive & ar, NameValuePair<T> const & t )
  {
    ar.setNextName( t.name );
    ar( t.value );
  }

  template <class T> inline
  void CEREAL_LOAD_FUNCTION_NAME( JSONInputArchive & ar, NameValuePair<T> & t )
  {
    ar.setNextName( t.name );
    ar( t.value );
  }

  //! Saving for nullptr to JSON
  inline
  void CEREAL_SAVE_FUNCTION_NAME(JSONOutputArchive & ar, std::nullptr_t const & t)
  {
    ar.saveValue( t );
  }

  //! Loading arithmetic from JSON
  inline
  void CEREAL_LOAD_FUNCTION_NAME(JSONInputArchive & ar, std::nullptr_t & t)
  {
    ar.loadValue( t );
  }

  //! Saving for arithmetic to JSON
  template <class T, traits::EnableIf<std::is_arithmetic<T>::value> = traits::sfinae> inline
  void CEREAL_SAVE_FUNCTION_NAME(JSONOutputArchive & ar, T const & t)
  {
    ar.saveValue( t );
  }

  //! Loading arithmetic from JSON
  template <class T, traits::EnableIf<std::is_arithmetic<T>::value> = traits::sfinae> inline
  void CEREAL_LOAD_FUNCTION_NAME(JSONInputArchive & ar, T & t)
  {
    ar.loadValue( t );
  }

  //! saving string to JSON
  template<class CharT, class Traits, class Alloc> inline
  void CEREAL_SAVE_FUNCTION_NAME(JSONOutputArchive & ar, std::basic_string<CharT, Traits, Alloc> const & str)
  {
    ar.saveValue( str );
  }

  //! loading string from JSON
  template<class CharT, class Traits, class Alloc> inline
  void CEREAL_LOAD_FUNCTION_NAME(JSONInputArchive & ar, std::basic_string<CharT, Traits, Alloc> & str)
  {
    ar.loadValue( str );
  }

  // ######################################################################
  //! Saving SizeTags to JSON
  template <class T> inline
  void CEREAL_SAVE_FUNCTION_NAME( JSONOutputArchive &, SizeTag<T> const & )
  {
    // nothing to do here, we don't explicitly save the size
  }

  //! Loading SizeTags from JSON
  template <class T> inline
  void CEREAL_LOAD_FUNCTION_NAME( JSONInputArchive & ar, SizeTag<T> & st )
  {
    ar.loadSize( st.size );
  }
} // namespace cereal

// register archives for polymorphic support
CEREAL_REGISTER_ARCHIVE(cereal::JSONInputArchive)
CEREAL_REGISTER_ARCHIVE(cereal::JSONOutputArchive)

// tie input and output archives together
CEREAL_SETUP_ARCHIVE_TRAITS(cereal::JSONInputArchive, cereal::JSONOutputArchive)

#endif // CEREAL_ARCHIVES_JSON_HPP_
```
