## Supplemental code for "The motif composition of variable-number tandem repeats impacts gene expression": portable_binary.hpp

```
/*! \file binary.hpp
    \brief Binary input and output archives */
/*
  Copyright (c) 2014, Randolph Voorhies, Shane Grant
  All rights reserved.

  Redistribution and use in source and binary forms, with or without
  modification, are permitted provided that the following conditions are met:
      * Redistributions of source code must retain the above copyright
        notice, this list of conditions and the following disclaimer.
      * Redistributions in binary form must reproduce the above copyright
        notice, this list of conditions and the following disclaimer in the
        documentation and/or other materials provided with the distribution.
      * Neither the name of cereal nor the
        names of its contributors may be used to endorse or promote products
        derived from this software without specific prior written permission.

  THIS SOFTWARE IS PROVIDED BY THE COPYRIGHT HOLDERS AND CONTRIBUTORS "AS IS" AND
  ANY EXPRESS OR IMPLIED WARRANTIES, INCLUDING, BUT NOT LIMITED TO, THE IMPLIED
  WARRANTIES OF MERCHANTABILITY AND FITNESS FOR A PARTICULAR PURPOSE ARE
  DISCLAIMED. IN NO EVENT SHALL RANDOLPH VOORHIES OR SHANE GRANT BE LIABLE FOR ANY
  DIRECT, INDIRECT, INCIDENTAL, SPECIAL, EXEMPLARY, OR CONSEQUENTIAL DAMAGES
  (INCLUDING, BUT NOT LIMITED TO, PROCUREMENT OF SUBSTITUTE GOODS OR SERVICES;
  LOSS OF USE, DATA, OR PROFITS; OR BUSINESS INTERRUPTION) HOWEVER CAUSED AND
  ON ANY THEORY OF LIABILITY, WHETHER IN CONTRACT, STRICT LIABILITY, OR TORT
  (INCLUDING NEGLIGENCE OR OTHERWISE) ARISING IN ANY WAY OUT OF THE USE OF THIS
  SOFTWARE, EVEN IF ADVISED OF THE POSSIBILITY OF SUCH DAMAGE.
*/
#ifndef CEREAL_ARCHIVES_PORTABLE_BINARY_HPP_
#define CEREAL_ARCHIVES_PORTABLE_BINARY_HPP_

#include "cereal/cereal.hpp"
#include <sstream>
#include <limits>

namespace cereal
{
  namespace portable_binary_detail
  {
    //! Returns true if the current machine is little endian
    /*! @ingroup Internal */
    inline std::uint8_t is_little_endian()
    {
      static std::int32_t test = 1;
      return *reinterpret_cast<std::int8_t*>( &test ) == 1;
    }

    //! Swaps the order of bytes for some chunk of memory
    /*! @param data The data as a uint8_t pointer
        @tparam DataSize The true size of the data
        @ingroup Internal */
    template <std::size_t DataSize>
    inline void swap_bytes( std::uint8_t * data )
    {
      for( std::size_t i = 0, end = DataSize / 2; i < end; ++i )
        std::swap( data[i], data[DataSize - i - 1] );
    }
  } // end namespace portable_binary_detail

  // ######################################################################
  //! An output archive designed to save data in a compact binary representation portable over different architectures
  /*! This archive outputs data to a stream in an extremely compact binary
      representation with as little extra metadata as possible.

      This archive will record the endianness of the data as well as the desired in/out endianness
      and assuming that the user takes care of ensuring serialized types are the same size
      across machines, is portable over different architectures.

      When using a binary archive and a file stream, you must use the
      std::ios::binary format flag to avoid having your data altered
      inadvertently.

      \warning This archive has not been thoroughly tested across different architectures.
               Please report any issues, optimizations, or feature requests at
               <a href="www.github.com/USCiLab/cereal">the project github</a>.

    \ingroup Archives */
  class PortableBinaryOutputArchive : public OutputArchive<PortableBinaryOutputArchive, AllowEmptyClassElision>
  {
    public:
      //! A class containing various advanced options for the PortableBinaryOutput archive
      class Options
      {
        public:
          //! Represents desired endianness
          enum class Endianness : std::uint8_t
          { big, little };

          //! Default options, preserve system endianness
          static Options Default(){ return Options(); }

          //! Save as little endian
          static Options LittleEndian(){ return Options( Endianness::little ); }

          //! Save as big endian
          static Options BigEndian(){ return Options( Endianness::big ); }

          //! Specify specific options for the PortableBinaryOutputArchive
          /*! @param outputEndian The desired endianness of saved (output) data */
          explicit Options( Endianness outputEndian = getEndianness() ) :
            itsOutputEndianness( outputEndian ) { }

        private:
          //! Gets the endianness of the system
          inline static Endianness getEndianness()
          { return portable_binary_detail::is_little_endian() ? Endianness::little : Endianness::big; }

          //! Checks if Options is set for little endian
          inline std::uint8_t is_little_endian() const
          { return itsOutputEndianness == Endianness::little; }

          friend class PortableBinaryOutputArchive;
          Endianness itsOutputEndianness;
      };

      //! Construct, outputting to the provided stream
      /*! @param stream The stream to output to. Should be opened with std::ios::binary flag.
          @param options The PortableBinary specific options to use.  See the Options struct
                         for the values of default parameters */
      PortableBinaryOutputArchive(std::ostream & stream, Options const & options = Options::Default()) :
        OutputArchive<PortableBinaryOutputArchive, AllowEmptyClassElision>(this),
        itsStream(stream),
        itsConvertEndianness( portable_binary_detail::is_little_endian() ^ options.is_little_endian() )
      {
        this->operator()( options.is_little_endian() );
      }

      ~PortableBinaryOutputArchive() CEREAL_NOEXCEPT = default;

      //! Writes size bytes of data to the output stream
      template <std::streamsize DataSize> inline
      void saveBinary( const void * data, std::streamsize size )
      {
        std::streamsize writtenSize = 0;

        if( itsConvertEndianness )
        {
          for( std::streamsize i = 0; i < size; i += DataSize )
            for( std::streamsize j = 0; j < DataSize; ++j )
              writtenSize += itsStream.rdbuf()->sputn( reinterpret_cast<const char*>( data ) + DataSize - j - 1 + i, 1 );
        }
        else
          writtenSize = itsStream.rdbuf()->sputn( reinterpret_cast<const char*>( data ), size );

        if(writtenSize != size)
          throw Exception("Failed to write " + std::to_string(size) + " bytes to output stream! Wrote " + std::to_string(writtenSize));
      }

    private:
      std::ostream & itsStream;
      const uint8_t itsConvertEndianness; //!< If set to true, we will need to swap bytes upon saving
  };

  // ######################################################################
  //! An input archive designed to load data saved using PortableBinaryOutputArchive
  /*! This archive outputs data to a stream in an extremely compact binary
      representation with as little extra metadata as possible.

      This archive will load the endianness of the serialized data and
      if necessary transform it to match that of the local machine.  This comes
      at a significant performance cost compared to non portable archives if
      the transformation is necessary, and also causes a small performance hit
      even if it is not necessary.

      It is recommended to use portable archives only if you know that you will
      be sending binary data to machines with different endianness.

      The archive will do nothing to ensure types are the same size - that is
      the responsibility of the user.

      When using a binary archive and a file stream, you must use the
      std::ios::binary format flag to avoid having your data altered
      inadvertently.

      \warning This archive has not been thoroughly tested across different architectures.
               Please report any issues, optimizations, or feature requests at
               <a href="www.github.com/USCiLab/cereal">the project github</a>.

    \ingroup Archives */
  class PortableBinaryInputArchive : public InputArchive<PortableBinaryInputArchive, AllowEmptyClassElision>
  {
    public:
      //! A class containing various advanced options for the PortableBinaryInput archive
      class Options
      {
        public:
          //! Represents desired endianness
          enum class Endianness : std::uint8_t
          { big, little };

          //! Default options, preserve system endianness
          static Options Default(){ return Options(); }

          //! Load into little endian
          static Options LittleEndian(){ return Options( Endianness::little ); }

          //! Load into big endian
          static Options BigEndian(){ return Options( Endianness::big ); }

          //! Specify specific options for the PortableBinaryInputArchive
          /*! @param inputEndian The desired endianness of loaded (input) data */
          explicit Options( Endianness inputEndian = getEndianness() ) :
            itsInputEndianness( inputEndian ) { }

        private:
          //! Gets the endianness of the system
          inline static Endianness getEndianness()
          { return portable_binary_detail::is_little_endian() ? Endianness::little : Endianness::big; }

          //! Checks if Options is set for little endian
          inline std::uint8_t is_little_endian() const
          { return itsInputEndianness == Endianness::little; }

          friend class PortableBinaryInputArchive;
          Endianness itsInputEndianness;
      };

      //! Construct, loading from the provided stream
      /*! @param stream The stream to read from. Should be opened with std::ios::binary flag.
          @param options The PortableBinary specific options to use.  See the Options struct
                         for the values of default parameters */
      PortableBinaryInputArchive(std::istream & stream, Options const & options = Options::Default()) :
        InputArchive<PortableBinaryInputArchive, AllowEmptyClassElision>(this),
        itsStream(stream),
        itsConvertEndianness( false )
      {
        uint8_t streamLittleEndian;
        this->operator()( streamLittleEndian );
        itsConvertEndianness = options.is_little_endian() ^ streamLittleEndian;
      }

      ~PortableBinaryInputArchive() CEREAL_NOEXCEPT = default;

      //! Reads size bytes of data from the input stream
      /*! @param data The data to save
          @param size The number of bytes in the data
          @tparam DataSize T The size of the actual type of the data elements being loaded */
      template <std::streamsize DataSize> inline
      void loadBinary( void * const data, std::streamsize size )
      {
        // load data
        auto const readSize = itsStream.rdbuf()->sgetn( reinterpret_cast<char*>( data ), size );

        if(readSize != size)
          throw Exception("Failed to read " + std::to_string(size) + " bytes from input stream! Read " + std::to_string(readSize));

        // flip bits if needed
        if( itsConvertEndianness )
        {
          std::uint8_t * ptr = reinterpret_cast<std::uint8_t*>( data );
          for( std::streamsize i = 0; i < size; i += DataSize )
            portable_binary_detail::swap_bytes<DataSize>( ptr + i );
        }
      }

    private:
      std::istream & itsStream;
      uint8_t itsConvertEndianness; //!< If set to true, we will need to swap bytes upon loading
  };

  // ######################################################################
  // Common BinaryArchive serialization functions

  //! Saving for POD types to portable binary
  template<class T> inline
  typename std::enable_if<std::is_arithmetic<T>::value, void>::type
  CEREAL_SAVE_FUNCTION_NAME(PortableBinaryOutputArchive & ar, T const & t)
  {
    static_assert( !std::is_floating_point<T>::value ||
                   (std::is_floating_point<T>::value && std::numeric_limits<T>::is_iec559),
                   "Portable binary only supports IEEE 754 standardized floating point" );
    ar.template saveBinary<sizeof(T)>(std::addressof(t), sizeof(t));
  }

  //! Loading for POD types from portable binary
  template<class T> inline
  typename std::enable_if<std::is_arithmetic<T>::value, void>::type
  CEREAL_LOAD_FUNCTION_NAME(PortableBinaryInputArchive & ar, T & t)
  {
    static_assert( !std::is_floating_point<T>::value ||
                   (std::is_floating_point<T>::value && std::numeric_limits<T>::is_iec559),
                   "Portable binary only supports IEEE 754 standardized floating point" );
    ar.template loadBinary<sizeof(T)>(std::addressof(t), sizeof(t));
  }

  //! Serializing NVP types to portable binary
  template <class Archive, class T> inline
  CEREAL_ARCHIVE_RESTRICT(PortableBinaryInputArchive, PortableBinaryOutputArchive)
  CEREAL_SERIALIZE_FUNCTION_NAME( Archive & ar, NameValuePair<T> & t )
  {
    ar( t.value );
  }

  //! Serializing SizeTags to portable binary
  template <class Archive, class T> inline
  CEREAL_ARCHIVE_RESTRICT(PortableBinaryInputArchive, PortableBinaryOutputArchive)
  CEREAL_SERIALIZE_FUNCTION_NAME( Archive & ar, SizeTag<T> & t )
  {
    ar( t.size );
  }

  //! Saving binary data to portable binary
  template <class T> inline
  void CEREAL_SAVE_FUNCTION_NAME(PortableBinaryOutputArchive & ar, BinaryData<T> const & bd)
  {
    typedef typename std::remove_pointer<T>::type TT;
    static_assert( !std::is_floating_point<TT>::value ||
                   (std::is_floating_point<TT>::value && std::numeric_limits<TT>::is_iec559),
                   "Portable binary only supports IEEE 754 standardized floating point" );

    ar.template saveBinary<sizeof(TT)>( bd.data, static_cast<std::streamsize>( bd.size ) );
  }

  //! Loading binary data from portable binary
  template <class T> inline
  void CEREAL_LOAD_FUNCTION_NAME(PortableBinaryInputArchive & ar, BinaryData<T> & bd)
  {
    typedef typename std::remove_pointer<T>::type TT;
    static_assert( !std::is_floating_point<TT>::value ||
                   (std::is_floating_point<TT>::value && std::numeric_limits<TT>::is_iec559),
                   "Portable binary only supports IEEE 754 standardized floating point" );

    ar.template loadBinary<sizeof(TT)>( bd.data, static_cast<std::streamsize>( bd.size ) );
  }
} // namespace cereal

// register archives for polymorphic support
CEREAL_REGISTER_ARCHIVE(cereal::PortableBinaryOutputArchive)
CEREAL_REGISTER_ARCHIVE(cereal::PortableBinaryInputArchive)

// tie input and output archives together
CEREAL_SETUP_ARCHIVE_TRAITS(cereal::PortableBinaryInputArchive, cereal::PortableBinaryOutputArchive)

#endif // CEREAL_ARCHIVES_PORTABLE_BINARY_HPP_
```
