## Supplemental code for "The motif composition of variable-number tandem repeats impacts gene expression": xml.hpp

```
/*! \file xml.hpp
    \brief XML input and output archives */
/*
  Copyright (c) 2014, Randolph Voorhies, Shane Grant
  All rights reserved.

  Redistribution and use in source and binary forms, with or without
  modification, are permitted provided that the following conditions are met:
      * Redistributions of source code must retain the above copyright
        notice, this list of conditions and the following disclaimer.
      * Redistributions in binary form must reproduce the above copyright
        notice, this list of conditions and the following disclaimer in the
        documentation and/or other materials provided with the distribution.
      * Neither the name of cereal nor the
        names of its contributors may be used to endorse or promote products
        derived from this software without specific prior written permission.

  THIS SOFTWARE IS PROVIDED BY THE COPYRIGHT HOLDERS AND CONTRIBUTORS "AS IS" AND
  ANY EXPRESS OR IMPLIED WARRANTIES, INCLUDING, BUT NOT LIMITED TO, THE IMPLIED
  WARRANTIES OF MERCHANTABILITY AND FITNESS FOR A PARTICULAR PURPOSE ARE
  DISCLAIMED. IN NO EVENT SHALL RANDOLPH VOORHIES OR SHANE GRANT BE LIABLE FOR ANY
  DIRECT, INDIRECT, INCIDENTAL, SPECIAL, EXEMPLARY, OR CONSEQUENTIAL DAMAGES
  (INCLUDING, BUT NOT LIMITED TO, PROCUREMENT OF SUBSTITUTE GOODS OR SERVICES;
  LOSS OF USE, DATA, OR PROFITS; OR BUSINESS INTERRUPTION) HOWEVER CAUSED AND
  ON ANY THEORY OF LIABILITY, WHETHER IN CONTRACT, STRICT LIABILITY, OR TORT
  (INCLUDING NEGLIGENCE OR OTHERWISE) ARISING IN ANY WAY OUT OF THE USE OF THIS
  SOFTWARE, EVEN IF ADVISED OF THE POSSIBILITY OF SUCH DAMAGE.
*/
#ifndef CEREAL_ARCHIVES_XML_HPP_
#define CEREAL_ARCHIVES_XML_HPP_
#include "cereal/cereal.hpp"
#include "cereal/details/util.hpp"

#include "cereal/external/rapidxml/rapidxml.hpp"
#include "cereal/external/rapidxml/rapidxml_print.hpp"
#include "cereal/external/base64.hpp"

#include <sstream>
#include <stack>
#include <vector>
#include <limits>
#include <string>
#include <cstring>
#include <cmath>

namespace cereal
{
  namespace xml_detail
  {
    #ifndef CEREAL_XML_STRING_VALUE
    //! The default name for the root node in a cereal xml archive.
    /*! You can define CEREAL_XML_STRING_VALUE to be different assuming you do so
        before this file is included. */
    #define CEREAL_XML_STRING_VALUE "cereal"
    #endif // CEREAL_XML_STRING_VALUE

    //! The name given to the root node in a cereal xml archive
    static const char * CEREAL_XML_STRING = CEREAL_XML_STRING_VALUE;

    //! Returns true if the character is whitespace
    inline bool isWhitespace( char c )
    {
      return c == ' ' || c == '\t' || c == '\n' || c == '\r';
    }
  }

  // ######################################################################
  //! An output archive designed to save data to XML
  /*! This archive uses RapidXML to build an in memory XML tree of the
      data it serializes before outputting it to its stream upon destruction.
      This archive should be used in an RAII fashion, letting
      the automatic destruction of the object cause the flush to its stream.

      XML archives provides a human readable output but at decreased
      performance (both in time and space) compared to binary archives.

      XML benefits greatly from name-value pairs, which if present, will
      name the nodes in the output.  If these are not present, each level
      of the output tree will be given an automatically generated delimited name.

      The precision of the output archive controls the number of decimals output
      for floating point numbers and should be sufficiently large (i.e. at least 20)
      if there is a desire to have binary equality between the numbers output and
      those read in.  In general you should expect a loss of precision when going
      from floating point to text and back.

      XML archives can optionally print the type of everything they serialize, which
      adds an attribute to each node.

      XML archives do not output the size information for any dynamically sized structure
      and instead infer it from the number of children for a node.  This means that data
      can be hand edited for dynamic sized structures and will still be readable.  This
      is accomplished through the cereal::SizeTag object, which will also add an attribute
      to its parent field.
      \ingroup Archives */
  class XMLOutputArchive : public OutputArchive<XMLOutputArchive>, public traits::TextArchive
  {
    public:
      /*! @name Common Functionality
          Common use cases for directly interacting with an XMLOutputArchive */
      //! @{

      //! A class containing various advanced options for the XML archive
      /*! Options can either be directly passed to the constructor, or chained using the
          modifier functions for an interface analogous to named parameters */
      class Options
      {
        public:
          //! Default options
          static Options Default(){ return Options(); }

          //! Specify specific options for the XMLOutputArchive
          /*! @param precision_ The precision used for floating point numbers
              @param indent_ Whether to indent each line of XML
              @param outputType_ Whether to output the type of each serialized object as an attribute
              @param sizeAttributes_ Whether dynamically sized containers output the size=dynamic attribute */
          explicit Options( int precision_ = std::numeric_limits<double>::max_digits10,
                            bool indent_ = true,
                            bool outputType_ = false,
                            bool sizeAttributes_ = true ) :
            itsPrecision( precision_ ),
            itsIndent( indent_ ),
            itsOutputType( outputType_ ),
            itsSizeAttributes( sizeAttributes_ )
          { }

          /*! @name Option Modifiers
              An interface for setting option settings analogous to named parameters.

              @code{cpp}
              cereal::XMLOutputArchive ar( myStream,
                                           cereal::XMLOutputArchive::Options()
                                           .indent(true)
                                           .sizeAttributes(false) );
              @endcode
              */
          //! @{

          //! Sets the precision used for floaing point numbers
          Options & precision( int value ){ itsPrecision = value; return * this; }
          //! Whether to indent each line of XML
          Options & indent( bool enable ){ itsIndent = enable; return *this; }
          //! Whether to output the type of each serialized object as an attribute
          Options & outputType( bool enable ){ itsOutputType = enable; return *this; }
          //! Whether dynamically sized containers (e.g. vector) output the size=dynamic attribute
          Options & sizeAttributes( bool enable ){ itsSizeAttributes = enable; return *this; }

          //! @}

        private:
          friend class XMLOutputArchive;
          int itsPrecision;
          bool itsIndent;
          bool itsOutputType;
          bool itsSizeAttributes;
      };

      //! Construct, outputting to the provided stream upon destruction
      /*! @param stream  The stream to output to.  Note that XML is only guaranteed to flush
                         its output to the stream upon destruction.
          @param options The XML specific options to use.  See the Options struct
                         for the values of default parameters */
      XMLOutputArchive( std::ostream & stream, Options const & options = Options::Default() ) :
        OutputArchive<XMLOutputArchive>(this),
        itsStream(stream),
        itsOutputType( options.itsOutputType ),
        itsIndent( options.itsIndent ),
        itsSizeAttributes(options.itsSizeAttributes)
      {
        // rapidxml will delete all allocations when xml_document is cleared
        auto node = itsXML.allocate_node( rapidxml::node_declaration );
        node->append_attribute( itsXML.allocate_attribute( "version", "1.0" ) );
        node->append_attribute( itsXML.allocate_attribute( "encoding", "utf-8" ) );
        itsXML.append_node( node );

        // allocate root node
        auto root = itsXML.allocate_node( rapidxml::node_element, xml_detail::CEREAL_XML_STRING );
        itsXML.append_node( root );
        itsNodes.emplace( root );

        // set attributes on the streams
        itsStream << std::boolalpha;
        itsStream.precision( options.itsPrecision );
        itsOS << std::boolalpha;
        itsOS.precision( options.itsPrecision );
      }

      //! Destructor, flushes the XML
      ~XMLOutputArchive() CEREAL_NOEXCEPT
      {
        const int flags = itsIndent ? 0x0 : rapidxml::print_no_indenting;
        rapidxml::print( itsStream, itsXML, flags );
        itsXML.clear();
      }

      //! Saves some binary data, encoded as a base64 string, with an optional name
      /*! This can be called directly by users and it will automatically create a child node for
          the current XML node, populate it with a base64 encoded string, and optionally name
          it.  The node will be finished after it has been populated.  */
      void saveBinaryValue( const void * data, size_t size, const char * name = nullptr )
      {
        itsNodes.top().name = name;

        startNode();

        auto base64string = base64::encode( reinterpret_cast<const unsigned char *>( data ), size );
        saveValue( base64string );

        if( itsOutputType )
          itsNodes.top().node->append_attribute( itsXML.allocate_attribute( "type", "cereal binary data" ) );

        finishNode();
      }

      //! @}
      /*! @name Internal Functionality
          Functionality designed for use by those requiring control over the inner mechanisms of
          the XMLOutputArchive */
      //! @{

      //! Creates a new node that is a child of the node at the top of the stack
      /*! Nodes will be given a name that has either been pre-set by a name value pair,
          or generated based upon a counter unique to the parent node.  If you want to
          give a node a specific name, use setNextName prior to calling startNode.

          The node will then be pushed onto the node stack. */
      void startNode()
      {
        // generate a name for this new node
        const auto nameString = itsNodes.top().getValueName();

        // allocate strings for all of the data in the XML object
        auto namePtr = itsXML.allocate_string( nameString.data(), nameString.length() + 1 );

        // insert into the XML
        auto node = itsXML.allocate_node( rapidxml::node_element, namePtr, nullptr, nameString.size() );
        itsNodes.top().node->append_node( node );
        itsNodes.emplace( node );
      }

      //! Designates the most recently added node as finished
      void finishNode()
      {
        itsNodes.pop();
      }

      //! Sets the name for the next node created with startNode
      void setNextName( const char * name )
      {
        itsNodes.top().name = name;
      }

      //! Saves some data, encoded as a string, into the current top level node
      /*! The data will be be named with the most recent name if one exists,
          otherwise it will be given some default delimited value that depends upon
          the parent node */
      template <class T> inline
      void saveValue( T const & value )
      {
        itsOS.clear(); itsOS.seekp( 0, std::ios::beg );
        itsOS << value << std::ends;

        auto strValue = itsOS.str();

        // itsOS.str() may contain data from previous calls after the first '\0' that was just inserted
        // and this data is counted in the length call. We make sure to remove that section so that the
        // whitespace validation is done properly
        strValue.resize(std::strlen(strValue.c_str()));

        // If the first or last character is a whitespace, add xml:space attribute
        const auto len = strValue.length();
        if ( len > 0 && ( xml_detail::isWhitespace( strValue[0] ) || xml_detail::isWhitespace( strValue[len - 1] ) ) )
        {
          itsNodes.top().node->append_attribute( itsXML.allocate_attribute( "xml:space", "preserve" ) );
        }

        // allocate strings for all of the data in the XML object
        auto dataPtr = itsXML.allocate_string(strValue.c_str(), strValue.length() + 1 );

        // insert into the XML
        itsNodes.top().node->append_node( itsXML.allocate_node( rapidxml::node_data, nullptr, dataPtr ) );
      }

      //! Overload for uint8_t prevents them from being serialized as characters
      void saveValue( uint8_t const & value )
      {
        saveValue( static_cast<uint32_t>( value ) );
      }

      //! Overload for int8_t prevents them from being serialized as characters
      void saveValue( int8_t const & value )
      {
        saveValue( static_cast<int32_t>( value ) );
      }

      //! Causes the type to be appended as an attribute to the most recently made node if output type is set to true
      template <class T> inline
      void insertType()
      {
        if( !itsOutputType )
          return;

        // generate a name for this new node
        const auto nameString = util::demangledName<T>();

        // allocate strings for all of the data in the XML object
        auto namePtr = itsXML.allocate_string( nameString.data(), nameString.length() + 1 );

        itsNodes.top().node->append_attribute( itsXML.allocate_attribute( "type", namePtr ) );
      }

      //! Appends an attribute to the current top level node
      void appendAttribute( const char * name, const char * value )
      {
        auto namePtr =  itsXML.allocate_string( name );
        auto valuePtr = itsXML.allocate_string( value );
        itsNodes.top().node->append_attribute( itsXML.allocate_attribute( namePtr, valuePtr ) );
      }

      bool hasSizeAttributes() const { return itsSizeAttributes; }

    protected:
      //! A struct that contains metadata about a node
      struct NodeInfo
      {
        NodeInfo( rapidxml::xml_node<> * n = nullptr,
                  const char * nm = nullptr ) :
          node( n ),
          counter( 0 ),
          name( nm )
        { }

        rapidxml::xml_node<> * node; //!< A pointer to this node
        size_t counter;              //!< The counter for naming child nodes
        const char * name;           //!< The name for the next child node

        //! Gets the name for the next child node created from this node
        /*! The name will be automatically generated using the counter if
            a name has not been previously set.  If a name has been previously
            set, that name will be returned only once */
        std::string getValueName()
        {
          if( name )
          {
            auto n = name;
            name = nullptr;
            return {n};
          }
          else
            return "value" + std::to_string( counter++ ) + "\0";
        }
      }; // NodeInfo

      //! @}

    private:
      std::ostream & itsStream;        //!< The output stream
      rapidxml::xml_document<> itsXML; //!< The XML document
      std::stack<NodeInfo> itsNodes;   //!< A stack of nodes added to the document
      std::ostringstream itsOS;        //!< Used to format strings internally
      bool itsOutputType;              //!< Controls whether type information is printed
      bool itsIndent;                  //!< Controls whether indenting is used
      bool itsSizeAttributes;          //!< Controls whether lists have a size attribute
  }; // XMLOutputArchive

  // ######################################################################
  //! An output archive designed to load data from XML
  /*! This archive uses RapidXML to build an in memory XML tree of the
      data in the stream it is given before loading any types serialized.

      As with the output XML archive, the preferred way to use this archive is in
      an RAII fashion, ensuring its destruction after all data has been read.

      Input XML should have been produced by the XMLOutputArchive.  Data can
      only be added to dynamically sized containers - the input archive will
      determine their size by looking at the number of child nodes.  Data that
      did not originate from an XMLOutputArchive is not officially supported,
      but may be possible to use if properly formatted.

      The XMLInputArchive does not require that nodes are loaded in the same
      order they were saved by XMLOutputArchive.  Using name value pairs (NVPs),
      it is possible to load in an out of order fashion or otherwise skip/select
      specific nodes to load.

      The default behavior of the input archive is to read sequentially starting
      with the first node and exploring its children.  When a given NVP does
      not match the read in name for a node, the archive will search for that
      node at the current level and load it if it exists.  After loading an out of
      order node, the archive will then proceed back to loading sequentially from
      its new position.

      Consider this simple example where loading of some data is skipped:

      @code{cpp}
      // imagine the input file has someData(1-9) saved in order at the top level node
      ar( someData1, someData2, someData3 );        // XML loads in the order it sees in the file
      ar( cereal::make_nvp( "hello", someData6 ) ); // NVP given does not
                                                    // match expected NVP name, so we search
                                                    // for the given NVP and load that value
      ar( someData7, someData8, someData9 );        // with no NVP given, loading resumes at its
                                                    // current location, proceeding sequentially
      @endcode

      \ingroup Archives */
  class XMLInputArchive : public InputArchive<XMLInputArchive>, public traits::TextArchive
  {
    public:
      /*! @name Common Functionality
          Common use cases for directly interacting with an XMLInputArchive */
      //! @{

      //! Construct, reading in from the provided stream
      /*! Reads in an entire XML document from some stream and parses it as soon
          as serialization starts

          @param stream The stream to read from.  Can be a stringstream or a file. */
      XMLInputArchive( std::istream & stream ) :
        InputArchive<XMLInputArchive>( this ),
        itsData( std::istreambuf_iterator<char>( stream ), std::istreambuf_iterator<char>() )
      {
        try
        {
          itsData.push_back('\0'); // rapidxml will do terrible things without the data being null terminated
          itsXML.parse<rapidxml::parse_trim_whitespace | rapidxml::parse_no_data_nodes | rapidxml::parse_declaration_node>( reinterpret_cast<char *>( itsData.data() ) );
        }
        catch( rapidxml::parse_error const & )
        {
          //std::cerr << "-----Original-----" << std::endl;
          //stream.seekg(0);
          //std::cout << std::string( std::istreambuf_iterator<char>( stream ), std::istreambuf_iterator<char>() ) << std::endl;

          //std::cerr << "-----Error-----" << std::endl;
          //std::cerr << e.what() << std::endl;
          //std::cerr << e.where<char>() << std::endl;
          throw Exception("XML Parsing failed - likely due to invalid characters or invalid naming");
        }

        // Parse the root
        auto root = itsXML.first_node( xml_detail::CEREAL_XML_STRING );
        if( root == nullptr )
          throw Exception("Could not detect cereal root node - likely due to empty or invalid input");
        else
          itsNodes.emplace( root );
      }

      ~XMLInputArchive() CEREAL_NOEXCEPT = default;

      //! Loads some binary data, encoded as a base64 string, optionally specified by some name
      /*! This will automatically start and finish a node to load the data, and can be called directly by
          users.

          Note that this follows the same ordering rules specified in the class description in regards
          to loading in/out of order */
      void loadBinaryValue( void * data, size_t size, const char * name = nullptr )
      {
        setNextName( name );
        startNode();

        std::string encoded;
        loadValue( encoded );

        auto decoded = base64::decode( encoded );

        if( size != decoded.size() )
          throw Exception("Decoded binary data size does not match specified size");

        std::memcpy( data, decoded.data(), decoded.size() );

        finishNode();
      }

      //! @}
      /*! @name Internal Functionality
          Functionality designed for use by those requiring control over the inner mechanisms of
          the XMLInputArchive */
      //! @{

      //! Prepares to start reading the next node
      /*! This places the next node to be parsed onto the nodes stack.

          By default our strategy is to start with the document root node and then
          recursively iterate through all children in the order they show up in the document.
          We don't need to know NVPs do to this; we'll just blindly load in the order things appear in.

          We check to see if the specified NVP matches what the next automatically loaded node is.  If they
          match, we just continue as normal, going in order.  If they don't match, we attempt to find a node
          named after the NVP that is being loaded.  If that NVP does not exist, we throw an exception. */
      void startNode()
      {
        auto next = itsNodes.top().child; // By default we would move to the next child node
        auto const expectedName = itsNodes.top().name; // this is the expected name from the NVP, if provided

        // If we were given an NVP name, look for it in the current level of the document.
        //    We only need to do this if either we have exhausted the siblings of the current level or
        //    the NVP name does not match the name of the node we would normally read next
        if( expectedName && ( next == nullptr || std::strcmp( next->name(), expectedName ) != 0 ) )
        {
          next = itsNodes.top().search( expectedName );

          if( next == nullptr )
            throw Exception("XML Parsing failed - provided NVP (" + std::string(expectedName) + ") not found");
        }

        itsNodes.emplace( next );
      }

      //! Finishes reading the current node
      void finishNode()
      {
        // remove current
        itsNodes.pop();

        // advance parent
        itsNodes.top().advance();

        // Reset name
        itsNodes.top().name = nullptr;
      }

      //! Retrieves the current node name
      //! will return @c nullptr if the node does not have a name
      const char * getNodeName() const
      {
        return itsNodes.top().getChildName();
      }

      //! Sets the name for the next node created with startNode
      void setNextName( const char * name )
      {
        itsNodes.top().name = name;
      }

      //! Loads a bool from the current top node
      template <class T, traits::EnableIf<std::is_unsigned<T>::value,
                                          std::is_same<T, bool>::value> = traits::sfinae> inline
      void loadValue( T & value )
      {
        std::istringstream is( itsNodes.top().node->value() );
        is.setf( std::ios::boolalpha );
        is >> value;
      }

      //! Loads a char (signed or unsigned) from the current top node
      template <class T, traits::EnableIf<std::is_integral<T>::value,
                                          !std::is_same<T, bool>::value,
                                          sizeof(T) == sizeof(char)> = traits::sfinae> inline
      void loadValue( T & value )
      {
        value = *reinterpret_cast<T*>( itsNodes.top().node->value() );
      }

      //! Load an int8_t from the current top node (ensures we parse entire number)
      void loadValue( int8_t & value )
      {
        int32_t val; loadValue( val ); value = static_cast<int8_t>( val );
      }

      //! Load a uint8_t from the current top node (ensures we parse entire number)
      void loadValue( uint8_t & value )
      {
        uint32_t val; loadValue( val ); value = static_cast<uint8_t>( val );
      }

      //! Loads a type best represented as an unsigned long from the current top node
      template <class T, traits::EnableIf<std::is_unsigned<T>::value,
                                          !std::is_same<T, bool>::value,
                                          !std::is_same<T, char>::value,
                                          !std::is_same<T, unsigned char>::value,
                                          sizeof(T) < sizeof(long long)> = traits::sfinae> inline
      void loadValue( T & value )
      {
        value = static_cast<T>( std::stoul( itsNodes.top().node->value() ) );
      }

      //! Loads a type best represented as an unsigned long long from the current top node
      template <class T, traits::EnableIf<std::is_unsigned<T>::value,
                                          !std::is_same<T, bool>::value,
                                          sizeof(T) >= sizeof(long long)> = traits::sfinae> inline
      void loadValue( T & value )
      {
        value = static_cast<T>( std::stoull( itsNodes.top().node->value() ) );
      }

      //! Loads a type best represented as an int from the current top node
      template <class T, traits::EnableIf<std::is_signed<T>::value,
                                          !std::is_same<T, char>::value,
                                          sizeof(T) <= sizeof(int)> = traits::sfinae> inline
      void loadValue( T & value )
      {
        value = static_cast<T>( std::stoi( itsNodes.top().node->value() ) );
      }

      //! Loads a type best represented as a long from the current top node
      template <class T, traits::EnableIf<std::is_signed<T>::value,
                                          (sizeof(T) > sizeof(int)),
                                          sizeof(T) <= sizeof(long)> = traits::sfinae> inline
      void loadValue( T & value )
      {
        value = static_cast<T>( std::stol( itsNodes.top().node->value() ) );
      }

      //! Loads a type best represented as a long long from the current top node
      template <class T, traits::EnableIf<std::is_signed<T>::value,
                                          (sizeof(T) > sizeof(long)),
                                          sizeof(T) <= sizeof(long long)> = traits::sfinae> inline
      void loadValue( T & value )
      {
        value = static_cast<T>( std::stoll( itsNodes.top().node->value() ) );
      }

      //! Loads a type best represented as a float from the current top node
      void loadValue( float & value )
      {
        try
        {
          value = std::stof( itsNodes.top().node->value() );
        }
        catch( std::out_of_range const & )
        {
          // special case for denormalized values
          std::istringstream is( itsNodes.top().node->value() );
          is >> value;
          if( std::fpclassify( value ) != FP_SUBNORMAL )
            throw;
        }
      }

      //! Loads a type best represented as a double from the current top node
      void loadValue( double & value )
      {
        try
        {
          value = std::stod( itsNodes.top().node->value() );
        }
        catch( std::out_of_range const & )
        {
          // special case for denormalized values
          std::istringstream is( itsNodes.top().node->value() );
          is >> value;
          if( std::fpclassify( value ) != FP_SUBNORMAL )
            throw;
        }
      }

      //! Loads a type best represented as a long double from the current top node
      void loadValue( long double & value )
      {
        try
        {
          value = std::stold( itsNodes.top().node->value() );
        }
        catch( std::out_of_range const & )
        {
          // special case for denormalized values
          std::istringstream is( itsNodes.top().node->value() );
          is >> value;
          if( std::fpclassify( value ) != FP_SUBNORMAL )
            throw;
        }
      }

      //! Loads a string from the current node from the current top node
      template<class CharT, class Traits, class Alloc> inline
      void loadValue( std::basic_string<CharT, Traits, Alloc> & str )
      {
        std::basic_istringstream<CharT, Traits> is( itsNodes.top().node->value() );

        str.assign( std::istreambuf_iterator<CharT, Traits>( is ),
                    std::istreambuf_iterator<CharT, Traits>() );
      }

      //! Loads the size of the current top node
      template <class T> inline
      void loadSize( T & value )
      {
        value = getNumChildren( itsNodes.top().node );
      }

    protected:
      //! Gets the number of children (usually interpreted as size) for the specified node
      static size_t getNumChildren( rapidxml::xml_node<> * node )
      {
        size_t size = 0;
        node = node->first_node(); // get first child

        while( node != nullptr )
        {
          ++size;
          node = node->next_sibling();
        }

        return size;
      }

      //! A struct that contains metadata about a node
      /*! Keeps track of some top level node, its number of
          remaining children, and the current active child node */
      struct NodeInfo
      {
        NodeInfo( rapidxml::xml_node<> * n = nullptr ) :
          node( n ),
          child( n->first_node() ),
          size( XMLInputArchive::getNumChildren( n ) ),
          name( nullptr )
        { }

        //! Advances to the next sibling node of the child
        /*! If this is the last sibling child will be null after calling */
        void advance()
        {
          if( size > 0 )
          {
            --size;
            child = child->next_sibling();
          }
        }

        //! Searches for a child with the given name in this node
        /*! @param searchName The name to search for (must be null terminated)
            @return The node if found, nullptr otherwise */
        rapidxml::xml_node<> * search( const char * searchName )
        {
          if( searchName )
          {
            size_t new_size = XMLInputArchive::getNumChildren( node );
            const size_t name_size = rapidxml::internal::measure( searchName );

            for( auto new_child = node->first_node(); new_child != nullptr; new_child = new_child->next_sibling() )
            {
              if( rapidxml::internal::compare( new_child->name(), new_child->name_size(), searchName, name_size, true ) )
              {
                size = new_size;
                child = new_child;

                return new_child;
              }
              --new_size;
            }
          }

          return nullptr;
        }

        //! Returns the actual name of the next child node, if it exists
        const char * getChildName() const
        {
          return child ? child->name() : nullptr;
        }

        rapidxml::xml_node<> * node;  //!< A pointer to this node
        rapidxml::xml_node<> * child; //!< A pointer to its current child
        size_t size;                  //!< The remaining number of children for this node
        const char * name;            //!< The NVP name for next child node
      }; // NodeInfo

      //! @}

    private:
      std::vector<char> itsData;       //!< The raw data loaded
      rapidxml::xml_document<> itsXML; //!< The XML document
      std::stack<NodeInfo> itsNodes;   //!< A stack of nodes read from the document
  };

  // ######################################################################
  // XMLArchive prologue and epilogue functions
  // ######################################################################

  // ######################################################################
  //! Prologue for NVPs for XML output archives
  /*! NVPs do not start or finish nodes - they just set up the names */
  template <class T> inline
  void prologue( XMLOutputArchive &, NameValuePair<T> const & )
  { }

  //! Prologue for NVPs for XML input archives
  template <class T> inline
  void prologue( XMLInputArchive &, NameValuePair<T> const & )
  { }

  // ######################################################################
  //! Epilogue for NVPs for XML output archives
  /*! NVPs do not start or finish nodes - they just set up the names */
  template <class T> inline
  void epilogue( XMLOutputArchive &, NameValuePair<T> const & )
  { }

  //! Epilogue for NVPs for XML input archives
  template <class T> inline
  void epilogue( XMLInputArchive &, NameValuePair<T> const & )
  { }

  // ######################################################################
  //! Prologue for deferred data for XML archives
  /*! Do nothing for the defer wrapper */
  template <class T> inline
  void prologue( XMLOutputArchive &, DeferredData<T> const & )
  { }

  //! Prologue for deferred data for XML archives
  template <class T> inline
  void prologue( XMLInputArchive &, DeferredData<T> const & )
  { }

  // ######################################################################
  //! Epilogue for deferred for XML archives
  /*! NVPs do not start or finish nodes - they just set up the names */
  template <class T> inline
  void epilogue( XMLOutputArchive &, DeferredData<T> const & )
  { }

  //! Epilogue for deferred for XML archives
  /*! Do nothing for the defer wrapper */
  template <class T> inline
  void epilogue( XMLInputArchive &, DeferredData<T> const & )
  { }

  // ######################################################################
  //! Prologue for SizeTags for XML output archives
  /*! SizeTags do not start or finish nodes */
  template <class T> inline
  void prologue( XMLOutputArchive & ar, SizeTag<T> const & )
  {
      if (ar.hasSizeAttributes())
      {
          ar.appendAttribute("size", "dynamic");
      }
  }

  template <class T> inline
  void prologue( XMLInputArchive &, SizeTag<T> const & )
  { }

  //! Epilogue for SizeTags for XML output archives
  /*! SizeTags do not start or finish nodes */
  template <class T> inline
  void epilogue( XMLOutputArchive &, SizeTag<T> const & )
  { }

  template <class T> inline
  void epilogue( XMLInputArchive &, SizeTag<T> const & )
  { }

  // ######################################################################
  //! Prologue for all other types for XML output archives (except minimal types)
  /*! Starts a new node, named either automatically or by some NVP,
      that may be given data by the type about to be archived

      Minimal types do not start or end nodes */
  template <class T, traits::DisableIf<traits::has_minimal_base_class_serialization<T, traits::has_minimal_output_serialization, XMLOutputArchive>::value ||
                                       traits::has_minimal_output_serialization<T, XMLOutputArchive>::value> = traits::sfinae> inline
  void prologue( XMLOutputArchive & ar, T const & )
  {
    ar.startNode();
    ar.insertType<T>();
  }

  //! Prologue for all other types for XML input archives (except minimal types)
  template <class T, traits::DisableIf<traits::has_minimal_base_class_serialization<T, traits::has_minimal_input_serialization, XMLInputArchive>::value ||
                                       traits::has_minimal_input_serialization<T, XMLInputArchive>::value> = traits::sfinae> inline
  void prologue( XMLInputArchive & ar, T const & )
  {
    ar.startNode();
  }

  // ######################################################################
  //! Epilogue for all other types other for XML output archives (except minimal types)
  /*! Finishes the node created in the prologue

      Minimal types do not start or end nodes */
  template <class T, traits::DisableIf<traits::has_minimal_base_class_serialization<T, traits::has_minimal_output_serialization, XMLOutputArchive>::value ||
                                       traits::has_minimal_output_serialization<T, XMLOutputArchive>::value> = traits::sfinae> inline
  void epilogue( XMLOutputArchive & ar, T const & )
  {
    ar.finishNode();
  }

  //! Epilogue for all other types other for XML output archives (except minimal types)
  template <class T, traits::DisableIf<traits::has_minimal_base_class_serialization<T, traits::has_minimal_input_serialization, XMLInputArchive>::value ||
                                       traits::has_minimal_input_serialization<T, XMLInputArchive>::value> = traits::sfinae> inline
  void epilogue( XMLInputArchive & ar, T const & )
  {
    ar.finishNode();
  }

  // ######################################################################
  // Common XMLArchive serialization functions
  // ######################################################################

  //! Saving NVP types to XML
  template <class T> inline
  void CEREAL_SAVE_FUNCTION_NAME( XMLOutputArchive & ar, NameValuePair<T> const & t )
  {
    ar.setNextName( t.name );
    ar( t.value );
  }

  //! Loading NVP types from XML
  template <class T> inline
  void CEREAL_LOAD_FUNCTION_NAME( XMLInputArchive & ar, NameValuePair<T> & t )
  {
    ar.setNextName( t.name );
    ar( t.value );
  }

  // ######################################################################
  //! Saving SizeTags to XML
  template <class T> inline
  void CEREAL_SAVE_FUNCTION_NAME( XMLOutputArchive &, SizeTag<T> const & )
  { }

  //! Loading SizeTags from XML
  template <class T> inline
  void CEREAL_LOAD_FUNCTION_NAME( XMLInputArchive & ar, SizeTag<T> & st )
  {
    ar.loadSize( st.size );
  }

  // ######################################################################
  //! Saving for POD types to xml
  template <class T, traits::EnableIf<std::is_arithmetic<T>::value> = traits::sfinae> inline
  void CEREAL_SAVE_FUNCTION_NAME(XMLOutputArchive & ar, T const & t)
  {
    ar.saveValue( t );
  }

  //! Loading for POD types from xml
  template <class T, traits::EnableIf<std::is_arithmetic<T>::value> = traits::sfinae> inline
  void CEREAL_LOAD_FUNCTION_NAME(XMLInputArchive & ar, T & t)
  {
    ar.loadValue( t );
  }

  // ######################################################################
  //! saving string to xml
  template<class CharT, class Traits, class Alloc> inline
  void CEREAL_SAVE_FUNCTION_NAME(XMLOutputArchive & ar, std::basic_string<CharT, Traits, Alloc> const & str)
  {
    ar.saveValue( str );
  }

  //! loading string from xml
  template<class CharT, class Traits, class Alloc> inline
  void CEREAL_LOAD_FUNCTION_NAME(XMLInputArchive & ar, std::basic_string<CharT, Traits, Alloc> & str)
  {
    ar.loadValue( str );
  }
} // namespace cereal

// register archives for polymorphic support
CEREAL_REGISTER_ARCHIVE(cereal::XMLOutputArchive)
CEREAL_REGISTER_ARCHIVE(cereal::XMLInputArchive)

// tie input and output archives together
CEREAL_SETUP_ARCHIVE_TRAITS(cereal::XMLInputArchive, cereal::XMLOutputArchive)

#endif // CEREAL_ARCHIVES_XML_HPP_
```
