## Supplemental code for "The motif composition of variable-number tandem repeats impacts gene expression": cereal.hpp

```
/*! \file cereal.hpp
    \brief Main cereal functionality */
/*
  Copyright (c) 2014, Randolph Voorhies, Shane Grant
  All rights reserved.

  Redistribution and use in source and binary forms, with or without
  modification, are permitted provided that the following conditions are met:
      * Redistributions of source code must retain the above copyright
        notice, this list of conditions and the following disclaimer.
      * Redistributions in binary form must reproduce the above copyright
        notice, this list of conditions and the following disclaimer in the
        documentation and/or other materials provided with the distribution.
      * Neither the name of cereal nor the
        names of its contributors may be used to endorse or promote products
        derived from this software without specific prior written permission.

  THIS SOFTWARE IS PROVIDED BY THE COPYRIGHT HOLDERS AND CONTRIBUTORS "AS IS" AND
  ANY EXPRESS OR IMPLIED WARRANTIES, INCLUDING, BUT NOT LIMITED TO, THE IMPLIED
  WARRANTIES OF MERCHANTABILITY AND FITNESS FOR A PARTICULAR PURPOSE ARE
  DISCLAIMED. IN NO EVENT SHALL RANDOLPH VOORHIES OR SHANE GRANT BE LIABLE FOR ANY
  DIRECT, INDIRECT, INCIDENTAL, SPECIAL, EXEMPLARY, OR CONSEQUENTIAL DAMAGES
  (INCLUDING, BUT NOT LIMITED TO, PROCUREMENT OF SUBSTITUTE GOODS OR SERVICES;
  LOSS OF USE, DATA, OR PROFITS; OR BUSINESS INTERRUPTION) HOWEVER CAUSED AND
  ON ANY THEORY OF LIABILITY, WHETHER IN CONTRACT, STRICT LIABILITY, OR TORT
  (INCLUDING NEGLIGENCE OR OTHERWISE) ARISING IN ANY WAY OUT OF THE USE OF THIS
  SOFTWARE, EVEN IF ADVISED OF THE POSSIBILITY OF SUCH DAMAGE.
*/
#ifndef CEREAL_CEREAL_HPP_
#define CEREAL_CEREAL_HPP_

#include <type_traits>
#include <string>
#include <memory>
#include <functional>
#include <unordered_map>
#include <unordered_set>
#include <vector>
#include <cstddef>
#include <cstdint>
#include <functional>

#include "cereal/macros.hpp"
#include "cereal/details/traits.hpp"
#include "cereal/details/helpers.hpp"
#include "cereal/types/base_class.hpp"

namespace cereal
{
  // ######################################################################
  //! Creates a name value pair
  /*! @relates NameValuePair
      @ingroup Utility */
  template <class T> inline
  NameValuePair<T> make_nvp( std::string const & name, T && value )
  {
    return {name.c_str(), std::forward<T>(value)};
  }

  //! Creates a name value pair
  /*! @relates NameValuePair
      @ingroup Utility */
  template <class T> inline
  NameValuePair<T> make_nvp( const char * name, T && value )
  {
    return {name, std::forward<T>(value)};
  }

  //! Creates a name value pair for the variable T with the same name as the variable
  /*! @relates NameValuePair
      @ingroup Utility */
  #define CEREAL_NVP(T) ::cereal::make_nvp(#T, T)

  // ######################################################################
  //! Convenience function to create binary data for both const and non const pointers
  /*! @param data Pointer to beginning of the data
      @param size The size in bytes of the data
      @relates BinaryData
      @ingroup Utility */
  template <class T> inline
  BinaryData<T> binary_data( T && data, size_t size )
  {
    return {std::forward<T>(data), size};
  }

  // ######################################################################
  //! Creates a size tag from some variable.
  /*! Will normally be used to serialize size (e.g. size()) information for
      variable size containers.  If you have a variable sized container,
      the very first thing it serializes should be its size, wrapped in
      a SizeTag.

      @relates SizeTag
      @ingroup Utility */
  template <class T> inline
  SizeTag<T> make_size_tag( T && sz )
  {
    return {std::forward<T>(sz)};
  }

  // ######################################################################
  //! Marks data for deferred serialization
  /*! cereal performs a recursive depth-first traversal of data it serializes. When
      serializing smart pointers to large, nested, or cyclical data structures, it
      is possible to encounter a stack overflow from excessive recursion when following
      a chain of pointers.

      Deferment can help in these situations if the data can be serialized separately from
      the pointers used to traverse the structure. For example, a graph structure can have its
      nodes serialized before its edges:

      @code{.cpp}
      struct MyEdge
      {
        std::shared_ptr<MyNode> connection;
        int some_value;

        template<class Archive>
        void serialize(Archive & archive)
        {
          // when we serialize an edge, we'll defer serializing the associated node
          archive( cereal::defer( connection ),
                   some_value );
        }
      };

      struct MyGraphStructure
      {
        std::vector<MyEdge> edges;
        std::vector<MyNodes> nodes;

        template<class Archive>
        void serialize(Archive & archive)
        {
          // because of the deferment, we ensure all nodes are fully serialized
          // before any connection pointers to those nodes are serialized
          archive( edges, nodes );

          // we have to explicitly inform the archive when it is safe to serialize
          // the deferred data
          archive.serializeDeferments();
        }
      };
      @endcode

      @relates DeferredData
      @ingroup Utility */
  template <class T> inline
  DeferredData<T> defer( T && value )
  {
    return {std::forward<T>(value)};
  }

  // ######################################################################
  //! Called before a type is serialized to set up any special archive state
  //! for processing some type
  /*! If designing a serializer that needs to set up any kind of special
      state or output extra information for a type, specialize this function
      for the archive type and the types that require the extra information.
      @ingroup Internal */
  template <class Archive, class T> inline
  void prologue( Archive & /* archive */, T const & /* data */)
  { }

  //! Called after a type is serialized to tear down any special archive state
  //! for processing some type
  /*! @ingroup Internal */
  template <class Archive, class T> inline
  void epilogue( Archive & /* archive */, T const & /* data */)
  { }

  // ######################################################################
  //! Special flags for archives
  /*! AllowEmptyClassElision
        This allows for empty classes to be serialized even if they do not provide
        a serialization function.  Classes with no data members are considered to be
        empty.  Be warned that if this is enabled and you attempt to serialize an
        empty class with improperly formed serialize or load/save functions, no
        static error will occur - the error will propogate silently and your
        intended serialization functions may not be called.  You can manually
        ensure that your classes that have custom serialization are correct
        by using the traits is_output_serializable and is_input_serializable
        in cereal/details/traits.hpp.
      @ingroup Internal */
  enum Flags { AllowEmptyClassElision = 1 };

  // ######################################################################
  //! Registers a specific Archive type with cereal
  /*! This registration should be done once per archive.  A good place to
      put this is immediately following the definition of your archive.
      Archive registration is only strictly necessary if you wish to
      support pointers to polymorphic data types.  All archives that
      come with cereal are already registered.
      @ingroup Internal */
  #define CEREAL_REGISTER_ARCHIVE(Archive)                              \
  namespace cereal { namespace detail {                                 \
  template <class T, class BindingTag>                                  \
  typename polymorphic_serialization_support<Archive, T>::type          \
  instantiate_polymorphic_binding( T*, Archive*, BindingTag, adl_tag ); \
  } } /* end namespaces */

  //! Helper macro to omit unused warning
  #if defined(__GNUC__)
    // GCC / clang don't want the function
    #define CEREAL_UNUSED_FUNCTION
  #else
    #define CEREAL_UNUSED_FUNCTION static void unused() { (void)version; }
  #endif

  // ######################################################################
  //! Defines a class version for some type
  /*! Versioning information is optional and adds some small amount of
      overhead to serialization.  This overhead will occur both in terms of
      space in the archive (the version information for each class will be
      stored exactly once) as well as runtime (versioned serialization functions
      must check to see if they need to load or store version information).

      Versioning is useful if you plan on fundamentally changing the way some
      type is serialized in the future.  Versioned serialization functions
      cannot be used to load non-versioned data.

      By default, all types have an assumed version value of zero.  By
      using this macro, you may change the version number associated with
      some type.  cereal will then use this value as a second parameter
      to your serialization functions.

      The interface for the serialization functions is nearly identical
      to non-versioned serialization with the addition of a second parameter,
      const std::uint32_t version, which will be supplied with the correct
      version number.  Serializing the version number on a save happens
      automatically.

      Versioning cannot be mixed with non-versioned serialization functions.
      Having both types will result result in a compile time error.  Data
      serialized without versioning cannot be loaded by a serialization
      function with added versioning support.

      Example interface for versioning on a non-member serialize function:

      @code{cpp}
      CEREAL_CLASS_VERSION( Mytype, 77 ); // register class version

      template <class Archive>
      void serialize( Archive & ar, Mytype & t, const std::uint32_t version )
      {
        // When performing a load, the version associated with the class
        // is whatever it was when that data was originally serialized
        //
        // When we save, we'll use the version that is defined in the macro

        if( version >= some_number )
          // do this
        else
          // do that
      }
      @endcode

      Interfaces for other forms of serialization functions is similar.  This
      macro should be placed at global scope.
      @ingroup Utility */

  //! On C++17, define the StaticObject as inline to merge the definitions across TUs
  //! This prevents multiple definition errors when this macro appears in a header file
  //! included in multiple TUs.
  #ifdef CEREAL_HAS_CPP17
  #define CEREAL_CLASS_VERSION(TYPE, VERSION_NUMBER)                             \
  namespace cereal { namespace detail {                                          \
    template <> struct Version<TYPE>                                             \
    {                                                                            \
      static std::uint32_t registerVersion()                                     \
      {                                                                          \
        ::cereal::detail::StaticObject<Versions>::getInstance().mapping.emplace( \
             std::type_index(typeid(TYPE)).hash_code(), VERSION_NUMBER );        \
        return VERSION_NUMBER;                                                   \
      }                                                                          \
      static inline const std::uint32_t version = registerVersion();             \
      CEREAL_UNUSED_FUNCTION                                                     \
    }; /* end Version */                                                         \
  } } // end namespaces
  #else
  #define CEREAL_CLASS_VERSION(TYPE, VERSION_NUMBER)                             \
  namespace cereal { namespace detail {                                          \
    template <> struct Version<TYPE>                                             \
    {                                                                            \
      static const std::uint32_t version;                                        \
      static std::uint32_t registerVersion()                                     \
      {                                                                          \
        ::cereal::detail::StaticObject<Versions>::getInstance().mapping.emplace( \
             std::type_index(typeid(TYPE)).hash_code(), VERSION_NUMBER );        \
        return VERSION_NUMBER;                                                   \
      }                                                                          \
      CEREAL_UNUSED_FUNCTION                                                     \
    }; /* end Version */                                                         \
    const std::uint32_t Version<TYPE>::version =                                 \
      Version<TYPE>::registerVersion();                                          \
  } } // end namespaces

  #endif

  // ######################################################################
  //! The base output archive class
  /*! This is the base output archive for all output archives.  If you create
      a custom archive class, it should derive from this, passing itself as
      a template parameter for the ArchiveType.

      The base class provides all of the functionality necessary to
      properly forward data to the correct serialization functions.

      Individual archives should use a combination of prologue and
      epilogue functions together with specializations of serialize, save,
      and load to alter the functionality of their serialization.

      @tparam ArchiveType The archive type that derives from OutputArchive
      @tparam Flags Flags to control advanced functionality.  See the Flags
                    enum for more information.
      @ingroup Internal */
  template<class ArchiveType, std::uint32_t Flags = 0>
  class OutputArchive : public detail::OutputArchiveBase
  {
    public:
      //! Construct the output archive
      /*! @param derived A pointer to the derived ArchiveType (pass this from the derived archive) */
      OutputArchive(ArchiveType * const derived) : self(derived), itsCurrentPointerId(1), itsCurrentPolymorphicTypeId(1)
      { }

      OutputArchive & operator=( OutputArchive const & ) = delete;

      //! Serializes all passed in data
      /*! This is the primary interface for serializing data with an archive */
      template <class ... Types> inline
      ArchiveType & operator()( Types && ... args )
      {
        self->process( std::forward<Types>( args )... );
        return *self;
      }

      //! Serializes any data marked for deferment using defer
      /*! This will cause any data wrapped in DeferredData to be immediately serialized */
      void serializeDeferments()
      {
        for( auto & deferment : itsDeferments )
          deferment();
      }

      /*! @name Boost Transition Layer
          Functionality that mirrors the syntax for Boost.  This is useful if you are transitioning
          a large project from Boost to cereal.  The preferred interface for cereal is using operator(). */
      //! @{

      //! Indicates this archive is not intended for loading
      /*! This ensures compatibility with boost archive types.  If you are transitioning
          from boost, you can check this value within a member or external serialize function
          (i.e., Archive::is_loading::value) to disable behavior specific to loading, until
          you can transition to split save/load or save_minimal/load_minimal functions */
      using is_loading = std::false_type;

      //! Indicates this archive is intended for saving
      /*! This ensures compatibility with boost archive types.  If you are transitioning
          from boost, you can check this value within a member or external serialize function
          (i.e., Archive::is_saving::value) to enable behavior specific to loading, until
          you can transition to split save/load or save_minimal/load_minimal functions */
      using is_saving = std::true_type;

      //! Serializes passed in data
      /*! This is a boost compatability layer and is not the preferred way of using
          cereal.  If you are transitioning from boost, use this until you can
          transition to the operator() overload */
      template <class T> inline
      ArchiveType & operator&( T && arg )
      {
        self->process( std::forward<T>( arg ) );
        return *self;
      }

      //! Serializes passed in data
      /*! This is a boost compatability layer and is not the preferred way of using
          cereal.  If you are transitioning from boost, use this until you can
          transition to the operator() overload */
      template <class T> inline
      ArchiveType & operator<<( T && arg )
      {
        self->process( std::forward<T>( arg ) );
        return *self;
      }

      //! @}

      //! Registers a shared pointer with the archive
      /*! This function is used to track shared pointer targets to prevent
          unnecessary saves from taking place if multiple shared pointers
          point to the same data.

          @internal
          @param sharedPointer The shared pointer itself (the adress is taked via get()).
                               The archive takes a copy to prevent the memory location to be freed
                               as long as the address is used as id. This is needed to prevent CVE-2020-11105.
          @return A key that uniquely identifies the pointer */
      inline std::uint32_t registerSharedPointer(const std::shared_ptr<const void>& sharedPointer)
      {
        void const * addr = sharedPointer.get();

        // Handle null pointers by just returning 0
        if(addr == 0) return 0;
        itsSharedPointerStorage.push_back(sharedPointer);

        auto id = itsSharedPointerMap.find( addr );
        if( id == itsSharedPointerMap.end() )
        {
          auto ptrId = itsCurrentPointerId++;
          itsSharedPointerMap.insert( {addr, ptrId} );
          return ptrId | detail::msb_32bit; // mask MSB to be 1
        }
        else
          return id->second;
      }

      //! Registers a polymorphic type name with the archive
      /*! This function is used to track polymorphic types to prevent
          unnecessary saves of identifying strings used by the polymorphic
          support functionality.

          @internal
          @param name The name to associate with a polymorphic type
          @return A key that uniquely identifies the polymorphic type name */
      inline std::uint32_t registerPolymorphicType( char const * name )
      {
        auto id = itsPolymorphicTypeMap.find( name );
        if( id == itsPolymorphicTypeMap.end() )
        {
          auto polyId = itsCurrentPolymorphicTypeId++;
          itsPolymorphicTypeMap.insert( {name, polyId} );
          return polyId | detail::msb_32bit; // mask MSB to be 1
        }
        else
          return id->second;
      }

    private:
      //! Serializes data after calling prologue, then calls epilogue
      template <class T> inline
      void process( T && head )
      {
        prologue( *self, head );
        self->processImpl( head );
        epilogue( *self, head );
      }

      //! Unwinds to process all data
      template <class T, class ... Other> inline
      void process( T && head, Other && ... tail )
      {
        self->process( std::forward<T>( head ) );
        self->process( std::forward<Other>( tail )... );
      }

      //! Serialization of a virtual_base_class wrapper
      /*! \sa virtual_base_class */
      template <class T> inline
      ArchiveType & processImpl(virtual_base_class<T> const & b)
      {
        traits::detail::base_class_id id(b.base_ptr);
        if(itsBaseClassSet.count(id) == 0)
        {
          itsBaseClassSet.insert(id);
          self->processImpl( *b.base_ptr );
        }
        return *self;
      }

      //! Serialization of a base_class wrapper
      /*! \sa base_class */
      template <class T> inline
      ArchiveType & processImpl(base_class<T> const & b)
      {
        self->processImpl( *b.base_ptr );
        return *self;
      }

      std::vector<std::function<void(void)>> itsDeferments;

      template <class T> inline
      ArchiveType & processImpl(DeferredData<T> const & d)
      {
        std::function<void(void)> deferment( [this, d](){ self->process( d.value ); } );
        itsDeferments.emplace_back( std::move(deferment) );

        return *self;
      }

      //! Helper macro that expands the requirements for activating an overload
      /*! Requirements:
            Has the requested serialization function
            Does not have version and unversioned at the same time
            Is output serializable AND
              is specialized for this type of function OR
              has no specialization at all */
      #define PROCESS_IF(name)                                                             \
      traits::EnableIf<traits::has_##name<T, ArchiveType>::value,                          \
                       !traits::has_invalid_output_versioning<T, ArchiveType>::value,      \
                       (traits::is_output_serializable<T, ArchiveType>::value &&           \
                        (traits::is_specialized_##name<T, ArchiveType>::value ||           \
                         !traits::is_specialized<T, ArchiveType>::value))> = traits::sfinae

      //! Member serialization
      template <class T, PROCESS_IF(member_serialize)> inline
      ArchiveType & processImpl(T const & t)
      {
        access::member_serialize(*self, const_cast<T &>(t));
        return *self;
      }

      //! Non member serialization
      template <class T, PROCESS_IF(non_member_serialize)> inline
      ArchiveType & processImpl(T const & t)
      {
        CEREAL_SERIALIZE_FUNCTION_NAME(*self, const_cast<T &>(t));
        return *self;
      }

      //! Member split (save)
      template <class T, PROCESS_IF(member_save)> inline
      ArchiveType & processImpl(T const & t)
      {
        access::member_save(*self, t);
        return *self;
      }

      //! Non member split (save)
      template <class T, PROCESS_IF(non_member_save)> inline
      ArchiveType & processImpl(T const & t)
      {
        CEREAL_SAVE_FUNCTION_NAME(*self, t);
        return *self;
      }

      //! Member split (save_minimal)
      template <class T, PROCESS_IF(member_save_minimal)> inline
      ArchiveType & processImpl(T const & t)
      {
        self->process( access::member_save_minimal(*self, t) );
        return *self;
      }

      //! Non member split (save_minimal)
      template <class T, PROCESS_IF(non_member_save_minimal)> inline
      ArchiveType & processImpl(T const & t)
      {
        self->process( CEREAL_SAVE_MINIMAL_FUNCTION_NAME(*self, t) );
        return *self;
      }

      //! Empty class specialization
      template <class T, traits::EnableIf<(Flags & AllowEmptyClassElision),
                                          !traits::is_output_serializable<T, ArchiveType>::value,
                                          std::is_empty<T>::value> = traits::sfinae> inline
      ArchiveType & processImpl(T const &)
      {
        return *self;
      }

      //! No matching serialization
      /*! Invalid if we have invalid output versioning or
          we are not output serializable, and either
          don't allow empty class ellision or allow it but are not serializing an empty class */
      template <class T, traits::EnableIf<traits::has_invalid_output_versioning<T, ArchiveType>::value ||
                                          (!traits::is_output_serializable<T, ArchiveType>::value &&
                                           (!(Flags & AllowEmptyClassElision) || ((Flags & AllowEmptyClassElision) && !std::is_empty<T>::value)))> = traits::sfinae> inline
      ArchiveType & processImpl(T const &)
      {
        static_assert(traits::detail::count_output_serializers<T, ArchiveType>::value != 0,
            "cereal could not find any output serialization functions for the provided type and archive combination. \n\n "
            "Types must either have a serialize function, load/save pair, or load_minimal/save_minimal pair (you may not mix these). \n "
            "Serialize functions generally have the following signature: \n\n "
            "template<class Archive> \n "
            "  void serialize(Archive & ar) \n "
            "  { \n "
            "    ar( member1, member2, member3 ); \n "
            "  } \n\n " );

        static_assert(traits::detail::count_output_serializers<T, ArchiveType>::value < 2,
            "cereal found more than one compatible output serialization function for the provided type and archive combination. \n\n "
            "Types must either have a serialize function, load/save pair, or load_minimal/save_minimal pair (you may not mix these). \n "
            "Use specialization (see access.hpp) if you need to disambiguate between serialize vs load/save functions.  \n "
            "Note that serialization functions can be inherited which may lead to the aforementioned ambiguities. \n "
            "In addition, you may not mix versioned with non-versioned serialization functions. \n\n ");

        return *self;
      }

      //! Registers a class version with the archive and serializes it if necessary
      /*! If this is the first time this class has been serialized, we will record its
          version number and serialize that.

          @tparam T The type of the class being serialized */
      template <class T> inline
      std::uint32_t registerClassVersion()
      {
        static const auto hash = std::type_index(typeid(T)).hash_code();
        const auto insertResult = itsVersionedTypes.insert( hash );
        const auto lock = detail::StaticObject<detail::Versions>::lock();
        const auto version =
          detail::StaticObject<detail::Versions>::getInstance().find( hash, detail::Version<T>::version );

        if( insertResult.second ) // insertion took place, serialize the version number
          process( make_nvp<ArchiveType>("cereal_class_version", version) );

        return version;
      }

      //! Member serialization
      /*! Versioning implementation */
      template <class T, PROCESS_IF(member_versioned_serialize)> inline
      ArchiveType & processImpl(T const & t)
      {
        access::member_serialize(*self, const_cast<T &>(t), registerClassVersion<T>());
        return *self;
      }

      //! Non member serialization
      /*! Versioning implementation */
      template <class T, PROCESS_IF(non_member_versioned_serialize)> inline
      ArchiveType & processImpl(T const & t)
      {
        CEREAL_SERIALIZE_FUNCTION_NAME(*self, const_cast<T &>(t), registerClassVersion<T>());
        return *self;
      }

      //! Member split (save)
      /*! Versioning implementation */
      template <class T, PROCESS_IF(member_versioned_save)> inline
      ArchiveType & processImpl(T const & t)
      {
        access::member_save(*self, t, registerClassVersion<T>());
        return *self;
      }

      //! Non member split (save)
      /*! Versioning implementation */
      template <class T, PROCESS_IF(non_member_versioned_save)> inline
      ArchiveType & processImpl(T const & t)
      {
        CEREAL_SAVE_FUNCTION_NAME(*self, t, registerClassVersion<T>());
        return *self;
      }

      //! Member split (save_minimal)
      /*! Versioning implementation */
      template <class T, PROCESS_IF(member_versioned_save_minimal)> inline
      ArchiveType & processImpl(T const & t)
      {
        self->process( access::member_save_minimal(*self, t, registerClassVersion<T>()) );
        return *self;
      }

      //! Non member split (save_minimal)
      /*! Versioning implementation */
      template <class T, PROCESS_IF(non_member_versioned_save_minimal)> inline
      ArchiveType & processImpl(T const & t)
      {
        self->process( CEREAL_SAVE_MINIMAL_FUNCTION_NAME(*self, t, registerClassVersion<T>()) );
        return *self;
      }

    #undef PROCESS_IF

    private:
      ArchiveType * const self;

      //! A set of all base classes that have been serialized
      std::unordered_set<traits::detail::base_class_id, traits::detail::base_class_id_hash> itsBaseClassSet;

      //! Maps from addresses to pointer ids
      std::unordered_map<void const *, std::uint32_t> itsSharedPointerMap;

      //! Copy of shared pointers used in #itsSharedPointerMap to make sure they are kept alive
      //  during lifetime of itsSharedPointerMap to prevent CVE-2020-11105.
      std::vector<std::shared_ptr<const void>> itsSharedPointerStorage;

      //! The id to be given to the next pointer
      std::uint32_t itsCurrentPointerId;

      //! Maps from polymorphic type name strings to ids
      std::unordered_map<char const *, std::uint32_t> itsPolymorphicTypeMap;

      //! The id to be given to the next polymorphic type name
      std::uint32_t itsCurrentPolymorphicTypeId;

      //! Keeps track of classes that have versioning information associated with them
      std::unordered_set<size_type> itsVersionedTypes;
  }; // class OutputArchive

  // ######################################################################
  //! The base input archive class
  /*! This is the base input archive for all input archives.  If you create
      a custom archive class, it should derive from this, passing itself as
      a template parameter for the ArchiveType.

      The base class provides all of the functionality necessary to
      properly forward data to the correct serialization functions.

      Individual archives should use a combination of prologue and
      epilogue functions together with specializations of serialize, save,
      and load to alter the functionality of their serialization.

      @tparam ArchiveType The archive type that derives from InputArchive
      @tparam Flags Flags to control advanced functionality.  See the Flags
                    enum for more information.
      @ingroup Internal */
  template<class ArchiveType, std::uint32_t Flags = 0>
  class InputArchive : public detail::InputArchiveBase
  {
    public:
      //! Construct the output archive
      /*! @param derived A pointer to the derived ArchiveType (pass this from the derived archive) */
      InputArchive(ArchiveType * const derived) :
        self(derived),
        itsBaseClassSet(),
        itsSharedPointerMap(),
        itsPolymorphicTypeMap(),
        itsVersionedTypes()
      { }

      InputArchive & operator=( InputArchive const & ) = delete;

      //! Serializes all passed in data
      /*! This is the primary interface for serializing data with an archive */
      template <class ... Types> inline
      ArchiveType & operator()( Types && ... args )
      {
        process( std::forward<Types>( args )... );
        return *self;
      }

      //! Serializes any data marked for deferment using defer
      /*! This will cause any data wrapped in DeferredData to be immediately serialized */
      void serializeDeferments()
      {
        for( auto & deferment : itsDeferments )
          deferment();
      }

      /*! @name Boost Transition Layer
          Functionality that mirrors the syntax for Boost.  This is useful if you are transitioning
          a large project from Boost to cereal.  The preferred interface for cereal is using operator(). */
      //! @{

      //! Indicates this archive is intended for loading
      /*! This ensures compatibility with boost archive types.  If you are transitioning
          from boost, you can check this value within a member or external serialize function
          (i.e., Archive::is_loading::value) to enable behavior specific to loading, until
          you can transition to split save/load or save_minimal/load_minimal functions */
      using is_loading = std::true_type;

      //! Indicates this archive is not intended for saving
      /*! This ensures compatibility with boost archive types.  If you are transitioning
          from boost, you can check this value within a member or external serialize function
          (i.e., Archive::is_saving::value) to disable behavior specific to loading, until
          you can transition to split save/load or save_minimal/load_minimal functions */
      using is_saving = std::false_type;

      //! Serializes passed in data
      /*! This is a boost compatability layer and is not the preferred way of using
          cereal.  If you are transitioning from boost, use this until you can
          transition to the operator() overload */
      template <class T> inline
      ArchiveType & operator&( T && arg )
      {
        self->process( std::forward<T>( arg ) );
        return *self;
      }

      //! Serializes passed in data
      /*! This is a boost compatability layer and is not the preferred way of using
          cereal.  If you are transitioning from boost, use this until you can
          transition to the operator() overload */
      template <class T> inline
      ArchiveType & operator>>( T && arg )
      {
        self->process( std::forward<T>( arg ) );
        return *self;
      }

      //! @}

      //! Retrieves a shared pointer given a unique key for it
      /*! This is used to retrieve a previously registered shared_ptr
          which has already been loaded.

          @internal
          @param id The unique id that was serialized for the pointer
          @return A shared pointer to the data
          @throw Exception if the id does not exist */
      inline std::shared_ptr<void> getSharedPointer(std::uint32_t const id)
      {
        if(id == 0) return std::shared_ptr<void>(nullptr);

        auto iter = itsSharedPointerMap.find( id );
        if(iter == itsSharedPointerMap.end())
          throw Exception("Error while trying to deserialize a smart pointer. Could not find id " + std::to_string(id));

        return iter->second;
      }

      //! Registers a shared pointer to its unique identifier
      /*! After a shared pointer has been allocated for the first time, it should
          be registered with its loaded id for future references to it.

          @internal
          @param id The unique identifier for the shared pointer
          @param ptr The actual shared pointer */
      inline void registerSharedPointer(std::uint32_t const id, std::shared_ptr<void> ptr)
      {
        std::uint32_t const stripped_id = id & ~detail::msb_32bit;
        itsSharedPointerMap[stripped_id] = ptr;
      }

      //! Retrieves the string for a polymorphic type given a unique key for it
      /*! This is used to retrieve a string previously registered during
          a polymorphic load.

          @internal
          @param id The unique id that was serialized for the polymorphic type
          @return The string identifier for the tyep */
      inline std::string getPolymorphicName(std::uint32_t const id)
      {
        auto name = itsPolymorphicTypeMap.find( id );
        if(name == itsPolymorphicTypeMap.end())
        {
          throw Exception("Error while trying to deserialize a polymorphic pointer. Could not find type id " + std::to_string(id));
        }
        return name->second;
      }

      //! Registers a polymorphic name string to its unique identifier
      /*! After a polymorphic type has been loaded for the first time, it should
          be registered with its loaded id for future references to it.

          @internal
          @param id The unique identifier for the polymorphic type
          @param name The name associated with the tyep */
      inline void registerPolymorphicName(std::uint32_t const id, std::string const & name)
      {
        std::uint32_t const stripped_id = id & ~detail::msb_32bit;
        itsPolymorphicTypeMap.insert( {stripped_id, name} );
      }

    private:
      //! Serializes data after calling prologue, then calls epilogue
      template <class T> inline
      void process( T && head )
      {
        prologue( *self, head );
        self->processImpl( head );
        epilogue( *self, head );
      }

      //! Unwinds to process all data
      template <class T, class ... Other> inline
      void process( T && head, Other && ... tail )
      {
        process( std::forward<T>( head ) );
        process( std::forward<Other>( tail )... );
      }

      //! Serialization of a virtual_base_class wrapper
      /*! \sa virtual_base_class */
      template <class T> inline
      ArchiveType & processImpl(virtual_base_class<T> & b)
      {
        traits::detail::base_class_id id(b.base_ptr);
        if(itsBaseClassSet.count(id) == 0)
        {
          itsBaseClassSet.insert(id);
          self->processImpl( *b.base_ptr );
        }
        return *self;
      }

      //! Serialization of a base_class wrapper
      /*! \sa base_class */
      template <class T> inline
      ArchiveType & processImpl(base_class<T> & b)
      {
        self->processImpl( *b.base_ptr );
        return *self;
      }

      std::vector<std::function<void(void)>> itsDeferments;

      template <class T> inline
      ArchiveType & processImpl(DeferredData<T> const & d)
      {
        std::function<void(void)> deferment( [this, d](){ self->process( d.value ); } );
        itsDeferments.emplace_back( std::move(deferment) );

        return *self;
      }

      //! Helper macro that expands the requirements for activating an overload
      /*! Requirements:
            Has the requested serialization function
            Does not have version and unversioned at the same time
            Is input serializable AND
              is specialized for this type of function OR
              has no specialization at all */
      #define PROCESS_IF(name)                                                              \
      traits::EnableIf<traits::has_##name<T, ArchiveType>::value,                           \
                       !traits::has_invalid_input_versioning<T, ArchiveType>::value,        \
                       (traits::is_input_serializable<T, ArchiveType>::value &&             \
                        (traits::is_specialized_##name<T, ArchiveType>::value ||            \
                         !traits::is_specialized<T, ArchiveType>::value))> = traits::sfinae

      //! Member serialization
      template <class T, PROCESS_IF(member_serialize)> inline
      ArchiveType & processImpl(T & t)
      {
        access::member_serialize(*self, t);
        return *self;
      }

      //! Non member serialization
      template <class T, PROCESS_IF(non_member_serialize)> inline
      ArchiveType & processImpl(T & t)
      {
        CEREAL_SERIALIZE_FUNCTION_NAME(*self, t);
        return *self;
      }

      //! Member split (load)
      template <class T, PROCESS_IF(member_load)> inline
      ArchiveType & processImpl(T & t)
      {
        access::member_load(*self, t);
        return *self;
      }

      //! Non member split (load)
      template <class T, PROCESS_IF(non_member_load)> inline
      ArchiveType & processImpl(T & t)
      {
        CEREAL_LOAD_FUNCTION_NAME(*self, t);
        return *self;
      }

      //! Member split (load_minimal)
      template <class T, PROCESS_IF(member_load_minimal)> inline
      ArchiveType & processImpl(T & t)
      {
        using OutArchiveType = typename traits::detail::get_output_from_input<ArchiveType>::type;
        typename traits::has_member_save_minimal<T, OutArchiveType>::type value;
        self->process( value );
        access::member_load_minimal(*self, t, value);
        return *self;
      }

      //! Non member split (load_minimal)
      template <class T, PROCESS_IF(non_member_load_minimal)> inline
      ArchiveType & processImpl(T & t)
      {
        using OutArchiveType = typename traits::detail::get_output_from_input<ArchiveType>::type;
        typename traits::has_non_member_save_minimal<T, OutArchiveType>::type value;
        self->process( value );
        CEREAL_LOAD_MINIMAL_FUNCTION_NAME(*self, t, value);
        return *self;
      }

      //! Empty class specialization
      template <class T, traits::EnableIf<(Flags & AllowEmptyClassElision),
                                          !traits::is_input_serializable<T, ArchiveType>::value,
                                          std::is_empty<T>::value> = traits::sfinae> inline
      ArchiveType & processImpl(T const &)
      {
        return *self;
      }

      //! No matching serialization
      /*! Invalid if we have invalid input versioning or
          we are not input serializable, and either
          don't allow empty class ellision or allow it but are not serializing an empty class */
      template <class T, traits::EnableIf<traits::has_invalid_input_versioning<T, ArchiveType>::value ||
                                          (!traits::is_input_serializable<T, ArchiveType>::value &&
                                           (!(Flags & AllowEmptyClassElision) || ((Flags & AllowEmptyClassElision) && !std::is_empty<T>::value)))> = traits::sfinae> inline
      ArchiveType & processImpl(T const &)
      {
        static_assert(traits::detail::count_input_serializers<T, ArchiveType>::value != 0,
            "cereal could not find any input serialization functions for the provided type and archive combination. \n\n "
            "Types must either have a serialize function, load/save pair, or load_minimal/save_minimal pair (you may not mix these). \n "
            "Serialize functions generally have the following signature: \n\n "
            "template<class Archive> \n "
            "  void serialize(Archive & ar) \n "
            "  { \n "
            "    ar( member1, member2, member3 ); \n "
            "  } \n\n " );

        static_assert(traits::detail::count_input_serializers<T, ArchiveType>::value < 2,
            "cereal found more than one compatible input serialization function for the provided type and archive combination. \n\n "
            "Types must either have a serialize function, load/save pair, or load_minimal/save_minimal pair (you may not mix these). \n "
            "Use specialization (see access.hpp) if you need to disambiguate between serialize vs load/save functions.  \n "
            "Note that serialization functions can be inherited which may lead to the aforementioned ambiguities. \n "
            "In addition, you may not mix versioned with non-versioned serialization functions. \n\n ");

        return *self;
      }

      //! Befriend for versioning in load_and_construct
      template <class A, class B, bool C, bool D, bool E, bool F> friend struct detail::Construct;

      //! Registers a class version with the archive and serializes it if necessary
      /*! If this is the first time this class has been serialized, we will record its
          version number and serialize that.

          @tparam T The type of the class being serialized */
      template <class T> inline
      std::uint32_t loadClassVersion()
      {
        static const auto hash = std::type_index(typeid(T)).hash_code();
        auto lookupResult = itsVersionedTypes.find( hash );

        if( lookupResult != itsVersionedTypes.end() ) // already exists
          return lookupResult->second;
        else // need to load
        {
          std::uint32_t version;

          process( make_nvp<ArchiveType>("cereal_class_version", version) );
          itsVersionedTypes.emplace_hint( lookupResult, hash, version );

          return version;
        }
      }

      //! Member serialization
      /*! Versioning implementation */
      template <class T, PROCESS_IF(member_versioned_serialize)> inline
      ArchiveType & processImpl(T & t)
      {
        const auto version = loadClassVersion<T>();
        access::member_serialize(*self, t, version);
        return *self;
      }

      //! Non member serialization
      /*! Versioning implementation */
      template <class T, PROCESS_IF(non_member_versioned_serialize)> inline
      ArchiveType & processImpl(T & t)
      {
        const auto version = loadClassVersion<T>();
        CEREAL_SERIALIZE_FUNCTION_NAME(*self, t, version);
        return *self;
      }

      //! Member split (load)
      /*! Versioning implementation */
      template <class T, PROCESS_IF(member_versioned_load)> inline
      ArchiveType & processImpl(T & t)
      {
        const auto version = loadClassVersion<T>();
        access::member_load(*self, t, version);
        return *self;
      }

      //! Non member split (load)
      /*! Versioning implementation */
      template <class T, PROCESS_IF(non_member_versioned_load)> inline
      ArchiveType & processImpl(T & t)
      {
        const auto version = loadClassVersion<T>();
        CEREAL_LOAD_FUNCTION_NAME(*self, t, version);
        return *self;
      }

      //! Member split (load_minimal)
      /*! Versioning implementation */
      template <class T, PROCESS_IF(member_versioned_load_minimal)> inline
      ArchiveType & processImpl(T & t)
      {
        using OutArchiveType = typename traits::detail::get_output_from_input<ArchiveType>::type;
        const auto version = loadClassVersion<T>();
        typename traits::has_member_versioned_save_minimal<T, OutArchiveType>::type value;
        self->process(value);
        access::member_load_minimal(*self, t, value, version);
        return *self;
      }

      //! Non member split (load_minimal)
      /*! Versioning implementation */
      template <class T, PROCESS_IF(non_member_versioned_load_minimal)> inline
      ArchiveType & processImpl(T & t)
      {
        using OutArchiveType = typename traits::detail::get_output_from_input<ArchiveType>::type;
        const auto version = loadClassVersion<T>();
        typename traits::has_non_member_versioned_save_minimal<T, OutArchiveType>::type value;
        self->process(value);
        CEREAL_LOAD_MINIMAL_FUNCTION_NAME(*self, t, value, version);
        return *self;
      }

      #undef PROCESS_IF

    private:
      ArchiveType * const self;

      //! A set of all base classes that have been serialized
      std::unordered_set<traits::detail::base_class_id, traits::detail::base_class_id_hash> itsBaseClassSet;

      //! Maps from pointer ids to metadata
      std::unordered_map<std::uint32_t, std::shared_ptr<void>> itsSharedPointerMap;

      //! Maps from name ids to names
      std::unordered_map<std::uint32_t, std::string> itsPolymorphicTypeMap;

      //! Maps from type hash codes to version numbers
      std::unordered_map<std::size_t, std::uint32_t> itsVersionedTypes;
  }; // class InputArchive
} // namespace cereal

// This include needs to come after things such as binary_data, make_nvp, etc
#include "cereal/types/common.hpp"

#endif // CEREAL_CEREAL_HPP_
```
