## Supplemental code for "The motif composition of variable-number tandem repeats impacts gene expression": helpers.hpp

```
/*! \file helpers.hpp
    \brief Internal helper functionality
    \ingroup Internal */
/*
  Copyright (c) 2014, Randolph Voorhies, Shane Grant
  All rights reserved.

  Redistribution and use in source and binary forms, with or without
  modification, are permitted provided that the following conditions are met:
      * Redistributions of source code must retain the above copyright
        notice, this list of conditions and the following disclaimer.
      * Redistributions in binary form must reproduce the above copyright
        notice, this list of conditions and the following disclaimer in the
        documentation and/or other materials provided with the distribution.
      * Neither the name of cereal nor the
        names of its contributors may be used to endorse or promote products
        derived from this software without specific prior written permission.

  THIS SOFTWARE IS PROVIDED BY THE COPYRIGHT HOLDERS AND CONTRIBUTORS "AS IS" AND
  ANY EXPRESS OR IMPLIED WARRANTIES, INCLUDING, BUT NOT LIMITED TO, THE IMPLIED
  WARRANTIES OF MERCHANTABILITY AND FITNESS FOR A PARTICULAR PURPOSE ARE
  DISCLAIMED. IN NO EVENT SHALL RANDOLPH VOORHIES OR SHANE GRANT BE LIABLE FOR ANY
  DIRECT, INDIRECT, INCIDENTAL, SPECIAL, EXEMPLARY, OR CONSEQUENTIAL DAMAGES
  (INCLUDING, BUT NOT LIMITED TO, PROCUREMENT OF SUBSTITUTE GOODS OR SERVICES;
  LOSS OF USE, DATA, OR PROFITS; OR BUSINESS INTERRUPTION) HOWEVER CAUSED AND
  ON ANY THEORY OF LIABILITY, WHETHER IN CONTRACT, STRICT LIABILITY, OR TORT
  (INCLUDING NEGLIGENCE OR OTHERWISE) ARISING IN ANY WAY OUT OF THE USE OF THIS
  SOFTWARE, EVEN IF ADVISED OF THE POSSIBILITY OF SUCH DAMAGE.
*/
#ifndef CEREAL_DETAILS_HELPERS_HPP_
#define CEREAL_DETAILS_HELPERS_HPP_

#include <type_traits>
#include <cstdint>
#include <utility>
#include <memory>
#include <unordered_map>
#include <stdexcept>

#include "cereal/macros.hpp"
#include "cereal/details/static_object.hpp"

namespace cereal
{
  // ######################################################################
  //! An exception class thrown when things go wrong at runtime
  /*! @ingroup Utility */
  struct Exception : public std::runtime_error
  {
    explicit Exception( const std::string & what_ ) : std::runtime_error(what_) {}
    explicit Exception( const char * what_ ) : std::runtime_error(what_) {}
  };

  // ######################################################################
  //! The size type used by cereal
  /*! To ensure compatability between 32, 64, etc bit machines, we need to use
      a fixed size type instead of size_t, which may vary from machine to
      machine.

      The default value for CEREAL_SIZE_TYPE is specified in cereal/macros.hpp */
  using size_type = CEREAL_SIZE_TYPE;

  // forward decls
  class BinaryOutputArchive;
  class BinaryInputArchive;

  // ######################################################################
  namespace detail
  {
    struct NameValuePairCore {}; //!< Traits struct for NVPs
    struct DeferredDataCore {}; //!< Traits struct for DeferredData
  }

  // ######################################################################
  //! For holding name value pairs
  /*! This pairs a name (some string) with some value such that an archive
      can potentially take advantage of the pairing.

      In serialization functions, NameValuePairs are usually created like so:
      @code{.cpp}
      struct MyStruct
      {
        int a, b, c, d, e;

        template<class Archive>
        void serialize(Archive & archive)
        {
          archive( CEREAL_NVP(a),
                   CEREAL_NVP(b),
                   CEREAL_NVP(c),
                   CEREAL_NVP(d),
                   CEREAL_NVP(e) );
        }
      };
      @endcode

      Alternatively, you can give you data members custom names like so:
      @code{.cpp}
      struct MyStruct
      {
        int a, b, my_embarrassing_variable_name, d, e;

        template<class Archive>
        void serialize(Archive & archive)
        {
          archive( CEREAL_NVP(a),
                   CEREAL_NVP(b),
                   cereal::make_nvp("var", my_embarrassing_variable_name) );
                   CEREAL_NVP(d),
                   CEREAL_NVP(e) );
        }
      };
      @endcode

      There is a slight amount of overhead to creating NameValuePairs, so there
      is a third method which will elide the names when they are not used by
      the Archive:

      @code{.cpp}
      struct MyStruct
      {
        int a, b;

        template<class Archive>
        void serialize(Archive & archive)
        {
          archive( cereal::make_nvp<Archive>(a),
                   cereal::make_nvp<Archive>(b) );
        }
      };
      @endcode

      This third method is generally only used when providing generic type
      support.  Users writing their own serialize functions will normally
      explicitly control whether they want to use NVPs or not.

      @internal */
  template <class T>
  class NameValuePair : detail::NameValuePairCore
  {
    private:
      // If we get passed an array, keep the type as is, otherwise store
      // a reference if we were passed an l value reference, else copy the value
      using Type = typename std::conditional<std::is_array<typename std::remove_reference<T>::type>::value,
                                             typename std::remove_cv<T>::type,
                                             typename std::conditional<std::is_lvalue_reference<T>::value,
                                                                       T,
                                                                       typename std::decay<T>::type>::type>::type;

      // prevent nested nvps
      static_assert( !std::is_base_of<detail::NameValuePairCore, T>::value,
                     "Cannot pair a name to a NameValuePair" );

      NameValuePair & operator=( NameValuePair const & ) = delete;

    public:
      //! Constructs a new NameValuePair
      /*! @param n The name of the pair
          @param v The value to pair.  Ideally this should be an l-value reference so that
                   the value can be both loaded and saved to.  If you pass an r-value reference,
                   the NameValuePair will store a copy of it instead of a reference.  Thus you should
                   only pass r-values in cases where this makes sense, such as the result of some
                   size() call.
          @internal */
      NameValuePair( char const * n, T && v ) : name(n), value(std::forward<T>(v)) {}

      char const * name;
      Type value;
  };

  //! A specialization of make_nvp<> that simply forwards the value for binary archives
  /*! @relates NameValuePair
      @internal */
  template<class Archive, class T> inline
  typename
  std::enable_if<std::is_same<Archive, ::cereal::BinaryInputArchive>::value ||
                 std::is_same<Archive, ::cereal::BinaryOutputArchive>::value,
  T && >::type
  make_nvp( const char *, T && value )
  {
    return std::forward<T>(value);
  }

  //! A specialization of make_nvp<> that actually creates an nvp for non-binary archives
  /*! @relates NameValuePair
      @internal */
  template<class Archive, class T> inline
  typename
  std::enable_if<!std::is_same<Archive, ::cereal::BinaryInputArchive>::value &&
                 !std::is_same<Archive, ::cereal::BinaryOutputArchive>::value,
  NameValuePair<T> >::type
  make_nvp( const char * name, T && value)
  {
    return {name, std::forward<T>(value)};
  }

  //! Convenience for creating a templated NVP
  /*! For use in internal generic typing functions which have an
      Archive type declared
      @internal */
  #define CEREAL_NVP_(name, value) ::cereal::make_nvp<Archive>(name, value)

  // ######################################################################
  //! A wrapper around data that can be serialized in a binary fashion
  /*! This class is used to demarcate data that can safely be serialized
      as a binary chunk of data.  Individual archives can then choose how
      best represent this during serialization.

      @internal */
  template <class T>
  struct BinaryData
  {
    //! Internally store the pointer as a void *, keeping const if created with
    //! a const pointer
    using PT = typename std::conditional<std::is_const<typename std::remove_pointer<typename std::remove_reference<T>::type>::type>::value,
                                         const void *,
                                         void *>::type;

    BinaryData( T && d, uint64_t s ) : data(std::forward<T>(d)), size(s) {}

    PT data;       //!< pointer to beginning of data
    uint64_t size; //!< size in bytes
  };

  // ######################################################################
  //! A wrapper around data that should be serialized after all non-deferred data
  /*! This class is used to demarcate data that can only be safely serialized after
      any data not wrapped in this class.

      @internal */
  template <class T>
  class DeferredData : detail::DeferredDataCore
  {
    private:
      // If we get passed an array, keep the type as is, otherwise store
      // a reference if we were passed an l value reference, else copy the value
      using Type = typename std::conditional<std::is_array<typename std::remove_reference<T>::type>::value,
                                             typename std::remove_cv<T>::type,
                                             typename std::conditional<std::is_lvalue_reference<T>::value,
                                                                       T,
                                                                       typename std::decay<T>::type>::type>::type;

      // prevent nested nvps
      static_assert( !std::is_base_of<detail::DeferredDataCore, T>::value,
                     "Cannot defer DeferredData" );

      DeferredData & operator=( DeferredData const & ) = delete;

    public:
      //! Constructs a new NameValuePair
      /*! @param v The value to defer.  Ideally this should be an l-value reference so that
                   the value can be both loaded and saved to.  If you pass an r-value reference,
                   the DeferredData will store a copy of it instead of a reference.  Thus you should
                   only pass r-values in cases where this makes sense, such as the result of some
                   size() call.
          @internal */
      DeferredData( T && v ) : value(std::forward<T>(v)) {}

      Type value;
  };

  // ######################################################################
  namespace detail
  {
    // base classes for type checking
    /* The rtti virtual function only exists to enable an archive to
       be used in a polymorphic fashion, if necessary.  See the
       archive adapters for an example of this */
    class OutputArchiveBase
    {
      public:
        OutputArchiveBase() = default;
        OutputArchiveBase( OutputArchiveBase && ) CEREAL_NOEXCEPT {}
        OutputArchiveBase & operator=( OutputArchiveBase && ) CEREAL_NOEXCEPT { return *this; }
        virtual ~OutputArchiveBase() CEREAL_NOEXCEPT = default;

      private:
        virtual void rtti() {}
    };

    class InputArchiveBase
    {
      public:
        InputArchiveBase() = default;
        InputArchiveBase( InputArchiveBase && ) CEREAL_NOEXCEPT {}
        InputArchiveBase & operator=( InputArchiveBase && ) CEREAL_NOEXCEPT { return *this; }
        virtual ~InputArchiveBase() CEREAL_NOEXCEPT = default;

      private:
        virtual void rtti() {}
    };

    // forward decls for polymorphic support
    template <class Archive, class T> struct polymorphic_serialization_support;
    struct adl_tag;

    // used during saving pointers
    static const uint32_t msb_32bit  = 0x80000000;
    static const int32_t msb2_32bit = 0x40000000;
  }

  // ######################################################################
  //! A wrapper around size metadata
  /*! This class provides a way for archives to have more flexibility over how
      they choose to serialize size metadata for containers.  For some archive
      types, the size may be implicitly encoded in the output (e.g. JSON) and
      not need an explicit entry.  Specializing serialize or load/save for
      your archive and SizeTags allows you to choose what happens.

      @internal */
  template <class T>
  class SizeTag
  {
    private:
      // Store a reference if passed an lvalue reference, otherwise
      // make a copy of the data
      using Type = typename std::conditional<std::is_lvalue_reference<T>::value,
                                             T,
                                             typename std::decay<T>::type>::type;

      SizeTag & operator=( SizeTag const & ) = delete;

    public:
      SizeTag( T && sz ) : size(std::forward<T>(sz)) {}

      Type size;
  };

  // ######################################################################
  //! A wrapper around a key and value for serializing data into maps.
  /*! This class just provides a grouping of keys and values into a struct for
      human readable archives. For example, XML archives will use this wrapper
      to write maps like so:

      @code{.xml}
      <mymap>
        <item0>
          <key>MyFirstKey</key>
          <value>MyFirstValue</value>
        </item0>
        <item1>
          <key>MySecondKey</key>
          <value>MySecondValue</value>
        </item1>
      </mymap>
      @endcode

      \sa make_map_item
      @internal */
  template <class Key, class Value>
  struct MapItem
  {
    using KeyType = typename std::conditional<
      std::is_lvalue_reference<Key>::value,
      Key,
      typename std::decay<Key>::type>::type;

    using ValueType = typename std::conditional<
      std::is_lvalue_reference<Value>::value,
      Value,
      typename std::decay<Value>::type>::type;

    //! Construct a MapItem from a key and a value
    /*! @internal */
    MapItem( Key && key_, Value && value_ ) : key(std::forward<Key>(key_)), value(std::forward<Value>(value_)) {}

    MapItem & operator=( MapItem const & ) = delete;

    KeyType key;
    ValueType value;

    //! Serialize the MapItem with the NVPs "key" and "value"
    template <class Archive> inline
    void CEREAL_SERIALIZE_FUNCTION_NAME(Archive & archive)
    {
      archive( make_nvp<Archive>("key",   key),
               make_nvp<Archive>("value", value) );
    }
  };

  //! Create a MapItem so that human readable archives will group keys and values together
  /*! @internal
      @relates MapItem */
  template <class KeyType, class ValueType> inline
  MapItem<KeyType, ValueType> make_map_item(KeyType && key, ValueType && value)
  {
    return {std::forward<KeyType>(key), std::forward<ValueType>(value)};
  }

  namespace detail
  {
    //! Tag for Version, which due to its anonymous namespace, becomes a different
    //! type in each translation unit
    /*! This allows CEREAL_CLASS_VERSION to be safely called in a header file */
    namespace{ struct version_binding_tag {}; }

    // ######################################################################
    //! Version information class
    /*! This is the base case for classes that have not been explicitly
        registered */
    template <class T, class BindingTag = version_binding_tag> struct Version
    {
      static const std::uint32_t version = 0;
      // we don't need to explicitly register these types since they
      // always get a version number of 0
    };

    //! Holds all registered version information
    struct Versions
    {
      std::unordered_map<std::size_t, std::uint32_t> mapping;

      std::uint32_t find( std::size_t hash, std::uint32_t version )
      {
        const auto result = mapping.emplace( hash, version );
        return result.first->second;
      }
    }; // struct Versions
  } // namespace detail
} // namespace cereal

#endif // CEREAL_DETAILS_HELPERS_HPP_
```
