## Supplemental code for "The motif composition of variable-number tandem repeats impacts gene expression": polymorphic_impl.hpp

```
/*! \file polymorphic_impl.hpp
    \brief Internal polymorphism support
    \ingroup Internal */
/*
  Copyright (c) 2014, Randolph Voorhies, Shane Grant
  All rights reserved.

  Redistribution and use in source and binary forms, with or without
  modification, are permitted provided that the following conditions are met:
      * Redistributions of source code must retain the above copyright
        notice, this list of conditions and the following disclaimer.
      * Redistributions in binary form must reproduce the above copyright
        notice, this list of conditions and the following disclaimer in the
        documentation and/or other materials provided with the distribution.
      * Neither the name of cereal nor the
        names of its contributors may be used to endorse or promote products
        derived from this software without specific prior written permission.

  THIS SOFTWARE IS PROVIDED BY THE COPYRIGHT HOLDERS AND CONTRIBUTORS "AS IS" AND
  ANY EXPRESS OR IMPLIED WARRANTIES, INCLUDING, BUT NOT LIMITED TO, THE IMPLIED
  WARRANTIES OF MERCHANTABILITY AND FITNESS FOR A PARTICULAR PURPOSE ARE
  DISCLAIMED. IN NO EVENT SHALL RANDOLPH VOORHIES OR SHANE GRANT BE LIABLE FOR ANY
  DIRECT, INDIRECT, INCIDENTAL, SPECIAL, EXEMPLARY, OR CONSEQUENTIAL DAMAGES
  (INCLUDING, BUT NOT LIMITED TO, PROCUREMENT OF SUBSTITUTE GOODS OR SERVICES;
  LOSS OF USE, DATA, OR PROFITS; OR BUSINESS INTERRUPTION) HOWEVER CAUSED AND
  ON ANY THEORY OF LIABILITY, WHETHER IN CONTRACT, STRICT LIABILITY, OR TORT
  (INCLUDING NEGLIGENCE OR OTHERWISE) ARISING IN ANY WAY OUT OF THE USE OF THIS
  SOFTWARE, EVEN IF ADVISED OF THE POSSIBILITY OF SUCH DAMAGE.
*/

/* This code is heavily inspired by the boost serialization implementation by the following authors

   (C) Copyright 2002 Robert Ramey - http://www.rrsd.com .
   Use, modification and distribution is subject to the Boost Software
   License, Version 1.0. (See http://www.boost.org/LICENSE_1_0.txt)

    See http://www.boost.org for updates, documentation, and revision history.

   (C) Copyright 2006 David Abrahams - http://www.boost.org.

   See /boost/serialization/export.hpp, /boost/archive/detail/register_archive.hpp,
   and /boost/serialization/void_cast.hpp for their implementation. Additional details
   found in other files split across serialization and archive.
*/
#ifndef CEREAL_DETAILS_POLYMORPHIC_IMPL_HPP_
#define CEREAL_DETAILS_POLYMORPHIC_IMPL_HPP_

#include "cereal/details/polymorphic_impl_fwd.hpp"
#include "cereal/details/static_object.hpp"
#include "cereal/types/memory.hpp"
#include "cereal/types/string.hpp"
#include <functional>
#include <typeindex>
#include <map>
#include <limits>
#include <set>
#include <stack>

//! Helper macro to omit unused warning
#if defined(__GNUC__)
  // GCC / clang don't want the function
  #define CEREAL_BIND_TO_ARCHIVES_UNUSED_FUNCTION
#else
  #define CEREAL_BIND_TO_ARCHIVES_UNUSED_FUNCTION static void unused() { (void)b; }
#endif

//! Binds a polymorhic type to all registered archives
/*! This binds a polymorphic type to all compatible registered archives that
    have been registered with CEREAL_REGISTER_ARCHIVE.  This must be called
    after all archives are registered (usually after the archives themselves
    have been included). */
#ifdef CEREAL_HAS_CPP17
#define CEREAL_BIND_TO_ARCHIVES(...)                                     \
    namespace cereal {                                                   \
    namespace detail {                                                   \
    template<>                                                           \
    struct init_binding<__VA_ARGS__> {                                   \
        static inline bind_to_archives<__VA_ARGS__> const & b=           \
        ::cereal::detail::StaticObject<                                  \
            bind_to_archives<__VA_ARGS__>                                \
        >::getInstance().bind();                                         \
        CEREAL_BIND_TO_ARCHIVES_UNUSED_FUNCTION                          \
    };                                                                   \
    }} /* end namespaces */
#else
#define CEREAL_BIND_TO_ARCHIVES(...)                                     \
    namespace cereal {                                                   \
    namespace detail {                                                   \
    template<>                                                           \
    struct init_binding<__VA_ARGS__> {                                   \
        static bind_to_archives<__VA_ARGS__> const& b;                   \
        CEREAL_BIND_TO_ARCHIVES_UNUSED_FUNCTION                          \
    };                                                                   \
    bind_to_archives<__VA_ARGS__> const & init_binding<__VA_ARGS__>::b = \
        ::cereal::detail::StaticObject<                                  \
            bind_to_archives<__VA_ARGS__>                                \
        >::getInstance().bind();                                         \
    }} /* end namespaces */
#endif

namespace cereal
{
  /* Polymorphic casting support */
  namespace detail
  {
    //! Base type for polymorphic void casting
    /*! Contains functions for casting between registered base and derived types.

        This is necessary so that cereal can properly cast between polymorphic types
        even though void pointers are used, which normally have no type information.
        Runtime type information is used instead to index a compile-time made mapping
        that can perform the proper cast. In the case of multiple levels of inheritance,
        cereal will attempt to find the shortest path by using registered relationships to
        perform the cast.

        This class will be allocated as a StaticObject and only referenced by pointer,
        allowing a templated derived version of it to define strongly typed functions
        that cast between registered base and derived types. */
    struct PolymorphicCaster
    {
      PolymorphicCaster() = default;
      PolymorphicCaster( const PolymorphicCaster & ) = default;
      PolymorphicCaster & operator=( const PolymorphicCaster & ) = default;
      PolymorphicCaster( PolymorphicCaster && ) CEREAL_NOEXCEPT {}
      PolymorphicCaster & operator=( PolymorphicCaster && ) CEREAL_NOEXCEPT { return *this; }
      virtual ~PolymorphicCaster() CEREAL_NOEXCEPT = default;

      //! Downcasts to the proper derived type
      virtual void const * downcast( void const * const ptr ) const = 0;
      //! Upcast to proper base type
      virtual void * upcast( void * const ptr ) const = 0;
      //! Upcast to proper base type, shared_ptr version
      virtual std::shared_ptr<void> upcast( std::shared_ptr<void> const & ptr ) const = 0;
    };

    //! Holds registered mappings between base and derived types for casting
    /*! This will be allocated as a StaticObject and holds a map containing
        all registered mappings between base and derived types. */
    struct PolymorphicCasters
    {
      //! Maps from a derived type index to a set of chainable casters
      using DerivedCasterMap = std::unordered_map<std::type_index, std::vector<PolymorphicCaster const *>>;
      //! Maps from base type index to a map from derived type index to caster
      std::unordered_map<std::type_index, DerivedCasterMap> map;

      std::multimap<std::type_index, std::type_index> reverseMap;

      //! Error message used for unregistered polymorphic casts
      #define UNREGISTERED_POLYMORPHIC_CAST_EXCEPTION(LoadSave)                                                                                                                \
        throw cereal::Exception("Trying to " #LoadSave " a registered polymorphic type with an unregistered polymorphic cast.\n"                                               \
                                "Could not find a path to a base class (" + util::demangle(baseInfo.name()) + ") for type: " + ::cereal::util::demangledName<Derived>() + "\n" \
                                "Make sure you either serialize the base class at some point via cereal::base_class or cereal::virtual_base_class.\n"                          \
                                "Alternatively, manually register the association with CEREAL_REGISTER_POLYMORPHIC_RELATION.");

      //! Checks if the mapping object that can perform the upcast or downcast exists, and returns it if so
      /*! Uses the type index from the base and derived class to find the matching
          registered caster. If no matching caster exists, the bool in the pair will be false and the vector
          reference should not be used. */
      static std::pair<bool, std::vector<PolymorphicCaster const *> const &>
      lookup_if_exists( std::type_index const & baseIndex, std::type_index const & derivedIndex )
      {
        // First phase of lookup - match base type index
        auto const & baseMap = StaticObject<PolymorphicCasters>::getInstance().map;
        auto baseIter = baseMap.find( baseIndex );
        if (baseIter == baseMap.end())
          return {false, {}};

        // Second phase - find a match from base to derived
        auto const & derivedMap = baseIter->second;
        auto derivedIter = derivedMap.find( derivedIndex );
        if (derivedIter == derivedMap.end())
          return {false, {}};

        return {true, derivedIter->second};
      }

      //! Gets the mapping object that can perform the upcast or downcast
      /*! Uses the type index from the base and derived class to find the matching
          registered caster. If no matching caster exists, calls the exception function.

          The returned PolymorphicCaster is capable of upcasting or downcasting between the two types. */
      template <class F> inline
      static std::vector<PolymorphicCaster const *> const & lookup( std::type_index const & baseIndex, std::type_index const & derivedIndex, F && exceptionFunc )
      {
        // First phase of lookup - match base type index
        auto const & baseMap = StaticObject<PolymorphicCasters>::getInstance().map;
        auto baseIter = baseMap.find( baseIndex );
        if( baseIter == baseMap.end() )
          exceptionFunc();

        // Second phase - find a match from base to derived
        auto const & derivedMap = baseIter->second;
        auto derivedIter = derivedMap.find( derivedIndex );
        if( derivedIter == derivedMap.end() )
          exceptionFunc();

        return derivedIter->second;
      }

      //! Performs a downcast to the derived type using a registered mapping
      template <class Derived> inline
      static const Derived * downcast( const void * dptr, std::type_info const & baseInfo )
      {
        auto const & mapping = lookup( baseInfo, typeid(Derived), [&](){ UNREGISTERED_POLYMORPHIC_CAST_EXCEPTION(save) } );

        for( auto const * dmap : mapping )
          dptr = dmap->downcast( dptr );

        return static_cast<Derived const *>( dptr );
      }

      //! Performs an upcast to the registered base type using the given a derived type
      /*! The return is untyped because the final casting to the base type must happen in the polymorphic
          serialization function, where the type is known at compile time */
      template <class Derived> inline
      static void * upcast( Derived * const dptr, std::type_info const & baseInfo )
      {
        auto const & mapping = lookup( baseInfo, typeid(Derived), [&](){ UNREGISTERED_POLYMORPHIC_CAST_EXCEPTION(load) } );

        void * uptr = dptr;
        for( auto mIter = mapping.rbegin(), mEnd = mapping.rend(); mIter != mEnd; ++mIter )
          uptr = (*mIter)->upcast( uptr );

        return uptr;
      }

      //! Upcasts for shared pointers
      template <class Derived> inline
      static std::shared_ptr<void> upcast( std::shared_ptr<Derived> const & dptr, std::type_info const & baseInfo )
      {
        auto const & mapping = lookup( baseInfo, typeid(Derived), [&](){ UNREGISTERED_POLYMORPHIC_CAST_EXCEPTION(load) } );

        std::shared_ptr<void> uptr = dptr;
        for( auto mIter = mapping.rbegin(), mEnd = mapping.rend(); mIter != mEnd; ++mIter )
          uptr = (*mIter)->upcast( uptr );

        return uptr;
      }

      #undef UNREGISTERED_POLYMORPHIC_CAST_EXCEPTION
    };

    #ifdef CEREAL_OLDER_GCC
      #define CEREAL_EMPLACE_MAP(map, key, value)                     \
      map.insert( std::make_pair(std::move(key), std::move(value)) );
    #else // NOT CEREAL_OLDER_GCC
      #define CEREAL_EMPLACE_MAP(map, key, value)                     \
      map.emplace( key, value );
    #endif // NOT_CEREAL_OLDER_GCC

    //! Strongly typed derivation of PolymorphicCaster
    template <class Base, class Derived>
    struct PolymorphicVirtualCaster : PolymorphicCaster
    {
      //! Inserts an entry in the polymorphic casting map for this pairing
      /*! Creates an explicit mapping between Base and Derived in both upwards and
          downwards directions, allowing void pointers to either to be properly cast
          assuming dynamic type information is available */
      PolymorphicVirtualCaster()
      {
        const auto baseKey = std::type_index(typeid(Base));
        const auto derivedKey = std::type_index(typeid(Derived));

        // First insert the relation Base->Derived
        const auto lock = StaticObject<PolymorphicCasters>::lock();
        auto & baseMap = StaticObject<PolymorphicCasters>::getInstance().map;

        {
          auto & derivedMap = baseMap.insert( {baseKey, PolymorphicCasters::DerivedCasterMap{}} ).first->second;
          auto & derivedVec = derivedMap.insert( {derivedKey, {}} ).first->second;
          derivedVec.push_back( this );
        }

        // Insert reverse relation Derived->Base
        auto & reverseMap = StaticObject<PolymorphicCasters>::getInstance().reverseMap;
        CEREAL_EMPLACE_MAP(reverseMap, derivedKey, baseKey);

        // Find all chainable unregistered relations
        /* The strategy here is to process only the nodes in the class hierarchy graph that have been
           affected by the new insertion. The aglorithm iteratively processes a node an ensures that it
           is updated with all new shortest length paths. It then rocesses the parents of the active node,
           with the knowledge that all children have already been processed.

           Note that for the following, we'll use the nomenclature of parent and child to not confuse with
           the inserted base derived relationship */
        {
          // Checks whether there is a path from parent->child and returns a <dist, path> pair
          // dist is set to MAX if the path does not exist
          auto checkRelation = [](std::type_index const & parentInfo, std::type_index const & childInfo) ->
            std::pair<size_t, std::vector<PolymorphicCaster const *> const &>
          {
            auto result = PolymorphicCasters::lookup_if_exists( parentInfo, childInfo );
            if( result.first )
            {
              auto const & path = result.second;
              return {path.size(), path};
            }
            else
              return {(std::numeric_limits<size_t>::max)(), {}};
          };

          std::stack<std::type_index>         parentStack;      // Holds the parent nodes to be processed
          std::vector<std::type_index> dirtySet;                // Marks child nodes that have been changed
          std::unordered_set<std::type_index> processedParents; // Marks parent nodes that have been processed

          // Checks if a child has been marked dirty
          auto isDirty = [&](std::type_index const & c)
          {
            auto const dirtySetSize = dirtySet.size();
            for( size_t i = 0; i < dirtySetSize; ++i )
              if( dirtySet[i] == c )
                return true;

            return false;
          };

          // Begin processing the base key and mark derived as dirty
          parentStack.push( baseKey );
          dirtySet.emplace_back( derivedKey );

          while( !parentStack.empty() )
          {
            using Relations = std::unordered_multimap<std::type_index, std::pair<std::type_index, std::vector<PolymorphicCaster const *>>>;
            Relations unregisteredRelations; // Defer insertions until after main loop to prevent iterator invalidation

            const auto parent = parentStack.top();
            parentStack.pop();

            // Update paths to all children marked dirty
            for( auto const & childPair : baseMap[parent] )
            {
              const auto child = childPair.first;
              if( isDirty( child ) && baseMap.count( child ) )
              {
                auto parentChildPath = checkRelation( parent, child );

                // Search all paths from the child to its own children (finalChild),
                // looking for a shorter parth from parent to finalChild
                for( auto const & finalChildPair : baseMap[child] )
                {
                  const auto finalChild = finalChildPair.first;

                  auto parentFinalChildPath = checkRelation( parent, finalChild );
                  auto childFinalChildPath  = checkRelation( child, finalChild );

                  const size_t newLength = 1u + parentChildPath.first;

                  if( newLength < parentFinalChildPath.first )
                  {
                    std::vector<PolymorphicCaster const *> path = parentChildPath.second;
                    path.insert( path.end(), childFinalChildPath.second.begin(), childFinalChildPath.second.end() );

                    // Check to see if we have a previous uncommitted path in unregisteredRelations
                    // that is shorter. If so, ignore this path
                    auto hintRange = unregisteredRelations.equal_range( parent );
                    auto hint = hintRange.first;
                    for( ; hint != hintRange.second; ++hint )
                      if( hint->second.first == finalChild )
                        break;

                    const bool uncommittedExists = hint != unregisteredRelations.end();
                    if( uncommittedExists && (hint->second.second.size() <= newLength) )
                      continue;

                    auto newPath = std::pair<std::type_index, std::vector<PolymorphicCaster const *>>{finalChild, std::move(path)};

                    // Insert the new path if it doesn't exist, otherwise this will just lookup where to do the
                    // replacement
                    #ifdef CEREAL_OLDER_GCC
                    auto old = unregisteredRelations.insert( hint, std::make_pair(parent, newPath) );
                    #else // NOT CEREAL_OLDER_GCC
                    auto old = unregisteredRelations.emplace_hint( hint, parent, newPath );
                    #endif // NOT CEREAL_OLDER_GCC

                    // If there was an uncommitted path, we need to perform a replacement
                    if( uncommittedExists )
                      old->second = newPath;
                  }
                } // end loop over child's children
              } // end if dirty and child has children
            } // end loop over children

            // Insert chained relations
            for( auto const & it : unregisteredRelations )
            {
              auto & derivedMap = baseMap.find( it.first )->second;
              derivedMap[it.second.first] = it.second.second;
              CEREAL_EMPLACE_MAP(reverseMap, it.second.first, it.first );
            }

            // Mark current parent as modified
            dirtySet.emplace_back( parent );

            // Insert all parents of the current parent node that haven't yet been processed
            auto parentRange = reverseMap.equal_range( parent );
            for( auto pIter = parentRange.first; pIter != parentRange.second; ++pIter )
            {
              const auto pParent = pIter->second;
              if( !processedParents.count( pParent ) )
              {
                parentStack.push( pParent );
                processedParents.insert( pParent );
              }
            }
          } // end loop over parent stack
        } // end chainable relations
      } // end PolymorphicVirtualCaster()

      #undef CEREAL_EMPLACE_MAP

      //! Performs the proper downcast with the templated types
      void const * downcast( void const * const ptr ) const override
      {
        return dynamic_cast<Derived const*>( static_cast<Base const*>( ptr ) );
      }

      //! Performs the proper upcast with the templated types
      void * upcast( void * const ptr ) const override
      {
        return dynamic_cast<Base*>( static_cast<Derived*>( ptr ) );
      }

      //! Performs the proper upcast with the templated types (shared_ptr version)
      std::shared_ptr<void> upcast( std::shared_ptr<void> const & ptr ) const override
      {
        return std::dynamic_pointer_cast<Base>( std::static_pointer_cast<Derived>( ptr ) );
      }
    };

    //! Registers a polymorphic casting relation between a Base and Derived type
    /*! Registering a relation allows cereal to properly cast between the two types
        given runtime type information and void pointers.

        Registration happens automatically via cereal::base_class and cereal::virtual_base_class
        instantiations. For cases where neither is called, see the CEREAL_REGISTER_POLYMORPHIC_RELATION
        macro */
    template <class Base, class Derived>
    struct RegisterPolymorphicCaster
    {
      static PolymorphicCaster const * bind( std::true_type /* is_polymorphic<Base> */)
      {
        return &StaticObject<PolymorphicVirtualCaster<Base, Derived>>::getInstance();
      }

      static PolymorphicCaster const * bind( std::false_type /* is_polymorphic<Base> */ )
      { return nullptr; }

      //! Performs registration (binding) between Base and Derived
      /*! If the type is not polymorphic, nothing will happen */
      static PolymorphicCaster const * bind()
      { return bind( typename std::is_polymorphic<Base>::type() ); }
    };
  }

  /* General polymorphism support */
  namespace detail
  {
    //! Binds a compile time type with a user defined string
    template <class T>
    struct binding_name {};

    //! A structure holding a map from type_indices to output serializer functions
    /*! A static object of this map should be created for each registered archive
        type, containing entries for every registered type that describe how to
        properly cast the type to its real type in polymorphic scenarios for
        shared_ptr, weak_ptr, and unique_ptr. */
    template <class Archive>
    struct OutputBindingMap
    {
      //! A serializer function
      /*! Serializer functions return nothing and take an archive as
          their first parameter (will be cast properly inside the function,
          a pointer to actual data (contents of smart_ptr's get() function)
          as their second parameter, and the type info of the owning smart_ptr
          as their final parameter */
      typedef std::function<void(void*, void const *, std::type_info const &)> Serializer;

      //! Struct containing the serializer functions for all pointer types
      struct Serializers
      {
        Serializer shared_ptr, //!< Serializer function for shared/weak pointers
                   unique_ptr; //!< Serializer function for unique pointers
      };

      //! A map of serializers for pointers of all registered types
      std::map<std::type_index, Serializers> map;
    };

    //! An empty noop deleter
    template<class T> struct EmptyDeleter { void operator()(T *) const {} };

    //! A structure holding a map from type name strings to input serializer functions
    /*! A static object of this map should be created for each registered archive
        type, containing entries for every registered type that describe how to
        properly cast the type to its real type in polymorphic scenarios for
        shared_ptr, weak_ptr, and unique_ptr. */
    template <class Archive>
    struct InputBindingMap
    {
      //! Shared ptr serializer function
      /*! Serializer functions return nothing and take an archive as
          their first parameter (will be cast properly inside the function,
          a shared_ptr (or unique_ptr for the unique case) of any base
          type, and the type id of said base type as the third parameter.
          Internally it will properly be loaded and cast to the correct type. */
      typedef std::function<void(void*, std::shared_ptr<void> &, std::type_info const &)> SharedSerializer;
      //! Unique ptr serializer function
      typedef std::function<void(void*, std::unique_ptr<void, EmptyDeleter<void>> &, std::type_info const &)> UniqueSerializer;

      //! Struct containing the serializer functions for all pointer types
      struct Serializers
      {
        SharedSerializer shared_ptr; //!< Serializer function for shared/weak pointers
        UniqueSerializer unique_ptr; //!< Serializer function for unique pointers
      };

      //! A map of serializers for pointers of all registered types
      std::map<std::string, Serializers> map;
    };

    // forward decls for archives from cereal.hpp
    class InputArchiveBase;
    class OutputArchiveBase;

    //! Creates a binding (map entry) between an input archive type and a polymorphic type
    /*! Bindings are made when types are registered, assuming that at least one
        archive has already been registered.  When this struct is created,
        it will insert (at run time) an entry into a map that properly handles
        casting for serializing polymorphic objects */
    template <class Archive, class T> struct InputBindingCreator
    {
      //! Initialize the binding
      InputBindingCreator()
      {
        auto & map = StaticObject<InputBindingMap<Archive>>::getInstance().map;
        auto lock = StaticObject<InputBindingMap<Archive>>::lock();
        auto key = std::string(binding_name<T>::name());
        auto lb = map.lower_bound(key);

        if (lb != map.end() && lb->first == key)
          return;

        typename InputBindingMap<Archive>::Serializers serializers;

        serializers.shared_ptr =
          [](void * arptr, std::shared_ptr<void> & dptr, std::type_info const & baseInfo)
          {
            Archive & ar = *static_cast<Archive*>(arptr);
            std::shared_ptr<T> ptr;

            ar( CEREAL_NVP_("ptr_wrapper", ::cereal::memory_detail::make_ptr_wrapper(ptr)) );

            dptr = PolymorphicCasters::template upcast<T>( ptr, baseInfo );
          };

        serializers.unique_ptr =
          [](void * arptr, std::unique_ptr<void, EmptyDeleter<void>> & dptr, std::type_info const & baseInfo)
          {
            Archive & ar = *static_cast<Archive*>(arptr);
            std::unique_ptr<T> ptr;

            ar( CEREAL_NVP_("ptr_wrapper", ::cereal::memory_detail::make_ptr_wrapper(ptr)) );

            dptr.reset( PolymorphicCasters::template upcast<T>( ptr.release(), baseInfo ));
          };

        map.insert( lb, { std::move(key), std::move(serializers) } );
      }
    };

    //! Creates a binding (map entry) between an output archive type and a polymorphic type
    /*! Bindings are made when types are registered, assuming that at least one
        archive has already been registered.  When this struct is created,
        it will insert (at run time) an entry into a map that properly handles
        casting for serializing polymorphic objects */
    template <class Archive, class T> struct OutputBindingCreator
    {
      //! Writes appropriate metadata to the archive for this polymorphic type
      static void writeMetadata(Archive & ar)
      {
        // Register the polymorphic type name with the archive, and get the id
        char const * name = binding_name<T>::name();
        std::uint32_t id = ar.registerPolymorphicType(name);

        // Serialize the id
        ar( CEREAL_NVP_("polymorphic_id", id) );

        // If the msb of the id is 1, then the type name is new, and we should serialize it
        if( id & detail::msb_32bit )
        {
          std::string namestring(name);
          ar( CEREAL_NVP_("polymorphic_name", namestring) );
        }
      }

      //! Holds a properly typed shared_ptr to the polymorphic type
      class PolymorphicSharedPointerWrapper
      {
        public:
          /*! Wrap a raw polymorphic pointer in a shared_ptr to its true type

              The wrapped pointer will not be responsible for ownership of the held pointer
              so it will not attempt to destroy it; instead the refcount of the wrapped
              pointer will be tied to a fake 'ownership pointer' that will do nothing
              when it ultimately goes out of scope.

              The main reason for doing this, other than not to destroy the true object
              with our wrapper pointer, is to avoid meddling with the internal reference
              count in a polymorphic type that inherits from std::enable_shared_from_this.

              @param dptr A void pointer to the contents of the shared_ptr to serialize */
          PolymorphicSharedPointerWrapper( T const * dptr ) : refCount(), wrappedPtr( refCount, dptr )
          { }

          //! Get the wrapped shared_ptr */
          inline std::shared_ptr<T const> const & operator()() const { return wrappedPtr; }

        private:
          std::shared_ptr<void> refCount;      //!< The ownership pointer
          std::shared_ptr<T const> wrappedPtr; //!< The wrapped pointer
      };

      //! Does the actual work of saving a polymorphic shared_ptr
      /*! This function will properly create a shared_ptr from the void * that is passed in
          before passing it to the archive for serialization.

          In addition, this will also preserve the state of any internal enable_shared_from_this mechanisms

          @param ar The archive to serialize to
          @param dptr Pointer to the actual data held by the shared_ptr */
      static inline void savePolymorphicSharedPtr( Archive & ar, T const * dptr, std::true_type /* has_shared_from_this */ )
      {
        ::cereal::memory_detail::EnableSharedStateHelper<T> state( const_cast<T *>(dptr) );
        PolymorphicSharedPointerWrapper psptr( dptr );
        ar( CEREAL_NVP_("ptr_wrapper", memory_detail::make_ptr_wrapper( psptr() ) ) );
      }

      //! Does the actual work of saving a polymorphic shared_ptr
      /*! This function will properly create a shared_ptr from the void * that is passed in
          before passing it to the archive for serialization.

          This version is for types that do not inherit from std::enable_shared_from_this.

          @param ar The archive to serialize to
          @param dptr Pointer to the actual data held by the shared_ptr */
      static inline void savePolymorphicSharedPtr( Archive & ar, T const * dptr, std::false_type /* has_shared_from_this */ )
      {
        PolymorphicSharedPointerWrapper psptr( dptr );
        ar( CEREAL_NVP_("ptr_wrapper", memory_detail::make_ptr_wrapper( psptr() ) ) );
      }

      //! Initialize the binding
      OutputBindingCreator()
      {
        auto & map = StaticObject<OutputBindingMap<Archive>>::getInstance().map;
        auto key = std::type_index(typeid(T));
        auto lb = map.lower_bound(key);

        if (lb != map.end() && lb->first == key)
          return;

        typename OutputBindingMap<Archive>::Serializers serializers;

        serializers.shared_ptr =
          [&](void * arptr, void const * dptr, std::type_info const & baseInfo)
          {
            Archive & ar = *static_cast<Archive*>(arptr);
            writeMetadata(ar);

            auto ptr = PolymorphicCasters::template downcast<T>( dptr, baseInfo );

            #if defined(_MSC_VER) && _MSC_VER < 1916 && !defined(__clang__)
            savePolymorphicSharedPtr( ar, ptr, ::cereal::traits::has_shared_from_this<T>::type() ); // MSVC doesn't like typename here
            #else // not _MSC_VER
            savePolymorphicSharedPtr( ar, ptr, typename ::cereal::traits::has_shared_from_this<T>::type() );
            #endif // _MSC_VER
          };

        serializers.unique_ptr =
          [&](void * arptr, void const * dptr, std::type_info const & baseInfo)
          {
            Archive & ar = *static_cast<Archive*>(arptr);
            writeMetadata(ar);

            std::unique_ptr<T const, EmptyDeleter<T const>> const ptr( PolymorphicCasters::template downcast<T>( dptr, baseInfo ) );

            ar( CEREAL_NVP_("ptr_wrapper", memory_detail::make_ptr_wrapper(ptr)) );
          };

        map.insert( { std::move(key), std::move(serializers) } );
      }
    };

    //! Used to help out argument dependent lookup for finding potential overloads
    //! of instantiate_polymorphic_binding
    struct adl_tag {};

    //! Tag for init_binding, bind_to_archives and instantiate_polymorphic_binding.
    //! For C++14 and below, we must instantiate a unique StaticObject per TU that is
    //! otherwise identical -- otherwise we get multiple definition problems (ODR violations).
    //! To achieve this, put a tag in an anonymous namespace and use it as a template argument.
    //!
    //! For C++17, we can use static inline global variables to unify these definitions across
    //! all TUs in the same shared object (DLL).  The tag is therefore not necessary.
    //! For convenience, keep it to not complicate other code, but don't put it in
    //! an anonymous namespace.  Now the template instantiations will correspond
    //! to the same type, and since they are marked inline with C++17, they will be merged
    //! across all TUs.
#ifdef CEREAL_HAS_CPP17
    struct polymorphic_binding_tag {};
#else
    namespace { struct polymorphic_binding_tag {}; }
#endif


    //! Causes the static object bindings between an archive type and a serializable type T
    template <class Archive, class T>
    struct create_bindings
    {
      static const InputBindingCreator<Archive, T> &
      load(std::true_type)
      {
        return cereal::detail::StaticObject<InputBindingCreator<Archive, T>>::getInstance();
      }

      static const OutputBindingCreator<Archive, T> &
      save(std::true_type)
      {
        return cereal::detail::StaticObject<OutputBindingCreator<Archive, T>>::getInstance();
      }

      inline static void load(std::false_type) {}
      inline static void save(std::false_type) {}
    };

    //! When specialized, causes the compiler to instantiate its parameter
    template <void(*)()>
    struct instantiate_function {};

    /*! This struct is used as the return type of instantiate_polymorphic_binding
        for specific Archive types.  When the compiler looks for overloads of
        instantiate_polymorphic_binding, it will be forced to instantiate this
        struct during overload resolution, even though it will not be part of a valid
        overload */
    template <class Archive, class T>
    struct polymorphic_serialization_support
    {
      #if defined(_MSC_VER) && !defined(__INTEL_COMPILER)
      //! Creates the appropriate bindings depending on whether the archive supports
      //! saving or loading
      virtual CEREAL_DLL_EXPORT void instantiate() CEREAL_USED;
      #else // NOT _MSC_VER
      //! Creates the appropriate bindings depending on whether the archive supports
      //! saving or loading
      static CEREAL_DLL_EXPORT void instantiate() CEREAL_USED;
      //! This typedef causes the compiler to instantiate this static function
      typedef instantiate_function<instantiate> unused;
      #endif // _MSC_VER
    };

    // instantiate implementation
    template <class Archive, class T>
    CEREAL_DLL_EXPORT void polymorphic_serialization_support<Archive,T>::instantiate()
    {
      create_bindings<Archive,T>::save( std::integral_constant<bool,
                                          std::is_base_of<detail::OutputArchiveBase, Archive>::value &&
                                          traits::is_output_serializable<T, Archive>::value>{} );

      create_bindings<Archive,T>::load( std::integral_constant<bool,
                                          std::is_base_of<detail::InputArchiveBase, Archive>::value &&
                                          traits::is_input_serializable<T, Archive>::value>{} );
    }

    //! Begins the binding process of a type to all registered archives
    /*! Archives need to be registered prior to this struct being instantiated via
        the CEREAL_REGISTER_ARCHIVE macro.  Overload resolution will then force
        several static objects to be made that allow us to bind together all
        registered archive types with the parameter type T. */
    template <class T, class Tag = polymorphic_binding_tag>
    struct bind_to_archives
    {
      //! Binding for non abstract types
      void bind(std::false_type) const
      {
        instantiate_polymorphic_binding(static_cast<T*>(nullptr), 0, Tag{}, adl_tag{});
      }

      //! Binding for abstract types
      void bind(std::true_type) const
      { }

      //! Binds the type T to all registered archives
      /*! If T is abstract, we will not serialize it and thus
          do not need to make a binding */
      bind_to_archives const & bind() const
      {
        static_assert( std::is_polymorphic<T>::value,
                       "Attempting to register non polymorphic type" );
        bind( std::is_abstract<T>() );
        return *this;
      }
    };

    //! Used to hide the static object used to bind T to registered archives
    template <class T, class Tag = polymorphic_binding_tag>
    struct init_binding;

    //! Base case overload for instantiation
    /*! This will end up always being the best overload due to the second
        parameter always being passed as an int.  All other overloads will
        accept pointers to archive types and have lower precedence than int.

        Since the compiler needs to check all possible overloads, the
        other overloads created via CEREAL_REGISTER_ARCHIVE, which will have
        lower precedence due to requring a conversion from int to (Archive*),
        will cause their return types to be instantiated through the static object
        mechanisms even though they are never called.

        See the documentation for the other functions to try and understand this */
    template <class T, typename BindingTag>
    void instantiate_polymorphic_binding( T*, int, BindingTag, adl_tag ) {}
  } // namespace detail
} // namespace cereal

#endif // CEREAL_DETAILS_POLYMORPHIC_IMPL_HPP_
```
