## Supplemental code for "The motif composition of variable-number tandem repeats impacts gene expression": static_object.hpp

```
/*! \file static_object.hpp
    \brief Internal polymorphism static object support
    \ingroup Internal */
/*
  Copyright (c) 2014, Randolph Voorhies, Shane Grant
  All rights reserved.
  Redistribution and use in source and binary forms, with or without
  modification, are permitted provided that the following conditions are met:
      * Redistributions of source code must retain the above copyright
        notice, this list of conditions and the following disclaimer.
      * Redistributions in binary form must reproduce the above copyright
        notice, this list of conditions and the following disclaimer in the
        documentation and/or other materials provided with the distribution.
      * Neither the name of cereal nor the
        names of its contributors may be used to endorse or promote products
        derived from this software without specific prior written permission.
  THIS SOFTWARE IS PROVIDED BY THE COPYRIGHT HOLDERS AND CONTRIBUTORS "AS IS" AND
  ANY EXPRESS OR IMPLIED WARRANTIES, INCLUDING, BUT NOT LIMITED TO, THE IMPLIED
  WARRANTIES OF MERCHANTABILITY AND FITNESS FOR A PARTICULAR PURPOSE ARE
  DISCLAIMED. IN NO EVENT SHALL RANDOLPH VOORHIES OR SHANE GRANT BE LIABLE FOR ANY
  DIRECT, INDIRECT, INCIDENTAL, SPECIAL, EXEMPLARY, OR CONSEQUENTIAL DAMAGES
  (INCLUDING, BUT NOT LIMITED TO, PROCUREMENT OF SUBSTITUTE GOODS OR SERVICES;
  LOSS OF USE, DATA, OR PROFITS; OR BUSINESS INTERRUPTION) HOWEVER CAUSED AND
  ON ANY THEORY OF LIABILITY, WHETHER IN CONTRACT, STRICT LIABILITY, OR TORT
  (INCLUDING NEGLIGENCE OR OTHERWISE) ARISING IN ANY WAY OUT OF THE USE OF THIS
  SOFTWARE, EVEN IF ADVISED OF THE POSSIBILITY OF SUCH DAMAGE.
*/
#ifndef CEREAL_DETAILS_STATIC_OBJECT_HPP_
#define CEREAL_DETAILS_STATIC_OBJECT_HPP_

#include "cereal/macros.hpp"

#if CEREAL_THREAD_SAFE
#include <mutex>
#endif

//! Prevent link optimization from removing non-referenced static objects
/*! Especially for polymorphic support, we create static objects which
    may not ever be explicitly referenced.  Most linkers will detect this
    and remove the code causing various unpleasant runtime errors.  These
    macros, adopted from Boost (see force_include.hpp) prevent this
    (C) Copyright 2002 Robert Ramey - http://www.rrsd.com .
    Use, modification and distribution is subject to the Boost Software
    License, Version 1.0. (See accompanying file LICENSE_1_0.txt or copy at
    http://www.boost.org/LICENSE_1_0.txt) */

#if defined(_MSC_VER) && !defined(__clang__)
#   define CEREAL_DLL_EXPORT __declspec(dllexport)
#   define CEREAL_USED
#else // clang or gcc
#   define CEREAL_DLL_EXPORT __attribute__ ((visibility("default")))
#   define CEREAL_USED __attribute__ ((__used__))
#endif

namespace cereal
{
  namespace detail
  {
    //! A static, pre-execution object
    /*! This class will create a single copy (singleton) of some
        type and ensures that merely referencing this type will
        cause it to be instantiated and initialized pre-execution.
        For example, this is used heavily in the polymorphic pointer
        serialization mechanisms to bind various archive types with
        different polymorphic classes */
    template <class T>
    class CEREAL_DLL_EXPORT StaticObject
    {
      private:

        static T & create()
        {
          static T t;
          //! Forces instantiation at pre-execution time
          (void)instance;
          return t;
        }

        StaticObject( StaticObject const & /*other*/ ) {}

      public:
        static T & getInstance()
        {
          return create();
        }

        //! A class that acts like std::lock_guard
        class LockGuard
        {
          #if CEREAL_THREAD_SAFE
          public:
            LockGuard(std::mutex & m) : lock(m) {}
          private:
            std::unique_lock<std::mutex> lock;
          #else
          public:
            LockGuard() = default;
            LockGuard(LockGuard const &) = default; // prevents implicit copy ctor warning
            ~LockGuard() CEREAL_NOEXCEPT {} // prevents variable not used
          #endif
        };

        //! Attempts to lock this static object for the current scope
        /*! @note This function is a no-op if cereal is not compiled with
                  thread safety enabled (CEREAL_THREAD_SAFE = 1).

            This function returns an object that holds a lock for
            this StaticObject that will release its lock upon destruction. This
            call will block until the lock is available. */
        static LockGuard lock()
        {
          #if CEREAL_THREAD_SAFE
          static std::mutex instanceMutex;
          return LockGuard{instanceMutex};
          #else
          return LockGuard{};
          #endif
        }

      private:
        static T & instance;
    };

    template <class T> T & StaticObject<T>::instance = StaticObject<T>::create();
  } // namespace detail
} // namespace cereal

#endif // CEREAL_DETAILS_STATIC_OBJECT_HPP_
```
