## Supplemental code for "The motif composition of variable-number tandem repeats impacts gene expression": traits.hpp

```
/*! \file traits.hpp
    \brief Internal type trait support
    \ingroup Internal */
/*
  Copyright (c) 2014, Randolph Voorhies, Shane Grant
  All rights reserved.

  Redistribution and use in source and binary forms, with or without
  modification, are permitted provided that the following conditions are met:
      * Redistributions of source code must retain the above copyright
        notice, this list of conditions and the following disclaimer.
      * Redistributions in binary form must reproduce the above copyright
        notice, this list of conditions and the following disclaimer in the
        documentation and/or other materials provided with the distribution.
      * Neither the name of cereal nor the
        names of its contributors may be used to endorse or promote products
        derived from this software without specific prior written permission.

  THIS SOFTWARE IS PROVIDED BY THE COPYRIGHT HOLDERS AND CONTRIBUTORS "AS IS" AND
  ANY EXPRESS OR IMPLIED WARRANTIES, INCLUDING, BUT NOT LIMITED TO, THE IMPLIED
  WARRANTIES OF MERCHANTABILITY AND FITNESS FOR A PARTICULAR PURPOSE ARE
  DISCLAIMED. IN NO EVENT SHALL RANDOLPH VOORHIES OR SHANE GRANT BE LIABLE FOR ANY
  DIRECT, INDIRECT, INCIDENTAL, SPECIAL, EXEMPLARY, OR CONSEQUENTIAL DAMAGES
  (INCLUDING, BUT NOT LIMITED TO, PROCUREMENT OF SUBSTITUTE GOODS OR SERVICES;
  LOSS OF USE, DATA, OR PROFITS; OR BUSINESS INTERRUPTION) HOWEVER CAUSED AND
  ON ANY THEORY OF LIABILITY, WHETHER IN CONTRACT, STRICT LIABILITY, OR TORT
  (INCLUDING NEGLIGENCE OR OTHERWISE) ARISING IN ANY WAY OUT OF THE USE OF THIS
  SOFTWARE, EVEN IF ADVISED OF THE POSSIBILITY OF SUCH DAMAGE.
*/
#ifndef CEREAL_DETAILS_TRAITS_HPP_
#define CEREAL_DETAILS_TRAITS_HPP_

#ifndef __clang__
#if (__GNUC__ == 4 && __GNUC_MINOR__ <= 7)
#define CEREAL_OLDER_GCC
#endif // gcc 4.7 or earlier
#endif // __clang__

#include <type_traits>
#include <typeindex>

#include "cereal/macros.hpp"
#include "cereal/access.hpp"

namespace cereal
{
  namespace traits
  {
    using yes = std::true_type;
    using no  = std::false_type;

    namespace detail
    {
      // ######################################################################
      //! Used to delay a static_assert until template instantiation
      template <class T>
      struct delay_static_assert : std::false_type {};

      // ######################################################################
      // SFINAE Helpers
      #ifdef CEREAL_OLDER_GCC // when VS supports better SFINAE, we can use this as the default
      template<typename> struct Void { typedef void type; };
      #endif // CEREAL_OLDER_GCC

      //! Return type for SFINAE Enablers
      enum class sfinae {};

      // ######################################################################
      // Helper functionality for boolean integral constants and Enable/DisableIf
      template <bool H, bool ... T> struct meta_bool_and : std::integral_constant<bool, H && meta_bool_and<T...>::value> {};
      template <bool B> struct meta_bool_and<B> : std::integral_constant<bool, B> {};

      template <bool H, bool ... T> struct meta_bool_or : std::integral_constant<bool, H || meta_bool_or<T...>::value> {};
      template <bool B> struct meta_bool_or<B> : std::integral_constant<bool, B> {};

      // workaround needed due to bug in MSVC 2013, see
      // http://connect.microsoft.com/VisualStudio/feedback/details/800231/c-11-alias-template-issue
      template <bool ... Conditions>
      struct EnableIfHelper : std::enable_if<meta_bool_and<Conditions...>::value, sfinae> {};

      template <bool ... Conditions>
      struct DisableIfHelper : std::enable_if<!meta_bool_or<Conditions...>::value, sfinae> {};
    } // namespace detail

    //! Used as the default value for EnableIf and DisableIf template parameters
    /*! @relates EnableIf
        @relates DisableIf */
    static const detail::sfinae sfinae = {};

    // ######################################################################
    //! Provides a way to enable a function if conditions are met
    /*! This is intended to be used in a near identical fashion to std::enable_if
        while being significantly easier to read at the cost of not allowing for as
        complicated of a condition.

        This will compile (allow the function) if every condition evaluates to true.
        at compile time.  This should be used with SFINAE to ensure that at least
        one other candidate function works when one fails due to an EnableIf.

        This should be used as the las template parameter to a function as
        an unnamed parameter with a default value of cereal::traits::sfinae:

        @code{cpp}
        // using by making the last template argument variadic
        template <class T, EnableIf<std::is_same<T, bool>::value> = sfinae>
        void func(T t );
        @endcode

        Note that this performs a logical AND of all conditions, so you will need
        to construct more complicated requirements with this fact in mind.

        @relates DisableIf
        @relates sfinae
        @tparam Conditions The conditions which will be logically ANDed to enable the function. */
    template <bool ... Conditions>
    using EnableIf = typename detail::EnableIfHelper<Conditions...>::type;

    // ######################################################################
    //! Provides a way to disable a function if conditions are met
    /*! This is intended to be used in a near identical fashion to std::enable_if
        while being significantly easier to read at the cost of not allowing for as
        complicated of a condition.

        This will compile (allow the function) if every condition evaluates to false.
        This should be used with SFINAE to ensure that at least one other candidate
        function works when one fails due to a DisableIf.

        This should be used as the las template parameter to a function as
        an unnamed parameter with a default value of cereal::traits::sfinae:

        @code{cpp}
        // using by making the last template argument variadic
        template <class T, DisableIf<std::is_same<T, bool>::value> = sfinae>
        void func(T t );
        @endcode

        This is often used in conjunction with EnableIf to form an enable/disable pair of
        overloads.

        Note that this performs a logical AND of all conditions, so you will need
        to construct more complicated requirements with this fact in mind.  If all conditions
        hold, the function will be disabled.

        @relates EnableIf
        @relates sfinae
        @tparam Conditions The conditions which will be logically ANDed to disable the function. */
    template <bool ... Conditions>
    using DisableIf = typename detail::DisableIfHelper<Conditions...>::type;

    // ######################################################################
    namespace detail
    {
      template <class InputArchive>
      struct get_output_from_input : no
      {
        static_assert( detail::delay_static_assert<InputArchive>::value,
            "Could not find an associated output archive for input archive." );
      };

      template <class OutputArchive>
      struct get_input_from_output : no
      {
        static_assert( detail::delay_static_assert<OutputArchive>::value,
            "Could not find an associated input archive for output archive." );
      };
    }

    //! Sets up traits that relate an input archive to an output archive
    #define CEREAL_SETUP_ARCHIVE_TRAITS(InputArchive, OutputArchive)  \
    namespace cereal { namespace traits { namespace detail {          \
      template <> struct get_output_from_input<InputArchive>          \
      { using type = OutputArchive; };                                \
      template <> struct get_input_from_output<OutputArchive>         \
      { using type = InputArchive; }; } } } /* end namespaces */

    // ######################################################################
    //! Used to convert a MAKE_HAS_XXX macro into a versioned variant
    #define CEREAL_MAKE_VERSIONED_TEST ,0

    // ######################################################################
    //! Creates a test for whether a non const member function exists
    /*! This creates a class derived from std::integral_constant that will be true if
        the type has the proper member function for the given archive.

        @param name The name of the function to test for (e.g. serialize, load, save)
        @param test_name The name to give the test for the function being tested for (e.g. serialize, versioned_serialize)
        @param versioned Either blank or the macro CEREAL_MAKE_VERSIONED_TEST */
    #ifdef CEREAL_OLDER_GCC
    #define CEREAL_MAKE_HAS_MEMBER_TEST(name, test_name, versioned)                                                                         \
    template <class T, class A, class SFINAE = void>                                                                                        \
    struct has_member_##test_name : no {};                                                                                                  \
    template <class T, class A>                                                                                                             \
    struct has_member_##test_name<T, A,                                                                                                     \
      typename detail::Void< decltype( cereal::access::member_##name( std::declval<A&>(), std::declval<T&>() versioned ) ) >::type> : yes {}
    #else // NOT CEREAL_OLDER_GCC
    #define CEREAL_MAKE_HAS_MEMBER_TEST(name, test_name, versioned)                                                                     \
    namespace detail                                                                                                                    \
    {                                                                                                                                   \
      template <class T, class A>                                                                                                       \
      struct has_member_##name##_##versioned##_impl                                                                                     \
      {                                                                                                                                 \
        template <class TT, class AA>                                                                                                   \
        static auto test(int) -> decltype( cereal::access::member_##name( std::declval<AA&>(), std::declval<TT&>() versioned ), yes()); \
        template <class, class>                                                                                                         \
        static no test(...);                                                                                                            \
        static const bool value = std::is_same<decltype(test<T, A>(0)), yes>::value;                                                    \
      };                                                                                                                                \
    } /* end namespace detail */                                                                                                        \
    template <class T, class A>                                                                                                         \
    struct has_member_##test_name : std::integral_constant<bool, detail::has_member_##name##_##versioned##_impl<T, A>::value> {}
    #endif // NOT CEREAL_OLDER_GCC

    // ######################################################################
    //! Creates a test for whether a non const non-member function exists
    /*! This creates a class derived from std::integral_constant that will be true if
        the type has the proper non-member function for the given archive. */
    #define CEREAL_MAKE_HAS_NON_MEMBER_TEST(test_name, func, versioned)                                                         \
    namespace detail                                                                                                            \
    {                                                                                                                           \
      template <class T, class A>                                                                                               \
      struct has_non_member_##test_name##_impl                                                                                  \
      {                                                                                                                         \
        template <class TT, class AA>                                                                                           \
        static auto test(int) -> decltype( func( std::declval<AA&>(), std::declval<TT&>() versioned ), yes());                  \
        template <class, class>                                                                                                 \
        static no test( ... );                                                                                                  \
        static const bool value = std::is_same<decltype( test<T, A>( 0 ) ), yes>::value;                                        \
      };                                                                                                                        \
    } /* end namespace detail */                                                                                                \
    template <class T, class A>                                                                                                 \
    struct has_non_member_##test_name : std::integral_constant<bool, detail::has_non_member_##test_name##_impl<T, A>::value> {}

    // ######################################################################
    // Member Serialize
    CEREAL_MAKE_HAS_MEMBER_TEST(serialize, serialize,);

    // ######################################################################
    // Member Serialize (versioned)
    CEREAL_MAKE_HAS_MEMBER_TEST(serialize, versioned_serialize, CEREAL_MAKE_VERSIONED_TEST);

    // ######################################################################
    // Non Member Serialize
    CEREAL_MAKE_HAS_NON_MEMBER_TEST(serialize, CEREAL_SERIALIZE_FUNCTION_NAME,);

    // ######################################################################
    // Non Member Serialize (versioned)
    CEREAL_MAKE_HAS_NON_MEMBER_TEST(versioned_serialize, CEREAL_SERIALIZE_FUNCTION_NAME, CEREAL_MAKE_VERSIONED_TEST);

    // ######################################################################
    // Member Load
    CEREAL_MAKE_HAS_MEMBER_TEST(load, load,);

    // ######################################################################
    // Member Load (versioned)
    CEREAL_MAKE_HAS_MEMBER_TEST(load, versioned_load, CEREAL_MAKE_VERSIONED_TEST);

    // ######################################################################
    // Non Member Load
    CEREAL_MAKE_HAS_NON_MEMBER_TEST(load, CEREAL_LOAD_FUNCTION_NAME,);

    // ######################################################################
    // Non Member Load (versioned)
    CEREAL_MAKE_HAS_NON_MEMBER_TEST(versioned_load, CEREAL_LOAD_FUNCTION_NAME, CEREAL_MAKE_VERSIONED_TEST);

    // ######################################################################
    #undef CEREAL_MAKE_HAS_NON_MEMBER_TEST
    #undef CEREAL_MAKE_HAS_MEMBER_TEST

    // ######################################################################
    //! Creates a test for whether a member save function exists
    /*! This creates a class derived from std::integral_constant that will be true if
        the type has the proper member function for the given archive.

        @param test_name The name to give the test (e.g. save or versioned_save)
        @param versioned Either blank or the macro CEREAL_MAKE_VERSIONED_TEST */
    #ifdef CEREAL_OLDER_GCC
    #define CEREAL_MAKE_HAS_MEMBER_SAVE_IMPL(test_name, versioned)                                                                  \
    namespace detail                                                                                                                \
    {                                                                                                                               \
    template <class T, class A>                                                                                                     \
    struct has_member_##test_name##_impl                                                                                            \
      {                                                                                                                             \
        template <class TT, class AA, class SFINAE = void> struct test : no {};                                                     \
        template <class TT, class AA>                                                                                               \
        struct test<TT, AA,                                                                                                         \
          typename detail::Void< decltype( cereal::access::member_save( std::declval<AA&>(),                                        \
                                                                        std::declval<TT const &>() versioned ) ) >::type> : yes {}; \
        static const bool value = test<T, A>();                                                                                     \
                                                                                                                                    \
        template <class TT, class AA, class SFINAE = void> struct test2 : no {};                                                    \
        template <class TT, class AA>                                                                                               \
        struct test2<TT, AA,                                                                                                        \
          typename detail::Void< decltype( cereal::access::member_save_non_const(                                                   \
                                            std::declval<AA&>(),                                                                    \
                                            std::declval<typename std::remove_const<TT>::type&>() versioned ) ) >::type> : yes {};  \
        static const bool not_const_type = test2<T, A>();                                                                           \
      };                                                                                                                            \
    } /* end namespace detail */
    #else /* NOT CEREAL_OLDER_GCC =================================== */
    #define CEREAL_MAKE_HAS_MEMBER_SAVE_IMPL(test_name, versioned)                                                                  \
    namespace detail                                                                                                                \
    {                                                                                                                               \
    template <class T, class A>                                                                                                     \
    struct has_member_##test_name##_impl                                                                                            \
      {                                                                                                                             \
        template <class TT, class AA>                                                                                               \
        static auto test(int) -> decltype( cereal::access::member_save( std::declval<AA&>(),                                        \
                                                                        std::declval<TT const &>() versioned ), yes());             \
        template <class, class> static no test(...);                                                                                \
        static const bool value = std::is_same<decltype(test<T, A>(0)), yes>::value;                                                \
                                                                                                                                    \
        template <class TT, class AA>                                                                                               \
        static auto test2(int) -> decltype( cereal::access::member_save_non_const(                                                  \
                                              std::declval<AA &>(),                                                                 \
                                              std::declval<typename std::remove_const<TT>::type&>() versioned ), yes());            \
        template <class, class> static no test2(...);                                                                               \
        static const bool not_const_type = std::is_same<decltype(test2<T, A>(0)), yes>::value;                                      \
      };                                                                                                                            \
    } /* end namespace detail */
    #endif /* NOT CEREAL_OLDER_GCC */

    // ######################################################################
    // Member Save
    CEREAL_MAKE_HAS_MEMBER_SAVE_IMPL(save, )

    template <class T, class A>
    struct has_member_save : std::integral_constant<bool, detail::has_member_save_impl<T, A>::value>
    {
      typedef typename detail::has_member_save_impl<T, A> check;
      static_assert( check::value || !check::not_const_type,
        "cereal detected a non-const save. \n "
        "save member functions must always be const" );
    };

    // ######################################################################
    // Member Save (versioned)
    CEREAL_MAKE_HAS_MEMBER_SAVE_IMPL(versioned_save, CEREAL_MAKE_VERSIONED_TEST)

    template <class T, class A>
    struct has_member_versioned_save : std::integral_constant<bool, detail::has_member_versioned_save_impl<T, A>::value>
    {
      typedef typename detail::has_member_versioned_save_impl<T, A> check;
      static_assert( check::value || !check::not_const_type,
        "cereal detected a versioned non-const save. \n "
        "save member functions must always be const" );
    };

    // ######################################################################
    #undef CEREAL_MAKE_HAS_MEMBER_SAVE_IMPL

    // ######################################################################
    //! Creates a test for whether a non-member save function exists
    /*! This creates a class derived from std::integral_constant that will be true if
        the type has the proper non-member function for the given archive.

        @param test_name The name to give the test (e.g. save or versioned_save)
        @param versioned Either blank or the macro CEREAL_MAKE_VERSIONED_TEST */
    #define CEREAL_MAKE_HAS_NON_MEMBER_SAVE_TEST(test_name, versioned)                                                       \
    namespace detail                                                                                                         \
    {                                                                                                                        \
      template <class T, class A>                                                                                            \
      struct has_non_member_##test_name##_impl                                                                               \
      {                                                                                                                      \
        template <class TT, class AA>                                                                                        \
        static auto test(int) -> decltype( CEREAL_SAVE_FUNCTION_NAME(                                                        \
                                              std::declval<AA&>(),                                                           \
                                              std::declval<TT const &>() versioned ), yes());                                \
        template <class, class> static no test(...);                                                                         \
        static const bool value = std::is_same<decltype(test<T, A>(0)), yes>::value;                                         \
                                                                                                                             \
        template <class TT, class AA>                                                                                        \
        static auto test2(int) -> decltype( CEREAL_SAVE_FUNCTION_NAME(                                                       \
                                              std::declval<AA &>(),                                                          \
                                              std::declval<typename std::remove_const<TT>::type&>() versioned ), yes());     \
        template <class, class> static no test2(...);                                                                        \
        static const bool not_const_type = std::is_same<decltype(test2<T, A>(0)), yes>::value;                               \
      };                                                                                                                     \
    } /* end namespace detail */                                                                                             \
                                                                                                                             \
    template <class T, class A>                                                                                              \
    struct has_non_member_##test_name : std::integral_constant<bool, detail::has_non_member_##test_name##_impl<T, A>::value> \
    {                                                                                                                        \
      using check = typename detail::has_non_member_##test_name##_impl<T, A>;                                                \
      static_assert( check::value || !check::not_const_type,                                                                 \
        "cereal detected a non-const type parameter in non-member " #test_name ". \n "                                       \
        #test_name " non-member functions must always pass their types as const" );                                          \
    };

    // ######################################################################
    // Non Member Save
    CEREAL_MAKE_HAS_NON_MEMBER_SAVE_TEST(save, )

    // ######################################################################
    // Non Member Save (versioned)
    CEREAL_MAKE_HAS_NON_MEMBER_SAVE_TEST(versioned_save, CEREAL_MAKE_VERSIONED_TEST)

    // ######################################################################
    #undef CEREAL_MAKE_HAS_NON_MEMBER_SAVE_TEST

    // ######################################################################
    // Minimal Utilities
    namespace detail
    {
      // Determines if the provided type is an std::string
      template <class> struct is_string : std::false_type {};

      template <class CharT, class Traits, class Alloc>
      struct is_string<std::basic_string<CharT, Traits, Alloc>> : std::true_type {};
    }

    // Determines if the type is valid for use with a minimal serialize function
    template <class T>
    struct is_minimal_type : std::integral_constant<bool,
      detail::is_string<T>::value || std::is_arithmetic<T>::value> {};

    // ######################################################################
    //! Creates implementation details for whether a member save_minimal function exists
    /*! This creates a class derived from std::integral_constant that will be true if
        the type has the proper member function for the given archive.

        @param test_name The name to give the test (e.g. save_minimal or versioned_save_minimal)
        @param versioned Either blank or the macro CEREAL_MAKE_VERSIONED_TEST */
    #ifdef CEREAL_OLDER_GCC
    #define CEREAL_MAKE_HAS_MEMBER_SAVE_MINIMAL_IMPL(test_name, versioned)                                                                        \
    namespace detail                                                                                                                              \
    {                                                                                                                                             \
      template <class T, class A>                                                                                                                 \
      struct has_member_##test_name##_impl                                                                                                        \
      {                                                                                                                                           \
        template <class TT, class AA, class SFINAE = void> struct test : no {};                                                                   \
        template <class TT, class AA>                                                                                                             \
        struct test<TT, AA, typename detail::Void< decltype(                                                                                      \
            cereal::access::member_save_minimal( std::declval<AA const &>(),                                                                      \
                                                 std::declval<TT const &>() versioned ) ) >::type> : yes {};                                      \
                                                                                                                                                  \
        static const bool value = test<T, A>();                                                                                                   \
                                                                                                                                                  \
        template <class TT, class AA, class SFINAE = void> struct test2 : no {};                                                                  \
        template <class TT, class AA>                                                                                                             \
        struct test2<TT, AA, typename detail::Void< decltype(                                                                                     \
            cereal::access::member_save_minimal_non_const( std::declval<AA const &>(),                                                            \
                                                           std::declval<typename std::remove_const<TT>::type&>() versioned ) ) >::type> : yes {}; \
        static const bool not_const_type = test2<T, A>();                                                                                         \
                                                                                                                                                  \
        static const bool valid = value || !not_const_type;                                                                                       \
      };                                                                                                                                          \
    } /* end namespace detail */
    #else /* NOT CEREAL_OLDER_GCC =================================== */
    #define CEREAL_MAKE_HAS_MEMBER_SAVE_MINIMAL_IMPL(test_name, versioned)                     \
    namespace detail                                                                           \
    {                                                                                          \
      template <class T, class A>                                                              \
      struct has_member_##test_name##_impl                                                     \
      {                                                                                        \
        template <class TT, class AA>                                                          \
        static auto test(int) -> decltype( cereal::access::member_save_minimal(                \
              std::declval<AA const &>(),                                                      \
              std::declval<TT const &>() versioned ), yes());                                  \
        template <class, class> static no test(...);                                           \
        static const bool value = std::is_same<decltype(test<T, A>(0)), yes>::value;           \
                                                                                               \
        template <class TT, class AA>                                                          \
        static auto test2(int) -> decltype( cereal::access::member_save_minimal_non_const(     \
              std::declval<AA const &>(),                                                      \
              std::declval<typename std::remove_const<TT>::type&>() versioned ), yes());       \
        template <class, class> static no test2(...);                                          \
        static const bool not_const_type = std::is_same<decltype(test2<T, A>(0)), yes>::value; \
                                                                                               \
        static const bool valid = value || !not_const_type;                                    \
      };                                                                                       \
    } /* end namespace detail */
    #endif // NOT CEREAL_OLDER_GCC

    // ######################################################################
    //! Creates helpers for minimal save functions
    /*! The get_member_*_type structs allow access to the return type of a save_minimal,
        assuming that the function actually exists.  If the function does not
        exist, the type will be void.

        @param test_name The name to give the test (e.g. save_minimal or versioned_save_minimal)
        @param versioned Either blank or the macro CEREAL_MAKE_VERSIONED_TEST */
    #define CEREAL_MAKE_HAS_MEMBER_SAVE_MINIMAL_HELPERS_IMPL(test_name, versioned)                           \
    namespace detail                                                                                         \
    {                                                                                                        \
      template <class T, class A, bool Valid>                                                                \
      struct get_member_##test_name##_type { using type = void; };                                           \
                                                                                                             \
      template <class T, class A>                                                                            \
      struct get_member_##test_name##_type<T, A, true>                                                       \
      {                                                                                                      \
        using type = decltype( cereal::access::member_save_minimal( std::declval<A const &>(),               \
                                                                    std::declval<T const &>() versioned ) ); \
      };                                                                                                     \
    } /* end namespace detail */

    // ######################################################################
    //! Creates a test for whether a member save_minimal function exists
    /*! This creates a class derived from std::integral_constant that will be true if
        the type has the proper member function for the given archive.

        @param test_name The name to give the test (e.g. save_minimal or versioned_save_minimal) */
    #define CEREAL_MAKE_HAS_MEMBER_SAVE_MINIMAL_TEST(test_name)                                                      \
    template <class T, class A>                                                                                      \
    struct has_member_##test_name : std::integral_constant<bool, detail::has_member_##test_name##_impl<T, A>::value> \
    {                                                                                                                \
      using check = typename detail::has_member_##test_name##_impl<T, A>;                                            \
      static_assert( check::valid,                                                                                   \
        "cereal detected a non-const member " #test_name ". \n "                                                     \
        #test_name " member functions must always be const" );                                                       \
                                                                                                                     \
      using type = typename detail::get_member_##test_name##_type<T, A, check::value>::type;                         \
      static_assert( (check::value && is_minimal_type<type>::value) || !check::value,                                \
        "cereal detected a member " #test_name " with an invalid return type. \n "                                   \
        "return type must be arithmetic or string" );                                                                \
    };

    // ######################################################################
    // Member Save Minimal
    CEREAL_MAKE_HAS_MEMBER_SAVE_MINIMAL_IMPL(save_minimal, )
    CEREAL_MAKE_HAS_MEMBER_SAVE_MINIMAL_HELPERS_IMPL(save_minimal, )
    CEREAL_MAKE_HAS_MEMBER_SAVE_MINIMAL_TEST(save_minimal)

    // ######################################################################
    // Member Save Minimal (versioned)
    CEREAL_MAKE_HAS_MEMBER_SAVE_MINIMAL_IMPL(versioned_save_minimal, CEREAL_MAKE_VERSIONED_TEST)
    CEREAL_MAKE_HAS_MEMBER_SAVE_MINIMAL_HELPERS_IMPL(versioned_save_minimal, CEREAL_MAKE_VERSIONED_TEST)
    CEREAL_MAKE_HAS_MEMBER_SAVE_MINIMAL_TEST(versioned_save_minimal)

    // ######################################################################
    #undef CEREAL_MAKE_HAS_MEMBER_SAVE_MINIMAL_IMPL
    #undef CEREAL_MAKE_HAS_MEMBER_SAVE_MINIMAL_HELPERS_IMPL
    #undef CEREAL_MAKE_HAS_MEMBER_SAVE_MINIMAL_TEST

    // ######################################################################
    //! Creates a test for whether a non-member save_minimal function exists
    /*! This creates a class derived from std::integral_constant that will be true if
        the type has the proper member function for the given archive.

        @param test_name The name to give the test (e.g. save_minimal or versioned_save_minimal)
        @param versioned Either blank or the macro CEREAL_MAKE_VERSIONED_TEST */
    #define CEREAL_MAKE_HAS_NON_MEMBER_SAVE_MINIMAL_TEST(test_name, versioned)                                               \
    namespace detail                                                                                                         \
    {                                                                                                                        \
      template <class T, class A>                                                                                            \
      struct has_non_member_##test_name##_impl                                                                               \
      {                                                                                                                      \
        template <class TT, class AA>                                                                                        \
        static auto test(int) -> decltype( CEREAL_SAVE_MINIMAL_FUNCTION_NAME(                                                \
              std::declval<AA const &>(),                                                                                    \
              std::declval<TT const &>() versioned ), yes());                                                                \
        template <class, class> static no test(...);                                                                         \
        static const bool value = std::is_same<decltype(test<T, A>(0)), yes>::value;                                         \
                                                                                                                             \
        template <class TT, class AA>                                                                                        \
        static auto test2(int) -> decltype( CEREAL_SAVE_MINIMAL_FUNCTION_NAME(                                               \
              std::declval<AA const &>(),                                                                                    \
              std::declval<typename std::remove_const<TT>::type&>() versioned ), yes());                                     \
        template <class, class> static no test2(...);                                                                        \
        static const bool not_const_type = std::is_same<decltype(test2<T, A>(0)), yes>::value;                               \
                                                                                                                             \
        static const bool valid = value || !not_const_type;                                                                  \
      };                                                                                                                     \
                                                                                                                             \
      template <class T, class A, bool Valid>                                                                                \
      struct get_non_member_##test_name##_type { using type = void; };                                                       \
                                                                                                                             \
      template <class T, class A>                                                                                            \
      struct get_non_member_##test_name##_type <T, A, true>                                                                  \
      {                                                                                                                      \
        using type = decltype( CEREAL_SAVE_MINIMAL_FUNCTION_NAME( std::declval<A const &>(),                                 \
                                                                  std::declval<T const &>() versioned ) );                   \
      };                                                                                                                     \
    } /* end namespace detail */                                                                                             \
                                                                                                                             \
    template <class T, class A>                                                                                              \
    struct has_non_member_##test_name : std::integral_constant<bool, detail::has_non_member_##test_name##_impl<T, A>::value> \
    {                                                                                                                        \
      using check = typename detail::has_non_member_##test_name##_impl<T, A>;                                                \
      static_assert( check::valid,                                                                                           \
        "cereal detected a non-const type parameter in non-member " #test_name ". \n "                                       \
        #test_name " non-member functions must always pass their types as const" );                                          \
                                                                                                                             \
      using type = typename detail::get_non_member_##test_name##_type<T, A, check::value>::type;                             \
      static_assert( (check::value && is_minimal_type<type>::value) || !check::value,                                        \
        "cereal detected a non-member " #test_name " with an invalid return type. \n "                                       \
        "return type must be arithmetic or string" );                                                                        \
    };

    // ######################################################################
    // Non-Member Save Minimal
    CEREAL_MAKE_HAS_NON_MEMBER_SAVE_MINIMAL_TEST(save_minimal, )

    // ######################################################################
    // Non-Member Save Minimal (versioned)
    CEREAL_MAKE_HAS_NON_MEMBER_SAVE_MINIMAL_TEST(versioned_save_minimal, CEREAL_MAKE_VERSIONED_TEST)

    // ######################################################################
    #undef CEREAL_MAKE_HAS_NON_MEMBER_SAVE_MINIMAL_TEST

    // ######################################################################
    // Load Minimal Utilities
    namespace detail
    {
      //! Used to help strip away conversion wrappers
      /*! If someone writes a non-member load/save minimal function that accepts its
          parameter as some generic template type and needs to perform trait checks
          on that type, our NoConvert wrappers will interfere with this.  Using
          the struct strip_minmal, users can strip away our wrappers to get to
          the underlying type, allowing traits to work properly */
      struct NoConvertBase {};

      //! A struct that prevents implicit conversion
      /*! Any type instantiated with this struct will be unable to implicitly convert
          to another type.  Is designed to only allow conversion to Source const &.

          @tparam Source the type of the original source */
      template <class Source>
      struct NoConvertConstRef : NoConvertBase
      {
        using type = Source; //!< Used to get underlying type easily

        template <class Dest, class = typename std::enable_if<std::is_same<Source, Dest>::value>::type>
        operator Dest () = delete;

        //! only allow conversion if the types are the same and we are converting into a const reference
        template <class Dest, class = typename std::enable_if<std::is_same<Source, Dest>::value>::type>
        operator Dest const & ();
      };

      //! A struct that prevents implicit conversion
      /*! Any type instantiated with this struct will be unable to implicitly convert
          to another type.  Is designed to only allow conversion to Source &.

          @tparam Source the type of the original source */
      template <class Source>
      struct NoConvertRef : NoConvertBase
      {
        using type = Source; //!< Used to get underlying type easily

        template <class Dest, class = typename std::enable_if<std::is_same<Source, Dest>::value>::type>
        operator Dest () = delete;

        #ifdef __clang__
        template <class Dest, class = typename std::enable_if<std::is_same<Source, Dest>::value>::type>
        operator Dest const & () = delete;
        #endif // __clang__

        //! only allow conversion if the types are the same and we are converting into a const reference
        template <class Dest, class = typename std::enable_if<std::is_same<Source, Dest>::value>::type>
        operator Dest & ();
      };

      //! A type that can implicitly convert to anything else
      struct AnyConvert
      {
        template <class Dest>
        operator Dest & ();

        template <class Dest>
        operator Dest const & () const;
      };
    } // namespace detail

    // ######################################################################
    //! Creates a test for whether a member load_minimal function exists
    /*! This creates a class derived from std::integral_constant that will be true if
        the type has the proper member function for the given archive.

        Our strategy here is to first check if a function matching the signature more or less exists
        (allow anything like load_minimal(xxx) using AnyConvert, and then secondly enforce
        that it has the correct signature using NoConvertConstRef

        @param test_name The name to give the test (e.g. load_minimal or versioned_load_minimal)
        @param versioned Either blank or the macro CEREAL_MAKE_VERSIONED_TEST */
    #ifdef CEREAL_OLDER_GCC
    #define CEREAL_MAKE_HAS_MEMBER_LOAD_MINIMAL_IMPL(test_name, versioned)                                                    \
    namespace detail                                                                                                          \
    {                                                                                                                         \
      template <class T, class A, class SFINAE = void> struct has_member_##test_name##_impl : no {};                          \
      template <class T, class A>                                                                                             \
      struct has_member_##test_name##_impl<T, A, typename detail::Void< decltype(                                             \
          cereal::access::member_load_minimal( std::declval<A const &>(),                                                     \
                                               std::declval<T &>(), AnyConvert() versioned ) ) >::type> : yes {};             \
                                                                                                                              \
        template <class T, class A, class U, class SFINAE = void> struct has_member_##test_name##_type_impl : no {};          \
        template <class T, class A, class U>                                                                                  \
        struct has_member_##test_name##_type_impl<T, A, U, typename detail::Void< decltype(                                   \
            cereal::access::member_load_minimal( std::declval<A const &>(),                                                   \
                                                 std::declval<T &>(), NoConvertConstRef<U>() versioned ) ) >::type> : yes {}; \
    } /* end namespace detail */
    #else /* NOT CEREAL_OLDER_GCC =================================== */
    #define CEREAL_MAKE_HAS_MEMBER_LOAD_MINIMAL_IMPL(test_name, versioned)              \
    namespace detail                                                                    \
    {                                                                                   \
      template <class T, class A>                                                       \
      struct has_member_##test_name##_impl                                              \
      {                                                                                 \
        template <class TT, class AA>                                                   \
        static auto test(int) -> decltype( cereal::access::member_load_minimal(         \
              std::declval<AA const &>(),                                               \
              std::declval<TT &>(), AnyConvert() versioned ), yes());                   \
        template <class, class> static no test(...);                                    \
        static const bool value = std::is_same<decltype(test<T, A>(0)), yes>::value;    \
      };                                                                                \
      template <class T, class A, class U>                                              \
      struct has_member_##test_name##_type_impl                                         \
      {                                                                                 \
        template <class TT, class AA, class UU>                                         \
        static auto test(int) -> decltype( cereal::access::member_load_minimal(         \
              std::declval<AA const &>(),                                               \
              std::declval<TT &>(), NoConvertConstRef<UU>() versioned ), yes());        \
        template <class, class, class> static no test(...);                             \
        static const bool value = std::is_same<decltype(test<T, A, U>(0)), yes>::value; \
                                                                                        \
      };                                                                                \
    } /* end namespace detail */
    #endif // NOT CEREAL_OLDER_GCC

    // ######################################################################
    //! Creates helpers for minimal load functions
    /*! The has_member_*_wrapper structs ensure that the load and save types for the
        requested function type match appropriately.

        @param load_test_name The name to give the test (e.g. load_minimal or versioned_load_minimal)
        @param save_test_name The name to give the test (e.g. save_minimal or versioned_save_minimal,
                              should match the load name.
        @param save_test_prefix The name to give the test (e.g. save_minimal or versioned_save_minimal,
                              should match the load name, without the trailing "_minimal" (e.g.
                              save or versioned_save).  Needed because the preprocessor is an abomination.
        @param versioned Either blank or the macro CEREAL_MAKE_VERSIONED_TEST */
    #define CEREAL_MAKE_HAS_MEMBER_LOAD_MINIMAL_HELPERS_IMPL(load_test_name, save_test_name, save_test_prefix, versioned) \
    namespace detail                                                                                                      \
    {                                                                                                                     \
      template <class T, class A, bool Valid>                                                                             \
      struct has_member_##load_test_name##_wrapper : std::false_type {};                                                  \
                                                                                                                          \
      template <class T, class A>                                                                                         \
      struct has_member_##load_test_name##_wrapper<T, A, true>                                                            \
      {                                                                                                                   \
        using AOut = typename detail::get_output_from_input<A>::type;                                                     \
                                                                                                                          \
        static_assert( has_member_##save_test_prefix##_minimal<T, AOut>::value,                                           \
          "cereal detected member " #load_test_name " but no valid member " #save_test_name ". \n "                       \
          "cannot evaluate correctness of " #load_test_name " without valid " #save_test_name "." );                      \
                                                                                                                          \
        using SaveType = typename detail::get_member_##save_test_prefix##_minimal_type<T, AOut, true>::type;              \
        const static bool value = has_member_##load_test_name##_impl<T, A>::value;                                        \
        const static bool valid = has_member_##load_test_name##_type_impl<T, A, SaveType>::value;                         \
                                                                                                                          \
        static_assert( valid || !value, "cereal detected different or invalid types in corresponding member "             \
            #load_test_name " and " #save_test_name " functions. \n "                                                     \
            "the paramater to " #load_test_name " must be a constant reference to the type that "                         \
            #save_test_name " returns." );                                                                                \
      };                                                                                                                  \
    } /* end namespace detail */

    // ######################################################################
    //! Creates a test for whether a member load_minimal function exists
    /*! This creates a class derived from std::integral_constant that will be true if
        the type has the proper member function for the given archive.

        @param load_test_name The name to give the test (e.g. load_minimal or versioned_load_minimal)
        @param load_test_prefix The above parameter minus the trailing "_minimal" */
    #define CEREAL_MAKE_HAS_MEMBER_LOAD_MINIMAL_TEST(load_test_name, load_test_prefix)                                         \
    template <class T, class A>                                                                                                \
    struct has_member_##load_test_prefix##_minimal : std::integral_constant<bool,                                              \
      detail::has_member_##load_test_name##_wrapper<T, A, detail::has_member_##load_test_name##_impl<T, A>::value>::value> {};

    // ######################################################################
    // Member Load Minimal
    CEREAL_MAKE_HAS_MEMBER_LOAD_MINIMAL_IMPL(load_minimal, )
    CEREAL_MAKE_HAS_MEMBER_LOAD_MINIMAL_HELPERS_IMPL(load_minimal, save_minimal, save, )
    CEREAL_MAKE_HAS_MEMBER_LOAD_MINIMAL_TEST(load_minimal, load)

    // ######################################################################
    // Member Load Minimal (versioned)
    CEREAL_MAKE_HAS_MEMBER_LOAD_MINIMAL_IMPL(versioned_load_minimal, CEREAL_MAKE_VERSIONED_TEST)
    CEREAL_MAKE_HAS_MEMBER_LOAD_MINIMAL_HELPERS_IMPL(versioned_load_minimal, versioned_save_minimal, versioned_save, CEREAL_MAKE_VERSIONED_TEST)
    CEREAL_MAKE_HAS_MEMBER_LOAD_MINIMAL_TEST(versioned_load_minimal, versioned_load)

    // ######################################################################
    #undef CEREAL_MAKE_HAS_MEMBER_LOAD_MINIMAL_IMPL
    #undef CEREAL_MAKE_HAS_MEMBER_LOAD_MINIMAL_HELPERS_IMPL
    #undef CEREAL_MAKE_HAS_MEMBER_LOAD_MINIMAL_TEST

    // ######################################################################
    // Non-Member Load Minimal
    namespace detail
    {
      #ifdef CEREAL_OLDER_GCC
      void CEREAL_LOAD_MINIMAL_FUNCTION_NAME(); // prevents nonsense complaining about not finding this
      void CEREAL_SAVE_MINIMAL_FUNCTION_NAME();
      #endif // CEREAL_OLDER_GCC
    } // namespace detail

    // ######################################################################
    //! Creates a test for whether a non-member load_minimal function exists
    /*! This creates a class derived from std::integral_constant that will be true if
        the type has the proper member function for the given archive.

        See notes from member load_minimal implementation.

        Note that there should be an additional const check on load_minimal after the valid check,
        but this currently interferes with many valid uses of minimal serialization.  It has been
        removed (see #565 on github) and previously was:

        @code
        static_assert( check::const_valid || !check::exists,
            "cereal detected an invalid serialization type parameter in non-member " #test_name ".  "
            #test_name " non-member functions must accept their serialization type by non-const reference" );
        @endcode

        See #132, #436, #263, and #565 on https://github.com/USCiLab/cereal for more details.

        @param test_name The name to give the test (e.g. load_minimal or versioned_load_minimal)
        @param save_name The corresponding name the save test would have (e.g. save_minimal or versioned_save_minimal)
        @param versioned Either blank or the macro CEREAL_MAKE_VERSIONED_TEST */
    #define CEREAL_MAKE_HAS_NON_MEMBER_LOAD_MINIMAL_TEST(test_name, save_name, versioned)                                    \
    namespace detail                                                                                                         \
    {                                                                                                                        \
      template <class T, class A, class U = void>                                                                            \
      struct has_non_member_##test_name##_impl                                                                               \
      {                                                                                                                      \
        template <class TT, class AA>                                                                                        \
        static auto test(int) -> decltype( CEREAL_LOAD_MINIMAL_FUNCTION_NAME(                                                \
              std::declval<AA const &>(), std::declval<TT&>(), AnyConvert() versioned ), yes() );                            \
        template <class, class> static no test( ... );                                                                       \
        static const bool exists = std::is_same<decltype( test<T, A>( 0 ) ), yes>::value;                                    \
                                                                                                                             \
        template <class TT, class AA, class UU>                                                                              \
        static auto test2(int) -> decltype( CEREAL_LOAD_MINIMAL_FUNCTION_NAME(                                               \
              std::declval<AA const &>(), std::declval<TT&>(), NoConvertConstRef<UU>() versioned ), yes() );                 \
        template <class, class, class> static no test2( ... );                                                               \
        static const bool valid = std::is_same<decltype( test2<T, A, U>( 0 ) ), yes>::value;                                 \
                                                                                                                             \
        template <class TT, class AA>                                                                                        \
        static auto test3(int) -> decltype( CEREAL_LOAD_MINIMAL_FUNCTION_NAME(                                               \
              std::declval<AA const &>(), NoConvertRef<TT>(), AnyConvert() versioned ), yes() );                             \
        template <class, class> static no test3( ... );                                                                      \
        static const bool const_valid = std::is_same<decltype( test3<T, A>( 0 ) ), yes>::value;                              \
      };                                                                                                                     \
                                                                                                                             \
      template <class T, class A, bool Valid>                                                                                \
      struct has_non_member_##test_name##_wrapper : std::false_type {};                                                      \
                                                                                                                             \
      template <class T, class A>                                                                                            \
      struct has_non_member_##test_name##_wrapper<T, A, true>                                                                \
      {                                                                                                                      \
        using AOut = typename detail::get_output_from_input<A>::type;                                                        \
                                                                                                                             \
        static_assert( detail::has_non_member_##save_name##_impl<T, AOut>::valid,                                            \
          "cereal detected non-member " #test_name " but no valid non-member " #save_name ". \n "                            \
          "cannot evaluate correctness of " #test_name " without valid " #save_name "." );                                   \
                                                                                                                             \
        using SaveType = typename detail::get_non_member_##save_name##_type<T, AOut, true>::type;                            \
        using check = has_non_member_##test_name##_impl<T, A, SaveType>;                                                     \
        static const bool value = check::exists;                                                                             \
                                                                                                                             \
        static_assert( check::valid || !check::exists, "cereal detected different types in corresponding non-member "        \
            #test_name " and " #save_name " functions. \n "                                                                  \
            "the paramater to " #test_name " must be a constant reference to the type that " #save_name " returns." );       \
      };                                                                                                                     \
    } /* namespace detail */                                                                                                 \
                                                                                                                             \
    template <class T, class A>                                                                                              \
    struct has_non_member_##test_name : std::integral_constant<bool,                                                         \
      detail::has_non_member_##test_name##_wrapper<T, A, detail::has_non_member_##test_name##_impl<T, A>::exists>::value> {};

    // ######################################################################
    // Non-Member Load Minimal
    CEREAL_MAKE_HAS_NON_MEMBER_LOAD_MINIMAL_TEST(load_minimal, save_minimal, )

    // ######################################################################
    // Non-Member Load Minimal (versioned)
    CEREAL_MAKE_HAS_NON_MEMBER_LOAD_MINIMAL_TEST(versioned_load_minimal, versioned_save_minimal, CEREAL_MAKE_VERSIONED_TEST)

    // ######################################################################
    #undef CEREAL_MAKE_HAS_NON_MEMBER_LOAD_MINIMAL_TEST

    // ######################################################################
    namespace detail
    {
      // const stripped away before reaching here, prevents errors on conversion from
      // construct<const T> to construct<T>
      template<typename T, typename A>
      struct has_member_load_and_construct_impl : std::integral_constant<bool,
        std::is_same<decltype( access::load_and_construct<T>( std::declval<A&>(), std::declval< ::cereal::construct<T>&>() ) ), void>::value>
      { };

      template<typename T, typename A>
      struct has_member_versioned_load_and_construct_impl : std::integral_constant<bool,
        std::is_same<decltype( access::load_and_construct<T>( std::declval<A&>(), std::declval< ::cereal::construct<T>&>(), 0 ) ), void>::value>
      { };
    } // namespace detail

    //! Member load and construct check
    template<typename T, typename A>
    struct has_member_load_and_construct : detail::has_member_load_and_construct_impl<typename std::remove_const<T>::type, A>
    { };

    //! Member load and construct check (versioned)
    template<typename T, typename A>
    struct has_member_versioned_load_and_construct : detail::has_member_versioned_load_and_construct_impl<typename std::remove_const<T>::type, A>
    { };

    // ######################################################################
    //! Creates a test for whether a non-member load_and_construct specialization exists
    /*! This creates a class derived from std::integral_constant that will be true if
        the type has the proper non-member function for the given archive. */
    #define CEREAL_MAKE_HAS_NON_MEMBER_LOAD_AND_CONSTRUCT_TEST(test_name, versioned)                                            \
    namespace detail                                                                                                            \
    {                                                                                                                           \
      template <class T, class A>                                                                                               \
      struct has_non_member_##test_name##_impl                                                                                  \
      {                                                                                                                         \
        template <class TT, class AA>                                                                                           \
        static auto test(int) -> decltype( LoadAndConstruct<TT>::load_and_construct(                                            \
                                           std::declval<AA&>(), std::declval< ::cereal::construct<TT>&>() versioned ), yes());  \
        template <class, class>                                                                                                 \
        static no test( ... );                                                                                                  \
        static const bool value = std::is_same<decltype( test<T, A>( 0 ) ), yes>::value;                                        \
      };                                                                                                                        \
    } /* end namespace detail */                                                                                                \
    template <class T, class A>                                                                                                 \
    struct has_non_member_##test_name :                                                                                         \
      std::integral_constant<bool, detail::has_non_member_##test_name##_impl<typename std::remove_const<T>::type, A>::value> {};

    // ######################################################################
    //! Non member load and construct check
    CEREAL_MAKE_HAS_NON_MEMBER_LOAD_AND_CONSTRUCT_TEST(load_and_construct, )

    // ######################################################################
    //! Non member load and construct check (versioned)
    CEREAL_MAKE_HAS_NON_MEMBER_LOAD_AND_CONSTRUCT_TEST(versioned_load_and_construct, CEREAL_MAKE_VERSIONED_TEST)

    // ######################################################################
    //! Has either a member or non member load and construct
    template<typename T, typename A>
    struct has_load_and_construct : std::integral_constant<bool,
      has_member_load_and_construct<T, A>::value || has_non_member_load_and_construct<T, A>::value ||
      has_member_versioned_load_and_construct<T, A>::value || has_non_member_versioned_load_and_construct<T, A>::value>
    { };

    // ######################################################################
    #undef CEREAL_MAKE_HAS_NON_MEMBER_LOAD_AND_CONSTRUCT_TEST

    // ######################################################################
    // End of serialization existence tests
    #undef CEREAL_MAKE_VERSIONED_TEST

    // ######################################################################
    template <class T, class InputArchive, class OutputArchive>
    struct has_member_split : std::integral_constant<bool,
      (has_member_load<T, InputArchive>::value && has_member_save<T, OutputArchive>::value) ||
      (has_member_versioned_load<T, InputArchive>::value && has_member_versioned_save<T, OutputArchive>::value)> {};

    // ######################################################################
    template <class T, class InputArchive, class OutputArchive>
    struct has_non_member_split : std::integral_constant<bool,
      (has_non_member_load<T, InputArchive>::value && has_non_member_save<T, OutputArchive>::value) ||
      (has_non_member_versioned_load<T, InputArchive>::value && has_non_member_versioned_save<T, OutputArchive>::value)> {};

    // ######################################################################
    template <class T, class OutputArchive>
    struct has_invalid_output_versioning : std::integral_constant<bool,
      (has_member_versioned_save<T, OutputArchive>::value && has_member_save<T, OutputArchive>::value) ||
      (has_non_member_versioned_save<T, OutputArchive>::value && has_non_member_save<T, OutputArchive>::value) ||
      (has_member_versioned_serialize<T, OutputArchive>::value && has_member_serialize<T, OutputArchive>::value) ||
      (has_non_member_versioned_serialize<T, OutputArchive>::value && has_non_member_serialize<T, OutputArchive>::value) ||
      (has_member_versioned_save_minimal<T, OutputArchive>::value && has_member_save_minimal<T, OutputArchive>::value) ||
      (has_non_member_versioned_save_minimal<T, OutputArchive>::value &&  has_non_member_save_minimal<T, OutputArchive>::value)> {};

    // ######################################################################
    template <class T, class InputArchive>
    struct has_invalid_input_versioning : std::integral_constant<bool,
      (has_member_versioned_load<T, InputArchive>::value && has_member_load<T, InputArchive>::value) ||
      (has_non_member_versioned_load<T, InputArchive>::value && has_non_member_load<T, InputArchive>::value) ||
      (has_member_versioned_serialize<T, InputArchive>::value && has_member_serialize<T, InputArchive>::value) ||
      (has_non_member_versioned_serialize<T, InputArchive>::value && has_non_member_serialize<T, InputArchive>::value) ||
      (has_member_versioned_load_minimal<T, InputArchive>::value && has_member_load_minimal<T, InputArchive>::value) ||
      (has_non_member_versioned_load_minimal<T, InputArchive>::value &&  has_non_member_load_minimal<T, InputArchive>::value)> {};

    // ######################################################################
    namespace detail
    {
      //! Create a test for a cereal::specialization entry
      #define CEREAL_MAKE_IS_SPECIALIZED_IMPL(name)                                          \
      template <class T, class A>                                                            \
      struct is_specialized_##name : std::integral_constant<bool,                            \
        !std::is_base_of<std::false_type, specialize<A, T, specialization::name>>::value> {}

      CEREAL_MAKE_IS_SPECIALIZED_IMPL(member_serialize);
      CEREAL_MAKE_IS_SPECIALIZED_IMPL(member_load_save);
      CEREAL_MAKE_IS_SPECIALIZED_IMPL(member_load_save_minimal);
      CEREAL_MAKE_IS_SPECIALIZED_IMPL(non_member_serialize);
      CEREAL_MAKE_IS_SPECIALIZED_IMPL(non_member_load_save);
      CEREAL_MAKE_IS_SPECIALIZED_IMPL(non_member_load_save_minimal);

      #undef CEREAL_MAKE_IS_SPECIALIZED_IMPL

      //! Number of specializations detected
      template <class T, class A>
      struct count_specializations : std::integral_constant<int,
        is_specialized_member_serialize<T, A>::value +
        is_specialized_member_load_save<T, A>::value +
        is_specialized_member_load_save_minimal<T, A>::value +
        is_specialized_non_member_serialize<T, A>::value +
        is_specialized_non_member_load_save<T, A>::value +
        is_specialized_non_member_load_save_minimal<T, A>::value> {};
    } // namespace detail

    //! Check if any specialization exists for a type
    template <class T, class A>
    struct is_specialized : std::integral_constant<bool,
      detail::is_specialized_member_serialize<T, A>::value ||
      detail::is_specialized_member_load_save<T, A>::value ||
      detail::is_specialized_member_load_save_minimal<T, A>::value ||
      detail::is_specialized_non_member_serialize<T, A>::value ||
      detail::is_specialized_non_member_load_save<T, A>::value ||
      detail::is_specialized_non_member_load_save_minimal<T, A>::value>
    {
      static_assert(detail::count_specializations<T, A>::value <= 1, "More than one explicit specialization detected for type.");
    };

    //! Create the static assertion for some specialization
    /*! This assertion will fail if the type is indeed specialized and does not have the appropriate
        type of serialization functions */
    #define CEREAL_MAKE_IS_SPECIALIZED_ASSERT(name, versioned_name, print_name, spec_name)                      \
    static_assert( (is_specialized<T, A>::value && detail::is_specialized_##spec_name<T, A>::value &&           \
                   (has_##name<T, A>::value || has_##versioned_name<T, A>::value))                              \
                   || !(is_specialized<T, A>::value && detail::is_specialized_##spec_name<T, A>::value),        \
                   "cereal detected " #print_name " specialization but no " #print_name " serialize function" )

    //! Generates a test for specialization for versioned and unversioned functions
    /*! This creates checks that can be queried to see if a given type of serialization function
        has been specialized for this type */
    #define CEREAL_MAKE_IS_SPECIALIZED(name, versioned_name, spec_name)                     \
    template <class T, class A>                                                             \
    struct is_specialized_##name : std::integral_constant<bool,                             \
      is_specialized<T, A>::value && detail::is_specialized_##spec_name<T, A>::value>       \
    { CEREAL_MAKE_IS_SPECIALIZED_ASSERT(name, versioned_name, name, spec_name); };          \
    template <class T, class A>                                                             \
    struct is_specialized_##versioned_name : std::integral_constant<bool,                   \
      is_specialized<T, A>::value && detail::is_specialized_##spec_name<T, A>::value>       \
    { CEREAL_MAKE_IS_SPECIALIZED_ASSERT(name, versioned_name, versioned_name, spec_name); }

    CEREAL_MAKE_IS_SPECIALIZED(member_serialize, member_versioned_serialize, member_serialize);
    CEREAL_MAKE_IS_SPECIALIZED(non_member_serialize, non_member_versioned_serialize, non_member_serialize);

    CEREAL_MAKE_IS_SPECIALIZED(member_save, member_versioned_save, member_load_save);
    CEREAL_MAKE_IS_SPECIALIZED(non_member_save, non_member_versioned_save, non_member_load_save);
    CEREAL_MAKE_IS_SPECIALIZED(member_load, member_versioned_load, member_load_save);
    CEREAL_MAKE_IS_SPECIALIZED(non_member_load, non_member_versioned_load, non_member_load_save);

    CEREAL_MAKE_IS_SPECIALIZED(member_save_minimal, member_versioned_save_minimal, member_load_save_minimal);
    CEREAL_MAKE_IS_SPECIALIZED(non_member_save_minimal, non_member_versioned_save_minimal, non_member_load_save_minimal);
    CEREAL_MAKE_IS_SPECIALIZED(member_load_minimal, member_versioned_load_minimal, member_load_save_minimal);
    CEREAL_MAKE_IS_SPECIALIZED(non_member_load_minimal, non_member_versioned_load_minimal, non_member_load_save_minimal);

    #undef CEREAL_MAKE_IS_SPECIALIZED_ASSERT
    #undef CEREAL_MAKE_IS_SPECIALIZED

    // ######################################################################
    // detects if a type has any active minimal output serialization
    template <class T, class OutputArchive>
    struct has_minimal_output_serialization : std::integral_constant<bool,
      is_specialized_member_save_minimal<T, OutputArchive>::value ||
      ((has_member_save_minimal<T, OutputArchive>::value ||
        has_non_member_save_minimal<T, OutputArchive>::value ||
        has_member_versioned_save_minimal<T, OutputArchive>::value ||
        has_non_member_versioned_save_minimal<T, OutputArchive>::value) &&
       !(is_specialized_member_serialize<T, OutputArchive>::value ||
         is_specialized_member_save<T, OutputArchive>::value))> {};

    // ######################################################################
    // detects if a type has any active minimal input serialization
    template <class T, class InputArchive>
    struct has_minimal_input_serialization : std::integral_constant<bool,
      is_specialized_member_load_minimal<T, InputArchive>::value ||
      ((has_member_load_minimal<T, InputArchive>::value ||
        has_non_member_load_minimal<T, InputArchive>::value ||
        has_member_versioned_load_minimal<T, InputArchive>::value ||
        has_non_member_versioned_load_minimal<T, InputArchive>::value) &&
       !(is_specialized_member_serialize<T, InputArchive>::value ||
         is_specialized_member_load<T, InputArchive>::value))> {};

    // ######################################################################
    namespace detail
    {
      //! The number of output serialization functions available
      /*! If specialization is being used, we'll count only those; otherwise we'll count everything */
      template <class T, class OutputArchive>
      struct count_output_serializers : std::integral_constant<int,
        count_specializations<T, OutputArchive>::value ? count_specializations<T, OutputArchive>::value :
        has_member_save<T, OutputArchive>::value +
        has_non_member_save<T, OutputArchive>::value +
        has_member_serialize<T, OutputArchive>::value +
        has_non_member_serialize<T, OutputArchive>::value +
        has_member_save_minimal<T, OutputArchive>::value +
        has_non_member_save_minimal<T, OutputArchive>::value +
        /*-versioned---------------------------------------------------------*/
        has_member_versioned_save<T, OutputArchive>::value +
        has_non_member_versioned_save<T, OutputArchive>::value +
        has_member_versioned_serialize<T, OutputArchive>::value +
        has_non_member_versioned_serialize<T, OutputArchive>::value +
        has_member_versioned_save_minimal<T, OutputArchive>::value +
        has_non_member_versioned_save_minimal<T, OutputArchive>::value> {};
    }

    template <class T, class OutputArchive>
    struct is_output_serializable : std::integral_constant<bool,
      detail::count_output_serializers<T, OutputArchive>::value == 1> {};

    // ######################################################################
    namespace detail
    {
      //! The number of input serialization functions available
      /*! If specialization is being used, we'll count only those; otherwise we'll count everything */
      template <class T, class InputArchive>
      struct count_input_serializers : std::integral_constant<int,
        count_specializations<T, InputArchive>::value ? count_specializations<T, InputArchive>::value :
        has_member_load<T, InputArchive>::value +
        has_non_member_load<T, InputArchive>::value +
        has_member_serialize<T, InputArchive>::value +
        has_non_member_serialize<T, InputArchive>::value +
        has_member_load_minimal<T, InputArchive>::value +
        has_non_member_load_minimal<T, InputArchive>::value +
        /*-versioned---------------------------------------------------------*/
        has_member_versioned_load<T, InputArchive>::value +
        has_non_member_versioned_load<T, InputArchive>::value +
        has_member_versioned_serialize<T, InputArchive>::value +
        has_non_member_versioned_serialize<T, InputArchive>::value +
        has_member_versioned_load_minimal<T, InputArchive>::value +
        has_non_member_versioned_load_minimal<T, InputArchive>::value> {};
    }

    template <class T, class InputArchive>
    struct is_input_serializable : std::integral_constant<bool,
      detail::count_input_serializers<T, InputArchive>::value == 1> {};

    // ######################################################################
    // Base Class Support
    namespace detail
    {
      struct base_class_id
      {
        template<class T>
          base_class_id(T const * const t) :
          type(typeid(T)),
          ptr(t),
          hash(std::hash<std::type_index>()(typeid(T)) ^ (std::hash<void const *>()(t) << 1))
          { }

          bool operator==(base_class_id const & other) const
          { return (type == other.type) && (ptr == other.ptr); }

          std::type_index type;
          void const * ptr;
          size_t hash;
      };
      struct base_class_id_hash { size_t operator()(base_class_id const & id) const { return id.hash; }  };
    } // namespace detail

    namespace detail
    {
      //! Common base type for base class casting
      struct BaseCastBase {};

      template <class>
      struct get_base_class;

      template <template<typename> class Cast, class Base>
      struct get_base_class<Cast<Base>>
      {
        using type = Base;
      };

      //! Base class cast, behave as the test
      template <class Cast, template<class, class> class Test, class Archive,
                bool IsBaseCast = std::is_base_of<BaseCastBase, Cast>::value>
      struct has_minimal_base_class_serialization_impl : Test<typename get_base_class<Cast>::type, Archive>
      { };

      //! Not a base class cast
      template <class Cast, template<class, class> class Test, class Archive>
      struct has_minimal_base_class_serialization_impl<Cast,Test, Archive, false> : std::false_type
      { };
    }

    //! Checks to see if the base class used in a cast has a minimal serialization
    /*! @tparam Cast Either base_class or virtual_base_class wrapped type
        @tparam Test A has_minimal test (for either input or output)
        @tparam Archive The archive to use with the test */
    template <class Cast, template<class, class> class Test, class Archive>
    struct has_minimal_base_class_serialization : detail::has_minimal_base_class_serialization_impl<Cast, Test, Archive>
    { };


    // ######################################################################
    namespace detail
    {
      struct shared_from_this_wrapper
      {
        template <class U>
        static auto (check)( U const & t ) -> decltype( ::cereal::access::shared_from_this(t), std::true_type() );

        static auto (check)( ... ) -> decltype( std::false_type() );

        template <class U>
        static auto get( U const & t ) -> decltype( t.shared_from_this() );
      };
    }

    //! Determine if T or any base class of T has inherited from std::enable_shared_from_this
    template<class T>
    struct has_shared_from_this : decltype((detail::shared_from_this_wrapper::check)(std::declval<T>()))
    { };

    //! Get the type of the base class of T which inherited from std::enable_shared_from_this
    template <class T>
    struct get_shared_from_this_base
    {
      private:
        using PtrType = decltype(detail::shared_from_this_wrapper::get(std::declval<T>()));
      public:
        //! The type of the base of T that inherited from std::enable_shared_from_this
        using type = typename std::decay<typename PtrType::element_type>::type;
    };

    // ######################################################################
    //! Extracts the true type from something possibly wrapped in a cereal NoConvert
    /*! Internally cereal uses some wrapper classes to test the validity of non-member
        minimal load and save functions.  This can interfere with user type traits on
        templated load and save minimal functions.  To get to the correct underlying type,
        users should use strip_minimal when performing any enable_if type type trait checks.

        See the enum serialization in types/common.hpp for an example of using this */
    template <class T, bool IsCerealMinimalTrait = std::is_base_of<detail::NoConvertBase, T>::value>
    struct strip_minimal
    {
      using type = T;
    };

    //! Specialization for types wrapped in a NoConvert
    template <class T>
    struct strip_minimal<T, true>
    {
      using type = typename T::type;
    };

    // ######################################################################
    //! Determines whether the class T can be default constructed by cereal::access
    template <class T>
    struct is_default_constructible
    {
      #ifdef CEREAL_OLDER_GCC
      template <class TT, class SFINAE = void>
      struct test : no {};
      template <class TT>
      struct test<TT, typename detail::Void< decltype( cereal::access::construct<TT>() ) >::type> : yes {};
      static const bool value = test<T>();
      #else // NOT CEREAL_OLDER_GCC =========================================
      template <class TT>
      static auto test(int) -> decltype( cereal::access::construct<TT>(), yes());
      template <class>
      static no test(...);
      static const bool value = std::is_same<decltype(test<T>(0)), yes>::value;
      #endif // NOT CEREAL_OLDER_GCC
    };

    // ######################################################################
    namespace detail
    {
      //! Removes all qualifiers and minimal wrappers from an archive
      template <class A>
      using decay_archive = typename std::decay<typename strip_minimal<A>::type>::type;
    }

    //! Checks if the provided archive type is equal to some cereal archive type
    /*! This automatically does things such as std::decay and removing any other wrappers that may be
        on the Archive template parameter.

        Example use:
        @code{cpp}
        // example use to disable a serialization function
        template <class Archive, EnableIf<cereal::traits::is_same_archive<Archive, cereal::BinaryOutputArchive>::value> = sfinae>
        void save( Archive & ar, MyType const & mt );
        @endcode */
    template <class ArchiveT, class CerealArchiveT>
    struct is_same_archive : std::integral_constant<bool,
      std::is_same<detail::decay_archive<ArchiveT>, CerealArchiveT>::value>
    { };

    // ######################################################################
    //! A macro to use to restrict which types of archives your function will work for.
    /*! This requires you to have a template class parameter named Archive and replaces the void return
        type for your function.

        INTYPE refers to the input archive type you wish to restrict on.
        OUTTYPE refers to the output archive type you wish to restrict on.

        For example, if we want to limit a serialize to only work with binary serialization:

        @code{.cpp}
        template <class Archive>
        CEREAL_ARCHIVE_RESTRICT(BinaryInputArchive, BinaryOutputArchive)
        serialize( Archive & ar, MyCoolType & m )
        {
          ar & m;
        }
        @endcode

        If you need to do more restrictions in your enable_if, you will need to do this by hand.
     */
    #define CEREAL_ARCHIVE_RESTRICT(INTYPE, OUTTYPE) \
    typename std::enable_if<cereal::traits::is_same_archive<Archive, INTYPE>::value || cereal::traits::is_same_archive<Archive, OUTTYPE>::value, void>::type

    //! Type traits only struct used to mark an archive as human readable (text based)
    /*! Archives that wish to identify as text based/human readable should inherit from
        this struct */
    struct TextArchive {};

    //! Checks if an archive is a text archive (human readable)
    template <class A>
    struct is_text_archive : std::integral_constant<bool,
      std::is_base_of<TextArchive, detail::decay_archive<A>>::value>
    { };
  } // namespace traits

  // ######################################################################
  namespace detail
  {
    template <class T, class A,
              bool Member = traits::has_member_load_and_construct<T, A>::value,
              bool MemberVersioned = traits::has_member_versioned_load_and_construct<T, A>::value,
              bool NonMember = traits::has_non_member_load_and_construct<T, A>::value,
              bool NonMemberVersioned = traits::has_non_member_versioned_load_and_construct<T, A>::value>
    struct Construct
    {
      static_assert( cereal::traits::detail::delay_static_assert<T>::value,
        "cereal found more than one compatible load_and_construct function for the provided type and archive combination. \n\n "
        "Types must either have a member load_and_construct function or a non-member specialization of LoadAndConstruct (you may not mix these). \n "
        "In addition, you may not mix versioned with non-versioned load_and_construct functions. \n\n " );
      static T * load_andor_construct( A & /*ar*/, construct<T> & /*construct*/ )
      { return nullptr; }
    };

    // no load and construct case
    template <class T, class A>
    struct Construct<T, A, false, false, false, false>
    {
      static_assert( ::cereal::traits::is_default_constructible<T>::value,
                     "Trying to serialize a an object with no default constructor. \n\n "
                     "Types must either be default constructible or define either a member or non member Construct function. \n "
                     "Construct functions generally have the signature: \n\n "
                     "template <class Archive> \n "
                     "static void load_and_construct(Archive & ar, cereal::construct<T> & construct) \n "
                     "{ \n "
                     "  var a; \n "
                     "  ar( a ) \n "
                     "  construct( a ); \n "
                     "} \n\n" );
      static T * load_andor_construct()
      { return ::cereal::access::construct<T>(); }
    };

    // member non-versioned
    template <class T, class A>
    struct Construct<T, A, true, false, false, false>
    {
      static void load_andor_construct( A & ar, construct<T> & construct )
      {
        access::load_and_construct<T>( ar, construct );
      }
    };

    // member versioned
    template <class T, class A>
    struct Construct<T, A, false, true, false, false>
    {
      static void load_andor_construct( A & ar, construct<T> & construct )
      {
        const auto version = ar.template loadClassVersion<T>();
        access::load_and_construct<T>( ar, construct, version );
      }
    };

    // non-member non-versioned
    template <class T, class A>
    struct Construct<T, A, false, false, true, false>
    {
      static void load_andor_construct( A & ar, construct<T> & construct )
      {
        LoadAndConstruct<T>::load_and_construct( ar, construct );
      }
    };

    // non-member versioned
    template <class T, class A>
    struct Construct<T, A, false, false, false, true>
    {
      static void load_andor_construct( A & ar, construct<T> & construct )
      {
        const auto version = ar.template loadClassVersion<T>();
        LoadAndConstruct<T>::load_and_construct( ar, construct, version );
      }
    };
  } // namespace detail
} // namespace cereal

#endif // CEREAL_DETAILS_TRAITS_HPP_
```
