## Supplemental code for "The motif composition of variable-number tandem repeats impacts gene expression": base64.hpp

```
/*
   Copyright (C) 2004-2008 René Nyffenegger

   This source code is provided 'as-is', without any express or implied
   warranty. In no event will the author be held liable for any damages
   arising from the use of this software.

   Permission is granted to anyone to use this software for any purpose,
   including commercial applications, and to alter it and redistribute it
   freely, subject to the following restrictions:

   1. The origin of this source code must not be misrepresented; you must not
      claim that you wrote the original source code. If you use this source code
      in a product, an acknowledgment in the product documentation would be
      appreciated but is not required.

   2. Altered source versions must be plainly marked as such, and must not be
      misrepresented as being the original source code.

   3. This notice may not be removed or altered from any source distribution.

   René Nyffenegger
*/

#ifndef CEREAL_EXTERNAL_BASE64_HPP_
#define CEREAL_EXTERNAL_BASE64_HPP_

#ifdef __GNUC__
#pragma GCC diagnostic push
#pragma GCC diagnostic ignored "-Wconversion"
#endif

#include <string>

namespace cereal
{
  namespace base64
  {
    static const std::string chars =
      "ABCDEFGHIJKLMNOPQRSTUVWXYZ"
      "abcdefghijklmnopqrstuvwxyz"
      "0123456789+/";

    static inline bool is_base64(unsigned char c) {
      return (isalnum(c) || (c == '+') || (c == '/'));
    }

    inline std::string encode(unsigned char const* bytes_to_encode, size_t in_len) {
      std::string ret;
      int i = 0;
      int j = 0;
      unsigned char char_array_3[3];
      unsigned char char_array_4[4];

      while (in_len--) {
        char_array_3[i++] = *(bytes_to_encode++);
        if (i == 3) {
          char_array_4[0] = static_cast<unsigned char>((char_array_3[0] & 0xfc) >> 2);
          char_array_4[1] = static_cast<unsigned char>( ( ( char_array_3[0] & 0x03 ) << 4 ) + ( ( char_array_3[1] & 0xf0 ) >> 4 ) );
          char_array_4[2] = static_cast<unsigned char>( ( ( char_array_3[1] & 0x0f ) << 2 ) + ( ( char_array_3[2] & 0xc0 ) >> 6 ) );
          char_array_4[3] = static_cast<unsigned char>( char_array_3[2] & 0x3f );

          for(i = 0; (i <4) ; i++)
            ret += chars[char_array_4[i]];
          i = 0;
        }
      }

      if (i)
      {
        for(j = i; j < 3; j++)
          char_array_3[j] = '\0';

        char_array_4[0] = (char_array_3[0] & 0xfc) >> 2;
        char_array_4[1] = ((char_array_3[0] & 0x03) << 4) + ((char_array_3[1] & 0xf0) >> 4);
        char_array_4[2] = ((char_array_3[1] & 0x0f) << 2) + ((char_array_3[2] & 0xc0) >> 6);
        char_array_4[3] = char_array_3[2] & 0x3f;

        for (j = 0; (j < i + 1); j++)
          ret += chars[char_array_4[j]];

        while((i++ < 3))
          ret += '=';
      }

      return ret;
    }

    inline std::string decode(std::string const& encoded_string) {
      size_t in_len = encoded_string.size();
      size_t i = 0;
      size_t j = 0;
      int in_ = 0;
      unsigned char char_array_4[4], char_array_3[3];
      std::string ret;

      while (in_len-- && ( encoded_string[in_] != '=') && is_base64(encoded_string[in_])) {
        char_array_4[i++] = encoded_string[in_]; in_++;
        if (i ==4) {
          for (i = 0; i <4; i++)
            char_array_4[i] = static_cast<unsigned char>(chars.find( char_array_4[i] ));

          char_array_3[0] = (char_array_4[0] << 2) + ((char_array_4[1] & 0x30) >> 4);
          char_array_3[1] = ((char_array_4[1] & 0xf) << 4) + ((char_array_4[2] & 0x3c) >> 2);
          char_array_3[2] = ((char_array_4[2] & 0x3) << 6) + char_array_4[3];

          for (i = 0; (i < 3); i++)
            ret += char_array_3[i];
          i = 0;
        }
      }

      if (i) {
        for (j = i; j <4; j++)
          char_array_4[j] = 0;

        for (j = 0; j <4; j++)
          char_array_4[j] = static_cast<unsigned char>(chars.find( char_array_4[j] ));

        char_array_3[0] = (char_array_4[0] << 2) + ((char_array_4[1] & 0x30) >> 4);
        char_array_3[1] = ((char_array_4[1] & 0xf) << 4) + ((char_array_4[2] & 0x3c) >> 2);
        char_array_3[2] = ((char_array_4[2] & 0x3) << 6) + char_array_4[3];

        for (j = 0; (j < i - 1); j++) ret += char_array_3[j];
      }

      return ret;
    }
  } // namespace base64
} // namespace cereal
#ifdef __GNUC__
#pragma GCC diagnostic pop
#endif
#endif // CEREAL_EXTERNAL_BASE64_HPP_
```
