## Supplemental code for "The motif composition of variable-number tandem repeats impacts gene expression": allocators.h

```
// Tencent is pleased to support the open source community by making RapidJSON available.
// 
// Copyright (C) 2015 THL A29 Limited, a Tencent company, and Milo Yip. All rights reserved.
//
// Licensed under the MIT License (the "License"); you may not use this file except
// in compliance with the License. You may obtain a copy of the License at
//
// http://opensource.org/licenses/MIT
//
// Unless required by applicable law or agreed to in writing, software distributed 
// under the License is distributed on an "AS IS" BASIS, WITHOUT WARRANTIES OR 
// CONDITIONS OF ANY KIND, either express or implied. See the License for the 
// specific language governing permissions and limitations under the License.

#ifndef CEREAL_RAPIDJSON_ALLOCATORS_H_
#define CEREAL_RAPIDJSON_ALLOCATORS_H_

#include "rapidjson.h"

CEREAL_RAPIDJSON_NAMESPACE_BEGIN

///////////////////////////////////////////////////////////////////////////////
// Allocator

/*! \class rapidjson::Allocator
    \brief Concept for allocating, resizing and freeing memory block.
    
    Note that Malloc() and Realloc() are non-static but Free() is static.
    
    So if an allocator need to support Free(), it needs to put its pointer in 
    the header of memory block.

\code
concept Allocator {
    static const bool kNeedFree;    //!< Whether this allocator needs to call Free().

    // Allocate a memory block.
    // \param size of the memory block in bytes.
    // \returns pointer to the memory block.
    void* Malloc(size_t size);

    // Resize a memory block.
    // \param originalPtr The pointer to current memory block. Null pointer is permitted.
    // \param originalSize The current size in bytes. (Design issue: since some allocator may not book-keep this, explicitly pass to it can save memory.)
    // \param newSize the new size in bytes.
    void* Realloc(void* originalPtr, size_t originalSize, size_t newSize);

    // Free a memory block.
    // \param pointer to the memory block. Null pointer is permitted.
    static void Free(void *ptr);
};
\endcode
*/


/*! \def CEREAL_RAPIDJSON_ALLOCATOR_DEFAULT_CHUNK_CAPACITY
    \ingroup CEREAL_RAPIDJSON_CONFIG
    \brief User-defined kDefaultChunkCapacity definition.

    User can define this as any \c size that is a power of 2.
*/

#ifndef CEREAL_RAPIDJSON_ALLOCATOR_DEFAULT_CHUNK_CAPACITY
#define CEREAL_RAPIDJSON_ALLOCATOR_DEFAULT_CHUNK_CAPACITY (64 * 1024)
#endif


///////////////////////////////////////////////////////////////////////////////
// CrtAllocator

//! C-runtime library allocator.
/*! This class is just wrapper for standard C library memory routines.
    \note implements Allocator concept
*/
class CrtAllocator {
public:
    static const bool kNeedFree = true;
    void* Malloc(size_t size) { 
        if (size) //  behavior of malloc(0) is implementation defined.
            return std::malloc(size);
        else
            return NULL; // standardize to returning NULL.
    }
    void* Realloc(void* originalPtr, size_t originalSize, size_t newSize) {
        (void)originalSize;
        if (newSize == 0) {
            std::free(originalPtr);
            return NULL;
        }
        return std::realloc(originalPtr, newSize);
    }
    static void Free(void *ptr) { std::free(ptr); }
};

///////////////////////////////////////////////////////////////////////////////
// MemoryPoolAllocator

//! Default memory allocator used by the parser and DOM.
/*! This allocator allocate memory blocks from pre-allocated memory chunks. 

    It does not free memory blocks. And Realloc() only allocate new memory.

    The memory chunks are allocated by BaseAllocator, which is CrtAllocator by default.

    User may also supply a buffer as the first chunk.

    If the user-buffer is full then additional chunks are allocated by BaseAllocator.

    The user-buffer is not deallocated by this allocator.

    \tparam BaseAllocator the allocator type for allocating memory chunks. Default is CrtAllocator.
    \note implements Allocator concept
*/
template <typename BaseAllocator = CrtAllocator>
class MemoryPoolAllocator {
public:
    static const bool kNeedFree = false;    //!< Tell users that no need to call Free() with this allocator. (concept Allocator)

    //! Constructor with chunkSize.
    /*! \param chunkSize The size of memory chunk. The default is kDefaultChunkSize.
        \param baseAllocator The allocator for allocating memory chunks.
    */
    MemoryPoolAllocator(size_t chunkSize = kDefaultChunkCapacity, BaseAllocator* baseAllocator = 0) : 
        chunkHead_(0), chunk_capacity_(chunkSize), userBuffer_(0), baseAllocator_(baseAllocator), ownBaseAllocator_(0)
    {
    }

    //! Constructor with user-supplied buffer.
    /*! The user buffer will be used firstly. When it is full, memory pool allocates new chunk with chunk size.

        The user buffer will not be deallocated when this allocator is destructed.

        \param buffer User supplied buffer.
        \param size Size of the buffer in bytes. It must at least larger than sizeof(ChunkHeader).
        \param chunkSize The size of memory chunk. The default is kDefaultChunkSize.
        \param baseAllocator The allocator for allocating memory chunks.
    */
    MemoryPoolAllocator(void *buffer, size_t size, size_t chunkSize = kDefaultChunkCapacity, BaseAllocator* baseAllocator = 0) :
        chunkHead_(0), chunk_capacity_(chunkSize), userBuffer_(buffer), baseAllocator_(baseAllocator), ownBaseAllocator_(0)
    {
        CEREAL_RAPIDJSON_ASSERT(buffer != 0);
        CEREAL_RAPIDJSON_ASSERT(size > sizeof(ChunkHeader));
        chunkHead_ = reinterpret_cast<ChunkHeader*>(buffer);
        chunkHead_->capacity = size - sizeof(ChunkHeader);
        chunkHead_->size = 0;
        chunkHead_->next = 0;
    }

    //! Destructor.
    /*! This deallocates all memory chunks, excluding the user-supplied buffer.
    */
    ~MemoryPoolAllocator() {
        Clear();
        CEREAL_RAPIDJSON_DELETE(ownBaseAllocator_);
    }

    //! Deallocates all memory chunks, excluding the user-supplied buffer.
    void Clear() {
        while (chunkHead_ && chunkHead_ != userBuffer_) {
            ChunkHeader* next = chunkHead_->next;
            baseAllocator_->Free(chunkHead_);
            chunkHead_ = next;
        }
        if (chunkHead_ && chunkHead_ == userBuffer_)
            chunkHead_->size = 0; // Clear user buffer
    }

    //! Computes the total capacity of allocated memory chunks.
    /*! \return total capacity in bytes.
    */
    size_t Capacity() const {
        size_t capacity = 0;
        for (ChunkHeader* c = chunkHead_; c != 0; c = c->next)
            capacity += c->capacity;
        return capacity;
    }

    //! Computes the memory blocks allocated.
    /*! \return total used bytes.
    */
    size_t Size() const {
        size_t size = 0;
        for (ChunkHeader* c = chunkHead_; c != 0; c = c->next)
            size += c->size;
        return size;
    }

    //! Allocates a memory block. (concept Allocator)
    void* Malloc(size_t size) {
        if (!size)
            return NULL;

        size = CEREAL_RAPIDJSON_ALIGN(size);
        if (chunkHead_ == 0 || chunkHead_->size + size > chunkHead_->capacity)
            if (!AddChunk(chunk_capacity_ > size ? chunk_capacity_ : size))
                return NULL;

        void *buffer = reinterpret_cast<char *>(chunkHead_) + CEREAL_RAPIDJSON_ALIGN(sizeof(ChunkHeader)) + chunkHead_->size;
        chunkHead_->size += size;
        return buffer;
    }

    //! Resizes a memory block (concept Allocator)
    void* Realloc(void* originalPtr, size_t originalSize, size_t newSize) {
        if (originalPtr == 0)
            return Malloc(newSize);

        if (newSize == 0)
            return NULL;

        originalSize = CEREAL_RAPIDJSON_ALIGN(originalSize);
        newSize = CEREAL_RAPIDJSON_ALIGN(newSize);

        // Do not shrink if new size is smaller than original
        if (originalSize >= newSize)
            return originalPtr;

        // Simply expand it if it is the last allocation and there is sufficient space
        if (originalPtr == reinterpret_cast<char *>(chunkHead_) + CEREAL_RAPIDJSON_ALIGN(sizeof(ChunkHeader)) + chunkHead_->size - originalSize) {
            size_t increment = static_cast<size_t>(newSize - originalSize);
            if (chunkHead_->size + increment <= chunkHead_->capacity) {
                chunkHead_->size += increment;
                return originalPtr;
            }
        }

        // Realloc process: allocate and copy memory, do not free original buffer.
        if (void* newBuffer = Malloc(newSize)) {
            if (originalSize)
                std::memcpy(newBuffer, originalPtr, originalSize);
            return newBuffer;
        }
        else
            return NULL;
    }

    //! Frees a memory block (concept Allocator)
    static void Free(void *ptr) { (void)ptr; } // Do nothing

private:
    //! Copy constructor is not permitted.
    MemoryPoolAllocator(const MemoryPoolAllocator& rhs) /* = delete */;
    //! Copy assignment operator is not permitted.
    MemoryPoolAllocator& operator=(const MemoryPoolAllocator& rhs) /* = delete */;

    //! Creates a new chunk.
    /*! \param capacity Capacity of the chunk in bytes.
        \return true if success.
    */
    bool AddChunk(size_t capacity) {
        if (!baseAllocator_)
            ownBaseAllocator_ = baseAllocator_ = CEREAL_RAPIDJSON_NEW(BaseAllocator)();
        if (ChunkHeader* chunk = reinterpret_cast<ChunkHeader*>(baseAllocator_->Malloc(CEREAL_RAPIDJSON_ALIGN(sizeof(ChunkHeader)) + capacity))) {
            chunk->capacity = capacity;
            chunk->size = 0;
            chunk->next = chunkHead_;
            chunkHead_ =  chunk;
            return true;
        }
        else
            return false;
    }

    static const int kDefaultChunkCapacity = CEREAL_RAPIDJSON_ALLOCATOR_DEFAULT_CHUNK_CAPACITY; //!< Default chunk capacity.

    //! Chunk header for perpending to each chunk.
    /*! Chunks are stored as a singly linked list.
    */
    struct ChunkHeader {
        size_t capacity;    //!< Capacity of the chunk in bytes (excluding the header itself).
        size_t size;        //!< Current size of allocated memory in bytes.
        ChunkHeader *next;  //!< Next chunk in the linked list.
    };

    ChunkHeader *chunkHead_;    //!< Head of the chunk linked-list. Only the head chunk serves allocation.
    size_t chunk_capacity_;     //!< The minimum capacity of chunk when they are allocated.
    void *userBuffer_;          //!< User supplied buffer.
    BaseAllocator* baseAllocator_;  //!< base allocator for allocating memory chunks.
    BaseAllocator* ownBaseAllocator_;   //!< base allocator created by this object.
};

CEREAL_RAPIDJSON_NAMESPACE_END

#endif // CEREAL_RAPIDJSON_ENCODINGS_H_
```
