## Supplemental code for "The motif composition of variable-number tandem repeats impacts gene expression": cursorstreamwrapper.h

```
// Tencent is pleased to support the open source community by making RapidJSON available.
//
// Copyright (C) 2015 THL A29 Limited, a Tencent company, and Milo Yip. All rights reserved.
//
// Licensed under the MIT License (the "License"); you may not use this file except
// in compliance with the License. You may obtain a copy of the License at
//
// http://opensource.org/licenses/MIT
//
// Unless required by applicable law or agreed to in writing, software distributed
// under the License is distributed on an "AS IS" BASIS, WITHOUT WARRANTIES OR
// CONDITIONS OF ANY KIND, either express or implied. See the License for the
// specific language governing permissions and limitations under the License.

#ifndef CEREAL_RAPIDJSON_CURSORSTREAMWRAPPER_H_
#define CEREAL_RAPIDJSON_CURSORSTREAMWRAPPER_H_

#include "stream.h"

#if defined(__GNUC__)
CEREAL_RAPIDJSON_DIAG_PUSH
CEREAL_RAPIDJSON_DIAG_OFF(effc++)
#endif

#if defined(_MSC_VER) && _MSC_VER <= 1800
CEREAL_RAPIDJSON_DIAG_PUSH
CEREAL_RAPIDJSON_DIAG_OFF(4702)  // unreachable code
CEREAL_RAPIDJSON_DIAG_OFF(4512)  // assignment operator could not be generated
#endif

CEREAL_RAPIDJSON_NAMESPACE_BEGIN


//! Cursor stream wrapper for counting line and column number if error exists.
/*!
    \tparam InputStream     Any stream that implements Stream Concept
*/
template <typename InputStream, typename Encoding = UTF8<> >
class CursorStreamWrapper : public GenericStreamWrapper<InputStream, Encoding> {
public:
    typedef typename Encoding::Ch Ch;

    CursorStreamWrapper(InputStream& is):
        GenericStreamWrapper<InputStream, Encoding>(is), line_(1), col_(0) {}

    // counting line and column number
    Ch Take() {
        Ch ch = this->is_.Take();
        if(ch == '\n') {
            line_ ++;
            col_ = 0;
        } else {
            col_ ++;
        }
        return ch;
    }

    //! Get the error line number, if error exists.
    size_t GetLine() const { return line_; }
    //! Get the error column number, if error exists.
    size_t GetColumn() const { return col_; }

private:
    size_t line_;   //!< Current Line
    size_t col_;    //!< Current Column
};

#if defined(_MSC_VER) && _MSC_VER <= 1800
CEREAL_RAPIDJSON_DIAG_POP
#endif

#if defined(__GNUC__)
CEREAL_RAPIDJSON_DIAG_POP
#endif

CEREAL_RAPIDJSON_NAMESPACE_END

#endif // CEREAL_RAPIDJSON_CURSORSTREAMWRAPPER_H_
```
