## Supplemental code for "The motif composition of variable-number tandem repeats impacts gene expression": document.h

```
// Tencent is pleased to support the open source community by making RapidJSON available.
//
// Copyright (C) 2015 THL A29 Limited, a Tencent company, and Milo Yip. All rights reserved.
//
// Licensed under the MIT License (the "License"); you may not use this file except
// in compliance with the License. You may obtain a copy of the License at
//
// http://opensource.org/licenses/MIT
//
// Unless required by applicable law or agreed to in writing, software distributed
// under the License is distributed on an "AS IS" BASIS, WITHOUT WARRANTIES OR
// CONDITIONS OF ANY KIND, either express or implied. See the License for the
// specific language governing permissions and limitations under the License.

#ifndef CEREAL_RAPIDJSON_DOCUMENT_H_
#define CEREAL_RAPIDJSON_DOCUMENT_H_

/*! \file document.h */

#include "reader.h"
#include "internal/meta.h"
#include "internal/strfunc.h"
#include "memorystream.h"
#include "encodedstream.h"
#include <new>      // placement new
#include <limits>
#ifdef __cpp_lib_three_way_comparison
#include <compare>
#endif

CEREAL_RAPIDJSON_DIAG_PUSH
#ifdef __clang__
CEREAL_RAPIDJSON_DIAG_OFF(padded)
CEREAL_RAPIDJSON_DIAG_OFF(switch-enum)
CEREAL_RAPIDJSON_DIAG_OFF(c++98-compat)
#elif defined(_MSC_VER)
CEREAL_RAPIDJSON_DIAG_OFF(4127) // conditional expression is constant
CEREAL_RAPIDJSON_DIAG_OFF(4244) // conversion from kXxxFlags to 'uint16_t', possible loss of data
#endif

#ifdef __GNUC__
CEREAL_RAPIDJSON_DIAG_OFF(effc++)
#endif // __GNUC__

#ifndef CEREAL_RAPIDJSON_NOMEMBERITERATORCLASS
#include <iterator> // std::random_access_iterator_tag
#endif

#if CEREAL_RAPIDJSON_HAS_CXX11_RVALUE_REFS
#include <utility> // std::move
#endif

CEREAL_RAPIDJSON_NAMESPACE_BEGIN

// Forward declaration.
template <typename Encoding, typename Allocator>
class GenericValue;

template <typename Encoding, typename Allocator, typename StackAllocator>
class GenericDocument;

//! Name-value pair in a JSON object value.
/*!
    This class was internal to GenericValue. It used to be a inner struct.
    But a compiler (IBM XL C/C++ for AIX) have reported to have problem with that so it moved as a namespace scope struct.
    https://code.google.com/p/rapidjson/issues/detail?id=64
*/
template <typename Encoding, typename Allocator>
struct GenericMember {
    GenericValue<Encoding, Allocator> name;     //!< name of member (must be a string)
    GenericValue<Encoding, Allocator> value;    //!< value of member.

    // swap() for std::sort() and other potential use in STL.
    friend inline void swap(GenericMember& a, GenericMember& b) CEREAL_RAPIDJSON_NOEXCEPT {
        a.name.Swap(b.name);
        a.value.Swap(b.value);
    }
};

///////////////////////////////////////////////////////////////////////////////
// GenericMemberIterator

#ifndef CEREAL_RAPIDJSON_NOMEMBERITERATORCLASS

//! (Constant) member iterator for a JSON object value
/*!
    \tparam Const Is this a constant iterator?
    \tparam Encoding    Encoding of the value. (Even non-string values need to have the same encoding in a document)
    \tparam Allocator   Allocator type for allocating memory of object, array and string.

    This class implements a Random Access Iterator for GenericMember elements
    of a GenericValue, see ISO/IEC 14882:2003(E) C++ standard, 24.1 [lib.iterator.requirements].

    \note This iterator implementation is mainly intended to avoid implicit
        conversions from iterator values to \c NULL,
        e.g. from GenericValue::FindMember.

    \note Define \c CEREAL_RAPIDJSON_NOMEMBERITERATORCLASS to fall back to a
        pointer-based implementation, if your platform doesn't provide
        the C++ <iterator> header.

    \see GenericMember, GenericValue::MemberIterator, GenericValue::ConstMemberIterator
 */
template <bool Const, typename Encoding, typename Allocator>
class GenericMemberIterator {

    friend class GenericValue<Encoding,Allocator>;
    template <bool, typename, typename> friend class GenericMemberIterator;

    typedef GenericMember<Encoding,Allocator> PlainType;
    typedef typename internal::MaybeAddConst<Const,PlainType>::Type ValueType;

public:
    //! Iterator type itself
    typedef GenericMemberIterator Iterator;
    //! Constant iterator type
    typedef GenericMemberIterator<true,Encoding,Allocator>  ConstIterator;
    //! Non-constant iterator type
    typedef GenericMemberIterator<false,Encoding,Allocator> NonConstIterator;

    /** \name std::iterator_traits support */
    //@{
    typedef ValueType      value_type;
    typedef ValueType *    pointer;
    typedef ValueType &    reference;
    typedef std::ptrdiff_t difference_type;
    typedef std::random_access_iterator_tag iterator_category;
    //@}

    //! Pointer to (const) GenericMember
    typedef pointer         Pointer;
    //! Reference to (const) GenericMember
    typedef reference       Reference;
    //! Signed integer type (e.g. \c ptrdiff_t)
    typedef difference_type DifferenceType;

    //! Default constructor (singular value)
    /*! Creates an iterator pointing to no element.
        \note All operations, except for comparisons, are undefined on such values.
     */
    GenericMemberIterator() : ptr_() {}

    //! Iterator conversions to more const
    /*!
        \param it (Non-const) iterator to copy from

        Allows the creation of an iterator from another GenericMemberIterator
        that is "less const".  Especially, creating a non-constant iterator
        from a constant iterator are disabled:
        \li const -> non-const (not ok)
        \li const -> const (ok)
        \li non-const -> const (ok)
        \li non-const -> non-const (ok)

        \note If the \c Const template parameter is already \c false, this
            constructor effectively defines a regular copy-constructor.
            Otherwise, the copy constructor is implicitly defined.
    */
    GenericMemberIterator(const NonConstIterator & it) : ptr_(it.ptr_) {}
    Iterator& operator=(const NonConstIterator & it) { ptr_ = it.ptr_; return *this; }

    //! @name stepping
    //@{
    Iterator& operator++(){ ++ptr_; return *this; }
    Iterator& operator--(){ --ptr_; return *this; }
    Iterator  operator++(int){ Iterator old(*this); ++ptr_; return old; }
    Iterator  operator--(int){ Iterator old(*this); --ptr_; return old; }
    //@}

    //! @name increment/decrement
    //@{
    Iterator operator+(DifferenceType n) const { return Iterator(ptr_+n); }
    Iterator operator-(DifferenceType n) const { return Iterator(ptr_-n); }

    Iterator& operator+=(DifferenceType n) { ptr_+=n; return *this; }
    Iterator& operator-=(DifferenceType n) { ptr_-=n; return *this; }
    //@}

    //! @name relations
    //@{
    template <bool Const_> bool operator==(const GenericMemberIterator<Const_, Encoding, Allocator>& that) const { return ptr_ == that.ptr_; }
    template <bool Const_> bool operator!=(const GenericMemberIterator<Const_, Encoding, Allocator>& that) const { return ptr_ != that.ptr_; }
    template <bool Const_> bool operator<=(const GenericMemberIterator<Const_, Encoding, Allocator>& that) const { return ptr_ <= that.ptr_; }
    template <bool Const_> bool operator>=(const GenericMemberIterator<Const_, Encoding, Allocator>& that) const { return ptr_ >= that.ptr_; }
    template <bool Const_> bool operator< (const GenericMemberIterator<Const_, Encoding, Allocator>& that) const { return ptr_ < that.ptr_; }
    template <bool Const_> bool operator> (const GenericMemberIterator<Const_, Encoding, Allocator>& that) const { return ptr_ > that.ptr_; }

#ifdef __cpp_lib_three_way_comparison
    template <bool Const_> std::strong_ordering operator<=>(const GenericMemberIterator<Const_, Encoding, Allocator>& that) const { return ptr_ <=> that.ptr_; }
#endif
    //@}

    //! @name dereference
    //@{
    Reference operator*() const { return *ptr_; }
    Pointer   operator->() const { return ptr_; }
    Reference operator[](DifferenceType n) const { return ptr_[n]; }
    //@}

    //! Distance
    DifferenceType operator-(ConstIterator that) const { return ptr_-that.ptr_; }

private:
    //! Internal constructor from plain pointer
    explicit GenericMemberIterator(Pointer p) : ptr_(p) {}

    Pointer ptr_; //!< raw pointer
};

#else // CEREAL_RAPIDJSON_NOMEMBERITERATORCLASS

// class-based member iterator implementation disabled, use plain pointers

template <bool Const, typename Encoding, typename Allocator>
class GenericMemberIterator;

//! non-const GenericMemberIterator
template <typename Encoding, typename Allocator>
class GenericMemberIterator<false,Encoding,Allocator> {
    //! use plain pointer as iterator type
    typedef GenericMember<Encoding,Allocator>* Iterator;
};
//! const GenericMemberIterator
template <typename Encoding, typename Allocator>
class GenericMemberIterator<true,Encoding,Allocator> {
    //! use plain const pointer as iterator type
    typedef const GenericMember<Encoding,Allocator>* Iterator;
};

#endif // CEREAL_RAPIDJSON_NOMEMBERITERATORCLASS

///////////////////////////////////////////////////////////////////////////////
// GenericStringRef

//! Reference to a constant string (not taking a copy)
/*!
    \tparam CharType character type of the string

    This helper class is used to automatically infer constant string
    references for string literals, especially from \c const \b (!)
    character arrays.

    The main use is for creating JSON string values without copying the
    source string via an \ref Allocator.  This requires that the referenced
    string pointers have a sufficient lifetime, which exceeds the lifetime
    of the associated GenericValue.

    \b Example
    \code
    Value v("foo");   // ok, no need to copy & calculate length
    const char foo[] = "foo";
    v.SetString(foo); // ok

    const char* bar = foo;
    // Value x(bar); // not ok, can't rely on bar's lifetime
    Value x(StringRef(bar)); // lifetime explicitly guaranteed by user
    Value y(StringRef(bar, 3));  // ok, explicitly pass length
    \endcode

    \see StringRef, GenericValue::SetString
*/
template<typename CharType>
struct GenericStringRef {
    typedef CharType Ch; //!< character type of the string

    //! Create string reference from \c const character array
#ifndef __clang__ // -Wdocumentation
    /*!
        This constructor implicitly creates a constant string reference from
        a \c const character array.  It has better performance than
        \ref StringRef(const CharType*) by inferring the string \ref length
        from the array length, and also supports strings containing null
        characters.

        \tparam N length of the string, automatically inferred

        \param str Constant character array, lifetime assumed to be longer
            than the use of the string in e.g. a GenericValue

        \post \ref s == str

        \note Constant complexity.
        \note There is a hidden, private overload to disallow references to
            non-const character arrays to be created via this constructor.
            By this, e.g. function-scope arrays used to be filled via
            \c snprintf are excluded from consideration.
            In such cases, the referenced string should be \b copied to the
            GenericValue instead.
     */
#endif
    template<SizeType N>
    GenericStringRef(const CharType (&str)[N]) CEREAL_RAPIDJSON_NOEXCEPT
        : s(str), length(N-1) {}

    //! Explicitly create string reference from \c const character pointer
#ifndef __clang__ // -Wdocumentation
    /*!
        This constructor can be used to \b explicitly  create a reference to
        a constant string pointer.

        \see StringRef(const CharType*)

        \param str Constant character pointer, lifetime assumed to be longer
            than the use of the string in e.g. a GenericValue

        \post \ref s == str

        \note There is a hidden, private overload to disallow references to
            non-const character arrays to be created via this constructor.
            By this, e.g. function-scope arrays used to be filled via
            \c snprintf are excluded from consideration.
            In such cases, the referenced string should be \b copied to the
            GenericValue instead.
     */
#endif
    explicit GenericStringRef(const CharType* str)
        : s(str), length(NotNullStrLen(str)) {}

    //! Create constant string reference from pointer and length
#ifndef __clang__ // -Wdocumentation
    /*! \param str constant string, lifetime assumed to be longer than the use of the string in e.g. a GenericValue
        \param len length of the string, excluding the trailing NULL terminator

        \post \ref s == str && \ref length == len
        \note Constant complexity.
     */
#endif
    GenericStringRef(const CharType* str, SizeType len)
        : s(CEREAL_RAPIDJSON_LIKELY(str) ? str : emptyString), length(len) { CEREAL_RAPIDJSON_ASSERT(str != 0 || len == 0u); }

    GenericStringRef(const GenericStringRef& rhs) : s(rhs.s), length(rhs.length) {}

    //! implicit conversion to plain CharType pointer
    operator const Ch *() const { return s; }

    const Ch* const s; //!< plain CharType pointer
    const SizeType length; //!< length of the string (excluding the trailing NULL terminator)

private:
    SizeType NotNullStrLen(const CharType* str) {
        CEREAL_RAPIDJSON_ASSERT(str != 0);
        return internal::StrLen(str);
    }

    /// Empty string - used when passing in a NULL pointer
    static const Ch emptyString[];

    //! Disallow construction from non-const array
    template<SizeType N>
    GenericStringRef(CharType (&str)[N]) /* = delete */;
    //! Copy assignment operator not permitted - immutable type
    GenericStringRef& operator=(const GenericStringRef& rhs) /* = delete */;
};

template<typename CharType>
const CharType GenericStringRef<CharType>::emptyString[] = { CharType() };

//! Mark a character pointer as constant string
/*! Mark a plain character pointer as a "string literal".  This function
    can be used to avoid copying a character string to be referenced as a
    value in a JSON GenericValue object, if the string's lifetime is known
    to be valid long enough.
    \tparam CharType Character type of the string
    \param str Constant string, lifetime assumed to be longer than the use of the string in e.g. a GenericValue
    \return GenericStringRef string reference object
    \relatesalso GenericStringRef

    \see GenericValue::GenericValue(StringRefType), GenericValue::operator=(StringRefType), GenericValue::SetString(StringRefType), GenericValue::PushBack(StringRefType, Allocator&), GenericValue::AddMember
*/
template<typename CharType>
inline GenericStringRef<CharType> StringRef(const CharType* str) {
    return GenericStringRef<CharType>(str);
}

//! Mark a character pointer as constant string
/*! Mark a plain character pointer as a "string literal".  This function
    can be used to avoid copying a character string to be referenced as a
    value in a JSON GenericValue object, if the string's lifetime is known
    to be valid long enough.

    This version has better performance with supplied length, and also
    supports string containing null characters.

    \tparam CharType character type of the string
    \param str Constant string, lifetime assumed to be longer than the use of the string in e.g. a GenericValue
    \param length The length of source string.
    \return GenericStringRef string reference object
    \relatesalso GenericStringRef
*/
template<typename CharType>
inline GenericStringRef<CharType> StringRef(const CharType* str, size_t length) {
    return GenericStringRef<CharType>(str, SizeType(length));
}

#if CEREAL_RAPIDJSON_HAS_STDSTRING
//! Mark a string object as constant string
/*! Mark a string object (e.g. \c std::string) as a "string literal".
    This function can be used to avoid copying a string to be referenced as a
    value in a JSON GenericValue object, if the string's lifetime is known
    to be valid long enough.

    \tparam CharType character type of the string
    \param str Constant string, lifetime assumed to be longer than the use of the string in e.g. a GenericValue
    \return GenericStringRef string reference object
    \relatesalso GenericStringRef
    \note Requires the definition of the preprocessor symbol \ref CEREAL_RAPIDJSON_HAS_STDSTRING.
*/
template<typename CharType>
inline GenericStringRef<CharType> StringRef(const std::basic_string<CharType>& str) {
    return GenericStringRef<CharType>(str.data(), SizeType(str.size()));
}
#endif

///////////////////////////////////////////////////////////////////////////////
// GenericValue type traits
namespace internal {

template <typename T, typename Encoding = void, typename Allocator = void>
struct IsGenericValueImpl : FalseType {};

// select candidates according to nested encoding and allocator types
template <typename T> struct IsGenericValueImpl<T, typename Void<typename T::EncodingType>::Type, typename Void<typename T::AllocatorType>::Type>
    : IsBaseOf<GenericValue<typename T::EncodingType, typename T::AllocatorType>, T>::Type {};

// helper to match arbitrary GenericValue instantiations, including derived classes
template <typename T> struct IsGenericValue : IsGenericValueImpl<T>::Type {};

} // namespace internal

///////////////////////////////////////////////////////////////////////////////
// TypeHelper

namespace internal {

template <typename ValueType, typename T>
struct TypeHelper {};

template<typename ValueType>
struct TypeHelper<ValueType, bool> {
    static bool Is(const ValueType& v) { return v.IsBool(); }
    static bool Get(const ValueType& v) { return v.GetBool(); }
    static ValueType& Set(ValueType& v, bool data) { return v.SetBool(data); }
    static ValueType& Set(ValueType& v, bool data, typename ValueType::AllocatorType&) { return v.SetBool(data); }
};

template<typename ValueType>
struct TypeHelper<ValueType, int> {
    static bool Is(const ValueType& v) { return v.IsInt(); }
    static int Get(const ValueType& v) { return v.GetInt(); }
    static ValueType& Set(ValueType& v, int data) { return v.SetInt(data); }
    static ValueType& Set(ValueType& v, int data, typename ValueType::AllocatorType&) { return v.SetInt(data); }
};

template<typename ValueType>
struct TypeHelper<ValueType, unsigned> {
    static bool Is(const ValueType& v) { return v.IsUint(); }
    static unsigned Get(const ValueType& v) { return v.GetUint(); }
    static ValueType& Set(ValueType& v, unsigned data) { return v.SetUint(data); }
    static ValueType& Set(ValueType& v, unsigned data, typename ValueType::AllocatorType&) { return v.SetUint(data); }
};

#ifdef _MSC_VER
CEREAL_RAPIDJSON_STATIC_ASSERT(sizeof(long) == sizeof(int));
template<typename ValueType>
struct TypeHelper<ValueType, long> {
    static bool Is(const ValueType& v) { return v.IsInt(); }
    static long Get(const ValueType& v) { return v.GetInt(); }
    static ValueType& Set(ValueType& v, long data) { return v.SetInt(data); }
    static ValueType& Set(ValueType& v, long data, typename ValueType::AllocatorType&) { return v.SetInt(data); }
};

CEREAL_RAPIDJSON_STATIC_ASSERT(sizeof(unsigned long) == sizeof(unsigned));
template<typename ValueType>
struct TypeHelper<ValueType, unsigned long> {
    static bool Is(const ValueType& v) { return v.IsUint(); }
    static unsigned long Get(const ValueType& v) { return v.GetUint(); }
    static ValueType& Set(ValueType& v, unsigned long data) { return v.SetUint(data); }
    static ValueType& Set(ValueType& v, unsigned long data, typename ValueType::AllocatorType&) { return v.SetUint(data); }
};
#endif

template<typename ValueType>
struct TypeHelper<ValueType, int64_t> {
    static bool Is(const ValueType& v) { return v.IsInt64(); }
    static int64_t Get(const ValueType& v) { return v.GetInt64(); }
    static ValueType& Set(ValueType& v, int64_t data) { return v.SetInt64(data); }
    static ValueType& Set(ValueType& v, int64_t data, typename ValueType::AllocatorType&) { return v.SetInt64(data); }
};

template<typename ValueType>
struct TypeHelper<ValueType, uint64_t> {
    static bool Is(const ValueType& v) { return v.IsUint64(); }
    static uint64_t Get(const ValueType& v) { return v.GetUint64(); }
    static ValueType& Set(ValueType& v, uint64_t data) { return v.SetUint64(data); }
    static ValueType& Set(ValueType& v, uint64_t data, typename ValueType::AllocatorType&) { return v.SetUint64(data); }
};

template<typename ValueType>
struct TypeHelper<ValueType, double> {
    static bool Is(const ValueType& v) { return v.IsDouble(); }
    static double Get(const ValueType& v) { return v.GetDouble(); }
    static ValueType& Set(ValueType& v, double data) { return v.SetDouble(data); }
    static ValueType& Set(ValueType& v, double data, typename ValueType::AllocatorType&) { return v.SetDouble(data); }
};

template<typename ValueType>
struct TypeHelper<ValueType, float> {
    static bool Is(const ValueType& v) { return v.IsFloat(); }
    static float Get(const ValueType& v) { return v.GetFloat(); }
    static ValueType& Set(ValueType& v, float data) { return v.SetFloat(data); }
    static ValueType& Set(ValueType& v, float data, typename ValueType::AllocatorType&) { return v.SetFloat(data); }
};

template<typename ValueType>
struct TypeHelper<ValueType, const typename ValueType::Ch*> {
    typedef const typename ValueType::Ch* StringType;
    static bool Is(const ValueType& v) { return v.IsString(); }
    static StringType Get(const ValueType& v) { return v.GetString(); }
    static ValueType& Set(ValueType& v, const StringType data) { return v.SetString(typename ValueType::StringRefType(data)); }
    static ValueType& Set(ValueType& v, const StringType data, typename ValueType::AllocatorType& a) { return v.SetString(data, a); }
};

#if CEREAL_RAPIDJSON_HAS_STDSTRING
template<typename ValueType>
struct TypeHelper<ValueType, std::basic_string<typename ValueType::Ch> > {
    typedef std::basic_string<typename ValueType::Ch> StringType;
    static bool Is(const ValueType& v) { return v.IsString(); }
    static StringType Get(const ValueType& v) { return StringType(v.GetString(), v.GetStringLength()); }
    static ValueType& Set(ValueType& v, const StringType& data, typename ValueType::AllocatorType& a) { return v.SetString(data, a); }
};
#endif

template<typename ValueType>
struct TypeHelper<ValueType, typename ValueType::Array> {
    typedef typename ValueType::Array ArrayType;
    static bool Is(const ValueType& v) { return v.IsArray(); }
    static ArrayType Get(ValueType& v) { return v.GetArray(); }
    static ValueType& Set(ValueType& v, ArrayType data) { return v = data; }
    static ValueType& Set(ValueType& v, ArrayType data, typename ValueType::AllocatorType&) { return v = data; }
};

template<typename ValueType>
struct TypeHelper<ValueType, typename ValueType::ConstArray> {
    typedef typename ValueType::ConstArray ArrayType;
    static bool Is(const ValueType& v) { return v.IsArray(); }
    static ArrayType Get(const ValueType& v) { return v.GetArray(); }
};

template<typename ValueType>
struct TypeHelper<ValueType, typename ValueType::Object> {
    typedef typename ValueType::Object ObjectType;
    static bool Is(const ValueType& v) { return v.IsObject(); }
    static ObjectType Get(ValueType& v) { return v.GetObject(); }
    static ValueType& Set(ValueType& v, ObjectType data) { return v = data; }
    static ValueType& Set(ValueType& v, ObjectType data, typename ValueType::AllocatorType&) { return v = data; }
};

template<typename ValueType>
struct TypeHelper<ValueType, typename ValueType::ConstObject> {
    typedef typename ValueType::ConstObject ObjectType;
    static bool Is(const ValueType& v) { return v.IsObject(); }
    static ObjectType Get(const ValueType& v) { return v.GetObject(); }
};

} // namespace internal

// Forward declarations
template <bool, typename> class GenericArray;
template <bool, typename> class GenericObject;

///////////////////////////////////////////////////////////////////////////////
// GenericValue

//! Represents a JSON value. Use Value for UTF8 encoding and default allocator.
/*!
    A JSON value can be one of 7 types. This class is a variant type supporting
    these types.

    Use the Value if UTF8 and default allocator

    \tparam Encoding    Encoding of the value. (Even non-string values need to have the same encoding in a document)
    \tparam Allocator   Allocator type for allocating memory of object, array and string.
*/
template <typename Encoding, typename Allocator = MemoryPoolAllocator<> >
class GenericValue {
public:
    //! Name-value pair in an object.
    typedef GenericMember<Encoding, Allocator> Member;
    typedef Encoding EncodingType;                  //!< Encoding type from template parameter.
    typedef Allocator AllocatorType;                //!< Allocator type from template parameter.
    typedef typename Encoding::Ch Ch;               //!< Character type derived from Encoding.
    typedef GenericStringRef<Ch> StringRefType;     //!< Reference to a constant string
    typedef typename GenericMemberIterator<false,Encoding,Allocator>::Iterator MemberIterator;  //!< Member iterator for iterating in object.
    typedef typename GenericMemberIterator<true,Encoding,Allocator>::Iterator ConstMemberIterator;  //!< Constant member iterator for iterating in object.
    typedef GenericValue* ValueIterator;            //!< Value iterator for iterating in array.
    typedef const GenericValue* ConstValueIterator; //!< Constant value iterator for iterating in array.
    typedef GenericValue<Encoding, Allocator> ValueType;    //!< Value type of itself.
    typedef GenericArray<false, ValueType> Array;
    typedef GenericArray<true, ValueType> ConstArray;
    typedef GenericObject<false, ValueType> Object;
    typedef GenericObject<true, ValueType> ConstObject;

    //!@name Constructors and destructor.
    //@{

    //! Default constructor creates a null value.
    GenericValue() CEREAL_RAPIDJSON_NOEXCEPT : data_() { data_.f.flags = kNullFlag; }

#if CEREAL_RAPIDJSON_HAS_CXX11_RVALUE_REFS
    //! Move constructor in C++11
    GenericValue(GenericValue&& rhs) CEREAL_RAPIDJSON_NOEXCEPT : data_(rhs.data_) {
        rhs.data_.f.flags = kNullFlag; // give up contents
    }
#endif

private:
    //! Copy constructor is not permitted.
    GenericValue(const GenericValue& rhs);

#if CEREAL_RAPIDJSON_HAS_CXX11_RVALUE_REFS
    //! Moving from a GenericDocument is not permitted.
    template <typename StackAllocator>
    GenericValue(GenericDocument<Encoding,Allocator,StackAllocator>&& rhs);

    //! Move assignment from a GenericDocument is not permitted.
    template <typename StackAllocator>
    GenericValue& operator=(GenericDocument<Encoding,Allocator,StackAllocator>&& rhs);
#endif

public:

    //! Constructor with JSON value type.
    /*! This creates a Value of specified type with default content.
        \param type Type of the value.
        \note Default content for number is zero.
    */
    explicit GenericValue(Type type) CEREAL_RAPIDJSON_NOEXCEPT : data_() {
        static const uint16_t defaultFlags[] = {
            kNullFlag, kFalseFlag, kTrueFlag, kObjectFlag, kArrayFlag, kShortStringFlag,
            kNumberAnyFlag
        };
        CEREAL_RAPIDJSON_NOEXCEPT_ASSERT(type >= kNullType && type <= kNumberType);
        data_.f.flags = defaultFlags[type];

        // Use ShortString to store empty string.
        if (type == kStringType)
            data_.ss.SetLength(0);
    }

    //! Explicit copy constructor (with allocator)
    /*! Creates a copy of a Value by using the given Allocator
        \tparam SourceAllocator allocator of \c rhs
        \param rhs Value to copy from (read-only)
        \param allocator Allocator for allocating copied elements and buffers. Commonly use GenericDocument::GetAllocator().
        \param copyConstStrings Force copying of constant strings (e.g. referencing an in-situ buffer)
        \see CopyFrom()
    */
    template <typename SourceAllocator>
    GenericValue(const GenericValue<Encoding,SourceAllocator>& rhs, Allocator& allocator, bool copyConstStrings = false) {
        switch (rhs.GetType()) {
        case kObjectType: {
                SizeType count = rhs.data_.o.size;
                Member* lm = reinterpret_cast<Member*>(allocator.Malloc(count * sizeof(Member)));
                const typename GenericValue<Encoding,SourceAllocator>::Member* rm = rhs.GetMembersPointer();
                for (SizeType i = 0; i < count; i++) {
                    new (&lm[i].name) GenericValue(rm[i].name, allocator, copyConstStrings);
                    new (&lm[i].value) GenericValue(rm[i].value, allocator, copyConstStrings);
                }
                data_.f.flags = kObjectFlag;
                data_.o.size = data_.o.capacity = count;
                SetMembersPointer(lm);
            }
            break;
        case kArrayType: {
                SizeType count = rhs.data_.a.size;
                GenericValue* le = reinterpret_cast<GenericValue*>(allocator.Malloc(count * sizeof(GenericValue)));
                const GenericValue<Encoding,SourceAllocator>* re = rhs.GetElementsPointer();
                for (SizeType i = 0; i < count; i++)
                    new (&le[i]) GenericValue(re[i], allocator, copyConstStrings);
                data_.f.flags = kArrayFlag;
                data_.a.size = data_.a.capacity = count;
                SetElementsPointer(le);
            }
            break;
        case kStringType:
            if (rhs.data_.f.flags == kConstStringFlag && !copyConstStrings) {
                data_.f.flags = rhs.data_.f.flags;
                data_  = *reinterpret_cast<const Data*>(&rhs.data_);
            }
            else
                SetStringRaw(StringRef(rhs.GetString(), rhs.GetStringLength()), allocator);
            break;
        default:
            data_.f.flags = rhs.data_.f.flags;
            data_  = *reinterpret_cast<const Data*>(&rhs.data_);
            break;
        }
    }

    //! Constructor for boolean value.
    /*! \param b Boolean value
        \note This constructor is limited to \em real boolean values and rejects
            implicitly converted types like arbitrary pointers.  Use an explicit cast
            to \c bool, if you want to construct a boolean JSON value in such cases.
     */
#ifndef CEREAL_RAPIDJSON_DOXYGEN_RUNNING // hide SFINAE from Doxygen
    template <typename T>
    explicit GenericValue(T b, CEREAL_RAPIDJSON_ENABLEIF((internal::IsSame<bool, T>))) CEREAL_RAPIDJSON_NOEXCEPT  // See #472
#else
    explicit GenericValue(bool b) CEREAL_RAPIDJSON_NOEXCEPT
#endif
        : data_() {
            // safe-guard against failing SFINAE
            CEREAL_RAPIDJSON_STATIC_ASSERT((internal::IsSame<bool,T>::Value));
            data_.f.flags = b ? kTrueFlag : kFalseFlag;
    }

    //! Constructor for int value.
    explicit GenericValue(int i) CEREAL_RAPIDJSON_NOEXCEPT : data_() {
        data_.n.i64 = i;
        data_.f.flags = (i >= 0) ? (kNumberIntFlag | kUintFlag | kUint64Flag) : kNumberIntFlag;
    }

    //! Constructor for unsigned value.
    explicit GenericValue(unsigned u) CEREAL_RAPIDJSON_NOEXCEPT : data_() {
        data_.n.u64 = u;
        data_.f.flags = (u & 0x80000000) ? kNumberUintFlag : (kNumberUintFlag | kIntFlag | kInt64Flag);
    }

    //! Constructor for int64_t value.
    explicit GenericValue(int64_t i64) CEREAL_RAPIDJSON_NOEXCEPT : data_() {
        data_.n.i64 = i64;
        data_.f.flags = kNumberInt64Flag;
        if (i64 >= 0) {
            data_.f.flags |= kNumberUint64Flag;
            if (!(static_cast<uint64_t>(i64) & CEREAL_RAPIDJSON_UINT64_C2(0xFFFFFFFF, 0x00000000)))
                data_.f.flags |= kUintFlag;
            if (!(static_cast<uint64_t>(i64) & CEREAL_RAPIDJSON_UINT64_C2(0xFFFFFFFF, 0x80000000)))
                data_.f.flags |= kIntFlag;
        }
        else if (i64 >= static_cast<int64_t>(CEREAL_RAPIDJSON_UINT64_C2(0xFFFFFFFF, 0x80000000)))
            data_.f.flags |= kIntFlag;
    }

    //! Constructor for uint64_t value.
    explicit GenericValue(uint64_t u64) CEREAL_RAPIDJSON_NOEXCEPT : data_() {
        data_.n.u64 = u64;
        data_.f.flags = kNumberUint64Flag;
        if (!(u64 & CEREAL_RAPIDJSON_UINT64_C2(0x80000000, 0x00000000)))
            data_.f.flags |= kInt64Flag;
        if (!(u64 & CEREAL_RAPIDJSON_UINT64_C2(0xFFFFFFFF, 0x00000000)))
            data_.f.flags |= kUintFlag;
        if (!(u64 & CEREAL_RAPIDJSON_UINT64_C2(0xFFFFFFFF, 0x80000000)))
            data_.f.flags |= kIntFlag;
    }

    //! Constructor for double value.
    explicit GenericValue(double d) CEREAL_RAPIDJSON_NOEXCEPT : data_() { data_.n.d = d; data_.f.flags = kNumberDoubleFlag; }

    //! Constructor for float value.
    explicit GenericValue(float f) CEREAL_RAPIDJSON_NOEXCEPT : data_() { data_.n.d = static_cast<double>(f); data_.f.flags = kNumberDoubleFlag; }

    //! Constructor for constant string (i.e. do not make a copy of string)
    GenericValue(const Ch* s, SizeType length) CEREAL_RAPIDJSON_NOEXCEPT : data_() { SetStringRaw(StringRef(s, length)); }

    //! Constructor for constant string (i.e. do not make a copy of string)
    explicit GenericValue(StringRefType s) CEREAL_RAPIDJSON_NOEXCEPT : data_() { SetStringRaw(s); }

    //! Constructor for copy-string (i.e. do make a copy of string)
    GenericValue(const Ch* s, SizeType length, Allocator& allocator) : data_() { SetStringRaw(StringRef(s, length), allocator); }

    //! Constructor for copy-string (i.e. do make a copy of string)
    GenericValue(const Ch*s, Allocator& allocator) : data_() { SetStringRaw(StringRef(s), allocator); }

#if CEREAL_RAPIDJSON_HAS_STDSTRING
    //! Constructor for copy-string from a string object (i.e. do make a copy of string)
    /*! \note Requires the definition of the preprocessor symbol \ref CEREAL_RAPIDJSON_HAS_STDSTRING.
     */
    GenericValue(const std::basic_string<Ch>& s, Allocator& allocator) : data_() { SetStringRaw(StringRef(s), allocator); }
#endif

    //! Constructor for Array.
    /*!
        \param a An array obtained by \c GetArray().
        \note \c Array is always pass-by-value.
        \note the source array is moved into this value and the sourec array becomes empty.
    */
    GenericValue(Array a) CEREAL_RAPIDJSON_NOEXCEPT : data_(a.value_.data_) {
        a.value_.data_ = Data();
        a.value_.data_.f.flags = kArrayFlag;
    }

    //! Constructor for Object.
    /*!
        \param o An object obtained by \c GetObject().
        \note \c Object is always pass-by-value.
        \note the source object is moved into this value and the sourec object becomes empty.
    */
    GenericValue(Object o) CEREAL_RAPIDJSON_NOEXCEPT : data_(o.value_.data_) {
        o.value_.data_ = Data();
        o.value_.data_.f.flags = kObjectFlag;
    }

    //! Destructor.
    /*! Need to destruct elements of array, members of object, or copy-string.
    */
    ~GenericValue() {
        if (Allocator::kNeedFree) { // Shortcut by Allocator's trait
            switch(data_.f.flags) {
            case kArrayFlag:
                {
                    GenericValue* e = GetElementsPointer();
                    for (GenericValue* v = e; v != e + data_.a.size; ++v)
                        v->~GenericValue();
                    Allocator::Free(e);
                }
                break;

            case kObjectFlag:
                for (MemberIterator m = MemberBegin(); m != MemberEnd(); ++m)
                    m->~Member();
                Allocator::Free(GetMembersPointer());
                break;

            case kCopyStringFlag:
                Allocator::Free(const_cast<Ch*>(GetStringPointer()));
                break;

            default:
                break;  // Do nothing for other types.
            }
        }
    }

    //@}

    //!@name Assignment operators
    //@{

    //! Assignment with move semantics.
    /*! \param rhs Source of the assignment. It will become a null value after assignment.
    */
    GenericValue& operator=(GenericValue& rhs) CEREAL_RAPIDJSON_NOEXCEPT {
        if (CEREAL_RAPIDJSON_LIKELY(this != &rhs)) {
            this->~GenericValue();
            RawAssign(rhs);
        }
        return *this;
    }

#if CEREAL_RAPIDJSON_HAS_CXX11_RVALUE_REFS
    //! Move assignment in C++11
    GenericValue& operator=(GenericValue&& rhs) CEREAL_RAPIDJSON_NOEXCEPT {
        return *this = rhs.Move();
    }
#endif

    //! Assignment of constant string reference (no copy)
    /*! \param str Constant string reference to be assigned
        \note This overload is needed to avoid clashes with the generic primitive type assignment overload below.
        \see GenericStringRef, operator=(T)
    */
    GenericValue& operator=(StringRefType str) CEREAL_RAPIDJSON_NOEXCEPT {
        GenericValue s(str);
        return *this = s;
    }

    //! Assignment with primitive types.
    /*! \tparam T Either \ref Type, \c int, \c unsigned, \c int64_t, \c uint64_t
        \param value The value to be assigned.

        \note The source type \c T explicitly disallows all pointer types,
            especially (\c const) \ref Ch*.  This helps avoiding implicitly
            referencing character strings with insufficient lifetime, use
            \ref SetString(const Ch*, Allocator&) (for copying) or
            \ref StringRef() (to explicitly mark the pointer as constant) instead.
            All other pointer types would implicitly convert to \c bool,
            use \ref SetBool() instead.
    */
    template <typename T>
    CEREAL_RAPIDJSON_DISABLEIF_RETURN((internal::IsPointer<T>), (GenericValue&))
    operator=(T value) {
        GenericValue v(value);
        return *this = v;
    }

    //! Deep-copy assignment from Value
    /*! Assigns a \b copy of the Value to the current Value object
        \tparam SourceAllocator Allocator type of \c rhs
        \param rhs Value to copy from (read-only)
        \param allocator Allocator to use for copying
        \param copyConstStrings Force copying of constant strings (e.g. referencing an in-situ buffer)
     */
    template <typename SourceAllocator>
    GenericValue& CopyFrom(const GenericValue<Encoding, SourceAllocator>& rhs, Allocator& allocator, bool copyConstStrings = false) {
        CEREAL_RAPIDJSON_ASSERT(static_cast<void*>(this) != static_cast<void const*>(&rhs));
        this->~GenericValue();
        new (this) GenericValue(rhs, allocator, copyConstStrings);
        return *this;
    }

    //! Exchange the contents of this value with those of other.
    /*!
        \param other Another value.
        \note Constant complexity.
    */
    GenericValue& Swap(GenericValue& other) CEREAL_RAPIDJSON_NOEXCEPT {
        GenericValue temp;
        temp.RawAssign(*this);
        RawAssign(other);
        other.RawAssign(temp);
        return *this;
    }

    //! free-standing swap function helper
    /*!
        Helper function to enable support for common swap implementation pattern based on \c std::swap:
        \code
        void swap(MyClass& a, MyClass& b) {
            using std::swap;
            swap(a.value, b.value);
            // ...
        }
        \endcode
        \see Swap()
     */
    friend inline void swap(GenericValue& a, GenericValue& b) CEREAL_RAPIDJSON_NOEXCEPT { a.Swap(b); }

    //! Prepare Value for move semantics
    /*! \return *this */
    GenericValue& Move() CEREAL_RAPIDJSON_NOEXCEPT { return *this; }
    //@}

    //!@name Equal-to and not-equal-to operators
    //@{
    //! Equal-to operator
    /*!
        \note If an object contains duplicated named member, comparing equality with any object is always \c false.
        \note Complexity is quadratic in Object's member number and linear for the rest (number of all values in the subtree and total lengths of all strings).
    */
    template <typename SourceAllocator>
    bool operator==(const GenericValue<Encoding, SourceAllocator>& rhs) const {
        typedef GenericValue<Encoding, SourceAllocator> RhsType;
        if (GetType() != rhs.GetType())
            return false;

        switch (GetType()) {
        case kObjectType: // Warning: O(n^2) inner-loop
            if (data_.o.size != rhs.data_.o.size)
                return false;
            for (ConstMemberIterator lhsMemberItr = MemberBegin(); lhsMemberItr != MemberEnd(); ++lhsMemberItr) {
                typename RhsType::ConstMemberIterator rhsMemberItr = rhs.FindMember(lhsMemberItr->name);
                if (rhsMemberItr == rhs.MemberEnd() || lhsMemberItr->value != rhsMemberItr->value)
                    return false;
            }
            return true;

        case kArrayType:
            if (data_.a.size != rhs.data_.a.size)
                return false;
            for (SizeType i = 0; i < data_.a.size; i++)
                if ((*this)[i] != rhs[i])
                    return false;
            return true;

        case kStringType:
            return StringEqual(rhs);

        case kNumberType:
            if (IsDouble() || rhs.IsDouble()) {
                double a = GetDouble();     // May convert from integer to double.
                double b = rhs.GetDouble(); // Ditto
                return a >= b && a <= b;    // Prevent -Wfloat-equal
            }
            else
                return data_.n.u64 == rhs.data_.n.u64;

        default:
            return true;
        }
    }

    //! Equal-to operator with const C-string pointer
    bool operator==(const Ch* rhs) const { return *this == GenericValue(StringRef(rhs)); }

#if CEREAL_RAPIDJSON_HAS_STDSTRING
    //! Equal-to operator with string object
    /*! \note Requires the definition of the preprocessor symbol \ref CEREAL_RAPIDJSON_HAS_STDSTRING.
     */
    bool operator==(const std::basic_string<Ch>& rhs) const { return *this == GenericValue(StringRef(rhs)); }
#endif

    //! Equal-to operator with primitive types
    /*! \tparam T Either \ref Type, \c int, \c unsigned, \c int64_t, \c uint64_t, \c double, \c true, \c false
    */
    template <typename T> CEREAL_RAPIDJSON_DISABLEIF_RETURN((internal::OrExpr<internal::IsPointer<T>,internal::IsGenericValue<T> >), (bool)) operator==(const T& rhs) const { return *this == GenericValue(rhs); }

    //! Not-equal-to operator
    /*! \return !(*this == rhs)
     */
    template <typename SourceAllocator>
    bool operator!=(const GenericValue<Encoding, SourceAllocator>& rhs) const { return !(*this == rhs); }

    //! Not-equal-to operator with const C-string pointer
    bool operator!=(const Ch* rhs) const { return !(*this == rhs); }

    //! Not-equal-to operator with arbitrary types
    /*! \return !(*this == rhs)
     */
    template <typename T> CEREAL_RAPIDJSON_DISABLEIF_RETURN((internal::IsGenericValue<T>), (bool)) operator!=(const T& rhs) const { return !(*this == rhs); }

    //! Equal-to operator with arbitrary types (symmetric version)
    /*! \return (rhs == lhs)
     */
    template <typename T> friend CEREAL_RAPIDJSON_DISABLEIF_RETURN((internal::IsGenericValue<T>), (bool)) operator==(const T& lhs, const GenericValue& rhs) { return rhs == lhs; }

    //! Not-Equal-to operator with arbitrary types (symmetric version)
    /*! \return !(rhs == lhs)
     */
    template <typename T> friend CEREAL_RAPIDJSON_DISABLEIF_RETURN((internal::IsGenericValue<T>), (bool)) operator!=(const T& lhs, const GenericValue& rhs) { return !(rhs == lhs); }
    //@}

    //!@name Type
    //@{

    Type GetType()  const { return static_cast<Type>(data_.f.flags & kTypeMask); }
    bool IsNull()   const { return data_.f.flags == kNullFlag; }
    bool IsFalse()  const { return data_.f.flags == kFalseFlag; }
    bool IsTrue()   const { return data_.f.flags == kTrueFlag; }
    bool IsBool()   const { return (data_.f.flags & kBoolFlag) != 0; }
    bool IsObject() const { return data_.f.flags == kObjectFlag; }
    bool IsArray()  const { return data_.f.flags == kArrayFlag; }
    bool IsNumber() const { return (data_.f.flags & kNumberFlag) != 0; }
    bool IsInt()    const { return (data_.f.flags & kIntFlag) != 0; }
    bool IsUint()   const { return (data_.f.flags & kUintFlag) != 0; }
    bool IsInt64()  const { return (data_.f.flags & kInt64Flag) != 0; }
    bool IsUint64() const { return (data_.f.flags & kUint64Flag) != 0; }
    bool IsDouble() const { return (data_.f.flags & kDoubleFlag) != 0; }
    bool IsString() const { return (data_.f.flags & kStringFlag) != 0; }

    // Checks whether a number can be losslessly converted to a double.
    bool IsLosslessDouble() const {
        if (!IsNumber()) return false;
        if (IsUint64()) {
            uint64_t u = GetUint64();
            volatile double d = static_cast<double>(u);
            return (d >= 0.0)
                && (d < static_cast<double>((std::numeric_limits<uint64_t>::max)()))
                && (u == static_cast<uint64_t>(d));
        }
        if (IsInt64()) {
            int64_t i = GetInt64();
            volatile double d = static_cast<double>(i);
            return (d >= static_cast<double>((std::numeric_limits<int64_t>::min)()))
                && (d < static_cast<double>((std::numeric_limits<int64_t>::max)()))
                && (i == static_cast<int64_t>(d));
        }
        return true; // double, int, uint are always lossless
    }

    // Checks whether a number is a float (possible lossy).
    bool IsFloat() const  {
        if ((data_.f.flags & kDoubleFlag) == 0)
            return false;
        double d = GetDouble();
        return d >= -3.4028234e38 && d <= 3.4028234e38;
    }
    // Checks whether a number can be losslessly converted to a float.
    bool IsLosslessFloat() const {
        if (!IsNumber()) return false;
        double a = GetDouble();
        if (a < static_cast<double>(-(std::numeric_limits<float>::max)())
                || a > static_cast<double>((std::numeric_limits<float>::max)()))
            return false;
        double b = static_cast<double>(static_cast<float>(a));
        return a >= b && a <= b;    // Prevent -Wfloat-equal
    }

    //@}

    //!@name Null
    //@{

    GenericValue& SetNull() { this->~GenericValue(); new (this) GenericValue(); return *this; }

    //@}

    //!@name Bool
    //@{

    bool GetBool() const { CEREAL_RAPIDJSON_ASSERT(IsBool()); return data_.f.flags == kTrueFlag; }
    //!< Set boolean value
    /*! \post IsBool() == true */
    GenericValue& SetBool(bool b) { this->~GenericValue(); new (this) GenericValue(b); return *this; }

    //@}

    //!@name Object
    //@{

    //! Set this value as an empty object.
    /*! \post IsObject() == true */
    GenericValue& SetObject() { this->~GenericValue(); new (this) GenericValue(kObjectType); return *this; }

    //! Get the number of members in the object.
    SizeType MemberCount() const { CEREAL_RAPIDJSON_ASSERT(IsObject()); return data_.o.size; }

    //! Get the capacity of object.
    SizeType MemberCapacity() const { CEREAL_RAPIDJSON_ASSERT(IsObject()); return data_.o.capacity; }

    //! Check whether the object is empty.
    bool ObjectEmpty() const { CEREAL_RAPIDJSON_ASSERT(IsObject()); return data_.o.size == 0; }

    //! Get a value from an object associated with the name.
    /*! \pre IsObject() == true
        \tparam T Either \c Ch or \c const \c Ch (template used for disambiguation with \ref operator[](SizeType))
        \note In version 0.1x, if the member is not found, this function returns a null value. This makes issue 7.
        Since 0.2, if the name is not correct, it will assert.
        If user is unsure whether a member exists, user should use HasMember() first.
        A better approach is to use FindMember().
        \note Linear time complexity.
    */
    template <typename T>
    CEREAL_RAPIDJSON_DISABLEIF_RETURN((internal::NotExpr<internal::IsSame<typename internal::RemoveConst<T>::Type, Ch> >),(GenericValue&)) operator[](T* name) {
        GenericValue n(StringRef(name));
        return (*this)[n];
    }
    template <typename T>
    CEREAL_RAPIDJSON_DISABLEIF_RETURN((internal::NotExpr<internal::IsSame<typename internal::RemoveConst<T>::Type, Ch> >),(const GenericValue&)) operator[](T* name) const { return const_cast<GenericValue&>(*this)[name]; }

    //! Get a value from an object associated with the name.
    /*! \pre IsObject() == true
        \tparam SourceAllocator Allocator of the \c name value

        \note Compared to \ref operator[](T*), this version is faster because it does not need a StrLen().
        And it can also handle strings with embedded null characters.

        \note Linear time complexity.
    */
    template <typename SourceAllocator>
    GenericValue& operator[](const GenericValue<Encoding, SourceAllocator>& name) {
        MemberIterator member = FindMember(name);
        if (member != MemberEnd())
            return member->value;
        else {
            CEREAL_RAPIDJSON_ASSERT(false);    // see above note

            // This will generate -Wexit-time-destructors in clang
            // static GenericValue NullValue;
            // return NullValue;

            // Use static buffer and placement-new to prevent destruction
            static char buffer[sizeof(GenericValue)];
            return *new (buffer) GenericValue();
        }
    }
    template <typename SourceAllocator>
    const GenericValue& operator[](const GenericValue<Encoding, SourceAllocator>& name) const { return const_cast<GenericValue&>(*this)[name]; }

#if CEREAL_RAPIDJSON_HAS_STDSTRING
    //! Get a value from an object associated with name (string object).
    GenericValue& operator[](const std::basic_string<Ch>& name) { return (*this)[GenericValue(StringRef(name))]; }
    const GenericValue& operator[](const std::basic_string<Ch>& name) const { return (*this)[GenericValue(StringRef(name))]; }
#endif

    //! Const member iterator
    /*! \pre IsObject() == true */
    ConstMemberIterator MemberBegin() const { CEREAL_RAPIDJSON_ASSERT(IsObject()); return ConstMemberIterator(GetMembersPointer()); }
    //! Const \em past-the-end member iterator
    /*! \pre IsObject() == true */
    ConstMemberIterator MemberEnd() const   { CEREAL_RAPIDJSON_ASSERT(IsObject()); return ConstMemberIterator(GetMembersPointer() + data_.o.size); }
    //! Member iterator
    /*! \pre IsObject() == true */
    MemberIterator MemberBegin()            { CEREAL_RAPIDJSON_ASSERT(IsObject()); return MemberIterator(GetMembersPointer()); }
    //! \em Past-the-end member iterator
    /*! \pre IsObject() == true */
    MemberIterator MemberEnd()              { CEREAL_RAPIDJSON_ASSERT(IsObject()); return MemberIterator(GetMembersPointer() + data_.o.size); }

    //! Request the object to have enough capacity to store members.
    /*! \param newCapacity  The capacity that the object at least need to have.
        \param allocator    Allocator for reallocating memory. It must be the same one as used before. Commonly use GenericDocument::GetAllocator().
        \return The value itself for fluent API.
        \note Linear time complexity.
    */
    GenericValue& MemberReserve(SizeType newCapacity, Allocator &allocator) {
        CEREAL_RAPIDJSON_ASSERT(IsObject());
        if (newCapacity > data_.o.capacity) {
            SetMembersPointer(reinterpret_cast<Member*>(allocator.Realloc(GetMembersPointer(), data_.o.capacity * sizeof(Member), newCapacity * sizeof(Member))));
            data_.o.capacity = newCapacity;
        }
        return *this;
    }

    //! Check whether a member exists in the object.
    /*!
        \param name Member name to be searched.
        \pre IsObject() == true
        \return Whether a member with that name exists.
        \note It is better to use FindMember() directly if you need the obtain the value as well.
        \note Linear time complexity.
    */
    bool HasMember(const Ch* name) const { return FindMember(name) != MemberEnd(); }

#if CEREAL_RAPIDJSON_HAS_STDSTRING
    //! Check whether a member exists in the object with string object.
    /*!
        \param name Member name to be searched.
        \pre IsObject() == true
        \return Whether a member with that name exists.
        \note It is better to use FindMember() directly if you need the obtain the value as well.
        \note Linear time complexity.
    */
    bool HasMember(const std::basic_string<Ch>& name) const { return FindMember(name) != MemberEnd(); }
#endif

    //! Check whether a member exists in the object with GenericValue name.
    /*!
        This version is faster because it does not need a StrLen(). It can also handle string with null character.
        \param name Member name to be searched.
        \pre IsObject() == true
        \return Whether a member with that name exists.
        \note It is better to use FindMember() directly if you need the obtain the value as well.
        \note Linear time complexity.
    */
    template <typename SourceAllocator>
    bool HasMember(const GenericValue<Encoding, SourceAllocator>& name) const { return FindMember(name) != MemberEnd(); }

    //! Find member by name.
    /*!
        \param name Member name to be searched.
        \pre IsObject() == true
        \return Iterator to member, if it exists.
            Otherwise returns \ref MemberEnd().

        \note Earlier versions of Rapidjson returned a \c NULL pointer, in case
            the requested member doesn't exist. For consistency with e.g.
            \c std::map, this has been changed to MemberEnd() now.
        \note Linear time complexity.
    */
    MemberIterator FindMember(const Ch* name) {
        GenericValue n(StringRef(name));
        return FindMember(n);
    }

    ConstMemberIterator FindMember(const Ch* name) const { return const_cast<GenericValue&>(*this).FindMember(name); }

    //! Find member by name.
    /*!
        This version is faster because it does not need a StrLen(). It can also handle string with null character.
        \param name Member name to be searched.
        \pre IsObject() == true
        \return Iterator to member, if it exists.
            Otherwise returns \ref MemberEnd().

        \note Earlier versions of Rapidjson returned a \c NULL pointer, in case
            the requested member doesn't exist. For consistency with e.g.
            \c std::map, this has been changed to MemberEnd() now.
        \note Linear time complexity.
    */
    template <typename SourceAllocator>
    MemberIterator FindMember(const GenericValue<Encoding, SourceAllocator>& name) {
        CEREAL_RAPIDJSON_ASSERT(IsObject());
        CEREAL_RAPIDJSON_ASSERT(name.IsString());
        MemberIterator member = MemberBegin();
        for ( ; member != MemberEnd(); ++member)
            if (name.StringEqual(member->name))
                break;
        return member;
    }
    template <typename SourceAllocator> ConstMemberIterator FindMember(const GenericValue<Encoding, SourceAllocator>& name) const { return const_cast<GenericValue&>(*this).FindMember(name); }

#if CEREAL_RAPIDJSON_HAS_STDSTRING
    //! Find member by string object name.
    /*!
        \param name Member name to be searched.
        \pre IsObject() == true
        \return Iterator to member, if it exists.
            Otherwise returns \ref MemberEnd().
    */
    MemberIterator FindMember(const std::basic_string<Ch>& name) { return FindMember(GenericValue(StringRef(name))); }
    ConstMemberIterator FindMember(const std::basic_string<Ch>& name) const { return FindMember(GenericValue(StringRef(name))); }
#endif

    //! Add a member (name-value pair) to the object.
    /*! \param name A string value as name of member.
        \param value Value of any type.
        \param allocator    Allocator for reallocating memory. It must be the same one as used before. Commonly use GenericDocument::GetAllocator().
        \return The value itself for fluent API.
        \note The ownership of \c name and \c value will be transferred to this object on success.
        \pre  IsObject() && name.IsString()
        \post name.IsNull() && value.IsNull()
        \note Amortized Constant time complexity.
    */
    GenericValue& AddMember(GenericValue& name, GenericValue& value, Allocator& allocator) {
        CEREAL_RAPIDJSON_ASSERT(IsObject());
        CEREAL_RAPIDJSON_ASSERT(name.IsString());

        ObjectData& o = data_.o;
        if (o.size >= o.capacity)
            MemberReserve(o.capacity == 0 ? kDefaultObjectCapacity : (o.capacity + (o.capacity + 1) / 2), allocator);
        Member* members = GetMembersPointer();
        members[o.size].name.RawAssign(name);
        members[o.size].value.RawAssign(value);
        o.size++;
        return *this;
    }

    //! Add a constant string value as member (name-value pair) to the object.
    /*! \param name A string value as name of member.
        \param value constant string reference as value of member.
        \param allocator    Allocator for reallocating memory. It must be the same one as used before. Commonly use GenericDocument::GetAllocator().
        \return The value itself for fluent API.
        \pre  IsObject()
        \note This overload is needed to avoid clashes with the generic primitive type AddMember(GenericValue&,T,Allocator&) overload below.
        \note Amortized Constant time complexity.
    */
    GenericValue& AddMember(GenericValue& name, StringRefType value, Allocator& allocator) {
        GenericValue v(value);
        return AddMember(name, v, allocator);
    }

#if CEREAL_RAPIDJSON_HAS_STDSTRING
    //! Add a string object as member (name-value pair) to the object.
    /*! \param name A string value as name of member.
        \param value constant string reference as value of member.
        \param allocator    Allocator for reallocating memory. It must be the same one as used before. Commonly use GenericDocument::GetAllocator().
        \return The value itself for fluent API.
        \pre  IsObject()
        \note This overload is needed to avoid clashes with the generic primitive type AddMember(GenericValue&,T,Allocator&) overload below.
        \note Amortized Constant time complexity.
    */
    GenericValue& AddMember(GenericValue& name, std::basic_string<Ch>& value, Allocator& allocator) {
        GenericValue v(value, allocator);
        return AddMember(name, v, allocator);
    }
#endif

    //! Add any primitive value as member (name-value pair) to the object.
    /*! \tparam T Either \ref Type, \c int, \c unsigned, \c int64_t, \c uint64_t
        \param name A string value as name of member.
        \param value Value of primitive type \c T as value of member
        \param allocator Allocator for reallocating memory. Commonly use GenericDocument::GetAllocator().
        \return The value itself for fluent API.
        \pre  IsObject()

        \note The source type \c T explicitly disallows all pointer types,
            especially (\c const) \ref Ch*.  This helps avoiding implicitly
            referencing character strings with insufficient lifetime, use
            \ref AddMember(StringRefType, GenericValue&, Allocator&) or \ref
            AddMember(StringRefType, StringRefType, Allocator&).
            All other pointer types would implicitly convert to \c bool,
            use an explicit cast instead, if needed.
        \note Amortized Constant time complexity.
    */
    template <typename T>
    CEREAL_RAPIDJSON_DISABLEIF_RETURN((internal::OrExpr<internal::IsPointer<T>, internal::IsGenericValue<T> >), (GenericValue&))
    AddMember(GenericValue& name, T value, Allocator& allocator) {
        GenericValue v(value);
        return AddMember(name, v, allocator);
    }

#if CEREAL_RAPIDJSON_HAS_CXX11_RVALUE_REFS
    GenericValue& AddMember(GenericValue&& name, GenericValue&& value, Allocator& allocator) {
        return AddMember(name, value, allocator);
    }
    GenericValue& AddMember(GenericValue&& name, GenericValue& value, Allocator& allocator) {
        return AddMember(name, value, allocator);
    }
    GenericValue& AddMember(GenericValue& name, GenericValue&& value, Allocator& allocator) {
        return AddMember(name, value, allocator);
    }
    GenericValue& AddMember(StringRefType name, GenericValue&& value, Allocator& allocator) {
        GenericValue n(name);
        return AddMember(n, value, allocator);
    }
#endif // CEREAL_RAPIDJSON_HAS_CXX11_RVALUE_REFS


    //! Add a member (name-value pair) to the object.
    /*! \param name A constant string reference as name of member.
        \param value Value of any type.
        \param allocator    Allocator for reallocating memory. It must be the same one as used before. Commonly use GenericDocument::GetAllocator().
        \return The value itself for fluent API.
        \note The ownership of \c value will be transferred to this object on success.
        \pre  IsObject()
        \post value.IsNull()
        \note Amortized Constant time complexity.
    */
    GenericValue& AddMember(StringRefType name, GenericValue& value, Allocator& allocator) {
        GenericValue n(name);
        return AddMember(n, value, allocator);
    }

    //! Add a constant string value as member (name-value pair) to the object.
    /*! \param name A constant string reference as name of member.
        \param value constant string reference as value of member.
        \param allocator    Allocator for reallocating memory. It must be the same one as used before. Commonly use GenericDocument::GetAllocator().
        \return The value itself for fluent API.
        \pre  IsObject()
        \note This overload is needed to avoid clashes with the generic primitive type AddMember(StringRefType,T,Allocator&) overload below.
        \note Amortized Constant time complexity.
    */
    GenericValue& AddMember(StringRefType name, StringRefType value, Allocator& allocator) {
        GenericValue v(value);
        return AddMember(name, v, allocator);
    }

    //! Add any primitive value as member (name-value pair) to the object.
    /*! \tparam T Either \ref Type, \c int, \c unsigned, \c int64_t, \c uint64_t
        \param name A constant string reference as name of member.
        \param value Value of primitive type \c T as value of member
        \param allocator Allocator for reallocating memory. Commonly use GenericDocument::GetAllocator().
        \return The value itself for fluent API.
        \pre  IsObject()

        \note The source type \c T explicitly disallows all pointer types,
            especially (\c const) \ref Ch*.  This helps avoiding implicitly
            referencing character strings with insufficient lifetime, use
            \ref AddMember(StringRefType, GenericValue&, Allocator&) or \ref
            AddMember(StringRefType, StringRefType, Allocator&).
            All other pointer types would implicitly convert to \c bool,
            use an explicit cast instead, if needed.
        \note Amortized Constant time complexity.
    */
    template <typename T>
    CEREAL_RAPIDJSON_DISABLEIF_RETURN((internal::OrExpr<internal::IsPointer<T>, internal::IsGenericValue<T> >), (GenericValue&))
    AddMember(StringRefType name, T value, Allocator& allocator) {
        GenericValue n(name);
        return AddMember(n, value, allocator);
    }

    //! Remove all members in the object.
    /*! This function do not deallocate memory in the object, i.e. the capacity is unchanged.
        \note Linear time complexity.
    */
    void RemoveAllMembers() {
        CEREAL_RAPIDJSON_ASSERT(IsObject());
        for (MemberIterator m = MemberBegin(); m != MemberEnd(); ++m)
            m->~Member();
        data_.o.size = 0;
    }

    //! Remove a member in object by its name.
    /*! \param name Name of member to be removed.
        \return Whether the member existed.
        \note This function may reorder the object members. Use \ref
            EraseMember(ConstMemberIterator) if you need to preserve the
            relative order of the remaining members.
        \note Linear time complexity.
    */
    bool RemoveMember(const Ch* name) {
        GenericValue n(StringRef(name));
        return RemoveMember(n);
    }

#if CEREAL_RAPIDJSON_HAS_STDSTRING
    bool RemoveMember(const std::basic_string<Ch>& name) { return RemoveMember(GenericValue(StringRef(name))); }
#endif

    template <typename SourceAllocator>
    bool RemoveMember(const GenericValue<Encoding, SourceAllocator>& name) {
        MemberIterator m = FindMember(name);
        if (m != MemberEnd()) {
            RemoveMember(m);
            return true;
        }
        else
            return false;
    }

    //! Remove a member in object by iterator.
    /*! \param m member iterator (obtained by FindMember() or MemberBegin()).
        \return the new iterator after removal.
        \note This function may reorder the object members. Use \ref
            EraseMember(ConstMemberIterator) if you need to preserve the
            relative order of the remaining members.
        \note Constant time complexity.
    */
    MemberIterator RemoveMember(MemberIterator m) {
        CEREAL_RAPIDJSON_ASSERT(IsObject());
        CEREAL_RAPIDJSON_ASSERT(data_.o.size > 0);
        CEREAL_RAPIDJSON_ASSERT(GetMembersPointer() != 0);
        CEREAL_RAPIDJSON_ASSERT(m >= MemberBegin() && m < MemberEnd());

        MemberIterator last(GetMembersPointer() + (data_.o.size - 1));
        if (data_.o.size > 1 && m != last)
            *m = *last; // Move the last one to this place
        else
            m->~Member(); // Only one left, just destroy
        --data_.o.size;
        return m;
    }

    //! Remove a member from an object by iterator.
    /*! \param pos iterator to the member to remove
        \pre IsObject() == true && \ref MemberBegin() <= \c pos < \ref MemberEnd()
        \return Iterator following the removed element.
            If the iterator \c pos refers to the last element, the \ref MemberEnd() iterator is returned.
        \note This function preserves the relative order of the remaining object
            members. If you do not need this, use the more efficient \ref RemoveMember(MemberIterator).
        \note Linear time complexity.
    */
    MemberIterator EraseMember(ConstMemberIterator pos) {
        return EraseMember(pos, pos +1);
    }

    //! Remove members in the range [first, last) from an object.
    /*! \param first iterator to the first member to remove
        \param last  iterator following the last member to remove
        \pre IsObject() == true && \ref MemberBegin() <= \c first <= \c last <= \ref MemberEnd()
        \return Iterator following the last removed element.
        \note This function preserves the relative order of the remaining object
            members.
        \note Linear time complexity.
    */
    MemberIterator EraseMember(ConstMemberIterator first, ConstMemberIterator last) {
        CEREAL_RAPIDJSON_ASSERT(IsObject());
        CEREAL_RAPIDJSON_ASSERT(data_.o.size > 0);
        CEREAL_RAPIDJSON_ASSERT(GetMembersPointer() != 0);
        CEREAL_RAPIDJSON_ASSERT(first >= MemberBegin());
        CEREAL_RAPIDJSON_ASSERT(first <= last);
        CEREAL_RAPIDJSON_ASSERT(last <= MemberEnd());

        MemberIterator pos = MemberBegin() + (first - MemberBegin());
        for (MemberIterator itr = pos; itr != last; ++itr)
            itr->~Member();
        std::memmove(static_cast<void*>(&*pos), &*last, static_cast<size_t>(MemberEnd() - last) * sizeof(Member));
        data_.o.size -= static_cast<SizeType>(last - first);
        return pos;
    }

    //! Erase a member in object by its name.
    /*! \param name Name of member to be removed.
        \return Whether the member existed.
        \note Linear time complexity.
    */
    bool EraseMember(const Ch* name) {
        GenericValue n(StringRef(name));
        return EraseMember(n);
    }

#if CEREAL_RAPIDJSON_HAS_STDSTRING
    bool EraseMember(const std::basic_string<Ch>& name) { return EraseMember(GenericValue(StringRef(name))); }
#endif

    template <typename SourceAllocator>
    bool EraseMember(const GenericValue<Encoding, SourceAllocator>& name) {
        MemberIterator m = FindMember(name);
        if (m != MemberEnd()) {
            EraseMember(m);
            return true;
        }
        else
            return false;
    }

    Object GetObject() { CEREAL_RAPIDJSON_ASSERT(IsObject()); return Object(*this); }
    ConstObject GetObject() const { CEREAL_RAPIDJSON_ASSERT(IsObject()); return ConstObject(*this); }

    //@}

    //!@name Array
    //@{

    //! Set this value as an empty array.
    /*! \post IsArray == true */
    GenericValue& SetArray() { this->~GenericValue(); new (this) GenericValue(kArrayType); return *this; }

    //! Get the number of elements in array.
    SizeType Size() const { CEREAL_RAPIDJSON_ASSERT(IsArray()); return data_.a.size; }

    //! Get the capacity of array.
    SizeType Capacity() const { CEREAL_RAPIDJSON_ASSERT(IsArray()); return data_.a.capacity; }

    //! Check whether the array is empty.
    bool Empty() const { CEREAL_RAPIDJSON_ASSERT(IsArray()); return data_.a.size == 0; }

    //! Remove all elements in the array.
    /*! This function do not deallocate memory in the array, i.e. the capacity is unchanged.
        \note Linear time complexity.
    */
    void Clear() {
        CEREAL_RAPIDJSON_ASSERT(IsArray());
        GenericValue* e = GetElementsPointer();
        for (GenericValue* v = e; v != e + data_.a.size; ++v)
            v->~GenericValue();
        data_.a.size = 0;
    }

    //! Get an element from array by index.
    /*! \pre IsArray() == true
        \param index Zero-based index of element.
        \see operator[](T*)
    */
    GenericValue& operator[](SizeType index) {
        CEREAL_RAPIDJSON_ASSERT(IsArray());
        CEREAL_RAPIDJSON_ASSERT(index < data_.a.size);
        return GetElementsPointer()[index];
    }
    const GenericValue& operator[](SizeType index) const { return const_cast<GenericValue&>(*this)[index]; }

    //! Element iterator
    /*! \pre IsArray() == true */
    ValueIterator Begin() { CEREAL_RAPIDJSON_ASSERT(IsArray()); return GetElementsPointer(); }
    //! \em Past-the-end element iterator
    /*! \pre IsArray() == true */
    ValueIterator End() { CEREAL_RAPIDJSON_ASSERT(IsArray()); return GetElementsPointer() + data_.a.size; }
    //! Constant element iterator
    /*! \pre IsArray() == true */
    ConstValueIterator Begin() const { return const_cast<GenericValue&>(*this).Begin(); }
    //! Constant \em past-the-end element iterator
    /*! \pre IsArray() == true */
    ConstValueIterator End() const { return const_cast<GenericValue&>(*this).End(); }

    //! Request the array to have enough capacity to store elements.
    /*! \param newCapacity  The capacity that the array at least need to have.
        \param allocator    Allocator for reallocating memory. It must be the same one as used before. Commonly use GenericDocument::GetAllocator().
        \return The value itself for fluent API.
        \note Linear time complexity.
    */
    GenericValue& Reserve(SizeType newCapacity, Allocator &allocator) {
        CEREAL_RAPIDJSON_ASSERT(IsArray());
        if (newCapacity > data_.a.capacity) {
            SetElementsPointer(reinterpret_cast<GenericValue*>(allocator.Realloc(GetElementsPointer(), data_.a.capacity * sizeof(GenericValue), newCapacity * sizeof(GenericValue))));
            data_.a.capacity = newCapacity;
        }
        return *this;
    }

    //! Append a GenericValue at the end of the array.
    /*! \param value        Value to be appended.
        \param allocator    Allocator for reallocating memory. It must be the same one as used before. Commonly use GenericDocument::GetAllocator().
        \pre IsArray() == true
        \post value.IsNull() == true
        \return The value itself for fluent API.
        \note The ownership of \c value will be transferred to this array on success.
        \note If the number of elements to be appended is known, calls Reserve() once first may be more efficient.
        \note Amortized constant time complexity.
    */
    GenericValue& PushBack(GenericValue& value, Allocator& allocator) {
        CEREAL_RAPIDJSON_ASSERT(IsArray());
        if (data_.a.size >= data_.a.capacity)
            Reserve(data_.a.capacity == 0 ? kDefaultArrayCapacity : (data_.a.capacity + (data_.a.capacity + 1) / 2), allocator);
        GetElementsPointer()[data_.a.size++].RawAssign(value);
        return *this;
    }

#if CEREAL_RAPIDJSON_HAS_CXX11_RVALUE_REFS
    GenericValue& PushBack(GenericValue&& value, Allocator& allocator) {
        return PushBack(value, allocator);
    }
#endif // CEREAL_RAPIDJSON_HAS_CXX11_RVALUE_REFS

    //! Append a constant string reference at the end of the array.
    /*! \param value        Constant string reference to be appended.
        \param allocator    Allocator for reallocating memory. It must be the same one used previously. Commonly use GenericDocument::GetAllocator().
        \pre IsArray() == true
        \return The value itself for fluent API.
        \note If the number of elements to be appended is known, calls Reserve() once first may be more efficient.
        \note Amortized constant time complexity.
        \see GenericStringRef
    */
    GenericValue& PushBack(StringRefType value, Allocator& allocator) {
        return (*this).template PushBack<StringRefType>(value, allocator);
    }

    //! Append a primitive value at the end of the array.
    /*! \tparam T Either \ref Type, \c int, \c unsigned, \c int64_t, \c uint64_t
        \param value Value of primitive type T to be appended.
        \param allocator    Allocator for reallocating memory. It must be the same one as used before. Commonly use GenericDocument::GetAllocator().
        \pre IsArray() == true
        \return The value itself for fluent API.
        \note If the number of elements to be appended is known, calls Reserve() once first may be more efficient.

        \note The source type \c T explicitly disallows all pointer types,
            especially (\c const) \ref Ch*.  This helps avoiding implicitly
            referencing character strings with insufficient lifetime, use
            \ref PushBack(GenericValue&, Allocator&) or \ref
            PushBack(StringRefType, Allocator&).
            All other pointer types would implicitly convert to \c bool,
            use an explicit cast instead, if needed.
        \note Amortized constant time complexity.
    */
    template <typename T>
    CEREAL_RAPIDJSON_DISABLEIF_RETURN((internal::OrExpr<internal::IsPointer<T>, internal::IsGenericValue<T> >), (GenericValue&))
    PushBack(T value, Allocator& allocator) {
        GenericValue v(value);
        return PushBack(v, allocator);
    }

    //! Remove the last element in the array.
    /*!
        \note Constant time complexity.
    */
    GenericValue& PopBack() {
        CEREAL_RAPIDJSON_ASSERT(IsArray());
        CEREAL_RAPIDJSON_ASSERT(!Empty());
        GetElementsPointer()[--data_.a.size].~GenericValue();
        return *this;
    }

    //! Remove an element of array by iterator.
    /*!
        \param pos iterator to the element to remove
        \pre IsArray() == true && \ref Begin() <= \c pos < \ref End()
        \return Iterator following the removed element. If the iterator pos refers to the last element, the End() iterator is returned.
        \note Linear time complexity.
    */
    ValueIterator Erase(ConstValueIterator pos) {
        return Erase(pos, pos + 1);
    }

    //! Remove elements in the range [first, last) of the array.
    /*!
        \param first iterator to the first element to remove
        \param last  iterator following the last element to remove
        \pre IsArray() == true && \ref Begin() <= \c first <= \c last <= \ref End()
        \return Iterator following the last removed element.
        \note Linear time complexity.
    */
    ValueIterator Erase(ConstValueIterator first, ConstValueIterator last) {
        CEREAL_RAPIDJSON_ASSERT(IsArray());
        CEREAL_RAPIDJSON_ASSERT(data_.a.size > 0);
        CEREAL_RAPIDJSON_ASSERT(GetElementsPointer() != 0);
        CEREAL_RAPIDJSON_ASSERT(first >= Begin());
        CEREAL_RAPIDJSON_ASSERT(first <= last);
        CEREAL_RAPIDJSON_ASSERT(last <= End());
        ValueIterator pos = Begin() + (first - Begin());
        for (ValueIterator itr = pos; itr != last; ++itr)
            itr->~GenericValue();
        std::memmove(static_cast<void*>(pos), last, static_cast<size_t>(End() - last) * sizeof(GenericValue));
        data_.a.size -= static_cast<SizeType>(last - first);
        return pos;
    }

    Array GetArray() { CEREAL_RAPIDJSON_ASSERT(IsArray()); return Array(*this); }
    ConstArray GetArray() const { CEREAL_RAPIDJSON_ASSERT(IsArray()); return ConstArray(*this); }

    //@}

    //!@name Number
    //@{

    int GetInt() const          { CEREAL_RAPIDJSON_ASSERT(data_.f.flags & kIntFlag);   return data_.n.i.i;   }
    unsigned GetUint() const    { CEREAL_RAPIDJSON_ASSERT(data_.f.flags & kUintFlag);  return data_.n.u.u;   }
    int64_t GetInt64() const    { CEREAL_RAPIDJSON_ASSERT(data_.f.flags & kInt64Flag); return data_.n.i64; }
    uint64_t GetUint64() const  { CEREAL_RAPIDJSON_ASSERT(data_.f.flags & kUint64Flag); return data_.n.u64; }

    //! Get the value as double type.
    /*! \note If the value is 64-bit integer type, it may lose precision. Use \c IsLosslessDouble() to check whether the converison is lossless.
    */
    double GetDouble() const {
        CEREAL_RAPIDJSON_ASSERT(IsNumber());
        if ((data_.f.flags & kDoubleFlag) != 0)                return data_.n.d;   // exact type, no conversion.
        if ((data_.f.flags & kIntFlag) != 0)                   return data_.n.i.i; // int -> double
        if ((data_.f.flags & kUintFlag) != 0)                  return data_.n.u.u; // unsigned -> double
        if ((data_.f.flags & kInt64Flag) != 0)                 return static_cast<double>(data_.n.i64); // int64_t -> double (may lose precision)
        CEREAL_RAPIDJSON_ASSERT((data_.f.flags & kUint64Flag) != 0);  return static_cast<double>(data_.n.u64); // uint64_t -> double (may lose precision)
    }

    //! Get the value as float type.
    /*! \note If the value is 64-bit integer type, it may lose precision. Use \c IsLosslessFloat() to check whether the converison is lossless.
    */
    float GetFloat() const {
        return static_cast<float>(GetDouble());
    }

    GenericValue& SetInt(int i)             { this->~GenericValue(); new (this) GenericValue(i);    return *this; }
    GenericValue& SetUint(unsigned u)       { this->~GenericValue(); new (this) GenericValue(u);    return *this; }
    GenericValue& SetInt64(int64_t i64)     { this->~GenericValue(); new (this) GenericValue(i64);  return *this; }
    GenericValue& SetUint64(uint64_t u64)   { this->~GenericValue(); new (this) GenericValue(u64);  return *this; }
    GenericValue& SetDouble(double d)       { this->~GenericValue(); new (this) GenericValue(d);    return *this; }
    GenericValue& SetFloat(float f)         { this->~GenericValue(); new (this) GenericValue(static_cast<double>(f)); return *this; }

    //@}

    //!@name String
    //@{

    const Ch* GetString() const { CEREAL_RAPIDJSON_ASSERT(IsString()); return (data_.f.flags & kInlineStrFlag) ? data_.ss.str : GetStringPointer(); }

    //! Get the length of string.
    /*! Since rapidjson permits "\\u0000" in the json string, strlen(v.GetString()) may not equal to v.GetStringLength().
    */
    SizeType GetStringLength() const { CEREAL_RAPIDJSON_ASSERT(IsString()); return ((data_.f.flags & kInlineStrFlag) ? (data_.ss.GetLength()) : data_.s.length); }

    //! Set this value as a string without copying source string.
    /*! This version has better performance with supplied length, and also support string containing null character.
        \param s source string pointer.
        \param length The length of source string, excluding the trailing null terminator.
        \return The value itself for fluent API.
        \post IsString() == true && GetString() == s && GetStringLength() == length
        \see SetString(StringRefType)
    */
    GenericValue& SetString(const Ch* s, SizeType length) { return SetString(StringRef(s, length)); }

    //! Set this value as a string without copying source string.
    /*! \param s source string reference
        \return The value itself for fluent API.
        \post IsString() == true && GetString() == s && GetStringLength() == s.length
    */
    GenericValue& SetString(StringRefType s) { this->~GenericValue(); SetStringRaw(s); return *this; }

    //! Set this value as a string by copying from source string.
    /*! This version has better performance with supplied length, and also support string containing null character.
        \param s source string.
        \param length The length of source string, excluding the trailing null terminator.
        \param allocator Allocator for allocating copied buffer. Commonly use GenericDocument::GetAllocator().
        \return The value itself for fluent API.
        \post IsString() == true && GetString() != s && strcmp(GetString(),s) == 0 && GetStringLength() == length
    */
    GenericValue& SetString(const Ch* s, SizeType length, Allocator& allocator) { return SetString(StringRef(s, length), allocator); }

    //! Set this value as a string by copying from source string.
    /*! \param s source string.
        \param allocator Allocator for allocating copied buffer. Commonly use GenericDocument::GetAllocator().
        \return The value itself for fluent API.
        \post IsString() == true && GetString() != s && strcmp(GetString(),s) == 0 && GetStringLength() == length
    */
    GenericValue& SetString(const Ch* s, Allocator& allocator) { return SetString(StringRef(s), allocator); }

    //! Set this value as a string by copying from source string.
    /*! \param s source string reference
        \param allocator Allocator for allocating copied buffer. Commonly use GenericDocument::GetAllocator().
        \return The value itself for fluent API.
        \post IsString() == true && GetString() != s.s && strcmp(GetString(),s) == 0 && GetStringLength() == length
    */
    GenericValue& SetString(StringRefType s, Allocator& allocator) { this->~GenericValue(); SetStringRaw(s, allocator); return *this; }

#if CEREAL_RAPIDJSON_HAS_STDSTRING
    //! Set this value as a string by copying from source string.
    /*! \param s source string.
        \param allocator Allocator for allocating copied buffer. Commonly use GenericDocument::GetAllocator().
        \return The value itself for fluent API.
        \post IsString() == true && GetString() != s.data() && strcmp(GetString(),s.data() == 0 && GetStringLength() == s.size()
        \note Requires the definition of the preprocessor symbol \ref CEREAL_RAPIDJSON_HAS_STDSTRING.
    */
    GenericValue& SetString(const std::basic_string<Ch>& s, Allocator& allocator) { return SetString(StringRef(s), allocator); }
#endif

    //@}

    //!@name Array
    //@{

    //! Templated version for checking whether this value is type T.
    /*!
        \tparam T Either \c bool, \c int, \c unsigned, \c int64_t, \c uint64_t, \c double, \c float, \c const \c char*, \c std::basic_string<Ch>
    */
    template <typename T>
    bool Is() const { return internal::TypeHelper<ValueType, T>::Is(*this); }

    template <typename T>
    T Get() const { return internal::TypeHelper<ValueType, T>::Get(*this); }

    template <typename T>
    T Get() { return internal::TypeHelper<ValueType, T>::Get(*this); }

    template<typename T>
    ValueType& Set(const T& data) { return internal::TypeHelper<ValueType, T>::Set(*this, data); }

    template<typename T>
    ValueType& Set(const T& data, AllocatorType& allocator) { return internal::TypeHelper<ValueType, T>::Set(*this, data, allocator); }

    //@}

    //! Generate events of this value to a Handler.
    /*! This function adopts the GoF visitor pattern.
        Typical usage is to output this JSON value as JSON text via Writer, which is a Handler.
        It can also be used to deep clone this value via GenericDocument, which is also a Handler.
        \tparam Handler type of handler.
        \param handler An object implementing concept Handler.
    */
    template <typename Handler>
    bool Accept(Handler& handler) const {
        switch(GetType()) {
        case kNullType:     return handler.Null();
        case kFalseType:    return handler.Bool(false);
        case kTrueType:     return handler.Bool(true);

        case kObjectType:
            if (CEREAL_RAPIDJSON_UNLIKELY(!handler.StartObject()))
                return false;
            for (ConstMemberIterator m = MemberBegin(); m != MemberEnd(); ++m) {
                CEREAL_RAPIDJSON_ASSERT(m->name.IsString()); // User may change the type of name by MemberIterator.
                if (CEREAL_RAPIDJSON_UNLIKELY(!handler.Key(m->name.GetString(), m->name.GetStringLength(), (m->name.data_.f.flags & kCopyFlag) != 0)))
                    return false;
                if (CEREAL_RAPIDJSON_UNLIKELY(!m->value.Accept(handler)))
                    return false;
            }
            return handler.EndObject(data_.o.size);

        case kArrayType:
            if (CEREAL_RAPIDJSON_UNLIKELY(!handler.StartArray()))
                return false;
            for (const GenericValue* v = Begin(); v != End(); ++v)
                if (CEREAL_RAPIDJSON_UNLIKELY(!v->Accept(handler)))
                    return false;
            return handler.EndArray(data_.a.size);

        case kStringType:
            return handler.String(GetString(), GetStringLength(), (data_.f.flags & kCopyFlag) != 0);

        default:
            CEREAL_RAPIDJSON_ASSERT(GetType() == kNumberType);
            if (IsDouble())         return handler.Double(data_.n.d);
            else if (IsInt())       return handler.Int(data_.n.i.i);
            else if (IsUint())      return handler.Uint(data_.n.u.u);
            else if (IsInt64())     return handler.Int64(data_.n.i64);
            else                    return handler.Uint64(data_.n.u64);
        }
    }

private:
    template <typename, typename> friend class GenericValue;
    template <typename, typename, typename> friend class GenericDocument;

    enum {
        kBoolFlag       = 0x0008,
        kNumberFlag     = 0x0010,
        kIntFlag        = 0x0020,
        kUintFlag       = 0x0040,
        kInt64Flag      = 0x0080,
        kUint64Flag     = 0x0100,
        kDoubleFlag     = 0x0200,
        kStringFlag     = 0x0400,
        kCopyFlag       = 0x0800,
        kInlineStrFlag  = 0x1000,

        // Initial flags of different types.
        kNullFlag = kNullType,
        kTrueFlag = kTrueType | kBoolFlag,
        kFalseFlag = kFalseType | kBoolFlag,
        kNumberIntFlag = kNumberType | kNumberFlag | kIntFlag | kInt64Flag,
        kNumberUintFlag = kNumberType | kNumberFlag | kUintFlag | kUint64Flag | kInt64Flag,
        kNumberInt64Flag = kNumberType | kNumberFlag | kInt64Flag,
        kNumberUint64Flag = kNumberType | kNumberFlag | kUint64Flag,
        kNumberDoubleFlag = kNumberType | kNumberFlag | kDoubleFlag,
        kNumberAnyFlag = kNumberType | kNumberFlag | kIntFlag | kInt64Flag | kUintFlag | kUint64Flag | kDoubleFlag,
        kConstStringFlag = kStringType | kStringFlag,
        kCopyStringFlag = kStringType | kStringFlag | kCopyFlag,
        kShortStringFlag = kStringType | kStringFlag | kCopyFlag | kInlineStrFlag,
        kObjectFlag = kObjectType,
        kArrayFlag = kArrayType,

        kTypeMask = 0x07
    };

    static const SizeType kDefaultArrayCapacity = 16;
    static const SizeType kDefaultObjectCapacity = 16;

    struct Flag {
#if CEREAL_RAPIDJSON_48BITPOINTER_OPTIMIZATION
        char payload[sizeof(SizeType) * 2 + 6];     // 2 x SizeType + lower 48-bit pointer
#elif CEREAL_RAPIDJSON_64BIT
        char payload[sizeof(SizeType) * 2 + sizeof(void*) + 6]; // 6 padding bytes
#else
        char payload[sizeof(SizeType) * 2 + sizeof(void*) + 2]; // 2 padding bytes
#endif
        uint16_t flags;
    };

    struct String {
        SizeType length;
        SizeType hashcode;  //!< reserved
        const Ch* str;
    };  // 12 bytes in 32-bit mode, 16 bytes in 64-bit mode

    // implementation detail: ShortString can represent zero-terminated strings up to MaxSize chars
    // (excluding the terminating zero) and store a value to determine the length of the contained
    // string in the last character str[LenPos] by storing "MaxSize - length" there. If the string
    // to store has the maximal length of MaxSize then str[LenPos] will be 0 and therefore act as
    // the string terminator as well. For getting the string length back from that value just use
    // "MaxSize - str[LenPos]".
    // This allows to store 13-chars strings in 32-bit mode, 21-chars strings in 64-bit mode,
    // 13-chars strings for CEREAL_RAPIDJSON_48BITPOINTER_OPTIMIZATION=1 inline (for `UTF8`-encoded strings).
    struct ShortString {
        enum { MaxChars = sizeof(static_cast<Flag*>(0)->payload) / sizeof(Ch), MaxSize = MaxChars - 1, LenPos = MaxSize };
        Ch str[MaxChars];

        inline static bool Usable(SizeType len) { return                       (MaxSize >= len); }
        inline void     SetLength(SizeType len) { str[LenPos] = static_cast<Ch>(MaxSize -  len); }
        inline SizeType GetLength() const       { return  static_cast<SizeType>(MaxSize -  str[LenPos]); }
    };  // at most as many bytes as "String" above => 12 bytes in 32-bit mode, 16 bytes in 64-bit mode

    // By using proper binary layout, retrieval of different integer types do not need conversions.
    union Number {
#if CEREAL_RAPIDJSON_ENDIAN == CEREAL_RAPIDJSON_LITTLEENDIAN
        struct I {
            int i;
            char padding[4];
        }i;
        struct U {
            unsigned u;
            char padding2[4];
        }u;
#else
        struct I {
            char padding[4];
            int i;
        }i;
        struct U {
            char padding2[4];
            unsigned u;
        }u;
#endif
        int64_t i64;
        uint64_t u64;
        double d;
    };  // 8 bytes

    struct ObjectData {
        SizeType size;
        SizeType capacity;
        Member* members;
    };  // 12 bytes in 32-bit mode, 16 bytes in 64-bit mode

    struct ArrayData {
        SizeType size;
        SizeType capacity;
        GenericValue* elements;
    };  // 12 bytes in 32-bit mode, 16 bytes in 64-bit mode

    union Data {
        String s;
        ShortString ss;
        Number n;
        ObjectData o;
        ArrayData a;
        Flag f;
    };  // 16 bytes in 32-bit mode, 24 bytes in 64-bit mode, 16 bytes in 64-bit with CEREAL_RAPIDJSON_48BITPOINTER_OPTIMIZATION

    CEREAL_RAPIDJSON_FORCEINLINE const Ch* GetStringPointer() const { return CEREAL_RAPIDJSON_GETPOINTER(Ch, data_.s.str); }
    CEREAL_RAPIDJSON_FORCEINLINE const Ch* SetStringPointer(const Ch* str) { return CEREAL_RAPIDJSON_SETPOINTER(Ch, data_.s.str, str); }
    CEREAL_RAPIDJSON_FORCEINLINE GenericValue* GetElementsPointer() const { return CEREAL_RAPIDJSON_GETPOINTER(GenericValue, data_.a.elements); }
    CEREAL_RAPIDJSON_FORCEINLINE GenericValue* SetElementsPointer(GenericValue* elements) { return CEREAL_RAPIDJSON_SETPOINTER(GenericValue, data_.a.elements, elements); }
    CEREAL_RAPIDJSON_FORCEINLINE Member* GetMembersPointer() const { return CEREAL_RAPIDJSON_GETPOINTER(Member, data_.o.members); }
    CEREAL_RAPIDJSON_FORCEINLINE Member* SetMembersPointer(Member* members) { return CEREAL_RAPIDJSON_SETPOINTER(Member, data_.o.members, members); }

    // Initialize this value as array with initial data, without calling destructor.
    void SetArrayRaw(GenericValue* values, SizeType count, Allocator& allocator) {
        data_.f.flags = kArrayFlag;
        if (count) {
            GenericValue* e = static_cast<GenericValue*>(allocator.Malloc(count * sizeof(GenericValue)));
            SetElementsPointer(e);
            std::memcpy(static_cast<void*>(e), values, count * sizeof(GenericValue));
        }
        else
            SetElementsPointer(0);
        data_.a.size = data_.a.capacity = count;
    }

    //! Initialize this value as object with initial data, without calling destructor.
    void SetObjectRaw(Member* members, SizeType count, Allocator& allocator) {
        data_.f.flags = kObjectFlag;
        if (count) {
            Member* m = static_cast<Member*>(allocator.Malloc(count * sizeof(Member)));
            SetMembersPointer(m);
            std::memcpy(static_cast<void*>(m), members, count * sizeof(Member));
        }
        else
            SetMembersPointer(0);
        data_.o.size = data_.o.capacity = count;
    }

    //! Initialize this value as constant string, without calling destructor.
    void SetStringRaw(StringRefType s) CEREAL_RAPIDJSON_NOEXCEPT {
        data_.f.flags = kConstStringFlag;
        SetStringPointer(s);
        data_.s.length = s.length;
    }

    //! Initialize this value as copy string with initial data, without calling destructor.
    void SetStringRaw(StringRefType s, Allocator& allocator) {
        Ch* str = 0;
        if (ShortString::Usable(s.length)) {
            data_.f.flags = kShortStringFlag;
            data_.ss.SetLength(s.length);
            str = data_.ss.str;
        } else {
            data_.f.flags = kCopyStringFlag;
            data_.s.length = s.length;
            str = static_cast<Ch *>(allocator.Malloc((s.length + 1) * sizeof(Ch)));
            SetStringPointer(str);
        }
        std::memcpy(str, s, s.length * sizeof(Ch));
        str[s.length] = '\0';
    }

    //! Assignment without calling destructor
    void RawAssign(GenericValue& rhs) CEREAL_RAPIDJSON_NOEXCEPT {
        data_ = rhs.data_;
        // data_.f.flags = rhs.data_.f.flags;
        rhs.data_.f.flags = kNullFlag;
    }

    template <typename SourceAllocator>
    bool StringEqual(const GenericValue<Encoding, SourceAllocator>& rhs) const {
        CEREAL_RAPIDJSON_ASSERT(IsString());
        CEREAL_RAPIDJSON_ASSERT(rhs.IsString());

        const SizeType len1 = GetStringLength();
        const SizeType len2 = rhs.GetStringLength();
        if(len1 != len2) { return false; }

        const Ch* const str1 = GetString();
        const Ch* const str2 = rhs.GetString();
        if(str1 == str2) { return true; } // fast path for constant string

        return (std::memcmp(str1, str2, sizeof(Ch) * len1) == 0);
    }

    Data data_;
};

//! GenericValue with UTF8 encoding
typedef GenericValue<UTF8<> > Value;

///////////////////////////////////////////////////////////////////////////////
// GenericDocument

//! A document for parsing JSON text as DOM.
/*!
    \note implements Handler concept
    \tparam Encoding Encoding for both parsing and string storage.
    \tparam Allocator Allocator for allocating memory for the DOM
    \tparam StackAllocator Allocator for allocating memory for stack during parsing.
    \warning Although GenericDocument inherits from GenericValue, the API does \b not provide any virtual functions, especially no virtual destructor.  To avoid memory leaks, do not \c delete a GenericDocument object via a pointer to a GenericValue.
*/
template <typename Encoding, typename Allocator = MemoryPoolAllocator<>, typename StackAllocator = CrtAllocator>
class GenericDocument : public GenericValue<Encoding, Allocator> {
public:
    typedef typename Encoding::Ch Ch;                       //!< Character type derived from Encoding.
    typedef GenericValue<Encoding, Allocator> ValueType;    //!< Value type of the document.
    typedef Allocator AllocatorType;                        //!< Allocator type from template parameter.

    //! Constructor
    /*! Creates an empty document of specified type.
        \param type             Mandatory type of object to create.
        \param allocator        Optional allocator for allocating memory.
        \param stackCapacity    Optional initial capacity of stack in bytes.
        \param stackAllocator   Optional allocator for allocating memory for stack.
    */
    explicit GenericDocument(Type type, Allocator* allocator = 0, size_t stackCapacity = kDefaultStackCapacity, StackAllocator* stackAllocator = 0) :
        GenericValue<Encoding, Allocator>(type),  allocator_(allocator), ownAllocator_(0), stack_(stackAllocator, stackCapacity), parseResult_()
    {
        if (!allocator_)
            ownAllocator_ = allocator_ = CEREAL_RAPIDJSON_NEW(Allocator)();
    }

    //! Constructor
    /*! Creates an empty document which type is Null.
        \param allocator        Optional allocator for allocating memory.
        \param stackCapacity    Optional initial capacity of stack in bytes.
        \param stackAllocator   Optional allocator for allocating memory for stack.
    */
    GenericDocument(Allocator* allocator = 0, size_t stackCapacity = kDefaultStackCapacity, StackAllocator* stackAllocator = 0) :
        allocator_(allocator), ownAllocator_(0), stack_(stackAllocator, stackCapacity), parseResult_()
    {
        if (!allocator_)
            ownAllocator_ = allocator_ = CEREAL_RAPIDJSON_NEW(Allocator)();
    }

#if CEREAL_RAPIDJSON_HAS_CXX11_RVALUE_REFS
    //! Move constructor in C++11
    GenericDocument(GenericDocument&& rhs) CEREAL_RAPIDJSON_NOEXCEPT
        : ValueType(std::forward<ValueType>(rhs)), // explicit cast to avoid prohibited move from Document
          allocator_(rhs.allocator_),
          ownAllocator_(rhs.ownAllocator_),
          stack_(std::move(rhs.stack_)),
          parseResult_(rhs.parseResult_)
    {
        rhs.allocator_ = 0;
        rhs.ownAllocator_ = 0;
        rhs.parseResult_ = ParseResult();
    }
#endif

    ~GenericDocument() {
        Destroy();
    }

#if CEREAL_RAPIDJSON_HAS_CXX11_RVALUE_REFS
    //! Move assignment in C++11
    GenericDocument& operator=(GenericDocument&& rhs) CEREAL_RAPIDJSON_NOEXCEPT
    {
        // The cast to ValueType is necessary here, because otherwise it would
        // attempt to call GenericValue's templated assignment operator.
        ValueType::operator=(std::forward<ValueType>(rhs));

        // Calling the destructor here would prematurely call stack_'s destructor
        Destroy();

        allocator_ = rhs.allocator_;
        ownAllocator_ = rhs.ownAllocator_;
        stack_ = std::move(rhs.stack_);
        parseResult_ = rhs.parseResult_;

        rhs.allocator_ = 0;
        rhs.ownAllocator_ = 0;
        rhs.parseResult_ = ParseResult();

        return *this;
    }
#endif

    //! Exchange the contents of this document with those of another.
    /*!
        \param rhs Another document.
        \note Constant complexity.
        \see GenericValue::Swap
    */
    GenericDocument& Swap(GenericDocument& rhs) CEREAL_RAPIDJSON_NOEXCEPT {
        ValueType::Swap(rhs);
        stack_.Swap(rhs.stack_);
        internal::Swap(allocator_, rhs.allocator_);
        internal::Swap(ownAllocator_, rhs.ownAllocator_);
        internal::Swap(parseResult_, rhs.parseResult_);
        return *this;
    }

    // Allow Swap with ValueType.
    // Refer to Effective C++ 3rd Edition/Item 33: Avoid hiding inherited names.
    using ValueType::Swap;

    //! free-standing swap function helper
    /*!
        Helper function to enable support for common swap implementation pattern based on \c std::swap:
        \code
        void swap(MyClass& a, MyClass& b) {
            using std::swap;
            swap(a.doc, b.doc);
            // ...
        }
        \endcode
        \see Swap()
     */
    friend inline void swap(GenericDocument& a, GenericDocument& b) CEREAL_RAPIDJSON_NOEXCEPT { a.Swap(b); }

    //! Populate this document by a generator which produces SAX events.
    /*! \tparam Generator A functor with <tt>bool f(Handler)</tt> prototype.
        \param g Generator functor which sends SAX events to the parameter.
        \return The document itself for fluent API.
    */
    template <typename Generator>
    GenericDocument& Populate(Generator& g) {
        ClearStackOnExit scope(*this);
        if (g(*this)) {
            CEREAL_RAPIDJSON_ASSERT(stack_.GetSize() == sizeof(ValueType)); // Got one and only one root object
            ValueType::operator=(*stack_.template Pop<ValueType>(1));// Move value from stack to document
        }
        return *this;
    }

    //!@name Parse from stream
    //!@{

    //! Parse JSON text from an input stream (with Encoding conversion)
    /*! \tparam parseFlags Combination of \ref ParseFlag.
        \tparam SourceEncoding Encoding of input stream
        \tparam InputStream Type of input stream, implementing Stream concept
        \param is Input stream to be parsed.
        \return The document itself for fluent API.
    */
    template <unsigned parseFlags, typename SourceEncoding, typename InputStream>
    GenericDocument& ParseStream(InputStream& is) {
        GenericReader<SourceEncoding, Encoding, StackAllocator> reader(
            stack_.HasAllocator() ? &stack_.GetAllocator() : 0);
        ClearStackOnExit scope(*this);
        parseResult_ = reader.template Parse<parseFlags>(is, *this);
        if (parseResult_) {
            CEREAL_RAPIDJSON_ASSERT(stack_.GetSize() == sizeof(ValueType)); // Got one and only one root object
            ValueType::operator=(*stack_.template Pop<ValueType>(1));// Move value from stack to document
        }
        return *this;
    }

    //! Parse JSON text from an input stream
    /*! \tparam parseFlags Combination of \ref ParseFlag.
        \tparam InputStream Type of input stream, implementing Stream concept
        \param is Input stream to be parsed.
        \return The document itself for fluent API.
    */
    template <unsigned parseFlags, typename InputStream>
    GenericDocument& ParseStream(InputStream& is) {
        return ParseStream<parseFlags, Encoding, InputStream>(is);
    }

    //! Parse JSON text from an input stream (with \ref kParseDefaultFlags)
    /*! \tparam InputStream Type of input stream, implementing Stream concept
        \param is Input stream to be parsed.
        \return The document itself for fluent API.
    */
    template <typename InputStream>
    GenericDocument& ParseStream(InputStream& is) {
        return ParseStream<kParseDefaultFlags, Encoding, InputStream>(is);
    }
    //!@}

    //!@name Parse in-place from mutable string
    //!@{

    //! Parse JSON text from a mutable string
    /*! \tparam parseFlags Combination of \ref ParseFlag.
        \param str Mutable zero-terminated string to be parsed.
        \return The document itself for fluent API.
    */
    template <unsigned parseFlags>
    GenericDocument& ParseInsitu(Ch* str) {
        GenericInsituStringStream<Encoding> s(str);
        return ParseStream<parseFlags | kParseInsituFlag>(s);
    }

    //! Parse JSON text from a mutable string (with \ref kParseDefaultFlags)
    /*! \param str Mutable zero-terminated string to be parsed.
        \return The document itself for fluent API.
    */
    GenericDocument& ParseInsitu(Ch* str) {
        return ParseInsitu<kParseDefaultFlags>(str);
    }
    //!@}

    //!@name Parse from read-only string
    //!@{

    //! Parse JSON text from a read-only string (with Encoding conversion)
    /*! \tparam parseFlags Combination of \ref ParseFlag (must not contain \ref kParseInsituFlag).
        \tparam SourceEncoding Transcoding from input Encoding
        \param str Read-only zero-terminated string to be parsed.
    */
    template <unsigned parseFlags, typename SourceEncoding>
    GenericDocument& Parse(const typename SourceEncoding::Ch* str) {
        CEREAL_RAPIDJSON_ASSERT(!(parseFlags & kParseInsituFlag));
        GenericStringStream<SourceEncoding> s(str);
        return ParseStream<parseFlags, SourceEncoding>(s);
    }

    //! Parse JSON text from a read-only string
    /*! \tparam parseFlags Combination of \ref ParseFlag (must not contain \ref kParseInsituFlag).
        \param str Read-only zero-terminated string to be parsed.
    */
    template <unsigned parseFlags>
    GenericDocument& Parse(const Ch* str) {
        return Parse<parseFlags, Encoding>(str);
    }

    //! Parse JSON text from a read-only string (with \ref kParseDefaultFlags)
    /*! \param str Read-only zero-terminated string to be parsed.
    */
    GenericDocument& Parse(const Ch* str) {
        return Parse<kParseDefaultFlags>(str);
    }

    template <unsigned parseFlags, typename SourceEncoding>
    GenericDocument& Parse(const typename SourceEncoding::Ch* str, size_t length) {
        CEREAL_RAPIDJSON_ASSERT(!(parseFlags & kParseInsituFlag));
        MemoryStream ms(reinterpret_cast<const char*>(str), length * sizeof(typename SourceEncoding::Ch));
        EncodedInputStream<SourceEncoding, MemoryStream> is(ms);
        ParseStream<parseFlags, SourceEncoding>(is);
        return *this;
    }

    template <unsigned parseFlags>
    GenericDocument& Parse(const Ch* str, size_t length) {
        return Parse<parseFlags, Encoding>(str, length);
    }

    GenericDocument& Parse(const Ch* str, size_t length) {
        return Parse<kParseDefaultFlags>(str, length);
    }

#if CEREAL_RAPIDJSON_HAS_STDSTRING
    template <unsigned parseFlags, typename SourceEncoding>
    GenericDocument& Parse(const std::basic_string<typename SourceEncoding::Ch>& str) {
        // c_str() is constant complexity according to standard. Should be faster than Parse(const char*, size_t)
        return Parse<parseFlags, SourceEncoding>(str.c_str());
    }

    template <unsigned parseFlags>
    GenericDocument& Parse(const std::basic_string<Ch>& str) {
        return Parse<parseFlags, Encoding>(str.c_str());
    }

    GenericDocument& Parse(const std::basic_string<Ch>& str) {
        return Parse<kParseDefaultFlags>(str);
    }
#endif // CEREAL_RAPIDJSON_HAS_STDSTRING

    //!@}

    //!@name Handling parse errors
    //!@{

    //! Whether a parse error has occurred in the last parsing.
    bool HasParseError() const { return parseResult_.IsError(); }

    //! Get the \ref ParseErrorCode of last parsing.
    ParseErrorCode GetParseError() const { return parseResult_.Code(); }

    //! Get the position of last parsing error in input, 0 otherwise.
    size_t GetErrorOffset() const { return parseResult_.Offset(); }

    //! Implicit conversion to get the last parse result
#ifndef __clang // -Wdocumentation
    /*! \return \ref ParseResult of the last parse operation

        \code
          Document doc;
          ParseResult ok = doc.Parse(json);
          if (!ok)
            printf( "JSON parse error: %s (%u)\n", GetParseError_En(ok.Code()), ok.Offset());
        \endcode
     */
#endif
    operator ParseResult() const { return parseResult_; }
    //!@}

    //! Get the allocator of this document.
    Allocator& GetAllocator() {
        CEREAL_RAPIDJSON_ASSERT(allocator_);
        return *allocator_;
    }

    //! Get the capacity of stack in bytes.
    size_t GetStackCapacity() const { return stack_.GetCapacity(); }

private:
    // clear stack on any exit from ParseStream, e.g. due to exception
    struct ClearStackOnExit {
        explicit ClearStackOnExit(GenericDocument& d) : d_(d) {}
        ~ClearStackOnExit() { d_.ClearStack(); }
    private:
        ClearStackOnExit(const ClearStackOnExit&);
        ClearStackOnExit& operator=(const ClearStackOnExit&);
        GenericDocument& d_;
    };

    // callers of the following private Handler functions
    // template <typename,typename,typename> friend class GenericReader; // for parsing
    template <typename, typename> friend class GenericValue; // for deep copying

public:
    // Implementation of Handler
    bool Null() { new (stack_.template Push<ValueType>()) ValueType(); return true; }
    bool Bool(bool b) { new (stack_.template Push<ValueType>()) ValueType(b); return true; }
    bool Int(int i) { new (stack_.template Push<ValueType>()) ValueType(i); return true; }
    bool Uint(unsigned i) { new (stack_.template Push<ValueType>()) ValueType(i); return true; }
    bool Int64(int64_t i) { new (stack_.template Push<ValueType>()) ValueType(i); return true; }
    bool Uint64(uint64_t i) { new (stack_.template Push<ValueType>()) ValueType(i); return true; }
    bool Double(double d) { new (stack_.template Push<ValueType>()) ValueType(d); return true; }

    bool RawNumber(const Ch* str, SizeType length, bool copy) {
        if (copy)
            new (stack_.template Push<ValueType>()) ValueType(str, length, GetAllocator());
        else
            new (stack_.template Push<ValueType>()) ValueType(str, length);
        return true;
    }

    bool String(const Ch* str, SizeType length, bool copy) {
        if (copy)
            new (stack_.template Push<ValueType>()) ValueType(str, length, GetAllocator());
        else
            new (stack_.template Push<ValueType>()) ValueType(str, length);
        return true;
    }

    bool StartObject() { new (stack_.template Push<ValueType>()) ValueType(kObjectType); return true; }

    bool Key(const Ch* str, SizeType length, bool copy) { return String(str, length, copy); }

    bool EndObject(SizeType memberCount) {
        typename ValueType::Member* members = stack_.template Pop<typename ValueType::Member>(memberCount);
        stack_.template Top<ValueType>()->SetObjectRaw(members, memberCount, GetAllocator());
        return true;
    }

    bool StartArray() { new (stack_.template Push<ValueType>()) ValueType(kArrayType); return true; }

    bool EndArray(SizeType elementCount) {
        ValueType* elements = stack_.template Pop<ValueType>(elementCount);
        stack_.template Top<ValueType>()->SetArrayRaw(elements, elementCount, GetAllocator());
        return true;
    }

private:
    //! Prohibit copying
    GenericDocument(const GenericDocument&);
    //! Prohibit assignment
    GenericDocument& operator=(const GenericDocument&);

    void ClearStack() {
        if (Allocator::kNeedFree)
            while (stack_.GetSize() > 0)    // Here assumes all elements in stack array are GenericValue (Member is actually 2 GenericValue objects)
                (stack_.template Pop<ValueType>(1))->~ValueType();
        else
            stack_.Clear();
        stack_.ShrinkToFit();
    }

    void Destroy() {
        CEREAL_RAPIDJSON_DELETE(ownAllocator_);
    }

    static const size_t kDefaultStackCapacity = 1024;
    Allocator* allocator_;
    Allocator* ownAllocator_;
    internal::Stack<StackAllocator> stack_;
    ParseResult parseResult_;
};

//! GenericDocument with UTF8 encoding
typedef GenericDocument<UTF8<> > Document;

//! Helper class for accessing Value of array type.
/*!
    Instance of this helper class is obtained by \c GenericValue::GetArray().
    In addition to all APIs for array type, it provides range-based for loop if \c CEREAL_RAPIDJSON_HAS_CXX11_RANGE_FOR=1.
*/
template <bool Const, typename ValueT>
class GenericArray {
public:
    typedef GenericArray<true, ValueT> ConstArray;
    typedef GenericArray<false, ValueT> Array;
    typedef ValueT PlainType;
    typedef typename internal::MaybeAddConst<Const,PlainType>::Type ValueType;
    typedef ValueType* ValueIterator;  // This may be const or non-const iterator
    typedef const ValueT* ConstValueIterator;
    typedef typename ValueType::AllocatorType AllocatorType;
    typedef typename ValueType::StringRefType StringRefType;

    template <typename, typename>
    friend class GenericValue;

    GenericArray(const GenericArray& rhs) : value_(rhs.value_) {}
    GenericArray& operator=(const GenericArray& rhs) { value_ = rhs.value_; return *this; }
    ~GenericArray() {}

    SizeType Size() const { return value_.Size(); }
    SizeType Capacity() const { return value_.Capacity(); }
    bool Empty() const { return value_.Empty(); }
    void Clear() const { value_.Clear(); }
    ValueType& operator[](SizeType index) const {  return value_[index]; }
    ValueIterator Begin() const { return value_.Begin(); }
    ValueIterator End() const { return value_.End(); }
    GenericArray Reserve(SizeType newCapacity, AllocatorType &allocator) const { value_.Reserve(newCapacity, allocator); return *this; }
    GenericArray PushBack(ValueType& value, AllocatorType& allocator) const { value_.PushBack(value, allocator); return *this; }
#if CEREAL_RAPIDJSON_HAS_CXX11_RVALUE_REFS
    GenericArray PushBack(ValueType&& value, AllocatorType& allocator) const { value_.PushBack(value, allocator); return *this; }
#endif // CEREAL_RAPIDJSON_HAS_CXX11_RVALUE_REFS
    GenericArray PushBack(StringRefType value, AllocatorType& allocator) const { value_.PushBack(value, allocator); return *this; }
    template <typename T> CEREAL_RAPIDJSON_DISABLEIF_RETURN((internal::OrExpr<internal::IsPointer<T>, internal::IsGenericValue<T> >), (const GenericArray&)) PushBack(T value, AllocatorType& allocator) const { value_.PushBack(value, allocator); return *this; }
    GenericArray PopBack() const { value_.PopBack(); return *this; }
    ValueIterator Erase(ConstValueIterator pos) const { return value_.Erase(pos); }
    ValueIterator Erase(ConstValueIterator first, ConstValueIterator last) const { return value_.Erase(first, last); }

#if CEREAL_RAPIDJSON_HAS_CXX11_RANGE_FOR
    ValueIterator begin() const { return value_.Begin(); }
    ValueIterator end() const { return value_.End(); }
#endif

private:
    GenericArray();
    GenericArray(ValueType& value) : value_(value) {}
    ValueType& value_;
};

//! Helper class for accessing Value of object type.
/*!
    Instance of this helper class is obtained by \c GenericValue::GetObject().
    In addition to all APIs for array type, it provides range-based for loop if \c CEREAL_RAPIDJSON_HAS_CXX11_RANGE_FOR=1.
*/
template <bool Const, typename ValueT>
class GenericObject {
public:
    typedef GenericObject<true, ValueT> ConstObject;
    typedef GenericObject<false, ValueT> Object;
    typedef ValueT PlainType;
    typedef typename internal::MaybeAddConst<Const,PlainType>::Type ValueType;
    typedef GenericMemberIterator<Const, typename ValueT::EncodingType, typename ValueT::AllocatorType> MemberIterator;  // This may be const or non-const iterator
    typedef GenericMemberIterator<true, typename ValueT::EncodingType, typename ValueT::AllocatorType> ConstMemberIterator;
    typedef typename ValueType::AllocatorType AllocatorType;
    typedef typename ValueType::StringRefType StringRefType;
    typedef typename ValueType::EncodingType EncodingType;
    typedef typename ValueType::Ch Ch;

    template <typename, typename>
    friend class GenericValue;

    GenericObject(const GenericObject& rhs) : value_(rhs.value_) {}
    GenericObject& operator=(const GenericObject& rhs) { value_ = rhs.value_; return *this; }
    ~GenericObject() {}

    SizeType MemberCount() const { return value_.MemberCount(); }
    SizeType MemberCapacity() const { return value_.MemberCapacity(); }
    bool ObjectEmpty() const { return value_.ObjectEmpty(); }
    template <typename T> ValueType& operator[](T* name) const { return value_[name]; }
    template <typename SourceAllocator> ValueType& operator[](const GenericValue<EncodingType, SourceAllocator>& name) const { return value_[name]; }
#if CEREAL_RAPIDJSON_HAS_STDSTRING
    ValueType& operator[](const std::basic_string<Ch>& name) const { return value_[name]; }
#endif
    MemberIterator MemberBegin() const { return value_.MemberBegin(); }
    MemberIterator MemberEnd() const { return value_.MemberEnd(); }
    GenericObject MemberReserve(SizeType newCapacity, AllocatorType &allocator) const { value_.MemberReserve(newCapacity, allocator); return *this; }
    bool HasMember(const Ch* name) const { return value_.HasMember(name); }
#if CEREAL_RAPIDJSON_HAS_STDSTRING
    bool HasMember(const std::basic_string<Ch>& name) const { return value_.HasMember(name); }
#endif
    template <typename SourceAllocator> bool HasMember(const GenericValue<EncodingType, SourceAllocator>& name) const { return value_.HasMember(name); }
    MemberIterator FindMember(const Ch* name) const { return value_.FindMember(name); }
    template <typename SourceAllocator> MemberIterator FindMember(const GenericValue<EncodingType, SourceAllocator>& name) const { return value_.FindMember(name); }
#if CEREAL_RAPIDJSON_HAS_STDSTRING
    MemberIterator FindMember(const std::basic_string<Ch>& name) const { return value_.FindMember(name); }
#endif
    GenericObject AddMember(ValueType& name, ValueType& value, AllocatorType& allocator) const { value_.AddMember(name, value, allocator); return *this; }
    GenericObject AddMember(ValueType& name, StringRefType value, AllocatorType& allocator) const { value_.AddMember(name, value, allocator); return *this; }
#if CEREAL_RAPIDJSON_HAS_STDSTRING
    GenericObject AddMember(ValueType& name, std::basic_string<Ch>& value, AllocatorType& allocator) const { value_.AddMember(name, value, allocator); return *this; }
#endif
    template <typename T> CEREAL_RAPIDJSON_DISABLEIF_RETURN((internal::OrExpr<internal::IsPointer<T>, internal::IsGenericValue<T> >), (ValueType&)) AddMember(ValueType& name, T value, AllocatorType& allocator) const { value_.AddMember(name, value, allocator); return *this; }
#if CEREAL_RAPIDJSON_HAS_CXX11_RVALUE_REFS
    GenericObject AddMember(ValueType&& name, ValueType&& value, AllocatorType& allocator) const { value_.AddMember(name, value, allocator); return *this; }
    GenericObject AddMember(ValueType&& name, ValueType& value, AllocatorType& allocator) const { value_.AddMember(name, value, allocator); return *this; }
    GenericObject AddMember(ValueType& name, ValueType&& value, AllocatorType& allocator) const { value_.AddMember(name, value, allocator); return *this; }
    GenericObject AddMember(StringRefType name, ValueType&& value, AllocatorType& allocator) const { value_.AddMember(name, value, allocator); return *this; }
#endif // CEREAL_RAPIDJSON_HAS_CXX11_RVALUE_REFS
    GenericObject AddMember(StringRefType name, ValueType& value, AllocatorType& allocator) const { value_.AddMember(name, value, allocator); return *this; }
    GenericObject AddMember(StringRefType name, StringRefType value, AllocatorType& allocator) const { value_.AddMember(name, value, allocator); return *this; }
    template <typename T> CEREAL_RAPIDJSON_DISABLEIF_RETURN((internal::OrExpr<internal::IsPointer<T>, internal::IsGenericValue<T> >), (GenericObject)) AddMember(StringRefType name, T value, AllocatorType& allocator) const { value_.AddMember(name, value, allocator); return *this; }
    void RemoveAllMembers() { value_.RemoveAllMembers(); }
    bool RemoveMember(const Ch* name) const { return value_.RemoveMember(name); }
#if CEREAL_RAPIDJSON_HAS_STDSTRING
    bool RemoveMember(const std::basic_string<Ch>& name) const { return value_.RemoveMember(name); }
#endif
    template <typename SourceAllocator> bool RemoveMember(const GenericValue<EncodingType, SourceAllocator>& name) const { return value_.RemoveMember(name); }
    MemberIterator RemoveMember(MemberIterator m) const { return value_.RemoveMember(m); }
    MemberIterator EraseMember(ConstMemberIterator pos) const { return value_.EraseMember(pos); }
    MemberIterator EraseMember(ConstMemberIterator first, ConstMemberIterator last) const { return value_.EraseMember(first, last); }
    bool EraseMember(const Ch* name) const { return value_.EraseMember(name); }
#if CEREAL_RAPIDJSON_HAS_STDSTRING
    bool EraseMember(const std::basic_string<Ch>& name) const { return EraseMember(ValueType(StringRef(name))); }
#endif
    template <typename SourceAllocator> bool EraseMember(const GenericValue<EncodingType, SourceAllocator>& name) const { return value_.EraseMember(name); }

#if CEREAL_RAPIDJSON_HAS_CXX11_RANGE_FOR
    MemberIterator begin() const { return value_.MemberBegin(); }
    MemberIterator end() const { return value_.MemberEnd(); }
#endif

private:
    GenericObject();
    GenericObject(ValueType& value) : value_(value) {}
    ValueType& value_;
};

CEREAL_RAPIDJSON_NAMESPACE_END
CEREAL_RAPIDJSON_DIAG_POP

#endif // CEREAL_RAPIDJSON_DOCUMENT_H_
```
