## Supplemental code for "The motif composition of variable-number tandem repeats impacts gene expression": encodedstream.h

```
// Tencent is pleased to support the open source community by making RapidJSON available.
// 
// Copyright (C) 2015 THL A29 Limited, a Tencent company, and Milo Yip. All rights reserved.
//
// Licensed under the MIT License (the "License"); you may not use this file except
// in compliance with the License. You may obtain a copy of the License at
//
// http://opensource.org/licenses/MIT
//
// Unless required by applicable law or agreed to in writing, software distributed 
// under the License is distributed on an "AS IS" BASIS, WITHOUT WARRANTIES OR 
// CONDITIONS OF ANY KIND, either express or implied. See the License for the 
// specific language governing permissions and limitations under the License.

#ifndef CEREAL_RAPIDJSON_ENCODEDSTREAM_H_
#define CEREAL_RAPIDJSON_ENCODEDSTREAM_H_

#include "stream.h"
#include "memorystream.h"

#ifdef __GNUC__
CEREAL_RAPIDJSON_DIAG_PUSH
CEREAL_RAPIDJSON_DIAG_OFF(effc++)
#endif

#ifdef __clang__
CEREAL_RAPIDJSON_DIAG_PUSH
CEREAL_RAPIDJSON_DIAG_OFF(padded)
#endif

CEREAL_RAPIDJSON_NAMESPACE_BEGIN

//! Input byte stream wrapper with a statically bound encoding.
/*!
    \tparam Encoding The interpretation of encoding of the stream. Either UTF8, UTF16LE, UTF16BE, UTF32LE, UTF32BE.
    \tparam InputByteStream Type of input byte stream. For example, FileReadStream.
*/
template <typename Encoding, typename InputByteStream>
class EncodedInputStream {
    CEREAL_RAPIDJSON_STATIC_ASSERT(sizeof(typename InputByteStream::Ch) == 1);
public:
    typedef typename Encoding::Ch Ch;

    EncodedInputStream(InputByteStream& is) : is_(is) { 
        current_ = Encoding::TakeBOM(is_);
    }

    Ch Peek() const { return current_; }
    Ch Take() { Ch c = current_; current_ = Encoding::Take(is_); return c; }
    size_t Tell() const { return is_.Tell(); }

    // Not implemented
    void Put(Ch) { CEREAL_RAPIDJSON_ASSERT(false); }
    void Flush() { CEREAL_RAPIDJSON_ASSERT(false); } 
    Ch* PutBegin() { CEREAL_RAPIDJSON_ASSERT(false); return 0; }
    size_t PutEnd(Ch*) { CEREAL_RAPIDJSON_ASSERT(false); return 0; }

private:
    EncodedInputStream(const EncodedInputStream&);
    EncodedInputStream& operator=(const EncodedInputStream&);

    InputByteStream& is_;
    Ch current_;
};

//! Specialized for UTF8 MemoryStream.
template <>
class EncodedInputStream<UTF8<>, MemoryStream> {
public:
    typedef UTF8<>::Ch Ch;

    EncodedInputStream(MemoryStream& is) : is_(is) {
        if (static_cast<unsigned char>(is_.Peek()) == 0xEFu) is_.Take();
        if (static_cast<unsigned char>(is_.Peek()) == 0xBBu) is_.Take();
        if (static_cast<unsigned char>(is_.Peek()) == 0xBFu) is_.Take();
    }
    Ch Peek() const { return is_.Peek(); }
    Ch Take() { return is_.Take(); }
    size_t Tell() const { return is_.Tell(); }

    // Not implemented
    void Put(Ch) {}
    void Flush() {} 
    Ch* PutBegin() { return 0; }
    size_t PutEnd(Ch*) { return 0; }

    MemoryStream& is_;

private:
    EncodedInputStream(const EncodedInputStream&);
    EncodedInputStream& operator=(const EncodedInputStream&);
};

//! Output byte stream wrapper with statically bound encoding.
/*!
    \tparam Encoding The interpretation of encoding of the stream. Either UTF8, UTF16LE, UTF16BE, UTF32LE, UTF32BE.
    \tparam OutputByteStream Type of input byte stream. For example, FileWriteStream.
*/
template <typename Encoding, typename OutputByteStream>
class EncodedOutputStream {
    CEREAL_RAPIDJSON_STATIC_ASSERT(sizeof(typename OutputByteStream::Ch) == 1);
public:
    typedef typename Encoding::Ch Ch;

    EncodedOutputStream(OutputByteStream& os, bool putBOM = true) : os_(os) { 
        if (putBOM)
            Encoding::PutBOM(os_);
    }

    void Put(Ch c) { Encoding::Put(os_, c);  }
    void Flush() { os_.Flush(); }

    // Not implemented
    Ch Peek() const { CEREAL_RAPIDJSON_ASSERT(false); return 0;}
    Ch Take() { CEREAL_RAPIDJSON_ASSERT(false); return 0;}
    size_t Tell() const { CEREAL_RAPIDJSON_ASSERT(false);  return 0; }
    Ch* PutBegin() { CEREAL_RAPIDJSON_ASSERT(false); return 0; }
    size_t PutEnd(Ch*) { CEREAL_RAPIDJSON_ASSERT(false); return 0; }

private:
    EncodedOutputStream(const EncodedOutputStream&);
    EncodedOutputStream& operator=(const EncodedOutputStream&);

    OutputByteStream& os_;
};

#define CEREAL_RAPIDJSON_ENCODINGS_FUNC(x) UTF8<Ch>::x, UTF16LE<Ch>::x, UTF16BE<Ch>::x, UTF32LE<Ch>::x, UTF32BE<Ch>::x

//! Input stream wrapper with dynamically bound encoding and automatic encoding detection.
/*!
    \tparam CharType Type of character for reading.
    \tparam InputByteStream type of input byte stream to be wrapped.
*/
template <typename CharType, typename InputByteStream>
class AutoUTFInputStream {
    CEREAL_RAPIDJSON_STATIC_ASSERT(sizeof(typename InputByteStream::Ch) == 1);
public:
    typedef CharType Ch;

    //! Constructor.
    /*!
        \param is input stream to be wrapped.
        \param type UTF encoding type if it is not detected from the stream.
    */
    AutoUTFInputStream(InputByteStream& is, UTFType type = kUTF8) : is_(&is), type_(type), hasBOM_(false) {
        CEREAL_RAPIDJSON_ASSERT(type >= kUTF8 && type <= kUTF32BE);        
        DetectType();
        static const TakeFunc f[] = { CEREAL_RAPIDJSON_ENCODINGS_FUNC(Take) };
        takeFunc_ = f[type_];
        current_ = takeFunc_(*is_);
    }

    UTFType GetType() const { return type_; }
    bool HasBOM() const { return hasBOM_; }

    Ch Peek() const { return current_; }
    Ch Take() { Ch c = current_; current_ = takeFunc_(*is_); return c; }
    size_t Tell() const { return is_->Tell(); }

    // Not implemented
    void Put(Ch) { CEREAL_RAPIDJSON_ASSERT(false); }
    void Flush() { CEREAL_RAPIDJSON_ASSERT(false); } 
    Ch* PutBegin() { CEREAL_RAPIDJSON_ASSERT(false); return 0; }
    size_t PutEnd(Ch*) { CEREAL_RAPIDJSON_ASSERT(false); return 0; }

private:
    AutoUTFInputStream(const AutoUTFInputStream&);
    AutoUTFInputStream& operator=(const AutoUTFInputStream&);

    // Detect encoding type with BOM or RFC 4627
    void DetectType() {
        // BOM (Byte Order Mark):
        // 00 00 FE FF  UTF-32BE
        // FF FE 00 00  UTF-32LE
        // FE FF        UTF-16BE
        // FF FE        UTF-16LE
        // EF BB BF     UTF-8

        const unsigned char* c = reinterpret_cast<const unsigned char *>(is_->Peek4());
        if (!c)
            return;

        unsigned bom = static_cast<unsigned>(c[0] | (c[1] << 8) | (c[2] << 16) | (c[3] << 24));
        hasBOM_ = false;
        if (bom == 0xFFFE0000)                  { type_ = kUTF32BE; hasBOM_ = true; is_->Take(); is_->Take(); is_->Take(); is_->Take(); }
        else if (bom == 0x0000FEFF)             { type_ = kUTF32LE; hasBOM_ = true; is_->Take(); is_->Take(); is_->Take(); is_->Take(); }
        else if ((bom & 0xFFFF) == 0xFFFE)      { type_ = kUTF16BE; hasBOM_ = true; is_->Take(); is_->Take();                           }
        else if ((bom & 0xFFFF) == 0xFEFF)      { type_ = kUTF16LE; hasBOM_ = true; is_->Take(); is_->Take();                           }
        else if ((bom & 0xFFFFFF) == 0xBFBBEF)  { type_ = kUTF8;    hasBOM_ = true; is_->Take(); is_->Take(); is_->Take();              }

        // RFC 4627: Section 3
        // "Since the first two characters of a JSON text will always be ASCII
        // characters [RFC0020], it is possible to determine whether an octet
        // stream is UTF-8, UTF-16 (BE or LE), or UTF-32 (BE or LE) by looking
        // at the pattern of nulls in the first four octets."
        // 00 00 00 xx  UTF-32BE
        // 00 xx 00 xx  UTF-16BE
        // xx 00 00 00  UTF-32LE
        // xx 00 xx 00  UTF-16LE
        // xx xx xx xx  UTF-8

        if (!hasBOM_) {
            int pattern = (c[0] ? 1 : 0) | (c[1] ? 2 : 0) | (c[2] ? 4 : 0) | (c[3] ? 8 : 0);
            switch (pattern) {
            case 0x08: type_ = kUTF32BE; break;
            case 0x0A: type_ = kUTF16BE; break;
            case 0x01: type_ = kUTF32LE; break;
            case 0x05: type_ = kUTF16LE; break;
            case 0x0F: type_ = kUTF8;    break;
            default: break; // Use type defined by user.
            }
        }

        // Runtime check whether the size of character type is sufficient. It only perform checks with assertion.
        if (type_ == kUTF16LE || type_ == kUTF16BE) CEREAL_RAPIDJSON_ASSERT(sizeof(Ch) >= 2);
        if (type_ == kUTF32LE || type_ == kUTF32BE) CEREAL_RAPIDJSON_ASSERT(sizeof(Ch) >= 4);
    }

    typedef Ch (*TakeFunc)(InputByteStream& is);
    InputByteStream* is_;
    UTFType type_;
    Ch current_;
    TakeFunc takeFunc_;
    bool hasBOM_;
};

//! Output stream wrapper with dynamically bound encoding and automatic encoding detection.
/*!
    \tparam CharType Type of character for writing.
    \tparam OutputByteStream type of output byte stream to be wrapped.
*/
template <typename CharType, typename OutputByteStream>
class AutoUTFOutputStream {
    CEREAL_RAPIDJSON_STATIC_ASSERT(sizeof(typename OutputByteStream::Ch) == 1);
public:
    typedef CharType Ch;

    //! Constructor.
    /*!
        \param os output stream to be wrapped.
        \param type UTF encoding type.
        \param putBOM Whether to write BOM at the beginning of the stream.
    */
    AutoUTFOutputStream(OutputByteStream& os, UTFType type, bool putBOM) : os_(&os), type_(type) {
        CEREAL_RAPIDJSON_ASSERT(type >= kUTF8 && type <= kUTF32BE);

        // Runtime check whether the size of character type is sufficient. It only perform checks with assertion.
        if (type_ == kUTF16LE || type_ == kUTF16BE) CEREAL_RAPIDJSON_ASSERT(sizeof(Ch) >= 2);
        if (type_ == kUTF32LE || type_ == kUTF32BE) CEREAL_RAPIDJSON_ASSERT(sizeof(Ch) >= 4);

        static const PutFunc f[] = { CEREAL_RAPIDJSON_ENCODINGS_FUNC(Put) };
        putFunc_ = f[type_];

        if (putBOM)
            PutBOM();
    }

    UTFType GetType() const { return type_; }

    void Put(Ch c) { putFunc_(*os_, c); }
    void Flush() { os_->Flush(); } 

    // Not implemented
    Ch Peek() const { CEREAL_RAPIDJSON_ASSERT(false); return 0;}
    Ch Take() { CEREAL_RAPIDJSON_ASSERT(false); return 0;}
    size_t Tell() const { CEREAL_RAPIDJSON_ASSERT(false); return 0; }
    Ch* PutBegin() { CEREAL_RAPIDJSON_ASSERT(false); return 0; }
    size_t PutEnd(Ch*) { CEREAL_RAPIDJSON_ASSERT(false); return 0; }

private:
    AutoUTFOutputStream(const AutoUTFOutputStream&);
    AutoUTFOutputStream& operator=(const AutoUTFOutputStream&);

    void PutBOM() { 
        typedef void (*PutBOMFunc)(OutputByteStream&);
        static const PutBOMFunc f[] = { CEREAL_RAPIDJSON_ENCODINGS_FUNC(PutBOM) };
        f[type_](*os_);
    }

    typedef void (*PutFunc)(OutputByteStream&, Ch);

    OutputByteStream* os_;
    UTFType type_;
    PutFunc putFunc_;
};

#undef CEREAL_RAPIDJSON_ENCODINGS_FUNC

CEREAL_RAPIDJSON_NAMESPACE_END

#ifdef __clang__
CEREAL_RAPIDJSON_DIAG_POP
#endif

#ifdef __GNUC__
CEREAL_RAPIDJSON_DIAG_POP
#endif

#endif // CEREAL_RAPIDJSON_FILESTREAM_H_
```
