## Supplemental code for "The motif composition of variable-number tandem repeats impacts gene expression": encodings.h

```
// Tencent is pleased to support the open source community by making RapidJSON available.
// 
// Copyright (C) 2015 THL A29 Limited, a Tencent company, and Milo Yip. All rights reserved.
//
// Licensed under the MIT License (the "License"); you may not use this file except
// in compliance with the License. You may obtain a copy of the License at
//
// http://opensource.org/licenses/MIT
//
// Unless required by applicable law or agreed to in writing, software distributed 
// under the License is distributed on an "AS IS" BASIS, WITHOUT WARRANTIES OR 
// CONDITIONS OF ANY KIND, either express or implied. See the License for the 
// specific language governing permissions and limitations under the License.

#ifndef CEREAL_RAPIDJSON_ENCODINGS_H_
#define CEREAL_RAPIDJSON_ENCODINGS_H_

#include "rapidjson.h"

#if defined(_MSC_VER) && !defined(__clang__)
CEREAL_RAPIDJSON_DIAG_PUSH
CEREAL_RAPIDJSON_DIAG_OFF(4244) // conversion from 'type1' to 'type2', possible loss of data
CEREAL_RAPIDJSON_DIAG_OFF(4702)  // unreachable code
#elif defined(__GNUC__)
CEREAL_RAPIDJSON_DIAG_PUSH
CEREAL_RAPIDJSON_DIAG_OFF(effc++)
CEREAL_RAPIDJSON_DIAG_OFF(overflow)
#endif

CEREAL_RAPIDJSON_NAMESPACE_BEGIN

///////////////////////////////////////////////////////////////////////////////
// Encoding

/*! \class rapidjson::Encoding
    \brief Concept for encoding of Unicode characters.

\code
concept Encoding {
    typename Ch;    //! Type of character. A "character" is actually a code unit in unicode's definition.

    enum { supportUnicode = 1 }; // or 0 if not supporting unicode

    //! \brief Encode a Unicode codepoint to an output stream.
    //! \param os Output stream.
    //! \param codepoint An unicode codepoint, ranging from 0x0 to 0x10FFFF inclusively.
    template<typename OutputStream>
    static void Encode(OutputStream& os, unsigned codepoint);

    //! \brief Decode a Unicode codepoint from an input stream.
    //! \param is Input stream.
    //! \param codepoint Output of the unicode codepoint.
    //! \return true if a valid codepoint can be decoded from the stream.
    template <typename InputStream>
    static bool Decode(InputStream& is, unsigned* codepoint);

    //! \brief Validate one Unicode codepoint from an encoded stream.
    //! \param is Input stream to obtain codepoint.
    //! \param os Output for copying one codepoint.
    //! \return true if it is valid.
    //! \note This function just validating and copying the codepoint without actually decode it.
    template <typename InputStream, typename OutputStream>
    static bool Validate(InputStream& is, OutputStream& os);

    // The following functions are deal with byte streams.

    //! Take a character from input byte stream, skip BOM if exist.
    template <typename InputByteStream>
    static CharType TakeBOM(InputByteStream& is);

    //! Take a character from input byte stream.
    template <typename InputByteStream>
    static Ch Take(InputByteStream& is);

    //! Put BOM to output byte stream.
    template <typename OutputByteStream>
    static void PutBOM(OutputByteStream& os);

    //! Put a character to output byte stream.
    template <typename OutputByteStream>
    static void Put(OutputByteStream& os, Ch c);
};
\endcode
*/

///////////////////////////////////////////////////////////////////////////////
// UTF8

//! UTF-8 encoding.
/*! http://en.wikipedia.org/wiki/UTF-8
    http://tools.ietf.org/html/rfc3629
    \tparam CharType Code unit for storing 8-bit UTF-8 data. Default is char.
    \note implements Encoding concept
*/
template<typename CharType = char>
struct UTF8 {
    typedef CharType Ch;

    enum { supportUnicode = 1 };

    template<typename OutputStream>
    static void Encode(OutputStream& os, unsigned codepoint) {
        if (codepoint <= 0x7F) 
            os.Put(static_cast<Ch>(codepoint & 0xFF));
        else if (codepoint <= 0x7FF) {
            os.Put(static_cast<Ch>(0xC0 | ((codepoint >> 6) & 0xFF)));
            os.Put(static_cast<Ch>(0x80 | ((codepoint & 0x3F))));
        }
        else if (codepoint <= 0xFFFF) {
            os.Put(static_cast<Ch>(0xE0 | ((codepoint >> 12) & 0xFF)));
            os.Put(static_cast<Ch>(0x80 | ((codepoint >> 6) & 0x3F)));
            os.Put(static_cast<Ch>(0x80 | (codepoint & 0x3F)));
        }
        else {
            CEREAL_RAPIDJSON_ASSERT(codepoint <= 0x10FFFF);
            os.Put(static_cast<Ch>(0xF0 | ((codepoint >> 18) & 0xFF)));
            os.Put(static_cast<Ch>(0x80 | ((codepoint >> 12) & 0x3F)));
            os.Put(static_cast<Ch>(0x80 | ((codepoint >> 6) & 0x3F)));
            os.Put(static_cast<Ch>(0x80 | (codepoint & 0x3F)));
        }
    }

    template<typename OutputStream>
    static void EncodeUnsafe(OutputStream& os, unsigned codepoint) {
        if (codepoint <= 0x7F) 
            PutUnsafe(os, static_cast<Ch>(codepoint & 0xFF));
        else if (codepoint <= 0x7FF) {
            PutUnsafe(os, static_cast<Ch>(0xC0 | ((codepoint >> 6) & 0xFF)));
            PutUnsafe(os, static_cast<Ch>(0x80 | ((codepoint & 0x3F))));
        }
        else if (codepoint <= 0xFFFF) {
            PutUnsafe(os, static_cast<Ch>(0xE0 | ((codepoint >> 12) & 0xFF)));
            PutUnsafe(os, static_cast<Ch>(0x80 | ((codepoint >> 6) & 0x3F)));
            PutUnsafe(os, static_cast<Ch>(0x80 | (codepoint & 0x3F)));
        }
        else {
            CEREAL_RAPIDJSON_ASSERT(codepoint <= 0x10FFFF);
            PutUnsafe(os, static_cast<Ch>(0xF0 | ((codepoint >> 18) & 0xFF)));
            PutUnsafe(os, static_cast<Ch>(0x80 | ((codepoint >> 12) & 0x3F)));
            PutUnsafe(os, static_cast<Ch>(0x80 | ((codepoint >> 6) & 0x3F)));
            PutUnsafe(os, static_cast<Ch>(0x80 | (codepoint & 0x3F)));
        }
    }

    template <typename InputStream>
    static bool Decode(InputStream& is, unsigned* codepoint) {
#define CEREAL_RAPIDJSON_COPY() c = is.Take(); *codepoint = (*codepoint << 6) | (static_cast<unsigned char>(c) & 0x3Fu)
#define CEREAL_RAPIDJSON_TRANS(mask) result &= ((GetRange(static_cast<unsigned char>(c)) & mask) != 0)
#define CEREAL_RAPIDJSON_TAIL() CEREAL_RAPIDJSON_COPY(); CEREAL_RAPIDJSON_TRANS(0x70)
        typename InputStream::Ch c = is.Take();
        if (!(c & 0x80)) {
            *codepoint = static_cast<unsigned char>(c);
            return true;
        }

        unsigned char type = GetRange(static_cast<unsigned char>(c));
        if (type >= 32) {
            *codepoint = 0;
        } else {
            *codepoint = (0xFFu >> type) & static_cast<unsigned char>(c);
        }
        bool result = true;
        switch (type) {
        case 2: CEREAL_RAPIDJSON_TAIL(); return result;
        case 3: CEREAL_RAPIDJSON_TAIL(); CEREAL_RAPIDJSON_TAIL(); return result;
        case 4: CEREAL_RAPIDJSON_COPY(); CEREAL_RAPIDJSON_TRANS(0x50); CEREAL_RAPIDJSON_TAIL(); return result;
        case 5: CEREAL_RAPIDJSON_COPY(); CEREAL_RAPIDJSON_TRANS(0x10); CEREAL_RAPIDJSON_TAIL(); CEREAL_RAPIDJSON_TAIL(); return result;
        case 6: CEREAL_RAPIDJSON_TAIL(); CEREAL_RAPIDJSON_TAIL(); CEREAL_RAPIDJSON_TAIL(); return result;
        case 10: CEREAL_RAPIDJSON_COPY(); CEREAL_RAPIDJSON_TRANS(0x20); CEREAL_RAPIDJSON_TAIL(); return result;
        case 11: CEREAL_RAPIDJSON_COPY(); CEREAL_RAPIDJSON_TRANS(0x60); CEREAL_RAPIDJSON_TAIL(); CEREAL_RAPIDJSON_TAIL(); return result;
        default: return false;
        }
#undef CEREAL_RAPIDJSON_COPY
#undef CEREAL_RAPIDJSON_TRANS
#undef CEREAL_RAPIDJSON_TAIL
    }

    template <typename InputStream, typename OutputStream>
    static bool Validate(InputStream& is, OutputStream& os) {
#define CEREAL_RAPIDJSON_COPY() os.Put(c = is.Take())
#define CEREAL_RAPIDJSON_TRANS(mask) result &= ((GetRange(static_cast<unsigned char>(c)) & mask) != 0)
#define CEREAL_RAPIDJSON_TAIL() CEREAL_RAPIDJSON_COPY(); CEREAL_RAPIDJSON_TRANS(0x70)
        Ch c;
        CEREAL_RAPIDJSON_COPY();
        if (!(c & 0x80))
            return true;

        bool result = true;
        switch (GetRange(static_cast<unsigned char>(c))) {
        case 2: CEREAL_RAPIDJSON_TAIL(); return result;
        case 3: CEREAL_RAPIDJSON_TAIL(); CEREAL_RAPIDJSON_TAIL(); return result;
        case 4: CEREAL_RAPIDJSON_COPY(); CEREAL_RAPIDJSON_TRANS(0x50); CEREAL_RAPIDJSON_TAIL(); return result;
        case 5: CEREAL_RAPIDJSON_COPY(); CEREAL_RAPIDJSON_TRANS(0x10); CEREAL_RAPIDJSON_TAIL(); CEREAL_RAPIDJSON_TAIL(); return result;
        case 6: CEREAL_RAPIDJSON_TAIL(); CEREAL_RAPIDJSON_TAIL(); CEREAL_RAPIDJSON_TAIL(); return result;
        case 10: CEREAL_RAPIDJSON_COPY(); CEREAL_RAPIDJSON_TRANS(0x20); CEREAL_RAPIDJSON_TAIL(); return result;
        case 11: CEREAL_RAPIDJSON_COPY(); CEREAL_RAPIDJSON_TRANS(0x60); CEREAL_RAPIDJSON_TAIL(); CEREAL_RAPIDJSON_TAIL(); return result;
        default: return false;
        }
#undef CEREAL_RAPIDJSON_COPY
#undef CEREAL_RAPIDJSON_TRANS
#undef CEREAL_RAPIDJSON_TAIL
    }

    static unsigned char GetRange(unsigned char c) {
        // Referring to DFA of http://bjoern.hoehrmann.de/utf-8/decoder/dfa/
        // With new mapping 1 -> 0x10, 7 -> 0x20, 9 -> 0x40, such that AND operation can test multiple types.
        static const unsigned char type[] = {
            0,0,0,0,0,0,0,0,0,0,0,0,0,0,0,0,  0,0,0,0,0,0,0,0,0,0,0,0,0,0,0,0,
            0,0,0,0,0,0,0,0,0,0,0,0,0,0,0,0,  0,0,0,0,0,0,0,0,0,0,0,0,0,0,0,0,
            0,0,0,0,0,0,0,0,0,0,0,0,0,0,0,0,  0,0,0,0,0,0,0,0,0,0,0,0,0,0,0,0,
            0,0,0,0,0,0,0,0,0,0,0,0,0,0,0,0,  0,0,0,0,0,0,0,0,0,0,0,0,0,0,0,0,
            0x10,0x10,0x10,0x10,0x10,0x10,0x10,0x10,0x10,0x10,0x10,0x10,0x10,0x10,0x10,0x10,
            0x40,0x40,0x40,0x40,0x40,0x40,0x40,0x40,0x40,0x40,0x40,0x40,0x40,0x40,0x40,0x40,
            0x20,0x20,0x20,0x20,0x20,0x20,0x20,0x20,0x20,0x20,0x20,0x20,0x20,0x20,0x20,0x20,
            0x20,0x20,0x20,0x20,0x20,0x20,0x20,0x20,0x20,0x20,0x20,0x20,0x20,0x20,0x20,0x20,
            8,8,2,2,2,2,2,2,2,2,2,2,2,2,2,2,  2,2,2,2,2,2,2,2,2,2,2,2,2,2,2,2,
            10,3,3,3,3,3,3,3,3,3,3,3,3,4,3,3, 11,6,6,6,5,8,8,8,8,8,8,8,8,8,8,8,
        };
        return type[c];
    }

    template <typename InputByteStream>
    static CharType TakeBOM(InputByteStream& is) {
        CEREAL_RAPIDJSON_STATIC_ASSERT(sizeof(typename InputByteStream::Ch) == 1);
        typename InputByteStream::Ch c = Take(is);
        if (static_cast<unsigned char>(c) != 0xEFu) return c;
        c = is.Take();
        if (static_cast<unsigned char>(c) != 0xBBu) return c;
        c = is.Take();
        if (static_cast<unsigned char>(c) != 0xBFu) return c;
        c = is.Take();
        return c;
    }

    template <typename InputByteStream>
    static Ch Take(InputByteStream& is) {
        CEREAL_RAPIDJSON_STATIC_ASSERT(sizeof(typename InputByteStream::Ch) == 1);
        return static_cast<Ch>(is.Take());
    }

    template <typename OutputByteStream>
    static void PutBOM(OutputByteStream& os) {
        CEREAL_RAPIDJSON_STATIC_ASSERT(sizeof(typename OutputByteStream::Ch) == 1);
        os.Put(static_cast<typename OutputByteStream::Ch>(0xEFu));
        os.Put(static_cast<typename OutputByteStream::Ch>(0xBBu));
        os.Put(static_cast<typename OutputByteStream::Ch>(0xBFu));
    }

    template <typename OutputByteStream>
    static void Put(OutputByteStream& os, Ch c) {
        CEREAL_RAPIDJSON_STATIC_ASSERT(sizeof(typename OutputByteStream::Ch) == 1);
        os.Put(static_cast<typename OutputByteStream::Ch>(c));
    }
};

///////////////////////////////////////////////////////////////////////////////
// UTF16

//! UTF-16 encoding.
/*! http://en.wikipedia.org/wiki/UTF-16
    http://tools.ietf.org/html/rfc2781
    \tparam CharType Type for storing 16-bit UTF-16 data. Default is wchar_t. C++11 may use char16_t instead.
    \note implements Encoding concept

    \note For in-memory access, no need to concern endianness. The code units and code points are represented by CPU's endianness.
    For streaming, use UTF16LE and UTF16BE, which handle endianness.
*/
template<typename CharType = wchar_t>
struct UTF16 {
    typedef CharType Ch;
    CEREAL_RAPIDJSON_STATIC_ASSERT(sizeof(Ch) >= 2);

    enum { supportUnicode = 1 };

    template<typename OutputStream>
    static void Encode(OutputStream& os, unsigned codepoint) {
        CEREAL_RAPIDJSON_STATIC_ASSERT(sizeof(typename OutputStream::Ch) >= 2);
        if (codepoint <= 0xFFFF) {
            CEREAL_RAPIDJSON_ASSERT(codepoint < 0xD800 || codepoint > 0xDFFF); // Code point itself cannot be surrogate pair 
            os.Put(static_cast<typename OutputStream::Ch>(codepoint));
        }
        else {
            CEREAL_RAPIDJSON_ASSERT(codepoint <= 0x10FFFF);
            unsigned v = codepoint - 0x10000;
            os.Put(static_cast<typename OutputStream::Ch>((v >> 10) | 0xD800));
            os.Put(static_cast<typename OutputStream::Ch>((v & 0x3FF) | 0xDC00));
        }
    }


    template<typename OutputStream>
    static void EncodeUnsafe(OutputStream& os, unsigned codepoint) {
        CEREAL_RAPIDJSON_STATIC_ASSERT(sizeof(typename OutputStream::Ch) >= 2);
        if (codepoint <= 0xFFFF) {
            CEREAL_RAPIDJSON_ASSERT(codepoint < 0xD800 || codepoint > 0xDFFF); // Code point itself cannot be surrogate pair 
            PutUnsafe(os, static_cast<typename OutputStream::Ch>(codepoint));
        }
        else {
            CEREAL_RAPIDJSON_ASSERT(codepoint <= 0x10FFFF);
            unsigned v = codepoint - 0x10000;
            PutUnsafe(os, static_cast<typename OutputStream::Ch>((v >> 10) | 0xD800));
            PutUnsafe(os, static_cast<typename OutputStream::Ch>((v & 0x3FF) | 0xDC00));
        }
    }

    template <typename InputStream>
    static bool Decode(InputStream& is, unsigned* codepoint) {
        CEREAL_RAPIDJSON_STATIC_ASSERT(sizeof(typename InputStream::Ch) >= 2);
        typename InputStream::Ch c = is.Take();
        if (c < 0xD800 || c > 0xDFFF) {
            *codepoint = static_cast<unsigned>(c);
            return true;
        }
        else if (c <= 0xDBFF) {
            *codepoint = (static_cast<unsigned>(c) & 0x3FF) << 10;
            c = is.Take();
            *codepoint |= (static_cast<unsigned>(c) & 0x3FF);
            *codepoint += 0x10000;
            return c >= 0xDC00 && c <= 0xDFFF;
        }
        return false;
    }

    template <typename InputStream, typename OutputStream>
    static bool Validate(InputStream& is, OutputStream& os) {
        CEREAL_RAPIDJSON_STATIC_ASSERT(sizeof(typename InputStream::Ch) >= 2);
        CEREAL_RAPIDJSON_STATIC_ASSERT(sizeof(typename OutputStream::Ch) >= 2);
        typename InputStream::Ch c;
        os.Put(static_cast<typename OutputStream::Ch>(c = is.Take()));
        if (c < 0xD800 || c > 0xDFFF)
            return true;
        else if (c <= 0xDBFF) {
            os.Put(c = is.Take());
            return c >= 0xDC00 && c <= 0xDFFF;
        }
        return false;
    }
};

//! UTF-16 little endian encoding.
template<typename CharType = wchar_t>
struct UTF16LE : UTF16<CharType> {
    template <typename InputByteStream>
    static CharType TakeBOM(InputByteStream& is) {
        CEREAL_RAPIDJSON_STATIC_ASSERT(sizeof(typename InputByteStream::Ch) == 1);
        CharType c = Take(is);
        return static_cast<uint16_t>(c) == 0xFEFFu ? Take(is) : c;
    }

    template <typename InputByteStream>
    static CharType Take(InputByteStream& is) {
        CEREAL_RAPIDJSON_STATIC_ASSERT(sizeof(typename InputByteStream::Ch) == 1);
        unsigned c = static_cast<uint8_t>(is.Take());
        c |= static_cast<unsigned>(static_cast<uint8_t>(is.Take())) << 8;
        return static_cast<CharType>(c);
    }

    template <typename OutputByteStream>
    static void PutBOM(OutputByteStream& os) {
        CEREAL_RAPIDJSON_STATIC_ASSERT(sizeof(typename OutputByteStream::Ch) == 1);
        os.Put(static_cast<typename OutputByteStream::Ch>(0xFFu));
        os.Put(static_cast<typename OutputByteStream::Ch>(0xFEu));
    }

    template <typename OutputByteStream>
    static void Put(OutputByteStream& os, CharType c) {
        CEREAL_RAPIDJSON_STATIC_ASSERT(sizeof(typename OutputByteStream::Ch) == 1);
        os.Put(static_cast<typename OutputByteStream::Ch>(static_cast<unsigned>(c) & 0xFFu));
        os.Put(static_cast<typename OutputByteStream::Ch>((static_cast<unsigned>(c) >> 8) & 0xFFu));
    }
};

//! UTF-16 big endian encoding.
template<typename CharType = wchar_t>
struct UTF16BE : UTF16<CharType> {
    template <typename InputByteStream>
    static CharType TakeBOM(InputByteStream& is) {
        CEREAL_RAPIDJSON_STATIC_ASSERT(sizeof(typename InputByteStream::Ch) == 1);
        CharType c = Take(is);
        return static_cast<uint16_t>(c) == 0xFEFFu ? Take(is) : c;
    }

    template <typename InputByteStream>
    static CharType Take(InputByteStream& is) {
        CEREAL_RAPIDJSON_STATIC_ASSERT(sizeof(typename InputByteStream::Ch) == 1);
        unsigned c = static_cast<unsigned>(static_cast<uint8_t>(is.Take())) << 8;
        c |= static_cast<unsigned>(static_cast<uint8_t>(is.Take()));
        return static_cast<CharType>(c);
    }

    template <typename OutputByteStream>
    static void PutBOM(OutputByteStream& os) {
        CEREAL_RAPIDJSON_STATIC_ASSERT(sizeof(typename OutputByteStream::Ch) == 1);
        os.Put(static_cast<typename OutputByteStream::Ch>(0xFEu));
        os.Put(static_cast<typename OutputByteStream::Ch>(0xFFu));
    }

    template <typename OutputByteStream>
    static void Put(OutputByteStream& os, CharType c) {
        CEREAL_RAPIDJSON_STATIC_ASSERT(sizeof(typename OutputByteStream::Ch) == 1);
        os.Put(static_cast<typename OutputByteStream::Ch>((static_cast<unsigned>(c) >> 8) & 0xFFu));
        os.Put(static_cast<typename OutputByteStream::Ch>(static_cast<unsigned>(c) & 0xFFu));
    }
};

///////////////////////////////////////////////////////////////////////////////
// UTF32

//! UTF-32 encoding. 
/*! http://en.wikipedia.org/wiki/UTF-32
    \tparam CharType Type for storing 32-bit UTF-32 data. Default is unsigned. C++11 may use char32_t instead.
    \note implements Encoding concept

    \note For in-memory access, no need to concern endianness. The code units and code points are represented by CPU's endianness.
    For streaming, use UTF32LE and UTF32BE, which handle endianness.
*/
template<typename CharType = unsigned>
struct UTF32 {
    typedef CharType Ch;
    CEREAL_RAPIDJSON_STATIC_ASSERT(sizeof(Ch) >= 4);

    enum { supportUnicode = 1 };

    template<typename OutputStream>
    static void Encode(OutputStream& os, unsigned codepoint) {
        CEREAL_RAPIDJSON_STATIC_ASSERT(sizeof(typename OutputStream::Ch) >= 4);
        CEREAL_RAPIDJSON_ASSERT(codepoint <= 0x10FFFF);
        os.Put(codepoint);
    }

    template<typename OutputStream>
    static void EncodeUnsafe(OutputStream& os, unsigned codepoint) {
        CEREAL_RAPIDJSON_STATIC_ASSERT(sizeof(typename OutputStream::Ch) >= 4);
        CEREAL_RAPIDJSON_ASSERT(codepoint <= 0x10FFFF);
        PutUnsafe(os, codepoint);
    }

    template <typename InputStream>
    static bool Decode(InputStream& is, unsigned* codepoint) {
        CEREAL_RAPIDJSON_STATIC_ASSERT(sizeof(typename InputStream::Ch) >= 4);
        Ch c = is.Take();
        *codepoint = c;
        return c <= 0x10FFFF;
    }

    template <typename InputStream, typename OutputStream>
    static bool Validate(InputStream& is, OutputStream& os) {
        CEREAL_RAPIDJSON_STATIC_ASSERT(sizeof(typename InputStream::Ch) >= 4);
        Ch c;
        os.Put(c = is.Take());
        return c <= 0x10FFFF;
    }
};

//! UTF-32 little endian enocoding.
template<typename CharType = unsigned>
struct UTF32LE : UTF32<CharType> {
    template <typename InputByteStream>
    static CharType TakeBOM(InputByteStream& is) {
        CEREAL_RAPIDJSON_STATIC_ASSERT(sizeof(typename InputByteStream::Ch) == 1);
        CharType c = Take(is);
        return static_cast<uint32_t>(c) == 0x0000FEFFu ? Take(is) : c;
    }

    template <typename InputByteStream>
    static CharType Take(InputByteStream& is) {
        CEREAL_RAPIDJSON_STATIC_ASSERT(sizeof(typename InputByteStream::Ch) == 1);
        unsigned c = static_cast<uint8_t>(is.Take());
        c |= static_cast<unsigned>(static_cast<uint8_t>(is.Take())) << 8;
        c |= static_cast<unsigned>(static_cast<uint8_t>(is.Take())) << 16;
        c |= static_cast<unsigned>(static_cast<uint8_t>(is.Take())) << 24;
        return static_cast<CharType>(c);
    }

    template <typename OutputByteStream>
    static void PutBOM(OutputByteStream& os) {
        CEREAL_RAPIDJSON_STATIC_ASSERT(sizeof(typename OutputByteStream::Ch) == 1);
        os.Put(static_cast<typename OutputByteStream::Ch>(0xFFu));
        os.Put(static_cast<typename OutputByteStream::Ch>(0xFEu));
        os.Put(static_cast<typename OutputByteStream::Ch>(0x00u));
        os.Put(static_cast<typename OutputByteStream::Ch>(0x00u));
    }

    template <typename OutputByteStream>
    static void Put(OutputByteStream& os, CharType c) {
        CEREAL_RAPIDJSON_STATIC_ASSERT(sizeof(typename OutputByteStream::Ch) == 1);
        os.Put(static_cast<typename OutputByteStream::Ch>(c & 0xFFu));
        os.Put(static_cast<typename OutputByteStream::Ch>((c >> 8) & 0xFFu));
        os.Put(static_cast<typename OutputByteStream::Ch>((c >> 16) & 0xFFu));
        os.Put(static_cast<typename OutputByteStream::Ch>((c >> 24) & 0xFFu));
    }
};

//! UTF-32 big endian encoding.
template<typename CharType = unsigned>
struct UTF32BE : UTF32<CharType> {
    template <typename InputByteStream>
    static CharType TakeBOM(InputByteStream& is) {
        CEREAL_RAPIDJSON_STATIC_ASSERT(sizeof(typename InputByteStream::Ch) == 1);
        CharType c = Take(is);
        return static_cast<uint32_t>(c) == 0x0000FEFFu ? Take(is) : c; 
    }

    template <typename InputByteStream>
    static CharType Take(InputByteStream& is) {
        CEREAL_RAPIDJSON_STATIC_ASSERT(sizeof(typename InputByteStream::Ch) == 1);
        unsigned c = static_cast<unsigned>(static_cast<uint8_t>(is.Take())) << 24;
        c |= static_cast<unsigned>(static_cast<uint8_t>(is.Take())) << 16;
        c |= static_cast<unsigned>(static_cast<uint8_t>(is.Take())) << 8;
        c |= static_cast<unsigned>(static_cast<uint8_t>(is.Take()));
        return static_cast<CharType>(c);
    }

    template <typename OutputByteStream>
    static void PutBOM(OutputByteStream& os) {
        CEREAL_RAPIDJSON_STATIC_ASSERT(sizeof(typename OutputByteStream::Ch) == 1);
        os.Put(static_cast<typename OutputByteStream::Ch>(0x00u));
        os.Put(static_cast<typename OutputByteStream::Ch>(0x00u));
        os.Put(static_cast<typename OutputByteStream::Ch>(0xFEu));
        os.Put(static_cast<typename OutputByteStream::Ch>(0xFFu));
    }

    template <typename OutputByteStream>
    static void Put(OutputByteStream& os, CharType c) {
        CEREAL_RAPIDJSON_STATIC_ASSERT(sizeof(typename OutputByteStream::Ch) == 1);
        os.Put(static_cast<typename OutputByteStream::Ch>((c >> 24) & 0xFFu));
        os.Put(static_cast<typename OutputByteStream::Ch>((c >> 16) & 0xFFu));
        os.Put(static_cast<typename OutputByteStream::Ch>((c >> 8) & 0xFFu));
        os.Put(static_cast<typename OutputByteStream::Ch>(c & 0xFFu));
    }
};

///////////////////////////////////////////////////////////////////////////////
// ASCII

//! ASCII encoding.
/*! http://en.wikipedia.org/wiki/ASCII
    \tparam CharType Code unit for storing 7-bit ASCII data. Default is char.
    \note implements Encoding concept
*/
template<typename CharType = char>
struct ASCII {
    typedef CharType Ch;

    enum { supportUnicode = 0 };

    template<typename OutputStream>
    static void Encode(OutputStream& os, unsigned codepoint) {
        CEREAL_RAPIDJSON_ASSERT(codepoint <= 0x7F);
        os.Put(static_cast<Ch>(codepoint & 0xFF));
    }

    template<typename OutputStream>
    static void EncodeUnsafe(OutputStream& os, unsigned codepoint) {
        CEREAL_RAPIDJSON_ASSERT(codepoint <= 0x7F);
        PutUnsafe(os, static_cast<Ch>(codepoint & 0xFF));
    }

    template <typename InputStream>
    static bool Decode(InputStream& is, unsigned* codepoint) {
        uint8_t c = static_cast<uint8_t>(is.Take());
        *codepoint = c;
        return c <= 0X7F;
    }

    template <typename InputStream, typename OutputStream>
    static bool Validate(InputStream& is, OutputStream& os) {
        uint8_t c = static_cast<uint8_t>(is.Take());
        os.Put(static_cast<typename OutputStream::Ch>(c));
        return c <= 0x7F;
    }

    template <typename InputByteStream>
    static CharType TakeBOM(InputByteStream& is) {
        CEREAL_RAPIDJSON_STATIC_ASSERT(sizeof(typename InputByteStream::Ch) == 1);
        uint8_t c = static_cast<uint8_t>(Take(is));
        return static_cast<Ch>(c);
    }

    template <typename InputByteStream>
    static Ch Take(InputByteStream& is) {
        CEREAL_RAPIDJSON_STATIC_ASSERT(sizeof(typename InputByteStream::Ch) == 1);
        return static_cast<Ch>(is.Take());
    }

    template <typename OutputByteStream>
    static void PutBOM(OutputByteStream& os) {
        CEREAL_RAPIDJSON_STATIC_ASSERT(sizeof(typename OutputByteStream::Ch) == 1);
        (void)os;
    }

    template <typename OutputByteStream>
    static void Put(OutputByteStream& os, Ch c) {
        CEREAL_RAPIDJSON_STATIC_ASSERT(sizeof(typename OutputByteStream::Ch) == 1);
        os.Put(static_cast<typename OutputByteStream::Ch>(c));
    }
};

///////////////////////////////////////////////////////////////////////////////
// AutoUTF

//! Runtime-specified UTF encoding type of a stream.
enum UTFType {
    kUTF8 = 0,      //!< UTF-8.
    kUTF16LE = 1,   //!< UTF-16 little endian.
    kUTF16BE = 2,   //!< UTF-16 big endian.
    kUTF32LE = 3,   //!< UTF-32 little endian.
    kUTF32BE = 4    //!< UTF-32 big endian.
};

//! Dynamically select encoding according to stream's runtime-specified UTF encoding type.
/*! \note This class can be used with AutoUTFInputtStream and AutoUTFOutputStream, which provides GetType().
*/
template<typename CharType>
struct AutoUTF {
    typedef CharType Ch;

    enum { supportUnicode = 1 };

#define CEREAL_RAPIDJSON_ENCODINGS_FUNC(x) UTF8<Ch>::x, UTF16LE<Ch>::x, UTF16BE<Ch>::x, UTF32LE<Ch>::x, UTF32BE<Ch>::x

    template<typename OutputStream>
    static CEREAL_RAPIDJSON_FORCEINLINE void Encode(OutputStream& os, unsigned codepoint) {
        typedef void (*EncodeFunc)(OutputStream&, unsigned);
        static const EncodeFunc f[] = { CEREAL_RAPIDJSON_ENCODINGS_FUNC(Encode) };
        (*f[os.GetType()])(os, codepoint);
    }

    template<typename OutputStream>
    static CEREAL_RAPIDJSON_FORCEINLINE void EncodeUnsafe(OutputStream& os, unsigned codepoint) {
        typedef void (*EncodeFunc)(OutputStream&, unsigned);
        static const EncodeFunc f[] = { CEREAL_RAPIDJSON_ENCODINGS_FUNC(EncodeUnsafe) };
        (*f[os.GetType()])(os, codepoint);
    }

    template <typename InputStream>
    static CEREAL_RAPIDJSON_FORCEINLINE bool Decode(InputStream& is, unsigned* codepoint) {
        typedef bool (*DecodeFunc)(InputStream&, unsigned*);
        static const DecodeFunc f[] = { CEREAL_RAPIDJSON_ENCODINGS_FUNC(Decode) };
        return (*f[is.GetType()])(is, codepoint);
    }

    template <typename InputStream, typename OutputStream>
    static CEREAL_RAPIDJSON_FORCEINLINE bool Validate(InputStream& is, OutputStream& os) {
        typedef bool (*ValidateFunc)(InputStream&, OutputStream&);
        static const ValidateFunc f[] = { CEREAL_RAPIDJSON_ENCODINGS_FUNC(Validate) };
        return (*f[is.GetType()])(is, os);
    }

#undef CEREAL_RAPIDJSON_ENCODINGS_FUNC
};

///////////////////////////////////////////////////////////////////////////////
// Transcoder

//! Encoding conversion.
template<typename SourceEncoding, typename TargetEncoding>
struct Transcoder {
    //! Take one Unicode codepoint from source encoding, convert it to target encoding and put it to the output stream.
    template<typename InputStream, typename OutputStream>
    static CEREAL_RAPIDJSON_FORCEINLINE bool Transcode(InputStream& is, OutputStream& os) {
        unsigned codepoint;
        if (!SourceEncoding::Decode(is, &codepoint))
            return false;
        TargetEncoding::Encode(os, codepoint);
        return true;
    }

    template<typename InputStream, typename OutputStream>
    static CEREAL_RAPIDJSON_FORCEINLINE bool TranscodeUnsafe(InputStream& is, OutputStream& os) {
        unsigned codepoint;
        if (!SourceEncoding::Decode(is, &codepoint))
            return false;
        TargetEncoding::EncodeUnsafe(os, codepoint);
        return true;
    }

    //! Validate one Unicode codepoint from an encoded stream.
    template<typename InputStream, typename OutputStream>
    static CEREAL_RAPIDJSON_FORCEINLINE bool Validate(InputStream& is, OutputStream& os) {
        return Transcode(is, os);   // Since source/target encoding is different, must transcode.
    }
};

// Forward declaration.
template<typename Stream>
inline void PutUnsafe(Stream& stream, typename Stream::Ch c);

//! Specialization of Transcoder with same source and target encoding.
template<typename Encoding>
struct Transcoder<Encoding, Encoding> {
    template<typename InputStream, typename OutputStream>
    static CEREAL_RAPIDJSON_FORCEINLINE bool Transcode(InputStream& is, OutputStream& os) {
        os.Put(is.Take());  // Just copy one code unit. This semantic is different from primary template class.
        return true;
    }
    
    template<typename InputStream, typename OutputStream>
    static CEREAL_RAPIDJSON_FORCEINLINE bool TranscodeUnsafe(InputStream& is, OutputStream& os) {
        PutUnsafe(os, is.Take());  // Just copy one code unit. This semantic is different from primary template class.
        return true;
    }
    
    template<typename InputStream, typename OutputStream>
    static CEREAL_RAPIDJSON_FORCEINLINE bool Validate(InputStream& is, OutputStream& os) {
        return Encoding::Validate(is, os);  // source/target encoding are the same
    }
};

CEREAL_RAPIDJSON_NAMESPACE_END

#if defined(__GNUC__) || (defined(_MSC_VER) && !defined(__clang__))
CEREAL_RAPIDJSON_DIAG_POP
#endif

#endif // CEREAL_RAPIDJSON_ENCODINGS_H_
```
