## Supplemental code for "The motif composition of variable-number tandem repeats impacts gene expression": en.h

```
// Tencent is pleased to support the open source community by making RapidJSON available.
// 
// Copyright (C) 2015 THL A29 Limited, a Tencent company, and Milo Yip. All rights reserved.
//
// Licensed under the MIT License (the "License"); you may not use this file except
// in compliance with the License. You may obtain a copy of the License at
//
// http://opensource.org/licenses/MIT
//
// Unless required by applicable law or agreed to in writing, software distributed 
// under the License is distributed on an "AS IS" BASIS, WITHOUT WARRANTIES OR 
// CONDITIONS OF ANY KIND, either express or implied. See the License for the 
// specific language governing permissions and limitations under the License.

#ifndef CEREAL_RAPIDJSON_ERROR_EN_H_
#define CEREAL_RAPIDJSON_ERROR_EN_H_

#include "error.h"

#ifdef __clang__
CEREAL_RAPIDJSON_DIAG_PUSH
CEREAL_RAPIDJSON_DIAG_OFF(switch-enum)
CEREAL_RAPIDJSON_DIAG_OFF(covered-switch-default)
#endif

CEREAL_RAPIDJSON_NAMESPACE_BEGIN

//! Maps error code of parsing into error message.
/*!
    \ingroup CEREAL_RAPIDJSON_ERRORS
    \param parseErrorCode Error code obtained in parsing.
    \return the error message.
    \note User can make a copy of this function for localization.
        Using switch-case is safer for future modification of error codes.
*/
inline const CEREAL_RAPIDJSON_ERROR_CHARTYPE* GetParseError_En(ParseErrorCode parseErrorCode) {
    switch (parseErrorCode) {
        case kParseErrorNone:                           return CEREAL_RAPIDJSON_ERROR_STRING("No error.");

        case kParseErrorDocumentEmpty:                  return CEREAL_RAPIDJSON_ERROR_STRING("The document is empty.");
        case kParseErrorDocumentRootNotSingular:        return CEREAL_RAPIDJSON_ERROR_STRING("The document root must not be followed by other values.");
    
        case kParseErrorValueInvalid:                   return CEREAL_RAPIDJSON_ERROR_STRING("Invalid value.");
    
        case kParseErrorObjectMissName:                 return CEREAL_RAPIDJSON_ERROR_STRING("Missing a name for object member.");
        case kParseErrorObjectMissColon:                return CEREAL_RAPIDJSON_ERROR_STRING("Missing a colon after a name of object member.");
        case kParseErrorObjectMissCommaOrCurlyBracket:  return CEREAL_RAPIDJSON_ERROR_STRING("Missing a comma or '}' after an object member.");
    
        case kParseErrorArrayMissCommaOrSquareBracket:  return CEREAL_RAPIDJSON_ERROR_STRING("Missing a comma or ']' after an array element.");

        case kParseErrorStringUnicodeEscapeInvalidHex:  return CEREAL_RAPIDJSON_ERROR_STRING("Incorrect hex digit after \\u escape in string.");
        case kParseErrorStringUnicodeSurrogateInvalid:  return CEREAL_RAPIDJSON_ERROR_STRING("The surrogate pair in string is invalid.");
        case kParseErrorStringEscapeInvalid:            return CEREAL_RAPIDJSON_ERROR_STRING("Invalid escape character in string.");
        case kParseErrorStringMissQuotationMark:        return CEREAL_RAPIDJSON_ERROR_STRING("Missing a closing quotation mark in string.");
        case kParseErrorStringInvalidEncoding:          return CEREAL_RAPIDJSON_ERROR_STRING("Invalid encoding in string.");

        case kParseErrorNumberTooBig:                   return CEREAL_RAPIDJSON_ERROR_STRING("Number too big to be stored in double.");
        case kParseErrorNumberMissFraction:             return CEREAL_RAPIDJSON_ERROR_STRING("Miss fraction part in number.");
        case kParseErrorNumberMissExponent:             return CEREAL_RAPIDJSON_ERROR_STRING("Miss exponent in number.");

        case kParseErrorTermination:                    return CEREAL_RAPIDJSON_ERROR_STRING("Terminate parsing due to Handler error.");
        case kParseErrorUnspecificSyntaxError:          return CEREAL_RAPIDJSON_ERROR_STRING("Unspecific syntax error.");

        default:                                        return CEREAL_RAPIDJSON_ERROR_STRING("Unknown error.");
    }
}

CEREAL_RAPIDJSON_NAMESPACE_END

#ifdef __clang__
CEREAL_RAPIDJSON_DIAG_POP
#endif

#endif // CEREAL_RAPIDJSON_ERROR_EN_H_
```
