## Supplemental code for "The motif composition of variable-number tandem repeats impacts gene expression": error.h

```
// Tencent is pleased to support the open source community by making RapidJSON available.
// 
// Copyright (C) 2015 THL A29 Limited, a Tencent company, and Milo Yip. All rights reserved.
//
// Licensed under the MIT License (the "License"); you may not use this file except
// in compliance with the License. You may obtain a copy of the License at
//
// http://opensource.org/licenses/MIT
//
// Unless required by applicable law or agreed to in writing, software distributed 
// under the License is distributed on an "AS IS" BASIS, WITHOUT WARRANTIES OR 
// CONDITIONS OF ANY KIND, either express or implied. See the License for the 
// specific language governing permissions and limitations under the License.

#ifndef CEREAL_RAPIDJSON_ERROR_ERROR_H_
#define CEREAL_RAPIDJSON_ERROR_ERROR_H_

#include "../rapidjson.h"

#ifdef __clang__
CEREAL_RAPIDJSON_DIAG_PUSH
CEREAL_RAPIDJSON_DIAG_OFF(padded)
#endif

/*! \file error.h */

/*! \defgroup CEREAL_RAPIDJSON_ERRORS RapidJSON error handling */

///////////////////////////////////////////////////////////////////////////////
// CEREAL_RAPIDJSON_ERROR_CHARTYPE

//! Character type of error messages.
/*! \ingroup CEREAL_RAPIDJSON_ERRORS
    The default character type is \c char.
    On Windows, user can define this macro as \c TCHAR for supporting both
    unicode/non-unicode settings.
*/
#ifndef CEREAL_RAPIDJSON_ERROR_CHARTYPE
#define CEREAL_RAPIDJSON_ERROR_CHARTYPE char
#endif

///////////////////////////////////////////////////////////////////////////////
// CEREAL_RAPIDJSON_ERROR_STRING

//! Macro for converting string literial to \ref CEREAL_RAPIDJSON_ERROR_CHARTYPE[].
/*! \ingroup CEREAL_RAPIDJSON_ERRORS
    By default this conversion macro does nothing.
    On Windows, user can define this macro as \c _T(x) for supporting both
    unicode/non-unicode settings.
*/
#ifndef CEREAL_RAPIDJSON_ERROR_STRING
#define CEREAL_RAPIDJSON_ERROR_STRING(x) x
#endif

CEREAL_RAPIDJSON_NAMESPACE_BEGIN

///////////////////////////////////////////////////////////////////////////////
// ParseErrorCode

//! Error code of parsing.
/*! \ingroup CEREAL_RAPIDJSON_ERRORS
    \see GenericReader::Parse, GenericReader::GetParseErrorCode
*/
enum ParseErrorCode {
    kParseErrorNone = 0,                        //!< No error.

    kParseErrorDocumentEmpty,                   //!< The document is empty.
    kParseErrorDocumentRootNotSingular,         //!< The document root must not follow by other values.

    kParseErrorValueInvalid,                    //!< Invalid value.

    kParseErrorObjectMissName,                  //!< Missing a name for object member.
    kParseErrorObjectMissColon,                 //!< Missing a colon after a name of object member.
    kParseErrorObjectMissCommaOrCurlyBracket,   //!< Missing a comma or '}' after an object member.

    kParseErrorArrayMissCommaOrSquareBracket,   //!< Missing a comma or ']' after an array element.

    kParseErrorStringUnicodeEscapeInvalidHex,   //!< Incorrect hex digit after \\u escape in string.
    kParseErrorStringUnicodeSurrogateInvalid,   //!< The surrogate pair in string is invalid.
    kParseErrorStringEscapeInvalid,             //!< Invalid escape character in string.
    kParseErrorStringMissQuotationMark,         //!< Missing a closing quotation mark in string.
    kParseErrorStringInvalidEncoding,           //!< Invalid encoding in string.

    kParseErrorNumberTooBig,                    //!< Number too big to be stored in double.
    kParseErrorNumberMissFraction,              //!< Miss fraction part in number.
    kParseErrorNumberMissExponent,              //!< Miss exponent in number.

    kParseErrorTermination,                     //!< Parsing was terminated.
    kParseErrorUnspecificSyntaxError            //!< Unspecific syntax error.
};

//! Result of parsing (wraps ParseErrorCode)
/*!
    \ingroup CEREAL_RAPIDJSON_ERRORS
    \code
        Document doc;
        ParseResult ok = doc.Parse("[42]");
        if (!ok) {
            fprintf(stderr, "JSON parse error: %s (%u)",
                    GetParseError_En(ok.Code()), ok.Offset());
            exit(EXIT_FAILURE);
        }
    \endcode
    \see GenericReader::Parse, GenericDocument::Parse
*/
struct ParseResult {
    //!! Unspecified boolean type
    typedef bool (ParseResult::*BooleanType)() const;
public:
    //! Default constructor, no error.
    ParseResult() : code_(kParseErrorNone), offset_(0) {}
    //! Constructor to set an error.
    ParseResult(ParseErrorCode code, size_t offset) : code_(code), offset_(offset) {}

    //! Get the error code.
    ParseErrorCode Code() const { return code_; }
    //! Get the error offset, if \ref IsError(), 0 otherwise.
    size_t Offset() const { return offset_; }

    //! Explicit conversion to \c bool, returns \c true, iff !\ref IsError().
    operator BooleanType() const { return !IsError() ? &ParseResult::IsError : NULL; }
    //! Whether the result is an error.
    bool IsError() const { return code_ != kParseErrorNone; }

    bool operator==(const ParseResult& that) const { return code_ == that.code_; }
    bool operator==(ParseErrorCode code) const { return code_ == code; }
    friend bool operator==(ParseErrorCode code, const ParseResult & err) { return code == err.code_; }

    bool operator!=(const ParseResult& that) const { return !(*this == that); }
    bool operator!=(ParseErrorCode code) const { return !(*this == code); }
    friend bool operator!=(ParseErrorCode code, const ParseResult & err) { return err != code; }

    //! Reset error code.
    void Clear() { Set(kParseErrorNone); }
    //! Update error code and offset.
    void Set(ParseErrorCode code, size_t offset = 0) { code_ = code; offset_ = offset; }

private:
    ParseErrorCode code_;
    size_t offset_;
};

//! Function pointer type of GetParseError().
/*! \ingroup CEREAL_RAPIDJSON_ERRORS

    This is the prototype for \c GetParseError_X(), where \c X is a locale.
    User can dynamically change locale in runtime, e.g.:
\code
    GetParseErrorFunc GetParseError = GetParseError_En; // or whatever
    const CEREAL_RAPIDJSON_ERROR_CHARTYPE* s = GetParseError(document.GetParseErrorCode());
\endcode
*/
typedef const CEREAL_RAPIDJSON_ERROR_CHARTYPE* (*GetParseErrorFunc)(ParseErrorCode);

CEREAL_RAPIDJSON_NAMESPACE_END

#ifdef __clang__
CEREAL_RAPIDJSON_DIAG_POP
#endif

#endif // CEREAL_RAPIDJSON_ERROR_ERROR_H_
```
