## Supplemental code for "The motif composition of variable-number tandem repeats impacts gene expression": filereadstream.h

```
// Tencent is pleased to support the open source community by making RapidJSON available.
// 
// Copyright (C) 2015 THL A29 Limited, a Tencent company, and Milo Yip. All rights reserved.
//
// Licensed under the MIT License (the "License"); you may not use this file except
// in compliance with the License. You may obtain a copy of the License at
//
// http://opensource.org/licenses/MIT
//
// Unless required by applicable law or agreed to in writing, software distributed 
// under the License is distributed on an "AS IS" BASIS, WITHOUT WARRANTIES OR 
// CONDITIONS OF ANY KIND, either express or implied. See the License for the 
// specific language governing permissions and limitations under the License.

#ifndef CEREAL_RAPIDJSON_FILEREADSTREAM_H_
#define CEREAL_RAPIDJSON_FILEREADSTREAM_H_

#include "stream.h"
#include <cstdio>

#ifdef __clang__
CEREAL_RAPIDJSON_DIAG_PUSH
CEREAL_RAPIDJSON_DIAG_OFF(padded)
CEREAL_RAPIDJSON_DIAG_OFF(unreachable-code)
CEREAL_RAPIDJSON_DIAG_OFF(missing-noreturn)
#endif

CEREAL_RAPIDJSON_NAMESPACE_BEGIN

//! File byte stream for input using fread().
/*!
    \note implements Stream concept
*/
class FileReadStream {
public:
    typedef char Ch;    //!< Character type (byte).

    //! Constructor.
    /*!
        \param fp File pointer opened for read.
        \param buffer user-supplied buffer.
        \param bufferSize size of buffer in bytes. Must >=4 bytes.
    */
    FileReadStream(std::FILE* fp, char* buffer, size_t bufferSize) : fp_(fp), buffer_(buffer), bufferSize_(bufferSize), bufferLast_(0), current_(buffer_), readCount_(0), count_(0), eof_(false) { 
        CEREAL_RAPIDJSON_ASSERT(fp_ != 0);
        CEREAL_RAPIDJSON_ASSERT(bufferSize >= 4);
        Read();
    }

    Ch Peek() const { return *current_; }
    Ch Take() { Ch c = *current_; Read(); return c; }
    size_t Tell() const { return count_ + static_cast<size_t>(current_ - buffer_); }

    // Not implemented
    void Put(Ch) { CEREAL_RAPIDJSON_ASSERT(false); }
    void Flush() { CEREAL_RAPIDJSON_ASSERT(false); } 
    Ch* PutBegin() { CEREAL_RAPIDJSON_ASSERT(false); return 0; }
    size_t PutEnd(Ch*) { CEREAL_RAPIDJSON_ASSERT(false); return 0; }

    // For encoding detection only.
    const Ch* Peek4() const {
        return (current_ + 4 - !eof_ <= bufferLast_) ? current_ : 0;
    }

private:
    void Read() {
        if (current_ < bufferLast_)
            ++current_;
        else if (!eof_) {
            count_ += readCount_;
            readCount_ = std::fread(buffer_, 1, bufferSize_, fp_);
            bufferLast_ = buffer_ + readCount_ - 1;
            current_ = buffer_;

            if (readCount_ < bufferSize_) {
                buffer_[readCount_] = '\0';
                ++bufferLast_;
                eof_ = true;
            }
        }
    }

    std::FILE* fp_;
    Ch *buffer_;
    size_t bufferSize_;
    Ch *bufferLast_;
    Ch *current_;
    size_t readCount_;
    size_t count_;  //!< Number of characters read
    bool eof_;
};

CEREAL_RAPIDJSON_NAMESPACE_END

#ifdef __clang__
CEREAL_RAPIDJSON_DIAG_POP
#endif

#endif // CEREAL_RAPIDJSON_FILESTREAM_H_
```
