## Supplemental code for "The motif composition of variable-number tandem repeats impacts gene expression": filewritestream.h

```
// Tencent is pleased to support the open source community by making RapidJSON available.
// 
// Copyright (C) 2015 THL A29 Limited, a Tencent company, and Milo Yip. All rights reserved.
//
// Licensed under the MIT License (the "License"); you may not use this file except
// in compliance with the License. You may obtain a copy of the License at
//
// http://opensource.org/licenses/MIT
//
// Unless required by applicable law or agreed to in writing, software distributed 
// under the License is distributed on an "AS IS" BASIS, WITHOUT WARRANTIES OR 
// CONDITIONS OF ANY KIND, either express or implied. See the License for the 
// specific language governing permissions and limitations under the License.

#ifndef CEREAL_RAPIDJSON_FILEWRITESTREAM_H_
#define CEREAL_RAPIDJSON_FILEWRITESTREAM_H_

#include "stream.h"
#include <cstdio>

#ifdef __clang__
CEREAL_RAPIDJSON_DIAG_PUSH
CEREAL_RAPIDJSON_DIAG_OFF(unreachable-code)
#endif

CEREAL_RAPIDJSON_NAMESPACE_BEGIN

//! Wrapper of C file stream for output using fwrite().
/*!
    \note implements Stream concept
*/
class FileWriteStream {
public:
    typedef char Ch;    //!< Character type. Only support char.

    FileWriteStream(std::FILE* fp, char* buffer, size_t bufferSize) : fp_(fp), buffer_(buffer), bufferEnd_(buffer + bufferSize), current_(buffer_) { 
        CEREAL_RAPIDJSON_ASSERT(fp_ != 0);
    }

    void Put(char c) { 
        if (current_ >= bufferEnd_)
            Flush();

        *current_++ = c;
    }

    void PutN(char c, size_t n) {
        size_t avail = static_cast<size_t>(bufferEnd_ - current_);
        while (n > avail) {
            std::memset(current_, c, avail);
            current_ += avail;
            Flush();
            n -= avail;
            avail = static_cast<size_t>(bufferEnd_ - current_);
        }

        if (n > 0) {
            std::memset(current_, c, n);
            current_ += n;
        }
    }

    void Flush() {
        if (current_ != buffer_) {
            size_t result = std::fwrite(buffer_, 1, static_cast<size_t>(current_ - buffer_), fp_);
            if (result < static_cast<size_t>(current_ - buffer_)) {
                // failure deliberately ignored at this time
                // added to avoid warn_unused_result build errors
            }
            current_ = buffer_;
        }
    }

    // Not implemented
    char Peek() const { CEREAL_RAPIDJSON_ASSERT(false); return 0; }
    char Take() { CEREAL_RAPIDJSON_ASSERT(false); return 0; }
    size_t Tell() const { CEREAL_RAPIDJSON_ASSERT(false); return 0; }
    char* PutBegin() { CEREAL_RAPIDJSON_ASSERT(false); return 0; }
    size_t PutEnd(char*) { CEREAL_RAPIDJSON_ASSERT(false); return 0; }

private:
    // Prohibit copy constructor & assignment operator.
    FileWriteStream(const FileWriteStream&);
    FileWriteStream& operator=(const FileWriteStream&);

    std::FILE* fp_;
    char *buffer_;
    char *bufferEnd_;
    char *current_;
};

//! Implement specialized version of PutN() with memset() for better performance.
template<>
inline void PutN(FileWriteStream& stream, char c, size_t n) {
    stream.PutN(c, n);
}

CEREAL_RAPIDJSON_NAMESPACE_END

#ifdef __clang__
CEREAL_RAPIDJSON_DIAG_POP
#endif

#endif // CEREAL_RAPIDJSON_FILESTREAM_H_
```
