## Supplemental code for "The motif composition of variable-number tandem repeats impacts gene expression": fwd.h

```
// Tencent is pleased to support the open source community by making RapidJSON available.
// 
// Copyright (C) 2015 THL A29 Limited, a Tencent company, and Milo Yip. All rights reserved.
//
// Licensed under the MIT License (the "License"); you may not use this file except
// in compliance with the License. You may obtain a copy of the License at
//
// http://opensource.org/licenses/MIT
//
// Unless required by applicable law or agreed to in writing, software distributed 
// under the License is distributed on an "AS IS" BASIS, WITHOUT WARRANTIES OR 
// CONDITIONS OF ANY KIND, either express or implied. See the License for the 
// specific language governing permissions and limitations under the License.

#ifndef CEREAL_RAPIDJSON_FWD_H_
#define CEREAL_RAPIDJSON_FWD_H_

#include "rapidjson.h"

CEREAL_RAPIDJSON_NAMESPACE_BEGIN

// encodings.h

template<typename CharType> struct UTF8;
template<typename CharType> struct UTF16;
template<typename CharType> struct UTF16BE;
template<typename CharType> struct UTF16LE;
template<typename CharType> struct UTF32;
template<typename CharType> struct UTF32BE;
template<typename CharType> struct UTF32LE;
template<typename CharType> struct ASCII;
template<typename CharType> struct AutoUTF;

template<typename SourceEncoding, typename TargetEncoding>
struct Transcoder;

// allocators.h

class CrtAllocator;

template <typename BaseAllocator>
class MemoryPoolAllocator;

// stream.h

template <typename Encoding>
struct GenericStringStream;

typedef GenericStringStream<UTF8<char> > StringStream;

template <typename Encoding>
struct GenericInsituStringStream;

typedef GenericInsituStringStream<UTF8<char> > InsituStringStream;

// stringbuffer.h

template <typename Encoding, typename Allocator>
class GenericStringBuffer;

typedef GenericStringBuffer<UTF8<char>, CrtAllocator> StringBuffer;

// filereadstream.h

class FileReadStream;

// filewritestream.h

class FileWriteStream;

// memorybuffer.h

template <typename Allocator>
struct GenericMemoryBuffer;

typedef GenericMemoryBuffer<CrtAllocator> MemoryBuffer;

// memorystream.h

struct MemoryStream;

// reader.h

template<typename Encoding, typename Derived>
struct BaseReaderHandler;

template <typename SourceEncoding, typename TargetEncoding, typename StackAllocator>
class GenericReader;

typedef GenericReader<UTF8<char>, UTF8<char>, CrtAllocator> Reader;

// writer.h

template<typename OutputStream, typename SourceEncoding, typename TargetEncoding, typename StackAllocator, unsigned writeFlags>
class Writer;

// prettywriter.h

template<typename OutputStream, typename SourceEncoding, typename TargetEncoding, typename StackAllocator, unsigned writeFlags>
class PrettyWriter;

// document.h

template <typename Encoding, typename Allocator> 
struct GenericMember;

template <bool Const, typename Encoding, typename Allocator>
class GenericMemberIterator;

template<typename CharType>
struct GenericStringRef;

template <typename Encoding, typename Allocator> 
class GenericValue;

typedef GenericValue<UTF8<char>, MemoryPoolAllocator<CrtAllocator> > Value;

template <typename Encoding, typename Allocator, typename StackAllocator>
class GenericDocument;

typedef GenericDocument<UTF8<char>, MemoryPoolAllocator<CrtAllocator>, CrtAllocator> Document;

// pointer.h

template <typename ValueType, typename Allocator>
class GenericPointer;

typedef GenericPointer<Value, CrtAllocator> Pointer;

// schema.h

template <typename SchemaDocumentType>
class IGenericRemoteSchemaDocumentProvider;

template <typename ValueT, typename Allocator>
class GenericSchemaDocument;

typedef GenericSchemaDocument<Value, CrtAllocator> SchemaDocument;
typedef IGenericRemoteSchemaDocumentProvider<SchemaDocument> IRemoteSchemaDocumentProvider;

template <
    typename SchemaDocumentType,
    typename OutputHandler,
    typename StateAllocator>
class GenericSchemaValidator;

typedef GenericSchemaValidator<SchemaDocument, BaseReaderHandler<UTF8<char>, void>, CrtAllocator> SchemaValidator;

CEREAL_RAPIDJSON_NAMESPACE_END

#endif // CEREAL_RAPIDJSON_RAPIDJSONFWD_H_
```
