## Supplemental code for "The motif composition of variable-number tandem repeats impacts gene expression": biginteger.h

```
// Tencent is pleased to support the open source community by making RapidJSON available.
// 
// Copyright (C) 2015 THL A29 Limited, a Tencent company, and Milo Yip. All rights reserved.
//
// Licensed under the MIT License (the "License"); you may not use this file except
// in compliance with the License. You may obtain a copy of the License at
//
// http://opensource.org/licenses/MIT
//
// Unless required by applicable law or agreed to in writing, software distributed 
// under the License is distributed on an "AS IS" BASIS, WITHOUT WARRANTIES OR 
// CONDITIONS OF ANY KIND, either express or implied. See the License for the 
// specific language governing permissions and limitations under the License.

#ifndef CEREAL_RAPIDJSON_BIGINTEGER_H_
#define CEREAL_RAPIDJSON_BIGINTEGER_H_

#include "../rapidjson.h"

#if defined(_MSC_VER) && !__INTEL_COMPILER && defined(_M_AMD64)
#include <intrin.h> // for _umul128
#pragma intrinsic(_umul128)
#endif

CEREAL_RAPIDJSON_NAMESPACE_BEGIN
namespace internal {

class BigInteger {
public:
    typedef uint64_t Type;

    BigInteger(const BigInteger& rhs) : count_(rhs.count_) {
        std::memcpy(digits_, rhs.digits_, count_ * sizeof(Type));
    }

    explicit BigInteger(uint64_t u) : count_(1) {
        digits_[0] = u;
    }

    BigInteger(const char* decimals, size_t length) : count_(1) {
        CEREAL_RAPIDJSON_ASSERT(length > 0);
        digits_[0] = 0;
        size_t i = 0;
        const size_t kMaxDigitPerIteration = 19;  // 2^64 = 18446744073709551616 > 10^19
        while (length >= kMaxDigitPerIteration) {
            AppendDecimal64(decimals + i, decimals + i + kMaxDigitPerIteration);
            length -= kMaxDigitPerIteration;
            i += kMaxDigitPerIteration;
        }

        if (length > 0)
            AppendDecimal64(decimals + i, decimals + i + length);
    }
    
    BigInteger& operator=(const BigInteger &rhs)
    {
        if (this != &rhs) {
            count_ = rhs.count_;
            std::memcpy(digits_, rhs.digits_, count_ * sizeof(Type));
        }
        return *this;
    }
    
    BigInteger& operator=(uint64_t u) {
        digits_[0] = u;            
        count_ = 1;
        return *this;
    }

    BigInteger& operator+=(uint64_t u) {
        Type backup = digits_[0];
        digits_[0] += u;
        for (size_t i = 0; i < count_ - 1; i++) {
            if (digits_[i] >= backup)
                return *this; // no carry
            backup = digits_[i + 1];
            digits_[i + 1] += 1;
        }

        // Last carry
        if (digits_[count_ - 1] < backup)
            PushBack(1);

        return *this;
    }

    BigInteger& operator*=(uint64_t u) {
        if (u == 0) return *this = 0;
        if (u == 1) return *this;
        if (*this == 1) return *this = u;

        uint64_t k = 0;
        for (size_t i = 0; i < count_; i++) {
            uint64_t hi;
            digits_[i] = MulAdd64(digits_[i], u, k, &hi);
            k = hi;
        }
        
        if (k > 0)
            PushBack(k);

        return *this;
    }

    BigInteger& operator*=(uint32_t u) {
        if (u == 0) return *this = 0;
        if (u == 1) return *this;
        if (*this == 1) return *this = u;

        uint64_t k = 0;
        for (size_t i = 0; i < count_; i++) {
            const uint64_t c = digits_[i] >> 32;
            const uint64_t d = digits_[i] & 0xFFFFFFFF;
            const uint64_t uc = u * c;
            const uint64_t ud = u * d;
            const uint64_t p0 = ud + k;
            const uint64_t p1 = uc + (p0 >> 32);
            digits_[i] = (p0 & 0xFFFFFFFF) | (p1 << 32);
            k = p1 >> 32;
        }
        
        if (k > 0)
            PushBack(k);

        return *this;
    }

    BigInteger& operator<<=(size_t shift) {
        if (IsZero() || shift == 0) return *this;

        size_t offset = shift / kTypeBit;
        size_t interShift = shift % kTypeBit;
        CEREAL_RAPIDJSON_ASSERT(count_ + offset <= kCapacity);

        if (interShift == 0) {
            std::memmove(digits_ + offset, digits_, count_ * sizeof(Type));
            count_ += offset;
        }
        else {
            digits_[count_] = 0;
            for (size_t i = count_; i > 0; i--)
                digits_[i + offset] = (digits_[i] << interShift) | (digits_[i - 1] >> (kTypeBit - interShift));
            digits_[offset] = digits_[0] << interShift;
            count_ += offset;
            if (digits_[count_])
                count_++;
        }

        std::memset(digits_, 0, offset * sizeof(Type));

        return *this;
    }

    bool operator==(const BigInteger& rhs) const {
        return count_ == rhs.count_ && std::memcmp(digits_, rhs.digits_, count_ * sizeof(Type)) == 0;
    }

    bool operator==(const Type rhs) const {
        return count_ == 1 && digits_[0] == rhs;
    }

    BigInteger& MultiplyPow5(unsigned exp) {
        static const uint32_t kPow5[12] = {
            5,
            5 * 5,
            5 * 5 * 5,
            5 * 5 * 5 * 5,
            5 * 5 * 5 * 5 * 5,
            5 * 5 * 5 * 5 * 5 * 5,
            5 * 5 * 5 * 5 * 5 * 5 * 5,
            5 * 5 * 5 * 5 * 5 * 5 * 5 * 5,
            5 * 5 * 5 * 5 * 5 * 5 * 5 * 5 * 5,
            5 * 5 * 5 * 5 * 5 * 5 * 5 * 5 * 5 * 5,
            5 * 5 * 5 * 5 * 5 * 5 * 5 * 5 * 5 * 5 * 5,
            5 * 5 * 5 * 5 * 5 * 5 * 5 * 5 * 5 * 5 * 5 * 5
        };
        if (exp == 0) return *this;
        for (; exp >= 27; exp -= 27) *this *= CEREAL_RAPIDJSON_UINT64_C2(0X6765C793, 0XFA10079D); // 5^27
        for (; exp >= 13; exp -= 13) *this *= static_cast<uint32_t>(1220703125u); // 5^13
        if (exp > 0)                 *this *= kPow5[exp - 1];
        return *this;
    }

    // Compute absolute difference of this and rhs.
    // Assume this != rhs
    bool Difference(const BigInteger& rhs, BigInteger* out) const {
        int cmp = Compare(rhs);
        CEREAL_RAPIDJSON_ASSERT(cmp != 0);
        const BigInteger *a, *b;  // Makes a > b
        bool ret;
        if (cmp < 0) { a = &rhs; b = this; ret = true; }
        else         { a = this; b = &rhs; ret = false; }

        Type borrow = 0;
        for (size_t i = 0; i < a->count_; i++) {
            Type d = a->digits_[i] - borrow;
            if (i < b->count_)
                d -= b->digits_[i];
            borrow = (d > a->digits_[i]) ? 1 : 0;
            out->digits_[i] = d;
            if (d != 0)
                out->count_ = i + 1;
        }

        return ret;
    }

    int Compare(const BigInteger& rhs) const {
        if (count_ != rhs.count_)
            return count_ < rhs.count_ ? -1 : 1;

        for (size_t i = count_; i-- > 0;)
            if (digits_[i] != rhs.digits_[i])
                return digits_[i] < rhs.digits_[i] ? -1 : 1;

        return 0;
    }

    size_t GetCount() const { return count_; }
    Type GetDigit(size_t index) const { CEREAL_RAPIDJSON_ASSERT(index < count_); return digits_[index]; }
    bool IsZero() const { return count_ == 1 && digits_[0] == 0; }

private:
    void AppendDecimal64(const char* begin, const char* end) {
        uint64_t u = ParseUint64(begin, end);
        if (IsZero())
            *this = u;
        else {
            unsigned exp = static_cast<unsigned>(end - begin);
            (MultiplyPow5(exp) <<= exp) += u;   // *this = *this * 10^exp + u
        }
    }

    void PushBack(Type digit) {
        CEREAL_RAPIDJSON_ASSERT(count_ < kCapacity);
        digits_[count_++] = digit;
    }

    static uint64_t ParseUint64(const char* begin, const char* end) {
        uint64_t r = 0;
        for (const char* p = begin; p != end; ++p) {
            CEREAL_RAPIDJSON_ASSERT(*p >= '0' && *p <= '9');
            r = r * 10u + static_cast<unsigned>(*p - '0');
        }
        return r;
    }

    // Assume a * b + k < 2^128
    static uint64_t MulAdd64(uint64_t a, uint64_t b, uint64_t k, uint64_t* outHigh) {
#if defined(_MSC_VER) && defined(_M_AMD64)
        uint64_t low = _umul128(a, b, outHigh) + k;
        if (low < k)
            (*outHigh)++;
        return low;
#elif (__GNUC__ > 4 || (__GNUC__ == 4 && __GNUC_MINOR__ >= 6)) && defined(__x86_64__)
        __extension__ typedef unsigned __int128 uint128;
        uint128 p = static_cast<uint128>(a) * static_cast<uint128>(b);
        p += k;
        *outHigh = static_cast<uint64_t>(p >> 64);
        return static_cast<uint64_t>(p);
#else
        const uint64_t a0 = a & 0xFFFFFFFF, a1 = a >> 32, b0 = b & 0xFFFFFFFF, b1 = b >> 32;
        uint64_t x0 = a0 * b0, x1 = a0 * b1, x2 = a1 * b0, x3 = a1 * b1;
        x1 += (x0 >> 32); // can't give carry
        x1 += x2;
        if (x1 < x2)
            x3 += (static_cast<uint64_t>(1) << 32);
        uint64_t lo = (x1 << 32) + (x0 & 0xFFFFFFFF);
        uint64_t hi = x3 + (x1 >> 32);

        lo += k;
        if (lo < k)
            hi++;
        *outHigh = hi;
        return lo;
#endif
    }

    static const size_t kBitCount = 3328;  // 64bit * 54 > 10^1000
    static const size_t kCapacity = kBitCount / sizeof(Type);
    static const size_t kTypeBit = sizeof(Type) * 8;

    Type digits_[kCapacity];
    size_t count_;
};

} // namespace internal
CEREAL_RAPIDJSON_NAMESPACE_END

#endif // CEREAL_RAPIDJSON_BIGINTEGER_H_
```
