## Supplemental code for "The motif composition of variable-number tandem repeats impacts gene expression": diyfp.h

```
// Tencent is pleased to support the open source community by making RapidJSON available.
//
// Copyright (C) 2015 THL A29 Limited, a Tencent company, and Milo Yip. All rights reserved.
//
// Licensed under the MIT License (the "License"); you may not use this file except
// in compliance with the License. You may obtain a copy of the License at
//
// http://opensource.org/licenses/MIT
//
// Unless required by applicable law or agreed to in writing, software distributed
// under the License is distributed on an "AS IS" BASIS, WITHOUT WARRANTIES OR
// CONDITIONS OF ANY KIND, either express or implied. See the License for the
// specific language governing permissions and limitations under the License.

// This is a C++ header-only implementation of Grisu2 algorithm from the publication:
// Loitsch, Florian. "Printing floating-point numbers quickly and accurately with
// integers." ACM Sigplan Notices 45.6 (2010): 233-243.

#ifndef CEREAL_RAPIDJSON_DIYFP_H_
#define CEREAL_RAPIDJSON_DIYFP_H_

#include "../rapidjson.h"
#include <limits>

#if defined(_MSC_VER) && defined(_M_AMD64) && !defined(__INTEL_COMPILER)
#include <intrin.h>
#pragma intrinsic(_BitScanReverse64)
#pragma intrinsic(_umul128)
#endif

CEREAL_RAPIDJSON_NAMESPACE_BEGIN
namespace internal {

#ifdef __GNUC__
CEREAL_RAPIDJSON_DIAG_PUSH
CEREAL_RAPIDJSON_DIAG_OFF(effc++)
#endif

#ifdef __clang__
CEREAL_RAPIDJSON_DIAG_PUSH
CEREAL_RAPIDJSON_DIAG_OFF(padded)
#endif

struct DiyFp {
    DiyFp() : f(), e() {}

    DiyFp(uint64_t fp, int exp) : f(fp), e(exp) {}

    explicit DiyFp(double d) {
        union {
            double d;
            uint64_t u64;
        } u = { d };

        int biased_e = static_cast<int>((u.u64 & kDpExponentMask) >> kDpSignificandSize);
        uint64_t significand = (u.u64 & kDpSignificandMask);
        if (biased_e != 0) {
            f = significand + kDpHiddenBit;
            e = biased_e - kDpExponentBias;
        }
        else {
            f = significand;
            e = kDpMinExponent + 1;
        }
    }

    DiyFp operator-(const DiyFp& rhs) const {
        return DiyFp(f - rhs.f, e);
    }

    DiyFp operator*(const DiyFp& rhs) const {
#if defined(_MSC_VER) && defined(_M_AMD64)
        uint64_t h;
        uint64_t l = _umul128(f, rhs.f, &h);
        if (l & (uint64_t(1) << 63)) // rounding
            h++;
        return DiyFp(h, e + rhs.e + 64);
#elif (__GNUC__ > 4 || (__GNUC__ == 4 && __GNUC_MINOR__ >= 6)) && defined(__x86_64__)
        __extension__ typedef unsigned __int128 uint128;
        uint128 p = static_cast<uint128>(f) * static_cast<uint128>(rhs.f);
        uint64_t h = static_cast<uint64_t>(p >> 64);
        uint64_t l = static_cast<uint64_t>(p);
        if (l & (uint64_t(1) << 63)) // rounding
            h++;
        return DiyFp(h, e + rhs.e + 64);
#else
        const uint64_t M32 = 0xFFFFFFFF;
        const uint64_t a = f >> 32;
        const uint64_t b = f & M32;
        const uint64_t c = rhs.f >> 32;
        const uint64_t d = rhs.f & M32;
        const uint64_t ac = a * c;
        const uint64_t bc = b * c;
        const uint64_t ad = a * d;
        const uint64_t bd = b * d;
        uint64_t tmp = (bd >> 32) + (ad & M32) + (bc & M32);
        tmp += 1U << 31;  /// mult_round
        return DiyFp(ac + (ad >> 32) + (bc >> 32) + (tmp >> 32), e + rhs.e + 64);
#endif
    }

    DiyFp Normalize() const {
        CEREAL_RAPIDJSON_ASSERT(f != 0); // https://stackoverflow.com/a/26809183/291737
#if defined(_MSC_VER) && defined(_M_AMD64)
        unsigned long index;
        _BitScanReverse64(&index, f);
        return DiyFp(f << (63 - index), e - (63 - index));
#elif defined(__GNUC__) && __GNUC__ >= 4
        int s = __builtin_clzll(f);
        return DiyFp(f << s, e - s);
#else
        DiyFp res = *this;
        while (!(res.f & (static_cast<uint64_t>(1) << 63))) {
            res.f <<= 1;
            res.e--;
        }
        return res;
#endif
    }

    DiyFp NormalizeBoundary() const {
        DiyFp res = *this;
        while (!(res.f & (kDpHiddenBit << 1))) {
            res.f <<= 1;
            res.e--;
        }
        res.f <<= (kDiySignificandSize - kDpSignificandSize - 2);
        res.e = res.e - (kDiySignificandSize - kDpSignificandSize - 2);
        return res;
    }

    void NormalizedBoundaries(DiyFp* minus, DiyFp* plus) const {
        DiyFp pl = DiyFp((f << 1) + 1, e - 1).NormalizeBoundary();
        DiyFp mi = (f == kDpHiddenBit) ? DiyFp((f << 2) - 1, e - 2) : DiyFp((f << 1) - 1, e - 1);
        mi.f <<= mi.e - pl.e;
        mi.e = pl.e;
        *plus = pl;
        *minus = mi;
    }

    double ToDouble() const {
        union {
            double d;
            uint64_t u64;
        }u;
        CEREAL_RAPIDJSON_ASSERT(f <= kDpHiddenBit + kDpSignificandMask);
        if (e < kDpDenormalExponent) {
            // Underflow.
            return 0.0;
        }
        if (e >= kDpMaxExponent) {
            // Overflow.
            return std::numeric_limits<double>::infinity();
        }
        const uint64_t be = (e == kDpDenormalExponent && (f & kDpHiddenBit) == 0) ? 0 :
            static_cast<uint64_t>(e + kDpExponentBias);
        u.u64 = (f & kDpSignificandMask) | (be << kDpSignificandSize);
        return u.d;
    }

    static const int kDiySignificandSize = 64;
    static const int kDpSignificandSize = 52;
    static const int kDpExponentBias = 0x3FF + kDpSignificandSize;
    static const int kDpMaxExponent = 0x7FF - kDpExponentBias;
    static const int kDpMinExponent = -kDpExponentBias;
    static const int kDpDenormalExponent = -kDpExponentBias + 1;
    static const uint64_t kDpExponentMask = CEREAL_RAPIDJSON_UINT64_C2(0x7FF00000, 0x00000000);
    static const uint64_t kDpSignificandMask = CEREAL_RAPIDJSON_UINT64_C2(0x000FFFFF, 0xFFFFFFFF);
    static const uint64_t kDpHiddenBit = CEREAL_RAPIDJSON_UINT64_C2(0x00100000, 0x00000000);

    uint64_t f;
    int e;
};

inline DiyFp GetCachedPowerByIndex(size_t index) {
    // 10^-348, 10^-340, ..., 10^340
    static const uint64_t kCachedPowers_F[] = {
        CEREAL_RAPIDJSON_UINT64_C2(0xfa8fd5a0, 0x081c0288), CEREAL_RAPIDJSON_UINT64_C2(0xbaaee17f, 0xa23ebf76),
        CEREAL_RAPIDJSON_UINT64_C2(0x8b16fb20, 0x3055ac76), CEREAL_RAPIDJSON_UINT64_C2(0xcf42894a, 0x5dce35ea),
        CEREAL_RAPIDJSON_UINT64_C2(0x9a6bb0aa, 0x55653b2d), CEREAL_RAPIDJSON_UINT64_C2(0xe61acf03, 0x3d1a45df),
        CEREAL_RAPIDJSON_UINT64_C2(0xab70fe17, 0xc79ac6ca), CEREAL_RAPIDJSON_UINT64_C2(0xff77b1fc, 0xbebcdc4f),
        CEREAL_RAPIDJSON_UINT64_C2(0xbe5691ef, 0x416bd60c), CEREAL_RAPIDJSON_UINT64_C2(0x8dd01fad, 0x907ffc3c),
        CEREAL_RAPIDJSON_UINT64_C2(0xd3515c28, 0x31559a83), CEREAL_RAPIDJSON_UINT64_C2(0x9d71ac8f, 0xada6c9b5),
        CEREAL_RAPIDJSON_UINT64_C2(0xea9c2277, 0x23ee8bcb), CEREAL_RAPIDJSON_UINT64_C2(0xaecc4991, 0x4078536d),
        CEREAL_RAPIDJSON_UINT64_C2(0x823c1279, 0x5db6ce57), CEREAL_RAPIDJSON_UINT64_C2(0xc2109436, 0x4dfb5637),
        CEREAL_RAPIDJSON_UINT64_C2(0x9096ea6f, 0x3848984f), CEREAL_RAPIDJSON_UINT64_C2(0xd77485cb, 0x25823ac7),
        CEREAL_RAPIDJSON_UINT64_C2(0xa086cfcd, 0x97bf97f4), CEREAL_RAPIDJSON_UINT64_C2(0xef340a98, 0x172aace5),
        CEREAL_RAPIDJSON_UINT64_C2(0xb23867fb, 0x2a35b28e), CEREAL_RAPIDJSON_UINT64_C2(0x84c8d4df, 0xd2c63f3b),
        CEREAL_RAPIDJSON_UINT64_C2(0xc5dd4427, 0x1ad3cdba), CEREAL_RAPIDJSON_UINT64_C2(0x936b9fce, 0xbb25c996),
        CEREAL_RAPIDJSON_UINT64_C2(0xdbac6c24, 0x7d62a584), CEREAL_RAPIDJSON_UINT64_C2(0xa3ab6658, 0x0d5fdaf6),
        CEREAL_RAPIDJSON_UINT64_C2(0xf3e2f893, 0xdec3f126), CEREAL_RAPIDJSON_UINT64_C2(0xb5b5ada8, 0xaaff80b8),
        CEREAL_RAPIDJSON_UINT64_C2(0x87625f05, 0x6c7c4a8b), CEREAL_RAPIDJSON_UINT64_C2(0xc9bcff60, 0x34c13053),
        CEREAL_RAPIDJSON_UINT64_C2(0x964e858c, 0x91ba2655), CEREAL_RAPIDJSON_UINT64_C2(0xdff97724, 0x70297ebd),
        CEREAL_RAPIDJSON_UINT64_C2(0xa6dfbd9f, 0xb8e5b88f), CEREAL_RAPIDJSON_UINT64_C2(0xf8a95fcf, 0x88747d94),
        CEREAL_RAPIDJSON_UINT64_C2(0xb9447093, 0x8fa89bcf), CEREAL_RAPIDJSON_UINT64_C2(0x8a08f0f8, 0xbf0f156b),
        CEREAL_RAPIDJSON_UINT64_C2(0xcdb02555, 0x653131b6), CEREAL_RAPIDJSON_UINT64_C2(0x993fe2c6, 0xd07b7fac),
        CEREAL_RAPIDJSON_UINT64_C2(0xe45c10c4, 0x2a2b3b06), CEREAL_RAPIDJSON_UINT64_C2(0xaa242499, 0x697392d3),
        CEREAL_RAPIDJSON_UINT64_C2(0xfd87b5f2, 0x8300ca0e), CEREAL_RAPIDJSON_UINT64_C2(0xbce50864, 0x92111aeb),
        CEREAL_RAPIDJSON_UINT64_C2(0x8cbccc09, 0x6f5088cc), CEREAL_RAPIDJSON_UINT64_C2(0xd1b71758, 0xe219652c),
        CEREAL_RAPIDJSON_UINT64_C2(0x9c400000, 0x00000000), CEREAL_RAPIDJSON_UINT64_C2(0xe8d4a510, 0x00000000),
        CEREAL_RAPIDJSON_UINT64_C2(0xad78ebc5, 0xac620000), CEREAL_RAPIDJSON_UINT64_C2(0x813f3978, 0xf8940984),
        CEREAL_RAPIDJSON_UINT64_C2(0xc097ce7b, 0xc90715b3), CEREAL_RAPIDJSON_UINT64_C2(0x8f7e32ce, 0x7bea5c70),
        CEREAL_RAPIDJSON_UINT64_C2(0xd5d238a4, 0xabe98068), CEREAL_RAPIDJSON_UINT64_C2(0x9f4f2726, 0x179a2245),
        CEREAL_RAPIDJSON_UINT64_C2(0xed63a231, 0xd4c4fb27), CEREAL_RAPIDJSON_UINT64_C2(0xb0de6538, 0x8cc8ada8),
        CEREAL_RAPIDJSON_UINT64_C2(0x83c7088e, 0x1aab65db), CEREAL_RAPIDJSON_UINT64_C2(0xc45d1df9, 0x42711d9a),
        CEREAL_RAPIDJSON_UINT64_C2(0x924d692c, 0xa61be758), CEREAL_RAPIDJSON_UINT64_C2(0xda01ee64, 0x1a708dea),
        CEREAL_RAPIDJSON_UINT64_C2(0xa26da399, 0x9aef774a), CEREAL_RAPIDJSON_UINT64_C2(0xf209787b, 0xb47d6b85),
        CEREAL_RAPIDJSON_UINT64_C2(0xb454e4a1, 0x79dd1877), CEREAL_RAPIDJSON_UINT64_C2(0x865b8692, 0x5b9bc5c2),
        CEREAL_RAPIDJSON_UINT64_C2(0xc83553c5, 0xc8965d3d), CEREAL_RAPIDJSON_UINT64_C2(0x952ab45c, 0xfa97a0b3),
        CEREAL_RAPIDJSON_UINT64_C2(0xde469fbd, 0x99a05fe3), CEREAL_RAPIDJSON_UINT64_C2(0xa59bc234, 0xdb398c25),
        CEREAL_RAPIDJSON_UINT64_C2(0xf6c69a72, 0xa3989f5c), CEREAL_RAPIDJSON_UINT64_C2(0xb7dcbf53, 0x54e9bece),
        CEREAL_RAPIDJSON_UINT64_C2(0x88fcf317, 0xf22241e2), CEREAL_RAPIDJSON_UINT64_C2(0xcc20ce9b, 0xd35c78a5),
        CEREAL_RAPIDJSON_UINT64_C2(0x98165af3, 0x7b2153df), CEREAL_RAPIDJSON_UINT64_C2(0xe2a0b5dc, 0x971f303a),
        CEREAL_RAPIDJSON_UINT64_C2(0xa8d9d153, 0x5ce3b396), CEREAL_RAPIDJSON_UINT64_C2(0xfb9b7cd9, 0xa4a7443c),
        CEREAL_RAPIDJSON_UINT64_C2(0xbb764c4c, 0xa7a44410), CEREAL_RAPIDJSON_UINT64_C2(0x8bab8eef, 0xb6409c1a),
        CEREAL_RAPIDJSON_UINT64_C2(0xd01fef10, 0xa657842c), CEREAL_RAPIDJSON_UINT64_C2(0x9b10a4e5, 0xe9913129),
        CEREAL_RAPIDJSON_UINT64_C2(0xe7109bfb, 0xa19c0c9d), CEREAL_RAPIDJSON_UINT64_C2(0xac2820d9, 0x623bf429),
        CEREAL_RAPIDJSON_UINT64_C2(0x80444b5e, 0x7aa7cf85), CEREAL_RAPIDJSON_UINT64_C2(0xbf21e440, 0x03acdd2d),
        CEREAL_RAPIDJSON_UINT64_C2(0x8e679c2f, 0x5e44ff8f), CEREAL_RAPIDJSON_UINT64_C2(0xd433179d, 0x9c8cb841),
        CEREAL_RAPIDJSON_UINT64_C2(0x9e19db92, 0xb4e31ba9), CEREAL_RAPIDJSON_UINT64_C2(0xeb96bf6e, 0xbadf77d9),
        CEREAL_RAPIDJSON_UINT64_C2(0xaf87023b, 0x9bf0ee6b)
    };
    static const int16_t kCachedPowers_E[] = {
        -1220, -1193, -1166, -1140, -1113, -1087, -1060, -1034, -1007,  -980,
        -954,  -927,  -901,  -874,  -847,  -821,  -794,  -768,  -741,  -715,
        -688,  -661,  -635,  -608,  -582,  -555,  -529,  -502,  -475,  -449,
        -422,  -396,  -369,  -343,  -316,  -289,  -263,  -236,  -210,  -183,
        -157,  -130,  -103,   -77,   -50,   -24,     3,    30,    56,    83,
        109,   136,   162,   189,   216,   242,   269,   295,   322,   348,
        375,   402,   428,   455,   481,   508,   534,   561,   588,   614,
        641,   667,   694,   720,   747,   774,   800,   827,   853,   880,
        907,   933,   960,   986,  1013,  1039,  1066
    };
    CEREAL_RAPIDJSON_ASSERT(index < 87);
    return DiyFp(kCachedPowers_F[index], kCachedPowers_E[index]);
}

inline DiyFp GetCachedPower(int e, int* K) {

    //int k = static_cast<int>(ceil((-61 - e) * 0.30102999566398114)) + 374;
    double dk = (-61 - e) * 0.30102999566398114 + 347;  // dk must be positive, so can do ceiling in positive
    int k = static_cast<int>(dk);
    if (dk - k > 0.0)
        k++;

    unsigned index = static_cast<unsigned>((k >> 3) + 1);
    *K = -(-348 + static_cast<int>(index << 3));    // decimal exponent no need lookup table

    return GetCachedPowerByIndex(index);
}

inline DiyFp GetCachedPower10(int exp, int *outExp) {
    CEREAL_RAPIDJSON_ASSERT(exp >= -348);
    unsigned index = static_cast<unsigned>(exp + 348) / 8u;
    *outExp = -348 + static_cast<int>(index) * 8;
    return GetCachedPowerByIndex(index);
}

#ifdef __GNUC__
CEREAL_RAPIDJSON_DIAG_POP
#endif

#ifdef __clang__
CEREAL_RAPIDJSON_DIAG_POP
CEREAL_RAPIDJSON_DIAG_OFF(padded)
#endif

} // namespace internal
CEREAL_RAPIDJSON_NAMESPACE_END

#endif // CEREAL_RAPIDJSON_DIYFP_H_
```
