## Supplemental code for "The motif composition of variable-number tandem repeats impacts gene expression": dtoa.h

```
// Tencent is pleased to support the open source community by making RapidJSON available.
// 
// Copyright (C) 2015 THL A29 Limited, a Tencent company, and Milo Yip. All rights reserved.
//
// Licensed under the MIT License (the "License"); you may not use this file except
// in compliance with the License. You may obtain a copy of the License at
//
// http://opensource.org/licenses/MIT
//
// Unless required by applicable law or agreed to in writing, software distributed 
// under the License is distributed on an "AS IS" BASIS, WITHOUT WARRANTIES OR 
// CONDITIONS OF ANY KIND, either express or implied. See the License for the 
// specific language governing permissions and limitations under the License.

// This is a C++ header-only implementation of Grisu2 algorithm from the publication:
// Loitsch, Florian. "Printing floating-point numbers quickly and accurately with
// integers." ACM Sigplan Notices 45.6 (2010): 233-243.

#ifndef CEREAL_RAPIDJSON_DTOA_
#define CEREAL_RAPIDJSON_DTOA_

#include "itoa.h" // GetDigitsLut()
#include "diyfp.h"
#include "ieee754.h"

CEREAL_RAPIDJSON_NAMESPACE_BEGIN
namespace internal {

#ifdef __GNUC__
CEREAL_RAPIDJSON_DIAG_PUSH
CEREAL_RAPIDJSON_DIAG_OFF(effc++)
CEREAL_RAPIDJSON_DIAG_OFF(array-bounds) // some gcc versions generate wrong warnings https://gcc.gnu.org/bugzilla/show_bug.cgi?id=59124
#endif

inline void GrisuRound(char* buffer, int len, uint64_t delta, uint64_t rest, uint64_t ten_kappa, uint64_t wp_w) {
    while (rest < wp_w && delta - rest >= ten_kappa &&
           (rest + ten_kappa < wp_w ||  /// closer
            wp_w - rest > rest + ten_kappa - wp_w)) {
        buffer[len - 1]--;
        rest += ten_kappa;
    }
}

inline int CountDecimalDigit32(uint32_t n) {
    // Simple pure C++ implementation was faster than __builtin_clz version in this situation.
    if (n < 10) return 1;
    if (n < 100) return 2;
    if (n < 1000) return 3;
    if (n < 10000) return 4;
    if (n < 100000) return 5;
    if (n < 1000000) return 6;
    if (n < 10000000) return 7;
    if (n < 100000000) return 8;
    // Will not reach 10 digits in DigitGen()
    //if (n < 1000000000) return 9;
    //return 10;
    return 9;
}

inline void DigitGen(const DiyFp& W, const DiyFp& Mp, uint64_t delta, char* buffer, int* len, int* K) {
    static const uint32_t kPow10[] = { 1, 10, 100, 1000, 10000, 100000, 1000000, 10000000, 100000000, 1000000000 };
    const DiyFp one(uint64_t(1) << -Mp.e, Mp.e);
    const DiyFp wp_w = Mp - W;
    uint32_t p1 = static_cast<uint32_t>(Mp.f >> -one.e);
    uint64_t p2 = Mp.f & (one.f - 1);
    int kappa = CountDecimalDigit32(p1); // kappa in [0, 9]
    *len = 0;

    while (kappa > 0) {
        uint32_t d = 0;
        switch (kappa) {
            case  9: d = p1 /  100000000; p1 %=  100000000; break;
            case  8: d = p1 /   10000000; p1 %=   10000000; break;
            case  7: d = p1 /    1000000; p1 %=    1000000; break;
            case  6: d = p1 /     100000; p1 %=     100000; break;
            case  5: d = p1 /      10000; p1 %=      10000; break;
            case  4: d = p1 /       1000; p1 %=       1000; break;
            case  3: d = p1 /        100; p1 %=        100; break;
            case  2: d = p1 /         10; p1 %=         10; break;
            case  1: d = p1;              p1 =           0; break;
            default:;
        }
        if (d || *len)
            buffer[(*len)++] = static_cast<char>('0' + static_cast<char>(d));
        kappa--;
        uint64_t tmp = (static_cast<uint64_t>(p1) << -one.e) + p2;
        if (tmp <= delta) {
            *K += kappa;
            GrisuRound(buffer, *len, delta, tmp, static_cast<uint64_t>(kPow10[kappa]) << -one.e, wp_w.f);
            return;
        }
    }

    // kappa = 0
    for (;;) {
        p2 *= 10;
        delta *= 10;
        char d = static_cast<char>(p2 >> -one.e);
        if (d || *len)
            buffer[(*len)++] = static_cast<char>('0' + d);
        p2 &= one.f - 1;
        kappa--;
        if (p2 < delta) {
            *K += kappa;
            int index = -kappa;
            GrisuRound(buffer, *len, delta, p2, one.f, wp_w.f * (index < 9 ? kPow10[index] : 0));
            return;
        }
    }
}

inline void Grisu2(double value, char* buffer, int* length, int* K) {
    const DiyFp v(value);
    DiyFp w_m, w_p;
    v.NormalizedBoundaries(&w_m, &w_p);

    const DiyFp c_mk = GetCachedPower(w_p.e, K);
    const DiyFp W = v.Normalize() * c_mk;
    DiyFp Wp = w_p * c_mk;
    DiyFp Wm = w_m * c_mk;
    Wm.f++;
    Wp.f--;
    DigitGen(W, Wp, Wp.f - Wm.f, buffer, length, K);
}

inline char* WriteExponent(int K, char* buffer) {
    if (K < 0) {
        *buffer++ = '-';
        K = -K;
    }

    if (K >= 100) {
        *buffer++ = static_cast<char>('0' + static_cast<char>(K / 100));
        K %= 100;
        const char* d = GetDigitsLut() + K * 2;
        *buffer++ = d[0];
        *buffer++ = d[1];
    }
    else if (K >= 10) {
        const char* d = GetDigitsLut() + K * 2;
        *buffer++ = d[0];
        *buffer++ = d[1];
    }
    else
        *buffer++ = static_cast<char>('0' + static_cast<char>(K));

    return buffer;
}

inline char* Prettify(char* buffer, int length, int k, int maxDecimalPlaces) {
    const int kk = length + k;  // 10^(kk-1) <= v < 10^kk

    if (0 <= k && kk <= 21) {
        // 1234e7 -> 12340000000
        for (int i = length; i < kk; i++)
            buffer[i] = '0';
        buffer[kk] = '.';
        buffer[kk + 1] = '0';
        return &buffer[kk + 2];
    }
    else if (0 < kk && kk <= 21) {
        // 1234e-2 -> 12.34
        std::memmove(&buffer[kk + 1], &buffer[kk], static_cast<size_t>(length - kk));
        buffer[kk] = '.';
        if (0 > k + maxDecimalPlaces) {
            // When maxDecimalPlaces = 2, 1.2345 -> 1.23, 1.102 -> 1.1
            // Remove extra trailing zeros (at least one) after truncation.
            for (int i = kk + maxDecimalPlaces; i > kk + 1; i--)
                if (buffer[i] != '0')
                    return &buffer[i + 1];
            return &buffer[kk + 2]; // Reserve one zero
        }
        else
            return &buffer[length + 1];
    }
    else if (-6 < kk && kk <= 0) {
        // 1234e-6 -> 0.001234
        const int offset = 2 - kk;
        std::memmove(&buffer[offset], &buffer[0], static_cast<size_t>(length));
        buffer[0] = '0';
        buffer[1] = '.';
        for (int i = 2; i < offset; i++)
            buffer[i] = '0';
        if (length - kk > maxDecimalPlaces) {
            // When maxDecimalPlaces = 2, 0.123 -> 0.12, 0.102 -> 0.1
            // Remove extra trailing zeros (at least one) after truncation.
            for (int i = maxDecimalPlaces + 1; i > 2; i--)
                if (buffer[i] != '0')
                    return &buffer[i + 1];
            return &buffer[3]; // Reserve one zero
        }
        else
            return &buffer[length + offset];
    }
    else if (kk < -maxDecimalPlaces) {
        // Truncate to zero
        buffer[0] = '0';
        buffer[1] = '.';
        buffer[2] = '0';
        return &buffer[3];
    }
    else if (length == 1) {
        // 1e30
        buffer[1] = 'e';
        return WriteExponent(kk - 1, &buffer[2]);
    }
    else {
        // 1234e30 -> 1.234e33
        std::memmove(&buffer[2], &buffer[1], static_cast<size_t>(length - 1));
        buffer[1] = '.';
        buffer[length + 1] = 'e';
        return WriteExponent(kk - 1, &buffer[0 + length + 2]);
    }
}

inline char* dtoa(double value, char* buffer, int maxDecimalPlaces = 324) {
    CEREAL_RAPIDJSON_ASSERT(maxDecimalPlaces >= 1);
    Double d(value);
    if (d.IsZero()) {
        if (d.Sign())
            *buffer++ = '-';     // -0.0, Issue #289
        buffer[0] = '0';
        buffer[1] = '.';
        buffer[2] = '0';
        return &buffer[3];
    }
    else {
        if (value < 0) {
            *buffer++ = '-';
            value = -value;
        }
        int length, K;
        Grisu2(value, buffer, &length, &K);
        return Prettify(buffer, length, K, maxDecimalPlaces);
    }
}

#ifdef __GNUC__
CEREAL_RAPIDJSON_DIAG_POP
#endif

} // namespace internal
CEREAL_RAPIDJSON_NAMESPACE_END

#endif // CEREAL_RAPIDJSON_DTOA_
```
