## Supplemental code for "The motif composition of variable-number tandem repeats impacts gene expression": ieee754.h

```
// Tencent is pleased to support the open source community by making RapidJSON available.
// 
// Copyright (C) 2015 THL A29 Limited, a Tencent company, and Milo Yip. All rights reserved.
//
// Licensed under the MIT License (the "License"); you may not use this file except
// in compliance with the License. You may obtain a copy of the License at
//
// http://opensource.org/licenses/MIT
//
// Unless required by applicable law or agreed to in writing, software distributed 
// under the License is distributed on an "AS IS" BASIS, WITHOUT WARRANTIES OR 
// CONDITIONS OF ANY KIND, either express or implied. See the License for the 
// specific language governing permissions and limitations under the License.

#ifndef CEREAL_RAPIDJSON_IEEE754_
#define CEREAL_RAPIDJSON_IEEE754_

#include "../rapidjson.h"

CEREAL_RAPIDJSON_NAMESPACE_BEGIN
namespace internal {

class Double {
public:
    Double() {}
    Double(double d) : d_(d) {}
    Double(uint64_t u) : u_(u) {}

    double Value() const { return d_; }
    uint64_t Uint64Value() const { return u_; }

    double NextPositiveDouble() const {
        CEREAL_RAPIDJSON_ASSERT(!Sign());
        return Double(u_ + 1).Value();
    }

    bool Sign() const { return (u_ & kSignMask) != 0; }
    uint64_t Significand() const { return u_ & kSignificandMask; }
    int Exponent() const { return static_cast<int>(((u_ & kExponentMask) >> kSignificandSize) - kExponentBias); }

    bool IsNan() const { return (u_ & kExponentMask) == kExponentMask && Significand() != 0; }
    bool IsInf() const { return (u_ & kExponentMask) == kExponentMask && Significand() == 0; }
    bool IsNanOrInf() const { return (u_ & kExponentMask) == kExponentMask; }
    bool IsNormal() const { return (u_ & kExponentMask) != 0 || Significand() == 0; }
    bool IsZero() const { return (u_ & (kExponentMask | kSignificandMask)) == 0; }

    uint64_t IntegerSignificand() const { return IsNormal() ? Significand() | kHiddenBit : Significand(); }
    int IntegerExponent() const { return (IsNormal() ? Exponent() : kDenormalExponent) - kSignificandSize; }
    uint64_t ToBias() const { return (u_ & kSignMask) ? ~u_ + 1 : u_ | kSignMask; }

    static int EffectiveSignificandSize(int order) {
        if (order >= -1021)
            return 53;
        else if (order <= -1074)
            return 0;
        else
            return order + 1074;
    }

private:
    static const int kSignificandSize = 52;
    static const int kExponentBias = 0x3FF;
    static const int kDenormalExponent = 1 - kExponentBias;
    static const uint64_t kSignMask = CEREAL_RAPIDJSON_UINT64_C2(0x80000000, 0x00000000);
    static const uint64_t kExponentMask = CEREAL_RAPIDJSON_UINT64_C2(0x7FF00000, 0x00000000);
    static const uint64_t kSignificandMask = CEREAL_RAPIDJSON_UINT64_C2(0x000FFFFF, 0xFFFFFFFF);
    static const uint64_t kHiddenBit = CEREAL_RAPIDJSON_UINT64_C2(0x00100000, 0x00000000);

    union {
        double d_;
        uint64_t u_;
    };
};

} // namespace internal
CEREAL_RAPIDJSON_NAMESPACE_END

#endif // CEREAL_RAPIDJSON_IEEE754_
```
