## Supplemental code for "The motif composition of variable-number tandem repeats impacts gene expression": itoa.h

```
// Tencent is pleased to support the open source community by making RapidJSON available.
//
// Copyright (C) 2015 THL A29 Limited, a Tencent company, and Milo Yip. All rights reserved.
//
// Licensed under the MIT License (the "License"); you may not use this file except
// in compliance with the License. You may obtain a copy of the License at
//
// http://opensource.org/licenses/MIT
//
// Unless required by applicable law or agreed to in writing, software distributed
// under the License is distributed on an "AS IS" BASIS, WITHOUT WARRANTIES OR
// CONDITIONS OF ANY KIND, either express or implied. See the License for the
// specific language governing permissions and limitations under the License.

#ifndef CEREAL_RAPIDJSON_ITOA_
#define CEREAL_RAPIDJSON_ITOA_

#include "../rapidjson.h"

CEREAL_RAPIDJSON_NAMESPACE_BEGIN
namespace internal {

inline const char* GetDigitsLut() {
    static const char cDigitsLut[200] = {
        '0','0','0','1','0','2','0','3','0','4','0','5','0','6','0','7','0','8','0','9',
        '1','0','1','1','1','2','1','3','1','4','1','5','1','6','1','7','1','8','1','9',
        '2','0','2','1','2','2','2','3','2','4','2','5','2','6','2','7','2','8','2','9',
        '3','0','3','1','3','2','3','3','3','4','3','5','3','6','3','7','3','8','3','9',
        '4','0','4','1','4','2','4','3','4','4','4','5','4','6','4','7','4','8','4','9',
        '5','0','5','1','5','2','5','3','5','4','5','5','5','6','5','7','5','8','5','9',
        '6','0','6','1','6','2','6','3','6','4','6','5','6','6','6','7','6','8','6','9',
        '7','0','7','1','7','2','7','3','7','4','7','5','7','6','7','7','7','8','7','9',
        '8','0','8','1','8','2','8','3','8','4','8','5','8','6','8','7','8','8','8','9',
        '9','0','9','1','9','2','9','3','9','4','9','5','9','6','9','7','9','8','9','9'
    };
    return cDigitsLut;
}

inline char* u32toa(uint32_t value, char* buffer) {
    CEREAL_RAPIDJSON_ASSERT(buffer != 0);

    const char* cDigitsLut = GetDigitsLut();

    if (value < 10000) {
        const uint32_t d1 = (value / 100) << 1;
        const uint32_t d2 = (value % 100) << 1;

        if (value >= 1000)
            *buffer++ = cDigitsLut[d1];
        if (value >= 100)
            *buffer++ = cDigitsLut[d1 + 1];
        if (value >= 10)
            *buffer++ = cDigitsLut[d2];
        *buffer++ = cDigitsLut[d2 + 1];
    }
    else if (value < 100000000) {
        // value = bbbbcccc
        const uint32_t b = value / 10000;
        const uint32_t c = value % 10000;

        const uint32_t d1 = (b / 100) << 1;
        const uint32_t d2 = (b % 100) << 1;

        const uint32_t d3 = (c / 100) << 1;
        const uint32_t d4 = (c % 100) << 1;

        if (value >= 10000000)
            *buffer++ = cDigitsLut[d1];
        if (value >= 1000000)
            *buffer++ = cDigitsLut[d1 + 1];
        if (value >= 100000)
            *buffer++ = cDigitsLut[d2];
        *buffer++ = cDigitsLut[d2 + 1];

        *buffer++ = cDigitsLut[d3];
        *buffer++ = cDigitsLut[d3 + 1];
        *buffer++ = cDigitsLut[d4];
        *buffer++ = cDigitsLut[d4 + 1];
    }
    else {
        // value = aabbbbcccc in decimal

        const uint32_t a = value / 100000000; // 1 to 42
        value %= 100000000;

        if (a >= 10) {
            const unsigned i = a << 1;
            *buffer++ = cDigitsLut[i];
            *buffer++ = cDigitsLut[i + 1];
        }
        else
            *buffer++ = static_cast<char>('0' + static_cast<char>(a));

        const uint32_t b = value / 10000; // 0 to 9999
        const uint32_t c = value % 10000; // 0 to 9999

        const uint32_t d1 = (b / 100) << 1;
        const uint32_t d2 = (b % 100) << 1;

        const uint32_t d3 = (c / 100) << 1;
        const uint32_t d4 = (c % 100) << 1;

        *buffer++ = cDigitsLut[d1];
        *buffer++ = cDigitsLut[d1 + 1];
        *buffer++ = cDigitsLut[d2];
        *buffer++ = cDigitsLut[d2 + 1];
        *buffer++ = cDigitsLut[d3];
        *buffer++ = cDigitsLut[d3 + 1];
        *buffer++ = cDigitsLut[d4];
        *buffer++ = cDigitsLut[d4 + 1];
    }
    return buffer;
}

inline char* i32toa(int32_t value, char* buffer) {
    CEREAL_RAPIDJSON_ASSERT(buffer != 0);
    uint32_t u = static_cast<uint32_t>(value);
    if (value < 0) {
        *buffer++ = '-';
        u = ~u + 1;
    }

    return u32toa(u, buffer);
}

inline char* u64toa(uint64_t value, char* buffer) {
    CEREAL_RAPIDJSON_ASSERT(buffer != 0);
    const char* cDigitsLut = GetDigitsLut();
    const uint64_t  kTen8 = 100000000;
    const uint64_t  kTen9 = kTen8 * 10;
    const uint64_t kTen10 = kTen8 * 100;
    const uint64_t kTen11 = kTen8 * 1000;
    const uint64_t kTen12 = kTen8 * 10000;
    const uint64_t kTen13 = kTen8 * 100000;
    const uint64_t kTen14 = kTen8 * 1000000;
    const uint64_t kTen15 = kTen8 * 10000000;
    const uint64_t kTen16 = kTen8 * kTen8;

    if (value < kTen8) {
        uint32_t v = static_cast<uint32_t>(value);
        if (v < 10000) {
            const uint32_t d1 = (v / 100) << 1;
            const uint32_t d2 = (v % 100) << 1;

            if (v >= 1000)
                *buffer++ = cDigitsLut[d1];
            if (v >= 100)
                *buffer++ = cDigitsLut[d1 + 1];
            if (v >= 10)
                *buffer++ = cDigitsLut[d2];
            *buffer++ = cDigitsLut[d2 + 1];
        }
        else {
            // value = bbbbcccc
            const uint32_t b = v / 10000;
            const uint32_t c = v % 10000;

            const uint32_t d1 = (b / 100) << 1;
            const uint32_t d2 = (b % 100) << 1;

            const uint32_t d3 = (c / 100) << 1;
            const uint32_t d4 = (c % 100) << 1;

            if (value >= 10000000)
                *buffer++ = cDigitsLut[d1];
            if (value >= 1000000)
                *buffer++ = cDigitsLut[d1 + 1];
            if (value >= 100000)
                *buffer++ = cDigitsLut[d2];
            *buffer++ = cDigitsLut[d2 + 1];

            *buffer++ = cDigitsLut[d3];
            *buffer++ = cDigitsLut[d3 + 1];
            *buffer++ = cDigitsLut[d4];
            *buffer++ = cDigitsLut[d4 + 1];
        }
    }
    else if (value < kTen16) {
        const uint32_t v0 = static_cast<uint32_t>(value / kTen8);
        const uint32_t v1 = static_cast<uint32_t>(value % kTen8);

        const uint32_t b0 = v0 / 10000;
        const uint32_t c0 = v0 % 10000;

        const uint32_t d1 = (b0 / 100) << 1;
        const uint32_t d2 = (b0 % 100) << 1;

        const uint32_t d3 = (c0 / 100) << 1;
        const uint32_t d4 = (c0 % 100) << 1;

        const uint32_t b1 = v1 / 10000;
        const uint32_t c1 = v1 % 10000;

        const uint32_t d5 = (b1 / 100) << 1;
        const uint32_t d6 = (b1 % 100) << 1;

        const uint32_t d7 = (c1 / 100) << 1;
        const uint32_t d8 = (c1 % 100) << 1;

        if (value >= kTen15)
            *buffer++ = cDigitsLut[d1];
        if (value >= kTen14)
            *buffer++ = cDigitsLut[d1 + 1];
        if (value >= kTen13)
            *buffer++ = cDigitsLut[d2];
        if (value >= kTen12)
            *buffer++ = cDigitsLut[d2 + 1];
        if (value >= kTen11)
            *buffer++ = cDigitsLut[d3];
        if (value >= kTen10)
            *buffer++ = cDigitsLut[d3 + 1];
        if (value >= kTen9)
            *buffer++ = cDigitsLut[d4];

        *buffer++ = cDigitsLut[d4 + 1];
        *buffer++ = cDigitsLut[d5];
        *buffer++ = cDigitsLut[d5 + 1];
        *buffer++ = cDigitsLut[d6];
        *buffer++ = cDigitsLut[d6 + 1];
        *buffer++ = cDigitsLut[d7];
        *buffer++ = cDigitsLut[d7 + 1];
        *buffer++ = cDigitsLut[d8];
        *buffer++ = cDigitsLut[d8 + 1];
    }
    else {
        const uint32_t a = static_cast<uint32_t>(value / kTen16); // 1 to 1844
        value %= kTen16;

        if (a < 10)
            *buffer++ = static_cast<char>('0' + static_cast<char>(a));
        else if (a < 100) {
            const uint32_t i = a << 1;
            *buffer++ = cDigitsLut[i];
            *buffer++ = cDigitsLut[i + 1];
        }
        else if (a < 1000) {
            *buffer++ = static_cast<char>('0' + static_cast<char>(a / 100));

            const uint32_t i = (a % 100) << 1;
            *buffer++ = cDigitsLut[i];
            *buffer++ = cDigitsLut[i + 1];
        }
        else {
            const uint32_t i = (a / 100) << 1;
            const uint32_t j = (a % 100) << 1;
            *buffer++ = cDigitsLut[i];
            *buffer++ = cDigitsLut[i + 1];
            *buffer++ = cDigitsLut[j];
            *buffer++ = cDigitsLut[j + 1];
        }

        const uint32_t v0 = static_cast<uint32_t>(value / kTen8);
        const uint32_t v1 = static_cast<uint32_t>(value % kTen8);

        const uint32_t b0 = v0 / 10000;
        const uint32_t c0 = v0 % 10000;

        const uint32_t d1 = (b0 / 100) << 1;
        const uint32_t d2 = (b0 % 100) << 1;

        const uint32_t d3 = (c0 / 100) << 1;
        const uint32_t d4 = (c0 % 100) << 1;

        const uint32_t b1 = v1 / 10000;
        const uint32_t c1 = v1 % 10000;

        const uint32_t d5 = (b1 / 100) << 1;
        const uint32_t d6 = (b1 % 100) << 1;

        const uint32_t d7 = (c1 / 100) << 1;
        const uint32_t d8 = (c1 % 100) << 1;

        *buffer++ = cDigitsLut[d1];
        *buffer++ = cDigitsLut[d1 + 1];
        *buffer++ = cDigitsLut[d2];
        *buffer++ = cDigitsLut[d2 + 1];
        *buffer++ = cDigitsLut[d3];
        *buffer++ = cDigitsLut[d3 + 1];
        *buffer++ = cDigitsLut[d4];
        *buffer++ = cDigitsLut[d4 + 1];
        *buffer++ = cDigitsLut[d5];
        *buffer++ = cDigitsLut[d5 + 1];
        *buffer++ = cDigitsLut[d6];
        *buffer++ = cDigitsLut[d6 + 1];
        *buffer++ = cDigitsLut[d7];
        *buffer++ = cDigitsLut[d7 + 1];
        *buffer++ = cDigitsLut[d8];
        *buffer++ = cDigitsLut[d8 + 1];
    }

    return buffer;
}

inline char* i64toa(int64_t value, char* buffer) {
    CEREAL_RAPIDJSON_ASSERT(buffer != 0);
    uint64_t u = static_cast<uint64_t>(value);
    if (value < 0) {
        *buffer++ = '-';
        u = ~u + 1;
    }

    return u64toa(u, buffer);
}

} // namespace internal
CEREAL_RAPIDJSON_NAMESPACE_END

#endif // CEREAL_RAPIDJSON_ITOA_
```
