## Supplemental code for "The motif composition of variable-number tandem repeats impacts gene expression": meta.h

```
// Tencent is pleased to support the open source community by making RapidJSON available.
// 
// Copyright (C) 2015 THL A29 Limited, a Tencent company, and Milo Yip. All rights reserved.
//
// Licensed under the MIT License (the "License"); you may not use this file except
// in compliance with the License. You may obtain a copy of the License at
//
// http://opensource.org/licenses/MIT
//
// Unless required by applicable law or agreed to in writing, software distributed 
// under the License is distributed on an "AS IS" BASIS, WITHOUT WARRANTIES OR 
// CONDITIONS OF ANY KIND, either express or implied. See the License for the 
// specific language governing permissions and limitations under the License.

#ifndef CEREAL_RAPIDJSON_INTERNAL_META_H_
#define CEREAL_RAPIDJSON_INTERNAL_META_H_

#include "../rapidjson.h"

#ifdef __GNUC__
CEREAL_RAPIDJSON_DIAG_PUSH
CEREAL_RAPIDJSON_DIAG_OFF(effc++)
#endif

#if defined(_MSC_VER) && !defined(__clang__)
CEREAL_RAPIDJSON_DIAG_PUSH
CEREAL_RAPIDJSON_DIAG_OFF(6334)
#endif

#if CEREAL_RAPIDJSON_HAS_CXX11_TYPETRAITS
#include <type_traits>
#endif

//@cond CEREAL_RAPIDJSON_INTERNAL
CEREAL_RAPIDJSON_NAMESPACE_BEGIN
namespace internal {

// Helper to wrap/convert arbitrary types to void, useful for arbitrary type matching
template <typename T> struct Void { typedef void Type; };

///////////////////////////////////////////////////////////////////////////////
// BoolType, TrueType, FalseType
//
template <bool Cond> struct BoolType {
    static const bool Value = Cond;
    typedef BoolType Type;
};
typedef BoolType<true> TrueType;
typedef BoolType<false> FalseType;


///////////////////////////////////////////////////////////////////////////////
// SelectIf, BoolExpr, NotExpr, AndExpr, OrExpr
//

template <bool C> struct SelectIfImpl { template <typename T1, typename T2> struct Apply { typedef T1 Type; }; };
template <> struct SelectIfImpl<false> { template <typename T1, typename T2> struct Apply { typedef T2 Type; }; };
template <bool C, typename T1, typename T2> struct SelectIfCond : SelectIfImpl<C>::template Apply<T1,T2> {};
template <typename C, typename T1, typename T2> struct SelectIf : SelectIfCond<C::Value, T1, T2> {};

template <bool Cond1, bool Cond2> struct AndExprCond : FalseType {};
template <> struct AndExprCond<true, true> : TrueType {};
template <bool Cond1, bool Cond2> struct OrExprCond : TrueType {};
template <> struct OrExprCond<false, false> : FalseType {};

template <typename C> struct BoolExpr : SelectIf<C,TrueType,FalseType>::Type {};
template <typename C> struct NotExpr  : SelectIf<C,FalseType,TrueType>::Type {};
template <typename C1, typename C2> struct AndExpr : AndExprCond<C1::Value, C2::Value>::Type {};
template <typename C1, typename C2> struct OrExpr  : OrExprCond<C1::Value, C2::Value>::Type {};


///////////////////////////////////////////////////////////////////////////////
// AddConst, MaybeAddConst, RemoveConst
template <typename T> struct AddConst { typedef const T Type; };
template <bool Constify, typename T> struct MaybeAddConst : SelectIfCond<Constify, const T, T> {};
template <typename T> struct RemoveConst { typedef T Type; };
template <typename T> struct RemoveConst<const T> { typedef T Type; };


///////////////////////////////////////////////////////////////////////////////
// IsSame, IsConst, IsMoreConst, IsPointer
//
template <typename T, typename U> struct IsSame : FalseType {};
template <typename T> struct IsSame<T, T> : TrueType {};

template <typename T> struct IsConst : FalseType {};
template <typename T> struct IsConst<const T> : TrueType {};

template <typename CT, typename T>
struct IsMoreConst
    : AndExpr<IsSame<typename RemoveConst<CT>::Type, typename RemoveConst<T>::Type>,
              BoolType<IsConst<CT>::Value >= IsConst<T>::Value> >::Type {};

template <typename T> struct IsPointer : FalseType {};
template <typename T> struct IsPointer<T*> : TrueType {};

///////////////////////////////////////////////////////////////////////////////
// IsBaseOf
//
#if CEREAL_RAPIDJSON_HAS_CXX11_TYPETRAITS

template <typename B, typename D> struct IsBaseOf
    : BoolType< ::std::is_base_of<B,D>::value> {};

#else // simplified version adopted from Boost

template<typename B, typename D> struct IsBaseOfImpl {
    CEREAL_RAPIDJSON_STATIC_ASSERT(sizeof(B) != 0);
    CEREAL_RAPIDJSON_STATIC_ASSERT(sizeof(D) != 0);

    typedef char (&Yes)[1];
    typedef char (&No) [2];

    template <typename T>
    static Yes Check(const D*, T);
    static No  Check(const B*, int);

    struct Host {
        operator const B*() const;
        operator const D*();
    };

    enum { Value = (sizeof(Check(Host(), 0)) == sizeof(Yes)) };
};

template <typename B, typename D> struct IsBaseOf
    : OrExpr<IsSame<B, D>, BoolExpr<IsBaseOfImpl<B, D> > >::Type {};

#endif // CEREAL_RAPIDJSON_HAS_CXX11_TYPETRAITS


//////////////////////////////////////////////////////////////////////////
// EnableIf / DisableIf
//
template <bool Condition, typename T = void> struct EnableIfCond  { typedef T Type; };
template <typename T> struct EnableIfCond<false, T> { /* empty */ };

template <bool Condition, typename T = void> struct DisableIfCond { typedef T Type; };
template <typename T> struct DisableIfCond<true, T> { /* empty */ };

template <typename Condition, typename T = void>
struct EnableIf : EnableIfCond<Condition::Value, T> {};

template <typename Condition, typename T = void>
struct DisableIf : DisableIfCond<Condition::Value, T> {};

// SFINAE helpers
struct SfinaeTag {};
template <typename T> struct RemoveSfinaeTag;
template <typename T> struct RemoveSfinaeTag<SfinaeTag&(*)(T)> { typedef T Type; };

#define CEREAL_RAPIDJSON_REMOVEFPTR_(type) \
    typename ::CEREAL_RAPIDJSON_NAMESPACE::internal::RemoveSfinaeTag \
        < ::CEREAL_RAPIDJSON_NAMESPACE::internal::SfinaeTag&(*) type>::Type

#define CEREAL_RAPIDJSON_ENABLEIF(cond) \
    typename ::CEREAL_RAPIDJSON_NAMESPACE::internal::EnableIf \
        <CEREAL_RAPIDJSON_REMOVEFPTR_(cond)>::Type * = NULL

#define CEREAL_RAPIDJSON_DISABLEIF(cond) \
    typename ::CEREAL_RAPIDJSON_NAMESPACE::internal::DisableIf \
        <CEREAL_RAPIDJSON_REMOVEFPTR_(cond)>::Type * = NULL

#define CEREAL_RAPIDJSON_ENABLEIF_RETURN(cond,returntype) \
    typename ::CEREAL_RAPIDJSON_NAMESPACE::internal::EnableIf \
        <CEREAL_RAPIDJSON_REMOVEFPTR_(cond), \
         CEREAL_RAPIDJSON_REMOVEFPTR_(returntype)>::Type

#define CEREAL_RAPIDJSON_DISABLEIF_RETURN(cond,returntype) \
    typename ::CEREAL_RAPIDJSON_NAMESPACE::internal::DisableIf \
        <CEREAL_RAPIDJSON_REMOVEFPTR_(cond), \
         CEREAL_RAPIDJSON_REMOVEFPTR_(returntype)>::Type

} // namespace internal
CEREAL_RAPIDJSON_NAMESPACE_END
//@endcond

#if defined(_MSC_VER) && !defined(__clang__)
CEREAL_RAPIDJSON_DIAG_POP
#endif

#ifdef __GNUC__
CEREAL_RAPIDJSON_DIAG_POP
#endif

#endif // CEREAL_RAPIDJSON_INTERNAL_META_H_
```
