## Supplemental code for "The motif composition of variable-number tandem repeats impacts gene expression": pow10.h

```
// Tencent is pleased to support the open source community by making RapidJSON available.
// 
// Copyright (C) 2015 THL A29 Limited, a Tencent company, and Milo Yip. All rights reserved.
//
// Licensed under the MIT License (the "License"); you may not use this file except
// in compliance with the License. You may obtain a copy of the License at
//
// http://opensource.org/licenses/MIT
//
// Unless required by applicable law or agreed to in writing, software distributed 
// under the License is distributed on an "AS IS" BASIS, WITHOUT WARRANTIES OR 
// CONDITIONS OF ANY KIND, either express or implied. See the License for the 
// specific language governing permissions and limitations under the License.

#ifndef CEREAL_RAPIDJSON_POW10_
#define CEREAL_RAPIDJSON_POW10_

#include "../rapidjson.h"

CEREAL_RAPIDJSON_NAMESPACE_BEGIN
namespace internal {

//! Computes integer powers of 10 in double (10.0^n).
/*! This function uses lookup table for fast and accurate results.
    \param n non-negative exponent. Must <= 308.
    \return 10.0^n
*/
inline double Pow10(int n) {
    static const double e[] = { // 1e-0...1e308: 309 * 8 bytes = 2472 bytes
        1e+0,  
        1e+1,  1e+2,  1e+3,  1e+4,  1e+5,  1e+6,  1e+7,  1e+8,  1e+9,  1e+10, 1e+11, 1e+12, 1e+13, 1e+14, 1e+15, 1e+16, 1e+17, 1e+18, 1e+19, 1e+20, 
        1e+21, 1e+22, 1e+23, 1e+24, 1e+25, 1e+26, 1e+27, 1e+28, 1e+29, 1e+30, 1e+31, 1e+32, 1e+33, 1e+34, 1e+35, 1e+36, 1e+37, 1e+38, 1e+39, 1e+40,
        1e+41, 1e+42, 1e+43, 1e+44, 1e+45, 1e+46, 1e+47, 1e+48, 1e+49, 1e+50, 1e+51, 1e+52, 1e+53, 1e+54, 1e+55, 1e+56, 1e+57, 1e+58, 1e+59, 1e+60,
        1e+61, 1e+62, 1e+63, 1e+64, 1e+65, 1e+66, 1e+67, 1e+68, 1e+69, 1e+70, 1e+71, 1e+72, 1e+73, 1e+74, 1e+75, 1e+76, 1e+77, 1e+78, 1e+79, 1e+80,
        1e+81, 1e+82, 1e+83, 1e+84, 1e+85, 1e+86, 1e+87, 1e+88, 1e+89, 1e+90, 1e+91, 1e+92, 1e+93, 1e+94, 1e+95, 1e+96, 1e+97, 1e+98, 1e+99, 1e+100,
        1e+101,1e+102,1e+103,1e+104,1e+105,1e+106,1e+107,1e+108,1e+109,1e+110,1e+111,1e+112,1e+113,1e+114,1e+115,1e+116,1e+117,1e+118,1e+119,1e+120,
        1e+121,1e+122,1e+123,1e+124,1e+125,1e+126,1e+127,1e+128,1e+129,1e+130,1e+131,1e+132,1e+133,1e+134,1e+135,1e+136,1e+137,1e+138,1e+139,1e+140,
        1e+141,1e+142,1e+143,1e+144,1e+145,1e+146,1e+147,1e+148,1e+149,1e+150,1e+151,1e+152,1e+153,1e+154,1e+155,1e+156,1e+157,1e+158,1e+159,1e+160,
        1e+161,1e+162,1e+163,1e+164,1e+165,1e+166,1e+167,1e+168,1e+169,1e+170,1e+171,1e+172,1e+173,1e+174,1e+175,1e+176,1e+177,1e+178,1e+179,1e+180,
        1e+181,1e+182,1e+183,1e+184,1e+185,1e+186,1e+187,1e+188,1e+189,1e+190,1e+191,1e+192,1e+193,1e+194,1e+195,1e+196,1e+197,1e+198,1e+199,1e+200,
        1e+201,1e+202,1e+203,1e+204,1e+205,1e+206,1e+207,1e+208,1e+209,1e+210,1e+211,1e+212,1e+213,1e+214,1e+215,1e+216,1e+217,1e+218,1e+219,1e+220,
        1e+221,1e+222,1e+223,1e+224,1e+225,1e+226,1e+227,1e+228,1e+229,1e+230,1e+231,1e+232,1e+233,1e+234,1e+235,1e+236,1e+237,1e+238,1e+239,1e+240,
        1e+241,1e+242,1e+243,1e+244,1e+245,1e+246,1e+247,1e+248,1e+249,1e+250,1e+251,1e+252,1e+253,1e+254,1e+255,1e+256,1e+257,1e+258,1e+259,1e+260,
        1e+261,1e+262,1e+263,1e+264,1e+265,1e+266,1e+267,1e+268,1e+269,1e+270,1e+271,1e+272,1e+273,1e+274,1e+275,1e+276,1e+277,1e+278,1e+279,1e+280,
        1e+281,1e+282,1e+283,1e+284,1e+285,1e+286,1e+287,1e+288,1e+289,1e+290,1e+291,1e+292,1e+293,1e+294,1e+295,1e+296,1e+297,1e+298,1e+299,1e+300,
        1e+301,1e+302,1e+303,1e+304,1e+305,1e+306,1e+307,1e+308
    };
    CEREAL_RAPIDJSON_ASSERT(n >= 0 && n <= 308);
    return e[n];
}

} // namespace internal
CEREAL_RAPIDJSON_NAMESPACE_END

#endif // CEREAL_RAPIDJSON_POW10_
```
