## Supplemental code for "The motif composition of variable-number tandem repeats impacts gene expression": regex.h

```
// Tencent is pleased to support the open source community by making RapidJSON available.
// 
// Copyright (C) 2015 THL A29 Limited, a Tencent company, and Milo Yip. All rights reserved.
//
// Licensed under the MIT License (the "License"); you may not use this file except
// in compliance with the License. You may obtain a copy of the License at
//
// http://opensource.org/licenses/MIT
//
// Unless required by applicable law or agreed to in writing, software distributed 
// under the License is distributed on an "AS IS" BASIS, WITHOUT WARRANTIES OR 
// CONDITIONS OF ANY KIND, either express or implied. See the License for the 
// specific language governing permissions and limitations under the License.

#ifndef CEREAL_RAPIDJSON_INTERNAL_REGEX_H_
#define CEREAL_RAPIDJSON_INTERNAL_REGEX_H_

#include "../allocators.h"
#include "../stream.h"
#include "stack.h"

#ifdef __clang__
CEREAL_RAPIDJSON_DIAG_PUSH
CEREAL_RAPIDJSON_DIAG_OFF(padded)
CEREAL_RAPIDJSON_DIAG_OFF(switch-enum)
CEREAL_RAPIDJSON_DIAG_OFF(implicit-fallthrough)
#elif defined(_MSC_VER)
CEREAL_RAPIDJSON_DIAG_PUSH
CEREAL_RAPIDJSON_DIAG_OFF(4512) // assignment operator could not be generated
#endif

#ifdef __GNUC__
CEREAL_RAPIDJSON_DIAG_PUSH
CEREAL_RAPIDJSON_DIAG_OFF(effc++)
#if __GNUC__ >= 7
CEREAL_RAPIDJSON_DIAG_OFF(implicit-fallthrough)
#endif
#endif

#ifndef CEREAL_RAPIDJSON_REGEX_VERBOSE
#define CEREAL_RAPIDJSON_REGEX_VERBOSE 0
#endif

CEREAL_RAPIDJSON_NAMESPACE_BEGIN
namespace internal {

///////////////////////////////////////////////////////////////////////////////
// DecodedStream

template <typename SourceStream, typename Encoding>
class DecodedStream {
public:
    DecodedStream(SourceStream& ss) : ss_(ss), codepoint_() { Decode(); }
    unsigned Peek() { return codepoint_; }
    unsigned Take() {
        unsigned c = codepoint_;
        if (c) // No further decoding when '\0'
            Decode();
        return c;
    }

private:
    void Decode() {
        if (!Encoding::Decode(ss_, &codepoint_))
            codepoint_ = 0;
    }

    SourceStream& ss_;
    unsigned codepoint_;
};

///////////////////////////////////////////////////////////////////////////////
// GenericRegex

static const SizeType kRegexInvalidState = ~SizeType(0);  //!< Represents an invalid index in GenericRegex::State::out, out1
static const SizeType kRegexInvalidRange = ~SizeType(0);

template <typename Encoding, typename Allocator>
class GenericRegexSearch;

//! Regular expression engine with subset of ECMAscript grammar.
/*!
    Supported regular expression syntax:
    - \c ab     Concatenation
    - \c a|b    Alternation
    - \c a?     Zero or one
    - \c a*     Zero or more
    - \c a+     One or more
    - \c a{3}   Exactly 3 times
    - \c a{3,}  At least 3 times
    - \c a{3,5} 3 to 5 times
    - \c (ab)   Grouping
    - \c ^a     At the beginning
    - \c a$     At the end
    - \c .      Any character
    - \c [abc]  Character classes
    - \c [a-c]  Character class range
    - \c [a-z0-9_] Character class combination
    - \c [^abc] Negated character classes
    - \c [^a-c] Negated character class range
    - \c [\b]   Backspace (U+0008)
    - \c \\| \\\\ ...  Escape characters
    - \c \\f Form feed (U+000C)
    - \c \\n Line feed (U+000A)
    - \c \\r Carriage return (U+000D)
    - \c \\t Tab (U+0009)
    - \c \\v Vertical tab (U+000B)

    \note This is a Thompson NFA engine, implemented with reference to 
        Cox, Russ. "Regular Expression Matching Can Be Simple And Fast (but is slow in Java, Perl, PHP, Python, Ruby,...).", 
        https://swtch.com/~rsc/regexp/regexp1.html 
*/
template <typename Encoding, typename Allocator = CrtAllocator>
class GenericRegex {
public:
    typedef Encoding EncodingType;
    typedef typename Encoding::Ch Ch;
    template <typename, typename> friend class GenericRegexSearch;

    GenericRegex(const Ch* source, Allocator* allocator = 0) : 
        ownAllocator_(allocator ? 0 : CEREAL_RAPIDJSON_NEW(Allocator)()), allocator_(allocator ? allocator : ownAllocator_), 
        states_(allocator_, 256), ranges_(allocator_, 256), root_(kRegexInvalidState), stateCount_(), rangeCount_(), 
        anchorBegin_(), anchorEnd_()
    {
        GenericStringStream<Encoding> ss(source);
        DecodedStream<GenericStringStream<Encoding>, Encoding> ds(ss);
        Parse(ds);
    }

    ~GenericRegex()
    {
        CEREAL_RAPIDJSON_DELETE(ownAllocator_);
    }

    bool IsValid() const {
        return root_ != kRegexInvalidState;
    }

private:
    enum Operator {
        kZeroOrOne,
        kZeroOrMore,
        kOneOrMore,
        kConcatenation,
        kAlternation,
        kLeftParenthesis
    };

    static const unsigned kAnyCharacterClass = 0xFFFFFFFF;   //!< For '.'
    static const unsigned kRangeCharacterClass = 0xFFFFFFFE;
    static const unsigned kRangeNegationFlag = 0x80000000;

    struct Range {
        unsigned start; // 
        unsigned end;
        SizeType next;
    };

    struct State {
        SizeType out;     //!< Equals to kInvalid for matching state
        SizeType out1;    //!< Equals to non-kInvalid for split
        SizeType rangeStart;
        unsigned codepoint;
    };

    struct Frag {
        Frag(SizeType s, SizeType o, SizeType m) : start(s), out(o), minIndex(m) {}
        SizeType start;
        SizeType out; //!< link-list of all output states
        SizeType minIndex;
    };

    State& GetState(SizeType index) {
        CEREAL_RAPIDJSON_ASSERT(index < stateCount_);
        return states_.template Bottom<State>()[index];
    }

    const State& GetState(SizeType index) const {
        CEREAL_RAPIDJSON_ASSERT(index < stateCount_);
        return states_.template Bottom<State>()[index];
    }

    Range& GetRange(SizeType index) {
        CEREAL_RAPIDJSON_ASSERT(index < rangeCount_);
        return ranges_.template Bottom<Range>()[index];
    }

    const Range& GetRange(SizeType index) const {
        CEREAL_RAPIDJSON_ASSERT(index < rangeCount_);
        return ranges_.template Bottom<Range>()[index];
    }

    template <typename InputStream>
    void Parse(DecodedStream<InputStream, Encoding>& ds) {
        Stack<Allocator> operandStack(allocator_, 256);    // Frag
        Stack<Allocator> operatorStack(allocator_, 256);   // Operator
        Stack<Allocator> atomCountStack(allocator_, 256);  // unsigned (Atom per parenthesis)

        *atomCountStack.template Push<unsigned>() = 0;

        unsigned codepoint;
        while (ds.Peek() != 0) {
            switch (codepoint = ds.Take()) {
                case '^':
                    anchorBegin_ = true;
                    break;

                case '$':
                    anchorEnd_ = true;
                    break;

                case '|':
                    while (!operatorStack.Empty() && *operatorStack.template Top<Operator>() < kAlternation)
                        if (!Eval(operandStack, *operatorStack.template Pop<Operator>(1)))
                            return;
                    *operatorStack.template Push<Operator>() = kAlternation;
                    *atomCountStack.template Top<unsigned>() = 0;
                    break;

                case '(':
                    *operatorStack.template Push<Operator>() = kLeftParenthesis;
                    *atomCountStack.template Push<unsigned>() = 0;
                    break;

                case ')':
                    while (!operatorStack.Empty() && *operatorStack.template Top<Operator>() != kLeftParenthesis)
                        if (!Eval(operandStack, *operatorStack.template Pop<Operator>(1)))
                            return;
                    if (operatorStack.Empty())
                        return;
                    operatorStack.template Pop<Operator>(1);
                    atomCountStack.template Pop<unsigned>(1);
                    ImplicitConcatenation(atomCountStack, operatorStack);
                    break;

                case '?':
                    if (!Eval(operandStack, kZeroOrOne))
                        return;
                    break;

                case '*':
                    if (!Eval(operandStack, kZeroOrMore))
                        return;
                    break;

                case '+':
                    if (!Eval(operandStack, kOneOrMore))
                        return;
                    break;

                case '{':
                    {
                        unsigned n, m;
                        if (!ParseUnsigned(ds, &n))
                            return;

                        if (ds.Peek() == ',') {
                            ds.Take();
                            if (ds.Peek() == '}')
                                m = kInfinityQuantifier;
                            else if (!ParseUnsigned(ds, &m) || m < n)
                                return;
                        }
                        else
                            m = n;

                        if (!EvalQuantifier(operandStack, n, m) || ds.Peek() != '}')
                            return;
                        ds.Take();
                    }
                    break;

                case '.':
                    PushOperand(operandStack, kAnyCharacterClass);
                    ImplicitConcatenation(atomCountStack, operatorStack);
                    break;

                case '[':
                    {
                        SizeType range;
                        if (!ParseRange(ds, &range))
                            return;
                        SizeType s = NewState(kRegexInvalidState, kRegexInvalidState, kRangeCharacterClass);
                        GetState(s).rangeStart = range;
                        *operandStack.template Push<Frag>() = Frag(s, s, s);
                    }
                    ImplicitConcatenation(atomCountStack, operatorStack);
                    break;

                case '\\': // Escape character
                    if (!CharacterEscape(ds, &codepoint))
                        return; // Unsupported escape character
                    // fall through to default

                default: // Pattern character
                    PushOperand(operandStack, codepoint);
                    ImplicitConcatenation(atomCountStack, operatorStack);
            }
        }

        while (!operatorStack.Empty())
            if (!Eval(operandStack, *operatorStack.template Pop<Operator>(1)))
                return;

        // Link the operand to matching state.
        if (operandStack.GetSize() == sizeof(Frag)) {
            Frag* e = operandStack.template Pop<Frag>(1);
            Patch(e->out, NewState(kRegexInvalidState, kRegexInvalidState, 0));
            root_ = e->start;

#if CEREAL_RAPIDJSON_REGEX_VERBOSE
            printf("root: %d\n", root_);
            for (SizeType i = 0; i < stateCount_ ; i++) {
                State& s = GetState(i);
                printf("[%2d] out: %2d out1: %2d c: '%c'\n", i, s.out, s.out1, (char)s.codepoint);
            }
            printf("\n");
#endif
        }
    }

    SizeType NewState(SizeType out, SizeType out1, unsigned codepoint) {
        State* s = states_.template Push<State>();
        s->out = out;
        s->out1 = out1;
        s->codepoint = codepoint;
        s->rangeStart = kRegexInvalidRange;
        return stateCount_++;
    }

    void PushOperand(Stack<Allocator>& operandStack, unsigned codepoint) {
        SizeType s = NewState(kRegexInvalidState, kRegexInvalidState, codepoint);
        *operandStack.template Push<Frag>() = Frag(s, s, s);
    }

    void ImplicitConcatenation(Stack<Allocator>& atomCountStack, Stack<Allocator>& operatorStack) {
        if (*atomCountStack.template Top<unsigned>())
            *operatorStack.template Push<Operator>() = kConcatenation;
        (*atomCountStack.template Top<unsigned>())++;
    }

    SizeType Append(SizeType l1, SizeType l2) {
        SizeType old = l1;
        while (GetState(l1).out != kRegexInvalidState)
            l1 = GetState(l1).out;
        GetState(l1).out = l2;
        return old;
    }

    void Patch(SizeType l, SizeType s) {
        for (SizeType next; l != kRegexInvalidState; l = next) {
            next = GetState(l).out;
            GetState(l).out = s;
        }
    }

    bool Eval(Stack<Allocator>& operandStack, Operator op) {
        switch (op) {
            case kConcatenation:
                CEREAL_RAPIDJSON_ASSERT(operandStack.GetSize() >= sizeof(Frag) * 2);
                {
                    Frag e2 = *operandStack.template Pop<Frag>(1);
                    Frag e1 = *operandStack.template Pop<Frag>(1);
                    Patch(e1.out, e2.start);
                    *operandStack.template Push<Frag>() = Frag(e1.start, e2.out, Min(e1.minIndex, e2.minIndex));
                }
                return true;

            case kAlternation:
                if (operandStack.GetSize() >= sizeof(Frag) * 2) {
                    Frag e2 = *operandStack.template Pop<Frag>(1);
                    Frag e1 = *operandStack.template Pop<Frag>(1);
                    SizeType s = NewState(e1.start, e2.start, 0);
                    *operandStack.template Push<Frag>() = Frag(s, Append(e1.out, e2.out), Min(e1.minIndex, e2.minIndex));
                    return true;
                }
                return false;

            case kZeroOrOne:
                if (operandStack.GetSize() >= sizeof(Frag)) {
                    Frag e = *operandStack.template Pop<Frag>(1);
                    SizeType s = NewState(kRegexInvalidState, e.start, 0);
                    *operandStack.template Push<Frag>() = Frag(s, Append(e.out, s), e.minIndex);
                    return true;
                }
                return false;

            case kZeroOrMore:
                if (operandStack.GetSize() >= sizeof(Frag)) {
                    Frag e = *operandStack.template Pop<Frag>(1);
                    SizeType s = NewState(kRegexInvalidState, e.start, 0);
                    Patch(e.out, s);
                    *operandStack.template Push<Frag>() = Frag(s, s, e.minIndex);
                    return true;
                }
                return false;

            case kOneOrMore:
                if (operandStack.GetSize() >= sizeof(Frag)) {
                    Frag e = *operandStack.template Pop<Frag>(1);
                    SizeType s = NewState(kRegexInvalidState, e.start, 0);
                    Patch(e.out, s);
                    *operandStack.template Push<Frag>() = Frag(e.start, s, e.minIndex);
                    return true;
                }
                return false;

            default: 
                // syntax error (e.g. unclosed kLeftParenthesis)
                return false;
        }
    }

    bool EvalQuantifier(Stack<Allocator>& operandStack, unsigned n, unsigned m) {
        CEREAL_RAPIDJSON_ASSERT(n <= m);
        CEREAL_RAPIDJSON_ASSERT(operandStack.GetSize() >= sizeof(Frag));

        if (n == 0) {
            if (m == 0)                             // a{0} not support
                return false;
            else if (m == kInfinityQuantifier)
                Eval(operandStack, kZeroOrMore);    // a{0,} -> a*
            else {
                Eval(operandStack, kZeroOrOne);         // a{0,5} -> a?
                for (unsigned i = 0; i < m - 1; i++)
                    CloneTopOperand(operandStack);      // a{0,5} -> a? a? a? a? a?
                for (unsigned i = 0; i < m - 1; i++)
                    Eval(operandStack, kConcatenation); // a{0,5} -> a?a?a?a?a?
            }
            return true;
        }

        for (unsigned i = 0; i < n - 1; i++)        // a{3} -> a a a
            CloneTopOperand(operandStack);

        if (m == kInfinityQuantifier)
            Eval(operandStack, kOneOrMore);         // a{3,} -> a a a+
        else if (m > n) {
            CloneTopOperand(operandStack);          // a{3,5} -> a a a a
            Eval(operandStack, kZeroOrOne);         // a{3,5} -> a a a a?
            for (unsigned i = n; i < m - 1; i++)
                CloneTopOperand(operandStack);      // a{3,5} -> a a a a? a?
            for (unsigned i = n; i < m; i++)
                Eval(operandStack, kConcatenation); // a{3,5} -> a a aa?a?
        }

        for (unsigned i = 0; i < n - 1; i++)
            Eval(operandStack, kConcatenation);     // a{3} -> aaa, a{3,} -> aaa+, a{3.5} -> aaaa?a?

        return true;
    }

    static SizeType Min(SizeType a, SizeType b) { return a < b ? a : b; }

    void CloneTopOperand(Stack<Allocator>& operandStack) {
        const Frag src = *operandStack.template Top<Frag>(); // Copy constructor to prevent invalidation
        SizeType count = stateCount_ - src.minIndex; // Assumes top operand contains states in [src->minIndex, stateCount_)
        State* s = states_.template Push<State>(count);
        memcpy(s, &GetState(src.minIndex), count * sizeof(State));
        for (SizeType j = 0; j < count; j++) {
            if (s[j].out != kRegexInvalidState)
                s[j].out += count;
            if (s[j].out1 != kRegexInvalidState)
                s[j].out1 += count;
        }
        *operandStack.template Push<Frag>() = Frag(src.start + count, src.out + count, src.minIndex + count);
        stateCount_ += count;
    }

    template <typename InputStream>
    bool ParseUnsigned(DecodedStream<InputStream, Encoding>& ds, unsigned* u) {
        unsigned r = 0;
        if (ds.Peek() < '0' || ds.Peek() > '9')
            return false;
        while (ds.Peek() >= '0' && ds.Peek() <= '9') {
            if (r >= 429496729 && ds.Peek() > '5') // 2^32 - 1 = 4294967295
                return false; // overflow
            r = r * 10 + (ds.Take() - '0');
        }
        *u = r;
        return true;
    }

    template <typename InputStream>
    bool ParseRange(DecodedStream<InputStream, Encoding>& ds, SizeType* range) {
        bool isBegin = true;
        bool negate = false;
        int step = 0;
        SizeType start = kRegexInvalidRange;
        SizeType current = kRegexInvalidRange;
        unsigned codepoint;
        while ((codepoint = ds.Take()) != 0) {
            if (isBegin) {
                isBegin = false;
                if (codepoint == '^') {
                    negate = true;
                    continue;
                }
            }

            switch (codepoint) {
            case ']':
                if (start == kRegexInvalidRange)
                    return false;   // Error: nothing inside []
                if (step == 2) { // Add trailing '-'
                    SizeType r = NewRange('-');
                    CEREAL_RAPIDJSON_ASSERT(current != kRegexInvalidRange);
                    GetRange(current).next = r;
                }
                if (negate)
                    GetRange(start).start |= kRangeNegationFlag;
                *range = start;
                return true;

            case '\\':
                if (ds.Peek() == 'b') {
                    ds.Take();
                    codepoint = 0x0008; // Escape backspace character
                }
                else if (!CharacterEscape(ds, &codepoint))
                    return false;
                // fall through to default

            default:
                switch (step) {
                case 1:
                    if (codepoint == '-') {
                        step++;
                        break;
                    }
                    // fall through to step 0 for other characters

                case 0:
                    {
                        SizeType r = NewRange(codepoint);
                        if (current != kRegexInvalidRange)
                            GetRange(current).next = r;
                        if (start == kRegexInvalidRange)
                            start = r;
                        current = r;
                    }
                    step = 1;
                    break;

                default:
                    CEREAL_RAPIDJSON_ASSERT(step == 2);
                    GetRange(current).end = codepoint;
                    step = 0;
                }
            }
        }
        return false;
    }
    
    SizeType NewRange(unsigned codepoint) {
        Range* r = ranges_.template Push<Range>();
        r->start = r->end = codepoint;
        r->next = kRegexInvalidRange;
        return rangeCount_++;
    }

    template <typename InputStream>
    bool CharacterEscape(DecodedStream<InputStream, Encoding>& ds, unsigned* escapedCodepoint) {
        unsigned codepoint;
        switch (codepoint = ds.Take()) {
            case '^':
            case '$':
            case '|':
            case '(':
            case ')':
            case '?':
            case '*':
            case '+':
            case '.':
            case '[':
            case ']':
            case '{':
            case '}':
            case '\\':
                *escapedCodepoint = codepoint; return true;
            case 'f': *escapedCodepoint = 0x000C; return true;
            case 'n': *escapedCodepoint = 0x000A; return true;
            case 'r': *escapedCodepoint = 0x000D; return true;
            case 't': *escapedCodepoint = 0x0009; return true;
            case 'v': *escapedCodepoint = 0x000B; return true;
            default:
                return false; // Unsupported escape character
        }
    }

    Allocator* ownAllocator_;
    Allocator* allocator_;
    Stack<Allocator> states_;
    Stack<Allocator> ranges_;
    SizeType root_;
    SizeType stateCount_;
    SizeType rangeCount_;

    static const unsigned kInfinityQuantifier = ~0u;

    // For SearchWithAnchoring()
    bool anchorBegin_;
    bool anchorEnd_;
};

template <typename RegexType, typename Allocator = CrtAllocator>
class GenericRegexSearch {
public:
    typedef typename RegexType::EncodingType Encoding;
    typedef typename Encoding::Ch Ch;

    GenericRegexSearch(const RegexType& regex, Allocator* allocator = 0) : 
        regex_(regex), allocator_(allocator), ownAllocator_(0),
        state0_(allocator, 0), state1_(allocator, 0), stateSet_()
    {
        CEREAL_RAPIDJSON_ASSERT(regex_.IsValid());
        if (!allocator_)
            ownAllocator_ = allocator_ = CEREAL_RAPIDJSON_NEW(Allocator)();
        stateSet_ = static_cast<unsigned*>(allocator_->Malloc(GetStateSetSize()));
        state0_.template Reserve<SizeType>(regex_.stateCount_);
        state1_.template Reserve<SizeType>(regex_.stateCount_);
    }

    ~GenericRegexSearch() {
        Allocator::Free(stateSet_);
        CEREAL_RAPIDJSON_DELETE(ownAllocator_);
    }

    template <typename InputStream>
    bool Match(InputStream& is) {
        return SearchWithAnchoring(is, true, true);
    }

    bool Match(const Ch* s) {
        GenericStringStream<Encoding> is(s);
        return Match(is);
    }

    template <typename InputStream>
    bool Search(InputStream& is) {
        return SearchWithAnchoring(is, regex_.anchorBegin_, regex_.anchorEnd_);
    }

    bool Search(const Ch* s) {
        GenericStringStream<Encoding> is(s);
        return Search(is);
    }

private:
    typedef typename RegexType::State State;
    typedef typename RegexType::Range Range;

    template <typename InputStream>
    bool SearchWithAnchoring(InputStream& is, bool anchorBegin, bool anchorEnd) {
        DecodedStream<InputStream, Encoding> ds(is);

        state0_.Clear();
        Stack<Allocator> *current = &state0_, *next = &state1_;
        const size_t stateSetSize = GetStateSetSize();
        std::memset(stateSet_, 0, stateSetSize);

        bool matched = AddState(*current, regex_.root_);
        unsigned codepoint;
        while (!current->Empty() && (codepoint = ds.Take()) != 0) {
            std::memset(stateSet_, 0, stateSetSize);
            next->Clear();
            matched = false;
            for (const SizeType* s = current->template Bottom<SizeType>(); s != current->template End<SizeType>(); ++s) {
                const State& sr = regex_.GetState(*s);
                if (sr.codepoint == codepoint ||
                    sr.codepoint == RegexType::kAnyCharacterClass || 
                    (sr.codepoint == RegexType::kRangeCharacterClass && MatchRange(sr.rangeStart, codepoint)))
                {
                    matched = AddState(*next, sr.out) || matched;
                    if (!anchorEnd && matched)
                        return true;
                }
                if (!anchorBegin)
                    AddState(*next, regex_.root_);
            }
            internal::Swap(current, next);
        }

        return matched;
    }

    size_t GetStateSetSize() const {
        return (regex_.stateCount_ + 31) / 32 * 4;
    }

    // Return whether the added states is a match state
    bool AddState(Stack<Allocator>& l, SizeType index) {
        CEREAL_RAPIDJSON_ASSERT(index != kRegexInvalidState);

        const State& s = regex_.GetState(index);
        if (s.out1 != kRegexInvalidState) { // Split
            bool matched = AddState(l, s.out);
            return AddState(l, s.out1) || matched;
        }
        else if (!(stateSet_[index >> 5] & (1u << (index & 31)))) {
            stateSet_[index >> 5] |= (1u << (index & 31));
            *l.template PushUnsafe<SizeType>() = index;
        }
        return s.out == kRegexInvalidState; // by using PushUnsafe() above, we can ensure s is not validated due to reallocation.
    }

    bool MatchRange(SizeType rangeIndex, unsigned codepoint) const {
        bool yes = (regex_.GetRange(rangeIndex).start & RegexType::kRangeNegationFlag) == 0;
        while (rangeIndex != kRegexInvalidRange) {
            const Range& r = regex_.GetRange(rangeIndex);
            if (codepoint >= (r.start & ~RegexType::kRangeNegationFlag) && codepoint <= r.end)
                return yes;
            rangeIndex = r.next;
        }
        return !yes;
    }

    const RegexType& regex_;
    Allocator* allocator_;
    Allocator* ownAllocator_;
    Stack<Allocator> state0_;
    Stack<Allocator> state1_;
    uint32_t* stateSet_;
};

typedef GenericRegex<UTF8<> > Regex;
typedef GenericRegexSearch<Regex> RegexSearch;

} // namespace internal
CEREAL_RAPIDJSON_NAMESPACE_END

#ifdef __GNUC__
CEREAL_RAPIDJSON_DIAG_POP
#endif

#if defined(__clang__) || defined(_MSC_VER)
CEREAL_RAPIDJSON_DIAG_POP
#endif

#endif // CEREAL_RAPIDJSON_INTERNAL_REGEX_H_
```
