## Supplemental code for "The motif composition of variable-number tandem repeats impacts gene expression": stack.h

```
// Tencent is pleased to support the open source community by making RapidJSON available.
// 
// Copyright (C) 2015 THL A29 Limited, a Tencent company, and Milo Yip. All rights reserved.
//
// Licensed under the MIT License (the "License"); you may not use this file except
// in compliance with the License. You may obtain a copy of the License at
//
// http://opensource.org/licenses/MIT
//
// Unless required by applicable law or agreed to in writing, software distributed 
// under the License is distributed on an "AS IS" BASIS, WITHOUT WARRANTIES OR 
// CONDITIONS OF ANY KIND, either express or implied. See the License for the 
// specific language governing permissions and limitations under the License.

#ifndef CEREAL_RAPIDJSON_INTERNAL_STACK_H_
#define CEREAL_RAPIDJSON_INTERNAL_STACK_H_

#include "../allocators.h"
#include "swap.h"
#include <cstddef>

#if defined(__clang__)
CEREAL_RAPIDJSON_DIAG_PUSH
CEREAL_RAPIDJSON_DIAG_OFF(c++98-compat)
#endif

CEREAL_RAPIDJSON_NAMESPACE_BEGIN
namespace internal {

///////////////////////////////////////////////////////////////////////////////
// Stack

//! A type-unsafe stack for storing different types of data.
/*! \tparam Allocator Allocator for allocating stack memory.
*/
template <typename Allocator>
class Stack {
public:
    // Optimization note: Do not allocate memory for stack_ in constructor.
    // Do it lazily when first Push() -> Expand() -> Resize().
    Stack(Allocator* allocator, size_t stackCapacity) : allocator_(allocator), ownAllocator_(0), stack_(0), stackTop_(0), stackEnd_(0), initialCapacity_(stackCapacity) {
    }

#if CEREAL_RAPIDJSON_HAS_CXX11_RVALUE_REFS
    Stack(Stack&& rhs)
        : allocator_(rhs.allocator_),
          ownAllocator_(rhs.ownAllocator_),
          stack_(rhs.stack_),
          stackTop_(rhs.stackTop_),
          stackEnd_(rhs.stackEnd_),
          initialCapacity_(rhs.initialCapacity_)
    {
        rhs.allocator_ = 0;
        rhs.ownAllocator_ = 0;
        rhs.stack_ = 0;
        rhs.stackTop_ = 0;
        rhs.stackEnd_ = 0;
        rhs.initialCapacity_ = 0;
    }
#endif

    ~Stack() {
        Destroy();
    }

#if CEREAL_RAPIDJSON_HAS_CXX11_RVALUE_REFS
    Stack& operator=(Stack&& rhs) {
        if (&rhs != this)
        {
            Destroy();

            allocator_ = rhs.allocator_;
            ownAllocator_ = rhs.ownAllocator_;
            stack_ = rhs.stack_;
            stackTop_ = rhs.stackTop_;
            stackEnd_ = rhs.stackEnd_;
            initialCapacity_ = rhs.initialCapacity_;

            rhs.allocator_ = 0;
            rhs.ownAllocator_ = 0;
            rhs.stack_ = 0;
            rhs.stackTop_ = 0;
            rhs.stackEnd_ = 0;
            rhs.initialCapacity_ = 0;
        }
        return *this;
    }
#endif

    void Swap(Stack& rhs) CEREAL_RAPIDJSON_NOEXCEPT {
        internal::Swap(allocator_, rhs.allocator_);
        internal::Swap(ownAllocator_, rhs.ownAllocator_);
        internal::Swap(stack_, rhs.stack_);
        internal::Swap(stackTop_, rhs.stackTop_);
        internal::Swap(stackEnd_, rhs.stackEnd_);
        internal::Swap(initialCapacity_, rhs.initialCapacity_);
    }

    void Clear() { stackTop_ = stack_; }

    void ShrinkToFit() { 
        if (Empty()) {
            // If the stack is empty, completely deallocate the memory.
            Allocator::Free(stack_); // NOLINT (+clang-analyzer-unix.Malloc)
            stack_ = 0;
            stackTop_ = 0;
            stackEnd_ = 0;
        }
        else
            Resize(GetSize());
    }

    // Optimization note: try to minimize the size of this function for force inline.
    // Expansion is run very infrequently, so it is moved to another (probably non-inline) function.
    template<typename T>
    CEREAL_RAPIDJSON_FORCEINLINE void Reserve(size_t count = 1) {
         // Expand the stack if needed
        if (CEREAL_RAPIDJSON_UNLIKELY(static_cast<std::ptrdiff_t>(sizeof(T) * count) > (stackEnd_ - stackTop_)))
            Expand<T>(count);
    }

    template<typename T>
    CEREAL_RAPIDJSON_FORCEINLINE T* Push(size_t count = 1) {
        Reserve<T>(count);
        return PushUnsafe<T>(count);
    }

    template<typename T>
    CEREAL_RAPIDJSON_FORCEINLINE T* PushUnsafe(size_t count = 1) {
        CEREAL_RAPIDJSON_ASSERT(stackTop_);
        CEREAL_RAPIDJSON_ASSERT(static_cast<std::ptrdiff_t>(sizeof(T) * count) <= (stackEnd_ - stackTop_));
        T* ret = reinterpret_cast<T*>(stackTop_);
        stackTop_ += sizeof(T) * count;
        return ret;
    }

    template<typename T>
    T* Pop(size_t count) {
        CEREAL_RAPIDJSON_ASSERT(GetSize() >= count * sizeof(T));
        stackTop_ -= count * sizeof(T);
        return reinterpret_cast<T*>(stackTop_);
    }

    template<typename T>
    T* Top() { 
        CEREAL_RAPIDJSON_ASSERT(GetSize() >= sizeof(T));
        return reinterpret_cast<T*>(stackTop_ - sizeof(T));
    }

    template<typename T>
    const T* Top() const {
        CEREAL_RAPIDJSON_ASSERT(GetSize() >= sizeof(T));
        return reinterpret_cast<T*>(stackTop_ - sizeof(T));
    }

    template<typename T>
    T* End() { return reinterpret_cast<T*>(stackTop_); }

    template<typename T>
    const T* End() const { return reinterpret_cast<T*>(stackTop_); }

    template<typename T>
    T* Bottom() { return reinterpret_cast<T*>(stack_); }

    template<typename T>
    const T* Bottom() const { return reinterpret_cast<T*>(stack_); }

    bool HasAllocator() const {
        return allocator_ != 0;
    }

    Allocator& GetAllocator() {
        CEREAL_RAPIDJSON_ASSERT(allocator_);
        return *allocator_;
    }

    bool Empty() const { return stackTop_ == stack_; }
    size_t GetSize() const { return static_cast<size_t>(stackTop_ - stack_); }
    size_t GetCapacity() const { return static_cast<size_t>(stackEnd_ - stack_); }

private:
    template<typename T>
    void Expand(size_t count) {
        // Only expand the capacity if the current stack exists. Otherwise just create a stack with initial capacity.
        size_t newCapacity;
        if (stack_ == 0) {
            if (!allocator_)
                ownAllocator_ = allocator_ = CEREAL_RAPIDJSON_NEW(Allocator)();
            newCapacity = initialCapacity_;
        } else {
            newCapacity = GetCapacity();
            newCapacity += (newCapacity + 1) / 2;
        }
        size_t newSize = GetSize() + sizeof(T) * count;
        if (newCapacity < newSize)
            newCapacity = newSize;

        Resize(newCapacity);
    }

    void Resize(size_t newCapacity) {
        const size_t size = GetSize();  // Backup the current size
        stack_ = static_cast<char*>(allocator_->Realloc(stack_, GetCapacity(), newCapacity));
        stackTop_ = stack_ + size;
        stackEnd_ = stack_ + newCapacity;
    }

    void Destroy() {
        Allocator::Free(stack_);
        CEREAL_RAPIDJSON_DELETE(ownAllocator_); // Only delete if it is owned by the stack
    }

    // Prohibit copy constructor & assignment operator.
    Stack(const Stack&);
    Stack& operator=(const Stack&);

    Allocator* allocator_;
    Allocator* ownAllocator_;
    char *stack_;
    char *stackTop_;
    char *stackEnd_;
    size_t initialCapacity_;
};

} // namespace internal
CEREAL_RAPIDJSON_NAMESPACE_END

#if defined(__clang__)
CEREAL_RAPIDJSON_DIAG_POP
#endif

#endif // CEREAL_RAPIDJSON_STACK_H_
```
