## Supplemental code for "The motif composition of variable-number tandem repeats impacts gene expression": strfunc.h

```
// Tencent is pleased to support the open source community by making RapidJSON available.
// 
// Copyright (C) 2015 THL A29 Limited, a Tencent company, and Milo Yip. All rights reserved.
//
// Licensed under the MIT License (the "License"); you may not use this file except
// in compliance with the License. You may obtain a copy of the License at
//
// http://opensource.org/licenses/MIT
//
// Unless required by applicable law or agreed to in writing, software distributed 
// under the License is distributed on an "AS IS" BASIS, WITHOUT WARRANTIES OR 
// CONDITIONS OF ANY KIND, either express or implied. See the License for the 
// specific language governing permissions and limitations under the License.

#ifndef CEREAL_RAPIDJSON_INTERNAL_STRFUNC_H_
#define CEREAL_RAPIDJSON_INTERNAL_STRFUNC_H_

#include "../stream.h"
#include <cwchar>

CEREAL_RAPIDJSON_NAMESPACE_BEGIN
namespace internal {

//! Custom strlen() which works on different character types.
/*! \tparam Ch Character type (e.g. char, wchar_t, short)
    \param s Null-terminated input string.
    \return Number of characters in the string. 
    \note This has the same semantics as strlen(), the return value is not number of Unicode codepoints.
*/
template <typename Ch>
inline SizeType StrLen(const Ch* s) {
    CEREAL_RAPIDJSON_ASSERT(s != 0);
    const Ch* p = s;
    while (*p) ++p;
    return SizeType(p - s);
}

template <>
inline SizeType StrLen(const char* s) {
    return SizeType(std::strlen(s));
}

template <>
inline SizeType StrLen(const wchar_t* s) {
    return SizeType(std::wcslen(s));
}

//! Returns number of code points in a encoded string.
template<typename Encoding>
bool CountStringCodePoint(const typename Encoding::Ch* s, SizeType length, SizeType* outCount) {
    CEREAL_RAPIDJSON_ASSERT(s != 0);
    CEREAL_RAPIDJSON_ASSERT(outCount != 0);
    GenericStringStream<Encoding> is(s);
    const typename Encoding::Ch* end = s + length;
    SizeType count = 0;
    while (is.src_ < end) {
        unsigned codepoint;
        if (!Encoding::Decode(is, &codepoint))
            return false;
        count++;
    }
    *outCount = count;
    return true;
}

} // namespace internal
CEREAL_RAPIDJSON_NAMESPACE_END

#endif // CEREAL_RAPIDJSON_INTERNAL_STRFUNC_H_
```
