## Supplemental code for "The motif composition of variable-number tandem repeats impacts gene expression": strtod.h

```
// Tencent is pleased to support the open source community by making RapidJSON available.
// 
// Copyright (C) 2015 THL A29 Limited, a Tencent company, and Milo Yip. All rights reserved.
//
// Licensed under the MIT License (the "License"); you may not use this file except
// in compliance with the License. You may obtain a copy of the License at
//
// http://opensource.org/licenses/MIT
//
// Unless required by applicable law or agreed to in writing, software distributed 
// under the License is distributed on an "AS IS" BASIS, WITHOUT WARRANTIES OR 
// CONDITIONS OF ANY KIND, either express or implied. See the License for the 
// specific language governing permissions and limitations under the License.

#ifndef CEREAL_RAPIDJSON_STRTOD_
#define CEREAL_RAPIDJSON_STRTOD_

#include "ieee754.h"
#include "biginteger.h"
#include "diyfp.h"
#include "pow10.h"
#include <climits>
#include <limits>

CEREAL_RAPIDJSON_NAMESPACE_BEGIN
namespace internal {

inline double FastPath(double significand, int exp) {
    if (exp < -308)
        return 0.0;
    else if (exp >= 0)
        return significand * internal::Pow10(exp);
    else
        return significand / internal::Pow10(-exp);
}

inline double StrtodNormalPrecision(double d, int p) {
    if (p < -308) {
        // Prevent expSum < -308, making Pow10(p) = 0
        d = FastPath(d, -308);
        d = FastPath(d, p + 308);
    }
    else
        d = FastPath(d, p);
    return d;
}

template <typename T>
inline T Min3(T a, T b, T c) {
    T m = a;
    if (m > b) m = b;
    if (m > c) m = c;
    return m;
}

inline int CheckWithinHalfULP(double b, const BigInteger& d, int dExp) {
    const Double db(b);
    const uint64_t bInt = db.IntegerSignificand();
    const int bExp = db.IntegerExponent();
    const int hExp = bExp - 1;

    int dS_Exp2 = 0, dS_Exp5 = 0, bS_Exp2 = 0, bS_Exp5 = 0, hS_Exp2 = 0, hS_Exp5 = 0;

    // Adjust for decimal exponent
    if (dExp >= 0) {
        dS_Exp2 += dExp;
        dS_Exp5 += dExp;
    }
    else {
        bS_Exp2 -= dExp;
        bS_Exp5 -= dExp;
        hS_Exp2 -= dExp;
        hS_Exp5 -= dExp;
    }

    // Adjust for binary exponent
    if (bExp >= 0)
        bS_Exp2 += bExp;
    else {
        dS_Exp2 -= bExp;
        hS_Exp2 -= bExp;
    }

    // Adjust for half ulp exponent
    if (hExp >= 0)
        hS_Exp2 += hExp;
    else {
        dS_Exp2 -= hExp;
        bS_Exp2 -= hExp;
    }

    // Remove common power of two factor from all three scaled values
    int common_Exp2 = Min3(dS_Exp2, bS_Exp2, hS_Exp2);
    dS_Exp2 -= common_Exp2;
    bS_Exp2 -= common_Exp2;
    hS_Exp2 -= common_Exp2;

    BigInteger dS = d;
    dS.MultiplyPow5(static_cast<unsigned>(dS_Exp5)) <<= static_cast<unsigned>(dS_Exp2);

    BigInteger bS(bInt);
    bS.MultiplyPow5(static_cast<unsigned>(bS_Exp5)) <<= static_cast<unsigned>(bS_Exp2);

    BigInteger hS(1);
    hS.MultiplyPow5(static_cast<unsigned>(hS_Exp5)) <<= static_cast<unsigned>(hS_Exp2);

    BigInteger delta(0);
    dS.Difference(bS, &delta);

    return delta.Compare(hS);
}

inline bool StrtodFast(double d, int p, double* result) {
    // Use fast path for string-to-double conversion if possible
    // see http://www.exploringbinary.com/fast-path-decimal-to-floating-point-conversion/
    if (p > 22  && p < 22 + 16) {
        // Fast Path Cases In Disguise
        d *= internal::Pow10(p - 22);
        p = 22;
    }

    if (p >= -22 && p <= 22 && d <= 9007199254740991.0) { // 2^53 - 1
        *result = FastPath(d, p);
        return true;
    }
    else
        return false;
}

// Compute an approximation and see if it is within 1/2 ULP
inline bool StrtodDiyFp(const char* decimals, int dLen, int dExp, double* result) {
    uint64_t significand = 0;
    int i = 0;   // 2^64 - 1 = 18446744073709551615, 1844674407370955161 = 0x1999999999999999    
    for (; i < dLen; i++) {
        if (significand  >  CEREAL_RAPIDJSON_UINT64_C2(0x19999999, 0x99999999) ||
            (significand == CEREAL_RAPIDJSON_UINT64_C2(0x19999999, 0x99999999) && decimals[i] > '5'))
            break;
        significand = significand * 10u + static_cast<unsigned>(decimals[i] - '0');
    }
    
    if (i < dLen && decimals[i] >= '5') // Rounding
        significand++;

    int remaining = dLen - i;
    const int kUlpShift = 3;
    const int kUlp = 1 << kUlpShift;
    int64_t error = (remaining == 0) ? 0 : kUlp / 2;

    DiyFp v(significand, 0);
    v = v.Normalize();
    error <<= -v.e;

    dExp += remaining;

    int actualExp;
    DiyFp cachedPower = GetCachedPower10(dExp, &actualExp);
    if (actualExp != dExp) {
        static const DiyFp kPow10[] = {
            DiyFp(CEREAL_RAPIDJSON_UINT64_C2(0xa0000000, 0x00000000), -60),  // 10^1
            DiyFp(CEREAL_RAPIDJSON_UINT64_C2(0xc8000000, 0x00000000), -57),  // 10^2
            DiyFp(CEREAL_RAPIDJSON_UINT64_C2(0xfa000000, 0x00000000), -54),  // 10^3
            DiyFp(CEREAL_RAPIDJSON_UINT64_C2(0x9c400000, 0x00000000), -50),  // 10^4
            DiyFp(CEREAL_RAPIDJSON_UINT64_C2(0xc3500000, 0x00000000), -47),  // 10^5
            DiyFp(CEREAL_RAPIDJSON_UINT64_C2(0xf4240000, 0x00000000), -44),  // 10^6
            DiyFp(CEREAL_RAPIDJSON_UINT64_C2(0x98968000, 0x00000000), -40)   // 10^7
        };
        int adjustment = dExp - actualExp;
        CEREAL_RAPIDJSON_ASSERT(adjustment >= 1 && adjustment < 8);
        v = v * kPow10[adjustment - 1];
        if (dLen + adjustment > 19) // has more digits than decimal digits in 64-bit
            error += kUlp / 2;
    }

    v = v * cachedPower;

    error += kUlp + (error == 0 ? 0 : 1);

    const int oldExp = v.e;
    v = v.Normalize();
    error <<= oldExp - v.e;

    const int effectiveSignificandSize = Double::EffectiveSignificandSize(64 + v.e);
    int precisionSize = 64 - effectiveSignificandSize;
    if (precisionSize + kUlpShift >= 64) {
        int scaleExp = (precisionSize + kUlpShift) - 63;
        v.f >>= scaleExp;
        v.e += scaleExp; 
        error = (error >> scaleExp) + 1 + kUlp;
        precisionSize -= scaleExp;
    }

    DiyFp rounded(v.f >> precisionSize, v.e + precisionSize);
    const uint64_t precisionBits = (v.f & ((uint64_t(1) << precisionSize) - 1)) * kUlp;
    const uint64_t halfWay = (uint64_t(1) << (precisionSize - 1)) * kUlp;
    if (precisionBits >= halfWay + static_cast<unsigned>(error)) {
        rounded.f++;
        if (rounded.f & (DiyFp::kDpHiddenBit << 1)) { // rounding overflows mantissa (issue #340)
            rounded.f >>= 1;
            rounded.e++;
        }
    }

    *result = rounded.ToDouble();

    return halfWay - static_cast<unsigned>(error) >= precisionBits || precisionBits >= halfWay + static_cast<unsigned>(error);
}

inline double StrtodBigInteger(double approx, const char* decimals, int dLen, int dExp) {
    CEREAL_RAPIDJSON_ASSERT(dLen >= 0);
    const BigInteger dInt(decimals, static_cast<unsigned>(dLen));
    Double a(approx);
    int cmp = CheckWithinHalfULP(a.Value(), dInt, dExp);
    if (cmp < 0)
        return a.Value();  // within half ULP
    else if (cmp == 0) {
        // Round towards even
        if (a.Significand() & 1)
            return a.NextPositiveDouble();
        else
            return a.Value();
    }
    else // adjustment
        return a.NextPositiveDouble();
}

inline double StrtodFullPrecision(double d, int p, const char* decimals, size_t length, size_t decimalPosition, int exp) {
    CEREAL_RAPIDJSON_ASSERT(d >= 0.0);
    CEREAL_RAPIDJSON_ASSERT(length >= 1);

    double result = 0.0;
    if (StrtodFast(d, p, &result))
        return result;

    CEREAL_RAPIDJSON_ASSERT(length <= INT_MAX);
    int dLen = static_cast<int>(length);

    CEREAL_RAPIDJSON_ASSERT(length >= decimalPosition);
    CEREAL_RAPIDJSON_ASSERT(length - decimalPosition <= INT_MAX);
    int dExpAdjust = static_cast<int>(length - decimalPosition);

    CEREAL_RAPIDJSON_ASSERT(exp >= INT_MIN + dExpAdjust);
    int dExp = exp - dExpAdjust;

    // Make sure length+dExp does not overflow
    CEREAL_RAPIDJSON_ASSERT(dExp <= INT_MAX - dLen);

    // Trim leading zeros
    while (dLen > 0 && *decimals == '0') {
        dLen--;
        decimals++;
    }

    // Trim trailing zeros
    while (dLen > 0 && decimals[dLen - 1] == '0') {
        dLen--;
        dExp++;
    }

    if (dLen == 0) { // Buffer only contains zeros.
        return 0.0;
    }

    // Trim right-most digits
    const int kMaxDecimalDigit = 767 + 1;
    if (dLen > kMaxDecimalDigit) {
        dExp += dLen - kMaxDecimalDigit;
        dLen = kMaxDecimalDigit;
    }

    // If too small, underflow to zero.
    // Any x <= 10^-324 is interpreted as zero.
    if (dLen + dExp <= -324)
        return 0.0;

    // If too large, overflow to infinity.
    // Any x >= 10^309 is interpreted as +infinity.
    if (dLen + dExp > 309)
        return std::numeric_limits<double>::infinity();

    if (StrtodDiyFp(decimals, dLen, dExp, &result))
        return result;

    // Use approximation from StrtodDiyFp and make adjustment with BigInteger comparison
    return StrtodBigInteger(result, decimals, dLen, dExp);
}

} // namespace internal
CEREAL_RAPIDJSON_NAMESPACE_END

#endif // CEREAL_RAPIDJSON_STRTOD_
```
