## Supplemental code for "The motif composition of variable-number tandem repeats impacts gene expression": swap.h

```
// Tencent is pleased to support the open source community by making RapidJSON available.
//
// Copyright (C) 2015 THL A29 Limited, a Tencent company, and Milo Yip. All rights reserved.
//
// Licensed under the MIT License (the "License"); you may not use this file except
// in compliance with the License. You may obtain a copy of the License at
//
// http://opensource.org/licenses/MIT
//
// Unless required by applicable law or agreed to in writing, software distributed
// under the License is distributed on an "AS IS" BASIS, WITHOUT WARRANTIES OR
// CONDITIONS OF ANY KIND, either express or implied. See the License for the
// specific language governing permissions and limitations under the License.

#ifndef CEREAL_RAPIDJSON_INTERNAL_SWAP_H_
#define CEREAL_RAPIDJSON_INTERNAL_SWAP_H_

#include "../rapidjson.h"

#if defined(__clang__)
CEREAL_RAPIDJSON_DIAG_PUSH
CEREAL_RAPIDJSON_DIAG_OFF(c++98-compat)
#endif

CEREAL_RAPIDJSON_NAMESPACE_BEGIN
namespace internal {

//! Custom swap() to avoid dependency on C++ <algorithm> header
/*! \tparam T Type of the arguments to swap, should be instantiated with primitive C++ types only.
    \note This has the same semantics as std::swap().
*/
template <typename T>
inline void Swap(T& a, T& b) CEREAL_RAPIDJSON_NOEXCEPT {
    T tmp = a;
        a = b;
        b = tmp;
}

} // namespace internal
CEREAL_RAPIDJSON_NAMESPACE_END

#if defined(__clang__)
CEREAL_RAPIDJSON_DIAG_POP
#endif

#endif // CEREAL_RAPIDJSON_INTERNAL_SWAP_H_
```
