## Supplemental code for "The motif composition of variable-number tandem repeats impacts gene expression": memorybuffer.h

```
// Tencent is pleased to support the open source community by making RapidJSON available.
// 
// Copyright (C) 2015 THL A29 Limited, a Tencent company, and Milo Yip. All rights reserved.
//
// Licensed under the MIT License (the "License"); you may not use this file except
// in compliance with the License. You may obtain a copy of the License at
//
// http://opensource.org/licenses/MIT
//
// Unless required by applicable law or agreed to in writing, software distributed 
// under the License is distributed on an "AS IS" BASIS, WITHOUT WARRANTIES OR 
// CONDITIONS OF ANY KIND, either express or implied. See the License for the 
// specific language governing permissions and limitations under the License.

#ifndef CEREAL_RAPIDJSON_MEMORYBUFFER_H_
#define CEREAL_RAPIDJSON_MEMORYBUFFER_H_

#include "stream.h"
#include "internal/stack.h"

CEREAL_RAPIDJSON_NAMESPACE_BEGIN

//! Represents an in-memory output byte stream.
/*!
    This class is mainly for being wrapped by EncodedOutputStream or AutoUTFOutputStream.

    It is similar to FileWriteBuffer but the destination is an in-memory buffer instead of a file.

    Differences between MemoryBuffer and StringBuffer:
    1. StringBuffer has Encoding but MemoryBuffer is only a byte buffer. 
    2. StringBuffer::GetString() returns a null-terminated string. MemoryBuffer::GetBuffer() returns a buffer without terminator.

    \tparam Allocator type for allocating memory buffer.
    \note implements Stream concept
*/
template <typename Allocator = CrtAllocator>
struct GenericMemoryBuffer {
    typedef char Ch; // byte

    GenericMemoryBuffer(Allocator* allocator = 0, size_t capacity = kDefaultCapacity) : stack_(allocator, capacity) {}

    void Put(Ch c) { *stack_.template Push<Ch>() = c; }
    void Flush() {}

    void Clear() { stack_.Clear(); }
    void ShrinkToFit() { stack_.ShrinkToFit(); }
    Ch* Push(size_t count) { return stack_.template Push<Ch>(count); }
    void Pop(size_t count) { stack_.template Pop<Ch>(count); }

    const Ch* GetBuffer() const {
        return stack_.template Bottom<Ch>();
    }

    size_t GetSize() const { return stack_.GetSize(); }

    static const size_t kDefaultCapacity = 256;
    mutable internal::Stack<Allocator> stack_;
};

typedef GenericMemoryBuffer<> MemoryBuffer;

//! Implement specialized version of PutN() with memset() for better performance.
template<>
inline void PutN(MemoryBuffer& memoryBuffer, char c, size_t n) {
    std::memset(memoryBuffer.stack_.Push<char>(n), c, n * sizeof(c));
}

CEREAL_RAPIDJSON_NAMESPACE_END

#endif // CEREAL_RAPIDJSON_MEMORYBUFFER_H_
```
