## Supplemental code for "The motif composition of variable-number tandem repeats impacts gene expression": memorystream.h

```
// Tencent is pleased to support the open source community by making RapidJSON available.
// 
// Copyright (C) 2015 THL A29 Limited, a Tencent company, and Milo Yip. All rights reserved.
//
// Licensed under the MIT License (the "License"); you may not use this file except
// in compliance with the License. You may obtain a copy of the License at
//
// http://opensource.org/licenses/MIT
//
// Unless required by applicable law or agreed to in writing, software distributed 
// under the License is distributed on an "AS IS" BASIS, WITHOUT WARRANTIES OR 
// CONDITIONS OF ANY KIND, either express or implied. See the License for the 
// specific language governing permissions and limitations under the License.

#ifndef CEREAL_RAPIDJSON_MEMORYSTREAM_H_
#define CEREAL_RAPIDJSON_MEMORYSTREAM_H_

#include "stream.h"

#ifdef __clang__
CEREAL_RAPIDJSON_DIAG_PUSH
CEREAL_RAPIDJSON_DIAG_OFF(unreachable-code)
CEREAL_RAPIDJSON_DIAG_OFF(missing-noreturn)
#endif

CEREAL_RAPIDJSON_NAMESPACE_BEGIN

//! Represents an in-memory input byte stream.
/*!
    This class is mainly for being wrapped by EncodedInputStream or AutoUTFInputStream.

    It is similar to FileReadBuffer but the source is an in-memory buffer instead of a file.

    Differences between MemoryStream and StringStream:
    1. StringStream has encoding but MemoryStream is a byte stream.
    2. MemoryStream needs size of the source buffer and the buffer don't need to be null terminated. StringStream assume null-terminated string as source.
    3. MemoryStream supports Peek4() for encoding detection. StringStream is specified with an encoding so it should not have Peek4().
    \note implements Stream concept
*/
struct MemoryStream {
    typedef char Ch; // byte

    MemoryStream(const Ch *src, size_t size) : src_(src), begin_(src), end_(src + size), size_(size) {}

    Ch Peek() const { return CEREAL_RAPIDJSON_UNLIKELY(src_ == end_) ? '\0' : *src_; }
    Ch Take() { return CEREAL_RAPIDJSON_UNLIKELY(src_ == end_) ? '\0' : *src_++; }
    size_t Tell() const { return static_cast<size_t>(src_ - begin_); }

    Ch* PutBegin() { CEREAL_RAPIDJSON_ASSERT(false); return 0; }
    void Put(Ch) { CEREAL_RAPIDJSON_ASSERT(false); }
    void Flush() { CEREAL_RAPIDJSON_ASSERT(false); }
    size_t PutEnd(Ch*) { CEREAL_RAPIDJSON_ASSERT(false); return 0; }

    // For encoding detection only.
    const Ch* Peek4() const {
        return Tell() + 4 <= size_ ? src_ : 0;
    }

    const Ch* src_;     //!< Current read position.
    const Ch* begin_;   //!< Original head of the string.
    const Ch* end_;     //!< End of stream.
    size_t size_;       //!< Size of the stream.
};

CEREAL_RAPIDJSON_NAMESPACE_END

#ifdef __clang__
CEREAL_RAPIDJSON_DIAG_POP
#endif

#endif // CEREAL_RAPIDJSON_MEMORYBUFFER_H_
```
