## Supplemental code for "The motif composition of variable-number tandem repeats impacts gene expression": inttypes.h

```
// ISO C9x  compliant inttypes.h for Microsoft Visual Studio
// Based on ISO/IEC 9899:TC2 Committee draft (May 6, 2005) WG14/N1124 
// 
//  Copyright (c) 2006-2013 Alexander Chemeris
// 
// Redistribution and use in source and binary forms, with or without
// modification, are permitted provided that the following conditions are met:
// 
//   1. Redistributions of source code must retain the above copyright notice,
//      this list of conditions and the following disclaimer.
// 
//   2. Redistributions in binary form must reproduce the above copyright
//      notice, this list of conditions and the following disclaimer in the
//      documentation and/or other materials provided with the distribution.
// 
//   3. Neither the name of the product nor the names of its contributors may
//      be used to endorse or promote products derived from this software
//      without specific prior written permission.
// 
// THIS SOFTWARE IS PROVIDED BY THE AUTHOR ``AS IS'' AND ANY EXPRESS OR IMPLIED
// WARRANTIES, INCLUDING, BUT NOT LIMITED TO, THE IMPLIED WARRANTIES OF
// MERCHANTABILITY AND FITNESS FOR A PARTICULAR PURPOSE ARE DISCLAIMED. IN NO
// EVENT SHALL THE AUTHOR BE LIABLE FOR ANY DIRECT, INDIRECT, INCIDENTAL,
// SPECIAL, EXEMPLARY, OR CONSEQUENTIAL DAMAGES (INCLUDING, BUT NOT LIMITED TO,
// PROCUREMENT OF SUBSTITUTE GOODS OR SERVICES; LOSS OF USE, DATA, OR PROFITS;
// OR BUSINESS INTERRUPTION) HOWEVER CAUSED AND ON ANY THEORY OF LIABILITY, 
// WHETHER IN CONTRACT, STRICT LIABILITY, OR TORT (INCLUDING NEGLIGENCE OR
// OTHERWISE) ARISING IN ANY WAY OUT OF THE USE OF THIS SOFTWARE, EVEN IF
// ADVISED OF THE POSSIBILITY OF SUCH DAMAGE.
// 
///////////////////////////////////////////////////////////////////////////////

// The above software in this distribution may have been modified by 
// THL A29 Limited ("Tencent Modifications"). 
// All Tencent Modifications are Copyright (C) 2015 THL A29 Limited.

#ifndef _MSC_VER // [
#error "Use this header only with Microsoft Visual C++ compilers!"
#endif // _MSC_VER ]

#ifndef _MSC_INTTYPES_H_ // [
#define _MSC_INTTYPES_H_

#if _MSC_VER > 1000
#pragma once
#endif

#include "stdint.h"

// miloyip: VC supports inttypes.h since VC2013
#if _MSC_VER >= 1800
#include <inttypes.h>
#else

// 7.8 Format conversion of integer types

typedef struct {
   intmax_t quot;
   intmax_t rem;
} imaxdiv_t;

// 7.8.1 Macros for format specifiers

#if !defined(__cplusplus) || defined(__STDC_FORMAT_MACROS) // [   See footnote 185 at page 198

// The fprintf macros for signed integers are:
#define PRId8       "d"
#define PRIi8       "i"
#define PRIdLEAST8  "d"
#define PRIiLEAST8  "i"
#define PRIdFAST8   "d"
#define PRIiFAST8   "i"

#define PRId16       "hd"
#define PRIi16       "hi"
#define PRIdLEAST16  "hd"
#define PRIiLEAST16  "hi"
#define PRIdFAST16   "hd"
#define PRIiFAST16   "hi"

#define PRId32       "I32d"
#define PRIi32       "I32i"
#define PRIdLEAST32  "I32d"
#define PRIiLEAST32  "I32i"
#define PRIdFAST32   "I32d"
#define PRIiFAST32   "I32i"

#define PRId64       "I64d"
#define PRIi64       "I64i"
#define PRIdLEAST64  "I64d"
#define PRIiLEAST64  "I64i"
#define PRIdFAST64   "I64d"
#define PRIiFAST64   "I64i"

#define PRIdMAX     "I64d"
#define PRIiMAX     "I64i"

#define PRIdPTR     "Id"
#define PRIiPTR     "Ii"

// The fprintf macros for unsigned integers are:
#define PRIo8       "o"
#define PRIu8       "u"
#define PRIx8       "x"
#define PRIX8       "X"
#define PRIoLEAST8  "o"
#define PRIuLEAST8  "u"
#define PRIxLEAST8  "x"
#define PRIXLEAST8  "X"
#define PRIoFAST8   "o"
#define PRIuFAST8   "u"
#define PRIxFAST8   "x"
#define PRIXFAST8   "X"

#define PRIo16       "ho"
#define PRIu16       "hu"
#define PRIx16       "hx"
#define PRIX16       "hX"
#define PRIoLEAST16  "ho"
#define PRIuLEAST16  "hu"
#define PRIxLEAST16  "hx"
#define PRIXLEAST16  "hX"
#define PRIoFAST16   "ho"
#define PRIuFAST16   "hu"
#define PRIxFAST16   "hx"
#define PRIXFAST16   "hX"

#define PRIo32       "I32o"
#define PRIu32       "I32u"
#define PRIx32       "I32x"
#define PRIX32       "I32X"
#define PRIoLEAST32  "I32o"
#define PRIuLEAST32  "I32u"
#define PRIxLEAST32  "I32x"
#define PRIXLEAST32  "I32X"
#define PRIoFAST32   "I32o"
#define PRIuFAST32   "I32u"
#define PRIxFAST32   "I32x"
#define PRIXFAST32   "I32X"

#define PRIo64       "I64o"
#define PRIu64       "I64u"
#define PRIx64       "I64x"
#define PRIX64       "I64X"
#define PRIoLEAST64  "I64o"
#define PRIuLEAST64  "I64u"
#define PRIxLEAST64  "I64x"
#define PRIXLEAST64  "I64X"
#define PRIoFAST64   "I64o"
#define PRIuFAST64   "I64u"
#define PRIxFAST64   "I64x"
#define PRIXFAST64   "I64X"

#define PRIoMAX     "I64o"
#define PRIuMAX     "I64u"
#define PRIxMAX     "I64x"
#define PRIXMAX     "I64X"

#define PRIoPTR     "Io"
#define PRIuPTR     "Iu"
#define PRIxPTR     "Ix"
#define PRIXPTR     "IX"

// The fscanf macros for signed integers are:
#define SCNd8       "d"
#define SCNi8       "i"
#define SCNdLEAST8  "d"
#define SCNiLEAST8  "i"
#define SCNdFAST8   "d"
#define SCNiFAST8   "i"

#define SCNd16       "hd"
#define SCNi16       "hi"
#define SCNdLEAST16  "hd"
#define SCNiLEAST16  "hi"
#define SCNdFAST16   "hd"
#define SCNiFAST16   "hi"

#define SCNd32       "ld"
#define SCNi32       "li"
#define SCNdLEAST32  "ld"
#define SCNiLEAST32  "li"
#define SCNdFAST32   "ld"
#define SCNiFAST32   "li"

#define SCNd64       "I64d"
#define SCNi64       "I64i"
#define SCNdLEAST64  "I64d"
#define SCNiLEAST64  "I64i"
#define SCNdFAST64   "I64d"
#define SCNiFAST64   "I64i"

#define SCNdMAX     "I64d"
#define SCNiMAX     "I64i"

#ifdef _WIN64 // [
#  define SCNdPTR     "I64d"
#  define SCNiPTR     "I64i"
#else  // _WIN64 ][
#  define SCNdPTR     "ld"
#  define SCNiPTR     "li"
#endif  // _WIN64 ]

// The fscanf macros for unsigned integers are:
#define SCNo8       "o"
#define SCNu8       "u"
#define SCNx8       "x"
#define SCNX8       "X"
#define SCNoLEAST8  "o"
#define SCNuLEAST8  "u"
#define SCNxLEAST8  "x"
#define SCNXLEAST8  "X"
#define SCNoFAST8   "o"
#define SCNuFAST8   "u"
#define SCNxFAST8   "x"
#define SCNXFAST8   "X"

#define SCNo16       "ho"
#define SCNu16       "hu"
#define SCNx16       "hx"
#define SCNX16       "hX"
#define SCNoLEAST16  "ho"
#define SCNuLEAST16  "hu"
#define SCNxLEAST16  "hx"
#define SCNXLEAST16  "hX"
#define SCNoFAST16   "ho"
#define SCNuFAST16   "hu"
#define SCNxFAST16   "hx"
#define SCNXFAST16   "hX"

#define SCNo32       "lo"
#define SCNu32       "lu"
#define SCNx32       "lx"
#define SCNX32       "lX"
#define SCNoLEAST32  "lo"
#define SCNuLEAST32  "lu"
#define SCNxLEAST32  "lx"
#define SCNXLEAST32  "lX"
#define SCNoFAST32   "lo"
#define SCNuFAST32   "lu"
#define SCNxFAST32   "lx"
#define SCNXFAST32   "lX"

#define SCNo64       "I64o"
#define SCNu64       "I64u"
#define SCNx64       "I64x"
#define SCNX64       "I64X"
#define SCNoLEAST64  "I64o"
#define SCNuLEAST64  "I64u"
#define SCNxLEAST64  "I64x"
#define SCNXLEAST64  "I64X"
#define SCNoFAST64   "I64o"
#define SCNuFAST64   "I64u"
#define SCNxFAST64   "I64x"
#define SCNXFAST64   "I64X"

#define SCNoMAX     "I64o"
#define SCNuMAX     "I64u"
#define SCNxMAX     "I64x"
#define SCNXMAX     "I64X"

#ifdef _WIN64 // [
#  define SCNoPTR     "I64o"
#  define SCNuPTR     "I64u"
#  define SCNxPTR     "I64x"
#  define SCNXPTR     "I64X"
#else  // _WIN64 ][
#  define SCNoPTR     "lo"
#  define SCNuPTR     "lu"
#  define SCNxPTR     "lx"
#  define SCNXPTR     "lX"
#endif  // _WIN64 ]

#endif // __STDC_FORMAT_MACROS ]

// 7.8.2 Functions for greatest-width integer types

// 7.8.2.1 The imaxabs function
#define imaxabs _abs64

// 7.8.2.2 The imaxdiv function

// This is modified version of div() function from Microsoft's div.c found
// in %MSVC.NET%\crt\src\div.c
#ifdef STATIC_IMAXDIV // [
static
#else // STATIC_IMAXDIV ][
_inline
#endif // STATIC_IMAXDIV ]
imaxdiv_t __cdecl imaxdiv(intmax_t numer, intmax_t denom)
{
   imaxdiv_t result;

   result.quot = numer / denom;
   result.rem = numer % denom;

   if (numer < 0 && result.rem > 0) {
      // did division wrong; must fix up
      ++result.quot;
      result.rem -= denom;
   }

   return result;
}

// 7.8.2.3 The strtoimax and strtoumax functions
#define strtoimax _strtoi64
#define strtoumax _strtoui64

// 7.8.2.4 The wcstoimax and wcstoumax functions
#define wcstoimax _wcstoi64
#define wcstoumax _wcstoui64

#endif // _MSC_VER >= 1800

#endif // _MSC_INTTYPES_H_ ]
```
