## Supplemental code for "The motif composition of variable-number tandem repeats impacts gene expression": stdint.h

```
// ISO C9x  compliant stdint.h for Microsoft Visual Studio
// Based on ISO/IEC 9899:TC2 Committee draft (May 6, 2005) WG14/N1124 
// 
//  Copyright (c) 2006-2013 Alexander Chemeris
// 
// Redistribution and use in source and binary forms, with or without
// modification, are permitted provided that the following conditions are met:
// 
//   1. Redistributions of source code must retain the above copyright notice,
//      this list of conditions and the following disclaimer.
// 
//   2. Redistributions in binary form must reproduce the above copyright
//      notice, this list of conditions and the following disclaimer in the
//      documentation and/or other materials provided with the distribution.
// 
//   3. Neither the name of the product nor the names of its contributors may
//      be used to endorse or promote products derived from this software
//      without specific prior written permission.
// 
// THIS SOFTWARE IS PROVIDED BY THE AUTHOR ``AS IS'' AND ANY EXPRESS OR IMPLIED
// WARRANTIES, INCLUDING, BUT NOT LIMITED TO, THE IMPLIED WARRANTIES OF
// MERCHANTABILITY AND FITNESS FOR A PARTICULAR PURPOSE ARE DISCLAIMED. IN NO
// EVENT SHALL THE AUTHOR BE LIABLE FOR ANY DIRECT, INDIRECT, INCIDENTAL,
// SPECIAL, EXEMPLARY, OR CONSEQUENTIAL DAMAGES (INCLUDING, BUT NOT LIMITED TO,
// PROCUREMENT OF SUBSTITUTE GOODS OR SERVICES; LOSS OF USE, DATA, OR PROFITS;
// OR BUSINESS INTERRUPTION) HOWEVER CAUSED AND ON ANY THEORY OF LIABILITY, 
// WHETHER IN CONTRACT, STRICT LIABILITY, OR TORT (INCLUDING NEGLIGENCE OR
// OTHERWISE) ARISING IN ANY WAY OUT OF THE USE OF THIS SOFTWARE, EVEN IF
// ADVISED OF THE POSSIBILITY OF SUCH DAMAGE.
// 
///////////////////////////////////////////////////////////////////////////////

// The above software in this distribution may have been modified by 
// THL A29 Limited ("Tencent Modifications"). 
// All Tencent Modifications are Copyright (C) 2015 THL A29 Limited.

#ifndef _MSC_VER // [
#error "Use this header only with Microsoft Visual C++ compilers!"
#endif // _MSC_VER ]

#ifndef _MSC_STDINT_H_ // [
#define _MSC_STDINT_H_

#if _MSC_VER > 1000
#pragma once
#endif

// miloyip: Originally Visual Studio 2010 uses its own stdint.h. However it generates warning with INT64_C(), so change to use this file for vs2010.
#if _MSC_VER >= 1600 // [
#include <stdint.h>

#if !defined(__cplusplus) || defined(__STDC_CONSTANT_MACROS) // [   See footnote 224 at page 260

#undef INT8_C
#undef INT16_C
#undef INT32_C
#undef INT64_C
#undef UINT8_C
#undef UINT16_C
#undef UINT32_C
#undef UINT64_C

// 7.18.4.1 Macros for minimum-width integer constants

#define INT8_C(val)  val##i8
#define INT16_C(val) val##i16
#define INT32_C(val) val##i32
#define INT64_C(val) val##i64

#define UINT8_C(val)  val##ui8
#define UINT16_C(val) val##ui16
#define UINT32_C(val) val##ui32
#define UINT64_C(val) val##ui64

// 7.18.4.2 Macros for greatest-width integer constants
// These #ifndef's are needed to prevent collisions with <boost/cstdint.hpp>.
// Check out Issue 9 for the details.
#ifndef INTMAX_C //   [
#  define INTMAX_C   INT64_C
#endif // INTMAX_C    ]
#ifndef UINTMAX_C //  [
#  define UINTMAX_C  UINT64_C
#endif // UINTMAX_C   ]

#endif // __STDC_CONSTANT_MACROS ]

#else // ] _MSC_VER >= 1700 [

#include <limits.h>

// For Visual Studio 6 in C++ mode and for many Visual Studio versions when
// compiling for ARM we have to wrap <wchar.h> include with 'extern "C++" {}'
// or compiler would give many errors like this:
//   error C2733: second C linkage of overloaded function 'wmemchr' not allowed
#if defined(__cplusplus) && !defined(_M_ARM)
extern "C" {
#endif
#  include <wchar.h>
#if defined(__cplusplus) && !defined(_M_ARM)
}
#endif

// Define _W64 macros to mark types changing their size, like intptr_t.
#ifndef _W64
#  if !defined(__midl) && (defined(_X86_) || defined(_M_IX86)) && _MSC_VER >= 1300
#     define _W64 __w64
#  else
#     define _W64
#  endif
#endif


// 7.18.1 Integer types

// 7.18.1.1 Exact-width integer types

// Visual Studio 6 and Embedded Visual C++ 4 doesn't
// realize that, e.g. char has the same size as __int8
// so we give up on __intX for them.
#if (_MSC_VER < 1300)
   typedef signed char       int8_t;
   typedef signed short      int16_t;
   typedef signed int        int32_t;
   typedef unsigned char     uint8_t;
   typedef unsigned short    uint16_t;
   typedef unsigned int      uint32_t;
#else
   typedef signed __int8     int8_t;
   typedef signed __int16    int16_t;
   typedef signed __int32    int32_t;
   typedef unsigned __int8   uint8_t;
   typedef unsigned __int16  uint16_t;
   typedef unsigned __int32  uint32_t;
#endif
typedef signed __int64       int64_t;
typedef unsigned __int64     uint64_t;


// 7.18.1.2 Minimum-width integer types
typedef int8_t    int_least8_t;
typedef int16_t   int_least16_t;
typedef int32_t   int_least32_t;
typedef int64_t   int_least64_t;
typedef uint8_t   uint_least8_t;
typedef uint16_t  uint_least16_t;
typedef uint32_t  uint_least32_t;
typedef uint64_t  uint_least64_t;

// 7.18.1.3 Fastest minimum-width integer types
typedef int8_t    int_fast8_t;
typedef int16_t   int_fast16_t;
typedef int32_t   int_fast32_t;
typedef int64_t   int_fast64_t;
typedef uint8_t   uint_fast8_t;
typedef uint16_t  uint_fast16_t;
typedef uint32_t  uint_fast32_t;
typedef uint64_t  uint_fast64_t;

// 7.18.1.4 Integer types capable of holding object pointers
#ifdef _WIN64 // [
   typedef signed __int64    intptr_t;
   typedef unsigned __int64  uintptr_t;
#else // _WIN64 ][
   typedef _W64 signed int   intptr_t;
   typedef _W64 unsigned int uintptr_t;
#endif // _WIN64 ]

// 7.18.1.5 Greatest-width integer types
typedef int64_t   intmax_t;
typedef uint64_t  uintmax_t;


// 7.18.2 Limits of specified-width integer types

#if !defined(__cplusplus) || defined(__STDC_LIMIT_MACROS) // [   See footnote 220 at page 257 and footnote 221 at page 259

// 7.18.2.1 Limits of exact-width integer types
#define INT8_MIN     ((int8_t)_I8_MIN)
#define INT8_MAX     _I8_MAX
#define INT16_MIN    ((int16_t)_I16_MIN)
#define INT16_MAX    _I16_MAX
#define INT32_MIN    ((int32_t)_I32_MIN)
#define INT32_MAX    _I32_MAX
#define INT64_MIN    ((int64_t)_I64_MIN)
#define INT64_MAX    _I64_MAX
#define UINT8_MAX    _UI8_MAX
#define UINT16_MAX   _UI16_MAX
#define UINT32_MAX   _UI32_MAX
#define UINT64_MAX   _UI64_MAX

// 7.18.2.2 Limits of minimum-width integer types
#define INT_LEAST8_MIN    INT8_MIN
#define INT_LEAST8_MAX    INT8_MAX
#define INT_LEAST16_MIN   INT16_MIN
#define INT_LEAST16_MAX   INT16_MAX
#define INT_LEAST32_MIN   INT32_MIN
#define INT_LEAST32_MAX   INT32_MAX
#define INT_LEAST64_MIN   INT64_MIN
#define INT_LEAST64_MAX   INT64_MAX
#define UINT_LEAST8_MAX   UINT8_MAX
#define UINT_LEAST16_MAX  UINT16_MAX
#define UINT_LEAST32_MAX  UINT32_MAX
#define UINT_LEAST64_MAX  UINT64_MAX

// 7.18.2.3 Limits of fastest minimum-width integer types
#define INT_FAST8_MIN    INT8_MIN
#define INT_FAST8_MAX    INT8_MAX
#define INT_FAST16_MIN   INT16_MIN
#define INT_FAST16_MAX   INT16_MAX
#define INT_FAST32_MIN   INT32_MIN
#define INT_FAST32_MAX   INT32_MAX
#define INT_FAST64_MIN   INT64_MIN
#define INT_FAST64_MAX   INT64_MAX
#define UINT_FAST8_MAX   UINT8_MAX
#define UINT_FAST16_MAX  UINT16_MAX
#define UINT_FAST32_MAX  UINT32_MAX
#define UINT_FAST64_MAX  UINT64_MAX

// 7.18.2.4 Limits of integer types capable of holding object pointers
#ifdef _WIN64 // [
#  define INTPTR_MIN   INT64_MIN
#  define INTPTR_MAX   INT64_MAX
#  define UINTPTR_MAX  UINT64_MAX
#else // _WIN64 ][
#  define INTPTR_MIN   INT32_MIN
#  define INTPTR_MAX   INT32_MAX
#  define UINTPTR_MAX  UINT32_MAX
#endif // _WIN64 ]

// 7.18.2.5 Limits of greatest-width integer types
#define INTMAX_MIN   INT64_MIN
#define INTMAX_MAX   INT64_MAX
#define UINTMAX_MAX  UINT64_MAX

// 7.18.3 Limits of other integer types

#ifdef _WIN64 // [
#  define PTRDIFF_MIN  _I64_MIN
#  define PTRDIFF_MAX  _I64_MAX
#else  // _WIN64 ][
#  define PTRDIFF_MIN  _I32_MIN
#  define PTRDIFF_MAX  _I32_MAX
#endif  // _WIN64 ]

#define SIG_ATOMIC_MIN  INT_MIN
#define SIG_ATOMIC_MAX  INT_MAX

#ifndef SIZE_MAX // [
#  ifdef _WIN64 // [
#     define SIZE_MAX  _UI64_MAX
#  else // _WIN64 ][
#     define SIZE_MAX  _UI32_MAX
#  endif // _WIN64 ]
#endif // SIZE_MAX ]

// WCHAR_MIN and WCHAR_MAX are also defined in <wchar.h>
#ifndef WCHAR_MIN // [
#  define WCHAR_MIN  0
#endif  // WCHAR_MIN ]
#ifndef WCHAR_MAX // [
#  define WCHAR_MAX  _UI16_MAX
#endif  // WCHAR_MAX ]

#define WINT_MIN  0
#define WINT_MAX  _UI16_MAX

#endif // __STDC_LIMIT_MACROS ]


// 7.18.4 Limits of other integer types

#if !defined(__cplusplus) || defined(__STDC_CONSTANT_MACROS) // [   See footnote 224 at page 260

// 7.18.4.1 Macros for minimum-width integer constants

#define INT8_C(val)  val##i8
#define INT16_C(val) val##i16
#define INT32_C(val) val##i32
#define INT64_C(val) val##i64

#define UINT8_C(val)  val##ui8
#define UINT16_C(val) val##ui16
#define UINT32_C(val) val##ui32
#define UINT64_C(val) val##ui64

// 7.18.4.2 Macros for greatest-width integer constants
// These #ifndef's are needed to prevent collisions with <boost/cstdint.hpp>.
// Check out Issue 9 for the details.
#ifndef INTMAX_C //   [
#  define INTMAX_C   INT64_C
#endif // INTMAX_C    ]
#ifndef UINTMAX_C //  [
#  define UINTMAX_C  UINT64_C
#endif // UINTMAX_C   ]

#endif // __STDC_CONSTANT_MACROS ]

#endif // _MSC_VER >= 1600 ]

#endif // _MSC_STDINT_H_ ]
```
