## Supplemental code for "The motif composition of variable-number tandem repeats impacts gene expression": ostreamwrapper.h

```
// Tencent is pleased to support the open source community by making RapidJSON available.
// 
// Copyright (C) 2015 THL A29 Limited, a Tencent company, and Milo Yip. All rights reserved.
//
// Licensed under the MIT License (the "License"); you may not use this file except
// in compliance with the License. You may obtain a copy of the License at
//
// http://opensource.org/licenses/MIT
//
// Unless required by applicable law or agreed to in writing, software distributed 
// under the License is distributed on an "AS IS" BASIS, WITHOUT WARRANTIES OR 
// CONDITIONS OF ANY KIND, either express or implied. See the License for the 
// specific language governing permissions and limitations under the License.

#ifndef CEREAL_RAPIDJSON_OSTREAMWRAPPER_H_
#define CEREAL_RAPIDJSON_OSTREAMWRAPPER_H_

#include "stream.h"
#include <iosfwd>

#ifdef __clang__
CEREAL_RAPIDJSON_DIAG_PUSH
CEREAL_RAPIDJSON_DIAG_OFF(padded)
#endif

CEREAL_RAPIDJSON_NAMESPACE_BEGIN

//! Wrapper of \c std::basic_ostream into RapidJSON's Stream concept.
/*!
    The classes can be wrapped including but not limited to:

    - \c std::ostringstream
    - \c std::stringstream
    - \c std::wpstringstream
    - \c std::wstringstream
    - \c std::ifstream
    - \c std::fstream
    - \c std::wofstream
    - \c std::wfstream

    \tparam StreamType Class derived from \c std::basic_ostream.
*/
   
template <typename StreamType>
class BasicOStreamWrapper {
public:
    typedef typename StreamType::char_type Ch;
    BasicOStreamWrapper(StreamType& stream) : stream_(stream) {}

    void Put(Ch c) {
        stream_.put(c);
    }

    void Flush() {
        stream_.flush();
    }

    // Not implemented
    char Peek() const { CEREAL_RAPIDJSON_ASSERT(false); return 0; }
    char Take() { CEREAL_RAPIDJSON_ASSERT(false); return 0; }
    size_t Tell() const { CEREAL_RAPIDJSON_ASSERT(false); return 0; }
    char* PutBegin() { CEREAL_RAPIDJSON_ASSERT(false); return 0; }
    size_t PutEnd(char*) { CEREAL_RAPIDJSON_ASSERT(false); return 0; }

private:
    BasicOStreamWrapper(const BasicOStreamWrapper&);
    BasicOStreamWrapper& operator=(const BasicOStreamWrapper&);

    StreamType& stream_;
};

typedef BasicOStreamWrapper<std::ostream> OStreamWrapper;
typedef BasicOStreamWrapper<std::wostream> WOStreamWrapper;

#ifdef __clang__
CEREAL_RAPIDJSON_DIAG_POP
#endif

CEREAL_RAPIDJSON_NAMESPACE_END

#endif // CEREAL_RAPIDJSON_OSTREAMWRAPPER_H_
```
