## Supplemental code for "The motif composition of variable-number tandem repeats impacts gene expression": pointer.h

```
// Tencent is pleased to support the open source community by making RapidJSON available.
// 
// Copyright (C) 2015 THL A29 Limited, a Tencent company, and Milo Yip. All rights reserved.
//
// Licensed under the MIT License (the "License"); you may not use this file except
// in compliance with the License. You may obtain a copy of the License at
//
// http://opensource.org/licenses/MIT
//
// Unless required by applicable law or agreed to in writing, software distributed 
// under the License is distributed on an "AS IS" BASIS, WITHOUT WARRANTIES OR 
// CONDITIONS OF ANY KIND, either express or implied. See the License for the 
// specific language governing permissions and limitations under the License.

#ifndef CEREAL_RAPIDJSON_POINTER_H_
#define CEREAL_RAPIDJSON_POINTER_H_

#include "document.h"
#include "internal/itoa.h"

#ifdef __clang__
CEREAL_RAPIDJSON_DIAG_PUSH
CEREAL_RAPIDJSON_DIAG_OFF(switch-enum)
#elif defined(_MSC_VER)
CEREAL_RAPIDJSON_DIAG_PUSH
CEREAL_RAPIDJSON_DIAG_OFF(4512) // assignment operator could not be generated
#endif

CEREAL_RAPIDJSON_NAMESPACE_BEGIN

static const SizeType kPointerInvalidIndex = ~SizeType(0);  //!< Represents an invalid index in GenericPointer::Token

//! Error code of parsing.
/*! \ingroup CEREAL_RAPIDJSON_ERRORS
    \see GenericPointer::GenericPointer, GenericPointer::GetParseErrorCode
*/
enum PointerParseErrorCode {
    kPointerParseErrorNone = 0,                     //!< The parse is successful

    kPointerParseErrorTokenMustBeginWithSolidus,    //!< A token must begin with a '/'
    kPointerParseErrorInvalidEscape,                //!< Invalid escape
    kPointerParseErrorInvalidPercentEncoding,       //!< Invalid percent encoding in URI fragment
    kPointerParseErrorCharacterMustPercentEncode    //!< A character must percent encoded in URI fragment
};

///////////////////////////////////////////////////////////////////////////////
// GenericPointer

//! Represents a JSON Pointer. Use Pointer for UTF8 encoding and default allocator.
/*!
    This class implements RFC 6901 "JavaScript Object Notation (JSON) Pointer" 
    (https://tools.ietf.org/html/rfc6901).

    A JSON pointer is for identifying a specific value in a JSON document
    (GenericDocument). It can simplify coding of DOM tree manipulation, because it
    can access multiple-level depth of DOM tree with single API call.

    After it parses a string representation (e.g. "/foo/0" or URI fragment 
    representation (e.g. "#/foo/0") into its internal representation (tokens),
    it can be used to resolve a specific value in multiple documents, or sub-tree 
    of documents.

    Contrary to GenericValue, Pointer can be copy constructed and copy assigned.
    Apart from assignment, a Pointer cannot be modified after construction.

    Although Pointer is very convenient, please aware that constructing Pointer
    involves parsing and dynamic memory allocation. A special constructor with user-
    supplied tokens eliminates these.

    GenericPointer depends on GenericDocument and GenericValue.
    
    \tparam ValueType The value type of the DOM tree. E.g. GenericValue<UTF8<> >
    \tparam Allocator The allocator type for allocating memory for internal representation.
    
    \note GenericPointer uses same encoding of ValueType.
    However, Allocator of GenericPointer is independent of Allocator of Value.
*/
template <typename ValueType, typename Allocator = CrtAllocator>
class GenericPointer {
public:
    typedef typename ValueType::EncodingType EncodingType;  //!< Encoding type from Value
    typedef typename ValueType::Ch Ch;                      //!< Character type from Value

    //! A token is the basic units of internal representation.
    /*!
        A JSON pointer string representation "/foo/123" is parsed to two tokens: 
        "foo" and 123. 123 will be represented in both numeric form and string form.
        They are resolved according to the actual value type (object or array).

        For token that are not numbers, or the numeric value is out of bound
        (greater than limits of SizeType), they are only treated as string form
        (i.e. the token's index will be equal to kPointerInvalidIndex).

        This struct is public so that user can create a Pointer without parsing and 
        allocation, using a special constructor.
    */
    struct Token {
        const Ch* name;             //!< Name of the token. It has null character at the end but it can contain null character.
        SizeType length;            //!< Length of the name.
        SizeType index;             //!< A valid array index, if it is not equal to kPointerInvalidIndex.
    };

    //!@name Constructors and destructor.
    //@{

    //! Default constructor.
    GenericPointer(Allocator* allocator = 0) : allocator_(allocator), ownAllocator_(), nameBuffer_(), tokens_(), tokenCount_(), parseErrorOffset_(), parseErrorCode_(kPointerParseErrorNone) {}

    //! Constructor that parses a string or URI fragment representation.
    /*!
        \param source A null-terminated, string or URI fragment representation of JSON pointer.
        \param allocator User supplied allocator for this pointer. If no allocator is provided, it creates a self-owned one.
    */
    explicit GenericPointer(const Ch* source, Allocator* allocator = 0) : allocator_(allocator), ownAllocator_(), nameBuffer_(), tokens_(), tokenCount_(), parseErrorOffset_(), parseErrorCode_(kPointerParseErrorNone) {
        Parse(source, internal::StrLen(source));
    }

#if CEREAL_RAPIDJSON_HAS_STDSTRING
    //! Constructor that parses a string or URI fragment representation.
    /*!
        \param source A string or URI fragment representation of JSON pointer.
        \param allocator User supplied allocator for this pointer. If no allocator is provided, it creates a self-owned one.
        \note Requires the definition of the preprocessor symbol \ref CEREAL_RAPIDJSON_HAS_STDSTRING.
    */
    explicit GenericPointer(const std::basic_string<Ch>& source, Allocator* allocator = 0) : allocator_(allocator), ownAllocator_(), nameBuffer_(), tokens_(), tokenCount_(), parseErrorOffset_(), parseErrorCode_(kPointerParseErrorNone) {
        Parse(source.c_str(), source.size());
    }
#endif

    //! Constructor that parses a string or URI fragment representation, with length of the source string.
    /*!
        \param source A string or URI fragment representation of JSON pointer.
        \param length Length of source.
        \param allocator User supplied allocator for this pointer. If no allocator is provided, it creates a self-owned one.
        \note Slightly faster than the overload without length.
    */
    GenericPointer(const Ch* source, size_t length, Allocator* allocator = 0) : allocator_(allocator), ownAllocator_(), nameBuffer_(), tokens_(), tokenCount_(), parseErrorOffset_(), parseErrorCode_(kPointerParseErrorNone) {
        Parse(source, length);
    }

    //! Constructor with user-supplied tokens.
    /*!
        This constructor let user supplies const array of tokens.
        This prevents the parsing process and eliminates allocation.
        This is preferred for memory constrained environments.

        \param tokens An constant array of tokens representing the JSON pointer.
        \param tokenCount Number of tokens.

        \b Example
        \code
        #define NAME(s) { s, sizeof(s) / sizeof(s[0]) - 1, kPointerInvalidIndex }
        #define INDEX(i) { #i, sizeof(#i) - 1, i }

        static const Pointer::Token kTokens[] = { NAME("foo"), INDEX(123) };
        static const Pointer p(kTokens, sizeof(kTokens) / sizeof(kTokens[0]));
        // Equivalent to static const Pointer p("/foo/123");

        #undef NAME
        #undef INDEX
        \endcode
    */
    GenericPointer(const Token* tokens, size_t tokenCount) : allocator_(), ownAllocator_(), nameBuffer_(), tokens_(const_cast<Token*>(tokens)), tokenCount_(tokenCount), parseErrorOffset_(), parseErrorCode_(kPointerParseErrorNone) {}

    //! Copy constructor.
    GenericPointer(const GenericPointer& rhs) : allocator_(rhs.allocator_), ownAllocator_(), nameBuffer_(), tokens_(), tokenCount_(), parseErrorOffset_(), parseErrorCode_(kPointerParseErrorNone) {
        *this = rhs;
    }

    //! Copy constructor.
    GenericPointer(const GenericPointer& rhs, Allocator* allocator) : allocator_(allocator), ownAllocator_(), nameBuffer_(), tokens_(), tokenCount_(), parseErrorOffset_(), parseErrorCode_(kPointerParseErrorNone) {
        *this = rhs;
    }

    //! Destructor.
    ~GenericPointer() {
        if (nameBuffer_)    // If user-supplied tokens constructor is used, nameBuffer_ is nullptr and tokens_ are not deallocated.
            Allocator::Free(tokens_);
        CEREAL_RAPIDJSON_DELETE(ownAllocator_);
    }

    //! Assignment operator.
    GenericPointer& operator=(const GenericPointer& rhs) {
        if (this != &rhs) {
            // Do not delete ownAllcator
            if (nameBuffer_)
                Allocator::Free(tokens_);

            tokenCount_ = rhs.tokenCount_;
            parseErrorOffset_ = rhs.parseErrorOffset_;
            parseErrorCode_ = rhs.parseErrorCode_;

            if (rhs.nameBuffer_)
                CopyFromRaw(rhs); // Normally parsed tokens.
            else {
                tokens_ = rhs.tokens_; // User supplied const tokens.
                nameBuffer_ = 0;
            }
        }
        return *this;
    }

    //! Swap the content of this pointer with an other.
    /*!
        \param other The pointer to swap with.
        \note Constant complexity.
    */
    GenericPointer& Swap(GenericPointer& other) CEREAL_RAPIDJSON_NOEXCEPT {
        internal::Swap(allocator_, other.allocator_);
        internal::Swap(ownAllocator_, other.ownAllocator_);
        internal::Swap(nameBuffer_, other.nameBuffer_);
        internal::Swap(tokens_, other.tokens_);
        internal::Swap(tokenCount_, other.tokenCount_);
        internal::Swap(parseErrorOffset_, other.parseErrorOffset_);
        internal::Swap(parseErrorCode_, other.parseErrorCode_);
        return *this;
    }

    //! free-standing swap function helper
    /*!
        Helper function to enable support for common swap implementation pattern based on \c std::swap:
        \code
        void swap(MyClass& a, MyClass& b) {
            using std::swap;
            swap(a.pointer, b.pointer);
            // ...
        }
        \endcode
        \see Swap()
     */
    friend inline void swap(GenericPointer& a, GenericPointer& b) CEREAL_RAPIDJSON_NOEXCEPT { a.Swap(b); }

    //@}

    //!@name Append token
    //@{

    //! Append a token and return a new Pointer
    /*!
        \param token Token to be appended.
        \param allocator Allocator for the newly return Pointer.
        \return A new Pointer with appended token.
    */
    GenericPointer Append(const Token& token, Allocator* allocator = 0) const {
        GenericPointer r;
        r.allocator_ = allocator;
        Ch *p = r.CopyFromRaw(*this, 1, token.length + 1);
        std::memcpy(p, token.name, (token.length + 1) * sizeof(Ch));
        r.tokens_[tokenCount_].name = p;
        r.tokens_[tokenCount_].length = token.length;
        r.tokens_[tokenCount_].index = token.index;
        return r;
    }

    //! Append a name token with length, and return a new Pointer
    /*!
        \param name Name to be appended.
        \param length Length of name.
        \param allocator Allocator for the newly return Pointer.
        \return A new Pointer with appended token.
    */
    GenericPointer Append(const Ch* name, SizeType length, Allocator* allocator = 0) const {
        Token token = { name, length, kPointerInvalidIndex };
        return Append(token, allocator);
    }

    //! Append a name token without length, and return a new Pointer
    /*!
        \param name Name (const Ch*) to be appended.
        \param allocator Allocator for the newly return Pointer.
        \return A new Pointer with appended token.
    */
    template <typename T>
    CEREAL_RAPIDJSON_DISABLEIF_RETURN((internal::NotExpr<internal::IsSame<typename internal::RemoveConst<T>::Type, Ch> >), (GenericPointer))
    Append(T* name, Allocator* allocator = 0) const {
        return Append(name, internal::StrLen(name), allocator);
    }

#if CEREAL_RAPIDJSON_HAS_STDSTRING
    //! Append a name token, and return a new Pointer
    /*!
        \param name Name to be appended.
        \param allocator Allocator for the newly return Pointer.
        \return A new Pointer with appended token.
    */
    GenericPointer Append(const std::basic_string<Ch>& name, Allocator* allocator = 0) const {
        return Append(name.c_str(), static_cast<SizeType>(name.size()), allocator);
    }
#endif

    //! Append a index token, and return a new Pointer
    /*!
        \param index Index to be appended.
        \param allocator Allocator for the newly return Pointer.
        \return A new Pointer with appended token.
    */
    GenericPointer Append(SizeType index, Allocator* allocator = 0) const {
        char buffer[21];
        char* end = sizeof(SizeType) == 4 ? internal::u32toa(index, buffer) : internal::u64toa(index, buffer);
        SizeType length = static_cast<SizeType>(end - buffer);
        buffer[length] = '\0';

        if (sizeof(Ch) == 1) {
            Token token = { reinterpret_cast<Ch*>(buffer), length, index };
            return Append(token, allocator);
        }
        else {
            Ch name[21];
            for (size_t i = 0; i <= length; i++)
                name[i] = static_cast<Ch>(buffer[i]);
            Token token = { name, length, index };
            return Append(token, allocator);
        }
    }

    //! Append a token by value, and return a new Pointer
    /*!
        \param token token to be appended.
        \param allocator Allocator for the newly return Pointer.
        \return A new Pointer with appended token.
    */
    GenericPointer Append(const ValueType& token, Allocator* allocator = 0) const {
        if (token.IsString())
            return Append(token.GetString(), token.GetStringLength(), allocator);
        else {
            CEREAL_RAPIDJSON_ASSERT(token.IsUint64());
            CEREAL_RAPIDJSON_ASSERT(token.GetUint64() <= SizeType(~0));
            return Append(static_cast<SizeType>(token.GetUint64()), allocator);
        }
    }

    //!@name Handling Parse Error
    //@{

    //! Check whether this is a valid pointer.
    bool IsValid() const { return parseErrorCode_ == kPointerParseErrorNone; }

    //! Get the parsing error offset in code unit.
    size_t GetParseErrorOffset() const { return parseErrorOffset_; }

    //! Get the parsing error code.
    PointerParseErrorCode GetParseErrorCode() const { return parseErrorCode_; }

    //@}

    //! Get the allocator of this pointer.
    Allocator& GetAllocator() { return *allocator_; }

    //!@name Tokens
    //@{

    //! Get the token array (const version only).
    const Token* GetTokens() const { return tokens_; }

    //! Get the number of tokens.
    size_t GetTokenCount() const { return tokenCount_; }

    //@}

    //!@name Equality/inequality operators
    //@{

    //! Equality operator.
    /*!
        \note When any pointers are invalid, always returns false.
    */
    bool operator==(const GenericPointer& rhs) const {
        if (!IsValid() || !rhs.IsValid() || tokenCount_ != rhs.tokenCount_)
            return false;

        for (size_t i = 0; i < tokenCount_; i++) {
            if (tokens_[i].index != rhs.tokens_[i].index ||
                tokens_[i].length != rhs.tokens_[i].length || 
                (tokens_[i].length != 0 && std::memcmp(tokens_[i].name, rhs.tokens_[i].name, sizeof(Ch)* tokens_[i].length) != 0))
            {
                return false;
            }
        }

        return true;
    }

    //! Inequality operator.
    /*!
        \note When any pointers are invalid, always returns true.
    */
    bool operator!=(const GenericPointer& rhs) const { return !(*this == rhs); }

    //! Less than operator.
    /*!
        \note Invalid pointers are always greater than valid ones.
    */
    bool operator<(const GenericPointer& rhs) const {
        if (!IsValid())
            return false;
        if (!rhs.IsValid())
            return true;

        if (tokenCount_ != rhs.tokenCount_)
            return tokenCount_ < rhs.tokenCount_;

        for (size_t i = 0; i < tokenCount_; i++) {
            if (tokens_[i].index != rhs.tokens_[i].index)
                return tokens_[i].index < rhs.tokens_[i].index;

            if (tokens_[i].length != rhs.tokens_[i].length)
                return tokens_[i].length < rhs.tokens_[i].length;

            if (int cmp = std::memcmp(tokens_[i].name, rhs.tokens_[i].name, sizeof(Ch) * tokens_[i].length))
                return cmp < 0;
        }

        return false;
    }

    //@}

    //!@name Stringify
    //@{

    //! Stringify the pointer into string representation.
    /*!
        \tparam OutputStream Type of output stream.
        \param os The output stream.
    */
    template<typename OutputStream>
    bool Stringify(OutputStream& os) const {
        return Stringify<false, OutputStream>(os);
    }

    //! Stringify the pointer into URI fragment representation.
    /*!
        \tparam OutputStream Type of output stream.
        \param os The output stream.
    */
    template<typename OutputStream>
    bool StringifyUriFragment(OutputStream& os) const {
        return Stringify<true, OutputStream>(os);
    }

    //@}

    //!@name Create value
    //@{

    //! Create a value in a subtree.
    /*!
        If the value is not exist, it creates all parent values and a JSON Null value.
        So it always succeed and return the newly created or existing value.

        Remind that it may change types of parents according to tokens, so it 
        potentially removes previously stored values. For example, if a document 
        was an array, and "/foo" is used to create a value, then the document 
        will be changed to an object, and all existing array elements are lost.

        \param root Root value of a DOM subtree to be resolved. It can be any value other than document root.
        \param allocator Allocator for creating the values if the specified value or its parents are not exist.
        \param alreadyExist If non-null, it stores whether the resolved value is already exist.
        \return The resolved newly created (a JSON Null value), or already exists value.
    */
    ValueType& Create(ValueType& root, typename ValueType::AllocatorType& allocator, bool* alreadyExist = 0) const {
        CEREAL_RAPIDJSON_ASSERT(IsValid());
        ValueType* v = &root;
        bool exist = true;
        for (const Token *t = tokens_; t != tokens_ + tokenCount_; ++t) {
            if (v->IsArray() && t->name[0] == '-' && t->length == 1) {
                v->PushBack(ValueType().Move(), allocator);
                v = &((*v)[v->Size() - 1]);
                exist = false;
            }
            else {
                if (t->index == kPointerInvalidIndex) { // must be object name
                    if (!v->IsObject())
                        v->SetObject(); // Change to Object
                }
                else { // object name or array index
                    if (!v->IsArray() && !v->IsObject())
                        v->SetArray(); // Change to Array
                }

                if (v->IsArray()) {
                    if (t->index >= v->Size()) {
                        v->Reserve(t->index + 1, allocator);
                        while (t->index >= v->Size())
                            v->PushBack(ValueType().Move(), allocator);
                        exist = false;
                    }
                    v = &((*v)[t->index]);
                }
                else {
                    typename ValueType::MemberIterator m = v->FindMember(GenericStringRef<Ch>(t->name, t->length));
                    if (m == v->MemberEnd()) {
                        v->AddMember(ValueType(t->name, t->length, allocator).Move(), ValueType().Move(), allocator);
                        v = &(--v->MemberEnd())->value; // Assumes AddMember() appends at the end
                        exist = false;
                    }
                    else
                        v = &m->value;
                }
            }
        }

        if (alreadyExist)
            *alreadyExist = exist;

        return *v;
    }

    //! Creates a value in a document.
    /*!
        \param document A document to be resolved.
        \param alreadyExist If non-null, it stores whether the resolved value is already exist.
        \return The resolved newly created, or already exists value.
    */
    template <typename stackAllocator>
    ValueType& Create(GenericDocument<EncodingType, typename ValueType::AllocatorType, stackAllocator>& document, bool* alreadyExist = 0) const {
        return Create(document, document.GetAllocator(), alreadyExist);
    }

    //@}

    //!@name Query value
    //@{

    //! Query a value in a subtree.
    /*!
        \param root Root value of a DOM sub-tree to be resolved. It can be any value other than document root.
        \param unresolvedTokenIndex If the pointer cannot resolve a token in the pointer, this parameter can obtain the index of unresolved token.
        \return Pointer to the value if it can be resolved. Otherwise null.

        \note
        There are only 3 situations when a value cannot be resolved:
        1. A value in the path is not an array nor object.
        2. An object value does not contain the token.
        3. A token is out of range of an array value.

        Use unresolvedTokenIndex to retrieve the token index.
    */
    ValueType* Get(ValueType& root, size_t* unresolvedTokenIndex = 0) const {
        CEREAL_RAPIDJSON_ASSERT(IsValid());
        ValueType* v = &root;
        for (const Token *t = tokens_; t != tokens_ + tokenCount_; ++t) {
            switch (v->GetType()) {
            case kObjectType:
                {
                    typename ValueType::MemberIterator m = v->FindMember(GenericStringRef<Ch>(t->name, t->length));
                    if (m == v->MemberEnd())
                        break;
                    v = &m->value;
                }
                continue;
            case kArrayType:
                if (t->index == kPointerInvalidIndex || t->index >= v->Size())
                    break;
                v = &((*v)[t->index]);
                continue;
            default:
                break;
            }

            // Error: unresolved token
            if (unresolvedTokenIndex)
                *unresolvedTokenIndex = static_cast<size_t>(t - tokens_);
            return 0;
        }
        return v;
    }

    //! Query a const value in a const subtree.
    /*!
        \param root Root value of a DOM sub-tree to be resolved. It can be any value other than document root.
        \return Pointer to the value if it can be resolved. Otherwise null.
    */
    const ValueType* Get(const ValueType& root, size_t* unresolvedTokenIndex = 0) const { 
        return Get(const_cast<ValueType&>(root), unresolvedTokenIndex);
    }

    //@}

    //!@name Query a value with default
    //@{

    //! Query a value in a subtree with default value.
    /*!
        Similar to Get(), but if the specified value do not exists, it creates all parents and clone the default value.
        So that this function always succeed.

        \param root Root value of a DOM sub-tree to be resolved. It can be any value other than document root.
        \param defaultValue Default value to be cloned if the value was not exists.
        \param allocator Allocator for creating the values if the specified value or its parents are not exist.
        \see Create()
    */
    ValueType& GetWithDefault(ValueType& root, const ValueType& defaultValue, typename ValueType::AllocatorType& allocator) const {
        bool alreadyExist;
        ValueType& v = Create(root, allocator, &alreadyExist);
        return alreadyExist ? v : v.CopyFrom(defaultValue, allocator);
    }

    //! Query a value in a subtree with default null-terminated string.
    ValueType& GetWithDefault(ValueType& root, const Ch* defaultValue, typename ValueType::AllocatorType& allocator) const {
        bool alreadyExist;
        ValueType& v = Create(root, allocator, &alreadyExist);
        return alreadyExist ? v : v.SetString(defaultValue, allocator);
    }

#if CEREAL_RAPIDJSON_HAS_STDSTRING
    //! Query a value in a subtree with default std::basic_string.
    ValueType& GetWithDefault(ValueType& root, const std::basic_string<Ch>& defaultValue, typename ValueType::AllocatorType& allocator) const {
        bool alreadyExist;
        ValueType& v = Create(root, allocator, &alreadyExist);
        return alreadyExist ? v : v.SetString(defaultValue, allocator);
    }
#endif

    //! Query a value in a subtree with default primitive value.
    /*!
        \tparam T Either \ref Type, \c int, \c unsigned, \c int64_t, \c uint64_t, \c bool
    */
    template <typename T>
    CEREAL_RAPIDJSON_DISABLEIF_RETURN((internal::OrExpr<internal::IsPointer<T>, internal::IsGenericValue<T> >), (ValueType&))
    GetWithDefault(ValueType& root, T defaultValue, typename ValueType::AllocatorType& allocator) const {
        return GetWithDefault(root, ValueType(defaultValue).Move(), allocator);
    }

    //! Query a value in a document with default value.
    template <typename stackAllocator>
    ValueType& GetWithDefault(GenericDocument<EncodingType, typename ValueType::AllocatorType, stackAllocator>& document, const ValueType& defaultValue) const {
        return GetWithDefault(document, defaultValue, document.GetAllocator());
    }

    //! Query a value in a document with default null-terminated string.
    template <typename stackAllocator>
    ValueType& GetWithDefault(GenericDocument<EncodingType, typename ValueType::AllocatorType, stackAllocator>& document, const Ch* defaultValue) const {
        return GetWithDefault(document, defaultValue, document.GetAllocator());
    }
    
#if CEREAL_RAPIDJSON_HAS_STDSTRING
    //! Query a value in a document with default std::basic_string.
    template <typename stackAllocator>
    ValueType& GetWithDefault(GenericDocument<EncodingType, typename ValueType::AllocatorType, stackAllocator>& document, const std::basic_string<Ch>& defaultValue) const {
        return GetWithDefault(document, defaultValue, document.GetAllocator());
    }
#endif

    //! Query a value in a document with default primitive value.
    /*!
        \tparam T Either \ref Type, \c int, \c unsigned, \c int64_t, \c uint64_t, \c bool
    */
    template <typename T, typename stackAllocator>
    CEREAL_RAPIDJSON_DISABLEIF_RETURN((internal::OrExpr<internal::IsPointer<T>, internal::IsGenericValue<T> >), (ValueType&))
    GetWithDefault(GenericDocument<EncodingType, typename ValueType::AllocatorType, stackAllocator>& document, T defaultValue) const {
        return GetWithDefault(document, defaultValue, document.GetAllocator());
    }

    //@}

    //!@name Set a value
    //@{

    //! Set a value in a subtree, with move semantics.
    /*!
        It creates all parents if they are not exist or types are different to the tokens.
        So this function always succeeds but potentially remove existing values.

        \param root Root value of a DOM sub-tree to be resolved. It can be any value other than document root.
        \param value Value to be set.
        \param allocator Allocator for creating the values if the specified value or its parents are not exist.
        \see Create()
    */
    ValueType& Set(ValueType& root, ValueType& value, typename ValueType::AllocatorType& allocator) const {
        return Create(root, allocator) = value;
    }

    //! Set a value in a subtree, with copy semantics.
    ValueType& Set(ValueType& root, const ValueType& value, typename ValueType::AllocatorType& allocator) const {
        return Create(root, allocator).CopyFrom(value, allocator);
    }

    //! Set a null-terminated string in a subtree.
    ValueType& Set(ValueType& root, const Ch* value, typename ValueType::AllocatorType& allocator) const {
        return Create(root, allocator) = ValueType(value, allocator).Move();
    }

#if CEREAL_RAPIDJSON_HAS_STDSTRING
    //! Set a std::basic_string in a subtree.
    ValueType& Set(ValueType& root, const std::basic_string<Ch>& value, typename ValueType::AllocatorType& allocator) const {
        return Create(root, allocator) = ValueType(value, allocator).Move();
    }
#endif

    //! Set a primitive value in a subtree.
    /*!
        \tparam T Either \ref Type, \c int, \c unsigned, \c int64_t, \c uint64_t, \c bool
    */
    template <typename T>
    CEREAL_RAPIDJSON_DISABLEIF_RETURN((internal::OrExpr<internal::IsPointer<T>, internal::IsGenericValue<T> >), (ValueType&))
    Set(ValueType& root, T value, typename ValueType::AllocatorType& allocator) const {
        return Create(root, allocator) = ValueType(value).Move();
    }

    //! Set a value in a document, with move semantics.
    template <typename stackAllocator>
    ValueType& Set(GenericDocument<EncodingType, typename ValueType::AllocatorType, stackAllocator>& document, ValueType& value) const {
        return Create(document) = value;
    }

    //! Set a value in a document, with copy semantics.
    template <typename stackAllocator>
    ValueType& Set(GenericDocument<EncodingType, typename ValueType::AllocatorType, stackAllocator>& document, const ValueType& value) const {
        return Create(document).CopyFrom(value, document.GetAllocator());
    }

    //! Set a null-terminated string in a document.
    template <typename stackAllocator>
    ValueType& Set(GenericDocument<EncodingType, typename ValueType::AllocatorType, stackAllocator>& document, const Ch* value) const {
        return Create(document) = ValueType(value, document.GetAllocator()).Move();
    }

#if CEREAL_RAPIDJSON_HAS_STDSTRING
    //! Sets a std::basic_string in a document.
    template <typename stackAllocator>
    ValueType& Set(GenericDocument<EncodingType, typename ValueType::AllocatorType, stackAllocator>& document, const std::basic_string<Ch>& value) const {
        return Create(document) = ValueType(value, document.GetAllocator()).Move();
    }
#endif

    //! Set a primitive value in a document.
    /*!
    \tparam T Either \ref Type, \c int, \c unsigned, \c int64_t, \c uint64_t, \c bool
    */
    template <typename T, typename stackAllocator>
    CEREAL_RAPIDJSON_DISABLEIF_RETURN((internal::OrExpr<internal::IsPointer<T>, internal::IsGenericValue<T> >), (ValueType&))
        Set(GenericDocument<EncodingType, typename ValueType::AllocatorType, stackAllocator>& document, T value) const {
            return Create(document) = value;
    }

    //@}

    //!@name Swap a value
    //@{

    //! Swap a value with a value in a subtree.
    /*!
        It creates all parents if they are not exist or types are different to the tokens.
        So this function always succeeds but potentially remove existing values.

        \param root Root value of a DOM sub-tree to be resolved. It can be any value other than document root.
        \param value Value to be swapped.
        \param allocator Allocator for creating the values if the specified value or its parents are not exist.
        \see Create()
    */
    ValueType& Swap(ValueType& root, ValueType& value, typename ValueType::AllocatorType& allocator) const {
        return Create(root, allocator).Swap(value);
    }

    //! Swap a value with a value in a document.
    template <typename stackAllocator>
    ValueType& Swap(GenericDocument<EncodingType, typename ValueType::AllocatorType, stackAllocator>& document, ValueType& value) const {
        return Create(document).Swap(value);
    }

    //@}

    //! Erase a value in a subtree.
    /*!
        \param root Root value of a DOM sub-tree to be resolved. It can be any value other than document root.
        \return Whether the resolved value is found and erased.

        \note Erasing with an empty pointer \c Pointer(""), i.e. the root, always fail and return false.
    */
    bool Erase(ValueType& root) const {
        CEREAL_RAPIDJSON_ASSERT(IsValid());
        if (tokenCount_ == 0) // Cannot erase the root
            return false;

        ValueType* v = &root;
        const Token* last = tokens_ + (tokenCount_ - 1);
        for (const Token *t = tokens_; t != last; ++t) {
            switch (v->GetType()) {
            case kObjectType:
                {
                    typename ValueType::MemberIterator m = v->FindMember(GenericStringRef<Ch>(t->name, t->length));
                    if (m == v->MemberEnd())
                        return false;
                    v = &m->value;
                }
                break;
            case kArrayType:
                if (t->index == kPointerInvalidIndex || t->index >= v->Size())
                    return false;
                v = &((*v)[t->index]);
                break;
            default:
                return false;
            }
        }

        switch (v->GetType()) {
        case kObjectType:
            return v->EraseMember(GenericStringRef<Ch>(last->name, last->length));
        case kArrayType:
            if (last->index == kPointerInvalidIndex || last->index >= v->Size())
                return false;
            v->Erase(v->Begin() + last->index);
            return true;
        default:
            return false;
        }
    }

private:
    //! Clone the content from rhs to this.
    /*!
        \param rhs Source pointer.
        \param extraToken Extra tokens to be allocated.
        \param extraNameBufferSize Extra name buffer size (in number of Ch) to be allocated.
        \return Start of non-occupied name buffer, for storing extra names.
    */
    Ch* CopyFromRaw(const GenericPointer& rhs, size_t extraToken = 0, size_t extraNameBufferSize = 0) {
        if (!allocator_) // allocator is independently owned.
            ownAllocator_ = allocator_ = CEREAL_RAPIDJSON_NEW(Allocator)();

        size_t nameBufferSize = rhs.tokenCount_; // null terminators for tokens
        for (Token *t = rhs.tokens_; t != rhs.tokens_ + rhs.tokenCount_; ++t)
            nameBufferSize += t->length;

        tokenCount_ = rhs.tokenCount_ + extraToken;
        tokens_ = static_cast<Token *>(allocator_->Malloc(tokenCount_ * sizeof(Token) + (nameBufferSize + extraNameBufferSize) * sizeof(Ch)));
        nameBuffer_ = reinterpret_cast<Ch *>(tokens_ + tokenCount_);
        if (rhs.tokenCount_ > 0) {
            std::memcpy(tokens_, rhs.tokens_, rhs.tokenCount_ * sizeof(Token));
        }
        if (nameBufferSize > 0) {
            std::memcpy(nameBuffer_, rhs.nameBuffer_, nameBufferSize * sizeof(Ch));
        }

        // Adjust pointers to name buffer
        std::ptrdiff_t diff = nameBuffer_ - rhs.nameBuffer_;
        for (Token *t = tokens_; t != tokens_ + rhs.tokenCount_; ++t)
            t->name += diff;

        return nameBuffer_ + nameBufferSize;
    }

    //! Check whether a character should be percent-encoded.
    /*!
        According to RFC 3986 2.3 Unreserved Characters.
        \param c The character (code unit) to be tested.
    */
    bool NeedPercentEncode(Ch c) const {
        return !((c >= '0' && c <= '9') || (c >= 'A' && c <='Z') || (c >= 'a' && c <= 'z') || c == '-' || c == '.' || c == '_' || c =='~');
    }

    //! Parse a JSON String or its URI fragment representation into tokens.
#ifndef __clang__ // -Wdocumentation
    /*!
        \param source Either a JSON Pointer string, or its URI fragment representation. Not need to be null terminated.
        \param length Length of the source string.
        \note Source cannot be JSON String Representation of JSON Pointer, e.g. In "/\u0000", \u0000 will not be unescaped.
    */
#endif
    void Parse(const Ch* source, size_t length) {
        CEREAL_RAPIDJSON_ASSERT(source != NULL);
        CEREAL_RAPIDJSON_ASSERT(nameBuffer_ == 0);
        CEREAL_RAPIDJSON_ASSERT(tokens_ == 0);

        // Create own allocator if user did not supply.
        if (!allocator_)
            ownAllocator_ = allocator_ = CEREAL_RAPIDJSON_NEW(Allocator)();

        // Count number of '/' as tokenCount
        tokenCount_ = 0;
        for (const Ch* s = source; s != source + length; s++) 
            if (*s == '/')
                tokenCount_++;

        Token* token = tokens_ = static_cast<Token *>(allocator_->Malloc(tokenCount_ * sizeof(Token) + length * sizeof(Ch)));
        Ch* name = nameBuffer_ = reinterpret_cast<Ch *>(tokens_ + tokenCount_);
        size_t i = 0;

        // Detect if it is a URI fragment
        bool uriFragment = false;
        if (source[i] == '#') {
            uriFragment = true;
            i++;
        }

        if (i != length && source[i] != '/') {
            parseErrorCode_ = kPointerParseErrorTokenMustBeginWithSolidus;
            goto error;
        }

        while (i < length) {
            CEREAL_RAPIDJSON_ASSERT(source[i] == '/');
            i++; // consumes '/'

            token->name = name;
            bool isNumber = true;

            while (i < length && source[i] != '/') {
                Ch c = source[i];
                if (uriFragment) {
                    // Decoding percent-encoding for URI fragment
                    if (c == '%') {
                        PercentDecodeStream is(&source[i], source + length);
                        GenericInsituStringStream<EncodingType> os(name);
                        Ch* begin = os.PutBegin();
                        if (!Transcoder<UTF8<>, EncodingType>().Validate(is, os) || !is.IsValid()) {
                            parseErrorCode_ = kPointerParseErrorInvalidPercentEncoding;
                            goto error;
                        }
                        size_t len = os.PutEnd(begin);
                        i += is.Tell() - 1;
                        if (len == 1)
                            c = *name;
                        else {
                            name += len;
                            isNumber = false;
                            i++;
                            continue;
                        }
                    }
                    else if (NeedPercentEncode(c)) {
                        parseErrorCode_ = kPointerParseErrorCharacterMustPercentEncode;
                        goto error;
                    }
                }

                i++;
                
                // Escaping "~0" -> '~', "~1" -> '/'
                if (c == '~') {
                    if (i < length) {
                        c = source[i];
                        if (c == '0')       c = '~';
                        else if (c == '1')  c = '/';
                        else {
                            parseErrorCode_ = kPointerParseErrorInvalidEscape;
                            goto error;
                        }
                        i++;
                    }
                    else {
                        parseErrorCode_ = kPointerParseErrorInvalidEscape;
                        goto error;
                    }
                }

                // First check for index: all of characters are digit
                if (c < '0' || c > '9')
                    isNumber = false;

                *name++ = c;
            }
            token->length = static_cast<SizeType>(name - token->name);
            if (token->length == 0)
                isNumber = false;
            *name++ = '\0'; // Null terminator

            // Second check for index: more than one digit cannot have leading zero
            if (isNumber && token->length > 1 && token->name[0] == '0')
                isNumber = false;

            // String to SizeType conversion
            SizeType n = 0;
            if (isNumber) {
                for (size_t j = 0; j < token->length; j++) {
                    SizeType m = n * 10 + static_cast<SizeType>(token->name[j] - '0');
                    if (m < n) {   // overflow detection
                        isNumber = false;
                        break;
                    }
                    n = m;
                }
            }

            token->index = isNumber ? n : kPointerInvalidIndex;
            token++;
        }

        CEREAL_RAPIDJSON_ASSERT(name <= nameBuffer_ + length); // Should not overflow buffer
        parseErrorCode_ = kPointerParseErrorNone;
        return;

    error:
        Allocator::Free(tokens_);
        nameBuffer_ = 0;
        tokens_ = 0;
        tokenCount_ = 0;
        parseErrorOffset_ = i;
        return;
    }

    //! Stringify to string or URI fragment representation.
    /*!
        \tparam uriFragment True for stringifying to URI fragment representation. False for string representation.
        \tparam OutputStream type of output stream.
        \param os The output stream.
    */
    template<bool uriFragment, typename OutputStream>
    bool Stringify(OutputStream& os) const {
        CEREAL_RAPIDJSON_ASSERT(IsValid());

        if (uriFragment)
            os.Put('#');

        for (Token *t = tokens_; t != tokens_ + tokenCount_; ++t) {
            os.Put('/');
            for (size_t j = 0; j < t->length; j++) {
                Ch c = t->name[j];
                if (c == '~') {
                    os.Put('~');
                    os.Put('0');
                }
                else if (c == '/') {
                    os.Put('~');
                    os.Put('1');
                }
                else if (uriFragment && NeedPercentEncode(c)) { 
                    // Transcode to UTF8 sequence
                    GenericStringStream<typename ValueType::EncodingType> source(&t->name[j]);
                    PercentEncodeStream<OutputStream> target(os);
                    if (!Transcoder<EncodingType, UTF8<> >().Validate(source, target))
                        return false;
                    j += source.Tell() - 1;
                }
                else
                    os.Put(c);
            }
        }
        return true;
    }

    //! A helper stream for decoding a percent-encoded sequence into code unit.
    /*!
        This stream decodes %XY triplet into code unit (0-255).
        If it encounters invalid characters, it sets output code unit as 0 and 
        mark invalid, and to be checked by IsValid().
    */
    class PercentDecodeStream {
    public:
        typedef typename ValueType::Ch Ch;

        //! Constructor
        /*!
            \param source Start of the stream
            \param end Past-the-end of the stream.
        */
        PercentDecodeStream(const Ch* source, const Ch* end) : src_(source), head_(source), end_(end), valid_(true) {}

        Ch Take() {
            if (*src_ != '%' || src_ + 3 > end_) { // %XY triplet
                valid_ = false;
                return 0;
            }
            src_++;
            Ch c = 0;
            for (int j = 0; j < 2; j++) {
                c = static_cast<Ch>(c << 4);
                Ch h = *src_;
                if      (h >= '0' && h <= '9') c = static_cast<Ch>(c + h - '0');
                else if (h >= 'A' && h <= 'F') c = static_cast<Ch>(c + h - 'A' + 10);
                else if (h >= 'a' && h <= 'f') c = static_cast<Ch>(c + h - 'a' + 10);
                else {
                    valid_ = false;
                    return 0;
                }
                src_++;
            }
            return c;
        }

        size_t Tell() const { return static_cast<size_t>(src_ - head_); }
        bool IsValid() const { return valid_; }

    private:
        const Ch* src_;     //!< Current read position.
        const Ch* head_;    //!< Original head of the string.
        const Ch* end_;     //!< Past-the-end position.
        bool valid_;        //!< Whether the parsing is valid.
    };

    //! A helper stream to encode character (UTF-8 code unit) into percent-encoded sequence.
    template <typename OutputStream>
    class PercentEncodeStream {
    public:
        PercentEncodeStream(OutputStream& os) : os_(os) {}
        void Put(char c) { // UTF-8 must be byte
            unsigned char u = static_cast<unsigned char>(c);
            static const char hexDigits[16] = { '0', '1', '2', '3', '4', '5', '6', '7', '8', '9', 'A', 'B', 'C', 'D', 'E', 'F' };
            os_.Put('%');
            os_.Put(static_cast<typename OutputStream::Ch>(hexDigits[u >> 4]));
            os_.Put(static_cast<typename OutputStream::Ch>(hexDigits[u & 15]));
        }
    private:
        OutputStream& os_;
    };

    Allocator* allocator_;                  //!< The current allocator. It is either user-supplied or equal to ownAllocator_.
    Allocator* ownAllocator_;               //!< Allocator owned by this Pointer.
    Ch* nameBuffer_;                        //!< A buffer containing all names in tokens.
    Token* tokens_;                         //!< A list of tokens.
    size_t tokenCount_;                     //!< Number of tokens in tokens_.
    size_t parseErrorOffset_;               //!< Offset in code unit when parsing fail.
    PointerParseErrorCode parseErrorCode_;  //!< Parsing error code.
};

//! GenericPointer for Value (UTF-8, default allocator).
typedef GenericPointer<Value> Pointer;

//!@name Helper functions for GenericPointer
//@{

//////////////////////////////////////////////////////////////////////////////

template <typename T>
typename T::ValueType& CreateValueByPointer(T& root, const GenericPointer<typename T::ValueType>& pointer, typename T::AllocatorType& a) {
    return pointer.Create(root, a);
}

template <typename T, typename CharType, size_t N>
typename T::ValueType& CreateValueByPointer(T& root, const CharType(&source)[N], typename T::AllocatorType& a) {
    return GenericPointer<typename T::ValueType>(source, N - 1).Create(root, a);
}

// No allocator parameter

template <typename DocumentType>
typename DocumentType::ValueType& CreateValueByPointer(DocumentType& document, const GenericPointer<typename DocumentType::ValueType>& pointer) {
    return pointer.Create(document);
}

template <typename DocumentType, typename CharType, size_t N>
typename DocumentType::ValueType& CreateValueByPointer(DocumentType& document, const CharType(&source)[N]) {
    return GenericPointer<typename DocumentType::ValueType>(source, N - 1).Create(document);
}

//////////////////////////////////////////////////////////////////////////////

template <typename T>
typename T::ValueType* GetValueByPointer(T& root, const GenericPointer<typename T::ValueType>& pointer, size_t* unresolvedTokenIndex = 0) {
    return pointer.Get(root, unresolvedTokenIndex);
}

template <typename T>
const typename T::ValueType* GetValueByPointer(const T& root, const GenericPointer<typename T::ValueType>& pointer, size_t* unresolvedTokenIndex = 0) {
    return pointer.Get(root, unresolvedTokenIndex);
}

template <typename T, typename CharType, size_t N>
typename T::ValueType* GetValueByPointer(T& root, const CharType (&source)[N], size_t* unresolvedTokenIndex = 0) {
    return GenericPointer<typename T::ValueType>(source, N - 1).Get(root, unresolvedTokenIndex);
}

template <typename T, typename CharType, size_t N>
const typename T::ValueType* GetValueByPointer(const T& root, const CharType(&source)[N], size_t* unresolvedTokenIndex = 0) {
    return GenericPointer<typename T::ValueType>(source, N - 1).Get(root, unresolvedTokenIndex);
}

//////////////////////////////////////////////////////////////////////////////

template <typename T>
typename T::ValueType& GetValueByPointerWithDefault(T& root, const GenericPointer<typename T::ValueType>& pointer, const typename T::ValueType& defaultValue, typename T::AllocatorType& a) {
    return pointer.GetWithDefault(root, defaultValue, a);
}

template <typename T>
typename T::ValueType& GetValueByPointerWithDefault(T& root, const GenericPointer<typename T::ValueType>& pointer, const typename T::Ch* defaultValue, typename T::AllocatorType& a) {
    return pointer.GetWithDefault(root, defaultValue, a);
}

#if CEREAL_RAPIDJSON_HAS_STDSTRING
template <typename T>
typename T::ValueType& GetValueByPointerWithDefault(T& root, const GenericPointer<typename T::ValueType>& pointer, const std::basic_string<typename T::Ch>& defaultValue, typename T::AllocatorType& a) {
    return pointer.GetWithDefault(root, defaultValue, a);
}
#endif

template <typename T, typename T2>
CEREAL_RAPIDJSON_DISABLEIF_RETURN((internal::OrExpr<internal::IsPointer<T2>, internal::IsGenericValue<T2> >), (typename T::ValueType&))
GetValueByPointerWithDefault(T& root, const GenericPointer<typename T::ValueType>& pointer, T2 defaultValue, typename T::AllocatorType& a) {
    return pointer.GetWithDefault(root, defaultValue, a);
}

template <typename T, typename CharType, size_t N>
typename T::ValueType& GetValueByPointerWithDefault(T& root, const CharType(&source)[N], const typename T::ValueType& defaultValue, typename T::AllocatorType& a) {
    return GenericPointer<typename T::ValueType>(source, N - 1).GetWithDefault(root, defaultValue, a);
}

template <typename T, typename CharType, size_t N>
typename T::ValueType& GetValueByPointerWithDefault(T& root, const CharType(&source)[N], const typename T::Ch* defaultValue, typename T::AllocatorType& a) {
    return GenericPointer<typename T::ValueType>(source, N - 1).GetWithDefault(root, defaultValue, a);
}

#if CEREAL_RAPIDJSON_HAS_STDSTRING
template <typename T, typename CharType, size_t N>
typename T::ValueType& GetValueByPointerWithDefault(T& root, const CharType(&source)[N], const std::basic_string<typename T::Ch>& defaultValue, typename T::AllocatorType& a) {
    return GenericPointer<typename T::ValueType>(source, N - 1).GetWithDefault(root, defaultValue, a);
}
#endif

template <typename T, typename CharType, size_t N, typename T2>
CEREAL_RAPIDJSON_DISABLEIF_RETURN((internal::OrExpr<internal::IsPointer<T2>, internal::IsGenericValue<T2> >), (typename T::ValueType&))
GetValueByPointerWithDefault(T& root, const CharType(&source)[N], T2 defaultValue, typename T::AllocatorType& a) {
    return GenericPointer<typename T::ValueType>(source, N - 1).GetWithDefault(root, defaultValue, a);
}

// No allocator parameter

template <typename DocumentType>
typename DocumentType::ValueType& GetValueByPointerWithDefault(DocumentType& document, const GenericPointer<typename DocumentType::ValueType>& pointer, const typename DocumentType::ValueType& defaultValue) {
    return pointer.GetWithDefault(document, defaultValue);
}

template <typename DocumentType>
typename DocumentType::ValueType& GetValueByPointerWithDefault(DocumentType& document, const GenericPointer<typename DocumentType::ValueType>& pointer, const typename DocumentType::Ch* defaultValue) {
    return pointer.GetWithDefault(document, defaultValue);
}

#if CEREAL_RAPIDJSON_HAS_STDSTRING
template <typename DocumentType>
typename DocumentType::ValueType& GetValueByPointerWithDefault(DocumentType& document, const GenericPointer<typename DocumentType::ValueType>& pointer, const std::basic_string<typename DocumentType::Ch>& defaultValue) {
    return pointer.GetWithDefault(document, defaultValue);
}
#endif

template <typename DocumentType, typename T2>
CEREAL_RAPIDJSON_DISABLEIF_RETURN((internal::OrExpr<internal::IsPointer<T2>, internal::IsGenericValue<T2> >), (typename DocumentType::ValueType&))
GetValueByPointerWithDefault(DocumentType& document, const GenericPointer<typename DocumentType::ValueType>& pointer, T2 defaultValue) {
    return pointer.GetWithDefault(document, defaultValue);
}

template <typename DocumentType, typename CharType, size_t N>
typename DocumentType::ValueType& GetValueByPointerWithDefault(DocumentType& document, const CharType(&source)[N], const typename DocumentType::ValueType& defaultValue) {
    return GenericPointer<typename DocumentType::ValueType>(source, N - 1).GetWithDefault(document, defaultValue);
}

template <typename DocumentType, typename CharType, size_t N>
typename DocumentType::ValueType& GetValueByPointerWithDefault(DocumentType& document, const CharType(&source)[N], const typename DocumentType::Ch* defaultValue) {
    return GenericPointer<typename DocumentType::ValueType>(source, N - 1).GetWithDefault(document, defaultValue);
}

#if CEREAL_RAPIDJSON_HAS_STDSTRING
template <typename DocumentType, typename CharType, size_t N>
typename DocumentType::ValueType& GetValueByPointerWithDefault(DocumentType& document, const CharType(&source)[N], const std::basic_string<typename DocumentType::Ch>& defaultValue) {
    return GenericPointer<typename DocumentType::ValueType>(source, N - 1).GetWithDefault(document, defaultValue);
}
#endif

template <typename DocumentType, typename CharType, size_t N, typename T2>
CEREAL_RAPIDJSON_DISABLEIF_RETURN((internal::OrExpr<internal::IsPointer<T2>, internal::IsGenericValue<T2> >), (typename DocumentType::ValueType&))
GetValueByPointerWithDefault(DocumentType& document, const CharType(&source)[N], T2 defaultValue) {
    return GenericPointer<typename DocumentType::ValueType>(source, N - 1).GetWithDefault(document, defaultValue);
}

//////////////////////////////////////////////////////////////////////////////

template <typename T>
typename T::ValueType& SetValueByPointer(T& root, const GenericPointer<typename T::ValueType>& pointer, typename T::ValueType& value, typename T::AllocatorType& a) {
    return pointer.Set(root, value, a);
}

template <typename T>
typename T::ValueType& SetValueByPointer(T& root, const GenericPointer<typename T::ValueType>& pointer, const typename T::ValueType& value, typename T::AllocatorType& a) {
    return pointer.Set(root, value, a);
}

template <typename T>
typename T::ValueType& SetValueByPointer(T& root, const GenericPointer<typename T::ValueType>& pointer, const typename T::Ch* value, typename T::AllocatorType& a) {
    return pointer.Set(root, value, a);
}

#if CEREAL_RAPIDJSON_HAS_STDSTRING
template <typename T>
typename T::ValueType& SetValueByPointer(T& root, const GenericPointer<typename T::ValueType>& pointer, const std::basic_string<typename T::Ch>& value, typename T::AllocatorType& a) {
    return pointer.Set(root, value, a);
}
#endif

template <typename T, typename T2>
CEREAL_RAPIDJSON_DISABLEIF_RETURN((internal::OrExpr<internal::IsPointer<T2>, internal::IsGenericValue<T2> >), (typename T::ValueType&))
SetValueByPointer(T& root, const GenericPointer<typename T::ValueType>& pointer, T2 value, typename T::AllocatorType& a) {
    return pointer.Set(root, value, a);
}

template <typename T, typename CharType, size_t N>
typename T::ValueType& SetValueByPointer(T& root, const CharType(&source)[N], typename T::ValueType& value, typename T::AllocatorType& a) {
    return GenericPointer<typename T::ValueType>(source, N - 1).Set(root, value, a);
}

template <typename T, typename CharType, size_t N>
typename T::ValueType& SetValueByPointer(T& root, const CharType(&source)[N], const typename T::ValueType& value, typename T::AllocatorType& a) {
    return GenericPointer<typename T::ValueType>(source, N - 1).Set(root, value, a);
}

template <typename T, typename CharType, size_t N>
typename T::ValueType& SetValueByPointer(T& root, const CharType(&source)[N], const typename T::Ch* value, typename T::AllocatorType& a) {
    return GenericPointer<typename T::ValueType>(source, N - 1).Set(root, value, a);
}

#if CEREAL_RAPIDJSON_HAS_STDSTRING
template <typename T, typename CharType, size_t N>
typename T::ValueType& SetValueByPointer(T& root, const CharType(&source)[N], const std::basic_string<typename T::Ch>& value, typename T::AllocatorType& a) {
    return GenericPointer<typename T::ValueType>(source, N - 1).Set(root, value, a);
}
#endif

template <typename T, typename CharType, size_t N, typename T2>
CEREAL_RAPIDJSON_DISABLEIF_RETURN((internal::OrExpr<internal::IsPointer<T2>, internal::IsGenericValue<T2> >), (typename T::ValueType&))
SetValueByPointer(T& root, const CharType(&source)[N], T2 value, typename T::AllocatorType& a) {
    return GenericPointer<typename T::ValueType>(source, N - 1).Set(root, value, a);
}

// No allocator parameter

template <typename DocumentType>
typename DocumentType::ValueType& SetValueByPointer(DocumentType& document, const GenericPointer<typename DocumentType::ValueType>& pointer, typename DocumentType::ValueType& value) {
    return pointer.Set(document, value);
}

template <typename DocumentType>
typename DocumentType::ValueType& SetValueByPointer(DocumentType& document, const GenericPointer<typename DocumentType::ValueType>& pointer, const typename DocumentType::ValueType& value) {
    return pointer.Set(document, value);
}

template <typename DocumentType>
typename DocumentType::ValueType& SetValueByPointer(DocumentType& document, const GenericPointer<typename DocumentType::ValueType>& pointer, const typename DocumentType::Ch* value) {
    return pointer.Set(document, value);
}

#if CEREAL_RAPIDJSON_HAS_STDSTRING
template <typename DocumentType>
typename DocumentType::ValueType& SetValueByPointer(DocumentType& document, const GenericPointer<typename DocumentType::ValueType>& pointer, const std::basic_string<typename DocumentType::Ch>& value) {
    return pointer.Set(document, value);
}
#endif

template <typename DocumentType, typename T2>
CEREAL_RAPIDJSON_DISABLEIF_RETURN((internal::OrExpr<internal::IsPointer<T2>, internal::IsGenericValue<T2> >), (typename DocumentType::ValueType&))
SetValueByPointer(DocumentType& document, const GenericPointer<typename DocumentType::ValueType>& pointer, T2 value) {
    return pointer.Set(document, value);
}

template <typename DocumentType, typename CharType, size_t N>
typename DocumentType::ValueType& SetValueByPointer(DocumentType& document, const CharType(&source)[N], typename DocumentType::ValueType& value) {
    return GenericPointer<typename DocumentType::ValueType>(source, N - 1).Set(document, value);
}

template <typename DocumentType, typename CharType, size_t N>
typename DocumentType::ValueType& SetValueByPointer(DocumentType& document, const CharType(&source)[N], const typename DocumentType::ValueType& value) {
    return GenericPointer<typename DocumentType::ValueType>(source, N - 1).Set(document, value);
}

template <typename DocumentType, typename CharType, size_t N>
typename DocumentType::ValueType& SetValueByPointer(DocumentType& document, const CharType(&source)[N], const typename DocumentType::Ch* value) {
    return GenericPointer<typename DocumentType::ValueType>(source, N - 1).Set(document, value);
}

#if CEREAL_RAPIDJSON_HAS_STDSTRING
template <typename DocumentType, typename CharType, size_t N>
typename DocumentType::ValueType& SetValueByPointer(DocumentType& document, const CharType(&source)[N], const std::basic_string<typename DocumentType::Ch>& value) {
    return GenericPointer<typename DocumentType::ValueType>(source, N - 1).Set(document, value);
}
#endif

template <typename DocumentType, typename CharType, size_t N, typename T2>
CEREAL_RAPIDJSON_DISABLEIF_RETURN((internal::OrExpr<internal::IsPointer<T2>, internal::IsGenericValue<T2> >), (typename DocumentType::ValueType&))
SetValueByPointer(DocumentType& document, const CharType(&source)[N], T2 value) {
    return GenericPointer<typename DocumentType::ValueType>(source, N - 1).Set(document, value);
}

//////////////////////////////////////////////////////////////////////////////

template <typename T>
typename T::ValueType& SwapValueByPointer(T& root, const GenericPointer<typename T::ValueType>& pointer, typename T::ValueType& value, typename T::AllocatorType& a) {
    return pointer.Swap(root, value, a);
}

template <typename T, typename CharType, size_t N>
typename T::ValueType& SwapValueByPointer(T& root, const CharType(&source)[N], typename T::ValueType& value, typename T::AllocatorType& a) {
    return GenericPointer<typename T::ValueType>(source, N - 1).Swap(root, value, a);
}

template <typename DocumentType>
typename DocumentType::ValueType& SwapValueByPointer(DocumentType& document, const GenericPointer<typename DocumentType::ValueType>& pointer, typename DocumentType::ValueType& value) {
    return pointer.Swap(document, value);
}

template <typename DocumentType, typename CharType, size_t N>
typename DocumentType::ValueType& SwapValueByPointer(DocumentType& document, const CharType(&source)[N], typename DocumentType::ValueType& value) {
    return GenericPointer<typename DocumentType::ValueType>(source, N - 1).Swap(document, value);
}

//////////////////////////////////////////////////////////////////////////////

template <typename T>
bool EraseValueByPointer(T& root, const GenericPointer<typename T::ValueType>& pointer) {
    return pointer.Erase(root);
}

template <typename T, typename CharType, size_t N>
bool EraseValueByPointer(T& root, const CharType(&source)[N]) {
    return GenericPointer<typename T::ValueType>(source, N - 1).Erase(root);
}

//@}

CEREAL_RAPIDJSON_NAMESPACE_END

#if defined(__clang__) || defined(_MSC_VER)
CEREAL_RAPIDJSON_DIAG_POP
#endif

#endif // CEREAL_RAPIDJSON_POINTER_H_
```
