## Supplemental code for "The motif composition of variable-number tandem repeats impacts gene expression": prettywriter.h

```
// Tencent is pleased to support the open source community by making RapidJSON available.
// 
// Copyright (C) 2015 THL A29 Limited, a Tencent company, and Milo Yip. All rights reserved.
//
// Licensed under the MIT License (the "License"); you may not use this file except
// in compliance with the License. You may obtain a copy of the License at
//
// http://opensource.org/licenses/MIT
//
// Unless required by applicable law or agreed to in writing, software distributed 
// under the License is distributed on an "AS IS" BASIS, WITHOUT WARRANTIES OR 
// CONDITIONS OF ANY KIND, either express or implied. See the License for the 
// specific language governing permissions and limitations under the License.

#ifndef CEREAL_RAPIDJSON_PRETTYWRITER_H_
#define CEREAL_RAPIDJSON_PRETTYWRITER_H_

#include "writer.h"

#ifdef __GNUC__
CEREAL_RAPIDJSON_DIAG_PUSH
CEREAL_RAPIDJSON_DIAG_OFF(effc++)
#endif

#if defined(__clang__)
CEREAL_RAPIDJSON_DIAG_PUSH
CEREAL_RAPIDJSON_DIAG_OFF(c++98-compat)
#endif

CEREAL_RAPIDJSON_NAMESPACE_BEGIN

//! Combination of PrettyWriter format flags.
/*! \see PrettyWriter::SetFormatOptions
 */
enum PrettyFormatOptions {
    kFormatDefault = 0,         //!< Default pretty formatting.
    kFormatSingleLineArray = 1  //!< Format arrays on a single line.
};

//! Writer with indentation and spacing.
/*!
    \tparam OutputStream Type of output os.
    \tparam SourceEncoding Encoding of source string.
    \tparam TargetEncoding Encoding of output stream.
    \tparam StackAllocator Type of allocator for allocating memory of stack.
*/
template<typename OutputStream, typename SourceEncoding = UTF8<>, typename TargetEncoding = UTF8<>, typename StackAllocator = CrtAllocator, unsigned writeFlags = kWriteDefaultFlags>
class PrettyWriter : public Writer<OutputStream, SourceEncoding, TargetEncoding, StackAllocator, writeFlags> {
public:
    typedef Writer<OutputStream, SourceEncoding, TargetEncoding, StackAllocator, writeFlags> Base;
    typedef typename Base::Ch Ch;

    //! Constructor
    /*! \param os Output stream.
        \param allocator User supplied allocator. If it is null, it will create a private one.
        \param levelDepth Initial capacity of stack.
    */
    explicit PrettyWriter(OutputStream& os, StackAllocator* allocator = 0, size_t levelDepth = Base::kDefaultLevelDepth) : 
        Base(os, allocator, levelDepth), indentChar_(' '), indentCharCount_(4), formatOptions_(kFormatDefault) {}


    explicit PrettyWriter(StackAllocator* allocator = 0, size_t levelDepth = Base::kDefaultLevelDepth) : 
        Base(allocator, levelDepth), indentChar_(' '), indentCharCount_(4) {}

#if CEREAL_RAPIDJSON_HAS_CXX11_RVALUE_REFS
    PrettyWriter(PrettyWriter&& rhs) :
        Base(std::forward<PrettyWriter>(rhs)), indentChar_(rhs.indentChar_), indentCharCount_(rhs.indentCharCount_), formatOptions_(rhs.formatOptions_) {}
#endif

    //! Set custom indentation.
    /*! \param indentChar       Character for indentation. Must be whitespace character (' ', '\\t', '\\n', '\\r').
        \param indentCharCount  Number of indent characters for each indentation level.
        \note The default indentation is 4 spaces.
    */
    PrettyWriter& SetIndent(Ch indentChar, unsigned indentCharCount) {
        CEREAL_RAPIDJSON_ASSERT(indentChar == ' ' || indentChar == '\t' || indentChar == '\n' || indentChar == '\r');
        indentChar_ = indentChar;
        indentCharCount_ = indentCharCount;
        return *this;
    }

    //! Set pretty writer formatting options.
    /*! \param options Formatting options.
    */
    PrettyWriter& SetFormatOptions(PrettyFormatOptions options) {
        formatOptions_ = options;
        return *this;
    }

    /*! @name Implementation of Handler
        \see Handler
    */
    //@{

    bool Null()                 { PrettyPrefix(kNullType);   return Base::EndValue(Base::WriteNull()); }
    bool Bool(bool b)           { PrettyPrefix(b ? kTrueType : kFalseType); return Base::EndValue(Base::WriteBool(b)); }
    bool Int(int i)             { PrettyPrefix(kNumberType); return Base::EndValue(Base::WriteInt(i)); }
    bool Uint(unsigned u)       { PrettyPrefix(kNumberType); return Base::EndValue(Base::WriteUint(u)); }
    bool Int64(int64_t i64)     { PrettyPrefix(kNumberType); return Base::EndValue(Base::WriteInt64(i64)); }
    bool Uint64(uint64_t u64)   { PrettyPrefix(kNumberType); return Base::EndValue(Base::WriteUint64(u64));  }
    bool Double(double d)       { PrettyPrefix(kNumberType); return Base::EndValue(Base::WriteDouble(d)); }

    bool RawNumber(const Ch* str, SizeType length, bool copy = false) {
        CEREAL_RAPIDJSON_ASSERT(str != 0);
        (void)copy;
        PrettyPrefix(kNumberType);
        return Base::EndValue(Base::WriteString(str, length));
    }

    bool String(const Ch* str, SizeType length, bool copy = false) {
        CEREAL_RAPIDJSON_ASSERT(str != 0);
        (void)copy;
        PrettyPrefix(kStringType);
        return Base::EndValue(Base::WriteString(str, length));
    }

#if CEREAL_RAPIDJSON_HAS_STDSTRING
    bool String(const std::basic_string<Ch>& str) {
        return String(str.data(), SizeType(str.size()));
    }
#endif

    bool StartObject() {
        PrettyPrefix(kObjectType);
        new (Base::level_stack_.template Push<typename Base::Level>()) typename Base::Level(false);
        return Base::WriteStartObject();
    }

    bool Key(const Ch* str, SizeType length, bool copy = false) { return String(str, length, copy); }

#if CEREAL_RAPIDJSON_HAS_STDSTRING
    bool Key(const std::basic_string<Ch>& str) {
        return Key(str.data(), SizeType(str.size()));
    }
#endif
	
    bool EndObject(SizeType memberCount = 0) {
        (void)memberCount;
        CEREAL_RAPIDJSON_ASSERT(Base::level_stack_.GetSize() >= sizeof(typename Base::Level)); // not inside an Object
        CEREAL_RAPIDJSON_ASSERT(!Base::level_stack_.template Top<typename Base::Level>()->inArray); // currently inside an Array, not Object
        CEREAL_RAPIDJSON_ASSERT(0 == Base::level_stack_.template Top<typename Base::Level>()->valueCount % 2); // Object has a Key without a Value
       
        bool empty = Base::level_stack_.template Pop<typename Base::Level>(1)->valueCount == 0;

        if (!empty) {
            Base::os_->Put('\n');
            WriteIndent();
        }
        bool ret = Base::EndValue(Base::WriteEndObject());
        (void)ret;
        CEREAL_RAPIDJSON_ASSERT(ret == true);
        if (Base::level_stack_.Empty()) // end of json text
            Base::Flush();
        return true;
    }

    bool StartArray() {
        PrettyPrefix(kArrayType);
        new (Base::level_stack_.template Push<typename Base::Level>()) typename Base::Level(true);
        return Base::WriteStartArray();
    }

    bool EndArray(SizeType memberCount = 0) {
        (void)memberCount;
        CEREAL_RAPIDJSON_ASSERT(Base::level_stack_.GetSize() >= sizeof(typename Base::Level));
        CEREAL_RAPIDJSON_ASSERT(Base::level_stack_.template Top<typename Base::Level>()->inArray);
        bool empty = Base::level_stack_.template Pop<typename Base::Level>(1)->valueCount == 0;

        if (!empty && !(formatOptions_ & kFormatSingleLineArray)) {
            Base::os_->Put('\n');
            WriteIndent();
        }
        bool ret = Base::EndValue(Base::WriteEndArray());
        (void)ret;
        CEREAL_RAPIDJSON_ASSERT(ret == true);
        if (Base::level_stack_.Empty()) // end of json text
            Base::Flush();
        return true;
    }

    //@}

    /*! @name Convenience extensions */
    //@{

    //! Simpler but slower overload.
    bool String(const Ch* str) { return String(str, internal::StrLen(str)); }
    bool Key(const Ch* str) { return Key(str, internal::StrLen(str)); }

    //@}

    //! Write a raw JSON value.
    /*!
        For user to write a stringified JSON as a value.

        \param json A well-formed JSON value. It should not contain null character within [0, length - 1] range.
        \param length Length of the json.
        \param type Type of the root of json.
        \note When using PrettyWriter::RawValue(), the result json may not be indented correctly.
    */
    bool RawValue(const Ch* json, size_t length, Type type) {
        CEREAL_RAPIDJSON_ASSERT(json != 0);
        PrettyPrefix(type);
        return Base::EndValue(Base::WriteRawValue(json, length));
    }

protected:
    void PrettyPrefix(Type type) {
        (void)type;
        if (Base::level_stack_.GetSize() != 0) { // this value is not at root
            typename Base::Level* level = Base::level_stack_.template Top<typename Base::Level>();

            if (level->inArray) {
                if (level->valueCount > 0) {
                    Base::os_->Put(','); // add comma if it is not the first element in array
                    if (formatOptions_ & kFormatSingleLineArray)
                        Base::os_->Put(' ');
                }

                if (!(formatOptions_ & kFormatSingleLineArray)) {
                    Base::os_->Put('\n');
                    WriteIndent();
                }
            }
            else {  // in object
                if (level->valueCount > 0) {
                    if (level->valueCount % 2 == 0) {
                        Base::os_->Put(',');
                        Base::os_->Put('\n');
                    }
                    else {
                        Base::os_->Put(':');
                        Base::os_->Put(' ');
                    }
                }
                else
                    Base::os_->Put('\n');

                if (level->valueCount % 2 == 0)
                    WriteIndent();
            }
            if (!level->inArray && level->valueCount % 2 == 0)
                CEREAL_RAPIDJSON_ASSERT(type == kStringType);  // if it's in object, then even number should be a name
            level->valueCount++;
        }
        else {
            CEREAL_RAPIDJSON_ASSERT(!Base::hasRoot_);  // Should only has one and only one root.
            Base::hasRoot_ = true;
        }
    }

    void WriteIndent()  {
        size_t count = (Base::level_stack_.GetSize() / sizeof(typename Base::Level)) * indentCharCount_;
        PutN(*Base::os_, static_cast<typename OutputStream::Ch>(indentChar_), count);
    }

    Ch indentChar_;
    unsigned indentCharCount_;
    PrettyFormatOptions formatOptions_;

private:
    // Prohibit copy constructor & assignment operator.
    PrettyWriter(const PrettyWriter&);
    PrettyWriter& operator=(const PrettyWriter&);
};

CEREAL_RAPIDJSON_NAMESPACE_END

#if defined(__clang__)
CEREAL_RAPIDJSON_DIAG_POP
#endif

#ifdef __GNUC__
CEREAL_RAPIDJSON_DIAG_POP
#endif

#endif // CEREAL_RAPIDJSON_CEREAL_RAPIDJSON_H_
```
