## Supplemental code for "The motif composition of variable-number tandem repeats impacts gene expression": rapidjson.h

```
// Tencent is pleased to support the open source community by making RapidJSON available.
// 
// Copyright (C) 2015 THL A29 Limited, a Tencent company, and Milo Yip. All rights reserved.
//
// Licensed under the MIT License (the "License"); you may not use this file except
// in compliance with the License. You may obtain a copy of the License at
//
// http://opensource.org/licenses/MIT
//
// Unless required by applicable law or agreed to in writing, software distributed 
// under the License is distributed on an "AS IS" BASIS, WITHOUT WARRANTIES OR 
// CONDITIONS OF ANY KIND, either express or implied. See the License for the 
// specific language governing permissions and limitations under the License.

#ifndef CEREAL_RAPIDJSON_CEREAL_RAPIDJSON_H_
#define CEREAL_RAPIDJSON_CEREAL_RAPIDJSON_H_

/*!\file rapidjson.h
    \brief common definitions and configuration
    
    \see CEREAL_RAPIDJSON_CONFIG
 */

/*! \defgroup CEREAL_RAPIDJSON_CONFIG RapidJSON configuration
    \brief Configuration macros for library features

    Some RapidJSON features are configurable to adapt the library to a wide
    variety of platforms, environments and usage scenarios.  Most of the
    features can be configured in terms of overridden or predefined
    preprocessor macros at compile-time.

    Some additional customization is available in the \ref CEREAL_RAPIDJSON_ERRORS APIs.

    \note These macros should be given on the compiler command-line
          (where applicable)  to avoid inconsistent values when compiling
          different translation units of a single application.
 */

#include <cstdlib>  // malloc(), realloc(), free(), size_t
#include <cstring>  // memset(), memcpy(), memmove(), memcmp()

///////////////////////////////////////////////////////////////////////////////
// CEREAL_RAPIDJSON_VERSION_STRING
//
// ALWAYS synchronize the following 3 macros with corresponding variables in /CMakeLists.txt.
//

//!@cond CEREAL_RAPIDJSON_HIDDEN_FROM_DOXYGEN
// token stringification
#define CEREAL_RAPIDJSON_STRINGIFY(x) CEREAL_RAPIDJSON_DO_STRINGIFY(x)
#define CEREAL_RAPIDJSON_DO_STRINGIFY(x) #x

// token concatenation
#define CEREAL_RAPIDJSON_JOIN(X, Y) CEREAL_RAPIDJSON_DO_JOIN(X, Y)
#define CEREAL_RAPIDJSON_DO_JOIN(X, Y) CEREAL_RAPIDJSON_DO_JOIN2(X, Y)
#define CEREAL_RAPIDJSON_DO_JOIN2(X, Y) X##Y
//!@endcond

/*! \def CEREAL_RAPIDJSON_MAJOR_VERSION
    \ingroup CEREAL_RAPIDJSON_CONFIG
    \brief Major version of RapidJSON in integer.
*/
/*! \def CEREAL_RAPIDJSON_MINOR_VERSION
    \ingroup CEREAL_RAPIDJSON_CONFIG
    \brief Minor version of RapidJSON in integer.
*/
/*! \def CEREAL_RAPIDJSON_PATCH_VERSION
    \ingroup CEREAL_RAPIDJSON_CONFIG
    \brief Patch version of RapidJSON in integer.
*/
/*! \def CEREAL_RAPIDJSON_VERSION_STRING
    \ingroup CEREAL_RAPIDJSON_CONFIG
    \brief Version of RapidJSON in "<major>.<minor>.<patch>" string format.
*/
#define CEREAL_RAPIDJSON_MAJOR_VERSION 1
#define CEREAL_RAPIDJSON_MINOR_VERSION 1
#define CEREAL_RAPIDJSON_PATCH_VERSION 0
#define CEREAL_RAPIDJSON_VERSION_STRING \
    CEREAL_RAPIDJSON_STRINGIFY(CEREAL_RAPIDJSON_MAJOR_VERSION.CEREAL_RAPIDJSON_MINOR_VERSION.CEREAL_RAPIDJSON_PATCH_VERSION)

///////////////////////////////////////////////////////////////////////////////
// CEREAL_RAPIDJSON_NAMESPACE_(BEGIN|END)
/*! \def CEREAL_RAPIDJSON_NAMESPACE
    \ingroup CEREAL_RAPIDJSON_CONFIG
    \brief   provide custom rapidjson namespace

    In order to avoid symbol clashes and/or "One Definition Rule" errors
    between multiple inclusions of (different versions of) RapidJSON in
    a single binary, users can customize the name of the main RapidJSON
    namespace.

    In case of a single nesting level, defining \c CEREAL_RAPIDJSON_NAMESPACE
    to a custom name (e.g. \c MyRapidJSON) is sufficient.  If multiple
    levels are needed, both \ref CEREAL_RAPIDJSON_NAMESPACE_BEGIN and \ref
    CEREAL_RAPIDJSON_NAMESPACE_END need to be defined as well:

    \code
    // in some .cpp file
    #define CEREAL_RAPIDJSON_NAMESPACE my::rapidjson
    #define CEREAL_RAPIDJSON_NAMESPACE_BEGIN namespace my { namespace rapidjson {
    #define CEREAL_RAPIDJSON_NAMESPACE_END   } }
    #include "rapidjson/..."
    \endcode

    \see rapidjson
 */
/*! \def CEREAL_RAPIDJSON_NAMESPACE_BEGIN
    \ingroup CEREAL_RAPIDJSON_CONFIG
    \brief   provide custom rapidjson namespace (opening expression)
    \see CEREAL_RAPIDJSON_NAMESPACE
*/
/*! \def CEREAL_RAPIDJSON_NAMESPACE_END
    \ingroup CEREAL_RAPIDJSON_CONFIG
    \brief   provide custom rapidjson namespace (closing expression)
    \see CEREAL_RAPIDJSON_NAMESPACE
*/
#ifndef CEREAL_RAPIDJSON_NAMESPACE
#define CEREAL_RAPIDJSON_NAMESPACE rapidjson
#endif
#ifndef CEREAL_RAPIDJSON_NAMESPACE_BEGIN
#define CEREAL_RAPIDJSON_NAMESPACE_BEGIN namespace CEREAL_RAPIDJSON_NAMESPACE {
#endif
#ifndef CEREAL_RAPIDJSON_NAMESPACE_END
#define CEREAL_RAPIDJSON_NAMESPACE_END }
#endif

///////////////////////////////////////////////////////////////////////////////
// CEREAL_RAPIDJSON_HAS_STDSTRING

#ifndef CEREAL_RAPIDJSON_HAS_STDSTRING
#ifdef CEREAL_RAPIDJSON_DOXYGEN_RUNNING
#define CEREAL_RAPIDJSON_HAS_STDSTRING 1 // force generation of documentation
#else
#define CEREAL_RAPIDJSON_HAS_STDSTRING 0 // no std::string support by default
#endif
/*! \def CEREAL_RAPIDJSON_HAS_STDSTRING
    \ingroup CEREAL_RAPIDJSON_CONFIG
    \brief Enable RapidJSON support for \c std::string

    By defining this preprocessor symbol to \c 1, several convenience functions for using
    \ref rapidjson::GenericValue with \c std::string are enabled, especially
    for construction and comparison.

    \hideinitializer
*/
#endif // !defined(CEREAL_RAPIDJSON_HAS_STDSTRING)

#if CEREAL_RAPIDJSON_HAS_STDSTRING
#include <string>
#endif // CEREAL_RAPIDJSON_HAS_STDSTRING

///////////////////////////////////////////////////////////////////////////////
// CEREAL_RAPIDJSON_NO_INT64DEFINE

/*! \def CEREAL_RAPIDJSON_NO_INT64DEFINE
    \ingroup CEREAL_RAPIDJSON_CONFIG
    \brief Use external 64-bit integer types.

    RapidJSON requires the 64-bit integer types \c int64_t and  \c uint64_t types
    to be available at global scope.

    If users have their own definition, define CEREAL_RAPIDJSON_NO_INT64DEFINE to
    prevent RapidJSON from defining its own types.
*/
#ifndef CEREAL_RAPIDJSON_NO_INT64DEFINE
//!@cond CEREAL_RAPIDJSON_HIDDEN_FROM_DOXYGEN
#if defined(_MSC_VER) && (_MSC_VER < 1800)	// Visual Studio 2013
#include "msinttypes/stdint.h"
#include "msinttypes/inttypes.h"
#else
// Other compilers should have this.
#include <stdint.h>
#include <inttypes.h>
#endif
//!@endcond
#ifdef CEREAL_RAPIDJSON_DOXYGEN_RUNNING
#define CEREAL_RAPIDJSON_NO_INT64DEFINE
#endif
#endif // CEREAL_RAPIDJSON_NO_INT64TYPEDEF

///////////////////////////////////////////////////////////////////////////////
// CEREAL_RAPIDJSON_FORCEINLINE

#ifndef CEREAL_RAPIDJSON_FORCEINLINE
//!@cond CEREAL_RAPIDJSON_HIDDEN_FROM_DOXYGEN
#if defined(_MSC_VER) && defined(NDEBUG)
#define CEREAL_RAPIDJSON_FORCEINLINE __forceinline
#elif defined(__GNUC__) && __GNUC__ >= 4 && defined(NDEBUG)
#define CEREAL_RAPIDJSON_FORCEINLINE __attribute__((always_inline))
#else
#define CEREAL_RAPIDJSON_FORCEINLINE
#endif
//!@endcond
#endif // CEREAL_RAPIDJSON_FORCEINLINE

///////////////////////////////////////////////////////////////////////////////
// CEREAL_RAPIDJSON_ENDIAN
#define CEREAL_RAPIDJSON_LITTLEENDIAN  0   //!< Little endian machine
#define CEREAL_RAPIDJSON_BIGENDIAN     1   //!< Big endian machine

//! Endianness of the machine.
/*!
    \def CEREAL_RAPIDJSON_ENDIAN
    \ingroup CEREAL_RAPIDJSON_CONFIG

    GCC 4.6 provided macro for detecting endianness of the target machine. But other
    compilers may not have this. User can define CEREAL_RAPIDJSON_ENDIAN to either
    \ref CEREAL_RAPIDJSON_LITTLEENDIAN or \ref CEREAL_RAPIDJSON_BIGENDIAN.

    Default detection implemented with reference to
    \li https://gcc.gnu.org/onlinedocs/gcc-4.6.0/cpp/Common-Predefined-Macros.html
    \li http://www.boost.org/doc/libs/1_42_0/boost/detail/endian.hpp
*/
#ifndef CEREAL_RAPIDJSON_ENDIAN
// Detect with GCC 4.6's macro
#  ifdef __BYTE_ORDER__
#    if __BYTE_ORDER__ == __ORDER_LITTLE_ENDIAN__
#      define CEREAL_RAPIDJSON_ENDIAN CEREAL_RAPIDJSON_LITTLEENDIAN
#    elif __BYTE_ORDER__ == __ORDER_BIG_ENDIAN__
#      define CEREAL_RAPIDJSON_ENDIAN CEREAL_RAPIDJSON_BIGENDIAN
#    else
#      error Unknown machine endianness detected. User needs to define CEREAL_RAPIDJSON_ENDIAN.
#    endif // __BYTE_ORDER__
// Detect with GLIBC's endian.h
#  elif defined(__GLIBC__)
#    include <endian.h>
#    if (__BYTE_ORDER == __LITTLE_ENDIAN)
#      define CEREAL_RAPIDJSON_ENDIAN CEREAL_RAPIDJSON_LITTLEENDIAN
#    elif (__BYTE_ORDER == __BIG_ENDIAN)
#      define CEREAL_RAPIDJSON_ENDIAN CEREAL_RAPIDJSON_BIGENDIAN
#    else
#      error Unknown machine endianness detected. User needs to define CEREAL_RAPIDJSON_ENDIAN.
#   endif // __GLIBC__
// Detect with _LITTLE_ENDIAN and _BIG_ENDIAN macro
#  elif defined(_LITTLE_ENDIAN) && !defined(_BIG_ENDIAN)
#    define CEREAL_RAPIDJSON_ENDIAN CEREAL_RAPIDJSON_LITTLEENDIAN
#  elif defined(_BIG_ENDIAN) && !defined(_LITTLE_ENDIAN)
#    define CEREAL_RAPIDJSON_ENDIAN CEREAL_RAPIDJSON_BIGENDIAN
// Detect with architecture macros
#  elif defined(__sparc) || defined(__sparc__) || defined(_POWER) || defined(__powerpc__) || defined(__ppc__) || defined(__hpux) || defined(__hppa) || defined(_MIPSEB) || defined(_POWER) || defined(__s390__)
#    define CEREAL_RAPIDJSON_ENDIAN CEREAL_RAPIDJSON_BIGENDIAN
#  elif defined(__i386__) || defined(__alpha__) || defined(__ia64) || defined(__ia64__) || defined(_M_IX86) || defined(_M_IA64) || defined(_M_ALPHA) || defined(__amd64) || defined(__amd64__) || defined(_M_AMD64) || defined(__x86_64) || defined(__x86_64__) || defined(_M_X64) || defined(__bfin__)
#    define CEREAL_RAPIDJSON_ENDIAN CEREAL_RAPIDJSON_LITTLEENDIAN
#  elif defined(_MSC_VER) && (defined(_M_ARM) || defined(_M_ARM64))
#    define CEREAL_RAPIDJSON_ENDIAN CEREAL_RAPIDJSON_LITTLEENDIAN
#  elif defined(CEREAL_RAPIDJSON_DOXYGEN_RUNNING)
#    define CEREAL_RAPIDJSON_ENDIAN
#  else
#    error Unknown machine endianness detected. User needs to define CEREAL_RAPIDJSON_ENDIAN.   
#  endif
#endif // CEREAL_RAPIDJSON_ENDIAN

///////////////////////////////////////////////////////////////////////////////
// CEREAL_RAPIDJSON_64BIT

//! Whether using 64-bit architecture
#ifndef CEREAL_RAPIDJSON_64BIT
#if defined(__LP64__) || (defined(__x86_64__) && defined(__ILP32__)) || defined(_WIN64) || defined(__EMSCRIPTEN__)
#define CEREAL_RAPIDJSON_64BIT 1
#else
#define CEREAL_RAPIDJSON_64BIT 0
#endif
#endif // CEREAL_RAPIDJSON_64BIT

///////////////////////////////////////////////////////////////////////////////
// CEREAL_RAPIDJSON_ALIGN

//! Data alignment of the machine.
/*! \ingroup CEREAL_RAPIDJSON_CONFIG
    \param x pointer to align

    Some machines require strict data alignment. The default is 8 bytes.
    User can customize by defining the CEREAL_RAPIDJSON_ALIGN function macro.
*/
#ifndef CEREAL_RAPIDJSON_ALIGN
#define CEREAL_RAPIDJSON_ALIGN(x) (((x) + static_cast<size_t>(7u)) & ~static_cast<size_t>(7u))
#endif

///////////////////////////////////////////////////////////////////////////////
// CEREAL_RAPIDJSON_UINT64_C2

//! Construct a 64-bit literal by a pair of 32-bit integer.
/*!
    64-bit literal with or without ULL suffix is prone to compiler warnings.
    UINT64_C() is C macro which cause compilation problems.
    Use this macro to define 64-bit constants by a pair of 32-bit integer.
*/
#ifndef CEREAL_RAPIDJSON_UINT64_C2
#define CEREAL_RAPIDJSON_UINT64_C2(high32, low32) ((static_cast<uint64_t>(high32) << 32) | static_cast<uint64_t>(low32))
#endif

///////////////////////////////////////////////////////////////////////////////
// CEREAL_RAPIDJSON_48BITPOINTER_OPTIMIZATION

//! Use only lower 48-bit address for some pointers.
/*!
    \ingroup CEREAL_RAPIDJSON_CONFIG

    This optimization uses the fact that current X86-64 architecture only implement lower 48-bit virtual address.
    The higher 16-bit can be used for storing other data.
    \c GenericValue uses this optimization to reduce its size form 24 bytes to 16 bytes in 64-bit architecture.
*/
#ifndef CEREAL_RAPIDJSON_48BITPOINTER_OPTIMIZATION
#if defined(__amd64__) || defined(__amd64) || defined(__x86_64__) || defined(__x86_64) || defined(_M_X64) || defined(_M_AMD64)
#define CEREAL_RAPIDJSON_48BITPOINTER_OPTIMIZATION 1
#else
#define CEREAL_RAPIDJSON_48BITPOINTER_OPTIMIZATION 0
#endif
#endif // CEREAL_RAPIDJSON_48BITPOINTER_OPTIMIZATION

#if CEREAL_RAPIDJSON_48BITPOINTER_OPTIMIZATION == 1
#if CEREAL_RAPIDJSON_64BIT != 1
#error CEREAL_RAPIDJSON_48BITPOINTER_OPTIMIZATION can only be set to 1 when CEREAL_RAPIDJSON_64BIT=1
#endif
#define CEREAL_RAPIDJSON_SETPOINTER(type, p, x) (p = reinterpret_cast<type *>((reinterpret_cast<uintptr_t>(p) & static_cast<uintptr_t>(CEREAL_RAPIDJSON_UINT64_C2(0xFFFF0000, 0x00000000))) | reinterpret_cast<uintptr_t>(reinterpret_cast<const void*>(x))))
#define CEREAL_RAPIDJSON_GETPOINTER(type, p) (reinterpret_cast<type *>(reinterpret_cast<uintptr_t>(p) & static_cast<uintptr_t>(CEREAL_RAPIDJSON_UINT64_C2(0x0000FFFF, 0xFFFFFFFF))))
#else
#define CEREAL_RAPIDJSON_SETPOINTER(type, p, x) (p = (x))
#define CEREAL_RAPIDJSON_GETPOINTER(type, p) (p)
#endif

///////////////////////////////////////////////////////////////////////////////
// CEREAL_RAPIDJSON_SSE2/CEREAL_RAPIDJSON_SSE42/CEREAL_RAPIDJSON_NEON/CEREAL_RAPIDJSON_SIMD

/*! \def CEREAL_RAPIDJSON_SIMD
    \ingroup CEREAL_RAPIDJSON_CONFIG
    \brief Enable SSE2/SSE4.2/Neon optimization.

    RapidJSON supports optimized implementations for some parsing operations
    based on the SSE2, SSE4.2 or NEon SIMD extensions on modern Intel
    or ARM compatible processors.

    To enable these optimizations, three different symbols can be defined;
    \code
    // Enable SSE2 optimization.
    #define CEREAL_RAPIDJSON_SSE2

    // Enable SSE4.2 optimization.
    #define CEREAL_RAPIDJSON_SSE42
    \endcode

    // Enable ARM Neon optimization.
    #define CEREAL_RAPIDJSON_NEON
    \endcode

    \c CEREAL_RAPIDJSON_SSE42 takes precedence over SSE2, if both are defined.

    If any of these symbols is defined, RapidJSON defines the macro
    \c CEREAL_RAPIDJSON_SIMD to indicate the availability of the optimized code.
*/
#if defined(CEREAL_RAPIDJSON_SSE2) || defined(CEREAL_RAPIDJSON_SSE42) \
    || defined(CEREAL_RAPIDJSON_NEON) || defined(CEREAL_RAPIDJSON_DOXYGEN_RUNNING)
#define CEREAL_RAPIDJSON_SIMD
#endif

///////////////////////////////////////////////////////////////////////////////
// CEREAL_RAPIDJSON_NO_SIZETYPEDEFINE

#ifndef CEREAL_RAPIDJSON_NO_SIZETYPEDEFINE
/*! \def CEREAL_RAPIDJSON_NO_SIZETYPEDEFINE
    \ingroup CEREAL_RAPIDJSON_CONFIG
    \brief User-provided \c SizeType definition.

    In order to avoid using 32-bit size types for indexing strings and arrays,
    define this preprocessor symbol and provide the type rapidjson::SizeType
    before including RapidJSON:
    \code
    #define CEREAL_RAPIDJSON_NO_SIZETYPEDEFINE
    namespace rapidjson { typedef ::std::size_t SizeType; }
    #include "rapidjson/..."
    \endcode

    \see rapidjson::SizeType
*/
#ifdef CEREAL_RAPIDJSON_DOXYGEN_RUNNING
#define CEREAL_RAPIDJSON_NO_SIZETYPEDEFINE
#endif
CEREAL_RAPIDJSON_NAMESPACE_BEGIN
//! Size type (for string lengths, array sizes, etc.)
/*! RapidJSON uses 32-bit array/string indices even on 64-bit platforms,
    instead of using \c size_t. Users may override the SizeType by defining
    \ref CEREAL_RAPIDJSON_NO_SIZETYPEDEFINE.
*/
typedef unsigned SizeType;
CEREAL_RAPIDJSON_NAMESPACE_END
#endif

// always import std::size_t to rapidjson namespace
CEREAL_RAPIDJSON_NAMESPACE_BEGIN
using std::size_t;
CEREAL_RAPIDJSON_NAMESPACE_END

///////////////////////////////////////////////////////////////////////////////
// CEREAL_RAPIDJSON_ASSERT

//! Assertion.
/*! \ingroup CEREAL_RAPIDJSON_CONFIG
    By default, rapidjson uses C \c assert() for internal assertions.
    User can override it by defining CEREAL_RAPIDJSON_ASSERT(x) macro.

    \note Parsing errors are handled and can be customized by the
          \ref CEREAL_RAPIDJSON_ERRORS APIs.
*/
#ifndef CEREAL_RAPIDJSON_ASSERT
#include <cassert>
#define CEREAL_RAPIDJSON_ASSERT(x) assert(x)
#endif // CEREAL_RAPIDJSON_ASSERT

///////////////////////////////////////////////////////////////////////////////
// CEREAL_RAPIDJSON_STATIC_ASSERT

// Prefer C++11 static_assert, if available
#ifndef CEREAL_RAPIDJSON_STATIC_ASSERT
#if __cplusplus >= 201103L || ( defined(_MSC_VER) && _MSC_VER >= 1800 )
#define CEREAL_RAPIDJSON_STATIC_ASSERT(x) \
   static_assert(x, CEREAL_RAPIDJSON_STRINGIFY(x))
#endif // C++11
#endif // CEREAL_RAPIDJSON_STATIC_ASSERT

// Adopt C++03 implementation from boost
#ifndef CEREAL_RAPIDJSON_STATIC_ASSERT
#ifndef __clang__
//!@cond CEREAL_RAPIDJSON_HIDDEN_FROM_DOXYGEN
#endif
CEREAL_RAPIDJSON_NAMESPACE_BEGIN
template <bool x> struct STATIC_ASSERTION_FAILURE;
template <> struct STATIC_ASSERTION_FAILURE<true> { enum { value = 1 }; };
template <size_t x> struct StaticAssertTest {};
CEREAL_RAPIDJSON_NAMESPACE_END

#if defined(__GNUC__) || defined(__clang__)
#define CEREAL_RAPIDJSON_STATIC_ASSERT_UNUSED_ATTRIBUTE __attribute__((unused))
#else
#define CEREAL_RAPIDJSON_STATIC_ASSERT_UNUSED_ATTRIBUTE 
#endif
#ifndef __clang__
//!@endcond
#endif

/*! \def CEREAL_RAPIDJSON_STATIC_ASSERT
    \brief (Internal) macro to check for conditions at compile-time
    \param x compile-time condition
    \hideinitializer
 */
#define CEREAL_RAPIDJSON_STATIC_ASSERT(x) \
    typedef ::CEREAL_RAPIDJSON_NAMESPACE::StaticAssertTest< \
      sizeof(::CEREAL_RAPIDJSON_NAMESPACE::STATIC_ASSERTION_FAILURE<bool(x) >)> \
    CEREAL_RAPIDJSON_JOIN(StaticAssertTypedef, __LINE__) CEREAL_RAPIDJSON_STATIC_ASSERT_UNUSED_ATTRIBUTE
#endif // CEREAL_RAPIDJSON_STATIC_ASSERT

///////////////////////////////////////////////////////////////////////////////
// CEREAL_RAPIDJSON_LIKELY, CEREAL_RAPIDJSON_UNLIKELY

//! Compiler branching hint for expression with high probability to be true.
/*!
    \ingroup CEREAL_RAPIDJSON_CONFIG
    \param x Boolean expression likely to be true.
*/
#ifndef CEREAL_RAPIDJSON_LIKELY
#if defined(__GNUC__) || defined(__clang__)
#define CEREAL_RAPIDJSON_LIKELY(x) __builtin_expect(!!(x), 1)
#else
#define CEREAL_RAPIDJSON_LIKELY(x) (x)
#endif
#endif

//! Compiler branching hint for expression with low probability to be true.
/*!
    \ingroup CEREAL_RAPIDJSON_CONFIG
    \param x Boolean expression unlikely to be true.
*/
#ifndef CEREAL_RAPIDJSON_UNLIKELY
#if defined(__GNUC__) || defined(__clang__)
#define CEREAL_RAPIDJSON_UNLIKELY(x) __builtin_expect(!!(x), 0)
#else
#define CEREAL_RAPIDJSON_UNLIKELY(x) (x)
#endif
#endif

///////////////////////////////////////////////////////////////////////////////
// Helpers

//!@cond CEREAL_RAPIDJSON_HIDDEN_FROM_DOXYGEN

#define CEREAL_RAPIDJSON_MULTILINEMACRO_BEGIN do {  
#define CEREAL_RAPIDJSON_MULTILINEMACRO_END \
} while((void)0, 0)

// adopted from Boost
#define CEREAL_RAPIDJSON_VERSION_CODE(x,y,z) \
  (((x)*100000) + ((y)*100) + (z))

///////////////////////////////////////////////////////////////////////////////
// CEREAL_RAPIDJSON_DIAG_PUSH/POP, CEREAL_RAPIDJSON_DIAG_OFF

#if defined(__GNUC__)
#define CEREAL_RAPIDJSON_GNUC \
    CEREAL_RAPIDJSON_VERSION_CODE(__GNUC__,__GNUC_MINOR__,__GNUC_PATCHLEVEL__)
#endif

#if defined(__clang__) || (defined(CEREAL_RAPIDJSON_GNUC) && CEREAL_RAPIDJSON_GNUC >= CEREAL_RAPIDJSON_VERSION_CODE(4,2,0))

#define CEREAL_RAPIDJSON_PRAGMA(x) _Pragma(CEREAL_RAPIDJSON_STRINGIFY(x))
#define CEREAL_RAPIDJSON_DIAG_PRAGMA(x) CEREAL_RAPIDJSON_PRAGMA(GCC diagnostic x)
#define CEREAL_RAPIDJSON_DIAG_OFF(x) \
    CEREAL_RAPIDJSON_DIAG_PRAGMA(ignored CEREAL_RAPIDJSON_STRINGIFY(CEREAL_RAPIDJSON_JOIN(-W,x)))

// push/pop support in Clang and GCC>=4.6
#if defined(__clang__) || (defined(CEREAL_RAPIDJSON_GNUC) && CEREAL_RAPIDJSON_GNUC >= CEREAL_RAPIDJSON_VERSION_CODE(4,6,0))
#define CEREAL_RAPIDJSON_DIAG_PUSH CEREAL_RAPIDJSON_DIAG_PRAGMA(push)
#define CEREAL_RAPIDJSON_DIAG_POP  CEREAL_RAPIDJSON_DIAG_PRAGMA(pop)
#else // GCC >= 4.2, < 4.6
#define CEREAL_RAPIDJSON_DIAG_PUSH /* ignored */
#define CEREAL_RAPIDJSON_DIAG_POP /* ignored */
#endif

#elif defined(_MSC_VER)

// pragma (MSVC specific)
#define CEREAL_RAPIDJSON_PRAGMA(x) __pragma(x)
#define CEREAL_RAPIDJSON_DIAG_PRAGMA(x) CEREAL_RAPIDJSON_PRAGMA(warning(x))

#define CEREAL_RAPIDJSON_DIAG_OFF(x) CEREAL_RAPIDJSON_DIAG_PRAGMA(disable: x)
#define CEREAL_RAPIDJSON_DIAG_PUSH CEREAL_RAPIDJSON_DIAG_PRAGMA(push)
#define CEREAL_RAPIDJSON_DIAG_POP  CEREAL_RAPIDJSON_DIAG_PRAGMA(pop)

#else

#define CEREAL_RAPIDJSON_DIAG_OFF(x) /* ignored */
#define CEREAL_RAPIDJSON_DIAG_PUSH   /* ignored */
#define CEREAL_RAPIDJSON_DIAG_POP    /* ignored */

#endif // CEREAL_RAPIDJSON_DIAG_*

///////////////////////////////////////////////////////////////////////////////
// C++11 features

#ifndef CEREAL_RAPIDJSON_HAS_CXX11_RVALUE_REFS
#if defined(__clang__)
#if __has_feature(cxx_rvalue_references) && \
    (defined(_MSC_VER) || defined(_LIBCPP_VERSION) || defined(__GLIBCXX__) && __GLIBCXX__ >= 20080306)
#define CEREAL_RAPIDJSON_HAS_CXX11_RVALUE_REFS 1
#else
#define CEREAL_RAPIDJSON_HAS_CXX11_RVALUE_REFS 0
#endif
#elif (defined(CEREAL_RAPIDJSON_GNUC) && (CEREAL_RAPIDJSON_GNUC >= CEREAL_RAPIDJSON_VERSION_CODE(4,3,0)) && defined(__GXX_EXPERIMENTAL_CXX0X__)) || \
      (defined(_MSC_VER) && _MSC_VER >= 1600) || \
      (defined(__SUNPRO_CC) && __SUNPRO_CC >= 0x5140 && defined(__GXX_EXPERIMENTAL_CXX0X__))

#define CEREAL_RAPIDJSON_HAS_CXX11_RVALUE_REFS 1
#else
#define CEREAL_RAPIDJSON_HAS_CXX11_RVALUE_REFS 0
#endif
#endif // CEREAL_RAPIDJSON_HAS_CXX11_RVALUE_REFS

#ifndef CEREAL_RAPIDJSON_HAS_CXX11_NOEXCEPT
#if defined(__clang__)
#define CEREAL_RAPIDJSON_HAS_CXX11_NOEXCEPT __has_feature(cxx_noexcept)
#elif (defined(CEREAL_RAPIDJSON_GNUC) && (CEREAL_RAPIDJSON_GNUC >= CEREAL_RAPIDJSON_VERSION_CODE(4,6,0)) && defined(__GXX_EXPERIMENTAL_CXX0X__)) || \
    (defined(_MSC_VER) && _MSC_VER >= 1900) || \
    (defined(__SUNPRO_CC) && __SUNPRO_CC >= 0x5140 && defined(__GXX_EXPERIMENTAL_CXX0X__))
#define CEREAL_RAPIDJSON_HAS_CXX11_NOEXCEPT 1
#else
#define CEREAL_RAPIDJSON_HAS_CXX11_NOEXCEPT 0
#endif
#endif
#if CEREAL_RAPIDJSON_HAS_CXX11_NOEXCEPT
#define CEREAL_RAPIDJSON_NOEXCEPT noexcept
#else
#define CEREAL_RAPIDJSON_NOEXCEPT /* noexcept */
#endif // CEREAL_RAPIDJSON_HAS_CXX11_NOEXCEPT

// no automatic detection, yet
#ifndef CEREAL_RAPIDJSON_HAS_CXX11_TYPETRAITS
#if (defined(_MSC_VER) && _MSC_VER >= 1700)
#define CEREAL_RAPIDJSON_HAS_CXX11_TYPETRAITS 1
#else
#define CEREAL_RAPIDJSON_HAS_CXX11_TYPETRAITS 0
#endif
#endif

#ifndef CEREAL_RAPIDJSON_HAS_CXX11_RANGE_FOR
#if defined(__clang__)
#define CEREAL_RAPIDJSON_HAS_CXX11_RANGE_FOR __has_feature(cxx_range_for)
#elif (defined(CEREAL_RAPIDJSON_GNUC) && (CEREAL_RAPIDJSON_GNUC >= CEREAL_RAPIDJSON_VERSION_CODE(4,6,0)) && defined(__GXX_EXPERIMENTAL_CXX0X__)) || \
      (defined(_MSC_VER) && _MSC_VER >= 1700) || \
      (defined(__SUNPRO_CC) && __SUNPRO_CC >= 0x5140 && defined(__GXX_EXPERIMENTAL_CXX0X__))
#define CEREAL_RAPIDJSON_HAS_CXX11_RANGE_FOR 1
#else
#define CEREAL_RAPIDJSON_HAS_CXX11_RANGE_FOR 0
#endif
#endif // CEREAL_RAPIDJSON_HAS_CXX11_RANGE_FOR

//!@endcond

//! Assertion (in non-throwing contexts).
 /*! \ingroup CEREAL_RAPIDJSON_CONFIG
    Some functions provide a \c noexcept guarantee, if the compiler supports it.
    In these cases, the \ref CEREAL_RAPIDJSON_ASSERT macro cannot be overridden to
    throw an exception.  This macro adds a separate customization point for
    such cases.

    Defaults to C \c assert() (as \ref CEREAL_RAPIDJSON_ASSERT), if \c noexcept is
    supported, and to \ref CEREAL_RAPIDJSON_ASSERT otherwise.
 */

///////////////////////////////////////////////////////////////////////////////
// CEREAL_RAPIDJSON_NOEXCEPT_ASSERT

#ifndef CEREAL_RAPIDJSON_NOEXCEPT_ASSERT
#ifdef CEREAL_RAPIDJSON_ASSERT_THROWS
#if CEREAL_RAPIDJSON_HAS_CXX11_NOEXCEPT
#define CEREAL_RAPIDJSON_NOEXCEPT_ASSERT(x)
#else
#define CEREAL_RAPIDJSON_NOEXCEPT_ASSERT(x) CEREAL_RAPIDJSON_ASSERT(x)
#endif // CEREAL_RAPIDJSON_HAS_CXX11_NOEXCEPT
#else
#define CEREAL_RAPIDJSON_NOEXCEPT_ASSERT(x) CEREAL_RAPIDJSON_ASSERT(x)
#endif // CEREAL_RAPIDJSON_ASSERT_THROWS
#endif // CEREAL_RAPIDJSON_NOEXCEPT_ASSERT

///////////////////////////////////////////////////////////////////////////////
// new/delete

#ifndef CEREAL_RAPIDJSON_NEW
///! customization point for global \c new
#define CEREAL_RAPIDJSON_NEW(TypeName) new TypeName
#endif
#ifndef CEREAL_RAPIDJSON_DELETE
///! customization point for global \c delete
#define CEREAL_RAPIDJSON_DELETE(x) delete x
#endif

///////////////////////////////////////////////////////////////////////////////
// Type

/*! \namespace rapidjson
    \brief main RapidJSON namespace
    \see CEREAL_RAPIDJSON_NAMESPACE
*/
CEREAL_RAPIDJSON_NAMESPACE_BEGIN

//! Type of JSON value
enum Type {
    kNullType = 0,      //!< null
    kFalseType = 1,     //!< false
    kTrueType = 2,      //!< true
    kObjectType = 3,    //!< object
    kArrayType = 4,     //!< array 
    kStringType = 5,    //!< string
    kNumberType = 6     //!< number
};

CEREAL_RAPIDJSON_NAMESPACE_END

#endif // CEREAL_RAPIDJSON_CEREAL_RAPIDJSON_H_
```
