## Supplemental code for "The motif composition of variable-number tandem repeats impacts gene expression": reader.h

```
// Tencent is pleased to support the open source community by making RapidJSON available.
//
// Copyright (C) 2015 THL A29 Limited, a Tencent company, and Milo Yip. All rights reserved.
//
// Licensed under the MIT License (the "License"); you may not use this file except
// in compliance with the License. You may obtain a copy of the License at
//
// http://opensource.org/licenses/MIT
//
// Unless required by applicable law or agreed to in writing, software distributed
// under the License is distributed on an "AS IS" BASIS, WITHOUT WARRANTIES OR
// CONDITIONS OF ANY KIND, either express or implied. See the License for the
// specific language governing permissions and limitations under the License.

#ifndef CEREAL_RAPIDJSON_READER_H_
#define CEREAL_RAPIDJSON_READER_H_

/*! \file reader.h */

#include "allocators.h"
#include "stream.h"
#include "encodedstream.h"
#include "internal/meta.h"
#include "internal/stack.h"
#include "internal/strtod.h"
#include <limits>

#if defined(CEREAL_RAPIDJSON_SIMD) && defined(_MSC_VER)
#include <intrin.h>
#pragma intrinsic(_BitScanForward)
#endif
#ifdef CEREAL_RAPIDJSON_SSE42
#include <nmmintrin.h>
#elif defined(CEREAL_RAPIDJSON_SSE2)
#include <emmintrin.h>
#elif defined(CEREAL_RAPIDJSON_NEON)
#include <arm_neon.h>
#endif

#ifdef __clang__
CEREAL_RAPIDJSON_DIAG_PUSH
CEREAL_RAPIDJSON_DIAG_OFF(old-style-cast)
CEREAL_RAPIDJSON_DIAG_OFF(padded)
CEREAL_RAPIDJSON_DIAG_OFF(switch-enum)
#elif defined(_MSC_VER)
CEREAL_RAPIDJSON_DIAG_PUSH
CEREAL_RAPIDJSON_DIAG_OFF(4127)  // conditional expression is constant
CEREAL_RAPIDJSON_DIAG_OFF(4702)  // unreachable code
#endif

#ifdef __GNUC__
CEREAL_RAPIDJSON_DIAG_PUSH
CEREAL_RAPIDJSON_DIAG_OFF(effc++)
#endif

//!@cond CEREAL_RAPIDJSON_HIDDEN_FROM_DOXYGEN
#define CEREAL_RAPIDJSON_NOTHING /* deliberately empty */
#ifndef CEREAL_RAPIDJSON_PARSE_ERROR_EARLY_RETURN
#define CEREAL_RAPIDJSON_PARSE_ERROR_EARLY_RETURN(value) \
    CEREAL_RAPIDJSON_MULTILINEMACRO_BEGIN \
    if (CEREAL_RAPIDJSON_UNLIKELY(HasParseError())) { return value; } \
    CEREAL_RAPIDJSON_MULTILINEMACRO_END
#endif
#define CEREAL_RAPIDJSON_PARSE_ERROR_EARLY_RETURN_VOID \
    CEREAL_RAPIDJSON_PARSE_ERROR_EARLY_RETURN(CEREAL_RAPIDJSON_NOTHING)
//!@endcond

/*! \def CEREAL_RAPIDJSON_PARSE_ERROR_NORETURN
    \ingroup CEREAL_RAPIDJSON_ERRORS
    \brief Macro to indicate a parse error.
    \param parseErrorCode \ref rapidjson::ParseErrorCode of the error
    \param offset  position of the error in JSON input (\c size_t)

    This macros can be used as a customization point for the internal
    error handling mechanism of RapidJSON.

    A common usage model is to throw an exception instead of requiring the
    caller to explicitly check the \ref rapidjson::GenericReader::Parse's
    return value:

    \code
    #define CEREAL_RAPIDJSON_PARSE_ERROR_NORETURN(parseErrorCode,offset) \
       throw ParseException(parseErrorCode, #parseErrorCode, offset)

    #include <stdexcept>               // std::runtime_error
    #include "rapidjson/error/error.h" // rapidjson::ParseResult

    struct ParseException : std::runtime_error, rapidjson::ParseResult {
      ParseException(rapidjson::ParseErrorCode code, const char* msg, size_t offset)
        : std::runtime_error(msg), ParseResult(code, offset) {}
    };

    #include "rapidjson/reader.h"
    \endcode

    \see CEREAL_RAPIDJSON_PARSE_ERROR, rapidjson::GenericReader::Parse
 */
#ifndef CEREAL_RAPIDJSON_PARSE_ERROR_NORETURN
#define CEREAL_RAPIDJSON_PARSE_ERROR_NORETURN(parseErrorCode, offset) \
    CEREAL_RAPIDJSON_MULTILINEMACRO_BEGIN \
    CEREAL_RAPIDJSON_ASSERT(!HasParseError()); /* Error can only be assigned once */ \
    SetParseError(parseErrorCode, offset); \
    CEREAL_RAPIDJSON_MULTILINEMACRO_END
#endif

/*! \def CEREAL_RAPIDJSON_PARSE_ERROR
    \ingroup CEREAL_RAPIDJSON_ERRORS
    \brief (Internal) macro to indicate and handle a parse error.
    \param parseErrorCode \ref rapidjson::ParseErrorCode of the error
    \param offset  position of the error in JSON input (\c size_t)

    Invokes CEREAL_RAPIDJSON_PARSE_ERROR_NORETURN and stops the parsing.

    \see CEREAL_RAPIDJSON_PARSE_ERROR_NORETURN
    \hideinitializer
 */
#ifndef CEREAL_RAPIDJSON_PARSE_ERROR
#define CEREAL_RAPIDJSON_PARSE_ERROR(parseErrorCode, offset) \
    CEREAL_RAPIDJSON_MULTILINEMACRO_BEGIN \
    CEREAL_RAPIDJSON_PARSE_ERROR_NORETURN(parseErrorCode, offset); \
    CEREAL_RAPIDJSON_PARSE_ERROR_EARLY_RETURN_VOID; \
    CEREAL_RAPIDJSON_MULTILINEMACRO_END
#endif

#include "error/error.h" // ParseErrorCode, ParseResult

CEREAL_RAPIDJSON_NAMESPACE_BEGIN

///////////////////////////////////////////////////////////////////////////////
// ParseFlag

/*! \def CEREAL_RAPIDJSON_PARSE_DEFAULT_FLAGS
    \ingroup CEREAL_RAPIDJSON_CONFIG
    \brief User-defined kParseDefaultFlags definition.

    User can define this as any \c ParseFlag combinations.
*/
#ifndef CEREAL_RAPIDJSON_PARSE_DEFAULT_FLAGS
#define CEREAL_RAPIDJSON_PARSE_DEFAULT_FLAGS kParseNoFlags
#endif

//! Combination of parseFlags
/*! \see Reader::Parse, Document::Parse, Document::ParseInsitu, Document::ParseStream
 */
enum ParseFlag {
    kParseNoFlags = 0,              //!< No flags are set.
    kParseInsituFlag = 1,           //!< In-situ(destructive) parsing.
    kParseValidateEncodingFlag = 2, //!< Validate encoding of JSON strings.
    kParseIterativeFlag = 4,        //!< Iterative(constant complexity in terms of function call stack size) parsing.
    kParseStopWhenDoneFlag = 8,     //!< After parsing a complete JSON root from stream, stop further processing the rest of stream. When this flag is used, parser will not generate kParseErrorDocumentRootNotSingular error.
    kParseFullPrecisionFlag = 16,   //!< Parse number in full precision (but slower).
    kParseCommentsFlag = 32,        //!< Allow one-line (//) and multi-line (/**/) comments.
    kParseNumbersAsStringsFlag = 64,    //!< Parse all numbers (ints/doubles) as strings.
    kParseTrailingCommasFlag = 128, //!< Allow trailing commas at the end of objects and arrays.
    kParseNanAndInfFlag = 256,      //!< Allow parsing NaN, Inf, Infinity, -Inf and -Infinity as doubles.
    kParseDefaultFlags = CEREAL_RAPIDJSON_PARSE_DEFAULT_FLAGS  //!< Default parse flags. Can be customized by defining CEREAL_RAPIDJSON_PARSE_DEFAULT_FLAGS
};

///////////////////////////////////////////////////////////////////////////////
// Handler

/*! \class rapidjson::Handler
    \brief Concept for receiving events from GenericReader upon parsing.
    The functions return true if no error occurs. If they return false,
    the event publisher should terminate the process.
\code
concept Handler {
    typename Ch;

    bool Null();
    bool Bool(bool b);
    bool Int(int i);
    bool Uint(unsigned i);
    bool Int64(int64_t i);
    bool Uint64(uint64_t i);
    bool Double(double d);
    /// enabled via kParseNumbersAsStringsFlag, string is not null-terminated (use length)
    bool RawNumber(const Ch* str, SizeType length, bool copy);
    bool String(const Ch* str, SizeType length, bool copy);
    bool StartObject();
    bool Key(const Ch* str, SizeType length, bool copy);
    bool EndObject(SizeType memberCount);
    bool StartArray();
    bool EndArray(SizeType elementCount);
};
\endcode
*/
///////////////////////////////////////////////////////////////////////////////
// BaseReaderHandler

//! Default implementation of Handler.
/*! This can be used as base class of any reader handler.
    \note implements Handler concept
*/
template<typename Encoding = UTF8<>, typename Derived = void>
struct BaseReaderHandler {
    typedef typename Encoding::Ch Ch;

    typedef typename internal::SelectIf<internal::IsSame<Derived, void>, BaseReaderHandler, Derived>::Type Override;

    bool Default() { return true; }
    bool Null() { return static_cast<Override&>(*this).Default(); }
    bool Bool(bool) { return static_cast<Override&>(*this).Default(); }
    bool Int(int) { return static_cast<Override&>(*this).Default(); }
    bool Uint(unsigned) { return static_cast<Override&>(*this).Default(); }
    bool Int64(int64_t) { return static_cast<Override&>(*this).Default(); }
    bool Uint64(uint64_t) { return static_cast<Override&>(*this).Default(); }
    bool Double(double) { return static_cast<Override&>(*this).Default(); }
    /// enabled via kParseNumbersAsStringsFlag, string is not null-terminated (use length)
    bool RawNumber(const Ch* str, SizeType len, bool copy) { return static_cast<Override&>(*this).String(str, len, copy); }
    bool String(const Ch*, SizeType, bool) { return static_cast<Override&>(*this).Default(); }
    bool StartObject() { return static_cast<Override&>(*this).Default(); }
    bool Key(const Ch* str, SizeType len, bool copy) { return static_cast<Override&>(*this).String(str, len, copy); }
    bool EndObject(SizeType) { return static_cast<Override&>(*this).Default(); }
    bool StartArray() { return static_cast<Override&>(*this).Default(); }
    bool EndArray(SizeType) { return static_cast<Override&>(*this).Default(); }
};

///////////////////////////////////////////////////////////////////////////////
// StreamLocalCopy

namespace internal {

template<typename Stream, int = StreamTraits<Stream>::copyOptimization>
class StreamLocalCopy;

//! Do copy optimization.
template<typename Stream>
class StreamLocalCopy<Stream, 1> {
public:
    StreamLocalCopy(Stream& original) : s(original), original_(original) {}
    ~StreamLocalCopy() { original_ = s; }

    Stream s;

private:
    StreamLocalCopy& operator=(const StreamLocalCopy&) /* = delete */;

    Stream& original_;
};

//! Keep reference.
template<typename Stream>
class StreamLocalCopy<Stream, 0> {
public:
    StreamLocalCopy(Stream& original) : s(original) {}

    Stream& s;

private:
    StreamLocalCopy& operator=(const StreamLocalCopy&) /* = delete */;
};

} // namespace internal

///////////////////////////////////////////////////////////////////////////////
// SkipWhitespace

//! Skip the JSON white spaces in a stream.
/*! \param is A input stream for skipping white spaces.
    \note This function has SSE2/SSE4.2 specialization.
*/
template<typename InputStream>
void SkipWhitespace(InputStream& is) {
    internal::StreamLocalCopy<InputStream> copy(is);
    InputStream& s(copy.s);

    typename InputStream::Ch c;
    while ((c = s.Peek()) == ' ' || c == '\n' || c == '\r' || c == '\t')
        s.Take();
}

inline const char* SkipWhitespace(const char* p, const char* end) {
    while (p != end && (*p == ' ' || *p == '\n' || *p == '\r' || *p == '\t'))
        ++p;
    return p;
}

#ifdef CEREAL_RAPIDJSON_SSE42
//! Skip whitespace with SSE 4.2 pcmpistrm instruction, testing 16 8-byte characters at once.
inline const char *SkipWhitespace_SIMD(const char* p) {
    // Fast return for single non-whitespace
    if (*p == ' ' || *p == '\n' || *p == '\r' || *p == '\t')
        ++p;
    else
        return p;

    // 16-byte align to the next boundary
    const char* nextAligned = reinterpret_cast<const char*>((reinterpret_cast<size_t>(p) + 15) & static_cast<size_t>(~15));
    while (p != nextAligned)
        if (*p == ' ' || *p == '\n' || *p == '\r' || *p == '\t')
            ++p;
        else
            return p;

    // The rest of string using SIMD
    static const char whitespace[16] = " \n\r\t";
    const __m128i w = _mm_loadu_si128(reinterpret_cast<const __m128i *>(&whitespace[0]));

    for (;; p += 16) {
        const __m128i s = _mm_load_si128(reinterpret_cast<const __m128i *>(p));
        const int r = _mm_cmpistri(w, s, _SIDD_UBYTE_OPS | _SIDD_CMP_EQUAL_ANY | _SIDD_LEAST_SIGNIFICANT | _SIDD_NEGATIVE_POLARITY);
        if (r != 16)    // some of characters is non-whitespace
            return p + r;
    }
}

inline const char *SkipWhitespace_SIMD(const char* p, const char* end) {
    // Fast return for single non-whitespace
    if (p != end && (*p == ' ' || *p == '\n' || *p == '\r' || *p == '\t'))
        ++p;
    else
        return p;

    // The middle of string using SIMD
    static const char whitespace[16] = " \n\r\t";
    const __m128i w = _mm_loadu_si128(reinterpret_cast<const __m128i *>(&whitespace[0]));

    for (; p <= end - 16; p += 16) {
        const __m128i s = _mm_loadu_si128(reinterpret_cast<const __m128i *>(p));
        const int r = _mm_cmpistri(w, s, _SIDD_UBYTE_OPS | _SIDD_CMP_EQUAL_ANY | _SIDD_LEAST_SIGNIFICANT | _SIDD_NEGATIVE_POLARITY);
        if (r != 16)    // some of characters is non-whitespace
            return p + r;
    }

    return SkipWhitespace(p, end);
}

#elif defined(CEREAL_RAPIDJSON_SSE2)

//! Skip whitespace with SSE2 instructions, testing 16 8-byte characters at once.
inline const char *SkipWhitespace_SIMD(const char* p) {
    // Fast return for single non-whitespace
    if (*p == ' ' || *p == '\n' || *p == '\r' || *p == '\t')
        ++p;
    else
        return p;

    // 16-byte align to the next boundary
    const char* nextAligned = reinterpret_cast<const char*>((reinterpret_cast<size_t>(p) + 15) & static_cast<size_t>(~15));
    while (p != nextAligned)
        if (*p == ' ' || *p == '\n' || *p == '\r' || *p == '\t')
            ++p;
        else
            return p;

    // The rest of string
    #define C16(c) { c, c, c, c, c, c, c, c, c, c, c, c, c, c, c, c }
    static const char whitespaces[4][16] = { C16(' '), C16('\n'), C16('\r'), C16('\t') };
    #undef C16

    const __m128i w0 = _mm_loadu_si128(reinterpret_cast<const __m128i *>(&whitespaces[0][0]));
    const __m128i w1 = _mm_loadu_si128(reinterpret_cast<const __m128i *>(&whitespaces[1][0]));
    const __m128i w2 = _mm_loadu_si128(reinterpret_cast<const __m128i *>(&whitespaces[2][0]));
    const __m128i w3 = _mm_loadu_si128(reinterpret_cast<const __m128i *>(&whitespaces[3][0]));

    for (;; p += 16) {
        const __m128i s = _mm_load_si128(reinterpret_cast<const __m128i *>(p));
        __m128i x = _mm_cmpeq_epi8(s, w0);
        x = _mm_or_si128(x, _mm_cmpeq_epi8(s, w1));
        x = _mm_or_si128(x, _mm_cmpeq_epi8(s, w2));
        x = _mm_or_si128(x, _mm_cmpeq_epi8(s, w3));
        unsigned short r = static_cast<unsigned short>(~_mm_movemask_epi8(x));
        if (r != 0) {   // some of characters may be non-whitespace
#ifdef _MSC_VER         // Find the index of first non-whitespace
            unsigned long offset;
            _BitScanForward(&offset, r);
            return p + offset;
#else
            return p + __builtin_ffs(r) - 1;
#endif
        }
    }
}

inline const char *SkipWhitespace_SIMD(const char* p, const char* end) {
    // Fast return for single non-whitespace
    if (p != end && (*p == ' ' || *p == '\n' || *p == '\r' || *p == '\t'))
        ++p;
    else
        return p;

    // The rest of string
    #define C16(c) { c, c, c, c, c, c, c, c, c, c, c, c, c, c, c, c }
    static const char whitespaces[4][16] = { C16(' '), C16('\n'), C16('\r'), C16('\t') };
    #undef C16

    const __m128i w0 = _mm_loadu_si128(reinterpret_cast<const __m128i *>(&whitespaces[0][0]));
    const __m128i w1 = _mm_loadu_si128(reinterpret_cast<const __m128i *>(&whitespaces[1][0]));
    const __m128i w2 = _mm_loadu_si128(reinterpret_cast<const __m128i *>(&whitespaces[2][0]));
    const __m128i w3 = _mm_loadu_si128(reinterpret_cast<const __m128i *>(&whitespaces[3][0]));

    for (; p <= end - 16; p += 16) {
        const __m128i s = _mm_loadu_si128(reinterpret_cast<const __m128i *>(p));
        __m128i x = _mm_cmpeq_epi8(s, w0);
        x = _mm_or_si128(x, _mm_cmpeq_epi8(s, w1));
        x = _mm_or_si128(x, _mm_cmpeq_epi8(s, w2));
        x = _mm_or_si128(x, _mm_cmpeq_epi8(s, w3));
        unsigned short r = static_cast<unsigned short>(~_mm_movemask_epi8(x));
        if (r != 0) {   // some of characters may be non-whitespace
#ifdef _MSC_VER         // Find the index of first non-whitespace
            unsigned long offset;
            _BitScanForward(&offset, r);
            return p + offset;
#else
            return p + __builtin_ffs(r) - 1;
#endif
        }
    }

    return SkipWhitespace(p, end);
}

#elif defined(CEREAL_RAPIDJSON_NEON)

//! Skip whitespace with ARM Neon instructions, testing 16 8-byte characters at once.
inline const char *SkipWhitespace_SIMD(const char* p) {
    // Fast return for single non-whitespace
    if (*p == ' ' || *p == '\n' || *p == '\r' || *p == '\t')
        ++p;
    else
        return p;

    // 16-byte align to the next boundary
    const char* nextAligned = reinterpret_cast<const char*>((reinterpret_cast<size_t>(p) + 15) & static_cast<size_t>(~15));
    while (p != nextAligned)
        if (*p == ' ' || *p == '\n' || *p == '\r' || *p == '\t')
            ++p;
        else
            return p;

    const uint8x16_t w0 = vmovq_n_u8(' ');
    const uint8x16_t w1 = vmovq_n_u8('\n');
    const uint8x16_t w2 = vmovq_n_u8('\r');
    const uint8x16_t w3 = vmovq_n_u8('\t');

    for (;; p += 16) {
        const uint8x16_t s = vld1q_u8(reinterpret_cast<const uint8_t *>(p));
        uint8x16_t x = vceqq_u8(s, w0);
        x = vorrq_u8(x, vceqq_u8(s, w1));
        x = vorrq_u8(x, vceqq_u8(s, w2));
        x = vorrq_u8(x, vceqq_u8(s, w3));

        x = vmvnq_u8(x);                       // Negate
        x = vrev64q_u8(x);                     // Rev in 64
        uint64_t low = vgetq_lane_u64(reinterpret_cast<uint64x2_t>(x), 0);   // extract
        uint64_t high = vgetq_lane_u64(reinterpret_cast<uint64x2_t>(x), 1);  // extract

        if (low == 0) {
            if (high != 0) {
                int lz =__builtin_clzll(high);;
                return p + 8 + (lz >> 3);
            }
        } else {
            int lz = __builtin_clzll(low);;
            return p + (lz >> 3);
        }
    }
}

inline const char *SkipWhitespace_SIMD(const char* p, const char* end) {
    // Fast return for single non-whitespace
    if (p != end && (*p == ' ' || *p == '\n' || *p == '\r' || *p == '\t'))
        ++p;
    else
        return p;

    const uint8x16_t w0 = vmovq_n_u8(' ');
    const uint8x16_t w1 = vmovq_n_u8('\n');
    const uint8x16_t w2 = vmovq_n_u8('\r');
    const uint8x16_t w3 = vmovq_n_u8('\t');

    for (; p <= end - 16; p += 16) {
        const uint8x16_t s = vld1q_u8(reinterpret_cast<const uint8_t *>(p));
        uint8x16_t x = vceqq_u8(s, w0);
        x = vorrq_u8(x, vceqq_u8(s, w1));
        x = vorrq_u8(x, vceqq_u8(s, w2));
        x = vorrq_u8(x, vceqq_u8(s, w3));

        x = vmvnq_u8(x);                       // Negate
        x = vrev64q_u8(x);                     // Rev in 64
        uint64_t low = vgetq_lane_u64(reinterpret_cast<uint64x2_t>(x), 0);   // extract
        uint64_t high = vgetq_lane_u64(reinterpret_cast<uint64x2_t>(x), 1);  // extract

        if (low == 0) {
            if (high != 0) {
                int lz = __builtin_clzll(high);
                return p + 8 + (lz >> 3);
            }
        } else {
            int lz = __builtin_clzll(low);
            return p + (lz >> 3);
        }
    }

    return SkipWhitespace(p, end);
}

#endif // CEREAL_RAPIDJSON_NEON

#ifdef CEREAL_RAPIDJSON_SIMD
//! Template function specialization for InsituStringStream
template<> inline void SkipWhitespace(InsituStringStream& is) {
    is.src_ = const_cast<char*>(SkipWhitespace_SIMD(is.src_));
}

//! Template function specialization for StringStream
template<> inline void SkipWhitespace(StringStream& is) {
    is.src_ = SkipWhitespace_SIMD(is.src_);
}

template<> inline void SkipWhitespace(EncodedInputStream<UTF8<>, MemoryStream>& is) {
    is.is_.src_ = SkipWhitespace_SIMD(is.is_.src_, is.is_.end_);
}
#endif // CEREAL_RAPIDJSON_SIMD

///////////////////////////////////////////////////////////////////////////////
// GenericReader

//! SAX-style JSON parser. Use \ref Reader for UTF8 encoding and default allocator.
/*! GenericReader parses JSON text from a stream, and send events synchronously to an
    object implementing Handler concept.

    It needs to allocate a stack for storing a single decoded string during
    non-destructive parsing.

    For in-situ parsing, the decoded string is directly written to the source
    text string, no temporary buffer is required.

    A GenericReader object can be reused for parsing multiple JSON text.

    \tparam SourceEncoding Encoding of the input stream.
    \tparam TargetEncoding Encoding of the parse output.
    \tparam StackAllocator Allocator type for stack.
*/
template <typename SourceEncoding, typename TargetEncoding, typename StackAllocator = CrtAllocator>
class GenericReader {
public:
    typedef typename SourceEncoding::Ch Ch; //!< SourceEncoding character type

    //! Constructor.
    /*! \param stackAllocator Optional allocator for allocating stack memory. (Only use for non-destructive parsing)
        \param stackCapacity stack capacity in bytes for storing a single decoded string.  (Only use for non-destructive parsing)
    */
    GenericReader(StackAllocator* stackAllocator = 0, size_t stackCapacity = kDefaultStackCapacity) :
        stack_(stackAllocator, stackCapacity), parseResult_(), state_(IterativeParsingStartState) {}

    //! Parse JSON text.
    /*! \tparam parseFlags Combination of \ref ParseFlag.
        \tparam InputStream Type of input stream, implementing Stream concept.
        \tparam Handler Type of handler, implementing Handler concept.
        \param is Input stream to be parsed.
        \param handler The handler to receive events.
        \return Whether the parsing is successful.
    */
    template <unsigned parseFlags, typename InputStream, typename Handler>
    ParseResult Parse(InputStream& is, Handler& handler) {
        if (parseFlags & kParseIterativeFlag)
            return IterativeParse<parseFlags>(is, handler);

        parseResult_.Clear();

        ClearStackOnExit scope(*this);

        SkipWhitespaceAndComments<parseFlags>(is);
        CEREAL_RAPIDJSON_PARSE_ERROR_EARLY_RETURN(parseResult_);

        if (CEREAL_RAPIDJSON_UNLIKELY(is.Peek() == '\0')) {
            CEREAL_RAPIDJSON_PARSE_ERROR_NORETURN(kParseErrorDocumentEmpty, is.Tell());
            CEREAL_RAPIDJSON_PARSE_ERROR_EARLY_RETURN(parseResult_);
        }
        else {
            ParseValue<parseFlags>(is, handler);
            CEREAL_RAPIDJSON_PARSE_ERROR_EARLY_RETURN(parseResult_);

            if (!(parseFlags & kParseStopWhenDoneFlag)) {
                SkipWhitespaceAndComments<parseFlags>(is);
                CEREAL_RAPIDJSON_PARSE_ERROR_EARLY_RETURN(parseResult_);

                if (CEREAL_RAPIDJSON_UNLIKELY(is.Peek() != '\0')) {
                    CEREAL_RAPIDJSON_PARSE_ERROR_NORETURN(kParseErrorDocumentRootNotSingular, is.Tell());
                    CEREAL_RAPIDJSON_PARSE_ERROR_EARLY_RETURN(parseResult_);
                }
            }
        }

        return parseResult_;
    }

    //! Parse JSON text (with \ref kParseDefaultFlags)
    /*! \tparam InputStream Type of input stream, implementing Stream concept
        \tparam Handler Type of handler, implementing Handler concept.
        \param is Input stream to be parsed.
        \param handler The handler to receive events.
        \return Whether the parsing is successful.
    */
    template <typename InputStream, typename Handler>
    ParseResult Parse(InputStream& is, Handler& handler) {
        return Parse<kParseDefaultFlags>(is, handler);
    }

    //! Initialize JSON text token-by-token parsing
    /*!
     */
    void IterativeParseInit() {
        parseResult_.Clear();
        state_ = IterativeParsingStartState;
    }

    //! Parse one token from JSON text
    /*! \tparam InputStream Type of input stream, implementing Stream concept
        \tparam Handler Type of handler, implementing Handler concept.
        \param is Input stream to be parsed.
        \param handler The handler to receive events.
        \return Whether the parsing is successful.
     */
    template <unsigned parseFlags, typename InputStream, typename Handler>
    bool IterativeParseNext(InputStream& is, Handler& handler) {
        while (CEREAL_RAPIDJSON_LIKELY(is.Peek() != '\0')) {
            SkipWhitespaceAndComments<parseFlags>(is);

            Token t = Tokenize(is.Peek());
            IterativeParsingState n = Predict(state_, t);
            IterativeParsingState d = Transit<parseFlags>(state_, t, n, is, handler);

            // If we've finished or hit an error...
            if (CEREAL_RAPIDJSON_UNLIKELY(IsIterativeParsingCompleteState(d))) {
                // Report errors.
                if (d == IterativeParsingErrorState) {
                    HandleError(state_, is);
                    return false;
                }

                // Transition to the finish state.
                CEREAL_RAPIDJSON_ASSERT(d == IterativeParsingFinishState);
                state_ = d;

                // If StopWhenDone is not set...
                if (!(parseFlags & kParseStopWhenDoneFlag)) {
                    // ... and extra non-whitespace data is found...
                    SkipWhitespaceAndComments<parseFlags>(is);
                    if (is.Peek() != '\0') {
                        // ... this is considered an error.
                        HandleError(state_, is);
                        return false;
                    }
                }

                // Success! We are done!
                return true;
            }

            // Transition to the new state.
            state_ = d;

            // If we parsed anything other than a delimiter, we invoked the handler, so we can return true now.
            if (!IsIterativeParsingDelimiterState(n))
                return true;
        }

        // We reached the end of file.
        stack_.Clear();

        if (state_ != IterativeParsingFinishState) {
            HandleError(state_, is);
            return false;
        }

        return true;
    }

    //! Check if token-by-token parsing JSON text is complete
    /*! \return Whether the JSON has been fully decoded.
     */
    CEREAL_RAPIDJSON_FORCEINLINE bool IterativeParseComplete() const {
        return IsIterativeParsingCompleteState(state_);
    }

    //! Whether a parse error has occurred in the last parsing.
    bool HasParseError() const { return parseResult_.IsError(); }

    //! Get the \ref ParseErrorCode of last parsing.
    ParseErrorCode GetParseErrorCode() const { return parseResult_.Code(); }

    //! Get the position of last parsing error in input, 0 otherwise.
    size_t GetErrorOffset() const { return parseResult_.Offset(); }

protected:
    void SetParseError(ParseErrorCode code, size_t offset) { parseResult_.Set(code, offset); }

private:
    // Prohibit copy constructor & assignment operator.
    GenericReader(const GenericReader&);
    GenericReader& operator=(const GenericReader&);

    void ClearStack() { stack_.Clear(); }

    // clear stack on any exit from ParseStream, e.g. due to exception
    struct ClearStackOnExit {
        explicit ClearStackOnExit(GenericReader& r) : r_(r) {}
        ~ClearStackOnExit() { r_.ClearStack(); }
    private:
        GenericReader& r_;
        ClearStackOnExit(const ClearStackOnExit&);
        ClearStackOnExit& operator=(const ClearStackOnExit&);
    };

    template<unsigned parseFlags, typename InputStream>
    void SkipWhitespaceAndComments(InputStream& is) {
        SkipWhitespace(is);

        if (parseFlags & kParseCommentsFlag) {
            while (CEREAL_RAPIDJSON_UNLIKELY(Consume(is, '/'))) {
                if (Consume(is, '*')) {
                    while (true) {
                        if (CEREAL_RAPIDJSON_UNLIKELY(is.Peek() == '\0'))
                            CEREAL_RAPIDJSON_PARSE_ERROR(kParseErrorUnspecificSyntaxError, is.Tell());
                        else if (Consume(is, '*')) {
                            if (Consume(is, '/'))
                                break;
                        }
                        else
                            is.Take();
                    }
                }
                else if (CEREAL_RAPIDJSON_LIKELY(Consume(is, '/')))
                    while (is.Peek() != '\0' && is.Take() != '\n') {}
                else
                    CEREAL_RAPIDJSON_PARSE_ERROR(kParseErrorUnspecificSyntaxError, is.Tell());

                SkipWhitespace(is);
            }
        }
    }

    // Parse object: { string : value, ... }
    template<unsigned parseFlags, typename InputStream, typename Handler>
    void ParseObject(InputStream& is, Handler& handler) {
        CEREAL_RAPIDJSON_ASSERT(is.Peek() == '{');
        is.Take();  // Skip '{'

        if (CEREAL_RAPIDJSON_UNLIKELY(!handler.StartObject()))
            CEREAL_RAPIDJSON_PARSE_ERROR(kParseErrorTermination, is.Tell());

        SkipWhitespaceAndComments<parseFlags>(is);
        CEREAL_RAPIDJSON_PARSE_ERROR_EARLY_RETURN_VOID;

        if (Consume(is, '}')) {
            if (CEREAL_RAPIDJSON_UNLIKELY(!handler.EndObject(0)))  // empty object
                CEREAL_RAPIDJSON_PARSE_ERROR(kParseErrorTermination, is.Tell());
            return;
        }

        for (SizeType memberCount = 0;;) {
            if (CEREAL_RAPIDJSON_UNLIKELY(is.Peek() != '"'))
                CEREAL_RAPIDJSON_PARSE_ERROR(kParseErrorObjectMissName, is.Tell());

            ParseString<parseFlags>(is, handler, true);
            CEREAL_RAPIDJSON_PARSE_ERROR_EARLY_RETURN_VOID;

            SkipWhitespaceAndComments<parseFlags>(is);
            CEREAL_RAPIDJSON_PARSE_ERROR_EARLY_RETURN_VOID;

            if (CEREAL_RAPIDJSON_UNLIKELY(!Consume(is, ':')))
                CEREAL_RAPIDJSON_PARSE_ERROR(kParseErrorObjectMissColon, is.Tell());

            SkipWhitespaceAndComments<parseFlags>(is);
            CEREAL_RAPIDJSON_PARSE_ERROR_EARLY_RETURN_VOID;

            ParseValue<parseFlags>(is, handler);
            CEREAL_RAPIDJSON_PARSE_ERROR_EARLY_RETURN_VOID;

            SkipWhitespaceAndComments<parseFlags>(is);
            CEREAL_RAPIDJSON_PARSE_ERROR_EARLY_RETURN_VOID;

            ++memberCount;

            switch (is.Peek()) {
                case ',':
                    is.Take();
                    SkipWhitespaceAndComments<parseFlags>(is);
                    CEREAL_RAPIDJSON_PARSE_ERROR_EARLY_RETURN_VOID;
                    break;
                case '}':
                    is.Take();
                    if (CEREAL_RAPIDJSON_UNLIKELY(!handler.EndObject(memberCount)))
                        CEREAL_RAPIDJSON_PARSE_ERROR(kParseErrorTermination, is.Tell());
                    return;
                default:
                    CEREAL_RAPIDJSON_PARSE_ERROR(kParseErrorObjectMissCommaOrCurlyBracket, is.Tell()); break; // This useless break is only for making warning and coverage happy
            }

            if (parseFlags & kParseTrailingCommasFlag) {
                if (is.Peek() == '}') {
                    if (CEREAL_RAPIDJSON_UNLIKELY(!handler.EndObject(memberCount)))
                        CEREAL_RAPIDJSON_PARSE_ERROR(kParseErrorTermination, is.Tell());
                    is.Take();
                    return;
                }
            }
        }
    }

    // Parse array: [ value, ... ]
    template<unsigned parseFlags, typename InputStream, typename Handler>
    void ParseArray(InputStream& is, Handler& handler) {
        CEREAL_RAPIDJSON_ASSERT(is.Peek() == '[');
        is.Take();  // Skip '['

        if (CEREAL_RAPIDJSON_UNLIKELY(!handler.StartArray()))
            CEREAL_RAPIDJSON_PARSE_ERROR(kParseErrorTermination, is.Tell());

        SkipWhitespaceAndComments<parseFlags>(is);
        CEREAL_RAPIDJSON_PARSE_ERROR_EARLY_RETURN_VOID;

        if (Consume(is, ']')) {
            if (CEREAL_RAPIDJSON_UNLIKELY(!handler.EndArray(0))) // empty array
                CEREAL_RAPIDJSON_PARSE_ERROR(kParseErrorTermination, is.Tell());
            return;
        }

        for (SizeType elementCount = 0;;) {
            ParseValue<parseFlags>(is, handler);
            CEREAL_RAPIDJSON_PARSE_ERROR_EARLY_RETURN_VOID;

            ++elementCount;
            SkipWhitespaceAndComments<parseFlags>(is);
            CEREAL_RAPIDJSON_PARSE_ERROR_EARLY_RETURN_VOID;

            if (Consume(is, ',')) {
                SkipWhitespaceAndComments<parseFlags>(is);
                CEREAL_RAPIDJSON_PARSE_ERROR_EARLY_RETURN_VOID;
            }
            else if (Consume(is, ']')) {
                if (CEREAL_RAPIDJSON_UNLIKELY(!handler.EndArray(elementCount)))
                    CEREAL_RAPIDJSON_PARSE_ERROR(kParseErrorTermination, is.Tell());
                return;
            }
            else
                CEREAL_RAPIDJSON_PARSE_ERROR(kParseErrorArrayMissCommaOrSquareBracket, is.Tell());

            if (parseFlags & kParseTrailingCommasFlag) {
                if (is.Peek() == ']') {
                    if (CEREAL_RAPIDJSON_UNLIKELY(!handler.EndArray(elementCount)))
                        CEREAL_RAPIDJSON_PARSE_ERROR(kParseErrorTermination, is.Tell());
                    is.Take();
                    return;
                }
            }
        }
    }

    template<unsigned parseFlags, typename InputStream, typename Handler>
    void ParseNull(InputStream& is, Handler& handler) {
        CEREAL_RAPIDJSON_ASSERT(is.Peek() == 'n');
        is.Take();

        if (CEREAL_RAPIDJSON_LIKELY(Consume(is, 'u') && Consume(is, 'l') && Consume(is, 'l'))) {
            if (CEREAL_RAPIDJSON_UNLIKELY(!handler.Null()))
                CEREAL_RAPIDJSON_PARSE_ERROR(kParseErrorTermination, is.Tell());
        }
        else
            CEREAL_RAPIDJSON_PARSE_ERROR(kParseErrorValueInvalid, is.Tell());
    }

    template<unsigned parseFlags, typename InputStream, typename Handler>
    void ParseTrue(InputStream& is, Handler& handler) {
        CEREAL_RAPIDJSON_ASSERT(is.Peek() == 't');
        is.Take();

        if (CEREAL_RAPIDJSON_LIKELY(Consume(is, 'r') && Consume(is, 'u') && Consume(is, 'e'))) {
            if (CEREAL_RAPIDJSON_UNLIKELY(!handler.Bool(true)))
                CEREAL_RAPIDJSON_PARSE_ERROR(kParseErrorTermination, is.Tell());
        }
        else
            CEREAL_RAPIDJSON_PARSE_ERROR(kParseErrorValueInvalid, is.Tell());
    }

    template<unsigned parseFlags, typename InputStream, typename Handler>
    void ParseFalse(InputStream& is, Handler& handler) {
        CEREAL_RAPIDJSON_ASSERT(is.Peek() == 'f');
        is.Take();

        if (CEREAL_RAPIDJSON_LIKELY(Consume(is, 'a') && Consume(is, 'l') && Consume(is, 's') && Consume(is, 'e'))) {
            if (CEREAL_RAPIDJSON_UNLIKELY(!handler.Bool(false)))
                CEREAL_RAPIDJSON_PARSE_ERROR(kParseErrorTermination, is.Tell());
        }
        else
            CEREAL_RAPIDJSON_PARSE_ERROR(kParseErrorValueInvalid, is.Tell());
    }

    template<typename InputStream>
    CEREAL_RAPIDJSON_FORCEINLINE static bool Consume(InputStream& is, typename InputStream::Ch expect) {
        if (CEREAL_RAPIDJSON_LIKELY(is.Peek() == expect)) {
            is.Take();
            return true;
        }
        else
            return false;
    }

    // Helper function to parse four hexadecimal digits in \uXXXX in ParseString().
    template<typename InputStream>
    unsigned ParseHex4(InputStream& is, size_t escapeOffset) {
        unsigned codepoint = 0;
        for (int i = 0; i < 4; i++) {
            Ch c = is.Peek();
            codepoint <<= 4;
            codepoint += static_cast<unsigned>(c);
            if (c >= '0' && c <= '9')
                codepoint -= '0';
            else if (c >= 'A' && c <= 'F')
                codepoint -= 'A' - 10;
            else if (c >= 'a' && c <= 'f')
                codepoint -= 'a' - 10;
            else {
                CEREAL_RAPIDJSON_PARSE_ERROR_NORETURN(kParseErrorStringUnicodeEscapeInvalidHex, escapeOffset);
                CEREAL_RAPIDJSON_PARSE_ERROR_EARLY_RETURN(0);
            }
            is.Take();
        }
        return codepoint;
    }

    template <typename CharType>
    class StackStream {
    public:
        typedef CharType Ch;

        StackStream(internal::Stack<StackAllocator>& stack) : stack_(stack), length_(0) {}
        CEREAL_RAPIDJSON_FORCEINLINE void Put(Ch c) {
            *stack_.template Push<Ch>() = c;
            ++length_;
        }

        CEREAL_RAPIDJSON_FORCEINLINE void* Push(SizeType count) {
            length_ += count;
            return stack_.template Push<Ch>(count);
        }

        size_t Length() const { return length_; }

        Ch* Pop() {
            return stack_.template Pop<Ch>(length_);
        }

    private:
        StackStream(const StackStream&);
        StackStream& operator=(const StackStream&);

        internal::Stack<StackAllocator>& stack_;
        SizeType length_;
    };

    // Parse string and generate String event. Different code paths for kParseInsituFlag.
    template<unsigned parseFlags, typename InputStream, typename Handler>
    void ParseString(InputStream& is, Handler& handler, bool isKey = false) {
        internal::StreamLocalCopy<InputStream> copy(is);
        InputStream& s(copy.s);

        CEREAL_RAPIDJSON_ASSERT(s.Peek() == '\"');
        s.Take();  // Skip '\"'

        bool success = false;
        if (parseFlags & kParseInsituFlag) {
            typename InputStream::Ch *head = s.PutBegin();
            ParseStringToStream<parseFlags, SourceEncoding, SourceEncoding>(s, s);
            CEREAL_RAPIDJSON_PARSE_ERROR_EARLY_RETURN_VOID;
            size_t length = s.PutEnd(head) - 1;
            CEREAL_RAPIDJSON_ASSERT(length <= 0xFFFFFFFF);
            const typename TargetEncoding::Ch* const str = reinterpret_cast<typename TargetEncoding::Ch*>(head);
            success = (isKey ? handler.Key(str, SizeType(length), false) : handler.String(str, SizeType(length), false));
        }
        else {
            StackStream<typename TargetEncoding::Ch> stackStream(stack_);
            ParseStringToStream<parseFlags, SourceEncoding, TargetEncoding>(s, stackStream);
            CEREAL_RAPIDJSON_PARSE_ERROR_EARLY_RETURN_VOID;
            SizeType length = static_cast<SizeType>(stackStream.Length()) - 1;
            const typename TargetEncoding::Ch* const str = stackStream.Pop();
            success = (isKey ? handler.Key(str, length, true) : handler.String(str, length, true));
        }
        if (CEREAL_RAPIDJSON_UNLIKELY(!success))
            CEREAL_RAPIDJSON_PARSE_ERROR(kParseErrorTermination, s.Tell());
    }

    // Parse string to an output is
    // This function handles the prefix/suffix double quotes, escaping, and optional encoding validation.
    template<unsigned parseFlags, typename SEncoding, typename TEncoding, typename InputStream, typename OutputStream>
    CEREAL_RAPIDJSON_FORCEINLINE void ParseStringToStream(InputStream& is, OutputStream& os) {
//!@cond CEREAL_RAPIDJSON_HIDDEN_FROM_DOXYGEN
#define Z16 0,0,0,0,0,0,0,0,0,0,0,0,0,0,0,0
        static const char escape[256] = {
            Z16, Z16, 0, 0,'\"', 0, 0, 0, 0, 0, 0, 0, 0, 0, 0, 0, 0,'/',
            Z16, Z16, 0, 0, 0, 0, 0, 0, 0, 0, 0, 0, 0, 0,'\\', 0, 0, 0,
            0, 0,'\b', 0, 0, 0,'\f', 0, 0, 0, 0, 0, 0, 0,'\n', 0,
            0, 0,'\r', 0,'\t', 0, 0, 0, 0, 0, 0, 0, 0, 0, 0, 0,
            Z16, Z16, Z16, Z16, Z16, Z16, Z16, Z16
        };
#undef Z16
//!@endcond

        for (;;) {
            // Scan and copy string before "\\\"" or < 0x20. This is an optional optimzation.
            if (!(parseFlags & kParseValidateEncodingFlag))
                ScanCopyUnescapedString(is, os);

            Ch c = is.Peek();
            if (CEREAL_RAPIDJSON_UNLIKELY(c == '\\')) {    // Escape
                size_t escapeOffset = is.Tell();    // For invalid escaping, report the initial '\\' as error offset
                is.Take();
                Ch e = is.Peek();
                if ((sizeof(Ch) == 1 || unsigned(e) < 256) && CEREAL_RAPIDJSON_LIKELY(escape[static_cast<unsigned char>(e)])) {
                    is.Take();
                    os.Put(static_cast<typename TEncoding::Ch>(escape[static_cast<unsigned char>(e)]));
                }
                else if (CEREAL_RAPIDJSON_LIKELY(e == 'u')) {    // Unicode
                    is.Take();
                    unsigned codepoint = ParseHex4(is, escapeOffset);
                    CEREAL_RAPIDJSON_PARSE_ERROR_EARLY_RETURN_VOID;
                    if (CEREAL_RAPIDJSON_UNLIKELY(codepoint >= 0xD800 && codepoint <= 0xDBFF)) {
                        // Handle UTF-16 surrogate pair
                        if (CEREAL_RAPIDJSON_UNLIKELY(!Consume(is, '\\') || !Consume(is, 'u')))
                            CEREAL_RAPIDJSON_PARSE_ERROR(kParseErrorStringUnicodeSurrogateInvalid, escapeOffset);
                        unsigned codepoint2 = ParseHex4(is, escapeOffset);
                        CEREAL_RAPIDJSON_PARSE_ERROR_EARLY_RETURN_VOID;
                        if (CEREAL_RAPIDJSON_UNLIKELY(codepoint2 < 0xDC00 || codepoint2 > 0xDFFF))
                            CEREAL_RAPIDJSON_PARSE_ERROR(kParseErrorStringUnicodeSurrogateInvalid, escapeOffset);
                        codepoint = (((codepoint - 0xD800) << 10) | (codepoint2 - 0xDC00)) + 0x10000;
                    }
                    TEncoding::Encode(os, codepoint);
                }
                else
                    CEREAL_RAPIDJSON_PARSE_ERROR(kParseErrorStringEscapeInvalid, escapeOffset);
            }
            else if (CEREAL_RAPIDJSON_UNLIKELY(c == '"')) {    // Closing double quote
                is.Take();
                os.Put('\0');   // null-terminate the string
                return;
            }
            else if (CEREAL_RAPIDJSON_UNLIKELY(static_cast<unsigned>(c) < 0x20)) { // RFC 4627: unescaped = %x20-21 / %x23-5B / %x5D-10FFFF
                if (c == '\0')
                    CEREAL_RAPIDJSON_PARSE_ERROR(kParseErrorStringMissQuotationMark, is.Tell());
                else
                    CEREAL_RAPIDJSON_PARSE_ERROR(kParseErrorStringInvalidEncoding, is.Tell());
            }
            else {
                size_t offset = is.Tell();
                if (CEREAL_RAPIDJSON_UNLIKELY((parseFlags & kParseValidateEncodingFlag ?
                    !Transcoder<SEncoding, TEncoding>::Validate(is, os) :
                    !Transcoder<SEncoding, TEncoding>::Transcode(is, os))))
                    CEREAL_RAPIDJSON_PARSE_ERROR(kParseErrorStringInvalidEncoding, offset);
            }
        }
    }

    template<typename InputStream, typename OutputStream>
    static CEREAL_RAPIDJSON_FORCEINLINE void ScanCopyUnescapedString(InputStream&, OutputStream&) {
            // Do nothing for generic version
    }

#if defined(CEREAL_RAPIDJSON_SSE2) || defined(CEREAL_RAPIDJSON_SSE42)
    // StringStream -> StackStream<char>
    static CEREAL_RAPIDJSON_FORCEINLINE void ScanCopyUnescapedString(StringStream& is, StackStream<char>& os) {
        const char* p = is.src_;

        // Scan one by one until alignment (unaligned load may cross page boundary and cause crash)
        const char* nextAligned = reinterpret_cast<const char*>((reinterpret_cast<size_t>(p) + 15) & static_cast<size_t>(~15));
        while (p != nextAligned)
            if (CEREAL_RAPIDJSON_UNLIKELY(*p == '\"') || CEREAL_RAPIDJSON_UNLIKELY(*p == '\\') || CEREAL_RAPIDJSON_UNLIKELY(static_cast<unsigned>(*p) < 0x20)) {
                is.src_ = p;
                return;
            }
            else
                os.Put(*p++);

        // The rest of string using SIMD
        static const char dquote[16] = { '\"', '\"', '\"', '\"', '\"', '\"', '\"', '\"', '\"', '\"', '\"', '\"', '\"', '\"', '\"', '\"' };
        static const char bslash[16] = { '\\', '\\', '\\', '\\', '\\', '\\', '\\', '\\', '\\', '\\', '\\', '\\', '\\', '\\', '\\', '\\' };
        static const char space[16]  = { 0x1F, 0x1F, 0x1F, 0x1F, 0x1F, 0x1F, 0x1F, 0x1F, 0x1F, 0x1F, 0x1F, 0x1F, 0x1F, 0x1F, 0x1F, 0x1F };
        const __m128i dq = _mm_loadu_si128(reinterpret_cast<const __m128i *>(&dquote[0]));
        const __m128i bs = _mm_loadu_si128(reinterpret_cast<const __m128i *>(&bslash[0]));
        const __m128i sp = _mm_loadu_si128(reinterpret_cast<const __m128i *>(&space[0]));

        for (;; p += 16) {
            const __m128i s = _mm_load_si128(reinterpret_cast<const __m128i *>(p));
            const __m128i t1 = _mm_cmpeq_epi8(s, dq);
            const __m128i t2 = _mm_cmpeq_epi8(s, bs);
            const __m128i t3 = _mm_cmpeq_epi8(_mm_max_epu8(s, sp), sp); // s < 0x20 <=> max(s, 0x1F) == 0x1F
            const __m128i x = _mm_or_si128(_mm_or_si128(t1, t2), t3);
            unsigned short r = static_cast<unsigned short>(_mm_movemask_epi8(x));
            if (CEREAL_RAPIDJSON_UNLIKELY(r != 0)) {   // some of characters is escaped
                SizeType length;
    #ifdef _MSC_VER         // Find the index of first escaped
                unsigned long offset;
                _BitScanForward(&offset, r);
                length = offset;
    #else
                length = static_cast<SizeType>(__builtin_ffs(r) - 1);
    #endif
                if (length != 0) {
                    char* q = reinterpret_cast<char*>(os.Push(length));
                    for (size_t i = 0; i < length; i++)
                        q[i] = p[i];

                    p += length;
                }
                break;
            }
            _mm_storeu_si128(reinterpret_cast<__m128i *>(os.Push(16)), s);
        }

        is.src_ = p;
    }

    // InsituStringStream -> InsituStringStream
    static CEREAL_RAPIDJSON_FORCEINLINE void ScanCopyUnescapedString(InsituStringStream& is, InsituStringStream& os) {
        CEREAL_RAPIDJSON_ASSERT(&is == &os);
        (void)os;

        if (is.src_ == is.dst_) {
            SkipUnescapedString(is);
            return;
        }

        char* p = is.src_;
        char *q = is.dst_;

        // Scan one by one until alignment (unaligned load may cross page boundary and cause crash)
        const char* nextAligned = reinterpret_cast<const char*>((reinterpret_cast<size_t>(p) + 15) & static_cast<size_t>(~15));
        while (p != nextAligned)
            if (CEREAL_RAPIDJSON_UNLIKELY(*p == '\"') || CEREAL_RAPIDJSON_UNLIKELY(*p == '\\') || CEREAL_RAPIDJSON_UNLIKELY(static_cast<unsigned>(*p) < 0x20)) {
                is.src_ = p;
                is.dst_ = q;
                return;
            }
            else
                *q++ = *p++;

        // The rest of string using SIMD
        static const char dquote[16] = { '\"', '\"', '\"', '\"', '\"', '\"', '\"', '\"', '\"', '\"', '\"', '\"', '\"', '\"', '\"', '\"' };
        static const char bslash[16] = { '\\', '\\', '\\', '\\', '\\', '\\', '\\', '\\', '\\', '\\', '\\', '\\', '\\', '\\', '\\', '\\' };
        static const char space[16] = { 0x1F, 0x1F, 0x1F, 0x1F, 0x1F, 0x1F, 0x1F, 0x1F, 0x1F, 0x1F, 0x1F, 0x1F, 0x1F, 0x1F, 0x1F, 0x1F };
        const __m128i dq = _mm_loadu_si128(reinterpret_cast<const __m128i *>(&dquote[0]));
        const __m128i bs = _mm_loadu_si128(reinterpret_cast<const __m128i *>(&bslash[0]));
        const __m128i sp = _mm_loadu_si128(reinterpret_cast<const __m128i *>(&space[0]));

        for (;; p += 16, q += 16) {
            const __m128i s = _mm_load_si128(reinterpret_cast<const __m128i *>(p));
            const __m128i t1 = _mm_cmpeq_epi8(s, dq);
            const __m128i t2 = _mm_cmpeq_epi8(s, bs);
            const __m128i t3 = _mm_cmpeq_epi8(_mm_max_epu8(s, sp), sp); // s < 0x20 <=> max(s, 0x1F) == 0x1F
            const __m128i x = _mm_or_si128(_mm_or_si128(t1, t2), t3);
            unsigned short r = static_cast<unsigned short>(_mm_movemask_epi8(x));
            if (CEREAL_RAPIDJSON_UNLIKELY(r != 0)) {   // some of characters is escaped
                size_t length;
#ifdef _MSC_VER         // Find the index of first escaped
                unsigned long offset;
                _BitScanForward(&offset, r);
                length = offset;
#else
                length = static_cast<size_t>(__builtin_ffs(r) - 1);
#endif
                for (const char* pend = p + length; p != pend; )
                    *q++ = *p++;
                break;
            }
            _mm_storeu_si128(reinterpret_cast<__m128i *>(q), s);
        }

        is.src_ = p;
        is.dst_ = q;
    }

    // When read/write pointers are the same for insitu stream, just skip unescaped characters
    static CEREAL_RAPIDJSON_FORCEINLINE void SkipUnescapedString(InsituStringStream& is) {
        CEREAL_RAPIDJSON_ASSERT(is.src_ == is.dst_);
        char* p = is.src_;

        // Scan one by one until alignment (unaligned load may cross page boundary and cause crash)
        const char* nextAligned = reinterpret_cast<const char*>((reinterpret_cast<size_t>(p) + 15) & static_cast<size_t>(~15));
        for (; p != nextAligned; p++)
            if (CEREAL_RAPIDJSON_UNLIKELY(*p == '\"') || CEREAL_RAPIDJSON_UNLIKELY(*p == '\\') || CEREAL_RAPIDJSON_UNLIKELY(static_cast<unsigned>(*p) < 0x20)) {
                is.src_ = is.dst_ = p;
                return;
            }

        // The rest of string using SIMD
        static const char dquote[16] = { '\"', '\"', '\"', '\"', '\"', '\"', '\"', '\"', '\"', '\"', '\"', '\"', '\"', '\"', '\"', '\"' };
        static const char bslash[16] = { '\\', '\\', '\\', '\\', '\\', '\\', '\\', '\\', '\\', '\\', '\\', '\\', '\\', '\\', '\\', '\\' };
        static const char space[16] = { 0x1F, 0x1F, 0x1F, 0x1F, 0x1F, 0x1F, 0x1F, 0x1F, 0x1F, 0x1F, 0x1F, 0x1F, 0x1F, 0x1F, 0x1F, 0x1F };
        const __m128i dq = _mm_loadu_si128(reinterpret_cast<const __m128i *>(&dquote[0]));
        const __m128i bs = _mm_loadu_si128(reinterpret_cast<const __m128i *>(&bslash[0]));
        const __m128i sp = _mm_loadu_si128(reinterpret_cast<const __m128i *>(&space[0]));

        for (;; p += 16) {
            const __m128i s = _mm_load_si128(reinterpret_cast<const __m128i *>(p));
            const __m128i t1 = _mm_cmpeq_epi8(s, dq);
            const __m128i t2 = _mm_cmpeq_epi8(s, bs);
            const __m128i t3 = _mm_cmpeq_epi8(_mm_max_epu8(s, sp), sp); // s < 0x20 <=> max(s, 0x1F) == 0x1F
            const __m128i x = _mm_or_si128(_mm_or_si128(t1, t2), t3);
            unsigned short r = static_cast<unsigned short>(_mm_movemask_epi8(x));
            if (CEREAL_RAPIDJSON_UNLIKELY(r != 0)) {   // some of characters is escaped
                size_t length;
#ifdef _MSC_VER         // Find the index of first escaped
                unsigned long offset;
                _BitScanForward(&offset, r);
                length = offset;
#else
                length = static_cast<size_t>(__builtin_ffs(r) - 1);
#endif
                p += length;
                break;
            }
        }

        is.src_ = is.dst_ = p;
    }
#elif defined(CEREAL_RAPIDJSON_NEON)
    // StringStream -> StackStream<char>
    static CEREAL_RAPIDJSON_FORCEINLINE void ScanCopyUnescapedString(StringStream& is, StackStream<char>& os) {
        const char* p = is.src_;

        // Scan one by one until alignment (unaligned load may cross page boundary and cause crash)
        const char* nextAligned = reinterpret_cast<const char*>((reinterpret_cast<size_t>(p) + 15) & static_cast<size_t>(~15));
        while (p != nextAligned)
            if (CEREAL_RAPIDJSON_UNLIKELY(*p == '\"') || CEREAL_RAPIDJSON_UNLIKELY(*p == '\\') || CEREAL_RAPIDJSON_UNLIKELY(static_cast<unsigned>(*p) < 0x20)) {
                is.src_ = p;
                return;
            }
            else
                os.Put(*p++);

        // The rest of string using SIMD
        const uint8x16_t s0 = vmovq_n_u8('"');
        const uint8x16_t s1 = vmovq_n_u8('\\');
        const uint8x16_t s2 = vmovq_n_u8('\b');
        const uint8x16_t s3 = vmovq_n_u8(32);

        for (;; p += 16) {
            const uint8x16_t s = vld1q_u8(reinterpret_cast<const uint8_t *>(p));
            uint8x16_t x = vceqq_u8(s, s0);
            x = vorrq_u8(x, vceqq_u8(s, s1));
            x = vorrq_u8(x, vceqq_u8(s, s2));
            x = vorrq_u8(x, vcltq_u8(s, s3));

            x = vrev64q_u8(x);                     // Rev in 64
            uint64_t low = vgetq_lane_u64(reinterpret_cast<uint64x2_t>(x), 0);   // extract
            uint64_t high = vgetq_lane_u64(reinterpret_cast<uint64x2_t>(x), 1);  // extract

            SizeType length = 0;
            bool escaped = false;
            if (low == 0) {
                if (high != 0) {
                    unsigned lz = (unsigned)__builtin_clzll(high);;
                    length = 8 + (lz >> 3);
                    escaped = true;
                }
            } else {
                unsigned lz = (unsigned)__builtin_clzll(low);;
                length = lz >> 3;
                escaped = true;
            }
            if (CEREAL_RAPIDJSON_UNLIKELY(escaped)) {   // some of characters is escaped
                if (length != 0) {
                    char* q = reinterpret_cast<char*>(os.Push(length));
                    for (size_t i = 0; i < length; i++)
                        q[i] = p[i];

                    p += length;
                }
                break;
            }
            vst1q_u8(reinterpret_cast<uint8_t *>(os.Push(16)), s);
        }

        is.src_ = p;
    }

    // InsituStringStream -> InsituStringStream
    static CEREAL_RAPIDJSON_FORCEINLINE void ScanCopyUnescapedString(InsituStringStream& is, InsituStringStream& os) {
        CEREAL_RAPIDJSON_ASSERT(&is == &os);
        (void)os;

        if (is.src_ == is.dst_) {
            SkipUnescapedString(is);
            return;
        }

        char* p = is.src_;
        char *q = is.dst_;

        // Scan one by one until alignment (unaligned load may cross page boundary and cause crash)
        const char* nextAligned = reinterpret_cast<const char*>((reinterpret_cast<size_t>(p) + 15) & static_cast<size_t>(~15));
        while (p != nextAligned)
            if (CEREAL_RAPIDJSON_UNLIKELY(*p == '\"') || CEREAL_RAPIDJSON_UNLIKELY(*p == '\\') || CEREAL_RAPIDJSON_UNLIKELY(static_cast<unsigned>(*p) < 0x20)) {
                is.src_ = p;
                is.dst_ = q;
                return;
            }
            else
                *q++ = *p++;

        // The rest of string using SIMD
        const uint8x16_t s0 = vmovq_n_u8('"');
        const uint8x16_t s1 = vmovq_n_u8('\\');
        const uint8x16_t s2 = vmovq_n_u8('\b');
        const uint8x16_t s3 = vmovq_n_u8(32);

        for (;; p += 16, q += 16) {
            const uint8x16_t s = vld1q_u8(reinterpret_cast<uint8_t *>(p));
            uint8x16_t x = vceqq_u8(s, s0);
            x = vorrq_u8(x, vceqq_u8(s, s1));
            x = vorrq_u8(x, vceqq_u8(s, s2));
            x = vorrq_u8(x, vcltq_u8(s, s3));

            x = vrev64q_u8(x);                     // Rev in 64
            uint64_t low = vgetq_lane_u64(reinterpret_cast<uint64x2_t>(x), 0);   // extract
            uint64_t high = vgetq_lane_u64(reinterpret_cast<uint64x2_t>(x), 1);  // extract

            SizeType length = 0;
            bool escaped = false;
            if (low == 0) {
                if (high != 0) {
                    unsigned lz = (unsigned)__builtin_clzll(high);
                    length = 8 + (lz >> 3);
                    escaped = true;
                }
            } else {
                unsigned lz = (unsigned)__builtin_clzll(low);
                length = lz >> 3;
                escaped = true;
            }
            if (CEREAL_RAPIDJSON_UNLIKELY(escaped)) {   // some of characters is escaped
                for (const char* pend = p + length; p != pend; ) {
                    *q++ = *p++;
                }
                break;
            }
            vst1q_u8(reinterpret_cast<uint8_t *>(q), s);
        }

        is.src_ = p;
        is.dst_ = q;
    }

    // When read/write pointers are the same for insitu stream, just skip unescaped characters
    static CEREAL_RAPIDJSON_FORCEINLINE void SkipUnescapedString(InsituStringStream& is) {
        CEREAL_RAPIDJSON_ASSERT(is.src_ == is.dst_);
        char* p = is.src_;

        // Scan one by one until alignment (unaligned load may cross page boundary and cause crash)
        const char* nextAligned = reinterpret_cast<const char*>((reinterpret_cast<size_t>(p) + 15) & static_cast<size_t>(~15));
        for (; p != nextAligned; p++)
            if (CEREAL_RAPIDJSON_UNLIKELY(*p == '\"') || CEREAL_RAPIDJSON_UNLIKELY(*p == '\\') || CEREAL_RAPIDJSON_UNLIKELY(static_cast<unsigned>(*p) < 0x20)) {
                is.src_ = is.dst_ = p;
                return;
            }

        // The rest of string using SIMD
        const uint8x16_t s0 = vmovq_n_u8('"');
        const uint8x16_t s1 = vmovq_n_u8('\\');
        const uint8x16_t s2 = vmovq_n_u8('\b');
        const uint8x16_t s3 = vmovq_n_u8(32);

        for (;; p += 16) {
            const uint8x16_t s = vld1q_u8(reinterpret_cast<uint8_t *>(p));
            uint8x16_t x = vceqq_u8(s, s0);
            x = vorrq_u8(x, vceqq_u8(s, s1));
            x = vorrq_u8(x, vceqq_u8(s, s2));
            x = vorrq_u8(x, vcltq_u8(s, s3));

            x = vrev64q_u8(x);                     // Rev in 64
            uint64_t low = vgetq_lane_u64(reinterpret_cast<uint64x2_t>(x), 0);   // extract
            uint64_t high = vgetq_lane_u64(reinterpret_cast<uint64x2_t>(x), 1);  // extract

            if (low == 0) {
                if (high != 0) {
                    int lz = __builtin_clzll(high);
                    p += 8 + (lz >> 3);
                    break;
                }
            } else {
                int lz = __builtin_clzll(low);
                p += lz >> 3;
                break;
            }
        }

        is.src_ = is.dst_ = p;
    }
#endif // CEREAL_RAPIDJSON_NEON

    template<typename InputStream, bool backup, bool pushOnTake>
    class NumberStream;

    template<typename InputStream>
    class NumberStream<InputStream, false, false> {
    public:
        typedef typename InputStream::Ch Ch;

        NumberStream(GenericReader& reader, InputStream& s) : is(s) { (void)reader;  }

        CEREAL_RAPIDJSON_FORCEINLINE Ch Peek() const { return is.Peek(); }
        CEREAL_RAPIDJSON_FORCEINLINE Ch TakePush() { return is.Take(); }
        CEREAL_RAPIDJSON_FORCEINLINE Ch Take() { return is.Take(); }
		  CEREAL_RAPIDJSON_FORCEINLINE void Push(char) {}

        size_t Tell() { return is.Tell(); }
        size_t Length() { return 0; }
        const char* Pop() { return 0; }

    protected:
        NumberStream& operator=(const NumberStream&);

        InputStream& is;
    };

    template<typename InputStream>
    class NumberStream<InputStream, true, false> : public NumberStream<InputStream, false, false> {
        typedef NumberStream<InputStream, false, false> Base;
    public:
        NumberStream(GenericReader& reader, InputStream& s) : Base(reader, s), stackStream(reader.stack_) {}

        CEREAL_RAPIDJSON_FORCEINLINE Ch TakePush() {
            stackStream.Put(static_cast<char>(Base::is.Peek()));
            return Base::is.Take();
        }

        CEREAL_RAPIDJSON_FORCEINLINE void Push(char c) {
            stackStream.Put(c);
        }

        size_t Length() { return stackStream.Length(); }

        const char* Pop() {
            stackStream.Put('\0');
            return stackStream.Pop();
        }

    private:
        StackStream<char> stackStream;
    };

    template<typename InputStream>
    class NumberStream<InputStream, true, true> : public NumberStream<InputStream, true, false> {
        typedef NumberStream<InputStream, true, false> Base;
    public:
        NumberStream(GenericReader& reader, InputStream& is) : Base(reader, is) {}

        CEREAL_RAPIDJSON_FORCEINLINE Ch Take() { return Base::TakePush(); }
    };

    template<unsigned parseFlags, typename InputStream, typename Handler>
    void ParseNumber(InputStream& is, Handler& handler) {
        internal::StreamLocalCopy<InputStream> copy(is);
        NumberStream<InputStream,
            ((parseFlags & kParseNumbersAsStringsFlag) != 0) ?
                ((parseFlags & kParseInsituFlag) == 0) :
                ((parseFlags & kParseFullPrecisionFlag) != 0),
            (parseFlags & kParseNumbersAsStringsFlag) != 0 &&
                (parseFlags & kParseInsituFlag) == 0> s(*this, copy.s);

        size_t startOffset = s.Tell();
        double d = 0.0;
        bool useNanOrInf = false;

        // Parse minus
        bool minus = Consume(s, '-');

        // Parse int: zero / ( digit1-9 *DIGIT )
        unsigned i = 0;
        uint64_t i64 = 0;
        bool use64bit = false;
        int significandDigit = 0;
        if (CEREAL_RAPIDJSON_UNLIKELY(s.Peek() == '0')) {
            i = 0;
            s.TakePush();
        }
        else if (CEREAL_RAPIDJSON_LIKELY(s.Peek() >= '1' && s.Peek() <= '9')) {
            i = static_cast<unsigned>(s.TakePush() - '0');

            if (minus)
                while (CEREAL_RAPIDJSON_LIKELY(s.Peek() >= '0' && s.Peek() <= '9')) {
                    if (CEREAL_RAPIDJSON_UNLIKELY(i >= 214748364)) { // 2^31 = 2147483648
                        if (CEREAL_RAPIDJSON_LIKELY(i != 214748364 || s.Peek() > '8')) {
                            i64 = i;
                            use64bit = true;
                            break;
                        }
                    }
                    i = i * 10 + static_cast<unsigned>(s.TakePush() - '0');
                    significandDigit++;
                }
            else
                while (CEREAL_RAPIDJSON_LIKELY(s.Peek() >= '0' && s.Peek() <= '9')) {
                    if (CEREAL_RAPIDJSON_UNLIKELY(i >= 429496729)) { // 2^32 - 1 = 4294967295
                        if (CEREAL_RAPIDJSON_LIKELY(i != 429496729 || s.Peek() > '5')) {
                            i64 = i;
                            use64bit = true;
                            break;
                        }
                    }
                    i = i * 10 + static_cast<unsigned>(s.TakePush() - '0');
                    significandDigit++;
                }
        }
        // Parse NaN or Infinity here
        else if ((parseFlags & kParseNanAndInfFlag) && CEREAL_RAPIDJSON_LIKELY((s.Peek() == 'I' || s.Peek() == 'N'))) {
            if (Consume(s, 'N')) {
                if (Consume(s, 'a') && Consume(s, 'N')) {
                    d = std::numeric_limits<double>::quiet_NaN();
                    useNanOrInf = true;
                }
            }
            else if (CEREAL_RAPIDJSON_LIKELY(Consume(s, 'I'))) {
                if (Consume(s, 'n') && Consume(s, 'f')) {
                    d = (minus ? -std::numeric_limits<double>::infinity() : std::numeric_limits<double>::infinity());
                    useNanOrInf = true;

                    if (CEREAL_RAPIDJSON_UNLIKELY(s.Peek() == 'i' && !(Consume(s, 'i') && Consume(s, 'n')
                                                                && Consume(s, 'i') && Consume(s, 't') && Consume(s, 'y')))) {
                        CEREAL_RAPIDJSON_PARSE_ERROR(kParseErrorValueInvalid, s.Tell());
                    }
                }
            }

            if (CEREAL_RAPIDJSON_UNLIKELY(!useNanOrInf)) {
                CEREAL_RAPIDJSON_PARSE_ERROR(kParseErrorValueInvalid, s.Tell());
            }
        }
        else
            CEREAL_RAPIDJSON_PARSE_ERROR(kParseErrorValueInvalid, s.Tell());

        // Parse 64bit int
        bool useDouble = false;
        if (use64bit) {
            if (minus)
                while (CEREAL_RAPIDJSON_LIKELY(s.Peek() >= '0' && s.Peek() <= '9')) {
                     if (CEREAL_RAPIDJSON_UNLIKELY(i64 >= CEREAL_RAPIDJSON_UINT64_C2(0x0CCCCCCC, 0xCCCCCCCC))) // 2^63 = 9223372036854775808
                        if (CEREAL_RAPIDJSON_LIKELY(i64 != CEREAL_RAPIDJSON_UINT64_C2(0x0CCCCCCC, 0xCCCCCCCC) || s.Peek() > '8')) {
                            d = static_cast<double>(i64);
                            useDouble = true;
                            break;
                        }
                    i64 = i64 * 10 + static_cast<unsigned>(s.TakePush() - '0');
                    significandDigit++;
                }
            else
                while (CEREAL_RAPIDJSON_LIKELY(s.Peek() >= '0' && s.Peek() <= '9')) {
                    if (CEREAL_RAPIDJSON_UNLIKELY(i64 >= CEREAL_RAPIDJSON_UINT64_C2(0x19999999, 0x99999999))) // 2^64 - 1 = 18446744073709551615
                        if (CEREAL_RAPIDJSON_LIKELY(i64 != CEREAL_RAPIDJSON_UINT64_C2(0x19999999, 0x99999999) || s.Peek() > '5')) {
                            d = static_cast<double>(i64);
                            useDouble = true;
                            break;
                        }
                    i64 = i64 * 10 + static_cast<unsigned>(s.TakePush() - '0');
                    significandDigit++;
                }
        }

        // Force double for big integer
        if (useDouble) {
            while (CEREAL_RAPIDJSON_LIKELY(s.Peek() >= '0' && s.Peek() <= '9')) {
                d = d * 10 + (s.TakePush() - '0');
            }
        }

        // Parse frac = decimal-point 1*DIGIT
        int expFrac = 0;
        size_t decimalPosition;
        if (Consume(s, '.')) {
            decimalPosition = s.Length();

            if (CEREAL_RAPIDJSON_UNLIKELY(!(s.Peek() >= '0' && s.Peek() <= '9')))
                CEREAL_RAPIDJSON_PARSE_ERROR(kParseErrorNumberMissFraction, s.Tell());

            if (!useDouble) {
#if CEREAL_RAPIDJSON_64BIT
                // Use i64 to store significand in 64-bit architecture
                if (!use64bit)
                    i64 = i;

                while (CEREAL_RAPIDJSON_LIKELY(s.Peek() >= '0' && s.Peek() <= '9')) {
                    if (i64 > CEREAL_RAPIDJSON_UINT64_C2(0x1FFFFF, 0xFFFFFFFF)) // 2^53 - 1 for fast path
                        break;
                    else {
                        i64 = i64 * 10 + static_cast<unsigned>(s.TakePush() - '0');
                        --expFrac;
                        if (i64 != 0)
                            significandDigit++;
                    }
                }

                d = static_cast<double>(i64);
#else
                // Use double to store significand in 32-bit architecture
                d = static_cast<double>(use64bit ? i64 : i);
#endif
                useDouble = true;
            }

            while (CEREAL_RAPIDJSON_LIKELY(s.Peek() >= '0' && s.Peek() <= '9')) {
                if (significandDigit < 17) {
                    d = d * 10.0 + (s.TakePush() - '0');
                    --expFrac;
                    if (CEREAL_RAPIDJSON_LIKELY(d > 0.0))
                        significandDigit++;
                }
                else
                    s.TakePush();
            }
        }
        else
            decimalPosition = s.Length(); // decimal position at the end of integer.

        // Parse exp = e [ minus / plus ] 1*DIGIT
        int exp = 0;
        if (Consume(s, 'e') || Consume(s, 'E')) {
            if (!useDouble) {
                d = static_cast<double>(use64bit ? i64 : i);
                useDouble = true;
            }

            bool expMinus = false;
            if (Consume(s, '+'))
                ;
            else if (Consume(s, '-'))
                expMinus = true;

            if (CEREAL_RAPIDJSON_LIKELY(s.Peek() >= '0' && s.Peek() <= '9')) {
                exp = static_cast<int>(s.Take() - '0');
                if (expMinus) {
                    // (exp + expFrac) must not underflow int => we're detecting when -exp gets
                    // dangerously close to INT_MIN (a pessimistic next digit 9 would push it into
                    // underflow territory):
                    //
                    //        -(exp * 10 + 9) + expFrac >= INT_MIN
                    //   <=>  exp <= (expFrac - INT_MIN - 9) / 10
                    CEREAL_RAPIDJSON_ASSERT(expFrac <= 0);
                    int maxExp = (expFrac + 2147483639) / 10;

                    while (CEREAL_RAPIDJSON_LIKELY(s.Peek() >= '0' && s.Peek() <= '9')) {
                        exp = exp * 10 + static_cast<int>(s.Take() - '0');
                        if (CEREAL_RAPIDJSON_UNLIKELY(exp > maxExp)) {
                            while (CEREAL_RAPIDJSON_UNLIKELY(s.Peek() >= '0' && s.Peek() <= '9'))  // Consume the rest of exponent
                                s.Take();
                        }
                    }
                }
                else {  // positive exp
                    int maxExp = 308 - expFrac;
                    while (CEREAL_RAPIDJSON_LIKELY(s.Peek() >= '0' && s.Peek() <= '9')) {
                        exp = exp * 10 + static_cast<int>(s.Take() - '0');
                        if (CEREAL_RAPIDJSON_UNLIKELY(exp > maxExp))
                            CEREAL_RAPIDJSON_PARSE_ERROR(kParseErrorNumberTooBig, startOffset);
                    }
                }
            }
            else
                CEREAL_RAPIDJSON_PARSE_ERROR(kParseErrorNumberMissExponent, s.Tell());

            if (expMinus)
                exp = -exp;
        }

        // Finish parsing, call event according to the type of number.
        bool cont = true;

        if (parseFlags & kParseNumbersAsStringsFlag) {
            if (parseFlags & kParseInsituFlag) {
                s.Pop();  // Pop stack no matter if it will be used or not.
                typename InputStream::Ch* head = is.PutBegin();
                const size_t length = s.Tell() - startOffset;
                CEREAL_RAPIDJSON_ASSERT(length <= 0xFFFFFFFF);
                // unable to insert the \0 character here, it will erase the comma after this number
                const typename TargetEncoding::Ch* const str = reinterpret_cast<typename TargetEncoding::Ch*>(head);
                cont = handler.RawNumber(str, SizeType(length), false);
            }
            else {
                SizeType numCharsToCopy = static_cast<SizeType>(s.Length());
                StringStream srcStream(s.Pop());
                StackStream<typename TargetEncoding::Ch> dstStream(stack_);
                while (numCharsToCopy--) {
                    Transcoder<UTF8<>, TargetEncoding>::Transcode(srcStream, dstStream);
                }
                dstStream.Put('\0');
                const typename TargetEncoding::Ch* str = dstStream.Pop();
                const SizeType length = static_cast<SizeType>(dstStream.Length()) - 1;
                cont = handler.RawNumber(str, SizeType(length), true);
            }
        }
        else {
           size_t length = s.Length();
           const char* decimal = s.Pop();  // Pop stack no matter if it will be used or not.

           if (useDouble) {
               int p = exp + expFrac;
               if (parseFlags & kParseFullPrecisionFlag)
                   d = internal::StrtodFullPrecision(d, p, decimal, length, decimalPosition, exp);
               else
                   d = internal::StrtodNormalPrecision(d, p);

               // Use > max, instead of == inf, to fix bogus warning -Wfloat-equal
               if (d > (std::numeric_limits<double>::max)()) {
                   // Overflow
                   // TODO: internal::StrtodX should report overflow (or underflow)
                   CEREAL_RAPIDJSON_PARSE_ERROR(kParseErrorNumberTooBig, startOffset);
               }

               cont = handler.Double(minus ? -d : d);
           }
           else if (useNanOrInf) {
               cont = handler.Double(d);
           }
           else {
               if (use64bit) {
                   if (minus)
                       cont = handler.Int64(static_cast<int64_t>(~i64 + 1));
                   else
                       cont = handler.Uint64(i64);
               }
               else {
                   if (minus)
                       cont = handler.Int(static_cast<int32_t>(~i + 1));
                   else
                       cont = handler.Uint(i);
               }
           }
        }
        if (CEREAL_RAPIDJSON_UNLIKELY(!cont))
            CEREAL_RAPIDJSON_PARSE_ERROR(kParseErrorTermination, startOffset);
    }

    // Parse any JSON value
    template<unsigned parseFlags, typename InputStream, typename Handler>
    void ParseValue(InputStream& is, Handler& handler) {
        switch (is.Peek()) {
            case 'n': ParseNull  <parseFlags>(is, handler); break;
            case 't': ParseTrue  <parseFlags>(is, handler); break;
            case 'f': ParseFalse <parseFlags>(is, handler); break;
            case '"': ParseString<parseFlags>(is, handler); break;
            case '{': ParseObject<parseFlags>(is, handler); break;
            case '[': ParseArray <parseFlags>(is, handler); break;
            default :
                      ParseNumber<parseFlags>(is, handler);
                      break;

        }
    }

    // Iterative Parsing

    // States
    enum IterativeParsingState {
        IterativeParsingFinishState = 0, // sink states at top
        IterativeParsingErrorState,      // sink states at top
        IterativeParsingStartState,

        // Object states
        IterativeParsingObjectInitialState,
        IterativeParsingMemberKeyState,
        IterativeParsingMemberValueState,
        IterativeParsingObjectFinishState,

        // Array states
        IterativeParsingArrayInitialState,
        IterativeParsingElementState,
        IterativeParsingArrayFinishState,

        // Single value state
        IterativeParsingValueState,

        // Delimiter states (at bottom)
        IterativeParsingElementDelimiterState,
        IterativeParsingMemberDelimiterState,
        IterativeParsingKeyValueDelimiterState,

        cIterativeParsingStateCount
    };

    // Tokens
    enum Token {
        LeftBracketToken = 0,
        RightBracketToken,

        LeftCurlyBracketToken,
        RightCurlyBracketToken,

        CommaToken,
        ColonToken,

        StringToken,
        FalseToken,
        TrueToken,
        NullToken,
        NumberToken,

        kTokenCount
    };

    CEREAL_RAPIDJSON_FORCEINLINE Token Tokenize(Ch c) const {

//!@cond CEREAL_RAPIDJSON_HIDDEN_FROM_DOXYGEN
#define N NumberToken
#define N16 N,N,N,N,N,N,N,N,N,N,N,N,N,N,N,N
        // Maps from ASCII to Token
        static const unsigned char tokenMap[256] = {
            N16, // 00~0F
            N16, // 10~1F
            N, N, StringToken, N, N, N, N, N, N, N, N, N, CommaToken, N, N, N, // 20~2F
            N, N, N, N, N, N, N, N, N, N, ColonToken, N, N, N, N, N, // 30~3F
            N16, // 40~4F
            N, N, N, N, N, N, N, N, N, N, N, LeftBracketToken, N, RightBracketToken, N, N, // 50~5F
            N, N, N, N, N, N, FalseToken, N, N, N, N, N, N, N, NullToken, N, // 60~6F
            N, N, N, N, TrueToken, N, N, N, N, N, N, LeftCurlyBracketToken, N, RightCurlyBracketToken, N, N, // 70~7F
            N16, N16, N16, N16, N16, N16, N16, N16 // 80~FF
        };
#undef N
#undef N16
//!@endcond

        if (sizeof(Ch) == 1 || static_cast<unsigned>(c) < 256)
            return static_cast<Token>(tokenMap[static_cast<unsigned char>(c)]);
        else
            return NumberToken;
    }

    CEREAL_RAPIDJSON_FORCEINLINE IterativeParsingState Predict(IterativeParsingState state, Token token) const {
        // current state x one lookahead token -> new state
        static const char G[cIterativeParsingStateCount][kTokenCount] = {
            // Finish(sink state)
            {
                IterativeParsingErrorState, IterativeParsingErrorState, IterativeParsingErrorState, IterativeParsingErrorState, IterativeParsingErrorState,
                IterativeParsingErrorState, IterativeParsingErrorState, IterativeParsingErrorState, IterativeParsingErrorState, IterativeParsingErrorState,
                IterativeParsingErrorState
            },
            // Error(sink state)
            {
                IterativeParsingErrorState, IterativeParsingErrorState, IterativeParsingErrorState, IterativeParsingErrorState, IterativeParsingErrorState,
                IterativeParsingErrorState, IterativeParsingErrorState, IterativeParsingErrorState, IterativeParsingErrorState, IterativeParsingErrorState,
                IterativeParsingErrorState
            },
            // Start
            {
                IterativeParsingArrayInitialState,  // Left bracket
                IterativeParsingErrorState,         // Right bracket
                IterativeParsingObjectInitialState, // Left curly bracket
                IterativeParsingErrorState,         // Right curly bracket
                IterativeParsingErrorState,         // Comma
                IterativeParsingErrorState,         // Colon
                IterativeParsingValueState,         // String
                IterativeParsingValueState,         // False
                IterativeParsingValueState,         // True
                IterativeParsingValueState,         // Null
                IterativeParsingValueState          // Number
            },
            // ObjectInitial
            {
                IterativeParsingErrorState,         // Left bracket
                IterativeParsingErrorState,         // Right bracket
                IterativeParsingErrorState,         // Left curly bracket
                IterativeParsingObjectFinishState,  // Right curly bracket
                IterativeParsingErrorState,         // Comma
                IterativeParsingErrorState,         // Colon
                IterativeParsingMemberKeyState,     // String
                IterativeParsingErrorState,         // False
                IterativeParsingErrorState,         // True
                IterativeParsingErrorState,         // Null
                IterativeParsingErrorState          // Number
            },
            // MemberKey
            {
                IterativeParsingErrorState,             // Left bracket
                IterativeParsingErrorState,             // Right bracket
                IterativeParsingErrorState,             // Left curly bracket
                IterativeParsingErrorState,             // Right curly bracket
                IterativeParsingErrorState,             // Comma
                IterativeParsingKeyValueDelimiterState, // Colon
                IterativeParsingErrorState,             // String
                IterativeParsingErrorState,             // False
                IterativeParsingErrorState,             // True
                IterativeParsingErrorState,             // Null
                IterativeParsingErrorState              // Number
            },
            // MemberValue
            {
                IterativeParsingErrorState,             // Left bracket
                IterativeParsingErrorState,             // Right bracket
                IterativeParsingErrorState,             // Left curly bracket
                IterativeParsingObjectFinishState,      // Right curly bracket
                IterativeParsingMemberDelimiterState,   // Comma
                IterativeParsingErrorState,             // Colon
                IterativeParsingErrorState,             // String
                IterativeParsingErrorState,             // False
                IterativeParsingErrorState,             // True
                IterativeParsingErrorState,             // Null
                IterativeParsingErrorState              // Number
            },
            // ObjectFinish(sink state)
            {
                IterativeParsingErrorState, IterativeParsingErrorState, IterativeParsingErrorState, IterativeParsingErrorState, IterativeParsingErrorState,
                IterativeParsingErrorState, IterativeParsingErrorState, IterativeParsingErrorState, IterativeParsingErrorState, IterativeParsingErrorState,
                IterativeParsingErrorState
            },
            // ArrayInitial
            {
                IterativeParsingArrayInitialState,      // Left bracket(push Element state)
                IterativeParsingArrayFinishState,       // Right bracket
                IterativeParsingObjectInitialState,     // Left curly bracket(push Element state)
                IterativeParsingErrorState,             // Right curly bracket
                IterativeParsingErrorState,             // Comma
                IterativeParsingErrorState,             // Colon
                IterativeParsingElementState,           // String
                IterativeParsingElementState,           // False
                IterativeParsingElementState,           // True
                IterativeParsingElementState,           // Null
                IterativeParsingElementState            // Number
            },
            // Element
            {
                IterativeParsingErrorState,             // Left bracket
                IterativeParsingArrayFinishState,       // Right bracket
                IterativeParsingErrorState,             // Left curly bracket
                IterativeParsingErrorState,             // Right curly bracket
                IterativeParsingElementDelimiterState,  // Comma
                IterativeParsingErrorState,             // Colon
                IterativeParsingErrorState,             // String
                IterativeParsingErrorState,             // False
                IterativeParsingErrorState,             // True
                IterativeParsingErrorState,             // Null
                IterativeParsingErrorState              // Number
            },
            // ArrayFinish(sink state)
            {
                IterativeParsingErrorState, IterativeParsingErrorState, IterativeParsingErrorState, IterativeParsingErrorState, IterativeParsingErrorState,
                IterativeParsingErrorState, IterativeParsingErrorState, IterativeParsingErrorState, IterativeParsingErrorState, IterativeParsingErrorState,
                IterativeParsingErrorState
            },
            // Single Value (sink state)
            {
                IterativeParsingErrorState, IterativeParsingErrorState, IterativeParsingErrorState, IterativeParsingErrorState, IterativeParsingErrorState,
                IterativeParsingErrorState, IterativeParsingErrorState, IterativeParsingErrorState, IterativeParsingErrorState, IterativeParsingErrorState,
                IterativeParsingErrorState
            },
            // ElementDelimiter
            {
                IterativeParsingArrayInitialState,      // Left bracket(push Element state)
                IterativeParsingArrayFinishState,       // Right bracket
                IterativeParsingObjectInitialState,     // Left curly bracket(push Element state)
                IterativeParsingErrorState,             // Right curly bracket
                IterativeParsingErrorState,             // Comma
                IterativeParsingErrorState,             // Colon
                IterativeParsingElementState,           // String
                IterativeParsingElementState,           // False
                IterativeParsingElementState,           // True
                IterativeParsingElementState,           // Null
                IterativeParsingElementState            // Number
            },
            // MemberDelimiter
            {
                IterativeParsingErrorState,         // Left bracket
                IterativeParsingErrorState,         // Right bracket
                IterativeParsingErrorState,         // Left curly bracket
                IterativeParsingObjectFinishState,  // Right curly bracket
                IterativeParsingErrorState,         // Comma
                IterativeParsingErrorState,         // Colon
                IterativeParsingMemberKeyState,     // String
                IterativeParsingErrorState,         // False
                IterativeParsingErrorState,         // True
                IterativeParsingErrorState,         // Null
                IterativeParsingErrorState          // Number
            },
            // KeyValueDelimiter
            {
                IterativeParsingArrayInitialState,      // Left bracket(push MemberValue state)
                IterativeParsingErrorState,             // Right bracket
                IterativeParsingObjectInitialState,     // Left curly bracket(push MemberValue state)
                IterativeParsingErrorState,             // Right curly bracket
                IterativeParsingErrorState,             // Comma
                IterativeParsingErrorState,             // Colon
                IterativeParsingMemberValueState,       // String
                IterativeParsingMemberValueState,       // False
                IterativeParsingMemberValueState,       // True
                IterativeParsingMemberValueState,       // Null
                IterativeParsingMemberValueState        // Number
            },
        }; // End of G

        return static_cast<IterativeParsingState>(G[state][token]);
    }

    // Make an advance in the token stream and state based on the candidate destination state which was returned by Transit().
    // May return a new state on state pop.
    template <unsigned parseFlags, typename InputStream, typename Handler>
    CEREAL_RAPIDJSON_FORCEINLINE IterativeParsingState Transit(IterativeParsingState src, Token token, IterativeParsingState dst, InputStream& is, Handler& handler) {
        (void)token;

        switch (dst) {
        case IterativeParsingErrorState:
            return dst;

        case IterativeParsingObjectInitialState:
        case IterativeParsingArrayInitialState:
        {
            // Push the state(Element or MemeberValue) if we are nested in another array or value of member.
            // In this way we can get the correct state on ObjectFinish or ArrayFinish by frame pop.
            IterativeParsingState n = src;
            if (src == IterativeParsingArrayInitialState || src == IterativeParsingElementDelimiterState)
                n = IterativeParsingElementState;
            else if (src == IterativeParsingKeyValueDelimiterState)
                n = IterativeParsingMemberValueState;
            // Push current state.
            *stack_.template Push<SizeType>(1) = n;
            // Initialize and push the member/element count.
            *stack_.template Push<SizeType>(1) = 0;
            // Call handler
            bool hr = (dst == IterativeParsingObjectInitialState) ? handler.StartObject() : handler.StartArray();
            // On handler short circuits the parsing.
            if (!hr) {
                CEREAL_RAPIDJSON_PARSE_ERROR_NORETURN(kParseErrorTermination, is.Tell());
                return IterativeParsingErrorState;
            }
            else {
                is.Take();
                return dst;
            }
        }

        case IterativeParsingMemberKeyState:
            ParseString<parseFlags>(is, handler, true);
            if (HasParseError())
                return IterativeParsingErrorState;
            else
                return dst;

        case IterativeParsingKeyValueDelimiterState:
            CEREAL_RAPIDJSON_ASSERT(token == ColonToken);
            is.Take();
            return dst;

        case IterativeParsingMemberValueState:
            // Must be non-compound value. Or it would be ObjectInitial or ArrayInitial state.
            ParseValue<parseFlags>(is, handler);
            if (HasParseError()) {
                return IterativeParsingErrorState;
            }
            return dst;

        case IterativeParsingElementState:
            // Must be non-compound value. Or it would be ObjectInitial or ArrayInitial state.
            ParseValue<parseFlags>(is, handler);
            if (HasParseError()) {
                return IterativeParsingErrorState;
            }
            return dst;

        case IterativeParsingMemberDelimiterState:
        case IterativeParsingElementDelimiterState:
            is.Take();
            // Update member/element count.
            *stack_.template Top<SizeType>() = *stack_.template Top<SizeType>() + 1;
            return dst;

        case IterativeParsingObjectFinishState:
        {
            // Transit from delimiter is only allowed when trailing commas are enabled
            if (!(parseFlags & kParseTrailingCommasFlag) && src == IterativeParsingMemberDelimiterState) {
                CEREAL_RAPIDJSON_PARSE_ERROR_NORETURN(kParseErrorObjectMissName, is.Tell());
                return IterativeParsingErrorState;
            }
            // Get member count.
            SizeType c = *stack_.template Pop<SizeType>(1);
            // If the object is not empty, count the last member.
            if (src == IterativeParsingMemberValueState)
                ++c;
            // Restore the state.
            IterativeParsingState n = static_cast<IterativeParsingState>(*stack_.template Pop<SizeType>(1));
            // Transit to Finish state if this is the topmost scope.
            if (n == IterativeParsingStartState)
                n = IterativeParsingFinishState;
            // Call handler
            bool hr = handler.EndObject(c);
            // On handler short circuits the parsing.
            if (!hr) {
                CEREAL_RAPIDJSON_PARSE_ERROR_NORETURN(kParseErrorTermination, is.Tell());
                return IterativeParsingErrorState;
            }
            else {
                is.Take();
                return n;
            }
        }

        case IterativeParsingArrayFinishState:
        {
            // Transit from delimiter is only allowed when trailing commas are enabled
            if (!(parseFlags & kParseTrailingCommasFlag) && src == IterativeParsingElementDelimiterState) {
                CEREAL_RAPIDJSON_PARSE_ERROR_NORETURN(kParseErrorValueInvalid, is.Tell());
                return IterativeParsingErrorState;
            }
            // Get element count.
            SizeType c = *stack_.template Pop<SizeType>(1);
            // If the array is not empty, count the last element.
            if (src == IterativeParsingElementState)
                ++c;
            // Restore the state.
            IterativeParsingState n = static_cast<IterativeParsingState>(*stack_.template Pop<SizeType>(1));
            // Transit to Finish state if this is the topmost scope.
            if (n == IterativeParsingStartState)
                n = IterativeParsingFinishState;
            // Call handler
            bool hr = handler.EndArray(c);
            // On handler short circuits the parsing.
            if (!hr) {
                CEREAL_RAPIDJSON_PARSE_ERROR_NORETURN(kParseErrorTermination, is.Tell());
                return IterativeParsingErrorState;
            }
            else {
                is.Take();
                return n;
            }
        }

        default:
            // This branch is for IterativeParsingValueState actually.
            // Use `default:` rather than
            // `case IterativeParsingValueState:` is for code coverage.

            // The IterativeParsingStartState is not enumerated in this switch-case.
            // It is impossible for that case. And it can be caught by following assertion.

            // The IterativeParsingFinishState is not enumerated in this switch-case either.
            // It is a "derivative" state which cannot triggered from Predict() directly.
            // Therefore it cannot happen here. And it can be caught by following assertion.
            CEREAL_RAPIDJSON_ASSERT(dst == IterativeParsingValueState);

            // Must be non-compound value. Or it would be ObjectInitial or ArrayInitial state.
            ParseValue<parseFlags>(is, handler);
            if (HasParseError()) {
                return IterativeParsingErrorState;
            }
            return IterativeParsingFinishState;
        }
    }

    template <typename InputStream>
    void HandleError(IterativeParsingState src, InputStream& is) {
        if (HasParseError()) {
            // Error flag has been set.
            return;
        }

        switch (src) {
        case IterativeParsingStartState:            CEREAL_RAPIDJSON_PARSE_ERROR(kParseErrorDocumentEmpty, is.Tell()); return;
        case IterativeParsingFinishState:           CEREAL_RAPIDJSON_PARSE_ERROR(kParseErrorDocumentRootNotSingular, is.Tell()); return;
        case IterativeParsingObjectInitialState:
        case IterativeParsingMemberDelimiterState:  CEREAL_RAPIDJSON_PARSE_ERROR(kParseErrorObjectMissName, is.Tell()); return;
        case IterativeParsingMemberKeyState:        CEREAL_RAPIDJSON_PARSE_ERROR(kParseErrorObjectMissColon, is.Tell()); return;
        case IterativeParsingMemberValueState:      CEREAL_RAPIDJSON_PARSE_ERROR(kParseErrorObjectMissCommaOrCurlyBracket, is.Tell()); return;
        case IterativeParsingKeyValueDelimiterState:
        case IterativeParsingArrayInitialState:
        case IterativeParsingElementDelimiterState: CEREAL_RAPIDJSON_PARSE_ERROR(kParseErrorValueInvalid, is.Tell()); return;
        default: CEREAL_RAPIDJSON_ASSERT(src == IterativeParsingElementState); CEREAL_RAPIDJSON_PARSE_ERROR(kParseErrorArrayMissCommaOrSquareBracket, is.Tell()); return;
        }
    }

    CEREAL_RAPIDJSON_FORCEINLINE bool IsIterativeParsingDelimiterState(IterativeParsingState s) const {
        return s >= IterativeParsingElementDelimiterState;
    }

    CEREAL_RAPIDJSON_FORCEINLINE bool IsIterativeParsingCompleteState(IterativeParsingState s) const {
        return s <= IterativeParsingErrorState;
    }

    template <unsigned parseFlags, typename InputStream, typename Handler>
    ParseResult IterativeParse(InputStream& is, Handler& handler) {
        parseResult_.Clear();
        ClearStackOnExit scope(*this);
        IterativeParsingState state = IterativeParsingStartState;

        SkipWhitespaceAndComments<parseFlags>(is);
        CEREAL_RAPIDJSON_PARSE_ERROR_EARLY_RETURN(parseResult_);
        while (is.Peek() != '\0') {
            Token t = Tokenize(is.Peek());
            IterativeParsingState n = Predict(state, t);
            IterativeParsingState d = Transit<parseFlags>(state, t, n, is, handler);

            if (d == IterativeParsingErrorState) {
                HandleError(state, is);
                break;
            }

            state = d;

            // Do not further consume streams if a root JSON has been parsed.
            if ((parseFlags & kParseStopWhenDoneFlag) && state == IterativeParsingFinishState)
                break;

            SkipWhitespaceAndComments<parseFlags>(is);
            CEREAL_RAPIDJSON_PARSE_ERROR_EARLY_RETURN(parseResult_);
        }

        // Handle the end of file.
        if (state != IterativeParsingFinishState)
            HandleError(state, is);

        return parseResult_;
    }

    static const size_t kDefaultStackCapacity = 256;    //!< Default stack capacity in bytes for storing a single decoded string.
    internal::Stack<StackAllocator> stack_;  //!< A stack for storing decoded string temporarily during non-destructive parsing.
    ParseResult parseResult_;
    IterativeParsingState state_;
}; // class GenericReader

//! Reader with UTF8 encoding and default allocator.
typedef GenericReader<UTF8<>, UTF8<> > Reader;

CEREAL_RAPIDJSON_NAMESPACE_END

#if defined(__clang__) || defined(_MSC_VER)
CEREAL_RAPIDJSON_DIAG_POP
#endif


#ifdef __GNUC__
CEREAL_RAPIDJSON_DIAG_POP
#endif

#endif // CEREAL_RAPIDJSON_READER_H_
```
