## Supplemental code for "The motif composition of variable-number tandem repeats impacts gene expression": schema.h

```
// Tencent is pleased to support the open source community by making RapidJSON available->
// 
// Copyright (C) 2015 THL A29 Limited, a Tencent company, and Milo Yip-> All rights reserved->
//
// Licensed under the MIT License (the "License"); you may not use this file except
// in compliance with the License-> You may obtain a copy of the License at
//
// http://opensource->org/licenses/MIT
//
// Unless required by applicable law or agreed to in writing, software distributed 
// under the License is distributed on an "AS IS" BASIS, WITHOUT WARRANTIES OR 
// CONDITIONS OF ANY KIND, either express or implied-> See the License for the 
// specific language governing permissions and limitations under the License->

#ifndef CEREAL_RAPIDJSON_SCHEMA_H_
#define CEREAL_RAPIDJSON_SCHEMA_H_

#include "document.h"
#include "pointer.h"
#include "stringbuffer.h"
#include <cmath> // abs, floor

#if !defined(CEREAL_RAPIDJSON_SCHEMA_USE_INTERNALREGEX)
#define CEREAL_RAPIDJSON_SCHEMA_USE_INTERNALREGEX 1
#else
#define CEREAL_RAPIDJSON_SCHEMA_USE_INTERNALREGEX 0
#endif

#if !CEREAL_RAPIDJSON_SCHEMA_USE_INTERNALREGEX && defined(CEREAL_RAPIDJSON_SCHEMA_USE_STDREGEX) && (__cplusplus >=201103L || (defined(_MSC_VER) && _MSC_VER >= 1800))
#define CEREAL_RAPIDJSON_SCHEMA_USE_STDREGEX 1
#else
#define CEREAL_RAPIDJSON_SCHEMA_USE_STDREGEX 0
#endif

#if CEREAL_RAPIDJSON_SCHEMA_USE_INTERNALREGEX
#include "internal/regex.h"
#elif CEREAL_RAPIDJSON_SCHEMA_USE_STDREGEX
#include <regex>
#endif

#if CEREAL_RAPIDJSON_SCHEMA_USE_INTERNALREGEX || CEREAL_RAPIDJSON_SCHEMA_USE_STDREGEX
#define CEREAL_RAPIDJSON_SCHEMA_HAS_REGEX 1
#else
#define CEREAL_RAPIDJSON_SCHEMA_HAS_REGEX 0
#endif

#ifndef CEREAL_RAPIDJSON_SCHEMA_VERBOSE
#define CEREAL_RAPIDJSON_SCHEMA_VERBOSE 0
#endif

#if CEREAL_RAPIDJSON_SCHEMA_VERBOSE
#include "stringbuffer.h"
#endif

CEREAL_RAPIDJSON_DIAG_PUSH

#if defined(__GNUC__)
CEREAL_RAPIDJSON_DIAG_OFF(effc++)
#endif

#ifdef __clang__
CEREAL_RAPIDJSON_DIAG_OFF(weak-vtables)
CEREAL_RAPIDJSON_DIAG_OFF(exit-time-destructors)
CEREAL_RAPIDJSON_DIAG_OFF(c++98-compat-pedantic)
CEREAL_RAPIDJSON_DIAG_OFF(variadic-macros)
#elif defined(_MSC_VER)
CEREAL_RAPIDJSON_DIAG_OFF(4512) // assignment operator could not be generated
#endif

CEREAL_RAPIDJSON_NAMESPACE_BEGIN

///////////////////////////////////////////////////////////////////////////////
// Verbose Utilities

#if CEREAL_RAPIDJSON_SCHEMA_VERBOSE

namespace internal {

inline void PrintInvalidKeyword(const char* keyword) {
    printf("Fail keyword: %s\n", keyword);
}

inline void PrintInvalidKeyword(const wchar_t* keyword) {
    wprintf(L"Fail keyword: %ls\n", keyword);
}

inline void PrintInvalidDocument(const char* document) {
    printf("Fail document: %s\n\n", document);
}

inline void PrintInvalidDocument(const wchar_t* document) {
    wprintf(L"Fail document: %ls\n\n", document);
}

inline void PrintValidatorPointers(unsigned depth, const char* s, const char* d) {
    printf("S: %*s%s\nD: %*s%s\n\n", depth * 4, " ", s, depth * 4, " ", d);
}

inline void PrintValidatorPointers(unsigned depth, const wchar_t* s, const wchar_t* d) {
    wprintf(L"S: %*ls%ls\nD: %*ls%ls\n\n", depth * 4, L" ", s, depth * 4, L" ", d);
}

} // namespace internal

#endif // CEREAL_RAPIDJSON_SCHEMA_VERBOSE

///////////////////////////////////////////////////////////////////////////////
// CEREAL_RAPIDJSON_INVALID_KEYWORD_RETURN

#if CEREAL_RAPIDJSON_SCHEMA_VERBOSE
#define CEREAL_RAPIDJSON_INVALID_KEYWORD_VERBOSE(keyword) internal::PrintInvalidKeyword(keyword)
#else
#define CEREAL_RAPIDJSON_INVALID_KEYWORD_VERBOSE(keyword)
#endif

#define CEREAL_RAPIDJSON_INVALID_KEYWORD_RETURN(keyword)\
CEREAL_RAPIDJSON_MULTILINEMACRO_BEGIN\
    context.invalidKeyword = keyword.GetString();\
    CEREAL_RAPIDJSON_INVALID_KEYWORD_VERBOSE(keyword.GetString());\
    return false;\
CEREAL_RAPIDJSON_MULTILINEMACRO_END

///////////////////////////////////////////////////////////////////////////////
// Forward declarations

template <typename ValueType, typename Allocator>
class GenericSchemaDocument;

namespace internal {

template <typename SchemaDocumentType>
class Schema;

///////////////////////////////////////////////////////////////////////////////
// ISchemaValidator

class ISchemaValidator {
public:
    virtual ~ISchemaValidator() {}
    virtual bool IsValid() const = 0;
};

///////////////////////////////////////////////////////////////////////////////
// ISchemaStateFactory

template <typename SchemaType>
class ISchemaStateFactory {
public:
    virtual ~ISchemaStateFactory() {}
    virtual ISchemaValidator* CreateSchemaValidator(const SchemaType&) = 0;
    virtual void DestroySchemaValidator(ISchemaValidator* validator) = 0;
    virtual void* CreateHasher() = 0;
    virtual uint64_t GetHashCode(void* hasher) = 0;
    virtual void DestroryHasher(void* hasher) = 0;
    virtual void* MallocState(size_t size) = 0;
    virtual void FreeState(void* p) = 0;
};

///////////////////////////////////////////////////////////////////////////////
// IValidationErrorHandler

template <typename SchemaType>
class IValidationErrorHandler {
public:
    typedef typename SchemaType::Ch Ch;
    typedef typename SchemaType::SValue SValue;

    virtual ~IValidationErrorHandler() {}

    virtual void NotMultipleOf(int64_t actual, const SValue& expected) = 0;
    virtual void NotMultipleOf(uint64_t actual, const SValue& expected) = 0;
    virtual void NotMultipleOf(double actual, const SValue& expected) = 0;
    virtual void AboveMaximum(int64_t actual, const SValue& expected, bool exclusive) = 0;
    virtual void AboveMaximum(uint64_t actual, const SValue& expected, bool exclusive) = 0;
    virtual void AboveMaximum(double actual, const SValue& expected, bool exclusive) = 0;
    virtual void BelowMinimum(int64_t actual, const SValue& expected, bool exclusive) = 0;
    virtual void BelowMinimum(uint64_t actual, const SValue& expected, bool exclusive) = 0;
    virtual void BelowMinimum(double actual, const SValue& expected, bool exclusive) = 0;

    virtual void TooLong(const Ch* str, SizeType length, SizeType expected) = 0;
    virtual void TooShort(const Ch* str, SizeType length, SizeType expected) = 0;
    virtual void DoesNotMatch(const Ch* str, SizeType length) = 0;

    virtual void DisallowedItem(SizeType index) = 0;
    virtual void TooFewItems(SizeType actualCount, SizeType expectedCount) = 0;
    virtual void TooManyItems(SizeType actualCount, SizeType expectedCount) = 0;
    virtual void DuplicateItems(SizeType index1, SizeType index2) = 0;

    virtual void TooManyProperties(SizeType actualCount, SizeType expectedCount) = 0;
    virtual void TooFewProperties(SizeType actualCount, SizeType expectedCount) = 0;
    virtual void StartMissingProperties() = 0;
    virtual void AddMissingProperty(const SValue& name) = 0;
    virtual bool EndMissingProperties() = 0;
    virtual void PropertyViolations(ISchemaValidator** subvalidators, SizeType count) = 0;
    virtual void DisallowedProperty(const Ch* name, SizeType length) = 0;

    virtual void StartDependencyErrors() = 0;
    virtual void StartMissingDependentProperties() = 0;
    virtual void AddMissingDependentProperty(const SValue& targetName) = 0;
    virtual void EndMissingDependentProperties(const SValue& sourceName) = 0;
    virtual void AddDependencySchemaError(const SValue& souceName, ISchemaValidator* subvalidator) = 0;
    virtual bool EndDependencyErrors() = 0;

    virtual void DisallowedValue() = 0;
    virtual void StartDisallowedType() = 0;
    virtual void AddExpectedType(const typename SchemaType::ValueType& expectedType) = 0;
    virtual void EndDisallowedType(const typename SchemaType::ValueType& actualType) = 0;
    virtual void NotAllOf(ISchemaValidator** subvalidators, SizeType count) = 0;
    virtual void NoneOf(ISchemaValidator** subvalidators, SizeType count) = 0;
    virtual void NotOneOf(ISchemaValidator** subvalidators, SizeType count) = 0;
    virtual void Disallowed() = 0;
};


///////////////////////////////////////////////////////////////////////////////
// Hasher

// For comparison of compound value
template<typename Encoding, typename Allocator>
class Hasher {
public:
    typedef typename Encoding::Ch Ch;

    Hasher(Allocator* allocator = 0, size_t stackCapacity = kDefaultSize) : stack_(allocator, stackCapacity) {}

    bool Null() { return WriteType(kNullType); }
    bool Bool(bool b) { return WriteType(b ? kTrueType : kFalseType); }
    bool Int(int i) { Number n; n.u.i = i; n.d = static_cast<double>(i); return WriteNumber(n); }
    bool Uint(unsigned u) { Number n; n.u.u = u; n.d = static_cast<double>(u); return WriteNumber(n); }
    bool Int64(int64_t i) { Number n; n.u.i = i; n.d = static_cast<double>(i); return WriteNumber(n); }
    bool Uint64(uint64_t u) { Number n; n.u.u = u; n.d = static_cast<double>(u); return WriteNumber(n); }
    bool Double(double d) { 
        Number n; 
        if (d < 0) n.u.i = static_cast<int64_t>(d);
        else       n.u.u = static_cast<uint64_t>(d); 
        n.d = d;
        return WriteNumber(n);
    }

    bool RawNumber(const Ch* str, SizeType len, bool) {
        WriteBuffer(kNumberType, str, len * sizeof(Ch));
        return true;
    }

    bool String(const Ch* str, SizeType len, bool) {
        WriteBuffer(kStringType, str, len * sizeof(Ch));
        return true;
    }

    bool StartObject() { return true; }
    bool Key(const Ch* str, SizeType len, bool copy) { return String(str, len, copy); }
    bool EndObject(SizeType memberCount) { 
        uint64_t h = Hash(0, kObjectType);
        uint64_t* kv = stack_.template Pop<uint64_t>(memberCount * 2);
        for (SizeType i = 0; i < memberCount; i++)
            h ^= Hash(kv[i * 2], kv[i * 2 + 1]);  // Use xor to achieve member order insensitive
        *stack_.template Push<uint64_t>() = h;
        return true;
    }
    
    bool StartArray() { return true; }
    bool EndArray(SizeType elementCount) { 
        uint64_t h = Hash(0, kArrayType);
        uint64_t* e = stack_.template Pop<uint64_t>(elementCount);
        for (SizeType i = 0; i < elementCount; i++)
            h = Hash(h, e[i]); // Use hash to achieve element order sensitive
        *stack_.template Push<uint64_t>() = h;
        return true;
    }

    bool IsValid() const { return stack_.GetSize() == sizeof(uint64_t); }

    uint64_t GetHashCode() const {
        CEREAL_RAPIDJSON_ASSERT(IsValid());
        return *stack_.template Top<uint64_t>();
    }

private:
    static const size_t kDefaultSize = 256;
    struct Number {
        union U {
            uint64_t u;
            int64_t i;
        }u;
        double d;
    };

    bool WriteType(Type type) { return WriteBuffer(type, 0, 0); }
    
    bool WriteNumber(const Number& n) { return WriteBuffer(kNumberType, &n, sizeof(n)); }
    
    bool WriteBuffer(Type type, const void* data, size_t len) {
        // FNV-1a from http://isthe.com/chongo/tech/comp/fnv/
        uint64_t h = Hash(CEREAL_RAPIDJSON_UINT64_C2(0x84222325, 0xcbf29ce4), type);
        const unsigned char* d = static_cast<const unsigned char*>(data);
        for (size_t i = 0; i < len; i++)
            h = Hash(h, d[i]);
        *stack_.template Push<uint64_t>() = h;
        return true;
    }

    static uint64_t Hash(uint64_t h, uint64_t d) {
        static const uint64_t kPrime = CEREAL_RAPIDJSON_UINT64_C2(0x00000100, 0x000001b3);
        h ^= d;
        h *= kPrime;
        return h;
    }

    Stack<Allocator> stack_;
};

///////////////////////////////////////////////////////////////////////////////
// SchemaValidationContext

template <typename SchemaDocumentType>
struct SchemaValidationContext {
    typedef Schema<SchemaDocumentType> SchemaType;
    typedef ISchemaStateFactory<SchemaType> SchemaValidatorFactoryType;
    typedef IValidationErrorHandler<SchemaType> ErrorHandlerType;
    typedef typename SchemaType::ValueType ValueType;
    typedef typename ValueType::Ch Ch;

    enum PatternValidatorType {
        kPatternValidatorOnly,
        kPatternValidatorWithProperty,
        kPatternValidatorWithAdditionalProperty
    };

    SchemaValidationContext(SchemaValidatorFactoryType& f, ErrorHandlerType& eh, const SchemaType* s) :
        factory(f),
        error_handler(eh),
        schema(s),
        valueSchema(),
        invalidKeyword(),
        hasher(),
        arrayElementHashCodes(),
        validators(),
        validatorCount(),
        patternPropertiesValidators(),
        patternPropertiesValidatorCount(),
        patternPropertiesSchemas(),
        patternPropertiesSchemaCount(),
        valuePatternValidatorType(kPatternValidatorOnly),
        propertyExist(),
        inArray(false),
        valueUniqueness(false),
        arrayUniqueness(false)
    {
    }

    ~SchemaValidationContext() {
        if (hasher)
            factory.DestroryHasher(hasher);
        if (validators) {
            for (SizeType i = 0; i < validatorCount; i++)
                factory.DestroySchemaValidator(validators[i]);
            factory.FreeState(validators);
        }
        if (patternPropertiesValidators) {
            for (SizeType i = 0; i < patternPropertiesValidatorCount; i++)
                factory.DestroySchemaValidator(patternPropertiesValidators[i]);
            factory.FreeState(patternPropertiesValidators);
        }
        if (patternPropertiesSchemas)
            factory.FreeState(patternPropertiesSchemas);
        if (propertyExist)
            factory.FreeState(propertyExist);
    }

    SchemaValidatorFactoryType& factory;
    ErrorHandlerType& error_handler;
    const SchemaType* schema;
    const SchemaType* valueSchema;
    const Ch* invalidKeyword;
    void* hasher; // Only validator access
    void* arrayElementHashCodes; // Only validator access this
    ISchemaValidator** validators;
    SizeType validatorCount;
    ISchemaValidator** patternPropertiesValidators;
    SizeType patternPropertiesValidatorCount;
    const SchemaType** patternPropertiesSchemas;
    SizeType patternPropertiesSchemaCount;
    PatternValidatorType valuePatternValidatorType;
    PatternValidatorType objectPatternValidatorType;
    SizeType arrayElementIndex;
    bool* propertyExist;
    bool inArray;
    bool valueUniqueness;
    bool arrayUniqueness;
};

///////////////////////////////////////////////////////////////////////////////
// Schema

template <typename SchemaDocumentType>
class Schema {
public:
    typedef typename SchemaDocumentType::ValueType ValueType;
    typedef typename SchemaDocumentType::AllocatorType AllocatorType;
    typedef typename SchemaDocumentType::PointerType PointerType;
    typedef typename ValueType::EncodingType EncodingType;
    typedef typename EncodingType::Ch Ch;
    typedef SchemaValidationContext<SchemaDocumentType> Context;
    typedef Schema<SchemaDocumentType> SchemaType;
    typedef GenericValue<EncodingType, AllocatorType> SValue;
    typedef IValidationErrorHandler<Schema> ErrorHandler;
    friend class GenericSchemaDocument<ValueType, AllocatorType>;

    Schema(SchemaDocumentType* schemaDocument, const PointerType& p, const ValueType& value, const ValueType& document, AllocatorType* allocator) :
        allocator_(allocator),
        uri_(schemaDocument->GetURI(), *allocator),
        pointer_(p, allocator),
        typeless_(schemaDocument->GetTypeless()),
        enum_(),
        enumCount_(),
        not_(),
        type_((1 << kTotalSchemaType) - 1), // typeless
        validatorCount_(),
        notValidatorIndex_(),
        properties_(),
        additionalPropertiesSchema_(),
        patternProperties_(),
        patternPropertyCount_(),
        propertyCount_(),
        minProperties_(),
        maxProperties_(SizeType(~0)),
        additionalProperties_(true),
        hasDependencies_(),
        hasRequired_(),
        hasSchemaDependencies_(),
        additionalItemsSchema_(),
        itemsList_(),
        itemsTuple_(),
        itemsTupleCount_(),
        minItems_(),
        maxItems_(SizeType(~0)),
        additionalItems_(true),
        uniqueItems_(false),
        pattern_(),
        minLength_(0),
        maxLength_(~SizeType(0)),
        exclusiveMinimum_(false),
        exclusiveMaximum_(false),
        defaultValueLength_(0)
    {
        typedef typename SchemaDocumentType::ValueType ValueType;
        typedef typename ValueType::ConstValueIterator ConstValueIterator;
        typedef typename ValueType::ConstMemberIterator ConstMemberIterator;

        if (!value.IsObject())
            return;

        if (const ValueType* v = GetMember(value, GetTypeString())) {
            type_ = 0;
            if (v->IsString())
                AddType(*v);
            else if (v->IsArray())
                for (ConstValueIterator itr = v->Begin(); itr != v->End(); ++itr)
                    AddType(*itr);
        }

        if (const ValueType* v = GetMember(value, GetEnumString()))
            if (v->IsArray() && v->Size() > 0) {
                enum_ = static_cast<uint64_t*>(allocator_->Malloc(sizeof(uint64_t) * v->Size()));
                for (ConstValueIterator itr = v->Begin(); itr != v->End(); ++itr) {
                    typedef Hasher<EncodingType, MemoryPoolAllocator<> > EnumHasherType;
                    char buffer[256u + 24];
                    MemoryPoolAllocator<> hasherAllocator(buffer, sizeof(buffer));
                    EnumHasherType h(&hasherAllocator, 256);
                    itr->Accept(h);
                    enum_[enumCount_++] = h.GetHashCode();
                }
            }

        if (schemaDocument) {
            AssignIfExist(allOf_, *schemaDocument, p, value, GetAllOfString(), document);
            AssignIfExist(anyOf_, *schemaDocument, p, value, GetAnyOfString(), document);
            AssignIfExist(oneOf_, *schemaDocument, p, value, GetOneOfString(), document);
        }

        if (const ValueType* v = GetMember(value, GetNotString())) {
            schemaDocument->CreateSchema(&not_, p.Append(GetNotString(), allocator_), *v, document);
            notValidatorIndex_ = validatorCount_;
            validatorCount_++;
        }

        // Object

        const ValueType* properties = GetMember(value, GetPropertiesString());
        const ValueType* required = GetMember(value, GetRequiredString());
        const ValueType* dependencies = GetMember(value, GetDependenciesString());
        {
            // Gather properties from properties/required/dependencies
            SValue allProperties(kArrayType);

            if (properties && properties->IsObject())
                for (ConstMemberIterator itr = properties->MemberBegin(); itr != properties->MemberEnd(); ++itr)
                    AddUniqueElement(allProperties, itr->name);
            
            if (required && required->IsArray())
                for (ConstValueIterator itr = required->Begin(); itr != required->End(); ++itr)
                    if (itr->IsString())
                        AddUniqueElement(allProperties, *itr);

            if (dependencies && dependencies->IsObject())
                for (ConstMemberIterator itr = dependencies->MemberBegin(); itr != dependencies->MemberEnd(); ++itr) {
                    AddUniqueElement(allProperties, itr->name);
                    if (itr->value.IsArray())
                        for (ConstValueIterator i = itr->value.Begin(); i != itr->value.End(); ++i)
                            if (i->IsString())
                                AddUniqueElement(allProperties, *i);
                }

            if (allProperties.Size() > 0) {
                propertyCount_ = allProperties.Size();
                properties_ = static_cast<Property*>(allocator_->Malloc(sizeof(Property) * propertyCount_));
                for (SizeType i = 0; i < propertyCount_; i++) {
                    new (&properties_[i]) Property();
                    properties_[i].name = allProperties[i];
                    properties_[i].schema = typeless_;
                }
            }
        }

        if (properties && properties->IsObject()) {
            PointerType q = p.Append(GetPropertiesString(), allocator_);
            for (ConstMemberIterator itr = properties->MemberBegin(); itr != properties->MemberEnd(); ++itr) {
                SizeType index;
                if (FindPropertyIndex(itr->name, &index))
                    schemaDocument->CreateSchema(&properties_[index].schema, q.Append(itr->name, allocator_), itr->value, document);
            }
        }

        if (const ValueType* v = GetMember(value, GetPatternPropertiesString())) {
            PointerType q = p.Append(GetPatternPropertiesString(), allocator_);
            patternProperties_ = static_cast<PatternProperty*>(allocator_->Malloc(sizeof(PatternProperty) * v->MemberCount()));
            patternPropertyCount_ = 0;

            for (ConstMemberIterator itr = v->MemberBegin(); itr != v->MemberEnd(); ++itr) {
                new (&patternProperties_[patternPropertyCount_]) PatternProperty();
                patternProperties_[patternPropertyCount_].pattern = CreatePattern(itr->name);
                schemaDocument->CreateSchema(&patternProperties_[patternPropertyCount_].schema, q.Append(itr->name, allocator_), itr->value, document);
                patternPropertyCount_++;
            }
        }

        if (required && required->IsArray())
            for (ConstValueIterator itr = required->Begin(); itr != required->End(); ++itr)
                if (itr->IsString()) {
                    SizeType index;
                    if (FindPropertyIndex(*itr, &index)) {
                        properties_[index].required = true;
                        hasRequired_ = true;
                    }
                }

        if (dependencies && dependencies->IsObject()) {
            PointerType q = p.Append(GetDependenciesString(), allocator_);
            hasDependencies_ = true;
            for (ConstMemberIterator itr = dependencies->MemberBegin(); itr != dependencies->MemberEnd(); ++itr) {
                SizeType sourceIndex;
                if (FindPropertyIndex(itr->name, &sourceIndex)) {
                    if (itr->value.IsArray()) {
                        properties_[sourceIndex].dependencies = static_cast<bool*>(allocator_->Malloc(sizeof(bool) * propertyCount_));
                        std::memset(properties_[sourceIndex].dependencies, 0, sizeof(bool)* propertyCount_);
                        for (ConstValueIterator targetItr = itr->value.Begin(); targetItr != itr->value.End(); ++targetItr) {
                            SizeType targetIndex;
                            if (FindPropertyIndex(*targetItr, &targetIndex))
                                properties_[sourceIndex].dependencies[targetIndex] = true;
                        }
                    }
                    else if (itr->value.IsObject()) {
                        hasSchemaDependencies_ = true;
                        schemaDocument->CreateSchema(&properties_[sourceIndex].dependenciesSchema, q.Append(itr->name, allocator_), itr->value, document);
                        properties_[sourceIndex].dependenciesValidatorIndex = validatorCount_;
                        validatorCount_++;
                    }
                }
            }
        }

        if (const ValueType* v = GetMember(value, GetAdditionalPropertiesString())) {
            if (v->IsBool())
                additionalProperties_ = v->GetBool();
            else if (v->IsObject())
                schemaDocument->CreateSchema(&additionalPropertiesSchema_, p.Append(GetAdditionalPropertiesString(), allocator_), *v, document);
        }

        AssignIfExist(minProperties_, value, GetMinPropertiesString());
        AssignIfExist(maxProperties_, value, GetMaxPropertiesString());

        // Array
        if (const ValueType* v = GetMember(value, GetItemsString())) {
            PointerType q = p.Append(GetItemsString(), allocator_);
            if (v->IsObject()) // List validation
                schemaDocument->CreateSchema(&itemsList_, q, *v, document);
            else if (v->IsArray()) { // Tuple validation
                itemsTuple_ = static_cast<const Schema**>(allocator_->Malloc(sizeof(const Schema*) * v->Size()));
                SizeType index = 0;
                for (ConstValueIterator itr = v->Begin(); itr != v->End(); ++itr, index++)
                    schemaDocument->CreateSchema(&itemsTuple_[itemsTupleCount_++], q.Append(index, allocator_), *itr, document);
            }
        }

        AssignIfExist(minItems_, value, GetMinItemsString());
        AssignIfExist(maxItems_, value, GetMaxItemsString());

        if (const ValueType* v = GetMember(value, GetAdditionalItemsString())) {
            if (v->IsBool())
                additionalItems_ = v->GetBool();
            else if (v->IsObject())
                schemaDocument->CreateSchema(&additionalItemsSchema_, p.Append(GetAdditionalItemsString(), allocator_), *v, document);
        }

        AssignIfExist(uniqueItems_, value, GetUniqueItemsString());

        // String
        AssignIfExist(minLength_, value, GetMinLengthString());
        AssignIfExist(maxLength_, value, GetMaxLengthString());

        if (const ValueType* v = GetMember(value, GetPatternString()))
            pattern_ = CreatePattern(*v);

        // Number
        if (const ValueType* v = GetMember(value, GetMinimumString()))
            if (v->IsNumber())
                minimum_.CopyFrom(*v, *allocator_);

        if (const ValueType* v = GetMember(value, GetMaximumString()))
            if (v->IsNumber())
                maximum_.CopyFrom(*v, *allocator_);

        AssignIfExist(exclusiveMinimum_, value, GetExclusiveMinimumString());
        AssignIfExist(exclusiveMaximum_, value, GetExclusiveMaximumString());

        if (const ValueType* v = GetMember(value, GetMultipleOfString()))
            if (v->IsNumber() && v->GetDouble() > 0.0)
                multipleOf_.CopyFrom(*v, *allocator_);

        // Default
        if (const ValueType* v = GetMember(value, GetDefaultValueString()))
            if (v->IsString())
                defaultValueLength_ = v->GetStringLength();

    }

    ~Schema() {
        AllocatorType::Free(enum_);
        if (properties_) {
            for (SizeType i = 0; i < propertyCount_; i++)
                properties_[i].~Property();
            AllocatorType::Free(properties_);
        }
        if (patternProperties_) {
            for (SizeType i = 0; i < patternPropertyCount_; i++)
                patternProperties_[i].~PatternProperty();
            AllocatorType::Free(patternProperties_);
        }
        AllocatorType::Free(itemsTuple_);
#if CEREAL_RAPIDJSON_SCHEMA_HAS_REGEX
        if (pattern_) {
            pattern_->~RegexType();
            AllocatorType::Free(pattern_);
        }
#endif
    }

    const SValue& GetURI() const {
        return uri_;
    }

    const PointerType& GetPointer() const {
        return pointer_;
    }

    bool BeginValue(Context& context) const {
        if (context.inArray) {
            if (uniqueItems_)
                context.valueUniqueness = true;

            if (itemsList_)
                context.valueSchema = itemsList_;
            else if (itemsTuple_) {
                if (context.arrayElementIndex < itemsTupleCount_)
                    context.valueSchema = itemsTuple_[context.arrayElementIndex];
                else if (additionalItemsSchema_)
                    context.valueSchema = additionalItemsSchema_;
                else if (additionalItems_)
                    context.valueSchema = typeless_;
                else {
                    context.error_handler.DisallowedItem(context.arrayElementIndex);
                    CEREAL_RAPIDJSON_INVALID_KEYWORD_RETURN(GetItemsString());
                }
            }
            else
                context.valueSchema = typeless_;

            context.arrayElementIndex++;
        }
        return true;
    }

    CEREAL_RAPIDJSON_FORCEINLINE bool EndValue(Context& context) const {
        if (context.patternPropertiesValidatorCount > 0) {
            bool otherValid = false;
            SizeType count = context.patternPropertiesValidatorCount;
            if (context.objectPatternValidatorType != Context::kPatternValidatorOnly)
                otherValid = context.patternPropertiesValidators[--count]->IsValid();

            bool patternValid = true;
            for (SizeType i = 0; i < count; i++)
                if (!context.patternPropertiesValidators[i]->IsValid()) {
                    patternValid = false;
                    break;
                }

            if (context.objectPatternValidatorType == Context::kPatternValidatorOnly) {
                if (!patternValid) {
                    context.error_handler.PropertyViolations(context.patternPropertiesValidators, count);
                    CEREAL_RAPIDJSON_INVALID_KEYWORD_RETURN(GetPatternPropertiesString());
                }
            }
            else if (context.objectPatternValidatorType == Context::kPatternValidatorWithProperty) {
                if (!patternValid || !otherValid) {
                    context.error_handler.PropertyViolations(context.patternPropertiesValidators, count + 1);
                    CEREAL_RAPIDJSON_INVALID_KEYWORD_RETURN(GetPatternPropertiesString());
                }
            }
            else if (!patternValid && !otherValid) { // kPatternValidatorWithAdditionalProperty)
                context.error_handler.PropertyViolations(context.patternPropertiesValidators, count + 1);
                CEREAL_RAPIDJSON_INVALID_KEYWORD_RETURN(GetPatternPropertiesString());
            }
        }

        if (enum_) {
            const uint64_t h = context.factory.GetHashCode(context.hasher);
            for (SizeType i = 0; i < enumCount_; i++)
                if (enum_[i] == h)
                    goto foundEnum;
            context.error_handler.DisallowedValue();
            CEREAL_RAPIDJSON_INVALID_KEYWORD_RETURN(GetEnumString());
            foundEnum:;
        }

        if (allOf_.schemas)
            for (SizeType i = allOf_.begin; i < allOf_.begin + allOf_.count; i++)
                if (!context.validators[i]->IsValid()) {
                    context.error_handler.NotAllOf(&context.validators[allOf_.begin], allOf_.count);
                    CEREAL_RAPIDJSON_INVALID_KEYWORD_RETURN(GetAllOfString());
                }
        
        if (anyOf_.schemas) {
            for (SizeType i = anyOf_.begin; i < anyOf_.begin + anyOf_.count; i++)
                if (context.validators[i]->IsValid())
                    goto foundAny;
            context.error_handler.NoneOf(&context.validators[anyOf_.begin], anyOf_.count);
            CEREAL_RAPIDJSON_INVALID_KEYWORD_RETURN(GetAnyOfString());
            foundAny:;
        }

        if (oneOf_.schemas) {
            bool oneValid = false;
            for (SizeType i = oneOf_.begin; i < oneOf_.begin + oneOf_.count; i++)
                if (context.validators[i]->IsValid()) {
                    if (oneValid) {
                        context.error_handler.NotOneOf(&context.validators[oneOf_.begin], oneOf_.count);
                        CEREAL_RAPIDJSON_INVALID_KEYWORD_RETURN(GetOneOfString());
                    } else
                        oneValid = true;
                }
            if (!oneValid) {
                context.error_handler.NotOneOf(&context.validators[oneOf_.begin], oneOf_.count);
                CEREAL_RAPIDJSON_INVALID_KEYWORD_RETURN(GetOneOfString());
            }
        }

        if (not_ && context.validators[notValidatorIndex_]->IsValid()) {
            context.error_handler.Disallowed();
            CEREAL_RAPIDJSON_INVALID_KEYWORD_RETURN(GetNotString());
        }

        return true;
    }

    bool Null(Context& context) const {
        if (!(type_ & (1 << kNullSchemaType))) {
            DisallowedType(context, GetNullString());
            CEREAL_RAPIDJSON_INVALID_KEYWORD_RETURN(GetTypeString());
        }
        return CreateParallelValidator(context);
    }
    
    bool Bool(Context& context, bool) const {
        if (!(type_ & (1 << kBooleanSchemaType))) {
            DisallowedType(context, GetBooleanString());
            CEREAL_RAPIDJSON_INVALID_KEYWORD_RETURN(GetTypeString());
        }
        return CreateParallelValidator(context);
    }

    bool Int(Context& context, int i) const {
        if (!CheckInt(context, i))
            return false;
        return CreateParallelValidator(context);
    }

    bool Uint(Context& context, unsigned u) const {
        if (!CheckUint(context, u))
            return false;
        return CreateParallelValidator(context);
    }

    bool Int64(Context& context, int64_t i) const {
        if (!CheckInt(context, i))
            return false;
        return CreateParallelValidator(context);
    }

    bool Uint64(Context& context, uint64_t u) const {
        if (!CheckUint(context, u))
            return false;
        return CreateParallelValidator(context);
    }

    bool Double(Context& context, double d) const {
        if (!(type_ & (1 << kNumberSchemaType))) {
            DisallowedType(context, GetNumberString());
            CEREAL_RAPIDJSON_INVALID_KEYWORD_RETURN(GetTypeString());
        }

        if (!minimum_.IsNull() && !CheckDoubleMinimum(context, d))
            return false;

        if (!maximum_.IsNull() && !CheckDoubleMaximum(context, d))
            return false;
        
        if (!multipleOf_.IsNull() && !CheckDoubleMultipleOf(context, d))
            return false;
        
        return CreateParallelValidator(context);
    }
    
    bool String(Context& context, const Ch* str, SizeType length, bool) const {
        if (!(type_ & (1 << kStringSchemaType))) {
            DisallowedType(context, GetStringString());
            CEREAL_RAPIDJSON_INVALID_KEYWORD_RETURN(GetTypeString());
        }

        if (minLength_ != 0 || maxLength_ != SizeType(~0)) {
            SizeType count;
            if (internal::CountStringCodePoint<EncodingType>(str, length, &count)) {
                if (count < minLength_) {
                    context.error_handler.TooShort(str, length, minLength_);
                    CEREAL_RAPIDJSON_INVALID_KEYWORD_RETURN(GetMinLengthString());
                }
                if (count > maxLength_) {
                    context.error_handler.TooLong(str, length, maxLength_);
                    CEREAL_RAPIDJSON_INVALID_KEYWORD_RETURN(GetMaxLengthString());
                }
            }
        }

        if (pattern_ && !IsPatternMatch(pattern_, str, length)) {
            context.error_handler.DoesNotMatch(str, length);
            CEREAL_RAPIDJSON_INVALID_KEYWORD_RETURN(GetPatternString());
        }

        return CreateParallelValidator(context);
    }

    bool StartObject(Context& context) const {
        if (!(type_ & (1 << kObjectSchemaType))) {
            DisallowedType(context, GetObjectString());
            CEREAL_RAPIDJSON_INVALID_KEYWORD_RETURN(GetTypeString());
        }

        if (hasDependencies_ || hasRequired_) {
            context.propertyExist = static_cast<bool*>(context.factory.MallocState(sizeof(bool) * propertyCount_));
            std::memset(context.propertyExist, 0, sizeof(bool) * propertyCount_);
        }

        if (patternProperties_) { // pre-allocate schema array
            SizeType count = patternPropertyCount_ + 1; // extra for valuePatternValidatorType
            context.patternPropertiesSchemas = static_cast<const SchemaType**>(context.factory.MallocState(sizeof(const SchemaType*) * count));
            context.patternPropertiesSchemaCount = 0;
            std::memset(context.patternPropertiesSchemas, 0, sizeof(SchemaType*) * count);
        }

        return CreateParallelValidator(context);
    }
    
    bool Key(Context& context, const Ch* str, SizeType len, bool) const {
        if (patternProperties_) {
            context.patternPropertiesSchemaCount = 0;
            for (SizeType i = 0; i < patternPropertyCount_; i++)
                if (patternProperties_[i].pattern && IsPatternMatch(patternProperties_[i].pattern, str, len)) {
                    context.patternPropertiesSchemas[context.patternPropertiesSchemaCount++] = patternProperties_[i].schema;
                    context.valueSchema = typeless_;
                }
        }

        SizeType index;
        if (FindPropertyIndex(ValueType(str, len).Move(), &index)) {
            if (context.patternPropertiesSchemaCount > 0) {
                context.patternPropertiesSchemas[context.patternPropertiesSchemaCount++] = properties_[index].schema;
                context.valueSchema = typeless_;
                context.valuePatternValidatorType = Context::kPatternValidatorWithProperty;
            }
            else
                context.valueSchema = properties_[index].schema;

            if (context.propertyExist)
                context.propertyExist[index] = true;

            return true;
        }

        if (additionalPropertiesSchema_) {
            if (additionalPropertiesSchema_ && context.patternPropertiesSchemaCount > 0) {
                context.patternPropertiesSchemas[context.patternPropertiesSchemaCount++] = additionalPropertiesSchema_;
                context.valueSchema = typeless_;
                context.valuePatternValidatorType = Context::kPatternValidatorWithAdditionalProperty;
            }
            else
                context.valueSchema = additionalPropertiesSchema_;
            return true;
        }
        else if (additionalProperties_) {
            context.valueSchema = typeless_;
            return true;
        }

        if (context.patternPropertiesSchemaCount == 0) { // patternProperties are not additional properties
            context.error_handler.DisallowedProperty(str, len);
            CEREAL_RAPIDJSON_INVALID_KEYWORD_RETURN(GetAdditionalPropertiesString());
        }

        return true;
    }

    bool EndObject(Context& context, SizeType memberCount) const {
        if (hasRequired_) {
            context.error_handler.StartMissingProperties();
            for (SizeType index = 0; index < propertyCount_; index++)
                if (properties_[index].required && !context.propertyExist[index])
                    if (properties_[index].schema->defaultValueLength_ == 0 )
                        context.error_handler.AddMissingProperty(properties_[index].name);
            if (context.error_handler.EndMissingProperties())
                CEREAL_RAPIDJSON_INVALID_KEYWORD_RETURN(GetRequiredString());
        }

        if (memberCount < minProperties_) {
            context.error_handler.TooFewProperties(memberCount, minProperties_);
            CEREAL_RAPIDJSON_INVALID_KEYWORD_RETURN(GetMinPropertiesString());
        }

        if (memberCount > maxProperties_) {
            context.error_handler.TooManyProperties(memberCount, maxProperties_);
            CEREAL_RAPIDJSON_INVALID_KEYWORD_RETURN(GetMaxPropertiesString());
        }

        if (hasDependencies_) {
            context.error_handler.StartDependencyErrors();
            for (SizeType sourceIndex = 0; sourceIndex < propertyCount_; sourceIndex++) {
                const Property& source = properties_[sourceIndex];
                if (context.propertyExist[sourceIndex]) {
                    if (source.dependencies) {
                        context.error_handler.StartMissingDependentProperties();
                        for (SizeType targetIndex = 0; targetIndex < propertyCount_; targetIndex++)
                            if (source.dependencies[targetIndex] && !context.propertyExist[targetIndex])
                                context.error_handler.AddMissingDependentProperty(properties_[targetIndex].name);
                        context.error_handler.EndMissingDependentProperties(source.name);
                    }
                    else if (source.dependenciesSchema) {
                        ISchemaValidator* dependenciesValidator = context.validators[source.dependenciesValidatorIndex];
                        if (!dependenciesValidator->IsValid())
                            context.error_handler.AddDependencySchemaError(source.name, dependenciesValidator);
                    }
                }
            }
            if (context.error_handler.EndDependencyErrors())
                CEREAL_RAPIDJSON_INVALID_KEYWORD_RETURN(GetDependenciesString());
        }

        return true;
    }

    bool StartArray(Context& context) const {
        if (!(type_ & (1 << kArraySchemaType))) {
            DisallowedType(context, GetArrayString());
            CEREAL_RAPIDJSON_INVALID_KEYWORD_RETURN(GetTypeString());
        }

        context.arrayElementIndex = 0;
        context.inArray = true;

        return CreateParallelValidator(context);
    }

    bool EndArray(Context& context, SizeType elementCount) const {
        context.inArray = false;
        
        if (elementCount < minItems_) {
            context.error_handler.TooFewItems(elementCount, minItems_);
            CEREAL_RAPIDJSON_INVALID_KEYWORD_RETURN(GetMinItemsString());
        }
        
        if (elementCount > maxItems_) {
            context.error_handler.TooManyItems(elementCount, maxItems_);
            CEREAL_RAPIDJSON_INVALID_KEYWORD_RETURN(GetMaxItemsString());
        }

        return true;
    }

    // Generate functions for string literal according to Ch
#define CEREAL_RAPIDJSON_STRING_(name, ...) \
    static const ValueType& Get##name##String() {\
        static const Ch s[] = { __VA_ARGS__, '\0' };\
        static const ValueType v(s, static_cast<SizeType>(sizeof(s) / sizeof(Ch) - 1));\
        return v;\
    }

    CEREAL_RAPIDJSON_STRING_(Null, 'n', 'u', 'l', 'l')
    CEREAL_RAPIDJSON_STRING_(Boolean, 'b', 'o', 'o', 'l', 'e', 'a', 'n')
    CEREAL_RAPIDJSON_STRING_(Object, 'o', 'b', 'j', 'e', 'c', 't')
    CEREAL_RAPIDJSON_STRING_(Array, 'a', 'r', 'r', 'a', 'y')
    CEREAL_RAPIDJSON_STRING_(String, 's', 't', 'r', 'i', 'n', 'g')
    CEREAL_RAPIDJSON_STRING_(Number, 'n', 'u', 'm', 'b', 'e', 'r')
    CEREAL_RAPIDJSON_STRING_(Integer, 'i', 'n', 't', 'e', 'g', 'e', 'r')
    CEREAL_RAPIDJSON_STRING_(Type, 't', 'y', 'p', 'e')
    CEREAL_RAPIDJSON_STRING_(Enum, 'e', 'n', 'u', 'm')
    CEREAL_RAPIDJSON_STRING_(AllOf, 'a', 'l', 'l', 'O', 'f')
    CEREAL_RAPIDJSON_STRING_(AnyOf, 'a', 'n', 'y', 'O', 'f')
    CEREAL_RAPIDJSON_STRING_(OneOf, 'o', 'n', 'e', 'O', 'f')
    CEREAL_RAPIDJSON_STRING_(Not, 'n', 'o', 't')
    CEREAL_RAPIDJSON_STRING_(Properties, 'p', 'r', 'o', 'p', 'e', 'r', 't', 'i', 'e', 's')
    CEREAL_RAPIDJSON_STRING_(Required, 'r', 'e', 'q', 'u', 'i', 'r', 'e', 'd')
    CEREAL_RAPIDJSON_STRING_(Dependencies, 'd', 'e', 'p', 'e', 'n', 'd', 'e', 'n', 'c', 'i', 'e', 's')
    CEREAL_RAPIDJSON_STRING_(PatternProperties, 'p', 'a', 't', 't', 'e', 'r', 'n', 'P', 'r', 'o', 'p', 'e', 'r', 't', 'i', 'e', 's')
    CEREAL_RAPIDJSON_STRING_(AdditionalProperties, 'a', 'd', 'd', 'i', 't', 'i', 'o', 'n', 'a', 'l', 'P', 'r', 'o', 'p', 'e', 'r', 't', 'i', 'e', 's')
    CEREAL_RAPIDJSON_STRING_(MinProperties, 'm', 'i', 'n', 'P', 'r', 'o', 'p', 'e', 'r', 't', 'i', 'e', 's')
    CEREAL_RAPIDJSON_STRING_(MaxProperties, 'm', 'a', 'x', 'P', 'r', 'o', 'p', 'e', 'r', 't', 'i', 'e', 's')
    CEREAL_RAPIDJSON_STRING_(Items, 'i', 't', 'e', 'm', 's')
    CEREAL_RAPIDJSON_STRING_(MinItems, 'm', 'i', 'n', 'I', 't', 'e', 'm', 's')
    CEREAL_RAPIDJSON_STRING_(MaxItems, 'm', 'a', 'x', 'I', 't', 'e', 'm', 's')
    CEREAL_RAPIDJSON_STRING_(AdditionalItems, 'a', 'd', 'd', 'i', 't', 'i', 'o', 'n', 'a', 'l', 'I', 't', 'e', 'm', 's')
    CEREAL_RAPIDJSON_STRING_(UniqueItems, 'u', 'n', 'i', 'q', 'u', 'e', 'I', 't', 'e', 'm', 's')
    CEREAL_RAPIDJSON_STRING_(MinLength, 'm', 'i', 'n', 'L', 'e', 'n', 'g', 't', 'h')
    CEREAL_RAPIDJSON_STRING_(MaxLength, 'm', 'a', 'x', 'L', 'e', 'n', 'g', 't', 'h')
    CEREAL_RAPIDJSON_STRING_(Pattern, 'p', 'a', 't', 't', 'e', 'r', 'n')
    CEREAL_RAPIDJSON_STRING_(Minimum, 'm', 'i', 'n', 'i', 'm', 'u', 'm')
    CEREAL_RAPIDJSON_STRING_(Maximum, 'm', 'a', 'x', 'i', 'm', 'u', 'm')
    CEREAL_RAPIDJSON_STRING_(ExclusiveMinimum, 'e', 'x', 'c', 'l', 'u', 's', 'i', 'v', 'e', 'M', 'i', 'n', 'i', 'm', 'u', 'm')
    CEREAL_RAPIDJSON_STRING_(ExclusiveMaximum, 'e', 'x', 'c', 'l', 'u', 's', 'i', 'v', 'e', 'M', 'a', 'x', 'i', 'm', 'u', 'm')
    CEREAL_RAPIDJSON_STRING_(MultipleOf, 'm', 'u', 'l', 't', 'i', 'p', 'l', 'e', 'O', 'f')
    CEREAL_RAPIDJSON_STRING_(DefaultValue, 'd', 'e', 'f', 'a', 'u', 'l', 't')

#undef CEREAL_RAPIDJSON_STRING_

private:
    enum SchemaValueType {
        kNullSchemaType,
        kBooleanSchemaType,
        kObjectSchemaType,
        kArraySchemaType,
        kStringSchemaType,
        kNumberSchemaType,
        kIntegerSchemaType,
        kTotalSchemaType
    };

#if CEREAL_RAPIDJSON_SCHEMA_USE_INTERNALREGEX
        typedef internal::GenericRegex<EncodingType, AllocatorType> RegexType;
#elif CEREAL_RAPIDJSON_SCHEMA_USE_STDREGEX
        typedef std::basic_regex<Ch> RegexType;
#else
        typedef char RegexType;
#endif

    struct SchemaArray {
        SchemaArray() : schemas(), count() {}
        ~SchemaArray() { AllocatorType::Free(schemas); }
        const SchemaType** schemas;
        SizeType begin; // begin index of context.validators
        SizeType count;
    };

    template <typename V1, typename V2>
    void AddUniqueElement(V1& a, const V2& v) {
        for (typename V1::ConstValueIterator itr = a.Begin(); itr != a.End(); ++itr)
            if (*itr == v)
                return;
        V1 c(v, *allocator_);
        a.PushBack(c, *allocator_);
    }

    static const ValueType* GetMember(const ValueType& value, const ValueType& name) {
        typename ValueType::ConstMemberIterator itr = value.FindMember(name);
        return itr != value.MemberEnd() ? &(itr->value) : 0;
    }

    static void AssignIfExist(bool& out, const ValueType& value, const ValueType& name) {
        if (const ValueType* v = GetMember(value, name))
            if (v->IsBool())
                out = v->GetBool();
    }

    static void AssignIfExist(SizeType& out, const ValueType& value, const ValueType& name) {
        if (const ValueType* v = GetMember(value, name))
            if (v->IsUint64() && v->GetUint64() <= SizeType(~0))
                out = static_cast<SizeType>(v->GetUint64());
    }

    void AssignIfExist(SchemaArray& out, SchemaDocumentType& schemaDocument, const PointerType& p, const ValueType& value, const ValueType& name, const ValueType& document) {
        if (const ValueType* v = GetMember(value, name)) {
            if (v->IsArray() && v->Size() > 0) {
                PointerType q = p.Append(name, allocator_);
                out.count = v->Size();
                out.schemas = static_cast<const Schema**>(allocator_->Malloc(out.count * sizeof(const Schema*)));
                memset(out.schemas, 0, sizeof(Schema*)* out.count);
                for (SizeType i = 0; i < out.count; i++)
                    schemaDocument.CreateSchema(&out.schemas[i], q.Append(i, allocator_), (*v)[i], document);
                out.begin = validatorCount_;
                validatorCount_ += out.count;
            }
        }
    }

#if CEREAL_RAPIDJSON_SCHEMA_USE_INTERNALREGEX
    template <typename ValueType>
    RegexType* CreatePattern(const ValueType& value) {
        if (value.IsString()) {
            RegexType* r = new (allocator_->Malloc(sizeof(RegexType))) RegexType(value.GetString(), allocator_);
            if (!r->IsValid()) {
                r->~RegexType();
                AllocatorType::Free(r);
                r = 0;
            }
            return r;
        }
        return 0;
    }

    static bool IsPatternMatch(const RegexType* pattern, const Ch *str, SizeType) {
        GenericRegexSearch<RegexType> rs(*pattern);
        return rs.Search(str);
    }
#elif CEREAL_RAPIDJSON_SCHEMA_USE_STDREGEX
    template <typename ValueType>
    RegexType* CreatePattern(const ValueType& value) {
        if (value.IsString()) {
            RegexType *r = static_cast<RegexType*>(allocator_->Malloc(sizeof(RegexType)));
            try {
                return new (r) RegexType(value.GetString(), std::size_t(value.GetStringLength()), std::regex_constants::ECMAScript);
            }
            catch (const std::regex_error&) {
                AllocatorType::Free(r);
            }
        }
        return 0;
    }

    static bool IsPatternMatch(const RegexType* pattern, const Ch *str, SizeType length) {
        std::match_results<const Ch*> r;
        return std::regex_search(str, str + length, r, *pattern);
    }
#else
    template <typename ValueType>
    RegexType* CreatePattern(const ValueType&) { return 0; }

    static bool IsPatternMatch(const RegexType*, const Ch *, SizeType) { return true; }
#endif // CEREAL_RAPIDJSON_SCHEMA_USE_STDREGEX

    void AddType(const ValueType& type) {
        if      (type == GetNullString()   ) type_ |= 1 << kNullSchemaType;
        else if (type == GetBooleanString()) type_ |= 1 << kBooleanSchemaType;
        else if (type == GetObjectString() ) type_ |= 1 << kObjectSchemaType;
        else if (type == GetArrayString()  ) type_ |= 1 << kArraySchemaType;
        else if (type == GetStringString() ) type_ |= 1 << kStringSchemaType;
        else if (type == GetIntegerString()) type_ |= 1 << kIntegerSchemaType;
        else if (type == GetNumberString() ) type_ |= (1 << kNumberSchemaType) | (1 << kIntegerSchemaType);
    }

    bool CreateParallelValidator(Context& context) const {
        if (enum_ || context.arrayUniqueness)
            context.hasher = context.factory.CreateHasher();

        if (validatorCount_) {
            CEREAL_RAPIDJSON_ASSERT(context.validators == 0);
            context.validators = static_cast<ISchemaValidator**>(context.factory.MallocState(sizeof(ISchemaValidator*) * validatorCount_));
            context.validatorCount = validatorCount_;

            if (allOf_.schemas)
                CreateSchemaValidators(context, allOf_);

            if (anyOf_.schemas)
                CreateSchemaValidators(context, anyOf_);
            
            if (oneOf_.schemas)
                CreateSchemaValidators(context, oneOf_);
            
            if (not_)
                context.validators[notValidatorIndex_] = context.factory.CreateSchemaValidator(*not_);
            
            if (hasSchemaDependencies_) {
                for (SizeType i = 0; i < propertyCount_; i++)
                    if (properties_[i].dependenciesSchema)
                        context.validators[properties_[i].dependenciesValidatorIndex] = context.factory.CreateSchemaValidator(*properties_[i].dependenciesSchema);
            }
        }

        return true;
    }

    void CreateSchemaValidators(Context& context, const SchemaArray& schemas) const {
        for (SizeType i = 0; i < schemas.count; i++)
            context.validators[schemas.begin + i] = context.factory.CreateSchemaValidator(*schemas.schemas[i]);
    }

    // O(n)
    bool FindPropertyIndex(const ValueType& name, SizeType* outIndex) const {
        SizeType len = name.GetStringLength();
        const Ch* str = name.GetString();
        for (SizeType index = 0; index < propertyCount_; index++)
            if (properties_[index].name.GetStringLength() == len && 
                (std::memcmp(properties_[index].name.GetString(), str, sizeof(Ch) * len) == 0))
            {
                *outIndex = index;
                return true;
            }
        return false;
    }

    bool CheckInt(Context& context, int64_t i) const {
        if (!(type_ & ((1 << kIntegerSchemaType) | (1 << kNumberSchemaType)))) {
            DisallowedType(context, GetIntegerString());
            CEREAL_RAPIDJSON_INVALID_KEYWORD_RETURN(GetTypeString());
        }

        if (!minimum_.IsNull()) {
            if (minimum_.IsInt64()) {
                if (exclusiveMinimum_ ? i <= minimum_.GetInt64() : i < minimum_.GetInt64()) {
                    context.error_handler.BelowMinimum(i, minimum_, exclusiveMinimum_);
                    CEREAL_RAPIDJSON_INVALID_KEYWORD_RETURN(GetMinimumString());
                }
            }
            else if (minimum_.IsUint64()) {
                context.error_handler.BelowMinimum(i, minimum_, exclusiveMinimum_);
                CEREAL_RAPIDJSON_INVALID_KEYWORD_RETURN(GetMinimumString()); // i <= max(int64_t) < minimum.GetUint64()
            }
            else if (!CheckDoubleMinimum(context, static_cast<double>(i)))
                return false;
        }

        if (!maximum_.IsNull()) {
            if (maximum_.IsInt64()) {
                if (exclusiveMaximum_ ? i >= maximum_.GetInt64() : i > maximum_.GetInt64()) {
                    context.error_handler.AboveMaximum(i, maximum_, exclusiveMaximum_);
                    CEREAL_RAPIDJSON_INVALID_KEYWORD_RETURN(GetMaximumString());
                }
            }
            else if (maximum_.IsUint64()) { }
                /* do nothing */ // i <= max(int64_t) < maximum_.GetUint64()
            else if (!CheckDoubleMaximum(context, static_cast<double>(i)))
                return false;
        }

        if (!multipleOf_.IsNull()) {
            if (multipleOf_.IsUint64()) {
                if (static_cast<uint64_t>(i >= 0 ? i : -i) % multipleOf_.GetUint64() != 0) {
                    context.error_handler.NotMultipleOf(i, multipleOf_);
                    CEREAL_RAPIDJSON_INVALID_KEYWORD_RETURN(GetMultipleOfString());
                }
            }
            else if (!CheckDoubleMultipleOf(context, static_cast<double>(i)))
                return false;
        }

        return true;
    }

    bool CheckUint(Context& context, uint64_t i) const {
        if (!(type_ & ((1 << kIntegerSchemaType) | (1 << kNumberSchemaType)))) {
            DisallowedType(context, GetIntegerString());
            CEREAL_RAPIDJSON_INVALID_KEYWORD_RETURN(GetTypeString());
        }

        if (!minimum_.IsNull()) {
            if (minimum_.IsUint64()) {
                if (exclusiveMinimum_ ? i <= minimum_.GetUint64() : i < minimum_.GetUint64()) {
                    context.error_handler.BelowMinimum(i, minimum_, exclusiveMinimum_);
                    CEREAL_RAPIDJSON_INVALID_KEYWORD_RETURN(GetMinimumString());
                }
            }
            else if (minimum_.IsInt64())
                /* do nothing */; // i >= 0 > minimum.Getint64()
            else if (!CheckDoubleMinimum(context, static_cast<double>(i)))
                return false;
        }

        if (!maximum_.IsNull()) {
            if (maximum_.IsUint64()) {
                if (exclusiveMaximum_ ? i >= maximum_.GetUint64() : i > maximum_.GetUint64()) {
                    context.error_handler.AboveMaximum(i, maximum_, exclusiveMaximum_);
                    CEREAL_RAPIDJSON_INVALID_KEYWORD_RETURN(GetMaximumString());
                }
            }
            else if (maximum_.IsInt64()) {
                context.error_handler.AboveMaximum(i, maximum_, exclusiveMaximum_);
                CEREAL_RAPIDJSON_INVALID_KEYWORD_RETURN(GetMaximumString()); // i >= 0 > maximum_
            }
            else if (!CheckDoubleMaximum(context, static_cast<double>(i)))
                return false;
        }

        if (!multipleOf_.IsNull()) {
            if (multipleOf_.IsUint64()) {
                if (i % multipleOf_.GetUint64() != 0) {
                    context.error_handler.NotMultipleOf(i, multipleOf_);
                    CEREAL_RAPIDJSON_INVALID_KEYWORD_RETURN(GetMultipleOfString());
                }
            }
            else if (!CheckDoubleMultipleOf(context, static_cast<double>(i)))
                return false;
        }

        return true;
    }

    bool CheckDoubleMinimum(Context& context, double d) const {
        if (exclusiveMinimum_ ? d <= minimum_.GetDouble() : d < minimum_.GetDouble()) {
            context.error_handler.BelowMinimum(d, minimum_, exclusiveMinimum_);
            CEREAL_RAPIDJSON_INVALID_KEYWORD_RETURN(GetMinimumString());
        }
        return true;
    }

    bool CheckDoubleMaximum(Context& context, double d) const {
        if (exclusiveMaximum_ ? d >= maximum_.GetDouble() : d > maximum_.GetDouble()) {
            context.error_handler.AboveMaximum(d, maximum_, exclusiveMaximum_);
            CEREAL_RAPIDJSON_INVALID_KEYWORD_RETURN(GetMaximumString());
        }
        return true;
    }

    bool CheckDoubleMultipleOf(Context& context, double d) const {
        double a = std::abs(d), b = std::abs(multipleOf_.GetDouble());
        double q = std::floor(a / b);
        double r = a - q * b;
        if (r > 0.0) {
            context.error_handler.NotMultipleOf(d, multipleOf_);
            CEREAL_RAPIDJSON_INVALID_KEYWORD_RETURN(GetMultipleOfString());
        }
        return true;
    }

    void DisallowedType(Context& context, const ValueType& actualType) const {
        ErrorHandler& eh = context.error_handler;
        eh.StartDisallowedType();

        if (type_ & (1 << kNullSchemaType)) eh.AddExpectedType(GetNullString());
        if (type_ & (1 << kBooleanSchemaType)) eh.AddExpectedType(GetBooleanString());
        if (type_ & (1 << kObjectSchemaType)) eh.AddExpectedType(GetObjectString());
        if (type_ & (1 << kArraySchemaType)) eh.AddExpectedType(GetArrayString());
        if (type_ & (1 << kStringSchemaType)) eh.AddExpectedType(GetStringString());

        if (type_ & (1 << kNumberSchemaType)) eh.AddExpectedType(GetNumberString());
        else if (type_ & (1 << kIntegerSchemaType)) eh.AddExpectedType(GetIntegerString());

        eh.EndDisallowedType(actualType);
    }

    struct Property {
        Property() : schema(), dependenciesSchema(), dependenciesValidatorIndex(), dependencies(), required(false) {}
        ~Property() { AllocatorType::Free(dependencies); }
        SValue name;
        const SchemaType* schema;
        const SchemaType* dependenciesSchema;
        SizeType dependenciesValidatorIndex;
        bool* dependencies;
        bool required;
    };

    struct PatternProperty {
        PatternProperty() : schema(), pattern() {}
        ~PatternProperty() { 
            if (pattern) {
                pattern->~RegexType();
                AllocatorType::Free(pattern);
            }
        }
        const SchemaType* schema;
        RegexType* pattern;
    };

    AllocatorType* allocator_;
    SValue uri_;
    PointerType pointer_;
    const SchemaType* typeless_;
    uint64_t* enum_;
    SizeType enumCount_;
    SchemaArray allOf_;
    SchemaArray anyOf_;
    SchemaArray oneOf_;
    const SchemaType* not_;
    unsigned type_; // bitmask of kSchemaType
    SizeType validatorCount_;
    SizeType notValidatorIndex_;

    Property* properties_;
    const SchemaType* additionalPropertiesSchema_;
    PatternProperty* patternProperties_;
    SizeType patternPropertyCount_;
    SizeType propertyCount_;
    SizeType minProperties_;
    SizeType maxProperties_;
    bool additionalProperties_;
    bool hasDependencies_;
    bool hasRequired_;
    bool hasSchemaDependencies_;

    const SchemaType* additionalItemsSchema_;
    const SchemaType* itemsList_;
    const SchemaType** itemsTuple_;
    SizeType itemsTupleCount_;
    SizeType minItems_;
    SizeType maxItems_;
    bool additionalItems_;
    bool uniqueItems_;

    RegexType* pattern_;
    SizeType minLength_;
    SizeType maxLength_;

    SValue minimum_;
    SValue maximum_;
    SValue multipleOf_;
    bool exclusiveMinimum_;
    bool exclusiveMaximum_;
    
    SizeType defaultValueLength_;
};

template<typename Stack, typename Ch>
struct TokenHelper {
    CEREAL_RAPIDJSON_FORCEINLINE static void AppendIndexToken(Stack& documentStack, SizeType index) {
        *documentStack.template Push<Ch>() = '/';
        char buffer[21];
        size_t length = static_cast<size_t>((sizeof(SizeType) == 4 ? u32toa(index, buffer) : u64toa(index, buffer)) - buffer);
        for (size_t i = 0; i < length; i++)
            *documentStack.template Push<Ch>() = static_cast<Ch>(buffer[i]);
    }
};

// Partial specialized version for char to prevent buffer copying.
template <typename Stack>
struct TokenHelper<Stack, char> {
    CEREAL_RAPIDJSON_FORCEINLINE static void AppendIndexToken(Stack& documentStack, SizeType index) {
        if (sizeof(SizeType) == 4) {
            char *buffer = documentStack.template Push<char>(1 + 10); // '/' + uint
            *buffer++ = '/';
            const char* end = internal::u32toa(index, buffer);
             documentStack.template Pop<char>(static_cast<size_t>(10 - (end - buffer)));
        }
        else {
            char *buffer = documentStack.template Push<char>(1 + 20); // '/' + uint64
            *buffer++ = '/';
            const char* end = internal::u64toa(index, buffer);
            documentStack.template Pop<char>(static_cast<size_t>(20 - (end - buffer)));
        }
    }
};

} // namespace internal

///////////////////////////////////////////////////////////////////////////////
// IGenericRemoteSchemaDocumentProvider

template <typename SchemaDocumentType>
class IGenericRemoteSchemaDocumentProvider {
public:
    typedef typename SchemaDocumentType::Ch Ch;

    virtual ~IGenericRemoteSchemaDocumentProvider() {}
    virtual const SchemaDocumentType* GetRemoteDocument(const Ch* uri, SizeType length) = 0;
};

///////////////////////////////////////////////////////////////////////////////
// GenericSchemaDocument

//! JSON schema document.
/*!
    A JSON schema document is a compiled version of a JSON schema.
    It is basically a tree of internal::Schema.

    \note This is an immutable class (i.e. its instance cannot be modified after construction).
    \tparam ValueT Type of JSON value (e.g. \c Value ), which also determine the encoding.
    \tparam Allocator Allocator type for allocating memory of this document.
*/
template <typename ValueT, typename Allocator = CrtAllocator>
class GenericSchemaDocument {
public:
    typedef ValueT ValueType;
    typedef IGenericRemoteSchemaDocumentProvider<GenericSchemaDocument> IRemoteSchemaDocumentProviderType;
    typedef Allocator AllocatorType;
    typedef typename ValueType::EncodingType EncodingType;
    typedef typename EncodingType::Ch Ch;
    typedef internal::Schema<GenericSchemaDocument> SchemaType;
    typedef GenericPointer<ValueType, Allocator> PointerType;
    typedef GenericValue<EncodingType, Allocator> URIType;
    friend class internal::Schema<GenericSchemaDocument>;
    template <typename, typename, typename>
    friend class GenericSchemaValidator;

    //! Constructor.
    /*!
        Compile a JSON document into schema document.

        \param document A JSON document as source.
        \param uri The base URI of this schema document for purposes of violation reporting.
        \param uriLength Length of \c name, in code points.
        \param remoteProvider An optional remote schema document provider for resolving remote reference. Can be null.
        \param allocator An optional allocator instance for allocating memory. Can be null.
    */
    explicit GenericSchemaDocument(const ValueType& document, const Ch* uri = 0, SizeType uriLength = 0,
        IRemoteSchemaDocumentProviderType* remoteProvider = 0, Allocator* allocator = 0) :
        remoteProvider_(remoteProvider),
        allocator_(allocator),
        ownAllocator_(),
        root_(),
        typeless_(),
        schemaMap_(allocator, kInitialSchemaMapSize),
        schemaRef_(allocator, kInitialSchemaRefSize)
    {
        if (!allocator_)
            ownAllocator_ = allocator_ = CEREAL_RAPIDJSON_NEW(Allocator)();

        Ch noUri[1] = {0};
        uri_.SetString(uri ? uri : noUri, uriLength, *allocator_);

        typeless_ = static_cast<SchemaType*>(allocator_->Malloc(sizeof(SchemaType)));
        new (typeless_) SchemaType(this, PointerType(), ValueType(kObjectType).Move(), ValueType(kObjectType).Move(), allocator_);

        // Generate root schema, it will call CreateSchema() to create sub-schemas,
        // And call AddRefSchema() if there are $ref.
        CreateSchemaRecursive(&root_, PointerType(), document, document);

        // Resolve $ref
        while (!schemaRef_.Empty()) {
            SchemaRefEntry* refEntry = schemaRef_.template Pop<SchemaRefEntry>(1);
            if (const SchemaType* s = GetSchema(refEntry->target)) {
                if (refEntry->schema)
                    *refEntry->schema = s;

                // Create entry in map if not exist
                if (!GetSchema(refEntry->source)) {
                    new (schemaMap_.template Push<SchemaEntry>()) SchemaEntry(refEntry->source, const_cast<SchemaType*>(s), false, allocator_);
                }
            }
            else if (refEntry->schema)
                *refEntry->schema = typeless_;

            refEntry->~SchemaRefEntry();
        }

        CEREAL_RAPIDJSON_ASSERT(root_ != 0);

        schemaRef_.ShrinkToFit(); // Deallocate all memory for ref
    }

#if CEREAL_RAPIDJSON_HAS_CXX11_RVALUE_REFS
    //! Move constructor in C++11
    GenericSchemaDocument(GenericSchemaDocument&& rhs) CEREAL_RAPIDJSON_NOEXCEPT :
        remoteProvider_(rhs.remoteProvider_),
        allocator_(rhs.allocator_),
        ownAllocator_(rhs.ownAllocator_),
        root_(rhs.root_),
        typeless_(rhs.typeless_),
        schemaMap_(std::move(rhs.schemaMap_)),
        schemaRef_(std::move(rhs.schemaRef_)),
        uri_(std::move(rhs.uri_))
    {
        rhs.remoteProvider_ = 0;
        rhs.allocator_ = 0;
        rhs.ownAllocator_ = 0;
        rhs.typeless_ = 0;
    }
#endif

    //! Destructor
    ~GenericSchemaDocument() {
        while (!schemaMap_.Empty())
            schemaMap_.template Pop<SchemaEntry>(1)->~SchemaEntry();

        if (typeless_) {
            typeless_->~SchemaType();
            Allocator::Free(typeless_);
        }

        CEREAL_RAPIDJSON_DELETE(ownAllocator_);
    }

    const URIType& GetURI() const { return uri_; }

    //! Get the root schema.
    const SchemaType& GetRoot() const { return *root_; }

private:
    //! Prohibit copying
    GenericSchemaDocument(const GenericSchemaDocument&);
    //! Prohibit assignment
    GenericSchemaDocument& operator=(const GenericSchemaDocument&);

    struct SchemaRefEntry {
        SchemaRefEntry(const PointerType& s, const PointerType& t, const SchemaType** outSchema, Allocator *allocator) : source(s, allocator), target(t, allocator), schema(outSchema) {}
        PointerType source;
        PointerType target;
        const SchemaType** schema;
    };

    struct SchemaEntry {
        SchemaEntry(const PointerType& p, SchemaType* s, bool o, Allocator* allocator) : pointer(p, allocator), schema(s), owned(o) {}
        ~SchemaEntry() {
            if (owned) {
                schema->~SchemaType();
                Allocator::Free(schema);
            }
        }
        PointerType pointer;
        SchemaType* schema;
        bool owned;
    };

    void CreateSchemaRecursive(const SchemaType** schema, const PointerType& pointer, const ValueType& v, const ValueType& document) {
        if (schema)
            *schema = typeless_;

        if (v.GetType() == kObjectType) {
            const SchemaType* s = GetSchema(pointer);
            if (!s)
                CreateSchema(schema, pointer, v, document);

            for (typename ValueType::ConstMemberIterator itr = v.MemberBegin(); itr != v.MemberEnd(); ++itr)
                CreateSchemaRecursive(0, pointer.Append(itr->name, allocator_), itr->value, document);
        }
        else if (v.GetType() == kArrayType)
            for (SizeType i = 0; i < v.Size(); i++)
                CreateSchemaRecursive(0, pointer.Append(i, allocator_), v[i], document);
    }

    void CreateSchema(const SchemaType** schema, const PointerType& pointer, const ValueType& v, const ValueType& document) {
        CEREAL_RAPIDJSON_ASSERT(pointer.IsValid());
        if (v.IsObject()) {
            if (!HandleRefSchema(pointer, schema, v, document)) {
                SchemaType* s = new (allocator_->Malloc(sizeof(SchemaType))) SchemaType(this, pointer, v, document, allocator_);
                new (schemaMap_.template Push<SchemaEntry>()) SchemaEntry(pointer, s, true, allocator_);
                if (schema)
                    *schema = s;
            }
        }
    }

    bool HandleRefSchema(const PointerType& source, const SchemaType** schema, const ValueType& v, const ValueType& document) {
        static const Ch kRefString[] = { '$', 'r', 'e', 'f', '\0' };
        static const ValueType kRefValue(kRefString, 4);

        typename ValueType::ConstMemberIterator itr = v.FindMember(kRefValue);
        if (itr == v.MemberEnd())
            return false;

        if (itr->value.IsString()) {
            SizeType len = itr->value.GetStringLength();
            if (len > 0) {
                const Ch* s = itr->value.GetString();
                SizeType i = 0;
                while (i < len && s[i] != '#') // Find the first #
                    i++;

                if (i > 0) { // Remote reference, resolve immediately
                    if (remoteProvider_) {
                        if (const GenericSchemaDocument* remoteDocument = remoteProvider_->GetRemoteDocument(s, i)) {
                            PointerType pointer(&s[i], len - i, allocator_);
                            if (pointer.IsValid()) {
                                if (const SchemaType* sc = remoteDocument->GetSchema(pointer)) {
                                    if (schema)
                                        *schema = sc;
                                    new (schemaMap_.template Push<SchemaEntry>()) SchemaEntry(source, const_cast<SchemaType*>(sc), false, allocator_);
                                    return true;
                                }
                            }
                        }
                    }
                }
                else if (s[i] == '#') { // Local reference, defer resolution
                    PointerType pointer(&s[i], len - i, allocator_);
                    if (pointer.IsValid()) {
                        if (const ValueType* nv = pointer.Get(document))
                            if (HandleRefSchema(source, schema, *nv, document))
                                return true;

                        new (schemaRef_.template Push<SchemaRefEntry>()) SchemaRefEntry(source, pointer, schema, allocator_);
                        return true;
                    }
                }
            }
        }
        return false;
    }

    const SchemaType* GetSchema(const PointerType& pointer) const {
        for (const SchemaEntry* target = schemaMap_.template Bottom<SchemaEntry>(); target != schemaMap_.template End<SchemaEntry>(); ++target)
            if (pointer == target->pointer)
                return target->schema;
        return 0;
    }

    PointerType GetPointer(const SchemaType* schema) const {
        for (const SchemaEntry* target = schemaMap_.template Bottom<SchemaEntry>(); target != schemaMap_.template End<SchemaEntry>(); ++target)
            if (schema == target->schema)
                return target->pointer;
        return PointerType();
    }

    const SchemaType* GetTypeless() const { return typeless_; }

    static const size_t kInitialSchemaMapSize = 64;
    static const size_t kInitialSchemaRefSize = 64;

    IRemoteSchemaDocumentProviderType* remoteProvider_;
    Allocator *allocator_;
    Allocator *ownAllocator_;
    const SchemaType* root_;                //!< Root schema.
    SchemaType* typeless_;
    internal::Stack<Allocator> schemaMap_;  // Stores created Pointer -> Schemas
    internal::Stack<Allocator> schemaRef_;  // Stores Pointer from $ref and schema which holds the $ref
    URIType uri_;
};

//! GenericSchemaDocument using Value type.
typedef GenericSchemaDocument<Value> SchemaDocument;
//! IGenericRemoteSchemaDocumentProvider using SchemaDocument.
typedef IGenericRemoteSchemaDocumentProvider<SchemaDocument> IRemoteSchemaDocumentProvider;

///////////////////////////////////////////////////////////////////////////////
// GenericSchemaValidator

//! JSON Schema Validator.
/*!
    A SAX style JSON schema validator.
    It uses a \c GenericSchemaDocument to validate SAX events.
    It delegates the incoming SAX events to an output handler.
    The default output handler does nothing.
    It can be reused multiple times by calling \c Reset().

    \tparam SchemaDocumentType Type of schema document.
    \tparam OutputHandler Type of output handler. Default handler does nothing.
    \tparam StateAllocator Allocator for storing the internal validation states.
*/
template <
    typename SchemaDocumentType,
    typename OutputHandler = BaseReaderHandler<typename SchemaDocumentType::SchemaType::EncodingType>,
    typename StateAllocator = CrtAllocator>
class GenericSchemaValidator :
    public internal::ISchemaStateFactory<typename SchemaDocumentType::SchemaType>, 
    public internal::ISchemaValidator,
    public internal::IValidationErrorHandler<typename SchemaDocumentType::SchemaType>
{
public:
    typedef typename SchemaDocumentType::SchemaType SchemaType;
    typedef typename SchemaDocumentType::PointerType PointerType;
    typedef typename SchemaType::EncodingType EncodingType;
    typedef typename SchemaType::SValue SValue;
    typedef typename EncodingType::Ch Ch;
    typedef GenericStringRef<Ch> StringRefType;
    typedef GenericValue<EncodingType, StateAllocator> ValueType;

    //! Constructor without output handler.
    /*!
        \param schemaDocument The schema document to conform to.
        \param allocator Optional allocator for storing internal validation states.
        \param schemaStackCapacity Optional initial capacity of schema path stack.
        \param documentStackCapacity Optional initial capacity of document path stack.
    */
    GenericSchemaValidator(
        const SchemaDocumentType& schemaDocument,
        StateAllocator* allocator = 0, 
        size_t schemaStackCapacity = kDefaultSchemaStackCapacity,
        size_t documentStackCapacity = kDefaultDocumentStackCapacity)
        :
        schemaDocument_(&schemaDocument),
        root_(schemaDocument.GetRoot()),
        stateAllocator_(allocator),
        ownStateAllocator_(0),
        schemaStack_(allocator, schemaStackCapacity),
        documentStack_(allocator, documentStackCapacity),
        outputHandler_(0),
        error_(kObjectType),
        currentError_(),
        missingDependents_(),
        valid_(true)
#if CEREAL_RAPIDJSON_SCHEMA_VERBOSE
        , depth_(0)
#endif
    {
    }

    //! Constructor with output handler.
    /*!
        \param schemaDocument The schema document to conform to.
        \param allocator Optional allocator for storing internal validation states.
        \param schemaStackCapacity Optional initial capacity of schema path stack.
        \param documentStackCapacity Optional initial capacity of document path stack.
    */
    GenericSchemaValidator(
        const SchemaDocumentType& schemaDocument,
        OutputHandler& outputHandler,
        StateAllocator* allocator = 0, 
        size_t schemaStackCapacity = kDefaultSchemaStackCapacity,
        size_t documentStackCapacity = kDefaultDocumentStackCapacity)
        :
        schemaDocument_(&schemaDocument),
        root_(schemaDocument.GetRoot()),
        stateAllocator_(allocator),
        ownStateAllocator_(0),
        schemaStack_(allocator, schemaStackCapacity),
        documentStack_(allocator, documentStackCapacity),
        outputHandler_(&outputHandler),
        error_(kObjectType),
        currentError_(),
        missingDependents_(),
        valid_(true)
#if CEREAL_RAPIDJSON_SCHEMA_VERBOSE
        , depth_(0)
#endif
    {
    }

    //! Destructor.
    ~GenericSchemaValidator() {
        Reset();
        CEREAL_RAPIDJSON_DELETE(ownStateAllocator_);
    }

    //! Reset the internal states.
    void Reset() {
        while (!schemaStack_.Empty())
            PopSchema();
        documentStack_.Clear();
        error_.SetObject();
        currentError_.SetNull();
        missingDependents_.SetNull();
        valid_ = true;
    }

    //! Checks whether the current state is valid.
    // Implementation of ISchemaValidator
    virtual bool IsValid() const { return valid_; }

    //! Gets the error object.
    ValueType& GetError() { return error_; }
    const ValueType& GetError() const { return error_; }

    //! Gets the JSON pointer pointed to the invalid schema.
    PointerType GetInvalidSchemaPointer() const {
        return schemaStack_.Empty() ? PointerType() : CurrentSchema().GetPointer();
    }

    //! Gets the keyword of invalid schema.
    const Ch* GetInvalidSchemaKeyword() const {
        return schemaStack_.Empty() ? 0 : CurrentContext().invalidKeyword;
    }

    //! Gets the JSON pointer pointed to the invalid value.
    PointerType GetInvalidDocumentPointer() const {
        if (documentStack_.Empty()) {
            return PointerType();
        }
        else {
            return PointerType(documentStack_.template Bottom<Ch>(), documentStack_.GetSize() / sizeof(Ch));
        }
    }

    void NotMultipleOf(int64_t actual, const SValue& expected) {
        AddNumberError(SchemaType::GetMultipleOfString(), ValueType(actual).Move(), expected);
    }
    void NotMultipleOf(uint64_t actual, const SValue& expected) {
        AddNumberError(SchemaType::GetMultipleOfString(), ValueType(actual).Move(), expected);
    }
    void NotMultipleOf(double actual, const SValue& expected) {
        AddNumberError(SchemaType::GetMultipleOfString(), ValueType(actual).Move(), expected);
    }
    void AboveMaximum(int64_t actual, const SValue& expected, bool exclusive) {
        AddNumberError(SchemaType::GetMaximumString(), ValueType(actual).Move(), expected,
            exclusive ? &SchemaType::GetExclusiveMaximumString : 0);
    }
    void AboveMaximum(uint64_t actual, const SValue& expected, bool exclusive) {
        AddNumberError(SchemaType::GetMaximumString(), ValueType(actual).Move(), expected,
            exclusive ? &SchemaType::GetExclusiveMaximumString : 0);
    }
    void AboveMaximum(double actual, const SValue& expected, bool exclusive) {
        AddNumberError(SchemaType::GetMaximumString(), ValueType(actual).Move(), expected,
            exclusive ? &SchemaType::GetExclusiveMaximumString : 0);
    }
    void BelowMinimum(int64_t actual, const SValue& expected, bool exclusive) {
        AddNumberError(SchemaType::GetMinimumString(), ValueType(actual).Move(), expected,
            exclusive ? &SchemaType::GetExclusiveMinimumString : 0);
    }
    void BelowMinimum(uint64_t actual, const SValue& expected, bool exclusive) {
        AddNumberError(SchemaType::GetMinimumString(), ValueType(actual).Move(), expected,
            exclusive ? &SchemaType::GetExclusiveMinimumString : 0);
    }
    void BelowMinimum(double actual, const SValue& expected, bool exclusive) {
        AddNumberError(SchemaType::GetMinimumString(), ValueType(actual).Move(), expected,
            exclusive ? &SchemaType::GetExclusiveMinimumString : 0);
    }

    void TooLong(const Ch* str, SizeType length, SizeType expected) {
        AddNumberError(SchemaType::GetMaxLengthString(),
            ValueType(str, length, GetStateAllocator()).Move(), SValue(expected).Move());
    }
    void TooShort(const Ch* str, SizeType length, SizeType expected) {
        AddNumberError(SchemaType::GetMinLengthString(),
            ValueType(str, length, GetStateAllocator()).Move(), SValue(expected).Move());
    }
    void DoesNotMatch(const Ch* str, SizeType length) {
        currentError_.SetObject();
        currentError_.AddMember(GetActualString(), ValueType(str, length, GetStateAllocator()).Move(), GetStateAllocator());
        AddCurrentError(SchemaType::GetPatternString());
    }

    void DisallowedItem(SizeType index) {
        currentError_.SetObject();
        currentError_.AddMember(GetDisallowedString(), ValueType(index).Move(), GetStateAllocator());
        AddCurrentError(SchemaType::GetAdditionalItemsString(), true);
    }
    void TooFewItems(SizeType actualCount, SizeType expectedCount) {
        AddNumberError(SchemaType::GetMinItemsString(),
            ValueType(actualCount).Move(), SValue(expectedCount).Move());
    }
    void TooManyItems(SizeType actualCount, SizeType expectedCount) {
        AddNumberError(SchemaType::GetMaxItemsString(),
            ValueType(actualCount).Move(), SValue(expectedCount).Move());
    }
    void DuplicateItems(SizeType index1, SizeType index2) {
        ValueType duplicates(kArrayType);
        duplicates.PushBack(index1, GetStateAllocator());
        duplicates.PushBack(index2, GetStateAllocator());
        currentError_.SetObject();
        currentError_.AddMember(GetDuplicatesString(), duplicates, GetStateAllocator());
        AddCurrentError(SchemaType::GetUniqueItemsString(), true);
    }

    void TooManyProperties(SizeType actualCount, SizeType expectedCount) {
        AddNumberError(SchemaType::GetMaxPropertiesString(),
            ValueType(actualCount).Move(), SValue(expectedCount).Move());
    }
    void TooFewProperties(SizeType actualCount, SizeType expectedCount) {
        AddNumberError(SchemaType::GetMinPropertiesString(),
            ValueType(actualCount).Move(), SValue(expectedCount).Move());
    }
    void StartMissingProperties() {
        currentError_.SetArray();
    }
    void AddMissingProperty(const SValue& name) {
        currentError_.PushBack(ValueType(name, GetStateAllocator()).Move(), GetStateAllocator());
    }
    bool EndMissingProperties() {
        if (currentError_.Empty())
            return false;
        ValueType error(kObjectType);
        error.AddMember(GetMissingString(), currentError_, GetStateAllocator());
        currentError_ = error;
        AddCurrentError(SchemaType::GetRequiredString());
        return true;
    }
    void PropertyViolations(ISchemaValidator** subvalidators, SizeType count) {
        for (SizeType i = 0; i < count; ++i)
            MergeError(static_cast<GenericSchemaValidator*>(subvalidators[i])->GetError());
    }
    void DisallowedProperty(const Ch* name, SizeType length) {
        currentError_.SetObject();
        currentError_.AddMember(GetDisallowedString(), ValueType(name, length, GetStateAllocator()).Move(), GetStateAllocator());
        AddCurrentError(SchemaType::GetAdditionalPropertiesString(), true);
    }

    void StartDependencyErrors() {
        currentError_.SetObject();
    }
    void StartMissingDependentProperties() {
        missingDependents_.SetArray();
    }
    void AddMissingDependentProperty(const SValue& targetName) {
        missingDependents_.PushBack(ValueType(targetName, GetStateAllocator()).Move(), GetStateAllocator());
    }
    void EndMissingDependentProperties(const SValue& sourceName) {
        if (!missingDependents_.Empty())
            currentError_.AddMember(ValueType(sourceName, GetStateAllocator()).Move(),
                missingDependents_, GetStateAllocator());
    }
    void AddDependencySchemaError(const SValue& sourceName, ISchemaValidator* subvalidator) {
        currentError_.AddMember(ValueType(sourceName, GetStateAllocator()).Move(),
            static_cast<GenericSchemaValidator*>(subvalidator)->GetError(), GetStateAllocator());
    }
    bool EndDependencyErrors() {
        if (currentError_.ObjectEmpty())
            return false;
        ValueType error(kObjectType);
        error.AddMember(GetErrorsString(), currentError_, GetStateAllocator());
        currentError_ = error;
        AddCurrentError(SchemaType::GetDependenciesString());
        return true;
    }

    void DisallowedValue() {
        currentError_.SetObject();
        AddCurrentError(SchemaType::GetEnumString());
    }
    void StartDisallowedType() {
        currentError_.SetArray();
    }
    void AddExpectedType(const typename SchemaType::ValueType& expectedType) {
        currentError_.PushBack(ValueType(expectedType, GetStateAllocator()).Move(), GetStateAllocator());
    }
    void EndDisallowedType(const typename SchemaType::ValueType& actualType) {
        ValueType error(kObjectType);
        error.AddMember(GetExpectedString(), currentError_, GetStateAllocator());
        error.AddMember(GetActualString(), ValueType(actualType, GetStateAllocator()).Move(), GetStateAllocator());
        currentError_ = error;
        AddCurrentError(SchemaType::GetTypeString());
    }
    void NotAllOf(ISchemaValidator** subvalidators, SizeType count) {
        for (SizeType i = 0; i < count; ++i) {
            MergeError(static_cast<GenericSchemaValidator*>(subvalidators[i])->GetError());
        }
    }
    void NoneOf(ISchemaValidator** subvalidators, SizeType count) {
        AddErrorArray(SchemaType::GetAnyOfString(), subvalidators, count);
    }
    void NotOneOf(ISchemaValidator** subvalidators, SizeType count) {
        AddErrorArray(SchemaType::GetOneOfString(), subvalidators, count);
    }
    void Disallowed() {
        currentError_.SetObject();
        AddCurrentError(SchemaType::GetNotString());
    }

#define CEREAL_RAPIDJSON_STRING_(name, ...) \
    static const StringRefType& Get##name##String() {\
        static const Ch s[] = { __VA_ARGS__, '\0' };\
        static const StringRefType v(s, static_cast<SizeType>(sizeof(s) / sizeof(Ch) - 1)); \
        return v;\
    }

    CEREAL_RAPIDJSON_STRING_(InstanceRef, 'i', 'n', 's', 't', 'a', 'n', 'c', 'e', 'R', 'e', 'f')
    CEREAL_RAPIDJSON_STRING_(SchemaRef, 's', 'c', 'h', 'e', 'm', 'a', 'R', 'e', 'f')
    CEREAL_RAPIDJSON_STRING_(Expected, 'e', 'x', 'p', 'e', 'c', 't', 'e', 'd')
    CEREAL_RAPIDJSON_STRING_(Actual, 'a', 'c', 't', 'u', 'a', 'l')
    CEREAL_RAPIDJSON_STRING_(Disallowed, 'd', 'i', 's', 'a', 'l', 'l', 'o', 'w', 'e', 'd')
    CEREAL_RAPIDJSON_STRING_(Missing, 'm', 'i', 's', 's', 'i', 'n', 'g')
    CEREAL_RAPIDJSON_STRING_(Errors, 'e', 'r', 'r', 'o', 'r', 's')
    CEREAL_RAPIDJSON_STRING_(Duplicates, 'd', 'u', 'p', 'l', 'i', 'c', 'a', 't', 'e', 's')

#undef CEREAL_RAPIDJSON_STRING_

#if CEREAL_RAPIDJSON_SCHEMA_VERBOSE
#define CEREAL_RAPIDJSON_SCHEMA_HANDLE_BEGIN_VERBOSE_() \
CEREAL_RAPIDJSON_MULTILINEMACRO_BEGIN\
    *documentStack_.template Push<Ch>() = '\0';\
    documentStack_.template Pop<Ch>(1);\
    internal::PrintInvalidDocument(documentStack_.template Bottom<Ch>());\
CEREAL_RAPIDJSON_MULTILINEMACRO_END
#else
#define CEREAL_RAPIDJSON_SCHEMA_HANDLE_BEGIN_VERBOSE_()
#endif

#define CEREAL_RAPIDJSON_SCHEMA_HANDLE_BEGIN_(method, arg1)\
    if (!valid_) return false; \
    if (!BeginValue() || !CurrentSchema().method arg1) {\
        CEREAL_RAPIDJSON_SCHEMA_HANDLE_BEGIN_VERBOSE_();\
        return valid_ = false;\
    }

#define CEREAL_RAPIDJSON_SCHEMA_HANDLE_PARALLEL_(method, arg2)\
    for (Context* context = schemaStack_.template Bottom<Context>(); context != schemaStack_.template End<Context>(); context++) {\
        if (context->hasher)\
            static_cast<HasherType*>(context->hasher)->method arg2;\
        if (context->validators)\
            for (SizeType i_ = 0; i_ < context->validatorCount; i_++)\
                static_cast<GenericSchemaValidator*>(context->validators[i_])->method arg2;\
        if (context->patternPropertiesValidators)\
            for (SizeType i_ = 0; i_ < context->patternPropertiesValidatorCount; i_++)\
                static_cast<GenericSchemaValidator*>(context->patternPropertiesValidators[i_])->method arg2;\
    }

#define CEREAL_RAPIDJSON_SCHEMA_HANDLE_END_(method, arg2)\
    return valid_ = EndValue() && (!outputHandler_ || outputHandler_->method arg2)

#define CEREAL_RAPIDJSON_SCHEMA_HANDLE_VALUE_(method, arg1, arg2) \
    CEREAL_RAPIDJSON_SCHEMA_HANDLE_BEGIN_   (method, arg1);\
    CEREAL_RAPIDJSON_SCHEMA_HANDLE_PARALLEL_(method, arg2);\
    CEREAL_RAPIDJSON_SCHEMA_HANDLE_END_     (method, arg2)

    bool Null()             { CEREAL_RAPIDJSON_SCHEMA_HANDLE_VALUE_(Null,   (CurrentContext()), ( )); }
    bool Bool(bool b)       { CEREAL_RAPIDJSON_SCHEMA_HANDLE_VALUE_(Bool,   (CurrentContext(), b), (b)); }
    bool Int(int i)         { CEREAL_RAPIDJSON_SCHEMA_HANDLE_VALUE_(Int,    (CurrentContext(), i), (i)); }
    bool Uint(unsigned u)   { CEREAL_RAPIDJSON_SCHEMA_HANDLE_VALUE_(Uint,   (CurrentContext(), u), (u)); }
    bool Int64(int64_t i)   { CEREAL_RAPIDJSON_SCHEMA_HANDLE_VALUE_(Int64,  (CurrentContext(), i), (i)); }
    bool Uint64(uint64_t u) { CEREAL_RAPIDJSON_SCHEMA_HANDLE_VALUE_(Uint64, (CurrentContext(), u), (u)); }
    bool Double(double d)   { CEREAL_RAPIDJSON_SCHEMA_HANDLE_VALUE_(Double, (CurrentContext(), d), (d)); }
    bool RawNumber(const Ch* str, SizeType length, bool copy)
                                    { CEREAL_RAPIDJSON_SCHEMA_HANDLE_VALUE_(String, (CurrentContext(), str, length, copy), (str, length, copy)); }
    bool String(const Ch* str, SizeType length, bool copy)
                                    { CEREAL_RAPIDJSON_SCHEMA_HANDLE_VALUE_(String, (CurrentContext(), str, length, copy), (str, length, copy)); }

    bool StartObject() {
        CEREAL_RAPIDJSON_SCHEMA_HANDLE_BEGIN_(StartObject, (CurrentContext()));
        CEREAL_RAPIDJSON_SCHEMA_HANDLE_PARALLEL_(StartObject, ());
        return valid_ = !outputHandler_ || outputHandler_->StartObject();
    }
    
    bool Key(const Ch* str, SizeType len, bool copy) {
        if (!valid_) return false;
        AppendToken(str, len);
        if (!CurrentSchema().Key(CurrentContext(), str, len, copy)) return valid_ = false;
        CEREAL_RAPIDJSON_SCHEMA_HANDLE_PARALLEL_(Key, (str, len, copy));
        return valid_ = !outputHandler_ || outputHandler_->Key(str, len, copy);
    }
    
    bool EndObject(SizeType memberCount) { 
        if (!valid_) return false;
        CEREAL_RAPIDJSON_SCHEMA_HANDLE_PARALLEL_(EndObject, (memberCount));
        if (!CurrentSchema().EndObject(CurrentContext(), memberCount)) return valid_ = false;
        CEREAL_RAPIDJSON_SCHEMA_HANDLE_END_(EndObject, (memberCount));
    }

    bool StartArray() {
        CEREAL_RAPIDJSON_SCHEMA_HANDLE_BEGIN_(StartArray, (CurrentContext()));
        CEREAL_RAPIDJSON_SCHEMA_HANDLE_PARALLEL_(StartArray, ());
        return valid_ = !outputHandler_ || outputHandler_->StartArray();
    }
    
    bool EndArray(SizeType elementCount) {
        if (!valid_) return false;
        CEREAL_RAPIDJSON_SCHEMA_HANDLE_PARALLEL_(EndArray, (elementCount));
        if (!CurrentSchema().EndArray(CurrentContext(), elementCount)) return valid_ = false;
        CEREAL_RAPIDJSON_SCHEMA_HANDLE_END_(EndArray, (elementCount));
    }

#undef CEREAL_RAPIDJSON_SCHEMA_HANDLE_BEGIN_VERBOSE_
#undef CEREAL_RAPIDJSON_SCHEMA_HANDLE_BEGIN_
#undef CEREAL_RAPIDJSON_SCHEMA_HANDLE_PARALLEL_
#undef CEREAL_RAPIDJSON_SCHEMA_HANDLE_VALUE_

    // Implementation of ISchemaStateFactory<SchemaType>
    virtual ISchemaValidator* CreateSchemaValidator(const SchemaType& root) {
        return new (GetStateAllocator().Malloc(sizeof(GenericSchemaValidator))) GenericSchemaValidator(*schemaDocument_, root, documentStack_.template Bottom<char>(), documentStack_.GetSize(),
#if CEREAL_RAPIDJSON_SCHEMA_VERBOSE
        depth_ + 1,
#endif
        &GetStateAllocator());
    }

    virtual void DestroySchemaValidator(ISchemaValidator* validator) {
        GenericSchemaValidator* v = static_cast<GenericSchemaValidator*>(validator);
        v->~GenericSchemaValidator();
        StateAllocator::Free(v);
    }

    virtual void* CreateHasher() {
        return new (GetStateAllocator().Malloc(sizeof(HasherType))) HasherType(&GetStateAllocator());
    }

    virtual uint64_t GetHashCode(void* hasher) {
        return static_cast<HasherType*>(hasher)->GetHashCode();
    }

    virtual void DestroryHasher(void* hasher) {
        HasherType* h = static_cast<HasherType*>(hasher);
        h->~HasherType();
        StateAllocator::Free(h);
    }

    virtual void* MallocState(size_t size) {
        return GetStateAllocator().Malloc(size);
    }

    virtual void FreeState(void* p) {
        StateAllocator::Free(p);
    }

private:
    typedef typename SchemaType::Context Context;
    typedef GenericValue<UTF8<>, StateAllocator> HashCodeArray;
    typedef internal::Hasher<EncodingType, StateAllocator> HasherType;

    GenericSchemaValidator( 
        const SchemaDocumentType& schemaDocument,
        const SchemaType& root,
        const char* basePath, size_t basePathSize,
#if CEREAL_RAPIDJSON_SCHEMA_VERBOSE
        unsigned depth,
#endif
        StateAllocator* allocator = 0,
        size_t schemaStackCapacity = kDefaultSchemaStackCapacity,
        size_t documentStackCapacity = kDefaultDocumentStackCapacity)
        :
        schemaDocument_(&schemaDocument),
        root_(root),
        stateAllocator_(allocator),
        ownStateAllocator_(0),
        schemaStack_(allocator, schemaStackCapacity),
        documentStack_(allocator, documentStackCapacity),
        outputHandler_(0),
        error_(kObjectType),
        currentError_(),
        missingDependents_(),
        valid_(true)
#if CEREAL_RAPIDJSON_SCHEMA_VERBOSE
        , depth_(depth)
#endif
    {
        if (basePath && basePathSize)
            memcpy(documentStack_.template Push<char>(basePathSize), basePath, basePathSize);
    }

    StateAllocator& GetStateAllocator() {
        if (!stateAllocator_)
            stateAllocator_ = ownStateAllocator_ = CEREAL_RAPIDJSON_NEW(StateAllocator)();
        return *stateAllocator_;
    }

    bool BeginValue() {
        if (schemaStack_.Empty())
            PushSchema(root_);
        else {
            if (CurrentContext().inArray)
                internal::TokenHelper<internal::Stack<StateAllocator>, Ch>::AppendIndexToken(documentStack_, CurrentContext().arrayElementIndex);

            if (!CurrentSchema().BeginValue(CurrentContext()))
                return false;

            SizeType count = CurrentContext().patternPropertiesSchemaCount;
            const SchemaType** sa = CurrentContext().patternPropertiesSchemas;
            typename Context::PatternValidatorType patternValidatorType = CurrentContext().valuePatternValidatorType;
            bool valueUniqueness = CurrentContext().valueUniqueness;
            CEREAL_RAPIDJSON_ASSERT(CurrentContext().valueSchema);
            PushSchema(*CurrentContext().valueSchema);

            if (count > 0) {
                CurrentContext().objectPatternValidatorType = patternValidatorType;
                ISchemaValidator**& va = CurrentContext().patternPropertiesValidators;
                SizeType& validatorCount = CurrentContext().patternPropertiesValidatorCount;
                va = static_cast<ISchemaValidator**>(MallocState(sizeof(ISchemaValidator*) * count));
                for (SizeType i = 0; i < count; i++)
                    va[validatorCount++] = CreateSchemaValidator(*sa[i]);
            }

            CurrentContext().arrayUniqueness = valueUniqueness;
        }
        return true;
    }

    bool EndValue() {
        if (!CurrentSchema().EndValue(CurrentContext()))
            return false;

#if CEREAL_RAPIDJSON_SCHEMA_VERBOSE
        GenericStringBuffer<EncodingType> sb;
        schemaDocument_->GetPointer(&CurrentSchema()).Stringify(sb);

        *documentStack_.template Push<Ch>() = '\0';
        documentStack_.template Pop<Ch>(1);
        internal::PrintValidatorPointers(depth_, sb.GetString(), documentStack_.template Bottom<Ch>());
#endif

        uint64_t h = CurrentContext().arrayUniqueness ? static_cast<HasherType*>(CurrentContext().hasher)->GetHashCode() : 0;
        
        PopSchema();

        if (!schemaStack_.Empty()) {
            Context& context = CurrentContext();
            if (context.valueUniqueness) {
                HashCodeArray* a = static_cast<HashCodeArray*>(context.arrayElementHashCodes);
                if (!a)
                    CurrentContext().arrayElementHashCodes = a = new (GetStateAllocator().Malloc(sizeof(HashCodeArray))) HashCodeArray(kArrayType);
                for (typename HashCodeArray::ConstValueIterator itr = a->Begin(); itr != a->End(); ++itr)
                    if (itr->GetUint64() == h) {
                        DuplicateItems(static_cast<SizeType>(itr - a->Begin()), a->Size());
                        CEREAL_RAPIDJSON_INVALID_KEYWORD_RETURN(SchemaType::GetUniqueItemsString());
                    }
                a->PushBack(h, GetStateAllocator());
            }
        }

        // Remove the last token of document pointer
        while (!documentStack_.Empty() && *documentStack_.template Pop<Ch>(1) != '/')
            ;

        return true;
    }

    void AppendToken(const Ch* str, SizeType len) {
        documentStack_.template Reserve<Ch>(1 + len * 2); // worst case all characters are escaped as two characters
        *documentStack_.template PushUnsafe<Ch>() = '/';
        for (SizeType i = 0; i < len; i++) {
            if (str[i] == '~') {
                *documentStack_.template PushUnsafe<Ch>() = '~';
                *documentStack_.template PushUnsafe<Ch>() = '0';
            }
            else if (str[i] == '/') {
                *documentStack_.template PushUnsafe<Ch>() = '~';
                *documentStack_.template PushUnsafe<Ch>() = '1';
            }
            else
                *documentStack_.template PushUnsafe<Ch>() = str[i];
        }
    }

    CEREAL_RAPIDJSON_FORCEINLINE void PushSchema(const SchemaType& schema) { new (schemaStack_.template Push<Context>()) Context(*this, *this, &schema); }
    
    CEREAL_RAPIDJSON_FORCEINLINE void PopSchema() {
        Context* c = schemaStack_.template Pop<Context>(1);
        if (HashCodeArray* a = static_cast<HashCodeArray*>(c->arrayElementHashCodes)) {
            a->~HashCodeArray();
            StateAllocator::Free(a);
        }
        c->~Context();
    }

    void AddErrorLocation(ValueType& result, bool parent) {
        GenericStringBuffer<EncodingType> sb;
        PointerType instancePointer = GetInvalidDocumentPointer();
        ((parent && instancePointer.GetTokenCount() > 0)
            ? PointerType(instancePointer.GetTokens(), instancePointer.GetTokenCount() - 1)
            : instancePointer).StringifyUriFragment(sb);
        ValueType instanceRef(sb.GetString(), static_cast<SizeType>(sb.GetSize() / sizeof(Ch)),
            GetStateAllocator());
        result.AddMember(GetInstanceRefString(), instanceRef, GetStateAllocator());
        sb.Clear();
        memcpy(sb.Push(CurrentSchema().GetURI().GetStringLength()),
            CurrentSchema().GetURI().GetString(),
            CurrentSchema().GetURI().GetStringLength() * sizeof(Ch));
        GetInvalidSchemaPointer().StringifyUriFragment(sb);
        ValueType schemaRef(sb.GetString(), static_cast<SizeType>(sb.GetSize() / sizeof(Ch)),
            GetStateAllocator());
        result.AddMember(GetSchemaRefString(), schemaRef, GetStateAllocator());
    }

    void AddError(ValueType& keyword, ValueType& error) {
        typename ValueType::MemberIterator member = error_.FindMember(keyword);
        if (member == error_.MemberEnd())
            error_.AddMember(keyword, error, GetStateAllocator());
        else {
            if (member->value.IsObject()) {
                ValueType errors(kArrayType);
                errors.PushBack(member->value, GetStateAllocator());
                member->value = errors;
            }
            member->value.PushBack(error, GetStateAllocator());
        }
    }

    void AddCurrentError(const typename SchemaType::ValueType& keyword, bool parent = false) {
        AddErrorLocation(currentError_, parent);
        AddError(ValueType(keyword, GetStateAllocator(), false).Move(), currentError_);
    }

    void MergeError(ValueType& other) {
        for (typename ValueType::MemberIterator it = other.MemberBegin(), end = other.MemberEnd(); it != end; ++it) {
            AddError(it->name, it->value);
        }
    }

    void AddNumberError(const typename SchemaType::ValueType& keyword, ValueType& actual, const SValue& expected,
        const typename SchemaType::ValueType& (*exclusive)() = 0) {
        currentError_.SetObject();
        currentError_.AddMember(GetActualString(), actual, GetStateAllocator());
        currentError_.AddMember(GetExpectedString(), ValueType(expected, GetStateAllocator()).Move(), GetStateAllocator());
        if (exclusive)
            currentError_.AddMember(ValueType(exclusive(), GetStateAllocator()).Move(), true, GetStateAllocator());
        AddCurrentError(keyword);
    }

    void AddErrorArray(const typename SchemaType::ValueType& keyword,
        ISchemaValidator** subvalidators, SizeType count) {
        ValueType errors(kArrayType);
        for (SizeType i = 0; i < count; ++i)
            errors.PushBack(static_cast<GenericSchemaValidator*>(subvalidators[i])->GetError(), GetStateAllocator());
        currentError_.SetObject();
        currentError_.AddMember(GetErrorsString(), errors, GetStateAllocator());
        AddCurrentError(keyword);
    }

    const SchemaType& CurrentSchema() const { return *schemaStack_.template Top<Context>()->schema; }
    Context& CurrentContext() { return *schemaStack_.template Top<Context>(); }
    const Context& CurrentContext() const { return *schemaStack_.template Top<Context>(); }

    static const size_t kDefaultSchemaStackCapacity = 1024;
    static const size_t kDefaultDocumentStackCapacity = 256;
    const SchemaDocumentType* schemaDocument_;
    const SchemaType& root_;
    StateAllocator* stateAllocator_;
    StateAllocator* ownStateAllocator_;
    internal::Stack<StateAllocator> schemaStack_;    //!< stack to store the current path of schema (BaseSchemaType *)
    internal::Stack<StateAllocator> documentStack_;  //!< stack to store the current path of validating document (Ch)
    OutputHandler* outputHandler_;
    ValueType error_;
    ValueType currentError_;
    ValueType missingDependents_;
    bool valid_;
#if CEREAL_RAPIDJSON_SCHEMA_VERBOSE
    unsigned depth_;
#endif
};

typedef GenericSchemaValidator<SchemaDocument> SchemaValidator;

///////////////////////////////////////////////////////////////////////////////
// SchemaValidatingReader

//! A helper class for parsing with validation.
/*!
    This helper class is a functor, designed as a parameter of \ref GenericDocument::Populate().

    \tparam parseFlags Combination of \ref ParseFlag.
    \tparam InputStream Type of input stream, implementing Stream concept.
    \tparam SourceEncoding Encoding of the input stream.
    \tparam SchemaDocumentType Type of schema document.
    \tparam StackAllocator Allocator type for stack.
*/
template <
    unsigned parseFlags,
    typename InputStream,
    typename SourceEncoding,
    typename SchemaDocumentType = SchemaDocument,
    typename StackAllocator = CrtAllocator>
class SchemaValidatingReader {
public:
    typedef typename SchemaDocumentType::PointerType PointerType;
    typedef typename InputStream::Ch Ch;
    typedef GenericValue<SourceEncoding, StackAllocator> ValueType;

    //! Constructor
    /*!
        \param is Input stream.
        \param sd Schema document.
    */
    SchemaValidatingReader(InputStream& is, const SchemaDocumentType& sd) : is_(is), sd_(sd), invalidSchemaKeyword_(), error_(kObjectType), isValid_(true) {}

    template <typename Handler>
    bool operator()(Handler& handler) {
        GenericReader<SourceEncoding, typename SchemaDocumentType::EncodingType, StackAllocator> reader;
        GenericSchemaValidator<SchemaDocumentType, Handler> validator(sd_, handler);
        parseResult_ = reader.template Parse<parseFlags>(is_, validator);

        isValid_ = validator.IsValid();
        if (isValid_) {
            invalidSchemaPointer_ = PointerType();
            invalidSchemaKeyword_ = 0;
            invalidDocumentPointer_ = PointerType();
            error_.SetObject();
        }
        else {
            invalidSchemaPointer_ = validator.GetInvalidSchemaPointer();
            invalidSchemaKeyword_ = validator.GetInvalidSchemaKeyword();
            invalidDocumentPointer_ = validator.GetInvalidDocumentPointer();
            error_.CopyFrom(validator.GetError(), allocator_);
        }

        return parseResult_;
    }

    const ParseResult& GetParseResult() const { return parseResult_; }
    bool IsValid() const { return isValid_; }
    const PointerType& GetInvalidSchemaPointer() const { return invalidSchemaPointer_; }
    const Ch* GetInvalidSchemaKeyword() const { return invalidSchemaKeyword_; }
    const PointerType& GetInvalidDocumentPointer() const { return invalidDocumentPointer_; }
    const ValueType& GetError() const { return error_; }

private:
    InputStream& is_;
    const SchemaDocumentType& sd_;

    ParseResult parseResult_;
    PointerType invalidSchemaPointer_;
    const Ch* invalidSchemaKeyword_;
    PointerType invalidDocumentPointer_;
    StackAllocator allocator_;
    ValueType error_;
    bool isValid_;
};

CEREAL_RAPIDJSON_NAMESPACE_END
CEREAL_RAPIDJSON_DIAG_POP

#endif // CEREAL_RAPIDJSON_SCHEMA_H_
```
