## Supplemental code for "The motif composition of variable-number tandem repeats impacts gene expression": stream.h

```
// Tencent is pleased to support the open source community by making RapidJSON available.
//
// Copyright (C) 2015 THL A29 Limited, a Tencent company, and Milo Yip. All rights reserved.
//
// Licensed under the MIT License (the "License"); you may not use this file except
// in compliance with the License. You may obtain a copy of the License at
//
// http://opensource.org/licenses/MIT
//
// Unless required by applicable law or agreed to in writing, software distributed
// under the License is distributed on an "AS IS" BASIS, WITHOUT WARRANTIES OR
// CONDITIONS OF ANY KIND, either express or implied. See the License for the
// specific language governing permissions and limitations under the License.

#include "rapidjson.h"

#ifndef CEREAL_RAPIDJSON_STREAM_H_
#define CEREAL_RAPIDJSON_STREAM_H_

#include "encodings.h"

CEREAL_RAPIDJSON_NAMESPACE_BEGIN

///////////////////////////////////////////////////////////////////////////////
//  Stream

/*! \class rapidjson::Stream
    \brief Concept for reading and writing characters.

    For read-only stream, no need to implement PutBegin(), Put(), Flush() and PutEnd().

    For write-only stream, only need to implement Put() and Flush().

\code
concept Stream {
    typename Ch;    //!< Character type of the stream.

    //! Read the current character from stream without moving the read cursor.
    Ch Peek() const;

    //! Read the current character from stream and moving the read cursor to next character.
    Ch Take();

    //! Get the current read cursor.
    //! \return Number of characters read from start.
    size_t Tell();

    //! Begin writing operation at the current read pointer.
    //! \return The begin writer pointer.
    Ch* PutBegin();

    //! Write a character.
    void Put(Ch c);

    //! Flush the buffer.
    void Flush();

    //! End the writing operation.
    //! \param begin The begin write pointer returned by PutBegin().
    //! \return Number of characters written.
    size_t PutEnd(Ch* begin);
}
\endcode
*/

//! Provides additional information for stream.
/*!
    By using traits pattern, this type provides a default configuration for stream.
    For custom stream, this type can be specialized for other configuration.
    See TEST(Reader, CustomStringStream) in readertest.cpp for example.
*/
template<typename Stream>
struct StreamTraits {
    //! Whether to make local copy of stream for optimization during parsing.
    /*!
        By default, for safety, streams do not use local copy optimization.
        Stream that can be copied fast should specialize this, like StreamTraits<StringStream>.
    */
    enum { copyOptimization = 0 };
};

//! Reserve n characters for writing to a stream.
template<typename Stream>
inline void PutReserve(Stream& stream, size_t count) {
    (void)stream;
    (void)count;
}

//! Write character to a stream, presuming buffer is reserved.
template<typename Stream>
inline void PutUnsafe(Stream& stream, typename Stream::Ch c) {
    stream.Put(c);
}

//! Put N copies of a character to a stream.
template<typename Stream, typename Ch>
inline void PutN(Stream& stream, Ch c, size_t n) {
    PutReserve(stream, n);
    for (size_t i = 0; i < n; i++)
        PutUnsafe(stream, c);
}

///////////////////////////////////////////////////////////////////////////////
// GenericStreamWrapper

//! A Stream Wrapper
/*! \tThis string stream is a wrapper for any stream by just forwarding any
    \treceived message to the origin stream.
    \note implements Stream concept
*/

#if defined(_MSC_VER) && _MSC_VER <= 1800
CEREAL_RAPIDJSON_DIAG_PUSH
CEREAL_RAPIDJSON_DIAG_OFF(4702)  // unreachable code
CEREAL_RAPIDJSON_DIAG_OFF(4512)  // assignment operator could not be generated
#endif

template <typename InputStream, typename Encoding = UTF8<> >
class GenericStreamWrapper {
public:
    typedef typename Encoding::Ch Ch;
    GenericStreamWrapper(InputStream& is): is_(is) {}

    Ch Peek() const { return is_.Peek(); }
    Ch Take() { return is_.Take(); }
    size_t Tell() { return is_.Tell(); }
    Ch* PutBegin() { return is_.PutBegin(); }
    void Put(Ch ch) { is_.Put(ch); }
    void Flush() { is_.Flush(); }
    size_t PutEnd(Ch* ch) { return is_.PutEnd(ch); }

    // wrapper for MemoryStream
    const Ch* Peek4() const { return is_.Peek4(); }

    // wrapper for AutoUTFInputStream
    UTFType GetType() const { return is_.GetType(); }
    bool HasBOM() const { return is_.HasBOM(); }

protected:
    InputStream& is_;
};

#if defined(_MSC_VER) && _MSC_VER <= 1800
CEREAL_RAPIDJSON_DIAG_POP
#endif

///////////////////////////////////////////////////////////////////////////////
// StringStream

//! Read-only string stream.
/*! \note implements Stream concept
*/
template <typename Encoding>
struct GenericStringStream {
    typedef typename Encoding::Ch Ch;

    GenericStringStream(const Ch *src) : src_(src), head_(src) {}

    Ch Peek() const { return *src_; }
    Ch Take() { return *src_++; }
    size_t Tell() const { return static_cast<size_t>(src_ - head_); }

    Ch* PutBegin() { CEREAL_RAPIDJSON_ASSERT(false); return 0; }
    void Put(Ch) { CEREAL_RAPIDJSON_ASSERT(false); }
    void Flush() { CEREAL_RAPIDJSON_ASSERT(false); }
    size_t PutEnd(Ch*) { CEREAL_RAPIDJSON_ASSERT(false); return 0; }

    const Ch* src_;     //!< Current read position.
    const Ch* head_;    //!< Original head of the string.
};

template <typename Encoding>
struct StreamTraits<GenericStringStream<Encoding> > {
    enum { copyOptimization = 1 };
};

//! String stream with UTF8 encoding.
typedef GenericStringStream<UTF8<> > StringStream;

///////////////////////////////////////////////////////////////////////////////
// InsituStringStream

//! A read-write string stream.
/*! This string stream is particularly designed for in-situ parsing.
    \note implements Stream concept
*/
template <typename Encoding>
struct GenericInsituStringStream {
    typedef typename Encoding::Ch Ch;

    GenericInsituStringStream(Ch *src) : src_(src), dst_(0), head_(src) {}

    // Read
    Ch Peek() { return *src_; }
    Ch Take() { return *src_++; }
    size_t Tell() { return static_cast<size_t>(src_ - head_); }

    // Write
    void Put(Ch c) { CEREAL_RAPIDJSON_ASSERT(dst_ != 0); *dst_++ = c; }

    Ch* PutBegin() { return dst_ = src_; }
    size_t PutEnd(Ch* begin) { return static_cast<size_t>(dst_ - begin); }
    void Flush() {}

    Ch* Push(size_t count) { Ch* begin = dst_; dst_ += count; return begin; }
    void Pop(size_t count) { dst_ -= count; }

    Ch* src_;
    Ch* dst_;
    Ch* head_;
};

template <typename Encoding>
struct StreamTraits<GenericInsituStringStream<Encoding> > {
    enum { copyOptimization = 1 };
};

//! Insitu string stream with UTF8 encoding.
typedef GenericInsituStringStream<UTF8<> > InsituStringStream;

CEREAL_RAPIDJSON_NAMESPACE_END

#endif // CEREAL_RAPIDJSON_STREAM_H_
```
