## Supplemental code for "The motif composition of variable-number tandem repeats impacts gene expression": stringbuffer.h

```
// Tencent is pleased to support the open source community by making RapidJSON available.
// 
// Copyright (C) 2015 THL A29 Limited, a Tencent company, and Milo Yip. All rights reserved.
//
// Licensed under the MIT License (the "License"); you may not use this file except
// in compliance with the License. You may obtain a copy of the License at
//
// http://opensource.org/licenses/MIT
//
// Unless required by applicable law or agreed to in writing, software distributed 
// under the License is distributed on an "AS IS" BASIS, WITHOUT WARRANTIES OR 
// CONDITIONS OF ANY KIND, either express or implied. See the License for the 
// specific language governing permissions and limitations under the License.

#ifndef CEREAL_RAPIDJSON_STRINGBUFFER_H_
#define CEREAL_RAPIDJSON_STRINGBUFFER_H_

#include "stream.h"
#include "internal/stack.h"

#if CEREAL_RAPIDJSON_HAS_CXX11_RVALUE_REFS
#include <utility> // std::move
#endif

#include "internal/stack.h"

#if defined(__clang__)
CEREAL_RAPIDJSON_DIAG_PUSH
CEREAL_RAPIDJSON_DIAG_OFF(c++98-compat)
#endif

CEREAL_RAPIDJSON_NAMESPACE_BEGIN

//! Represents an in-memory output stream.
/*!
    \tparam Encoding Encoding of the stream.
    \tparam Allocator type for allocating memory buffer.
    \note implements Stream concept
*/
template <typename Encoding, typename Allocator = CrtAllocator>
class GenericStringBuffer {
public:
    typedef typename Encoding::Ch Ch;

    GenericStringBuffer(Allocator* allocator = 0, size_t capacity = kDefaultCapacity) : stack_(allocator, capacity) {}

#if CEREAL_RAPIDJSON_HAS_CXX11_RVALUE_REFS
    GenericStringBuffer(GenericStringBuffer&& rhs) : stack_(std::move(rhs.stack_)) {}
    GenericStringBuffer& operator=(GenericStringBuffer&& rhs) {
        if (&rhs != this)
            stack_ = std::move(rhs.stack_);
        return *this;
    }
#endif

    void Put(Ch c) { *stack_.template Push<Ch>() = c; }
    void PutUnsafe(Ch c) { *stack_.template PushUnsafe<Ch>() = c; }
    void Flush() {}

    void Clear() { stack_.Clear(); }
    void ShrinkToFit() {
        // Push and pop a null terminator. This is safe.
        *stack_.template Push<Ch>() = '\0';
        stack_.ShrinkToFit();
        stack_.template Pop<Ch>(1);
    }

    void Reserve(size_t count) { stack_.template Reserve<Ch>(count); }
    Ch* Push(size_t count) { return stack_.template Push<Ch>(count); }
    Ch* PushUnsafe(size_t count) { return stack_.template PushUnsafe<Ch>(count); }
    void Pop(size_t count) { stack_.template Pop<Ch>(count); }

    const Ch* GetString() const {
        // Push and pop a null terminator. This is safe.
        *stack_.template Push<Ch>() = '\0';
        stack_.template Pop<Ch>(1);

        return stack_.template Bottom<Ch>();
    }

    //! Get the size of string in bytes in the string buffer.
    size_t GetSize() const { return stack_.GetSize(); }

    //! Get the length of string in Ch in the string buffer.
    size_t GetLength() const { return stack_.GetSize() / sizeof(Ch); }

    static const size_t kDefaultCapacity = 256;
    mutable internal::Stack<Allocator> stack_;

private:
    // Prohibit copy constructor & assignment operator.
    GenericStringBuffer(const GenericStringBuffer&);
    GenericStringBuffer& operator=(const GenericStringBuffer&);
};

//! String buffer with UTF8 encoding
typedef GenericStringBuffer<UTF8<> > StringBuffer;

template<typename Encoding, typename Allocator>
inline void PutReserve(GenericStringBuffer<Encoding, Allocator>& stream, size_t count) {
    stream.Reserve(count);
}

template<typename Encoding, typename Allocator>
inline void PutUnsafe(GenericStringBuffer<Encoding, Allocator>& stream, typename Encoding::Ch c) {
    stream.PutUnsafe(c);
}

//! Implement specialized version of PutN() with memset() for better performance.
template<>
inline void PutN(GenericStringBuffer<UTF8<> >& stream, char c, size_t n) {
    std::memset(stream.stack_.Push<char>(n), c, n * sizeof(c));
}

CEREAL_RAPIDJSON_NAMESPACE_END

#if defined(__clang__)
CEREAL_RAPIDJSON_DIAG_POP
#endif

#endif // CEREAL_RAPIDJSON_STRINGBUFFER_H_
```
