## Supplemental code for "The motif composition of variable-number tandem repeats impacts gene expression": writer.h

```
// Tencent is pleased to support the open source community by making RapidJSON available.
// 
// Copyright (C) 2015 THL A29 Limited, a Tencent company, and Milo Yip. All rights reserved.
//
// Licensed under the MIT License (the "License"); you may not use this file except
// in compliance with the License. You may obtain a copy of the License at
//
// http://opensource.org/licenses/MIT
//
// Unless required by applicable law or agreed to in writing, software distributed 
// under the License is distributed on an "AS IS" BASIS, WITHOUT WARRANTIES OR 
// CONDITIONS OF ANY KIND, either express or implied. See the License for the 
// specific language governing permissions and limitations under the License.

#ifndef CEREAL_RAPIDJSON_WRITER_H_
#define CEREAL_RAPIDJSON_WRITER_H_

#include "stream.h"
#include "internal/meta.h"
#include "internal/stack.h"
#include "internal/strfunc.h"
#include "internal/dtoa.h"
#include "internal/itoa.h"
#include "stringbuffer.h"
#include <new>      // placement new

#if defined(CEREAL_RAPIDJSON_SIMD) && defined(_MSC_VER)
#include <intrin.h>
#pragma intrinsic(_BitScanForward)
#endif
#ifdef CEREAL_RAPIDJSON_SSE42
#include <nmmintrin.h>
#elif defined(CEREAL_RAPIDJSON_SSE2)
#include <emmintrin.h>
#elif defined(CEREAL_RAPIDJSON_NEON)
#include <arm_neon.h>
#endif

#ifdef __clang__
CEREAL_RAPIDJSON_DIAG_PUSH
CEREAL_RAPIDJSON_DIAG_OFF(padded)
CEREAL_RAPIDJSON_DIAG_OFF(unreachable-code)
CEREAL_RAPIDJSON_DIAG_OFF(c++98-compat)
#elif defined(_MSC_VER)
CEREAL_RAPIDJSON_DIAG_PUSH
CEREAL_RAPIDJSON_DIAG_OFF(4127) // conditional expression is constant
#endif

CEREAL_RAPIDJSON_NAMESPACE_BEGIN

///////////////////////////////////////////////////////////////////////////////
// WriteFlag

/*! \def CEREAL_RAPIDJSON_WRITE_DEFAULT_FLAGS 
    \ingroup CEREAL_RAPIDJSON_CONFIG
    \brief User-defined kWriteDefaultFlags definition.

    User can define this as any \c WriteFlag combinations.
*/
#ifndef CEREAL_RAPIDJSON_WRITE_DEFAULT_FLAGS
#define CEREAL_RAPIDJSON_WRITE_DEFAULT_FLAGS kWriteNoFlags
#endif

//! Combination of writeFlags
enum WriteFlag {
    kWriteNoFlags = 0,              //!< No flags are set.
    kWriteValidateEncodingFlag = 1, //!< Validate encoding of JSON strings.
    kWriteNanAndInfFlag = 2,        //!< Allow writing of Infinity, -Infinity and NaN.
    kWriteDefaultFlags = CEREAL_RAPIDJSON_WRITE_DEFAULT_FLAGS  //!< Default write flags. Can be customized by defining CEREAL_RAPIDJSON_WRITE_DEFAULT_FLAGS
};

//! JSON writer
/*! Writer implements the concept Handler.
    It generates JSON text by events to an output os.

    User may programmatically calls the functions of a writer to generate JSON text.

    On the other side, a writer can also be passed to objects that generates events, 

    for example Reader::Parse() and Document::Accept().

    \tparam OutputStream Type of output stream.
    \tparam SourceEncoding Encoding of source string.
    \tparam TargetEncoding Encoding of output stream.
    \tparam StackAllocator Type of allocator for allocating memory of stack.
    \note implements Handler concept
*/
template<typename OutputStream, typename SourceEncoding = UTF8<>, typename TargetEncoding = UTF8<>, typename StackAllocator = CrtAllocator, unsigned writeFlags = kWriteDefaultFlags>
class Writer {
public:
    typedef typename SourceEncoding::Ch Ch;

    static const int kDefaultMaxDecimalPlaces = 324;

    //! Constructor
    /*! \param os Output stream.
        \param stackAllocator User supplied allocator. If it is null, it will create a private one.
        \param levelDepth Initial capacity of stack.
    */
    explicit
    Writer(OutputStream& os, StackAllocator* stackAllocator = 0, size_t levelDepth = kDefaultLevelDepth) : 
        os_(&os), level_stack_(stackAllocator, levelDepth * sizeof(Level)), maxDecimalPlaces_(kDefaultMaxDecimalPlaces), hasRoot_(false) {}

    explicit
    Writer(StackAllocator* allocator = 0, size_t levelDepth = kDefaultLevelDepth) :
        os_(0), level_stack_(allocator, levelDepth * sizeof(Level)), maxDecimalPlaces_(kDefaultMaxDecimalPlaces), hasRoot_(false) {}

#if CEREAL_RAPIDJSON_HAS_CXX11_RVALUE_REFS
    Writer(Writer&& rhs) :
        os_(rhs.os_), level_stack_(std::move(rhs.level_stack_)), maxDecimalPlaces_(rhs.maxDecimalPlaces_), hasRoot_(rhs.hasRoot_) {
        rhs.os_ = 0;
    }
#endif

    //! Reset the writer with a new stream.
    /*!
        This function reset the writer with a new stream and default settings,
        in order to make a Writer object reusable for output multiple JSONs.

        \param os New output stream.
        \code
        Writer<OutputStream> writer(os1);
        writer.StartObject();
        // ...
        writer.EndObject();

        writer.Reset(os2);
        writer.StartObject();
        // ...
        writer.EndObject();
        \endcode
    */
    void Reset(OutputStream& os) {
        os_ = &os;
        hasRoot_ = false;
        level_stack_.Clear();
    }

    //! Checks whether the output is a complete JSON.
    /*!
        A complete JSON has a complete root object or array.
    */
    bool IsComplete() const {
        return hasRoot_ && level_stack_.Empty();
    }

    int GetMaxDecimalPlaces() const {
        return maxDecimalPlaces_;
    }

    //! Sets the maximum number of decimal places for double output.
    /*!
        This setting truncates the output with specified number of decimal places.

        For example, 

        \code
        writer.SetMaxDecimalPlaces(3);
        writer.StartArray();
        writer.Double(0.12345);                 // "0.123"
        writer.Double(0.0001);                  // "0.0"
        writer.Double(1.234567890123456e30);    // "1.234567890123456e30" (do not truncate significand for positive exponent)
        writer.Double(1.23e-4);                 // "0.0"                  (do truncate significand for negative exponent)
        writer.EndArray();
        \endcode

        The default setting does not truncate any decimal places. You can restore to this setting by calling
        \code
        writer.SetMaxDecimalPlaces(Writer::kDefaultMaxDecimalPlaces);
        \endcode
    */
    void SetMaxDecimalPlaces(int maxDecimalPlaces) {
        maxDecimalPlaces_ = maxDecimalPlaces;
    }

    /*!@name Implementation of Handler
        \see Handler
    */
    //@{

    bool Null()                 { Prefix(kNullType);   return EndValue(WriteNull()); }
    bool Bool(bool b)           { Prefix(b ? kTrueType : kFalseType); return EndValue(WriteBool(b)); }
    bool Int(int i)             { Prefix(kNumberType); return EndValue(WriteInt(i)); }
    bool Uint(unsigned u)       { Prefix(kNumberType); return EndValue(WriteUint(u)); }
    bool Int64(int64_t i64)     { Prefix(kNumberType); return EndValue(WriteInt64(i64)); }
    bool Uint64(uint64_t u64)   { Prefix(kNumberType); return EndValue(WriteUint64(u64)); }

    //! Writes the given \c double value to the stream
    /*!
        \param d The value to be written.
        \return Whether it is succeed.
    */
    bool Double(double d)       { Prefix(kNumberType); return EndValue(WriteDouble(d)); }

    bool RawNumber(const Ch* str, SizeType length, bool copy = false) {
        CEREAL_RAPIDJSON_ASSERT(str != 0);
        (void)copy;
        Prefix(kNumberType);
        return EndValue(WriteString(str, length));
    }

    bool String(const Ch* str, SizeType length, bool copy = false) {
        CEREAL_RAPIDJSON_ASSERT(str != 0);
        (void)copy;
        Prefix(kStringType);
        return EndValue(WriteString(str, length));
    }

#if CEREAL_RAPIDJSON_HAS_STDSTRING
    bool String(const std::basic_string<Ch>& str) {
        return String(str.data(), SizeType(str.size()));
    }
#endif

    bool StartObject() {
        Prefix(kObjectType);
        new (level_stack_.template Push<Level>()) Level(false);
        return WriteStartObject();
    }

    bool Key(const Ch* str, SizeType length, bool copy = false) { return String(str, length, copy); }

#if CEREAL_RAPIDJSON_HAS_STDSTRING
    bool Key(const std::basic_string<Ch>& str)
    {
      return Key(str.data(), SizeType(str.size()));
    }
#endif
	
    bool EndObject(SizeType memberCount = 0) {
        (void)memberCount;
        CEREAL_RAPIDJSON_ASSERT(level_stack_.GetSize() >= sizeof(Level)); // not inside an Object
        CEREAL_RAPIDJSON_ASSERT(!level_stack_.template Top<Level>()->inArray); // currently inside an Array, not Object
        CEREAL_RAPIDJSON_ASSERT(0 == level_stack_.template Top<Level>()->valueCount % 2); // Object has a Key without a Value
        level_stack_.template Pop<Level>(1);
        return EndValue(WriteEndObject());
    }

    bool StartArray() {
        Prefix(kArrayType);
        new (level_stack_.template Push<Level>()) Level(true);
        return WriteStartArray();
    }

    bool EndArray(SizeType elementCount = 0) {
        (void)elementCount;
        CEREAL_RAPIDJSON_ASSERT(level_stack_.GetSize() >= sizeof(Level));
        CEREAL_RAPIDJSON_ASSERT(level_stack_.template Top<Level>()->inArray);
        level_stack_.template Pop<Level>(1);
        return EndValue(WriteEndArray());
    }
    //@}

    /*! @name Convenience extensions */
    //@{

    //! Simpler but slower overload.
    bool String(const Ch* const& str) { return String(str, internal::StrLen(str)); }
    bool Key(const Ch* const& str) { return Key(str, internal::StrLen(str)); }
    
    //@}

    //! Write a raw JSON value.
    /*!
        For user to write a stringified JSON as a value.

        \param json A well-formed JSON value. It should not contain null character within [0, length - 1] range.
        \param length Length of the json.
        \param type Type of the root of json.
    */
    bool RawValue(const Ch* json, size_t length, Type type) {
        CEREAL_RAPIDJSON_ASSERT(json != 0);
        Prefix(type);
        return EndValue(WriteRawValue(json, length));
    }

    //! Flush the output stream.
    /*!
        Allows the user to flush the output stream immediately.
     */
    void Flush() {
        os_->Flush();
    }

protected:
    //! Information for each nested level
    struct Level {
        Level(bool inArray_) : valueCount(0), inArray(inArray_) {}
        size_t valueCount;  //!< number of values in this level
        bool inArray;       //!< true if in array, otherwise in object
    };

    static const size_t kDefaultLevelDepth = 32;

    bool WriteNull()  {
        PutReserve(*os_, 4);
        PutUnsafe(*os_, 'n'); PutUnsafe(*os_, 'u'); PutUnsafe(*os_, 'l'); PutUnsafe(*os_, 'l'); return true;
    }

    bool WriteBool(bool b)  {
        if (b) {
            PutReserve(*os_, 4);
            PutUnsafe(*os_, 't'); PutUnsafe(*os_, 'r'); PutUnsafe(*os_, 'u'); PutUnsafe(*os_, 'e');
        }
        else {
            PutReserve(*os_, 5);
            PutUnsafe(*os_, 'f'); PutUnsafe(*os_, 'a'); PutUnsafe(*os_, 'l'); PutUnsafe(*os_, 's'); PutUnsafe(*os_, 'e');
        }
        return true;
    }

    bool WriteInt(int i) {
        char buffer[11];
        const char* end = internal::i32toa(i, buffer);
        PutReserve(*os_, static_cast<size_t>(end - buffer));
        for (const char* p = buffer; p != end; ++p)
            PutUnsafe(*os_, static_cast<typename OutputStream::Ch>(*p));
        return true;
    }

    bool WriteUint(unsigned u) {
        char buffer[10];
        const char* end = internal::u32toa(u, buffer);
        PutReserve(*os_, static_cast<size_t>(end - buffer));
        for (const char* p = buffer; p != end; ++p)
            PutUnsafe(*os_, static_cast<typename OutputStream::Ch>(*p));
        return true;
    }

    bool WriteInt64(int64_t i64) {
        char buffer[21];
        const char* end = internal::i64toa(i64, buffer);
        PutReserve(*os_, static_cast<size_t>(end - buffer));
        for (const char* p = buffer; p != end; ++p)
            PutUnsafe(*os_, static_cast<typename OutputStream::Ch>(*p));
        return true;
    }

    bool WriteUint64(uint64_t u64) {
        char buffer[20];
        char* end = internal::u64toa(u64, buffer);
        PutReserve(*os_, static_cast<size_t>(end - buffer));
        for (char* p = buffer; p != end; ++p)
            PutUnsafe(*os_, static_cast<typename OutputStream::Ch>(*p));
        return true;
    }

    bool WriteDouble(double d) {
        if (internal::Double(d).IsNanOrInf()) {
            if (!(writeFlags & kWriteNanAndInfFlag))
                return false;
            if (internal::Double(d).IsNan()) {
                PutReserve(*os_, 3);
                PutUnsafe(*os_, 'N'); PutUnsafe(*os_, 'a'); PutUnsafe(*os_, 'N');
                return true;
            }
            if (internal::Double(d).Sign()) {
                PutReserve(*os_, 9);
                PutUnsafe(*os_, '-');
            }
            else
                PutReserve(*os_, 8);
            PutUnsafe(*os_, 'I'); PutUnsafe(*os_, 'n'); PutUnsafe(*os_, 'f');
            PutUnsafe(*os_, 'i'); PutUnsafe(*os_, 'n'); PutUnsafe(*os_, 'i'); PutUnsafe(*os_, 't'); PutUnsafe(*os_, 'y');
            return true;
        }

        char buffer[25];
        char* end = internal::dtoa(d, buffer, maxDecimalPlaces_);
        PutReserve(*os_, static_cast<size_t>(end - buffer));
        for (char* p = buffer; p != end; ++p)
            PutUnsafe(*os_, static_cast<typename OutputStream::Ch>(*p));
        return true;
    }

    bool WriteString(const Ch* str, SizeType length)  {
        static const typename OutputStream::Ch hexDigits[16] = { '0', '1', '2', '3', '4', '5', '6', '7', '8', '9', 'A', 'B', 'C', 'D', 'E', 'F' };
        static const char escape[256] = {
#define Z16 0,0,0,0,0,0,0,0,0,0,0,0,0,0,0,0
            //0    1    2    3    4    5    6    7    8    9    A    B    C    D    E    F
            'u', 'u', 'u', 'u', 'u', 'u', 'u', 'u', 'b', 't', 'n', 'u', 'f', 'r', 'u', 'u', // 00
            'u', 'u', 'u', 'u', 'u', 'u', 'u', 'u', 'u', 'u', 'u', 'u', 'u', 'u', 'u', 'u', // 10
              0,   0, '"',   0,   0,   0,   0,   0,   0,   0,   0,   0,   0,   0,   0,   0, // 20
            Z16, Z16,                                                                       // 30~4F
              0,   0,   0,   0,   0,   0,   0,   0,   0,   0,   0,   0,'\\',   0,   0,   0, // 50
            Z16, Z16, Z16, Z16, Z16, Z16, Z16, Z16, Z16, Z16                                // 60~FF
#undef Z16
        };

        if (TargetEncoding::supportUnicode)
            PutReserve(*os_, 2 + length * 6); // "\uxxxx..."
        else
            PutReserve(*os_, 2 + length * 12);  // "\uxxxx\uyyyy..."

        PutUnsafe(*os_, '\"');
        GenericStringStream<SourceEncoding> is(str);
        while (ScanWriteUnescapedString(is, length)) {
            const Ch c = is.Peek();
            if (!TargetEncoding::supportUnicode && static_cast<unsigned>(c) >= 0x80) {
                // Unicode escaping
                unsigned codepoint;
                if (CEREAL_RAPIDJSON_UNLIKELY(!SourceEncoding::Decode(is, &codepoint)))
                    return false;
                PutUnsafe(*os_, '\\');
                PutUnsafe(*os_, 'u');
                if (codepoint <= 0xD7FF || (codepoint >= 0xE000 && codepoint <= 0xFFFF)) {
                    PutUnsafe(*os_, hexDigits[(codepoint >> 12) & 15]);
                    PutUnsafe(*os_, hexDigits[(codepoint >>  8) & 15]);
                    PutUnsafe(*os_, hexDigits[(codepoint >>  4) & 15]);
                    PutUnsafe(*os_, hexDigits[(codepoint      ) & 15]);
                }
                else {
                    CEREAL_RAPIDJSON_ASSERT(codepoint >= 0x010000 && codepoint <= 0x10FFFF);
                    // Surrogate pair
                    unsigned s = codepoint - 0x010000;
                    unsigned lead = (s >> 10) + 0xD800;
                    unsigned trail = (s & 0x3FF) + 0xDC00;
                    PutUnsafe(*os_, hexDigits[(lead >> 12) & 15]);
                    PutUnsafe(*os_, hexDigits[(lead >>  8) & 15]);
                    PutUnsafe(*os_, hexDigits[(lead >>  4) & 15]);
                    PutUnsafe(*os_, hexDigits[(lead      ) & 15]);
                    PutUnsafe(*os_, '\\');
                    PutUnsafe(*os_, 'u');
                    PutUnsafe(*os_, hexDigits[(trail >> 12) & 15]);
                    PutUnsafe(*os_, hexDigits[(trail >>  8) & 15]);
                    PutUnsafe(*os_, hexDigits[(trail >>  4) & 15]);
                    PutUnsafe(*os_, hexDigits[(trail      ) & 15]);                    
                }
            }
            else if ((sizeof(Ch) == 1 || static_cast<unsigned>(c) < 256) && CEREAL_RAPIDJSON_UNLIKELY(escape[static_cast<unsigned char>(c)]))  {
                is.Take();
                PutUnsafe(*os_, '\\');
                PutUnsafe(*os_, static_cast<typename OutputStream::Ch>(escape[static_cast<unsigned char>(c)]));
                if (escape[static_cast<unsigned char>(c)] == 'u') {
                    PutUnsafe(*os_, '0');
                    PutUnsafe(*os_, '0');
                    PutUnsafe(*os_, hexDigits[static_cast<unsigned char>(c) >> 4]);
                    PutUnsafe(*os_, hexDigits[static_cast<unsigned char>(c) & 0xF]);
                }
            }
            else if (CEREAL_RAPIDJSON_UNLIKELY(!(writeFlags & kWriteValidateEncodingFlag ? 
                Transcoder<SourceEncoding, TargetEncoding>::Validate(is, *os_) :
                Transcoder<SourceEncoding, TargetEncoding>::TranscodeUnsafe(is, *os_))))
                return false;
        }
        PutUnsafe(*os_, '\"');
        return true;
    }

    bool ScanWriteUnescapedString(GenericStringStream<SourceEncoding>& is, size_t length) {
        return CEREAL_RAPIDJSON_LIKELY(is.Tell() < length);
    }

    bool WriteStartObject() { os_->Put('{'); return true; }
    bool WriteEndObject()   { os_->Put('}'); return true; }
    bool WriteStartArray()  { os_->Put('['); return true; }
    bool WriteEndArray()    { os_->Put(']'); return true; }

    bool WriteRawValue(const Ch* json, size_t length) {
        PutReserve(*os_, length);
        GenericStringStream<SourceEncoding> is(json);
        while (CEREAL_RAPIDJSON_LIKELY(is.Tell() < length)) {
            CEREAL_RAPIDJSON_ASSERT(is.Peek() != '\0');
            if (CEREAL_RAPIDJSON_UNLIKELY(!(writeFlags & kWriteValidateEncodingFlag ? 
                Transcoder<SourceEncoding, TargetEncoding>::Validate(is, *os_) :
                Transcoder<SourceEncoding, TargetEncoding>::TranscodeUnsafe(is, *os_))))
                return false;
        }
        return true;
    }

    void Prefix(Type type) {
        (void)type;
        if (CEREAL_RAPIDJSON_LIKELY(level_stack_.GetSize() != 0)) { // this value is not at root
            Level* level = level_stack_.template Top<Level>();
            if (level->valueCount > 0) {
                if (level->inArray) 
                    os_->Put(','); // add comma if it is not the first element in array
                else  // in object
                    os_->Put((level->valueCount % 2 == 0) ? ',' : ':');
            }
            if (!level->inArray && level->valueCount % 2 == 0)
                CEREAL_RAPIDJSON_ASSERT(type == kStringType);  // if it's in object, then even number should be a name
            level->valueCount++;
        }
        else {
            CEREAL_RAPIDJSON_ASSERT(!hasRoot_);    // Should only has one and only one root.
            hasRoot_ = true;
        }
    }

    // Flush the value if it is the top level one.
    bool EndValue(bool ret) {
        if (CEREAL_RAPIDJSON_UNLIKELY(level_stack_.Empty()))   // end of json text
            Flush();
        return ret;
    }

    OutputStream* os_;
    internal::Stack<StackAllocator> level_stack_;
    int maxDecimalPlaces_;
    bool hasRoot_;

private:
    // Prohibit copy constructor & assignment operator.
    Writer(const Writer&);
    Writer& operator=(const Writer&);
};

// Full specialization for StringStream to prevent memory copying

template<>
inline bool Writer<StringBuffer>::WriteInt(int i) {
    char *buffer = os_->Push(11);
    const char* end = internal::i32toa(i, buffer);
    os_->Pop(static_cast<size_t>(11 - (end - buffer)));
    return true;
}

template<>
inline bool Writer<StringBuffer>::WriteUint(unsigned u) {
    char *buffer = os_->Push(10);
    const char* end = internal::u32toa(u, buffer);
    os_->Pop(static_cast<size_t>(10 - (end - buffer)));
    return true;
}

template<>
inline bool Writer<StringBuffer>::WriteInt64(int64_t i64) {
    char *buffer = os_->Push(21);
    const char* end = internal::i64toa(i64, buffer);
    os_->Pop(static_cast<size_t>(21 - (end - buffer)));
    return true;
}

template<>
inline bool Writer<StringBuffer>::WriteUint64(uint64_t u) {
    char *buffer = os_->Push(20);
    const char* end = internal::u64toa(u, buffer);
    os_->Pop(static_cast<size_t>(20 - (end - buffer)));
    return true;
}

template<>
inline bool Writer<StringBuffer>::WriteDouble(double d) {
    if (internal::Double(d).IsNanOrInf()) {
        // Note: This code path can only be reached if (CEREAL_RAPIDJSON_WRITE_DEFAULT_FLAGS & kWriteNanAndInfFlag).
        if (!(kWriteDefaultFlags & kWriteNanAndInfFlag))
            return false;
        if (internal::Double(d).IsNan()) {
            PutReserve(*os_, 3);
            PutUnsafe(*os_, 'N'); PutUnsafe(*os_, 'a'); PutUnsafe(*os_, 'N');
            return true;
        }
        if (internal::Double(d).Sign()) {
            PutReserve(*os_, 9);
            PutUnsafe(*os_, '-');
        }
        else
            PutReserve(*os_, 8);
        PutUnsafe(*os_, 'I'); PutUnsafe(*os_, 'n'); PutUnsafe(*os_, 'f');
        PutUnsafe(*os_, 'i'); PutUnsafe(*os_, 'n'); PutUnsafe(*os_, 'i'); PutUnsafe(*os_, 't'); PutUnsafe(*os_, 'y');
        return true;
    }
    
    char *buffer = os_->Push(25);
    char* end = internal::dtoa(d, buffer, maxDecimalPlaces_);
    os_->Pop(static_cast<size_t>(25 - (end - buffer)));
    return true;
}

#if defined(CEREAL_RAPIDJSON_SSE2) || defined(CEREAL_RAPIDJSON_SSE42)
template<>
inline bool Writer<StringBuffer>::ScanWriteUnescapedString(StringStream& is, size_t length) {
    if (length < 16)
        return CEREAL_RAPIDJSON_LIKELY(is.Tell() < length);

    if (!CEREAL_RAPIDJSON_LIKELY(is.Tell() < length))
        return false;

    const char* p = is.src_;
    const char* end = is.head_ + length;
    const char* nextAligned = reinterpret_cast<const char*>((reinterpret_cast<size_t>(p) + 15) & static_cast<size_t>(~15));
    const char* endAligned = reinterpret_cast<const char*>(reinterpret_cast<size_t>(end) & static_cast<size_t>(~15));
    if (nextAligned > end)
        return true;

    while (p != nextAligned)
        if (*p < 0x20 || *p == '\"' || *p == '\\') {
            is.src_ = p;
            return CEREAL_RAPIDJSON_LIKELY(is.Tell() < length);
        }
        else
            os_->PutUnsafe(*p++);

    // The rest of string using SIMD
    static const char dquote[16] = { '\"', '\"', '\"', '\"', '\"', '\"', '\"', '\"', '\"', '\"', '\"', '\"', '\"', '\"', '\"', '\"' };
    static const char bslash[16] = { '\\', '\\', '\\', '\\', '\\', '\\', '\\', '\\', '\\', '\\', '\\', '\\', '\\', '\\', '\\', '\\' };
    static const char space[16]  = { 0x1F, 0x1F, 0x1F, 0x1F, 0x1F, 0x1F, 0x1F, 0x1F, 0x1F, 0x1F, 0x1F, 0x1F, 0x1F, 0x1F, 0x1F, 0x1F };
    const __m128i dq = _mm_loadu_si128(reinterpret_cast<const __m128i *>(&dquote[0]));
    const __m128i bs = _mm_loadu_si128(reinterpret_cast<const __m128i *>(&bslash[0]));
    const __m128i sp = _mm_loadu_si128(reinterpret_cast<const __m128i *>(&space[0]));

    for (; p != endAligned; p += 16) {
        const __m128i s = _mm_load_si128(reinterpret_cast<const __m128i *>(p));
        const __m128i t1 = _mm_cmpeq_epi8(s, dq);
        const __m128i t2 = _mm_cmpeq_epi8(s, bs);
        const __m128i t3 = _mm_cmpeq_epi8(_mm_max_epu8(s, sp), sp); // s < 0x20 <=> max(s, 0x1F) == 0x1F
        const __m128i x = _mm_or_si128(_mm_or_si128(t1, t2), t3);
        unsigned short r = static_cast<unsigned short>(_mm_movemask_epi8(x));
        if (CEREAL_RAPIDJSON_UNLIKELY(r != 0)) {   // some of characters is escaped
            SizeType len;
#ifdef _MSC_VER         // Find the index of first escaped
            unsigned long offset;
            _BitScanForward(&offset, r);
            len = offset;
#else
            len = static_cast<SizeType>(__builtin_ffs(r) - 1);
#endif
            char* q = reinterpret_cast<char*>(os_->PushUnsafe(len));
            for (size_t i = 0; i < len; i++)
                q[i] = p[i];

            p += len;
            break;
        }
        _mm_storeu_si128(reinterpret_cast<__m128i *>(os_->PushUnsafe(16)), s);
    }

    is.src_ = p;
    return CEREAL_RAPIDJSON_LIKELY(is.Tell() < length);
}
#elif defined(CEREAL_RAPIDJSON_NEON)
template<>
inline bool Writer<StringBuffer>::ScanWriteUnescapedString(StringStream& is, size_t length) {
    if (length < 16)
        return CEREAL_RAPIDJSON_LIKELY(is.Tell() < length);

    if (!CEREAL_RAPIDJSON_LIKELY(is.Tell() < length))
        return false;

    const char* p = is.src_;
    const char* end = is.head_ + length;
    const char* nextAligned = reinterpret_cast<const char*>((reinterpret_cast<size_t>(p) + 15) & static_cast<size_t>(~15));
    const char* endAligned = reinterpret_cast<const char*>(reinterpret_cast<size_t>(end) & static_cast<size_t>(~15));
    if (nextAligned > end)
        return true;

    while (p != nextAligned)
        if (*p < 0x20 || *p == '\"' || *p == '\\') {
            is.src_ = p;
            return CEREAL_RAPIDJSON_LIKELY(is.Tell() < length);
        }
        else
            os_->PutUnsafe(*p++);

    // The rest of string using SIMD
    const uint8x16_t s0 = vmovq_n_u8('"');
    const uint8x16_t s1 = vmovq_n_u8('\\');
    const uint8x16_t s2 = vmovq_n_u8('\b');
    const uint8x16_t s3 = vmovq_n_u8(32);

    for (; p != endAligned; p += 16) {
        const uint8x16_t s = vld1q_u8(reinterpret_cast<const uint8_t *>(p));
        uint8x16_t x = vceqq_u8(s, s0);
        x = vorrq_u8(x, vceqq_u8(s, s1));
        x = vorrq_u8(x, vceqq_u8(s, s2));
        x = vorrq_u8(x, vcltq_u8(s, s3));

        x = vrev64q_u8(x);                     // Rev in 64
        uint64_t low = vgetq_lane_u64(reinterpret_cast<uint64x2_t>(x), 0);   // extract
        uint64_t high = vgetq_lane_u64(reinterpret_cast<uint64x2_t>(x), 1);  // extract

        SizeType len = 0;
        bool escaped = false;
        if (low == 0) {
            if (high != 0) {
                unsigned lz = (unsigned)__builtin_clzll(high);
                len = 8 + (lz >> 3);
                escaped = true;
            }
        } else {
            unsigned lz = (unsigned)__builtin_clzll(low);
            len = lz >> 3;
            escaped = true;
        }
        if (CEREAL_RAPIDJSON_UNLIKELY(escaped)) {   // some of characters is escaped
            char* q = reinterpret_cast<char*>(os_->PushUnsafe(len));
            for (size_t i = 0; i < len; i++)
                q[i] = p[i];

            p += len;
            break;
        }
        vst1q_u8(reinterpret_cast<uint8_t *>(os_->PushUnsafe(16)), s);
    }

    is.src_ = p;
    return CEREAL_RAPIDJSON_LIKELY(is.Tell() < length);
}
#endif // CEREAL_RAPIDJSON_NEON

CEREAL_RAPIDJSON_NAMESPACE_END

#if defined(_MSC_VER) || defined(__clang__)
CEREAL_RAPIDJSON_DIAG_POP
#endif

#endif // CEREAL_RAPIDJSON_CEREAL_RAPIDJSON_H_
```
