## Supplemental code for "The motif composition of variable-number tandem repeats impacts gene expression": license.txt

Use of this software is granted under one of the following two licenses,
to be chosen freely by the user.

1. Boost Software License - Version 1.0 - August 17th, 2003
===============================================================================

Copyright (c) 2006, 2007 Marcin Kalicinski

Permission is hereby granted, free of charge, to any person or organization
obtaining a copy of the software and accompanying documentation covered by
this license (the "Software") to use, reproduce, display, distribute,
execute, and transmit the Software, and to prepare derivative works of the
Software, and to permit third-parties to whom the Software is furnished to
do so, all subject to the following:

The copyright notices in the Software and this entire statement, including
the above license grant, this restriction and the following disclaimer,
must be included in all copies of the Software, in whole or in part, and
all derivative works of the Software, unless such copies or derivative
works are solely in the form of machine-executable object code generated by
a source language processor.

THE SOFTWARE IS PROVIDED "AS IS", WITHOUT WARRANTY OF ANY KIND, EXPRESS OR
IMPLIED, INCLUDING BUT NOT LIMITED TO THE WARRANTIES OF MERCHANTABILITY,
FITNESS FOR A PARTICULAR PURPOSE, TITLE AND NON-INFRINGEMENT. IN NO EVENT
SHALL THE COPYRIGHT HOLDERS OR ANYONE DISTRIBUTING THE SOFTWARE BE LIABLE
FOR ANY DAMAGES OR OTHER LIABILITY, WHETHER IN CONTRACT, TORT OR OTHERWISE,
ARISING FROM, OUT OF OR IN CONNECTION WITH THE SOFTWARE OR THE USE OR OTHER
DEALINGS IN THE SOFTWARE.

2. The MIT License
===============================================================================

Copyright (c) 2006, 2007 Marcin Kalicinski

Permission is hereby granted, free of charge, to any person obtaining a copy 
of this software and associated documentation files (the "Software"), to deal 
in the Software without restriction, including without limitation the rights 
to use, copy, modify, merge, publish, distribute, sublicense, and/or sell copies 
of the Software, and to permit persons to whom the Software is furnished to do so, 
subject to the following conditions:

The above copyright notice and this permission notice shall be included in all 
copies or substantial portions of the Software.

THE SOFTWARE IS PROVIDED "AS IS", WITHOUT WARRANTY OF ANY KIND, EXPRESS OR 
IMPLIED, INCLUDING BUT NOT LIMITED TO THE WARRANTIES OF MERCHANTABILITY, 
FITNESS FOR A PARTICULAR PURPOSE AND NONINFRINGEMENT. IN NO EVENT SHALL 
THE AUTHORS OR COPYRIGHT HOLDERS BE LIABLE FOR ANY CLAIM, DAMAGES OR OTHER 
LIABILITY, WHETHER IN AN ACTION OF CONTRACT, TORT OR OTHERWISE, ARISING FROM, 
OUT OF OR IN CONNECTION WITH THE SOFTWARE OR THE USE OR OTHER DEALINGS 
IN THE SOFTWARE.
