## Supplemental code for "The motif composition of variable-number tandem repeats impacts gene expression": manual.html

 

### RAPIDXML Manual

##### Version 1.13

*Copyright (C) 2006, 2009 Marcin Kalicinski*  
*See accompanying file license.txt for license information.*

---

#### Table of Contents

1. What is RapidXml?  
1.1 Dependencies And Compatibility  
1.2 Character Types And Encodings  
1.3 Error Handling  
1.4 Memory Allocation  
1.5 W3C Compliance  
1.6 API Design  
1.7 Reliability  
1.8 Acknowledgements  
2. Two Minute Tutorial  
2.1 Parsing  
2.2 Accessing The DOM Tree  
2.3 Modifying The DOM Tree  
2.4 Printing XML  
3. Differences From Regular XML Parsers  
3.1 Lifetime Of Source Text  
3.2 Ownership Of Strings  
3.3 Destructive Vs Non-Destructive Mode  
4. Performance  
4.1 Comparison With Other Parsers  
5. Reference  
  

#### 1. What is RapidXml?

RapidXml is an attempt to create the fastest XML DOM parser possible, while retaining useability, portability and reasonable W3C compatibility. It is an in-situ parser written in C++, with parsing speed approaching that of `strlen()` function executed on the same data.   
  
Entire parser is contained in a single header file, so no building or linking is neccesary. To use it you just need to copy `rapidxml.hpp` file to a convenient place (such as your project directory), and include it where needed. You may also want to use printing functions contained in header `rapidxml_print.hpp`.

##### 1.1 Dependencies And Compatibility

RapidXml has *no dependencies* other than a very small subset of standard C++ library (`<cassert>`, `<cstdlib>`, `<new>` and `<exception>`, unless exceptions are disabled). It should compile on any reasonably conformant compiler, and was tested on Visual C++ 2003, Visual C++ 2005, Visual C++ 2008, gcc 3, gcc 4, and Comeau 4.3.3. Care was taken that no warnings are produced on these compilers, even with highest warning levels enabled.

##### 1.2 Character Types And Encodings

RapidXml is character type agnostic, and can work both with narrow and wide characters. Current version does not fully support UTF-16 or UTF-32, so use of wide characters is somewhat incapacitated. However, it should succesfully parse `wchar_t` strings containing UTF-16 or UTF-32 if endianness of the data matches that of the machine. UTF-8 is fully supported, including all numeric character references, which are expanded into appropriate UTF-8 byte sequences (unless you enable parse\_no\_utf8 flag).   
  
Note that RapidXml performs no decoding - strings returned by name() and value() functions will contain text encoded using the same encoding as source file. Rapidxml understands and expands the following character references: `&apos; &amp; &quot; &lt; &gt; &#...;` Other character references are not expanded.

##### 1.3 Error Handling

By default, RapidXml uses C++ exceptions to report errors. If this behaviour is undesirable, RAPIDXML\_NO\_EXCEPTIONS can be defined to suppress exception code. See parse\_error class and parse\_error\_handler() function for more information.

##### 1.4 Memory Allocation

RapidXml uses a special memory pool object to allocate nodes and attributes, because direct allocation using `new` operator would be far too slow. Underlying memory allocations performed by the pool can be customized by use of memory\_pool::set\_allocator() function. See class memory\_pool for more information.

##### 1.5 W3C Compliance

RapidXml is not a W3C compliant parser, primarily because it ignores DOCTYPE declarations. There is a number of other, minor incompatibilities as well. Still, it can successfully parse and produce complete trees of all valid XML files in W3C conformance suite (over 1000 files specially designed to find flaws in XML processors). In destructive mode it performs whitespace normalization and character entity substitution for a small set of built-in entities.

##### 1.6 API Design

RapidXml API is minimalistic, to reduce code size as much as possible, and facilitate use in embedded environments. Additional convenience functions are provided in separate headers: `rapidxml_utils.hpp` and `rapidxml_print.hpp`. Contents of these headers is not an essential part of the library, and is currently not documented (otherwise than with comments in code).

##### 1.7 Reliability

RapidXml is very robust and comes with a large harness of unit tests. Special care has been taken to ensure stability of the parser no matter what source text is thrown at it. One of the unit tests produces 100,000 randomly corrupted variants of XML document, which (when uncorrupted) contains all constructs recognized by RapidXml. RapidXml passes this test when it correctly recognizes that errors have been introduced, and does not crash or loop indefinitely.   
  
Another unit test puts RapidXml head-to-head with another, well estabilished XML parser, and verifies that their outputs match across a wide variety of small and large documents.   
  
Yet another test feeds RapidXml with over 1000 test files from W3C compliance suite, and verifies that correct results are obtained. There are also additional tests that verify each API function separately, and test that various parsing modes work as expected.

##### 1.8 Acknowledgements

I would like to thank Arseny Kapoulkine for his work on pugixml, which was an inspiration for this project. Additional thanks go to Kristen Wegner for creating pugxml, from which pugixml was derived. Janusz Wohlfeil kindly ran RapidXml speed tests on hardware that I did not have access to, allowing me to expand performance comparison table.

#### 2. Two Minute Tutorial

##### 2.1 Parsing

The following code causes RapidXml to parse a zero-terminated string named `text`:

```
using namespace rapidxml;
xml_document<> doc;    // character type defaults to char
doc.parse<0>(text);    // 0 means default parse flags
```

`doc` object is now a root of DOM tree containing representation of the parsed XML. Because all RapidXml interface is contained inside namespace `rapidxml`, users must either bring contents of this namespace into scope, or fully qualify all the names. Class xml\_document represents a root of the DOM hierarchy. By means of public inheritance, it is also an xml\_node and a memory\_pool. Template parameter of xml\_document::parse() function is used to specify parsing flags, with which you can fine-tune behaviour of the parser. Note that flags must be a compile-time constant.

##### 2.2 Accessing The DOM Tree

To access the DOM tree, use methods of xml\_node and xml\_attribute classes:

```
cout << "Name of my first node is: " << doc.first_node()->name() << "\n";
xml_node<> *node = doc.first_node("foobar");
cout << "Node foobar has value " << node->value() << "\n";
for (xml_attribute<> *attr = node->first_attribute();
     attr; attr = attr->next_attribute())
{
    cout << "Node foobar has attribute " << attr->name() << " ";
    cout << "with value " << attr->value() << "\n";
}
```

##### 2.3 Modifying The DOM Tree

DOM tree produced by the parser is fully modifiable. Nodes and attributes can be added/removed, and their contents changed. The below example creates a HTML document, whose sole contents is a link to google.com website:

```
xml_document<> doc;
xml_node<> *node = doc.allocate_node(node_element, "a", "Google");
doc.append_node(node);
xml_attribute<> *attr = doc.allocate_attribute("href", "google.com");
node->append_attribute(attr);
```

One quirk is that nodes and attributes *do not own* the text of their names and values. This is because normally they only store pointers to the source text. So, when assigning a new name or value to the node, care must be taken to ensure proper lifetime of the string. The easiest way to achieve it is to allocate the string from the xml\_document memory pool. In the above example this is not necessary, because we are only assigning character constants. But the code below uses memory\_pool::allocate\_string() function to allocate node name (which will have the same lifetime as the document), and assigns it to a new node:

```
xml_document<> doc;
char *node_name = doc.allocate_string(name);        // Allocate string and copy name into it
xml_node<> *node = doc.allocate_node(node_element, node_name);  // Set node name to node_name
```

Check Reference section for description of the entire interface.

##### 2.4 Printing XML

You can print `xml_document` and `xml_node` objects into an XML string. Use print() function or operator <<, which are defined in `rapidxml_print.hpp` header.

```
using namespace rapidxml;
xml_document<> doc;    // character type defaults to char
// ... some code to fill the document

// Print to stream using operator <<
std::cout << doc;   

// Print to stream using print function, specifying printing flags
print(std::cout, doc, 0);   // 0 means default printing flags

// Print to string using output iterator
std::string s;
print(std::back_inserter(s), doc, 0);

// Print to memory buffer using output iterator
char buffer[4096];                      // You are responsible for making the buffer large enough!
char *end = print(buffer, doc, 0);      // end contains pointer to character after last printed character
*end = 0;                               // Add string terminator after XML
```

#### 3. Differences From Regular XML Parsers

RapidXml is an *in-situ parser*, which allows it to achieve very high parsing speed. In-situ means that parser does not make copies of strings. Instead, it places pointers to the *source text* in the DOM hierarchy.

##### 3.1 Lifetime Of Source Text

In-situ parsing requires that source text lives at least as long as the document object. If source text is destroyed, names and values of nodes in DOM tree will become destroyed as well. Additionally, whitespace processing, character entity translation, and zero-termination of strings require that source text be modified during parsing (but see non-destructive mode). This makes the text useless for further processing once it was parsed by RapidXml.   
  
In many cases however, these are not serious issues.

##### 3.2 Ownership Of Strings

Nodes and attributes produced by RapidXml do not own their name and value strings. They merely hold the pointers to them. This means you have to be careful when setting these values manually, by using xml\_base::name(const Ch \*) or xml\_base::value(const Ch \*) functions. Care must be taken to ensure that lifetime of the string passed is at least as long as lifetime of the node/attribute. The easiest way to achieve it is to allocate the string from memory\_pool owned by the document. Use memory\_pool::allocate\_string() function for this purpose.

##### 3.3 Destructive Vs Non-Destructive Mode

By default, the parser modifies source text during the parsing process. This is required to achieve character entity translation, whitespace normalization, and zero-termination of strings.   
  
In some cases this behaviour may be undesirable, for example if source text resides in read only memory, or is mapped to memory directly from file. By using appropriate parser flags (parse\_non\_destructive), source text modifications can be disabled. However, because RapidXml does in-situ parsing, it obviously has the following side-effects:

- no whitespace normalization is done
- no entity reference translation is done
- names and values are not zero-terminated, you must use xml\_base::name\_size() and xml\_base::value\_size() functions to tell where they end

#### 4. Performance

RapidXml achieves its speed through use of several techniques:

- In-situ parsing. When building DOM tree, RapidXml does not make copies of string data, such as node names and values. Instead, it stores pointers to interior of the source text.
- Use of template metaprogramming techniques. This allows it to move much of the work to compile time. Through magic of the templates, C++ compiler generates a separate copy of parsing code for any combination of parser flags you use. In each copy, all possible decisions are made at compile time and all unused code is omitted.
- Extensive use of lookup tables for parsing.
- Hand-tuned C++ with profiling done on several most popular CPUs.
This results in a very small and fast code: a parser which is custom tailored to exact needs with each invocation.

##### 4.1 Comparison With Other Parsers

The table below compares speed of RapidXml to some other parsers, and to `strlen()` function executed on the same data. On a modern CPU (as of 2007), you can expect parsing throughput to be close to 1 GB/s. As a rule of thumb, parsing speed is about 50-100x faster than Xerces DOM, 30-60x faster than TinyXml, 3-12x faster than pugxml, and about 5% - 30% faster than pugixml, the fastest XML parser I know of.

- The test file is a real-world, 50kB large, moderately dense XML file.
- All timing is done by using RDTSC instruction present in Pentium-compatible CPUs.
- No profile-guided optimizations are used.
- All parsers are running in their fastest modes.
- The results are given in CPU cycles per character, so frequency of CPUs is irrelevant.
- The results are minimum values from a large number of runs, to minimize effects of operating system activity, task switching, interrupt handling etc.
- A single parse of the test file takes about 1/10th of a millisecond, so with large number of runs there is a good chance of hitting at least one no-interrupt streak, and obtaining undisturbed results.

| Platform | Compiler | strlen() | RapidXml | pugixml 0.3 | pugxml | TinyXml |
| --- | --- | --- | --- | --- | --- | --- |
| Pentium 4 | MSVC 8.0 | 2.5 | 5.4 | 7.0 | 61.7 | 298.8 |
| Pentium 4 | gcc 4.1.1 | 0.8 | 6.1 | 9.5 | 67.0 | 413.2 |
| Core 2 | MSVC 8.0 | 1.0 | 4.5 | 5.0 | 24.6 | 154.8 |
| Core 2 | gcc 4.1.1 | 0.6 | 4.6 | 5.4 | 28.3 | 229.3 |
| Athlon XP | MSVC 8.0 | 3.1 | 7.7 | 8.0 | 25.5 | 182.6 |
| Athlon XP | gcc 4.1.1 | 0.9 | 8.2 | 9.2 | 33.7 | 265.2 |
| Pentium 3 | MSVC 8.0 | 2.0 | 6.3 | 7.0 | 30.9 | 211.9 |
| Pentium 3 | gcc 4.1.1 | 1.0 | 6.7 | 8.9 | 35.3 | 316.0 |

*(\*) All results are in CPU cycles per character of source text*

#### 5. Reference

This section lists all classes, functions, constants etc. and describes them in detail. 

class template rapidxml::memory\_pool

constructor memory\_pool()

destructor ~memory\_pool()

function allocate\_node(node\_type type, const Ch \*name=0, const Ch \*value=0, std::size\_t name\_size=0, std::size\_t value\_size=0)

function allocate\_attribute(const Ch \*name=0, const Ch \*value=0, std::size\_t name\_size=0, std::size\_t value\_size=0)

function allocate\_string(const Ch \*source=0, std::size\_t size=0)

function clone\_node(const xml\_node< Ch > \*source, xml\_node< Ch > \*result=0)

function clear()

function set\_allocator(alloc\_func \*af, free\_func \*ff)


class rapidxml::parse\_error

constructor parse\_error(const char \*what, void \*where)

function what() const

function where() const


class template rapidxml::xml\_attribute

constructor xml\_attribute()

function document() const

function previous\_attribute(const Ch \*name=0, std::size\_t name\_size=0, bool case\_sensitive=true) const

function next\_attribute(const Ch \*name=0, std::size\_t name\_size=0, bool case\_sensitive=true) const


class template rapidxml::xml\_base

constructor xml\_base()

function name() const

function name\_size() const

function value() const

function value\_size() const

function name(const Ch \*name, std::size\_t size)

function name(const Ch \*name)

function value(const Ch \*value, std::size\_t size)

function value(const Ch \*value)

function parent() const


class template rapidxml::xml\_document

constructor xml\_document()

function parse(Ch \*text)

function clear()


class template rapidxml::xml\_node

constructor xml\_node(node\_type type)

function type() const

function document() const

function first\_node(const Ch \*name=0, std::size\_t name\_size=0, bool case\_sensitive=true) const

function last\_node(const Ch \*name=0, std::size\_t name\_size=0, bool case\_sensitive=true) const

function previous\_sibling(const Ch \*name=0, std::size\_t name\_size=0, bool case\_sensitive=true) const

function next\_sibling(const Ch \*name=0, std::size\_t name\_size=0, bool case\_sensitive=true) const

function first\_attribute(const Ch \*name=0, std::size\_t name\_size=0, bool case\_sensitive=true) const

function last\_attribute(const Ch \*name=0, std::size\_t name\_size=0, bool case\_sensitive=true) const

function type(node\_type type)

function prepend\_node(xml\_node< Ch > \*child)

function append\_node(xml\_node< Ch > \*child)

function insert\_node(xml\_node< Ch > \*where, xml\_node< Ch > \*child)

function remove\_first\_node()

function remove\_last\_node()

function remove\_node(xml\_node< Ch > \*where)

function remove\_all\_nodes()

function prepend\_attribute(xml\_attribute< Ch > \*attribute)

function append\_attribute(xml\_attribute< Ch > \*attribute)

function insert\_attribute(xml\_attribute< Ch > \*where, xml\_attribute< Ch > \*attribute)

function remove\_first\_attribute()

function remove\_last\_attribute()

function remove\_attribute(xml\_attribute< Ch > \*where)

function remove\_all\_attributes()


namespace rapidxml

enum node\_type


function parse\_error\_handler(const char \*what, void \*where)

function print(OutIt out, const xml\_node< Ch > &node, int flags=0)

function print(std::basic\_ostream< Ch > &out, const xml\_node< Ch > &node, int flags=0)

function operator<<(std::basic\_ostream< Ch > &out, const xml\_node< Ch > &node)

constant parse\_no\_data\_nodes

constant parse\_no\_element\_values

constant parse\_no\_string\_terminators

constant parse\_no\_entity\_translation

constant parse\_no\_utf8

constant parse\_declaration\_node

constant parse\_comment\_nodes

constant parse\_doctype\_node

constant parse\_pi\_nodes

constant parse\_validate\_closing\_tags

constant parse\_trim\_whitespace

constant parse\_normalize\_whitespace

constant parse\_default

constant parse\_non\_destructive

constant parse\_fastest

constant parse\_full

constant print\_no\_indenting

---

##### class template rapidxml::memory\_pool

Defined in rapidxml.hpp  
Base class for
xml\_document

###### Description

This class is used by the parser to create new nodes and attributes, without overheads of dynamic memory allocation. In most cases, you will not need to use this class directly. However, if you need to create nodes manually or modify names/values of nodes, you are encouraged to use memory\_pool of relevant xml\_document to allocate the memory. Not only is this faster than allocating them by using `new` operator, but also their lifetime will be tied to the lifetime of document, possibly simplyfing memory management.   
  
Call allocate\_node() or allocate\_attribute() functions to obtain new nodes or attributes from the pool. You can also call allocate\_string() function to allocate strings. Such strings can then be used as names or values of nodes without worrying about their lifetime. Note that there is no `free()` function -- all allocations are freed at once when clear() function is called, or when the pool is destroyed.   
  
It is also possible to create a standalone memory\_pool, and use it to allocate nodes, whose lifetime will not be tied to any document.   
  
Pool maintains `RAPIDXML_STATIC_POOL_SIZE` bytes of statically allocated memory. Until static memory is exhausted, no dynamic memory allocations are done. When static memory is exhausted, pool allocates additional blocks of memory of size `RAPIDXML_DYNAMIC_POOL_SIZE` each, by using global `new[]` and `delete[]` operators. This behaviour can be changed by setting custom allocation routines. Use set\_allocator() function to set them.   
  
Allocations for nodes, attributes and strings are aligned at `RAPIDXML_ALIGNMENT` bytes. This value defaults to the size of pointer on target architecture.   
  
To obtain absolutely top performance from the parser, it is important that all nodes are allocated from a single, contiguous block of memory. Otherwise, cache misses when jumping between two (or more) disjoint blocks of memory can slow down parsing quite considerably. If required, you can tweak `RAPIDXML_STATIC_POOL_SIZE`, `RAPIDXML_DYNAMIC_POOL_SIZE` and `RAPIDXML_ALIGNMENT` to obtain best wasted memory to performance compromise. To do it, define their values before rapidxml.hpp file is included. 

###### Parameters

Ch
:   Character type of created nodes.

##### constructor memory\_pool::memory\_pool

###### Synopsis

`memory_pool();` 

###### Description

Constructs empty pool with default allocator functions. 

##### destructor memory\_pool::~memory\_pool

###### Synopsis

`~memory_pool();` 

###### Description

Destroys pool and frees all the memory. This causes memory occupied by nodes allocated by the pool to be freed. Nodes allocated from the pool are no longer valid. 

##### function memory\_pool::allocate\_node

###### Synopsis

`xml_node<Ch>* allocate_node(node_type type, const Ch *name=0, const Ch *value=0, std::size_t name_size=0, std::size_t value_size=0);` 

###### Description

Allocates a new node from the pool, and optionally assigns name and value to it. If the allocation request cannot be accomodated, this function will throw `std::bad_alloc`. If exceptions are disabled by defining RAPIDXML\_NO\_EXCEPTIONS, this function will call rapidxml::parse\_error\_handler() function. 

###### Parameters

type
:   Type of node to create.

name
:   Name to assign to the node, or 0 to assign no name.

value
:   Value to assign to the node, or 0 to assign no value.

name\_size
:   Size of name to assign, or 0 to automatically calculate size from name string.

value\_size
:   Size of value to assign, or 0 to automatically calculate size from value string.

###### Returns

Pointer to allocated node. This pointer will never be NULL.

##### function memory\_pool::allocate\_attribute

###### Synopsis

`xml_attribute<Ch>* allocate_attribute(const Ch *name=0, const Ch *value=0, std::size_t name_size=0, std::size_t value_size=0);` 

###### Description

Allocates a new attribute from the pool, and optionally assigns name and value to it. If the allocation request cannot be accomodated, this function will throw `std::bad_alloc`. If exceptions are disabled by defining RAPIDXML\_NO\_EXCEPTIONS, this function will call rapidxml::parse\_error\_handler() function. 

###### Parameters

name
:   Name to assign to the attribute, or 0 to assign no name.

value
:   Value to assign to the attribute, or 0 to assign no value.

name\_size
:   Size of name to assign, or 0 to automatically calculate size from name string.

value\_size
:   Size of value to assign, or 0 to automatically calculate size from value string.

###### Returns

Pointer to allocated attribute. This pointer will never be NULL.

##### function memory\_pool::allocate\_string

###### Synopsis

`Ch* allocate_string(const Ch *source=0, std::size_t size=0);` 

###### Description

Allocates a char array of given size from the pool, and optionally copies a given string to it. If the allocation request cannot be accomodated, this function will throw `std::bad_alloc`. If exceptions are disabled by defining RAPIDXML\_NO\_EXCEPTIONS, this function will call rapidxml::parse\_error\_handler() function. 

###### Parameters

source
:   String to initialize the allocated memory with, or 0 to not initialize it.

size
:   Number of characters to allocate, or zero to calculate it automatically from source string length; if size is 0, source string must be specified and null terminated.

###### Returns

Pointer to allocated char array. This pointer will never be NULL.

##### function memory\_pool::clone\_node

###### Synopsis

`xml_node<Ch>* clone_node(const xml_node< Ch > *source, xml_node< Ch > *result=0);` 

###### Description

Clones an xml\_node and its hierarchy of child nodes and attributes. Nodes and attributes are allocated from this memory pool. Names and values are not cloned, they are shared between the clone and the source. Result node can be optionally specified as a second parameter, in which case its contents will be replaced with cloned source node. This is useful when you want to clone entire document. 

###### Parameters

source
:   Node to clone.

result
:   Node to put results in, or 0 to automatically allocate result node

###### Returns

Pointer to cloned node. This pointer will never be NULL.

##### function memory\_pool::clear

###### Synopsis

`void clear();` 

###### Description

Clears the pool. This causes memory occupied by nodes allocated by the pool to be freed. Any nodes or strings allocated from the pool will no longer be valid. 

##### function memory\_pool::set\_allocator

###### Synopsis

`void set_allocator(alloc_func *af, free_func *ff);` 

###### Description

Sets or resets the user-defined memory allocation functions for the pool. This can only be called when no memory is allocated from the pool yet, otherwise results are undefined. Allocation function must not return invalid pointer on failure. It should either throw, stop the program, or use `longjmp()` function to pass control to other place of program. If it returns invalid pointer, results are undefined.   
  
User defined allocation functions must have the following forms:   
 `void *allocate(std::size_t size);   
void free(void *pointer);`   

###### Parameters

af
:   Allocation function, or 0 to restore default function

ff
:   Free function, or 0 to restore default function

##### class rapidxml::parse\_error

Defined in rapidxml.hpp  

###### Description

Parse error exception. This exception is thrown by the parser when an error occurs. Use what() function to get human-readable error message. Use where() function to get a pointer to position within source text where error was detected.   
  
If throwing exceptions by the parser is undesirable, it can be disabled by defining RAPIDXML\_NO\_EXCEPTIONS macro before rapidxml.hpp is included. This will cause the parser to call rapidxml::parse\_error\_handler() function instead of throwing an exception. This function must be defined by the user.   
  
This class derives from `std::exception` class. 

##### constructor parse\_error::parse\_error

###### Synopsis

`parse_error(const char *what, void *where);` 

###### Description

Constructs parse error. 

##### function parse\_error::what

###### Synopsis

`virtual const char* what() const;` 

###### Description

Gets human readable description of error. 

###### Returns

Pointer to null terminated description of the error.

##### function parse\_error::where

###### Synopsis

`Ch* where() const;` 

###### Description

Gets pointer to character data where error happened. Ch should be the same as char type of xml\_document that produced the error. 

###### Returns

Pointer to location within the parsed string where error occured.

##### class template rapidxml::xml\_attribute

Defined in rapidxml.hpp  
Inherits from
xml\_base   

###### Description

Class representing attribute node of XML document. Each attribute has name and value strings, which are available through name() and value() functions (inherited from xml\_base). Note that after parse, both name and value of attribute will point to interior of source text used for parsing. Thus, this text must persist in memory for the lifetime of attribute. 

###### Parameters

Ch
:   Character type to use.

##### constructor xml\_attribute::xml\_attribute

###### Synopsis

`xml_attribute();` 

###### Description

Constructs an empty attribute with the specified type. Consider using memory\_pool of appropriate xml\_document if allocating attributes manually. 

##### function xml\_attribute::document

###### Synopsis

`xml_document<Ch>* document() const;` 

###### Description

Gets document of which attribute is a child. 

###### Returns

Pointer to document that contains this attribute, or 0 if there is no parent document.

##### function xml\_attribute::previous\_attribute

###### Synopsis

`xml_attribute<Ch>* previous_attribute(const Ch *name=0, std::size_t name_size=0, bool case_sensitive=true) const;` 

###### Description

Gets previous attribute, optionally matching attribute name. 

###### Parameters

name
:   Name of attribute to find, or 0 to return previous attribute regardless of its name; this string doesn't have to be zero-terminated if name\_size is non-zero

name\_size
:   Size of name, in characters, or 0 to have size calculated automatically from string

case\_sensitive
:   Should name comparison be case-sensitive; non case-sensitive comparison works properly only for ASCII characters

###### Returns

Pointer to found attribute, or 0 if not found.

##### function xml\_attribute::next\_attribute

###### Synopsis

`xml_attribute<Ch>* next_attribute(const Ch *name=0, std::size_t name_size=0, bool case_sensitive=true) const;` 

###### Description

Gets next attribute, optionally matching attribute name. 

###### Parameters

name
:   Name of attribute to find, or 0 to return next attribute regardless of its name; this string doesn't have to be zero-terminated if name\_size is non-zero

name\_size
:   Size of name, in characters, or 0 to have size calculated automatically from string

case\_sensitive
:   Should name comparison be case-sensitive; non case-sensitive comparison works properly only for ASCII characters

###### Returns

Pointer to found attribute, or 0 if not found.

##### class template rapidxml::xml\_base

Defined in rapidxml.hpp  
Base class for
xml\_attribute xml\_node

###### Description

Base class for xml\_node and xml\_attribute implementing common functions: name(), name\_size(), value(), value\_size() and parent(). 

###### Parameters

Ch
:   Character type to use

##### constructor xml\_base::xml\_base

###### Synopsis

`xml_base();` 

##### function xml\_base::name

###### Synopsis

`Ch* name() const;` 

###### Description

Gets name of the node. Interpretation of name depends on type of node. Note that name will not be zero-terminated if rapidxml::parse\_no\_string\_terminators option was selected during parse.   
  
Use name\_size() function to determine length of the name. 

###### Returns

Name of node, or empty string if node has no name.

##### function xml\_base::name\_size

###### Synopsis

`std::size_t name_size() const;` 

###### Description

Gets size of node name, not including terminator character. This function works correctly irrespective of whether name is or is not zero terminated. 

###### Returns

Size of node name, in characters.

##### function xml\_base::value

###### Synopsis

`Ch* value() const;` 

###### Description

Gets value of node. Interpretation of value depends on type of node. Note that value will not be zero-terminated if rapidxml::parse\_no\_string\_terminators option was selected during parse.   
  
Use value\_size() function to determine length of the value. 

###### Returns

Value of node, or empty string if node has no value.

##### function xml\_base::value\_size

###### Synopsis

`std::size_t value_size() const;` 

###### Description

Gets size of node value, not including terminator character. This function works correctly irrespective of whether value is or is not zero terminated. 

###### Returns

Size of node value, in characters.

##### function xml\_base::name

###### Synopsis

`void name(const Ch *name, std::size_t size);` 

###### Description

Sets name of node to a non zero-terminated string. See Ownership Of Strings .   
  
Note that node does not own its name or value, it only stores a pointer to it. It will not delete or otherwise free the pointer on destruction. It is reponsibility of the user to properly manage lifetime of the string. The easiest way to achieve it is to use memory\_pool of the document to allocate the string - on destruction of the document the string will be automatically freed.   
  
Size of name must be specified separately, because name does not have to be zero terminated. Use name(const Ch \*) function to have the length automatically calculated (string must be zero terminated). 

###### Parameters

name
:   Name of node to set. Does not have to be zero terminated.

size
:   Size of name, in characters. This does not include zero terminator, if one is present.

##### function xml\_base::name

###### Synopsis

`void name(const Ch *name);` 

###### Description

Sets name of node to a zero-terminated string. See also Ownership Of Strings and xml\_node::name(const Ch \*, std::size\_t). 

###### Parameters

name
:   Name of node to set. Must be zero terminated.

##### function xml\_base::value

###### Synopsis

`void value(const Ch *value, std::size_t size);` 

###### Description

Sets value of node to a non zero-terminated string. See Ownership Of Strings .   
  
Note that node does not own its name or value, it only stores a pointer to it. It will not delete or otherwise free the pointer on destruction. It is reponsibility of the user to properly manage lifetime of the string. The easiest way to achieve it is to use memory\_pool of the document to allocate the string - on destruction of the document the string will be automatically freed.   
  
Size of value must be specified separately, because it does not have to be zero terminated. Use value(const Ch \*) function to have the length automatically calculated (string must be zero terminated).   
  
If an element has a child node of type node\_data, it will take precedence over element value when printing. If you want to manipulate data of elements using values, use parser flag rapidxml::parse\_no\_data\_nodes to prevent creation of data nodes by the parser. 

###### Parameters

value
:   value of node to set. Does not have to be zero terminated.

size
:   Size of value, in characters. This does not include zero terminator, if one is present.

##### function xml\_base::value

###### Synopsis

`void value(const Ch *value);` 

###### Description

Sets value of node to a zero-terminated string. See also Ownership Of Strings and xml\_node::value(const Ch \*, std::size\_t). 

###### Parameters

value
:   Vame of node to set. Must be zero terminated.

##### function xml\_base::parent

###### Synopsis

`xml_node<Ch>* parent() const;` 

###### Description

Gets node parent. 

###### Returns

Pointer to parent node, or 0 if there is no parent.

##### class template rapidxml::xml\_document

Defined in rapidxml.hpp  
Inherits from
xml\_node memory\_pool   

###### Description

This class represents root of the DOM hierarchy. It is also an xml\_node and a memory\_pool through public inheritance. Use parse() function to build a DOM tree from a zero-terminated XML text string. parse() function allocates memory for nodes and attributes by using functions of xml\_document, which are inherited from memory\_pool. To access root node of the document, use the document itself, as if it was an xml\_node. 

###### Parameters

Ch
:   Character type to use.

##### constructor xml\_document::xml\_document

###### Synopsis

`xml_document();` 

###### Description

Constructs empty XML document. 

##### function xml\_document::parse

###### Synopsis

`void parse(Ch *text);` 

###### Description

Parses zero-terminated XML string according to given flags. Passed string will be modified by the parser, unless rapidxml::parse\_non\_destructive flag is used. The string must persist for the lifetime of the document. In case of error, rapidxml::parse\_error exception will be thrown.   
  
If you want to parse contents of a file, you must first load the file into the memory, and pass pointer to its beginning. Make sure that data is zero-terminated.   
  
Document can be parsed into multiple times. Each new call to parse removes previous nodes and attributes (if any), but does not clear memory pool. 

###### Parameters

text
:   XML data to parse; pointer is non-const to denote fact that this data may be modified by the parser.

##### function xml\_document::clear

###### Synopsis

`void clear();` 

###### Description

Clears the document by deleting all nodes and clearing the memory pool. All nodes owned by document pool are destroyed. 

##### class template rapidxml::xml\_node

Defined in rapidxml.hpp  
Inherits from
xml\_base   
Base class for
xml\_document

###### Description

Class representing a node of XML document. Each node may have associated name and value strings, which are available through name() and value() functions. Interpretation of name and value depends on type of the node. Type of node can be determined by using type() function.   
  
Note that after parse, both name and value of node, if any, will point interior of source text used for parsing. Thus, this text must persist in the memory for the lifetime of node. 

###### Parameters

Ch
:   Character type to use.

##### constructor xml\_node::xml\_node

###### Synopsis

`xml_node(node_type type);` 

###### Description

Constructs an empty node with the specified type. Consider using memory\_pool of appropriate document to allocate nodes manually. 

###### Parameters

type
:   Type of node to construct.

##### function xml\_node::type

###### Synopsis

`node_type type() const;` 

###### Description

Gets type of node. 

###### Returns

Type of node.

##### function xml\_node::document

###### Synopsis

`xml_document<Ch>* document() const;` 

###### Description

Gets document of which node is a child. 

###### Returns

Pointer to document that contains this node, or 0 if there is no parent document.

##### function xml\_node::first\_node

###### Synopsis

`xml_node<Ch>* first_node(const Ch *name=0, std::size_t name_size=0, bool case_sensitive=true) const;` 

###### Description

Gets first child node, optionally matching node name. 

###### Parameters

name
:   Name of child to find, or 0 to return first child regardless of its name; this string doesn't have to be zero-terminated if name\_size is non-zero

name\_size
:   Size of name, in characters, or 0 to have size calculated automatically from string

case\_sensitive
:   Should name comparison be case-sensitive; non case-sensitive comparison works properly only for ASCII characters

###### Returns

Pointer to found child, or 0 if not found.

##### function xml\_node::last\_node

###### Synopsis

`xml_node<Ch>* last_node(const Ch *name=0, std::size_t name_size=0, bool case_sensitive=true) const;` 

###### Description

Gets last child node, optionally matching node name. Behaviour is undefined if node has no children. Use first\_node() to test if node has children. 

###### Parameters

name
:   Name of child to find, or 0 to return last child regardless of its name; this string doesn't have to be zero-terminated if name\_size is non-zero

name\_size
:   Size of name, in characters, or 0 to have size calculated automatically from string

case\_sensitive
:   Should name comparison be case-sensitive; non case-sensitive comparison works properly only for ASCII characters

###### Returns

Pointer to found child, or 0 if not found.

##### function xml\_node::previous\_sibling

###### Synopsis

`xml_node<Ch>* previous_sibling(const Ch *name=0, std::size_t name_size=0, bool case_sensitive=true) const;` 

###### Description

Gets previous sibling node, optionally matching node name. Behaviour is undefined if node has no parent. Use parent() to test if node has a parent. 

###### Parameters

name
:   Name of sibling to find, or 0 to return previous sibling regardless of its name; this string doesn't have to be zero-terminated if name\_size is non-zero

name\_size
:   Size of name, in characters, or 0 to have size calculated automatically from string

case\_sensitive
:   Should name comparison be case-sensitive; non case-sensitive comparison works properly only for ASCII characters

###### Returns

Pointer to found sibling, or 0 if not found.

##### function xml\_node::next\_sibling

###### Synopsis

`xml_node<Ch>* next_sibling(const Ch *name=0, std::size_t name_size=0, bool case_sensitive=true) const;` 

###### Description

Gets next sibling node, optionally matching node name. Behaviour is undefined if node has no parent. Use parent() to test if node has a parent. 

###### Parameters

name
:   Name of sibling to find, or 0 to return next sibling regardless of its name; this string doesn't have to be zero-terminated if name\_size is non-zero

name\_size
:   Size of name, in characters, or 0 to have size calculated automatically from string

case\_sensitive
:   Should name comparison be case-sensitive; non case-sensitive comparison works properly only for ASCII characters

###### Returns

Pointer to found sibling, or 0 if not found.

##### function xml\_node::first\_attribute

###### Synopsis

`xml_attribute<Ch>* first_attribute(const Ch *name=0, std::size_t name_size=0, bool case_sensitive=true) const;` 

###### Description

Gets first attribute of node, optionally matching attribute name. 

###### Parameters

name
:   Name of attribute to find, or 0 to return first attribute regardless of its name; this string doesn't have to be zero-terminated if name\_size is non-zero

name\_size
:   Size of name, in characters, or 0 to have size calculated automatically from string

case\_sensitive
:   Should name comparison be case-sensitive; non case-sensitive comparison works properly only for ASCII characters

###### Returns

Pointer to found attribute, or 0 if not found.

##### function xml\_node::last\_attribute

###### Synopsis

`xml_attribute<Ch>* last_attribute(const Ch *name=0, std::size_t name_size=0, bool case_sensitive=true) const;` 

###### Description

Gets last attribute of node, optionally matching attribute name. 

###### Parameters

name
:   Name of attribute to find, or 0 to return last attribute regardless of its name; this string doesn't have to be zero-terminated if name\_size is non-zero

name\_size
:   Size of name, in characters, or 0 to have size calculated automatically from string

case\_sensitive
:   Should name comparison be case-sensitive; non case-sensitive comparison works properly only for ASCII characters

###### Returns

Pointer to found attribute, or 0 if not found.

##### function xml\_node::type

###### Synopsis

`void type(node_type type);` 

###### Description

Sets type of node. 

###### Parameters

type
:   Type of node to set.

##### function xml\_node::prepend\_node

###### Synopsis

`void prepend_node(xml_node< Ch > *child);` 

###### Description

Prepends a new child node. The prepended child becomes the first child, and all existing children are moved one position back. 

###### Parameters

child
:   Node to prepend.

##### function xml\_node::append\_node

###### Synopsis

`void append_node(xml_node< Ch > *child);` 

###### Description

Appends a new child node. The appended child becomes the last child. 

###### Parameters

child
:   Node to append.

##### function xml\_node::insert\_node

###### Synopsis

`void insert_node(xml_node< Ch > *where, xml_node< Ch > *child);` 

###### Description

Inserts a new child node at specified place inside the node. All children after and including the specified node are moved one position back. 

###### Parameters

where
:   Place where to insert the child, or 0 to insert at the back.

child
:   Node to insert.

##### function xml\_node::remove\_first\_node

###### Synopsis

`void remove_first_node();` 

###### Description

Removes first child node. If node has no children, behaviour is undefined. Use first\_node() to test if node has children. 

##### function xml\_node::remove\_last\_node

###### Synopsis

`void remove_last_node();` 

###### Description

Removes last child of the node. If node has no children, behaviour is undefined. Use first\_node() to test if node has children. 

##### function xml\_node::remove\_node

###### Synopsis

`void remove_node(xml_node< Ch > *where);` 

###### Description

Removes specified child from the node. 

##### function xml\_node::remove\_all\_nodes

###### Synopsis

`void remove_all_nodes();` 

###### Description

Removes all child nodes (but not attributes). 

##### function xml\_node::prepend\_attribute

###### Synopsis

`void prepend_attribute(xml_attribute< Ch > *attribute);` 

###### Description

Prepends a new attribute to the node. 

###### Parameters

attribute
:   Attribute to prepend.

##### function xml\_node::append\_attribute

###### Synopsis

`void append_attribute(xml_attribute< Ch > *attribute);` 

###### Description

Appends a new attribute to the node. 

###### Parameters

attribute
:   Attribute to append.

##### function xml\_node::insert\_attribute

###### Synopsis

`void insert_attribute(xml_attribute< Ch > *where, xml_attribute< Ch > *attribute);` 

###### Description

Inserts a new attribute at specified place inside the node. All attributes after and including the specified attribute are moved one position back. 

###### Parameters

where
:   Place where to insert the attribute, or 0 to insert at the back.

attribute
:   Attribute to insert.

##### function xml\_node::remove\_first\_attribute

###### Synopsis

`void remove_first_attribute();` 

###### Description

Removes first attribute of the node. If node has no attributes, behaviour is undefined. Use first\_attribute() to test if node has attributes. 

##### function xml\_node::remove\_last\_attribute

###### Synopsis

`void remove_last_attribute();` 

###### Description

Removes last attribute of the node. If node has no attributes, behaviour is undefined. Use first\_attribute() to test if node has attributes. 

##### function xml\_node::remove\_attribute

###### Synopsis

`void remove_attribute(xml_attribute< Ch > *where);` 

###### Description

Removes specified attribute from node. 

###### Parameters

where
:   Pointer to attribute to be removed.

##### function xml\_node::remove\_all\_attributes

###### Synopsis

`void remove_all_attributes();` 

###### Description

Removes all attributes of node. 

##### enum node\_type

###### Description

Enumeration listing all node types produced by the parser. Use xml\_node::type() function to query node type. 

###### Values

node\_document
:   A document node. Name and value are empty.

node\_element
:   An element node. Name contains element name. Value contains text of first data node.

node\_data
:   A data node. Name is empty. Value contains data text.

node\_cdata
:   A CDATA node. Name is empty. Value contains data text.

node\_comment
:   A comment node. Name is empty. Value contains comment text.

node\_declaration
:   A declaration node. Name and value are empty. Declaration parameters (version, encoding and standalone) are in node attributes.

node\_doctype
:   A DOCTYPE node. Name is empty. Value contains DOCTYPE text.

node\_pi
:   A PI node. Name contains target. Value contains instructions.

##### function parse\_error\_handler

###### Synopsis

`void rapidxml::parse_error_handler(const char *what, void *where);` 

###### Description

When exceptions are disabled by defining RAPIDXML\_NO\_EXCEPTIONS, this function is called to notify user about the error. It must be defined by the user.   
  
This function cannot return. If it does, the results are undefined.   
  
A very simple definition might look like that: 
void rapidxml::parse\_error\_handler(const char \*what, void \*where)
{
std::cout << "Parse error: " << what << "\n";
std::abort();
}

###### Parameters

what
:   Human readable description of the error.

where
:   Pointer to character data where error was detected.

##### function print

###### Synopsis

`OutIt rapidxml::print(OutIt out, const xml_node< Ch > &node, int flags=0);` 

###### Description

Prints XML to given output iterator. 

###### Parameters

out
:   Output iterator to print to.

node
:   Node to be printed. Pass xml\_document to print entire document.

flags
:   Flags controlling how XML is printed.

###### Returns

Output iterator pointing to position immediately after last character of printed text.

##### function print

###### Synopsis

`std::basic_ostream<Ch>& rapidxml::print(std::basic_ostream< Ch > &out, const xml_node< Ch > &node, int flags=0);` 

###### Description

Prints XML to given output stream. 

###### Parameters

out
:   Output stream to print to.

node
:   Node to be printed. Pass xml\_document to print entire document.

flags
:   Flags controlling how XML is printed.

###### Returns

Output stream.

##### function operator<<

###### Synopsis

`std::basic_ostream<Ch>& rapidxml::operator<<(std::basic_ostream< Ch > &out, const xml_node< Ch > &node);` 

###### Description

Prints formatted XML to given output stream. Uses default printing flags. Use print() function to customize printing process. 

###### Parameters

out
:   Output stream to print to.

node
:   Node to be printed.

###### Returns

Output stream.

##### constant parse\_no\_data\_nodes

###### Synopsis

`const int parse_no_data_nodes
= 0x1;` 

###### Description

Parse flag instructing the parser to not create data nodes. Text of first data node will still be placed in value of parent element, unless rapidxml::parse\_no\_element\_values flag is also specified. Can be combined with other flags by use of | operator.   
  
See xml\_document::parse() function. 

##### constant parse\_no\_element\_values

###### Synopsis

`const int parse_no_element_values
= 0x2;` 

###### Description

Parse flag instructing the parser to not use text of first data node as a value of parent element. Can be combined with other flags by use of | operator. Note that child data nodes of element node take precendence over its value when printing. That is, if element has one or more child data nodes *and* a value, the value will be ignored. Use rapidxml::parse\_no\_data\_nodes flag to prevent creation of data nodes if you want to manipulate data using values of elements.   
  
See xml\_document::parse() function. 

##### constant parse\_no\_string\_terminators

###### Synopsis

`const int parse_no_string_terminators
= 0x4;` 

###### Description

Parse flag instructing the parser to not place zero terminators after strings in the source text. By default zero terminators are placed, modifying source text. Can be combined with other flags by use of | operator.   
  
See xml\_document::parse() function. 

##### constant parse\_no\_entity\_translation

###### Synopsis

`const int parse_no_entity_translation
= 0x8;` 

###### Description

Parse flag instructing the parser to not translate entities in the source text. By default entities are translated, modifying source text. Can be combined with other flags by use of | operator.   
  
See xml\_document::parse() function. 

##### constant parse\_no\_utf8

###### Synopsis

`const int parse_no_utf8
= 0x10;` 

###### Description

Parse flag instructing the parser to disable UTF-8 handling and assume plain 8 bit characters. By default, UTF-8 handling is enabled. Can be combined with other flags by use of | operator.   
  
See xml\_document::parse() function. 

##### constant parse\_declaration\_node

###### Synopsis

`const int parse_declaration_node
= 0x20;` 

###### Description

Parse flag instructing the parser to create XML declaration node. By default, declaration node is not created. Can be combined with other flags by use of | operator.   
  
See xml\_document::parse() function. 

##### constant parse\_comment\_nodes

###### Synopsis

`const int parse_comment_nodes
= 0x40;` 

###### Description

Parse flag instructing the parser to create comments nodes. By default, comment nodes are not created. Can be combined with other flags by use of | operator.   
  
See xml\_document::parse() function. 

##### constant parse\_doctype\_node

###### Synopsis

`const int parse_doctype_node
= 0x80;` 

###### Description

Parse flag instructing the parser to create DOCTYPE node. By default, doctype node is not created. Although W3C specification allows at most one DOCTYPE node, RapidXml will silently accept documents with more than one. Can be combined with other flags by use of | operator.   
  
See xml\_document::parse() function. 

##### constant parse\_pi\_nodes

###### Synopsis

`const int parse_pi_nodes
= 0x100;` 

###### Description

Parse flag instructing the parser to create PI nodes. By default, PI nodes are not created. Can be combined with other flags by use of | operator.   
  
See xml\_document::parse() function. 

##### constant parse\_validate\_closing\_tags

###### Synopsis

`const int parse_validate_closing_tags
= 0x200;` 

###### Description

Parse flag instructing the parser to validate closing tag names. If not set, name inside closing tag is irrelevant to the parser. By default, closing tags are not validated. Can be combined with other flags by use of | operator.   
  
See xml\_document::parse() function. 

##### constant parse\_trim\_whitespace

###### Synopsis

`const int parse_trim_whitespace
= 0x400;` 

###### Description

Parse flag instructing the parser to trim all leading and trailing whitespace of data nodes. By default, whitespace is not trimmed. This flag does not cause the parser to modify source text. Can be combined with other flags by use of | operator.   
  
See xml\_document::parse() function. 

##### constant parse\_normalize\_whitespace

###### Synopsis

`const int parse_normalize_whitespace
= 0x800;` 

###### Description

Parse flag instructing the parser to condense all whitespace runs of data nodes to a single space character. Trimming of leading and trailing whitespace of data is controlled by rapidxml::parse\_trim\_whitespace flag. By default, whitespace is not normalized. If this flag is specified, source text will be modified. Can be combined with other flags by use of | operator.   
  
See xml\_document::parse() function. 

##### constant parse\_default

###### Synopsis

`const int parse_default
= 0;` 

###### Description

Parse flags which represent default behaviour of the parser. This is always equal to 0, so that all other flags can be simply ored together. Normally there is no need to inconveniently disable flags by anding with their negated (~) values. This also means that meaning of each flag is a *negation* of the default setting. For example, if flag name is rapidxml::parse\_no\_utf8, it means that utf-8 is *enabled* by default, and using the flag will disable it.   
  
See xml\_document::parse() function. 

##### constant parse\_non\_destructive

###### Synopsis

`const int parse_non_destructive
= parse_no_string_terminators | parse_no_entity_translation;` 

###### Description

A combination of parse flags that forbids any modifications of the source text. This also results in faster parsing. However, note that the following will occur:

- names and values of nodes will not be zero terminated, you have to use xml\_base::name\_size() and xml\_base::value\_size() functions to determine where name and value ends
- entities will not be translated
- whitespace will not be normalized

See xml\_document::parse() function. 

##### constant parse\_fastest

###### Synopsis

`const int parse_fastest
= parse_non_destructive | parse_no_data_nodes;` 

###### Description

A combination of parse flags resulting in fastest possible parsing, without sacrificing important data.   
  
See xml\_document::parse() function. 

##### constant parse\_full

###### Synopsis

`const int parse_full
= parse_declaration_node | parse_comment_nodes | parse_doctype_node | parse_pi_nodes | parse_validate_closing_tags;` 

###### Description

A combination of parse flags resulting in largest amount of data being extracted. This usually results in slowest parsing.   
  
See xml\_document::parse() function. 

##### constant print\_no\_indenting

###### Synopsis

`const int print_no_indenting
= 0x1;` 

###### Description

Printer flag instructing the printer to suppress indenting of XML. See print() function. 
