## Supplemental code for "The motif composition of variable-number tandem repeats impacts gene expression": rapidxml.hpp

```
#ifndef CEREAL_RAPIDXML_HPP_INCLUDED
#define CEREAL_RAPIDXML_HPP_INCLUDED

// Copyright (C) 2006, 2009 Marcin Kalicinski
// Version 1.13
// Revision $DateTime: 2009/05/13 01:46:17 $

// If standard library is disabled, user must provide implementations of required functions and typedefs
#if !defined(CEREAL_RAPIDXML_NO_STDLIB)
    #include <cstdlib>      // For std::size_t
    #include <cassert>      // For assert
    #include <new>          // For placement new
#endif

// On MSVC, disable "conditional expression is constant" warning (level 4).
// This warning is almost impossible to avoid with certain types of templated code
#ifdef _MSC_VER
    #pragma warning(push)
    #pragma warning(disable:4127)   // Conditional expression is constant
    #pragma warning(disable:4100)   // unreferenced formal parameter
#endif

///////////////////////////////////////////////////////////////////////////
// CEREAL_RAPIDXML_PARSE_ERROR

#if defined(CEREAL_RAPIDXML_NO_EXCEPTIONS)

#define CEREAL_RAPIDXML_PARSE_ERROR(what, where) { parse_error_handler(what, where); assert(0); }

namespace cereal {
namespace rapidxml
{
    //! When exceptions are disabled by defining CEREAL_RAPIDXML_NO_EXCEPTIONS,
    //! this function is called to notify user about the error.
    //! It must be defined by the user.
    //! <br><br>
    //! This function cannot return. If it does, the results are undefined.
    //! <br><br>
    //! A very simple definition might look like that:
    //! <pre>
    //! void %rapidxml::%parse_error_handler(const char *what, void *where)
    //! {
    //!     std::cout << "Parse error: " << what << "\n";
    //!     std::abort();
    //! }
    //! </pre>
    //! \param what Human readable description of the error.
    //! \param where Pointer to character data where error was detected.
    void parse_error_handler(const char *what, void *where);
}
} // end namespace cereal

#else

#include <exception>    // For std::exception

#define CEREAL_RAPIDXML_PARSE_ERROR(what, where) throw parse_error(what, where)

namespace cereal {
namespace rapidxml
{

    //! Parse error exception.
    //! This exception is thrown by the parser when an error occurs.
    //! Use what() function to get human-readable error message.
    //! Use where() function to get a pointer to position within source text where error was detected.
    //! <br><br>
    //! If throwing exceptions by the parser is undesirable,
    //! it can be disabled by defining CEREAL_RAPIDXML_NO_EXCEPTIONS macro before rapidxml.hpp is included.
    //! This will cause the parser to call rapidxml::parse_error_handler() function instead of throwing an exception.
    //! This function must be defined by the user.
    //! <br><br>
    //! This class derives from <code>std::exception</code> class.
    class parse_error: public std::exception
    {

    public:

        //! Constructs parse error
        parse_error(const char *what_, void *where_)
            : m_what(what_)
            , m_where(where_)
        {
        }

        //! Gets human readable description of error.
        //! \return Pointer to null terminated description of the error.
        virtual const char *what() const CEREAL_NOEXCEPT override
        {
            return m_what;
        }

        //! Gets pointer to character data where error happened.
        //! Ch should be the same as char type of xml_document that produced the error.
        //! \return Pointer to location within the parsed string where error occured.
        template<class Ch>
        Ch *where() const
        {
            return reinterpret_cast<Ch *>(m_where);
        }

    private:

        const char *m_what;
        void *m_where;

    };
}
} // end namespace cereal

#endif

///////////////////////////////////////////////////////////////////////////
// Pool sizes

#ifndef CEREAL_RAPIDXML_STATIC_POOL_SIZE
    // Size of static memory block of memory_pool.
    // Define CEREAL_RAPIDXML_STATIC_POOL_SIZE before including rapidxml.hpp if you want to override the default value.
    // No dynamic memory allocations are performed by memory_pool until static memory is exhausted.
    #define CEREAL_RAPIDXML_STATIC_POOL_SIZE (64 * 1024)
#endif

#ifndef CEREAL_RAPIDXML_DYNAMIC_POOL_SIZE
    // Size of dynamic memory block of memory_pool.
    // Define CEREAL_RAPIDXML_DYNAMIC_POOL_SIZE before including rapidxml.hpp if you want to override the default value.
    // After the static block is exhausted, dynamic blocks with approximately this size are allocated by memory_pool.
    #define CEREAL_RAPIDXML_DYNAMIC_POOL_SIZE (64 * 1024)
#endif

#ifndef CEREAL_RAPIDXML_ALIGNMENT
    // Memory allocation alignment.
    // Define CEREAL_RAPIDXML_ALIGNMENT before including rapidxml.hpp if you want to override the default value, which is the size of pointer.
    // All memory allocations for nodes, attributes and strings will be aligned to this value.
    // This must be a power of 2 and at least 1, otherwise memory_pool will not work.
    #define CEREAL_RAPIDXML_ALIGNMENT sizeof(void *)
#endif

namespace cereal {
namespace rapidxml
{
    // Forward declarations
    template<class Ch> class xml_node;
    template<class Ch> class xml_attribute;
    template<class Ch> class xml_document;

    //! Enumeration listing all node types produced by the parser.
    //! Use xml_node::type() function to query node type.
    enum node_type
    {
        node_document,      //!< A document node. Name and value are empty.
        node_element,       //!< An element node. Name contains element name. Value contains text of first data node.
        node_data,          //!< A data node. Name is empty. Value contains data text.
        node_cdata,         //!< A CDATA node. Name is empty. Value contains data text.
        node_comment,       //!< A comment node. Name is empty. Value contains comment text.
        node_declaration,   //!< A declaration node. Name and value are empty. Declaration parameters (version, encoding and standalone) are in node attributes.
        node_doctype,       //!< A DOCTYPE node. Name is empty. Value contains DOCTYPE text.
        node_pi             //!< A PI node. Name contains target. Value contains instructions.
    };

    ///////////////////////////////////////////////////////////////////////
    // Parsing flags

    //! Parse flag instructing the parser to not create data nodes.
    //! Text of first data node will still be placed in value of parent element, unless rapidxml::parse_no_element_values flag is also specified.
    //! Can be combined with other flags by use of | operator.
    //! <br><br>
    //! See xml_document::parse() function.
    const int parse_no_data_nodes = 0x1;

    //! Parse flag instructing the parser to not use text of first data node as a value of parent element.
    //! Can be combined with other flags by use of | operator.
    //! Note that child data nodes of element node take precendence over its value when printing.
    //! That is, if element has one or more child data nodes <em>and</em> a value, the value will be ignored.
    //! Use rapidxml::parse_no_data_nodes flag to prevent creation of data nodes if you want to manipulate data using values of elements.
    //! <br><br>
    //! See xml_document::parse() function.
    const int parse_no_element_values = 0x2;

    //! Parse flag instructing the parser to not place zero terminators after strings in the source text.
    //! By default zero terminators are placed, modifying source text.
    //! Can be combined with other flags by use of | operator.
    //! <br><br>
    //! See xml_document::parse() function.
    const int parse_no_string_terminators = 0x4;

    //! Parse flag instructing the parser to not translate entities in the source text.
    //! By default entities are translated, modifying source text.
    //! Can be combined with other flags by use of | operator.
    //! <br><br>
    //! See xml_document::parse() function.
    const int parse_no_entity_translation = 0x8;

    //! Parse flag instructing the parser to disable UTF-8 handling and assume plain 8 bit characters.
    //! By default, UTF-8 handling is enabled.
    //! Can be combined with other flags by use of | operator.
    //! <br><br>
    //! See xml_document::parse() function.
    const int parse_no_utf8 = 0x10;

    //! Parse flag instructing the parser to create XML declaration node.
    //! By default, declaration node is not created.
    //! Can be combined with other flags by use of | operator.
    //! <br><br>
    //! See xml_document::parse() function.
    const int parse_declaration_node = 0x20;

    //! Parse flag instructing the parser to create comments nodes.
    //! By default, comment nodes are not created.
    //! Can be combined with other flags by use of | operator.
    //! <br><br>
    //! See xml_document::parse() function.
    const int parse_comment_nodes = 0x40;

    //! Parse flag instructing the parser to create DOCTYPE node.
    //! By default, doctype node is not created.
    //! Although W3C specification allows at most one DOCTYPE node, RapidXml will silently accept documents with more than one.
    //! Can be combined with other flags by use of | operator.
    //! <br><br>
    //! See xml_document::parse() function.
    const int parse_doctype_node = 0x80;

    //! Parse flag instructing the parser to create PI nodes.
    //! By default, PI nodes are not created.
    //! Can be combined with other flags by use of | operator.
    //! <br><br>
    //! See xml_document::parse() function.
    const int parse_pi_nodes = 0x100;

    //! Parse flag instructing the parser to validate closing tag names.
    //! If not set, name inside closing tag is irrelevant to the parser.
    //! By default, closing tags are not validated.
    //! Can be combined with other flags by use of | operator.
    //! <br><br>
    //! See xml_document::parse() function.
    const int parse_validate_closing_tags = 0x200;

    //! Parse flag instructing the parser to trim all leading and trailing whitespace of data nodes.
    //! By default, whitespace is not trimmed.
    //! This flag does not cause the parser to modify source text.
    //! Can be combined with other flags by use of | operator.
    //! <br><br>
    //! See xml_document::parse() function.
    const int parse_trim_whitespace = 0x400;

    //! Parse flag instructing the parser to condense all whitespace runs of data nodes to a single space character.
    //! Trimming of leading and trailing whitespace of data is controlled by rapidxml::parse_trim_whitespace flag.
    //! By default, whitespace is not normalized.
    //! If this flag is specified, source text will be modified.
    //! Can be combined with other flags by use of | operator.
    //! <br><br>
    //! See xml_document::parse() function.
    const int parse_normalize_whitespace = 0x800;

    // Compound flags

    //! Parse flags which represent default behaviour of the parser.
    //! This is always equal to 0, so that all other flags can be simply ored together.
    //! Normally there is no need to inconveniently disable flags by anding with their negated (~) values.
    //! This also means that meaning of each flag is a <i>negation</i> of the default setting.
    //! For example, if flag name is rapidxml::parse_no_utf8, it means that utf-8 is <i>enabled</i> by default,
    //! and using the flag will disable it.
    //! <br><br>
    //! See xml_document::parse() function.
    const int parse_default = 0;

    //! A combination of parse flags that forbids any modifications of the source text.
    //! This also results in faster parsing. However, note that the following will occur:
    //! <ul>
    //! <li>names and values of nodes will not be zero terminated, you have to use xml_base::name_size() and xml_base::value_size() functions to determine where name and value ends</li>
    //! <li>entities will not be translated</li>
    //! <li>whitespace will not be normalized</li>
    //! </ul>
    //! See xml_document::parse() function.
    const int parse_non_destructive = parse_no_string_terminators | parse_no_entity_translation;

    //! A combination of parse flags resulting in fastest possible parsing, without sacrificing important data.
    //! <br><br>
    //! See xml_document::parse() function.
    const int parse_fastest = parse_non_destructive | parse_no_data_nodes;

    //! A combination of parse flags resulting in largest amount of data being extracted.
    //! This usually results in slowest parsing.
    //! <br><br>
    //! See xml_document::parse() function.
    const int parse_full = parse_declaration_node | parse_comment_nodes | parse_doctype_node | parse_pi_nodes | parse_validate_closing_tags;

    ///////////////////////////////////////////////////////////////////////
    // Internals

    //! \cond internal
    namespace internal
    {

        // Struct that contains lookup tables for the parser
        // It must be a template to allow correct linking (because it has static data members, which are defined in a header file).
        template<int Dummy>
        struct lookup_tables
        {
            static const unsigned char lookup_whitespace[256];              // Whitespace table
            static const unsigned char lookup_node_name[256];               // Node name table
            static const unsigned char lookup_text[256];                    // Text table
            static const unsigned char lookup_text_pure_no_ws[256];         // Text table
            static const unsigned char lookup_text_pure_with_ws[256];       // Text table
            static const unsigned char lookup_attribute_name[256];          // Attribute name table
            static const unsigned char lookup_attribute_data_1[256];        // Attribute data table with single quote
            static const unsigned char lookup_attribute_data_1_pure[256];   // Attribute data table with single quote
            static const unsigned char lookup_attribute_data_2[256];        // Attribute data table with double quotes
            static const unsigned char lookup_attribute_data_2_pure[256];   // Attribute data table with double quotes
            static const unsigned char lookup_digits[256];                  // Digits
            static const unsigned char lookup_upcase[256];                  // To uppercase conversion table for ASCII characters
        };

        // Find length of the string
        template<class Ch>
        inline std::size_t measure(const Ch *p)
        {
            const Ch *tmp = p;
            while (*tmp)
                ++tmp;
            return static_cast<std::size_t>(tmp - p);
        }

        // Compare strings for equality
        template<class Ch>
        inline bool compare(const Ch *p1, std::size_t size1, const Ch *p2, std::size_t size2, bool case_sensitive)
        {
            if (size1 != size2)
                return false;
            if (case_sensitive)
            {
                for (const Ch *end = p1 + size1; p1 < end; ++p1, ++p2)
                    if (*p1 != *p2)
                        return false;
            }
            else
            {
                for (const Ch *end = p1 + size1; p1 < end; ++p1, ++p2)
                    if (lookup_tables<0>::lookup_upcase[static_cast<unsigned char>(*p1)] != lookup_tables<0>::lookup_upcase[static_cast<unsigned char>(*p2)])
                        return false;
            }
            return true;
        }

        template<class Ch>
        inline bool preserve_space(xml_node<Ch>* node)
        {
            const Ch preserve_value[] = { Ch('p'), Ch('r'), Ch('e'), Ch('s'), Ch('e'), Ch('r'), Ch('v'), Ch('e') };
            const xml_attribute<Ch>* space = node->first_attribute("xml:space");
            return space && internal::compare(space->value(), space->value_size(), preserve_value, sizeof(preserve_value) / sizeof(Ch), true);
        }
    }
    //! \endcond

    ///////////////////////////////////////////////////////////////////////
    // Memory pool

    //! This class is used by the parser to create new nodes and attributes, without overheads of dynamic memory allocation.
    //! In most cases, you will not need to use this class directly.
    //! However, if you need to create nodes manually or modify names/values of nodes,
    //! you are encouraged to use memory_pool of relevant xml_document to allocate the memory.
    //! Not only is this faster than allocating them by using <code>new</code> operator,
    //! but also their lifetime will be tied to the lifetime of document,
    //! possibly simplyfing memory management.
    //! <br><br>
    //! Call allocate_node() or allocate_attribute() functions to obtain new nodes or attributes from the pool.
    //! You can also call allocate_string() function to allocate strings.
    //! Such strings can then be used as names or values of nodes without worrying about their lifetime.
    //! Note that there is no <code>free()</code> function -- all allocations are freed at once when clear() function is called,
    //! or when the pool is destroyed.
    //! <br><br>
    //! It is also possible to create a standalone memory_pool, and use it
    //! to allocate nodes, whose lifetime will not be tied to any document.
    //! <br><br>
    //! Pool maintains <code>CEREAL_RAPIDXML_STATIC_POOL_SIZE</code> bytes of statically allocated memory.
    //! Until static memory is exhausted, no dynamic memory allocations are done.
    //! When static memory is exhausted, pool allocates additional blocks of memory of size <code>CEREAL_RAPIDXML_DYNAMIC_POOL_SIZE</code> each,
    //! by using global <code>new[]</code> and <code>delete[]</code> operators.
    //! This behaviour can be changed by setting custom allocation routines.
    //! Use set_allocator() function to set them.
    //! <br><br>
    //! Allocations for nodes, attributes and strings are aligned at <code>CEREAL_RAPIDXML_ALIGNMENT</code> bytes.
    //! This value defaults to the size of pointer on target architecture.
    //! <br><br>
    //! To obtain absolutely top performance from the parser,
    //! it is important that all nodes are allocated from a single, contiguous block of memory.
    //! Otherwise, cache misses when jumping between two (or more) disjoint blocks of memory can slow down parsing quite considerably.
    //! If required, you can tweak <code>CEREAL_RAPIDXML_STATIC_POOL_SIZE</code>, <code>CEREAL_RAPIDXML_DYNAMIC_POOL_SIZE</code> and <code>CEREAL_RAPIDXML_ALIGNMENT</code>
    //! to obtain best wasted memory to performance compromise.
    //! To do it, define their values before rapidxml.hpp file is included.
    //! \tparam Ch Character type of created nodes.
    template<class Ch = char>
    class memory_pool
    {

    public:

        //! \cond internal
        typedef void *(alloc_func)(std::size_t);       // Type of user-defined function used to allocate memory
        typedef void (free_func)(void *);              // Type of user-defined function used to free memory
        //! \endcond

        //! Constructs empty pool with default allocator functions.
        memory_pool()
            : m_alloc_func(0)
            , m_free_func(0)
        {
            init();
        }

        //! Destroys pool and frees all the memory.
        //! This causes memory occupied by nodes allocated by the pool to be freed.
        //! Nodes allocated from the pool are no longer valid.
        ~memory_pool()
        {
            clear();
        }

        //! Allocates a new node from the pool, and optionally assigns name and value to it.
        //! If the allocation request cannot be accomodated, this function will throw <code>std::bad_alloc</code>.
        //! If exceptions are disabled by defining CEREAL_RAPIDXML_NO_EXCEPTIONS, this function
        //! will call rapidxml::parse_error_handler() function.
        //! \param type Type of node to create.
        //! \param name Name to assign to the node, or 0 to assign no name.
        //! \param value Value to assign to the node, or 0 to assign no value.
        //! \param name_size Size of name to assign, or 0 to automatically calculate size from name string.
        //! \param value_size Size of value to assign, or 0 to automatically calculate size from value string.
        //! \return Pointer to allocated node. This pointer will never be NULL.
        xml_node<Ch> *allocate_node(node_type type,
                                    const Ch *name = 0, const Ch *value = 0,
                                    std::size_t name_size = 0, std::size_t value_size = 0)
        {
            void *memory = allocate_aligned(sizeof(xml_node<Ch>));
            xml_node<Ch> *node = new(memory) xml_node<Ch>(type);
            if (name)
            {
                if (name_size > 0)
                    node->name(name, name_size);
                else
                    node->name(name);
            }
            if (value)
            {
                if (value_size > 0)
                    node->value(value, value_size);
                else
                    node->value(value);
            }
            return node;
        }

        //! Allocates a new attribute from the pool, and optionally assigns name and value to it.
        //! If the allocation request cannot be accomodated, this function will throw <code>std::bad_alloc</code>.
        //! If exceptions are disabled by defining CEREAL_RAPIDXML_NO_EXCEPTIONS, this function
        //! will call rapidxml::parse_error_handler() function.
        //! \param name Name to assign to the attribute, or 0 to assign no name.
        //! \param value Value to assign to the attribute, or 0 to assign no value.
        //! \param name_size Size of name to assign, or 0 to automatically calculate size from name string.
        //! \param value_size Size of value to assign, or 0 to automatically calculate size from value string.
        //! \return Pointer to allocated attribute. This pointer will never be NULL.
        xml_attribute<Ch> *allocate_attribute(const Ch *name = 0, const Ch *value = 0,
                                              std::size_t name_size = 0, std::size_t value_size = 0)
        {
            void *memory = allocate_aligned(sizeof(xml_attribute<Ch>));
            xml_attribute<Ch> *attribute = new(memory) xml_attribute<Ch>;
            if (name)
            {
                if (name_size > 0)
                    attribute->name(name, name_size);
                else
                    attribute->name(name);
            }
            if (value)
            {
                if (value_size > 0)
                    attribute->value(value, value_size);
                else
                    attribute->value(value);
            }
            return attribute;
        }

        //! Allocates a char array of given size from the pool, and optionally copies a given string to it.
        //! If the allocation request cannot be accomodated, this function will throw <code>std::bad_alloc</code>.
        //! If exceptions are disabled by defining CEREAL_RAPIDXML_NO_EXCEPTIONS, this function
        //! will call rapidxml::parse_error_handler() function.
        //! \param source String to initialize the allocated memory with, or 0 to not initialize it.
        //! \param size Number of characters to allocate, or zero to calculate it automatically from source string length; if size is 0, source string must be specified and null terminated.
        //! \return Pointer to allocated char array. This pointer will never be NULL.
        Ch *allocate_string(const Ch *source = 0, std::size_t size = 0)
        {
            assert(source || size);     // Either source or size (or both) must be specified
            if (size == 0)
                size = internal::measure(source) + 1;
            Ch *result = static_cast<Ch *>(allocate_aligned(size * sizeof(Ch)));
            if (source)
                for (std::size_t i = 0; i < size; ++i)
                    result[i] = source[i];
            return result;
        }

        //! Clones an xml_node and its hierarchy of child nodes and attributes.
        //! Nodes and attributes are allocated from this memory pool.
        //! Names and values are not cloned, they are shared between the clone and the source.
        //! Result node can be optionally specified as a second parameter,
        //! in which case its contents will be replaced with cloned source node.
        //! This is useful when you want to clone entire document.
        //! \param source Node to clone.
        //! \param result Node to put results in, or 0 to automatically allocate result node
        //! \return Pointer to cloned node. This pointer will never be NULL.
        xml_node<Ch> *clone_node(const xml_node<Ch> *source, xml_node<Ch> *result = 0)
        {
            // Prepare result node
            if (result)
            {
                result->remove_all_attributes();
                result->remove_all_nodes();
                result->type(source->type());
            }
            else
                result = allocate_node(source->type());

            // Clone name and value
            result->name(source->name(), source->name_size());
            result->value(source->value(), source->value_size());

            // Clone child nodes and attributes
            for (xml_node<Ch> *child = source->first_node(); child; child = child->next_sibling())
                result->append_node(clone_node(child));
            for (xml_attribute<Ch> *attr = source->first_attribute(); attr; attr = attr->next_attribute())
                result->append_attribute(allocate_attribute(attr->name(), attr->value(), attr->name_size(), attr->value_size()));

            return result;
        }

        //! Clears the pool.
        //! This causes memory occupied by nodes allocated by the pool to be freed.
        //! Any nodes or strings allocated from the pool will no longer be valid.
        void clear()
        {
            while (m_begin != m_static_memory)
            {
                char *previous_begin = reinterpret_cast<header *>(align(m_begin))->previous_begin;
                if (m_free_func)
                    m_free_func(m_begin);
                else
                    delete[] m_begin;
                m_begin = previous_begin;
            }
            init();
        }

        //! Sets or resets the user-defined memory allocation functions for the pool.
        //! This can only be called when no memory is allocated from the pool yet, otherwise results are undefined.
        //! Allocation function must not return invalid pointer on failure. It should either throw,
        //! stop the program, or use <code>longjmp()</code> function to pass control to other place of program.
        //! If it returns invalid pointer, results are undefined.
        //! <br><br>
        //! User defined allocation functions must have the following forms:
        //! <br><code>
        //! <br>void *allocate(std::size_t size);
        //! <br>void free(void *pointer);
        //! </code><br>
        //! \param af Allocation function, or 0 to restore default function
        //! \param ff Free function, or 0 to restore default function
        void set_allocator(alloc_func *af, free_func *ff)
        {
            assert(m_begin == m_static_memory && m_ptr == align(m_begin));    // Verify that no memory is allocated yet
            m_alloc_func = af;
            m_free_func = ff;
        }

    private:

        struct header
        {
            char *previous_begin;
        };

        void init()
        {
            m_begin = m_static_memory;
            m_ptr = align(m_begin);
            m_end = m_static_memory + sizeof(m_static_memory);
        }

        char *align(char *ptr)
        {
            std::size_t alignment = ((CEREAL_RAPIDXML_ALIGNMENT - (std::size_t(ptr) & (CEREAL_RAPIDXML_ALIGNMENT - 1))) & (CEREAL_RAPIDXML_ALIGNMENT - 1));
            return ptr + alignment;
        }

        char *allocate_raw(std::size_t size)
        {
            // Allocate
            void *memory;
            if (m_alloc_func)   // Allocate memory using either user-specified allocation function or global operator new[]
            {
                memory = m_alloc_func(size);
                assert(memory); // Allocator is not allowed to return 0, on failure it must either throw, stop the program or use longjmp
            }
            else
            {
                memory = new char[size];
#ifdef CEREAL_RAPIDXML_NO_EXCEPTIONS
                if (!memory)            // If exceptions are disabled, verify memory allocation, because new will not be able to throw bad_alloc
                    CEREAL_RAPIDXML_PARSE_ERROR("out of memory", 0);
#endif
            }
            return static_cast<char *>(memory);
        }

        void *allocate_aligned(std::size_t size)
        {
            // Calculate aligned pointer
            char *result = align(m_ptr);

            // If not enough memory left in current pool, allocate a new pool
            if (result + size > m_end)
            {
                // Calculate required pool size (may be bigger than CEREAL_RAPIDXML_DYNAMIC_POOL_SIZE)
                std::size_t pool_size = CEREAL_RAPIDXML_DYNAMIC_POOL_SIZE;
                if (pool_size < size)
                    pool_size = size;

                // Allocate
                std::size_t alloc_size = sizeof(header) + (2 * CEREAL_RAPIDXML_ALIGNMENT - 2) + pool_size;     // 2 alignments required in worst case: one for header, one for actual allocation
                char *raw_memory = allocate_raw(alloc_size);

                // Setup new pool in allocated memory
                char *pool = align(raw_memory);
                header *new_header = reinterpret_cast<header *>(pool);
                new_header->previous_begin = m_begin;
                m_begin = raw_memory;
                m_ptr = pool + sizeof(header);
                m_end = raw_memory + alloc_size;

                // Calculate aligned pointer again using new pool
                result = align(m_ptr);
            }

            // Update pool and return aligned pointer
            m_ptr = result + size;
            return result;
        }

        char *m_begin;                                      // Start of raw memory making up current pool
        char *m_ptr;                                        // First free byte in current pool
        char *m_end;                                        // One past last available byte in current pool
        char m_static_memory[CEREAL_RAPIDXML_STATIC_POOL_SIZE];    // Static raw memory
        alloc_func *m_alloc_func;                           // Allocator function, or 0 if default is to be used
        free_func *m_free_func;                             // Free function, or 0 if default is to be used
    };

    ///////////////////////////////////////////////////////////////////////////
    // XML base

    //! Base class for xml_node and xml_attribute implementing common functions:
    //! name(), name_size(), value(), value_size() and parent().
    //! \tparam Ch Character type to use
    template<class Ch = char>
    class xml_base
    {

    public:

        ///////////////////////////////////////////////////////////////////////////
        // Construction & destruction

        // Construct a base with empty name, value and parent
        xml_base()
            : m_name(0)
            , m_value(0)
            , m_parent(0)
        {
        }

        ///////////////////////////////////////////////////////////////////////////
        // Node data access

        //! Gets name of the node.
        //! Interpretation of name depends on type of node.
        //! Note that name will not be zero-terminated if rapidxml::parse_no_string_terminators option was selected during parse.
        //! <br><br>
        //! Use name_size() function to determine length of the name.
        //! \return Name of node, or empty string if node has no name.
        Ch *name() const
        {
            return m_name ? m_name : nullstr();
        }

        //! Gets size of node name, not including terminator character.
        //! This function works correctly irrespective of whether name is or is not zero terminated.
        //! \return Size of node name, in characters.
        std::size_t name_size() const
        {
            return m_name ? m_name_size : 0;
        }

        //! Gets value of node.
        //! Interpretation of value depends on type of node.
        //! Note that value will not be zero-terminated if rapidxml::parse_no_string_terminators option was selected during parse.
        //! <br><br>
        //! Use value_size() function to determine length of the value.
        //! \return Value of node, or empty string if node has no value.
        Ch *value() const
        {
            return m_value ? m_value : nullstr();
        }

        //! Gets size of node value, not including terminator character.
        //! This function works correctly irrespective of whether value is or is not zero terminated.
        //! \return Size of node value, in characters.
        std::size_t value_size() const
        {
            return m_value ? m_value_size : 0;
        }

        ///////////////////////////////////////////////////////////////////////////
        // Node modification

        //! Sets name of node to a non zero-terminated string.
        //! See \ref ownership_of_strings.
        //! <br><br>
        //! Note that node does not own its name or value, it only stores a pointer to it.
        //! It will not delete or otherwise free the pointer on destruction.
        //! It is reponsibility of the user to properly manage lifetime of the string.
        //! The easiest way to achieve it is to use memory_pool of the document to allocate the string -
        //! on destruction of the document the string will be automatically freed.
        //! <br><br>
        //! Size of name must be specified separately, because name does not have to be zero terminated.
        //! Use name(const Ch *) function to have the length automatically calculated (string must be zero terminated).
        //! \param name_ Name of node to set. Does not have to be zero terminated.
        //! \param size Size of name, in characters. This does not include zero terminator, if one is present.
        void name(const Ch *name_, std::size_t size)
        {
            m_name = const_cast<Ch *>(name_);
            m_name_size = size;
        }

        //! Sets name of node to a zero-terminated string.
        //! See also \ref ownership_of_strings and xml_node::name(const Ch *, std::size_t).
        //! \param name_ Name of node to set. Must be zero terminated.
        void name(const Ch *name_)
        {
            this->name(name_, internal::measure(name_));
        }

        //! Sets value of node to a non zero-terminated string.
        //! See \ref ownership_of_strings.
        //! <br><br>
        //! Note that node does not own its name or value, it only stores a pointer to it.
        //! It will not delete or otherwise free the pointer on destruction.
        //! It is reponsibility of the user to properly manage lifetime of the string.
        //! The easiest way to achieve it is to use memory_pool of the document to allocate the string -
        //! on destruction of the document the string will be automatically freed.
        //! <br><br>
        //! Size of value must be specified separately, because it does not have to be zero terminated.
        //! Use value(const Ch *) function to have the length automatically calculated (string must be zero terminated).
        //! <br><br>
        //! If an element has a child node of type node_data, it will take precedence over element value when printing.
        //! If you want to manipulate data of elements using values, use parser flag rapidxml::parse_no_data_nodes to prevent creation of data nodes by the parser.
        //! \param value_ value of node to set. Does not have to be zero terminated.
        //! \param size Size of value, in characters. This does not include zero terminator, if one is present.
        void value(const Ch *value_, std::size_t size)
        {
            m_value = const_cast<Ch *>(value_);
            m_value_size = size;
        }

        //! Sets value of node to a zero-terminated string.
        //! See also \ref ownership_of_strings and xml_node::value(const Ch *, std::size_t).
        //! \param value_ Vame of node to set. Must be zero terminated.
        void value(const Ch *value_)
        {
            this->value(value_, internal::measure(value_));
        }

        ///////////////////////////////////////////////////////////////////////////
        // Related nodes access

        //! Gets node parent.
        //! \return Pointer to parent node, or 0 if there is no parent.
        xml_node<Ch> *parent() const
        {
            return m_parent;
        }

    protected:

        // Return empty string
        static Ch *nullstr()
        {
            static Ch zero = Ch('\0');
            return &zero;
        }

        Ch *m_name;                         // Name of node, or 0 if no name
        Ch *m_value;                        // Value of node, or 0 if no value
        std::size_t m_name_size;            // Length of node name, or undefined of no name
        std::size_t m_value_size;           // Length of node value, or undefined if no value
        xml_node<Ch> *m_parent;             // Pointer to parent node, or 0 if none

    };

    //! Class representing attribute node of XML document.
    //! Each attribute has name and value strings, which are available through name() and value() functions (inherited from xml_base).
    //! Note that after parse, both name and value of attribute will point to interior of source text used for parsing.
    //! Thus, this text must persist in memory for the lifetime of attribute.
    //! \tparam Ch Character type to use.
    template<class Ch = char>
    class xml_attribute: public xml_base<Ch>
    {

        friend class xml_node<Ch>;

    public:

        ///////////////////////////////////////////////////////////////////////////
        // Construction & destruction

        //! Constructs an empty attribute with the specified type.
        //! Consider using memory_pool of appropriate xml_document if allocating attributes manually.
        xml_attribute()
        {
        }

        ///////////////////////////////////////////////////////////////////////////
        // Related nodes access

        //! Gets document of which attribute is a child.
        //! \return Pointer to document that contains this attribute, or 0 if there is no parent document.
        xml_document<Ch> *document() const
        {
            if (xml_node<Ch> *node = this->parent())
            {
                while (node->parent())
                    node = node->parent();
                return node->type() == node_document ? static_cast<xml_document<Ch> *>(node) : 0;
            }
            else
                return 0;
        }

        //! Gets previous attribute, optionally matching attribute name.
        //! \param name Name of attribute to find, or 0 to return previous attribute regardless of its name; this string doesn't have to be zero-terminated if name_size is non-zero
        //! \param name_size Size of name, in characters, or 0 to have size calculated automatically from string
        //! \param case_sensitive Should name comparison be case-sensitive; non case-sensitive comparison works properly only for ASCII characters
        //! \return Pointer to found attribute, or 0 if not found.
        xml_attribute<Ch> *previous_attribute(const Ch *name = 0, std::size_t name_size = 0, bool case_sensitive = true) const
        {
            if (name)
            {
                if (name_size == 0)
                    name_size = internal::measure(name);
                for (xml_attribute<Ch> *attribute = m_prev_attribute; attribute; attribute = attribute->m_prev_attribute)
                    if (internal::compare(attribute->name(), attribute->name_size(), name, name_size, case_sensitive))
                        return attribute;
                return 0;
            }
            else
                return this->m_parent ? m_prev_attribute : 0;
        }

        //! Gets next attribute, optionally matching attribute name.
        //! \param name_ Name of attribute to find, or 0 to return next attribute regardless of its name; this string doesn't have to be zero-terminated if name_size is non-zero
        //! \param name_size_ Size of name, in characters, or 0 to have size calculated automatically from string
        //! \param case_sensitive Should name comparison be case-sensitive; non case-sensitive comparison works properly only for ASCII characters
        //! \return Pointer to found attribute, or 0 if not found.
        xml_attribute<Ch> *next_attribute(const Ch *name_ = 0, std::size_t name_size_ = 0, bool case_sensitive = true) const
        {
            if (name_)
            {
                if (name_size_ == 0)
                    name_size_ = internal::measure(name_);
                for (xml_attribute<Ch> *attribute = m_next_attribute; attribute; attribute = attribute->m_next_attribute)
                    if (internal::compare(attribute->name(), attribute->name_size(), name_, name_size_, case_sensitive))
                        return attribute;
                return 0;
            }
            else
                return this->m_parent ? m_next_attribute : 0;
        }

    private:

        xml_attribute<Ch> *m_prev_attribute;        // Pointer to previous sibling of attribute, or 0 if none; only valid if parent is non-zero
        xml_attribute<Ch> *m_next_attribute;        // Pointer to next sibling of attribute, or 0 if none; only valid if parent is non-zero

    };

    ///////////////////////////////////////////////////////////////////////////
    // XML node

    //! Class representing a node of XML document.
    //! Each node may have associated name and value strings, which are available through name() and value() functions.
    //! Interpretation of name and value depends on type of the node.
    //! Type of node can be determined by using type() function.
    //! <br><br>
    //! Note that after parse, both name and value of node, if any, will point interior of source text used for parsing.
    //! Thus, this text must persist in the memory for the lifetime of node.
    //! \tparam Ch Character type to use.
    template<class Ch = char>
    class xml_node: public xml_base<Ch>
    {

    public:

        ///////////////////////////////////////////////////////////////////////////
        // Construction & destruction

        //! Constructs an empty node with the specified type.
        //! Consider using memory_pool of appropriate document to allocate nodes manually.
        //! \param type_ Type of node to construct.
        xml_node(node_type type_)
            : m_type(type_)
            , m_first_node(0)
            , m_first_attribute(0)
        {
        }

        ///////////////////////////////////////////////////////////////////////////
        // Node data access

        //! Gets type of node.
        //! \return Type of node.
        node_type type() const
        {
            return m_type;
        }

        ///////////////////////////////////////////////////////////////////////////
        // Related nodes access

        //! Gets document of which node is a child.
        //! \return Pointer to document that contains this node, or 0 if there is no parent document.
        xml_document<Ch> *document() const
        {
            xml_node<Ch> *node = const_cast<xml_node<Ch> *>(this);
            while (node->parent())
                node = node->parent();
            return node->type() == node_document ? static_cast<xml_document<Ch> *>(node) : 0;
        }

        //! Gets first child node, optionally matching node name.
        //! \param name_ Name of child to find, or 0 to return first child regardless of its name; this string doesn't have to be zero-terminated if name_size is non-zero
        //! \param name_size_ Size of name, in characters, or 0 to have size calculated automatically from string
        //! \param case_sensitive Should name comparison be case-sensitive; non case-sensitive comparison works properly only for ASCII characters
        //! \return Pointer to found child, or 0 if not found.
        xml_node<Ch> *first_node(const Ch *name_ = 0, std::size_t name_size_ = 0, bool case_sensitive = true) const
        {
            if (name_)
            {
                if (name_size_ == 0)
                    name_size_ = internal::measure(name_);
                for (xml_node<Ch> *child = m_first_node; child; child = child->next_sibling())
                    if (internal::compare(child->name(), child->name_size(), name_, name_size_, case_sensitive))
                        return child;
                return 0;
            }
            else
                return m_first_node;
        }

        //! Gets last child node, optionally matching node name.
        //! Behaviour is undefined if node has no children.
        //! Use first_node() to test if node has children.
        //! \param name Name of child to find, or 0 to return last child regardless of its name; this string doesn't have to be zero-terminated if name_size is non-zero
        //! \param name_size Size of name, in characters, or 0 to have size calculated automatically from string
        //! \param case_sensitive Should name comparison be case-sensitive; non case-sensitive comparison works properly only for ASCII characters
        //! \return Pointer to found child, or 0 if not found.
        xml_node<Ch> *last_node(const Ch *name = 0, std::size_t name_size = 0, bool case_sensitive = true) const
        {
            assert(m_first_node);  // Cannot query for last child if node has no children
            if (name)
            {
                if (name_size == 0)
                    name_size = internal::measure(name);
                for (xml_node<Ch> *child = m_last_node; child; child = child->previous_sibling())
                    if (internal::compare(child->name(), child->name_size(), name, name_size, case_sensitive))
                        return child;
                return 0;
            }
            else
                return m_last_node;
        }

        //! Gets previous sibling node, optionally matching node name.
        //! Behaviour is undefined if node has no parent.
        //! Use parent() to test if node has a parent.
        //! \param name Name of sibling to find, or 0 to return previous sibling regardless of its name; this string doesn't have to be zero-terminated if name_size is non-zero
        //! \param name_size Size of name, in characters, or 0 to have size calculated automatically from string
        //! \param case_sensitive Should name comparison be case-sensitive; non case-sensitive comparison works properly only for ASCII characters
        //! \return Pointer to found sibling, or 0 if not found.
        xml_node<Ch> *previous_sibling(const Ch *name = 0, std::size_t name_size = 0, bool case_sensitive = true) const
        {
            assert(this->m_parent);     // Cannot query for siblings if node has no parent
            if (name)
            {
                if (name_size == 0)
                    name_size = internal::measure(name);
                for (xml_node<Ch> *sibling = m_prev_sibling; sibling; sibling = sibling->m_prev_sibling)
                    if (internal::compare(sibling->name(), sibling->name_size(), name, name_size, case_sensitive))
                        return sibling;
                return 0;
            }
            else
                return m_prev_sibling;
        }

        //! Gets next sibling node, optionally matching node name.
        //! Behaviour is undefined if node has no parent.
        //! Use parent() to test if node has a parent.
        //! \param name_ Name of sibling to find, or 0 to return next sibling regardless of its name; this string doesn't have to be zero-terminated if name_size is non-zero
        //! \param name_size_ Size of name, in characters, or 0 to have size calculated automatically from string
        //! \param case_sensitive Should name comparison be case-sensitive; non case-sensitive comparison works properly only for ASCII characters
        //! \return Pointer to found sibling, or 0 if not found.
        xml_node<Ch> *next_sibling(const Ch *name_ = 0, std::size_t name_size_ = 0, bool case_sensitive = true) const
        {
            assert(this->m_parent);     // Cannot query for siblings if node has no parent
            if (name_)
            {
                if (name_size_ == 0)
                    name_size_ = internal::measure(name_);
                for (xml_node<Ch> *sibling = m_next_sibling; sibling; sibling = sibling->m_next_sibling)
                    if (internal::compare(sibling->name(), sibling->name_size(), name_, name_size_, case_sensitive))
                        return sibling;
                return 0;
            }
            else
                return m_next_sibling;
        }

        //! Gets first attribute of node, optionally matching attribute name.
        //! \param name_ Name of attribute to find, or 0 to return first attribute regardless of its name; this string doesn't have to be zero-terminated if name_size is non-zero
        //! \param name_size_ Size of name, in characters, or 0 to have size calculated automatically from string
        //! \param case_sensitive Should name comparison be case-sensitive; non case-sensitive comparison works properly only for ASCII characters
        //! \return Pointer to found attribute, or 0 if not found.
        xml_attribute<Ch> *first_attribute(const Ch *name_ = 0, std::size_t name_size_ = 0, bool case_sensitive = true) const
        {
            if (name_)
            {
                if (name_size_ == 0)
                    name_size_ = internal::measure(name_);
                for (xml_attribute<Ch> *attribute = m_first_attribute; attribute; attribute = attribute->m_next_attribute)
                    if (internal::compare(attribute->name(), attribute->name_size(), name_, name_size_, case_sensitive))
                        return attribute;
                return 0;
            }
            else
                return m_first_attribute;
        }

        //! Gets last attribute of node, optionally matching attribute name.
        //! \param name Name of attribute to find, or 0 to return last attribute regardless of its name; this string doesn't have to be zero-terminated if name_size is non-zero
        //! \param name_size Size of name, in characters, or 0 to have size calculated automatically from string
        //! \param case_sensitive Should name comparison be case-sensitive; non case-sensitive comparison works properly only for ASCII characters
        //! \return Pointer to found attribute, or 0 if not found.
        xml_attribute<Ch> *last_attribute(const Ch *name = 0, std::size_t name_size = 0, bool case_sensitive = true) const
        {
            if (name)
            {
                if (name_size == 0)
                    name_size = internal::measure(name);
                for (xml_attribute<Ch> *attribute = m_last_attribute; attribute; attribute = attribute->m_prev_attribute)
                    if (internal::compare(attribute->name(), attribute->name_size(), name, name_size, case_sensitive))
                        return attribute;
                return 0;
            }
            else
                return m_first_attribute ? m_last_attribute : 0;
        }

        ///////////////////////////////////////////////////////////////////////////
        // Node modification

        //! Sets type of node.
        //! \param type_ Type of node to set.
        void type(node_type type_)
        {
            m_type = type_;
        }

        ///////////////////////////////////////////////////////////////////////////
        // Node manipulation

        //! Prepends a new child node.
        //! The prepended child becomes the first child, and all existing children are moved one position back.
        //! \param child Node to prepend.
        void prepend_node(xml_node<Ch> *child)
        {
            assert(child && !child->parent() && child->type() != node_document);
            if (first_node())
            {
                child->m_next_sibling = m_first_node;
                m_first_node->m_prev_sibling = child;
            }
            else
            {
                child->m_next_sibling = 0;
                m_last_node = child;
            }
            m_first_node = child;
            child->m_parent = this;
            child->m_prev_sibling = 0;
        }

        //! Appends a new child node.
        //! The appended child becomes the last child.
        //! \param child Node to append.
        void append_node(xml_node<Ch> *child)
        {
            assert(child && !child->parent() && child->type() != node_document);
            if (first_node())
            {
                child->m_prev_sibling = m_last_node;
                m_last_node->m_next_sibling = child;
            }
            else
            {
                child->m_prev_sibling = 0;
                m_first_node = child;
            }
            m_last_node = child;
            child->m_parent = this;
            child->m_next_sibling = 0;
        }

        //! Inserts a new child node at specified place inside the node.
        //! All children after and including the specified node are moved one position back.
        //! \param where Place where to insert the child, or 0 to insert at the back.
        //! \param child Node to insert.
        void insert_node(xml_node<Ch> *where, xml_node<Ch> *child)
        {
            assert(!where || where->parent() == this);
            assert(child && !child->parent() && child->type() != node_document);
            if (where == m_first_node)
                prepend_node(child);
            else if (where == 0)
                append_node(child);
            else
            {
                child->m_prev_sibling = where->m_prev_sibling;
                child->m_next_sibling = where;
                where->m_prev_sibling->m_next_sibling = child;
                where->m_prev_sibling = child;
                child->m_parent = this;
            }
        }

        //! Removes first child node.
        //! If node has no children, behaviour is undefined.
        //! Use first_node() to test if node has children.
        void remove_first_node()
        {
            assert(first_node());
            xml_node<Ch> *child = m_first_node;
            m_first_node = child->m_next_sibling;
            if (child->m_next_sibling)
                child->m_next_sibling->m_prev_sibling = 0;
            else
                m_last_node = 0;
            child->m_parent = 0;
        }

        //! Removes last child of the node.
        //! If node has no children, behaviour is undefined.
        //! Use first_node() to test if node has children.
        void remove_last_node()
        {
            assert(first_node());
            xml_node<Ch> *child = m_last_node;
            if (child->m_prev_sibling)
            {
                m_last_node = child->m_prev_sibling;
                child->m_prev_sibling->m_next_sibling = 0;
            }
            else
                m_first_node = 0;
            child->m_parent = 0;
        }

        //! Removes specified child from the node
        // \param where Pointer to child to be removed.
        void remove_node(xml_node<Ch> *where)
        {
            assert(where && where->parent() == this);
            assert(first_node());
            if (where == m_first_node)
                remove_first_node();
            else if (where == m_last_node)
                remove_last_node();
            else
            {
                where->m_prev_sibling->m_next_sibling = where->m_next_sibling;
                where->m_next_sibling->m_prev_sibling = where->m_prev_sibling;
                where->m_parent = 0;
            }
        }

        //! Removes all child nodes (but not attributes).
        void remove_all_nodes()
        {
            for (xml_node<Ch> *node = first_node(); node; node = node->m_next_sibling)
                node->m_parent = 0;
            m_first_node = 0;
        }

        //! Prepends a new attribute to the node.
        //! \param attribute Attribute to prepend.
        void prepend_attribute(xml_attribute<Ch> *attribute)
        {
            assert(attribute && !attribute->parent());
            if (first_attribute())
            {
                attribute->m_next_attribute = m_first_attribute;
                m_first_attribute->m_prev_attribute = attribute;
            }
            else
            {
                attribute->m_next_attribute = 0;
                m_last_attribute = attribute;
            }
            m_first_attribute = attribute;
            attribute->m_parent = this;
            attribute->m_prev_attribute = 0;
        }

        //! Appends a new attribute to the node.
        //! \param attribute Attribute to append.
        void append_attribute(xml_attribute<Ch> *attribute)
        {
            assert(attribute && !attribute->parent());
            if (first_attribute())
            {
                attribute->m_prev_attribute = m_last_attribute;
                m_last_attribute->m_next_attribute = attribute;
            }
            else
            {
                attribute->m_prev_attribute = 0;
                m_first_attribute = attribute;
            }
            m_last_attribute = attribute;
            attribute->m_parent = this;
            attribute->m_next_attribute = 0;
        }

        //! Inserts a new attribute at specified place inside the node.
        //! All attributes after and including the specified attribute are moved one position back.
        //! \param where Place where to insert the attribute, or 0 to insert at the back.
        //! \param attribute Attribute to insert.
        void insert_attribute(xml_attribute<Ch> *where, xml_attribute<Ch> *attribute)
        {
            assert(!where || where->parent() == this);
            assert(attribute && !attribute->parent());
            if (where == m_first_attribute)
                prepend_attribute(attribute);
            else if (where == 0)
                append_attribute(attribute);
            else
            {
                attribute->m_prev_attribute = where->m_prev_attribute;
                attribute->m_next_attribute = where;
                where->m_prev_attribute->m_next_attribute = attribute;
                where->m_prev_attribute = attribute;
                attribute->m_parent = this;
            }
        }

        //! Removes first attribute of the node.
        //! If node has no attributes, behaviour is undefined.
        //! Use first_attribute() to test if node has attributes.
        void remove_first_attribute()
        {
            assert(first_attribute());
            xml_attribute<Ch> *attribute = m_first_attribute;
            if (attribute->m_next_attribute)
            {
                attribute->m_next_attribute->m_prev_attribute = 0;
            }
            else
                m_last_attribute = 0;
            attribute->m_parent = 0;
            m_first_attribute = attribute->m_next_attribute;
        }

        //! Removes last attribute of the node.
        //! If node has no attributes, behaviour is undefined.
        //! Use first_attribute() to test if node has attributes.
        void remove_last_attribute()
        {
            assert(first_attribute());
            xml_attribute<Ch> *attribute = m_last_attribute;
            if (attribute->m_prev_attribute)
            {
                attribute->m_prev_attribute->m_next_attribute = 0;
                m_last_attribute = attribute->m_prev_attribute;
            }
            else
                m_first_attribute = 0;
            attribute->m_parent = 0;
        }

        //! Removes specified attribute from node.
        //! \param where Pointer to attribute to be removed.
        void remove_attribute(xml_attribute<Ch> *where)
        {
            assert(first_attribute() && where->parent() == this);
            if (where == m_first_attribute)
                remove_first_attribute();
            else if (where == m_last_attribute)
                remove_last_attribute();
            else
            {
                where->m_prev_attribute->m_next_attribute = where->m_next_attribute;
                where->m_next_attribute->m_prev_attribute = where->m_prev_attribute;
                where->m_parent = 0;
            }
        }

        //! Removes all attributes of node.
        void remove_all_attributes()
        {
            for (xml_attribute<Ch> *attribute = first_attribute(); attribute; attribute = attribute->m_next_attribute)
                attribute->m_parent = 0;
            m_first_attribute = 0;
        }

    private:

        ///////////////////////////////////////////////////////////////////////////
        // Restrictions

        // No copying
        xml_node(const xml_node &);
        void operator =(const xml_node &);

        ///////////////////////////////////////////////////////////////////////////
        // Data members

        // Note that some of the pointers below have UNDEFINED values if certain other pointers are 0.
        // This is required for maximum performance, as it allows the parser to omit initialization of
        // unneded/redundant values.
        //
        // The rules are as follows:
        // 1. first_node and first_attribute contain valid pointers, or 0 if node has no children/attributes respectively
        // 2. last_node and last_attribute are valid only if node has at least one child/attribute respectively, otherwise they contain garbage
        // 3. prev_sibling and next_sibling are valid only if node has a parent, otherwise they contain garbage

        node_type m_type;                       // Type of node; always valid
        xml_node<Ch> *m_first_node;             // Pointer to first child node, or 0 if none; always valid
        xml_node<Ch> *m_last_node;              // Pointer to last child node, or 0 if none; this value is only valid if m_first_node is non-zero
        xml_attribute<Ch> *m_first_attribute;   // Pointer to first attribute of node, or 0 if none; always valid
        xml_attribute<Ch> *m_last_attribute;    // Pointer to last attribute of node, or 0 if none; this value is only valid if m_first_attribute is non-zero
        xml_node<Ch> *m_prev_sibling;           // Pointer to previous sibling of node, or 0 if none; this value is only valid if m_parent is non-zero
        xml_node<Ch> *m_next_sibling;           // Pointer to next sibling of node, or 0 if none; this value is only valid if m_parent is non-zero

    };

    ///////////////////////////////////////////////////////////////////////////
    // XML document

    //! This class represents root of the DOM hierarchy.
    //! It is also an xml_node and a memory_pool through public inheritance.
    //! Use parse() function to build a DOM tree from a zero-terminated XML text string.
    //! parse() function allocates memory for nodes and attributes by using functions of xml_document,
    //! which are inherited from memory_pool.
    //! To access root node of the document, use the document itself, as if it was an xml_node.
    //! \tparam Ch Character type to use.
    template<class Ch = char>
    class xml_document: public xml_node<Ch>, public memory_pool<Ch>
    {

    public:

        //! Constructs empty XML document
        xml_document()
            : xml_node<Ch>(node_document)
        {
        }

        //! Parses zero-terminated XML string according to given flags.
        //! Passed string will be modified by the parser, unless rapidxml::parse_non_destructive flag is used.
        //! The string must persist for the lifetime of the document.
        //! In case of error, rapidxml::parse_error exception will be thrown.
        //! <br><br>
        //! If you want to parse contents of a file, you must first load the file into the memory, and pass pointer to its beginning.
        //! Make sure that data is zero-terminated.
        //! <br><br>
        //! Document can be parsed into multiple times.
        //! Each new call to parse removes previous nodes and attributes (if any), but does not clear memory pool.
        //! \param text XML data to parse; pointer is non-const to denote fact that this data may be modified by the parser.
        template<int Flags>
        void parse(Ch *text)
        {
            assert(text);

            // Remove current contents
            this->remove_all_nodes();
            this->remove_all_attributes();

            // Parse BOM, if any
            parse_bom<Flags>(text);

            // Parse children
            while (1)
            {
                // Skip whitespace before node
                skip<whitespace_pred, Flags>(text);
                if (*text == 0)
                    break;

                // Parse and append new child
                if (*text == Ch('<'))
                {
                    ++text;     // Skip '<'
                    if (xml_node<Ch> *node = parse_node<Flags>(text))
                        this->append_node(node);
                }
                else
                    CEREAL_RAPIDXML_PARSE_ERROR("expected <", text);
            }

        }

        //! Clears the document by deleting all nodes and clearing the memory pool.
        //! All nodes owned by document pool are destroyed.
        void clear()
        {
            this->remove_all_nodes();
            this->remove_all_attributes();
            memory_pool<Ch>::clear();
        }

    private:

        ///////////////////////////////////////////////////////////////////////
        // Internal character utility functions

        // Detect whitespace character
        struct whitespace_pred
        {
            static unsigned char test(Ch ch)
            {
                return internal::lookup_tables<0>::lookup_whitespace[static_cast<unsigned char>(ch)];
            }
        };

        // Detect node name character
        struct node_name_pred
        {
            static unsigned char test(Ch ch)
            {
                return internal::lookup_tables<0>::lookup_node_name[static_cast<unsigned char>(ch)];
            }
        };

        // Detect attribute name character
        struct attribute_name_pred
        {
            static unsigned char test(Ch ch)
            {
                return internal::lookup_tables<0>::lookup_attribute_name[static_cast<unsigned char>(ch)];
            }
        };

        // Detect text character (PCDATA)
        struct text_pred
        {
            static unsigned char test(Ch ch)
            {
                return internal::lookup_tables<0>::lookup_text[static_cast<unsigned char>(ch)];
            }
        };

        // Detect text character (PCDATA) that does not require processing
        struct text_pure_no_ws_pred
        {
            static unsigned char test(Ch ch)
            {
                return internal::lookup_tables<0>::lookup_text_pure_no_ws[static_cast<unsigned char>(ch)];
            }
        };

        // Detect text character (PCDATA) that does not require processing
        struct text_pure_with_ws_pred
        {
            static unsigned char test(Ch ch)
            {
                return internal::lookup_tables<0>::lookup_text_pure_with_ws[static_cast<unsigned char>(ch)];
            }
        };

        // Detect attribute value character
        template<Ch Quote>
        struct attribute_value_pred
        {
            static unsigned char test(Ch ch)
            {
                if (Quote == Ch('\''))
                    return internal::lookup_tables<0>::lookup_attribute_data_1[static_cast<unsigned char>(ch)];
                if (Quote == Ch('\"'))
                    return internal::lookup_tables<0>::lookup_attribute_data_2[static_cast<unsigned char>(ch)];
                return 0;       // Should never be executed, to avoid warnings on Comeau
            }
        };

        // Detect attribute value character
        template<Ch Quote>
        struct attribute_value_pure_pred
        {
            static unsigned char test(Ch ch)
            {
                if (Quote == Ch('\''))
                    return internal::lookup_tables<0>::lookup_attribute_data_1_pure[static_cast<unsigned char>(ch)];
                if (Quote == Ch('\"'))
                    return internal::lookup_tables<0>::lookup_attribute_data_2_pure[static_cast<unsigned char>(ch)];
                return 0;       // Should never be executed, to avoid warnings on Comeau
            }
        };

        // Insert coded character, using UTF8 or 8-bit ASCII
        template<int Flags>
        static void insert_coded_character(Ch *&text, unsigned long code)
        {
            if (Flags & parse_no_utf8)
            {
                // Insert 8-bit ASCII character
                // Todo: possibly verify that code is less than 256 and use replacement char otherwise?
                text[0] = static_cast<Ch>(code);
                text += 1;
            }
            else
            {
                // Insert UTF8 sequence
                if (code < 0x80)    // 1 byte sequence
                {
	                text[0] = static_cast<Ch>(code);
                    text += 1;
                }
                else if (code < 0x800)  // 2 byte sequence
                {
	                text[1] = static_cast<Ch>((code | 0x80) & 0xBF); code >>= 6;
	                text[0] = static_cast<Ch>(code | 0xC0);
                    text += 2;
                }
	            else if (code < 0x10000)    // 3 byte sequence
                {
	                text[2] = static_cast<Ch>((code | 0x80) & 0xBF); code >>= 6;
	                text[1] = static_cast<Ch>((code | 0x80) & 0xBF); code >>= 6;
	                text[0] = static_cast<Ch>(code | 0xE0);
                    text += 3;
                }
	            else if (code < 0x110000)   // 4 byte sequence
                {
	                text[3] = static_cast<Ch>((code | 0x80) & 0xBF); code >>= 6;
	                text[2] = static_cast<Ch>((code | 0x80) & 0xBF); code >>= 6;
	                text[1] = static_cast<Ch>((code | 0x80) & 0xBF); code >>= 6;
	                text[0] = static_cast<Ch>(code | 0xF0);
                    text += 4;
                }
                else    // Invalid, only codes up to 0x10FFFF are allowed in Unicode
                {
                    CEREAL_RAPIDXML_PARSE_ERROR("invalid numeric character entity", text);
                }
            }
        }

        // Skip characters until predicate evaluates to true
        template<class StopPred, int Flags>
        static void skip(Ch *&text)
        {
            Ch *tmp = text;
            while (StopPred::test(*tmp))
                ++tmp;
            text = tmp;
        }

        // Skip characters until predicate evaluates to true while doing the following:
        // - replacing XML character entity references with proper characters (&apos; &amp; &quot; &lt; &gt; &#...;)
        // - condensing whitespace sequences to single space character
        template<class StopPred, class StopPredPure, int Flags>
        static Ch *skip_and_expand_character_refs(Ch *&text, bool preserve_space)
        {
            // If entity translation, whitespace condense and whitespace trimming is disabled, use plain skip
            if (Flags & parse_no_entity_translation &&
                !(Flags & parse_normalize_whitespace) &&
                !(Flags & parse_trim_whitespace))
            {
                skip<StopPred, Flags>(text);
                return text;
            }

            // Use simple skip until first modification is detected
            skip<StopPredPure, Flags>(text);

            // Use translation skip
            Ch *src = text;
            Ch *dest = src;
            while (StopPred::test(*src))
            {
                // If entity translation is enabled
                if (!(Flags & parse_no_entity_translation))
                {
                    // Test if replacement is needed
                    if (src[0] == Ch('&'))
                    {
                        switch (src[1])
                        {

                        // &amp; &apos;
                        case Ch('a'):
                            if (src[2] == Ch('m') && src[3] == Ch('p') && src[4] == Ch(';'))
                            {
                                *dest = Ch('&');
                                ++dest;
                                src += 5;
                                continue;
                            }
                            if (src[2] == Ch('p') && src[3] == Ch('o') && src[4] == Ch('s') && src[5] == Ch(';'))
                            {
                                *dest = Ch('\'');
                                ++dest;
                                src += 6;
                                continue;
                            }
                            break;

                        // &quot;
                        case Ch('q'):
                            if (src[2] == Ch('u') && src[3] == Ch('o') && src[4] == Ch('t') && src[5] == Ch(';'))
                            {
                                *dest = Ch('"');
                                ++dest;
                                src += 6;
                                continue;
                            }
                            break;

                        // &gt;
                        case Ch('g'):
                            if (src[2] == Ch('t') && src[3] == Ch(';'))
                            {
                                *dest = Ch('>');
                                ++dest;
                                src += 4;
                                continue;
                            }
                            break;

                        // &lt;
                        case Ch('l'):
                            if (src[2] == Ch('t') && src[3] == Ch(';'))
                            {
                                *dest = Ch('<');
                                ++dest;
                                src += 4;
                                continue;
                            }
                            break;

                        // &#...; - assumes ASCII
                        case Ch('#'):
                            if (src[2] == Ch('x'))
                            {
                                unsigned long code = 0;
                                src += 3;   // Skip &#x
                                while (1)
                                {
                                    unsigned char digit = internal::lookup_tables<0>::lookup_digits[static_cast<unsigned char>(*src)];
                                    if (digit == 0xFF)
                                        break;
                                    code = code * 16 + digit;
                                    ++src;
                                }
                                insert_coded_character<Flags>(dest, code);    // Put character in output
                            }
                            else
                            {
                                unsigned long code = 0;
                                src += 2;   // Skip &#
                                while (1)
                                {
                                    unsigned char digit = internal::lookup_tables<0>::lookup_digits[static_cast<unsigned char>(*src)];
                                    if (digit == 0xFF)
                                        break;
                                    code = code * 10 + digit;
                                    ++src;
                                }
                                insert_coded_character<Flags>(dest, code);    // Put character in output
                            }
                            if (*src == Ch(';'))
                                ++src;
                            else
                                CEREAL_RAPIDXML_PARSE_ERROR("expected ;", src);
                            continue;

                        // Something else
                        default:
                            // Ignore, just copy '&' verbatim
                            break;

                        }
                    }
                }

                // If whitespace condensing is enabled
                if ((Flags & parse_normalize_whitespace) && !preserve_space)
                {
                    // Test if condensing is needed
                    if (whitespace_pred::test(*src))
                    {
                        *dest = Ch(' '); ++dest;    // Put single space in dest
                        ++src;                      // Skip first whitespace char
                        // Skip remaining whitespace chars
                        while (whitespace_pred::test(*src))
                            ++src;
                        continue;
                    }
                }

                // No replacement, only copy character
                *dest++ = *src++;

            }

            // Return new end
            text = src;
            return dest;

        }

        ///////////////////////////////////////////////////////////////////////
        // Internal parsing functions

        // Parse BOM, if any
        template<int Flags>
        void parse_bom(Ch *&text)
        {
            // UTF-8?
            if (static_cast<unsigned char>(text[0]) == 0xEF &&
                static_cast<unsigned char>(text[1]) == 0xBB &&
                static_cast<unsigned char>(text[2]) == 0xBF)
            {
                text += 3;      // Skup utf-8 bom
            }
        }

        // Parse XML declaration (<?xml...)
        template<int Flags>
        xml_node<Ch> *parse_xml_declaration(Ch *&text)
        {
            // If parsing of declaration is disabled
            if (!(Flags & parse_declaration_node))
            {
                // Skip until end of declaration
                while (text[0] != Ch('?') || text[1] != Ch('>'))
                {
                    if (!text[0])
                        CEREAL_RAPIDXML_PARSE_ERROR("unexpected end of data", text);
                    ++text;
                }
                text += 2;    // Skip '?>'
                return 0;
            }

            // Create declaration
            xml_node<Ch> *declaration = this->allocate_node(node_declaration);

            // Skip whitespace before attributes or ?>
            skip<whitespace_pred, Flags>(text);

            // Parse declaration attributes
            parse_node_attributes<Flags>(text, declaration);

            // Skip ?>
            if (text[0] != Ch('?') || text[1] != Ch('>'))
                CEREAL_RAPIDXML_PARSE_ERROR("expected ?>", text);
            text += 2;

            return declaration;
        }

        // Parse XML comment (<!--...)
        template<int Flags>
        xml_node<Ch> *parse_comment(Ch *&text)
        {
            // If parsing of comments is disabled
            if (!(Flags & parse_comment_nodes))
            {
                // Skip until end of comment
                while (text[0] != Ch('-') || text[1] != Ch('-') || text[2] != Ch('>'))
                {
                    if (!text[0])
                        CEREAL_RAPIDXML_PARSE_ERROR("unexpected end of data", text);
                    ++text;
                }
                text += 3;     // Skip '-->'
                return 0;      // Do not produce comment node
            }

            // Remember value start
            Ch *value_ = text;

            // Skip until end of comment
            while (text[0] != Ch('-') || text[1] != Ch('-') || text[2] != Ch('>'))
            {
                if (!text[0])
                    CEREAL_RAPIDXML_PARSE_ERROR("unexpected end of data", text);
                ++text;
            }

            // Create comment node
            xml_node<Ch> *comment = this->allocate_node(node_comment);
            comment->value(value_, static_cast<std::size_t>(text - value_));

            // Place zero terminator after comment value
            if (!(Flags & parse_no_string_terminators))
                *text = Ch('\0');

            text += 3;     // Skip '-->'
            return comment;
        }

        // Parse DOCTYPE
        template<int Flags>
        xml_node<Ch> *parse_doctype(Ch *&text)
        {
            // Remember value start
            Ch *value_ = text;

            // Skip to >
            while (*text != Ch('>'))
            {
                // Determine character type
                switch (*text)
                {

                // If '[' encountered, scan for matching ending ']' using naive algorithm with depth
                // This works for all W3C test files except for 2 most wicked
                case Ch('['):
                {
                    ++text;     // Skip '['
                    int depth = 1;
                    while (depth > 0)
                    {
                        switch (*text)
                        {
                            case Ch('['): ++depth; break;
                            case Ch(']'): --depth; break;
                            case 0: CEREAL_RAPIDXML_PARSE_ERROR("unexpected end of data", text);
                        }
                        ++text;
                    }
                    break;
                }

                // Error on end of text
                case Ch('\0'):
                    CEREAL_RAPIDXML_PARSE_ERROR("unexpected end of data", text);

                // Other character, skip it
                default:
                    ++text;

                }
            }

            // If DOCTYPE nodes enabled
            if (Flags & parse_doctype_node)
            {
                // Create a new doctype node
                xml_node<Ch> *doctype = this->allocate_node(node_doctype);
                doctype->value(value_, static_cast<std::size_t>(text - value_));

                // Place zero terminator after value
                if (!(Flags & parse_no_string_terminators))
                    *text = Ch('\0');

                text += 1;      // skip '>'
                return doctype;
            }
            else
            {
                text += 1;      // skip '>'
                return 0;
            }

        }

        // Parse PI
        template<int Flags>
        xml_node<Ch> *parse_pi(Ch *&text)
        {
            // If creation of PI nodes is enabled
            if (Flags & parse_pi_nodes)
            {
                // Create pi node
                xml_node<Ch> *pi = this->allocate_node(node_pi);

                // Extract PI target name
                Ch *name_ = text;
                skip<node_name_pred, Flags>(text);
                if (text == name_)
                    CEREAL_RAPIDXML_PARSE_ERROR("expected PI target", text);
                pi->name(name_, static_cast<std::size_t>(text - name_));

                // Skip whitespace between pi target and pi
                skip<whitespace_pred, Flags>(text);

                // Remember start of pi
                Ch *value_ = text;

                // Skip to '?>'
                while (text[0] != Ch('?') || text[1] != Ch('>'))
                {
                    if (*text == Ch('\0'))
                        CEREAL_RAPIDXML_PARSE_ERROR("unexpected end of data", text);
                    ++text;
                }

                // Set pi value (verbatim, no entity expansion or whitespace normalization)
                pi->value(value_, static_cast<std::size_t>(text - value_));

                // Place zero terminator after name and value
                if (!(Flags & parse_no_string_terminators))
                {
                    pi->name()[pi->name_size()] = Ch('\0');
                    pi->value()[pi->value_size()] = Ch('\0');
                }

                text += 2;                          // Skip '?>'
                return pi;
            }
            else
            {
                // Skip to '?>'
                while (text[0] != Ch('?') || text[1] != Ch('>'))
                {
                    if (*text == Ch('\0'))
                        CEREAL_RAPIDXML_PARSE_ERROR("unexpected end of data", text);
                    ++text;
                }
                text += 2;    // Skip '?>'
                return 0;
            }
        }

        // Parse and append data
        // Return character that ends data.
        // This is necessary because this character might have been overwritten by a terminating 0
        template<int Flags>
        Ch parse_and_append_data(xml_node<Ch> *node, Ch *&text, Ch *contents_start)
        {
            // Backup to contents start if whitespace trimming is disabled
            if (!(Flags & parse_trim_whitespace))
                text = contents_start;

            const bool preserve_space =  internal::preserve_space(node);

            // Skip until end of data
            Ch *value_ = text, *end;
            if ((Flags & parse_normalize_whitespace) && !preserve_space)
                end = skip_and_expand_character_refs<text_pred, text_pure_with_ws_pred, Flags>(text, false);
            else
                end = skip_and_expand_character_refs<text_pred, text_pure_no_ws_pred, Flags>(text, preserve_space);

            // Trim trailing whitespace if flag is set; leading was already trimmed by whitespace skip after >
            if ((Flags & parse_trim_whitespace) && !preserve_space)
            {
                if (Flags & parse_normalize_whitespace)
                {
                    // Whitespace is already condensed to single space characters by skipping function, so just trim 1 char off the end
                    if (*(end - 1) == Ch(' '))
                        --end;
                }
                else
                {
                    // Backup until non-whitespace character is found
                    while (whitespace_pred::test(*(end - 1)))
                        --end;
                }
            }

            // If characters are still left between end and value (this test is only necessary if normalization is enabled)
            // Create new data node
            if (!(Flags & parse_no_data_nodes))
            {
                xml_node<Ch> *data = this->allocate_node(node_data);
                data->value(value_, static_cast<std::size_t>(end - value_));
                node->append_node(data);
            }

            // Add data to parent node if no data exists yet
            if (!(Flags & parse_no_element_values))
                if (*node->value() == Ch('\0'))
                    node->value(value_, static_cast<std::size_t>(end - value_));

            // Place zero terminator after value
            if (!(Flags & parse_no_string_terminators))
            {
                Ch ch = *text;
                *end = Ch('\0');
                return ch;      // Return character that ends data; this is required because zero terminator overwritten it
            }

            // Return character that ends data
            return *text;
        }

        // Parse CDATA
        template<int Flags>
        xml_node<Ch> *parse_cdata(Ch *&text)
        {
            // If CDATA is disabled
            if (Flags & parse_no_data_nodes)
            {
                // Skip until end of cdata
                while (text[0] != Ch(']') || text[1] != Ch(']') || text[2] != Ch('>'))
                {
                    if (!text[0])
                        CEREAL_RAPIDXML_PARSE_ERROR("unexpected end of data", text);
                    ++text;
                }
                text += 3;      // Skip ]]>
                return 0;       // Do not produce CDATA node
            }

            // Skip until end of cdata
            Ch *value_ = text;
            while (text[0] != Ch(']') || text[1] != Ch(']') || text[2] != Ch('>'))
            {
                if (!text[0])
                    CEREAL_RAPIDXML_PARSE_ERROR("unexpected end of data", text);
                ++text;
            }

            // Create new cdata node
            xml_node<Ch> *cdata = this->allocate_node(node_cdata);
            cdata->value(value_, static_cast<std::size_t>(text - value_));

            // Place zero terminator after value
            if (!(Flags & parse_no_string_terminators))
                *text = Ch('\0');

            text += 3;      // Skip ]]>
            return cdata;
        }

        // Parse element node
        template<int Flags>
        xml_node<Ch> *parse_element(Ch *&text)
        {
            // Create element node
            xml_node<Ch> *element = this->allocate_node(node_element);

            // Extract element name
            Ch *name_ = text;
            skip<node_name_pred, Flags>(text);
            if (text == name_)
                CEREAL_RAPIDXML_PARSE_ERROR("expected element name", text);
            element->name(name_, static_cast<std::size_t>(text - name_));

            // Skip whitespace between element name and attributes or >
            skip<whitespace_pred, Flags>(text);

            // Parse attributes, if any
            parse_node_attributes<Flags>(text, element);

            // Determine ending type
            if (*text == Ch('>'))
            {
                ++text;
                parse_node_contents<Flags>(text, element);
            }
            else if (*text == Ch('/'))
            {
                ++text;
                if (*text != Ch('>'))
                    CEREAL_RAPIDXML_PARSE_ERROR("expected >", text);
                ++text;
            }
            else
                CEREAL_RAPIDXML_PARSE_ERROR("expected >", text);

            // Place zero terminator after name
            if (!(Flags & parse_no_string_terminators))
                element->name()[element->name_size()] = Ch('\0');

            // Return parsed element
            return element;
        }

        // Determine node type, and parse it
        template<int Flags>
        xml_node<Ch> *parse_node(Ch *&text)
        {
            // Parse proper node type
            switch (text[0])
            {

            // <...
            default:
                // Parse and append element node
                return parse_element<Flags>(text);

            // <?...
            case Ch('?'):
                ++text;     // Skip ?
                if ((text[0] == Ch('x') || text[0] == Ch('X')) &&
                    (text[1] == Ch('m') || text[1] == Ch('M')) &&
                    (text[2] == Ch('l') || text[2] == Ch('L')) &&
                    whitespace_pred::test(text[3]))
                {
                    // '<?xml ' - xml declaration
                    text += 4;      // Skip 'xml '
                    return parse_xml_declaration<Flags>(text);
                }
                else
                {
                    // Parse PI
                    return parse_pi<Flags>(text);
                }

            // <!...
            case Ch('!'):

                // Parse proper subset of <! node
                switch (text[1])
                {

                // <!-
                case Ch('-'):
                    if (text[2] == Ch('-'))
                    {
                        // '<!--' - xml comment
                        text += 3;     // Skip '!--'
                        return parse_comment<Flags>(text);
                    }
                    break;

                // <![
                case Ch('['):
                    if (text[2] == Ch('C') && text[3] == Ch('D') && text[4] == Ch('A') &&
                        text[5] == Ch('T') && text[6] == Ch('A') && text[7] == Ch('['))
                    {
                        // '<![CDATA[' - cdata
                        text += 8;     // Skip '![CDATA['
                        return parse_cdata<Flags>(text);
                    }
                    break;

                // <!D
                case Ch('D'):
                    if (text[2] == Ch('O') && text[3] == Ch('C') && text[4] == Ch('T') &&
                        text[5] == Ch('Y') && text[6] == Ch('P') && text[7] == Ch('E') &&
                        whitespace_pred::test(text[8]))
                    {
                        // '<!DOCTYPE ' - doctype
                        text += 9;      // skip '!DOCTYPE '
                        return parse_doctype<Flags>(text);
                    }

                }   // switch

                // Attempt to skip other, unrecognized node types starting with <!
                ++text;     // Skip !
                while (*text != Ch('>'))
                {
                    if (*text == 0)
                        CEREAL_RAPIDXML_PARSE_ERROR("unexpected end of data", text);
                    ++text;
                }
                ++text;     // Skip '>'
                return 0;   // No node recognized

            }
        }

        // Parse contents of the node - children, data etc.
        template<int Flags>
        void parse_node_contents(Ch *&text, xml_node<Ch> *node)
        {
            // For all children and text
            while (1)
            {
                // Skip whitespace between > and node contents
                Ch *contents_start = text;      // Store start of node contents before whitespace is skipped
                skip<whitespace_pred, Flags>(text);
                Ch next_char = *text;

            // After data nodes, instead of continuing the loop, control jumps here.
            // This is because zero termination inside parse_and_append_data() function
            // would wreak havoc with the above code.
            // Also, skipping whitespace after data nodes is unnecessary.
            after_data_node:

                // Determine what comes next: node closing, child node, data node, or 0?
                switch (next_char)
                {

                // Node closing or child node
                case Ch('<'):
                    if (text[1] == Ch('/'))
                    {
                        Ch *contents_end = 0;
                        if (internal::preserve_space(node))
                        {
                            contents_end = text;
                        }

                        // Node closing
                        text += 2;      // Skip '</'
                        if (Flags & parse_validate_closing_tags)
                        {
                            // Skip and validate closing tag name
                            Ch *closing_name = text;
                            skip<node_name_pred, Flags>(text);
                            if (!internal::compare(node->name(), node->name_size(), closing_name, static_cast<std::size_t>(text - closing_name), true))
                                CEREAL_RAPIDXML_PARSE_ERROR("invalid closing tag name", text);
                        }
                        else
                        {
                            // No validation, just skip name
                            skip<node_name_pred, Flags>(text);
                        }
                        // Skip remaining whitespace after node name
                        skip<whitespace_pred, Flags>(text);
                        if (*text != Ch('>'))
                            CEREAL_RAPIDXML_PARSE_ERROR("expected >", text);
                        ++text;     // Skip '>'

                        if (contents_end && contents_end != contents_start)
                        {
                            node->value(contents_start, static_cast<std::size_t>(contents_end - contents_start));
                            node->value()[node->value_size()] = Ch('\0');
                        }
                        return;     // Node closed, finished parsing contents
                    }
                    else
                    {
                        // Child node
                        ++text;     // Skip '<'
                        if (xml_node<Ch> *child = parse_node<Flags>(text))
                            node->append_node(child);
                    }
                    break;

                // End of data - error
                case Ch('\0'):
                    CEREAL_RAPIDXML_PARSE_ERROR("unexpected end of data", text);

                // Data node
                default:
                    next_char = parse_and_append_data<Flags>(node, text, contents_start);
                    goto after_data_node;   // Bypass regular processing after data nodes

                }
            }
        }

        // Parse XML attributes of the node
        template<int Flags>
        void parse_node_attributes(Ch *&text, xml_node<Ch> *node)
        {
            // For all attributes
            while (attribute_name_pred::test(*text))
            {
                // Extract attribute name
                Ch *name_ = text;
                ++text;     // Skip first character of attribute name
                skip<attribute_name_pred, Flags>(text);
                if (text == name_)
                    CEREAL_RAPIDXML_PARSE_ERROR("expected attribute name", name_);

                // Create new attribute
                xml_attribute<Ch> *attribute = this->allocate_attribute();
                attribute->name(name_, static_cast<std::size_t>(text - name_));
                node->append_attribute(attribute);

                // Skip whitespace after attribute name
                skip<whitespace_pred, Flags>(text);

                // Skip =
                if (*text != Ch('='))
                    CEREAL_RAPIDXML_PARSE_ERROR("expected =", text);
                ++text;

                // Add terminating zero after name
                if (!(Flags & parse_no_string_terminators))
                    attribute->name()[attribute->name_size()] = 0;

                // Skip whitespace after =
                skip<whitespace_pred, Flags>(text);

                // Skip quote and remember if it was ' or "
                Ch quote = *text;
                if (quote != Ch('\'') && quote != Ch('"'))
                    CEREAL_RAPIDXML_PARSE_ERROR("expected ' or \"", text);
                ++text;

                // Extract attribute value and expand char refs in it
                Ch *value_ = text, *end;
                const int AttFlags = Flags & ~parse_normalize_whitespace;   // No whitespace normalization in attributes
                if (quote == Ch('\''))
                    end = skip_and_expand_character_refs<attribute_value_pred<Ch('\'')>, attribute_value_pure_pred<Ch('\'')>, AttFlags>(text, false);
                else
                    end = skip_and_expand_character_refs<attribute_value_pred<Ch('"')>, attribute_value_pure_pred<Ch('"')>, AttFlags>(text, false);

                // Set attribute value
                attribute->value(value_, static_cast<std::size_t>(end - value_));

                // Make sure that end quote is present
                if (*text != quote)
                    CEREAL_RAPIDXML_PARSE_ERROR("expected ' or \"", text);
                ++text;     // Skip quote

                // Add terminating zero after value
                if (!(Flags & parse_no_string_terminators))
                    attribute->value()[attribute->value_size()] = 0;

                // Skip whitespace after attribute value
                skip<whitespace_pred, Flags>(text);
            }
        }

    };

    //! \cond internal
    namespace internal
    {

        // Whitespace (space \n \r \t)
        template<int Dummy>
        const unsigned char lookup_tables<Dummy>::lookup_whitespace[256] =
        {
          // 0   1   2   3   4   5   6   7   8   9   A   B   C   D   E   F
             0,  0,  0,  0,  0,  0,  0,  0,  0,  1,  1,  0,  0,  1,  0,  0,  // 0
             0,  0,  0,  0,  0,  0,  0,  0,  0,  0,  0,  0,  0,  0,  0,  0,  // 1
             1,  0,  0,  0,  0,  0,  0,  0,  0,  0,  0,  0,  0,  0,  0,  0,  // 2
             0,  0,  0,  0,  0,  0,  0,  0,  0,  0,  0,  0,  0,  0,  0,  0,  // 3
             0,  0,  0,  0,  0,  0,  0,  0,  0,  0,  0,  0,  0,  0,  0,  0,  // 4
             0,  0,  0,  0,  0,  0,  0,  0,  0,  0,  0,  0,  0,  0,  0,  0,  // 5
             0,  0,  0,  0,  0,  0,  0,  0,  0,  0,  0,  0,  0,  0,  0,  0,  // 6
             0,  0,  0,  0,  0,  0,  0,  0,  0,  0,  0,  0,  0,  0,  0,  0,  // 7
             0,  0,  0,  0,  0,  0,  0,  0,  0,  0,  0,  0,  0,  0,  0,  0,  // 8
             0,  0,  0,  0,  0,  0,  0,  0,  0,  0,  0,  0,  0,  0,  0,  0,  // 9
             0,  0,  0,  0,  0,  0,  0,  0,  0,  0,  0,  0,  0,  0,  0,  0,  // A
             0,  0,  0,  0,  0,  0,  0,  0,  0,  0,  0,  0,  0,  0,  0,  0,  // B
             0,  0,  0,  0,  0,  0,  0,  0,  0,  0,  0,  0,  0,  0,  0,  0,  // C
             0,  0,  0,  0,  0,  0,  0,  0,  0,  0,  0,  0,  0,  0,  0,  0,  // D
             0,  0,  0,  0,  0,  0,  0,  0,  0,  0,  0,  0,  0,  0,  0,  0,  // E
             0,  0,  0,  0,  0,  0,  0,  0,  0,  0,  0,  0,  0,  0,  0,  0   // F
        };

        // Node name (anything but space \n \r \t / > ? \0)
        template<int Dummy>
        const unsigned char lookup_tables<Dummy>::lookup_node_name[256] =
        {
          // 0   1   2   3   4   5   6   7   8   9   A   B   C   D   E   F
             0,  1,  1,  1,  1,  1,  1,  1,  1,  0,  0,  1,  1,  0,  1,  1,  // 0
             1,  1,  1,  1,  1,  1,  1,  1,  1,  1,  1,  1,  1,  1,  1,  1,  // 1
             0,  1,  1,  1,  1,  1,  1,  1,  1,  1,  1,  1,  1,  1,  1,  0,  // 2
             1,  1,  1,  1,  1,  1,  1,  1,  1,  1,  1,  1,  1,  1,  0,  0,  // 3
             1,  1,  1,  1,  1,  1,  1,  1,  1,  1,  1,  1,  1,  1,  1,  1,  // 4
             1,  1,  1,  1,  1,  1,  1,  1,  1,  1,  1,  1,  1,  1,  1,  1,  // 5
             1,  1,  1,  1,  1,  1,  1,  1,  1,  1,  1,  1,  1,  1,  1,  1,  // 6
             1,  1,  1,  1,  1,  1,  1,  1,  1,  1,  1,  1,  1,  1,  1,  1,  // 7
             1,  1,  1,  1,  1,  1,  1,  1,  1,  1,  1,  1,  1,  1,  1,  1,  // 8
             1,  1,  1,  1,  1,  1,  1,  1,  1,  1,  1,  1,  1,  1,  1,  1,  // 9
             1,  1,  1,  1,  1,  1,  1,  1,  1,  1,  1,  1,  1,  1,  1,  1,  // A
             1,  1,  1,  1,  1,  1,  1,  1,  1,  1,  1,  1,  1,  1,  1,  1,  // B
             1,  1,  1,  1,  1,  1,  1,  1,  1,  1,  1,  1,  1,  1,  1,  1,  // C
             1,  1,  1,  1,  1,  1,  1,  1,  1,  1,  1,  1,  1,  1,  1,  1,  // D
             1,  1,  1,  1,  1,  1,  1,  1,  1,  1,  1,  1,  1,  1,  1,  1,  // E
             1,  1,  1,  1,  1,  1,  1,  1,  1,  1,  1,  1,  1,  1,  1,  1   // F
        };

        // Text (i.e. PCDATA) (anything but < \0)
        template<int Dummy>
        const unsigned char lookup_tables<Dummy>::lookup_text[256] =
        {
          // 0   1   2   3   4   5   6   7   8   9   A   B   C   D   E   F
             0,  1,  1,  1,  1,  1,  1,  1,  1,  1,  1,  1,  1,  1,  1,  1,  // 0
             1,  1,  1,  1,  1,  1,  1,  1,  1,  1,  1,  1,  1,  1,  1,  1,  // 1
             1,  1,  1,  1,  1,  1,  1,  1,  1,  1,  1,  1,  1,  1,  1,  1,  // 2
             1,  1,  1,  1,  1,  1,  1,  1,  1,  1,  1,  1,  0,  1,  1,  1,  // 3
             1,  1,  1,  1,  1,  1,  1,  1,  1,  1,  1,  1,  1,  1,  1,  1,  // 4
             1,  1,  1,  1,  1,  1,  1,  1,  1,  1,  1,  1,  1,  1,  1,  1,  // 5
             1,  1,  1,  1,  1,  1,  1,  1,  1,  1,  1,  1,  1,  1,  1,  1,  // 6
             1,  1,  1,  1,  1,  1,  1,  1,  1,  1,  1,  1,  1,  1,  1,  1,  // 7
             1,  1,  1,  1,  1,  1,  1,  1,  1,  1,  1,  1,  1,  1,  1,  1,  // 8
             1,  1,  1,  1,  1,  1,  1,  1,  1,  1,  1,  1,  1,  1,  1,  1,  // 9
             1,  1,  1,  1,  1,  1,  1,  1,  1,  1,  1,  1,  1,  1,  1,  1,  // A
             1,  1,  1,  1,  1,  1,  1,  1,  1,  1,  1,  1,  1,  1,  1,  1,  // B
             1,  1,  1,  1,  1,  1,  1,  1,  1,  1,  1,  1,  1,  1,  1,  1,  // C
             1,  1,  1,  1,  1,  1,  1,  1,  1,  1,  1,  1,  1,  1,  1,  1,  // D
             1,  1,  1,  1,  1,  1,  1,  1,  1,  1,  1,  1,  1,  1,  1,  1,  // E
             1,  1,  1,  1,  1,  1,  1,  1,  1,  1,  1,  1,  1,  1,  1,  1   // F
        };

        // Text (i.e. PCDATA) that does not require processing when ws normalization is disabled
        // (anything but < \0 &)
        template<int Dummy>
        const unsigned char lookup_tables<Dummy>::lookup_text_pure_no_ws[256] =
        {
          // 0   1   2   3   4   5   6   7   8   9   A   B   C   D   E   F
             0,  1,  1,  1,  1,  1,  1,  1,  1,  1,  1,  1,  1,  1,  1,  1,  // 0
             1,  1,  1,  1,  1,  1,  1,  1,  1,  1,  1,  1,  1,  1,  1,  1,  // 1
             1,  1,  1,  1,  1,  1,  0,  1,  1,  1,  1,  1,  1,  1,  1,  1,  // 2
             1,  1,  1,  1,  1,  1,  1,  1,  1,  1,  1,  1,  0,  1,  1,  1,  // 3
             1,  1,  1,  1,  1,  1,  1,  1,  1,  1,  1,  1,  1,  1,  1,  1,  // 4
             1,  1,  1,  1,  1,  1,  1,  1,  1,  1,  1,  1,  1,  1,  1,  1,  // 5
             1,  1,  1,  1,  1,  1,  1,  1,  1,  1,  1,  1,  1,  1,  1,  1,  // 6
             1,  1,  1,  1,  1,  1,  1,  1,  1,  1,  1,  1,  1,  1,  1,  1,  // 7
             1,  1,  1,  1,  1,  1,  1,  1,  1,  1,  1,  1,  1,  1,  1,  1,  // 8
             1,  1,  1,  1,  1,  1,  1,  1,  1,  1,  1,  1,  1,  1,  1,  1,  // 9
             1,  1,  1,  1,  1,  1,  1,  1,  1,  1,  1,  1,  1,  1,  1,  1,  // A
             1,  1,  1,  1,  1,  1,  1,  1,  1,  1,  1,  1,  1,  1,  1,  1,  // B
             1,  1,  1,  1,  1,  1,  1,  1,  1,  1,  1,  1,  1,  1,  1,  1,  // C
             1,  1,  1,  1,  1,  1,  1,  1,  1,  1,  1,  1,  1,  1,  1,  1,  // D
             1,  1,  1,  1,  1,  1,  1,  1,  1,  1,  1,  1,  1,  1,  1,  1,  // E
             1,  1,  1,  1,  1,  1,  1,  1,  1,  1,  1,  1,  1,  1,  1,  1   // F
        };

        // Text (i.e. PCDATA) that does not require processing when ws normalizationis is enabled
        // (anything but < \0 & space \n \r \t)
        template<int Dummy>
        const unsigned char lookup_tables<Dummy>::lookup_text_pure_with_ws[256] =
        {
          // 0   1   2   3   4   5   6   7   8   9   A   B   C   D   E   F
             0,  1,  1,  1,  1,  1,  1,  1,  1,  0,  0,  1,  1,  0,  1,  1,  // 0
             1,  1,  1,  1,  1,  1,  1,  1,  1,  1,  1,  1,  1,  1,  1,  1,  // 1
             0,  1,  1,  1,  1,  1,  0,  1,  1,  1,  1,  1,  1,  1,  1,  1,  // 2
             1,  1,  1,  1,  1,  1,  1,  1,  1,  1,  1,  1,  0,  1,  1,  1,  // 3
             1,  1,  1,  1,  1,  1,  1,  1,  1,  1,  1,  1,  1,  1,  1,  1,  // 4
             1,  1,  1,  1,  1,  1,  1,  1,  1,  1,  1,  1,  1,  1,  1,  1,  // 5
             1,  1,  1,  1,  1,  1,  1,  1,  1,  1,  1,  1,  1,  1,  1,  1,  // 6
             1,  1,  1,  1,  1,  1,  1,  1,  1,  1,  1,  1,  1,  1,  1,  1,  // 7
             1,  1,  1,  1,  1,  1,  1,  1,  1,  1,  1,  1,  1,  1,  1,  1,  // 8
             1,  1,  1,  1,  1,  1,  1,  1,  1,  1,  1,  1,  1,  1,  1,  1,  // 9
             1,  1,  1,  1,  1,  1,  1,  1,  1,  1,  1,  1,  1,  1,  1,  1,  // A
             1,  1,  1,  1,  1,  1,  1,  1,  1,  1,  1,  1,  1,  1,  1,  1,  // B
             1,  1,  1,  1,  1,  1,  1,  1,  1,  1,  1,  1,  1,  1,  1,  1,  // C
             1,  1,  1,  1,  1,  1,  1,  1,  1,  1,  1,  1,  1,  1,  1,  1,  // D
             1,  1,  1,  1,  1,  1,  1,  1,  1,  1,  1,  1,  1,  1,  1,  1,  // E
             1,  1,  1,  1,  1,  1,  1,  1,  1,  1,  1,  1,  1,  1,  1,  1   // F
        };

        // Attribute name (anything but space \n \r \t / < > = ? ! \0)
        template<int Dummy>
        const unsigned char lookup_tables<Dummy>::lookup_attribute_name[256] =
        {
          // 0   1   2   3   4   5   6   7   8   9   A   B   C   D   E   F
             0,  1,  1,  1,  1,  1,  1,  1,  1,  0,  0,  1,  1,  0,  1,  1,  // 0
             1,  1,  1,  1,  1,  1,  1,  1,  1,  1,  1,  1,  1,  1,  1,  1,  // 1
             0,  0,  1,  1,  1,  1,  1,  1,  1,  1,  1,  1,  1,  1,  1,  0,  // 2
             1,  1,  1,  1,  1,  1,  1,  1,  1,  1,  1,  1,  0,  0,  0,  0,  // 3
             1,  1,  1,  1,  1,  1,  1,  1,  1,  1,  1,  1,  1,  1,  1,  1,  // 4
             1,  1,  1,  1,  1,  1,  1,  1,  1,  1,  1,  1,  1,  1,  1,  1,  // 5
             1,  1,  1,  1,  1,  1,  1,  1,  1,  1,  1,  1,  1,  1,  1,  1,  // 6
             1,  1,  1,  1,  1,  1,  1,  1,  1,  1,  1,  1,  1,  1,  1,  1,  // 7
             1,  1,  1,  1,  1,  1,  1,  1,  1,  1,  1,  1,  1,  1,  1,  1,  // 8
             1,  1,  1,  1,  1,  1,  1,  1,  1,  1,  1,  1,  1,  1,  1,  1,  // 9
             1,  1,  1,  1,  1,  1,  1,  1,  1,  1,  1,  1,  1,  1,  1,  1,  // A
             1,  1,  1,  1,  1,  1,  1,  1,  1,  1,  1,  1,  1,  1,  1,  1,  // B
             1,  1,  1,  1,  1,  1,  1,  1,  1,  1,  1,  1,  1,  1,  1,  1,  // C
             1,  1,  1,  1,  1,  1,  1,  1,  1,  1,  1,  1,  1,  1,  1,  1,  // D
             1,  1,  1,  1,  1,  1,  1,  1,  1,  1,  1,  1,  1,  1,  1,  1,  // E
             1,  1,  1,  1,  1,  1,  1,  1,  1,  1,  1,  1,  1,  1,  1,  1   // F
        };

        // Attribute data with single quote (anything but ' \0)
        template<int Dummy>
        const unsigned char lookup_tables<Dummy>::lookup_attribute_data_1[256] =
        {
          // 0   1   2   3   4   5   6   7   8   9   A   B   C   D   E   F
             0,  1,  1,  1,  1,  1,  1,  1,  1,  1,  1,  1,  1,  1,  1,  1,  // 0
             1,  1,  1,  1,  1,  1,  1,  1,  1,  1,  1,  1,  1,  1,  1,  1,  // 1
             1,  1,  1,  1,  1,  1,  1,  0,  1,  1,  1,  1,  1,  1,  1,  1,  // 2
             1,  1,  1,  1,  1,  1,  1,  1,  1,  1,  1,  1,  1,  1,  1,  1,  // 3
             1,  1,  1,  1,  1,  1,  1,  1,  1,  1,  1,  1,  1,  1,  1,  1,  // 4
             1,  1,  1,  1,  1,  1,  1,  1,  1,  1,  1,  1,  1,  1,  1,  1,  // 5
             1,  1,  1,  1,  1,  1,  1,  1,  1,  1,  1,  1,  1,  1,  1,  1,  // 6
             1,  1,  1,  1,  1,  1,  1,  1,  1,  1,  1,  1,  1,  1,  1,  1,  // 7
             1,  1,  1,  1,  1,  1,  1,  1,  1,  1,  1,  1,  1,  1,  1,  1,  // 8
             1,  1,  1,  1,  1,  1,  1,  1,  1,  1,  1,  1,  1,  1,  1,  1,  // 9
             1,  1,  1,  1,  1,  1,  1,  1,  1,  1,  1,  1,  1,  1,  1,  1,  // A
             1,  1,  1,  1,  1,  1,  1,  1,  1,  1,  1,  1,  1,  1,  1,  1,  // B
             1,  1,  1,  1,  1,  1,  1,  1,  1,  1,  1,  1,  1,  1,  1,  1,  // C
             1,  1,  1,  1,  1,  1,  1,  1,  1,  1,  1,  1,  1,  1,  1,  1,  // D
             1,  1,  1,  1,  1,  1,  1,  1,  1,  1,  1,  1,  1,  1,  1,  1,  // E
             1,  1,  1,  1,  1,  1,  1,  1,  1,  1,  1,  1,  1,  1,  1,  1   // F
        };

        // Attribute data with single quote that does not require processing (anything but ' \0 &)
        template<int Dummy>
        const unsigned char lookup_tables<Dummy>::lookup_attribute_data_1_pure[256] =
        {
          // 0   1   2   3   4   5   6   7   8   9   A   B   C   D   E   F
             0,  1,  1,  1,  1,  1,  1,  1,  1,  1,  1,  1,  1,  1,  1,  1,  // 0
             1,  1,  1,  1,  1,  1,  1,  1,  1,  1,  1,  1,  1,  1,  1,  1,  // 1
             1,  1,  1,  1,  1,  1,  0,  0,  1,  1,  1,  1,  1,  1,  1,  1,  // 2
             1,  1,  1,  1,  1,  1,  1,  1,  1,  1,  1,  1,  1,  1,  1,  1,  // 3
             1,  1,  1,  1,  1,  1,  1,  1,  1,  1,  1,  1,  1,  1,  1,  1,  // 4
             1,  1,  1,  1,  1,  1,  1,  1,  1,  1,  1,  1,  1,  1,  1,  1,  // 5
             1,  1,  1,  1,  1,  1,  1,  1,  1,  1,  1,  1,  1,  1,  1,  1,  // 6
             1,  1,  1,  1,  1,  1,  1,  1,  1,  1,  1,  1,  1,  1,  1,  1,  // 7
             1,  1,  1,  1,  1,  1,  1,  1,  1,  1,  1,  1,  1,  1,  1,  1,  // 8
             1,  1,  1,  1,  1,  1,  1,  1,  1,  1,  1,  1,  1,  1,  1,  1,  // 9
             1,  1,  1,  1,  1,  1,  1,  1,  1,  1,  1,  1,  1,  1,  1,  1,  // A
             1,  1,  1,  1,  1,  1,  1,  1,  1,  1,  1,  1,  1,  1,  1,  1,  // B
             1,  1,  1,  1,  1,  1,  1,  1,  1,  1,  1,  1,  1,  1,  1,  1,  // C
             1,  1,  1,  1,  1,  1,  1,  1,  1,  1,  1,  1,  1,  1,  1,  1,  // D
             1,  1,  1,  1,  1,  1,  1,  1,  1,  1,  1,  1,  1,  1,  1,  1,  // E
             1,  1,  1,  1,  1,  1,  1,  1,  1,  1,  1,  1,  1,  1,  1,  1   // F
        };

        // Attribute data with double quote (anything but " \0)
        template<int Dummy>
        const unsigned char lookup_tables<Dummy>::lookup_attribute_data_2[256] =
        {
          // 0   1   2   3   4   5   6   7   8   9   A   B   C   D   E   F
             0,  1,  1,  1,  1,  1,  1,  1,  1,  1,  1,  1,  1,  1,  1,  1,  // 0
             1,  1,  1,  1,  1,  1,  1,  1,  1,  1,  1,  1,  1,  1,  1,  1,  // 1
             1,  1,  0,  1,  1,  1,  1,  1,  1,  1,  1,  1,  1,  1,  1,  1,  // 2
             1,  1,  1,  1,  1,  1,  1,  1,  1,  1,  1,  1,  1,  1,  1,  1,  // 3
             1,  1,  1,  1,  1,  1,  1,  1,  1,  1,  1,  1,  1,  1,  1,  1,  // 4
             1,  1,  1,  1,  1,  1,  1,  1,  1,  1,  1,  1,  1,  1,  1,  1,  // 5
             1,  1,  1,  1,  1,  1,  1,  1,  1,  1,  1,  1,  1,  1,  1,  1,  // 6
             1,  1,  1,  1,  1,  1,  1,  1,  1,  1,  1,  1,  1,  1,  1,  1,  // 7
             1,  1,  1,  1,  1,  1,  1,  1,  1,  1,  1,  1,  1,  1,  1,  1,  // 8
             1,  1,  1,  1,  1,  1,  1,  1,  1,  1,  1,  1,  1,  1,  1,  1,  // 9
             1,  1,  1,  1,  1,  1,  1,  1,  1,  1,  1,  1,  1,  1,  1,  1,  // A
             1,  1,  1,  1,  1,  1,  1,  1,  1,  1,  1,  1,  1,  1,  1,  1,  // B
             1,  1,  1,  1,  1,  1,  1,  1,  1,  1,  1,  1,  1,  1,  1,  1,  // C
             1,  1,  1,  1,  1,  1,  1,  1,  1,  1,  1,  1,  1,  1,  1,  1,  // D
             1,  1,  1,  1,  1,  1,  1,  1,  1,  1,  1,  1,  1,  1,  1,  1,  // E
             1,  1,  1,  1,  1,  1,  1,  1,  1,  1,  1,  1,  1,  1,  1,  1   // F
        };

        // Attribute data with double quote that does not require processing (anything but " \0 &)
        template<int Dummy>
        const unsigned char lookup_tables<Dummy>::lookup_attribute_data_2_pure[256] =
        {
          // 0   1   2   3   4   5   6   7   8   9   A   B   C   D   E   F
             0,  1,  1,  1,  1,  1,  1,  1,  1,  1,  1,  1,  1,  1,  1,  1,  // 0
             1,  1,  1,  1,  1,  1,  1,  1,  1,  1,  1,  1,  1,  1,  1,  1,  // 1
             1,  1,  0,  1,  1,  1,  0,  1,  1,  1,  1,  1,  1,  1,  1,  1,  // 2
             1,  1,  1,  1,  1,  1,  1,  1,  1,  1,  1,  1,  1,  1,  1,  1,  // 3
             1,  1,  1,  1,  1,  1,  1,  1,  1,  1,  1,  1,  1,  1,  1,  1,  // 4
             1,  1,  1,  1,  1,  1,  1,  1,  1,  1,  1,  1,  1,  1,  1,  1,  // 5
             1,  1,  1,  1,  1,  1,  1,  1,  1,  1,  1,  1,  1,  1,  1,  1,  // 6
             1,  1,  1,  1,  1,  1,  1,  1,  1,  1,  1,  1,  1,  1,  1,  1,  // 7
             1,  1,  1,  1,  1,  1,  1,  1,  1,  1,  1,  1,  1,  1,  1,  1,  // 8
             1,  1,  1,  1,  1,  1,  1,  1,  1,  1,  1,  1,  1,  1,  1,  1,  // 9
             1,  1,  1,  1,  1,  1,  1,  1,  1,  1,  1,  1,  1,  1,  1,  1,  // A
             1,  1,  1,  1,  1,  1,  1,  1,  1,  1,  1,  1,  1,  1,  1,  1,  // B
             1,  1,  1,  1,  1,  1,  1,  1,  1,  1,  1,  1,  1,  1,  1,  1,  // C
             1,  1,  1,  1,  1,  1,  1,  1,  1,  1,  1,  1,  1,  1,  1,  1,  // D
             1,  1,  1,  1,  1,  1,  1,  1,  1,  1,  1,  1,  1,  1,  1,  1,  // E
             1,  1,  1,  1,  1,  1,  1,  1,  1,  1,  1,  1,  1,  1,  1,  1   // F
        };

        // Digits (dec and hex, 255 denotes end of numeric character reference)
        template<int Dummy>
        const unsigned char lookup_tables<Dummy>::lookup_digits[256] =
        {
          // 0   1   2   3   4   5   6   7   8   9   A   B   C   D   E   F
           255,255,255,255,255,255,255,255,255,255,255,255,255,255,255,255,  // 0
           255,255,255,255,255,255,255,255,255,255,255,255,255,255,255,255,  // 1
           255,255,255,255,255,255,255,255,255,255,255,255,255,255,255,255,  // 2
             0,  1,  2,  3,  4,  5,  6,  7,  8,  9,255,255,255,255,255,255,  // 3
           255, 10, 11, 12, 13, 14, 15,255,255,255,255,255,255,255,255,255,  // 4
           255,255,255,255,255,255,255,255,255,255,255,255,255,255,255,255,  // 5
           255, 10, 11, 12, 13, 14, 15,255,255,255,255,255,255,255,255,255,  // 6
           255,255,255,255,255,255,255,255,255,255,255,255,255,255,255,255,  // 7
           255,255,255,255,255,255,255,255,255,255,255,255,255,255,255,255,  // 8
           255,255,255,255,255,255,255,255,255,255,255,255,255,255,255,255,  // 9
           255,255,255,255,255,255,255,255,255,255,255,255,255,255,255,255,  // A
           255,255,255,255,255,255,255,255,255,255,255,255,255,255,255,255,  // B
           255,255,255,255,255,255,255,255,255,255,255,255,255,255,255,255,  // C
           255,255,255,255,255,255,255,255,255,255,255,255,255,255,255,255,  // D
           255,255,255,255,255,255,255,255,255,255,255,255,255,255,255,255,  // E
           255,255,255,255,255,255,255,255,255,255,255,255,255,255,255,255   // F
        };

        // Upper case conversion
        template<int Dummy>
        const unsigned char lookup_tables<Dummy>::lookup_upcase[256] =
        {
          // 0,  1,  2,  3,  4,  5,  6,  7,  8,  9,  A   B   C   D   E   F
           0,  1,  2,  3,  4,  5,  6,  7,  8,  9,  10, 11, 12, 13, 14, 15,   // 0
           16, 17, 18, 19, 20, 21, 22, 23, 24, 25, 26, 27, 28, 29, 30, 31,   // 1
           32, 33, 34, 35, 36, 37, 38, 39, 40, 41, 42, 43, 44, 45, 46, 47,   // 2
           48, 49, 50, 51, 52, 53, 54, 55, 56, 57, 58, 59, 60, 61, 62, 63,   // 3
           64, 65, 66, 67, 68, 69, 70, 71, 72, 73, 74, 75, 76, 77, 78, 79,   // 4
           80, 81, 82, 83, 84, 85, 86, 87, 88, 89, 90, 91, 92, 93, 94, 95,   // 5
           96, 65, 66, 67, 68, 69, 70, 71, 72, 73, 74, 75, 76, 77, 78, 79,   // 6
           80, 81, 82, 83, 84, 85, 86, 87, 88, 89, 90, 123,124,125,126,127,  // 7
           128,129,130,131,132,133,134,135,136,137,138,139,140,141,142,143,  // 8
           144,145,146,147,148,149,150,151,152,153,154,155,156,157,158,159,  // 9
           160,161,162,163,164,165,166,167,168,169,170,171,172,173,174,175,  // A
           176,177,178,179,180,181,182,183,184,185,186,187,188,189,190,191,  // B
           192,193,194,195,196,197,198,199,200,201,202,203,204,205,206,207,  // C
           208,209,210,211,212,213,214,215,216,217,218,219,220,221,222,223,  // D
           224,225,226,227,228,229,230,231,232,233,234,235,236,237,238,239,  // E
           240,241,242,243,244,245,246,247,248,249,250,251,252,253,254,255   // F
        };
    }
    //! \endcond

}
} // end namespace cereal

// Undefine internal macros
#undef CEREAL_RAPIDXML_PARSE_ERROR

// On MSVC, restore warnings state
#ifdef _MSC_VER
    #pragma warning(pop)
#endif

#endif
```
