## Supplemental code for "The motif composition of variable-number tandem repeats impacts gene expression": rapidxml_iterators.hpp

```
#ifndef CEREAL_RAPIDXML_ITERATORS_HPP_INCLUDED
#define CEREAL_RAPIDXML_ITERATORS_HPP_INCLUDED

// Copyright (C) 2006, 2009 Marcin Kalicinski
// Version 1.13
// Revision $DateTime: 2009/05/13 01:46:17 $

#include "rapidxml.hpp"

namespace cereal {
namespace rapidxml
{

    //! Iterator of child nodes of xml_node
    template<class Ch>
    class node_iterator
    {

    public:

        typedef typename xml_node<Ch> value_type;
        typedef typename xml_node<Ch> &reference;
        typedef typename xml_node<Ch> *pointer;
        typedef std::ptrdiff_t difference_type;
        typedef std::bidirectional_iterator_tag iterator_category;

        node_iterator()
            : m_node(0)
        {
        }

        node_iterator(xml_node<Ch> *node)
            : m_node(node->first_node())
        {
        }

        reference operator *() const
        {
            assert(m_node);
            return *m_node;
        }

        pointer operator->() const
        {
            assert(m_node);
            return m_node;
        }

        node_iterator& operator++()
        {
            assert(m_node);
            m_node = m_node->next_sibling();
            return *this;
        }

        node_iterator operator++(int)
        {
            node_iterator tmp = *this;
            ++this;
            return tmp;
        }

        node_iterator& operator--()
        {
            assert(m_node && m_node->previous_sibling());
            m_node = m_node->previous_sibling();
            return *this;
        }

        node_iterator operator--(int)
        {
            node_iterator tmp = *this;
            ++this;
            return tmp;
        }

        bool operator ==(const node_iterator<Ch> &rhs)
        {
            return m_node == rhs.m_node;
        }

        bool operator !=(const node_iterator<Ch> &rhs)
        {
            return m_node != rhs.m_node;
        }

    private:

        xml_node<Ch> *m_node;

    };

    //! Iterator of child attributes of xml_node
    template<class Ch>
    class attribute_iterator
    {

    public:

        typedef typename xml_attribute<Ch> value_type;
        typedef typename xml_attribute<Ch> &reference;
        typedef typename xml_attribute<Ch> *pointer;
        typedef std::ptrdiff_t difference_type;
        typedef std::bidirectional_iterator_tag iterator_category;

        attribute_iterator()
            : m_attribute(0)
        {
        }

        attribute_iterator(xml_node<Ch> *node)
            : m_attribute(node->first_attribute())
        {
        }

        reference operator *() const
        {
            assert(m_attribute);
            return *m_attribute;
        }

        pointer operator->() const
        {
            assert(m_attribute);
            return m_attribute;
        }

        attribute_iterator& operator++()
        {
            assert(m_attribute);
            m_attribute = m_attribute->next_attribute();
            return *this;
        }

        attribute_iterator operator++(int)
        {
            attribute_iterator tmp = *this;
            ++this;
            return tmp;
        }

        attribute_iterator& operator--()
        {
            assert(m_attribute && m_attribute->previous_attribute());
            m_attribute = m_attribute->previous_attribute();
            return *this;
        }

        attribute_iterator operator--(int)
        {
            attribute_iterator tmp = *this;
            ++this;
            return tmp;
        }

        bool operator ==(const attribute_iterator<Ch> &rhs)
        {
            return m_attribute == rhs.m_attribute;
        }

        bool operator !=(const attribute_iterator<Ch> &rhs)
        {
            return m_attribute != rhs.m_attribute;
        }

    private:

        xml_attribute<Ch> *m_attribute;

    };

}
} // namespace cereal

#endif
```
