## Supplemental code for "The motif composition of variable-number tandem repeats impacts gene expression": rapidxml_print.hpp

```
#ifndef CEREAL_RAPIDXML_PRINT_HPP_INCLUDED
#define CEREAL_RAPIDXML_PRINT_HPP_INCLUDED

// Copyright (C) 2006, 2009 Marcin Kalicinski
// Version 1.13
// Revision $DateTime: 2009/05/13 01:46:17 $

#include "rapidxml.hpp"

// Only include streams if not disabled
#ifndef CEREAL_RAPIDXML_NO_STREAMS
    #include <ostream>
    #include <iterator>
#endif

namespace cereal {
namespace rapidxml
{

    ///////////////////////////////////////////////////////////////////////
    // Printing flags

    const int print_no_indenting = 0x1;   //!< Printer flag instructing the printer to suppress indenting of XML. See print() function.

    ///////////////////////////////////////////////////////////////////////
    // Internal

    //! \cond internal
    namespace internal
    {

        ///////////////////////////////////////////////////////////////////////////
        // Internal character operations

        // Copy characters from given range to given output iterator
        template<class OutIt, class Ch>
        inline OutIt copy_chars(const Ch *begin, const Ch *end, OutIt out)
        {
            while (begin != end)
                *out++ = *begin++;
            return out;
        }

        // Copy characters from given range to given output iterator and expand
        // characters into references (&lt; &gt; &apos; &quot; &amp;)
        template<class OutIt, class Ch>
        inline OutIt copy_and_expand_chars(const Ch *begin, const Ch *end, Ch noexpand, OutIt out)
        {
            while (begin != end)
            {
                if (*begin == noexpand)
                {
                    *out++ = *begin;    // No expansion, copy character
                }
                else
                {
                    switch (*begin)
                    {
                    case Ch('<'):
                        *out++ = Ch('&'); *out++ = Ch('l'); *out++ = Ch('t'); *out++ = Ch(';');
                        break;
                    case Ch('>'):
                        *out++ = Ch('&'); *out++ = Ch('g'); *out++ = Ch('t'); *out++ = Ch(';');
                        break;
                    case Ch('\''):
                        *out++ = Ch('&'); *out++ = Ch('a'); *out++ = Ch('p'); *out++ = Ch('o'); *out++ = Ch('s'); *out++ = Ch(';');
                        break;
                    case Ch('"'):
                        *out++ = Ch('&'); *out++ = Ch('q'); *out++ = Ch('u'); *out++ = Ch('o'); *out++ = Ch('t'); *out++ = Ch(';');
                        break;
                    case Ch('&'):
                        *out++ = Ch('&'); *out++ = Ch('a'); *out++ = Ch('m'); *out++ = Ch('p'); *out++ = Ch(';');
                        break;
                    default:
                        *out++ = *begin;    // No expansion, copy character
                    }
                }
                ++begin;    // Step to next character
            }
            return out;
        }

        // Fill given output iterator with repetitions of the same character
        template<class OutIt, class Ch>
        inline OutIt fill_chars(OutIt out, int n, Ch ch)
        {
            for (int i = 0; i < n; ++i)
                *out++ = ch;
            return out;
        }

        // Find character
        template<class Ch, Ch ch>
        inline bool find_char(const Ch *begin, const Ch *end)
        {
            while (begin != end)
                if (*begin++ == ch)
                    return true;
            return false;
        }

        ///////////////////////////////////////////////////////////////////////////
        // Internal printing operations

        // Print node
        template<class OutIt, class Ch>
        inline OutIt print_node(OutIt out, const xml_node<Ch> *node, int flags, int indent);

        // Print children of the node
        template<class OutIt, class Ch>
        inline OutIt print_children(OutIt out, const xml_node<Ch> *node, int flags, int indent)
        {
            for (xml_node<Ch> *child = node->first_node(); child; child = child->next_sibling())
                out = print_node(out, child, flags, indent);
            return out;
        }

        // Print attributes of the node
        template<class OutIt, class Ch>
        inline OutIt print_attributes(OutIt out, const xml_node<Ch> *node, int /*flags*/)
        {
            for (xml_attribute<Ch> *attribute = node->first_attribute(); attribute; attribute = attribute->next_attribute())
            {
                if (attribute->name() && attribute->value())
                {
                    // Print attribute name
                    *out = Ch(' '), ++out;
                    out = copy_chars(attribute->name(), attribute->name() + attribute->name_size(), out);
                    *out = Ch('='), ++out;
                    // Print attribute value using appropriate quote type
                    if (find_char<Ch, Ch('"')>(attribute->value(), attribute->value() + attribute->value_size()))
                    {
                        *out = Ch('\''), ++out;
                        out = copy_and_expand_chars(attribute->value(), attribute->value() + attribute->value_size(), Ch('"'), out);
                        *out = Ch('\''), ++out;
                    }
                    else
                    {
                        *out = Ch('"'), ++out;
                        out = copy_and_expand_chars(attribute->value(), attribute->value() + attribute->value_size(), Ch('\''), out);
                        *out = Ch('"'), ++out;
                    }
                }
            }
            return out;
        }

        // Print data node
        template<class OutIt, class Ch>
        inline OutIt print_data_node(OutIt out, const xml_node<Ch> *node, int flags, int indent)
        {
            assert(node->type() == node_data);
            if (!(flags & print_no_indenting))
                out = fill_chars(out, indent, Ch('\t'));
            out = copy_and_expand_chars(node->value(), node->value() + node->value_size(), Ch(0), out);
            return out;
        }

        // Print data node
        template<class OutIt, class Ch>
        inline OutIt print_cdata_node(OutIt out, const xml_node<Ch> *node, int flags, int indent)
        {
            assert(node->type() == node_cdata);
            if (!(flags & print_no_indenting))
                out = fill_chars(out, indent, Ch('\t'));
            *out = Ch('<'); ++out;
            *out = Ch('!'); ++out;
            *out = Ch('['); ++out;
            *out = Ch('C'); ++out;
            *out = Ch('D'); ++out;
            *out = Ch('A'); ++out;
            *out = Ch('T'); ++out;
            *out = Ch('A'); ++out;
            *out = Ch('['); ++out;
            out = copy_chars(node->value(), node->value() + node->value_size(), out);
            *out = Ch(']'); ++out;
            *out = Ch(']'); ++out;
            *out = Ch('>'); ++out;
            return out;
        }

        // Print element node
        template<class OutIt, class Ch>
        inline OutIt print_element_node(OutIt out, const xml_node<Ch> *node, int flags, int indent)
        {
            assert(node->type() == node_element);

            // Print element name and attributes, if any
            if (!(flags & print_no_indenting))
                out = fill_chars(out, indent, Ch('\t'));
            *out = Ch('<'), ++out;
            out = copy_chars(node->name(), node->name() + node->name_size(), out);
            out = print_attributes(out, node, flags);

            // If node is childless
            if (node->value_size() == 0 && !node->first_node())
            {
                // Print childless node tag ending
                *out = Ch('/'), ++out;
                *out = Ch('>'), ++out;
            }
            else
            {
                // Print normal node tag ending
                *out = Ch('>'), ++out;

                // Test if node contains a single data node only (and no other nodes)
                xml_node<Ch> *child = node->first_node();
                if (!child)
                {
                    // If node has no children, only print its value without indenting
                    out = copy_and_expand_chars(node->value(), node->value() + node->value_size(), Ch(0), out);
                }
                else if (child->next_sibling() == 0 && child->type() == node_data)
                {
                    // If node has a sole data child, only print its value without indenting
                    out = copy_and_expand_chars(child->value(), child->value() + child->value_size(), Ch(0), out);
                }
                else
                {
                    // Print all children with full indenting
                    if (!(flags & print_no_indenting))
                        *out = Ch('\n'), ++out;
                    out = print_children(out, node, flags, indent + 1);
                    if (!(flags & print_no_indenting))
                        out = fill_chars(out, indent, Ch('\t'));
                }

                // Print node end
                *out = Ch('<'), ++out;
                *out = Ch('/'), ++out;
                out = copy_chars(node->name(), node->name() + node->name_size(), out);
                *out = Ch('>'), ++out;
            }
            return out;
        }

        // Print declaration node
        template<class OutIt, class Ch>
        inline OutIt print_declaration_node(OutIt out, const xml_node<Ch> *node, int flags, int indent)
        {
            // Print declaration start
            if (!(flags & print_no_indenting))
                out = fill_chars(out, indent, Ch('\t'));
            *out = Ch('<'), ++out;
            *out = Ch('?'), ++out;
            *out = Ch('x'), ++out;
            *out = Ch('m'), ++out;
            *out = Ch('l'), ++out;

            // Print attributes
            out = print_attributes(out, node, flags);

            // Print declaration end
            *out = Ch('?'), ++out;
            *out = Ch('>'), ++out;

            return out;
        }

        // Print comment node
        template<class OutIt, class Ch>
        inline OutIt print_comment_node(OutIt out, const xml_node<Ch> *node, int flags, int indent)
        {
            assert(node->type() == node_comment);
            if (!(flags & print_no_indenting))
                out = fill_chars(out, indent, Ch('\t'));
            *out = Ch('<'), ++out;
            *out = Ch('!'), ++out;
            *out = Ch('-'), ++out;
            *out = Ch('-'), ++out;
            out = copy_chars(node->value(), node->value() + node->value_size(), out);
            *out = Ch('-'), ++out;
            *out = Ch('-'), ++out;
            *out = Ch('>'), ++out;
            return out;
        }

        // Print doctype node
        template<class OutIt, class Ch>
        inline OutIt print_doctype_node(OutIt out, const xml_node<Ch> *node, int flags, int indent)
        {
            assert(node->type() == node_doctype);
            if (!(flags & print_no_indenting))
                out = fill_chars(out, indent, Ch('\t'));
            *out = Ch('<'), ++out;
            *out = Ch('!'), ++out;
            *out = Ch('D'), ++out;
            *out = Ch('O'), ++out;
            *out = Ch('C'), ++out;
            *out = Ch('T'), ++out;
            *out = Ch('Y'), ++out;
            *out = Ch('P'), ++out;
            *out = Ch('E'), ++out;
            *out = Ch(' '), ++out;
            out = copy_chars(node->value(), node->value() + node->value_size(), out);
            *out = Ch('>'), ++out;
            return out;
        }

        // Print pi node
        template<class OutIt, class Ch>
        inline OutIt print_pi_node(OutIt out, const xml_node<Ch> *node, int flags, int indent)
        {
            assert(node->type() == node_pi);
            if (!(flags & print_no_indenting))
                out = fill_chars(out, indent, Ch('\t'));
            *out = Ch('<'), ++out;
            *out = Ch('?'), ++out;
            out = copy_chars(node->name(), node->name() + node->name_size(), out);
            *out = Ch(' '), ++out;
            out = copy_chars(node->value(), node->value() + node->value_size(), out);
            *out = Ch('?'), ++out;
            *out = Ch('>'), ++out;
            return out;
        }

        // Print node
        template<class OutIt, class Ch>
        inline OutIt print_node(OutIt out, const xml_node<Ch> *node, int flags, int indent)
        {
            // Print proper node type
            switch (node->type())
            {

            // Document
            case node_document:
                out = print_children(out, node, flags, indent);
                break;

            // Element
            case node_element:
                out = print_element_node(out, node, flags, indent);
                break;

            // Data
            case node_data:
                out = print_data_node(out, node, flags, indent);
                break;

            // CDATA
            case node_cdata:
                out = print_cdata_node(out, node, flags, indent);
                break;

            // Declaration
            case node_declaration:
                out = print_declaration_node(out, node, flags, indent);
                break;

            // Comment
            case node_comment:
                out = print_comment_node(out, node, flags, indent);
                break;

            // Doctype
            case node_doctype:
                out = print_doctype_node(out, node, flags, indent);
                break;

            // Pi
            case node_pi:
                out = print_pi_node(out, node, flags, indent);
                break;

#ifndef __GNUC__
                // Unknown
            default:
                assert(0);
                break;
#endif
            }

            // If indenting not disabled, add line break after node
            if (!(flags & print_no_indenting))
                *out = Ch('\n'), ++out;

            // Return modified iterator
            return out;
        }

    }
    //! \endcond

    ///////////////////////////////////////////////////////////////////////////
    // Printing

    //! Prints XML to given output iterator.
    //! \param out Output iterator to print to.
    //! \param node Node to be printed. Pass xml_document to print entire document.
    //! \param flags Flags controlling how XML is printed.
    //! \return Output iterator pointing to position immediately after last character of printed text.
    template<class OutIt, class Ch>
    inline OutIt print(OutIt out, const xml_node<Ch> &node, int flags = 0)
    {
        return internal::print_node(out, &node, flags, 0);
    }

#ifndef CEREAL_RAPIDXML_NO_STREAMS

    //! Prints XML to given output stream.
    //! \param out Output stream to print to.
    //! \param node Node to be printed. Pass xml_document to print entire document.
    //! \param flags Flags controlling how XML is printed.
    //! \return Output stream.
    template<class Ch>
    inline std::basic_ostream<Ch> &print(std::basic_ostream<Ch> &out, const xml_node<Ch> &node, int flags = 0)
    {
        print(std::ostream_iterator<Ch>(out), node, flags);
        return out;
    }

    //! Prints formatted XML to given output stream. Uses default printing flags. Use print() function to customize printing process.
    //! \param out Output stream to print to.
    //! \param node Node to be printed.
    //! \return Output stream.
    template<class Ch>
    inline std::basic_ostream<Ch> &operator <<(std::basic_ostream<Ch> &out, const xml_node<Ch> &node)
    {
        return print(out, node);
    }

#endif

}
} // namespace cereal

#endif
```
