## Supplemental code for "The motif composition of variable-number tandem repeats impacts gene expression": rapidxml_utils.hpp

```
#ifndef CEREAL_RAPIDXML_UTILS_HPP_INCLUDED
#define CEREAL_RAPIDXML_UTILS_HPP_INCLUDED

// Copyright (C) 2006, 2009 Marcin Kalicinski
// Version 1.13
// Revision $DateTime: 2009/05/13 01:46:17 $
//! in certain simple scenarios. They should probably not be used if maximizing performance is the main objective.

#include "rapidxml.hpp"
#include <vector>
#include <string>
#include <fstream>
#include <stdexcept>

namespace cereal {
namespace rapidxml
{

    //! Represents data loaded from a file
    template<class Ch = char>
    class file
    {

    public:

        //! Loads file into the memory. Data will be automatically destroyed by the destructor.
        //! \param filename Filename to load.
        file(const char *filename)
        {
            using namespace std;

            // Open stream
            basic_ifstream<Ch> stream(filename, ios::binary);
            if (!stream)
                throw runtime_error(string("cannot open file ") + filename);
            stream.unsetf(ios::skipws);

            // Determine stream size
            stream.seekg(0, ios::end);
            size_t size = stream.tellg();
            stream.seekg(0);

            // Load data and add terminating 0
            m_data.resize(size + 1);
            stream.read(&m_data.front(), static_cast<streamsize>(size));
            m_data[size] = 0;
        }

        //! Loads file into the memory. Data will be automatically destroyed by the destructor
        //! \param stream Stream to load from
        file(std::basic_istream<Ch> &stream)
        {
            using namespace std;

            // Load data and add terminating 0
            stream.unsetf(ios::skipws);
            m_data.assign(istreambuf_iterator<Ch>(stream), istreambuf_iterator<Ch>());
            if (stream.fail() || stream.bad())
                throw runtime_error("error reading stream");
            m_data.push_back(0);
        }

        //! Gets file data.
        //! \return Pointer to data of file.
        Ch *data()
        {
            return &m_data.front();
        }

        //! Gets file data.
        //! \return Pointer to data of file.
        const Ch *data() const
        {
            return &m_data.front();
        }

        //! Gets file data size.
        //! \return Size of file data, in characters.
        std::size_t size() const
        {
            return m_data.size();
        }

    private:

        std::vector<Ch> m_data;   // File data

    };

    //! Counts children of node. Time complexity is O(n).
    //! \return Number of children of node
    template<class Ch>
    inline std::size_t count_children(xml_node<Ch> *node)
    {
        xml_node<Ch> *child = node->first_node();
        std::size_t count = 0;
        while (child)
        {
            ++count;
            child = child->next_sibling();
        }
        return count;
    }

    //! Counts attributes of node. Time complexity is O(n).
    //! \return Number of attributes of node
    template<class Ch>
    inline std::size_t count_attributes(xml_node<Ch> *node)
    {
        xml_attribute<Ch> *attr = node->first_attribute();
        std::size_t count = 0;
        while (attr)
        {
            ++count;
            attr = attr->next_attribute();
        }
        return count;
    }

}
} // namespace cereal

#endif
```
