## Supplemental code for "The motif composition of variable-number tandem repeats impacts gene expression": macros.hpp

```
/*! \file macros.hpp
    \brief Preprocessor macros that can customise the cereal library

    By default, cereal looks for serialization functions with very
    specific names, that is: serialize, load, save, load_minimal,
    or save_minimal.

    This file allows an advanced user to change these names to conform
    to some other style or preference.  This is implemented using
    preprocessor macros.

    As a result of this, in internal cereal code you will see macros
    used for these function names.  In user code, you should name
    the functions like you normally would and not use the macros
    to improve readability.
    \ingroup utility */
/*
  Copyright (c) 2014, Randolph Voorhies, Shane Grant
  All rights reserved.

  Redistribution and use in source and binary forms, with or without
  modification, are permitted provided that the following conditions are met:
      * Redistributions of source code must retain the above copyright
        notice, this list of conditions and the following disclaimer.
      * Redistributions in binary form must reproduce the above copyright
        notice, this list of conditions and the following disclaimer in the
        documentation and/or other materials provided with the distribution.
      * Neither the name of cereal nor the
        names of its contributors may be used to endorse or promote products
        derived from this software without specific prior written permission.

  THIS SOFTWARE IS PROVIDED BY THE COPYRIGHT HOLDERS AND CONTRIBUTORS "AS IS" AND
  ANY EXPRESS OR IMPLIED WARRANTIES, INCLUDING, BUT NOT LIMITED TO, THE IMPLIED
  WARRANTIES OF MERCHANTABILITY AND FITNESS FOR A PARTICULAR PURPOSE ARE
  DISCLAIMED. IN NO EVENT SHALL RANDOLPH VOORHIES OR SHANE GRANT BE LIABLE FOR ANY
  DIRECT, INDIRECT, INCIDENTAL, SPECIAL, EXEMPLARY, OR CONSEQUENTIAL DAMAGES
  (INCLUDING, BUT NOT LIMITED TO, PROCUREMENT OF SUBSTITUTE GOODS OR SERVICES;
  LOSS OF USE, DATA, OR PROFITS; OR BUSINESS INTERRUPTION) HOWEVER CAUSED AND
  ON ANY THEORY OF LIABILITY, WHETHER IN CONTRACT, STRICT LIABILITY, OR TORT
  (INCLUDING NEGLIGENCE OR OTHERWISE) ARISING IN ANY WAY OUT OF THE USE OF THIS
  SOFTWARE, EVEN IF ADVISED OF THE POSSIBILITY OF SUCH DAMAGE.
*/

#ifndef CEREAL_MACROS_HPP_
#define CEREAL_MACROS_HPP_

#ifndef CEREAL_THREAD_SAFE
//! Whether cereal should be compiled for a threaded environment
/*! This macro causes cereal to use mutexes to control access to
    global internal state in a thread safe manner.

    Note that even with this enabled you must still ensure that
    archives are accessed by only one thread at a time; it is safe
    to use multiple archives in parallel, but not to access one archive
    from many places simultaneously. */
#define CEREAL_THREAD_SAFE 0
#endif // CEREAL_THREAD_SAFE

#ifndef CEREAL_SIZE_TYPE
//! Determines the data type used for size_type
/*! cereal uses size_type to ensure that the serialized size of
    dynamic containers is compatible across different architectures
    (e.g. 32 vs 64 bit), which may use different underlying types for
    std::size_t.

    More information can be found in cereal/details/helpers.hpp.

    If you choose to modify this type, ensure that you use a fixed
    size type (e.g. uint32_t). */
#define CEREAL_SIZE_TYPE uint64_t
#endif // CEREAL_SIZE_TYPE

// ######################################################################
#ifndef CEREAL_SERIALIZE_FUNCTION_NAME
//! The serialization/deserialization function name to search for.
/*! You can define @c CEREAL_SERIALIZE_FUNCTION_NAME to be different assuming
    you do so before this file is included. */
#define CEREAL_SERIALIZE_FUNCTION_NAME serialize
#endif // CEREAL_SERIALIZE_FUNCTION_NAME

#ifndef CEREAL_LOAD_FUNCTION_NAME
//! The deserialization (load) function name to search for.
/*! You can define @c CEREAL_LOAD_FUNCTION_NAME to be different assuming you do so
    before this file is included. */
#define CEREAL_LOAD_FUNCTION_NAME load
#endif // CEREAL_LOAD_FUNCTION_NAME

#ifndef CEREAL_SAVE_FUNCTION_NAME
//! The serialization (save) function name to search for.
/*! You can define @c CEREAL_SAVE_FUNCTION_NAME to be different assuming you do so
    before this file is included. */
#define CEREAL_SAVE_FUNCTION_NAME save
#endif // CEREAL_SAVE_FUNCTION_NAME

#ifndef CEREAL_LOAD_MINIMAL_FUNCTION_NAME
//! The deserialization (load_minimal) function name to search for.
/*! You can define @c CEREAL_LOAD_MINIMAL_FUNCTION_NAME to be different assuming you do so
    before this file is included. */
#define CEREAL_LOAD_MINIMAL_FUNCTION_NAME load_minimal
#endif // CEREAL_LOAD_MINIMAL_FUNCTION_NAME

#ifndef CEREAL_SAVE_MINIMAL_FUNCTION_NAME
//! The serialization (save_minimal) function name to search for.
/*! You can define @c CEREAL_SAVE_MINIMAL_FUNCTION_NAME to be different assuming you do so
    before this file is included. */
#define CEREAL_SAVE_MINIMAL_FUNCTION_NAME save_minimal
#endif // CEREAL_SAVE_MINIMAL_FUNCTION_NAME

// ######################################################################
//! Defines the CEREAL_NOEXCEPT macro to use instead of noexcept
/*! If a compiler we support does not support noexcept, this macro
    will detect this and define CEREAL_NOEXCEPT as a no-op
    @internal */
#if !defined(CEREAL_HAS_NOEXCEPT)
  #if defined(__clang__)
    #if __has_feature(cxx_noexcept)
      #define CEREAL_HAS_NOEXCEPT
    #endif
  #else // NOT clang
    #if defined(__GXX_EXPERIMENTAL_CXX0X__) && __GNUC__ * 10 + __GNUC_MINOR__ >= 46 || \
        defined(_MSC_FULL_VER) && _MSC_FULL_VER >= 190023026
      #define CEREAL_HAS_NOEXCEPT
    #endif // end GCC/MSVC check
  #endif // end NOT clang block

  #ifndef CEREAL_NOEXCEPT
    #ifdef CEREAL_HAS_NOEXCEPT
      #define CEREAL_NOEXCEPT noexcept
    #else
      #define CEREAL_NOEXCEPT
    #endif // end CEREAL_HAS_NOEXCEPT
  #endif // end !defined(CEREAL_HAS_NOEXCEPT)
#endif // ifndef CEREAL_NOEXCEPT

// ######################################################################
//! Checks if C++17 is available
//! NOTE: clang v5 has a bug with inline variables, so disable C++17 on that compiler
#if (__cplusplus >= 201703L || (defined(_MSVC_LANG) && _MSVC_LANG >= 201703L)) \
    && (!defined(__clang__) || __clang_major__ > 5)
#define CEREAL_HAS_CPP17
#endif

//! Checks if C++14 is available
#if __cplusplus >= 201402L
#define CEREAL_HAS_CPP14
#endif

// ######################################################################
//! Defines the CEREAL_ALIGNOF macro to use instead of alignof
#if defined(_MSC_VER) && _MSC_VER < 1900
#define CEREAL_ALIGNOF __alignof
#else // not MSVC 2013 or older
#define CEREAL_ALIGNOF alignof
#endif // end MSVC check

#endif // CEREAL_MACROS_HPP_
```
