## Supplemental code for "The motif composition of variable-number tandem repeats impacts gene expression": specialize.hpp

```
/*! \file specialize.hpp
    \brief Serialization disambiguation */
/*
  Copyright (c) 2014, Randolph Voorhies, Shane Grant
  All rights reserved.

  Redistribution and use in source and binary forms, with or without
  modification, are permitted provided that the following conditions are met:
      * Redistributions of source code must retain the above copyright
        notice, this list of conditions and the following disclaimer.
      * Redistributions in binary form must reproduce the above copyright
        notice, this list of conditions and the following disclaimer in the
        documentation and/or other materials provided with the distribution.
      * Neither the name of cereal nor the
        names of its contributors may be used to endorse or promote products
        derived from this software without specific prior written permission.

  THIS SOFTWARE IS PROVIDED BY THE COPYRIGHT HOLDERS AND CONTRIBUTORS "AS IS" AND
  ANY EXPRESS OR IMPLIED WARRANTIES, INCLUDING, BUT NOT LIMITED TO, THE IMPLIED
  WARRANTIES OF MERCHANTABILITY AND FITNESS FOR A PARTICULAR PURPOSE ARE
  DISCLAIMED. IN NO EVENT SHALL RANDOLPH VOORHIES OR SHANE GRANT BE LIABLE FOR ANY
  DIRECT, INDIRECT, INCIDENTAL, SPECIAL, EXEMPLARY, OR CONSEQUENTIAL DAMAGES
  (INCLUDING, BUT NOT LIMITED TO, PROCUREMENT OF SUBSTITUTE GOODS OR SERVICES;
  LOSS OF USE, DATA, OR PROFITS; OR BUSINESS INTERRUPTION) HOWEVER CAUSED AND
  ON ANY THEORY OF LIABILITY, WHETHER IN CONTRACT, STRICT LIABILITY, OR TORT
  (INCLUDING NEGLIGENCE OR OTHERWISE) ARISING IN ANY WAY OUT OF THE USE OF THIS
  SOFTWARE, EVEN IF ADVISED OF THE POSSIBILITY OF SUCH DAMAGE.
*/

#ifndef CEREAL_SPECIALIZE_HPP_
#define CEREAL_SPECIALIZE_HPP_

namespace cereal
{
  // Forward declaration of access class that users can become friends with
  class access;

  // ######################################################################
  //! A specifier used in conjunction with cereal::specialize to disambiguate
  //! serialization in special cases
  /*! @relates specialize
      @ingroup Access */
  enum class specialization
  {
    member_serialize,            //!< Force the use of a member serialize function
    member_load_save,            //!< Force the use of a member load/save pair
    member_load_save_minimal,    //!< Force the use of a member minimal load/save pair
    non_member_serialize,        //!< Force the use of a non-member serialize function
    non_member_load_save,        //!< Force the use of a non-member load/save pair
    non_member_load_save_minimal //!< Force the use of a non-member minimal load/save pair
  };

  //! A class used to disambiguate cases where cereal cannot detect a unique way of serializing a class
  /*! cereal attempts to figure out which method of serialization (member vs. non-member serialize
      or load/save pair) at compile time.  If for some reason cereal cannot find a non-ambiguous way
      of serializing a type, it will produce a static assertion complaining about this.

      This can happen because you have both a serialize and load/save pair, or even because a base
      class has a serialize (public or private with friend access) and a derived class does not
      overwrite this due to choosing some other serialization type.

      Specializing this class will tell cereal to explicitly use the serialization type you specify
      and it will not complain about ambiguity in its compile time selection.  However, if cereal detects
      an ambiguity in specializations, it will continue to issue a static assertion.

      @code{.cpp}
      class MyParent
      {
        friend class cereal::access;
        template <class Archive>
        void serialize( Archive & ar ) {}
      };

      // Although serialize is private in MyParent, to cereal::access it will look public,
      // even through MyDerived
      class MyDerived : public MyParent
      {
        public:
          template <class Archive>
          void load( Archive & ar ) {}

          template <class Archive>
          void save( Archive & ar ) {}
      };

      // The load/save pair in MyDerived is ambiguous because serialize in MyParent can
      // be accessed from cereal::access.  This looks the same as making serialize public
      // in MyParent, making it seem as though MyDerived has both a serialize and a load/save pair.
      // cereal will complain about this at compile time unless we disambiguate:

      namespace cereal
      {
        // This struct specialization will tell cereal which is the right way to serialize the ambiguity
        template <class Archive> struct specialize<Archive, MyDerived, cereal::specialization::member_load_save> {};

        // If we only had a disambiguation for a specific archive type, it would look something like this
        template <> struct specialize<cereal::BinaryOutputArchive, MyDerived, cereal::specialization::member_load_save> {};
      }
      @endcode

      You can also choose to use the macros CEREAL_SPECIALIZE_FOR_ALL_ARCHIVES or
      CEREAL_SPECIALIZE_FOR_ARCHIVE if you want to type a little bit less.

      @tparam T The type to specialize the serialization for
      @tparam S The specialization type to use for T
      @ingroup Access */
  template <class Archive, class T, specialization S>
  struct specialize : public std::false_type {};

  //! Convenient macro for performing specialization for all archive types
  /*! This performs specialization for the specific type for all types of archives.
      This macro should be placed at the global namespace.

      @code{cpp}
      struct MyType {};
      CEREAL_SPECIALIZE_FOR_ALL_ARCHIVES( MyType, cereal::specialization::member_load_save );
      @endcode

      @relates specialize
      @ingroup Access */
  #define CEREAL_SPECIALIZE_FOR_ALL_ARCHIVES( Type, Specialization )                                \
  namespace cereal { template <class Archive> struct specialize<Archive, Type, Specialization> {}; }

  //! Convenient macro for performing specialization for a single archive type
  /*! This performs specialization for the specific type for a single type of archive.
      This macro should be placed at the global namespace.

      @code{cpp}
      struct MyType {};
      CEREAL_SPECIALIZE_FOR_ARCHIVE( cereal::XMLInputArchive, MyType, cereal::specialization::member_load_save );
      @endcode

      @relates specialize
      @ingroup Access */
  #define CEREAL_SPECIALIZE_FOR_ARCHIVE( Archive, Type, Specialization )               \
  namespace cereal { template <> struct specialize<Archive, Type, Specialization> {}; }
}

#endif
```
