## Supplemental code for "The motif composition of variable-number tandem repeats impacts gene expression": base_class.hpp

```
/*! \file base_class.hpp
    \brief Support for base classes (virtual and non-virtual)
    \ingroup OtherTypes */
/*
  Copyright (c) 2014, Randolph Voorhies, Shane Grant
  All rights reserved.

  Redistribution and use in source and binary forms, with or without
  modification, are permitted provided that the following conditions are met:
      * Redistributions of source code must retain the above copyright
        notice, this list of conditions and the following disclaimer.
      * Redistributions in binary form must reproduce the above copyright
        notice, this list of conditions and the following disclaimer in the
        documentation and/or other materials provided with the distribution.
      * Neither the name of cereal nor the
        names of its contributors may be used to endorse or promote products
        derived from this software without specific prior written permission.

  THIS SOFTWARE IS PROVIDED BY THE COPYRIGHT HOLDERS AND CONTRIBUTORS "AS IS" AND
  ANY EXPRESS OR IMPLIED WARRANTIES, INCLUDING, BUT NOT LIMITED TO, THE IMPLIED
  WARRANTIES OF MERCHANTABILITY AND FITNESS FOR A PARTICULAR PURPOSE ARE
  DISCLAIMED. IN NO EVENT SHALL RANDOLPH VOORHIES OR SHANE GRANT BE LIABLE FOR ANY
  DIRECT, INDIRECT, INCIDENTAL, SPECIAL, EXEMPLARY, OR CONSEQUENTIAL DAMAGES
  (INCLUDING, BUT NOT LIMITED TO, PROCUREMENT OF SUBSTITUTE GOODS OR SERVICES;
  LOSS OF USE, DATA, OR PROFITS; OR BUSINESS INTERRUPTION) HOWEVER CAUSED AND
  ON ANY THEORY OF LIABILITY, WHETHER IN CONTRACT, STRICT LIABILITY, OR TORT
  (INCLUDING NEGLIGENCE OR OTHERWISE) ARISING IN ANY WAY OUT OF THE USE OF THIS
  SOFTWARE, EVEN IF ADVISED OF THE POSSIBILITY OF SUCH DAMAGE.
*/
#ifndef CEREAL_TYPES_BASE_CLASS_HPP_
#define CEREAL_TYPES_BASE_CLASS_HPP_

#include "cereal/details/traits.hpp"
#include "cereal/details/polymorphic_impl_fwd.hpp"

namespace cereal
{
  namespace base_class_detail
  {
    //! Used to register polymorphic relations and avoid the need to include
    //! polymorphic.hpp when no polymorphism is used
    /*! @internal */
    template <class Base, class Derived, bool IsPolymorphic = std::is_polymorphic<Base>::value>
    struct RegisterPolymorphicBaseClass
    {
      static void bind()
      { }
    };

    //! Polymorphic version
    /*! @internal */
    template <class Base, class Derived>
    struct RegisterPolymorphicBaseClass<Base, Derived, true>
    {
      static void bind()
      { detail::RegisterPolymorphicCaster<Base, Derived>::bind(); }
    };
  }

  //! Casts a derived class to its non-virtual base class in a way that safely supports abstract classes
  /*! This should be used in cases when a derived type needs to serialize its base type. This is better than directly
      using static_cast, as it allows for serialization of pure virtual (abstract) base classes.

      This also automatically registers polymorphic relation between the base and derived class, assuming they
      are indeed polymorphic. Note this is not the same as polymorphic type registration. For more information
      see the documentation on polymorphism. If using a polymorphic class, be sure to include support for
      polymorphism (cereal/types/polymorphic.hpp).

      \sa virtual_base_class

      @code{.cpp}
      struct MyBase
      {
        int x;

        virtual void foo() = 0;

        template <class Archive>
        void serialize( Archive & ar )
        {
          ar( x );
        }
      };

      struct MyDerived : public MyBase //<-- Note non-virtual inheritance
      {
        int y;

        virtual void foo() {};

        template <class Archive>
        void serialize( Archive & ar )
        {
          ar( cereal::base_class<MyBase>(this) );
          ar( y );
        }
      };
      @endcode */
  template<class Base>
    struct base_class : private traits::detail::BaseCastBase
    {
      template<class Derived>
        base_class(Derived const * derived) :
          base_ptr(const_cast<Base*>(static_cast<Base const *>(derived)))
      {
        static_assert( std::is_base_of<Base, Derived>::value, "Can only use base_class on a valid base class" );
        base_class_detail::RegisterPolymorphicBaseClass<Base, Derived>::bind();
      }

        Base * base_ptr;
    };

  //! Casts a derived class to its virtual base class in a way that allows cereal to track inheritance
  /*! This should be used in cases when a derived type features virtual inheritance from some
      base type.  This allows cereal to track the inheritance and to avoid making duplicate copies
      during serialization.

      It is safe to use virtual_base_class in all circumstances for serializing base classes, even in cases
      where virtual inheritance does not take place, though it may be slightly faster to utilize
      cereal::base_class<> if you do not need to worry about virtual inheritance.

      This also automatically registers polymorphic relation between the base and derived class, assuming they
      are indeed polymorphic. Note this is not the same as polymorphic type registration. For more information
      see the documentation on polymorphism. If using a polymorphic class, be sure to include support for
      polymorphism (cereal/types/polymorphic.hpp).

      \sa base_class

      @code{.cpp}
      struct MyBase
      {
        int x;

        template <class Archive>
        void serialize( Archive & ar )
        {
          ar( x );
        }
      };

      struct MyLeft : virtual MyBase //<-- Note the virtual inheritance
      {
        int y;

        template <class Archive>
        void serialize( Archive & ar )
        {
          ar( cereal::virtual_base_class<MyBase>( this ) );
          ar( y );
        }
      };

      struct MyRight : virtual MyBase
      {
        int z;

        template <class Archive>
        void serialize( Archive & ar )
        {
          ar( cereal::virtual_base_clas<MyBase>( this ) );
          ar( z );
        }
      };

      // diamond virtual inheritance; contains one copy of each base class
      struct MyDerived : virtual MyLeft, virtual MyRight
      {
        int a;

        template <class Archive>
        void serialize( Archive & ar )
        {
          ar( cereal::virtual_base_class<MyLeft>( this ) );  // safely serialize data members in MyLeft
          ar( cereal::virtual_base_class<MyRight>( this ) ); // safely serialize data members in MyRight
          ar( a );

          // Because we used virtual_base_class, cereal will ensure that only one instance of MyBase is
          // serialized as we traverse the inheritance heirarchy. This means that there will be one copy
          // each of the variables x, y, z, and a

          // If we had chosen to use static_cast<> instead, cereal would perform no tracking and
          // assume that every base class should be serialized (in this case leading to a duplicate
          // serialization of MyBase due to diamond inheritance
      };
     }
     @endcode */
  template<class Base>
    struct virtual_base_class : private traits::detail::BaseCastBase
    {
      template<class Derived>
        virtual_base_class(Derived const * derived) :
          base_ptr(const_cast<Base*>(static_cast<Base const *>(derived)))
      {
        static_assert( std::is_base_of<Base, Derived>::value, "Can only use virtual_base_class on a valid base class" );
        base_class_detail::RegisterPolymorphicBaseClass<Base, Derived>::bind();
      }

        Base * base_ptr;
    };

} // namespace cereal

#endif // CEREAL_TYPES_BASE_CLASS_HPP_
```
