## Supplemental code for "The motif composition of variable-number tandem repeats impacts gene expression": boost_variant.hpp

```
/*! \file boost_variant.hpp
    \brief Support for boost::variant
    \ingroup OtherTypes */
/*
  Copyright (c) 2014, Randolph Voorhies, Shane Grant
  All rights reserved.

  Redistribution and use in source and binary forms, with or without
  modification, are permitted provided that the following conditions are met:
      * Redistributions of source code must retain the above copyright
        notice, this list of conditions and the following disclaimer.
      * Redistributions in binary form must reproduce the above copyright
        notice, this list of conditions and the following disclaimer in the
        documentation and/or other materials provided with the distribution.
      * Neither the name of cereal nor the
        names of its contributors may be used to endorse or promote products
        derived from this software without specific prior written permission.

  THIS SOFTWARE IS PROVIDED BY THE COPYRIGHT HOLDERS AND CONTRIBUTORS "AS IS" AND
  ANY EXPRESS OR IMPLIED WARRANTIES, INCLUDING, BUT NOT LIMITED TO, THE IMPLIED
  WARRANTIES OF MERCHANTABILITY AND FITNESS FOR A PARTICULAR PURPOSE ARE
  DISCLAIMED. IN NO EVENT SHALL RANDOLPH VOORHIES OR SHANE GRANT BE LIABLE FOR ANY
  DIRECT, INDIRECT, INCIDENTAL, SPECIAL, EXEMPLARY, OR CONSEQUENTIAL DAMAGES
  (INCLUDING, BUT NOT LIMITED TO, PROCUREMENT OF SUBSTITUTE GOODS OR SERVICES;
  LOSS OF USE, DATA, OR PROFITS; OR BUSINESS INTERRUPTION) HOWEVER CAUSED AND
  ON ANY THEORY OF LIABILITY, WHETHER IN CONTRACT, STRICT LIABILITY, OR TORT
  (INCLUDING NEGLIGENCE OR OTHERWISE) ARISING IN ANY WAY OUT OF THE USE OF THIS
  SOFTWARE, EVEN IF ADVISED OF THE POSSIBILITY OF SUCH DAMAGE.
*/
#ifndef CEREAL_TYPES_BOOST_VARIANT_HPP_
#define CEREAL_TYPES_BOOST_VARIANT_HPP_

//! @internal
#if defined(_MSC_VER) && _MSC_VER < 1911
#define CEREAL_CONSTEXPR_LAMBDA
#else // MSVC 2017 or newer, all other compilers
#define CEREAL_CONSTEXPR_LAMBDA constexpr
#endif

#include "cereal/cereal.hpp"
#include <boost/variant/variant_fwd.hpp>
#include <boost/variant/static_visitor.hpp>

namespace cereal
{
  namespace boost_variant_detail
  {
    //! @internal
    template <class Archive>
    struct variant_save_visitor : boost::static_visitor<>
    {
      variant_save_visitor(Archive & ar_) : ar(ar_) {}

      template<class T>
      void operator()(T const & value) const
      {
        ar( CEREAL_NVP_("data", value) );
      }

      Archive & ar;
    };

    //! @internal
    template <class Archive, class T>
    struct LoadAndConstructLoadWrapper
    {
      using ST = typename std::aligned_storage<sizeof(T), CEREAL_ALIGNOF(T)>::type;

      LoadAndConstructLoadWrapper() :
        construct( reinterpret_cast<T *>( &st ) )
      { }

      ~LoadAndConstructLoadWrapper()
      {
        if (construct.itsValid)
        {
          construct->~T();
        }
      }

      void CEREAL_SERIALIZE_FUNCTION_NAME( Archive & ar )
      {
        ::cereal::detail::Construct<T, Archive>::load_andor_construct( ar, construct );
      }

      ST st;
      ::cereal::construct<T> construct;
    };

    //! @internal
    template <class T> struct load_variant_wrapper;

    //! Avoid serializing variant void_ type
    /*! @internal */
    template <>
    struct load_variant_wrapper<boost::detail::variant::void_>
    {
      template <class Variant, class Archive>
      static void load_variant( Archive &, Variant & )
      { }
    };

    //! @internal
    template <class T>
    struct load_variant_wrapper
    {
      // default constructible
      template <class Archive, class Variant>
      static void load_variant_impl( Archive & ar, Variant & variant, std::true_type )
      {
        T value;
        ar( CEREAL_NVP_("data", value) );
        variant = std::move(value);
      }

      // not default constructible
      template<class Variant, class Archive>
      static void load_variant_impl(Archive & ar, Variant & variant, std::false_type )
      {
        LoadAndConstructLoadWrapper<Archive, T> loadWrapper;

        ar( CEREAL_NVP_("data", loadWrapper) );
        variant = std::move(*loadWrapper.construct.ptr());
      }

      //! @internal
      template<class Variant, class Archive>
      static void load_variant(Archive & ar, Variant & variant)
      {
        load_variant_impl( ar, variant, typename std::is_default_constructible<T>::type() );
      }
    };
  } // namespace boost_variant_detail

  //! Saving for boost::variant
  template <class Archive, typename ... VariantTypes> inline
  void CEREAL_SAVE_FUNCTION_NAME( Archive & ar, boost::variant<VariantTypes...> const & variant )
  {
    int32_t which = variant.which();
    ar( CEREAL_NVP_("which", which) );
    boost_variant_detail::variant_save_visitor<Archive> visitor(ar);
    variant.apply_visitor(visitor);
  }

  //! Loading for boost::variant
  template <class Archive, typename ... VariantTypes> inline
  void CEREAL_LOAD_FUNCTION_NAME( Archive & ar, boost::variant<VariantTypes...> & variant )
  {
    int32_t which;
    ar( CEREAL_NVP_("which", which) );

    using LoadFuncType = void(*)(Archive &, boost::variant<VariantTypes...> &);
    CEREAL_CONSTEXPR_LAMBDA LoadFuncType loadFuncArray[] = {&boost_variant_detail::load_variant_wrapper<VariantTypes>::load_variant...};

    if(which >= int32_t(sizeof(loadFuncArray)/sizeof(loadFuncArray[0])))
      throw Exception("Invalid 'which' selector when deserializing boost::variant");

    loadFuncArray[which](ar, variant);
  }
} // namespace cereal

#undef CEREAL_CONSTEXPR_LAMBDA

#endif // CEREAL_TYPES_BOOST_VARIANT_HPP_
```
