## Supplemental code for "The motif composition of variable-number tandem repeats impacts gene expression": common.hpp

```
/*! \file common.hpp
    \brief Support common types - always included automatically
    \ingroup OtherTypes */
/*
  Copyright (c) 2014, Randolph Voorhies, Shane Grant
  All rights reserved.

  Redistribution and use in source and binary forms, with or without
  modification, are permitted provided that the following conditions are met:
      * Redistributions of source code must retain the above copyright
        notice, this list of conditions and the following disclaimer.
      * Redistributions in binary form must reproduce the above copyright
        notice, this list of conditions and the following disclaimer in the
        documentation and/or other materials provided with the distribution.
      * Neither the name of cereal nor the
        names of its contributors may be used to endorse or promote products
        derived from this software without specific prior written permission.

  THIS SOFTWARE IS PROVIDED BY THE COPYRIGHT HOLDERS AND CONTRIBUTORS "AS IS" AND
  ANY EXPRESS OR IMPLIED WARRANTIES, INCLUDING, BUT NOT LIMITED TO, THE IMPLIED
  WARRANTIES OF MERCHANTABILITY AND FITNESS FOR A PARTICULAR PURPOSE ARE
  DISCLAIMED. IN NO EVENT SHALL RANDOLPH VOORHIES OR SHANE GRANT BE LIABLE FOR ANY
  DIRECT, INDIRECT, INCIDENTAL, SPECIAL, EXEMPLARY, OR CONSEQUENTIAL DAMAGES
  (INCLUDING, BUT NOT LIMITED TO, PROCUREMENT OF SUBSTITUTE GOODS OR SERVICES;
  LOSS OF USE, DATA, OR PROFITS; OR BUSINESS INTERRUPTION) HOWEVER CAUSED AND
  ON ANY THEORY OF LIABILITY, WHETHER IN CONTRACT, STRICT LIABILITY, OR TORT
  (INCLUDING NEGLIGENCE OR OTHERWISE) ARISING IN ANY WAY OUT OF THE USE OF THIS
  SOFTWARE, EVEN IF ADVISED OF THE POSSIBILITY OF SUCH DAMAGE.
*/
#ifndef CEREAL_TYPES_COMMON_HPP_
#define CEREAL_TYPES_COMMON_HPP_

#include "cereal/cereal.hpp"

namespace cereal
{
  namespace common_detail
  {
    //! Serialization for arrays if BinaryData is supported and we are arithmetic
    /*! @internal */
    template <class Archive, class T> inline
    void serializeArray( Archive & ar, T & array, std::true_type /* binary_supported */ )
    {
      ar( binary_data( array, sizeof(array) ) );
    }

    //! Serialization for arrays if BinaryData is not supported or we are not arithmetic
    /*! @internal */
    template <class Archive, class T> inline
    void serializeArray( Archive & ar, T & array, std::false_type /* binary_supported */ )
    {
      for( auto & i : array )
        ar( i );
    }

    namespace
    {
      //! Gets the underlying type of an enum
      /*! @internal */
      template <class T, bool IsEnum>
      struct enum_underlying_type : std::false_type {};

      //! Gets the underlying type of an enum
      /*! Specialization for when we actually have an enum
          @internal */
      template <class T>
      struct enum_underlying_type<T, true> { using type = typename std::underlying_type<T>::type; };
    } // anon namespace

    //! Checks if a type is an enum
    /*! This is needed over simply calling std::is_enum because the type
        traits checking at compile time will attempt to call something like
        load_minimal with a special NoConvertRef struct that wraps up the true type.

        This will strip away any of that and also expose the true underlying type.
        @internal */
    template <class T>
    class is_enum
    {
      private:
        using DecayedT  = typename std::decay<T>::type;
        using StrippedT = typename ::cereal::traits::strip_minimal<DecayedT>::type;

      public:
        static const bool value = std::is_enum<StrippedT>::value;
        using type = StrippedT;
        using base_type = typename enum_underlying_type<StrippedT, value>::type;
    };
  }

  //! Saving for enum types
  template <class Archive, class T> inline
  typename std::enable_if<common_detail::is_enum<T>::value,
                          typename common_detail::is_enum<T>::base_type>::type
  CEREAL_SAVE_MINIMAL_FUNCTION_NAME( Archive const &, T const & t )
  {
    return static_cast<typename common_detail::is_enum<T>::base_type>(t);
  }

  //! Loading for enum types
  template <class Archive, class T> inline
  typename std::enable_if<common_detail::is_enum<T>::value, void>::type
  CEREAL_LOAD_MINIMAL_FUNCTION_NAME( Archive const &, T && t,
                                     typename common_detail::is_enum<T>::base_type const & value )
  {
    t = reinterpret_cast<typename common_detail::is_enum<T>::type const &>( value );
  }

  //! Serialization for raw pointers
  /*! This exists only to throw a static_assert to let users know we don't support raw pointers. */
  template <class Archive, class T> inline
  void CEREAL_SERIALIZE_FUNCTION_NAME( Archive &, T * & )
  {
    static_assert(cereal::traits::detail::delay_static_assert<T>::value,
      "Cereal does not support serializing raw pointers - please use a smart pointer");
  }

  //! Serialization for C style arrays
  template <class Archive, class T> inline
  typename std::enable_if<std::is_array<T>::value, void>::type
  CEREAL_SERIALIZE_FUNCTION_NAME(Archive & ar, T & array)
  {
    common_detail::serializeArray( ar, array,
        std::integral_constant<bool, traits::is_output_serializable<BinaryData<T>, Archive>::value &&
                                     std::is_arithmetic<typename std::remove_all_extents<T>::type>::value>() );
  }
} // namespace cereal

#endif // CEREAL_TYPES_COMMON_HPP_
```
