## Supplemental code for "The motif composition of variable-number tandem repeats impacts gene expression": memory.hpp

```
/*! \file memory.hpp
    \brief Support for types found in \<memory\>
    \ingroup STLSupport */
/*
  Copyright (c) 2014, Randolph Voorhies, Shane Grant
  All rights reserved.

  Redistribution and use in source and binary forms, with or without
  modification, are permitted provided that the following conditions are met:
      * Redistributions of source code must retain the above copyright
        notice, this list of conditions and the following disclaimer.
      * Redistributions in binary form must reproduce the above copyright
        notice, this list of conditions and the following disclaimer in the
        documentation and/or other materials provided with the distribution.
      * Neither the name of cereal nor the
        names of its contributors may be used to endorse or promote products
        derived from this software without specific prior written permission.

  THIS SOFTWARE IS PROVIDED BY THE COPYRIGHT HOLDERS AND CONTRIBUTORS "AS IS" AND
  ANY EXPRESS OR IMPLIED WARRANTIES, INCLUDING, BUT NOT LIMITED TO, THE IMPLIED
  WARRANTIES OF MERCHANTABILITY AND FITNESS FOR A PARTICULAR PURPOSE ARE
  DISCLAIMED. IN NO EVENT SHALL RANDOLPH VOORHIES OR SHANE GRANT BE LIABLE FOR ANY
  DIRECT, INDIRECT, INCIDENTAL, SPECIAL, EXEMPLARY, OR CONSEQUENTIAL DAMAGES
  (INCLUDING, BUT NOT LIMITED TO, PROCUREMENT OF SUBSTITUTE GOODS OR SERVICES;
  LOSS OF USE, DATA, OR PROFITS; OR BUSINESS INTERRUPTION) HOWEVER CAUSED AND
  ON ANY THEORY OF LIABILITY, WHETHER IN CONTRACT, STRICT LIABILITY, OR TORT
  (INCLUDING NEGLIGENCE OR OTHERWISE) ARISING IN ANY WAY OUT OF THE USE OF THIS
  SOFTWARE, EVEN IF ADVISED OF THE POSSIBILITY OF SUCH DAMAGE.
*/
#ifndef CEREAL_TYPES_SHARED_PTR_HPP_
#define CEREAL_TYPES_SHARED_PTR_HPP_

#include "cereal/cereal.hpp"
#include <memory>
#include <cstring>

namespace cereal
{
  namespace memory_detail
  {
    //! A wrapper class to notify cereal that it is ok to serialize the contained pointer
    /*! This mechanism allows us to intercept and properly handle polymorphic pointers
        @internal */
    template<class T>
    struct PtrWrapper
    {
      PtrWrapper(T && p) : ptr(std::forward<T>(p)) {}
      T & ptr;

      PtrWrapper( PtrWrapper const & ) = default;
      PtrWrapper & operator=( PtrWrapper const & ) = delete;
    };

    //! Make a PtrWrapper
    /*! @internal */
    template<class T> inline
    PtrWrapper<T> make_ptr_wrapper(T && t)
    {
      return {std::forward<T>(t)};
    }

    //! A struct that acts as a wrapper around calling load_andor_construct
    /*! The purpose of this is to allow a load_and_construct call to properly enter into the
        'data' NVP of the ptr_wrapper
        @internal */
    template <class Archive, class T>
    struct LoadAndConstructLoadWrapper
    {
      LoadAndConstructLoadWrapper( T * ptr ) :
        construct( ptr )
      { }

      //! Constructor for embedding an early call for restoring shared_from_this
      template <class F>
      LoadAndConstructLoadWrapper( T * ptr, F && sharedFromThisFunc ) :
        construct( ptr, sharedFromThisFunc )
      { }

      inline void CEREAL_SERIALIZE_FUNCTION_NAME( Archive & ar )
      {
        ::cereal::detail::Construct<T, Archive>::load_andor_construct( ar, construct );
      }

      ::cereal::construct<T> construct;
    };

    //! A helper struct for saving and restoring the state of types that derive from
    //! std::enable_shared_from_this
    /*! This special struct is necessary because when a user uses load_and_construct,
        the weak_ptr (or whatever implementation defined variant) that allows
        enable_shared_from_this to function correctly will not be initialized properly.

        This internal weak_ptr can also be modified by the shared_ptr that is created
        during the serialization of a polymorphic pointer, where cereal creates a
        wrapper shared_ptr out of a void pointer to the real data.

        In the case of load_and_construct, this happens because it is the allocation
        of shared_ptr that perform this initialization, which we let happen on a buffer
        of memory (aligned_storage).  This buffer is then used for placement new
        later on, effectively overwriting any initialized weak_ptr with a default
        initialized one, eventually leading to issues when the user calls shared_from_this.

        To get around these issues, we will store the memory for the enable_shared_from_this
        portion of the class and replace it after whatever happens to modify it (e.g. the
        user performing construction or the wrapper shared_ptr in saving).

        Note that this goes into undefined behavior territory, but as of the initial writing
        of this, all standard library implementations of std::enable_shared_from_this are
        compatible with this memory manipulation. It is entirely possible that this may someday
        break or may not work with convoluted use cases.

        Example usage:

        @code{.cpp}
        T * myActualPointer;
        {
          EnableSharedStateHelper<T> helper( myActualPointer ); // save the state
          std::shared_ptr<T> myPtr( myActualPointer ); // modifies the internal weak_ptr
          // helper restores state when it goes out of scope
        }
        @endcode

        When possible, this is designed to be used in an RAII fashion - it will save state on
        construction and restore it on destruction. The restore can be done at an earlier time
        (e.g. after construct() is called in load_and_construct) in which case the destructor will
        do nothing. Performing the restore immediately following construct() allows a user to call
        shared_from_this within their load_and_construct function.

        @tparam T Type pointed to by shared_ptr
        @internal */
    template <class T>
    class EnableSharedStateHelper
    {
      // typedefs for parent type and storage type
      using BaseType = typename ::cereal::traits::get_shared_from_this_base<T>::type;
      using ParentType = std::enable_shared_from_this<BaseType>;
      using StorageType = typename std::aligned_storage<sizeof(ParentType), CEREAL_ALIGNOF(ParentType)>::type;

      public:
        //! Saves the state of some type inheriting from enable_shared_from_this
        /*! @param ptr The raw pointer held by the shared_ptr */
        inline EnableSharedStateHelper( T * ptr ) :
          itsPtr( static_cast<ParentType *>( ptr ) ),
          itsState(),
          itsRestored( false )
        {
          std::memcpy( &itsState, itsPtr, sizeof(ParentType) );
        }

        //! Restores the state of the held pointer (can only be done once)
        inline void restore()
        {
          if( !itsRestored )
          {
            // void * cast needed when type has no trivial copy-assignment
            std::memcpy( static_cast<void *>(itsPtr), &itsState, sizeof(ParentType) );
            itsRestored = true;
          }
        }

        //! Restores the state of the held pointer if not done previously
        inline ~EnableSharedStateHelper()
        {
          restore();
        }

      private:
        ParentType * itsPtr;
        StorageType itsState;
        bool itsRestored;
    }; // end EnableSharedStateHelper

    //! Performs loading and construction for a shared pointer that is derived from
    //! std::enable_shared_from_this
    /*! @param ar The archive
        @param ptr Raw pointer held by the shared_ptr
        @internal */
    template <class Archive, class T> inline
    void loadAndConstructSharedPtr( Archive & ar, T * ptr, std::true_type /* has_shared_from_this */ )
    {
      memory_detail::EnableSharedStateHelper<T> state( ptr );
      memory_detail::LoadAndConstructLoadWrapper<Archive, T> loadWrapper( ptr, [&](){ state.restore(); } );

      // let the user perform their initialization, shared state will be restored as soon as construct()
      // is called
      ar( CEREAL_NVP_("data", loadWrapper) );
    }

    //! Performs loading and construction for a shared pointer that is NOT derived from
    //! std::enable_shared_from_this
    /*! This is the typical case, where we simply pass the load wrapper to the
        archive.

        @param ar The archive
        @param ptr Raw pointer held by the shared_ptr
        @internal */
    template <class Archive, class T> inline
    void loadAndConstructSharedPtr( Archive & ar, T * ptr, std::false_type /* has_shared_from_this */ )
    {
      memory_detail::LoadAndConstructLoadWrapper<Archive, T> loadWrapper( ptr );
      ar( CEREAL_NVP_("data", loadWrapper) );
    }
  } // end namespace memory_detail

  //! Saving std::shared_ptr for non polymorphic types
  template <class Archive, class T> inline
  typename std::enable_if<!std::is_polymorphic<T>::value, void>::type
  CEREAL_SAVE_FUNCTION_NAME( Archive & ar, std::shared_ptr<T> const & ptr )
  {
    ar( CEREAL_NVP_("ptr_wrapper", memory_detail::make_ptr_wrapper( ptr )) );
  }

  //! Loading std::shared_ptr, case when no user load and construct for non polymorphic types
  template <class Archive, class T> inline
  typename std::enable_if<!std::is_polymorphic<T>::value, void>::type
  CEREAL_LOAD_FUNCTION_NAME( Archive & ar, std::shared_ptr<T> & ptr )
  {
    ar( CEREAL_NVP_("ptr_wrapper", memory_detail::make_ptr_wrapper( ptr )) );
  }

  //! Saving std::weak_ptr for non polymorphic types
  template <class Archive, class T> inline
  typename std::enable_if<!std::is_polymorphic<T>::value, void>::type
  CEREAL_SAVE_FUNCTION_NAME( Archive & ar, std::weak_ptr<T> const & ptr )
  {
    auto const sptr = ptr.lock();
    ar( CEREAL_NVP_("ptr_wrapper", memory_detail::make_ptr_wrapper( sptr )) );
  }

  //! Loading std::weak_ptr for non polymorphic types
  template <class Archive, class T> inline
  typename std::enable_if<!std::is_polymorphic<T>::value, void>::type
  CEREAL_LOAD_FUNCTION_NAME( Archive & ar, std::weak_ptr<T> & ptr )
  {
    std::shared_ptr<T> sptr;
    ar( CEREAL_NVP_("ptr_wrapper", memory_detail::make_ptr_wrapper( sptr )) );
    ptr = sptr;
  }

  //! Saving std::unique_ptr for non polymorphic types
  template <class Archive, class T, class D> inline
  typename std::enable_if<!std::is_polymorphic<T>::value, void>::type
  CEREAL_SAVE_FUNCTION_NAME( Archive & ar, std::unique_ptr<T, D> const & ptr )
  {
    ar( CEREAL_NVP_("ptr_wrapper", memory_detail::make_ptr_wrapper( ptr )) );
  }

  //! Loading std::unique_ptr, case when user provides load_and_construct for non polymorphic types
  template <class Archive, class T, class D> inline
  typename std::enable_if<!std::is_polymorphic<T>::value, void>::type
  CEREAL_LOAD_FUNCTION_NAME( Archive & ar, std::unique_ptr<T, D> & ptr )
  {
    ar( CEREAL_NVP_("ptr_wrapper", memory_detail::make_ptr_wrapper( ptr )) );
  }

  // ######################################################################
  // Pointer wrapper implementations follow below

  //! Saving std::shared_ptr (wrapper implementation)
  /*! @internal */
  template <class Archive, class T> inline
  void CEREAL_SAVE_FUNCTION_NAME( Archive & ar, memory_detail::PtrWrapper<std::shared_ptr<T> const &> const & wrapper )
  {
    auto & ptr = wrapper.ptr;

    uint32_t id = ar.registerSharedPointer( ptr );
    ar( CEREAL_NVP_("id", id) );

    if( id & detail::msb_32bit )
    {
      ar( CEREAL_NVP_("data", *ptr) );
    }
  }

  //! Loading std::shared_ptr, case when user load and construct (wrapper implementation)
  /*! @internal */
  template <class Archive, class T> inline
  typename std::enable_if<traits::has_load_and_construct<T, Archive>::value, void>::type
  CEREAL_LOAD_FUNCTION_NAME( Archive & ar, memory_detail::PtrWrapper<std::shared_ptr<T> &> & wrapper )
  {
    uint32_t id;

    ar( CEREAL_NVP_("id", id) );

    if( id & detail::msb_32bit )
    {
      // Storage type for the pointer - since we can't default construct this type,
      // we'll allocate it using std::aligned_storage and use a custom deleter
      using ST = typename std::aligned_storage<sizeof(T), CEREAL_ALIGNOF(T)>::type;

      // Valid flag - set to true once construction finishes
      //  This prevents us from calling the destructor on
      //  uninitialized data.
      auto valid = std::make_shared<bool>( false );

      // Allocate our storage, which we will treat as
      //  uninitialized until initialized with placement new
      using NonConstT = typename std::remove_const<T>::type;
      std::shared_ptr<NonConstT> ptr(reinterpret_cast<NonConstT *>(new ST()),
          [=]( NonConstT * t )
          {
            if( *valid )
              t->~T();

            delete reinterpret_cast<ST *>( t );
          } );

      // Register the pointer
      ar.registerSharedPointer( id, ptr );

      // Perform the actual loading and allocation
      memory_detail::loadAndConstructSharedPtr( ar, ptr.get(), typename ::cereal::traits::has_shared_from_this<NonConstT>::type() );

      // Mark pointer as valid (initialized)
      *valid = true;
      wrapper.ptr = std::move(ptr);
    }
    else
      wrapper.ptr = std::static_pointer_cast<T>(ar.getSharedPointer(id));
  }

  //! Loading std::shared_ptr, case when no user load and construct (wrapper implementation)
  /*! @internal */
  template <class Archive, class T> inline
  typename std::enable_if<!traits::has_load_and_construct<T, Archive>::value, void>::type
  CEREAL_LOAD_FUNCTION_NAME( Archive & ar, memory_detail::PtrWrapper<std::shared_ptr<T> &> & wrapper )
  {
    uint32_t id;

    ar( CEREAL_NVP_("id", id) );

    if( id & detail::msb_32bit )
    {
      using NonConstT = typename std::remove_const<T>::type;
      std::shared_ptr<NonConstT> ptr( detail::Construct<NonConstT, Archive>::load_andor_construct() );
      ar.registerSharedPointer( id, ptr );
      ar( CEREAL_NVP_("data", *ptr) );
      wrapper.ptr = std::move(ptr);
    }
    else
      wrapper.ptr = std::static_pointer_cast<T>(ar.getSharedPointer(id));
  }

  //! Saving std::unique_ptr (wrapper implementation)
  /*! @internal */
  template <class Archive, class T, class D> inline
  void CEREAL_SAVE_FUNCTION_NAME( Archive & ar, memory_detail::PtrWrapper<std::unique_ptr<T, D> const &> const & wrapper )
  {
    auto & ptr = wrapper.ptr;

    // unique_ptr get one byte of metadata which signifies whether they were a nullptr
    // 0 == nullptr
    // 1 == not null

    if( !ptr )
      ar( CEREAL_NVP_("valid", uint8_t(0)) );
    else
    {
      ar( CEREAL_NVP_("valid", uint8_t(1)) );
      ar( CEREAL_NVP_("data", *ptr) );
    }
  }

  //! Loading std::unique_ptr, case when user provides load_and_construct (wrapper implementation)
  /*! @internal */
  template <class Archive, class T, class D> inline
  typename std::enable_if<traits::has_load_and_construct<T, Archive>::value, void>::type
  CEREAL_LOAD_FUNCTION_NAME( Archive & ar, memory_detail::PtrWrapper<std::unique_ptr<T, D> &> & wrapper )
  {
    uint8_t isValid;
    ar( CEREAL_NVP_("valid", isValid) );

    auto & ptr = wrapper.ptr;

    if( isValid )
    {
      using NonConstT = typename std::remove_const<T>::type;
      // Storage type for the pointer - since we can't default construct this type,
      // we'll allocate it using std::aligned_storage
      using ST = typename std::aligned_storage<sizeof(NonConstT), CEREAL_ALIGNOF(NonConstT)>::type;

      // Allocate storage - note the ST type so that deleter is correct if
      //                    an exception is thrown before we are initialized
      std::unique_ptr<ST> stPtr( new ST() );

      // Use wrapper to enter into "data" nvp of ptr_wrapper
      memory_detail::LoadAndConstructLoadWrapper<Archive, NonConstT> loadWrapper( reinterpret_cast<NonConstT *>( stPtr.get() ) );

      // Initialize storage
      ar( CEREAL_NVP_("data", loadWrapper) );

      // Transfer ownership to correct unique_ptr type
      ptr.reset( reinterpret_cast<T *>( stPtr.release() ) );
    }
    else
      ptr.reset( nullptr );
  }

  //! Loading std::unique_ptr, case when no load_and_construct (wrapper implementation)
  /*! @internal */
  template <class Archive, class T, class D> inline
  typename std::enable_if<!traits::has_load_and_construct<T, Archive>::value, void>::type
  CEREAL_LOAD_FUNCTION_NAME( Archive & ar, memory_detail::PtrWrapper<std::unique_ptr<T, D> &> & wrapper )
  {
    uint8_t isValid;
    ar( CEREAL_NVP_("valid", isValid) );

    if( isValid )
    {
      using NonConstT = typename std::remove_const<T>::type;
      std::unique_ptr<NonConstT, D> ptr( detail::Construct<NonConstT, Archive>::load_andor_construct() );
      ar( CEREAL_NVP_( "data", *ptr ) );
      wrapper.ptr = std::move(ptr);
    }
    else
    {
      wrapper.ptr.reset( nullptr );
    }
  }
} // namespace cereal

// automatically include polymorphic support
#include "cereal/types/polymorphic.hpp"

#endif // CEREAL_TYPES_SHARED_PTR_HPP_
```
