## Supplemental code for "The motif composition of variable-number tandem repeats impacts gene expression": polymorphic.hpp

```
/*! \file polymorphic.hpp
    \brief Support for pointers to polymorphic base classes
    \ingroup OtherTypes */
/*
  Copyright (c) 2014, Randolph Voorhies, Shane Grant
  All rights reserved.

  Redistribution and use in source and binary forms, with or without
  modification, are permitted provided that the following conditions are met:
      * Redistributions of source code must retain the above copyright
        notice, this list of conditions and the following disclaimer.
      * Redistributions in binary form must reproduce the above copyright
        notice, this list of conditions and the following disclaimer in the
        documentation and/or other materials provided with the distribution.
      * Neither the name of cereal nor the
        names of its contributors may be used to endorse or promote products
        derived from this software without specific prior written permission.

  THIS SOFTWARE IS PROVIDED BY THE COPYRIGHT HOLDERS AND CONTRIBUTORS "AS IS" AND
  ANY EXPRESS OR IMPLIED WARRANTIES, INCLUDING, BUT NOT LIMITED TO, THE IMPLIED
  WARRANTIES OF MERCHANTABILITY AND FITNESS FOR A PARTICULAR PURPOSE ARE
  DISCLAIMED. IN NO EVENT SHALL RANDOLPH VOORHIES OR SHANE GRANT BE LIABLE FOR ANY
  DIRECT, INDIRECT, INCIDENTAL, SPECIAL, EXEMPLARY, OR CONSEQUENTIAL DAMAGES
  (INCLUDING, BUT NOT LIMITED TO, PROCUREMENT OF SUBSTITUTE GOODS OR SERVICES;
  LOSS OF USE, DATA, OR PROFITS; OR BUSINESS INTERRUPTION) HOWEVER CAUSED AND
  ON ANY THEORY OF LIABILITY, WHETHER IN CONTRACT, STRICT LIABILITY, OR TORT
  (INCLUDING NEGLIGENCE OR OTHERWISE) ARISING IN ANY WAY OUT OF THE USE OF THIS
  SOFTWARE, EVEN IF ADVISED OF THE POSSIBILITY OF SUCH DAMAGE.
*/
#ifndef CEREAL_TYPES_POLYMORPHIC_HPP_
#define CEREAL_TYPES_POLYMORPHIC_HPP_

#include "cereal/cereal.hpp"
#include "cereal/types/memory.hpp"

#include "cereal/details/util.hpp"
#include "cereal/details/helpers.hpp"
#include "cereal/details/traits.hpp"
#include "cereal/details/polymorphic_impl.hpp"

#if defined(_MSC_VER) && _MSC_VER < 1916
#define CEREAL_STATIC_CONSTEXPR static
#else
#define CEREAL_STATIC_CONSTEXPR static constexpr
#endif

//! Registers a derived polymorphic type with cereal
/*! Polymorphic types must be registered before smart
    pointers to them can be serialized.  Note that base
    classes do not need to be registered.

    Registering a type lets cereal know how to properly
    serialize it when a smart pointer to a base object is
    used in conjunction with a derived class.

    This assumes that all relevant archives have also
    previously been registered.  Registration for archives
    is usually done in the header file in which they are
    defined.  This means that type registration needs to
    happen after specific archives to be used are included.

    It is recommended that type registration be done in
    the header file in which the type is declared.

    Registration can also be placed in a source file,
    but this may require the use of the
    CEREAL_REGISTER_DYNAMIC_INIT macro (see below).

    Registration may be called repeatedly for the same
    type in different translation units to add support
    for additional archives if they are not initially
    available (included and registered).

    When building serialization support as a DLL on
    Windows, registration must happen in the header file.
    On Linux and Mac things should still work properly
    if placed in a source file, but see the above comments
    on registering in source files.

    Polymorphic support in cereal requires RTTI to be
    enabled */
#define CEREAL_REGISTER_TYPE(...)                                        \
  namespace cereal {                                                     \
  namespace detail {                                                     \
  template <>                                                            \
  struct binding_name<__VA_ARGS__>                                       \
  {                                                                      \
    CEREAL_STATIC_CONSTEXPR char const * name() { return #__VA_ARGS__; } \
  };                                                                     \
  } } /* end namespaces */                                               \
  CEREAL_BIND_TO_ARCHIVES(__VA_ARGS__)

//! Registers a polymorphic type with cereal, giving it a
//! user defined name
/*! In some cases the default name used with
    CEREAL_REGISTER_TYPE (the name of the type) may not be
    suitable.  This macro allows any name to be associated
    with the type.  The name should be unique */
#define CEREAL_REGISTER_TYPE_WITH_NAME(T, Name)                     \
  namespace cereal {                                                \
  namespace detail {                                                \
  template <>                                                       \
  struct binding_name<T>                                            \
  { CEREAL_STATIC_CONSTEXPR char const * name() { return Name; } }; \
  } } /* end namespaces */                                          \
  CEREAL_BIND_TO_ARCHIVES(T)

//! Registers the base-derived relationship for a polymorphic type
/*! When polymorphic serialization occurs, cereal needs to know how to
    properly cast between derived and base types for the polymorphic
    type. Normally this happens automatically whenever cereal::base_class
    or cereal::virtual_base_class are used to serialize a base class. In
    cases where neither of these is ever called but a base class still
    exists, this explicit registration is required.

    The Derived class should be the most derived type that will be serialized,
    and the Base type any possible base that has not been covered under a base
    class serialization that will be used to store a Derived pointer.

    Placement of this is the same as for CEREAL_REGISTER_TYPE. */
#define CEREAL_REGISTER_POLYMORPHIC_RELATION(Base, Derived)                     \
  namespace cereal {                                                            \
  namespace detail {                                                            \
  template <>                                                                   \
  struct PolymorphicRelation<Base, Derived>                                     \
  { static void bind() { RegisterPolymorphicCaster<Base, Derived>::bind(); } }; \
  } } /* end namespaces */

//! Adds a way to force initialization of a translation unit containing
//! calls to CEREAL_REGISTER_TYPE
/*! In C++, dynamic initialization of non-local variables of a translation
    unit may be deferred until "the first odr-use of any function or variable
    defined in the same translation unit as the variable to be initialized."

    Informally, odr-use means that your program takes the address of or binds
    a reference directly to an object, which must have a definition.

    Since polymorphic type support in cereal relies on the dynamic
    initialization of certain global objects happening before
    serialization is performed, it is important to ensure that something
    from files that call CEREAL_REGISTER_TYPE is odr-used before serialization
    occurs, otherwise the registration will never take place.  This may often
    be the case when serialization is built as a shared library external from
    your main program.

    This macro, with any name of your choosing, should be placed into the
    source file that contains calls to CEREAL_REGISTER_TYPE.

    Its counterpart, CEREAL_FORCE_DYNAMIC_INIT, should be placed in its
    associated header file such that it is included in the translation units
    (source files) in which you want the registration to appear.

    @relates CEREAL_FORCE_DYNAMIC_INIT
    */
#define CEREAL_REGISTER_DYNAMIC_INIT(LibName)                \
  namespace cereal {                                         \
  namespace detail {                                         \
    void CEREAL_DLL_EXPORT dynamic_init_dummy_##LibName() {} \
  } } /* end namespaces */

//! Forces dynamic initialization of polymorphic support in a
//! previously registered source file
/*! @sa CEREAL_REGISTER_DYNAMIC_INIT

    See CEREAL_REGISTER_DYNAMIC_INIT for detailed explanation
    of how this macro should be used.  The name used should
    match that for CEREAL_REGISTER_DYNAMIC_INIT. */
#define CEREAL_FORCE_DYNAMIC_INIT(LibName)                 \
  namespace cereal {                                       \
  namespace detail {                                       \
    void CEREAL_DLL_EXPORT dynamic_init_dummy_##LibName(); \
  } /* end detail */                                       \
  } /* end cereal */                                       \
  namespace {                                              \
    struct dynamic_init_##LibName {                        \
      dynamic_init_##LibName() {                           \
        ::cereal::detail::dynamic_init_dummy_##LibName();  \
      }                                                    \
    } dynamic_init_instance_##LibName;                     \
  } /* end anonymous namespace */

namespace cereal
{
  namespace polymorphic_detail
  {
    //! Error message used for unregistered polymorphic types
    /*! @internal */
    #define UNREGISTERED_POLYMORPHIC_EXCEPTION(LoadSave, Name)                                                                                      \
      throw cereal::Exception("Trying to " #LoadSave " an unregistered polymorphic type (" + Name + ").\n"                                          \
                              "Make sure your type is registered with CEREAL_REGISTER_TYPE and that the archive "                                   \
                              "you are using was included (and registered with CEREAL_REGISTER_ARCHIVE) prior to calling CEREAL_REGISTER_TYPE.\n"   \
                              "If your type is already registered and you still see this error, you may need to use CEREAL_REGISTER_DYNAMIC_INIT.");

    //! Get an input binding from the given archive by deserializing the type meta data
    /*! @internal */
    template<class Archive> inline
    typename ::cereal::detail::InputBindingMap<Archive>::Serializers getInputBinding(Archive & ar, std::uint32_t const nameid)
    {
      // If the nameid is zero, we serialized a null pointer
      if(nameid == 0)
      {
        typename ::cereal::detail::InputBindingMap<Archive>::Serializers emptySerializers;
        emptySerializers.shared_ptr = [](void*, std::shared_ptr<void> & ptr, std::type_info const &) { ptr.reset(); };
        emptySerializers.unique_ptr = [](void*, std::unique_ptr<void, ::cereal::detail::EmptyDeleter<void>> & ptr, std::type_info const &) { ptr.reset( nullptr ); };
        return emptySerializers;
      }

      std::string name;
      if(nameid & detail::msb_32bit)
      {
        ar( CEREAL_NVP_("polymorphic_name", name) );
        ar.registerPolymorphicName(nameid, name);
      }
      else
        name = ar.getPolymorphicName(nameid);

      auto const & bindingMap = detail::StaticObject<detail::InputBindingMap<Archive>>::getInstance().map;

      auto binding = bindingMap.find(name);
      if(binding == bindingMap.end())
        UNREGISTERED_POLYMORPHIC_EXCEPTION(load, name)
      return binding->second;
    }

    //! Serialize a shared_ptr if the 2nd msb in the nameid is set, and if we can actually construct the pointee
    /*! This check lets us try and skip doing polymorphic machinery if we can get away with
        using the derived class serialize function

        Note that on MSVC 2013 preview, is_default_constructible<T> returns true for abstract classes with
        default constructors, but on clang/gcc this will return false.  So we also need to check for that here.
        @internal */
    template<class Archive, class T> inline
    typename std::enable_if<(traits::is_default_constructible<T>::value
                             || traits::has_load_and_construct<T, Archive>::value)
                             && !std::is_abstract<T>::value, bool>::type
    serialize_wrapper(Archive & ar, std::shared_ptr<T> & ptr, std::uint32_t const nameid)
    {
      if(nameid & detail::msb2_32bit)
      {
        ar( CEREAL_NVP_("ptr_wrapper", memory_detail::make_ptr_wrapper(ptr)) );
        return true;
      }
      return false;
    }

    //! Serialize a unique_ptr if the 2nd msb in the nameid is set, and if we can actually construct the pointee
    /*! This check lets us try and skip doing polymorphic machinery if we can get away with
        using the derived class serialize function
        @internal */
    template<class Archive, class T, class D> inline
    typename std::enable_if<(traits::is_default_constructible<T>::value
                             || traits::has_load_and_construct<T, Archive>::value)
                             && !std::is_abstract<T>::value, bool>::type
    serialize_wrapper(Archive & ar, std::unique_ptr<T, D> & ptr, std::uint32_t const nameid)
    {
      if(nameid & detail::msb2_32bit)
      {
        ar( CEREAL_NVP_("ptr_wrapper", memory_detail::make_ptr_wrapper(ptr)) );
        return true;
      }
      return false;
    }

    //! Serialize a shared_ptr if the 2nd msb in the nameid is set, and if we can actually construct the pointee
    /*! This case is for when we can't actually construct the shared pointer.  Normally this would be caught
        as the pointer itself is serialized, but since this is a polymorphic pointer, if we tried to serialize
        the pointer we'd end up back here recursively.  So we have to catch the error here as well, if
        this was a polymorphic type serialized by its proper pointer type
        @internal */
    template<class Archive, class T> inline
    typename std::enable_if<(!traits::is_default_constructible<T>::value
                             && !traits::has_load_and_construct<T, Archive>::value)
                             || std::is_abstract<T>::value, bool>::type
    serialize_wrapper(Archive &, std::shared_ptr<T> &, std::uint32_t const nameid)
    {
      if(nameid & detail::msb2_32bit)
        throw cereal::Exception("Cannot load a polymorphic type that is not default constructable and does not have a load_and_construct function");
      return false;
    }

    //! Serialize a unique_ptr if the 2nd msb in the nameid is set, and if we can actually construct the pointee
    /*! This case is for when we can't actually construct the unique pointer.  Normally this would be caught
        as the pointer itself is serialized, but since this is a polymorphic pointer, if we tried to serialize
        the pointer we'd end up back here recursively.  So we have to catch the error here as well, if
        this was a polymorphic type serialized by its proper pointer type
        @internal */
    template<class Archive, class T, class D> inline
     typename std::enable_if<(!traits::is_default_constructible<T>::value
                               && !traits::has_load_and_construct<T, Archive>::value)
                               || std::is_abstract<T>::value, bool>::type
    serialize_wrapper(Archive &, std::unique_ptr<T, D> &, std::uint32_t const nameid)
    {
      if(nameid & detail::msb2_32bit)
        throw cereal::Exception("Cannot load a polymorphic type that is not default constructable and does not have a load_and_construct function");
      return false;
    }
  } // polymorphic_detail

  // ######################################################################
  // Pointer serialization for polymorphic types

  //! Saving std::shared_ptr for polymorphic types, abstract
  template <class Archive, class T> inline
  typename std::enable_if<std::is_polymorphic<T>::value && std::is_abstract<T>::value, void>::type
  CEREAL_SAVE_FUNCTION_NAME( Archive & ar, std::shared_ptr<T> const & ptr )
  {
    if(!ptr)
    {
      // same behavior as nullptr in memory implementation
      ar( CEREAL_NVP_("polymorphic_id", std::uint32_t(0)) );
      return;
    }

    std::type_info const & ptrinfo = typeid(*ptr.get());
    static std::type_info const & tinfo = typeid(T);
    // ptrinfo can never be equal to T info since we can't have an instance
    // of an abstract object
    //  this implies we need to do the lookup

    auto const & bindingMap = detail::StaticObject<detail::OutputBindingMap<Archive>>::getInstance().map;

    auto binding = bindingMap.find(std::type_index(ptrinfo));
    if(binding == bindingMap.end())
      UNREGISTERED_POLYMORPHIC_EXCEPTION(save, cereal::util::demangle(ptrinfo.name()))

    binding->second.shared_ptr(&ar, ptr.get(), tinfo);
  }

  //! Saving std::shared_ptr for polymorphic types, not abstract
  template <class Archive, class T> inline
  typename std::enable_if<std::is_polymorphic<T>::value && !std::is_abstract<T>::value, void>::type
  CEREAL_SAVE_FUNCTION_NAME( Archive & ar, std::shared_ptr<T> const & ptr )
  {
    if(!ptr)
    {
      // same behavior as nullptr in memory implementation
      ar( CEREAL_NVP_("polymorphic_id", std::uint32_t(0)) );
      return;
    }

    std::type_info const & ptrinfo = typeid(*ptr.get());
    static std::type_info const & tinfo = typeid(T);

    if(ptrinfo == tinfo)
    {
      // The 2nd msb signals that the following pointer does not need to be
      // cast with our polymorphic machinery
      ar( CEREAL_NVP_("polymorphic_id", detail::msb2_32bit) );

      ar( CEREAL_NVP_("ptr_wrapper", memory_detail::make_ptr_wrapper(ptr)) );

      return;
    }

    auto const & bindingMap = detail::StaticObject<detail::OutputBindingMap<Archive>>::getInstance().map;

    auto binding = bindingMap.find(std::type_index(ptrinfo));
    if(binding == bindingMap.end())
      UNREGISTERED_POLYMORPHIC_EXCEPTION(save, cereal::util::demangle(ptrinfo.name()))

    binding->second.shared_ptr(&ar, ptr.get(), tinfo);
  }

  //! Loading std::shared_ptr for polymorphic types
  template <class Archive, class T> inline
  typename std::enable_if<std::is_polymorphic<T>::value, void>::type
  CEREAL_LOAD_FUNCTION_NAME( Archive & ar, std::shared_ptr<T> & ptr )
  {
    std::uint32_t nameid;
    ar( CEREAL_NVP_("polymorphic_id", nameid) );

    // Check to see if we can skip all of this polymorphism business
    if(polymorphic_detail::serialize_wrapper(ar, ptr, nameid))
      return;

    auto binding = polymorphic_detail::getInputBinding(ar, nameid);
    std::shared_ptr<void> result;
    binding.shared_ptr(&ar, result, typeid(T));
    ptr = std::static_pointer_cast<T>(result);
  }

  //! Saving std::weak_ptr for polymorphic types
  template <class Archive, class T> inline
  typename std::enable_if<std::is_polymorphic<T>::value, void>::type
  CEREAL_SAVE_FUNCTION_NAME( Archive & ar, std::weak_ptr<T> const & ptr )
  {
    auto const sptr = ptr.lock();
    ar( CEREAL_NVP_("locked_ptr", sptr) );
  }

  //! Loading std::weak_ptr for polymorphic types
  template <class Archive, class T> inline
  typename std::enable_if<std::is_polymorphic<T>::value, void>::type
  CEREAL_LOAD_FUNCTION_NAME( Archive & ar, std::weak_ptr<T> & ptr )
  {
    std::shared_ptr<T> sptr;
    ar( CEREAL_NVP_("locked_ptr", sptr) );
    ptr = sptr;
  }

  //! Saving std::unique_ptr for polymorphic types that are abstract
  template <class Archive, class T, class D> inline
  typename std::enable_if<std::is_polymorphic<T>::value && std::is_abstract<T>::value, void>::type
  CEREAL_SAVE_FUNCTION_NAME( Archive & ar, std::unique_ptr<T, D> const & ptr )
  {
    if(!ptr)
    {
      // same behavior as nullptr in memory implementation
      ar( CEREAL_NVP_("polymorphic_id", std::uint32_t(0)) );
      return;
    }

    std::type_info const & ptrinfo = typeid(*ptr.get());
    static std::type_info const & tinfo = typeid(T);
    // ptrinfo can never be equal to T info since we can't have an instance
    // of an abstract object
    //  this implies we need to do the lookup

    auto const & bindingMap = detail::StaticObject<detail::OutputBindingMap<Archive>>::getInstance().map;

    auto binding = bindingMap.find(std::type_index(ptrinfo));
    if(binding == bindingMap.end())
      UNREGISTERED_POLYMORPHIC_EXCEPTION(save, cereal::util::demangle(ptrinfo.name()))

    binding->second.unique_ptr(&ar, ptr.get(), tinfo);
  }

  //! Saving std::unique_ptr for polymorphic types, not abstract
  template <class Archive, class T, class D> inline
  typename std::enable_if<std::is_polymorphic<T>::value && !std::is_abstract<T>::value, void>::type
  CEREAL_SAVE_FUNCTION_NAME( Archive & ar, std::unique_ptr<T, D> const & ptr )
  {
    if(!ptr)
    {
      // same behavior as nullptr in memory implementation
      ar( CEREAL_NVP_("polymorphic_id", std::uint32_t(0)) );
      return;
    }

    std::type_info const & ptrinfo = typeid(*ptr.get());
    static std::type_info const & tinfo = typeid(T);

    if(ptrinfo == tinfo)
    {
      // The 2nd msb signals that the following pointer does not need to be
      // cast with our polymorphic machinery
      ar( CEREAL_NVP_("polymorphic_id", detail::msb2_32bit) );

      ar( CEREAL_NVP_("ptr_wrapper", memory_detail::make_ptr_wrapper(ptr)) );

      return;
    }

    auto const & bindingMap = detail::StaticObject<detail::OutputBindingMap<Archive>>::getInstance().map;

    auto binding = bindingMap.find(std::type_index(ptrinfo));
    if(binding == bindingMap.end())
      UNREGISTERED_POLYMORPHIC_EXCEPTION(save, cereal::util::demangle(ptrinfo.name()))

    binding->second.unique_ptr(&ar, ptr.get(), tinfo);
  }

  //! Loading std::unique_ptr, case when user provides load_and_construct for polymorphic types
  template <class Archive, class T, class D> inline
  typename std::enable_if<std::is_polymorphic<T>::value, void>::type
  CEREAL_LOAD_FUNCTION_NAME( Archive & ar, std::unique_ptr<T, D> & ptr )
  {
    std::uint32_t nameid;
    ar( CEREAL_NVP_("polymorphic_id", nameid) );

    // Check to see if we can skip all of this polymorphism business
    if(polymorphic_detail::serialize_wrapper(ar, ptr, nameid))
      return;

    auto binding = polymorphic_detail::getInputBinding(ar, nameid);
    std::unique_ptr<void, ::cereal::detail::EmptyDeleter<void>> result;
    binding.unique_ptr(&ar, result, typeid(T));
    ptr.reset(static_cast<T*>(result.release()));
  }

  #undef UNREGISTERED_POLYMORPHIC_EXCEPTION
} // namespace cereal
#endif // CEREAL_TYPES_POLYMORPHIC_HPP_
```
