## Supplemental code for "The motif composition of variable-number tandem repeats impacts gene expression": string.hpp

```
/*! \file string.hpp
    \brief Support for types found in \<string\>
    \ingroup STLSupport */
/*
  Copyright (c) 2014, Randolph Voorhies, Shane Grant
  All rights reserved.

  Redistribution and use in source and binary forms, with or without
  modification, are permitted provided that the following conditions are met:
      * Redistributions of source code must retain the above copyright
        notice, this list of conditions and the following disclaimer.
      * Redistributions in binary form must reproduce the above copyright
        notice, this list of conditions and the following disclaimer in the
        documentation and/or other materials provided with the distribution.
      * Neither the name of cereal nor the
        names of its contributors may be used to endorse or promote products
        derived from this software without specific prior written permission.

  THIS SOFTWARE IS PROVIDED BY THE COPYRIGHT HOLDERS AND CONTRIBUTORS "AS IS" AND
  ANY EXPRESS OR IMPLIED WARRANTIES, INCLUDING, BUT NOT LIMITED TO, THE IMPLIED
  WARRANTIES OF MERCHANTABILITY AND FITNESS FOR A PARTICULAR PURPOSE ARE
  DISCLAIMED. IN NO EVENT SHALL RANDOLPH VOORHIES OR SHANE GRANT BE LIABLE FOR ANY
  DIRECT, INDIRECT, INCIDENTAL, SPECIAL, EXEMPLARY, OR CONSEQUENTIAL DAMAGES
  (INCLUDING, BUT NOT LIMITED TO, PROCUREMENT OF SUBSTITUTE GOODS OR SERVICES;
  LOSS OF USE, DATA, OR PROFITS; OR BUSINESS INTERRUPTION) HOWEVER CAUSED AND
  ON ANY THEORY OF LIABILITY, WHETHER IN CONTRACT, STRICT LIABILITY, OR TORT
  (INCLUDING NEGLIGENCE OR OTHERWISE) ARISING IN ANY WAY OUT OF THE USE OF THIS
  SOFTWARE, EVEN IF ADVISED OF THE POSSIBILITY OF SUCH DAMAGE.
*/
#ifndef CEREAL_TYPES_STRING_HPP_
#define CEREAL_TYPES_STRING_HPP_

#include "cereal/cereal.hpp"
#include <string>

namespace cereal
{
  //! Serialization for basic_string types, if binary data is supported
  template<class Archive, class CharT, class Traits, class Alloc> inline
  typename std::enable_if<traits::is_output_serializable<BinaryData<CharT>, Archive>::value, void>::type
  CEREAL_SAVE_FUNCTION_NAME(Archive & ar, std::basic_string<CharT, Traits, Alloc> const & str)
  {
    // Save number of chars + the data
    ar( make_size_tag( static_cast<size_type>(str.size()) ) );
    ar( binary_data( str.data(), str.size() * sizeof(CharT) ) );
  }

  //! Serialization for basic_string types, if binary data is supported
  template<class Archive, class CharT, class Traits, class Alloc> inline
  typename std::enable_if<traits::is_input_serializable<BinaryData<CharT>, Archive>::value, void>::type
  CEREAL_LOAD_FUNCTION_NAME(Archive & ar, std::basic_string<CharT, Traits, Alloc> & str)
  {
    size_type size;
    ar( make_size_tag( size ) );
    str.resize(static_cast<std::size_t>(size));
    ar( binary_data( const_cast<CharT *>( str.data() ), static_cast<std::size_t>(size) * sizeof(CharT) ) );
  }
} // namespace cereal

#endif // CEREAL_TYPES_STRING_HPP_
```
