## Supplemental code for "The motif composition of variable-number tandem repeats impacts gene expression": tuple.hpp

```
/*! \file tuple.hpp
    \brief Support for types found in \<tuple\>
    \ingroup STLSupport */
/*
  Copyright (c) 2014, Randolph Voorhies, Shane Grant
  All rights reserved.

  Redistribution and use in source and binary forms, with or without
  modification, are permitted provided that the following conditions are met:
      * Redistributions of source code must retain the above copyright
        notice, this list of conditions and the following disclaimer.
      * Redistributions in binary form must reproduce the above copyright
        notice, this list of conditions and the following disclaimer in the
        documentation and/or other materials provided with the distribution.
      * Neither the name of cereal nor the
        names of its contributors may be used to endorse or promote products
        derived from this software without specific prior written permission.

  THIS SOFTWARE IS PROVIDED BY THE COPYRIGHT HOLDERS AND CONTRIBUTORS "AS IS" AND
  ANY EXPRESS OR IMPLIED WARRANTIES, INCLUDING, BUT NOT LIMITED TO, THE IMPLIED
  WARRANTIES OF MERCHANTABILITY AND FITNESS FOR A PARTICULAR PURPOSE ARE
  DISCLAIMED. IN NO EVENT SHALL RANDOLPH VOORHIES OR SHANE GRANT BE LIABLE FOR ANY
  DIRECT, INDIRECT, INCIDENTAL, SPECIAL, EXEMPLARY, OR CONSEQUENTIAL DAMAGES
  (INCLUDING, BUT NOT LIMITED TO, PROCUREMENT OF SUBSTITUTE GOODS OR SERVICES;
  LOSS OF USE, DATA, OR PROFITS; OR BUSINESS INTERRUPTION) HOWEVER CAUSED AND
  ON ANY THEORY OF LIABILITY, WHETHER IN CONTRACT, STRICT LIABILITY, OR TORT
  (INCLUDING NEGLIGENCE OR OTHERWISE) ARISING IN ANY WAY OUT OF THE USE OF THIS
  SOFTWARE, EVEN IF ADVISED OF THE POSSIBILITY OF SUCH DAMAGE.
*/
#ifndef CEREAL_TYPES_TUPLE_HPP_
#define CEREAL_TYPES_TUPLE_HPP_

#include "cereal/cereal.hpp"
#include <tuple>

namespace cereal
{
  namespace tuple_detail
  {
    //! Creates a c string from a sequence of characters
    /*! The c string created will always be prefixed by "tuple_element"
        Based on code from: http://stackoverflow/a/20973438/710791
        @internal */
    template<char...Cs>
    struct char_seq_to_c_str
    {
      static const int size = 14;// Size of array for the word: tuple_element
      typedef const char (&arr_type)[sizeof...(Cs) + size];
      static const char str[sizeof...(Cs) + size];
    };

    // the word tuple_element plus a number
    //! @internal
    template<char...Cs>
    const char char_seq_to_c_str<Cs...>::str[sizeof...(Cs) + size] =
      {'t','u','p','l','e','_','e','l','e','m','e','n','t', Cs..., '\0'};

    //! Converts a number into a sequence of characters
    /*! @tparam Q The quotient of dividing the original number by 10
        @tparam R The remainder of dividing the original number by 10
        @tparam C The sequence built so far
        @internal */
    template <size_t Q, size_t R, char ... C>
    struct to_string_impl
    {
      using type = typename to_string_impl<Q/10, Q%10, static_cast<char>(R+std::size_t{'0'}), C...>::type;
    };

    //! Base case with no quotient
    /*! @internal */
    template <size_t R, char ... C>
    struct to_string_impl<0, R, C...>
    {
      using type = char_seq_to_c_str<static_cast<char>(R+std::size_t{'0'}), C...>;
    };

    //! Generates a c string for a given index of a tuple
    /*! Example use:
        @code{cpp}
        tuple_element_name<3>::c_str();// returns "tuple_element3"
        @endcode
        @internal */
    template<size_t T>
    struct tuple_element_name
    {
      using type = typename to_string_impl<T/10, T%10>::type;
      static const typename type::arr_type c_str(){ return type::str; }
    };

    // unwinds a tuple to save it
    //! @internal
    template <size_t Height>
    struct serialize
    {
      template <class Archive, class ... Types> inline
      static void apply( Archive & ar, std::tuple<Types...> & tuple )
      {
        serialize<Height - 1>::template apply( ar, tuple );
        ar( CEREAL_NVP_(tuple_element_name<Height - 1>::c_str(),
            std::get<Height - 1>( tuple )) );
      }
    };

    // Zero height specialization - nothing to do here
    //! @internal
    template <>
    struct serialize<0>
    {
      template <class Archive, class ... Types> inline
      static void apply( Archive &, std::tuple<Types...> & )
      { }
    };
  }

  //! Serializing for std::tuple
  template <class Archive, class ... Types> inline
  void CEREAL_SERIALIZE_FUNCTION_NAME( Archive & ar, std::tuple<Types...> & tuple )
  {
    tuple_detail::serialize<std::tuple_size<std::tuple<Types...>>::value>::template apply( ar, tuple );
  }
} // namespace cereal

#endif // CEREAL_TYPES_TUPLE_HPP_
```
