## Supplemental code for "The motif composition of variable-number tandem repeats impacts gene expression": valarray.hpp

```
/*! \file valarray.hpp
\brief Support for types found in \<valarray\>
\ingroup STLSupport */

/*
Copyright (c) 2014, Randolph Voorhies, Shane Grant
All rights reserved.

Redistribution and use in source and binary forms, with or without
modification, are permitted provided that the following conditions are met:
* Redistributions of source code must retain the above copyright
notice, this list of conditions and the following disclaimer.
* Redistributions in binary form must reproduce the above copyright
notice, this list of conditions and the following disclaimer in the
documentation and/or other materials provided with the distribution.
* Neither the name of cereal nor the
names of its contributors may be used to endorse or promote products
derived from this software without specific prior written permission.

THIS SOFTWARE IS PROVIDED BY THE COPYRIGHT HOLDERS AND CONTRIBUTORS "AS IS" AND
ANY EXPRESS OR IMPLIED WARRANTIES, INCLUDING, BUT NOT LIMITED TO, THE IMPLIED
WARRANTIES OF MERCHANTABILITY AND FITNESS FOR A PARTICULAR PURPOSE ARE
DISCLAIMED. IN NO EVENT SHALL RANDOLPH VOORHIES OR SHANE GRANT BE LIABLE FOR ANY
DIRECT, INDIRECT, INCIDENTAL, SPECIAL, EXEMPLARY, OR CONSEQUENTIAL DAMAGES
(INCLUDING, BUT NOT LIMITED TO, PROCUREMENT OF SUBSTITUTE GOODS OR SERVICES;
LOSS OF USE, DATA, OR PROFITS; OR BUSINESS INTERRUPTION) HOWEVER CAUSED AND
ON ANY THEORY OF LIABILITY, WHETHER IN CONTRACT, STRICT LIABILITY, OR TORT
(INCLUDING NEGLIGENCE OR OTHERWISE) ARISING IN ANY WAY OUT OF THE USE OF THIS
SOFTWARE, EVEN IF ADVISED OF THE POSSIBILITY OF SUCH DAMAGE.
*/

#ifndef CEREAL_TYPES_VALARRAY_HPP_
#define CEREAL_TYPES_VALARRAY_HPP_

#include "cereal/cereal.hpp"
#include <valarray>

namespace cereal
{
  //! Saving for std::valarray arithmetic types, using binary serialization, if supported
  template <class Archive, class T> inline
  typename std::enable_if<traits::is_output_serializable<BinaryData<T>, Archive>::value
                          && std::is_arithmetic<T>::value, void>::type
  CEREAL_SAVE_FUNCTION_NAME( Archive & ar, std::valarray<T> const & valarray )
  {
    ar( make_size_tag( static_cast<size_type>(valarray.size()) ) ); // number of elements
    ar( binary_data( &valarray[0], valarray.size() * sizeof(T) ) ); // &valarray[0] ok since guaranteed contiguous
  }

  //! Loading for std::valarray arithmetic types, using binary serialization, if supported
  template <class Archive, class T> inline
  typename std::enable_if<traits::is_input_serializable<BinaryData<T>, Archive>::value
                          && std::is_arithmetic<T>::value, void>::type
  CEREAL_LOAD_FUNCTION_NAME( Archive & ar, std::valarray<T> & valarray )
  {
    size_type valarraySize;
    ar( make_size_tag( valarraySize ) );

    valarray.resize( static_cast<std::size_t>( valarraySize ) );
    ar( binary_data( &valarray[0], static_cast<std::size_t>( valarraySize ) * sizeof(T) ) );
  }

  //! Saving for std::valarray all other types
  template <class Archive, class T> inline
  typename std::enable_if<!traits::is_output_serializable<BinaryData<T>, Archive>::value
                          || !std::is_arithmetic<T>::value, void>::type
  CEREAL_SAVE_FUNCTION_NAME( Archive & ar, std::valarray<T> const & valarray )
  {
    ar( make_size_tag( static_cast<size_type>(valarray.size()) ) ); // number of elements
    for(auto && v : valarray)
      ar(v);
  }

  //! Loading for std::valarray all other types
  template <class Archive, class T> inline
  typename std::enable_if<!traits::is_input_serializable<BinaryData<T>, Archive>::value
                          || !std::is_arithmetic<T>::value, void>::type
  CEREAL_LOAD_FUNCTION_NAME( Archive & ar, std::valarray<T> & valarray )
  {
    size_type valarraySize;
    ar( make_size_tag( valarraySize ) );

    valarray.resize( static_cast<size_t>( valarraySize ) );
    for(auto && v : valarray)
      ar(v);
  }
} // namespace cereal

#endif // CEREAL_TYPES_VALARRAY_HPP_
```
