## Supplemental code for "The motif composition of variable-number tandem repeats impacts gene expression": Makefile

```
PREFIX = /home/cmb-16/mjc/tsungyul/work/vntr/danbing-tk
TARGETS = bin/danbing-tk bin/vntr2kmers_thread bin/genPanKmers bin/ktools
TARGETSg = bin/danbing-tk_g bin/vntr2kmers_thread_g bin/genPanKmers_g bin/ktools_g
#TARGETS = aQueryFasta amphQueryFasta vntr2kmers kmer2dot seq2num num2seq rvseq bam2pe

CXX = g++ -std=c++11
LDFLAGS = -lpthread
dir_guard=@mkdir -p $(@D)

all: $(TARGETS)
allg: $(TARGETS) $(TARGETSg)


# dependencies between programs and .o files
bin/danbing-tk:	src/aQueryFasta_thread.cpp
	$(dir_guard)
	$(CXX) $(LDFLAGS) -O3 -I. -o bin/danbing-tk src/aQueryFasta_thread.cpp

bin/danbing-tk_g:	src/aQueryFasta_thread.cpp
	$(dir_guard)
	$(CXX) -g $(LDFLAGS) -I. -o bin/danbing-tk_g src/aQueryFasta_thread.cpp

bin/ktools:	src/kmertools.cpp
	$(dir_guard)
	$(CXX) -O3 -I. -o bin/ktools src/kmertools.cpp

bin/ktools_g:	src/kmertools.cpp
	$(dir_guard)
	$(CXX) -g -I. -o bin/ktools_g src/kmertools.cpp

bin/vntr2kmers_thread:	src/VNTR2kmers_thread.cpp
	$(dir_guard)
	$(CXX) -O3 -I. -o bin/vntr2kmers_thread src/VNTR2kmers_thread.cpp

bin/vntr2kmers_thread_g:	src/VNTR2kmers_thread.cpp
	$(dir_guard)
	$(CXX) -g -I. -o bin/vntr2kmers_thread_g src/VNTR2kmers_thread.cpp

bin/genPanKmers:	src/genPanKmers.cpp
	$(dir_guard)
	$(CXX) -O3 -I. -o bin/genPanKmers src/genPanKmers.cpp

bin/genPanKmers_g:	src/genPanKmers.cpp
	$(dir_guard)
	$(CXX) -g -I. -o bin/genPanKmers_g src/genPanKmers.cpp

bin/seq2num:	src/seq2num.cpp
	$(dir_guard)
	$(CXX) -O3 -o bin/seq2num src/seq2num.cpp

bin/num2seq:	src/num2seq.cpp
	$(dir_guard)
	$(CXX) -O3 -o bin/num2seq src/num2seq.cpp

bin/rvseq:		src/rvseq.cpp
	$(dir_guard)
	$(CXX) -O3 -o bin/rvseq src/rvseq.cpp

bin/bam2pe:		src/bam2pe.cpp
	$(dir_guard)
	$(CXX) -O3 -o bin/bam2pe src/bam2pe.cpp

#
# generic build rules
#

#$(TARGETS):
#	$(CXX) $^ -o $@

#.PHONY: install
#install: $(TARGETS)
#	mkdir -p $(PREFIX)/bin
#	cp $^ $(PREFIX)/bin/.

#uninstall: $(TARGETS)
#	for TARGET in $(TARGETS); do \
#		rm $(PREFIX)/bin/$$TARGET; \
#	done

clean:
	rm -f *.o *~ $(TARGETS)
```
