## Supplemental code for "The motif composition of variable-number tandem repeats impacts gene expression": GoodPanGenomeGraph.snakefile

```
import os
import numpy as np

configfile: "goodPanGenomeGraph.json"

srcdir = config["srcDir"]
indir = config["inputDir"]
outdir = config["outputDir"]

prune = config["pruning"] == "True"
gbpair = np.loadtxt(config["pairs"], dtype=object, ndmin=2)
genomes = gbpair[:,0].tolist()
bams = dict(gbpair) if prune else None
haps = ["0", "1"]
kmerTypes = ["tr", "ntr", "graph"]
binKmerTypes = ["graph.umap", "kmerDBi.umap", "kmerDBi.vv"]

aligner = config["AsmAligner"]
ksize = config["ksize"]
FS = int(config["flankSize"])
dist_merge = int(config["dist_merge"])
dist_scan = int(config["dist_scan"])
cth = config["countThreashold"]
rth = config["ratioThreashold"]
rstring = f'{rth*100:.0f}'
thcth = config["threadingCountThreshold"]
TRwindow = int(config["TRwindow"])
LB = int(config["sizeLowerBound"])
mbe_th1 = float(config["MBE_th1"])
mbe_th2 = float(config["MBE_th2"])
copts = config["clusterOpts"]


rule all:
    input:
        #chrsize = expand(outdir + "{genome}.{hap}.chrSize", genome=genomes, hap=haps),
        faAln = expand(outdir + "{genome}.{hap}.aln.foo", genome=genomes, hap=haps),
        lift = expand(outdir + "{genome}/lift.foo", genome=genomes),
        TRfa = expand(outdir + "{genome}.{hap}.tr.fasta", genome=genomes, hap=haps),
        #PBkmers = expand(outdir + "{genome}.PB.{kmerType}.kmers", genome=genomes, kmerType=kmerTypes),
        #rawPred = expand(outdir + "{genome}.rawLR.pred", genome=genomes),
        panKmers = expand(outdir + "pan.{kmerType}.kmers", kmerType=kmerTypes),
        binKmers = expand(outdir + "pan.{binKmerType}", binKmerType=binKmerTypes),
        #panILkmers = expand(outdir + "pan.{genome}.IL.tr.kmers", genome=genomes),
        #pred = expand(outdir + "{genome}.LR.pred", genome=genomes),
        bamcov = [outdir + "ctrl.cov"] if prune else []
        

rule IndexAsm:
    input:
        fa = expand(indir + "{{genome}}.{hap}.fa", hap=haps)
    output:
        chrsize = expand(outdir + "{{genome}}.{hap}.chrSize", hap=haps)
    resources:
        cores = 3,
        mem = 4,
    params:
        name = "IndexAsm",
        copts = copts,
        sd = srcdir,
        od = outdir,
        indir = indir,
        genomes = genomes,
    shell:"""
set -eu
cd {params.od}

ln -sf {input.fa} .
for hap in 0 1; do
    fa={params.indir}/{wildcards.genome}.$hap.fa
    ln -sf "$fa".fai .
done
"""

rule ComputeBamCoverage:
    input:
        ILbam = [bams[g] for g in genomes] if prune else [],
        #ILbam = expand(indir + "{genome}.final.cram", genome=genomes),
        #ILbai = expand(indir + "{genome}.final.cram.crai", genome=genomes)
    output:
        bamcov = outdir + "ctrl.cov"
    resources:
        cores = 4,
        mem = 4
    params:
        copts = copts,
        refctrl = config["refctrl"],
        genomes = genomes,
    shell:"""
set -eu

bams=( {input.ILbam} )
gi=0
for g in {params.genomes}; do
    samtools bedcov {params.refctrl} ${{bams[$gi]}} | awk '{{ print $4/($3-$2) }}' | tr '\n' '\t' | 
    awk -v g=$g -v gi=$gi 'BEGIN {{OFS="\t"}} {{$1=$1; print gi, g, $0}}'
    ((++gi))
done > {output.bamcov}
"""


def getMem():
    if aligner == "lra":
        return 60
    else:
        return 40

rule MapAsm2Ref:
    input:
        fa = indir + "{genome}.{hap}.fa",
    output:
        faAln = outdir + "{genome}.{hap}.aln.foo",
    resources:
        cores = 16,
        mem = lambda wildcards, attempt: getMem() + 20*(attempt-1),
    params:
        name = "MapAsm2Ref",
        faBam = outdir + "{genome}.{hap}.srt.bam",
        faPaf = outdir + "{genome}.{hap}.r2a.paf",
        od = outdir,
        copts = copts,
        ref = config["ref"],
        aligner = aligner
    shell:"""
ulimit -c 20000
set -eu
cd {params.od}

if [[ {params.aligner} == "lra" ]]; then
	lra align -t $(({resources.cores}-1)) -CONTIG {params.ref} {input.fa} -p s | samtools sort >{params.faBam} &&
	samtools index -@3 {params.faBam}
	touch {output.faAln}
elif [[ {params.aligner} == "minimap2" ]]; then
	minimap2 {input.fa} {params.ref} -t $(({resources.cores}-1)) -x asm5 -L -c --cs=long -o {params.faPaf}
	touch {output.faAln}
else
	echo "Invalid AsmAligner: {params.aligner}"
	exit 1
fi
"""


rule LiftTR:
    input:
        faAln = expand(outdir + "{{genome}}.{hap}.aln.foo", hap=haps)
    output:
        lift = outdir + "{genome}/lift.foo"
    resources:
        cores = 6,
        mem = lambda wildcards, attempt: 24 + 16*(attempt-1)
    params:
        name = "LiftTR",
        copts = copts,
        sd = srcdir,
        od = outdir,
        refTR = config["refTR"],
        aligner = aligner,
        faBam = expand(outdir + "{{genome}}.{hap}.srt.bam", hap=haps),
        faPaf = expand(outdir + "{{genome}}.{hap}.r2a.paf", hap=haps),
        LB = LB
    shell:"""
set -eu
ulimit -c 20000
mkdir -p {params.od}/{wildcards.genome}
cd {params.od}/{wildcards.genome}

### get asm TR regions
echo "Liftover asm regions"

bams=( {params.faBam} )
pafs=( {params.faPaf} )
cut -f 1-3 {params.refTR} > ref.bed
if [[ {params.aligner} == "lra" ]]; then
	for hap in 0 1; do
		{params.sd}/bin/samLiftover <(samtools view ${{bams[$hap]}}) ref.bed /dev/stdout --dir 1 --printNA |
		awk 'BEGIN {{OFS="\t"}} {{
			if ($3-$2 < {params.LB}) {{print "NA", "NA", "NA"}}
			else {{print $0}}
		}}' > tmp0.$hap.bed
	done
	{params.sd}/script/rmNAforBothBeds.py tmp0.?.bed tmp1.0.bed tmp1.1.bed
else
	cp ref.bed tmp0.m.bed
	for hap in 0 1; do
		paftools.js liftover -l 50 ${{pafs[$hap]}} ref.bed | sort -k1,1 -k2,2n -k3,3n > tmp0.l"$hap".bed
		{params.sd}/script/liftbed.clean.py tmp0.l"$hap".bed |
        sort -k1,1 -k2,2n -k3,3n |
        bedtools merge -c 1,4,5,6,7 -o count,collapse,collapse,collapse,collapse |
        awk '$4 == 1' | cut -f 1-3,5-8 >tmp0."$hap".bed
		bedtools map -c 4,5 -o collapse \
                -a tmp0.m.bed \
                -b <(awk 'BEGIN {{OFS="\t"}} {{print $4,$5,$6,$1":"$2":"$3,$7}}' tmp0.$hap.bed | sort -k1,1 -k2,2n -k3,3n ) > tmp0.m.bed.tmp &&
		mv tmp0.m.bed.tmp tmp0.m.bed
	done
	awk '$4 != "." && $6 != "." && $4 !~ /,/ && $6 !~ /,/' tmp0.m.bed > tmp0.mc.bed
	for hap in 0 1; do 
		cut -f 1,2,3,$(($hap*2+4)),$(($hap*2+5)) tmp0.mc.bed |
        tr ':' '\t' | awk 'BEGIN {{OFS="\t"}} {{print $4,$5,$6,$1,$2,$3,$7}}' > tmp1.$hap.bed
	done
    touch lift.foo
fi
"""

rule JointTRAnnotation:
    input:
        fa = expand(indir + "{genome}.{hap}.fa", genome=genomes, hap=haps),
        lift = expand(outdir + "{genome}/lift.foo", genome=genomes)
    output:
        mapping = outdir + "OrthoMap.v2.tsv",
        TRfa = expand(outdir + "{genome}.{hap}.tr.fasta", genome=genomes, hap=haps),
    resources:
        cores = 12,
        mem = lambda wildcards, attempt: 110
    params:
        name = "JointTRAnnotation",
        copts = copts,
        sd = srcdir,
        od = outdir,
        indir = indir,
        refTR = config["refTR"],
        ksize = ksize,
        FS = FS,
        dist_merge = dist_merge,
        dist_scan = dist_scan,
        LB = LB,
        TRwindow = TRwindow,
        th1 = mbe_th1,
        th2 = mbe_th2,
        pairs = config["pairs"],
        genomes = genomes
    shell:"""
set -eu
ulimit -c 20000
cd {params.od}

printf "Generating panbed"
cut -f 1-3 {params.refTR} >pan.tr.mbe.v0.bed
for g in {params.genomes}; do 
    printf "."
    bedtools map -c 1 -o count -a pan.tr.mbe.v0.bed -b <(cut -f 4-6 $g/tmp1.0.bed) >pan.tr.mbe.v0.bed.tmp && 
    mv pan.tr.mbe.v0.bed.tmp pan.tr.mbe.v0.bed
done
echo ""
mkdir -p MBE
{params.sd}/script/multiBoundaryExpansion.parallel.py {params.ksize} {params.dist_scan} {params.TRwindow} {params.pairs} pan.tr.mbe.v0.bed {params.th1} {params.th2} {resources.cores} {params.indir}
hi=0
for g in {params.genomes}; do
    for h in 0 1; do
        echo ">""$g"".""$h"
        cut -f $((4+4*hi))-$((6+4*hi)) pan.tr.mbe.v1.bed |
        awk 'BEGIN {{OFS="\t"}} {{print $0, NR-1}}' |
        grep -v "None" |
        sort -k1,1 -k2,2n -k3,3n >tmp.bed
        if [[ "$(cat tmp.bed | wc -l)" != "0" ]]; then
            bedtools merge -d {params.dist_merge} -c 4 -o collapse -i tmp.bed |
            cut -f 4 | {{ grep "," || true; }}
        fi
        ((++hi))
    done
done >mbe.m0.loci
rm tmp.bed
{params.sd}/script/mergeMBEbed.py {params.pairs} {params.th2}

### write fasta
echo "Fetching TR+flank" $(date)
hi=0
for g in {params.genomes}; do
    for h in 0 1; do
        cut -f $((4+4*hi))-$((6+4*hi)) pan.tr.mbe.v2.bed |
        grep -v "None" |
        awk 'BEGIN {{OFS="\t"}} {{
            $2=$2-{params.FS}
            $3=$3+{params.FS}
            print $0
        }}' |
        {params.sd}/script/SelectRegions.py /dev/stdin {params.indir}/"$g"."$h".fa /dev/stdout | 
        awk '{{if ($1 ~ />/) {{print}} else {{print toupper($0)}} }}' >"$g"."$h".tr.fasta
        ((++hi))
    done
done
"""


rule GenRawGenomeGraph:
    input:
        TRfa = expand(outdir + "{{genome}}.{hap}.tr.fasta", hap=haps),
        ILbam = lambda wildcards: [bams[wildcards.genome]] if prune else [],
        mapping = outdir + "OrthoMap.v2.tsv",
    output:
        rawPBkmers = expand(outdir + "{{genome}}.rawPB.{kmerType}.kmers", kmerType=kmerTypes),
        rawILkmers = [outdir + "{genome}.rawIL.tr.kmers"] if prune else []
    resources:
        cores = 24 if prune else 1,
        mem = lambda wildcards, attempt: 55 + 20*(attempt-1)
    params:
        name = "GenRawGenomeGraph",
        copts = copts,
        sd = srcdir,
        od = outdir,
        ksize = ksize,
        FS = FS,
        cth = cth,
        rth = rth,
        rstring = rstring,
        thcth = thcth,
        hi = lambda wildcards: 2*genomes.index(wildcards.genome),
        prune = int(prune)
    shell:"""
set -eu
ulimit -c 20000
cd {params.od}
module load gcc

{params.sd}/bin/vntr2kmers_thread -g -m <(cut -f $(({params.hi}+1)),$(({params.hi}+2)) {input.mapping}) -k {params.ksize} -fs {params.FS} -ntr {params.FS} -on {wildcards.genome}.rawPB -fa 2 {input.TRfa}

if [ {params.prune} == "1"  ]; then
    samtools fasta -@2 -n {input.ILbam} |
    {params.sd}/bin/bam2pe -fai /dev/stdin |
    {params.sd}/bin/danbing-tk -g {params.thcth} -k {params.ksize} -qs {params.od}/{wildcards.genome}.rawPB -fai /dev/stdin -o {wildcards.genome}.rawIL -p {resources.cores} -cth {params.cth} -rth {params.rth}
fi
"""


rule EvalRawGenomeGraph:
    input:
        rawPBkmers = expand(outdir + "{{genome}}.rawPB.{kmerType}.kmers", kmerType=kmerTypes),
        rawILkmers = outdir + "{genome}.rawIL.tr.kmers"
    output:
        rawPred = outdir + "{genome}.rawLR.pred"
    resources:
        cores = 12,
        mem = 8
    params:
        name = "EvalRawGenomeGraph",
        copts = copts,
        sd = srcdir,
        od = outdir,
    shell:"""
set -eu
ulimit -c 20000
cd {params.od}

{params.sd}/script/kmers.linreg.py --mode invalid --R2threshold -2 {wildcards.genome}.rawPB.tr.kmers {wildcards.genome}.rawIL.tr.kmers {wildcards.genome}.rawLR
"""


rule GenPrunedGenomeGraph:
    input:
        rawILkmers = outdir + "{genome}.rawIL.tr.kmers",
        TRfa = expand(outdir + "{{genome}}.{hap}.tr.fasta", hap=haps),
        mapping = outdir + "OrthoMap.v2.tsv",
    output:
        PBkmers = expand(outdir + "{{genome}}.PB.{kmerType}.kmers", kmerType=kmerTypes)
    resources:
        cores = 2,
        mem = 20
    params:
        name = "GenPrunedGenomeGraph",
        copts = copts,
        sd = srcdir,
        od = outdir,
        ksize = ksize,
        FS = FS,
        hi = lambda wildcards: 2*genomes.index(wildcards.genome)
    shell:"""
cd {params.od}
ulimit -c 20000

awk '$1 ~ />/ || $2 == 0' {input.rawILkmers} |
{params.sd}/bin/vntr2kmers_thread -g -p /dev/stdin -m <(cut -f $(({params.hi}+1)),$(({params.hi}+2)) {input.mapping}) -k {params.ksize} -fs {params.FS} -ntr {params.FS} -on {wildcards.genome}.PB -fa 2 {input.TRfa}
"""

def getRPGGin():
    if prune:
        return outdir + "{genome}.PB.{kmerType}.kmers"
    else:
        return outdir + "{genome}.rawPB.{kmerType}.kmers"

rule GenPanGenomeGraph:
    input:
        PBkmers = expand(getRPGGin() , genome=genomes, kmerType=kmerTypes),
    output:
        panKmers = expand(outdir + "pan.{kmerType}.kmers", kmerType=kmerTypes)
    resources:
        cores = 2,
        mem = lambda wildcards, attempt: 62+8*attempt
    params:
        name = "GenPanGenomeGraph",
        copts = copts,
        sd = srcdir,
        od = outdir,
        kmerpref = " ".join([f'{g}.PB' if prune else f'{g}.rawPB' for g in genomes])
    shell:"""
cd {params.od}
ulimit -c 20000
module load gcc

{params.sd}/bin/genPanKmers -o pan -m - -k {params.kmerpref}
"""

rule GenSerializedGraphAndIndex:
    input:
        panKmers = expand(outdir + "pan.{kmerType}.kmers", kmerType=kmerTypes)
    output:
        binKmers = expand(outdir + "pan.{binKmerType}", binKmerType=binKmerTypes)
    resources:
        cores = 2,
        mem = lambda wildcards, attempt: 90+20*(attempt-1)
    params:
        name = "GenSerializedGraphAndIndex",
        copts = copts,
        sd = srcdir,
        od = outdir,
        pref = f"{outdir}/pan"
    shell:"""
cd {params.od}
ulimit -c 20000
module load gcc

{params.sd}/bin/ktools serialize {params.pref}
{params.sd}/bin/ktools ksi pan.tr.kmers >{params.pref}.tr.ksi
"""
```
