## Supplemental code for "The motif composition of variable-number tandem repeats impacts gene expression": LeaveOneOut.snakefile

```
import os
import numpy as np

configfile: "leaveOneOut.json"

srcdir = config["srcDir"]
outdir = config["outputDir"]
pdir = config["pangenomeDir"]
#pindir = config["pangenomeInputDir"] /home/cmb-17/mjc/vntr_genotyping/rpgg_k21_84k/input/

genomes = np.loadtxt(config["genomefile"], dtype=object)
gbpair = np.loadtxt(config["LOOpairs"], dtype=object)
g2bam = dict(gbpair)
LOOgenomes = gbpair[:,0]
LOOmask = np.isin(genomes, LOOgenomes)
kmerTypes = ["tr", "ntr", "graph"]
covbed = config["covbed"]

ksize = config["ksize"]
cth = config["countThreashold"]
rth = config["ratioThreashold"]
rstring = f'{rth*100:.0f}'
thcth = config["threadingCountThreshold"]
copts = config["clusterOpts"]


localrules: all, all2LOO

rule all:
    input:
        mapping = outdir + "OrthoMap.v2.tsv",
        extFoo = expand(outdir + "checkpoint/{genome}.extract.foo", genome=LOOgenomes),
        panGTfoo = expand(outdir + "checkpoint/{genome}.pan.gt.foo", genome=LOOgenomes),
        LOOGTfoo = expand(outdir + "checkpoint/{genome}.LOO.gt.foo", genome=LOOgenomes),
        LRfoo = expand(outdir + "checkpoint/{genome}.rawLR.foo", genome=LOOgenomes),
        #LOOPBkmers = expand(outdir + "LOO.{genome}.PB.{kmerType}.kmers", genome=LOOgenomes, kmerType=kmerTypes),
        #LOOILkmers = expand(outdir + "LOO.{genome}.IL.tr.kmers", genome=LOOgenomes),
        #panILkmers = expand(outdir + "pan.{genome}.IL.tr.kmers", genome=genomes),
        #mappedILkmers = expand(outdir + "{genome}.mappedIL.tr.kmers", genome=genomes),
        #pickle = outdir + "analysis/step1_results.pickle",
        #relErr = outdir + "analysis/rel_err.txt",


rule all2LOO:
    input:
        mapping = pdir + "OrthoMap.v2.tsv",
    output:
        mapping = outdir + "OrthoMap.v2.tsv",
        #LOOmap = expand(outdir + "LOO.{genome}.mapping", genome=LOOgenomes),
    resources:
        cores = 1,
        mem = 4,
    priority: 100
    params:
        m = LOOmask,
        lgs = LOOgenomes,
        od = outdir,
    run:
        import numpy as np
        lgs = params.lgs
        od = params.od
        tbl = np.loadtxt(input.mapping, dtype=object)
        nloci, nh = tbl.shape
        m = np.repeat(params.m, 2)
        tbl = tbl[:,m]
        np.savetxt(output.mapping, tbl, fmt='%s', delimiter='\t')
        #ncol = len(lgs)
        #out = np.full([nloci, ncol], ".", dtype=object)
        #for i in range(ncol):
        #    lm = np.logical_or(tbl[:,2*i] != ".", tbl[:,2*i+1] != ".")
        #    out[lm,i] = np.nonzero(lm)[0]
        #    out[~lm,i] = "."
        #for i in range(ncol):
        #    g = lgs[i]
        #    om = np.ones(ncol, dtype=bool)
        #    om[i] = False
        #    np.savetxt(f"{od}/LOO.{g}.mapping", out[:,om], fmt='%s', delimiter="\t")


rule GenPGG:
    input:
        PBkmers = expand(pdir + "{genome}.rawPB.{kmerType}.kmers", genome=LOOgenomes, kmerType=kmerTypes),
    output:
        ofoo = outdir + "checkpoint/{genome}.genpgg.foo",
    resources:
        cores = 2,
        mem = 50,
    priority: 99
    params:
        copts = copts,
        sd = srcdir,
        od = outdir,
        kmerpref = lambda wildcards: " ".join([f'{pdir}/{g}.rawPB' for g in LOOgenomes])
    shell:"""
cd {params.od}
ulimit -c 20000
mkdir -p checkpoint

{params.sd}/bin/genPanKmers -o pan -m - -k {params.kmerpref}
{params.sd}/bin/ktools serialize pan
"""

rule GenLOOPGG:
    input:
        PBkmers = expand(pdir + "{genome}.rawPB.{kmerType}.kmers", genome=LOOgenomes, kmerType=kmerTypes),
        #mapping = outdir + "OrthoMap.v2.tsv",
        #LOOmap = outdir + "LOO.{genome}.mapping",
    output:
        ofoo = outdir + "checkpoint/{genome}.genloopgg.foo",
    resources:
        cores = 2,
        mem = 50,
    priority: 99
    params:
        copts = copts,
        sd = srcdir,
        od = outdir,
        kmerpref = lambda wildcards: " ".join([f'{pdir}/{g}.rawPB' for g in LOOgenomes if g != wildcards.genome])
    shell:"""
cd {params.od}
ulimit -c 20000
mkdir -p checkpoint

#{params.sd}/bin/genPanKmers -o LOO.{wildcards.genome}.PB -m - -k {params.kmerpref}
{params.sd}/bin/ktools serialize LOO.{wildcards.genome}.PB
touch {output.ofoo}
"""


rule Extract:
    input:
        ifoo = outdir + "checkpoint/{genome}.genpgg.foo",
        ILbam = lambda wildcards: g2bam[wildcards.genome],
    output:
        ofoo = outdir + "checkpoint/{genome}.extract.foo",
    resources:
        cores = 16,
        mem = lambda wildcards, attempt: 40 + 20*(attempt-1)
    priority: 98
    params:
        copts = copts,
        sd = srcdir,
        od = outdir,
        pd = pdir,
        ksize = ksize,
        cth = cth,
        rth = rth,
        rstring = rstring,
    shell:"""
set -eu 
ulimit -c 20000
cd {params.od}
mkdir -p checkpoint

samtools fasta -@2 -n {input.ILbam} |
{params.sd}/bin/danbing-tk -e 1 -k {params.ksize} -qs {params.od}/pan -fa /dev/stdin \
                           -p {resources.cores} -cth {params.cth} -rth {params.rth} | gzip >{wildcards.genome}.e{params.cth}.fa.gz
touch {output.ofoo}
"""


rule LOOGenotying:
    input:
        ifoo = outdir + "checkpoint/{genome}.extract.foo",
        pggfoo = outdir + "checkpoint/{genome}.genloopgg.foo",
    output:
        ofoo = outdir + "checkpoint/{genome}.LOO.gt.foo"
    resources:
        cores = 16,
        mem = lambda wildcards, attempt: 50 + 20*(attempt-1),
    priority: 97
    params:
        fagz = outdir + "{genome}.e" + str(cth) + ".fa.gz",
        copts = copts,
        sd = srcdir,
        od = outdir,
        ksize = ksize,
        cth = cth,
        rth = rth,
        rstring = rstring,
        thcth = thcth,
    shell:"""
set -eu
ulimit -c 20000
cd {params.od}

zcat {params.fagz} |
{params.sd}/bin/danbing-tk -gc {params.thcth} -k {params.ksize} -qs LOO.{wildcards.genome}.PB -fai /dev/stdin -o LOO.{wildcards.genome}.IL -p {resources.cores} -cth {params.cth} -rth {params.rth}
touch {output.ofoo}
"""


rule GenotypeSamples:
    input:
        ifoo = outdir + "checkpoint/{genome}.extract.foo",
    output:
        ofoo = outdir + "checkpoint/{genome}.pan.gt.foo"
    resources:
        cores = 16,
        mem = lambda wildcards, attempt: 50 + 20*(attempt-1)
    priority: 97
    params:
        fagz = outdir + "{genome}.e" + str(cth) + ".fa.gz",
        copts = copts,
        sd = srcdir,
        od = outdir,
        pd = pdir,
        ksize = ksize,
        cth = cth,
        rth = rth,
        rstring = rstring,
        thcth = thcth,
    shell:"""
set -eu
ulimit -c 20000
cd {params.od}

zcat {params.fagz} |
{params.sd}/bin/danbing-tk -gc {params.thcth} -k {params.ksize} -qs {params.od}/pan -fai /dev/stdin -o pan.{wildcards.genome}.IL -p {resources.cores} -cth {params.cth} -rth {params.rth}
touch {output.ofoo}
"""


rule EvalGenotypeQuality:
    input:
        #mapping = pdir + "OrthoMap.v2.tsv",
        ifoo = outdir + "checkpoint/{genome}.pan.gt.foo"
        #panILkmers = outdir + "pan.{genome}.IL.tr.kmers",
        #PBkmers = pdir + "{genome}.rawPB.tr.kmers",
    output:
        #mappedILkmers = outdir + "{genome}.mappedIL.tr.kmers",
        #pred = outdir + "{genome}.rawLR.pred",
        ofoo = outdir + "checkpoint/{genome}.rawLR.foo",
    resources:
        cores = 4,
        mem = lambda wildcards, attempt: 8*attempt,
    priority: 96
    params:
        copts = copts,
        sd = srcdir,
        od = outdir,
        pd = pdir,
    shell:"""
set -eu
ulimit -c 20000
cd {params.od}

{params.sd}/script/kmers.linreg.py --mapkmer --mode invalid --R2threshold -2 {params.pd}/{wildcards.genome}.rawPB.tr.kmers {params.od}/pan.{wildcards.genome}.IL.tr.kmers {wildcards.genome}.rawLR
touch {output.ofoo}
"""


rule PredictLength:
    input:
        mapping = outdir + "OrthoMap.v2.tsv",
        LOOILkmers = expand(outdir + "LOO.{genome}.IL.tr.kmers", genome=LOOgenomes),
        panILkmers = expand(outdir + "pan.{genome}.IL.tr.kmers", genome=genomes),
        pred = expand(outdir + "{genome}.LR.pred", genome=genomes),
    output:
        pickle = outdir + "analysis/step1_results.pickle",
        relErr = outdir + "analysis/rel_err.txt",
    resources:
        cores = 2,
        mem = 8,
    priority: 95
    params:
        copts = copts,
        sd = srcdir,
        od = outdir,
        pd = pdir,
        gf = genomes,
        covbed = covbed,
        LC = config["LOOconf"],
        SC = config["sampleConf"],
        badg = f'--badg {config["badgenome"]}' if config["badgenome"] else "",
    shell:"""
set -eu
ulimit -c 20000
mkdir -p {params.pd}/analysis
mkdir -p {params.od}/analysis
cd {params.od}

nloci=$(cat {input.mapping} | wc -l)
{params.sd}/script/kmc2length.py --genome {params.gf} --nloci $nloci --panmap {params.pd}/OrthoMap.v2.tsv \
								 --cov {params.pd}/ctrl.cov --covbed {params.covbed} \
                                 --LOOconf {params.LC} --sampleConf {params.SC} {params.badg}

{params.sd}/script/kmc2length.py --genome {params.gf} --nloci $nloci --panmap {params.pd}/OrthoMap.v2.tsv \
								 --cov {params.pd}/ctrl.cov --covbed {params.covbed} \
                                 --LOOconf {params.LC} --sampleConf {params.SC} {params.badg}

"""
```
