## Supplemental code for "The motif composition of variable-number tandem repeats impacts gene expression": RefGraph.snakefile

```
import os
import numpy as np

configfile: "refGraph.json"

srcdir = config["srcDir"]
indir = config["inputDir"]
outdir = config["outputDir"]
pdir = config["pangenomeDir"]

genomefile = config["genomefile"]
genomes = np.loadtxt(genomefile, dtype=object).reshape(-1).tolist()
ref = config["ref"]
refTR = config["refTR"]
refsize = config["refsize"]
kmerTypes = ["tr", "ntr", "graph"]

ksize = config["ksize"]
FS = config["flankSize"]
cth = config["countThreashold"]
rth = config["ratioThreashold"]
rstring = f'{rth*100:.0f}'
thcth = config["threadingCountThreshold"]
LB = config["sizeLowerBound"]
TRwindow = config["TRwindow"]
mbe_th1 = float(config["MBE_th1"])
mbe_th2 = float(config["MBE_th2"])
copts = config["clusterOpts"]


localrules: all, GenMap_v0_v2

rule all:
    input:
        MBEfoo = outdir + "MBE.foo",
        refkmers = expand(outdir + "hg38.{kmerType}.kmers", kmerType=kmerTypes),
        #TRfa = outdir + "ref.tr.fasta",
        #TRbed = outdir + "ref.bed",
        #panbed = outdir + "pan.tr.bed",
        #mapping = outdir + "refMap.tbl",
        #pankmers = expand(outdir + "pan.{kmerType}.kmers", kmerType=kmerTypes),
        #panILkmers = expand(outdir + "pan.{genome}.IL.tr.kmers", genome=genomes),
        #pred = expand(outdir + "{genome}.LR.pred", genome=genomes),
        

rule JointTRAnnotation:
    input:
        fa = ref,
    output:
        foo = outdir + "MBE.foo",
        TRfa = outdir + "hg38.tr.fasta",
        #mapping = outdir + "OrthoMap.v2.tsv",
        #TRfa = outdir + "ref.tr.fasta",
        #TRbed = outdir + "ref.bed",
    resources:
        cores = 6,
        mem = lambda wildcards, attempt: 40 + 20*(attempt-1)
    priority: 96
    params:
        copts = copts,
        sd = srcdir,
        od = outdir,
        refTR = refTR,
        ksize = ksize,
        FS = FS,
        LB = LB,
        TRwindow = TRwindow,
        th1 = mbe_th1,
        th2 = mbe_th2,
        #gf = genomefile,
        #genomes = genomes
    shell:"""
set -eu
ulimit -c 20000
cd {params.od}

g=hg38
ln -sf {input.fa} $g.0.fa
ln -sf {input.fa}.fai $g.0.fa.fai
ln -sf $g.0.fa $g.1.fa
ln -sf $g.0.fa.fai $g.1.fa.fai
mkdir -p $g
awk 'BEGIN {{OFS="\t"}} {{print $0, ".", ".", ".", "+"}}' {params.refTR} >$g/tmp1.0.bed
cp $g/tmp1.0.bed $g/tmp1.1.bed

echo "Generating panbed"
awk 'BEGIN {{OFS="\t"}} {{print $1, $2, $3, 1}}' {params.refTR} >pan.tr.mbe.v0.bed
{params.sd}/script/preMBE.py <(echo "hg38") pan.tr.mbe.v0.bed
{params.sd}/script/multiBoundaryExpansion.py
{params.sd}/script/writeMBEbed.py {params.th1} {params.th2}
hi=0
for h in 0 1; do
    echo ">""$g"".""$h"
    cut -f $((4+4*hi))-$((6+4*hi)) pan.tr.mbe.v1.bed |
    awk 'BEGIN {{OFS="\t"}} {{print $0, NR-1}}' |
    grep -v "None" |
    sort -k1,1 -k2,2n -k3,3n >tmp.bed
    if [[ "$(cat tmp.bed | wc -l)" != "0" ]]; then
        bedtools merge -d 700 -c 4 -o collapse -i tmp.bed |
        cut -f 4 | grep ","
    fi
    ((++hi))
done >mbe.m0.loci
rm tmp.bed
{params.sd}/script/mergeMBEbed.py

### write fasta
echo "Fetching TR+flank"
h=0
cut -f 4-6 pan.tr.mbe.v2.bed |
grep -v "None" |
awk 'BEGIN {{OFS="\t"}} {{
    $2=$2-700
    $3=$3+700
    print $0
}}' |
{params.sd}/script/SelectRegions.py /dev/stdin "$g"."$h".fa /dev/stdout |
awk '{{if ($1 ~ />/) {{print}} else {{print toupper($0)}} }}' >"$g".tr.fasta
rm hg38.?.fa*
touch MBE.foo
"""


rule GenMap_v0_v2:
    input:
        foo = outdir + "MBE.foo",
    output:
        mapping = outdir + "locusMap.v0.to.v2.txt",
    resources:
        cores = 1,
        mem = lambda wildcards, attempt: 4,
    params:
        copts = copts,
        sd = srcdir,
        od = outdir,
        ksize = ksize,
        FS = FS,
        graph = "hg38"
    run:
        import numpy as np
        nloci = np.loadtxt(f"{params.od}/pan.tr.mbe.v0.bed", usecols=1).size
        m21 = np.loadtxt(f"{params.od}/locusMap.v2.to.v1.txt", dtype=int)
        m10 = np.loadtxt(f"{params.od}/locusMap.v1.to.v0.txt", dtype=int)
        m02 = np.full(nloci, ".", dtype=object)
        m02[m10[m21]] = np.arange(m21.size)
        np.savetxt(f"{params.od}/locusMap.v0.to.v2.txt", m02, fmt='%s')    


rule GenRefGraph:
    input:
        TRfa = outdir + "hg38.tr.fasta",
        mapping = outdir + "locusMap.v0.to.v2.txt",
    output:
        refkmers = expand(outdir + "hg38.{kmerType}.kmers", kmerType=kmerTypes),
    resources:
        cores = 1,
        mem = lambda wildcards, attempt: 8*attempt,
    params:
        copts = copts,
        sd = srcdir,
        od = outdir,
        ksize = ksize,
        FS = FS,
        graph = "hg38"
    shell:"""
set -eu
ulimit -c 20000
cd {params.od}


{params.sd}/bin/vntr2kmers_thread -g -m {input.mapping} -k {params.ksize} -fs {params.FS} -ntr {params.FS} -o {params.graph} -fa 1 {input.TRfa}
"""


rule GenPanGraph:
    input:
        #TRbed = outdir + "hg38.bed",
        refkmers = expand(outdir + "hg38.{kmerType}.kmers", kmerType=kmerTypes),
    output:
        panbed = outdir + "pan.tr.bed",
        mapping = outdir + "refMap.tbl",
        pankmers = expand(outdir + "pan.{kmerType}.kmers", kmerType=kmerTypes),
    resources:
        cores = 1,
        mem = lambda wildcards, attempt: 8*attempt,
    params:
        copts = copts,
        sd = srcdir,
        od = outdir,
        refTR = config["refTR"],
        kmerpref = f'{outdir}/ref',
    shell:"""
cd {params.od}
ulimit -c 20000

awk 'BEGIN {{OFS="\t"}} {{$4=$4"\t"(NR-1); print $0}}' {params.refTR} > {output.panbed}
bedtools map -c 4 -o collapse -a {output.panbed} -b <(awk 'BEGIN {{OFS="\t"}} {{print $4, $5, $6, NR-1}}' {input.TRbed}) > {output.panbed}.tmp
mv {output.panbed}.tmp {output.panbed}

cut -f 6- {output.panbed} > {output.mapping}

{params.sd}/bin/genPanKmers -o pan -m {output.mapping} -k {params.kmerpref}
"""


rule GenotypeSamples:
    input:
        pankmers = expand(outdir + "pan.{kmerType}.kmers", kmerType=kmerTypes),
        ILbam = indir + "{genome}.IL.srt.bam",
        ILbai = indir + "{genome}.IL.srt.bam.bai",
    output:
        ILkmers = outdir + "pan.{genome}.IL.tr.kmers",
    resources:
        cores = 24,
        mem = 40,
    params:
        copts = copts,
        sd = srcdir,
        od = outdir,
        ksize = ksize,
        FS = FS,
        cth = cth,
        rth = rth,
        rstring = rstring,
        thcth = thcth,
        graph = "pan",
    shell:"""
set -eu
ulimit -c 20000
cd {params.od}

samtools fasta -@2 -n {input.ILbam} |
awk '{{if (substr($1,1,1) == ">") {{
        if (substr($1,length($1)-1,1) == "/") {{ print substr($1, 1, length($1)-2) }} else {{ print $1 }} }}
      else {{ print $1 }}
     }}' |
{params.sd}/bin/bam2pe -k {params.ksize} -fai /dev/stdin |
{params.sd}/bin/aQueryFasta_thread -g {params.thcth} -k {params.ksize} -qs {params.od}/{params.graph} -fai /dev/stdin -o {params.graph}.{wildcards.genome}.IL -p {resources.cores} -cth {params.cth} -rth {params.rth}
"""


rule EvalRefGraph:
    input:
        mapping = outdir + "locusMap.tbl",
        PBkmers = pdir + "{genome}.PB.tr.kmers",
        panILkmers = outdir + "pan.{genome}.IL.tr.kmers"
    output:
        PBkmers = indir + "{genome}.PB.tr.kmers",
        mappedILkmers = outdir + "{genome}.mappedIL.tr.kmers",
        pred = outdir + "{genome}.LR.pred",
    resources:
        cores = 12,
        mem = 8,
    params:
        copts = copts,
        sd = srcdir,
        od = outdir,
        indir = indir,
        gi = lambda wildcards: genomes.index(wildcards.genome),
    shell:"""
set -eu
ulimit -c 20000

cd {params.indir}
ln -s {input.PBkmers} .

cd {params.od}
{params.sd}/bin/mapkmers  {input.mapping}  {params.gi}  {input.panILkmers}  {input.PBkmers}  {wildcards.genome}.mappedIL.tr
{params.sd}/script/kmers.linreg.py --mode invalid --R2threshold -2 {input.PBkmers} {output.mappedILkmers} {wildcards.genome}.LR
"""
```
