## Supplemental code for "The motif composition of variable-number tandem repeats impacts gene expression": README.md

# danbing-tk
A toolkit for variable number tandem repeats (VNTRs) analysis, which enables:
1. building repeat-pan genome graphs (RPGG) for a set of "genotypable" VNTRs given haplotype-resolved assemblies and matched short-read seqeuncing (SRS) data (`danbing-tk build`),
2. genotyping each VNTR as a set of (*k*-mer, count) given SRS data (`danbing-tk align`), and
3. predicting VNTR length from the genotype (`danbing-tk predict`) given locus-specific sampling bias (LSB).

See [online manuscript](https://www.nature.com/articles/s41467-021-24378-0) for details.


## Download Releases
Each release comes with the lastest version of genotypable VNTRs, RPGG (and/or) precomputed LSB.

|                   | File                                                  | Input of           | Output of        |
|-------------------|-------------------------------------------------------|--------------------|------------------|
| Genotypable VNTRs | tr.good.bed                                           | danbing-tk build   |                  |
| Control regions   | ctrl.bed                                              | danbing-tk build   |                  |
| RPGG              | pan.(tr.kmers\|kmerDBi.umap\|kmerDBi.vv\|graph.umap)  | danbing-tk align   | danbing-tk build |
| Precomputed LSB   | LSB.tsv                                               | danbing-tk predict |                  |

- Release v1.3: Updated RPGG built from 35 HGSVC genomes.
- Release v1.0: VNTR summary statistics and eGene discoveries are also included. Example analyses such as differential length/motif analysis, eQTL mapping, VNTR locus QC, sample QC are also included.


## Building on Linux
```shell
git clone https://github.com/ChaissonLab/danbing-tk
cd danbing-tk && make
```

## danbing-tk align
Decompress the RPGG `RPGG.tar.gz` in your working directory.

An example usage to genotype SRS sample using the RPGG:

```shell
samtools fasta -@2 -n $SRS.bam |
/$PREFIX/danbing-tk/bin/danbing-tk -gc 80 -ae -kf 4 1 -cth 45 -o $OUT_PREF -k 21 -qs pan -fai /dev/stdin -p $THREADS | gzip >$OUT_PREF.aln.gz
```

`danbing-tk align` takes ~12 cpu hours to genotype a 30x SRS sample. This will generate `$OUT_PREF.tr.kmers` and `$OUT_PREF.aln.gz` output with format specified in [File Format](#file-format).

**Important note:** If outputs of `danbing-tk align` are intended to be used directly for downstream analyses e.g. association tests, please check the [distribution of LSB](#distribution-of-lsb) section below before running.


## danbing-tk build
### Install Dependencies
For users intended to use `danbing-tk align` only, this step is not required.

The `danbing-tk build` pipeline and `danbing-tk predict` require several external packages. It is recommended to install all requirements using conda as follows:

```bash
conda install -c conda-forge -c bioconda -c intel \
    snakemake=5.11.2 samtools=1.10 bedtools=2.29.2 minimap2=2.17 \
    scikit-learn=0.23.1 statsmodels=0.12.1 pysam=0.15.3
```

If the requirements are in conflict with existing packages, create a new environment specifically for danbing-tk with:
```
conda create -n $MY_ENVIRONMENT -c conda-forge -c bioconda -c intel \
    snakemake=5.11.2 samtools=1.10 bedtools=2.29.2 minimap2=2.17 \
    scikit-learn=0.23.1 statsmodels=0.12.1 pysam=0.15.3
```

To check if everything is configured properly:
1. Go to `/$PREFIX/danbing-tk/test/`
2. Replace `$PREFIX` in `goodPanGenomeGraph.json` and `input/genome.bam.tsv` with the path to danbing-tk
3. Run `snakemake -p -s ../pipeline/GoodPanGenomeGraph.snakefile -j 4 --rerun-incomplete --output-wait 3`

Tested on v1.0. 


### Running danbing-tk build
- Required inputs: 
	- haplotype-resolved assemblies (FASTA)
	- matched SRS data (BAM; optional)
	- reference genome (major chromosomes only without minor contigs)
	- tandem repeat regions (BED; available on [release page](https://github.com/ChaissonLab/danbing-tk/releases/tag/v1.0) or user-defined)

- Copy `/$PREFIX/danbing-tk/pipeline/goodPanGenomeGraph.json` to your working directory and edit accordingly. 

- A config file `/$INPUT_DIR/genome.bam.tsv` with two columns, one for genome name and one for bam file path, is required if SRS data is available for graph pruning, e.g.

    ```HG00514 /panfs/qcb-panasas/tsungyul/HG00514/HG00514.IL.srt.bam```
    
    Otherwise, set `pruning` in `goodPanGenomeGraph.json` to `False` and use a single column input for `genome.bam.tsv`.

- Run the snakemake pipline with:
```bash
snakemake -p -s /$PREFIX/danbing-tk/pipeline/GoodPanGenomeGraph.snakefile -j 40\
    --cluster "{params.copts} -c {resources.cores} --mem={resources.mem}G -k" \
    --rerun-incomplete --restart-times 1 --output-wait 30
```

Submitting jobs to cluster is preferred as `danbing-tk build` is compute-intensive, ~1200 cpu hours for the original dataset. Otherwise, remove `--cluster` and its parameters to run jobs locally.

## danbing-tk predict
Locus-specific sampling biases (LSB) at VNTR regions are critical for normalizing the sum of *k*-mer counts to VNTR length. We provided precomputed LSB at the VNTR regions for fast comparison, however this assumes the LSB of the dataset of interest is close enough to the dataset in the original paper. Please ensure this assumption is valid by running a joint PCA on the LSB of non-repetitive regions with the original dataset, provided in `LSB.tsv`. If this assumption failed, leave-one-out analysis (next section) on the dataset of interest is necessary to make accurate predictions. The following usage is for when the assumption holds.

Run `getCovByLocus.397.sh` on your SRS dataset.

Run length prediction with:

```bash
/$PREFIX/danbing-tk/script/kmc2length.py --outdir OUTDIR --ksize KSIZE --kmers KMERS --trbed
	TRBED --LSB LSB --cov COV --covbed COVBED
```

Length estimates are written to `estimated_TR_len.tsv`.

## Analysis
### Distribution of LSB
To ensure technical variations are consistent between samples and do not introduce bias to VNTR genotypes, or *k*-mer counts, in a subset of samples. It is necessary to ensure the distribution of LSB shows up in one cluster. An example analysis is shown in `LSB_analysis.ipynb` on the [release page](https://github.com/ChaissonLab/danbing-tk/releases/tag/v1.0) and `script/getCovByLocus.397.sh`.

### Leave-one-out analysis
To evaluate the quality of custom RPGG on matching SRS dataset, copy `/$PREFIX/danbing-tk/pipeline/leaveOneOut.snakefile` to your working directory and edit accordingly.
Run the snakemake pipleine with:

```bash
snakemake -p -s /$PREFIX/pipeline/LeaveOneOut.snakefile -j 40 --cluster \
    "{params.copts} -c {resources.cores} --mem={resources.mem}G -k" \
    --rerun-incomplete --restart-times 1 --output-wait 30
```

Submitting jobs to cluster is preferred as `danbing-tk build` is compute-intensive; otherwise, remove `--cluster` and its parameters to run jobs locally.


## File Format
- \*.graph.kmers
```
>locus i
kmer0	out_edges0
kmer1	out_edges1
...
>locus i+1
...
```
`out_edges` denotes the presence of T/G/C/A as the next nucleotide encoded with 4 bits.

- \*.(tr|ntr).kmers
```
>locus i
kmer0	kmer_count0
kmer1	kmer_count1
...
>locus i+1
...
```
The second field is optional.

- Alignment output (`-a` option)
	- Synopsis
		```
		<src> <dest> <read_name/0> <read_seq/0> <ops/0> <read_name/1> <read_seq/1> <ops/1>
		```
	- `src`: source locus of a read pair (for simulation only)
	- `dest`: aligned locus for the read pair
	- `ops`: operations to align the read to the graph
		- `=`: a match in the repeat
		- `.`: a match in the flank
		- `[A|C|G|T]`: a mismatch; letter in the graph is shown
		- `[0|1|2|3]`: a deletion; letter in the graph is shown as 0123 for ACGT, respectively.
		- `I`: an insertion in the read
		- `H`: a nucleotide in the homopolymer run
		- `*`: unalinged nucleotide
