## Supplemental code for "The motif composition of variable-number tandem repeats impacts gene expression": chrsize.sh

```
#!/usr/bin/env bash

awk 'BEGIN {OFS="\t"}
{
    if ($1 ~ />/) {
        if (size) {print name, size}
        name = substr($1,2,length($1)-1); size=0
    }
    else { size += length($1) }
}
END {
    print name, size
}' $1 | sort -k1,1
```
