## Supplemental code for "The motif composition of variable-number tandem repeats impacts gene expression": eqtl.noPerm.py

```
#! /usr/bin/env python3

import argparse 
import glob
import numpy as np
import matplotlib.pyplot as plt
import pickle
from sklearn.preprocessing import quantile_transform as qt
import statsmodels.api as sm
from statsmodels.stats.multitest import fdrcorrection as fdr
from scipy import stats

np.set_printoptions(precision=2, suppress=True, edgeitems=100)


# Gene-TR mapping relation
def getSize(f):
    bed = np.loadtxt(f, usecols=[1,2], dtype=int)
    return bed[:,1] - bed[:,0]


def getLociList():
    lociList = np.loadtxt(args.TRbed, dtype=object, usecols=[0,1,2])
    loci2ind = {}
    for ind, row in enumerate(lociList):
        loci2ind["_".join(row)] = ind
    return lociList, loci2ind


def indexGeneList(tissue):
    tisGeneList = np.loadtxt(f'{args.expDir}/{tissue}.v8.normalized_expression.bed.gz', dtype=object, skiprows=1, usecols=[3])      
    tisGene2ind = {}
    for ind, gene in enumerate(tisGeneList):
        tisGene2ind[gene] = ind        
    return tisGeneList, tisGene2ind


def getLocusi2tisGenei(tisGene2ind):
    locusi2tisGenei = {}
    ncomb = 0
    for row in TRxGene:
        locusname = "_".join(row[:-1])
        locusi = loci2ind[locusname]
        if row[-1] in tisGene2ind:
            if locusi not in locusi2tisGenei:
                locusi2tisGenei[locusi] = []
            locusi2tisGenei[locusi].append(tisGene2ind[row[-1]])
            ncomb += 1
    print(f'\t{len(locusi2tisGenei)} TRs')
    print(f'\t{ncomb} TR x Gene tests')
    return locusi2tisGenei


def getGenei2nloci(locusi2tisGenei):
    genei2nloci = {}
    for locusi, geneindices in locusi2tisGenei.items():
        for genei in geneindices:
            if genei not in genei2nloci:
                genei2nloci[genei] = 0
            genei2nloci[genei] += 1
    return genei2nloci


# expression matrix
def loadSNPPCinfo():
    if args.SNPPC is None:
        return None, None

    ndim = 838 # XXX
    tmp = np.loadtxt(args.SNPPC, usecols=np.arange(11), dtype=object)[:ndim] # XXX
    SNP_PCs = tmp[:,1:].astype(float)
    SNP_sampleList = [s.split("-")[-1] for s in tmp[:,0]]
    return SNP_PCs, SNP_sampleList


def getTisSNPResTpmMat(tissue, SNP_PCs, SNP_sampleList):
    # SNP PCs
    tmp = np.loadtxt(f'{args.expDir}/{tissue}.v8.normalized_expression.bed.gz', dtype=object, max_rows=1, comments="!")[4:]    
    tisSampleList = np.array([s[5:] for s in tmp])

    snpSample2ind = {}
    for sind, sample in enumerate(SNP_sampleList):
        snpSample2ind[sample] = sind

    sampleMap_tis2snp = np.zeros(tisSampleList.size, dtype=int)
    for ind in range(tisSampleList.size):
        sampleMap_tis2snp[ind] = snpSample2ind[tisSampleList[ind]]

    tisSNP_PCs = SNP_PCs[sampleMap_tis2snp]
    
    # GTEx PCs
    gtexPCs = np.loadtxt(f'{args.covDir}/{tissue}.v8.covariates.txt', dtype=object, skiprows=1)[:,1:].astype(float).T   
    
    C = np.hstack((gtexPCs, tisSNP_PCs))
    tisTpmMat = np.loadtxt(f'{args.expDir}/{tissue}.v8.normalized_expression.bed.gz', dtype=object, skiprows=1)[:,4:].astype(float).T        
    tisResTpmMat = (1 - C @ np.linalg.inv(C.T @ C) @ C.T) @ tisTpmMat
    return tisResTpmMat.T


# genotype matrix
def getGenotypeMat():
    genMat = np.zeros([nloci, nwgs], dtype=float)
    kmerfnames = glob.glob(f'{args.genDir}/*.tr.kmers')
    for fi, fname in enumerate(kmerfnames):
        print(".", end='', flush=True)
        if fi % 100 == 99: print("")
        with open(fname) as f:
            locusi = -1 # XXX was -14
            kms = 0
            for line in f:
                if line[0] == ">":
                    if locusi >= 0:
                        genMat[locusi, fi] = kms
                        kms = 0
                    locusi += 1
                else:
                    kms += int(line.split()[1])
            else:
                genMat[locusi, fi] = kms
    print("done reading genotypes", flush=True)
    return genMat


def processBamCov(bamcovmat, mth=1.2, sth=0.1):
    ctrlsize = getSize(args.ctrlbed)
    badmask = np.zeros_like(ctrlsize, dtype=bool)

    ### compute coverage for each locus; normalize wrt sample global coverage
    pnormcovmat = bamcovmat / (bamcovmat@ctrlsize / np.sum(ctrlsize))[:,None]

    ### check variance
    stds = np.std(pnormcovmat, axis=0)
    normstds = stds
    badmask = np.logical_or(badmask, normstds > sth)

    ### check if mean is biased
    mnormcov = np.mean(pnormcovmat, axis=0)
    badmask = np.logical_or(badmask, mnormcov > mth)    

    ### reject outliers
    pctrlsize = ctrlsize[~badmask]
    pcovmat = bamcovmat[:,~badmask]
    return pcovmat@pctrlsize / np.sum(pctrlsize)


def correctGenMat():
    gtexSex = np.loadtxt(args.phenotype, dtype=object, usecols=[0,1])[1:]    
    sample2sex = {}
    for i in range(gtexSex.shape[0]):
        sample = gtexSex[i,0].split("-")[1]
        sample2sex[sample] = int(gtexSex[i,1])
    print(len(sample2sex))
    print(genMat.shape)
    
    wgsSex = np.zeros_like(genomes, dtype=int)
    for ind, g in enumerate(genomes):
        wgsSex[ind] = sample2sex[g]
        
    covmat = np.loadtxt(f'{args.outDir}/ctrl.cov', dtype=object)
    gcov = processBamCov(covmat[:,2:].astype(float))
    
    normGenMat = genMat / gcov
    normGenMat[:args.NL1] /= 2
    normGenMat[args.NL1:args.NL2] /= wgsSex
    print(normGenMat.shape)
    return normGenMat


def getTissueGenMat(tissue):
    genoSample2ind = {}
    for ind, sample in enumerate(genomes):
        genoSample2ind[sample] = ind
        
    tmp = np.loadtxt(f'{args.expDir}/{tissue}.v8.normalized_expression.bed.gz', dtype=object, max_rows=1, comments="!")[4:]
    tisSampleList = np.array([s[5:] for s in tmp])
    sampleMap_tis2geno = np.zeros(tisSampleList.shape[0], dtype=int)
    for ind, sample in enumerate(tisSampleList):
        sampleMap_tis2geno[ind] = genoSample2ind[sample]
        
    return genMat[:,sampleMap_tis2geno]


# eQTL mapping
def runRegressionZ3(tisResTpmMat, tisGenMat, locusi2tisGenei, genei2nloci):
    outs = {}
    Y_zscore = (tisResTpmMat - np.mean(tisResTpmMat, axis=1)[:,None]) / np.std(tisResTpmMat, axis=1)[:,None]
    X_zscore = (tisGenMat - np.mean(tisGenMat, axis=1)[:,None]) / np.std(tisGenMat, axis=1)[:,None]
    for locusi, geneindices in locusi2tisGenei.items():
        for genei in geneindices:
            exps = Y_zscore[genei]
            genos = X_zscore[locusi]
            if not np.all(np.isfinite(exps)) or not np.all(np.isfinite(genos)):
                continue
            results = sm.OLS(exps, sm.add_constant(genos, prepend=True)).fit()
            p = results.pvalues[1] * genei2nloci[genei] # Bonferroni correction
            b = results.params[1]
            bse = results.bse[1]
            if genei in outs:
                if p < outs[genei][0]:
                    outs[genei] = (p, b, bse, locusi)
            else:
                outs[genei] = (p, b, bse, locusi)
                
    tiseGeneTR = np.array([[genei, v[-1]] for genei, v in outs.items()], dtype=int)
    stats = np.array([[*v[:-1]] for v in outs.values()])
    return tiseGeneTR, stats


def annotateGeneTR1(tissue, tisGeneList, genei2nloci, tiseGeneTR, stats, rejected, adjP):
    tiseGene = tisGeneList[tiseGeneTR[:,0]]
    numVar = np.zeros_like(tiseGene, dtype=int)
    for ind, genei in enumerate(tiseGeneTR[:,0]):
        numVar[ind] = genei2nloci[genei]
    tiseTR = tiseGeneTR[:,1]
    return np.hstack((tiseGene[:,None], numVar[:,None], tiseTR[:,None], stats, adjP[:,None]))[rejected]


def singleTissue_eGene_stat(tissue, SNP_PCs, SNP_sampleList):
    # establish mapping between TRs and Genes
    tisGeneList, tisGene2ind = indexGeneList(tissue)
    locusi2tisGenei = getLocusi2tisGenei(tisGene2ind)

    tisGenMat = getTissueGenMat(tissue) # genMat with samples missing in tpmMat removed
    print(f'\ttisGenMat {tisGenMat.shape}')

    if glob.glob(f'{args.resDir}/{tissue}.ResMat.pickle'):
        tisResTpmMat = pickle.load(open(f'{args.resDir}/{tissue}.ResMat.pickle', 'rb'))
    else:
        tisResTpmMat = getTisSNPResTpmMat(tissue, SNP_PCs, SNP_sampleList)
        pickle.dump(tisResTpmMat, open(f'{args.outDir}/{tissue}.ResMat.pickle', 'wb'))
    print(f'\ttisResTpmMat {tisResTpmMat.shape}')

    genei2nloci = getGenei2nloci(locusi2tisGenei) # count # of TRs mapped to each gene; used for Bonferroni correction

    tiseGeneTR, stats = runRegressionZ3(tisResTpmMat, tisGenMat, locusi2tisGenei, genei2nloci) # [genei, locusi], [p, b, bse]
    print(f'\t{tiseGeneTR.shape[0]} genes tested')

    rejected, adjP = fdr(stats[:,0]) 
    print(f'\t{np.sum(rejected)} tissue eGenes')

    eGeneStat = annotateGeneTR1(tissue, tisGeneList, genei2nloci, tiseGeneTR, stats, rejected, adjP)
    print(f'\t{eGeneStat.shape[0]} total eGenes')

    return eGeneStat


def writeAlleGeneTR():
    allGeneInfo = np.loadtxt(args.geneBed, dtype=object)[:,[3,4,0,1,2]]
    allGeneInfo[:,3:] = allGeneInfo[:,3:].astype(int)
    
    gene2allind = {}
    for i in range(allGeneInfo.shape[0]):
        gene2allind[allGeneInfo[i,0]] = i
        
    allTRinfo = np.loadtxt(args.TRbed, dtype=object, usecols=[0,1,2])
    allTRinfo[:,1:] = allTRinfo[:,1:].astype(int)
    
    SNP_PCs, SNP_sampleList = loadSNPPCinfo()
    tissues = np.loadtxt(args.tissues, dtype=object)
    
    for tissue in tissues:
        print("tissue: {}".format(tissue))
        
        out_ = singleTissue_eGene_stat(tissue, SNP_PCs, SNP_sampleList)
        N = out_.shape[0]
        
        outGeneIndices = np.zeros(N, dtype=int)
        for i in range(N):
            outGeneIndices[i] = gene2allind[out_[i,0]]
        outGeneInfo = allGeneInfo[outGeneIndices,1:]
        
        outTRinfo = allTRinfo[out_[:,2].astype(int)]
        
        out = np.hstack((out_[:,0:1], outGeneInfo, out_[:,1:2], outTRinfo, out_[:,2:]))
        out = out[np.argsort(out_[:,2])]
        np.savetxt(f'{args.outDir}/{tissue}.v8.egenes.txt', out, delimiter="\t", 
                   header="gene_id\tgene_name\tchr\tstart\tend\tnum_var\tTR_chr\tTR_start\tTR_end\tTR_locus\tpval_nominal\tslope\tslope_se\tqval",
                   fmt=['%s','%s','%s','%i','%i','%i','%s','%i','%i','%i','%.4e','%.4e','%.4e','%.4e']) # XXX pval_nominal, pval_adjusted


if __name__ == "__main__":
    
    ap = argparse.ArgumentParser(description=\
            "Run single tissue eQTL mapping; output egenes.txt and pairs.txt")

    ap.add_argument("--TRbed", help="bed file of TR regions", required=True) # /home/cmb-17/mjc/vntr_genotyping/goodPanGenomeGraph/input/tr.good.bed
    ap.add_argument("--geneBed", help="bed file of (gene regions, name, id)", required=True) # /home/cmb-17/mjc/vntr_genotyping/gtex/eqtl/input/genes_id_name.bed
    ap.add_argument("--pair", help="bed file of (gene,TR) pairs", required=True) # /home/cmb-17/mjc/vntr_genotyping/gtex/eqtl/gene.100k_window.tr.pair.bed
    ap.add_argument("--expDir", help="dir to expression matrices", required=True) # /home/cmb-17/mjc/vntr_genotyping/gtex/eqtl/input/GTEx_Analysis_v8_eQTL_expression_matrices
    ap.add_argument("--resDir", help="dir to residualized expression matrix pickles") # /home/cmb-17/mjc/vntr_genotyping/gtex/eqtl/ResMat/
    ap.add_argument("--genDir", help="dir to TR genotypes", required=True) # /home/cmb-17/mjc/vntr_genotyping/goodPanGenomeGraph/eqtl/genotype/
    ap.add_argument("--covDir", help="dir to GTEx covariates", required=True) # /home/cmb-17/mjc/vntr_genotyping/gtex/eqtl/input/GTEx_Analysis_v8_eQTL_covariates/
    ap.add_argument("--outDir", help="dir to output", required=True)
    ap.add_argument("--phenotype", help="GTEx phenotype annotations", required=True) # /home/cmb-17/mjc/vntr_genotyping/gtex/eqtl/input/GTEx_Analysis_v8_Annotations_SubjectPhenotypesDS.txt
    ap.add_argument("--genomes", help="file listing GTEx genomes", required=True) # /home/cmb-17/mjc/vntr_genotyping/gtex/genotype/genomes.txt
    ap.add_argument("--tissues", help="file listing GTEx tissues", required=True) # /home/cmb-17/mjc/vntr_genotyping/gtex/eqtl/input/GTEx_Analysis_v8_eQTL_expression_matrices/alltissue.txt
    ap.add_argument("--genMat", help="pickle of genotype matrix") # /home/cmb-17/mjc/vntr_genotyping/gtex/genotype/normedGenotype.29111x879.pickle.dat
    ap.add_argument("--NL1", help="number of TR loci in autosomes", required=True, type=int) # 28101
    ap.add_argument("--NL2", help="number of TR loci in autosomes plus X chr", required=True, type=int) # 29052
    ap.add_argument("--SNPPC", help="principal components of (GTEx,1KGP) SNP matrix") # /home/cmb-17/mjc/vntr_genotyping/gtex/variation/joint.pca.evec
    ap.add_argument("--ctrlbed", help="bed file of unique regions") # /home/cmb-17/mjc/vntr_genotyping/cmb-16/work/vntr/hapdb/a1_regions/ctrl/pan.fn0.bed.bak


    args = ap.parse_args()
    
    genomes = np.loadtxt(args.genomes, dtype=object)
    nwgs = genomes.size
    nloci = np.loadtxt(args.TRbed, usecols=[1]).size

    lociList, loci2ind = getLociList()

    TRxGene = np.loadtxt(args.pair, dtype=object, usecols=[5,6,7,3]) # XXX regenerate
            
    if args.genMat:
        print("reading genotype pickle")
        genMat = pickle.load(open(args.genMat, 'rb'))
    else:
        if glob.glob(f'{args.outDir}/rawGenotype.pickle'):
            genMat = pickle.load(open(f'{args.outDir}/rawGenotype.pickle', 'rb'))
        else:
            print("reading raw genotypes")
            genMat = getGenotypeMat()
            pickle.dump(genMat, open(f'{args.outDir}/rawGenotype.pickle', 'wb'))

        print("correcting genotypes")
        genMat = correctGenMat()
        pickle.dump(genMat, open(f'{args.outDir}/normedGenotype.pickle', 'wb'))

    print("starting eQTL mapping")
    writeAlleGeneTR()
```
