## Supplemental code for "The motif composition of variable-number tandem repeats impacts gene expression": fixMaskedFasta.py

```
#!/usr/bin/env python3

# the original file has missing empty loci and unexpeted short loci (< 4200 bp)

import argparse
import csv 
ap = argparse.ArgumentParser(description="fix missing titles and splitted lines in a fasta file")
ap.add_argument("th", help="length threshold for each locus", type=int)
ap.add_argument("fi", help="suffix of corrupted file e.g. combined-hap.fasta.masked for HG00514.h0.combined-hap.fasta.masked")
ap.add_argument("--fixTitle", help="fix missing titles as well, otherwise will fix splitted lines only", action='store_true')
ap.add_argument("--useCSV", help="use CSV file to configure haps", action='store_true')
ap.add_argument("--haps", help="specify hap to process")
args = ap.parse_args()
print(args)
th = args.th

fixInv = False # flag for fixing *.inv.fasta file
if args.fi.split('.')[-2:] == ["inv", "fasta"]:
    fixInv = True

# return next read title ending with '\n'
def mergeSplittedLines(fi, fout):
    line = fi.readline()
    seq = ""
    while line[0] != ">":
        seq += line.rstrip()
        line = fi.readline()

        if not line: # end of file
            if len(seq) >= th:
                fout.write(seq + "\n")
            return ""

    if len(seq) >= th:
        fout.write(seq + "\n")
    
    return line


# get haps
haps = []
if args.useCSV:
    with open("haplotypes.csv", newline="") as f:
        for row in csv.reader(f):
            if row[0] != "Name": haps.append(row[0])
else:
    haps = args.haps.split()
print(haps)


# fix file format
for hap in haps:
    print("processing ", hap)

    with open(hap+"."+args.fi+".fix" , 'w') as outf:
        with open(hap+"."+args.fi, 'r') as f2: # corrupted file
            ind = -1
            if not args.fixTitle:
                line2 = f2.readline()
                while line2:
                    ind += 1
                    outf.write(line2)
                    line2 = mergeSplittedLines(f2, outf)
                    
            else:
                with open(hap+".combined-hap.fasta", 'r') as f1: # reference
                    line2 = f2.readline()

                    for line1 in f1:
                        if not line1: break
                        if line1[0] == ">":
                            ind += 1

                            if fixInv:
                                #print('/'.join(line2.split('_')[:3]))
                                # if read title is found in reference, merge splitted lines of a read to a single read
                                if line1.rstrip() == '/'.join(line2.split('_')[:3]):
                                    while line1.rstrip() == '/'.join(line2.split('_')[:3]):
                                        outf.write(line1)
                                        line2 = mergeSplittedLines(f2, outf)
                                else:
                                    outf.write(line1)

                            else:
                                outf.write(line1)
                                # if read title is found in reference, merge splitted lines of a read to a single read
                                if line1.rstrip() == line2.split(':')[0]:
                                    line2 = mergeSplittedLines(f2, outf)

            print("finished at locus", ind)
```
