## Supplemental code for "The motif composition of variable-number tandem repeats impacts gene expression": getCovByLocus.397.sh

```
#!/usr/bin/env bash

#refctrl=/home/cmb-17/mjc/vntr_genotyping/cmb-16/work/vntr/hapdb/a1_regions/ctrl/pan.fn0.bed.bak
refctrl=/home/cmb-17/mjc/vntr_genotyping/cmb-16/work/vntr/hapdb/a1_regions/ctrl/pan.fn2.bed
out=ctrl.397.cov
datadir=/home/cmb-16/nobackups/mjc/data_download/phase3

gi=0
for cram in $datadir/*.cram; do
    g=$(basename $cram | cut -b 1-7)
    samtools bedcov $refctrl $cram | awk '{print $4/($3-$2)}' | tr '\n' '\t' |
    awk -v g=$g -v gi=$gi 'BEGIN {OFS="\t"} {$1=$1; print gi, g, $0}'
    ((++gi))
done > $out
```
