## Supplemental code for "The motif composition of variable-number tandem repeats impacts gene expression": individualExpansion.py

```
#!/usr/bin/env python3

import sys
import vntrutils as vu
import numpy as np
import pickle
from multiprocessing import Pool, Manager


def individualTRexpansion(seqs, poss, hap, locus, UB, ksize=21, zoomout=False, verbose=False):
    """plot loci w/ primary contamination only"""
    
    if locus not in poss[hap]: return False, False
    seq = seqs[hap][locus]
    pos = poss[hap][locus]
    
    start, end = pos
    TRsize = end - start
    kmers = vu.read2kmers(seq, ksize)
    kmersi = vu.getkmersindex(kmers)

    xs, ys, badfrom, badto = [], [], [], []
    badkmc = np.zeros(2, dtype=int)
    newregion = pos
    for kmer, indices in kmersi.items():
        if len(indices) > 1:

            # analyze contamination
            badkmc_, badindices = vu.getbadkmc_bothhaps(indices, indices, pos, pos, getindices=True)
            badkmc_ = badkmc_[:2]
            badindices = badindices[0]
            badkmc += badkmc_

            # compute new region to remove contamination
            if badindices:
                newregion = min(*(badindices), newregion[0]), max(max(badindices)+1, newregion[1]) # half open at the end

    while np.sum(badkmc) and start - newregion[0] < UB and newregion[1] - end < UB:
        if verbose: print(newregion, badkmc)
        prevregion = newregion
        badfrom, badto = [], []
        badkmc = np.zeros(2, dtype=int)
        for kmer, indices in kmersi.items():
            if len(indices) > 1:
                # analyze contamination
                badkmc_, badindices = vu.getbadkmc_bothhaps(indices, indices, prevregion, prevregion, getindices=True)
                badkmc_ = badkmc_[:2]
                badindices = badindices[0]
                badkmc += badkmc_

                # compute new region to remove contamination
                if badindices:
                    newregion = min(*(badindices), newregion[0]), max(max(badindices)+1, newregion[1]) # half open at the end

    if pos==newregion: return False, False, newregion
    else:
        if not np.any(badkmc): return True, True, newregion
        else: return True, False, newregion


def minibatch_both_hap_expansion(minibatch):
    [miniseqs, miniposs] = pickle.load(open("seqpos{}.pickle".format(minibatch), 'rb'))
    newposs = [{}, {}]
    stats = np.zeros((2,2), dtype=int)

    for hap in [0,1]:
        nexpanded, nresolved = 0, 0

        for locus in range(nloci):
            if locus % nprocess != minibatch: continue
            if locus % (nloci//100) == 0:
                print(".", end="", flush=True)
            if locus not in miniposs[0]: continue

            expanded, resolved, newposs[hap][locus] = individualTRexpansion(miniseqs, miniposs, hap, locus, TRWINDOW-FS, ksize=KSIZE)

            if expanded:
                nexpanded += 1
                nresolved += resolved
        
        stats[hap] = nexpanded, nresolved    
    return stats, newposs


if __name__ == "__main__":
    nprocess, KSIZE, FS, UB, TRWINDOW, nloci = [int(v) for v in sys.argv[1:7]]
    newposs = [{}, {}]
    stats = np.zeros((2,2), dtype=int)

    print("Running individualExpansion", end="")
    p = Pool(nprocess)
    results = p.map(minibatch_both_hap_expansion, list(range(nprocess)))
    p.close(); p.join()
    print()

    for i in range(nprocess):
        for hap in [0,1]:
            for k, v in results[i][1][hap].items():
                newposs[hap][k] = v
        stats += results[i][0]
    print(stats)

    pickle.dump([stats, newposs], open("newposs_stat.pickle", 'wb'))
```
