## Supplemental code for "The motif composition of variable-number tandem repeats impacts gene expression": jointExpansion.py

```
#!/usr/bin/env python3

import sys
import vntrutils as vu
import numpy as np
import pickle
from multiprocessing import Pool, Manager


def jointTRexpansion(seqs, poss, oldposs, locus, UB, ksize=21, ax=None, verbose=0):
    """plot loci w/ primary contamination only"""
    
    oldpos0 = oldposs[0][locus]
    oldpos1 = oldposs[1][locus]
    ostart0, oend0 = oldpos0[0], oldpos0[1]-ksize+1
    ostart1, oend1 = oldpos1[0], oldpos1[1]-ksize+1
    pos0 = poss[0][locus]
    pos1 = poss[1][locus]
    start0, end0 = pos0[0], pos0[1]-ksize+1
    start1, end1 = pos1[0], pos1[1]-ksize+1

    ctg0 = seqs[0][locus]
    ctg1 = seqs[1][locus]
    kmers0 = vu.read2kmers(ctg0, ksize, rightflank=ksize-1)
    kmers1 = vu.read2kmers(ctg1, ksize, rightflank=ksize-1)
    cokmers = (set(kmers0) & set(kmers1)) - set([0xffffffffffffffff])
    cokmersi = vu.getcokmersindex(cokmers, kmers0, kmers1)

    badkmc = np.zeros(4, dtype=int) # 0L, 0R, 1L, 1R
    newregion0, newregion1 = (start0, end0), (start1, end1)
    for kmer in cokmers:
        indices0 = cokmersi[0][kmer]
        indices1 = cokmersi[1][kmer]

        # analyze contamination
        badkmc_, badindices = vu.getbadkmc_bothhaps(indices0, indices1, newregion0, newregion1, getindices=True)
        badkmc += badkmc_

        # compute new region to remove contamination
        if badindices[0]:
            newregion0 = min(*(badindices[0]), newregion0[0]), max(max(badindices[0])+1, newregion0[1]) # half open at the end
            newregion1 = min(*(badindices[1]), newregion1[0]), max(max(badindices[1])+1, newregion1[1]) # half open at the end
            if verbose >= 2:
                print(badkmc_, badindices)

    es0 = (ostart0 - newregion0[0], newregion0[1] - oend0) # expansionsize h0
    es1 = (ostart1 - newregion1[0], newregion1[1] - oend1) # expansionsize h1
    if verbose and np.any(badkmc): print(es0, es1, badkmc)
    while np.any(badkmc) and es0[0] < UB and es0[1] < UB and es1[0] < UB and es1[1] < UB:
        prevregion0, prevregion1 = newregion0, newregion1
        badkmc = np.zeros(4, dtype=int) # 0L, 0R, 1L, 1R
        for kmer in cokmers:
            indices0 = cokmersi[0][kmer]
            indices1 = cokmersi[1][kmer]

            # analyze contamination
            badkmc_, badindices = vu.getbadkmc_bothhaps(indices0, indices1, prevregion0, prevregion1, getindices=True)
            badkmc += badkmc_

            # compute new region to remove contamination
            if badindices[0]:
                newregion0 = min(*(badindices[0]), newregion0[0]), max(max(badindices[0])+1, newregion0[1]) # half open at the end
                newregion1 = min(*(badindices[1]), newregion1[0]), max(max(badindices[1])+1, newregion1[1]) # half open at the end
                if verbose >= 2:
                    print(badindices)

        es0 = (ostart0 - newregion0[0], newregion0[1] - oend0)
        es1 = (ostart1 - newregion1[0], newregion1[1] - oend1)
        if verbose: print(es0, es1, badkmc)

    outregion0, outregion1 = (newregion0[0], newregion0[1]+ksize-1), (newregion1[0], newregion1[1]+ksize-1)
    if newregion0 == (start0, end0) and newregion1 == (start1, end1): 
        return False, None, pos0, pos1 # good locus after individualExpansion
    if not np.any(badkmc): 
        return True, True, outregion0, outregion1
    else:
        return True, False, outregion0, outregion1

def minibatch_jointExpansion(minibatch):
    [miniseqs, mininewposs, miniposs] = pickle.load(open("seqnewpos{}.pickle".format(minibatch), 'rb'))
    jointnewposs = [{}, {}]
    jointstats = np.zeros(2, dtype=int)
    nexpanded, nresolved = 0, 0

    for locus in range(nloci):
        if locus % nprocess != minibatch: continue
        if locus % (nloci//100) == 0:
            print(".", end="", flush=True)
        if locus not in miniposs[0]: continue

        expanded, resolved, newregion0, newregion1 = jointTRexpansion(miniseqs, mininewposs, miniposs, locus, TRWINDOW-FS, ksize=KSIZE)

        jointnewposs[0][locus] = newregion0
        jointnewposs[1][locus] = newregion1
        
        if expanded:
            nexpanded += 1
            nresolved += resolved
    
    jointstats = nexpanded, nresolved
    return jointstats, jointnewposs


if __name__ == "__main__":
    nprocess, KSIZE, FS, UB, TRWINDOW, nloci = [int(v) for v in sys.argv[1:7]]
    jointnewposs = [{}, {}]
    jointstats = np.zeros(2, dtype=int)
    
    print("Runnning JointExpansion", end="")
    p = Pool(nprocess)
    results = p.map(minibatch_jointExpansion, list(range(nprocess)))
    p.close(); p.join()
    print()
    
    for i in range(nprocess):
        for hap in [0,1]:
            for locus, newregion in results[i][1][hap].items():
                jointnewposs[hap][locus] = newregion
        jointstats += results[i][0]
    print(jointstats)

    pickle.dump([jointstats, jointnewposs], open("jointnewposs_stat.pickle", 'wb'))
```
