## Supplemental code for "The motif composition of variable-number tandem repeats impacts gene expression": kmc2length.LOO.py

```
#!/usr/bin/env python3

import os
import argparse
import numpy as np
import pandas as pd
import pickle
import warnings
from vntrutils import readKms


def loadbedsize(bed):
    tmp = np.loadtxt(bed, dtype=int, usecols=[1,2], ndmin=2)
    return tmp[:,1] - tmp[:,0]

def get1DIQRmask(data, whis=1.5):
    m = np.isfinite(data)
    q1s = np.quantile(data[m], 0.25)
    q3s = np.quantile(data[m], 0.75)
    kIQRs = (q3s - q1s) * whis
    return ~m | (data < (q1s - kIQRs)) | (data > (q3s + kIQRs))

def loadvntrmat(genomes, prefix="pan", suffix="IL.tr.kmers"):
    kmerfnames = {}
    for g in genomes:
        kmerfnames[g] = f'{prefix}.{g}.{suffix}'
        
    vntrmat = np.zeros([len(genomes), nloci], dtype=int)
    for gind, g in enumerate(genomes):
        readKms(kmerfnames[g], vntrmat[gind])
    return vntrmat

def processCtrlBamCov(covmat, whis=1.5):
    ctrlsize = loadbedsize(args.covbed)
    cov = covmat@ctrlsize / np.sum(ctrlsize)
    badmask = np.zeros_like(ctrlsize, dtype=bool)

    ### compute coverage for each locus; normalize wrt sample global coverage
    normcovmat = covmat / (covmat@ctrlsize / np.sum(ctrlsize))[:,None]

    ### check variance
    stds = np.std(normcovmat, axis=0)
    badmask = np.logical_or(badmask, get1DIQRmask(stds))

    ### check if mean is biased
    means = np.mean(normcovmat, axis=0)
    badmask = np.logical_or(badmask, get1DIQRmask(means))

    print(f'\t{np.sum(badmask)} out of {badmask.size} unique regions removed')

    ### reject outliers
    pctrlsize = ctrlsize[~badmask]
    pcovmat = covmat[:,~badmask]
    pcov = pcovmat@pctrlsize / np.sum(pctrlsize)

    ### covmat for nearest neighbor search
    normcovmat = covmat / cov[:,None]
    pnormcovmat = pcovmat / pcov[:,None]
    return pcov, pnormcovmat, normcovmat

def loadPanLen(wd, genomes):
    master = np.loadtxt(f"{args.pantr}", dtype=object, ndmin=2)[:,3:]
    master[master == "None"] = np.nan
    cm = np.tile(np.array([0,1,1,0], dtype=bool), 2*genomes.size)
    master = master[:,cm].astype(float)
    panlenmat = np.zeros([genomes.size, nloci])
    for gi, g in enumerate(genomes):
        l = master[:,[4*gi+1,4*gi+3]] - master[:,[4*gi+0,4*gi+2]]
        m = np.isfinite(l)
        panlenmat[gi] = np.nansum(l, axis=1) / np.sum(m, axis=1)
    return panlenmat

def getBiasMat(genomes):
    warnings.filterwarnings('ignore')
    biasmat = vntrmat / (np.clip(panlenmat-args.ksize+1, 0, None) * pbamcov[:,None])
    warnings.filterwarnings('default')
    return biasmat

def dumpStep1(wd):
    #df.to_csv(f'{wd}/step1_results.tsv', )
    with open(f'{wd}/step1_results.pickle', 'wb') as f:
        pickle.dump([pbamcov, pncovmat, ncovmat, panlenmat, biasmat], f)
    return

def loadStep1(wd):
    with open(f'{wd}/step1_results.pickle', 'rb') as f:
        pbamcov, pncovmat, ncovmat, panlenmat, biasmat = pickle.load(f)
    return pbamcov, pncovmat, ncovmat, panlenmat, biasmat

def matCorrelation(mat, n_cluster=4, alpha=0.01, reject=True, img=[1,1], outdir="./", annotateXY=None):
    warnings.filterwarnings('ignore')
    
    n = genomes.size
    stats = np.zeros([mat.shape[0], mat.shape[0]])
    for i in range(mat.shape[0]):
        for j in range(mat.shape[0]):
            if reject:
                bm = get1DIQRmask(mat[i]) | get1DIQRmask(mat[j]) | (mat[i] == 0) | (mat[j] == 0)
            else:
                bm = ~np.isfinite(mat[i]) | ~np.isfinite(mat[j]) | (mat[i] == 0) | (mat[j] == 0)
            gt, est = mat[i][~bm], mat[j][~bm]
            stats[i,j] = np.nanmean(np.abs(1 - gt/est))
            
    warnings.filterwarnings('default')
    return stats

def getBestUsingSeqrunPrior(srtgimat, gs):
    g2sr = dict(np.hstack((config["genome"].to_numpy()[:,None], config["sequencing_run"].to_numpy()[:,None])))
    badgis = np.nonzero(np.isin(gs, badg))[0]
    bestind = np.zeros(srtgimat.shape[0], dtype=int)
    for gi, srtgis in enumerate(srtgimat):
        run = g2sr[gs[gi]]
        for bidx in srtgis:
            # only use nearest neighbor from the same sequencing run, except samples from individual work
            # exclude bad genomes from nearest neighbor search
            if bidx not in badgis and (run == g2sr[gs[bidx]] or run == "individual"):
                bestind[gi] = bidx
                break
            else:
                continue
    return bestind

def lenPred(trlen, ilkms, bias, cov):
    est = np.full(trlen.size, np.nan)
    m = (trlen > 0) & (bias > 0) & np.isfinite(bias)
    est[m] = ilkms[m] / (cov * bias[m])
    est[est>=1] += (args.ksize - 1)
    est[est<1] *= args.ksize
    err = np.abs(est-trlen)/trlen
    return est, err

def BiasCorrectedLenPred(outdir="./"):
    ctrlcovmats = [ncovmat, pncovmat]
    methods = ["ncovmat", "pncovmat"]
    opts = {"alpha":1, "img":None}
    nplt = np.sum(LOOmask)
    ncol = 5
    nmethod = len(ctrlcovmats)
    ests = np.full([nmethod, nplt, nloci], np.nan)
    errs = np.full([nmethod, nplt, nloci], np.nan)
    for mi, covmat_ in enumerate(ctrlcovmats):
        covmat = covmat_[LOOmask]
        LOOlenmat = panlenmat[LOOmask]
        LOObiasmat = biasmat[LOOmask]
        LOOpbamcov = pbamcov[LOOmask]
        metric = matCorrelation(covmat, **opts)
        srtgimat = np.argsort(metric, axis=1)[:,1:]
        bestind = getBestUsingSeqrunPrior(srtgimat, LOOgenomes)
            
        for idx, bidx in enumerate(bestind):
            g, ghat = LOOgenomes[idx], LOOgenomes[bidx]
            trlen = LOOlenmat[idx]
            est, err = lenPred(LOOlenmat[idx], LOOmat[idx], LOObiasmat[bidx], LOOpbamcov[idx])
            e = np.nanmean(err)
            ests[mi,idx] = est
            errs[mi,idx] = err

        print(f"{methods[mi]} MAPE = {np.nanmean(errs):3f}")
    return ests, errs

def SaveEstErr(ests, errs, bestmethodi, outdir="./"):
    trid = [f"{r[0]}_{r[1]}_{r[2]}" for r in np.loadtxt(f"{args.pantr}", dtype=object, ndmin=2)[:,:3]]
    df = pd.DataFrame(errs[bestmethodi].T, columns=LOOgenomes, index=trid)
    df.to_csv(f'{outdir}/abs_percentage_err.tsv', sep="\t", na_rep="nan")
    df = pd.DataFrame(ests[bestmethodi].T, columns=LOOgenomes, index=trid)
    df.to_csv(f'{outdir}/estimated_TR_len.tsv', sep="\t", na_rep="nan")


if __name__ == "__main__":
    ap = argparse.ArgumentParser(description=\
            "The program process data in 2 steps\t"+\
            "  [1] Compute adjusted bam coverage and VNTR-specific biases\t"+\
            "  [2] Predict VNTR lengths from kmer genotype in leave-one-out setting\t"+
            "* Use --skip1 to avoid recomputing step 1")

    ap.add_argument("--ksize", help="kmer size of RPGG", type=int, required=True)
    ap.add_argument("--genome", help="genome config file", required=True)
    ap.add_argument("--nloci", help="number of VNTR loci", type=int, required=True)
    ap.add_argument("--pantr", help="locus mapping/coordinate file for all haplotypes")

    ap.add_argument("--skip1", help="skip step 1 and read from step1_results.pickle", action="store_true")
    ap.add_argument("--cov", help="bam coverage file; skipped if --skip1")
    ap.add_argument("--covbed", help="unique region bed file; skipped if --skip1")

    ap.add_argument("--config", help="config file for leave-one-out analysis, indicating the genomes to analyze, sequencing runs and outliers", required=True)
    ap.add_argument("--LOOpref", help="LOO kmer file prefix", nargs='?', default="")
    ap.add_argument("--LOOsuff", help="LOO kmer file suffix", nargs='?', default="IL.tr.kmers")
    args = ap.parse_args()
    wd = os.getcwd()


    ### step 1
    outdir = f'{wd}/analysis'
    genomes = np.loadtxt(args.genome, dtype=object, ndmin=1) # "/home/cmb-16/mjc/tsungyul/work/vntr/hapdb/config/genomes.txt"
    nloci = args.nloci

    if not args.skip1:
        print("1-1: processing bam coverage", flush=True)
        rawcovmat = np.loadtxt(args.cov, dtype=object, ndmin=2)
        pbamcov, pncovmat, ncovmat = processCtrlBamCov(rawcovmat[:,2:].astype(float))

        print("1-2: loading kmer genotypes", flush=True)
        vntrmat = loadvntrmat(genomes, prefix=f'{wd}/pan')

        print("1-3: computing VNTR-specific biases", flush=True)
        panlenmat = loadPanLen(wd, genomes)
        biasmat = getBiasMat(genomes)

        print("1-4: saving step 1 results", flush=True)
        dumpStep1(outdir)
    else:
        print("1-1: loading step 1 results", flush=True)
        pbamcov, pncovmat, ncovmat, panlenmat, biasmat = loadStep1(outdir)

    ### step 2
    print("2-1: loading leave-one-out kmer genotypes", flush=True)
    config = pd.read_csv(args.config, sep="\t")
    LOOmask = np.isin(genomes, config["genome"])
    LOOgenomes = genomes[LOOmask]
    LOOmat = loadvntrmat(genomes[LOOmask], prefix=f'{wd}/{args.LOOpref}' if args.LOOpref else f"{wd}/")

    print("2-2: computing nearest bias", flush=True)
    badg = genomes[LOOmask][np.nonzero(config["bad"].to_numpy())[0]]
    vntrstat = matCorrelation(biasmat, n_cluster=5, outdir=outdir)

    print("2-3: predicting VNTR lengths", flush=True)
    lenEstimates, lenPredErrs = BiasCorrectedLenPred(outdir=outdir)

    print("2-4: saving step 2 results", flush=True)
    SaveEstErr(lenEstimates, lenPredErrs, bestmethodi=0, outdir=outdir)  # ncovmat is empirically better than pncovmat
```
