## Supplemental code for "The motif composition of variable-number tandem repeats impacts gene expression": kmc2length.py

```
#!/usr/bin/env python3

import os
import argparse
import numpy as np
import pandas as pd
import warnings
from vntrutils import readKms
#import pickle


def get1DIQRmask(data, whis=1.5):
    m = np.isfinite(data)
    q1s = np.quantile(data[m], 0.25)
    q3s = np.quantile(data[m], 0.75)
    kIQRs = (q3s - q1s) * whis
    return ~m | (data < (q1s - kIQRs)) | (data > (q3s + kIQRs))

def loadvntrmat(fnames):
    vntrmat = np.zeros([fnames.size, nloci], dtype=int)
    for fi, f in enumerate(fnames):
        readKms(f, vntrmat[fi])
    return vntrmat

def processCtrlBamCov(covmat, whis=1.5):
    cov = covmat@ctrlsize / np.sum(ctrlsize)
    badmask = np.zeros_like(ctrlsize, dtype=bool)

    ### compute coverage for each locus; normalize wrt sample global coverage
    normcovmat = covmat / (covmat@ctrlsize / np.sum(ctrlsize))[:,None]

    ### check variance
    stds = np.std(normcovmat, axis=0)
    badmask = np.logical_or(badmask, get1DIQRmask(stds))

    ### check if mean is biased
    means = np.mean(normcovmat, axis=0)
    badmask = np.logical_or(badmask, get1DIQRmask(means))

    print(f'\t{np.sum(badmask)} out of {badmask.size} unique regions removed')

    ### reject outliers
    pctrlsize = ctrlsize[~badmask]
    pcovmat = covmat[:,~badmask]
    pcov = pcovmat@pctrlsize / np.sum(pctrlsize)

    ### covmat for nearest neighbor search
    normcovmat = covmat / cov[:,None]
    pnormcovmat = pcovmat / pcov[:,None]
    return pcov, pnormcovmat, normcovmat

def loadLSB(f):
    df = pd.read_csv(f, sep="\t", index_col=0)
    nloci0 = df.shape[0] - nloci
    ntrbiasmat_db = df.iloc[:nloci0].to_numpy().T
    trbiasmat_db = df.iloc[nloci0:].to_numpy().T
    dbgenomes = np.array(df.columns)
    cbed = np.array([v[4:].split("_") for v in df.index[:nloci0]], dtype=object)
    ctrlsize = cbed[:,2].astype(int) - cbed[:,1].astype(int)
    return trbiasmat_db, ntrbiasmat_db, dbgenomes, ctrlsize

def rowDistance(mat1, mat2, reject=True):
    # input: N1xL, N2xL matrix. output: N1xN2 distance matrix
    warnings.filterwarnings('ignore')
    
    n1, n2 = mat1.shape[0], mat2.shape[0]
    stats = np.zeros([n1, n2])
    for i in range(n1):
        for j in range(n2):
            if reject:
                bm = get1DIQRmask(mat1[i]) | get1DIQRmask(mat2[j]) | (mat1[i] == 0) | (mat2[j] == 0)
            else:
                bm = ~np.isfinite(mat1[i]) | ~np.isfinite(mat2[j]) | (mat1[i] == 0) | (mat2[j] == 0)
            gt, est = mat1[i][~bm], mat2[j][~bm]
            stats[i,j] = np.nanmean(np.abs(1 - gt/est))
            
    warnings.filterwarnings('default')
    return stats

def lenPred(ilkms, bias, cov):
    est = np.full(nloci, np.nan)
    m = (bias > 0) & np.isfinite(bias)
    est[m] = ilkms[m] / (cov * bias[m])
    est[est>=1] += (args.ksize - 1)
    est[est<1] *= args.ksize
    return est

def BiasCorrectedLenPred(outdir="./"):
    N = trmat.shape[0]
    ests = np.full([N, nloci], np.nan)
    dis = rowDistance(ntrbiasmat, ntrbiasmat_db) 
    bestids = np.argsort(dis, axis=1)[:,0]
    for idx, bidx in enumerate(bestids): # idx: sample index. bidx: index of best estimator in db
        ests[idx] = lenPred(trmat[idx], trbiasmat_db[bidx], pbamcov[idx])
    return ests

def SaveEstErr(ests, outdir="./"):
    trid = ["_".join(r) for r in trbed]
    df = pd.DataFrame(ests.T, index=trid)
    df.to_csv(f'{outdir}/estimated_TR_len.tsv', sep="\t", na_rep="nan")


if __name__ == "__main__":
    ap = argparse.ArgumentParser(description=\
            "Predict VNTR lengths from kmer genotype using precomputed locus-specific biases (LSB)\n"+\
            "** Please download precomputed LSB from the GitHub release page")

    ap.add_argument("--outdir", help="output directory for estimation/error table", required=True)
    ap.add_argument("--ksize", help="kmer size of RPGG", type=int, required=True)
    ap.add_argument("--kmers", help="file of sorted kmer file names", required=True)
    ap.add_argument("--trbed", help="VNTR bed file", required=True)
    ap.add_argument("--LSB", help="Precomputed non-TR and TR locus-specific sampling biases", required=True)
    ap.add_argument("--cov", help="bam coverage file", required=True)
    ap.add_argument("--covbed", help="unique region bed file", required=True)
    args = ap.parse_args()
    wd = os.getcwd()
    outdir = args.outdir

    print("Loading metadata and precomputed TR/NTR LSB", flush=True)
    trbed = np.loadtxt(args.trbed, dtype=object, ndmin=2)
    nloci = trbed.shape[0]
    trbiasmat_db, ntrbiasmat_db, dbgenomes, ctrlsize = loadLSB(args.LSB)

    print("Computing NTR LSB in current dataset", flush=True)
    rawcovmat = np.loadtxt(args.cov, dtype=object, ndmin=2)
    pbamcov, _, ntrbiasmat = processCtrlBamCov(rawcovmat[:,2:].astype(float))

    print("Loading genotype data", flush=True)
    trmat = loadvntrmat(np.loadtxt(args.kmers, dtype=object, ndmin=1))
    #with open("analysis/trmat.pickle", 'wb') as f:
    #    pickle.dump(trmat, f)
    #with open("analysis/trmat.pickle", 'rb') as f:
    #    trmat = pickle.load(f)

    print("Estimating VNTR Length", flush=True)
    lenEstimates = BiasCorrectedLenPred(outdir=outdir)

    print("Writing outputs", flush=True)
    SaveEstErr(lenEstimates, outdir=outdir)  # ncovmat is empirically better than pncovmat
```
