## Supplemental code for "The motif composition of variable-number tandem repeats impacts gene expression": kmers.linreg.py

```
#!/usr/bin/env python3

import argparse
import vntrutils as vu
import numpy as np
import statsmodels.api as sm
#from sklearn.linear_model import LinearRegression

ap = argparse.ArgumentParser(description="read *.kmers and output regression plots and prediction results")
ap.add_argument("pacbio", help="PB.tr.kmers with locus, kmer name, and kmer count information")
ap.add_argument("illumina", help="IL.tr.kmers", nargs='+')
ap.add_argument("out", help="output file prefix")
ap.add_argument("--index", help="tr.kmers with locus and kmer name info used for IL.tr.kmers when only count info is present", nargs='?', const="", default="")
ap.add_argument("--mapkmer", help="map pangenome kmers to genome kmers", action="store_true")
ap.add_argument("--mode", help="Outlier rejection mode in regression. Choose from 'invalid', 'invalid|zero', 'invalid|bad' or 'invalid|bad|zero' Default: invalid", nargs='?', const="invalid", default="invalid")
ap.add_argument("--combine", help="combine multiple IL.kmers when multiple IL.kmers are provided; will not perform regression. Default: False", action='store_true')
ap.add_argument("--plot", help="plot regression results of the loci specified.", nargs='?', const="", default="")
ap.add_argument("--threshold", help="rejecting outliers locating threshold*std away from the mean. Default: 10", type=int, nargs='?', const=10, default=10)
ap.add_argument("--R2threshold", help="plot summary report for loci with R^2 > threshold. Default: -1", type=float, nargs='?', const=-1, default=-1)
args = ap.parse_args()
print(args)
mapkmer = args.mapkmer
threshold = args.threshold
R2threshold = args.R2threshold
combine = args.combine and len(args.illumina) != 1
plotloci = set([int(v) for v in args.plot.split(",")]) if args.plot else set([])

print("reading illumina kmers")
y = {}
for fname in args.illumina:
    print("\treading", fname)
    if combine:
        vu.readKmerDict(fname, y)
    else:
        if args.index:
            vu.readKmersWithIndex(fname, args.index, y, sort=True, kmerName=mapkmer)
        else:
            vu.readKmers(fname, y, sort=True, kmerName=mapkmer)
if combine:
    vu.writeKmerDict(args.out, y)
    exit(0)

nloci = len(y)
print("#loci:", nloci)

print("reading pacbio kmers")
x = {}
vu.readKmers(args.pacbio, x, sort=True, kmerName=mapkmer)

data = {}
for k, v in y.items():
    if v.size and x[k].size:
        if mapkmer:
            m = np.isin(y[k][:,0], x[k][:,0])
            data[k] = np.column_stack((np.insert(x[k][:,1],0,0), np.insert(y[k][m,1],0,0)))
        else:
            data[k] = np.column_stack((np.insert(x[k],0,0), np.insert(y[k],0,0)))

results = np.zeros((nloci, 4))
for k, v in x.items():
    if v.size:
        if mapkmer:
            truth = np.sum(v[:,1])
        else:
            truth = np.sum(v)
        results[k,0] = truth

for k, v in data.items():
    slope, _, r2, pred = vu.PlotRegression(v[:,0:1], v[:,1:2], "Assembly kmer counts", "Read kmer counts", 
                                           f"locus {k}, n={v.shape[0]}", f"{args.out}.{k}", outlier=args.mode, pred=True)
    if k % 1000 == 0:
        print(".", end="", flush=True)
    results[k, 1:] = [pred, slope, r2]
print()

print("writing outputs")
np.savetxt(f'{args.out}.pred', results, fmt=['%i','%.1f','%.2f','%.4f'], delimiter="\t", header="TrueDosage\tPredDosage\tSlope\tr^2")
```
