## Supplemental code for "The motif composition of variable-number tandem repeats impacts gene expression": liftbed.clean.py

```
#!/usr/bin/env python3

import sys
import numpy as np
from collections import defaultdict

# Usage: program <liftbed>
if len(sys.argv) != 2:
    sys.exit("Usage: program <liftbed>\nLiftbed should be sorted. Filtered regions are printed to stdout")


# assign chrom mapped by contig based on majority vote
# fix edge cases of t5_t3
# fix edge case where a region is split into >2 segments
# Retain strand orientation tag
# split read may have different orientation
class DupInfo:
    def __init__(self):
        self.dup = False
        self.valid = True
        self.asm = ""
        self.regions = []
        self.start = -1
        self.end = -1
        self.strand = []
        
def cleanbed():
    r2a = defaultdict(DupInfo)
    for f1, f2, f3, f4, _, f6 in lb:
        r = "_".join(f4.split("_")[:3])
        f2, f3 = int(f2), int(f3)
        if r not in r2a:
            r2a[r].asm = f1
            r2a[r].regions.append((f2,f3))
            r2a[r].start = f2
            r2a[r].end = f3
            r2a[r].strand.append(f6)
        else:
            if r2a[r].valid:
                if r2a[r].asm == f1:
                    r2a[r].dup = True
                    r2a[r].regions.append((f2,f3))
                    r2a[r].strand.append(f6)
                    r2a[r].start = min(r2a[r].start, f2)
                    r2a[r].end = max(r2a[r].end, f3)
                else:
                    r2a[r].valid = False
                    
    a2ch = defaultdict(lambda: defaultdict(int))
    for f1, f2, f3, f4, _, f6 in lb:
        ch = f4.split("_")[0][3:]
        a2ch[f1][ch] += 1

    a2mc = {} # asm to major chrom
    for k0, v0 in a2ch.items():
        tc, mc, mr = 0, 0, ""
        for k1, v1 in v0.items():
            tc += v1
            if v1 > mc:
                mch = k1
                mc = v1
        if mc/tc >= 0.8: # check major mapped chrom freq
            a2mc[k0] = mch
                    
    # write clean bed
    s2i = {"+": 1, "-": -1}
    for k, v in r2a.items():
        rr = "\t".join(k.split("_"))
        if v.valid and v.asm in a2mc:
            ch = k.split("_")[0][3:]
            if ch == a2mc[v.asm]: # check if mapped chrom is the same as major mapped chrom
                strand = np.all(np.array(v.strand) == v.strand[0]) * s2i[v.strand[0]] # 1: plus strand, -1: minus strand, 0: mixed
                print(f'{v.asm}\t{v.start}\t{v.end}\t{rr}\t{strand}')


if __name__ == "__main__":
    lb = np.loadtxt(sys.argv[1], dtype=object, ndmin=2, comments=None)
    cleanbed()
```
