## Supplemental code for "The motif composition of variable-number tandem repeats impacts gene expression": mergeMBEbed.py

```
#!/usr/bin/env python3

import sys
import numpy as np

def parseMergeSet():
    ms = []
    bs = set()
    v2si = {}
    si = 0
    with open("mbe.m0.loci") as f:
        hap = ""
        for line in f:
            if line[0] == ">":
                hap = line.rstrip()[1:]
                continue
            seq = sorted([int(v) for v in line.rstrip().split(",")])
            skip = seq[0] in bs # check if good v reported by this hap is bad in another hap
            for i in range(1,len(seq)):
                skip |= seq[i] in bs
                if seq[i] != seq[i-1] + 1:
                    skip = True
                    for v in seq:
                        if v in v2si: # check if bad v reported by this hap is good in another hap
                            si_ = v2si[v]
                            if ms[si_] is not None:
                                for v_ in ms[si_]:
                                    bs.add(v_)
                                ms[si_] = None
                            v2si.pop(v)
                        bs.add(v)
                    print(f"Bad seq {seq} in {hap}")
                    break
            else:
                if skip:
                    for v in seq:
                        bs.add(v)
                        if v in v2si: # check if v was once reported good
                            si_ = v2si[v]
                            #if si_ == 12968: print("[2XXX]", seq, ms[si_])
                            ms[si_] = None
                            v2si.pop(v)
                    continue
                sis = set() # set index of existing v
                for v in seq:
                    if v in v2si:
                        sis.add(v2si[v])
                if len(sis) == 0: # make a new set
                    ms.append(set())
                    for v in seq:
                        ms[-1].add(v)
                        v2si[v] = si
                    si += 1
                else:
                    si_s = None
                    if len(sis) > 1:
                        vstr = f"{seq}"
                        for si_i in sis:
                            vstr += f"{ms[si_i]}"
                        print(f"[Note] {hap} induced merging across {vstr}", flush=True) # enable merging multiple sets
                    for si_i in sis:
                        if si_s is None:
                            si_s = si_i
                        else:
                            ms[si_s] |= ms[si_i]
                            for v in ms[si_i]:
                                v2si[v] = si_s
                            ms[si_i] = None
                    ms[si_s] |= set(seq)
                    for v in seq:
                        v2si[v] = si_s
    ms = np.array(ms, dtype=object)
    ms = ms[ms!=None].tolist()
    for i1s_ in ms:
        assert len(i1s_ & bs) == 0
    return ms, bs

def getdist(bed):
    out = []
    if int(bed[0,2]) == 1: # no inversion
        for i in range(bed.shape[0]-1):
            out.append(int(bed[i+1,0]) - int(bed[i,1]))
    else:
        for i in range(bed.shape[0]-1):
            out.append(int(bed[i,0]) - int(bed[i+1,1]))
    return out

def writeBed_MergeMBE(MAXSVLEN=10000):
    ms, bs = parseMergeSet()
    
    # QC on merging set
    panbed = np.loadtxt(f"pan.tr.mbe.v1.bed", dtype=object, ndmin=2, comments=None)
    i1togood = {}
    qcb = [] # QC bad
    for i1s_ in ms:
        i1s = sorted(list(i1s_))
        nm = len(i1s)-1
        dist = np.full([nm, 2*ng], np.nan)
        for hi in range(2*ng):
            if np.all(panbed[i1s,3+hi*4] != "None"):
                if np.any(panbed[i1s,3+hi*4] != panbed[i1s[0],3+hi*4]):
                    print(f"[Haplotype removed] merging across contigs: {hi}\t{i1s}\n {panbed[i1s,3+hi*4]}")
                else:
                    if np.any(panbed[i1s,6+hi*4] != panbed[i1s[0],6+hi*4]):
                        print(f"[Note] {i1s} mixed orientation")
                    dist[:,hi] = getdist(panbed[i1s,4+hi*4:7+hi*4])
        good = np.all(np.isfinite(dist), axis=0)
        if np.nanmax(dist) > MAXSVLEN:
            qcb.append(i1s_)
            print(f"[Loci removed] huge SV, {i1s}")
        elif np.sum(good)/(2*ng) < THRESH:
            qcb.append(i1s_)
            print(f"[Loci removed] QC failed {i1s}")
        else:
            i1togood[i1s[0]] = good # record hap to remove
    for i1s_ in qcb:
        assert i1s_ in ms
        ms.remove(i1s_)
        for i1 in i1s_:
            bs.add(i1)
    nmi = 0
    mis = set()
    for i1s_ in ms:
        nmi += len(i1s_)
        for i1 in i1s_:
            mis.add(i1)
        
    # fill v2 bed
    nloci1, _ = panbed.shape
    for i1s_ in ms:
        assert len(i1s_ & bs) == 0
    pv2bed = np.full([nloci1-nmi+len(ms)-len(bs), 3+2*ng*4], None, dtype=object)
    nloci2, _ = pv2bed.shape
    i2toi1 = set(list(range(nloci1))) - mis - bs | set([sorted(list(i1s_))[0] for i1s_ in ms])
    i2toi1 = sorted(list(i2toi1)) # map v2 index to v1
    assert nloci2 == len(i2toi1)
    i1toi2 = np.full(nloci1, None, dtype=object)
    i1toi2[i2toi1] = np.arange(nloci2) # map v1 index to v2
    pv2bed = panbed[i2toi1]
    for i1s_ in ms:
        i1s = sorted(list(i1s_))
        # fill ref
        i2 = i1toi2[i1s[0]]
        ids = i1s[0]
        ide = i1s[-1]+1
        refs = min([int(s) for s in panbed[ids:ide,1]])
        refe = max([int(e) for e in panbed[ids:ide,2]])
        pv2bed[i2,[1,2]] = [refs, refe]
        # fill asm
        for hi in range(2*ng):
            if not i1togood[i1s[0]][hi]: # bad hap to remove
                pv2bed[i2,3+hi*4:7+hi*4] = ["None"]*4
                continue
            asms = min([int(s) for s in panbed[ids:ide,4+hi*4]])
            asme = max([int(e) for e in panbed[ids:ide,5+hi*4]])
            pv2bed[i2,4+hi*4:6+hi*4] = [asms, asme]           
    np.savetxt("pan.tr.mbe.v2.bed", pv2bed, delimiter="\t", fmt='%s')
    
    # orthology map
    lmap = np.full([nloci2, 2*ng], ".", dtype=object)
    for hi in range(2*ng):
        m = pv2bed[:,3+4*hi] != "None"
        lmap[m,hi] = np.arange(np.sum(m))
    np.savetxt("OrthoMap.v2.tsv", lmap, delimiter="\t", fmt='%s')
    np.savetxt("locusMap.v2.to.v1.txt", i2toi1, fmt='%s')


if __name__ == "__main__":
    gs = np.loadtxt(sys.argv[1], usecols=0, dtype=object, ndmin=1)
    ng = gs.size
    #panmap = np.loadtxt(sys.argv[2], dtype=object, ndmin=2)[:,3:].astype(int)
    THRESH = float(sys.argv[2])
    writeBed_MergeMBE()
```
