## Supplemental code for "The motif composition of variable-number tandem repeats impacts gene expression": multiBoundaryExpansion.parallel.py

```
#!/usr/bin/env python3

import os
import sys
import numpy as np
from vntrutils import read2kmers
import pickle
from multiprocessing import Pool, Lock, Manager
from time import sleep

class expStat:
    def __init__(self, exp, fail, es, opos, npos):
        self.exp = exp
        self.fail = fail
        self.es = es
        self.opos = opos
        self.npos = npos
    #def __repr__(self):
    #    return f"expanded: {self.exp}\tfail: {self.fail}\tave. exp. size: {self.es:.0f}\nopos: {self.opos}\tnpos: {self.npos}\n"

def loadbeds(panmap):
    beds = np.full([nh, nloci, 4], None, dtype=object)
    for gi, g in enumerate(gs):
        m = panmap[:,gi]==1
        for h in [0,1]:
            hi = 2*gi + h
            bed = np.loadtxt(f"{g}/tmp1.{h}.bed", dtype=object, usecols=[0,1,2,6], ndmin=2, comments=None)
            if np.sum(m) != bed.shape[0]:
                raise ValueError(f"[Error] Inconsistent # of supports between {g}/tmp1.{h}.bed ({bed.shape[0]}) and column {gi+3} of pan.tr.mbe.v0.bed (np.sum(m))")
            beds[hi,m] = bed
        print(".", end="", flush=True)
    return beds

def make_process_pickle():
    for i in range(NPROCESS):
        ibeg, iend = i*bsize, min((i+1)*bsize, nloci)
        indices = np.nonzero(np.sum(panmap[ibeg:iend], axis=1)>0)[0] + ibeg
        with open(f"MBE/{i}.pickle", 'wb') as f:
            pickle.dump([beds[:,ibeg:iend,:], indices, ibeg], f)

def load_process_pickle(i):
    with open(f"MBE/{i}.pickle", 'rb') as f:
        obj = pickle.load(f)
    return obj

def get_ctg(fas, hi, hd):
    assert hd in hd2is[hi], f"{hi}, {gs[hi//2]}, {hi%2}, {hd}"
    i = hd2is[hi][hd]
    L, s, w = fais[hi][i]
    e = s + (L-1)//w + 1 + L
    fas[hi].seek(s, 0)
    return fas[hi].read(e-s).decode("utf-8").replace("\n","").upper(), L, s, e

def get_seq_pos(fas, beds, ctgs, hds, idx, ibeg):
    seqs, poss = [None] * nh, [None] * nh
    for hi in range(nh):
        hd = beds[hi,idx-ibeg,0]
        if hd is None: continue
        if hd != hds[hi]:
            hds[hi] = hd
            with locks[hi]:
                ctgs[hi], L, s, e = get_ctg(fas, hi, hd)
                if len(ctgs[hi]) != L:
                    raise ValueError(f"worker {ibeg//bsize}: {idx}.{hi} ctg actually len {len(ctgs[hi])} != theoretical len {L}; {fas[hi].tell()}, {s}, {e}")
        s, e = [int(v) for v in beds[hi,idx-ibeg,[1,2]]]
        assert s < e
        ns = s - TRWINDOW if s > TRWINDOW                 else 0
        ne = e + TRWINDOW if e + TRWINDOW < len(ctgs[hi]) else len(ctgs[hi])
        seqs[hi] = ctgs[hi][ns:ne]
        poss[hi] = (s-ns, e-ns) # start pos of TR relative to the left end of NTR
    return seqs, poss

def multipleBoundaryExpansion(seqs, poss, idx, ibeg, UB, ksize=21):
    trs = set()
    npos = poss.copy()
    vi = [hi for hi, v in enumerate(poss) if v is not None] # valid indices
    for hi in vi:
        seq = seqs[hi]
        s, e = npos[hi]
        tmp = read2kmers(seq, ksize, leftflank=s, rightflank=len(seq)-e)
        for kmer in tmp: # TR
            trs.add(kmer)
    
    # seq,dt can have distinct orientations
    # force kmers/noise to have the same orientation
    exp = False
    dt = np.zeros([nh,2], dtype=int) + FS # hap, delta start/end; initialized with flanksize
    kmers = np.full([nh, 2, FS], -1, dtype=int)
    fail = [False]*nh
    while True:
        noise = np.zeros([nh,2,FS], dtype=int) # hap, left/right flank, flank pos 
        for hi in vi:
            if fail[hi]:
                continue
            #inv = beds[hi,idx-ibeg,3] == "-"
            seq = seqs[hi]
            sl = len(seq)
            s, e = npos[hi]
            lf = [s-FS                    , e+FS-dt[hi,1]-ksize+1] # left flank of [left,right] delta_flank
            rf = [sl-s+FS-dt[hi,0]-ksize+1, sl-e-FS              ] # right flank of [left,right] delta_flank
            #d = -1 if inv else 1
            if lf[0] < 0 or rf[1] < 0: # indicate TR near breakpoint
                #print(f"{idx}.{hi}.m ",end="")
                fail[hi] = True
                continue
            assert lf[0] >= 0 and lf[1] >= 0 and rf[0] >= 0 and rf[1] >= 0, print(hi, idx, s, e, sl)
            for sfl in [0,1]: # seq left, right flank
                #kfl = -sfl-1 if inv else sfl # kmer flank
                if dt[hi,sfl]:
                    if sfl == 0: # (no-inv, left flank) or (inv, right flank)
                        ns = dt[hi,sfl]
                        ne = FS
                        os = 0
                        oe = FS-dt[hi,sfl]
                    else:
                        ns = 0
                        ne = FS-dt[hi,sfl]
                        os = dt[hi,sfl]
                        oe = FS
                    kmers[hi,sfl,ns:ne] = kmers[hi,sfl,os:oe]
                    if sfl == 0:
                        ns = 0 
                        ne = dt[hi,sfl]
                    else:
                        ns = FS-dt[hi,sfl]
                        ne = FS
                    kmers[hi,sfl,ns:ne] = read2kmers(seq, ksize, leftflank=lf[sfl], rightflank=rf[sfl])
                for ki, kmer in enumerate(kmers[hi,sfl]): # recheck even if not expanded
                    if kmer in trs:
                        noise[hi,sfl,ki] = 1
        if not np.any(noise) or np.all(fail):
            break
        else:
            exp = True
            dt = np.zeros([nh,2], dtype=int)
            for hi in vi:
                if fail[hi]:
                    continue
                #inv = beds[hi,idx-ibeg,3] == "-"
                if np.any(noise[hi,0]):
                    dt[hi,0] = FS - np.nonzero(noise[hi,0])[0][0]
                    for kmer in kmers[hi,0,-dt[hi,0]:]:
                        trs.add(kmer)
                if np.any(noise[hi,1]):
                    dt[hi,1] = np.nonzero(noise[hi,1])[0][-1] + 1
                    for kmer in kmers[hi,1,:dt[hi,1]]:
                        trs.add(kmer)
                #if inv:
                #    dt = dt[:,::-1]
                if np.any(noise[hi]):
                    npos[hi] = (npos[hi][0]-dt[hi,0], npos[hi][1]+dt[hi,1])
                    if poss[hi][0] - npos[hi][0] > UB or npos[hi][1] - poss[hi][1] > UB:
                        fail[hi] = True
                        #print(f"{idx}.{hi}.x ", end="")
            if np.all(fail):
                print(f"{idx}.X ", flush=True)
                break
    return exp, fail, npos               
            
def load_fais():
    fafns = [f"{g}.{h}.fa" for g in gs for h in [0,1]]
    locks = [Lock() for i in range(nh)]
    fais, hd2is = [], []
    for f in fafns:
        fais.append(np.loadtxt(f"{INDIR}/{f}.fai", dtype=object, ndmin=2, comments=None))
        d = {}
        for i, hd in enumerate(fais[-1][:,0]):
            d[hd] = i
        hd2is.append(d)
        fais[-1] = fais[-1][:,[1,2,3]].astype(int)
    return fais, hd2is, locks

def load_fas():
    return [open(f"{INDIR}/{g}.{h}.fa", 'rb') for g in gs for h in [0,1]]

def close_fas(fas):
    for fa in fas:
        fa.close()

def gwMBE(batch, stat):
    fas = load_fas()
    beds, indices, ibeg = load_process_pickle(batch) # partial table; full_table_index = partial_table_index + ibeg
    out = []
    ctgs, hds = [""]*nh, [""]*nh # contigs, headers
    for idx in indices:
        stat[0] += 1
        seqs, poss = get_seq_pos(fas, beds, ctgs, hds, idx, ibeg)
        expanded, failed, npos = multipleBoundaryExpansion(seqs, poss, idx, ibeg, TRWINDOW-FS, ksize=KSIZE)
        es = []
        if expanded:
            stat[1] += 1
            if not np.all(failed):
                for hi in [i for i, v in enumerate(npos) if v is not None]:
                    ops = poss[hi]
                    nps = npos[hi]
                    es.append(ops[0]-nps[0]+nps[1]-ops[1])
                es = np.average(es)
            else:
                stat[2] += 1
                es = -1
        else:
            es = 0
        out.append([idx, expStat(expanded, np.nonzero(failed)[0].tolist(), es, poss, npos)])
        if stat[0] % max(nloci//200, 1) == 0:
            print(f"n={stat[0]}({stat[0]/nloci:.1%}), n_expanded={stat[1]}, n_fail={stat[2]}", flush=True)
    close_fas(fas)
    return out

def writeBed_MBE(th1=0.1, th2=0.8):
    """
    th1: maximal fraction of haps failing MBE
    th2: minimal fraction of haps remaining
    """
    panmap = np.loadtxt(sys.argv[5], dtype=object, ndmin=2)[:,3:].astype(int)
    bs = set() # bad set
    for idx, expstat in idx2exp.items():
        if expstat.exp:
            if len(expstat.fail) == nh:
                bs.add(idx)
            else:
                ns = np.sum([v is not None for v in expstat.npos])
                nf = len(expstat.fail)
                #print(idx,nf/ns)
                if nf/ns > th1:
                    bs.add(idx)
    ns = ng * th2
    bs |= set(np.nonzero(np.sum(panmap, axis=1)<ns)[0].tolist())
    vi = sorted(list(set(range(nloci))-bs)) # valid indices
    np.savetxt("locusMap.v1.to.v0.txt", vi, fmt='%i')

    panbed = np.full([nloci, 3+nh*4], None, dtype=object)
    panbed[:,:3] = np.loadtxt(f"pan.tr.mbe.v0.bed", usecols=[0,1,2], dtype=object)
    for hi in range(nh):
        g = gs[hi//2]
        h = hi % 2
        bed = np.loadtxt(f"{g}/tmp1.{h}.bed", dtype=object, ndmin=2, comments=None) # iterate w/ genome index
        p2g = np.full(nloci, None, dtype=object) # map pan index to genome index
        p2g[panmap[:,hi//2]==1] = np.arange(bed.shape[0])
        f = open(f"{g}/tmp2.{h}.mbe.bed", 'w')
        for pid in vi:
            if idx2exp[pid].opos[hi] is None: # missing hap
                continue
            if hi in idx2exp[pid].fail: # MBE failed at this hap
                panbed[pid,3+hi*4:7+hi*4] = [None, None, None, None]
                continue
            gid = p2g[pid]
            os, oe = idx2exp[pid].opos[hi]
            ns, ne = idx2exp[pid].npos[hi]
            dt = [os-ns, ne-oe]
            s, e = int(bed[gid,1]), int(bed[gid,2])
            s -= dt[0]
            e += dt[1]
            f.write(f"{bed[gid,0]}\t{s}\t{e}\t{bed[gid,3]}\t{bed[gid,4]}\t{bed[gid,5]}\t{bed[gid,6]}\n")
            panbed[pid,3+hi*4:7+hi*4] = [bed[gid,0], s, e, bed[gid,6]]
        f.close()
    np.savetxt("pan.tr.mbe.v1.bed", panbed[vi], delimiter="\t", fmt='%s')

if __name__ == "__main__":
    KSIZE, FS, TRWINDOW = [int(sys.argv[i]) for i in range(1,4)]
    NPROCESS = int(sys.argv[8])
    INDIR = sys.argv[9]
    print("Loading metadata", flush=True)
    gs = np.loadtxt(sys.argv[4], usecols=0, dtype=object, ndmin=1)
    print("\tpanmap", flush=True)
    panmap = np.loadtxt(sys.argv[5], dtype=object, ndmin=2)[:,3:].astype(int)
    nloci, ng = panmap.shape
    nh = 2 * ng
    bsize = (nloci-1) // NPROCESS + 1
    print("\tbed's", flush=True, end="")
    beds = loadbeds(panmap)
    print("\n\tfasta's", flush=True)
    fais, hd2is, locks = load_fais()
    print("Making pickle for each process")
    make_process_pickle()
    del panmap
    del beds

    print("Running multi-boundary expansion", flush=True)
    os.system(f"taskset -p 0xffffffff {os.getpid()}")
    stat = Manager().list([0,0,0]) # ncase, nexp, nfail
    idx2exp = {}
    with Pool(NPROCESS) as pool:
        results = []
        for i in range(NPROCESS):
            results.append(pool.apply_async(gwMBE, args=(i,stat)))
        ps = set(list(range(NPROCESS)))
        while ps:
            done = []
            for i in ps:
                if results[i].ready():
                    print(f"[multiprocessing.Pool] worker {i} ended", flush=True)
                    for k, v in results[i].get():
                        idx2exp[k] = v
                    done.append(i)
            for i in done:
                ps.remove(i)
            sleep(5)
    print("Parallel computing results merged", flush=True)

    print("Dumping results", flush=True)
    with open("MBE/idx2exp.mbe.pickle", 'wb') as f:
        pickle.dump(idx2exp, f, protocol=pickle.HIGHEST_PROTOCOL)
    #with open("MBE/idx2exp.mbe.pickle", 'rb') as f:
    #    idx2exp = pickle.load(f)
    print("Writing new bed regions", flush=True)
    writeBed_MBE(float(sys.argv[6]), float(sys.argv[7]))
    print("Done", flush=True)
```
