## Supplemental code for "The motif composition of variable-number tandem repeats impacts gene expression": multiBoundaryExpansion.py

```
#!/usr/bin/env python3

import sys
import numpy as np
import vntrutils as vu
import pickle

class expStat:
    def __init__(self, exp, fail, es, opos, npos):
        self.exp = exp
        self.fail = fail
        self.es = es
        self.opos = opos
        self.npos = npos
    def __repr__(self):
        return f"expanded: {self.exp}\tfail: {self.fail}\tave. exp. size: {self.es:.0f}\nopos: {self.opos}\tnpos: {self.npos}\n"

def loadbeds(panmap):
    nloci, ng = panmap.shape
    beds = np.full([2*ng, nloci, 4], None, dtype=object)
    for gi, g in enumerate(gs):
        for h in [0,1]:
            hi = 2*gi + h
            m = panmap[:,gi]==1
            bed = np.loadtxt(f"{g}/tmp1.{h}.bed", dtype=object, usecols=[0,1,2,6], ndmin=2, comments=None)
            assert np.sum(m) == bed.shape[0], f"Inconsistent # of supports between {g}/tmp1.{h}.bed ({bed.shape[0]}) and column {gi+3} of pan.tr.mbe.v0.bed (np.sum(m))"
            beds[hi,m] = bed
    assert set(beds[:,:,-1].flatten().tolist()) == set(["-1","1",None]), print(set(beds[:,:,-1].flatten().tolist()) - set(["-1","1",None]))
    return beds

def get_ctg(hi, hd):
    i = hd2is[hi][hd]
    L, s, w = fais[hi][i]
    e = s + (L-1)//w + 1 + L
    fas[hi].seek(s)
    return fas[hi].read(e-s).decode("utf-8").replace("\n","")

def get_seq_pos(ctgs, hds, idx):
    nh = gs.size * 2
    seqs, poss = [None] * nh, [None] * nh
    for hi in range(nh):
        hd = beds[hi,idx,0]
        if hd is None: continue
        if hd != hds[hi]:
            hds[hi] = hd
            ctgs[hi] = get_ctg(hi, hd)
        s, e = [int(v) for v in beds[hi,idx,[1,2]]]
        assert s < e
        ns = s - TRWINDOW if s > TRWINDOW                 else 0
        ne = e + TRWINDOW if e + TRWINDOW < len(ctgs[hi]) else len(ctgs[hi])
        seqs[hi] = ctgs[hi][ns:ne]
        poss[hi] = (s-ns, e-ns) # start pos of TR relative to the left end of NTR
    return seqs, poss

def multipleBoundaryExpansion(seqs, poss, idx, UB, ksize=21):
    ng = gs.size
    trs = set()
    npos = poss.copy()
    vi = [hi for hi, v in enumerate(poss) if v is not None] # valid indices
    for hi in vi:
        seq = seqs[hi]
        s, e = npos[hi]
        tmp = vu.read2kmers(seq, ksize, leftflank=s, rightflank=len(seq)-e)
        for kmer in tmp: # TR
            trs.add(kmer)
    
    # seq,dt can have distinct orientations
    # force kmers/noise to have the same orientation
    exp = False
    dt = np.zeros([2*ng,2], dtype=int) + FS # hap, delta start/end; initialized with flanksize
    kmers = np.full([2*ng, 2, FS], -1, dtype=int)
    fail = np.zeros(2*ng, dtype=bool)
    while True:
        noise = np.zeros([2*ng,2,FS], dtype=int) # hap, left/right flank, flank pos 
        for hi in vi:
            if fail[hi]:
                continue
            inv = beds[hi,idx,3] == "-"
            seq = seqs[hi]
            sl = len(seq)
            s, e = npos[hi]
            lf = [s-FS                    , e+FS-dt[hi,1]-ksize+1] # left flank of [left,right] flank
            rf = [sl-s+FS-dt[hi,0]-ksize+1, sl-e-FS              ] # right flank of [left,right] flank
            d = -1 if inv else 1
            if lf[0] < 0 or rf[1] > sl: # indicate TR near breakpoint
                print(f"{idx}.{hi}.m ",end="")
                fail[hi] = True
                continue
            for sfl in [0,1]: # seq left, right flank
                kfl = -sfl-1 if inv else sfl # kmer flank
                if dt[hi,sfl]:
                    if int(inv) == sfl: # (no-inv, left flank) or (inv, right flank)
                        ns = dt[hi,sfl]
                        ne = FS
                        os = 0
                        oe = FS-dt[hi,sfl]
                    else:
                        ns = 0
                        ne = FS-dt[hi,sfl]
                        os = dt[hi,sfl]
                        oe = FS
                    kmers[hi,kfl,ns:ne] = kmers[hi,kfl,os:oe]
                    if int(inv) == sfl:
                        ns = 0 
                        ne = dt[hi,sfl]
                    else:
                        ns = FS-dt[hi,sfl]
                        ne = FS
                    kmers[hi,kfl,ns:ne:d] = vu.read2kmers(seq, ksize, leftflank=lf[kfl], rightflank=rf[kfl])
                for ki, kmer in enumerate(kmers[hi,kfl]): # recheck even if not expanded
                    if kmer in trs:
                        noise[hi,kfl,ki] = 1
        if not np.any(noise) or np.all(fail):
            break
        else:
            exp = True
            dt = np.zeros([2*ng,2], dtype=int)
            for hi in vi:
                if fail[hi]:
                    continue
                inv = beds[hi,idx,3] == "-"
                if np.any(noise[hi,0]):
                    dt[hi,0] = FS - np.nonzero(noise[hi,0])[0][0]
                    for kmer in kmers[hi,0,-dt[hi,0]:]:
                        trs.add(kmer)
                if np.any(noise[hi,1]):
                    dt[hi,1] = np.nonzero(noise[hi,1])[0][-1] + 1
                    for kmer in kmers[hi,1,:dt[hi,1]]:
                        trs.add(kmer)
                if inv:
                    dt = dt[:,::-1]
                if np.any(noise[hi]):
                    npos[hi] = (npos[hi][0]-dt[hi,0], npos[hi][1]+dt[hi,1])
                    if poss[hi][0] - npos[hi][0] > UB or npos[hi][1] - poss[hi][1] > UB:
                        fail[hi] = True
                        print(f"{idx}.{hi}.x ", end="")
            if np.all(fail):
                print(f"{idx}.X ", end="")
                break
    return exp, fail, npos               
            
def gwMBE(target=None):
    nloci, ng = panmap.shape
    out = {}
    indices = np.nonzero(np.sum(panmap, axis=1)>0)[0] if target is None else target
    ctgs, hds = [""]*2*ng, [""]*2*ng # contigs, headers
    ncase, nexp, nfail = 0, 0, 0
    for idx in indices:
        ncase += 1
        seqs, poss = get_seq_pos(ctgs, hds, idx)
        expanded, failed, npos = multipleBoundaryExpansion(seqs, poss, idx, TRWINDOW-FS, ksize=KSIZE)
        es = []
        if expanded:
            nexp += 1
            if not np.all(failed):
                for hi in [i for i, v in enumerate(npos) if v is not None]:
                    ops = poss[hi]
                    nps = npos[hi]
                    es.append(ops[0]-nps[0]+nps[1]-ops[1])
                es = np.average(es)
            else:
                nfail += 1
                es = -1
        else:
            es = 0
        out[idx] = expStat(expanded, failed, es, poss, npos)
        if ncase % 1000 == 0:
            print(f"\nn={ncase}, n_expanded={nexp}, n_fail={nfail}")
    return out

def load_fais():
    fafns = [f"{g}.{h}.fa" for g in gs for h in [0,1]]
    fas, fais, hd2is = [], [], []
    for f in fafns:
        fas.append(open(f, 'rb'))
        fais.append(np.loadtxt(f"{f}.fai", dtype=object, ndmin=2, comments=None))
        d = {}
        for i, hd in enumerate(fais[-1][:,0]):
            d[hd] = i
        hd2is.append(d)
        fais[-1] = fais[-1][:,[1,2,3]].astype(int)
    return fas, fais, hd2is

def close_fas():
    for fa in fas:
        fa.close()

def writeBed_MBE(th1=0.1, th2=0.8):
    nloci, ng = panmap.shape
    bs = set() # bad set
    for idx, expstat in idx2exp.items():
        if expstat.exp:
            if np.all(expstat.fail):
                bs.add(idx)
            else:
                ns = np.sum([v is not None for v in expstat.npos])
                nf = np.sum(expstat.fail)
                #print(idx,nf/ns)
                if nf/ns > th1:
                    bs.add(idx)
    ns = panmap.shape[1] * th2
    bs |= set(np.nonzero(np.sum(panmap, axis=1)<ns)[0].tolist())
    vi = sorted(list(set(range(nloci))-bs)) # valid indices
    np.savetxt("locusMap.v1.to.v0.txt", vi, fmt='%i')

    panbed = np.full([nloci, 3+2*ng*4], None, dtype=object)
    panbed[:,:3] = np.loadtxt(f"pan.tr.mbe.v0.bed", usecols=[0,1,2], dtype=object)
    for hi in range(2*ng):
        g = gs[hi//2]
        h = hi % 2
        bed = np.loadtxt(f"{g}/tmp1.{h}.bed", dtype=object, ndmin=2, comments=None) # iterate w/ genome index
        p2g = np.full(nloci, None, dtype=object) # map pan index to genome index
        p2g[panmap[:,hi//2]==1] = np.arange(bed.shape[0])
        f = open(f"{g}/tmp2.{h}.mbe.bed", 'w')
        for pid in vi:
            if idx2exp[pid].opos[hi] is None: # missing hap
                continue
            if idx2exp[pid].fail[hi]: # MBE failed at this hap
                panbed[pid,3+hi*4:7+hi*4] = [None, None, None, None]
                continue
            gid = p2g[pid]
            os, oe = idx2exp[pid].opos[hi]
            ns, ne = idx2exp[pid].npos[hi]
            dt = [os-ns, ne-oe]
            s, e = int(bed[gid,1]), int(bed[gid,2])
            s -= dt[0]
            e += dt[1]
            f.write(f"{bed[gid,0]}\t{s}\t{e}\t{bed[gid,3]}\t{bed[gid,4]}\t{bed[gid,5]}\t{bed[gid,6]}\n")
            panbed[pid,3+hi*4:7+hi*4] = [bed[gid,0], s, e, bed[gid,6]]
        f.close()
    np.savetxt("pan.tr.mbe.v1.bed", panbed[vi], delimiter="\t", fmt='%s')

if __name__ == "__main__":
    KSIZE, FS, TRWINDOW = [int(sys.argv[i]) for i in range(1,4)]
    print("Loading metadata")
    gs = np.loadtxt(sys.argv[4], usecols=0, dtype=object, ndmin=1)
    panmap = np.loadtxt(sys.argv[5], dtype=object, ndmin=2)[:,3:].astype(int)
    beds = loadbeds(panmap)
    fas, fais, hd2is = load_fais()
    print("Running multi-boundary expansion")
    idx2exp = gwMBE()
    print("\nDumping results")
    with open("idx2exp.mbe.pickle", 'wb') as f:
        pickle.dump(idx2exp, f)
    close_fas()
    print("Writing new bed regions")
    writeBed_MBE(float(sys.argv[6]), float(sys.argv[7]))
    print("Done")
```
