## Supplemental code for "The motif composition of variable-number tandem repeats impacts gene expression": PlotRegression.py

```
#!/usr/bin/env python3

import argparse
import vntrutils as vu
import numpy as np

#from scipy.optimize import minimize
#from scipy.misc import factorial

ap = argparse.ArgumentParser(description="plot the correlation given two files")
ap.add_argument("xFile", help="x as a single column of data")
ap.add_argument("yFile", help="y as a single column of data")
args = ap.parse_args()

x = np.loadtxt(args.xFile).reshape(-1,1)
y = np.loadtxt(args.yFile).reshape(-1,1)

vu.PlotRegression(x, y, args.xFile, args.yFile, fname=args.xFile+"."+args.yFile)
```
