## Supplemental code for "The motif composition of variable-number tandem repeats impacts gene expression": prepareIndividualDatasets.py

```
#!/usr/bin/env python3

import sys
import numpy as np
import pickle

nprocess, KSIZE, FS, UB, TRWINDOW, nloci = [int(v) for v in sys.argv[1:7]]
fastafiles = sys.argv[7:9]
beds = [np.loadtxt(bed, dtype=object, ndmin=2) for bed in sys.argv[9:11]]

def getName2loci(beds):
    name2loci = [{}, {}]
    for hap in range(2):
        for locus in range(nloci):
            name = beds[hap][locus,0]
            if name != "NA":
                if name not in name2loci[hap]:
                    name2loci[hap][name] = []
                name2loci[hap][name].append(locus)
    return name2loci

def getLociSeqFromCtg(hap, ctg, locus, beds, window=TRWINDOW):
    start = int(beds[hap][locus,1])
    end = int(beds[hap][locus,2])
    assert start < end
    start = start-window if start > window else 0
    end = end+window if end+window <= len(ctg) else len(ctg)
    return ctg[start:end]

def getLociPosFromCtg(hap, ctg, locus, beds, window=TRWINDOW):
    # TODO: this includes 1k flanking seq, although will be rescued later
    start = int(beds[hap][locus,1])
    end = int(beds[hap][locus,2])
    assert start < end
    NTRstart = start-window if start > window else 0
    return (start-NTRstart, end-NTRstart) # start pos of TR relative to the left end of NTR

def getLociSeqPos(fastafiles, beds, name2loci):
    lociSeqs = [{}, {}] # {locus : seq}
    lociPoss = [{}, {}] # {locus : (start,end)}
    for hap, fastafile in enumerate(fastafiles):
        print("hap ",hap)
        with open(fastafile) as f:
            ctg = ""
            getctg = False

            for line in f:
                if line[0] == '>':
                    if ctg:
                        for locus in name2loci[hap][name]:
                            lociSeqs[hap][locus] = getLociSeqFromCtg(hap, ctg, locus, beds)
                            lociPoss[hap][locus] = getLociPosFromCtg(hap, ctg, locus, beds)
                        ctg = ""
                        getctg = False

                    name = line.split()[0][1:]
                    if name in name2loci[hap]:
                        getctg = True
                else:
                    if getctg: ctg += line.rstrip()
            if ctg:
                for locus in name2loci[hap][name]:
                    lociSeqs[hap][locus] = getLociSeqFromCtg(hap, ctg, locus, beds)
                    lociPoss[hap][locus] = getLociPosFromCtg(hap, ctg, locus, beds)
    return lociSeqs, lociPoss

def prepareMinibatchData(seqs, poss):
    print("preparing minibatch datasets")
    for batch in range(nprocess):
        miniseqs, miniposs = [{}, {}], [{}, {}]
        for hap in [0,1]:
            for locus in range(nloci):
                if locus % nprocess == batch:
                    if locus in poss[hap]:
                        miniseqs[hap][locus] = seqs[hap][locus]
                        miniposs[hap][locus] = poss[hap][locus]
        pickle.dump([miniseqs, miniposs], open("seqpos{}.pickle".format(batch), 'wb'))

name2loci = getName2loci(beds)
seqs, poss = getLociSeqPos(fastafiles, beds, name2loci)
pickle.dump([seqs, poss], open("seqpos.pickle",'wb'))
prepareMinibatchData(seqs, poss)
```
