## Supplemental code for "The motif composition of variable-number tandem repeats impacts gene expression": prepareJointDatasets.py

```
#!/usr/bin/env python3


import sys
import numpy as np
import pickle

nprocess, KSIZE, FS, UB, TRWINDOW, nloci = [int(v) for v in sys.argv[1:7]]
[seqs, poss] = pickle.load(open("seqpos.pickle", 'rb'))
[_, newposs] = pickle.load(open("newposs_stat.pickle", 'rb'))

def prepareMinibatchData(seqs, poss, newposs):
    print("preparing minibatch datasets")
    for batch in range(nprocess):
        miniseqs, mininewposs, miniposs= [{}, {}], [{}, {}], [{}, {}]
        for hap in [0,1]:
            for locus in range(nloci):
                if locus % nprocess == batch:
                    if locus in poss[hap]:
                        miniseqs[hap][locus] = seqs[hap][locus]
                        mininewposs[hap][locus] = newposs[hap][locus]
                        miniposs[hap][locus] = poss[hap][locus]
        pickle.dump([miniseqs, mininewposs, miniposs], open("seqnewpos{}.pickle".format(batch), 'wb'))

prepareMinibatchData(seqs, poss, newposs)
```
