## Supplemental code for "The motif composition of variable-number tandem repeats impacts gene expression": rmLinebyIndFile.py

```
#!/usr/bin/env python3

import sys
import numpy as np

if len(sys.argv) == 1 or sys.argv[1] == "-h" or sys.argv[1] == "--help":
    print(
    """
    Remove line indices (0-based) specified in 'index.txt'
    usage: program [-k] index.txt inFile
           -k       Keep line indices in 'index.txt' instead of removing them.
    """)
    sys.exit()

rm = True
idxf = ""
infile = "" 
for i, v in enumerate(sys.argv):
    if i == 0:
        continue
    elif v == "-k":
        rm = False
    elif not idxf:
        idxf = v
    elif not infile:
        infile = v
    else:
        assert False, f"too many arguments {v}"
if not idxf:
    assert False, "index.txt not specified"
if not infile:
    assert False, "inFile not specified"

ids = set(np.loadtxt(idxf, dtype=int, ndmin=1).tolist())
with open(infile) as f:
    ind = 0
    for line in f:
        if (ind not in ids) == rm:
            print(line, end='')
        ind += 1
```
