## Supplemental code for "The motif composition of variable-number tandem repeats impacts gene expression": rmNAforBothBeds.py

```
#!/usr/bin/env python3

import sys

if len(sys.argv) != 5:
    print("usage: program in.bed0 in.bed1 out.bed0 out.bed1")
    exit

in0, in1, out0, out1 = sys.argv[1:]
badids = set()
with open(in0) as f:
    for ids, line in enumerate(f):
        if line[:2] == "NA":
            badids.add(ids)
with open(in1) as f:
    for ids, line in enumerate(f):
        if line[:2] == "NA":
            badids.add(ids)

with open(in0) as f:
    with open(out0, 'w') as fo:
        for ids, line in enumerate(f):
            if ids not in badids:
                fo.write(line)
with open(in1) as f:
    with open(out1, 'w') as fo:
        for ids, line in enumerate(f):
            if ids not in badids:
                fo.write(line)
```
