## Supplemental code for "The motif composition of variable-number tandem repeats impacts gene expression": SelectRegions.py

```
#!/usr/bin/env python
import pysam
import sys

if len(sys.argv) != 4:
    print("usage: program region.bed fasta region.fasta")
    sys.exit(1)

bedFile=open(sys.argv[1])
#asmFile=open(sys.argv[2])
asm = pysam.FastaFile(sys.argv[2])
outFile=open(sys.argv[3],'w')


for line in bedFile:
    vals = line.split()
    outFile.write(">"+":".join(vals)+"\n")
    if vals[0] == "NA": # NF does not have to be 6
        continue
    elif int(vals[1]) > int(vals[2]) or int(vals[1]) < 0 or int(vals[2]) < 0: # XXX use chrsize info, instead of only checking start pos
        # XXX print empty seq or not
        print("valError:\t", vals, file=sys.stderr)
        continue
    else:
#        sys.stderr.write("/".join(vals[0:3])+"\n")
        seq = asm.fetch(vals[0], int(vals[1]), int(vals[2]))
        outFile.write(seq + "\n")
```
