## Supplemental code for "The motif composition of variable-number tandem repeats impacts gene expression": writeBEbed.py

```
#!/usr/bin/env python3
import sys
import numpy as np
import pickle

def loadBE():
    BE = np.full(ncore, None, dtype=object)
    for i in range(ncore):
        with open(f"seqjointnewpos{i}.pickle", 'rb') as f:
            BE[i] = pickle.load(f) # seqs, jointnewposs, poss
    return BE

def loadQC():
    with open("QCresults.pickle", 'rb') as f:
        QCbadloci, QCunresolvedloci = pickle.load(f)
    return QCbadloci, QCunresolvedloci

def writeBEbeds():
    nloci = np.loadtxt(sys.argv[2], usecols=1).shape[0]
    beds = [np.loadtxt(sys.argv[h+2], dtype=object) for h in [0,1]]
    nbeds = [np.full([nloci,7], None, dtype=object) for h in [0,1]]
    for i in range(nloci):
        if i in QCbad:
            continue
        b = i%ncore
        rs = BEresults[b]
        for h in [0,1]:
            s, e = int(beds[h][i,1]), int(int(beds[h][i,2]))
            os, oe = rs[2][h][i]
            ns, ne = rs[1][h][i]
            e5, e3 = os - ns, ne - oe
            nbeds[h][i,0] = beds[h][i,0]
            nbeds[h][i,1] = s - e5
            nbeds[h][i,2] = e + e3
            nbeds[h][i,3] = beds[h][i,3]
            nbeds[h][i,4] = beds[h][i,4]
            nbeds[h][i,5] = beds[h][i,5]
            nbeds[h][i,6] = beds[h][i,6]
    for h in [0,1]:
        m = ~np.all(nbeds[h] == None, axis=1)
        np.savetxt(sys.argv[h+4], nbeds[h][m], fmt="%s", delimiter="\t")
            
ncore = int(sys.argv[1])
BEresults = loadBE()
QCbadloci, QCunresolvedloci = loadQC()
QCbad = set(list(QCbadloci.keys())) | QCunresolvedloci
writeBEbeds()
```
