## Supplemental code for "The motif composition of variable-number tandem repeats impacts gene expression": writeBoundaryExpandedBeds.py

```
#!/usr/bin/env python3

import sys
import numpy as np
import pickle

nprocess, KSIZE, FS, UB, TRWINDOW, nloci = [int(v) for v in sys.argv[1:7]]
beds = [np.loadtxt(bed, dtype=object, ndmin=2) for bed in sys.argv[7:9]]
chrsizefnames = sys.argv[9:11]
outbednames = sys.argv[11:13]


def loadchrsize(fnames):
    chrsizes = np.empty(len(fnames), dtype=object)
    for hap, fname in enumerate(fnames):
        chrsizes[hap] = {}
        with open(fname) as f:
            for line in f:
                fields = line.split()
                chrsizes[hap][fields[0]] = int(fields[1])
    return chrsizes

def filterInvalidLociSaveNewBed(beds, chrsizefnames, outbednames):
    """filter out loci within FS bp to breakpoints"""
    chrsizes = loadchrsize(chrsizefnames)
    for hap in [0,1]:
        for rind, row in enumerate(beds[hap]):
            cs = chrsizes[hap][row[0]]
            if row[1] < FS or row[2]+FS > cs:
                beds[hap][rind][0] = "NA"
        np.savetxt(outbednames[hap], beds[hap], fmt='%s', delimiter='\t')

def merge_full_overlap_region(srtbed):
    """ 
    field 2, 3 have to be ints 
    field 5, 6 have to be strings
    should not change reference contents, otherwise will affect mapping betw h0/h1 based on reference ordering
    """
    
    outbed = np.copy(srtbed)
    outbed[:,4:] = outbed[:,4:].astype(int)
    nloci = outbed.shape[0]
    non_overlap_mask = np.ones(nloci, dtype=bool)
    full_overlap_mask = np.zeros(nloci, dtype=bool)
    full_overlap_set = np.zeros(full_overlap_mask.size, dtype=int)
    overlap_counter = 0
    ind0, ind2 = 0, 0
    fulloverlap = True
    for ind1 in range(1, nloci):
        if ind1 < ind2: continue # skip loci already processed
        
        if outbed[ind0,0] != outbed[ind1,0]: # check contig name the same
            ind0 = ind1
        else:
            if outbed[ind1,1] > outbed[ind0,2]: # non-overlapping
                ind0 = ind1
            else:
                # find the right end of regions with the same start
                ind2 = ind1
                while outbed[ind0,1] == outbed[ind2,1] and outbed[ind0,0] == outbed[ind2,0]:
                    ind2 += 1
                    if ind2 == nloci: break
                e2 = np.amax(outbed[ind0:ind2,2])

                # check if overlapping regions are fully overlapping
                ind2 = ind1
                while outbed[ind2,2] <= e2 and outbed[ind0,0] == outbed[ind2,0]:
                    ind2 += 1
                    if ind2 >= nloci: break
                refs2 = np.amin(outbed[ind0:ind2,4])
                refe2 = np.amax(outbed[ind0:ind2,5])
                #print(refs2, refe2, outbed[ind0:ind2,4:])
                        
                if ind2 == nloci or outbed[ind0,0] != outbed[ind2,0]:
                    fulloverlap = True
                else:
                    fulloverlap = False if outbed[ind2,1] <= outbed[ind0,2] else True

                if fulloverlap:
                    # set locus ind0 as the full region, output locus ind0; discard loci [ind1:ind2)
                    overlap_counter += 1
                    full_overlap_set[ind0:ind2] = overlap_counter
                    full_overlap_mask[ind0] = True
                    non_overlap_mask[ind0:ind2] = False
                else:
                    # find all partial overlapping loci
                    while outbed[ind2,1] <= outbed[ind0,2] and outbed[ind0,0] == outbed[ind2,0]:
                        ind2 += 1
                        if ind2 >= nloci: break
                    non_overlap_mask[ind0:ind2] = False
                    
    print(np.sum(non_overlap_mask), np.sum(full_overlap_mask))
    return outbed, non_overlap_mask, full_overlap_mask, full_overlap_set

def writeNewBed(jointnewlociPoss, lociPoss, beds, QCbadloci, QCunresolvedloci):
    # remove loci failed the expansion steps
    goodmask_ref = np.ones(nloci, dtype=bool)
    for badlocus in QCbadloci.keys(): goodmask_ref[badlocus] = False
    for badlocus in QCunresolvedloci: goodmask_ref[badlocus] = False
    
    # record regions changed after expansion
    fixmask_ref = np.zeros(nloci, dtype=bool)
    for i in range(nloci):
        if i not in lociPoss[0]: continue
        fixmask_ref[i] = lociPoss[0][i] != jointnewlociPoss[0][i] or lociPoss[1][i] != jointnewlociPoss[1][i]
    fixmask_ref = fixmask_ref[goodmask_ref]

    outbeds_ass = [None, None]
    non_overlap_masks_ass = [None, None]
    full_overlap_masks_ass = [None, None]
    full_overlap_sets_ass = [None, None]
    ords_ref = [None, None]
    for hap in [0,1]:
        newbed_ref = np.copy(beds[hap])
        
        # compute new regions
        for locus in range(nloci):
            if goodmask_ref[locus]:
                pos = jointnewlociPoss[hap][locus]
                oldpos = lociPoss[hap][locus]
                old = beds[hap][locus]
                Lexp, Rexp = oldpos[0] - pos[0], pos[1] - oldpos[1]
                newbed_ref[locus,1:3] = int(old[1]) - Lexp, int(old[2]) + Rexp

        # sort based on assembly order
        # for mergin intervals
        newbed_ref = newbed_ref[goodmask_ref]
        ord_ass = np.lexsort((newbed_ref[:,2], newbed_ref[:,1], newbed_ref[:,0]))
        newbed_ass = newbed_ref[ord_ass]
        
        outbeds_ass[hap], non_overlap_masks_ass[hap], full_overlap_masks_ass[hap], full_overlap_sets_ass[hap] =\
                                                                        merge_full_overlap_region(newbed_ass)
        
        # sort based on ref order
        # for mapping loci between h0 and h1
        ords_ref[hap] = np.lexsort((outbeds_ass[hap][:,5], outbeds_ass[hap][:,4], outbeds_ass[hap][:,3]))
                
    outbeds_ref = [None, None]
    for hap in [0,1]:
        outbeds_ref[hap] = outbeds_ass[hap][ords_ref[hap]]
    
    # check merging consistency betw h0 and h1
    full_overlap_mask_ref = np.logical_and(full_overlap_masks_ass[0][ords_ref[0]], full_overlap_masks_ass[1][ords_ref[1]]) #XXX overlap set might differ
    full_overlap_sets_ref = [None, None]
    for hap in [0,1]:
        full_overlap_sets_ref[hap] = full_overlap_sets_ass[hap][ords_ref[hap]]
    total, good = 0, 0
    for i in range(1, np.amax(full_overlap_sets_ref)+1):
        mask0 = full_overlap_sets_ref[0] == i # loci in the same to-be-merged set in h0
        indices0 = np.nonzero(mask0)[0]
        total += np.sum(mask0)
        if not len(indices0): #XXX why zero
            mask1 = full_overlap_sets_ref[1] == i
            full_overlap_mask_ref[mask1] = False
        else:
            mask1 = full_overlap_sets_ref[1] == full_overlap_sets_ref[0][indices0[0]] # corresponding loci in h1
            if not np.all(mask0 == mask1):
                full_overlap_mask_ref[mask0] = False
                full_overlap_mask_ref[mask1] = False
            else:
                # check if duplicates are in the same chrom
                if np.any(np.concatenate((outbeds_ref[0][mask0,3], outbeds_ref[1][mask0,3])) != outbeds_ref[0][mask0,3][0]):
                    full_overlap_mask_ref[mask0] = False
                    full_overlap_mask_ref[mask1] = False
                else:
                    good += np.sum(mask0)
                    # consistent, update reference region
                    print(i, full_overlap_mask_ref[indices0], indices0, full_overlap_sets_ref[0][mask0], full_overlap_sets_ref[1][mask1])
                    ind = indices0[0]
                    masks = [mask0, mask1]
                    refs2 = np.amin(outbeds_ref[0][masks[0],4])
                    refe2 = np.amax(outbeds_ref[0][masks[0],5])
                    for hap in [0,1]:
                        e2 = np.amax(outbeds_ref[hap][masks[hap],2])
                        outbeds_ref[hap][ind,2] = e2
                        outbeds_ref[hap][ind,4:] = refs2, refe2
    
    print(total, good)
    non_overlap_mask_ref = np.logical_and(non_overlap_masks_ass[0][ords_ref[0]], non_overlap_masks_ass[1][ords_ref[1]])
    outmask_ref = np.logical_or(non_overlap_mask_ref, full_overlap_mask_ref)
    print(np.sum(full_overlap_mask_ref))
    
    completeoutbeds = [None, None]
    for hap in [0,1]:
        outbed_ref = outbeds_ref[hap][outmask_ref]
        outfixbit_ref = fixmask_ref[outmask_ref].astype(int)[:,None]
        outoverlapbit_ref = full_overlap_mask_ref[outmask_ref].astype(int)[:,None]
        completeoutbeds[hap] = np.hstack((outbed_ref, outfixbit_ref, outoverlapbit_ref))
        # correct ending pos considering kmer size = KSIZE
        completeoutbeds[hap][:,2] = completeoutbeds[hap][:,2]+(KSIZE-1)

    filterInvalidLociSaveNewBed(completeoutbeds, chrsizefnames, outbednames)

        
_, poss = pickle.load(open("pk.seqpos.dat", 'rb'))        
_, jointnewposs = pickle.load(open("pk.jointnewposs_stat.dat", 'rb'))
QCbadloci, QCunresolvedloci = pickle.load(open("pk.QCresults.dat", 'rb'))

writeNewBed(jointnewposs, poss, beds, QCbadloci, QCunresolvedloci)
```
