## Supplemental code for "The motif composition of variable-number tandem repeats impacts gene expression": aQueryFasta_thread.cpp

```
#include "aQueryFasta_thread.h"
//#include "/project/mchaisso_100/cmb-16/tsungyul/src/gperftools-2.9.1/src/gperftools/profiler.h"

#include <cstdlib>
#include <vector>
#include <string>
#include <iostream>
//#include <fstream>
#include <numeric>
#include <pthread.h>
#include <semaphore.h>
#include <fcntl.h>
#include <errno.h>
#include <ctime>
#include <cassert>
#include <cstring>
#include <algorithm>
#include <atomic>

using namespace std;

sem_t *semreader;
sem_t *semcount;
sem_t *semwriter;
bool testmode;
size_t ksize;
size_t N_FILTER = 4; // number of subsampled kmers
size_t NM_FILTER = 1; // minimal number of hits
int verbosity = 0;
const uint64_t NAN64 = 0xFFFFFFFFFFFFFFFF;
const uint32_t NAN32 = 0xFFFFFFFF;

typedef unordered_map<size_t, size_t> msa_umap; // dest_locus, counts
// src_locus -> {dest_locus -> [src_count, dest_count_uncorrected, dest_count_corrected]}
typedef unordered_map<size_t, unordered_map<size_t, std::tuple<size_t,size_t,size_t>>> err_umap;
typedef std::pair<uint8_t, uint8_t> PE_KMC; // pair-end kmer count // XXX not compatible with reads longer than 255 bp

struct EDIT {
	vector<char> ops1, ops2; // alignment operation for each end. M: match. S: skip. H: homopolymer. [ACGT]: edits.
	std::pair<size_t, size_t> map; // <src_locus, dest_locus>
};

struct asgn_t { // pe read assignment
	size_t idx = NAN32;
	size_t fc = 0, rc = 0;
};

void rand_str(char *dest, size_t length) {
    char charset[] = "0123456789"
                     "abcdefghijklmnopqrstuvwxyz"
                     "ABCDEFGHIJKLMNOPQRSTUVWXYZ";		
    while (length-- > 0) {
        size_t index = (double) rand() / RAND_MAX * (sizeof charset - 1);
        *dest++ = charset[index];
    }
    *dest = '\0';
}

bool subfilter(vector<size_t>& kmers1, vector<size_t>& kmers2, kmerIndex_uint32_umap& kmerDBi, size_t& nhash) {
	size_t L1 = kmers1.size(), L2 = kmers2.size();
	size_t S1 = L1 / (N_FILTER-1), S2 = L2 / (N_FILTER-1);
	size_t h1 = 0, h2 = 0;
	for (size_t i = 0; i < N_FILTER; ++i, ++nhash) {
		size_t i1 = (i != N_FILTER-1 ? i*S1 : L1-1); // XXX no need to use L1-1, i*S1 is okay
		h1 += kmerDBi.count(kmers1[i1]);
		if (h1 >= NM_FILTER) { break; }
	}
	if (h1 < NM_FILTER) { return true; }
	for (size_t i = 0; i < N_FILTER; ++i, ++nhash) {
		size_t i2 = (i != N_FILTER-1 ? i*S2 : L2-1);
		h2 += kmerDBi.count(kmers2[i2]);
		if (h2 >= NM_FILTER) { break; }
	}
	return h2 < NM_FILTER;
}

bool kfilter(vector<size_t>& kmers1, vector<size_t>& kmers2, vector<kmerIndex_uint32_umap::iterator>& its1, vector<kmerIndex_uint32_umap::iterator>& its2, kmerIndex_uint32_umap& kmerDBi, uint16_t Cthreshold, size_t& nhash) {
	size_t ns1 = 0, ns2 = 0;
    const size_t MAX_NS1 = kmers1.size() - Cthreshold;
    const size_t MAX_NS2 = kmers2.size() - Cthreshold;
    for (size_t kmer : kmers1) {
		++nhash;
        auto it = kmerDBi.find(kmer);
        if (it == kmerDBi.end()) { ++ns1; if (ns1 > MAX_NS1) { return true; } }
        else { its1.push_back(it); }
    }
    for (size_t kmer : kmers2) {
		++nhash;
        auto it = kmerDBi.find(kmer);
        if (it == kmerDBi.end()) { ++ns2; if (ns2 > MAX_NS2) { return true; } }
        else { its2.push_back(it); }
    }
	return false;
}

void getSortedIndex(vector<size_t>& data, vector<size_t>& indices) {
	std::iota(indices.begin(), indices.end(), 0);
	std::sort(indices.begin(), indices.end(), [&data](size_t ind1, size_t ind2) { return data[ind1] < data[ind2]; });
}

void getSortedIndex(vector<kmerIndex_uint32_umap::iterator>& data, vector<size_t>& indices) {
    std::iota(indices.begin(), indices.end(), 0);
    std::sort(indices.begin(), indices.end(), [&data](size_t ind1, size_t ind2) { return data[ind1]->first < data[ind2]->first; });
}

void countDupRemove(vector<kmerIndex_uint32_umap::iterator>& its, vector<kmerIndex_uint32_umap::iterator>& its_other, vector<PE_KMC>& dup) {
    // count the occurrence of kmers in each read
   	// Return:
    // 		its: unique entries only
    // 		its_other: empty
	// 		dup: count in <forward,reverse> strand for each entry in kmers
    vector<bool> orient(its.size(), 0);
    orient.resize(its.size() + its_other.size(), 1);
	its.insert(its.end(), its_other.begin(), its_other.end());
	its_other.clear();

    vector<size_t> indorder(its.size());
	getSortedIndex(its, indorder);
    // sort its and orient
    vector<kmerIndex_uint32_umap::iterator> old_its = its;
    vector<bool> old_orient = orient;
    for (size_t i = 0; i < its.size(); ++i) {
        its[i] = old_its[indorder[i]];
        orient[i] = old_orient[indorder[i]];
    }

    // iterate through its and count the occurrence in each read
    assert(its.size()); // XXX
    auto last = its[0];
	size_t itsize = 1;
    PE_KMC pe_kmc(0,0);
    (orient[0] ? ++pe_kmc.second : ++pe_kmc.first);
    for (size_t i = 1; i < its.size(); ++i) {
        if (last != its[i]) { 
            dup.push_back(pe_kmc);
            pe_kmc = std::make_pair(0,0);
            its[itsize] = its[i];
            last = its[i];
            ++itsize;
        }
        (orient[i] ? ++pe_kmc.second : ++pe_kmc.first); 
    }
    dup.push_back(pe_kmc);
    its.resize(itsize);
}

void countRemain(vector<PE_KMC>& dup, vector<size_t>& remain) {
    remain.resize(dup.size(), 0);
    size_t dupsum = std::accumulate(dup.begin(), dup.end(), 0, 
                                    [](size_t partialSum, PE_KMC pe_kmc) { return partialSum + pe_kmc.first + pe_kmc.second; });
    remain[0] = dupsum - dup[0].first - dup[0].second;
    for (size_t i = 1; i < remain.size()-1; ++i) {
        remain[i] = remain[i-1] - dup[i].first - dup[i].second;
    }
}

void fillstats(vector<uint32_t>& kmerDBi_vv, vector<kmerIndex_uint32_umap::iterator>& its, vector<kmerIndex_uint32_umap::iterator>& its_other, vector<PE_KMC>& dup, vector<size_t>& remain) {
    countDupRemove(its, its_other, dup); // count the occurrence of kmers in each read

    // get # of mapped loci for each kmer
    size_t nkmers = its.size();
    vector<size_t> nmappedloci(nkmers, 0); // XXX set to 1 and only change val when multiple loci
    for (size_t i = 0; i < nkmers; ++i) {
		uint32_t vi = its[i]->second;
		nmappedloci[i] = (vi % 2 ? kmerDBi_vv[vi>>1] : 1);
    }

    // sort kmers dup w.r.t. nmappedloci; remove entries w/o mapped locus
    vector<size_t> indorder(nmappedloci.size());
	getSortedIndex(nmappedloci, indorder);
    vector<kmerIndex_uint32_umap::iterator> old_its = its; // XXX reserve not copy
    vector<PE_KMC> old_dup = dup; // XXX reserve not copy
    for (size_t i = 0; i < nkmers; ++i) {
        its[i] = old_its[indorder[i]];
        dup[i] = old_dup[indorder[i]];
    }
    countRemain(dup, remain);
}

void updatetop2(size_t count_f, uint32_t ind, size_t count_r, asgn_t& top, asgn_t& second) { // for sorted_query algo
    if (count_f + count_r > top.fc + top.rc) {
        if (top.idx != ind) {
			second = top;
			top.idx = ind;
        }
		top.fc = count_f;
		top.rc = count_r;
    }
    else if (count_f + count_r > second.fc + second.rc) {
        if (second.idx != ind) {
            second.idx = ind;
        }
		second.fc = count_f;
		second.rc = count_r;
    }
}

inline bool get_cmp(asgn_t& top, asgn_t& second, size_t rem, float rth) {
	return float(top.fc + top.rc      )/(top.fc + top.rc + second.fc + second.rc + rem) <  rth and
           float(top.fc + top.rc + rem)/(top.fc + top.rc + second.fc + second.rc + rem) >= rth;
}

inline bool get_acm1(asgn_t& top, asgn_t& second, size_t rem, size_t cth) { // XXX speedup second is not used here
	// accumulate the score of the top locus to identify reads w/ score >= Cthreshold
	return (top.fc < cth and cth - top.fc <= rem) or (top.rc < cth and cth - top.rc <= rem);
}

inline bool get_acm2(asgn_t& top, asgn_t& second, size_t rem) { 
	// accumulate the scores of the top 2 loci to assign reads to the most similar locus
	return (top.fc + top.rc - second.fc - second.rc) < rem;
}

void find_matching_locus(vector<uint32_t>& kmerDBi_vv, vector<kmerIndex_uint32_umap::iterator>& its1, vector<uint32_t>& hits1, vector<uint32_t>& hits2, 
                         vector<PE_KMC>& dup, vector<size_t>& remain, asgn_t& top, asgn_t& second, uint16_t Cthreshold, float Rthreshold) {
    for (size_t i = 0; i < its1.size(); ++i) {
		uint32_t vi = its1[i]->second;
		if (vi % 2) {
			size_t j0 = (vi>>1) + 1;
			size_t j1 = j0 + kmerDBi_vv[vi>>1];
			for ( ; j0 < j1; ++j0) {
				uint32_t locus = kmerDBi_vv[j0];
				hits1[locus] += dup[i].first; // XXX speedup; try unordered_map for hits?
				hits2[locus] += dup[i].second;
				updatetop2(hits1[locus], locus, hits2[locus], top, second);
			}
		} else {
			uint32_t locus = vi >> 1;
			hits1[locus] += dup[i].first;
			hits2[locus] += dup[i].second;
			updatetop2(hits1[locus], locus, hits2[locus], top, second);
		}
        if (not get_acm2(top, second, remain[i])) { // second hit will not exceed top hit
            size_t j = i;
            //if (Rthreshold != 0.5) { // continue to count scores until PASS/FAIL can be determined
            //    while (get_cmp(top, second, remain[j], Rthreshold)) { // XXX second.idx might not be fixed yet
            //        unordered_set<uint32_t>& loci = its1[++j]->second;
            //        if (loci.count(top.idx)) {
            //            top.fc += dup[j].first;
            //            top.rc += dup[j].second;
            //        }
            //        else if (loci.count(second.idx)) {
            //            second.fc += dup[j].first;
            //            second.rc += dup[j].second;
            //        }
            //    }
            //}
            while (get_acm1(top, second, remain[j], Cthreshold)) { // XXX speedup? use graph to do the same thing
				if (++j >= its1.size()) { break; }
				uint32_t vj = its1[j]->second;
				if (vj % 2) {
					size_t j0 = (vj>>1) + 1;
					size_t j1 = j0 + kmerDBi_vv[vj>>1];
					for ( ; j0 < j1; ++j0) {
						uint32_t locus = kmerDBi_vv[j0];
						if (locus == top.idx) {
							top.fc += dup[j].first;
							top.rc += dup[j].second;
							break;
						}
					}
				} else {
					if ((vj >> 1) == top.idx) {
						top.fc += dup[j].first;
						top.rc += dup[j].second;
					}
				}
            }
            break;
        }
    }
}

size_t countHit(vector<uint32_t>& kmerDBi_vv, vector<kmerIndex_uint32_umap::iterator>& its1, vector<kmerIndex_uint32_umap::iterator>& its2, vector<uint32_t>& hits1, vector<uint32_t>& hits2,
                vector<PE_KMC>& dup, size_t nloci, uint16_t Cthreshold, float Rthreshold) {
	// pre-processing: sort kmer by # mapped loci XXX alternative: sort by frequncy in read
    vector<size_t> remain;
    fillstats(kmerDBi_vv, its1, its2, dup, remain);

    // for each kmer, increment counts of the mapped loci for each read
    // use "remain" to achieve early stopping
	asgn_t top, second;
	std::fill(hits1.begin(), hits1.end(), 0);
	std::fill(hits2.begin(), hits2.end(), 0);
	find_matching_locus(kmerDBi_vv, its1, hits1, hits2, dup, remain, top, second, Cthreshold, Rthreshold);
	if (top.fc >= Cthreshold and top.rc >= Cthreshold and 
	    float(top.fc + top.rc)/(top.fc + top.rc + second.fc + second.rc) >= Rthreshold and 
	    top.idx != NAN32) { 
		return top.idx;
	} 
	else { 
		return nloci;
	}
}

inline void prunePEinfo(string& title) {
	size_t len = title.size();
    if (title[len-2] == '/') {
        if (title[len-1] == '1' or title[len-1] == '2') {
			title = title.substr(0, len-2);
		}
    }
}

// skip step 1, read destLocus from >[READ_NAME]:[DEST_LOCUS]_[1|2]
void parseReadNames(vector<string>& titles, vector<size_t>& destLoci, size_t nReads_) {
	size_t ri = 0;
	for (size_t di = 0; di < nReads_/2; ++di) {
		size_t beg = titles[ri].size() - 1;
		size_t len = 1;
		while (titles[ri][--beg] != ':') { ++len; }
		destLoci[di] = stoul(titles[ri].substr(++beg, len));
		ri += 2;
	}
}

// simmode = 1; simmulated reads from TR only
template <typename ValueType>
void parseReadName(string& title, size_t readn, vector<ValueType>& loci, vector<size_t>& locusReadi) {
    string sep = ".";
    size_t first = title.find(sep);
    size_t newLocus = stoul(title.substr(1, first)); // skip the 1st '>' char
    if (readn == 0) {
        loci.push_back(newLocus);
    }
    else if (newLocus != loci.back()) {
        loci.push_back(newLocus);
        locusReadi.push_back(readn);
    }
}

// OBSOLETE. simmode = 2; simmulated reads from whole genome
template <typename ValueType>
void parseReadName(string& title, size_t readn, vector<size_t>& poss, vector<ValueType>& loci, vector<size_t>& locusReadi) {
    string sep = "_";
    size_t first = title.find(sep);
    size_t second = title.find(sep, first+1);
    float newLocus = stof(title.substr(first+1, second));
    if (readn == 0) {
        //size_t hap = stoi(title.substr(1, first)); // skip the 1st '>' char
        loci.push_back(newLocus);
        poss.push_back(stoul(title.substr(second+1, title.find(sep, second+1))));
    }
    else if (newLocus != loci.back()) {
        loci.push_back(newLocus);
        poss.push_back(stoul(title.substr(second+1, title.find(sep, second+1))));
        locusReadi.push_back(readn);
    }
}

// simmode = 2
void parseReadName(string& title, vector<std::pair<int, size_t>>& meta, size_t nloci) {
	// input: read name; vector of (read_locus, number_of_pe_reads)
	static const string sep = ":";
	size_t p1 = title.find(sep);
	size_t p2 = title.find(sep, p1+1);
	string v = title.substr(p1+1, p2-p1-1);
	size_t locus;
	if (v[0] == '.') { locus = nloci; }
	else { locus = stoul(v); }
	if (meta.size() == 0) { meta.push_back(std::make_pair(locus, 1)); }
	else {
		if (meta.back().first == locus) { ++meta.back().second; } // if locus is the same as the last read, increment number_of_pe_reads
		else { meta.push_back(std::make_pair(locus, meta.back().second+1)); } // else, append (read_locus, 1)
	}
}

void mapLocus(bool g2pan, vector<std::pair<int, size_t>>& meta, vector<size_t>& locusmap, size_t seqi, size_t& simi, size_t nloci, size_t& srcLocus) {
	if (simi == 0 or seqi / 2 >= meta[simi].second) { 
		if (seqi / 2 >= meta[simi].second) { ++simi; }
		if (meta[simi].first == nloci) { srcLocus = nloci; }
		else {
			if (g2pan) { 
				if (meta[simi].first >= locusmap.size()) {
					cerr << "read locus " << meta[simi].first << " > # of valid genome loci = " << locusmap.size() << endl;
					assert(false);
				}
				srcLocus = locusmap[meta[simi].first];
			} else {
				srcLocus = meta[simi].first;
			}
		}	
	}
}

template <typename T>
void printVec(vector<T>& vec) {
    for (auto v : vec) { cerr << v << ' '; }
    cerr << endl;
}

void getOutNodes(GraphType& g, size_t node, vector<size_t>& nnds, bool (&nnts)[4]) {
    // a node is a kmer and is not neccessarily canonical
    const static uint64_t mask = (1UL << 2*(ksize-1)) - 1;
	assert(g.count(node)); // prevents error from unclean graph XXX remove this after graph pruning code passes testing
	uint8_t nucBits = g[node]; // a 4-bit value that corresponds to the presence of trailing TGCA in downstream nodes
	size_t nnd = (node & mask) << 2;
	for (size_t i = 0; i < 4; ++i) {
		if (nucBits % 2) { nnds.push_back(nnd + i); }
		nnts[i] |= nucBits % 2; // CAUTION: OR operator
		nucBits >>= 1;
    }
}

void getNextNucs(GraphType& g, size_t node, bool (&nnts)[4]) {
    const static uint64_t mask = (1UL << 2*(ksize-1)) - 1;
	uint8_t nucBits;
	auto it = g.find(node);
	if (it != g.end()) {
		nucBits = it->second;
		for (size_t i = 0; i < 4; ++i) {
			nnts[i] = nucBits % 2; // CAUTION: assignment operator
			nucBits >>= 1;
		}
	}
}


struct thread_ext_t{
	// nem1: number of extended kmers starting with 1 substitution
	// nem2: number of extended kmers starting with 2 substitutions
	// nemi: number of extended kmers starting with 1 substitution + 1 insertion
	// nemd: number of extended kmers starting with 1 substitution + 1 deletion
	// ned1: number of extended kmers starting with 1 deletion
	// ned2: number of extended kmers starting with 2 deletions
	// nei1: number of extended kmers starting with 1 insertions
	// nei2: number of extended kmers starting with 2 insertions
	//                       type_shft
	size_t nem1[4]  = {}; //  0
	size_t nem2[16] = {}; //  1
	size_t nemi[4]  = {}; //  1
	size_t nemd[16] = {}; //  0
	size_t ned1[4]  = {}; // -1
	size_t ned2[16] = {}; // -1
	size_t nei1 = 0;      //  0
	size_t nei2 = 0;      //  1
	size_t score = 0;     // highest number of extended kmers
	size_t shft = 0;      // read index shifting = type_shft + score
	int dt_kmers = 0, dt_ki = 0;
	int dt_ops = 0;
	string edit = "";

	bool get_edit() {
		const char dels[4] = {'0','1','2','3'};
		if (nei1 > score) { score = nei1; shft = 0 + score; edit = "I"; }
		for (size_t i = 0; i < 4; ++i) { if (ned1[i] > score) { score = ned1[i]; shft = -1 + score; edit = string(1,dels[i]); } }
		for (size_t i = 0; i < 4; ++i) { if (nem1[i] > score) { score = nem1[i]; shft =  0 + score; edit = string(1,alphabet[i]); } }
		if (nei2 > score) { score = nei2; shft = 1 + score; edit = "II"; }
		for (size_t i = 0; i < 4; ++i) {
			if (nemi[i] > score) { score = nemi[i]; shft = 1 + score; edit = string(1,alphabet[i]) + "I"; }
			for (size_t j = 0; j < 4; ++j) {
				if (ned2[i*4+j] > score) { score = ned2[i*4+j]; shft = -1 + score; edit = string(1,dels[i])     + dels[j]; }
				if (nemd[i*4+j] > score) { score = nemd[i*4+j]; shft =  0 + score; edit = string(1,alphabet[i]) + dels[j]; }
				if (nem2[i*4+j] > score) { score = nem2[i*4+j]; shft =  1 + score; edit = string(1,alphabet[i]) + alphabet[j]; }
			}
		}
		return score > 0;
	}

	void edit_kmers(vector<size_t>& kmers, size_t& ki, bool aln, vector<char>& ops, size_t& opsi, kmer_aCount_umap& trKmers) {
		const static uint64_t mask = (1UL << 2*(ksize-1)) - 1;
		size_t nts[kmers.size() - ki];
		for (size_t i = ki; i < kmers.size(); ++i) {
			nts[i-ki] = kmers[i] % 4;
		}
		size_t nti = 0;
		for (size_t i = 0; i < edit.size(); ++i) {
			char c = edit[i];
			if (c == 'A' or c == 'C' or c == 'G' or c == 'T') { kmers[ki+i] = ((kmers[ki+i-1] & mask) << 2) + baseNumConversion[c]; ++nti; }
			else if (c == '0' or c == '1' or c == '2' or c == '3') { kmers[ki+i] = ((kmers[ki+i-1] & mask) << 2) + stoi(string(1,c)); ++dt_kmers; ++dt_ops; }
			else if (c == 'I') { --dt_ki; --dt_kmers; ++nti; }
			++dt_ki;
		}
		kmers.resize(kmers.size() + dt_kmers);
		ops.resize(ops.size() + dt_ops, '*');
		for (size_t i = ki+dt_ki; i < kmers.size(); ++i) {
			kmers[i] = ((kmers[i-1] & mask) << 2) + nts[nti++];
		}
		ki += dt_ki - 1;
		if (aln) { 
			for (size_t i = 0; i < edit.size(); ++i) { ops[opsi++] = edit[i]; } 
			for (size_t i = 0; i < score; ++i) { ops[opsi++] = trKmers.count(toCaKmer(kmers[++ki], ksize)) ? '=' : '.'; }
		}
	}
};

struct graph_triplet_t {
	bool mat[64] = {};

	void get_nnts(size_t i, bool (&nnts)[4]) { // CAUTION: OR operator
		for (size_t j = 0; j < 4; ++j) { for (size_t k = 0; k < 4; ++k) { nnts[j] |= mat[i*16 + j*4 + k]; } }
	}

	void get_nnts(size_t i, size_t j, bool (&nnts)[4]) { // CAUTION: OR operator
		for (size_t k = 0; k < 4; ++k) { nnts[k] |= mat[i*16 + j*4 + k]; }
	}
};

bool find_anchor(GraphType& g, vector<size_t>& kmers, bool aln, vector<char>& ops, size_t& nskip, size_t& pos, size_t& opsi, kmer_aCount_umap& trKmers, size_t& node) {
	while (not g.count(kmers[pos])) {
		++nskip;
        if (aln) { ops[opsi++] = '*'; }
        if (++pos >= kmers.size()) { return 0; }
    }
	node = kmers[pos];
    if (aln) { ops[opsi++] = trKmers.count(toCaKmer(node, ksize)) ? '=' : '.'; }
	return 1;
}

// 0: not feasible, 1: feasible, w/o correction, 2: feasible w/ correction
int isThreadFeasible(GraphType& g, string& seq, vector<size_t>& noncakmers, size_t thread_cth, bool correction, 
	bool aln, vector<char>& ops, kmer_aCount_umap& trKmers) {

    read2kmers(noncakmers, seq, ksize, 0, 0, false, true); // leftflank = 0, rightflank = 0, canonical = false, keepN = true
	vector<size_t> kmers(noncakmers.begin(), noncakmers.end());

	static const char nts[] = {'A', 'C', 'G', 'T', 'a', 'c', 'g', 't'};
    static const uint64_t mask = (1UL << 2*(ksize-1)) - 1;
    const size_t maxnskip = (kmers.size() >= thread_cth ? kmers.size() - thread_cth : 0);
    const size_t maxncorrection = 2;
    size_t i = 0, opsi = 0, nskip = 0, ncorrection = 0;
	size_t node = kmers[nskip];

	if (aln) { ops.resize(kmers.size(), '*'); }
	if (not find_anchor(g, kmers, aln, ops, nskip, i, opsi, trKmers, node)) { return 0; }
    for (i = i+1; i < kmers.size(); ++i) {
		if (kmers[i] == -1ULL) { // "N" in read
			if (aln) { ops[opsi++] = 'N'; }
			++nskip;
            if (nskip > maxnskip) { return 0; }
			continue;
		}
        if (kmers[i] == kmers[i-1]) { // skip homopolymer run
			if (aln) { ops[opsi++] = trKmers.count(toCaKmer(kmers[i], ksize)) ? 'H' : 'h'; }
            ++nskip;
			if (nskip > maxnskip) { return 0; }
            continue;
        }
		if (kmers[i-1] == -1ULL) {
			if (not find_anchor(g, kmers, aln, ops, nskip, i, opsi, trKmers, node)) { break; }
			else { 
				if (nskip > maxnskip) { return 0; }
				else { continue; }
			}
		}

        bool skip = true;
		bool nts0[4] = {};
		size_t nkmers = kmers.size();
		vector<size_t> nnds;
		getOutNodes(g, node, nnds, nts0);
		for (size_t nnd : nnds) {
			if (kmers[i] == nnd) { // matching node found
				node = nnd;
				skip = false;
				if (aln) { ops[opsi++] = trKmers.count(toCaKmer(kmers[i], ksize)) ? '=' : '.'; }
				break;
			}
		}
		if (not skip) { continue; }
        else { // read kmer has no matching node in the graph, try error correction
			if (i + 3 > nkmers) { 
				nskip += (nkmers - i);
				return (nskip <= maxnskip ? (ncorrection ? 2 : 1) : 0);
			}
            size_t oldnt = kmers[i] % 4;
			thread_ext_t txt;
			graph_triplet_t gnt3; // 3 consecutive nucleotides in the graph; 4x4x4 matrix

            if (correction and ncorrection < maxncorrection) {
				bool nts1[4] = {};
				bool nts2[4] = {};
				for (size_t node_i : nnds) {
					size_t nt0 = node_i % 4;
					vector<size_t> nodes_ip1;
					getOutNodes(g, node_i, nodes_ip1, nts1);
					for (size_t node_ip1 : nodes_ip1) {
						size_t nt1 = node_ip1 % 4;
						vector<size_t> nodes_ip2;
						getOutNodes(g, node_ip1, nodes_ip2, nts2);
						for (size_t node_ip2 : nodes_ip2) {
							size_t nt2 = node_ip2 % 4;
							gnt3.mat[nt0*4*4 + nt1*4 + nt2] = true;
						}
					}
				}
				
				// One mismatch: match at i+1 position
				if (nts1[kmers[i+1] % 4]) {
					for (size_t nt0 = 0; nt0 < 4; ++nt0) {
						if (not nts0[nt0]) { continue; }
						size_t crkmer = kmers[i] - oldnt + nt0; // corrected read kmer
						bool nnts[4] = {}; // next nucleotides
						gnt3.get_nnts(nt0, nnts);
						for (size_t j = 1; j < std::min(ksize, nkmers-i); ++j) {
							crkmer = ((crkmer & mask) << 2) + kmers[i+j] % 4;
							if (nnts[crkmer % 4]) {
								++txt.nem1[nt0];
								getNextNucs(g, crkmer, nnts);
							} else {
								break;
							}
						}
					}
				}
				// Two mismatches: match at i+2 position
				else if (nts2[kmers[i+2] % 4]) {
					for (size_t nt0 = 0; nt0 < 4; ++nt0) {
						if (not nts0[nt0]) { continue; }
						size_t crkmer0 = kmers[i] - oldnt + nt0; // corrected read kmer at i+0 positionA
						bool nnt0[4] = {}; // next nucleotides for node_{i+0}
						gnt3.get_nnts(nt0, nnt0);
						for (size_t nt1 = 0; nt1 < 4; ++nt1) {
							if (not nnt0[nt1]) { continue; }
							size_t crkmer1 = ((crkmer0 & mask) << 2) + nt1;
							bool nnt1[4] = {}; // next nucleotides for node_{i+1}
							gnt3.get_nnts(nt0, nt1, nnt1);
							for (size_t j = 2; j < std::min(ksize+1, nkmers-i); ++j) {
								crkmer1 = ((crkmer1 & mask) << 2) + kmers[i+j] % 4;
								if (nnt1[crkmer1 % 4]) {
									++txt.nem2[nt0*4 + nt1];
									getNextNucs(g, crkmer1, nnt1);
								} else {
									break;
								}
							}
						}
					}
				}
				// 1 substitution + 1 insersion
				if (nts1[kmers[i+2] % 4]) {
					for (size_t nt0 = 0; nt0 < 4; ++nt0) {
						if (not nts0[nt0]) { continue; }
						size_t crkmer = kmers[i] - oldnt + nt0;
						bool nnt0[4] = {};
						gnt3.get_nnts(nt0, nnt0);
						for (size_t j = 2; j < std::min(ksize+1, nkmers-i); ++j) {
							crkmer = ((crkmer & mask) << 2) + kmers[i+j] % 4;
							if (nnt0[crkmer % 4]) {
								++txt.nemi[nt0];
								getNextNucs(g, crkmer, nnt0);
							} else {
								break;
							}
						}
					}
				}
				// 1 substitution + 1 deletion
				if (nts2[kmers[i+1] % 4]) {
					for (size_t nt0 = 0; nt0 < 4; ++nt0) {
						if (not nts0[nt0]) { continue; }
						size_t crkmer0 = kmers[i] - oldnt + nt0;
						bool nnt0[4] = {};
						gnt3.get_nnts(nt0, nnt0);
						for (size_t nt1 = 0; nt1 < 4; ++nt1) {
							if (not nnt0[nt1]) { continue; }
							size_t crkmer1 = ((crkmer0 & mask) << 2) + nt1;
							bool nnt1[4] = {};
							gnt3.get_nnts(nt0, nt1, nnt1);
							for (size_t j = 1; j < std::min(ksize, nkmers-i); ++j) {
								crkmer1 = ((crkmer1 & mask) << 2) + kmers[i+j] % 4;
								if (nnt1[crkmer1 % 4]) {
									++txt.nemd[nt0*4 + nt1];
									getNextNucs(g, crkmer1, nnt1);
								} else {
									break;
								}
							}
						}
					}
				}
				// 1 insertion
				if (nts0[kmers[i+1] % 4]) {
					size_t crkmer = kmers[i-1];
					bool nnt0[4] = {nts0[0], nts0[1], nts0[2], nts0[3]};
					for (size_t j = 1; j < std::min(ksize, nkmers-i); ++j) {
						crkmer = ((crkmer & mask) << 2) + kmers[i+j] % 4;
						if (nnt0[crkmer % 4]) {
							++txt.nei1;
							getNextNucs(g, crkmer, nnt0);
						} else {
							break;
						}
					}
				}
				// 1 deletion
				if (nts1[kmers[i+0] % 4]) {
					for (size_t nt0 = 0; nt0 < 4; ++nt0) {
						if (not nts0[nt0]) { continue; }
						size_t crkmer = kmers[i] - oldnt + nt0;
						bool nnt0[4] = {};
						gnt3.get_nnts(nt0, nnt0);
						for (size_t j = 0; j < std::min(ksize-1, nkmers-i); ++j) {
							crkmer = ((crkmer & mask) << 2) + kmers[i+j] % 4;
							if (nnt0[crkmer % 4]) {
								++txt.ned1[nt0];
								getNextNucs(g, crkmer, nnt0);
							} else {
								break;
							}
						}
					}
				}
				// 2 insertions
				if (nts0[kmers[i+2] % 4]) {
					size_t crkmer = kmers[i-1];
					bool nnt0[4] = {nts0[0], nts0[1], nts0[2], nts0[3]};
					for (size_t j = 2; j < std::min(ksize+1, nkmers-i); ++j) {
						crkmer = ((crkmer & mask) << 2) + kmers[i+j] % 4;
						if (nnt0[crkmer % 4]) {
							++txt.nei2;
							getNextNucs(g, crkmer, nnt0);
						} else {
							break;
						}
					}
				}
				// 2 deletions
				if (nts2[kmers[i+0] % 4]) {
					for (size_t nt0 = 0; nt0 < 4; ++nt0) {
						if (not nts0[nt0]) { continue; }
						size_t crkmer0 = kmers[i] - oldnt + nt0;
						bool nnt0[4] = {};
						gnt3.get_nnts(nt0, nnt0);
						for (size_t nt1 = 0; nt1 < 4; ++nt1) {
							if (not nnt0[nt1]) { continue; }
							size_t crkmer1 = ((crkmer0 & mask) << 2) + nt1;
							bool nnt1[4] = {};
							gnt3.get_nnts(nt0, nt1, nnt1);
							for (size_t j = 0; j < std::min(ksize-1, nkmers-i); ++j) {
								crkmer1 = ((crkmer1 & mask) << 2) + kmers[i+j] % 4;
								if (nnt1[crkmer1 % 4]) {
									++txt.ned2[nt0*4 + nt1];
									getNextNucs(g, crkmer1, nnt1);
								} else {
									break;
								}
							}
						}
					}
				}
				skip = !txt.get_edit();
				// longer edits are treated with path-skipping and re-anchoring using find_anchor()
			}

			if (skip) {
				if (not find_anchor(g, kmers, aln, ops, nskip, i, opsi, trKmers, node)) { break; } // anchor can be arbitrary far from the last thread
				else { 
					if (nskip > maxnskip) { return 0; }
					else { continue; }
				}
            }
            else {
				// resize kmers & ops; shift i to the last kmer examined/edited; shift opsi to the next to be examined
				txt.edit_kmers(kmers, i, aln, ops, opsi, trKmers);
				node = kmers[i];
				++ncorrection;
            }
        }
    }
    return (nskip <= maxnskip and ncorrection <= maxncorrection ? (ncorrection ? 2 : 1) : 0);
}

void writeExtractedReads(int extractFasta, vector<string>& seqs, vector<string>& titles, vector<size_t>& extractindices, vector<size_t>& assignedloci) {
	for (size_t i = 0; i < extractindices.size(); ++i) {
		if (extractFasta == 1) { cout << titles[--extractindices[i]] << '\n'; } 
		else { cout << titles[--extractindices[i]] << ":" << assignedloci[i] << '\n'; }
		cout << seqs[extractindices[i]] << '\n';

		if (extractFasta == 1) { cout << titles[--extractindices[i]] << '\n'; } 
		else { cout << titles[--extractindices[i]] << ":" << assignedloci[i] << '\n'; }
		cout << seqs[extractindices[i]] << '\n';
	}
}

inline void writeOps(vector<char>& ops) {
	for (char c : ops) { cout << c; }
}

void writeAlignments(vector<string>& seqs, vector<string>& titles, vector<size_t>& alnindices, vector<EDIT>& sam) {
	for (size_t i = 0; i < sam.size(); ++i) {
		if (sam[i].map.first == -1ULL) { cout << '.' << '\t'; }
		else { cout << sam[i].map.first << '\t'; }
		cout << sam[i].map.second << '\t'
			 << titles[--alnindices[i]] << '\t'
			 << seqs[alnindices[i]] << '\t';
		writeOps(sam[i].ops2); // read2
		cout << '\t'
			 << titles[--alnindices[i]] << '\t'
			 << seqs[alnindices[i]] << '\t';
		writeOps(sam[i].ops1); // read1
		cout << '\n';
	}
}

void writeAlignments(vector<string>& seqs, vector<string>& titles, vector<size_t>& destLoci, vector<size_t>& alnindices, vector<EDIT>& sam) {
	for (size_t i = 0; i < sam.size(); ++i) {
		if (sam[i].map.first == -1ULL) { cout << '.' << '\t'; }
		else { cout << sam[i].map.first << '\t'; }
		cout << sam[i].map.second << '\t'
			 << titles[--alnindices[i]] << ':' << destLoci[alnindices[i]/2] << '\t'
			 << seqs[alnindices[i]] << '\t';
		writeOps(sam[i].ops2); // read2
		cout << '\t'
			 << titles[--alnindices[i]] << ':' << destLoci[alnindices[i]/2] << '\t'
			 << seqs[alnindices[i]] << '\t';
		writeOps(sam[i].ops1); // read1
		cout << '\n';
	}
}

class Counts {
public:
    bool interleaved, bait, threading, correction, aln, aln_minimal, g2pan, skip1;
    uint16_t Cthreshold, thread_cth;
    size_t *nReads, *nThreadingReads, *nFeasibleReads, *nSubFiltered, *nKmerFiltered;
    size_t nloci;
    float Rthreshold;
	unordered_map<string, string>* readDB;
	kmerIndex_uint32_umap* kmerDBi;
	vector<uint32_t>* kmerDBi_vv;
    vector<GraphType>* graphDB;
    vector<kmer_aCount_umap>* trResults;
    ifstream *in;
    // simmode only
    int simmode;
    vector<msa_umap>* msaStats;
    err_umap* errdb;
	vector<size_t>* locusmap;
    // extractFasta only
	int extractFasta;

    Counts(size_t nloci_) : nloci(nloci_) {}
};

class Threads {
public:
    vector<Counts> counts;
    Threads(size_t nproc, size_t nloci) : counts(nproc, Counts(nloci)) {}
};

template <typename ValueType>
void CountWords(void *data) {
    bool interleaved = ((Counts*)data)->interleaved;
    bool bait = ((Counts*)data)->bait;
    bool threading = ((Counts*)data)->threading;
    bool correction = ((Counts*)data)->correction;
    bool aln = ((Counts*)data)->aln;
    bool aln_minimal = ((Counts*)data)->aln_minimal;
	bool g2pan = ((Counts*)data)->g2pan;
	bool skip1 = ((Counts*)data)->skip1; // TODO new functionality: allow skipping step1 by reading assigned locus info from extracted reads
    int simmode = ((Counts*)data)->simmode;
    int extractFasta = ((Counts*)data)->extractFasta;
    size_t nReads_ = 0, nShort_ = 0, nThreadingReads_ = 0, nFeasibleReads_ = 0, nSubFiltered_ = 0, nKmerFiltered_ = 0;
    size_t& nReads = *((Counts*)data)->nReads;
    size_t& nThreadingReads = *((Counts*)data)->nThreadingReads;
    size_t& nFeasibleReads = *((Counts*)data)->nFeasibleReads;
    size_t& nSubFiltered = *((Counts*)data)->nSubFiltered;
    size_t& nKmerFiltered = *((Counts*)data)->nKmerFiltered;
    uint16_t Cthreshold = ((Counts*)data)->Cthreshold;
    uint16_t thread_cth = ((Counts*)data)->thread_cth;
    float Rthreshold = ((Counts*)data)->Rthreshold;
    const size_t nloci = ((Counts*)data)->nloci;
    const size_t readsPerBatch = 300000;
	const size_t minReadSize = Cthreshold + ksize - 1;
    ifstream *in = ((Counts*)data)->in;
    unordered_map<string, string>& readDB = *((Counts*)data)->readDB;
    kmerIndex_uint32_umap& kmerDBi = *((Counts*)data)->kmerDBi;
	vector<uint32_t>& kmerDBi_vv = *((Counts*)data)->kmerDBi_vv;
    vector<GraphType>& graphDB = *((Counts*)data)->graphDB;
    vector<kmer_aCount_umap>& trResults = *((Counts*)data)->trResults;
    vector<msa_umap>& msaStats = *((Counts*)data)->msaStats;
    err_umap& errdb = *((Counts*)data)->errdb;
	err_umap err;
	vector<size_t>& locusmap = *((Counts*)data)->locusmap;
	vector<string> seqs(readsPerBatch);
	vector<uint32_t> hits1(nloci+1,0), hits2(nloci+1,0);
	// extractFasta only
	vector<string> titles(readsPerBatch);
	vector<size_t> destLoci(readsPerBatch/2);
    // simmode only
    // loci: loci that are processed in this batch
    vector<ValueType> srcLoci;
    vector<size_t> poss;
    unordered_map<size_t, msa_umap> msa;

    while (true) {

        string title, title1, seq, seq1;
        // for simmode only
        // locusReadi: map locus to nReads_. 0th item = number of reads for 0th item in loci; last item = nReads_; has same length as loci
        size_t startpos;
        vector<size_t> locusReadi;
		vector<std::pair<int, size_t>> meta;
        // extractFasta only
        vector<size_t> extractindices, assignedloci;
		// aln only
		vector<size_t> alnindices;

        //
        // begin thread locking
        //
        sem_wait(semreader);

        nThreadingReads += nThreadingReads_;
        nFeasibleReads += nFeasibleReads_;
        nSubFiltered += nSubFiltered_;
        nKmerFiltered += nKmerFiltered_;
        nThreadingReads_ = 0;
        nFeasibleReads_ = 0;
		nSubFiltered_ = 0;
		nKmerFiltered_ = 0;
        nReads_ = 0;
		nShort_ = 0;

        if (simmode == 1 and srcLoci.size() != 0) {
            srcLoci.clear();
            msa.clear();
        }

        if (in->peek() == EOF) {
            sem_post(semreader);
            return;
        }

        while (nReads_ < readsPerBatch and in->peek() != EOF) {
            if (interleaved) {
                getline(*in, title);
                getline(*in, seq);
                getline(*in, title1);
                getline(*in, seq1);
            }
            else {
				bool se = true; // single end
				while (se) {
					getline(*in, title);
					getline(*in, seq);
					prunePEinfo(title);
					auto it = readDB.find(title);
					if (it != readDB.end()) {
						if (seq.size() < minReadSize or it->second.size() < minReadSize) { readDB.erase(title); continue; }
						title1 = title;
						seq1 = it->second;
						readDB.erase(title);
						se = false;
						break;
					} else {
						readDB[title] = seq;
					}
					if (in->peek() == EOF) { break; }
				}
				if (se and in->peek() == EOF) { break; }
            }
			
            if (simmode == 1) { parseReadName(title, nReads_, srcLoci, locusReadi); }
            else if (simmode == 2) { parseReadName(title, meta, nloci); }

			if (extractFasta or aln) { titles[nReads_] = title; }
            seqs[nReads_++] = seq;
			if (extractFasta or aln) { titles[nReads_] = title1; }
            seqs[nReads_++] = seq1;
        } 
        nReads += nReads_;

        if (simmode == 1) { locusReadi.push_back(nReads_); }
		if (skip1) { parseReadNames(titles, destLoci, nReads_); }

        cerr << "Buffered reading " << nReads_ << '\t' << nReads << '\t' << readDB.size() << endl;

        //
        // All done reading, release the thread lock.
        //
        sem_post(semreader);

        time_t time2 = time(nullptr);
        size_t seqi = 0;
		size_t nhash0 = 0, nhash1 = 0;
		size_t destLocus;
		// aln only TODO define SAM structure
		vector<EDIT> sam;
        // simmode only
        size_t simi = 0;
        ValueType srcLocus = -1;
        if (simmode == 1) { srcLocus = srcLoci[simi]; }

        while (seqi < nReads_) {

            vector<size_t> kmers1, kmers2;
			vector<kmerIndex_uint32_umap::iterator> its1, its2;
            vector<PE_KMC> dup;

            if (simmode == 1) {
                if (seqi >= locusReadi[simi]) {
                    ++simi;
                    srcLocus = srcLoci[simi];
                }
            }
			else if (simmode == 2) { mapLocus(g2pan, meta, locusmap, seqi, simi, nloci, srcLocus); }

            string& seq = seqs[seqi++];
            string& seq1 = seqs[seqi++];

			if (not skip1) {
				read2kmers(kmers1, seq, ksize); // stores numeric canonical kmers
				read2kmers(kmers2, seq1, ksize);
				if (not kmers1.size() or not kmers2.size()) { 
					++nShort_;
					if (verbosity >= 3) {
						cerr << titles[seqi-2] << ' ' << seq  << '\n'
							 << titles[seqi-1] << ' ' << seq1 << '\n';
					}
					continue; 
				}
				if (N_FILTER and NM_FILTER) {
					if (subfilter(kmers1, kmers2, kmerDBi, nhash0)) { 
						nSubFiltered_ += 2;
						continue;
					}
				}
				if (kfilter(kmers1, kmers2, its1, its2, kmerDBi, Cthreshold, nhash1)) {
					nKmerFiltered_ += 2;
					continue; 
				}
				destLoci[seqi/2 - 1] = countHit(kmerDBi_vv, its1, its2, hits1, hits2, dup, nloci, Cthreshold, Rthreshold);
			}
			destLocus = destLoci[seqi/2 - 1];

			if (destLocus != nloci) {
				int feasibility0 = 0, feasibility1 = 0;
				kmerCount_umap cakmers;
				EDIT edit;
				nThreadingReads_ += 2;

				if (threading) {
					vector<size_t> noncakmers0, noncakmers1;
					feasibility0 = isThreadFeasible(graphDB[destLocus], seq, noncakmers0, thread_cth, correction, aln, edit.ops1, trResults[destLocus]);
					feasibility1 = isThreadFeasible(graphDB[destLocus], seq1, noncakmers1, thread_cth, correction, aln, edit.ops2, trResults[destLocus]);
					if (feasibility0 and feasibility1) {
						noncaVec2CaUmap(noncakmers0, cakmers, ksize);
						noncaVec2CaUmap(noncakmers1, cakmers, ksize);
					}
					if (verbosity >= 3) { cerr << "Read threaded: " << feasibility0 << feasibility1 << endl; }
				}

				if ((threading and feasibility0 and feasibility1) or not threading) {
					kmer_aCount_umap &trKmers = trResults[destLocus];
					nFeasibleReads_ += 2;

					if (extractFasta) {
						// points to the next read pair 
						// i.e. to_be_extract_forward (seqi-2), to_be_extract_reverse (seqi-1)
						extractindices.push_back(seqi); 
						if (extractFasta == 2) {
							assignedloci.push_back(destLocus);
						}
					}
					else { // accumulate trKmers for output
						if (not threading) {
							for (size_t i = 0; i < its1.size(); ++i) {
								auto it = trKmers.find(its1[i]->first);
								if (it != trKmers.end()) { it->second += (dup[i].first + dup[i].second); }
							}
						}
						else {
							for (auto& p : cakmers) {
								auto it = trKmers.find(p.first);
								if (it != trKmers.end()) { it->second += p.second; }
							}
						}
					}
				}
				else { destLocus = nloci; } // removed by threading

				if (aln and threading) {
					if ((aln_minimal and srcLocus != nloci and destLocus != nloci) or (not aln_minimal and (srcLocus != nloci or destLocus != nloci))) {
						alnindices.push_back(seqi); // work the same as extractindices
						edit.map = std::make_pair(srcLocus, destLocus);
						sam.push_back(edit);
					}
				}
			}
        }

        if (extractFasta or aln) {
            // write reads or alignments to STDOUT
            // begin thread lock
            sem_wait(semwriter); 
			if (extractFasta) {
				writeExtractedReads(extractFasta, seqs, titles, extractindices, assignedloci);
			}
			else if (aln) {
				if (skip1) { writeAlignments(seqs, titles, alnindices, sam); }
				else { writeAlignments(seqs, titles, destLoci, alnindices, sam); }
			}
            sem_post(semwriter);
            //
            // end of thread lock
        }

        cerr << "Batch query in " << (time(nullptr) - time2) << " sec. " << 
		        nShort_ << "/" <<
		        (float)nhash0/nReads_ << "/" << 
		        (float)nhash1/(nReads_ - nSubFiltered_) << "/" << 
		        nSubFiltered_ << "/" << 
		        nKmerFiltered_ << "/" << 
		        nThreadingReads_ << "/" << 
		        nFeasibleReads_ << endl;
    }
}


int main(int argc, char* argv[]) {

    if (argc < 2) {
        cerr << endl
             << "Usage: danbing-tk [-v] [-e] [-g|-gc] [-a|-ae] [-kf] [-cth] [-o|-on] -k -qs <-fai|-fa> -p" << endl
             << "Options:" << endl
             //<< "  -b               (deprecated) Use baitDB to decrease ambiguous mapping" << endl
             //<< "  -t <INT>         Used trimmed pangenome graph e.g. \"-t 1\" for pan.*.trim1.kmers" << endl
             //<< "  -s <INT>         Run in simulation mode to write the origin and destination of mis-assigned reads to STDOUT" << endl
             //<< "                   Specify 1 for simulated reads from TR" << endl
             //<< "                   Specify 2 for simulated reads from whole genome" << endl
			 //<< "  -m <str>         locusMap.tbl, used for mapping g-locus to pan-locus in whole genome simulation" << endl
             //<< "  -au              Augmentation mode, use pruned kmers and augkmers" << endl
		     << "  -v <INT>         Verbosity: 0-3. Default: 0." << endl
             << "  -e <INT>         Write mapped reads to STDOUT in fasta format." << endl
             << "                   Specify 1 for keeping original read names. Will not write .kmers output." << endl
			 << "                   Specify 2 for appending assigned locus to each read name. Used to skip step1 for later queries." << endl
             << "  -g <INT>         Use graph threading algorithm w/o error correction" << endl
             << "  -gc <INT>        Use graph threading algorithm w/ error correction" << endl
             << "                   Discard pe reads if # of matching kmers < [INT]" << endl
			 << "  -a               Output alignments for all reads entering threading. Only work with -g or -gc." << endl
			 << "  -ae              Same as the -a option, but excluding unaligned reads in threading." << endl
			 << "  -kf <INT> <INT>  Parameters for kmer-based pre-filtering," << endl
			 << "                   optimized for 150bp paired-end reads." << endl
			 << "                   1st param: # of sub-sampled kmers. Default: 4." << endl
			 << "                   2nd param: minimal # of matches. Default: 1." << endl
             << "  -cth <INT>       Discard both pe reads if maxhit of one pe read is below this threshold." << endl
			 << "                   Will skip read filtering and run threading directly if not specified." << endl
             << "  -o <STR>         Output prefix" << endl
		     << "  -on <STR>        Same as the -o option, but write locus and kmer name as well" << endl
             << "  -k <INT>         Kmer size" << endl
             << "  -qs <STR>        Prefix for *.tr.kmers, *.ntr.kmers, *.graph.kmers files" << endl
             << "  -fai <STR>       Interleaved pair-end fasta file" << endl
             << "  -fa <STR>        Fasta file e.g. generated by samtools fasta -n" << endl
             << "                   Reads will be paired on the fly" << endl
             << "  -p <INT>         Use n threads." << endl
             //<< "  -rth <FLOAT>     Discard reads with maxhit/(maxhit+secondhit) below this threshold." << endl 
             //<< "                   Range [0.5, 1]. 1: does not allow noise. 0.5: no filtering." << endl
             << endl;
        return 0;
    }
   
    vector<string> args(argv, argv+argc);
    bool bait = false, aug = false, threading = false, correction = false, aln = false, aln_minimal=false, g2pan = false, skip1 = true, writeKmerName = false, interleaved;
    int simmode = 0, extractFasta = 0;
    size_t argi = 1, trim = 0, thread_cth = 0, Cthreshold = 0, nproc;
    float Rthreshold = 0.5;
    string trPrefix, trFname, fastxFname, outPrefix;
    ifstream fastxFile, trFile, augFile, baitFile, mapFile;
    ofstream outfile, baitOut;
    while (argi < argc) {
        if (args[argi] == "-b") {
            bait = true;
            baitFile.open("baitDB.kmers");
            assert(baitFile);
            baitFile.close();
        }
		else if (args[argi] == "-v") { verbosity = stoi(args[++argi]); }
        else if (args[argi] == "-e") { extractFasta = stoi(args[++argi]); }
        else if (args[argi] == "-t") { trim = stoi(args[++argi]); }
        else if (args[argi] == "-s") { simmode = stoi(args[++argi]); }
		else if (args[argi] == "-m") {
			g2pan = true;
			mapFile.open(args[++argi]);
			assert(mapFile);
		}
        else if (args[argi] == "-au") { aug = true; }
        else if (args[argi] == "-g" or args[argi] == "-gc") {
            if (args[argi] == "-gc") { correction = true; }
            threading = true;
            thread_cth = stoi(args[++argi]);
        }
		else if (args[argi] == "-a") { aln = true; }
		else if (args[argi] == "-ae") { aln = true; aln_minimal = true; }
		else if (args[argi] == "-kf") {
			N_FILTER = stoi(args[++argi]);
			NM_FILTER = stoi(args[++argi]);
		}
        else if (args[argi] == "-k") { ksize = stoi(args[++argi]); }
        else if (args[argi] == "-qs") {
            trPrefix = args[++argi];
            trFname = (trim ? trPrefix+".tr.trim"+std::to_string(trim)+".kmers" : trPrefix+".tr.kmers");
            trFile.open(trFname);
            assert(trFile);
            trFile.close();
            if (aug) {
                augFile.open(trPrefix+".tr.aug.kmers");
                assert(augFile);
                augFile.close();
            }
        }
        else if (args[argi] == "-fa" or args[argi] == "-fai") {
            interleaved = args[argi] == "-fai";
            fastxFname = args[++argi];
            fastxFile.open(fastxFname);
            assert(fastxFile);
        }
        else if (args[argi] == "-o" or args[argi] == "-on") {
			writeKmerName = args[argi] == "-on";
            outPrefix = args[++argi];
            outfile.open(outPrefix+".tr.kmers");
            assert(outfile);
            outfile.close();
            if (bait) {
                baitOut.open(outPrefix+".cntm");
                assert(baitOut);
                baitOut.close();
            }
        }
        else if (args[argi] == "-p") { nproc = stoi(args[++argi]); }
        else if (args[argi] == "-cth") { 
			Cthreshold = stoi(args[++argi]);
			skip1 = false;
		}
        else if (args[argi] == "-rth") {
            Rthreshold = stof(args[++argi]);
            assert(Rthreshold <= 1 and Rthreshold >= 0.5);
        }
        else { 
            cerr << "invalid option: " << args[argi] << endl;
			throw;
        }
        ++argi;
    }

    // report parameters
    cerr << "use baitDB: " << bait << endl
         << "extract fasta: " << extractFasta << endl
         << "interleaved: " << interleaved << endl
         << "sim mode: " << simmode << endl
         << "trim mode: " << trim << endl
         << "augmentation mode: " << aug << endl
         << "graph threading mode: " << threading << endl
         << "output alignment: " << aln << endl
         << "output successfully aligned reads only: " << aln_minimal << endl
         << "k: " << ksize << endl
         << "# of subsampled kmers in pre-filtering: " << N_FILTER << endl
         << "minimal # of matches in pre-filtering: " << NM_FILTER << endl
         << "Cthreshold: " << Cthreshold << endl
         << "Rthreshold: " << Rthreshold << endl
         << "threading Cthreshold: " << thread_cth << endl
         << (skip1 ? "Thread reads directly (step2: threading) without filtering (step1: kmer-based filtering)" : "Running both step1 (kmer-based filtering) and step2 (threading)") << endl
         << "fastx: " << fastxFname << endl
         << "query: " << trPrefix << ".(tr/ntr).kmers" << endl
         << endl
         << "total number of loci in " << trFname << ": ";
    size_t nloci = countLoci(trFname);
    cerr << nloci << endl;


    // read input files
    time_t time1 = time(nullptr);
    vector<kmer_aCount_umap> trKmerDB(nloci);
    vector<GraphType> graphDB(nloci);
    kmerIndex_uint32_umap kmerDBi;
	vector<uint32_t> kmerDBi_vv;

	unordered_map<string, string> readDB;
    vector<msa_umap> msaStats;
	err_umap errdb;
	vector<size_t> locusmap;

	if (extractFasta) { // step 1
		readBinaryIndex(kmerDBi, kmerDBi_vv, trPrefix);
		cerr << "deserialized index in " << (time(nullptr) - time1) << " sec." << endl;
		cerr << "# unique kmers in kmerDBi: " << kmerDBi.size() << endl;
	} else if (skip1) { // step 2
		readBinaryGraph(graphDB, trPrefix);
		readKmers(trKmerDB, trFname);
		cerr << "deserialized graph and read tr.kmers in " << (time(nullptr) - time1) << " sec." << endl;
	} else { // step 1+2
		readBinaryIndex(kmerDBi, kmerDBi_vv, trPrefix);
		readBinaryGraph(graphDB, trPrefix);
		readKmers(trKmerDB, trFname);
		cerr << "deserialized graph/index and read tr.kmers in " << (time(nullptr) - time1) << " sec." << endl;
		cerr << "# unique kmers in kmerDBi: " << kmerDBi.size() << endl;
	}

    // create data for each process
    cerr << "creating data for each process..." << endl;
    time1 = time(nullptr);
    Threads threaddata(nproc, nloci);
    size_t nReads = 0, nThreadingReads = 0, nFeasibleReads = 0, nSubFiltered = 0, nKmerFiltered = 0;
    for (size_t i = 0; i < nproc; ++i) {
        Counts &counts = threaddata.counts[i];

        counts.in = &fastxFile;
        counts.readDB = &readDB;
        counts.trResults = &trKmerDB;
        counts.graphDB = &graphDB;
        counts.kmerDBi = &kmerDBi;
		counts.kmerDBi_vv = &kmerDBi_vv;
        counts.nReads = &nReads;
        counts.nThreadingReads = &nThreadingReads;
        counts.nFeasibleReads = &nFeasibleReads;
        counts.nSubFiltered = &nSubFiltered;
        counts.nKmerFiltered = &nKmerFiltered;
        counts.msaStats = &msaStats;
        counts.errdb = &errdb;
		counts.locusmap = &locusmap;

        counts.interleaved = interleaved;
        counts.extractFasta = extractFasta;
        counts.bait = bait;
        counts.simmode = simmode;
        counts.threading = threading;
        counts.correction = correction;
		counts.aln = aln;
		counts.g2pan = g2pan;
		counts.skip1 = skip1;

        counts.Cthreshold = Cthreshold;
        counts.Rthreshold = Rthreshold;
        counts.thread_cth = thread_cth;
    }

    const int idLen=10;
    char id[idLen+1];
    id[idLen] = '\0';
    srand (time(NULL));
    rand_str(id, idLen);

    string readerName = string("/semreader_") + string(id);
    string countName  = string("/semcount_") + string(id);
    string semwriterName = string("/semwriter_") + string(id);

    semreader = sem_open(readerName.c_str(), O_CREAT, 0644, 1);
    if (semreader == NULL) {
        cerr << "ERROR opening semaphore. ERRNO " << errno << " " << readerName << endl;
        exit(1);
    }
    semcount = sem_open(countName.c_str(), O_CREAT, 0644, 1);
    if (semreader == NULL) {
        cerr << "ERROR opening semaphore. ERRNO " << errno << " " << countName << endl;
        exit(1);
    }
    semwriter = sem_open(semwriterName.c_str(), O_CREAT, 0644, 1);
    if (semwriter == NULL) {
        cerr << "ERROR opening semaphore. ERRNO " << errno << " " << semwriterName << endl;
        exit(1);
    }

	//ProfilerStart("prefilter.v10.prof");
    sem_init(semreader, 1, 1);
    sem_init(semcount, 1, 1);
    sem_init(semcount, 1, 1);

    pthread_attr_t *threadAttr = new pthread_attr_t[nproc];

    for (size_t t = 0; t < nproc; ++t) {
        pthread_attr_init(&threadAttr[t]);
    }
    pthread_t *threads = new pthread_t[nproc];

    // start computing
    for (size_t t = 0; t < nproc; ++t) {
		pthread_create(&threads[t], &threadAttr[t], (void* (*)(void*))CountWords<size_t>, &threaddata.counts[t]);
    }
    cerr << "threads created" << endl;
 
    for (size_t t = 0; t < nproc; ++t) {
        pthread_join(threads[t], NULL);
    }
	//ProfilerFlush();
	//ProfilerStop();

    cerr << nReads << " reads processed in total." << endl
	     << nSubFiltered << " reads removed by subsampled kmer-filter." << endl
	     << nKmerFiltered << " reads removed by kmer-filter." << endl
         << nThreadingReads << " reads entered threading step." << endl
         << nFeasibleReads << " reads passsed threading." << endl
         << "parallel query completed in " << (time(nullptr) - time1) << " sec." << endl;
    fastxFile.close();

    // write outputs
    if (not extractFasta) {
		cerr << "writing kmers..." << endl;
		if (writeKmerName) { writeKmersWithName(outPrefix+".tr", trKmerDB); }
		else { writeKmers(outPrefix+".tr", trKmerDB); }
	}

    cerr << "all done!" << endl;
    return 0;
}
```
