## Supplemental code for "The motif composition of variable-number tandem repeats impacts gene expression": aQueryFasta_thread.h

```
#ifndef A_QUERYFASTA_THREAD_H_
#define A_QUERYFASTA_THREAD_H_

#include "cereal/archives/binary.hpp"
#include "cereal/types/unordered_map.hpp"
#include "cereal/types/vector.hpp"

#include "stdlib.h"
#include <vector>
#include <string>
#include <iostream>
#include <fstream>
#include <sstream>
#include <numeric>
#include <fcntl.h>
#include <errno.h>
#include <time.h>
#include <map>
#include <unordered_map>
#include <set>
#include <unordered_set>
#include <cassert>
#include <cmath>
#include <algorithm>
#include <iomanip>
#include <atomic>

using namespace std;

//typedef unordered_set<size_t> kmer_set;
typedef unordered_map<size_t, uint32_t> kmerCount_umap; // assume count < (2^16 -1)
typedef unordered_map<size_t, atomic_size_t> kmer_aCount_umap;
typedef unordered_map<size_t, uint8_t> GraphType;
typedef unordered_map<size_t, unordered_set<uint32_t>> kmeruIndex_umap; // assume number of loci < (2^32 -1)
//typedef unordered_map<size_t, uint32_t> kmerIndex_uint32_umap; // assume number of loci < (2^32 -1)
typedef unordered_map<size_t, size_t> kmerIndex_uint32_umap;
typedef unordered_map<size_t, vector<uint16_t>> kmerAttr_dict;
typedef unordered_map<string, unordered_map<string, uint16_t>> adj_dict;
typedef unordered_map<string, unordered_map<string, vector<uint16_t>>> adjAttr_dict;
typedef unordered_map<size_t, unordered_map<size_t, uint16_t>> nuAdj_dict;
typedef unordered_map<size_t, unordered_map<size_t, vector<uint16_t>>> nuAdjAttr_dict;

//const unordered_map<char, size_t> base( {{'A', 0}, {'C', 1}, {'G', 2}, {'T', 3}});
//const char baseinv[] = {'A', 'C', 'G', 'T'};
//const unordered_map<char, char> Cbase( {{'A', 'T'}, {'C', 'G'}, {'G', 'C'}, {'T', 'A'}, {'N', 'N'}} );
const char alphabet[] = {'A', 'C', 'G', 'T'};
const char baseNumConversion[] = {
  'A','C','G','T',127,127,127,127,
  127,127,127,127,127,127,127,127,
  127,127,127,127,127,127,127,127,
  127,127,127,127,127,127,127,127,
  127,127,127,127,127,127,127,127,
  127,127,127,127,127,127,127,127,
  127,127,127,127,127,127,127,127,
  127,127,127,127,127,127,127,127,
  127, 0 ,127, 1 ,127,127,127, 2 ,
  127,127,127,127,127,127,127,127,
  127,127,127,127, 3 ,127,127,127,
  127,127,127,127,127,127,127,127,
  127, 0 ,127, 1 ,127,127,127, 2 ,
  127,127,127,127,127,127,127,127,
  127,127,127,127, 3 ,127,127,127,
};

const char baseComplement[] = {
    3,  2,  1,  0,127,127,127,127,
  127,127,127,127,127,127,127,127,
  127,127,127,127,127,127,127,127,
  127,127,127,127,127,127,127,127,
  127,127,127,127,127,127,127,127,
  127,127,127,127,127,127,127,127,
  127,127,127,127,127,127,127,127,
  127,127,127,127,127,127,127,127,
  127,'T',127,'G',127,127,127,'C',
  127,127,127,127,127,127,'N',127,
  127,127,127,127,'A',127,127,127,
  127,127,127,127,127,127,127,127,
  127,'t',127,'g',127,127,127,'c',
  127,127,127,127,127,127,'n',127,
  127,127,127,127,'a',127,127,127,
};
const unsigned char byteRC[]   = {
 255, 191, 127,  63, 239, 175, 111,  47, 223, 159,
  95,  31, 207, 143,  79,  15, 251, 187, 123,  59,
 235, 171, 107,  43, 219, 155,  91,  27, 203, 139,
  75,  11, 247, 183, 119,  55, 231, 167, 103,  39,
 215, 151,  87,  23, 199, 135,  71,   7, 243, 179,
 115,  51, 227, 163,  99,  35, 211, 147,  83,  19,
 195, 131,  67,   3, 254, 190, 126,  62, 238, 174,
 110,  46, 222, 158,  94,  30, 206, 142,  78,  14,
 250, 186, 122,  58, 234, 170, 106,  42, 218, 154,
  90,  26, 202, 138,  74,  10, 246, 182, 118,  54,
 230, 166, 102,  38, 214, 150,  86,  22, 198, 134,
  70,   6, 242, 178, 114,  50, 226, 162,  98,  34,
 210, 146,  82,  18, 194, 130,  66,   2, 253, 189,
 125,  61, 237, 173, 109,  45, 221, 157,  93,  29,
 205, 141,  77,  13, 249, 185, 121,  57, 233, 169,
 105,  41, 217, 153,  89,  25, 201, 137,  73,   9,
 245, 181, 117,  53, 229, 165, 101,  37, 213, 149,
  85,  21, 197, 133,  69,   5, 241, 177, 113,  49,
 225, 161,  97,  33, 209, 145,  81,  17, 193, 129,
  65,   1, 252, 188, 124,  60, 236, 172, 108,  44,
 220, 156,  92,  28, 204, 140,  76,  12, 248, 184,
 120,  56, 232, 168, 104,  40, 216, 152,  88,  24,
 200, 136,  72,   8, 244, 180, 116,  52, 228, 164,
 100,  36, 212, 148,  84,  20, 196, 132,  68,   4,
 240, 176, 112,  48, 224, 160,  96,  32, 208, 144,
  80,  16, 192, 128,  64,   0};


string decodeNumericSeq(size_t num, size_t k){
    string seq = "";
    for (size_t i = 0; i < k; ++i){
        seq = baseNumConversion[num % 4] + seq;
        num >>= 2;
    }
    return seq;
}
    
size_t encodeSeq(string& seq, size_t start, size_t k) { // no extra copy
    size_t numericSeq = 0;
    for (size_t i = start; i < start+k; ++i){
        numericSeq = (numericSeq<<2) + baseNumConversion[seq[i]];
    }
    return numericSeq;
}

size_t getNextKmer(size_t& kmer, size_t beg, string& read, size_t k){
    size_t rlen = read.size();
    if (beg + k > rlen){
        return rlen;
    }
    size_t validlen = 0;
    while (validlen != k){
        if (beg + k > rlen){
            return rlen;
        }
        if (find(alphabet, alphabet+4, read[beg + validlen]) == alphabet+4){
            beg = beg + validlen + 1;
            validlen = 0;
        } else {
            validlen += 1;
        }
    }
    kmer = encodeSeq(read, beg, k);
    return beg;
}

string getRC(const string &read) {
    string rcread;
    size_t rlen = read.size();
    rcread.resize(rlen);
    for (size_t i = 0; i < rlen; ++i) {
        rcread[i] = baseComplement[read[rlen - 1 - i]];
    }
    return rcread;
}

size_t getNuRC(size_t num, size_t k) {
    size_t num_rc = 0;
    while (k >= 4) { // convert a full byte
        num_rc <<= 8;
        num_rc += byteRC[num & 0xff];
        num >>= 8;
        k -= 4;
    }
    if (k > 0) { // convert remaining bits
        num_rc <<= (k<<1); // was num_rc <<= (k*2);
        num_rc += (byteRC[num] >> ((4-k)<<1)); // was num_rc += (byteRC[num] >> ((4-k)*2));
    }
    return num_rc;
}

inline size_t toCaKmer(size_t kmer, size_t k) {
	size_t rckmer = getNuRC(kmer, k);
	return kmer < rckmer ? kmer : rckmer;
}

template <typename T>
void buildNuKmers(T& kmers, string& read, size_t k, size_t leftflank = 0, size_t rightflank = 0, bool count = true) {
    size_t rlen = read.size();
    size_t mask = (1UL << 2*(k-1)) - 1;
    size_t beg, nbeg, canonicalkmer, kmer, rckmer;

    beg = getNextKmer(kmer, leftflank, read, k);
    if (beg == rlen){ return; }
    rckmer = getNuRC(kmer, k);
 
    for (size_t i = beg; i < rlen - k - rightflank + 1; ++i){
        if (kmer > rckmer) {
            canonicalkmer = rckmer;
        } else {
            canonicalkmer = kmer;
        }
        kmers[canonicalkmer] += (1 & count);

        if (find(alphabet, alphabet+4, read[i + k]) == alphabet+4){
            nbeg = getNextKmer(kmer, i+k+1, read, k);
            if (nbeg == rlen) { return; }
            rckmer = getNuRC(kmer, k);
            i = nbeg - 1;
        } else {
            kmer = ( (kmer & mask) << 2 ) + baseNumConversion[read[i + k]];
            rckmer = (rckmer >> 2) + ( (baseNumConversion[baseComplement[read[i + k]]] & mask) << (2*(k-1))); // XXX test correctness
        }
    }
}

void _buildKmerGraph(GraphType& g, string& read, size_t k, size_t leftflank, size_t rightflank, bool noselfloop) {
    const size_t rlen = read.size();
    const size_t mask = (1ULL << 2*(k-1)) - 1;

    size_t beg, nbeg, kmer;
    beg = getNextKmer(kmer, leftflank, read, k);
    if (beg != rlen){
        for (size_t i = beg; i < rlen - k - rightflank; ++i){
            if (std::find(alphabet, alphabet+4, read[i + k]) == alphabet+4){
                g[kmer] |= 0;
                nbeg = getNextKmer(kmer, i+k+1, read, k);
                if (nbeg == rlen) { break; }
                i = nbeg - 1;
            } else {
                size_t nextkmer = ((kmer & mask) << 2) + baseNumConversion[read[i + k]];
                bool valid = (not noselfloop) or (noselfloop and (kmer != nextkmer));
                g[kmer] |= ((1 & valid) << baseNumConversion[read[i + k]]);
                kmer = nextkmer;
            }
        }
        g[kmer] |= 0;
    }
}

void buildKmerGraph(GraphType& g, string& read, size_t k, size_t leftflank = 0, size_t rightflank = 0, bool noselfloop = true) {
    _buildKmerGraph(g, read, k, leftflank, rightflank, noselfloop);
    string rcread = getRC(read);
    _buildKmerGraph(g, rcread, k, rightflank, leftflank, noselfloop);
}

// invalid kmers are skipped by default unless keepN is set; input/output size differs
void read2kmers(vector<size_t>& kmers, string& read, size_t k, size_t leftflank = 0, size_t rightflank = 0, bool canonical = true, bool keepN = false) {
    const size_t rlen = read.size();
    const size_t mask = (1ULL << 2*(k-1)) - 1;
    size_t beg, nbeg, canonicalkmer, kmer, rckmer;

    beg = getNextKmer(kmer, leftflank, read, k);
    if (beg == rlen){ return; }
    rckmer = getNuRC(kmer, k);

    for (size_t i = beg; i < rlen - k - rightflank + 1; ++i){
        canonicalkmer = (kmer > rckmer ? rckmer : kmer);
        kmers.push_back(canonical ? canonicalkmer : kmer); // XXX speedup; resize not push_back

        if (std::find(alphabet, alphabet+4, read[i + k]) == alphabet+4){ // XXX speedup
            nbeg = getNextKmer(kmer, i+k+1, read, k);
			if (keepN) { for (size_t j = i+1; j < std::min(nbeg, rlen-k-rightflank+1); ++j) { kmers.push_back(-1); } } // XXX test
            if (nbeg == rlen) { return; }
            rckmer = getNuRC(kmer, k);
            i = nbeg - 1;
        } else {
            kmer = ( (kmer & mask) << 2 ) + baseNumConversion[read[i + k]];
            rckmer = (rckmer >> 2) + ( (baseNumConversion[baseComplement[read[i + k]]] & mask) << (2*(k-1)));
        }
    }
}

// Applied to a read pair to accumulate counts.
// Assuming maximal kmer counts from a VNTR read pair does not exceed 32 bits
void noncaVec2CaUmap(vector<size_t>& kmers, kmerCount_umap& out, size_t ksize) {
    size_t RCkmer;
    for (size_t kmer : kmers) {
		if (kmer == -1ULL) { continue; }
        RCkmer = getNuRC(kmer, ksize);
        ++out[(kmer <= RCkmer ? kmer : RCkmer)];
    }
}

size_t countLoci(string fname) {
    ifstream inf(fname);
    assert(inf);
    string line;
    size_t nloci = 0;
    while (getline(inf, line)) {
        if (line[0] == '>'){
            ++nloci;
        }
    }
    inf.close();
    return nloci;
}

size_t countBedLoci(string fname) {
    ifstream inf(fname);
    assert(inf);
    string line;
    size_t nloci = 0;
    while (getline(inf, line)) {
        ++nloci;
    }
    inf.close();
    return nloci;

}

// record kmerDB only
template <typename T>
void readKmers(T& kmerDB, string fname) {
    ifstream f(fname);
    assert(f);
    cerr <<"reading kmers from " << fname << endl;
    string line;
	size_t idx = -1;
	while (getline(f, line)) {
		if (line[0] == '>') { ++idx; }
		else { kmerDB[idx][stoul(line)] += 0; }
	}
	f.close();
}

template <typename T>
void readGraphKmers(T& kmerDB, string fname) {
    ifstream f(fname);
    assert(f);
    cerr <<"reading kmers from " << fname << endl;
	size_t idx = 0;
    string line;
    getline(f, line);
    while (true){
        if (f.peek() == EOF or f.peek() == '>'){
            ++idx;
            if (f.peek() == EOF){
                f.close();
                break;
            } else {
                getline(f, line);
            }
        } else {
            getline(f, line, '\t');
            size_t kmer = stoul(line);
            getline(f, line);
            size_t c = stoul(line);
			kmerDB[idx][kmer] |= c;
        }
    }
    f.close();
}

template <typename T>
void readKmersFile2DB(T& kmerDB, string fname, bool graph=false, bool count=true, size_t startInd=0, uint16_t threshold=0, uint16_t offset=0) {
    ifstream f(fname);
    assert(f);
    string line;
    getline(f, line);
    cerr <<"reading kmers from " << fname << endl;
    while (true){
        if (f.peek() == EOF or f.peek() == '>'){
            ++startInd;
            if (f.peek() == EOF){
                f.close();
                break;
            } else {
                getline(f, line);
            }
        } else {
            getline(f, line, '\t');
            size_t kmer = stoul(line);
            getline(f, line);
            size_t c = stoul(line);

            if (c < threshold) { continue; }
            if (count) {
				if (graph) {
					kmerDB[startInd][kmer] |= c;
				} else {
                	kmerDB[startInd][kmer] += (c + offset);
				}
            } else {
                kmerDB[startInd][kmer] += 0;
            }
        }
    }
    f.close();
}

// record kmerDB only; use orthology map to assign locus
template <typename T>
void mapKmersFile2DB(T& kmerDB, string fname, vector<bool>& omap, bool count=true, bool graph=false, uint16_t threshold=0, uint16_t offset=0) {
    ifstream f(fname);
    assert(f);
    string line;
    getline(f, line);
    cerr <<"reading kmers from " << fname << endl;
	int idx = -1;
    while (true){
        if (f.peek() == EOF or f.peek() == '>'){
            ++idx;
			while (not omap[idx]) { ++idx; }
            if (f.peek() == EOF){
                f.close();
                break;
            } else {
                getline(f, line);
            }
        } else {
            getline(f, line, '\t');
            size_t kmer = stoul(line);
            getline(f, line);
            size_t c = stoul(line);

            if (c < threshold) { continue; }
            if (count) {
				if (graph) {
					kmerDB[idx][kmer] |= c;
				} else {
                	kmerDB[idx][kmer] += (c + offset);
				}
            } else {
                kmerDB[idx][kmer] += 0;
            }
        }
    }
    f.close();
}

// record kmerIndex_dict kmerDBi only
//void readKmersFile2DBi(kmeruIndex_umap& kmerDBi, string fname, size_t startInd = 0, uint16_t threshold = 0) {
//    ifstream f(fname);
//    assert(f);
//    string line;
//    getline(f, line);
//    cerr <<"reading kmers from " << fname << endl;
//    while (true){
//        if (f.peek() == EOF or f.peek() == '>'){
//            ++startInd;
//            if (f.peek() == EOF){
//                f.close();
//                break;
//            } else {
//                getline(f, line);
//            }
//        } else {
//            getline(f, line, '\t');
//            size_t kmer = stoul(line);
//            getline(f, line);
//            size_t kmercount = stoul(line);
//
//            if (kmercount < threshold) { continue; }
//            if (kmerDBi[kmer].count(startInd) == 0) {
//                kmerDBi[kmer].insert(startInd);
//            }
//        }
//    }
//    f.close();
//}

void readBinaryIndex(kmerIndex_uint32_umap& kmerDBi, vector<uint32_t>& kmerDBi_vv, string& pref) {
	{
		cerr << "deserializing kmerDBi.umap" << endl;
		ifstream fin(pref+".kmerDBi.umap", ios::binary);
		assert(fin);
		cereal::BinaryInputArchive iarchive(fin);
		iarchive(kmerDBi);
	}
	{
		cerr << "deserializing kmerDBi.vv" << endl;
		ifstream fin(pref+".kmerDBi.vv", ios::binary);
		assert(fin);
		cereal::BinaryInputArchive iarchive(fin);
		iarchive(kmerDBi_vv);
	}
}

void readBinaryGraph(vector<GraphType>& graphDB, string& pref) {
	cerr << "deserializing graph.umap" << endl;
	ifstream fin(pref+".graph.umap", ios::binary);
	assert(fin);
	cereal::BinaryInputArchive iarchive(fin);
	iarchive(graphDB);
}


void readKmerIndex(kmerIndex_uint32_umap& kmerDBi, vector<vector<uint32_t>>& kmerDBi_vec, string fname) { // optimized version
    ifstream f(fname);
    assert(f);
    cerr <<"reading kmers from " << fname << endl;
	uint32_t idx = -1;
    uint32_t vsize = kmerDBi_vec.size();
    string line;
    while (getline(f, line)) {
        if (line[0] == '>') { ++idx; }
        else { 
			size_t kmer = stoul(line);
			auto it = kmerDBi.find(kmer);
            if (it != kmerDBi.end()) { // kmer is not unique
                uint32_t vi = it->second;
                if (vi % 2) { // kmer freq. >= 2 in current db; kmerDBi_vec[(vi>>1)] records the list of mapped loci
                    bool good = true;
                    for (uint32_t x : kmerDBi_vec[vi>>1]) { if (x == idx) { good = false; break; } }
                    if (good) { kmerDBi_vec[vi>>1].push_back(idx); }
                }
                else { // kmer freq. = 1 in current db; vi>>1 is the mapped locus.
                    if ((vi >> 1) != idx) {
                        kmerDBi_vec.push_back(vector<uint32_t>{vi>>1, idx});
                        it->second = ((vsize++) << 1) + 1;
                    }
                }
            } else {
                kmerDBi[kmer] = (idx << 1);
            }
		}
    }
    f.close();
}

void readKmersFile2DBi(kmerIndex_uint32_umap& kmerDBi, vector<vector<uint32_t>>& kmerDBi_vec, string fname) { // optimized version
    ifstream f(fname);
    assert(f);
    string line;
    getline(f, line);
    cerr <<"reading kmers from " << fname << " using optimized readKmersFile2DBi()" << endl;
	uint32_t vsize = kmerDBi_vec.size(), idx = 0;
    while (true){
        if (f.peek() == EOF or f.peek() == '>'){
            ++idx;
            if (f.peek() == EOF){
				f.close();
                break;
            } else {
                getline(f, line);
            }
        } else {
            getline(f, line, '\t');
            size_t kmer = stoul(line);
            getline(f, line);
            size_t kmercount = stoul(line);

			auto it = kmerDBi.find(kmer);
			if (it != kmerDBi.end()) { // kmer is not unique
				uint32_t vi = it->second;
				if (vi % 2) { // kmer freq. >= 2 in current db; kmerDBi_vec[(vi>>1)] records the list of mapped loci
					bool good = true;
					for (uint32_t x : kmerDBi_vec[vi>>1]) { if (x == idx) { good = false; break; } }
					if (good) { kmerDBi_vec[vi>>1].push_back(idx); }
				}
				else { // kmer freq. = 1 in current db; vi>>1 is the mapped locus.
					if ((vi >> 1) != idx) {
						kmerDBi_vec.push_back(vector<uint32_t>{vi>>1, idx});
						it->second = ((vsize++) << 1) + 1;
					}
				}
			} else {
				kmerDBi[kmer] = (idx << 1);
			}
        }
    }
    f.close();
}

// record kmerDB and kmerIndex_dict kmerDBi
template <typename T>
void readKmersWithIndex(T& kmerDB, kmerIndex_uint32_umap& kmerDBi, vector<vector<uint32_t>>& kmerDBi_vec, string fname) { // optimized version
    ifstream f(fname);
    assert(f);
    cerr <<"reading kmers from " << fname << endl;
    uint32_t idx = -1;
    uint32_t vsize = kmerDBi_vec.size();
    string line;
    while (getline(f, line)) {
        if (line[0] == '>') { ++idx; }
        else {
            size_t kmer = stoul(line);
            kmerDB[idx][kmer] += 0;

            auto it = kmerDBi.find(kmer);
            if (it != kmerDBi.end()) { // kmer is not unique
                uint32_t vi = it->second;
                if (vi % 2) { // kmer freq. >= 2 in current db; kmerDBi_vec[(vi>>1)] records the list of mapped loci
                    bool good = true;
                    for (uint32_t x : kmerDBi_vec[vi>>1]) { if (x == idx) { good = false; break; } }
                    if (good) { kmerDBi_vec[vi>>1].push_back(idx); }
                }
                else { // kmer freq. = 1 in current db; vi>>1 is the mapped locus.
                    if ((vi >> 1) != idx) {
                        kmerDBi_vec.push_back(vector<uint32_t>{vi>>1, idx});
                        it->second = ((vsize++) << 1) + 1;
                    }
                }
            } else {
                kmerDBi[kmer] = (idx << 1);
            }
        }
    }
    f.close();
}

template <typename T>
void readKmersFile(T& kmerDB, kmerIndex_uint32_umap& kmerDBi, vector<vector<uint32_t>>& kmerDBi_vec, string fname, bool count = true) { // optimized version
    ifstream f(fname);
    assert(f);
    string line;
    getline(f, line);
    cerr <<"reading kmers from " << fname << " using optimized readKmersFile()" << endl;
	uint32_t vsize = kmerDBi_vec.size(), idx = 0;
    while (true){
        if (f.peek() == EOF or f.peek() == '>'){
            ++idx;
            if (f.peek() == EOF){
				f.close();
                break;
            } else {
                getline(f, line);
            }
        } else {
            getline(f, line, '\t');
            size_t kmer = stoul(line);
            getline(f, line);
            size_t kmercount = stoul(line);

            if (count) {
                kmerDB[idx][kmer] += kmercount;
            } else {
                kmerDB[idx][kmer] += 0;
            }
			auto it = kmerDBi.find(kmer);
            if (it != kmerDBi.end()) { // kmer is not unique
                uint32_t vi = it->second;
                if (vi % 2) { // kmer freq. >= 2 in current db; kmerDBi_vec[(vi>>1)] records the list of mapped loci
					bool good = true;
                    for (uint32_t x : kmerDBi_vec[vi>>1]) { if (x == idx) { good = false; break; } }
                    if (good) { kmerDBi_vec[vi>>1].push_back(idx); }
                }
                else { // kmer freq. = 1 in current db; vi>>1 is the mapped locus.
					if ((vi >> 1) != idx) {
                        kmerDBi_vec.push_back(vector<uint32_t>{vi>>1, idx});
                        it->second = ((vsize++) << 1) + 1;
                    }
                }
            } else {
				kmerDBi[kmer] = (idx << 1);
            }
        }
    }
    f.close();
}

template <typename T>
void readKmersFile(T& kmerDB, kmeruIndex_umap& kmerDBi, string fname, size_t startInd = 0, bool count = true, uint16_t threshold = 0) {
    ifstream f(fname);
    assert(f);
    string line;
    getline(f, line);
    cerr <<"reading kmers from " << fname << endl;
    while (true){
        if (f.peek() == EOF or f.peek() == '>'){
            ++startInd;
            if (f.peek() == EOF){
                f.close();
                break;
            } else {
                getline(f, line);
            }
        } else {
            getline(f, line, '\t');
            size_t kmer = stoul(line);
            getline(f, line);
            size_t kmercount = stoul(line);

            if (kmercount < threshold) { continue; }
            if (count) {
                kmerDB[startInd][kmer] += kmercount;
            } else {
                kmerDB[startInd][kmer] += 0;
            }
            if (kmerDBi[kmer].count(startInd) == 0) {
                kmerDBi[kmer].insert(startInd);
            }
        }
    }
    f.close();
}

template <typename T>
void writeKmersWithName(string outfpref, T& kmerDB, size_t threshold = 0) {
    ofstream fout(outfpref+".kmers");
    assert(fout);
    for (size_t i = 0; i < kmerDB.size(); ++i) {
        fout << ">" << i <<"\n";
        for (auto &p : kmerDB[i]) {
            if (p.second < threshold) { continue; }
            fout << p.first << '\t' << (size_t)p.second << '\n';
        }
    }
    fout.close();
}

template <typename T>
void writeKmers(string outfpref, T& kmerDB, size_t threshold = 0) {
    ofstream fout(outfpref+".kmers");
    assert(fout);
    for (size_t i = 0; i < kmerDB.size(); ++i) {
        for (auto &p : kmerDB[i]) {
            if (p.second < threshold) { continue; }
            fout << (size_t)p.second << '\n';
        }
    }
    fout.close();
}

void writeKmers(string outfpref, vector<kmerAttr_dict>& kmerAttrDB) {
    ofstream fout(outfpref+".kmers");
    assert(fout);
    for (size_t i = 0; i < kmerAttrDB.size(); ++i) {
        fout << ">" << i <<"\n";
        for (auto &p : kmerAttrDB[i]) {
            fout << p.first;
            for (auto &q : p.second) {
                fout << '\t' << q;
            }
            fout << '\n';
        }
    }
    fout.close();
}

void readOrthoMap(string& mapf, vector<vector<bool>>& omap, size_t nhap) {
    ifstream fin(mapf);
    assert(fin);
    string line;
    size_t idx = 0;
    while (getline(fin, line)) {
        omap.push_back(vector<bool>(nhap));
        stringstream ss(line);
        for (size_t i = 0; i < nhap; ++i) {
            string v;
            ss >> v;
            omap[idx][i] = v != ".";
        }
        ++idx;
    }
    fin.close();
}

tuple<adj_dict, size_t> buildAdjDict(kmerCount_umap& kmers, size_t k) {
    adj_dict adj;
    string s, t;
    size_t max = 0, m;
    for (auto& p : kmers){
        s = decodeNumericSeq(p.first >> 2, k-1);
        t = decodeNumericSeq(p.first % (1UL << 2*(k-1)), k-1);
        m = kmers[p.first];
        adj[s][t] += m;
        if (m > max){
            max = m;
        }
    }
    return make_tuple(adj, max);
}

void writeDot(string outfpref, size_t i, adj_dict &adj) {
    ofstream fout;
    fout.open(outfpref + ".loci." + to_string(i) + ".dot");
    assert(fout.is_open());
    fout << "strict digraph \"\" {" << '\n';
    for (auto& p : adj){
        for (auto& q : p.second){
            fout << p.first << " -> " << q.first << " [Weight = \"   " << q.second << "\", ";
            fout << "penwidth = " << q.second << "];" << '\n';
        }
    }
    fout << "}";
    fout.close();
}

void writeDot(string outfpref, int i, adjAttr_dict &adjAttr) { // function polymorphism: adj with attributes information
    // can only compare 2 graphs at this moment
    ofstream fout;
    if (i == -1) {
        fout.open(outfpref + ".diff.dot");
    } else {
        fout.open(outfpref + "." + to_string(i) + ".unique.dot");
    }
    assert(fout.is_open());
    fout << "strict digraph \"\" {" << '\n';
    for (auto& p : adjAttr){
        for (auto& q : p.second){
            fout << p.first << " -> " << q.first << " [Weight = " << q.second[0];
            fout << ", Label = " << to_string(q.second[1]) << "];" << '\n';
        }
    }
    fout << "}";
    fout.close();
}

class DBG {
public:
    size_t nkmers;
    size_t setid = 0, nset = 0, maxcount = 0;

    adj_dict adj;
    adjAttr_dict adjAttr;
	
    unordered_map<string, size_t> sets;          // store which node belongs which set in adj
    vector<vector<string>> nodes;                // store what nodes are in each set
    vector<size_t> setsizes;                     // store the size (edge#) of each set in adj/adj_rc

    DBG(size_t nkmers_) : nkmers(nkmers_), nodes(nkmers_), setsizes(nkmers_) {}

    size_t getAdj(string &node, string &node_rc) {
        if (sets.count(node) == 1) 
            { return 1; }
        else if (sets.count(node_rc) == 1)
            { return 2; }
        else
            { return 0; }
    }
        
    void updatesets(string *s, string *t, int mode, int oldlabel = -1, int newlabel = -1) {

        if (mode == 2) {        // new set is created

            sets[*s] = setid;
            sets[*t] = setid;
            nodes[setid].push_back(*s);
            nodes[setid].push_back(*t);
            setsizes[setid] += 2;
            ++setid;
            ++nset;

        }
        else if (mode == 1) {    // s already exists; s -> t

            sets[*t] = sets[*s];
            nodes[sets[*t]].push_back(*t);
            setsizes[setid] += 1;

        }
        else if (mode == 0) {    // t already exists; s -> t

            sets[*s] = sets[*t];
            nodes[sets[*s]].push_back(*s);
            setsizes[setid] += 1;

        }
        else if (mode == -1) {                  // s, t in different sets of the same graph

            // change the label of the smaller set
            if (setsizes[sets[*s]] >= setsizes[sets[*t]]) {
                oldlabel = sets[*t];
                newlabel = sets[*s];
            }
            else {
                oldlabel = sets[*s];
                newlabel = sets[*t];
            }

            setsizes[newlabel] += setsizes[oldlabel];
            for (auto &node : nodes[oldlabel]) {
                sets[node] = newlabel;
            }

            nodes[newlabel].insert(nodes[newlabel].end(), 
                make_move_iterator(nodes[oldlabel].begin()),
                make_move_iterator(nodes[oldlabel].end()));
            nodes[oldlabel].clear();
            nset--;

        }
        else if (mode == -2) {                  // s, t in different sets and graphs

            setsizes[newlabel] += setsizes[oldlabel];
            for (auto &node : nodes[oldlabel]) {
                sets.erase(node);
                sets[getRC(node)] = newlabel;
            }
            // get reverse complement of each node
            vector<string> tmpnodes(nodes[oldlabel].size());
            for (size_t i = 0; i < nodes[oldlabel].size(); ++i) {
                tmpnodes[i] = getRC(nodes[oldlabel][i]);
            }
            // insert (t_rc, s_rc) as (source, target) in newlabel
            nodes[newlabel].insert(nodes[newlabel].end(), 
                make_move_iterator(tmpnodes.begin()),
                make_move_iterator(tmpnodes.end()));       
            nodes[oldlabel].clear();
            nset--;

    	}
    }

    void swapsubgraph(size_t label, bool isAttr = false) {
        if (isAttr) { // adj with attributes information
            adjAttr_dict tmpadjAttr;
            for (auto &node : nodes[label]) {
                string node_rc = getRC(node);
                if (adjAttr.count(node) != 0) { // equivelant to if (node is a source_node in adj)
                    for (auto &p : adjAttr[node]) {
                        string target_rc = getRC(p.first);
                        tmpadjAttr[target_rc][node_rc] = p.second;
                    }
                    adjAttr.erase(node);
                }
            }
            for (auto &p : tmpadjAttr) {
                for (auto &q : p.second) {
                    adjAttr[p.first][q.first] = q.second;
                }
            }
        }
        else {
            adj_dict tmpadj;
            for (auto &node : nodes[label]) {
                string node_rc = getRC(node);
                if (adj.count(node) != 0) { // equivelant to if (node is a source_node in adj)
                    for (auto &p : adj[node]) {
                        string target_rc = getRC(p.first);
                        tmpadj[target_rc][node_rc] = p.second;
                    }
                    adj.erase(node);
                }
            }
            for (auto &p : tmpadj) {
                for (auto &q : p.second) {
                    adj[p.first][q.first] += q.second;
                }
            }
        }
   }

    void updateDBG(int sInAdj, int tInAdj, string &s, string &t, string &s_rc, string &t_rc, uint16_t count) {

        if (sInAdj == 0 and tInAdj == 0) {		// s, t not in graph; create a new set
            updatesets(&s, &t, 2);
            adj[s][t] += count;
        }
        else if (sInAdj == 0 or tInAdj == 0) {		// either s or t in graph; expand existing set
            if (sInAdj == 1 or tInAdj == 1) {		// forward strand in adj
                if (tInAdj == 1)			// t is in the adj
                    { updatesets(&s, &t, 0); }
                else					// s is in the adj
                    { updatesets(&s, &t, 1); }
                adj[s][t] += count;
            }
            else {					// forward strand in adj_rc
                if (tInAdj == 2)
                    { updatesets(&t_rc, &s_rc, 1); }
                else
                    { updatesets(&t_rc, &s_rc, 0); }
                adj[t_rc][s_rc] += count;
            }
        }
        else if (sInAdj == tInAdj) {			// both s, t in the same graph
            if (sInAdj == 1) {						// s, t in adj
                if (sets[s] != sets[t])			// s, t in different sets
                    { updatesets(&s, &t, -1); }		// combine s, t sets 
            	else
                    { setsizes[sets[s]]++; }
                adj[s][t] += count;
            }
            if (sInAdj == 2)  {				// s, t in adj_rc
                if (sets[s_rc] != sets[t_rc])
                    { updatesets(&s_rc, &t_rc, -1); }
                else
                    { setsizes[sets[t_rc]]++; }
                adj[t_rc][s_rc] += count;
            }
        }
        else {						// s, t in different graphs
            if (sInAdj == 1) {				// s in adj, t in adj_rc
                if (sets[s] == sets[t_rc]) {		// s and t_rc in the same set; add edge
                    updatesets(&s, &t, 1);
                    adj[s][t] += count;
                }
                else {					// s and t_rc in different sets
                    size_t label1 = sets[s];
                    size_t label2 = sets[t_rc];
                    if (setsizes[label1] >= setsizes[label2]) {     // swap label2 subgraph and connect s with t
                        swapsubgraph(label2);
                        updatesets(NULL, NULL, -2, label2, label1); // merge label2 to label1
                        adj[s][t] += count;
                    }
                    else {
                        swapsubgraph(label1);
                        updatesets(NULL, NULL, -2, label1, label2);
                        adj[t_rc][s_rc] += count;
                    }
                }
            }
            else {                                      // sInAdj == 2
                if (sets[s_rc] == sets[t]) {
                    updatesets(&s, &t, 1);
                    adj[s][t] += count;
                }
                else {
                    size_t label1 = sets[s_rc];
                    size_t label2 = sets[t];
                    if (setsizes[label1] >= setsizes[label2]) {
                        swapsubgraph(label2);
                        updatesets(NULL, NULL, -2, label2, label1);
                        adj[t_rc][s_rc] += count;
                    }
                    else {
                        swapsubgraph(label1);
                        updatesets(NULL, NULL, -2, label1, label2);
                        adj[s][t] += count;
                    }
                }
            }
        }
    }

    void updateDBG(int sInAdj, int tInAdj, string &s, string &t, string &s_rc, string &t_rc, vector<uint16_t> &attr) {
    // function polymorphism: adj with attributes information

        if (sInAdj == 0 and tInAdj == 0) {		// s, t not in graph; create a new set
            updatesets(&s, &t, 2);
            adjAttr[s][t] = attr;                      // "=" sign only works when kmer size is odd; should implement elementwise += in the future
        }
        else if (sInAdj == 0 or tInAdj == 0) {		// either s or t in graph; expand existing set
            if (sInAdj == 1 or tInAdj == 1) {		// forward strand in adj
                if (tInAdj == 1)			// t is in the adj
                    { updatesets(&s, &t, 0); }
                else					// s is in the adj
                    { updatesets(&s, &t, 1); }
                adjAttr[s][t] = attr;
            }
            else {					// forward strand in adj_rc
                if (tInAdj == 2)
                    { updatesets(&t_rc, &s_rc, 1); }
                else
                    { updatesets(&t_rc, &s_rc, 0); }
                adjAttr[t_rc][s_rc] = attr;
            }
        }
        else if (sInAdj == tInAdj) {				// both s, t in the same graph
            if (sInAdj == 1) {					// s, t in adj
                if (sets[s] != sets[t])				// s, t in different sets
                    { updatesets(&s, &t, -1); }			// combine s, t sets 
            	else
                    { setsizes[sets[s]]++; }
                adjAttr[s][t] = attr;
            }
            if (sInAdj == 2)  {					// s, t in adj_rc
                if (sets[s_rc] != sets[t_rc])
                    { updatesets(&s_rc, &t_rc, -1); }
                else
                    { setsizes[sets[t_rc]]++; }
                adjAttr[t_rc][s_rc] = attr;
            }
        }
        else {								// s, t in different graphs
            if (sInAdj == 1) {						// s in adj, t in adj_rc
                if (sets[s] == sets[t_rc]) {		                // s and t_rc in the same set; add edge
                    updatesets(&s, &t, 1);
                    adjAttr[s][t] = attr;
                }
                else {							// s and t_rc in different sets
                    size_t label1 = sets[s];
                    size_t label2 = sets[t_rc];
                    if (setsizes[label1] >= setsizes[label2]) {         // swap label2 subgraph and connect s with t
                        swapsubgraph(label2, true);
                        updatesets(NULL, NULL, -2, label2, label1);     // merge label2 to label1
                        adjAttr[s][t] = attr;
                    }
                    else {
                        swapsubgraph(label1, true);
                        updatesets(NULL, NULL, -2, label1, label2);
                        adjAttr[t_rc][s_rc] = attr;
                    }
                }
            }
            else {                                          // sInAdj == 2
                if (sets[s_rc] == sets[t]) {
                    updatesets(&s, &t, 1);
                    adjAttr[s][t] = attr;
                }
                else {
                    size_t label1 = sets[s_rc];
                    size_t label2 = sets[t];
                    if (setsizes[label1] >= setsizes[label2]) {
                        swapsubgraph(label2, true);
                        updatesets(NULL, NULL, -2, label2, label1);
                        adjAttr[t_rc][s_rc] = attr;
                    }
                    else {
                        swapsubgraph(label1, true);
                        updatesets(NULL, NULL, -2, label1, label2);
                        adjAttr[s][t] = attr;
                    }
                }
            }
        }
    }

    void addkmer(const string &kmer, uint16_t count) {
        string s = kmer.substr(0, kmer.size() - 1);
        string t = kmer.substr(1, kmer.size() - 1);
        string s_rc = getRC(s);
        string t_rc = getRC(t);
        int sInAdj = getAdj(s, s_rc);
        int tInAdj = getAdj(t, t_rc);
        updateDBG(sInAdj, tInAdj, s, t, s_rc, t_rc, count);
        if (count > maxcount) { maxcount = count; }
    }

    void addkmer(const string &kmer, vector<uint16_t> &attr) { // function polymorphism: adj with attributes information
        string s = kmer.substr(0, kmer.size() - 1);
        string t = kmer.substr(1, kmer.size() - 1);
        string s_rc = getRC(s);
        string t_rc = getRC(t);
        int sInAdj = getAdj(s, s_rc);
        int tInAdj = getAdj(t, t_rc);
        updateDBG(sInAdj, tInAdj, s, t, s_rc, t_rc, attr);
        if (attr[0] > maxcount) { maxcount = attr[0]; }
    }

}; // class DBG


class BiDBG {
public:
    BiDBG(bool hasAttr_, size_t nkmers_, size_t ksize_) : hasAttr(hasAttr_), nkmers(nkmers_), ksize(ksize_) {}

    void addkmer(const string &kmer, uint16_t count) {
        string s, t, s_rc, t_rc;
        getNodes(kmer, s, t, s_rc, t_rc);
        adj[s][t] += count;
        adj[t_rc][s_rc] += count;
        if (count > maxcount) { maxcount = count; }
    }

    void addkmer(const string &kmer, vector<uint16_t> &attr) { // function polymorphism: adj with attributes information
        string s, t, s_rc, t_rc;
        getNodes(kmer, s, t, s_rc, t_rc);
        adjAttr[s][t] = attr;          // "=" sign works only when kmer size is odd
        adjAttr[t_rc][s_rc] = attr;
        if (attr[0] > maxcount) { maxcount = attr[0]; }
    }

    void addkmer(size_t kmer, uint16_t count) { // function polymorphism: numeric adj
        size_t s, t, s_rc, t_rc;
        getNodes(kmer, s, t, s_rc, t_rc);
        nuAdj[s][t] += count;
        nuAdj[t_rc][s_rc] += count;
        if (count > maxcount) { maxcount = count; }
    }

    void addkmer(size_t kmer, vector<uint16_t> &attr) { // function polymorphism: numeric adj with attributes information
        size_t s, t, s_rc, t_rc;
        getNodes(kmer, s, t, s_rc, t_rc);
        nuAdjAttr[s][t] = attr;          // "=" sign works only when kmer size is odd
        nuAdjAttr[t_rc][s_rc] = attr;
        if (attr[0] > maxcount) { maxcount = attr[0]; }
    }

    void getAdj(adj_dict& out) { out = adj; }

    void getAdj(adjAttr_dict& out) { out = adjAttr; }

    void getAdj(nuAdj_dict& out) { out = nuAdj; }

    void getAdj(nuAdjAttr_dict& out) { out = nuAdjAttr; }

    size_t getMaxCount() { return maxcount; }

    // TODO:
    // void checkGraphIntegrity () {}
    // check the number of subgraphs (<= 2)

private:
    size_t nkmers, ksize;
    size_t maxcount = 0;
    bool hasAttr; // if true: record other attributes other than counts

    adj_dict adj;
    adjAttr_dict adjAttr;
    nuAdj_dict nuAdj;
    nuAdjAttr_dict nuAdjAttr;

    void getNodes(const string& kmer, string& s, string& t, string& s_rc, string& t_rc) {
        s = kmer.substr(0, kmer.size() - 1);
        t = kmer.substr(1, kmer.size() - 1);
        s_rc = getRC(s);
        t_rc = getRC(t);
    }

    void getNodes(size_t kmer, size_t& s, size_t& t, size_t& s_rc, size_t& t_rc) {
        s = kmer >> 2;
        t = kmer % (1UL << (2*(ksize-1)));
        s_rc = getNuRC(s, ksize);
        t_rc = getNuRC(t, ksize);
    }

}; // class BiDBG

#endif
```
