## Supplemental code for "The motif composition of variable-number tandem repeats impacts gene expression": bam2pe.cpp

```
#include "aQueryFasta_thread.h"

#include <ctime>


struct Read {
    string seq;
    uint8_t start, len;
};

void prunePEinfo(string& title) {
	size_t endi = title.size();
	if (title[endi-2] == '/') {
		if (title[endi-1] == '1' or title[endi-1] == '2') {
			title = title.substr(0,endi-2);
		}
	}
}

int main(int argc, char* argv[]) {
    if (argc < 2) {
        cerr << endl
             << "Usage: bam2pe -fai <infile>" << endl << endl

             << "Options:" << endl
             << "  -fai     Input file from samtools fasta -n" << endl << endl;
        return 1;
    }
    vector<string> args(argv, argv+argc);

    size_t ind_fai = distance(args.begin(), find(args.begin(), args.begin()+argc, "-fai"));
    assert(ind_fai != argc);
    size_t ind_fname = ind_fai + 1;
    string fname = args[ind_fname];

    cerr << "fname: " << fname << endl;

    ifstream fin(fname);
    assert(fin);

    unordered_map<string, Read> readDB;
    size_t nread = 0, nPEread = 0;
    size_t batch = 10000000, PEbatch = 1000000;
    time_t time1 = time(nullptr);

	while (fin.good()) {
		string title, seq, qualtitle, qual;
		Read read;

		getline(fin, title);
		getline(fin, read.seq);
		prunePEinfo(title);

		read.start = 0;
		read.len = read.seq.size();

		if (readDB.count(title)) {
			Read& read1 = readDB[title];

			if (read.len < 1 or read1.len < 1) { continue; }
			else {
				// output PE reads
				cout << title << "_0\n";
				//cout << ">read " << to_string(nPEread) << "_0\n";
				cout << read.seq.substr(read.start, read.len) << '\n';
				cout << title << "_1\n";
				//cout << ">read " << to_string(nPEread) << "_1\n";
				cout << read1.seq.substr(read1.start, read1.len) << '\n';

				nPEread += 2;
				if (nPEread % PEbatch == 0) {
					cerr << "time: " << time(nullptr) - time1 << " sec. " << nPEread << " PE reads found \
							with " << readDB.size() << " reads in container " << endl;
					time1 = time(nullptr);
				}
				readDB.erase(title);
			}
		}
		else {
			readDB[title] = read;
		}
		nread++;
		if (nread % batch == 0) {
			cerr << nread << " reads processed" << endl;
		}
	}
	return 0;
}
```
