## Supplemental code for "The motif composition of variable-number tandem repeats impacts gene expression": genPanKmers.cpp

```
#include "aQueryFasta_thread.h"

#include <iostream>
#include <string>
#include <fstream>
#include <sstream>
#include <unordered_map>
#include <tuple>
#include <cmath>
#include <vector>
#include <cassert>
#include <algorithm>

using namespace std;

// skip content in fin until target locus and merge content in target locus with the content in graph
void readGraphLocus(GraphType& graph, ifstream& fin, size_t& current, size_t target) {
    string line;
    while (current < target) {
        getline(fin, line);
        if (line[0] == '>') { ++current; }
    }

    while (current == target) {
        if (fin.peek() == '>' or fin.peek() == EOF) { 
            getline(fin, line);
            ++current;
            return;
        }
        else {
            getline(fin, line, '\t');
            size_t kmer = stoul(line);
            getline(fin, line, '\n');
            uint8_t nucBits = stoul(line);
            graph[kmer] |= nucBits;
        }
    }
}

void readKmersLocus(kmerCount_umap& kmers, ifstream& fin, size_t& current, size_t target) {
    string line;
    while (current < target) {
        getline(fin, line);
        if (line[0] == '>') { ++current; }
    }

    while (current == target) {
        if (fin.peek() == '>' or fin.peek() == EOF) {
            getline(fin, line);
            ++current;
            return;
        }
        else {
            getline(fin, line, '\t');
            size_t kmer = stoul(line);
            getline(fin, line, '\n');
            kmers[kmer] = 0;
        }
    }
}

void getgmap(vector<vector<bool>>& omap, vector<bool>& gmap, vector<size_t> his) {
	for (size_t i = 0; i < gmap.size(); ++i) {
		bool good = false;
		for (size_t hi : his) {
			good |= omap[i][hi];
		}
		gmap[i] = good;
	}
}

int main(int argc, const char * argv[]) {

    if (argc < 2) {
        cerr << "usage: program  -o <output_prefix>  -m <mapping>  -k <kmer_file_prefixes>\n"
             << "  -m       if is '-', the program assumes no missing loci\n"
             << "           full path name for <mapping> is required in any case\n"
             << "  -k       requires PREFIX.TYPE.kmers\n"
             << "           TYPE = tr, ntr or graph\n"
             << "mapping file format:\n"
             << "   N columns; each column is a genome; order should be the same as specified in -k\n"
             << "   M rows; each row is a locus in the pan-genome (pan locus)\n"
             << "           NUMBER is the ordering of the pan locus in that genome\n"
             << "           '.' means the pan locus in missing that genome\n\n";
        return 0;
    }

    vector<string> args(argv, argv+argc);
    bool nomissing = false;
    size_t argi = 1, ngenome = argc-6, nloci;
    string indir, outpref, mapfname;
    vector<string> kmerpref(ngenome);

    while (argi < argc) {
        if (args[argi] == "-o") { outpref = args[++argi]; }
        else if (args[argi] == "-m") {
            mapfname = args[++argi];
            nomissing = (mapfname == "-");
        }
        else if (args[argi] == "-k") {
            kmerpref.assign(argv+argi+1, argv+argc);
            break;
        }
        else {
            cerr << "Error: invalid option " << args[argi] << '\n';
            return 1;
        }
        ++argi;
    }

	vector<vector<bool>> omap;
	if (not nomissing){
		readOrthoMap(mapfname, omap, 2*ngenome);
    	nloci = omap.size();
	} else {
		nloci = countLoci(kmerpref[0]+".tr.kmers");
	}
    cerr << "# loci in pangenome: " << nloci << endl
         << ngenome << " genomes to merge" << endl;

    vector<string> filetypes = {"tr", "ntr", "graph"};
    for (string& filetype : filetypes) {
        cerr << "merging " << filetype << ".kmers" << endl;

        bool graphmode = (filetype == "graph");
        vector<kmerCount_umap> kmersDB;
        vector<GraphType> graphDB;
        if (graphmode) { graphDB.resize(nloci); }
        else { kmersDB.resize(nloci); }

        for (size_t gi = 0; gi < ngenome; ++gi) {
			vector<bool> gmap(nloci, 1);
			if (not nomissing) {
				getgmap(omap, gmap, vector<size_t>{2*gi,2*gi+1});
				if (graphmode) {
					mapKmersFile2DB(graphDB, kmerpref[gi]+"."+filetype+".kmers", gmap, true);
				} else {
					mapKmersFile2DB(kmersDB, kmerpref[gi]+"."+filetype+".kmers", gmap);
				}
			}
			else {
				if (graphmode) {
					readKmersFile2DB(graphDB, kmerpref[gi]+"."+filetype+".kmers", true);
				} else {
					readKmersFile2DB(kmersDB, kmerpref[gi]+"."+filetype+".kmers");
				}
			}
        } 

        cerr << "writing " << filetype << ".kmers" << endl;
        if (graphmode) { 
			writeKmers(outpref+"."+filetype, graphDB);
		} else { 
			writeKmers(outpref+"."+filetype, kmersDB);
		}
    }
    
    

    return 0;
}
```
