## Supplemental code for "The motif composition of variable-number tandem repeats impacts gene expression": kmertools.cpp

```
#include "aQueryFasta_thread.h"
#include "cereal/archives/binary.hpp"
#include "cereal/types/unordered_map.hpp"
#include "cereal/types/vector.hpp"

#include <iostream>
#include <fstream>
#include <string>
#include <vector>
#include <cassert>
#include <ctime>


using std::cout;
using std::cerr;
using std::endl;
using std::ifstream;
using std::ofstream;
using std::string;
using std::vector;

int main (int argc, const char * argv[]) {
	
	if (argc == 1) {
		cerr << "Usage: ktools <commands> [options]" << endl << endl

		     << "Commands:" << endl
			 << "  ksi         generate ksi index for ktools sum" << endl
		     << "  sum         acculumate kmer counts for each locus" << endl
		     << "  serialize   generate kmer index using pan.(graph|ntr|tr).kmers" << endl << endl;
		return 0;
	}


	vector<string> args(argv, argv+argc);
	if (args[1] == "ksi") {
		if (argc == 2) {
			cerr << "Usage: ktools ksi <pan.tr.kmers> >$OUT.ksi" <<
			        "  Generate ksi index for ktools sum" << endl;
			return 0;
		}

		ifstream kmers(args[2]);
		string line;
		int idx = -1, nkmer = 0;
		while (getline(kmers, line)) {
			if (line[0] == '>') {
				++idx; 
				if (idx) { cout << nkmer << '\n'; }
			} else {
				++nkmer;
			}
		}
		if (idx) { cout << nkmer << '\n'; }
	}

	else if (args[1] == "sum") {
		if (argc == 2) {
			cerr << "Usage 1: ktools sum <.ksi> <.kmers> <out.kms>\n" <<
			        "  Read a single .kmers file and write a single column output.\n" <<
			        "Usage 2: ktools sum -f <.ksi> <.txt> <out.kms>\n" << 
			        "  Read all kmer files specified in .txt and output a kms table (row=sample, col=locus)." << endl;
			return 0;
		}

		if (args[2] == "-f") {
			ifstream ksif(args[3]);
            ifstream fofn(args[4]);
            ofstream fout(args[5]);
            assert(ksif);
            assert(fofn);
            assert(fout);

            vector<size_t> ksi;
            string line;
            while (getline(ksif, line)) {
                ksi.push_back(stoul(line));
            }
			ksif.close();
            cerr << ksi.size() << " loci in " << args[3] << endl;

			vector<string> kmerfs;
			while (getline(fofn, line)) {
				kmerfs.push_back(line);
			}
			fofn.close();
			cerr << kmerfs.size() << " samples in " << args[4] << endl;

			size_t ki;
			for (size_t fi = 0; fi < kmerfs.size(); ++fi) {
	            ifstream kmerf(kmerfs[fi]);
				assert(kmerf);
				size_t idx = 0, kms = 0;
				ki = 0;
				while (getline(kmerf, line)) {
					kms += stoul(line);
					++ki;
					while (ksi[idx] == ki) {
						++idx;
						if (idx != ksi.size()) { fout << kms << '\t'; kms = 0; }
						else { fout << kms << '\n'; break; } 
					}
				}
				kmerf.close();
			}
			fout.close();
            cerr << ki << " kmers processed in each file" << endl;
		} 
		else {
			ifstream ksif(args[2]);
			ifstream kmerf(args[3]);
			ofstream fout(args[4]);
			assert(ksif);
			assert(kmerf);
			assert(fout);

			vector<size_t> ksi;
			string line;
			while (getline(ksif, line)) {
				ksi.push_back(stoul(line));
			}
			cerr << ksi.size() << " loci in " << args[2] << endl;

			size_t idx = 0, ki = 0, kms = 0;
			while (getline(kmerf, line)) {
				kms += stoul(line);
				++ki;
				while (ksi[idx] == ki) { ++idx; fout << kms << '\n'; kms = 0; if (idx == ksi.size()) {break;}}
			}
			ksif.close();
			kmerf.close();
			fout.close();
			cerr << idx << " loci and " << ki << " kmers processed in " << args[3] << endl;
		}
	}
	else if (args[1] == "serialize") {
		if (argc == 2) {
			cerr << "Usage: ktools serialize <pref>" << endl << endl

			     << "  PREF     prefix of *.(graph|ntr|tr).kmers" << endl;
			return 0;
		}

        size_t nloci = countLoci(args[2]+".tr.kmers");
		//vector<kmer_aCount_umap> trKmerDB;
		vector<GraphType> graphDB(nloci);
        //{
        //    clock_t t = clock();
        //    readGraphKmers(trKmerDB, args[2]+".tr.kmers");
        //    cerr << "tr.kmers read in " << (float)(clock()-t) / CLOCKS_PER_SEC << " sec" << endl;
        //    cerr << "serializing tr.umap" << endl;
        //    ofstream fout(args[2]+".tr.umap", ios::binary);
        //    assert(fout);
        //    cereal::BinaryOutputArchive oarchive(fout);
        //    oarchive(trKmerDB);
        //}
        //{
		//	vector<kmer_aCount_umap> trKmerDB_copy;
        //    ifstream fin(args[2]+".tr.umap", ios::binary);
        //    assert(fin);
        //    cereal::BinaryInputArchive iarchive(fin);
        //    clock_t t = clock();
        //    iarchive(trKmerDB_copy);
        //    cerr << "tr.umap deserialized in " << (float)(clock()-t) / CLOCKS_PER_SEC << " sec" << endl;
        //    cerr << "validating tr.umap" << endl;
        //    for (size_t i = 0; i < nloci; ++i) {
        //        for (auto& p : trKmerDB[i]) {
        //            auto it = trKmerDB_copy[i].find(p.first);
        //            assert(it != trKmerDB_copy[i].end());
        //            assert(it->second == p.second);
        //        }
        //    }

        //}

        {
			clock_t t = clock();
            readGraphKmers(graphDB, args[2]+".graph.kmers");
			cerr << "graph.kmers read in " << (float)(clock()-t) / CLOCKS_PER_SEC << " sec" << endl;
            cerr << "serializing graph.umap" << endl;
            ofstream fout(args[2]+".graph.umap", ios::binary);
            assert(fout);
            cereal::BinaryOutputArchive oarchive(fout);
            oarchive(graphDB);
        }
		{
			vector<GraphType> graphDB_copy(nloci);
			ifstream fin(args[2]+".graph.umap", ios::binary);
            assert(fin);
            cereal::BinaryInputArchive iarchive(fin);
			clock_t t = clock();
            iarchive(graphDB_copy);
			cerr << "graph.umap deserialized in " << (float)(clock()-t) / CLOCKS_PER_SEC << " sec" << endl;
			cerr << "validating graph.umap" << endl;
			for (size_t i = 0; i < nloci; ++i) {
				for (auto& p : graphDB[i]) { 
					auto it = graphDB_copy[i].find(p.first);
					assert(it != graphDB_copy[i].end());
					assert(it->second == p.second);
				}
			}

		}

        kmerIndex_uint32_umap kmerDBi;
        vector<vector<uint32_t>> kmerDBi_vec;
		{
			clock_t t = clock();
			readKmerIndex(kmerDBi, kmerDBi_vec, args[2]+".tr.kmers");
			readKmerIndex(kmerDBi, kmerDBi_vec, args[2]+".ntr.kmers");
			cerr << "xtr.kmers read in " << (float)(clock()-t) / CLOCKS_PER_SEC << " sec" << endl;
		}

        cerr << "generating kmerDBi.vv (non-unique kmer container)" << endl;
        vector<uint32_t> vv;
        vector<uint32_t> vvi;
        for (vector<uint32_t>& v : kmerDBi_vec) {
            vvi.push_back(vv.size());
            vv.push_back(v.size());
            vv.insert(vv.end(), v.begin(), v.end());
        }

        cerr << "reindexing kmerDBi.umap (unique kmer container)" << endl;
        for (auto& p : kmerDBi) {
            if (p.second % 2) {
                assert(kmerDBi_vec[p.second>>1].size() == vv[vvi[p.second>>1]]);
                size_t i = 1;
                for (uint32_t v : kmerDBi_vec[p.second>>1]) { assert(v == vv[vvi[p.second>>1]+i]); ++i; }
                p.second = (vvi[p.second>>1] << 1) + 1;
            }
            //cout << p.first << '\t' << p.second << endl;
        }

        cerr << "serializing kmerDBi.umap" << endl;
        {
            ofstream fout(args[2]+".kmerDBi.umap", ios::binary);
            assert(fout);
            cereal::BinaryOutputArchive oarchive(fout);
            oarchive(kmerDBi);
        }

        cerr << "serializing kmerDBi.vv" << endl;
        {
            ofstream fout(args[2]+".kmerDBi.vv", ios::binary);
            assert(fout);
            cereal::BinaryOutputArchive oarchive(fout);
            oarchive(vv);
        }

		{
			kmerIndex_uint32_umap kmerDBi_copy;
			vector<uint32_t> vv_copy;
			clock_t t = clock();
			{
				cerr << "deserializing kmerDBi.umap" << endl;
				ifstream fin(args[2]+".kmerDBi.umap", ios::binary);
				assert(fin);
				cereal::BinaryInputArchive iarchive(fin);
				iarchive(kmerDBi_copy);
			}
			{
				cerr << "deserializing kmerDBi.vv" << endl;
				ifstream fin(args[2]+".kmerDBi.vv", ios::binary);
				assert(fin);
				cereal::BinaryInputArchive iarchive(fin);
				iarchive(vv_copy);
			}
			cerr << "kmerDBi.(umap|vv) deserialized in " << (float)(clock()-t) / CLOCKS_PER_SEC << " sec" << endl;

			cerr << "validating kmerDBi.umap" << endl;
			for (auto& p : kmerDBi) {
				auto it = kmerDBi_copy.find(p.first);
				assert(it != kmerDBi_copy.end());
				assert(it->second == p.second);
			}
			cerr << "validating kmerDBi.vv" << endl;
			for (size_t i = 0; i < vv.size(); ++i) { assert(vv[i] == vv_copy[i]); }
		}
		cerr << "Done!" << endl;

		// 1-step method
		//readKmersWithIndex(trKmerDB, kmerDBi, kmerDBi_vec, trFname);
		//readKmerIndex(kmerDBi, kmerDBi_vec, trPrefix+".ntr.kmers");
		//readGraphKmers(graphDB, trPrefix+".graph.kmers");


		// 2-step method
		//readKmerIndex(kmerDBi, kmerDBi_vec, trFname);
		//readKmerIndex(kmerDBi, kmerDBi_vec, trPrefix+".ntr.kmers");

		//readTRKmers(trKmerDB, trFname);
		//readGraphKmers(graphDB, trPrefix+".graph.kmers");


		//cerr << "# unique kmers in kmerDBi: " << kmerDBi.size() << '\n';
		//cerr << "# unique kmers in kmerDBi: " << kmerDBi.size() << '\n';
		//cerr << "read *.kmers file in " << (time(nullptr) - time1) << " sec." << endl;
	}
	else {
		cerr << "Unrecognized command " << args[1] << endl;
		return 0;
	}


	return 0;
}
```
