## Supplemental code for "The motif composition of variable-number tandem repeats impacts gene expression": mapkmers.cpp

```
#include "aQueryFasta_thread.h"

#include <iostream>
#include <string>
#include <fstream>
#include <sstream>
#include <unordered_map>
#include <tuple>
#include <cmath>
#include <vector>
#include <cassert>
#include <algorithm>

using namespace std;


int main(int argc, const char * argv[]) {

    if (argc < 2) {
        //cerr << "usage: program panbed pankmers kmers genome outpref\n";
        cerr << "usage: program panbed target_field pankmers kmers outpref\n\n"

             << "target_field   column index (0-indexed) to extract in panbed for locus mapping\n\n";
        return 0;
    }

    vector<string> args(argv, argv+argc);
    string panbedfname = args[1], pankmersfname = args[3], kmersfname = args[4], outpref = args[5];
    size_t tf = stoi(args[2]);

    ifstream panbedin(panbedfname);
    assert(panbedin);

    size_t npanloci = countBedLoci(panbedfname);
    size_t nloci = countLoci(kmersfname);
    vector<kmerCount_umap> pankmersDB(npanloci);
    vector<kmerCount_umap> kmersDB(nloci);

    ifstream panfin(pankmersfname), fin(kmersfname);
    assert(panfin and fin);
    panfin.close();
    fin.close();

    readKmersFile2DB(pankmersDB, pankmersfname);
    readKmersFile2DB(kmersDB, kmersfname, 0, false); // start from 0th locus, do not count

    cerr << "mapping kmers" << endl;
    string line;
    size_t panlocus = 0, locus = 0;
    while (getline(panbedin, line)) {
        stringstream ss(line);
        string tmp;

        for (size_t ind = 0; ind < tf; ++ind) { ss >> tmp; }
        ss >> tmp;
        if (tmp != ".") {
            locus = stoul(tmp);
            assert(locus < nloci);
            kmerCount_umap& kmers = kmersDB[locus];
            kmerCount_umap& pankmers = pankmersDB[panlocus];
            for (auto& p : kmers) {
                kmers[p.first] = pankmers[p.first];
            }
        }
        ++panlocus;
    }

    cerr << "writing kmers" << endl;
    writeKmers(outpref, kmersDB);

    return 0;
}
```
