## Supplemental code for "The motif composition of variable-number tandem repeats impacts gene expression": Num2seq.cpp

```
#include "nuQueryFasta.h"

int main (int argc, const char * argv[]) {
    
    if (argc < 2) {
        cerr << "usage: num2seq <sequence length> <numeric sequence>\n";
        cerr << "  e.g. num2seq 5 0\n";
        cerr << "       output: AAAAA\n";
        cerr << "       or pipe data through previous command\n";
        cerr << "  e.g. echo \"0\" | num2seq 5\n";
        cerr << "       output: AAAAA\n";
        exit(0);
    }

    vector<string> args(argv, argv+argc);
    size_t num;
    size_t k = stoi(args[1]);
    
    if (argc == 2) {
        while (cin >> num) {
            cout << decodeNumericSeq(num, k) << endl;
        }
        return 0;
    }

    if (args[2] == "/dev/stdin") {
        ifstream inf(args[2]);
        string line;
        while (true) {
            if (inf.peek() == EOF) { break; }
            getline(inf, line);
            num = stoul(line);
            cout << decodeNumericSeq(num, k) << endl;
        }
    }
    else {
        num = stoul(args[2]);
        cout << decodeNumericSeq(num, k) << endl;
    }


    return 0;
}
```
