## Supplemental code for "The motif composition of variable-number tandem repeats impacts gene expression": RVseq.cpp

```
#include "nuQueryFasta.h"

#include <iostream>

int main (int argc, const char * argv[]) {
	
	//if (argc < 2) {
	//	cerr << "usage: rvseq <sequence>\n";
	//	cerr << "  e.g. rvseq AAATTTGGGCCC\n";
	//	cerr << "       output: GGGCCCAAATTT\n\n";
	//}

	if (argc == 1) {
		string seq;
		while (cin >> seq) {
			cout << getRC(seq) << endl;
		}
		return 0;
	}

	string args1 = string(argv[1]);

    if (args1 == "/dev/stdin") {
        ifstream inf(args1);
        string line;
        while (true) {
            if (inf.peek() == EOF) { break; }
            getline(inf, line);
            cout << getRC(line) << endl;
        }
    }
    else {
		cout << getRC(args1) << endl;
	}


	return 0;
}
```
