## Supplemental code for "The motif composition of variable-number tandem repeats impacts gene expression": Seq2num.cpp

```
#include "nuQueryFasta.h"

#include <iostream>
#include <fstream>

int main (int argc, const char * argv[]) {
	
	//cout << "usage: input seqeunce e.g. ATCGGTG to get its numeric represnetation" << endl;
	//cout << "       where (A,C,G,T) = (0,1,2,3)" << endl;
	//cout << "       enter q to quit program" << endl << endl;

	string args1 = string(argv[1]);
	if (args1 == "/dev/stdin") {
		ifstream inf(args1);
		string line;
		while (true) {
			if (inf.peek() == EOF) { break; }
			getline(inf, line);
			cout << encodeSeq(line) << endl;
		}
	}
	else {
		cout << encodeSeq(args1) << endl;
	}
	return 0;
}
```
