## Supplemental code for "The motif composition of variable-number tandem repeats impacts gene expression": VNTR2kmers_thread.cpp

```
//
//  VNTR2kmers.cpp
//  convert VNTR loci to .kmer and generate .dot for corresponding DBG
//
//  Created by Tsung-Yu Lu on 6/5/18.
//  Copyright © 2018 Tsung-Yu Lu. All rights reserved.
//
#include "aQueryFasta_thread.h"

#include <iostream>
#include <string>
#include <fstream>
#include <sstream>
#include <unordered_map>
#include <tuple>
#include <cmath>
#include <vector>
#include <cassert>
#include <algorithm>

using namespace std;
size_t ksize;


void removeNodeFromGraph(size_t node, GraphType& graph) { // XXX test edge pruning 
	static const size_t MASK = (1UL << (2*ksize)) - 1 - 3; // keep k-1 prefix
	static const size_t PREF = 1UL << (2*(ksize-1));

    if (graph.count(node)) { // if the kmer is in the graph
        graph.erase(node); // remove the node

        uint8_t nucmask = 0xff - (1 << (node % 4));
        size_t km1mer = (node & MASK) >> 2;
        for (size_t nuc = 0; nuc < 4; ++nuc) { // check all possible upstream nodes
            size_t prevkmer = nuc * PREF + km1mer;
            if (graph.count(prevkmer)) { graph[prevkmer] &= nucmask; } // remove the edge that points from the upstream node
        }
    }
}


int main(int argc, const char * argv[]) {

    if (argc < 2){
        cerr << endl
             << "usage: vntr2kmers_thread [-th] [-g] [-p] [-m] -k -fs -ntr [-o|-on] -fa \n"
             << "  -th <INT>        Filter out kmers w/ count below this threshold. Default: 0, i.e. no filtering\n"
             << "  -g               output *graph.kmers for threading-based kmer query.\n"
             << "  -p <FILE>        Prune tr/graph kmers with the given kmers file.\n"
             << "  -m <FILE>        Use orthology map to merge haps.\n"
             << "  -k <INT>         Kmer size\n"
             << "  -fs <INT>        Length of flanking sequence in *.tr.fasta.\n"
             << "  -ntr <INT>       Length of desired NTR in *kmers.\n"
             << "  -o <STR>         Output file prefix" << endl
             << "  -on <STR>        Same as the -o option, but write locus and kmer name as well" << endl
             << "  -fa <n> <list>   Use specified *.fasta in the [list] instead of hapDB.\n"
             << "                   Count the first [n] files and build kmers for the rest\n\n";
        return 0;
    }
    vector<string> args(argv, argv+argc);
    bool genGraph = false, prune = false, usemap = false, writeKmerName = false;
    size_t argi = 1, threshold = 0, nhap = 0, NTRsize, fs, nfile2count, nloci;
    string pruneFname, outPref, mapf;
    vector<string> infnames;

    while (argi < argc) {
        if (args[argi] == "-th") { threshold = stoi(args[++argi]); }
        else if (args[argi] == "-g") { genGraph = true; }
        else if (args[argi] == "-p") {
            prune = true;
            pruneFname = args[++argi];
            ifstream tmp(pruneFname);
            assert(tmp);
            tmp.close();
        }
		else if (args[argi] == "-m") { 
			usemap = true;
			mapf = args[++argi];
		}
        else if (args[argi] == "-k") { ksize = stoi(args[++argi]); }
        else if (args[argi] == "-fs") { fs = stoi(args[++argi]); }
        else if (args[argi] == "-ntr") {
            NTRsize = stoi(args[++argi]);
            assert(fs >= NTRsize);
        }
        else if (args[argi] == "-o" or args[argi] == "-on") {
			writeKmerName = args[argi] == "-on";
            outPref = args[++argi];
            ofstream outf(outPref+".tr.kmers");
            assert(outf);
            outf.close();
        }
        else if (args[argi] == "-fa") {
            nfile2count = stoi(args[++argi]);
            while (++argi < argc) {
                ++nhap;
                infnames.push_back(args[argi]);
                ifstream inf(args[argi]);
                if (not inf) { cerr << args[argi] << endl; }
                assert(inf);
                inf.close();
            }
            nhap = infnames.size();
			if (not usemap) {
				cerr << "Not using orthology map, assuming all fasta files have the same number of loci\n"
				     << "Total number of loci in " << infnames[0] << ": ";
				nloci = countLoci(infnames[0]);
				cerr << nloci << endl;
			}
        }
        else {
            cerr << "invalid option" << endl;
            return 1;
        }
        ++argi;
    }
	vector<vector<bool>> omap;
	if (usemap) {
		readOrthoMap(mapf, omap, nhap);
		nloci = omap.size();
		cerr << "Using orthology map, total number of loci: " << nloci << endl;
	}
	

    // -----
    // open each file and create a kmer database for each loci
    // combine the kmer databases of the same loci across different files
    // -----
    vector<kmerCount_umap> TRkmersDB(nloci);
    vector<kmerCount_umap> NTRkmersDB(nloci);
    vector<GraphType> graphDB(nloci);
    for (size_t n = 0; n < nhap; ++n) {
        bool count = n < nfile2count;
        size_t locus = 0;
		
        string read, line;
        ifstream fin(infnames[n]);
        assert(fin.is_open());

        cerr << "building and counting " << infnames[n] << " kmers\n";
        while (getline(fin, line)) {
            if (line[0] != '>') {
                read += line;
            }
            if (fin.peek() == '>' or fin.peek() == EOF) {
                if (read != "") {
					if (usemap) { while (not omap[locus][n]) { ++locus; } }

                    size_t tr_l = fs;
                    size_t tr_r = fs;
                    size_t lntr_l = fs - NTRsize;
                    size_t lntr_r = read.size() - fs - (ksize-1); // seamless contenuation of kmers from ntr to tr
                    size_t rntr_l = read.size() - fs - (ksize-1); // seamless contenuation of kmers from tr to ntr
                    size_t rntr_r = fs - NTRsize;

                    buildNuKmers(TRkmersDB[locus], read, ksize, tr_l, tr_r, count); // (begin_pos, right_flank_size)
                    buildNuKmers(NTRkmersDB[locus], read, ksize, lntr_l, lntr_r, count);
                    buildNuKmers(NTRkmersDB[locus], read, ksize, rntr_l, rntr_r, count);
                    if (genGraph) {
                        buildKmerGraph(graphDB[locus], read, ksize); // no self loop
                    }
                }
                read = "";
                ++locus;
            }
        }
        fin.close();
    }

    if (prune) {
        cerr << "pruning unsupported kmers with " << pruneFname << endl;

        vector<kmerCount_umap> prunedkmersDB(nloci);
        readKmersFile2DB(prunedkmersDB, pruneFname);

        for (size_t locus = 0; locus < nloci; ++locus) {
            auto& TRkmers = TRkmersDB[locus];
            auto& prunedkmers = prunedkmersDB[locus];
            for (auto& p : prunedkmers) { TRkmers.erase(p.first); }

            if (genGraph) {
                auto& graph = graphDB[locus];
                for (auto& p : prunedkmers) {
                    removeNodeFromGraph(p.first, graph);
                    removeNodeFromGraph(getNuRC(p.first, ksize), graph);
                }
            }
        }
    }


    // -----
    // write kmers files for all kmer databases
    // -----
    cerr << "writing outputs" << endl;
    if (writeKmerName) {
        writeKmersWithName(outPref + ".tr", TRkmersDB, threshold);
        writeKmersWithName(outPref + ".ntr", NTRkmersDB, threshold);    
        if (genGraph)
            writeKmersWithName(outPref + ".graph", graphDB);
    }
    else {
        writeKmers(outPref + ".tr", TRkmersDB, threshold);
        writeKmers(outPref + ".ntr", NTRkmersDB, threshold);
        if (genGraph)
            writeKmers(outPref + ".graph", graphDB);
    }

    return 0;
}
```
