## Supplemental code for "The motif composition of variable-number tandem repeats impacts gene expression": emotif_single_tissue.gtex.py

```
#!/usr/bin/env python3

import sys
srcdir = "/project/mchaisso_100/cmb-16/tsungyul/work/vntr/danbing-tk/script"
sys.path.insert(0, srcdir)
import numpy as np
import pandas as pd
import vntrutils as vu
from collections import defaultdict
import pickle
import glob
import warnings
from datetime import datetime
import statsmodels.api as sm
from statsmodels.stats.multitest import fdrcorrection as fdr

KSIZE = 21

# Genotype matrix/index
def getGTmask():
    genMat = pickle.load(open(args.genMat, 'rb')).T.astype(float)
    badg = np.loadtxt(args.badg, dtype=object)
    gtSampleMask = ~np.isin(genomes, badg)
    return genMat, gtSampleMask

def loadGTindex():
    with open(args.gtidx, 'rb') as f:
        ccki_tr = pickle.load(f)[1]
    with open(args.gtmeta, 'rb') as f:
        _, ccks, cck_ns, _ = pickle.load(f)
    return ccki_tr, ccks, cck_ns

def load_filter():
    trfilter = np.all(pd.read_csv(args.filter, sep="\t", index_col=0).values[:,[3,4,5,6]].astype(bool).astype(int), axis=1)
    mtfilter = np.all(np.loadtxt(args.mape, dtype=object, skiprows=1)[:,3:].astype(int).astype(bool), axis=1)
    return trfilter, mtfilter


# Gene-TR mapping relation
def indexLociList():
    lociList = np.loadtxt(args.TRbed, dtype=object, usecols=[0,1,2])
    loci2ind = {}
    for ind, row in enumerate(lociList):
        loci2ind["_".join(row)] = ind
    return lociList, loci2ind

def indexGeneList(tissue):
    tisGeneList = np.loadtxt(f'{args.expDir}/{tissue}.v8.normalized_expression.bed.gz', dtype=object, skiprows=1, usecols=[3])
    tisGene2ind = {}
    for ind, gene in enumerate(tisGeneList):
        tisGene2ind[gene] = ind
    return tisGeneList, tisGene2ind


def indexPairs(tisGene2ind):
    tisGeneMask = np.array([gene in tisGene2ind for gene in TRxGene[:,3]], dtype=bool)

    pairs = np.empty([np.sum(tisGeneMask), 2], dtype=object)
    pairs[:,0] = ["_".join(locus) for locus in TRxGene[tisGeneMask,:3]]
    pairs[:,1] = TRxGene[tisGeneMask, 3]

    pairsi = np.array([[loci2ind[locus], tisGene2ind[gene]] for locus, gene in pairs])
    m0 = np.array([trfilter[tri] for tri in pairsi[:,0]])
    pairs = pairs[m0]
    pairsi = pairsi[m0]
    print(f'\t{len(set(pairsi[:,0].tolist()))} TRs')
    print(f'\t{len(set(pairsi[:,1].tolist()))} Genes')
    print(f'\t{pairsi.shape[0]} TR x Gene tests', flush=True)

    genei2nkmer = defaultdict(int)
    for locusi, genei in pairsi:
        si = ccki_tr[locusi-1] if locusi else 0
        ei = ccki_tr[locusi]
        genei2nkmer[genei] += np.sum(mtfilter[si:ei])
    print(f'\t{np.mean(list(genei2nkmer.values())):.1f} tests per gene', flush=True)

    return pairs, pairsi, genei2nkmer


# expression matrix
def loadSNPPCinfo(ndim=838):
    if args.SNPPC is None:
        return None, None

    tmp = np.loadtxt(args.SNPPC, usecols=np.arange(11), dtype=object)[:ndim] 
    SNP_PCs = tmp[:,1:].astype(float)
    SNP_sampleList = [s.split("-")[-1] for s in tmp[:,0]]
    return SNP_PCs, SNP_sampleList


def getTisSNPResTpmMat(tissue, SNP_PCs, SNP_sampleList):
    # SNP PCs
    tmp = np.loadtxt(f'{args.expDir}/{tissue}.v8.normalized_expression.bed.gz', dtype=object, max_rows=1, comments="!")[4:]
    tisSampleList = np.array([s[5:] for s in tmp])

    snpSample2ind = {}
    for sind, sample in enumerate(SNP_sampleList):
        snpSample2ind[sample] = sind

    sampleMap_tis2snp = np.zeros(tisSampleList.size, dtype=int)
    for ind in range(tisSampleList.size):
        sampleMap_tis2snp[ind] = snpSample2ind[tisSampleList[ind]]

    tisSNP_PCs = SNP_PCs[sampleMap_tis2snp]

    # GTEx PCs
    gtexPCs = np.loadtxt(f'{args.covDir}/{tissue}.v8.covariates.txt', dtype=object, skiprows=1)[:,1:].astype(float).T

    C = np.hstack((gtexPCs, tisSNP_PCs))
    tisTpmMat = np.loadtxt(f'{args.expDir}/{tissue}.v8.normalized_expression.bed.gz', dtype=object, skiprows=1)[:,4:].astype(float).T
    tisResTpmMat = (np.eye(C.shape[0]) - C @ np.linalg.inv(C.T @ C) @ C.T) @ tisTpmMat
    return tisResTpmMat.T


# genotype matrix
def getTissueGenMat(tissue):
    tmp = np.loadtxt(f'{args.expDir}/{tissue}.v8.normalized_expression.bed.gz', dtype=object, max_rows=1, comments="!")[4:]
    tisSampleList = np.array([s[5:] for s in tmp])
    mask = np.isin(genomes, tisSampleList)
    return genMat[:,mask&gtSampleMask], gtSampleMask[mask]


# eQTL mapping
def normalize_1d(arr):
    sd = np.std(arr)
    if np.isclose(sd, 0):
        return None
    else:
        return (arr - np.mean(arr)) / sd

def runRegressionZ8(tisResTpmMat, tisGenMat, pairsi, genei2nkmer, ccki_tr, allp, allb, OUT_THRESH=2):
    pairstats = np.full([pairsi.shape[0], 6], np.nan) # lead_TR, lead_p_nominal, lead_ki, p_nominal, slope, slope_se
    pairstats[:,0] = False
    leadP = {}
    mtfilter_outlier = np.copy(mtfilter)
    for pi, (locusi, genei) in enumerate(pairsi):
        if pi % 1000 == 0: print(".", end="", flush=True)

        ki_s = ccki_tr[locusi-1] if locusi != 0 else 0
        ki_e = ccki_tr[locusi]
        genos = np.copy(tisGenMat[ki_s:ki_e])
        m = np.any(np.isfinite(genos), axis=1)
        genos = genos[m]
        sdx = np.nanstd(genos, axis=1)
        mns = np.nanmean(genos, axis=1)
        m0 = ~np.isclose(sdx, 0)
        sdx = sdx[m0]
        mns = mns[m0]
        genos = genos[m0]
        m[m] = m0
        om = ((genos <= (mns + OUT_THRESH*sdx)[:,None]) & (genos >= (mns - OUT_THRESH*sdx)[:,None]))
        xs_ = (genos - mns[:,None]) / sdx[:,None]
        exps = tisResTpmMat[genei]
        ys_ = normalize_1d(exps)
        if not np.all(np.isfinite(ys_)): print("y", end="", flush=True); continue

        for i, ki in enumerate(np.arange(ki_s, ki_e)[m]):
            if not mtfilter[ki]:
                continue
            if not np.all(om[i]):
                xs = normalize_1d(xs_[i][om[i]])
                if xs is None:
                    continue
                ys = normalize_1d(ys_[om[i]])
            else:
                xs = xs_[i]
                ys = ys_
            if not np.any(xs): print("o", end="", flush=True); continue

            results = sm.OLS(ys, sm.add_constant(xs, prepend=True)).fit()
            p = results.pvalues[1]
            b = results.params[1]
            bse = results.bse[1]
            if p < pairstats[pi,3] or ~np.isfinite(pairstats[pi,3]):
                pairstats[pi,2:] = [ki, p, b, bse]
            if genei not in leadP:
                leadP[genei] = (p, pi)
            else:
                if p < leadP[genei][0]:
                    leadP[genei] = (p, pi)
            if genei not in allp: allp[genei] = {}
            if genei not in allb: allb[genei] = {}
            if locusi not in allp[genei]: allp[genei][locusi] = {}
            if locusi not in allb[genei]: allb[genei][locusi] = {}
            allp[genei][locusi][ki] = p
            allb[genei][locusi][ki] = (b, bse)

    nvar = np.zeros(pairsi.shape[0])
    for pi, (locusi, genei) in enumerate(pairsi):
        nvar[pi] = genei2nkmer[genei]
    for genei, (pmin, pi) in leadP.items():
        cisMask = pairsi[:,1] == pairsi[pi,1]
        pairstats[cisMask, 1] = pmin
        pairstats[pi,0] = True
    leadP_Bonf = (pairstats[:,3] * nvar)[pairstats[:,0].astype(bool)] # Bonferroni correction
    print("", flush=True)
    return pairstats, leadP_Bonf


def annotateGeneTR1(allGeneInfo, allTRinfo, gene2allind, genei2nkmer, pairs, pairsi, pairstats, rejected, qval):
    fulltbl = np.empty([pairs.shape[0], 19], dtype=object)

    fulltbl[:,:5] = np.array([allGeneInfo[gene2allind[gene]] for gene in pairs[:,1]], dtype=object) # gene_id, gene_name, chr, start, end
    fulltbl[:,5] = np.array([genei2nkmer[genei] for genei in pairsi[:,1]], dtype=object) # num_var
    fulltbl[:,6:9] = allTRinfo[pairsi[:,0]] # TR_chr, TR_start, TR_end
    fulltbl[:,9] = pairsi[:,0] # TR_locus
    fulltbl[:,10] = pairstats[:,2] # ki
    fulltbl[:,11] = False # eTR
    fulltbl[:,11][pairstats[:,0].astype(bool)] = rejected
    fulltbl[:,12] = pairstats[:,0].astype(bool) # lead_TR
    fulltbl[:,13:17] = pairstats[:,[1,3,4,5]] # lead_p_nominal, p_nominal, slope, slope_se
    fulltbl[:,17] = np.nan # qval
    fulltbl[:,17][pairstats[:,0].astype(bool)] = qval
    fulltbl[:,18] = [vu.decodeNumericString(ccks[int(ki)], KSIZE+cck_ns[int(ki)]-1) if np.isfinite(ki) else None for ki in pairstats[:,2]] # lead_kmer

    srti = np.argsort(fulltbl[:,9])
    return fulltbl[srti]


def singleTissue_eQTL(tissue, SNP_PCs, SNP_sampleList, allGeneInfo, allTRinfo, gene2allind, ccki_tr, fdr_cutoff=0.05):
    # establish mapping between TRs and Genes
    tisGeneList, tisGene2ind = indexGeneList(tissue)
    pairs, pairsi, genei2nkmer = indexPairs(tisGene2ind) # count # of TRs mapped to each gene; used for Bonferroni correction

    tisGenMat, tisGenMask = getTissueGenMat(tissue) # genMat with samples missing in tpmMat removed
    print(f'\ttisGenMat {tisGenMat.shape}', flush=True)

    if glob.glob(f'{args.resDir}/{tissue}.ResMat.pickle'):
        tisResTpmMat = pickle.load(open(f'{args.resDir}/{tissue}.ResMat.pickle', 'rb'))
    else:
        tisResTpmMat = getTisSNPResTpmMat(tissue, SNP_PCs, SNP_sampleList)
        pickle.dump(tisResTpmMat, open(f'{args.resDir}/{tissue}.ResMat.pickle', 'wb'))
    tisResTpmMat = tisResTpmMat[:,tisGenMask]
    print(f'\ttisResTpmMat {tisResTpmMat.shape}', flush=True)

    allp, allb = {}, {}
    pairstats, leadP_Bonf = runRegressionZ8(tisResTpmMat, tisGenMat, pairsi, genei2nkmer, ccki_tr, allp, allb) # [genei, locusi], [p, b, bse]
    rejected, qval = fdr(leadP_Bonf, alpha=fdr_cutoff)
    print(f'\t{np.sum(rejected)} tissue eGenes', flush=True)

    fulltbl = annotateGeneTR1(allGeneInfo, allTRinfo, gene2allind, genei2nkmer, pairs, pairsi, pairstats, rejected, qval)
    pickle.dump(allp, open(f'{args.outDir}/{tissue}.allp.pickle', 'wb'))
    pickle.dump(allb, open(f'{args.outDir}/{tissue}.allb.pickle', 'wb'))

    return fulltbl


def writeAlleGeneTR(TISSUE=None, fdr_cutoff=0.05):
    allGeneInfo = np.loadtxt(args.geneBed, dtype=object)[:,[3,4,0,1,2]]
    allGeneInfo[:,3:] = allGeneInfo[:,3:].astype(int)

    gene2allind = {}
    for i in range(allGeneInfo.shape[0]):
        gene2allind[allGeneInfo[i,0]] = i

    allTRinfo = np.loadtxt(args.TRbed, dtype=object, usecols=[0,1,2])
    allTRinfo[:,1:] = allTRinfo[:,1:].astype(int)

    SNP_PCs, SNP_sampleList = loadSNPPCinfo()
    tissues = np.loadtxt(args.tissues, dtype=object)

    for tissue in tissues:
        if TISSUE and TISSUE != tissue: continue
        print("tissue: {}".format(tissue), flush=True)

        fulltbl = singleTissue_eQTL(tissue, SNP_PCs, SNP_sampleList, allGeneInfo, allTRinfo, gene2allind, ccki_tr, fdr_cutoff)

        np.savetxt(f'{args.outDir}/{tissue}.v8.allGenes.txt', fulltbl, delimiter="\t",
                   header="gene_id\tgene_name\tchr\tstart\tend\tnum_var\tTR_chr\tTR_start\tTR_end\tTR_locus\tlead_ki\teTR\tlead_TR\tlead_p_nominal\tp_nominal\tslope\tslope_se\tqval\tlead_kmer",
                   fmt=['%s','%s','%s','%i','%i','%i','%s','%i','%i','%i','%s','%s','%s','%.4e','%.4e','%.4e','%.4e','%.4e','%s'])

        fulltbl = fulltbl[fulltbl[:,11].astype(bool)]
        np.savetxt(f'{args.outDir}/{tissue}.v8.egenes.txt', fulltbl, delimiter="\t",
                   header="gene_id\tgene_name\tchr\tstart\tend\tnum_var\tTR_chr\tTR_start\tTR_end\tTR_locus\tlead_ki\teTR\tlead_TR\tlead_p_nominal\tp_nominal\tslope\tslope_se\tqval\tlead_kmer",
                   fmt=['%s','%s','%s','%i','%i','%i','%s','%i','%i','%i','%s','%s','%s','%.4e','%.4e','%.4e','%.4e','%.4e','%s'])


class Args():
    def __init__(self):
        self.TRbed = "/project/mchaisso_100/cmb-17/vntr_genotyping/rpgg2_k21_84k/hgsvc/clinic/output/pan.tr.mbe.v2.bed"
        self.geneBed = "/project/mchaisso_100/cmb-17/vntr_genotyping/gtex/expression/genes_id_name.bed"
        self.pair = "/project/mchaisso_100/cmb-17/vntr_genotyping/rpgg2_k21_84k/hgsvc/gtex/emotif/input/gene.100k_window.tr.pair.bed"
        self.expDir = "/project/mchaisso_100/cmb-17/vntr_genotyping/gtex/expression/GTEx_Analysis_v8_eQTL_expression_matrices/"
        self.resDir = "/project/mchaisso_100/cmb-17/vntr_genotyping/gtex/eqtl/ResMat/"
        self.covDir = "/project/mchaisso_100/cmb-17/vntr_genotyping/gtex/expression/GTEx_Analysis_v8_eQTL_covariates/"
        self.outDir = "/project/mchaisso_100/cmb-17/vntr_genotyping/rpgg2_k21_84k/hgsvc/gtex/emotif/log10/test/"
        self.phenotype = "/project/mchaisso_100/cmb-17/vntr_genotyping/gtex/SubjectPhenotypesDS.txt"
        self.genomes = "/project/mchaisso_100/cmb-16/tsungyul/work/vntr/hapdb/config/genomes.gtex.txt"
        self.tissues = "/project/mchaisso_100/cmb-17/vntr_genotyping/gtex/expression/alltissue.txt"
        self.genMat = "/project/mchaisso_100/cmb-17/vntr_genotyping/rpgg2_k21_84k/hgsvc/gtex/gt_pickle/acgt_bc.v1.pickle"
        self.SNPPC = "/project/mchaisso_100/cmb-17/vntr_genotyping/gtex/variation/joint.pca.evec"
        self.badg = "/project/mchaisso_100/cmb-17/vntr_genotyping/rpgg2_k21_84k/hgsvc/gtex/input/bad.union_of_all_QC.txt"
        #self.master = "/project/mchaisso_100/cmb-17/vntr_genotyping/rpgg2_k21_84k/hgsvc/gtex/emotif/output5/Alltissue.egenes.tsv"
        self.gtidx = "/project/mchaisso_100/cmb-17/vntr_genotyping/rpgg2_k21_84k/hgsvc/clinic/cdbg/v1/ki_tr.ccki_tr.pickle"
        self.gtmeta = "/project/mchaisso_100/cmb-17/vntr_genotyping/rpgg2_k21_84k/hgsvc/clinic/cdbg/v1/ks.ccks.cck_ns.ki_map.pickle"
        self.filter = "/project/mchaisso_100/cmb-17/vntr_genotyping/rpgg2_k21_84k/hgsvc/clinic/analysis/80518_loci.alu.mismap_1.r2.lenvar.filter.tsv"
        self.mape = "/project/mchaisso_100/cmb-17/vntr_genotyping/rpgg2_k21_84k/hgsvc/clinic/analysis/mape.invariant.flank.alu.mismap_1.r2.v2.filter.tsv.gz"
args = Args()


print("loading metadata", flush=True)
genomes = np.loadtxt(args.genomes, dtype=object)
nwgs = genomes.size
nloci = np.loadtxt(args.TRbed, usecols=[1]).size
lociList, loci2ind = indexLociList()
TRxGene = np.loadtxt(args.pair, dtype=object, usecols=[5,6,7,3])
ccki_tr, ccks, cck_ns = loadGTindex()

print("reading genotype pickle", flush=True)
genMat, gtSampleMask = getGTmask()
print("reading genotype mape filter", flush=True)
trfilter, mtfilter = load_filter()

writeAlleGeneTR(fdr_cutoff=0.05, TISSUE=sys.argv[1])
```
