## Supplemental code for "The motif composition of variable-number tandem repeats impacts gene expression": fn1.pval.threshold.sh

```
#!/usr/bin/env bash
#SBATCH --ntasks=2
#SBATCH --time=168:00:00
#SBATCH --mem=16000
#SBATCH --partition=qcb
#SBATCH --account=mchaisso_100
#SBATCH -N 1
#SBATCH --job-name=pth
#SBATCH --output=slurm.%A_%a.%x.log
###SBATCH --constraint=xeon-2665,avx
###SBATCH --exclude=b10-10
###SBATCH --mail-type=ALL
###SBATCH --mail-user=
###SBATCH --array=0,1

#set -eu
set -e
source ~/.bashrc
conda activate nb1
#module load gcc usc

sd=/project/mchaisso_100/cmb-17/vntr_genotyping/rpgg2_k21_84k/hgsvc/gtex/emotif/log10
od=/project/mchaisso_100/cmb-17/vntr_genotyping/rpgg2_k21_84k/hgsvc/gtex/emotif/output10
ad=/project/mchaisso_100/cmb-17/vntr_genotyping/rpgg2_k21_84k/hgsvc/gtex/emotif/analysis10

date
$sd/fn1_.pval.threshold.py  $od  $ad
date
```
