## Supplemental code for "The motif composition of variable-number tandem repeats impacts gene expression": evntr_single_tissue.gtex.py

```
#!/usr/bin/env python3

import sys
import numpy as np
import pandas as pd
import pickle
import statsmodels.api as sm
from statsmodels.stats.multitest import fdrcorrection as fdr


# Genotype matrix/index
def getGTmask():
    with open(args.genMat, 'rb') as f:
        genMat = pickle.load(f)
    badg = np.loadtxt(args.badg, dtype=object)
    gtSampleMask = ~np.isin(genomes, badg)
    return genMat, gtSampleMask #gmMask

def load_filter():
    return np.all(pd.read_csv(args.filter, sep="\t", index_col=0).values[:,3:].astype(bool).astype(int), axis=1)


# Gene-TR mapping relation
def indexLociList():
    lociList = np.loadtxt(args.TRbed, dtype=object, usecols=[0,1,2])
    loci2ind = {}
    for ind, row in enumerate(lociList):
        loci2ind["_".join(row)] = ind
    return lociList, loci2ind

def indexGeneList(tissue):
    tisGeneList = np.loadtxt(f'{args.expDir}/{tissue}.v8.normalized_expression.bed.gz', dtype=object, skiprows=1, usecols=[3])
    tisGene2ind = {}
    for ind, gene in enumerate(tisGeneList):
        tisGene2ind[gene] = ind
    return tisGeneList, tisGene2ind


def indexPairs(tisGene2ind):
    tisGeneMask = np.array([gene in tisGene2ind for gene in TRxGene[:,3]], dtype=bool)

    pairs = np.empty([np.sum(tisGeneMask), 2], dtype=object)
    pairs[:,0] = ["_".join(locus) for locus in TRxGene[tisGeneMask,:3]]
    pairs[:,1] = TRxGene[tisGeneMask, 3]

    pairsi = np.array([[loci2ind[locus], tisGene2ind[gene]] for locus, gene in pairs])
    m0 = np.array([pass_filter[tri] for tri in pairsi[:,0]])
    pairs = pairs[m0]
    pairsi = pairsi[m0]
    print(f'\t{len(set(pairsi[:,0].tolist()))} TRs')
    print(f'\t{len(set(pairsi[:,1].tolist()))} Genes')
    print(f'\t{pairsi.shape[0]} TR x Gene tests', flush=True)

    genei2nloci = {}
    for locusi, genei in pairsi:
        if genei not in genei2nloci:
            genei2nloci[genei] = 1
        else:
            genei2nloci[genei] += 1
    print(f'\t{np.mean(list(genei2nloci.values())):.1f} tests per gene', flush=True)

    return pairs, pairsi, genei2nloci


# genotype matrix
def getTissueGenMat(tissue):
    tmp = np.loadtxt(f'{args.expDir}/{tissue}.v8.normalized_expression.bed.gz', dtype=object, max_rows=1, comments="!")[4:]
    tisSampleList = np.array([s[5:] for s in tmp])
    mask = np.isin(genomes, tisSampleList)
    return genMat[:,mask & gtSampleMask], gtSampleMask[mask]


# eQTL mapping
def runRegressionZ5(tisResTpmMat, tisGenMat, pairsi, genei2nloci, OUT_THRESH=3):
    """reject outliers in genotypes + qunatile normalize genotypes"""
    pairstats = np.full([pairsi.shape[0], 5], np.nan) # lead_TR, lead_p_nominal, p_nominal, slope, slope_se
    pairstats[:,0] = False
    leadP = {}

    for pi, (locusi, genei) in enumerate(pairsi):
        if pi % 1000 == 999: print(".", end="", flush=True)
        if not pass_filter[locusi]:
            genei2nloci[genei] -= 1
            continue
        genos = np.copy(tisGenMat[locusi])
        m0 = np.isfinite(genos)
        if not np.any(m0): continue
        exps = np.copy(tisResTpmMat[genei])
        xs = genos[m0]
        ys = exps[m0]
        sdx = np.std(xs)
        if np.isclose(sdx, 0): continue
        xs = (xs - np.mean(xs)) / sdx
        ys = (ys - np.mean(ys)) / np.std(ys)
        m = (xs < -OUT_THRESH) | (xs > OUT_THRESH)
        if np.any(m):
            xs, ys = xs[~m], ys[~m]
            sdx = np.std(xs)
            if np.isclose(sdx, 0): continue
            xs = (xs - np.mean(xs)) / sdx
            ys = (ys - np.mean(ys)) / np.std(ys)

        #if not np.any(genos): print("-", end=""); continue
        if not np.all(np.isfinite(xs)): print("-", end=""); continue
        if not np.all(np.isfinite(ys)): print("-", end=""); continue

        results = sm.OLS(ys, sm.add_constant(xs, prepend=True)).fit()
        p = results.pvalues[1]
        b = results.params[1]
        bse = results.bse[1]
        pairstats[pi,2:] = [p, b, bse]
        if genei not in leadP:
            leadP[genei] = (p, pi)
        else:
            if p < leadP[genei][0]:
                leadP[genei] = (p, pi)

    nvar = np.zeros(pairsi.shape[0])
    for pi, (locusi, genei) in enumerate(pairsi):
        nvar[pi] = genei2nloci[genei]
    for genei, (pmin, pi) in leadP.items():
        cisMask = pairsi[:,1] == pairsi[pi,1]
        pairstats[cisMask, 1] = pmin
        pairstats[pi,0] = True

    leadP_Bonf = (pairstats[:,2] * nvar)[pairstats[:,0].astype(bool)] # Bonferroni correction

    print("")
    return pairstats, leadP_Bonf


def annotateGeneTR1(allGeneInfo, allTRinfo, gene2allind, genei2nloci, pairs, pairsi, pairstats, rejected, qval):
    fulltbl = np.empty([pairs.shape[0], 17], dtype=object)

    fulltbl[:,:5] = np.array([allGeneInfo[gene2allind[gene]] for gene in pairs[:,1]], dtype=object) # gene_id, gene_name, chr, start, end
    fulltbl[:,5] = np.array([genei2nloci[genei] for genei in pairsi[:,1]], dtype=object) # num_var
    fulltbl[:,6:9] = allTRinfo[pairsi[:,0]] # TR_chr, TR_start, TR_end
    fulltbl[:,9] = pairsi[:,0] # TR_locus
    fulltbl[:,10] = False # eTR
    fulltbl[:,10][pairstats[:,0].astype(bool)] = rejected
    fulltbl[:,11] = pairstats[:,0].astype(bool) # lead_TR
    fulltbl[:,12:16] = pairstats[:,1:] # lead_p_nominal, p_nominal, slope, slope_se
    fulltbl[:,16] = np.nan # qval
    fulltbl[:,16][pairstats[:,0].astype(bool)] = qval

    srti = np.argsort(fulltbl[:,9])
    return fulltbl[srti]


def singleTissue_eQTL(tissue, allGeneInfo, allTRinfo, gene2allind, fdr_cutoff=0.05):
    # establish mapping between TRs and Genes
    tisGeneList, tisGene2ind = indexGeneList(tissue)
    pairs, pairsi, genei2nloci = indexPairs(tisGene2ind) # count # of TRs mapped to each gene; used for Bonferroni correction

    tisGenMat, tisGenMask = getTissueGenMat(tissue) # genMat with samples missing in tpmMat removed
    print(f'\ttisGenMat {tisGenMat.shape}', flush=True)

    tisResTpmMat = pickle.load(open(f'{args.resDir}/{tissue}.ResMat.pickle', 'rb'))[:,tisGenMask]
    print(f'\ttisResTpmMat {tisResTpmMat.shape}', flush=True)

    pairstats, leadP_Bonf = runRegressionZ5(tisResTpmMat, tisGenMat, pairsi, genei2nloci) # [genei, locusi], [p, b, bse]
    rejected, qval = fdr(leadP_Bonf, alpha=fdr_cutoff)
    print(f'\t{np.sum(rejected)} tissue eGenes', flush=True)

    fulltbl = annotateGeneTR1(allGeneInfo, allTRinfo, gene2allind, genei2nloci, pairs, pairsi, pairstats, rejected, qval)

    return fulltbl


def writeAlleGeneTR(TISSUE, fdr_cutoff=0.05):
    allGeneInfo = np.loadtxt(args.geneBed, dtype=object)[:,[3,4,0,1,2]]
    allGeneInfo[:,3:] = allGeneInfo[:,3:].astype(int)

    gene2allind = {}
    for i in range(allGeneInfo.shape[0]):
        gene2allind[allGeneInfo[i,0]] = i

    allTRinfo = np.loadtxt(args.TRbed, dtype=object, usecols=[0,1,2])
    allTRinfo[:,1:] = allTRinfo[:,1:].astype(int)

    tissues = np.loadtxt(args.tissues, dtype=object)

    for tissue in tissues:
        if tissue != TISSUE: continue
        print("tissue: {}".format(tissue), flush=True)

        fulltbl = singleTissue_eQTL(tissue, allGeneInfo, allTRinfo, gene2allind, fdr_cutoff)

        np.savetxt(f'{args.outDir}/{tissue}.v8.allGenes.txt', fulltbl, delimiter="\t",
                   header="gene_id\tgene_name\tchr\tstart\tend\tnum_var\tTR_chr\tTR_start\tTR_end\tTR_locus\teTR\tlead_TR\tlead_p_nominal\tp_nominal\tslope\tslope_se\tqval",
                   fmt=['%s','%s','%s','%i','%i','%i','%s','%i','%i','%i','%s','%s','%.4e','%.4e','%.4e','%.4e','%.4e'])

        fulltbl = fulltbl[fulltbl[:,10].astype(bool)]
        np.savetxt(f'{args.outDir}/{tissue}.v8.egenes.txt', fulltbl, delimiter="\t",
                   header="gene_id\tgene_name\tchr\tstart\tend\tnum_var\tTR_chr\tTR_start\tTR_end\tTR_locus\teTR\tlead_TR\tlead_p_nominal\tp_nominal\tslope\tslope_se\tqval",
                   fmt=['%s','%s','%s','%i','%i','%i','%s','%i','%i','%i','%s','%s','%.4e','%.4e','%.4e','%.4e','%.4e'])


class Args():
    def __init__(self):
        self.TRbed = "/project/mchaisso_100/cmb-17/vntr_genotyping/rpgg2_k21_84k/hgsvc/clinic/output/pan.tr.mbe.v2.bed"
        self.geneBed = "/project/mchaisso_100/cmb-17/vntr_genotyping/gtex/expression/genes_id_name.bed"
        self.pair = "/project/mchaisso_100/cmb-17/vntr_genotyping/rpgg2_k21_84k/hgsvc/gtex/emotif/input/gene.100k_window.tr.pair.bed"
        self.expDir = "/project/mchaisso_100/cmb-17/vntr_genotyping/gtex/expression/GTEx_Analysis_v8_eQTL_expression_matrices/"
        self.resDir = "/project/mchaisso_100/cmb-17/vntr_genotyping/gtex/eqtl/ResMat/"
        self.covDir = "/project/mchaisso_100/cmb-17/vntr_genotyping/gtex/expression/GTEx_Analysis_v8_eQTL_covariates/"
        self.outDir = "/project/mchaisso_100/cmb-17/vntr_genotyping/rpgg2_k21_84k/hgsvc/gtex/evntr/log7/test"
        self.phenotype = "/project/mchaisso_100/cmb-17/vntr_genotyping/gtex/SubjectPhenotypesDS.txt"
        self.genomes = "/project/mchaisso_100/cmb-16/tsungyul/work/vntr/hapdb/config/genomes.gtex.txt"
        self.tissues = "/project/mchaisso_100/cmb-17/vntr_genotyping/gtex/expression/alltissue.txt"
        self.genMat = "/project/mchaisso_100/cmb-17/vntr_genotyping/rpgg2_k21_84k/hgsvc/gtex/genotype/gtex.80518x879.akms_bc.v1.pickle"
        self.badg = "/project/mchaisso_100/cmb-17/vntr_genotyping/rpgg2_k21_84k/hgsvc/gtex/input/bad.union_of_all_QC.txt"
        self.filter = "/project/mchaisso_100/cmb-17/vntr_genotyping/rpgg2_k21_84k/hgsvc/clinic/analysis/80518_loci.alu.mismap_1.r2.lenvar.filter.tsv"
args = Args()

genomes = np.loadtxt(args.genomes, dtype=object)
nwgs = genomes.size
nloci = np.loadtxt(args.TRbed, usecols=[1]).size
lociList, loci2ind = indexLociList()
TRxGene = np.loadtxt(args.pair, dtype=object, usecols=[5,6,7,3])

print("Loading genotype pickle", flush=True)
genMat, gtSampleMask = getGTmask()
print("Reading VNTR filter", flush=True)
pass_filter = load_filter()

writeAlleGeneTR(sys.argv[1], fdr_cutoff=0.05)
```
