## Supplemental code for "The motif composition of variable-number tandem repeats impacts gene expression": fn0.evntr.sh

```
#!/usr/bin/env bash
#SBATCH --ntasks=4
#SBATCH --time=168:00:00
#SBATCH --mem=10000
#SBATCH --partition=qcb
#SBATCH --account=mchaisso_100
#SBATCH -N 1
#SBATCH --job-name=etr
#SBATCH --output=slurm.%A_%a.%x.log
###SBATCH --constraint=xeon-2665,avx
###SBATCH --exclude=b10-10
###SBATCH --mail-type=ALL
###SBATCH --mail-user=
###SBATCH --array=0,1

#set -eu
set -e
source ~/.bashrc
conda activate nb1
#module load gcc usc

ts=( $(cat /project/mchaisso_100/cmb-17/vntr_genotyping/gtex/expression/alltissue.txt) )
t=${ts[$SLURM_ARRAY_TASK_ID]}
sd=/project/mchaisso_100/cmb-17/vntr_genotyping/rpgg2_k21_84k/hgsvc/gtex/evntr/log7/test


date
$sd/evntr_single_tissue.gtex.py $t
date
```
