## Supplemental code for "The motif composition of variable-number tandem repeats impacts gene expression": fn1_.pval.threshold.py

```
#!/usr/bin/env python3

import sys
import numpy as np
import pickle
from statsmodels.stats.multitest import fdrcorrection as fdr

def get_pth():
    t2pth = {}
    for t in np.loadtxt("/project/mchaisso_100/cmb-17/vntr_genotyping/gtex/expression/alltissue.txt", dtype=object):
        ps = np.loadtxt(f"{OUTDIR}/{t}.v8.allGenes.txt", dtype=float, usecols=[13])
        rej, qval = fdr(ps)
        t2pth[t] = max(np.array(ps)[rej]) if np.any(rej) else np.nan
    return t2pth

def write_pickle():
    with open(f"{ANADIR}/nominal_p_genome_wide_cutoff.txt", 'w') as f:
        for t, pth in t2pth.items():
            f.write(f"{t}\t{pth:.5e}\n")

OUTDIR, ANADIR = sys.argv[1], sys.argv[2]
t2pth = get_pth()
write_pickle()
```
