## Supplemental code for "The motif composition of variable-number tandem repeats impacts gene expression": fn0-5.sh

```
#!/bin/bash
#SBATCH --ntasks=12
#SBATCH --time=120:00:00
#SBATCH --mem=36000
#SBATCH --partition=qcb
#SBATCH --account=mchaisso_100
#SBATCH -N 1
#SBATCH --job-name=fmap
#SBATCH --output=fn2.%x.%A_%a.log

source ~/.bashrc
module load gcc
set -eu
conda activate nb1

# sort based on data size
tissues=( $(ls -l ../input4/*.snp_emotif_tr*gz | awk 'BEGIN {OFS="\t"} {split($9,vs0,"."); split(vs0[3],vs1,"/"); print $5, vs1[3]}' | sort -k1,1nr | cut -f 2) )
#tissues=( $(cat /project/mchaisso_100/cmb-17/vntr_genotyping/gtex/expression/alltissue.txt) )
tis=${tissues[$SLURM_ARRAY_TASK_ID]}
motif_dir=/project/mchaisso_100/cmb-17/vntr_genotyping/rpgg2_k21_84k/hgsvc/gtex/emotif/output10
indir=/project/mchaisso_100/cmb-17/vntr_genotyping/rpgg2_k21_84k/hgsvc/gtex/susieR/input10
outdir=/project/mchaisso_100/cmb-17/vntr_genotyping/rpgg2_k21_84k/hgsvc/gtex/susieR/output10


date
./fn0-5_.0.py $tis  $motif_dir  $indir

date
vcf=/project/mchaisso_100/cmb-17/GTEx/variation/GTEx_Analysis_2017-06-05_v8_WholeGenomeSeq_838Indiv_Analysis_Freeze.vcf.gz
rgn=$indir/$tis.egenes.tss_cis_1Mb.bed
SNPgt=$indir/$tis.snp.gt.tsv.gz

date
module load bzip2 bcftools
bcftools view --threads 3 -Oz -R $rgn $vcf |
bcftools query -f '%CHROM %POS %REF %ALT{0}  [ %GT]\n' /dev/stdin |
awk 'BEGIN {OFS="\t"} {
        printf $1"_"$2"_"$3"_"$4"_b38\t"
        if ($5 == "0/0") {printf "0"}
        else if ($5 == "0/1") {printf "1"}
        else if ($5 == "1/1") {printf "2"}
        else if ($5 == "./.") {printf "3"}
        else {exit 1}
        for (i=6; i<=NF; ++i) {
                if ($i == "0/0") {printf "\t0"}
                else if ($i == "0/1") {printf "\t1"}
                else if ($i == "1/1") {printf "\t2"}
                else if ($i == "./.") {printf "\t3"}
                else {exit 1}
        }
        printf "\n"
}' | gzip >$SNPgt
module unload bzip2 bcftools

date
AKMS_BC_FN=/project/mchaisso_100/cmb-17/vntr_genotyping/rpgg2_k21_84k/hgsvc/gtex/genotype/gtex.80518x879.akms_bc.v1.pickle
ACGT_BC_FN=/project/mchaisso_100/cmb-17/vntr_genotyping/rpgg2_k21_84k/hgsvc/gtex/gt_pickle/acgt_bc.v1.pickle
FILTER_FN=/project/mchaisso_100/cmb-17/vntr_genotyping/rpgg2_k21_84k/hgsvc/clinic/analysis/80518_loci.alu.mismap_1.r2.v2.filter.tsv
./fn0-5_.2.py $tis  $motif_dir  $indir  $AKMS_BC_FN  $ACGT_BC_FN  $FILTER_FN

date
L=10
module load openblas/0.3.8 r/4.0.3 gsl
Rscript fn0-5_.3.R $tis  $L  $indir  $outdir
module unload openblas/0.3.8 r/4.0.3 gsl

date
mkdir -p $outdir/$tis
./fn0-5_.4.py $tis  $indir  $outdir  $L

date
anadir=/project/mchaisso_100/cmb-17/vntr_genotyping/rpgg2_k21_84k/hgsvc/gtex/susieR/analysis10
mkdir -p $indir/pickle
zcat $indir/$tis.snp_emotif_tr.gt.bed.gz |
awk 'BEGIN {OFS="\t"} {if (substr($4,1,1)=="t") {n=split($4,vs,"_"); if (n==6) {print vs[6],$5} else {print -vs[2], $5}} }' |
./fn0-5_.5.tmi2vari.py $tis  $indir
motif_anadir=/project/mchaisso_100/cmb-17/vntr_genotyping/rpgg2_k21_84k/hgsvc/gtex/emotif/analysis10/
vntr_dir=/project/mchaisso_100/cmb-17/vntr_genotyping/rpgg2_k21_84k/hgsvc/gtex/evntr/output6
vntr_anadir=/project/mchaisso_100/cmb-17/vntr_genotyping/rpgg2_k21_84k/hgsvc/gtex/evntr/analysis6/
./fn0-5_.5.py $tis  $motif_dir  $motif_anadir  $vntr_dir  $vntr_anadir  $indir  $outdir  $L

date
```
