## Supplemental code for "The motif composition of variable-number tandem repeats impacts gene expression": fn0-5_.0.py

```
#!/usr/bin/env python3

import sys
import numpy as np
import pandas as pd
from pybedtools import BedTool

def gen_tissue_egene_cis_bed():
    fai = "/project/mchaisso_100/cmb-16/tsungyul/work/vntr/datasets/reference/hg38_noalts/hg38.no_alts.fasta.fai"
    fn = f'/project/mchaisso_100/cmb-17/vntr_genotyping/gtex/expression/GTEx_Analysis_v8_eQTL_expression_matrices/{TIS}.v8.normalized_expression.bed.gz'
    tbed = np.loadtxt(fn, dtype=object, usecols=[0,1,2,3])
    tbed[:,1:3] = tbed[:,1:3].astype(int)
    ge2i = {}
    for i, ge in enumerate(tbed[:,3]):
        ge2i[ge] = i
    ges = set(np.loadtxt(f"{EMOTIF_DIR}/{TIS}.v8.egenes.txt", usecols=0, dtype=object).tolist())
    idxs = []
    for i, ge in enumerate(ges):
        if ge not in ge2i: continue
        idxs.append(ge2i[ge])
    ebed = tbed[idxs]
    i2 = np.argsort(ebed[:, 2].astype(int), kind='stable')
    i1 = np.argsort(ebed[i2, 1].astype(int), kind='stable')
    i0 = np.argsort(ebed[i2[i1], 0], kind='stable')
    srti = (i2[i1])[i0]
    ebed = ebed[srti]
    df = pd.DataFrame(ebed, columns=["chr","start","end","geneid"])
    df["gene_idx"] = np.arange(ebed.shape[0])
    BedTool.from_dataframe(df).sort().slop(g=fai, b=1000000).saveas(f"{FM_INDIR}/{TIS}.egenes.tss_cis_1Mb.bed")

TIS = sys.argv[1]
EMOTIF_DIR = sys.argv[2]
FM_INDIR = sys.argv[3]
gen_tissue_egene_cis_bed()
```
