## Supplemental code for "The motif composition of variable-number tandem repeats impacts gene expression": fn0-5_.2.filter_sample.py

```
#!/usr/bin/env python3

import sys
import numpy as np
import gzip

t = sys.argv[1]

badsamples = np.loadtxt("/project/mchaisso_100/cmb-17/vntr_genotyping/rpgg2_k21_84k/hgsvc/gtex/input/bad.union_of_all_QC.txt", dtype=object)
tmp = np.loadtxt(f'/project/mchaisso_100/cmb-17/vntr_genotyping/gtex/expression/GTEx_Analysis_v8_eQTL_expression_matrices/{t}.v8.normalized_expression.bed.gz', dtype=object, max_rows=1, comments="!")[4:]
tsamples = np.array([s[5:] for s in tmp])
vcf = "/project/mchaisso_100/cmb-17/GTEx/variation/GTEx_Analysis_2017-06-05_v8_WholeGenomeSeq_838Indiv_Analysis_Freeze.vcf.gz"
with gzip.open(vcf, 'rt') as f:
    for line in f:
        if line[:6] == "#CHROM":
            snpsamples = np.array([v[5:] for v in line.split()[9:]])
            break
btmask = np.isin(snpsamples, tsamples) & (~np.isin(snpsamples, badsamples))
mask = np.concatenate((np.ones(5, dtype=bool),btmask))
print(",".join((np.nonzero(mask)[0]+1).astype(str)))
```
