## Supplemental code for "The motif composition of variable-number tandem repeats impacts gene expression": fn0-5_.2.py

```
#!/usr/bin/env python3

import sys
import numpy as np
import pandas as pd
import pickle
import glob
import gzip
from pybedtools import BedTool
import pybedtools
import subprocess

def gen_tr_len_mt_bed(TIS):
    # meta
    vcf = "/project/mchaisso_100/cmb-17/GTEx/variation/GTEx_Analysis_2017-06-05_v8_WholeGenomeSeq_838Indiv_Analysis_Freeze.vcf.gz"
    with gzip.open(vcf, 'rt') as f:
        for line in f:
            if line[:6] == "#CHROM":
                snpsamples = np.array([v[5:] for v in line.split()[9:]])
                break
    nsample = snpsamples.size
    trsamples = np.loadtxt("/project/mchaisso_100/cmb-16/tsungyul/work/vntr/hapdb/config/genomes.gtex.txt", dtype=object)
    spm = np.isin(trsamples, snpsamples)

    # VNTR
    tbl = np.loadtxt(f"{EMOTIF_DIR}/{TIS}.v8.allGenes.txt", dtype=object)
    m = tbl[:,10] != "nan" # cleaning nan
    tbl = tbl[m]
    set_ki = set() # cleaning duplicate ki
    m = np.ones(tbl.shape[0], dtype=bool)
    for i, ki in enumerate(tbl[:,10].astype(float).astype(int)):
        if ki not in set_ki:
            set_ki.add(ki)
        else:
            m[i] = False
    tbl = tbl[m]
    kis = tbl[:,10].astype(float).astype(int)
    nki = kis.size
    trbed = np.loadtxt("/project/mchaisso_100/cmb-17/vntr_genotyping/rpgg2_k21_84k/hgsvc/clinic/output/pan.tr.mbe.v2.bed", usecols=[0,1,2], dtype=object)
    tr_filter = np.all(pd.read_csv(FILTER_FN, sep="\t", index_col=0).values[:,3:].astype(bool), axis=1)
    trbed = trbed[tr_filter]
    ntr = np.sum(tr_filter)
    tris = np.nonzero(tr_filter)[0]
    tri_kis = np.empty([nki+ntr, 2], dtype=int) # [tri, ki]
    gtbed = np.empty([nki+ntr, nsample+5], dtype=object) # [chr, start, end, variant_id, variant_index, gt[, ...]]
    # TR dosage
    gtbed[:ntr,0] = trbed[:,0]
    gtbed[:ntr,1] = trbed[:,1].astype(int)
    gtbed[:ntr,2] = gtbed[:ntr,1] + 1
    gtbed[:ntr,3] = [f"tr_{tris[i]}_{ch}_{s}_{e}" for i, (ch, s, e) in enumerate(trbed)]
    tri_kis[:ntr,0] = tris
    tri_kis[:ntr,1] = -1
    with open(AKMS_BC_FN, 'rb') as f:
        gtbed[:ntr,5:] = pickle.load(f)[:,spm][tr_filter]
    # motif dosage
    gtbed[ntr:,0] = tbl[:,6]
    gtbed[ntr:,1] = tbl[:,7].astype(int)
    gtbed[ntr:,2] = gtbed[ntr:,1] + 1
    gtbed[ntr:,3] = [f"tr_{int(tri)}_{ch}_{s}_{int(e)}_{ki}" for ch, s, e, tri, ki in zip(gtbed[ntr:,0], gtbed[ntr:,1], tbl[:,8], tbl[:,9], kis)]
    tri_kis[ntr:,0] = tbl[:,9].astype(int)
    tri_kis[ntr:,1] = kis
    with open(ACGT_BC_FN, 'rb') as f:
        gtbed[ntr:,5:] = pickle.load(f)[:,kis][spm].T
    # sort
    si1 = np.argsort(tri_kis[:,1], kind='stable')
    tri_kis_ = tri_kis[si1]
    si0 = np.argsort(tri_kis_[:,0], kind='stable')
    srti = si1[si0]
    gtbed = gtbed[srti]

    np.savetxt(f"{INDIR}/{TIS}.emotif_tr.bed.gz", gtbed, fmt='%s', delimiter="\t")

def gen_mapping_bed(TIS):
    gtfn = glob.glob(f"{INDIR}/{TIS}.snp_emotif_tr.gt.bed.gz")[0]
    genefn = f"{INDIR}/{TIS}.egenes.tss_cis_1Mb.bed"
    mapfn = f"{INDIR}/{TIS}.egenes.tss_cis_1Mb.map.snp_emotif_tr.bed"
    genebed = BedTool(genefn)
    genebed.map(c="5,5", o="min,max", b=gtfn).saveas(mapfn)

def gen_residualized_expression_mat(TIS):
    samples = np.loadtxt("/project/mchaisso_100/cmb-16/tsungyul/work/vntr/hapdb/config/genomes.gtex.txt", dtype=object)
    badsamples = np.loadtxt("/project/mchaisso_100/cmb-17/vntr_genotyping/rpgg2_k21_84k/hgsvc/gtex/input/bad.union_of_all_QC.txt", dtype=object)
    bmask = ~np.isin(samples, badsamples)
    expfn = f'/project/mchaisso_100/cmb-17/vntr_genotyping/gtex/expression/GTEx_Analysis_v8_eQTL_expression_matrices/{TIS}.v8.normalized_expression.bed.gz'
    tmp = np.loadtxt(expfn, dtype=object, max_rows=1, comments="!")[4:]
    tsamples = np.array([s[5:] for s in tmp])
    tmask = np.isin(samples, tsamples)
    btmask = bmask[tmask]

    mapfn = f"{INDIR}/{TIS}.egenes.tss_cis_1Mb.map.snp_emotif_tr.bed"
    egenes = np.loadtxt(mapfn, usecols=3, dtype=object)
    tgenebed = np.loadtxt(expfn, usecols=[0,1,2,3], dtype=object)
    tgenes = tgenebed[:,3]
    egmask = np.isin(tgenes, egenes)
    with open(f"/project/mchaisso_100/cmb-17/vntr_genotyping/gtex/eqtl/ResMat/{TIS}.ResMat.pickle", 'rb') as f:
        resmat = pickle.load(f)[:,btmask][egmask]
    print(resmat.shape, flush=True)
    nrow, ncol = resmat.shape
    resfn = f"{INDIR}/{TIS}.resmat.{nrow}egene_X_{ncol}sample.bed.gz"
    resbed = np.hstack((tgenebed[egmask], resmat))
    i2 = np.argsort(resbed[:,2].astype(int), kind='stable')
    i1 = np.argsort(resbed[:,1][i2].astype(int), kind='stable')
    i0 = np.argsort(resbed[:,0][i2[i1]], kind='stable')
    srti = (i2[i1])[i0]
    resbed = resbed[srti]
    np.savetxt(resfn, resbed, delimiter="\t", fmt='%s')

def preprocess():
    print("gen_tr_len_mt_bed", flush=True)
    gen_tr_len_mt_bed(TIS)
    print("gen_snp_motif_gt_bed", flush=True)
    subprocess.run(["bash", "fn0-5_.2.sh", INDIR, TIS])
    print("gen_mapping_bed", flush=True)
    gen_mapping_bed(TIS)
    print("gen_residualized_expression_mat", flush=True)
    gen_residualized_expression_mat(TIS)

TIS = sys.argv[1]
EMOTIF_DIR = sys.argv[2]
INDIR = sys.argv[3]
AKMS_BC_FN = sys.argv[4]
ACGT_BC_FN = sys.argv[5]
FILTER_FN = sys.argv[6]

preprocess()
```
