## Supplemental code for "The motif composition of variable-number tandem repeats impacts gene expression": fn0-5_.2.sh

```
#!/usr/bin/env bash

indir=$1
t=$2
cols=$(./fn0-5_.2.filter_sample.py $t)

cat <(zcat $indir/$t.snp.gt.tsv.gz | awk 'BEGIN {OFS="\t"} {split($1,vs1,"_"); $1=$1"\t."; print vs1[1], vs1[2]-1, vs1[2], $0}') \
    <(zcat $indir/$t.emotif_tr.bed.gz) |
sort -k1,1 -k2,2n -k3,3n --parallel=8 --buffer-size=24G --temporary-directory=/scratch2/tsungyul/tmp | 
awk 'BEGIN {OFS="\t"} {$5=NR-1; print $0}' | 
cut -f $cols | 
gzip >$indir/$t.snp_emotif_tr.gt.bed.gz
```
