## Supplemental code for "The motif composition of variable-number tandem repeats impacts gene expression": fn0-5_.3.R

```
library(susieR)
library(data.table)

t <- commandArgs(trailingOnly=TRUE)[1]
param_L <- as.integer(commandArgs(trailingOnly=TRUE)[2])
print(paste("tissue:", t, "L:", param_L))
indir <- commandArgs(trailingOnly=TRUE)[3]
od <- commandArgs(trailingOnly=TRUE)[4]
ofn <- paste(od, paste(t, ".pip.", param_L, ".txt", sep=""), sep="/")

print("buffered reading of gt")
gtfile <- paste(indir, paste(t, ".snp_emotif_tr.gt.bed.gz", sep=""), sep="/")
print(gtfile)
gtin <- file(gtfile, "r")
SI <- 1
EI <- 1
while (EI - SI < 500000) {
	line <- readLines(gtin, n=1)
	if (length(line) == 0) { break }
	vs <- strsplit(line, '\t')[[1]]
	if (EI == 1) {
		ncol <- 500000
		nrow <- length(vs) - 5
		gt <- matrix(numeric(nrow*ncol), nrow=nrow, ncol=ncol)
	}
	gt[ ,(EI-SI+1)] <- as.numeric(vs[-(1:5)])
	EI <- EI + 1
}

print("reading mapping")
mapfn <- paste(indir, paste(t, ".egenes.tss_cis_1Mb.map.snp_emotif_tr.bed", sep=""), sep="/")
mapping <- read.table(mapfn, sep="\t")

print("reading resmat")
resfn <- Sys.glob(paste(indir, paste(t, ".resmat.*egene_X_*sample.bed.gz", sep=""), sep="/"))
resmat <- t(as.matrix(read.table(gzfile(resfn), sep="\t")[ ,-(1:4)]))

ff <- FALSE
for (i in 1:dim(mapping)[1]) {
	yi <- as.integer(mapping[i,5]) + 1
	y <- scale(resmat[ ,yi])
	xsi <- as.integer(mapping[i,6]) + 1
	xei <- as.integer(mapping[i,7]) + 1

	# buffered reading
	if (xei >= EI) {
		print("buffered reading of gt")
		if (xsi < EI) {
			gt[ ,1:(EI-xsi)] <- gt[ ,(xsi-SI+1):(EI-SI)]
		} 
		else if (xsi > EI) {
			while (EI < xsi) {
				line <- readLines(gtin, n=1)
				if (length(line) == 0) { assert("buffered reading failed", FALSE) }
				EI <- EI + 1
			}
		}
		SI <- xsi
		while (EI - SI < 500000) {
			line <- readLines(gtin, n=1)
			if (length(line) == 0) { break }
			vs <- strsplit(line, '\t')[[1]]
			gt[ ,(EI-SI+1)] <- as.numeric(vs[-(1:5)])
			EI <- EI + 1
		}
	}

	X <- gt[ ,(xsi-SI+1):(xei-SI+1)]
	namask <- (X==3) | (is.na(X))
	for (j in 1:dim(X)[2]) { # replace missing gt with mean gt across samples
		if (sum(namask[ ,j])) { 
			X[namask[ ,j], j] <- mean(X[!namask[ ,j], j]) 
		}
	}
	X_ <- apply(X, 2, scale)
	nanmask <- !is.nan(X_[1, ])
	res <- susie(X_[ ,nanmask], y, L=param_L, na.rm=FALSE)
	nvar <- xei - xsi + 1
	pip <- matrix(NA, 1, nvar)
	pip[nanmask] <- res$pip
	cat(".")
	if (is.null(res$sets$cs$L1)) {
		fwrite(list(NA), ofn, append=ff, sep="\t", col.names=FALSE, na="nan")
	} else {
		inds <- c(1:nvar)
		L1 <- t(inds[nanmask][res$sets$cs$L1])
		fwrite(as.data.table(L1), ofn, append=ff, sep="\t", col.names=FALSE, na="nan")
	}
	ff <- TRUE
	fwrite(as.data.table(pip), ofn, append=ff, sep="\t", col.names=FALSE, na="nan")
}
cat("\n")
close(gtin)
```
