## Supplemental code for "The motif composition of variable-number tandem repeats impacts gene expression": fn0-5_.4.py

```
#!/usr/bin/env python3

import sys
import numpy as np
import pandas as pd
import matplotlib
import matplotlib.pyplot as plt
import glob
import statsmodels.api as sm
import gzip

matplotlib.rc('font', size=7)
matplotlib.rc('axes', titlesize=7)
matplotlib.rc('xtick', labelsize=5)
matplotlib.rc('ytick', labelsize=5)

def load_pip_cs_v1(t):
    n = 0
    n0 = 0
    pips = {}
    cs = {}
    with open(f"{OUTDIR}/{t}.pip.{FINEMAP_L}.txt") as f:
        nl = 0
        for line in f:
            if nl % 2:
                vs = np.array(line.split(), ndmin=1).astype(float)
                if np.any(vs):
                    pips[n] = vs
                else:
                    pips[n] = None
                    n0 += 1
                n += 1
            else:
                if line[0] == "n":
                    cs[n] = None
                else:
                    vs = np.array(line.split(), ndmin=1).astype(int)
                    cs[n] = vs - 1 # R script report 1-based index
            nl += 1
    print(n, n0)
    return pips, cs

def load_isTR_Motif(t):
    fn = f"{INDIR}/{t}.snp_emotif_tr.gt.bed.gz"
    isTR_Motif = []
    with gzip.open(fn, 'rt') as f:
        for line in f:
            vname = line.split()[3]
            if vname[0] == "t":
                if len(vname.split("_")) == 6:
                    isTR_Motif.append(2)
                else:
                    isTR_Motif.append(1)
            else:
                isTR_Motif.append(0)
    return np.array(isTR_Motif)

def load_mapping(t):
    fn = f"{INDIR}/{t}.egenes.tss_cis_1Mb.map.snp_emotif_tr.bed"
    mapping = np.loadtxt(fn, dtype=object)
    return mapping

def load_resmat(t):
    fn = glob.glob(f"{INDIR}/{t}.resmat.*egene_X_*sample.bed.gz")[0]
    ncol = np.loadtxt(fn, max_rows=1, dtype=object).size
    resmat = np.loadtxt(fn, usecols=np.arange(4,ncol))
    return resmat

def get_vari2triki(t):
    vari2triki = {}
    with gzip.open(f"{INDIR}/{t}.snp_emotif_tr.gt.bed.gz", 'rt') as f:
        for line in f:
            vs = line.split()
            if vs[3][0] == "t":
                vvs = vs[3].split("_")
                tri = int(vvs[1])
                ki = int(vvs[-1]) if len(vvs) == 6 else -1
                vari = int(vs[4])
                vari2triki[vari] = (tri,ki)
    return vari2triki

def zscore(arr):
    return (arr - np.mean(arr)) / np.std(arr)

def viz_causal_emotif_pip_summarize(t, pips, cs, isTR_Motif, mapping, resmat, mti2triki):
    od = f"{OUTDIR}/{t}"
    fn = f"{INDIR}/{t}.snp_emotif_tr.gt.bed.gz"    
    fin = gzip.open(fn, 'rt')
    SI, EI = 0, 0
    print("buffered reading", flush=True)
    while EI - SI < 500000:
        line = fin.readline()
        if not line: 
            fin.close()
            assert False
        vs = [float(v) for v in line.split()[5:]]
        if EI == 0:
            gt = np.zeros([len(vs), 500000])
        gt[:, EI-SI] = vs
        EI += 1
                    
    ks_cand = []
    for k, pip in pips.items():
        if pip is None: continue
        if not np.any(np.isfinite(pip)): continue
        if np.nanmax(pip) >= 0.8:
            si, ei = int(mapping[k,5]), int(mapping[k,6])+1
            istr_motif = isTR_Motif[si:ei]
            b_istr_motif = istr_motif.astype(bool)
            if not np.any(b_istr_motif): continue
            if not np.any(np.isfinite(pip[b_istr_motif])): continue
            maxmtpip = np.nanmax(pip[b_istr_motif])
            if maxmtpip < 0.8: continue
            if ei > EI:
                print("buffered reading", flush=True)
                if si < EI:
                    gt[:, :EI-si] = gt[:, si-SI:EI-SI]
                elif si > EI:
                    while EI < si:
                        line = fin.readline()
                        if not line: 
                            fin.close()
                            assert False
                        EI += 1
                SI = si
                while EI - SI < 500000:
                    line = fin.readline()
                    if not line: 
                        gt[:, EI-SI:] = 0
                        break
                    vs = [float(v) for v in line.split()[5:]]
                    gt[:, EI-SI] = vs
                    EI += 1
            
            cmtis = np.arange(pip.size)[b_istr_motif][np.nonzero(pip[b_istr_motif] >= 0.8)] + si # any emotif, not necessarily the original lead eMotif
            maxmti = np.arange(pip.size)[b_istr_motif][np.nanargmax(pip[b_istr_motif])] + si
            assert ei-si == pip.size
            assert maxmti < ei
            
            if cmtis.size:
                cm0 = istr_motif == 0
                cm1 = istr_motif == 1
                cm2 = istr_motif == 2
                plt.figure(figsize=(2.6,2.2), dpi=600)
                plt.plot(np.arange(pip.size)[cm0], pip[cm0], '.k', markersize=1, label="SNP")
                plt.plot(np.arange(pip.size)[cm1], pip[cm1], '.r', markersize=1, label="TR")
                plt.plot(np.arange(pip.size)[cm2], pip[cm2], '.b', markersize=1, label="eMotif")
                plt.xlabel("Variant index")
                plt.ylabel("PIP")
                plt.legend(fontsize=5)
                plt.tight_layout()
                plt.savefig(f"{od}/{k}.pip.pdf")
                plt.close()

            for cmti in cmtis:
                ge = mapping[k,3]
                tri, ki = mti2triki[cmti]
                print(f"candidate {ge}.{tri}.{ki} reported", flush=True)
                ks_cand.append(k)
                
                x = np.copy(gt[:, cmti-SI])
                m0 = ~np.isfinite(x)
                if np.any(m0):
                    x[m0] = np.mean(x[~m0])
                x = zscore(x)
                m = (x <= 2) & (x >= -2)
                x = zscore(x[m])
                yi = int(mapping[k,4])
                y = zscore(resmat[yi][m])
                res = sm.OLS(y, sm.add_constant(x)).fit()
                print(f"{ge}.{tri}.{ki}", k, si, ei, ei-si, cmti-si, mapping[k], f"{res.pvalues[1]:.4e}", f"{res.params[1]:.3f}", y.size, np.sum(m0), flush=True)
                
                plt.figure(figsize=(2.2,2), dpi=200)
                plt.plot(x, y, '.k', markersize=1, alpha=0.5)
                plt.xlabel("Norm. genotype")
                plt.ylabel("Norm. expression")
                plt.tight_layout()
                plt.savefig(f"{od}/{ge}.{tri}.{ki}.reg.pdf")
                plt.close()
    np.savetxt(f"{OUTDIR}/{t}.candidate_index.txt", ks_cand, fmt="%s")

    # Fraction of emotifs in the top credible set
    print("Computing summary stats", flush=True)
    nmap = mapping.shape[0]
    out = np.zeros([nmap, 8], dtype=object)
    for i in range(nmap):
        inds = cs[i]
        si = int(mapping[i,5])
        N0, N, maxsnppip, maxmtpip, pass_pip_thresh, tri, ki, cinds_str = np.nan, np.nan, np.nan, np.nan, False, np.nan, np.nan, "None"
        if inds is not None:
            N0 = np.sum(isTR_Motif[inds+si].astype(bool))
            N = inds.size
        pip = pips[i]
        if np.any(np.isfinite(pip)):
            istr_motif = isTR_Motif[si:si+pip.size]
            b_istr_motif = istr_motif.astype(bool)
            if np.any(pip[~b_istr_motif]):
                if np.any(np.isfinite(pip[~b_istr_motif])):
                    maxsnppip = np.nanmax(pip[~b_istr_motif])
            if np.any(b_istr_motif):
                if np.any(np.isfinite(pip[b_istr_motif])):
                    maxtrmtpip = np.nanmax(pip[b_istr_motif])
                    pass_pip_thresh = maxtrmtpip >= 0.8
                    maxtrmti = np.arange(pip.size)[b_istr_motif][np.nanargmax(pip[b_istr_motif])] + si # index for *.snp_emotif_tr.gt.bed.gz
                    tri, ki = vari2triki[maxtrmti]
                    cvaris = np.arange(pip.size)[b_istr_motif][np.nonzero(pip[b_istr_motif] >= 0.8)] + si
                    causal_inds = [vari2triki[cvari] for cvari in cvaris]
                    cinds_str = ";".join([f"({tri},{ki})" for tri, ki in causal_inds])
        out[i] = [N0, N, maxsnppip, maxtrmtpip, pass_pip_thresh, tri, ki, cinds_str]
    df = pd.DataFrame(out, columns=["|L1_emotif_tr|", "|L1|", "max(pip_snp)", "max(pip_emotif_tr)", "max(pip_emotif_tr)>=0.8", "max_pip_tri", "max_pip_ki", "all_pip>0.8_(tri,ki)"])
    df["|L1_emotif_tr|"] = df["|L1_emotif_tr|"].apply(lambda x: f"{x:.0f}")
    df["|L1|"] = df["|L1|"].apply(lambda x: f"{x:.0f}")
    df["max(pip_snp)"] = df["max(pip_snp)"].apply(lambda x: f"{x:.4e}")
    df["max(pip_emotif_tr)"] = df["max(pip_emotif_tr)"].apply(lambda x: f"{x:.4e}")
    df["max(pip_emotif_tr)>=0.8"] = df["max(pip_emotif_tr)>=0.8"].apply(lambda x: f"{bool(x)}")
    df["max_pip_tri"] = df["max_pip_tri"].apply(lambda x: f"{x:.0f}")
    df["max_pip_ki"] = df["max_pip_ki"].apply(lambda x: f"{x:.0f}")
    df.to_csv(f"{OUTDIR}/{t}.finemap.summary.tsv", sep="\t", na_rep="nan")


t = sys.argv[1]
INDIR = sys.argv[2]
OUTDIR = sys.argv[3]
FINEMAP_L = int(sys.argv[4])
print("loading pip & cs", flush=True)
pips, cs = load_pip_cs_v1(t)
print("loading isTR_Motif", flush=True)
isTR_Motif = load_isTR_Motif(t)
print("loading mapping", flush=True)
mapping = load_mapping(t)
print("loading resmat", flush=True)
resmat = load_resmat(t)
print("Computing vari2triki", flush=True)
vari2triki = get_vari2triki(t)
print("finding causal emotifs", flush=True)
viz_causal_emotif_pip_summarize(t, pips, cs, isTR_Motif, mapping, resmat, vari2triki)
```
