## Supplemental code for "The motif composition of variable-number tandem repeats impacts gene expression": fn0-5_.5.py

```
#!/usr/bin/env python3

import sys
import numpy as np
import pandas as pd
import pickle
import gzip
import vntrutils as vu
from collections import defaultdict

KSIZE = 21

def loadGTindex():
    with open("/project/mchaisso_100/cmb-17/vntr_genotyping/rpgg2_k21_84k/hgsvc/clinic/cdbg/v1/ks.ccks.cck_ns.ki_map.pickle", 'rb') as f:
        _, ccks, cck_ns, _ = pickle.load(f)
    return ccks, cck_ns
ccks, cck_ns = loadGTindex()

def get_tri2bed():
    tri2bed = {}
    trbed = np.loadtxt("/project/mchaisso_100/cmb-17/vntr_genotyping/rpgg2_k21_84k/hgsvc/clinic/output/pan.tr.mbe.v2.bed", usecols=[0,1,2], dtype=object)
    for i, (ch, s, e) in enumerate(trbed):
        tri2bed[i] = (ch,s,e)
    return tri2bed

def indexGeneList(tissue):
    tisGeneList = np.loadtxt(f'/project/mchaisso_100/cmb-17/vntr_genotyping/gtex/expression/GTEx_Analysis_v8_eQTL_expression_matrices/{tissue}.v8.normalized_expression.bed.gz', dtype=object, skiprows=1, usecols=[3])
    tisGene2ind = {}
    for ind, gene in enumerate(tisGeneList):
        tisGene2ind[gene] = ind
    return tisGeneList, tisGene2ind

def load_pips(t, OUTDIR):
    n = 0
    n0 = 0
    pips = {}
    with open(f"{OUTDIR}/{t}.pip.{FINEMAP_L}.txt") as f:
        nl = 0
        for line in f:
            if nl % 2:
                vs = np.array(line.split(), ndmin=1).astype(float)
                if np.any(vs):
                    pips[n] = vs
                else:
                    pips[n] = None
                    n0 += 1
                n += 1
            nl += 1
    return pips

class eGeneStat():
    def __init__(self):
        self.ch = []
        self.s = []
        self.e = []
        self.tri = []
        self.ki = []
        self.pip = []
        self.p = []
        self.b = []
        self.bse = []
        self.mtf = []
    def size(self):
        return len(self.ch)

def get_gi2tri2pb(tissue):
    tbl = np.loadtxt(f"{VNTR_DIR}/{tissue}.v8.allGenes.txt", dtype=object, usecols=[0,9,13,14,15])
    tisGeneList, tisGene2ind = indexGeneList(tissue)
    gi2tri2pb = defaultdict(dict)
    for ge, tri, p, b, bse in tbl:
        gi = tisGene2ind[ge]
        tri = int(tri)
        p, b, bse = float(p), float(b), float(bse)
        gi2tri2pb[gi][tri] = (p,b,bse)
    return gi2tri2pb

def validateFinemap(tis):
    fout = f"{OUTDIR}/{tis}.finemap.summary.validated.tsv"
    tri2bed = get_tri2bed()
    columns = ["tissue", "gene_id", "gene_name", "chr", "start", "end", "num_var", 
               "|L1_motif_tr|", "|L1|", "max(pip_snp)", "max(pip_motif_tr)", "num_emotif_tr", "TR_chr", "TR_start", "TR_end", "TR_locus", "ki", "pip", "min(p_nominal)", "p_nominal", "slope", "slope_se", "motif"]
    print(".", end="", flush=True)
    _, tisGene2ind = indexGeneList(tis)
    print("i",end="", flush=True)
    pips = load_pips(tis, OUTDIR)
    print("p", end="", flush=True)
    allp = pickle.load(open(f"{MOTIF_DIR}/{tis}.allp.pickle", 'rb'))
    print("q", end="", flush=True)
    allb = pickle.load(open(f"{MOTIF_DIR}/{tis}.allb.pickle", 'rb'))
    print("b", end="", flush=True)
    tmi2vari = pickle.load(open(f"{INDIR}/pickle/{tis}.tmi2vari.pickle", 'rb'))
    print("k", end="", flush=True)
    sis = np.loadtxt(f"{INDIR}/{tis}.egenes.tss_cis_1Mb.map.snp_emotif_tr.bed", usecols=5, dtype=int, ndmin=1)
    print("s", flush=True)
    pth = float(dict(np.loadtxt(f"{MOTIF_ANADIR}/nominal_p_genome_wide_cutoff.txt", dtype=object).tolist())[tis])
    pth_tr = float(dict(np.loadtxt(f"{VNTR_ANADIR}/nominal_p_genome_wide_cutoff.txt", dtype=object).tolist())[tis])
    gi2tri2pb = get_gi2tri2pb(tis)
    
    tbl1 = np.loadtxt(f"{MOTIF_DIR}/{tis}.v8.egenes.txt", dtype=object, ndmin=2, usecols=range(6))
    tbl1 = np.hstack((np.repeat([tis], tbl1.shape[0])[:,None], tbl1))
    tbl2 = pd.read_csv(f"{OUTDIR}/{tis}.finemap.summary.tsv", sep="\t", index_col=0, na_values="nan").to_numpy()
    mask = np.zeros(tbl2.shape[0], dtype=bool)
    for i, trikistr in enumerate(tbl2[:,7]):
        if type(trikistr) is str:
            mask[i] = True        
    df = pd.DataFrame(index=range(np.sum(mask)),columns=columns)
    df.iloc[:,:7] = tbl1[mask]
    for oi, (i, trikistr) in enumerate(zip(np.nonzero(mask)[0], tbl2[mask,7])):
        if type(trikistr) is str:
            egs = eGeneStat()
            #mask[i] = True # redundant
            trikis = trikistr.split(";")
            for triki in trikis:
                tri, ki = [int(float(v)) for v in triki[1:-1].split(",")]
                tmi = ki if ki >= 0 else -(tri+1)
                ge = tbl1[i,1]
                gi = tisGene2ind[ge]

                assert gi in allp
                if tri not in allp[gi]:
                    print(f"{tbl1[i,2]}\t{tri} not tested in eQTL mapping", flush=True)
                    continue
                if ki >= 0:
                    if ki not in allp[gi][tri]:
                        print(f"{tbl1[i,2]}\t{tri}\t{ki} not tested in eQTL mapping", flush=True)
                        continue
                    p = allp[gi][tri][ki]
                    b, bse = allb[gi][tri][ki]
                else:
                    p, b, bse = gi2tri2pb[gi][tri]
                if (ki >= 0 and p < pth) or (ki < 0 and p < pth_tr):
                    ch, s, e = tri2bed[tri]
                    si = sis[i]
                    vari = tmi2vari[tmi]
                    pip = pips[i][vari-si]
                    mtf = vu.decodeNumericString(ccks[ki], KSIZE+cck_ns[ki]-1) if ki >= 0 else "NA"
                    egs.ch.append(ch)
                    egs.s.append(s)
                    egs.e.append(e)
                    egs.tri.append(str(tri))
                    egs.ki.append(str(ki))
                    egs.pip.append(f"{pip:.3e}")
                    egs.p.append(f"{p:.4e}")
                    egs.b.append(f"{b:.4e}")
                    egs.bse.append(f"{bse:.4e}")
                    egs.mtf.append(mtf)
                else:
                    print(f"{tbl1[i,2]}\t{tri}\t{ki}\t{p:.2e}\t{b:.3f} not significant in eQTL mapping", flush=True)
            df["|L1_motif_tr|"][oi] = f"{float(tbl2[i,0]):.0f}"
            df["|L1|"][oi] = f"{float(tbl2[i,1]):.0f}"
            df["max(pip_snp)"][oi] = f"{float(tbl2[i,2]):.3e}"
            df["max(pip_motif_tr)"][oi] = f"{float(tbl2[i,3]):.3e}"
            df["num_emotif_tr"][oi] = egs.size()

            if egs.size():
                df["TR_chr"][oi] = ",".join(egs.ch)
                df["TR_start"][oi] = ",".join(egs.s)
                df["TR_end"][oi] = ",".join(egs.e)
                df["TR_locus"][oi] = ",".join(egs.tri)
                df["ki"][oi] = ",".join(egs.ki)
                df["pip"][oi] = ",".join(egs.pip)
                df["min(p_nominal)"][oi] = f"{min([float(v) for v in egs.p]):.4e}"
                df["p_nominal"][oi] = ",".join(egs.p)
                df["slope"][oi] = ",".join(egs.b)
                df["slope_se"][oi] = ",".join(egs.bse)
                df["motif"][oi] = ",".join(egs.mtf)
            
    df.to_csv(fout, sep="\t", na_rep=".")

tis = sys.argv[1]
MOTIF_DIR = sys.argv[2]
MOTIF_ANADIR = sys.argv[3]
VNTR_DIR = sys.argv[4]
VNTR_ANADIR = sys.argv[5]
INDIR = sys.argv[6]
OUTDIR = sys.argv[7]
FINEMAP_L = int(sys.argv[8])
validateFinemap(tis)
```
