## Supplemental code for "The motif composition of variable-number tandem repeats impacts gene expression": fn0-5_.5.tmi2vari.py

```
#!/usr/bin/env python3

import sys
import pickle

t = sys.argv[1]
INDIR = sys.argv[2]
# tmi: TR or motif index in RPGG
# vari: variant index in bed.gz file
tmi2vari = {}
with open("/dev/stdin") as f:
    for line in f:
        vs = line.split()
        if vs[0][0] == "-": # is TR dosage. Use -(tr_index + 1) to encode tr_index
            tmi = int(vs[0]) - 1
        else: # is Motif dosage
            tmi = int(vs[0])
        vari = int(vs[1])
        tmi2vari[tmi] = vari
pickle.dump(tmi2vari, open(f"{INDIR}/pickle/{t}.tmi2vari.pickle", 'wb'))
```
