## Supplemental code for "The motif composition of variable-number tandem repeats impacts gene expression": vntrutils.py

```
import numpy as np
#from sklearn.linear_model import LinearRegression
import statsmodels.api as sm
import matplotlib
import matplotlib.pyplot as plt

base = {'A':0, 'C':1,  'G':2,  'T':3}
baseinv = ['A', 'C', 'G', 'T']
baseNumConversion = [
 'A', 'C', 'G', 'T', 127, 127, 127, 127,
 127, 127, 127, 127, 127, 127, 127, 127,
 127, 127, 127, 127, 127, 127, 127, 127,
 127, 127, 127, 127, 127, 127, 127, 127,
 127, 127, 127, 127, 127, 127, 127, 127,
 127, 127, 127, 127, 127, 127, 127, 127,
 127, 127, 127, 127, 127, 127, 127, 127,
 127, 127, 127, 127, 127, 127, 127, 127,
 127,   0, 127,   1, 127, 127, 127,   2,
 127, 127, 127, 127, 127, 127, 127, 127,
 127, 127, 127, 127,   3, 127, 127, 127,
 127, 127, 127, 127, 127, 127, 127, 127,
 127,   0, 127,   1, 127, 127, 127,   2,
 127, 127, 127, 127, 127, 127, 127, 127,
 127, 127, 127, 127,   3, 127, 127, 127]
baseComplement = [
   3,   2,   1,   0, 127, 127, 127, 127,
 127, 127, 127, 127, 127, 127, 127, 127,
 127, 127, 127, 127, 127, 127, 127, 127,
 127, 127, 127, 127, 127, 127, 127, 127,
 127, 127, 127, 127, 127, 127, 127, 127,
 127, 127, 127, 127, 127, 127, 127, 127,
 127, 127, 127, 127, 127, 127, 127, 127,
 127, 127, 127, 127, 127, 127, 127, 127,
 127, 'T', 127, 'G', 127, 127, 127, 'C',
 127, 127, 127, 127, 127, 127, 'N', 127,
 127, 127, 127, 127, 'A', 127, 127, 127,
 127, 127, 127, 127, 127, 127, 127, 127,
 127, 't', 127, 'g', 127, 127, 127, 'c',
 127, 127, 127, 127, 127, 127, 'n', 127,
 127, 127, 127, 127, 'a', 127, 127, 127]
byteRC = [
 255, 191, 127,  63, 239, 175, 111,  47, 223, 159,
  95,  31, 207, 143,  79,  15, 251, 187, 123,  59,
 235, 171, 107,  43, 219, 155,  91,  27, 203, 139,
  75,  11, 247, 183, 119,  55, 231, 167, 103,  39,
 215, 151,  87,  23, 199, 135,  71,   7, 243, 179,
 115,  51, 227, 163,  99,  35, 211, 147,  83,  19,
 195, 131,  67,   3, 254, 190, 126,  62, 238, 174,
 110,  46, 222, 158,  94,  30, 206, 142,  78,  14,
 250, 186, 122,  58, 234, 170, 106,  42, 218, 154,
  90,  26, 202, 138,  74,  10, 246, 182, 118,  54,
 230, 166, 102,  38, 214, 150,  86,  22, 198, 134,
  70,   6, 242, 178, 114,  50, 226, 162,  98,  34,
 210, 146,  82,  18, 194, 130,  66,   2, 253, 189,
 125,  61, 237, 173, 109,  45, 221, 157,  93,  29,
 205, 141,  77,  13, 249, 185, 121,  57, 233, 169,
 105,  41, 217, 153,  89,  25, 201, 137,  73,   9,
 245, 181, 117,  53, 229, 165, 101,  37, 213, 149,
  85,  21, 197, 133,  69,   5, 241, 177, 113,  49,
 225, 161,  97,  33, 209, 145,  81,  17, 193, 129,
  65,   1, 252, 188, 124,  60, 236, 172, 108,  44,
 220, 156,  92,  28, 204, 140,  76,  12, 248, 184,
 120,  56, 232, 168, 104,  40, 216, 152,  88,  24,
 200, 136,  72,   8, 244, 180, 116,  52, 228, 164,
 100,  36, 212, 148,  84,  20, 196, 132,  68,   4,
 240, 176, 112,  48, 224, 160,  96,  32, 208, 144,
  80,  16, 192, 128,  64,   0]

def getRCstring(string):
    RCstring = np.empty(len(string), dtype='U')
    rlen = len(string)
    for i in range(rlen):
        RCstring[rlen-i-1] = baseComplement[ord(string[i])]

    return ''.join(RCstring)


def decodeNumericString(num, k):
    string = ''
    for i in range(k):
        string = baseinv[(num % 4)] + string
        num >>= 2

    return string


def getRCkmer(kmer, k):
    rckmer = 0
    while k >= 4:
        rckmer <<= 8
        rckmer += byteRC[kmer & 0xff]
        kmer >>= 8
        k -= 4
    if k > 0:
        rckmer <<= (k<<1)
        rckmer += (byteRC[kmer] >> ((4-k)<<1))
    return rckmer


def encodeString(string):
    """Direct encoding. Use string2CaKmer() for canonical encoding"""
    numericString = 0
    for i in range(len(string)):
        numericString = (numericString << 2) + base[string[i]]

    return numericString


def string2CaKmer(string):
    kmer = encodeString(string)
    rckmer = getRCkmer(kmer, len(string))
    return kmer if kmer <= rckmer else rckmer


def getNextKmer(beg, seq, k):
    if beg + k >= len(seq):
        return (len(seq), 0)

    validlen = 0
    while validlen != k:
        if beg + k >= len(seq):
            return (len(seq), 0)
        if seq[(beg + validlen)] not in base:
            beg = beg + validlen + 1
            validlen = 0
        else:
            validlen += 1

    return (beg, encodeString(seq[beg:beg + k]))


def buildNuKmers(read, k, leftflank=0, rightflank=0, count=True):
    rlen = len(read)
    mask = (1 << 2*(k-1)) - 1
    kmers = {}

    beg, kmer = getNextKmer(leftflank, read, k)
    if beg == rlen: return kmers
    rckmer = getRCkmer(kmer, k)

    it = iter(range(beg, rlen-k-rightflank+1))
    for i in it:
        canonicalkmer = rckmer if kmer > rckmer else kmer
        if canonicalkmer not in kmers: kmers[canonicalkmer] = 0
        kmers[canonicalkmer] += count

        if i + k >= rlen: return kmers
        if read[i + k] not in baseinv:
            nbeg, kmer = getNextKmer(i+k+1, read, k)
            rckmer = getRCkmer(kmer, k)
            for j in range(nbeg-i-1):
                next(it, None)
        else:
            kmer = ((kmer & mask) << 2) + base[read[i + k]]
            rckmer = (rckmer >> 2) + (3-base[read[i+k]] << 2*(k-1))

    return kmers

def read2kmers(read, k, leftflank=0, rightflank=0, dtype='uint64', canonical=True):
    """
    will return max_val_of_uint64 (-1) if there's 'N' within kmer
    """
    rlen = len(read)
    if rlen - k - leftflank - rightflank + 1 <= 0: return np.array([])

    mask = (1 << 2*(k-1)) - 1
    kmers = np.zeros(rlen-k-leftflank-rightflank+1, dtype=dtype) - 1

    beg, kmer = getNextKmer(leftflank, read, k)
    if beg == rlen: return kmers
    rckmer = getRCkmer(kmer, k)

    it = iter(range(beg, rlen-k-rightflank+1))
    for i in it:
        canonicalkmer = rckmer if kmer > rckmer else kmer
        kmers[i-beg] = canonicalkmer if canonical else kmer

        if i + k >= rlen: return kmers
        if read[i + k] not in baseinv:
            nbeg, kmer = getNextKmer(i+k+1, read, k)
            rckmer = getRCkmer(kmer, k)
            for j in range(nbeg-i-1):
                next(it, None)
        else:
            kmer = ((kmer & mask) << 2) + base[read[i + k]]
            rckmer = (rckmer >> 2) + (3-base[read[i+k]] << 2*(k-1))

    return kmers


def writeKmerDict(fname, kmerDB, precision=0):
    with open(fname+".kmers", 'w') as f:
        for locus, kmers in kmerDB.items():
            f.write(">{}\n".format(locus))
            for kmer, count in kmers.items():
                f.write("{:<21.0f}\t{:.{prec}f}\n".format(kmer, count, prec=precision))


def checkTable(table):
    if table.size:
        if np.any(table[:, 1]):
            return True
    return False


def assignNewTable(kmerDB, locus, table, sort, kmerName, threshold):
    if checkTable(table):
        table = table[(table[:, 1] >= threshold)]
        if sort:
            table = table[table[:, 0].argsort()]
        if kmerName:
            kmerDB[locus] = table
        else:
            kmerDB[locus] = table[:, 1]
    else:
        kmerDB[locus] = np.array([])


def IncrementKmerCount(kmerDB, locus, table, sort, kmerName):
    if checkTable(table):
        assert sort, "invalid argument: {'sort': False}"
        assert table.size == kmerDB[locus].size, "inconsistent table size"
        table = table[table[:, 0].argsort()]
        if kmerName:
            kmerDB[locus][:, 1] += table[:, 1]
        else:
            kmerDB[locus] += table[:, 1]
    else:
        return


def assignKmerTable(kmerDB, locus, table, sort, kmerName, threshold):
    table = np.array(table, dtype=int)
    if locus not in kmerDB:
        assignNewTable(kmerDB, locus, table, sort, kmerName, threshold)
    else:
        if kmerDB[locus].size:
            assert threshold == 0, "filtering while incrementing counts!"
            IncrementKmerCount(kmerDB, locus, table, sort, kmerName)
        else:
            assignNewTable(kmerDB, locus, table, sort, kmerName, threshold)


def readKmers(fname, kmerDB, sort=True, kmerName=False, threshold=0):
    """ read a kmer file as a table"""
    with open(fname) as f:
        table = []
        locus = 0
        f.readline()
        for line in f:
            if line[0] == '>':
                assignKmerTable(kmerDB, locus, table, sort, kmerName, threshold)
                table = []
                locus += 1
            else:
                table.append(line.split())
        else:
            assignKmerTable(kmerDB, locus, table, sort, kmerName, threshold)

def readKmersWithIndex(fn_kmc, fn_index, kmerDB, sort=True, kmerName=False, threshold=0):
    """ read kmer count and index files as a table"""
    f0 = open(fn_kmc)
    f1 = open(fn_index)
    table = []
    locus = 0
    f1.readline()
    for line in f1:
        if line[0] == '>':
            if len(table):
                table = np.array(table, dtype=int, ndmin=2)
                table[:,1] = [f0.readline().rstrip() for i in range(len(table))]
            assignKmerTable(kmerDB, locus, table, sort, kmerName, threshold)
            table = []
            locus += 1
        else:
            table.append([line.rstrip(), 0])
    else:
        if len(table):
            table = np.array(table, dtype=int, ndmin=2)
            table[:,1] = [f0.readline().rstrip() for i in range(len(table))]
        assignKmerTable(kmerDB, locus, table, sort, kmerName, threshold)
    f0.close()
    f1.close()

def readKmerDict(fname, kmerDB=None, threshold=0, checkkmer=False):
    """ read a kmer file as a dictionary """
    
    hasInput = False if kmerDB is None else True
    if not hasInput: kmerDB = {}

    with open(fname) as f:
        locus = -1
        for line in f:
            if line[0] == '>':
                locus += 1
                if locus not in kmerDB:
                    kmerDB[locus] = {}
            else:
                kmer, count = [int(v) for v in line.split()]
                if count >= threshold:
                    if kmer in kmerDB[locus]:
                        kmerDB[locus][kmer] += count
                    else:
                        if checkkmer: assert False
                        else: kmerDB[locus][kmer] = count

    if not hasInput: return kmerDB


def readKms(fin, ki_tr, out=None):
    """
    Read kmers and compute sum for each locus.
    Use out=ARRAY to change ARRAY inplace.
    REQUIRE ki_tr since danbing-tk v1.3 since .kmers file does not contain locus info.
        Use ki_tr=None for backward compatibility.
    """
    ndb = out is None
    if out is None:
        out = []
    with open(fin) as f:
        if ki_tr is None:
            idx = -1
            for line in f:
                if line[0] == ">":
                    idx += 1
                    if ndb:
                        out.append(0)
                else:
                    out[idx] += int(line.split()[1])
        else:
            idx = 0
            ki = -1
            out.append(0)
            for line in f:
                ki += 1
                if ki >= ki_tr[idx]:
                    idx += 1
                    if ndb:
                        out.append(0)
                out[idx] += int(line.rstrip())

    if ndb:
        return out


def filterKmers(fin, fout, indf):
    """filter kmers by indices listed in indf, and reindex w/o gap"""
    gl = set(np.loadtxt(indf, dtype=int).tolist()) # good loci
    outf = open(fout, 'w')
    with open(fin) as f: # input kmers file
        oi = -1 # old index
        ni = 0 # new index
        for line in f:
            if line[0] == ">":
                oi += 1
                out = oi in gl
                if out:
                    outf.write(f">{ni}\n")
                    ni += 1
            else:
                if out:
                    outf.write(line)
    outf.close()


def countLoci(fname):
    with open(fname) as f:
        nloci = 0
        for line in f:
            if line[0] == '>':
                nloci += 1

    return nloci


def readFasta(fname, nloci=0):
    if nloci == 0:
        nloci = countLoci(fname)
    seqDB = np.empty(nloci, dtype=object)
    with open(fname) as f:
        locus = 0
        seq = ''
        f.readline()
        for line in f:
            if line[0] == '>':
                seqDB[locus] = seq
                seq = ''
                locus += 1
            else:
                seq = line.rstrip()
        else:
            seqDB[locus] = seq

    return seqDB

def readFasta2dict(fname):
    seqDB = {}
    with open(fname) as f:
        locus = 0
        seq = ''
        f.readline()
        for line in f:
            if line[0] == '>':
                seqDB[locus] = seq
                seq = ''
                locus += 1
            else:
                seq = line.rstrip()
        else:
            seqDB[locus] = seq

    return seqDB

def RecursiveRejection(x, y, fit_intercept=False):
    if fit_intercept:
        reg = sm.OLS(y, sm.add_constant(x)).fit()
    else:
        reg = sm.OLS(y, x).fit()
    res = reg.resid
    #reg = LinearRegression(fit_intercept=False).fit(x, y)
    #res = y - reg.predict(x)
    m = np.mean(res)
    s = np.std(res)
    logic = (np.abs(res - m) < 10 * s)[:, 0]
    if np.sum(logic) == 0:
        print('all entries are rejected')
        return (x[logic], y[logic])
    elif not np.all(logic):
        return RecursiveRejection(x[logic], y[logic])
    else:
        return (x, y)


def RejectOutlier(x, y, rule):
    logic = np.isfinite(x)[:,0] & np.isfinite(y)[:,0]
    if rule == 0:
        return (x[logic], y[logic], 0)
    if rule == 1 or rule == 2:
        logic &= ((x != 0)[:,0]) & ((y != 0)[:,0])
        return rule == 1 and (x[logic], y[logic], 0)
    if rule == 2 or rule == 3:
        x, y = x[logic], y[logic]
        x0, y0 = RecursiveRejection(x, y)
        return (x0, y0, x.size - x0.size)


def PlotRegression(x, y, xlabel='data_X', ylabel='data_Y', title='', fname='', outlier='invalid', pred=False, fit_intercept=False):
    if title == '':
        title = xlabel + '.' + ylabel
    if not fname:
        fname = title
    outlierRule = {"invalid":0, "invalid|zero":1, "invalid|bad|zero":2, "invalid|bad":3}[outlier]
    x1, y1, nOut = RejectOutlier(x, y, outlierRule)
    if nOut:
        print('# of non-trivial outliers in', title, nOut)
    if not x1.size or not y1.size:
        x1, y1, nOut = RejectOutlier(x, y, 1)
    if not x1.size or not y1.size:
        if pred:
            return (0, 0, 0, 0)
        return (0, 0, 0)
    if fit_intercept:
        reg = sm.OLS(y1, sm.add_constant(x1)).fit()
        a, b = reg.params[1], reg.params[0]
    else:
        reg = sm.OLS(y1, x1).fit()
        a, b = reg.params[0], None
    #reg = (LinearRegression(fit_intercept=fit_intercept)).fit(x1, y1)
    #a, b = np.asscalar(reg.coef_), np.asscalar(reg.intercept_) if fit_intercept else reg.intercept_
    #rsquare = reg.score(x1, y1)
    r2 = reg.rsquared
    if pred:
        y1_proj = np.sum(y1) / a
    if pred:
        return (a, b, r2, y1_proj)
    else:
        return (a, b, r2)


#def KmersLinReg(PBfname, ILfname, out, threshold=10, R2threshold=0, plot=False):
#    print("reading illumina kmers")
#    y = {}
#    readKmers(ILfname, y, sort=True)
#    nloci = len(y)
#    print("#loci:", nloci)
#
#    print("reading pacbio kmers")
#    x = {}
#    readKmers(PBfname, x, sort=True)
#
#    data = {}
#    for k, v in y.items():
#        if v.size and x[k].size:
#            data[k] = np.column_stack((x[k], y[k]))
#
#    results = np.zeros((nloci, 4))
#    for k, v in x.items():
#        truth = np.sum(v) / 2              ## divide by 2 since diploid individual [!] might be incorrect for CHM1 & CHM13
#        results[k,0] = truth
#
#    for k, v in data.items():
#        if plot and k < 50:   ## only plot the first 50 loci
#            a, _, rsquare, pred = PlotRegression(v[:,0:1], v[:,1:2],
#                                        "PacBioEdgeWeights", "IlluminaEdgeWeights",
#                                        "locus."+str(k)+".Sample"+str(v.shape[0]), out+"."+str(k), outlier="strict", pred=True, plot=True)
#        else:
#            a, _, rsquare, pred = PlotRegression(v[:,0:1], v[:,1:2],
#                                        "PacBioEdgeWeights", "IlluminaEdgeWeights",
#                                        "locus."+str(k)+".Sample"+str(v.shape[0]), out+"."+str(k), outlier="strict", pred=True, plot=False)
#        if k % 1000 == 0:
#            print(str(k)+" loci processed")
#        # divide by 2 since diploid individual [!] might be incorrect for CHM1 & CHM13
#        # TODO should not treat missing hap as zero length
#        results[k, 1:] = [pred/2, a, rsquare]   
#
#    print("writing outputs")
#    np.savetxt(out+".strict.pred", results, fmt=['%8.0f','%8.0f','%8.2f','%8.4f'], header="TrueLen\t PredLen\t Slope\t R^2")
#
#    print("plotting summary report")
#    if R2threshold != -1:
#        logic = (results[:,3] > R2threshold)
#        results = results[logic]
#    PlotRegression(results[:,0:1], results[:,1:2], "TrueLength", "PredictedLength",
#                        title="True.PredictedLength.Sample"+str(nloci) ,fname='.'.join([out,"sum",str(R2threshold)]),
#                        outlier="strict", plot=True)

def getrectangle(xs, ys):
    return  [xs[0], xs[0], xs[1], xs[1], xs[0]], [ys[0], ys[1], ys[1], ys[0], ys[0]]    

def getkmersindex(kmers):
    kmersi = {}
    invalid = 0xffffffffffffffff
    for i, kmer in enumerate(kmers):
        if kmer == invalid: continue
        if kmer not in kmersi:
            kmersi[kmer] = []
        kmersi[kmer].append(i)
    return kmersi

def getcokmersindex(cokmers, kmers0, kmers1):
    kmersi = [{}, {}]
    invalid = 0xffffffffffffffff
    for hap, kmers in enumerate([kmers0, kmers1]):
        for i, kmer in enumerate(kmers):
            if kmer == invalid: continue
            if kmer not in cokmers: continue
            if kmer not in kmersi[hap]: 
                kmersi[hap][kmer] = []
            kmersi[hap][kmer].append(i)
    return kmersi

def inregion(x, y, region): # region = ([x0,x1), [y0,y1))
    return x >= region[0][0] and x < region[0][1] and y >= region[1][0] and y < region[1][1]

def binarysearch(vec, val):
    """ind returned satisfies vec[ind-1] <= val < vec[ind]"""
    nitem = len(vec)
    if nitem == 0: return 0

    Li = 0
    Ri = nitem
    Mi = nitem//2
    while True:
        if vec[Mi] > val: # left search
            if Mi == (Li+Mi)//2: return Mi
            Ri = Mi
            Mi = (Li+Mi)//2
        elif vec[Mi] < val: # right search
            if Mi == (Ri+Mi)//2: return Mi+1
            Li = Mi
            Mi = (Ri+Mi)//2
        else:
            return Mi+1

def getbadkmc_bothhaps(indices0, indices1, region0, region1, fs=700, getindices=False):
    s0, e0 = region0
    ss0, ee0 = s0-fs, e0+fs
    s1, e1 = region1
    ss1, ee1 = s1-fs, e1+fs
    badregions = [((ss0,s0), (s1,e1)), # 0L
                  ((e0,ee0), (s1,e1)), # 0R
                  ((s0,e0), (ss1,s1)), # 1L
                  ((s0,e0), (e1,ee1))] # 1R

    badkmc = np.zeros(4, dtype=int) # 0L, 0R, 1L, 1R
    badindices = [[], []] # [[x0,...], [y0,...]]

    for i0 in indices0:
        if i0 < ss0 or i0 > ee0: continue
        for i1 in indices1:
            if i1 < ss1 or i1 > ee1: continue
            
            for j, badregion in enumerate(badregions):
                if inregion(i0, i1, badregion):
                    badkmc[j] += 1
                    if getindices:
                        badindices[0].append(i0)
                        badindices[1].append(i1)
    
    return (badkmc, badindices) if getindices else badkmc

def plotCrossContamination(ctg0, ctg1, ksize=21, FS=700, ax=None, zoomoutsize=(0,0), offset=(0,0), showFS=True, 
                           silent=False, showcontam=True, reportcontam=False, reportbad=False, size=0.1):
    """plot loci w/ primary contamination only"""
    
    start0, end0 = FS, len(ctg0)-FS
    start1, end1 = FS, len(ctg1)-FS
    TRsize0 = end0 - start0
    TRsize1 = end1 - start1

    relpos0 = (start0, end0-ksize+1)
    relpos1 = (start1, end1-ksize+1)
    kmers0 = read2kmers(ctg0, ksize, leftflank=-zoomoutsize[0]+offset[0], rightflank=-zoomoutsize[0]-offset[0])
    kmers1 = read2kmers(ctg1, ksize, leftflank=-zoomoutsize[1]+offset[1], rightflank=-zoomoutsize[1]-offset[1])
    cokmers = (set(kmers0) & set(kmers1)) - set([0xffffffffffffffff])
    cokmersi = getcokmersindex(cokmers, kmers0, kmers1)

    xs, ys, badxs, badys = [], [], [], []
    badkmc = np.zeros(4, dtype=int) # 0L, 0R, 1L, 1R
    for kmer in cokmers:
        indices0 = cokmersi[0][kmer]
        indices1 = cokmersi[1][kmer]

        # analyze contamination
        badkmc_, badindices = getbadkmc_bothhaps(indices0, indices1, relpos0, relpos1, getindices=True)
        badkmc += badkmc_

        # for plotting
        if ax is not None:
            for i in indices0:
                for j in indices1:
                    xs.append(i)
                    ys.append(j)

            for i in badindices[0]: badxs.append(i)
            for j in badindices[1]: badys.append(j)
                
    if ax is not None:
        if not silent:
            ax.set_title("contam={}, {:.1f}%".format(badkmc, 100*np.sum(badkmc)/(TRsize0+TRsize1)))

        if showFS:
            # TR + NTR region
            xlines, ylines = getrectangle((relpos0[0]-FS, relpos0[1]+FS), relpos1)
            ax.plot(xlines, ylines, '-r', linewidth=1, alpha=0.8)
            xlines, ylines = getrectangle(relpos0, (relpos1[0]-FS, relpos1[1]+FS))
            ax.plot(xlines, ylines, '-r', linewidth=1, alpha=0.8)

        ax.scatter(xs, ys, c='k', s=size, linewidths=0)
        if showcontam:
            ax.plot(badxs, badys, '.r')

        xmax = end0 - start0 + 2*FS + 2*zoomoutsize[0] - ksize + 1
        ymax = end1 - start1 + 2*FS + 2*zoomoutsize[1] - ksize + 1
        ax.set_xlim((0, xmax))
        ax.set_ylim((0, ymax))
        ax.set_yticks((0, ymax//500*500))
        ax.set_yticklabels([0, ymax//500*500], rotation=90)

    if reportbad:
        return badkmc

    if reportcontam:
        return badxs, badys

def visSelfRepeat(seq, ksize=13, figsize=(8,6), dpi=100, size=0.1, FS=700):
    fig, ax = plt.subplots(1,1, figsize=figsize, dpi=dpi)
    plotCrossContamination(seq, seq, ksize=ksize, ax=ax, zoomoutsize=(0,0), offset=(0,0), 
                            silent=False, showcontam=True, reportcontam=True, size=size, FS=FS)
    plt.show(); plt.close()

def visPairedRepeat(seq1, seq2, ksize=21, figsize=(15,4), dpi=100, silent=False, showcontam=True, reportcontam=True, size=0.1, FS=700):
    fig, axes = plt.subplots(1, 3, figsize=figsize, dpi=dpi)
    for i, pair in enumerate([(seq1,seq1), (seq1,seq2), (seq2,seq2)]):
        plotCrossContamination(pair[0], pair[1], ksize=ksize, ax=axes[i], zoomoutsize=(0,0), offset=(0,0),
                               silent=silent, showcontam=showcontam, reportcontam=reportcontam, size=size, FS=FS)
    plt.show(); plt.close()
```
