## Supplemental Data S1 for "The motif composition of variable-number tandem repeats impacts gene expression": README.txt

Header of `vntr.statistics.tsv`:
	chr.raw:	TRF annotation for the chromosome of the TR
	start.raw:	TRF annotation for the start coordinate of the TR on hg38
	end.raw:	TRF annotation for the end coordinate of the TR on hg38
	chr:	danbing-tk annotation for the chromosome of the TR
	start:	danbing-tk annotation for the start coordinate of the TR on hg38
	end:	danbing-tk annotation for the end coordinate of the TR on hg38
	ref.len:	length of the TR on hg38
	QC.mobile:	pass QC filter: not overlapping RepeatMasker mobile elements
	n.hap:	num of supporting HGSVC haplotypes
	asm.len.mn:	mean(assembly TR size)
	asm.len.sd:	s.d.(assembly TR size)
	n.al.len:	num of alleles by TR size
	n.al.mtf:	num of alleles by motif composition
	mmr.ref:	mismap rate in hg38
	mmr.asm:	mean(mismap rate in assembly)
	aln-r2:	aln-r2
	QC.mmr.ref:	pass QC filter: `mmr.ref` < 1%
	QC.mmr.asm:	pass QC filter: `mmr.asm` < 5%
	QC.aln-r2:	pass QC filter: `aln-r2` > 0.5
	QC.all:	pass all QC filters
	n.kmer:	num of kmers in the TR
	n.motif:	num of motifs (unitigs in compact de Bruijn graph) in the TR
	b-r2.HG:	batch-r2 of TR length in HGSVC
	b-r2.mt.HG:	batch-r2 of motif in HGSVC
	b-r2.HP:	batch-r2 of TR length in HPRC
	b-r2.mt.HP:	batch-r2 of motif in HPRC
	is.eTR:	is an eTR in gene-level discoveries, due to length of motif
	is.eTR.len:	the eTR is associated with gene by length
	n.eGene:	num of genes associated with the TR
	n.eMotif:	num of eMotifs in the TR
	is.fm:	is a fine-mapped TR (susieR PIP > 0.8), due to length or motif
	is.fm.len:	is fine-mapped to the length of the TR
	n.fm.mtf:	num of fine-mapped motifs in the TR
	n.fm.gene:	num of genes associated with the fine-mapped variants
Columns in `VNTR_annot.hg38_raw.hg38_refined.35HGSVC_asm.bed`:
	col 1-3:	TRF annotation for the TR on hg38
	col 4-6:	danbing-tk annotation for the TR on hg38
	col 7-:	danbing-tk TR annotations for 35 HGSVC assemblies.
		Each assembly has 2 haplotypes, maternal first, then paternal, e.g. HG00096.0 and HG00096.1.
		Each haplotype has 4 fields: chr, start, end, strand.
		Strand: +/- means the TR sequence of the haplotype maps to the +/- strand of hg38.
		Assemblies are shown in order of the genome names as follows:
			HG00096
			HG00171
			HG00512
			HG00513
			HG00514
			HG00731
			HG00732
			HG00733
			HG00864
			HG01114
			HG01505
			HG01596
			HG02011
			HG02492
			HG02587
			HG02818
			HG03009
			HG03065
			HG03125
			HG03371
			HG03486
			HG03683
			HG03732
			NA12329
			NA12878
			NA18534
			NA18939
			NA19238
			NA19239
			NA19240
			NA19650
			NA19983
			NA20509
			NA20847
			NA24385
