## Supplemental Data S2 for "The motif composition of variable-number tandem repeats impacts gene expression": README.txt

Header explanation:
	tissue:	tissue where the gene is associated with the TR length in gene-level discoveries
	gene_id: GTEx Ensembl gene ID
	gene_name:	GTEx gene name
	chr:	chromosome of the gene
	start:	start coordinate of the gene on hg38
	end:	end coordinate of the gene on hg38
	num_var:	number of TRs tested for this gene
	TR_chr:	chromosome of the TR locus
	TR_start:	start choordinate of the TR locus on hg38
	TR_end:	end coordinate of the TR locus on hg38
	TR_locus:	index of the TR locus in our database, See the tables in Supplemental Data S1
	eTR:	the lead-TR is significant under gene-level discoveries
	lead_TR:	the TR has the lowest p-value among all TRs tested against this gene
	lead_p_nominal:	the min p-value among all TRs tested against this gene
	p_nominal:	the p-value of this TR
	slope:	the effect size of this TR
	slope_se:	the standard error of effect size of this TR
	qval:	the q-value from the Benjamini–Hochberg procedure
